# Nuclear Type I Myosins are Essential for Life and Genome Organization

**DOI:** 10.1101/2024.09.26.615191

**Authors:** Audrey Yi Tyan Peng, Jianhui Li, Brian C. Freeman

## Abstract

The active transport of large biomolecules within a cell is critical for homeostasis. While the cytoplasmic process is well-studied, how the spacing of nucleoplasmic cargo is coordinated is poorly understood. We investigated the impact of myosin motors in the nucleus of budding yeast. We found that life requires a nuclear type I myosin whereas the essential type II or V myosins were not requisite in the nucleus. Nuclear depletion of type I myosins triggered 3D genome disorganization, nucleolar disruption, broad gene expression changes, and nuclear membrane morphology collapse. Genome disorganization occurred first supporting a model where type I myosins actively maintain genome architecture that scaffolds nuclear membrane and nucleolar morphologies. Overall, nuclear myosin is critical for the form and function of the nucleus.

## Main Text

The motor proteins myosin, dynein, and kinesin are central to homeostasis for effectively moving and positioning the numerous large and complex biomolecules within the intricate and varying environments of eukaryotic cells (*1*). Myosin motors mobilize a broad array of cargoes participating in diverse processes including cytokinesis, organelle function, membrane dynamics, and cell polarization (*2*). To transport cargo, myosins generate ATP-dependent contractile forces to move along actin polymers (*2*). To travel long distances, myosins can rely on dynamic actin polymers comparable to the actin species observed in the nucleus (*3*, *4*). To handle the complexity of biological molecules requiring transport, the myosin family has evolved into a superfamily represented by 18 different classes with the largest and oldest subfamily being the type I myosins (*5*). Typically, myosins are considered strict cytosolic factors, yet recent studies show myosins contributing to diverse nuclear events including gene regulation, chromatin remodeling, and DNA repair (*6*). Despite these discoveries, our appreciation of the physiological relevance of nuclear myosin is limited.

The first myosin established to work in the nucleus was the B isoform of unconventional Myosin 1C, Nuclear Myosin I (NMI) (*7*). NMI has since been shown to contribute to RNA polymerases I and II activities, chromosome territory movements, and DNA repair (*8*).

For instance, human chromosomes 10, 18, and X reposition in a serum-dependent fashion unless myosin IC expression is reduced or actin polymerization is impaired (*9*). Similarly, directed long-range motion of an activated gene or repair of damaged DNA relies on NMI, though DNA damage repair also involves myosin IA and V (*10*, *11*).

Despite contributing to multiple, fundamental nuclear pathways, NMI is not broadly expressed in eukaryotes (*5*). Whether NMI is functionally replaced by other myosins or if nuclear myosin is optional, had not been established. Altogether, myosin I, II, Va, Vb, VI, X, XVI, and XVIII work in mammalian nuclei, which complicates our ability to fully understand the overall importance of nuclear myosin (*12*). To characterize the physiological relevance of myosin in the nucleus we exploited budding yeast where the myosin repertoire is comparatively rudimentary being comprised of 5 proteins representing myosin I, II, and V subfamilies (*5*). Of note, yeast does not express a B isoform of myosin I but rather has paralogs of myosin 1E/F encoded by *MYO3* and *MYO5* (*13*). In our previous work, we found that Myo3 mobilized a gene locus to the nuclear periphery following activation, yet another study disputed the need for Myo3 in gene motion (*4*, *14*). Hence, the role of yeast type I myosins in genome reorganization had been debated.

### Nuclear type I myosin is essential

The budding yeast genes encoding type II (*MYO1*) and V (*MYO2*) myosins are essential and the type I genes (*MYO3* and *MYO5*) are synthetically lethal including in our strain background (*15*-*17*; fig. S1A). To assess the importance of each myosin in the nucleus, we employed the anchor-away tactic to controllably deplete a target protein from the nucleus (*18*). In brief, we engineered yeast expressing each myosin as a fusion with the FKBP rapamycin binding (FRB) domain and the ribosomal subunit Rpl13A tagged with FKBP12. Addition of rapamycin to the media triggers tethering of the FRB fusion to the Rpl13A-FKBP12 protein thereby shuttling the complex out of the nucleus during the process of ribosome biogenesis (*18*). The nuclear depletion of either essential myosin, Myo1 or Myo2, did not impact growth, nor did the nuclear loss of the nonessential Myo3, Myo4, or Myo5 myosins (Fig. 1A). In contrast, shuttling out Myo3 in a *myo5*Δ background or Myo5 in *myo3*Δ resulted in a growth arrest phenotype in both a spot-test and liquid culture (Fig. 1B & S1B). Hence, a nuclear presence of type I myosin likely contributes to the synthetic lethal phenotype between Myo3 and Myo5. Of note, flow cytometry analysis indicated that nuclear depletion of type I myosin triggered cell cycle arrest regardless of stage unless the yeast were synchronized *a priori* then the cells arrested in G2/M (fig. S1D & S1E). Hereon, we will use the *myo5*Δ strain expressing Myo3-FRB and refer to it as Myo3-AA.

**Fig. 1.**
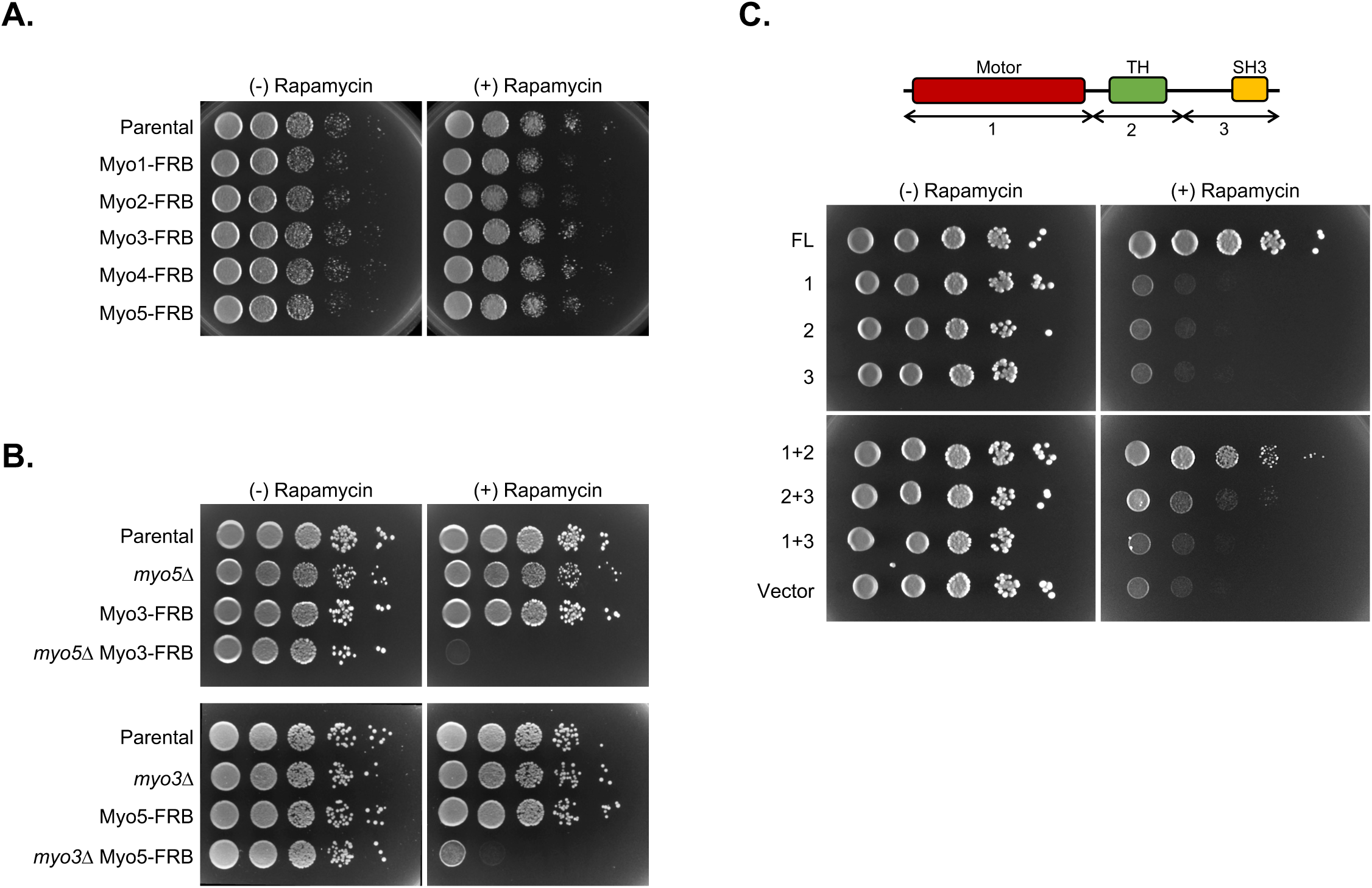
Nuclear type I myosin is essential. (**A**) The influence on cell viability of each myosin in the nucleus was checked using a combination of the anchor-away assay and spot-tests. Logarithmic phase parental yeast and strains expressing each myosin homolog as an FRB fusion (e.g., Myo1-FRB) were serially diluted and spotted onto media without and with rapamycin, as indicated. (**B**) The nuclear contributions to the synthetic lethal phenotype between the type I myosin paralogs, Myo3 and Myo5, were assessed with the anchor-away/spot-tests. Parental, *myo5Δ*, and *myo3Δ*, and strains expressing either Myo3 or Myo5 as an FRB fusion were serially diluted and spotted onto media without and with rapamycin, as marked. (**C**) The sufficiency of HA-tagged full-length (FL) Myo3 or truncated variants with the indicated combination of the amino-terminal motor (1), central tail homology (TH) (2), or carboxyl-terminal SH3 (3) domain(s) to support cell growth following the nuclear depletion of Myo3-FRB in *myo5Δ* cells was determined using the anchor-away/spot tests with cells grown on media without and with rapamycin, as indicated.

To further characterize the nuclear role of type I myosin, we tested which protein domains were required to support life. The E/F isoforms of type I myosins have 3 conserved domains including an amino-terminal motor, middle tail homology (TH) domain, and a carboxyl-terminal SH3 (*5*). Expression of full-length HA-tagged Myo3 in the Myo3-AA background rescued growth in the presence of rapamycin as did a fusion of the motor and TH domains, yet the other variants did not (Fig. 1C & S1C). Previously, we found that the motor and TH domains were sufficient to support localized genome reorganization in response to gene activation (*4*). Given our prior finding, we next examined whether nuclear type I myosin might have a general role in genome organization.

### Nuclear type I myosin is required for the maintenance of genome organization

To explore whether nuclear type I myosin influences 3D genome architecture, we applied the High-throughput Chromosome Conformation Capture (Hi-C) assay to Myo3-AA cells treated with DMSO or rapamycin for 2- or 5-hours. We plotted the chromosome contact frequency maps of each condition and identified changes in the maps by comparing the 2- and 5-hour patterns to the control map (Fig. 2A & S2A). Using averaged maps from biological triplicates, it was apparent that depletion of nuclear type I myosin resulted in extensive intra- and interchromosomal contact frequency differences across the genome that intensified with time (Fig. 2A). Importantly, the contact frequency differences were reproducible (fig. S2B), and the magnitude of the observed changes is comparable to the reported loss of the known genome organizers cohesin and chromatin remodelers (*19*). Yet, the breadth of contact changes were more expansive for nuclear myosin loss with the effects being apparent across the entire genome and not restricted to certain sites or by distance. Focusing on intrachromosomal contacts highlights the common decline in distant interactions and increase in local associations (Fig. 2B). Further assessment of the distance-based interactions found 100 kB to be the pivot point, as contacts generally increased below 100 kB and decreased above 100 kB for all the chromosomes apart from chromosome XII (Fig. 2C & S2C). The overall shift in contact frequencies suggests a possible collapse of the 3D genomic architecture where distal connections are lost and local contacts increased because of proximity as the genome seemingly folds in on itself.

**Fig 2.**
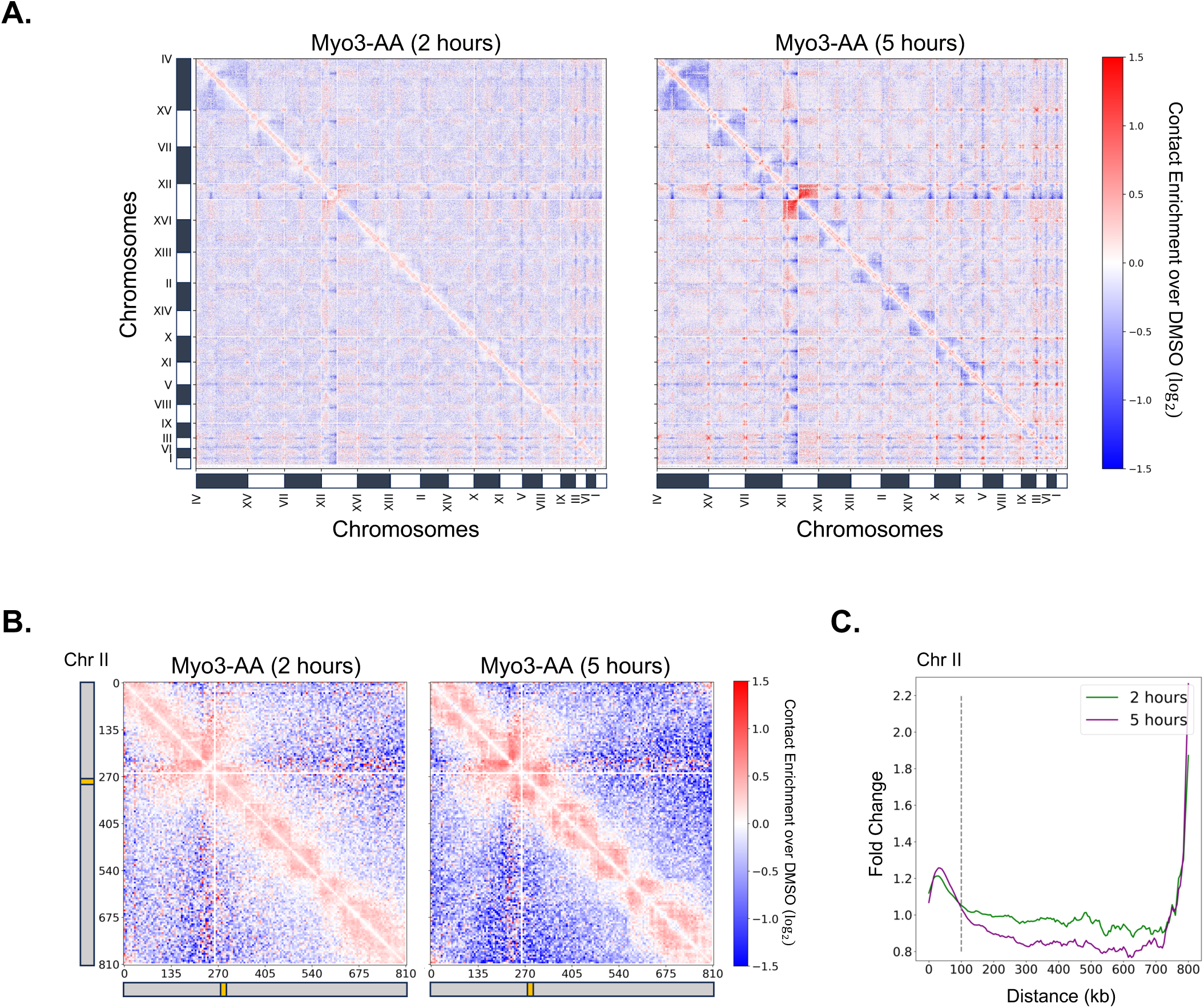
Genome organization is dependent on type I myosin in the nucleus. **(A)** The reliance of 3D genome architecture on nuclear type I myosin was determined by comparing the genome-wide contact frequency maps (5 kb resolution; fig. S2) of control and 2 or 5 h rapamycin treated Myo3-AA cells. The log_2_ ratio matrices are shown. (**B**) Chromosome localized changes were assessed using a log_2_ ratio matrices of contact maps (5 kb resolution) from Myo3-AA yeast treated for 2 or 5 h with rapamycin versus control (DMSO) samples. The results for chromosome II are shown. (**C**) The relationship between genomic contact frequency changes and DNA distance is shown for chromosome II with the 100 kb position of the chromosome marked with a grey dotted line.

The overall organization of budding yeast chromosomes is typically described as being in a Rabl configuration where centromeres group at the spindle pole body and telomeres form multiple clusters along the nuclear envelope (*20*, *21*). Depletion of nuclear type I myosin induced contact frequency changes in both of these Rabl hallmarks. The interchromosomal contacts between centromeres decreased (orange arrows fig. S2E and S2F upper panels) and intra- and interchromosomal interactions between telomeres increased (green arrows fig. S2E and S2F lower panels). Of note, as the centromere-to-centromere interactions declined a corresponding increase in centromere-to-noncentromere associations were observed, which is apparent by the rise in contacts running between centromeres (fig. S2E contacts between orange arrows). In addition to their Rabl-association, telomeres and centromeres are classic sites of heterochromatin formation along with ribosomal DNA (*21*).

### The nucleolus is dependent upon nuclear type I myosin

A prominent shift in genomic contacts occurred across Chromosome XII where the ribosomal DNA (rDNA) repeats are housed (Fig. 2A & 3A). Given the highly repetitive nature of rDNA, it is difficult to properly align the DNA sequence reads, and therefore the rDNA region is not mapped in a Hi-C assay (white lines in Fig. 3A). Since rDNA seeds and is encapsulated by the nucleolus, the Chromosome XII regions to the left and right of rDNA are typically well-separated (*22*). Yet, upon depletion of nuclear type I myosin, the contact frequencies between these two areas of chromosome XII increased with a corresponding decrease in interactions between the DNA on the same side of the rDNA repeats (Fig. 3A).

**Fig. 3.**
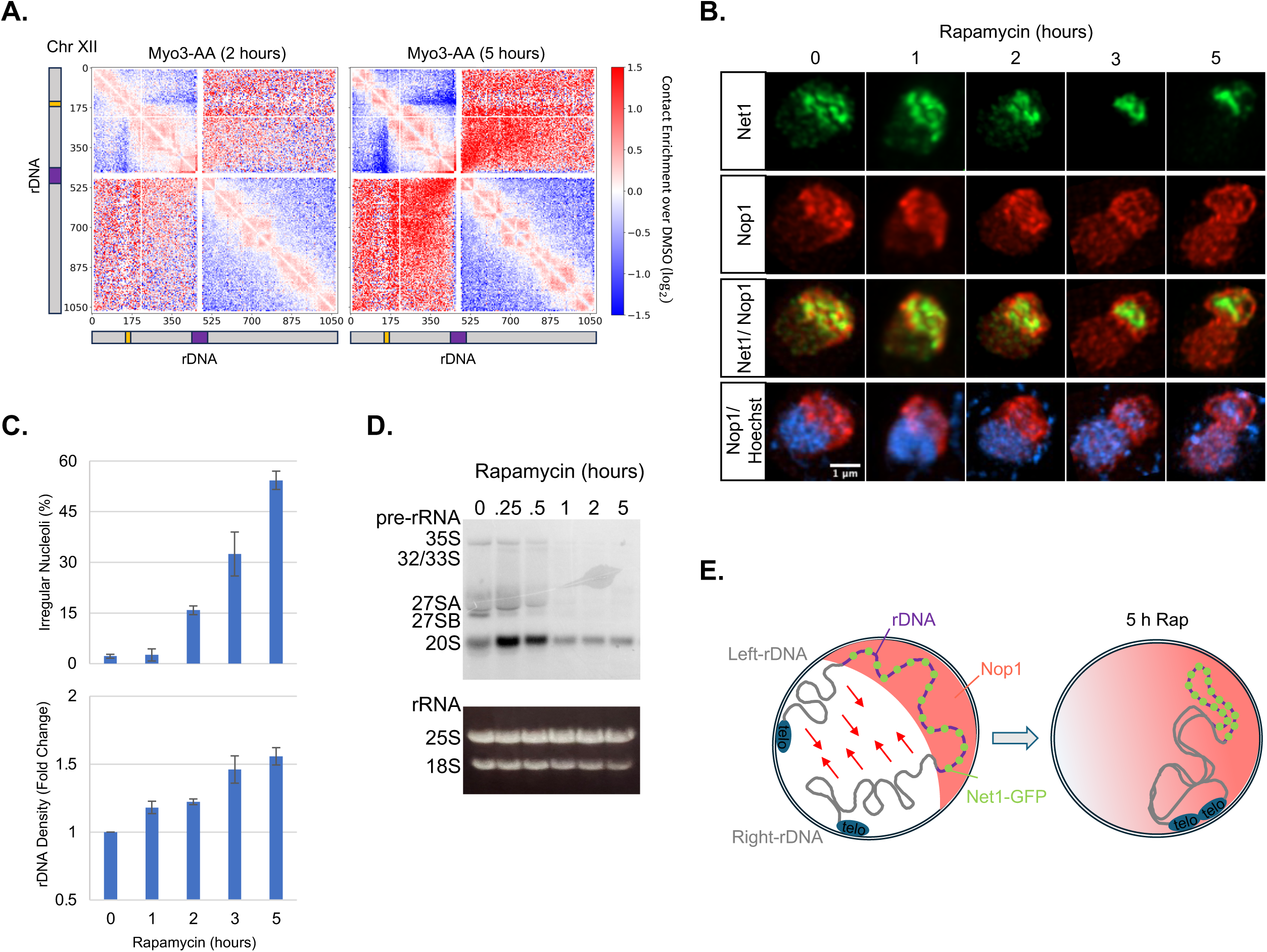
Nucleolar integrity relies on nuclear type I myosin. **(A)** The 3D architecture of chromosome XII, which houses the rDNA repeats (marked with purple box), was assessed with log_2_ ratio matrices of contact maps (5 kb resolution) for 2 or 5 h rapamycin treated Myo3-AA yeast versus DMSO control. (**B**) The rDNA repeats were visualized using the rDNA binding protein Net1 fused to GFP (green), the nucleolar space was followed by staining for the yeast fibrillarin homolog using an anti-Nop1 antibody (red), and the genomic DNA was detected using Hoechst stain (blue). (**C**) The time-dependent changes observed in nucleolar shape and rDNA density after rapamycin addition were quantified, as marked. A total of 50 cells were counted in triplicate. Error bars represent the standard error of the mean. (**D**) The production of pre-rRNAs was monitored after rapamycin addition for the indicated times in Myo3-AA cells by northern blotting using probes hybridizing to the ITS1 and ITS2 regions of pre-RNAs (upper panel). Mature rRNAs were detected by ethidium bromide staining (lower panel). (**E**) A model illustrating the changes to genome organization, rDNA configuration, and nucleolar integrity triggered by the nuclear depletion of type I myosin is presented. The nucleus on the left is undergoing nuclear depletion of type I myosin and the one on the right is after 5 hours of rapamycin treatment. The red arrows in the left nucleus represent the increasing interactions between the DNA of chromosome XII on either side of the rDNA repeats (purple) that occur as type I myosin is depleted from the nucleus and the genome disorganizes. The Net1-GFP (green circles) bound to the rDNA repeats, Nop1 immunofluorescence staining (coral area), and telomeric DNA repeats of chromosome XII (blue ovals) highlight additional outcomes of depleting nuclear type I myosin.

To interrogate the rDNA conformation, we visualized a GFP-Net1 fusion (Net1 binds to rDNA repeats (*23*)), and the nucleolar space was viewed using the yeast fibrillarin homolog Nop1. In DMSO-treated control cells, Net1 is dispersed throughout the nucleolus and closely demarcated by Nop1 (Fig. 3B). Following addition of rapamycin, the Net1 signal condensed and intensified consistent with a breakdown in the 3D organization of the rDNA (Fig. 3B). In contrast, the Nop1 signal spread during the time course eventually dispersing across the nucleus at the 5-hour time point (Fig. 3B). Quantification of the Nop1 and Net1 signals confirmed the disfigurement of the nucleolar space (Nop1) and condensation of the rDNA (Net1) (Fig. 3C). To assess if these changes affected nucleolar function, we analyzed rRNA biogenesis by northern blot analysis probing for the pre-rRNA spacer regions ITS1 and ITS2 (*24*). Following rapamycin addition, we observed a time-dependent declined production of nascent 35S and ITS1/2 pre-rRNAs (Fig. 3D). Hence, the form and function of nucleoli are reliant on nuclear type I myosin, as depletion of the motor from this central compartment triggers disorganization of the genome and disruption of the nucleolus (Fig. 3E).

### Changes in genome organization do not correlate with gene expression

The genomes of most, if not all, organisms are organized into higher-order structures (*25*). A prevailing concept for the 3D organization is to contribute to the transcriptional control of gene promoters by clustering coregulated loci into transcription factories (*26*). As the nuclear depletion of type I myosin had a genome-wide impact on organization (Fig. 2), we assessed whether cells mount a transcriptional response to the nuclear depletion of type I myosin and if any observed changes correlate with shifts in genomic architecture.

We subjected Myo3-AA cells to DMSO or rapamycin for 2- or 5-hours, collected RNA, and performed RNA-seq. At 2-hours, 398 genes were up-regulated ≥2-fold and 199 down-regulated ≥50% and at 5-hours, 609 were activated ≥2-fold and 368 decreased ≥50% (Fig. 4A). Gene Ontology (GO) analysis did not reveal enrichment in any specific category (fig. S3A). As myosin VI is known to alter the nuclear localization of RNA polymerase II in human cells (*27*), we examined the positioning of RNA polymerase II in Myo3-AA in the presence of rapamycin. No differences were apparent across the time course (fig. S3B). Next, we checked whether the gene regulatory fluctuations correlated with genome organization.

**Fig 4.**
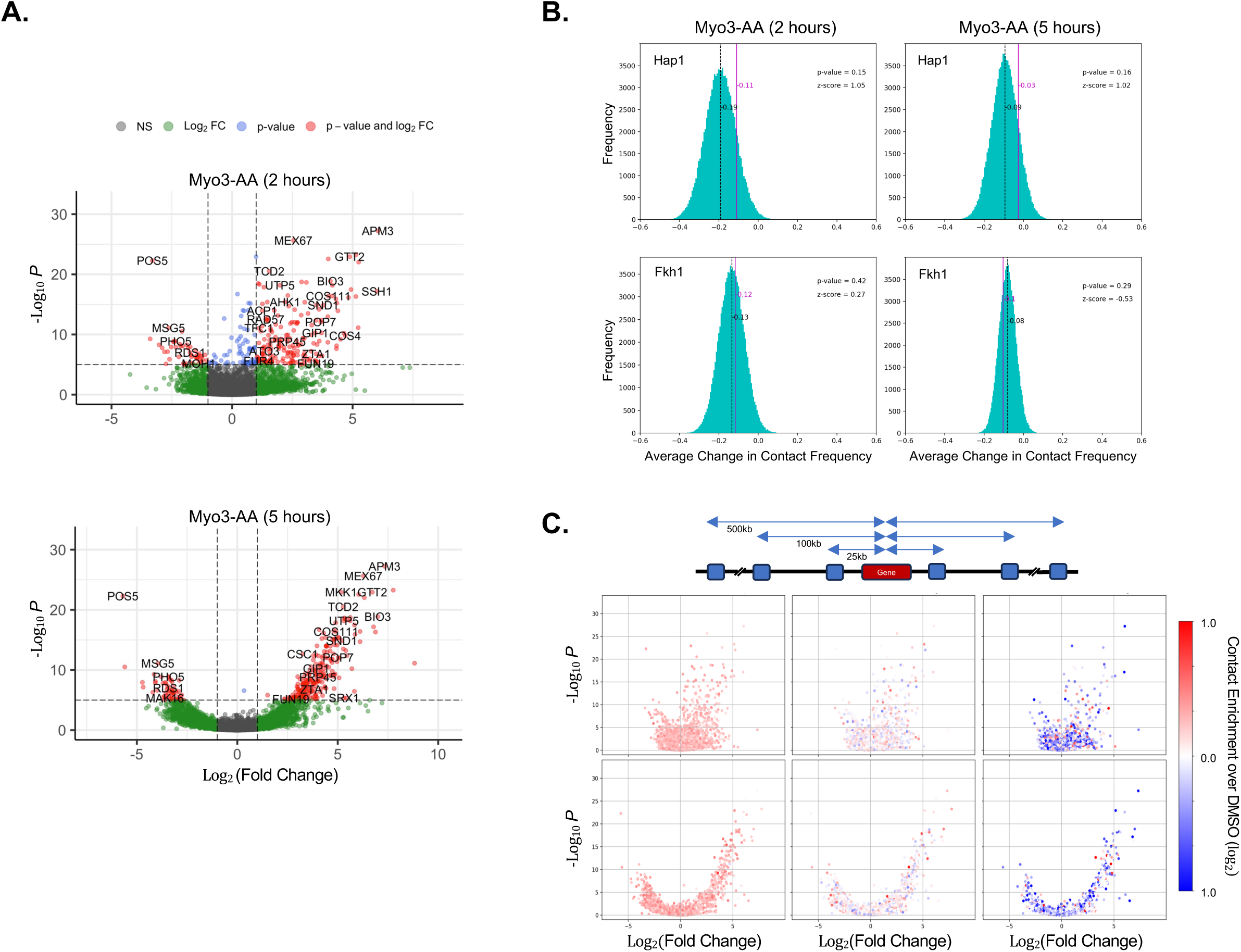
Genome disorganization and gene expression fluctuations are noncorrelative in response to the nuclear depletion of type I myosin. **(A)** Gene expression changes in Myo3-AA cells subjected to 2 (upper) or 5 (lower) hours of rapamycin. The data were normalized to control (DMSO) samples and visualized using volcano plots. (**B**) The distribution of contract frequency changes between Hap1- or Fkh1-bound (upper or lower panels, respectively) differentially expressed gene promoters and random genes are shown in teal and the average value is designated with the dotted black line in each. The purple lines represents the average contact frequency changes between Hap1 (upper) or Fkh1 (lower) regulated gene promoters. (**C**) Gene expression volcano plots of Myo3-AA cells treated with rapamycin for 2 or 5 hours were color-coded by the change in the log_2_ ratio of contact frequencies (5 kb window) between each gene and chromosome sites at the the set distances of 25, 100, or 500 kb from the gene of interest, as indicated.

An elementary assessment of the physical distribution of the differentially expressed genes across the chromosomes showed no set pattern including proximity to centromeres, telomeres, or rDNA (fig. S3C). We then checked the potential clustering of coregulated genes by determining the propensity of activated genes to gain genomic contacts (i.e., form a transcription factory) or for repressed genes to lose contacts (i.e., disperse from a factory). As a first step, we categorized the differentially expressed genes based upon the bound transcription factors at each locus using YEASTRACT+ (28). We applied permutation tests as an unbiased means to gauge whether genes associated with the same transcription factor interacted more/less with each other relative to random DNA. We examined transcription factors associated with genes that were activated or repressed following the nuclear depletion of type I myosin (Table S1 & S2). For illustrative purposes, we present the outcomes for Hap1 (associated with activated genes) and Fkh1 (associated with repressed genes). Significantly, the changes in contact frequencies between Hap1- or Fkh1-bound genes were similar to the differences in contacts with random genes (Fig. 4B).

As the exact determinants driving transcription factories are yet to be defined, we next took a more open-ended approach by checking for correlative changes in all contact frequencies with the expectation that up-regulated genes would gain interactions if a transcription factory were built, and down-regulated genes would lose contacts as a factory is dispersed. In assessing this concept, we considered our previous observation that the genome is seemingly folding in on itself, as distant contacts decline while local interactions increase (Fig. 2C), and therefore determined the fluctuations in contact frequencies for differentially expressed genes based on relative distances from each gene. We calculated changes to contact frequencies between each differentially expressed gene at distances of 25, 100, or 500 kB using a 5 kB interaction window and then overlayed the results onto the gene expression volcano plots (Fig. 4C). In general, the shifts in contact frequencies followed the global changes in genome organization (i.e., distal interactions decreased and proximal interactions increased) and did not correlate with gene expression changes. In brief, our data do not support the involvement of transcription factories in controlling yeast gene expression in reaction to nuclear type I myosin depletion.

### Collapse in nuclear morphology follows genome disorganization

The spatial architecture of mammalian chromatin is associated with the mechanical stability of the nucleus, with abnormal nuclear shapes linked to various human diseases (29). To determine if the genome disorganization is accompanied by nuclear morphology changes, we subjected Myo3-AA yeast expressing either GFP-Pho88 or GFP-Nup49, which mark the outer and nuclear membranes or just the nuclear membrane, respectively, to DMSO or rapamycin. We observed a time-dependent disfigurement of the nuclear membrane beginning ∼1 hour after rapamycin addition (Fig. 5A and S4A). By 5 hours post rapamycin addition, ∼50% of the nuclei were altered (Fig. 5B and S4B). Thus, nuclear morphology relies on nuclear type I myosin.

**Fig. 5.**
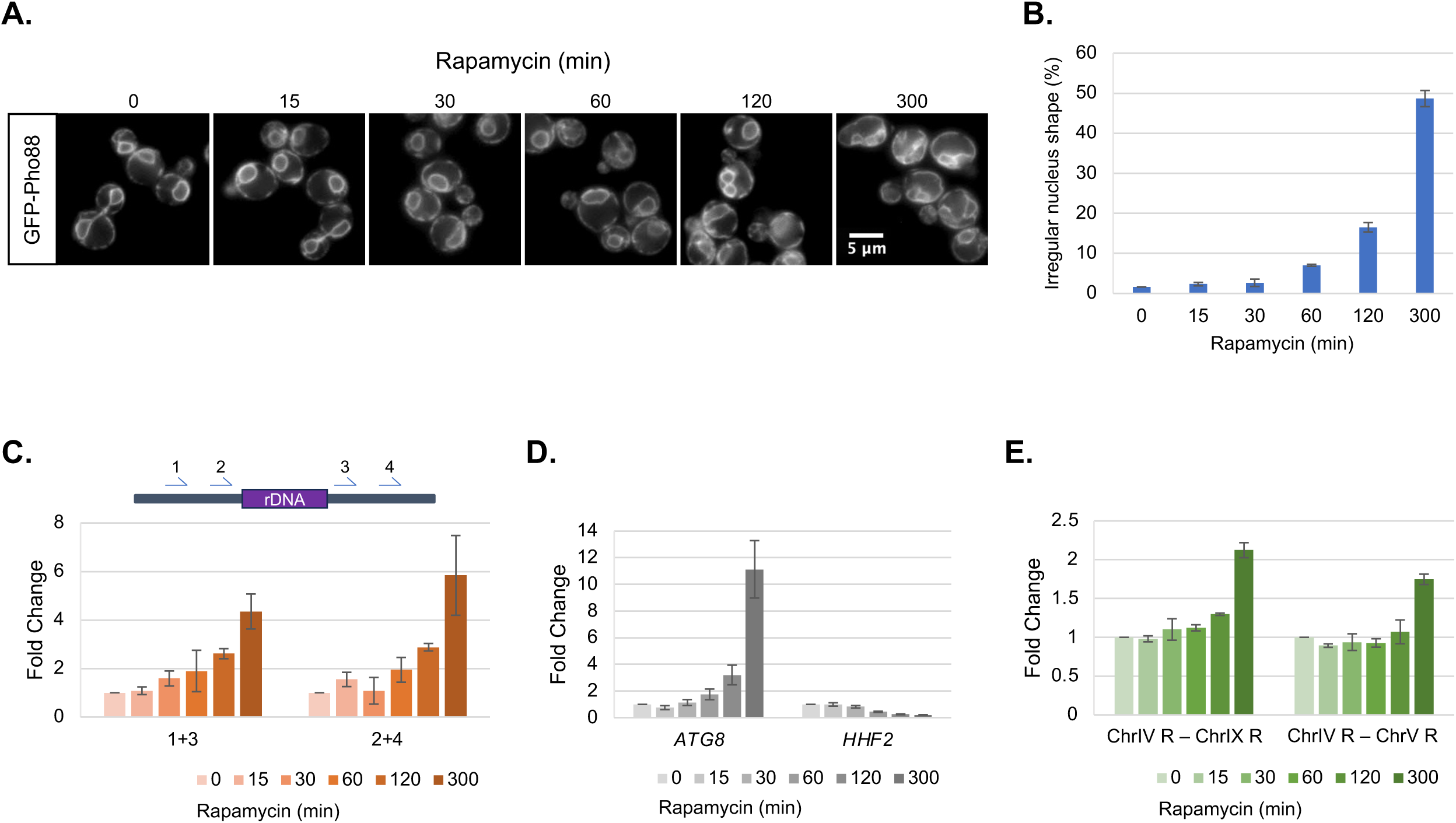
Nuclear envelope morphology is dependent on nuclear type I myosin. **(A)** The nuclear shape of Myo3-AA cells expressing the membrane marker GFP-Pho88 and treated with DMSO (control) or rapamycin for the indicated times was monitored by fluorescence microscopy. (**B**) The percent of cells with irregular nuclei was determined and quantified (50 per trial; n=3). (**C**) The time-dependent changes in genomic contacts between DNA sites on either side of the rDNA repeats following the addition of rapamycin to Myo3-AA yeast were assessed by 3C-qPCR using 2 different primer sets, as indicated. (**D**) The time-dependent changes in RNA levels at an induced (*ATG8*) and repressed (*HHF2*) gene target were determined by RT-qPCR following the addition of rapamycin to Myo3-AA cells at the marked time points. (**E**) The time-dependent changes in contact frequencies between the indicated chromosome telomeres were followed by 3C-qPCR following the addition of rapamycin to Myo3-AA cells. For all graphs, error bars represent standard error of the mean.

To further characterize the changes triggered by the nuclear depletion of type I myosin, we delineated the kinetics of gene expression, genome organization, and nuclear shape changes. The earliest detected shift was at the 15 min time point in genome reorganization around the rDNA repeats (Fig. 5C), followed by gene expression changes (Fig. 5D), then telomere clustering (Fig. 5E) in conjunction with nuclear membrane morphology collapse (Fig. 5A and S4). Thus, reorganization of the genome was the lead change (besides telomeres), followed by gene expression fluctuations, and then nuclear envelope shape deformation in conjunction with telomere grouping.

## Discussion

Our data reveal a broad influence of type I myosin on nuclear features including genome organization (Fig. 2), nucleolar integrity (Fig. 3), gene regulation (Fig. 4), and nuclear membrane morphology (Fig. 5). Kinetic analysis showed that genome architecture is a leading target of nuclear type I myosin with the exception of telomeres (Fig. 5). Given that chromatin entwined chromosomes are some of the largest biomolecules in a cell, it is unsurprising that these structures require motor driven control. Certainly, active chromosome transport following each cell division would be highly beneficial to establish the correct 3D genome organization by nominally fostering the directed placement of each chromosome region into a set position while also avoiding the entanglement of these long polymers (*30*). Yet, our data indicate that nuclear type I myosin is needed throughout the cell cycle and not just after cell division (Fig. 1 & S1), suggesting a persistent use of a type I motor. Our work opens up many interesting questions. Might nuclear myosin account for the variances in genome organization detected within single cells relative to the population average or is myosin resetting drifting architectures to a steady position (*25*)? Irrespective of the timing of motor activity, how are the actions of type I myosins coordinated? Do myosins rely on allosterically adjusted transcription factors to selectively control genome positioning (*4*), or is the epigenetic code, which correlates with 3D architecture (*31*), exploited by the motors to dictate genome positioning. Significantly, our work establishes that nuclear type I myosins are key factors for actively maintaining genome organization.

In addition to the general genome organization, we believe that the myosin-regulated architecture also contributes to nuclear envelope and nucleolar integrities. It is well established that rDNA is a central nucleolar component (*22*). Our data show that disrupting the configuration of rDNA correlates with the loss of nucleolar form and function (Fig. 3), suggesting that rDNA scaffolds nucleoli. Comparably, the 3D structure of the genome also appears to support the nuclear membrane, thereby, rationalizing why the envelope disfigures following depletion of nuclear myosin (Fig. 5). Alternatively, the telomeres, which tether to the nuclear membrane (*21*), might pull the envelope inward as the genome collapses. Yet not all nuclear activities correlated with shifts in genome organization. Unexpectedly, the robust fluctuations in gene expression triggered by the nuclear depletion of type I myosin were noncorrelative to changes in the genome architecture suggesting that yeast do not rely on transcription factories to mount this gene program (Fig. 4). Overall, it is apparent that having a type I myosin in the nucleus is critical to support the form and function of the nucleus.

Although nuclear myosin has long been recognized (*7*), how prominent a role the motor has in this compartment, had been argued (*14*). Minimally, our work shows that a nuclear type I myosin is required to support life (Fig. 1). It was, perhaps, unexpected that nuclear type I myosin would influence such a broad range of pathways, yet if we consider the minimal repertoire of myosins in budding yeast and all the reported roles of different myosins (e.g., I, II, V, VI, X, XVI, and XVIII) in the nuclei of larger eukaryotes (*12*), then it is plausible. For example, mammalian myosin VI was recently shown to maintain nucleolar structure without altering RNA polymerase I transcription yet NMI and Vb modulate RNA polymerase I activity without altering nucleoli (*32–34*). Hence, different mammalian myosins carry out similar roles performed by yeast type I myosins. We suspect that the evolution of the myosin family included expansion and specialization of myosins to efficiently control the higher-ordered complexities of nuclear features in larger eukaryotes. Our work is consistent with the yeast type I myosins, Myo3 and Myo5, serving as primordial myosin motors supporting central nuclear process including the 3D organization of the genome and full operation of the nucleolus and nucleus. Perhaps significantly, variants in the human myosins 1E/F homologs have been associated with a variety of diseases from cancers to kidney diseases yet the causative mechanism(s) for the pathologies often remain elusive (*35*). Our findings provide a new avenue by which myosin 1E/F motors might contribute to human health.

## Acknowledgments

We are grateful to Dr. Maya Schuldiner (Weizmann Institute of Science) for her kind gift of the amino-terminal GFP fusion library.

## Author contributions

Conceptualization: AYTP, BCF

Methodology: AYTP, JL, BCF

Investigation: AYTP, BCF

Visualization: AYTP, BCF

Funding acquisition: BCF

Project administration: BCF

Supervision: BCF

Writing – original draft: AYTP

Writing – review & editing: BCF

## Data and materials availability

Accession number: PRJNA1156761

Code: Github at https://github.com/audreypeng/Genome-Organization-Myosin.git

