## Supplemental Figures for "Nuclear Type I Myosins are Essential for Life and Genome Organization"

fig. S1

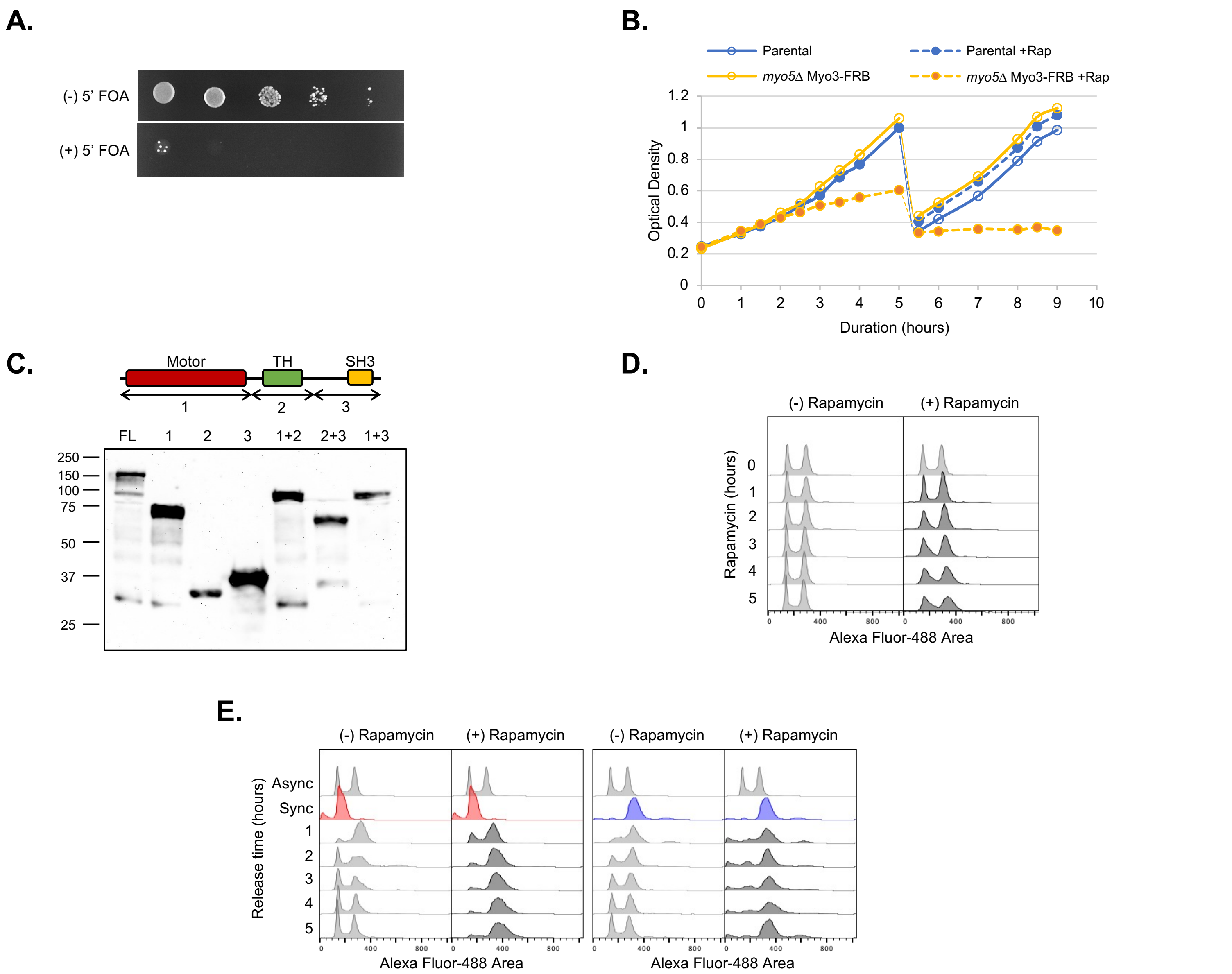

A.
(-) 5' FOA
(+) 5' FOA
B.
Parental
Parental +Rap
myo5Δ Myo3-FRB
myo5Δ Myo3-FRB +Rap
Optical Density
Duration (hours)
C.
Motor
TH
SH3
1
2
3
FL
1+2
2+3
1+3
250
150
100
75
50
37
25
D.
(-) Rapamycin
(+) Rapamycin
Rapamycin (hours)
Alexa Fluor-488 Area
E.
(-) Rapamycin
(+) Rapamycin
Async
Sync
Release time (hours)
Alexa Fluor-488 Area

fig. S2

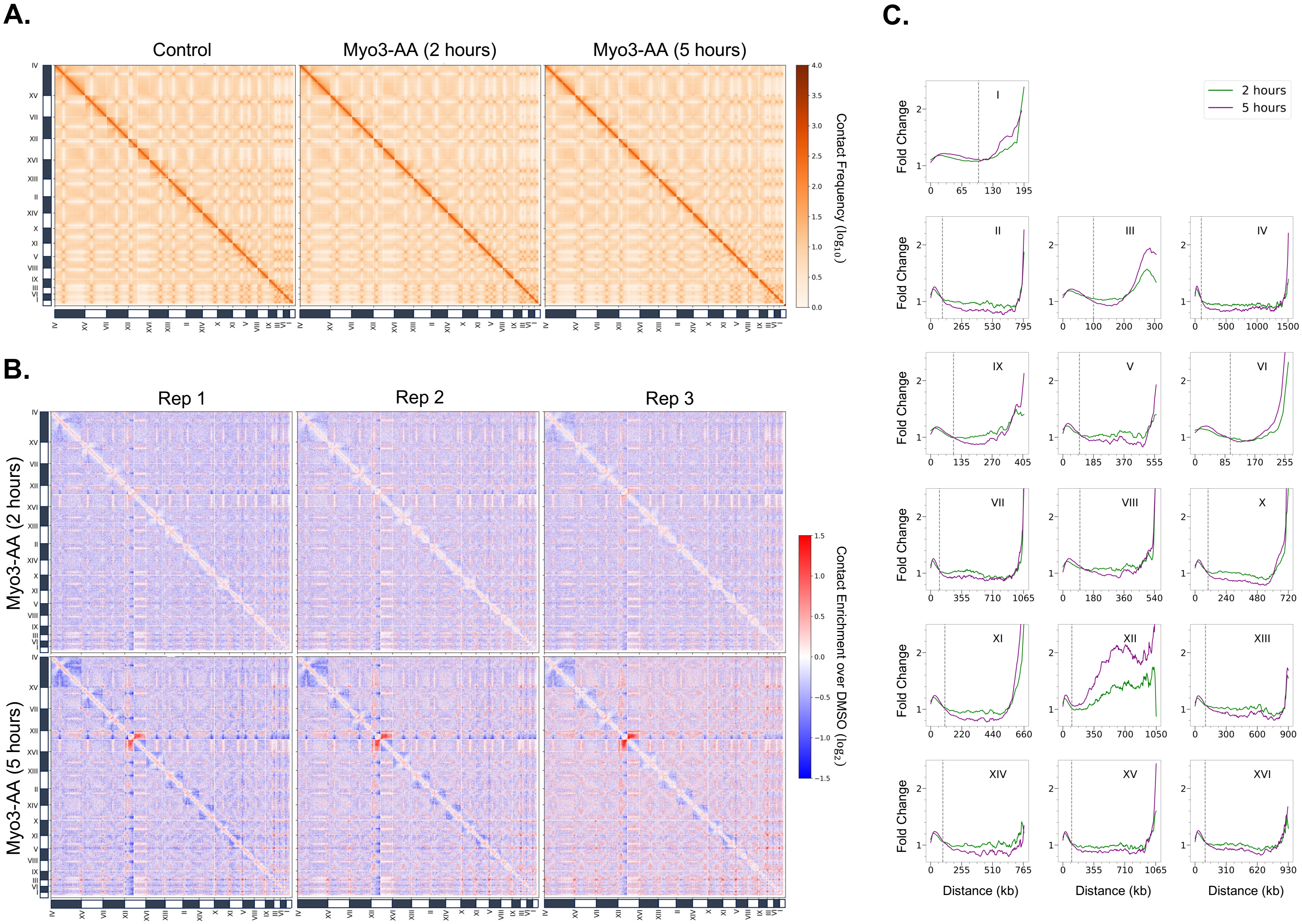
A.
Control
Myo3-AA (2 hours)
Myo3-AA (5 hours)
Contact Frequency ($log_{10}$)
B.
Rep 1
Rep 2
Rep 3
Myo3-AA (2 hours)
Myo3-AA (5 hours)
Contact Enrichment over DMSO ($log_2$)
C.
2 hours
5 hours
Fold Change
Distance (kb)
I
II
III
IV
IX
V
VI
VII
VIII
X
XI
XII
XIII
XIV
XV
XVI

fig. S2 continued

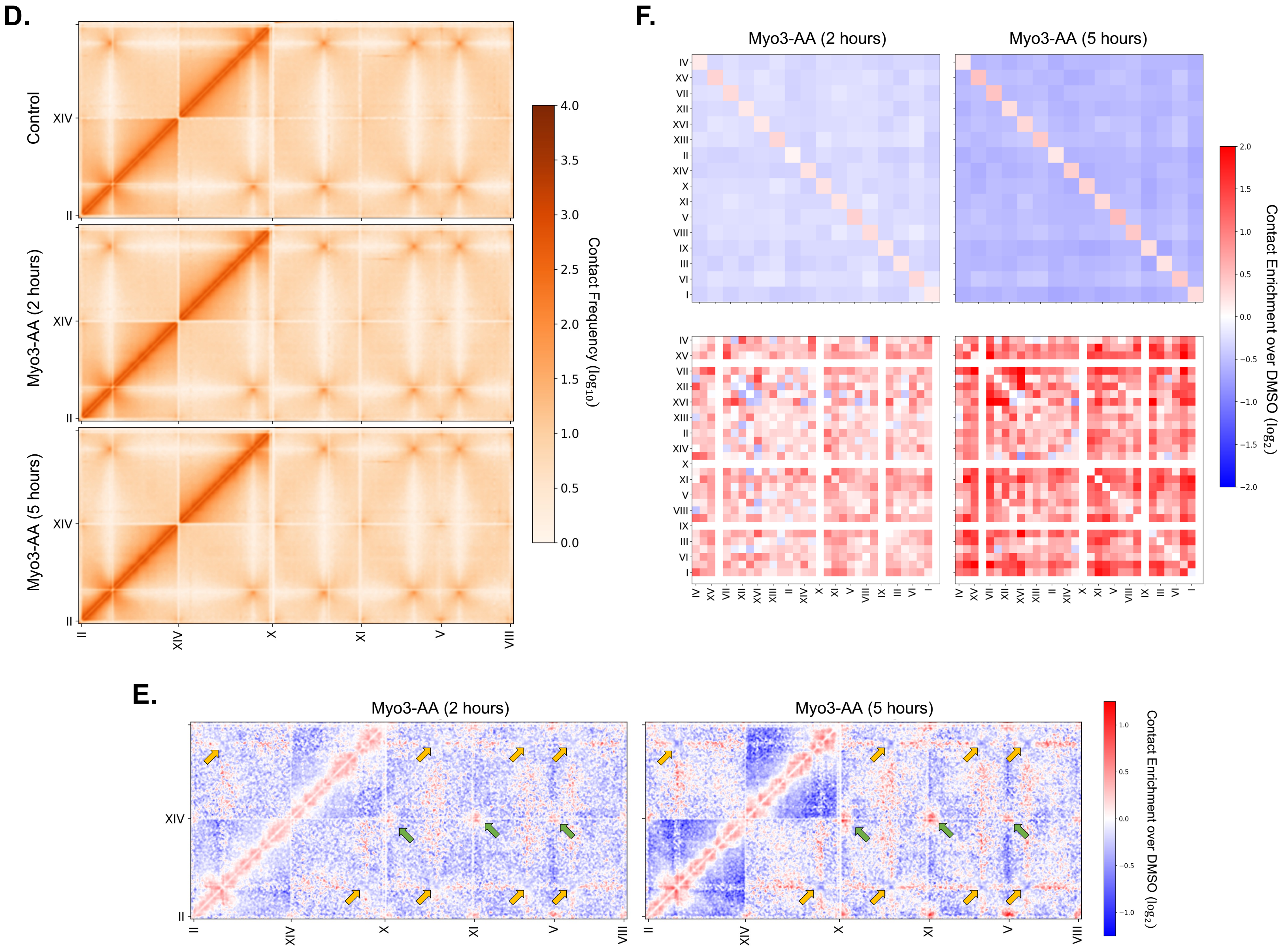

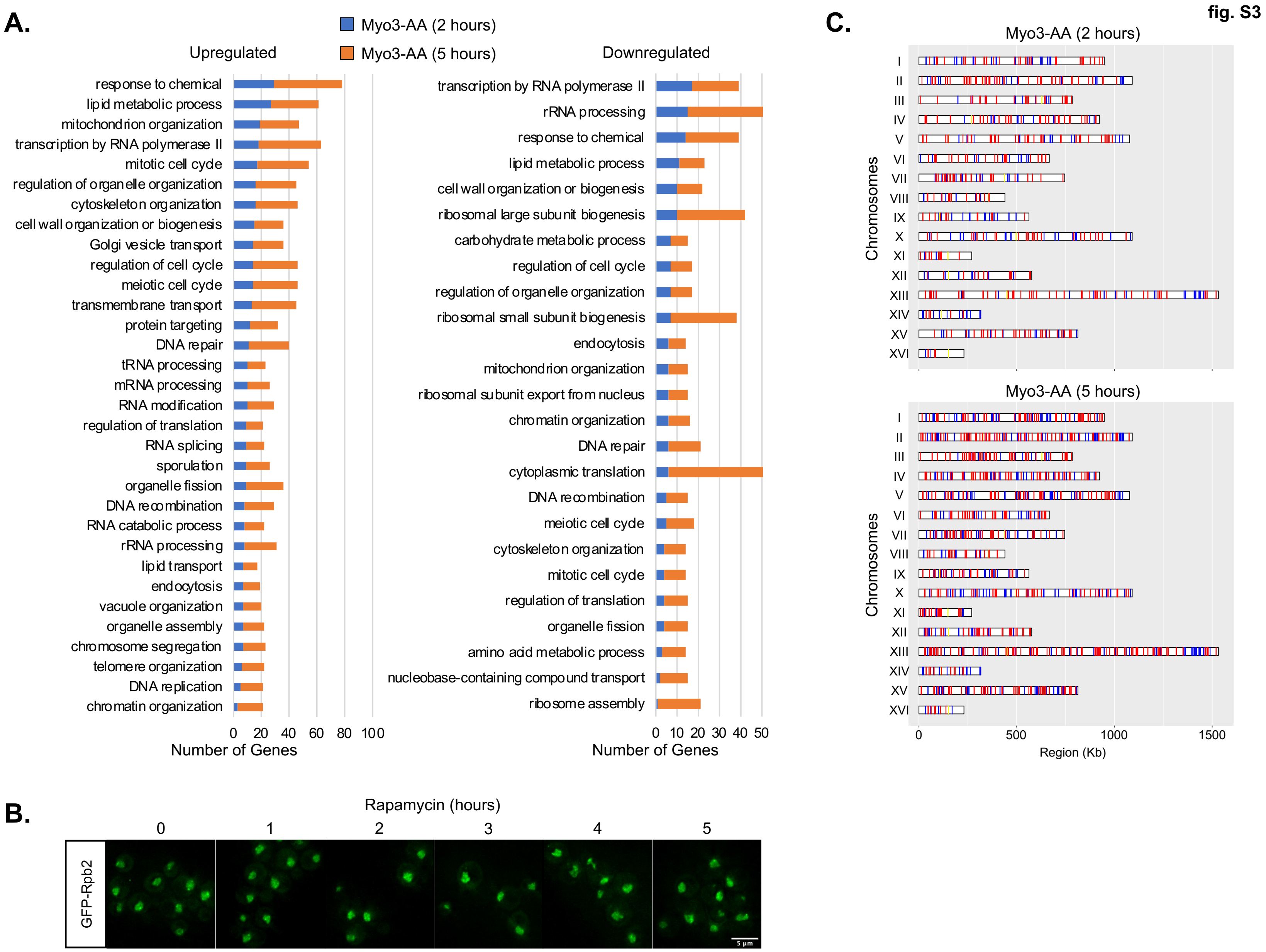

fig. S3
A.
Myo3-AA (2 hours)
Myo3-AA (5 hours)
Upregulated
response to chemical
lipid metabolic process
mitochondrion organization
transcription by RNA polymerase II
mitotic cell cycle
regulation of organelle organization
cytoskeleton organization
cell wall organization or biogenesis
Golgi vesicle transport
regulation of cell cycle
meiotic cell cycle
transmembrane transport
protein targeting
DNA repair
tRNA processing
mRNA processing
RNA modification
regulation of translation
RNA splicing
sporulation
organelle fission
DNA recombination
RNA catabolic process
rRNA processing
lipid transport
endocytosis
vacuole organization
organelle assembly
chromosome segregation
telomere organization
DNA replication
chromatin organization
0 20 40 60 80 100
Number of Genes
Downregulated
transcription by RNA polymerase II
rRNA processing
response to chemical
lipid metabolic process
cell wall organization or biogenesis
ribosomal large subunit biogenesis
carbohydrate metabolic process
regulation of cell cycle
regulation of organelle organization
ribosomal small subunit biogenesis
endocytosis
mitochondrion organization
ribosomal subunit export from nucleus
chromatin organization
DNA repair
cytoplasmic translation
DNA recombination
meiotic cell cycle
cytoskeleton organization
mitotic cell cycle
regulation of translation
organelle fission
amino acid metabolic process
nucleobase-containing compound transport
ribosome assembly
0 10 20 30 40 50
Number of Genes
B.
Rapamycin (hours)
0 1 2 3 4 5
GFP-Rpb2
C.
Myo3-AA (2 hours)
Chromosomes
I II III IV V VI VII VIII IX X XI XII XIII XIV XV XVI
Myo3-AA (5 hours)
Chromosomes
I II III IV V VI VII VIII IX X XI XII XIII XIV XV XVI
0 500 1000 1500
Region (Kb)

fig. S4

A.

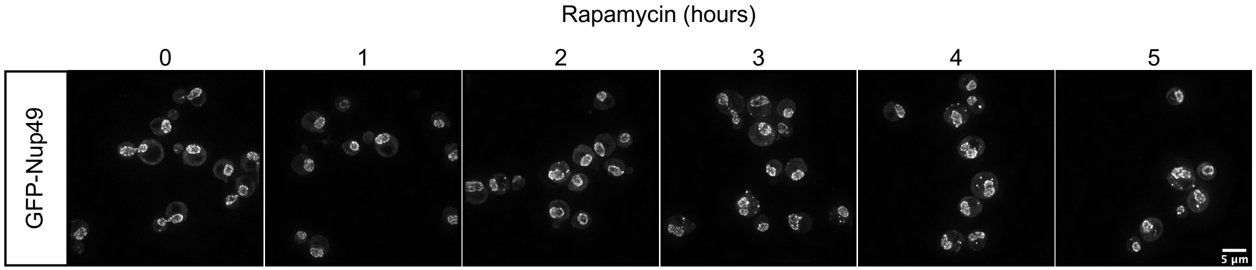
Rapamycin (hours)
0
1
2
3
4
5
GFP-Nup49
5 µm

B.

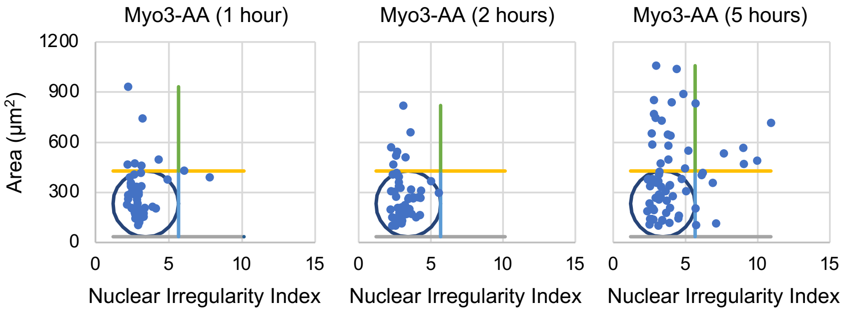
Myo3-AA (1 hour)
Myo3-AA (2 hours)
Myo3-AA (5 hours)
Area (µm²)
1200
900
600
300
0
0
5
10
15
Nuclear Irregularity Index

**fig. S1. Nuclear type I myosin is necessary for cell cycle progression.**

(**A**) The synthetic lethality of the type I myosin genes *MYO3* and *MYO5* was checked using a spot-test, *myo3*Δ*myo5*Δ transformants carrying a *URA3* marked Myo3 expression vector (P406-Myo3), and media with or without 5'-Fluorouracil (5'FOA), as indicated. (**B**) The influence of nuclear type I myosin on cell growth in liquid culture was assessed using parental or Myo3-AA cells in media with or without rapamycin, as marked. (**C**) The relative expression levels of the Myo3 variants was determined by immunoblot analysis using an α-HA antibody. (**D**) Flow cytometry was used to analyze the cell cycle status of Myo3-AA yeast grown in the absence and presence of rapamycin. (**E**) The influence of depleting type I myosin from the nucleus on the cell cycle progression of Myo3-AA cells synchronized in G2/M using Nocodazole or in G1 using a-factor was monitored by FACS after releasing the synchronized cells into media with or without rapamycin, as indicated.

**fig. S2. The influence of type I myosin on 3D chromosome organization is reproducible and genome wide.**

(**A**) The effect of nuclear type I myosin depletion on genome organization was determined using a Hi-C assay. Representative global contact frequency maps (5 kb resolution) of control (DMSO) or rapamycin treated Myo3-AA cells are shown using $\log_{10}$ matrices. The chromosome numbers are marked on the x- and y-axis. (**B**) Three biological replicates of the time-dependent changes in genomic contact frequencies (5 kb resolution) triggered by the nuclear depletion of type I myosin are shown. The $\log_2$ ratio matrices are provided comparing the 2 or 5 h rapamycin treated cells to the DMSO control. The chromosome numbers are marked on the x- and y-axis. (**C**) The increase or decrease in intra-molecular contact frequencies relative to distance across each chromosome, as indicated, were determined. The grey dotted line in each graph marks the 100 kb position along each chromosome. (**D**) The $\log_{10}$ contact frequency maps between chromosomes XIV and II versus II, XIV, X, XI, V, and VIII in presence of rapamycin treated Myo3-AA cells are shown. (**E**) To visualize the relative genome interactions near centromeres and telomeres the changes in contact frequencies are highlighted via a $\log_2$ ratio contact frequency map of Myo3-AA treated with rapamycin for 2- and 5-hours, as marked. The telomere-telomere interactions (green arrows) and interchromosomal centromere-centromere associations (orange arrows) are highlighted. (**F**) The contact frequencies between telomeres or centromeres were assessed using $\log_2$ ratio matrices plotting the contact frequency changes between all centromeres (upper 2 panels) or telomeres (lower 2 panels) at 2 or 5 h of rapamycin treatment. The chromosome numbers are marked on the x- and y-axis.

**fig. S3. Nuclear myosin I dependent genes are associated with diverse cellular processes and the genes are physically dispersed across the genome.**

(**A**) Genes displaying differential expression (increased ≥2-fold or ≤50% relative to control) after 2 (blue bars) or 5 (orange bars) hours of rapamycin treatment in Myo3-AA cells were partitioned into a variety of cellular processes using a GO Slim analysis. The downregulated and upregulated hits in each category were compiled and graphed, as marked. (**B**) To determine the influence of nuclear type I myosin on the nuclear distribution of RNA polymerase II, Myo3-AA cells expressing GFP-Rpb2 were treated with DMSO or rapamycin for the indicated times. (**C**) The physical distribution of the differentially expressed genes in Myo3-AA yeast was visualized using chromosome plots (chromosomes 1-16). The red lines mark the position of activated genes, blue lines are repressed genes, and yellow lines indicate the centromeres.

**fig. S4. Nuclear envelope morphology is dependent on the presence of type I myosin in the nucleus.**

(**A**) The shape of the nuclear envelope in Myo3-AA cells was monitored using a GFP fusion of the nuclear pore complex subunit Nup49 (GFP-Nup49). Myo3-AA cells were treated with DMSO (control) or rapamycin for the indicated time. The GFP-Nup49 signal was followed by fluorescence microscopy. (**B**) The presence of irregular shaped nuclei was determined using Myo3-AA yeast expressing 3xNLS-GFP and the Fiji plugin NII. The scatterplots for 1, 2, and 5 h rapamycin treated cells are shown.

### Materials and Methods

#### Cells, growth media, and conditions

*Saccharomyces cerevisiae* strains used in this study are described in Table S3. Yeast cultures were grown in complete (YPD) or minimal media supplemented with the necessary amino acids at 30°C unless otherwise noted. Engineered strains were created via standard procedure (*36*). Previously described amino-terminal GFP fusion strains (*37*) were used in the construction of anchor-away variants. For the anchor-away experiments, DMSO or Rapamycin (1 µg/ml final) was added for the indicated times. For cell cycle synchronization experiments, ethanol or a-Factor mating pheromone (5 µg/ml final) was added to the media for 2 hours to synchronize in G1, DMSO or Nocodazole (10 µg/ml final) was added to the media for 2 hours to synchronize in G2/M, and cells were then released into media with or without rapamycin. For experiments using 5' Fluorouracil, a final concentration of 0.1% was used.

#### Cell viability spot-test

A standard yeast spot-test assay was used to test cell viability by growing the cells in the appropriate media to mid-log phase, collecting by centrifugation, and resuspending in water to $O.D._{595}$ = 1. Cells were serially diluted in water ($10^0$ to $10^{-4}$) and spotted on to the described media.

#### Flow cytometry

Fluorescence-activated cell sorting (FACS) was accomplished as described (*38*). In brief, mid-log phase cells were fixed with methanol at -20°C overnight. DNA staining was accomplished with SYTOX Green (1 µM). Prior to sorting, stained cells were sonicated (20% output for 10 s). A BD LSR II Flow Cytometer (BD Biosciences) was used for sorting with a 488 nm excitation laser and analyzed using FlowJo 10.8.2.

High-throughput Chromosome Conformation Capture (Hi-C) and 3C assays

Hi-C DNA samples were generated by adapting a protocol from Dr. Romain Koszul (Institut Pasteur). Briefly, yeast cultures (150 ml) were grown to mid-log phase, an aliquot was used for Hi-C (125 ml) and the rest (25 ml) for RNA-seq. The Hi-C sample was fixed with 3% formaldehyde, quenched with glycine, and a cell pellet (125 mg) was treated with Zymolyase-100T (10 µg/ml) until 50-60% spheroplast formation was achieved. In situ genomic DNA digestion was done with DpnII (450 U New England Biolabs) and the overhangs were partially filled with Klenow and biotin-labelled dCTP (Invitrogen). Samples were then ligated using T4 DNA Ligase (480 U Thermo Fisher) for 3 hours at room temperature. DNA was isolated following Proteinase K treatment, phenol/chloroform extraction, and RNase treatment.

The Hi-C libraries were prepared using a modified version of an established protocol (*39*). In brief, the DNA samples were end-repaired and A-tailed following removal of the biotin at the non-ligated ends. The samples fractionated and enriched for biotinylated ligation products using Streptavidin C1 beads. Adapters (xGen Stubby; IDT) were ligated on using T4 Quick Ligase (NEB) and DNA was amplified with xGen UDI Primers via on-bead PCR (8 cycles total). The library was purified using AMPure XP beads (Beckman Coulter). Samples were paired-end sequenced using a NovaSeq 6000 SP (*40*).

Hi-C data analysis

*Alignment and matrix generation:* Hi-C contact frequency matrices at 5 kb resolution were generated using the Hi-C-pro pipeline (*41*) and SacCer3 Bowtie index. To normalize the depth of the generated AllValidPairs files, the read counts were limited by the file with the lowest read number using the Linux command -shuf before matrix building and ICE normalization. The files generated were then converted to hdf5 format using the HiCExplorer tool (*42*).

*Contact frequency matrix analysis:* The generated contact frequency matrices were imported into Python 3.9.13 as per the HiCExplorer 3.7.2 manual. Heatmap averages from biological triplicates were created, contact frequency versus genomic distance plots were generated, and Savitzky-Golay filter was used to smoothen the curve. For the analysis around centromeres, the centromere locations were based on SacCer3 and the average of 30 kb bins centered around each centromere were used. The telomere genomic interaction analysis used the average of 30 kb region at each chromosome end.

*Transcription hub/factory analysis:* Permutations tests were performed to assess whether gene promoters predicted to be bound by the same transcription factor interacted relative to random genes following activation or repression. The YEASTRACT+ database was used to predict gene promoters bound by the same transcription factors (*28*) and our RNA-Seq analysis (please see below) was used to identify activated and repressed genes. The genomic coordinates of the target genes were obtained with BiomaRt (*43*) and the corresponding 5 kb bins for each gene in the Hi-C matrices were used to compare changes in contact frequencies between the TFcategorized genes versus random sets of genes. For each test, 100,000 permutations were used to generate the null distribution, Z-scores, and p-values (Fig. 4B and Table S2). In addition to the permutations tests, the potential formation of transcription hubs/factories around activated genes or dispersal of hubs/factories at repressed genes was evaluated using gene expression Volcano plots generated with our RNA-Seq data (please see below). Here, the Volcano plots were color-coded to reflect the changes in contact frequency ($log_2$) between each target gene and any DNA locus at the set distances of 25, 100, or 500 kb using a 5 kb bin in the Hi-C matrices.

Chromosome Conformation Capture (3C) assay

The 3C experiments followed a modified, downscaled version of the described Hi-C protocol. In brief, 25 ml cultures were used. The ligation reactions used 60 U of T4 DNA Ligase (Thermo Fisher). After overnight Proteinase K digestion, phenol/chloroform extraction, and RNase treatment, the DNA was purified using a Monarch DNA cleanup column (NEB). The ARS504 locus was used as a control site for all 3C-qPCR reactions (*44*).

RT-qPCR and Northern blots

Total RNA was extracted from yeast cells using a standard hot phenol extraction method (*45*). RT-qPCR reactions were performed with the Power SYBR Green RNA-to-$C_T$ 1-Step Kit according to manufactures instructions (Applied Biosystems) using the listed oligonucleotide primers (Table S4).

Northern Blots were carried out using the NorthernMax-Gly Kit (Invitrogen), as per the manufacturer's protocol and the ITS1 and ITS2 probes were used to detect pre-rRNA species (*46*).

RNA-Seq

Libraries were constructed from total RNA using the Kapa Hyper Stranded mRNA library kit (Roche) and sequenced on the NovaSeq 6000 SP (*40*). Reads were aligned to the *Saccharomyces cerevisiae* (S288C) reference genome with STAR (*47*) and gene counts were generated using HTSeq count using the union mode (*48*). Differential gene expression analysis was done using RStudio 2022.12.0.353 using Deseq2 (*49*). Volcano plots were generated using the EnhancedVolcano package (*50*). Gene ontology analysis of differentially expressed genes was performed using the Saccharomyces Genome Database (SGD) Gene Ontology Slim Term Mapper. For gene position analysis (fig. S5A), chromosome plots were generated using a modified code from Biostars (https://www.biostars.org/p/269857/).

Fluorescence microscopy

*Live cell imaging:* Mid-log phase yeast cells were collected and washed once with complete synthetic media and resuspended in complete synthetic media. Images were acquired using a DeltaVision OMX microscope (GE Life Science) with 100x/1.40 NA oil immersion objective or Zeiss AxioObserver Z1 with 63x/1.4 NA.

*Fixed cell imaging:* Immunofluorescence staining was carried using a modified standard protocol (*51*). In brief, the secondary fixation was carried out for 2 h followed by spheroplasting with Zymolyase-20T (Nacalai Tesque) and cells were adhered to Poly-L-lysine coated 18x18mm coverslips. Mouse anti-Nop1p (Santa Cruz; sc-57940) and Goat anti-mouse IgG-CFL 594 (Santa Cruz; sc-362277) were used for immunostaining and Hoechst 3342 (BD Pharmingen) was used to visualize the DNA. Coverslips were mounted with SlowFade Diamond (Invitrogen). Airyscan microscopy was performed using a Zeiss LSM900 confocal laser scanning microscope equipped with an Airyscan detection unit. Imaging was done with a 63x/1.46 oil immersion objective using Immersol 518-F immersion media (ne = 1.518); Zeiss). All images were processed using Airyscan Joint Deconvolution with 12 iterations.

*Image analysis:* Images were processed in Fiji. For quantification of percentage of irregularly shaped nuclei and nucleoli, the multi-point tool in Fiji was employed. Nuclear irregularity index was measured using the NII plugin (*52*). To quantify Net1 density, a 2 µm diameter concentric circle was drawn around the Net1 cloud and the integrated density was determined.

**Table S1. Permutation test results assessing the contact frequency changes between associated transcription factors and activated genes.**

| Timepoint | TF | No. of genes | P-value | Z-score |
|---|---|---|---|---|
| 2 hours | Hap1 | 39 | 0.15 | 1.05 |
| | Spt23 | 98 | 0.4 | -0.24 |
| | Tos4 | 14 | 0.15 | 1.06 |
| 5 hours | Hap1 | 62 | 0.16 | 1.02 |
| | Spt23 | 146 | 0.17 | -0.91 |
| | Tos4 | 24 | 0.5 | 0.09 |
| | Ndd1 | 15 | 0.39 | 0.28 |

**Table S2. Permutation test results assessing the contact frequency changes between associated transcription factors and repressed genes.**

| Timepoint | TF | No. of genes | P-value | Z-score |
|---|---|---|---|---|
| 2 hours | Fkh1 | 69 | 0.42 | 0.27 |
| | Spt23 | 56 | 0.09 | -1.28 |
| | Ifh1 | 26 | 0.34 | -0.38 |
| 5 hours | Fkh1 | 183 | 0.29 | -0.53 |
| | Spt23 | 140 | 0.41 | 0.22 |
| | Ifh1 | 97 | 0.41 | -0.16 |
| | Hpc2 | 18 | 0.4 | 0.3 |

**Table S3. Yeast strains used in this study**

| Yeast strain | Genotype | Source |
|---|---|---|
| *myo3*Δ *myo5*Δ 3xHA-Myo3 | YPH499 *MATa myo5*Δ*::KANMX myo3*Δ*::LEU2MX* pRS406-ADH1-3xHA-Myo3 | This study |
| VDY1874 | W303 *MATα TPK1/2/3/as TOR1-1(S1972I)fpr1Δ::NATMX RPL13A-2xFKBP12–TRP1* | Solis et al., 2016 |
| Myo1-FRB | W303 *MATα TPK1/2/3/as TOR1-1(S1972I)fpr1Δ::NATMX RPL13A-2xFKBP12–TRP1 MYO1-FRB-HIS3MX* | This study |
| Myo2-FRB | W303 *MATα TPK1/2/3/as TOR1-1(S1972I)fpr1Δ::NATMX RPL13A-2xFKBP12–TRP1 MYO2-FRB-HIS3MX* | This study |
| Myo3-FRB | W303 *MATα TPK1/2/3/as TOR1-1(S1972I)fpr1Δ::NATMX RPL13A-2xFKBP12–TRP1 MYO3-FRB-HIS3MX* | This study |
| Myo4-FRB | W303 *MATα TPK1/2/3/as TOR1-1(S1972I)fpr1Δ::NATMX RPL13A-2xFKBP12–TRP1 MYO4-FRB-HIS3MX* | This study |
| Myo5-FRB | W303 *MATα TPK1/2/3/as TOR1-1(S1972I)fpr1Δ::NATMX RPL13A-2xFKBP12–TRP1 MYO5-FRB-HIS3MX* | This study |
| myo3Δ | W303 *MATα TPK1/2/3/as TOR1-1(S1972I)fpr1Δ::NATMX RPL13A-2xFKBP12–TRP1 myo3Δ::LEU2MX* | This study |
| Myo5Δ | W303 *MATα TPK1/2/3/as TOR1-1(S1972I)fpr1Δ::NATMX RPL13A-2xFKBP12–TRP1 myo5Δ::LEU2MX* | This study |
| myo3Δ Myo5-FRB | W303 *MATα TPK1/2/3/as TOR1-1(S1972I)fpr1Δ::NATMX RPL13A-2xFKBP12–TRP1 myo3Δ::LEU2MX MYO5-FRB-HIS3MX* | This study |
| myo5Δ Myo3-FRB 1 | W303 *MATα TPK1/2/3/as TOR1-1(S1972I)fpr1Δ::NATMX RPL13A-2xFKBP12–TRP1 myo5Δ::LEU2MX MYO3-FRB-HIS3MX* | This study |

| | | |
|---|---|---|
| myo5Δ Myo3-FRB 2 | W303 *MATα TPK1/2/3/as TOR1-1(S1972I)fpr1Δ::NATMX RPL13A-2xFKBP12–TRP1 myo5Δ::KANMX MYO3-FRB-HIS3MX* | This study |
| MYO3 FL | W303 *MATα TPK1/2/3/as TOR1-1(S1972I)fpr1Δ::NATMX RPL13A-2xFKBP12–TRP1 myo5Δ::KANMX MYO3-FRB-HIS3MX* pRS405-3xHA-MYO3 | This study, Wang et al., 2020 |
| MYO3-1 | W303 *MATα TPK1/2/3/as TOR1-1(S1972I)fpr1Δ::NATMX RPL13A-2xFKBP12–TRP1 myo5Δ::KANMX MYO3-FRB-HIS3MX* pRS405-3xHA-MYO3-1 | This study Wang et al., 2020 |
| MYO3-2 | W303 *MATα TPK1/2/3/as TOR1-1(S1972I)fpr1Δ::NATMX RPL13A-2xFKBP12–TRP1 myo5Δ::KANMX MYO3-FRB-HIS3MX* pRS405-3xHA-MYO3-2 | This study Wang et al., 2020 |
| MYO3-3 | W303 *MATα TPK1/2/3/as TOR1-1(S1972I)fpr1Δ::NATMX RPL13A-2xFKBP12–TRP1 myo5Δ::KANMX MYO3-FRB-HIS3MX* pRS405-3xHA-MYO3-3 | This study Wang et al., 2020 |
| MYO3-1+2 | W303 *MATα TPK1/2/3/as TOR1-1(S1972I)fpr1Δ::NATMX RPL13A-2xFKBP12–TRP1 myo5Δ::KANMX MYO3-FRB-HIS3MX* pRS405-3xHA-MYO3-1+2 | This study Wang et al., 2020 |
| MYO3-1+3 | W303 *MATα TPK1/2/3/as TOR1-1(S1972I)fpr1Δ::NATMX RPL13A-2xFKBP12–TRP1 myo5Δ::KANMX MYO3-FRB-HIS3MX* pRS405-3xHA-MYO3-1+3 | This study Wang et al., 2020 |
| MYO3-2+3 | W303 *MATα TPK1/2/3/as TOR1-1(S1972I)fpr1Δ::NATMX RPL13A-2xFKBP12–TRP1 myo5Δ::KANMX MYO3-FRB-HIS3MX* pRS405-3xHA-MYO3-2+3 | This study Wang et al., 2020 |
| GFP-NET1 | W303 *MATα TPK1/2/3/as TOR1-1(S1972I)fpr1Δ::NATMX RPL13A-2xFKBP12–TRP1 myo5Δ::KANMX MYO3-FRB-HIS3MX* URA-NOP1pr-GFP-NET1 | This study |

| | | |
|---|---|---|
| GFP-RPB2 | W303 *MATα TPK1/2/3/as TOR1-1(S1972I)fpr1Δ::NATMX RPL13A-2xFKBP12–TRP1 myo5Δ::KANMX MYO3-FRB-HIS3MX* URA-NOP1pr-GFP-RPB2 | This study |
| GFP-RPB2 | W303 *MATα TPK1/2/3/as TOR1-1(S1972I)fpr1Δ::NATMX RPL13A-2xFKBP12–TRP1 myo5Δ::KANMX MYO3-FRB-HIS3MX* URA-NOP1pr-GFP-RPB2 | This study |
| GFP-PHO88 | W303 *MATα TPK1/2/3/as TOR1-1(S1972I)fpr1Δ::NATMX RPL13A-2xFKBP12–TRP1 myo5Δ::LEU2MX MYO3-FRB-HIS3MX* URA-NOP1pr-GFP-PHO88 | This study |
| GFP-NUP49 | W303 *MATα TPK1/2/3/as TOR1-1(S1972I)fpr1Δ::NATMX RPL13A-2xFKBP12–TRP1 myo5Δ::LEU2MX MYO3-FRB-HIS3MX* URA-NOP1pr-GFP-NUP49 | This study |

**Table S4. Primers used in this study**

| Primer name | Sequence | Purpose |
|---|---|---|
| MYO1 C' tag pFA6 F | AAATATTGATAGTAACAATGCACAGAGTAAAATTT TCAGTcggatccccgggttaattaa | Strain construction |
| MYO1 C' tag pFA6 R | GGATATAAAGTCTTCCAAATTTTTAAAAAAAAGTTC GTTAgaattcgagctcgtttaaac | Strain construction |
| MYO2 C' tag pFA6 F | AGTTGACCTTGTTGCCCAACAAGTCGTTCAAGAC GGCCACcggatccccgggttaattaa | Strain construction |
| MYO2 C' tag pFA6 R | GCATTCATGTACAATTTTGTTTCTCGCGCCATCAG TTTTAgaattcgagctcgtttaaac | Strain construction |
| MYO3 C' tag pFA6 F | CGATGATGATAATGACGATGGCGATGATGATGAT GACTGGcggatccccgggttaattaa | Strain construction |
| MYO3 C' tag pFA6 R | TTCTATAAGAGACTAAACATCTATATATTCATGTTA ATTAgaattcgagctcgtttaaac | Strain construction |
| MYO4 C' tag pFA6 F | CTTAGCTACTGTCAGTAAAATTATAAAATTAGACA GAAAAcggatccccgggttaattaa | Strain construction |
| MYO4 C' tag pFA6 R | ACATATATACATATATGGGCGTATATTTACTTTGTT CTTAgaattcgagctcgtttaaac | Strain construction |
| MYO5 C' tag pFA6 F | TGACGAGGAGGCTAACGAAGATGAAGAGGAAGA TGATTGGcggatccccgggttaattaa | Strain construction |
| MYO5 C' tag pFA6 R | GCTCGTATAGAGTATATACTCGCTAAATACATTTT GATTAgaattcgagctcgtttaaac | Strain construction |
| MYO3 KO pFA6 F | AATCAAAAAAAGTAAAGAGAAATACTTATAACCGT CAATGcggatccccgggttaattaa | Strain construction |
| MYO3 KO pFA6 R | TTCTATAAGAGACTAAACATCTATATATTCATGTTA ATTAgaattcgagctcgtttaaac | Strain construction |
| MYO5 KO pFA6 F | GAACAGTGGCCAATACGAATTTAACCGCTTTATAG AAATGcggatccccgggttaattaa | Strain construction |
| MYO5 KO pFA6 R | GCTCGTATAGAGTATATACTCGCTAAATACATTTT GATTAgaattcgagctcgtttaaac | Strain construction |

| | | |
|---|---|---|
| NET1_GFPN_F | CTAGTGAAGAGTAGCTAGCACG | Strain construction |
| NET1_GFPN_R | GCAACAATTGAGAAGATTCGGC | Strain construction |
| RPB2_GFPN_F | CCTCAGGTTTACAGATAGTTCG | Strain construction |
| RPB2_GFPN_R | GGATATAACAGCCCACGAATC | Strain construction |
| PHO88_GFPN_F | TCTTCGACGTTTGCCACTGC | Strain construction |
| PHO88_GFPN_R | GTTCGACGTACTTCATGGTAGTC | Strain construction |
| GFPN_NUP49_F | GTTACTTGCTCTGCGGTGGC | Strain construction |
| GFPN_NUP49_R | ATCCTCCTGTAGATCCGGCTG | Strain construction |
| ITS1 | CGGTTTTAATTGTCCTA | Northern Blot |
| ITS2 | TGAGAAGGAAATGACGCT | Northern Blot |
| ATG8_F | AGTTCCTGCTGACCTTACCG | RT-qPCR |
| ATG8_R | ACCCGTCCTTATCCTTGTGT | RT-qPCR |
| HHF2_F | GCCAAGCGTCACAGAAAGAT | RT-qPCR |
| HHF2_R | GTCCCTGATGACGGATTCCA | RT-qPCR |
| ACT1_F | AACTGGGACGATATGGAAAA | RT-qPCR |
| ACT1_R | GAAGGCTGGAACGTTGAAAG | RT-qPCR |
| 1_RDN_ACS2 | CTTCCAGTAGCGATAACACAACTAG | 3C-qPCR |
| 1_RDN_PCD1 | GTTGGACTACATGACTATGGTAATC | 3C-qPCR |
| 9_RDN_MAS1 | CTAGAACACTTGGCTTTCAAGGGTA | 3C-qPCR |

| | | |
|---|---|---|
| 10_RDN_PUS5 | GATAGCGTACATAGAGTATTCACG | 3C-qPCR |
| TEL4R_PAU10 | CACTTGGCCCAATACTACATGTTC | 3C-qPCR |
| TEL9R_YIR043C | GTCTGGCCTACGAGTATCTTACATG | 3C-qPCR |
| TEL5R_YER188W | TACGCCAAGTCGGTCAGGTCTTAT | 3C-qPCR |
| ARS504_F | GTCAGACCTGTTCCTTTAAGAGG | 3C-qPCR |
| ARS504_R | CATACCCTCGGGTCAAACAC | 3C-qPCR |

Table S5. Differential gene expression analysis for Myo3-AA treated with rapamycin for 2 hours.

| Row.names | baseMean | log2FoldCha | lfcSE | stat | pvalue | padj | genename | chrom | start_pos | end_pos | strand |
|---|---|---|---|---|---|---|---|---|---|---|---|
| Q0158 | 6.52489728 | 1.30044067 | 1.17911538 | 2.46242714 | 0.29193808 | 0.46616698 | NA | Mito | 58009 | 62447 | 1 |
| YAL001C | 152.194332 | 2.5740339 | 0.76762264 | 11.4156737 | 0.00331985 | 0.01648534 | TFC3 | I | 147594 | 151166 | -1 |
| YAL002W | 78.7260454 | 0.11815865 | 0.69744289 | 0.72275892 | 0.69671457 | 0.81659461 | VPS8 | I | 143707 | 147531 | 1 |
| YAL003W | 117.199149 | -0.4918673 | 0.59563592 | 2.55984277 | 0.27805916 | 0.45261809 | EFB1 | I | 142174 | 143160 | 1 |
| YAL005C | 85.6274446 | 0.15958309 | 0.6531727 | 6.74773434 | 0.0342569 | 0.09906519 | SSA1 | I | 139503 | 141431 | -1 |
| YAL007C | 171.012771 | 0.39141258 | 0.50467907 | 20.9886448 | 2.77E-05 | 0.00031564 | ERP2 | I | 137698 | 138345 | -1 |
| YAL008W | 16.4436684 | 1.11627722 | 0.83648467 | 2.61332338 | 0.2707223 | 0.44417176 | FUN14 | I | 136914 | 137510 | 1 |
| YAL009W | 16.1660335 | -0.1488104 | 0.78011115 | 0.07451714 | 0.96342699 | 0.9730058 | SPO7 | I | 135854 | 136633 | 1 |
| YAL010C | 74.0582927 | -0.9091388 | 0.55566311 | 6.09173003 | 0.04755516 | 0.12572802 | MDM10 | I | 134184 | 135665 | -1 |
| YAL011W | 219.581007 | -0.425848 | 0.52146778 | 6.27233632 | 0.04344897 | 0.11810776 | SWC3 | I | 132199 | 134076 | 1 |
| YAL012W | 197.449702 | 0.27059142 | 0.49395316 | 4.18403485 | 0.12343786 | 0.25497866 | CYS3 | I | 130799 | 131983 | 1 |
| YAL013W | 173.375851 | 0.85553643 | 0.68760381 | 2.92971046 | 0.23111145 | 0.39792139 | DEP1 | I | 129270 | 130487 | 1 |
| YAL014C | 26.3703385 | 0.6835588 | 0.73213803 | 2.05406147 | 0.35806858 | 0.53120548 | SYN8 | I | 128252 | 129019 | -1 |
| YAL015C | 12.5995056 | 0.75894368 | 0.97317353 | 0.71046144 | 0.70101169 | 0.81801018 | NTG1 | I | 126903 | 128102 | -1 |
| YAL016W | 84.6687343 | 0.16012462 | 0.57865731 | 0.13350263 | 0.9354278 | 0.95803272 | TPD3 | I | 124879 | 126786 | 1 |
| YAL017W | 41.247613 | -0.2060059 | 0.67778 | 0.10729795 | 0.94776474 | 0.96435226 | PSK1 | I | 120225 | 124295 | 1 |
| YAL018C | 8.76638298 | -0.8616951 | 1.00597601 | 1.06893526 | 0.58598117 | 0.73378964 | LDS1 | I | 118564 | 119541 | -1 |
| YAL019W | 33.9776591 | 0.03923142 | 0.68818101 | 6.17671038 | 0.04557686 | 0.12191042 | FUN30 | I | 114919 | 118314 | 1 |
| YAL020C | 39.0307728 | 0.62927142 | 0.6708548 | 1.00110037 | 0.60619705 | 0.75088539 | ATS1 | I | 113614 | 114615 | -1 |
| YAL021C | 79.1822695 | 0.06459553 | 0.62030721 | 0.43478161 | 0.80461546 | 0.88279205 | CCR4 | I | 110846 | 113359 | -1 |
| YAL022C | 119.31372 | 0.09700411 | 0.62534661 | 14.6928354 | 0.0006449 | 0.00457593 | FUN26 | I | 108877 | 110430 | -1 |
| YAL023C | 85.7619927 | 0.94482946 | 0.58897977 | 3.64225043 | 0.16184354 | 0.31028944 | PMT2 | I | 106272 | 108551 | -1 |
| YAL024C | 13.6197097 | -0.8440935 | 1.05193959 | 4.03902184 | 0.13272036 | 0.26859521 | LTE1 | I | 101565 | 105872 | -1 |
| YAL025C | 102.736369 | -0.9255784 | 0.57872793 | 30.7178657 | 2.14E-07 | 4.63E-06 | MAK16 | I | 100225 | 101145 | -1 |
| YAL026C | 26.8507426 | 0.08625098 | 0.81847898 | 0.04998457 | 0.97531744 | 0.9809081 | DRS2 | I | 95630 | 99697 | -1 |
| YAL027W | 15.4075861 | 0.62661607 | 0.80459813 | 2.31500854 | 0.31426953 | 0.48639952 | SAW1 | I | 94687 | 95472 | 1 |
| YAL028W | 22.7228518 | -1.8803865 | 0.76654473 | 9.11580546 | 0.01048402 | 0.04090465 | FRT2 | I | 92900 | 94486 | 1 |
| YAL029C | 264.124837 | 2.39011139 | 0.89909486 | 7.8771885 | 0.01947557 | 0.06484392 | MYO4 | I | 87855 | 92270 | -1 |
| YAL030W | 174.597782 | 0.75075434 | 0.55758348 | 8.22967359 | 0.01632861 | 0.05693318 | SNC1 | I | 87286 | 87752 | 1 |
| YAL031C | 98.4896929 | -0.0421486 | 0.56764308 | 1.94896973 | 0.37738671 | 0.55108929 | GIP4 | I | 84749 | 87031 | -1 |
| YAL032C | 281.557756 | 2.29248678 | 0.5516531 | 48.3848365 | 3.11E-11 | 1.58E-09 | PRP45 | I | 83335 | 84474 | -1 |

| | | | | | | | | | | | |
|---|---|---|---|---|---|---|---|---|---|---|---|
| YAL033W | 52.7827913 | 1.02476483 | 0.66213079 | 2.54871713 | 0.27961026 | 0.45373975 | POP5 | I | 82706 | 83227 | 1 |
| YAL034C | 86.4634567 | 3.46830899 | 0.61258192 | 29.7533699 | 3.46E-07 | 6.96E-06 | FUN19 | I | 80710 | 81951 | -1 |
| YAL034W-A | 366.639954 | 3.43093035 | 0.63027522 | 32.5295341 | 8.64E-08 | 2.15E-06 | MTW1 | I | 79718 | 80587 | 1 |
| YAL035W | 690.188751 | -0.0568433 | 0.5181568 | 11.3865416 | 0.00336856 | 0.0166843 | FUN12 | I | 76427 | 79435 | 1 |
| YAL036C | 24.7767781 | -0.7915201 | 0.90483477 | 2.14681962 | 0.34184092 | 0.51556627 | RBG1 | I | 75043 | 76152 | -1 |
| YAL037C-A | 24876.3172 | 3.75737292 | 2.03422839 | 4.16720322 | 0.12448107 | 0.25658434 | NA | I | 73426 | 73518 | -1 |
| YAL037W | 182.809816 | -0.5329746 | 0.55425669 | 2.34600947 | 0.30943577 | 0.48239987 | NA | I | 74020 | 74823 | 1 |
| YAL038W | 1192.55953 | 0.55965223 | 0.47464606 | 8.09941568 | 0.01742747 | 0.05974247 | CDC19 | I | 71786 | 73288 | 1 |
| YAL039C | 185.960627 | -0.8842225 | 0.59578552 | 4.76423941 | 0.09235461 | 0.20685781 | CYC3 | I | 68716 | 69525 | -1 |
| YAL040C | 44.749424 | 0.0384695 | 0.61746271 | 4.66264229 | 0.09716729 | 0.21489634 | CLN3 | I | 65778 | 67520 | -1 |
| YAL041W | 143.741223 | 1.74857694 | 0.81593959 | 6.35754028 | 0.04163683 | 0.11447163 | CDC24 | I | 62840 | 65404 | 1 |
| YAL042W | 218.964554 | 0.43201691 | 0.59939448 | 25.776984 | 2.53E-06 | 3.94E-05 | ERV46 | I | 61316 | 62563 | 1 |
| YAL043C | 41.7538945 | 0.43987698 | 0.65588919 | 0.85382898 | 0.65251935 | 0.78302322 | PTA1 | I | 58695 | 61052 | -1 |
| YAL044C | 295.413083 | 2.61504165 | 0.88806424 | 9.89778291 | 0.00709127 | 0.0303238 | GCV3 | I | 57950 | 58462 | -1 |
| YAL044W-A | 15.1812068 | 0.50599437 | 0.89024041 | 2.70746223 | 0.25827481 | 0.43008401 | BOL1 | I | 57518 | 57850 | 1 |
| YAL046C | 43.4614896 | -1.5418298 | 0.67189732 | 16.7369544 | 0.00023207 | 0.00193893 | BOL3 | I | 57029 | 57385 | -1 |
| YAL047C | 19.8213171 | 0.51167546 | 0.76605444 | 0.52608763 | 0.76870821 | 0.85939768 | SPC72 | I | 54989 | 56857 | -1 |
| YAL048C | 60.3846288 | 0.25736752 | 0.55900816 | 0.56227894 | 0.75492304 | 0.85188158 | GEM1 | I | 52801 | 54789 | -1 |
| YAL049C | 40.0566149 | 2.39480393 | 0.88044625 | 8.288864 | 0.01585244 | 0.05596684 | AIM2 | I | 51855 | 52595 | -1 |
| YAL051W | 25.0791615 | 0.68588401 | 0.76407692 | 5.04199906 | 0.08037923 | 0.18705615 | OAF1 | I | 48564 | 51707 | 1 |
| YAL053W | 134.787587 | 2.63572203 | 0.54697189 | 23.0246777 | 1.00E-05 | 0.00013257 | FLC2 | I | 45899 | 48250 | 1 |
| YAL054C | 497.265684 | -0.8265229 | 0.49057639 | 5.4794021 | 0.06458965 | 0.15839648 | ACS1 | I | 42881 | 45022 | -1 |
| YAL055W | 66.0301892 | 1.49168925 | 1.0352318 | 2.23566208 | 0.32698825 | 0.49980517 | PEX22 | I | 42177 | 42719 | 1 |
| YAL056W | 22.520543 | 0.04698075 | 0.80518076 | 0.25185743 | 0.88167769 | 0.9300006 | GPB2 | I | 39259 | 41901 | 1 |
| YAL058W | 33.6448257 | -0.1786208 | 0.65392322 | 0.26338621 | 0.87660998 | 0.92721443 | CNE1 | I | 37464 | 38972 | 1 |
| YAL059W | 17.4722431 | -1.6160216 | 0.8351619 | 8.26261199 | 0.01606189 | 0.05643103 | ECM1 | I | 36509 | 37147 | 1 |
| YAL060W | 49.8893578 | -1.2426612 | 0.64590336 | 5.70830547 | 0.05760461 | 0.1457739 | BDH1 | I | 35155 | 36303 | 1 |
| YAL061W | 827.748389 | -1.6990371 | 0.54014129 | 13.3813506 | 0.00124244 | 0.00786166 | BDH2 | I | 33448 | 34701 | 1 |
| YAL062W | 1323.60444 | -0.2183306 | 0.51146542 | 1.89413611 | 0.38787659 | 0.56008824 | GDH3 | I | 31567 | 32940 | 1 |
| YAL063C | 1429.86781 | -0.2153104 | 0.5075828 | 0.17975256 | 0.91404426 | 0.94725547 | FLO9 | I | 24000 | 27968 | -1 |
| YAL063C-A | 22.9971074 | -0.6216644 | 0.7428179 | 0.71313174 | 0.70007636 | 0.81801018 | NA | I | 22395 | 22685 | -1 |
| YAL064C-A | 286.680621 | -1.3336781 | 0.52670589 | 8.06620496 | 0.01771927 | 0.06027884 | TDA8 | I | 13363 | 13743 | -1 |
| YAL064W-B | 34.2044202 | -0.7862817 | 0.66594344 | 3.25838733 | 0.19608762 | 0.35380098 | NA | I | 12046 | 12426 | 1 |

| | | | | | | | | | | | |
|---|---|---|---|---|---|---|---|---|---|---|---|
| YAL065C | 69.9340375 | -0.9916929 | 0.57981906 | 3.24044675 | 0.1978545 | 0.35610119 | NA | I | 11565 | 11951 | -1 |
| YAL067C | 148.676613 | -0.297003 | 0.54739016 | 2.90969397 | 0.23343608 | 0.40082886 | SEO1 | I | 7235 | 9016 | -1 |
| YAR002C-A | 25.4323182 | 0.32245166 | 0.67269316 | 1.55151091 | 0.46035587 | 0.62448065 | ERP1 | I | 154065 | 154724 | -1 |
| YAR002W | 97.1288927 | 0.76883559 | 0.64958507 | 6.30174738 | 0.0428147 | 0.11665747 | NUP60 | I | 152257 | 153876 | 1 |
| YAR003W | 31.4873408 | -0.2087569 | 0.65310525 | 0.52461139 | 0.76927583 | 0.85953436 | SWD1 | I | 155005 | 156285 | 1 |
| YAR007C | 217.684773 | 1.32651408 | 0.69707319 | 6.20963477 | 0.0448327 | 0.12051131 | RFA1 | I | 156754 | 158619 | -1 |
| YAR008W | 9.97066093 | -1.2477518 | 0.97761044 | 1.64471825 | 0.43939385 | 0.60805219 | SEN34 | I | 158966 | 159793 | 1 |
| YAR010C | 8.12399364 | -1.0892601 | 1.03674619 | 1.21944634 | 0.54350131 | 0.69699868 | NA | I | 164544 | 165866 | -1 |
| YAR014C | 49.9580748 | -0.6410288 | 0.60148208 | 3.78940721 | 0.1503629 | 0.2931317 | BUD14 | I | 166742 | 168871 | -1 |
| YAR015W | 88.8911506 | 0.06344534 | 0.60746682 | 19.0755842 | 7.21E-05 | 0.00071215 | ADE1 | I | 169375 | 170295 | 1 |
| YAR018C | 163.651055 | 0.1145507 | 0.5347128 | 0.63853774 | 0.72668014 | 0.83481707 | KIN3 | I | 170396 | 171703 | -1 |
| YAR019C | 67.7457183 | -0.6276871 | 0.60689385 | 3.65945777 | 0.16045706 | 0.30793716 | CDC15 | I | 172211 | 175135 | -1 |
| YAR023C | 83.6213596 | -0.8329767 | 0.6112855 | 9.25607291 | 0.00977393 | 0.03870798 | NA | I | 179281 | 179820 | -1 |
| YAR027W | 69.3232713 | 0.47991789 | 0.58893157 | 1.26988498 | 0.52996597 | 0.68432269 | UIP3 | I | 183770 | 184477 | 1 |
| YAR028W | 33.6169365 | -0.9263783 | 0.63234009 | 5.56038257 | 0.06202664 | 0.15354179 | NA | I | 184892 | 185596 | 1 |
| YAR029W | 35.7153511 | -1.321115 | 0.73985712 | 6.18521251 | 0.04538352 | 0.12165304 | NA | I | 186321 | 186545 | 1 |
| YAR031W | 441.518308 | -0.9795072 | 0.53591647 | 6.26412197 | 0.04362779 | 0.11842705 | PRM9 | I | 186836 | 187732 | 1 |
| YAR033W | 291.320116 | -1.3410094 | 0.53410655 | 6.69695967 | 0.03513773 | 0.10076644 | MST28 | I | 188107 | 188811 | 1 |
| YAR035C-A | 55.3368512 | 0.44039604 | 0.73802752 | 2.03780421 | 0.36099105 | 0.53387949 | NA | I | 192337 | 192417 | -1 |
| YAR035W | 120.492423 | -0.9873561 | 0.72910584 | 1.97912543 | 0.37173921 | 0.54514152 | YAT1 | I | 190193 | 192256 | 1 |
| YAR042W | 51.8015762 | 0.11323 | 0.61476302 | 0.30227492 | 0.85972951 | 0.91706593 | SWH1 | I | 192619 | 196185 | 1 |
| YAR050W | 305.750074 | -0.765098 | 0.54215214 | 2.76248446 | 0.25126623 | 0.42270232 | FLO1 | I | 203403 | 208016 | 1 |
| YAR061W | 17.5073638 | -1.1738068 | 0.79612582 | 9.46594497 | 0.00880027 | 0.03573389 | NA | I | 218140 | 219145 | 1 |
| YAR064W | 147.531005 | -1.0724421 | 0.52992936 | 5.58603885 | 0.06123604 | 0.15215948 | NA | I | 220198 | 220497 | 1 |
| YAR066W | 8.62362385 | -3.162205 | 1.30636511 | 6.14678384 | 0.04626396 | 0.12315786 | NA | I | 221049 | 221660 | 1 |
| YAR068W | 74.5682182 | -1.2212886 | 0.60616978 | 6.02355672 | 0.0492041 | 0.12908551 | NA | I | 222406 | 222891 | 1 |
| YBL001C | 53.1066746 | -0.2398103 | 0.63852231 | 3.89050831 | 0.14295089 | 0.2827761 | ECM15 | II | 237153 | 237467 | -1 |
| YBL002W | 56.0987531 | -2.1460553 | 0.75821677 | 19.5508116 | 5.68E-05 | 0.00057634 | HTB2 | II | 236492 | 236887 | 1 |
| YBL003C | 9.30485265 | -1.7674396 | 1.01744255 | 8.49506124 | 0.0142995 | 0.05164948 | HTA2 | II | 235394 | 235792 | -1 |
| YBL004W | 157.787747 | -0.6836591 | 0.59413274 | 1.9257807 | 0.38178779 | 0.55529381 | UTP20 | II | 227636 | 235117 | 1 |
| YBL005W | 3766.36863 | -0.9249149 | 0.48094964 | 7.24215194 | 0.02675387 | 0.08152891 | PDR3 | II | 217470 | 220400 | 1 |
| YBL005W-B | 8.43483097 | -0.0644296 | 1.00860525 | 0.2868475 | 0.86638686 | 0.92150571 | NA | II | 221330 | 226598 | 1 |
| YBL006C | 330.345265 | 0.74163974 | 0.55124394 | 19.7882846 | 5.05E-05 | 0.00051812 | LDB7 | II | 216587 | 217129 | -1 |

| | | | | | | | | | | | |
|---|---|---|---|---|---|---|---|---|---|---|---|
| YBL007C | 70.9587208 | 0.51992799 | 0.56855613 | 1.56572718 | 0.4570952 | 0.62243209 | SLA1 | II | 212632 | 216366 | -1 |
| YBL008W | 9.26943991 | -0.1767541 | 1.04488316 | 0.22319884 | 0.89440246 | 0.9350658 | HIR1 | II | 209653 | 212175 | 1 |
| YBL008W-A | 9.75644654 | 0.28914601 | 1.00737854 | 3.22021206 | 0.19986642 | 0.35894125 | NA | II | 209409 | 209648 | 1 |
| YBL009W | 81.3175795 | 0.19722345 | 0.56313635 | 0.52799488 | 0.7679755 | 0.85918999 | ALK2 | II | 207194 | 209224 | 1 |
| YBL010C | 11.6849997 | -0.1821959 | 0.89178906 | 0.25620379 | 0.87976373 | 0.92911891 | NA | II | 206107 | 206949 | -1 |
| YBL011W | 120.42145 | -0.9161992 | 0.50392135 | 7.62458665 | 0.02209744 | 0.07131784 | SCT1 | II | 203538 | 205817 | 1 |
| YBL013W | 45.950479 | -0.5081153 | 0.64831398 | 0.61788118 | 0.73422439 | 0.83775415 | FMT1 | II | 202056 | 203261 | 1 |
| YBL014C | 19.8043631 | -0.0882269 | 0.7436459 | 0.04845999 | 0.9760612 | 0.98148564 | RRN6 | II | 199064 | 201748 | -1 |
| YBL015W | 41.1753417 | -1.0917314 | 0.66877707 | 6.93193515 | 0.03124276 | 0.09229367 | ACH1 | II | 194122 | 195702 | 1 |
| YBL016W | 21.4586369 | -0.5677316 | 0.77169308 | 5.33290125 | 0.06949846 | 0.16745573 | FUS3 | II | 192451 | 193512 | 1 |
| YBL017C | 62.5472072 | 0.55127636 | 0.84419777 | 0.48857882 | 0.78326092 | 0.86895587 | PEP1 | II | 186844 | 191583 | -1 |
| YBL018C | 96.7110787 | 2.72331415 | 0.92745588 | 7.74443175 | 0.0208122 | 0.06800375 | POP8 | II | 185998 | 186474 | -1 |
| YBL019W | 428.27037 | -0.5185219 | 0.47174177 | 12.3005612 | 0.00213288 | 0.01172782 | APN2 | II | 184353 | 185915 | 1 |
| YBL020W | 32.5005079 | -0.5118226 | 0.65075561 | 0.91607105 | 0.632525 | 0.76955658 | RFT1 | II | 182401 | 184125 | 1 |
| YBL021C | 69.7223875 | -1.4641629 | 0.64039805 | 6.54887142 | 0.03783821 | 0.10666176 | HAP3 | II | 181660 | 182094 | -1 |
| YBL022C | 41.2835978 | 0.22466811 | 0.6665578 | 0.44984012 | 0.79858006 | 0.87821055 | PIM1 | II | 177874 | 181275 | -1 |
| YBL023C | 102.386519 | 0.35309032 | 0.52609713 | 4.40254901 | 0.11066203 | 0.23547709 | MCM2 | II | 174920 | 177526 | -1 |
| YBL024W | 43.4956088 | -1.0002344 | 0.87377845 | 6.22838802 | 0.04441429 | 0.11983849 | NCL1 | II | 172534 | 174588 | 1 |
| YBL025W | 112.266141 | -0.1830041 | 0.56181199 | 7.12594913 | 0.02835436 | 0.0851955 | RRN10 | II | 171481 | 171918 | 1 |
| YBL026W | 7.21219264 | 2.37841563 | 1.18893184 | 4.10672284 | 0.1283029 | 0.26194421 | LSM2 | II | 170623 | 171038 | 1 |
| YBL027W | 367.323685 | -0.2974564 | 0.61322348 | 7.19141098 | 0.02744132 | 0.08314728 | RPL19B | II | 168423 | 169376 | 1 |
| YBL028C | 25.7178279 | -1.5777235 | 0.72569304 | 11.9589722 | 0.00253013 | 0.01336627 | NA | II | 167518 | 167838 | -1 |
| YBL029C-A | 47.5392462 | -0.5546294 | 0.5992326 | 9.99336414 | 0.00676034 | 0.02921072 | NA | II | 164488 | 164772 | -1 |
| YBL029W | 1712.60259 | 1.3342865 | 0.54803774 | 14.3704963 | 0.00075768 | 0.00521018 | NA | II | 166134 | 167264 | 1 |
| YBL030C | 95.1418224 | -0.2157557 | 0.65196673 | 0.276045 | 0.87107909 | 0.92508216 | PET9 | II | 163041 | 163997 | -1 |
| YBL032W | 89.9842815 | -1.1828637 | 0.52350062 | 8.69119495 | 0.01296376 | 0.0478395 | HEK2 | II | 160184 | 161329 | 1 |
| YBL033C | 26.6861463 | 1.18319698 | 0.78288167 | 3.38265696 | 0.18427456 | 0.33979289 | RIB1 | II | 158656 | 159693 | -1 |
| YBL034C | 109.306107 | -0.7904752 | 0.58180682 | 22.4493111 | 1.33E-05 | 0.00016903 | STU1 | II | 153848 | 158389 | -1 |
| YBL035C | 317.806195 | 0.20466761 | 0.51401212 | 48.9089536 | 2.40E-11 | 1.30E-09 | POL12 | II | 151493 | 153610 | -1 |
| YBL036C | 32.439987 | 0.34285244 | 0.67107178 | 2.71430807 | 0.25739227 | 0.42958807 | NA | II | 150447 | 151220 | -1 |
| YBL037W | 21.1683494 | 0.83546786 | 0.79303951 | 1.78364015 | 0.40990901 | 0.58142409 | APL3 | II | 147209 | 150286 | 1 |
| YBL038W | 35.2141507 | 0.7809046 | 0.67316889 | 2.70934266 | 0.25803209 | 0.42992972 | MRPL16 | II | 146187 | 146885 | 1 |
| YBL039C | 54.7332205 | -1.5601353 | 0.70575793 | 5.74223134 | 0.0566357 | 0.14401437 | URA7 | II | 143989 | 145728 | -1 |

| | | | | | | | | | | | |
|---|---|---|---|---|---|---|---|---|---|---|---|
| YBL039W-B | 15.1852469 | -0.2014707 | 0.87588311 | 0.40524203 | 0.81658766 | 0.88956586 | NA | II | 143393 | 143572 | 1 |
| YBL040C | 222.504594 | 2.33941827 | 0.70084465 | 15.5111039 | 0.00042836 | 0.00324208 | ERD2 | II | 142112 | 142868 | -1 |
| YBL041W | 52.1723582 | 1.18936565 | 0.61873039 | 9.2774313 | 0.00967011 | 0.03840187 | PRE7 | II | 141247 | 141972 | 1 |
| YBL042C | 53.4812589 | -0.9785926 | 0.59639094 | 2.79206681 | 0.24757706 | 0.41832973 | FUI1 | II | 138341 | 140260 | -1 |
| YBL045C | 27.9930743 | -0.0168158 | 0.84215075 | 0.06654808 | 0.96727345 | 0.97486964 | COR1 | II | 134143 | 135516 | -1 |
| YBL046W | 24.1545039 | 1.07782737 | 0.78087418 | 2.00228486 | 0.36745941 | 0.54054623 | PSY4 | II | 132424 | 133749 | 1 |
| YBL047C | 113.041398 | 0.44175517 | 0.52179462 | 2.9115292 | 0.23322197 | 0.40069888 | EDE1 | II | 127895 | 132040 | -1 |
| YBL048W | 303.200081 | 0.52707647 | 0.78810554 | 12.2922292 | 0.00214179 | 0.01176561 | NA | II | 127299 | 127610 | 1 |
| YBL049W | 231.32859 | -1.382898 | 0.56457766 | 29.8097622 | 3.36E-07 | 6.83E-06 | MOH1 | II | 126828 | 127244 | 1 |
| YBL050W | 36.1125016 | 0.64840486 | 0.77664329 | 0.72594747 | 0.6956047 | 0.81611981 | SEC17 | II | 125125 | 126119 | 1 |
| YBL051C | 57.4560575 | 0.18708847 | 0.58271219 | 1.11412119 | 0.57289055 | 0.72277589 | PIN4 | II | 122753 | 124759 | -1 |
| YBL052C | 222.49205 | 0.72568076 | 0.55711012 | 42.2045152 | 6.85E-10 | 2.54E-08 | SAS3 | II | 119379 | 121874 | -1 |
| YBL054W | 60.7158248 | -0.1795349 | 0.63701372 | 10.8555403 | 0.00439288 | 0.02061165 | TOD6 | II | 117589 | 119166 | 1 |
| YBL055C | 452.452705 | -1.0922807 | 0.53164022 | 12.7963205 | 0.00166462 | 0.00978886 | NA | II | 115573 | 116829 | -1 |
| YBL056W | 301.265659 | 0.13873676 | 0.47603333 | 7.31504693 | 0.02579632 | 0.07948946 | PTC3 | II | 113762 | 115168 | 1 |
| YBL057C | 297.034107 | 1.92251571 | 0.84750345 | 7.27661744 | 0.02629678 | 0.08055998 | PTH2 | II | 112801 | 113427 | -1 |
| YBL058W | 71.9131435 | 0.45101302 | 0.5797616 | 0.60468581 | 0.73908458 | 0.84040603 | SHP1 | II | 111437 | 112708 | 1 |
| YBL059C-A | 10.3655581 | 0.68977566 | 0.93029556 | 3.68273679 | 0.15860025 | 0.30518293 | CMC2 | II | 110125 | 110539 | -1 |
| YBL059W | 106.594204 | -0.4690895 | 0.6100038 | 2.23885061 | 0.32646736 | 0.49927259 | NA | II | 110594 | 111244 | 1 |
| YBL060W | 224.055767 | -0.0660892 | 0.62185522 | 7.07499372 | 0.02908604 | 0.08680999 | YEL1 | II | 107932 | 109995 | 1 |
| YBL061C | 68.3948846 | 1.41282612 | 0.71429361 | 4.3733762 | 0.11228802 | 0.23858629 | SKT5 | II | 105316 | 107406 | -1 |
| YBL063W | 114.113116 | 0.09430194 | 0.54400258 | 15.4428098 | 0.00044324 | 0.00334159 | KIP1 | II | 101886 | 105221 | 1 |
| YBL064C | 12.5639633 | -0.0572677 | 0.88267673 | 0.27244411 | 0.87264884 | 0.92527295 | PRX1 | II | 100371 | 101156 | -1 |
| YBL066C | 227.590091 | 0.07424737 | 0.55919378 | 22.8304011 | 1.10E-05 | 0.00014379 | SEF1 | II | 96669 | 100115 | -1 |
| YBL067C | 40.0367263 | -0.0349883 | 0.60566155 | 3.07495386 | 0.21492268 | 0.3788135 | UBP13 | II | 93639 | 95882 | -1 |
| YBL069W | 144.723039 | -0.7143953 | 0.51931151 | 5.81016596 | 0.05474425 | 0.14006486 | AST1 | II | 90741 | 92030 | 1 |
| YBL071C | 1329.8121 | -0.5655316 | 0.50297492 | 4.1168539 | 0.12765462 | 0.26140797 | NA | II | 90223 | 90531 | -1 |
| YBL071W-A | 8.80941821 | -1.2695449 | 0.98488984 | 6.01916821 | 0.04931218 | 0.12931048 | KTI11 | II | 89978 | 90226 | 1 |
| YBL072C | 110.818765 | -0.5492519 | 0.60121864 | 16.9957962 | 0.0002039 | 0.00173612 | RPS8A | II | 88523 | 89125 | -1 |
| YBL075C | 32.4599416 | -0.6805011 | 0.76859997 | 1.69780288 | 0.42788473 | 0.59755248 | SSA3 | II | 84499 | 86448 | -1 |
| YBL076C | 102.332662 | -0.0394497 | 0.69925254 | 0.44529037 | 0.80039879 | 0.87954242 | ILS1 | II | 81043 | 84261 | -1 |
| YBL078C | 9.26171442 | 4.1988193 | 1.58530784 | 13.335567 | 0.00127121 | 0.00800643 | ATG8 | II | 80378 | 80731 | -1 |
| YBL079W | 67.4164512 | -0.6548634 | 0.6566961 | 29.4295085 | 4.07E-07 | 8.07E-06 | NUP170 | II | 75259 | 79767 | 1 |

| | | | | | | | | | | | |
|---|---|---|---|---|---|---|---|---|---|---|---|
| YBL080C | 117.420865 | 0.85357128 | 0.85355751 | 7.71033728 | 0.02117003 | 0.06905605 | PET112 | II | 73070 | 74695 | -1 |
| YBL081W | 96.7393112 | 1.07504219 | 0.75874341 | 4.02228974 | 0.13383536 | 0.26992447 | NA | II | 71866 | 72972 | 1 |
| YBL082C | 12.0103079 | -0.268045 | 0.89543838 | 0.31269134 | 0.8552635 | 0.91415464 | ALG3 | II | 69751 | 71127 | -1 |
| YBL084C | 8.01706199 | -1.6534438 | 1.17651429 | 3.46665665 | 0.17669533 | 0.32927775 | CDC27 | II | 67169 | 69445 | -1 |
| YBL085W | 24.9589338 | -0.0914846 | 0.72102587 | 3.66874054 | 0.15971405 | 0.3068163 | BOI1 | II | 63876 | 66818 | 1 |
| YBL086C | 23.4037512 | -0.0595673 | 0.7081346 | 1.10675015 | 0.57500584 | 0.72422968 | NA | II | 61202 | 62602 | -1 |
| YBL087C | 68.1187718 | -0.3688142 | 0.69368971 | 1.8757389 | 0.39146098 | 0.56270086 | RPL23A | II | 59822 | 60739 | -1 |
| YBL088C | 170.239302 | 0.00941643 | 0.5701547 | 4.600161 | 0.10025077 | 0.21944212 | TEL1 | II | 51019 | 59382 | -1 |
| YBL089W | 169.767331 | 1.11636967 | 0.68743592 | 3.34188847 | 0.1880694 | 0.34376539 | AVT5 | II | 49574 | 50953 | 1 |
| YBL090W | 21.3252378 | -1.1199586 | 0.85447605 | 2.11592003 | 0.3471633 | 0.52061007 | MRP21 | II | 48825 | 49358 | 1 |
| YBL091C | 86.2687368 | 0.30617221 | 0.59402439 | 3.74043188 | 0.15409038 | 0.2985646 | MAP2 | II | 47363 | 48628 | -1 |
| YBL091C-A | 207.301916 | -0.3362528 | 0.51047637 | 5.51527183 | 0.06344157 | 0.15629934 | SCS22 | II | 46565 | 47180 | -1 |
| YBL092W | 228.345945 | -0.6873362 | 0.58612259 | 22.0308759 | 1.64E-05 | 0.00020004 | RPL32 | II | 45978 | 46370 | 1 |
| YBL093C | 33.2344153 | 0.32930624 | 0.6457535 | 0.26435609 | 0.87618498 | 0.92712351 | ROX3 | II | 44256 | 44918 | -1 |
| YBL095W | 151.879964 | 1.06607814 | 0.56080485 | 3.88173361 | 0.14357944 | 0.28372866 | MRX3 | II | 43277 | 44089 | 1 |
| YBL097W | 27.4763209 | 0.1250408 | 0.74660994 | 0.28948797 | 0.86524378 | 0.92057359 | BRN1 | II | 40831 | 43095 | 1 |
| YBL098W | 15.5285733 | 0.06798518 | 0.96241365 | 1.02984144 | 0.59754797 | 0.74468418 | BNA4 | II | 39145 | 40527 | 1 |
| YBL099W | 55.9867717 | 0.53520951 | 0.63573248 | 1.8981586 | 0.38709726 | 0.55948406 | ATP1 | II | 37053 | 38690 | 1 |
| YBL100W-B | 14.6292161 | 1.74050517 | 0.886708 | 3.96893687 | 0.13745366 | 0.27509737 | NA | II | 29935 | 35248 | 1 |
| YBL100W-C | 144.709954 | 0.28455862 | 0.62649453 | 0.98468769 | 0.61119218 | 0.75489714 | NA | II | 28427 | 28546 | 1 |
| YBL101C | 72.7897334 | 1.27642681 | 0.61206602 | 4.60580758 | 0.09996814 | 0.21919967 | ECM21 | II | 24946 | 28299 | -1 |
| YBL102W | 14.1911084 | -1.0604172 | 0.86619094 | 7.5122462 | 0.02337418 | 0.07452452 | SFT2 | II | 24098 | 24745 | 1 |
| YBL103C | 22.2479841 | 0.03818366 | 0.80155582 | 4.56266755 | 0.10214787 | 0.22276062 | RTG3 | II | 22076 | 23536 | -1 |
| YBL104C | 31.1630661 | -0.1985008 | 0.72493066 | 3.89639134 | 0.14253101 | 0.28204189 | SEA4 | II | 18177 | 21293 | -1 |
| YBL105C | 31.6262671 | 0.90120319 | 0.71187091 | 2.35663212 | 0.30779661 | 0.48023239 | PKC1 | II | 14241 | 17696 | -1 |
| YBL106C | 1399.00522 | 0.92239994 | 0.47200953 | 12.7329993 | 0.00171816 | 0.00999815 | SRO77 | II | 10847 | 13879 | -1 |
| YBL107C | 18.6855308 | 0.46631224 | 0.76714846 | 0.39033667 | 0.82269616 | 0.89359303 | MIX23 | II | 9961 | 10551 | -1 |
| YBL111C | 14.091861 | -1.673971 | 0.86429608 | 4.7218338 | 0.09433369 | 0.20983176 | NA | II | 2907 | 5009 | -1 |
| YBL113C | 30.8960372 | -1.8180266 | 0.67699959 | 12.133227 | 0.00231901 | 0.01246712 | NA | II | 280 | 2658 | -1 |
| YBR001C | 199.054503 | 0.34335697 | 0.4949638 | 1.71754101 | 0.42368268 | 0.59440008 | NTH2 | II | 238941 | 241283 | -1 |
| YBR002C | 129.37105 | 1.22236496 | 0.6000835 | 26.1409825 | 2.11E-06 | 3.36E-05 | RER2 | II | 241708 | 242568 | -1 |
| YBR003W | 13.5564005 | 0.96159882 | 0.91457252 | 1.80998806 | 0.4045443 | 0.57550651 | COQ1 | II | 242809 | 244230 | 1 |
| YBR004C | 48.6459951 | 1.22708016 | 0.74946984 | 4.72930219 | 0.09398209 | 0.2093602 | GPI18 | II | 244366 | 245667 | -1 |

| | | | | | | | | | | | |
|---|---|---|---|---|---|---|---|---|---|---|---|
| YBR005W | 69.3974136 | -1.2547989 | 0.56144428 | 7.28507241 | 0.02618585 | 0.08039027 | RCR1 | II | 245906 | 246547 | 1 |
| YBR006W | 197.431846 | -0.955538 | 0.51990849 | 26.3373187 | 1.91E-06 | 3.15E-05 | UGA2 | II | 247010 | 248503 | 1 |
| YBR007C | 86.0059986 | -1.1404383 | 0.55753172 | 5.45980471 | 0.06522566 | 0.15948335 | DSF2 | II | 248806 | 251016 | -1 |
| YBR008C | 146.164609 | 0.68944533 | 0.60250821 | 1.67012904 | 0.43384649 | 0.60224674 | FLR1 | II | 252563 | 254209 | -1 |
| YBR009C | 118.531894 | -2.4511576 | 0.85145117 | 21.4571133 | 2.19E-05 | 0.0002568 | HHF2 | II | 255373 | 255684 | -1 |
| YBR010W | 97.8327738 | -1.9986095 | 0.60745967 | 22.0359264 | 1.64E-05 | 0.00019996 | HHT2 | II | 256331 | 256741 | 1 |
| YBR011C | 104.608298 | 0.05411616 | 0.63088439 | 2.66536803 | 0.26376835 | 0.43572423 | IPP1 | II | 257112 | 257975 | -1 |
| YBR014C | 13.4768211 | -0.9931587 | 0.86302502 | 2.43468659 | 0.29601555 | 0.46995614 | GRX7 | II | 266725 | 267336 | -1 |
| YBR015C | 28.9018869 | 0.14591791 | 0.69104079 | 4.59256369 | 0.10063232 | 0.21978918 | MNN2 | II | 267710 | 269503 | -1 |
| YBR016W | 120.602703 | 1.61932682 | 0.56588387 | 11.9442554 | 0.00254881 | 0.01344046 | NA | II | 270247 | 270633 | 1 |
| YBR017C | 66.2364598 | 0.84737852 | 0.61776653 | 4.50891672 | 0.10493036 | 0.22691719 | KAP104 | II | 270947 | 273703 | -1 |
| YBR019C | 1096.21125 | 0.05780036 | 0.56530781 | 0.01400937 | 0.99301979 | 0.99456575 | GAL10 | II | 276253 | 278352 | -1 |
| YBR021W | 124.651317 | 1.11234539 | 0.60966094 | 32.2013738 | 1.02E-07 | 2.48E-06 | FUR4 | II | 281443 | 283344 | 1 |
| YBR022W | 231.333759 | 1.13551968 | 0.5859149 | 6.24140653 | 0.04412613 | 0.11933221 | POA1 | II | 283738 | 284271 | 1 |
| YBR023C | 96.8530186 | 1.40665181 | 0.59227477 | 6.36520995 | 0.04147747 | 0.11428804 | CHS3 | II | 284428 | 287925 | -1 |
| YBR024W | 16.756137 | -0.7144008 | 0.76782917 | 4.77563996 | 0.09182966 | 0.20608283 | SCO2 | II | 289445 | 290350 | 1 |
| YBR025C | 116.807621 | -0.3859489 | 0.57882161 | 0.93796399 | 0.62563885 | 0.76430043 | OLA1 | II | 290681 | 291865 | -1 |
| YBR026C | 20.2491715 | 0.44445148 | 0.81821797 | 0.76804189 | 0.68111716 | 0.80384598 | ETR1 | II | 292877 | 294019 | -1 |
| YBR028C | 22.6467773 | -0.9333028 | 0.719703 | 1.83877443 | 0.39876332 | 0.56966189 | YPK3 | II | 294425 | 296002 | -1 |
| YBR029C | 28.5505022 | -0.8574214 | 0.67552212 | 3.53928631 | 0.17039378 | 0.32063048 | CDS1 | II | 296369 | 297742 | -1 |
| YBR030W | 42.0143239 | -1.4283388 | 0.64206441 | 9.78143041 | 0.00751605 | 0.03164938 | RKM3 | II | 298292 | 299950 | 1 |
| YBR031W | 53.6380664 | -0.387563 | 0.70735616 | 7.88036144 | 0.0194447 | 0.06477837 | RPL4A | II | 300166 | 301254 | 1 |
| YBR032W | 298.776708 | 0.30110957 | 0.56178753 | 1.37841317 | 0.50197418 | 0.66070255 | NA | II | 301519 | 301821 | 1 |
| YBR033W | 1841.15713 | -0.6433895 | 0.49775937 | 1.92887702 | 0.38119718 | 0.55483451 | EDS1 | II | 301944 | 304703 | 1 |
| YBR034C | 29.3050338 | 0.49511828 | 0.68938269 | 1.15630228 | 0.5609345 | 0.7122392 | HMT1 | II | 304930 | 305976 | -1 |
| YBR035C | 94.6979886 | -0.5797619 | 0.58444877 | 0.99138978 | 0.60914747 | 0.75306988 | PDX3 | II | 306269 | 306955 | -1 |
| YBR036C | 30.0687325 | 0.97928543 | 0.71908226 | 2.16639785 | 0.33851092 | 0.51174366 | CSG2 | II | 309081 | 310313 | -1 |
| YBR038W | 36.0318055 | -0.3328507 | 0.67840681 | 0.89362721 | 0.63966313 | 0.77534214 | CHS2 | II | 311898 | 314789 | 1 |
| YBR039W | 18.382251 | -0.5351585 | 0.88634135 | 2.05398635 | 0.35808203 | 0.53120548 | ATP3 | II | 315575 | 316510 | 1 |
| YBR041W | 83.6906172 | 1.1855047 | 0.57919718 | 15.6761362 | 0.00039443 | 0.00302885 | FAT1 | II | 318266 | 320275 | 1 |
| YBR042C | 240.702337 | 2.41071642 | 0.7677029 | 12.4347806 | 0.00199444 | 0.01112927 | CST26 | II | 320416 | 321609 | -1 |
| YBR043C | 60.1466405 | -0.3024098 | 0.59595955 | 5.46987457 | 0.06489808 | 0.15888366 | QDR3 | II | 321876 | 323945 | -1 |
| YBR044C | 49.1064942 | 0.71932115 | 0.5983439 | 1.42840495 | 0.48958241 | 0.65106127 | TCM62 | II | 324341 | 326059 | -1 |

| | | | | | | | | | | | |
|---|---|---|---|---|---|---|---|---|---|---|---|
| YBR045C | 664.991476 | 3.45053837 | 0.48367462 | 55.8194913 | 7.57E-13 | 5.41E-11 | GIP1 | II | 328173 | 330092 | -1 |
| YBR046C | 197.333702 | 3.49160285 | 0.59257828 | 38.1700814 | 5.15E-09 | 1.64E-07 | ZTA1 | II | 330507 | 331511 | -1 |
| YBR048W | 146.668327 | -0.5429127 | 0.68045205 | 18.0211897 | 0.00012211 | 0.00110991 | RPS11B | II | 332831 | 333812 | 1 |
| YBR049C | 115.166347 | -0.6865877 | 0.59050405 | 10.8300046 | 0.00444933 | 0.02079165 | REB1 | II | 334386 | 336818 | -1 |
| YBR050C | 80.620294 | 0.3598102 | 0.59812554 | 24.9539852 | 3.81E-06 | 5.66E-05 | REG2 | II | 337183 | 338199 | -1 |
| YBR052C | 13.7935772 | 0.27169621 | 0.84387193 | 0.12415145 | 0.93981171 | 0.96037943 | RFS1 | II | 338720 | 339352 | -1 |
| YBR053C | 80.3652142 | 0.71193309 | 0.6440129 | 17.1158279 | 0.00019202 | 0.00164466 | NA | II | 339675 | 340751 | -1 |
| YBR054W | 73.2061936 | -1.2322626 | 0.62411169 | 4.01230815 | 0.13450498 | 0.27098851 | YRO2 | II | 343101 | 344135 | 1 |
| YBR055C | 77.4420046 | 1.89374482 | 1.01117514 | 3.48445728 | 0.17512966 | 0.32688612 | PRP6 | II | 344602 | 347301 | -1 |
| YBR056W | 65.3642353 | 2.21254738 | 0.61223736 | 12.7116323 | 0.00173662 | 0.01007516 | NA | II | 347879 | 349384 | 1 |
| YBR056W-A | 82.332978 | 2.52779812 | 0.58527313 | 19.8164904 | 4.98E-05 | 0.00051177 | NA | II | 351255 | 351455 | 1 |
| YBR057C | 132.032365 | 1.33550085 | 0.55065952 | 36.3115969 | 1.30E-08 | 3.77E-07 | MUM2 | II | 352193 | 353293 | -1 |
| YBR058C | 28.4624625 | 0.13690921 | 0.76229218 | 0.0761986 | 0.96261735 | 0.97258112 | UBP14 | II | 353672 | 356017 | -1 |
| YBR058C-A | 145.863237 | -0.3868166 | 0.49785078 | 0.78822588 | 0.6742779 | 0.79886823 | TSC3 | II | 356324 | 356566 | -1 |
| YBR059C | 117.715 | -0.2324549 | 0.54057578 | 0.28396682 | 0.86763565 | 0.92227105 | AKL1 | II | 356861 | 360187 | -1 |
| YBR060C | 53.1128642 | -0.78839 | 0.58975409 | 2.58611939 | 0.27442983 | 0.44796975 | ORC2 | II | 360652 | 362514 | -1 |
| YBR061C | 20.8055825 | 0.01643943 | 0.79156027 | 3.22679308 | 0.19920984 | 0.35798416 | TRM7 | II | 364787 | 365719 | -1 |
| YBR062C | 85.6958833 | -1.3977257 | 0.52636655 | 16.0155032 | 0.00033287 | 0.00266206 | NA | II | 365976 | 366600 | -1 |
| YBR063C | 27.4086118 | -0.0175522 | 0.77015812 | 1.26810472 | 0.53043791 | 0.68477938 | NA | II | 366970 | 368184 | -1 |
| YBR065C | 24.1402283 | -0.5941452 | 0.77212471 | 7.01824775 | 0.02992312 | 0.08884865 | ECM2 | II | 368584 | 369678 | -1 |
| YBR066C | 273.271372 | 0.00885465 | 0.63339966 | 0.66597171 | 0.71678033 | 0.82837488 | NRG2 | II | 370037 | 370699 | -1 |
| YBR067C | 1218.53451 | 2.22226691 | 0.58725866 | 13.5142455 | 0.00116257 | 0.00745435 | TIP1 | II | 372103 | 372735 | -1 |
| YBR068C | 147.634629 | -0.2021928 | 0.50911846 | 1.68482801 | 0.43066963 | 0.59948398 | BAP2 | II | 373861 | 375690 | -1 |
| YBR069C | 544.691312 | -0.3517434 | 0.46865701 | 8.5079435 | 0.01420769 | 0.0513499 | TAT1 | II | 376574 | 378433 | -1 |
| YBR070C | 34.4064612 | -0.3082451 | 0.64615176 | 1.68990479 | 0.42957781 | 0.59890573 | ALG14 | II | 379221 | 379934 | -1 |
| YBR071W | 140.944152 | 0.8766774 | 0.52385976 | 4.92392579 | 0.08526742 | 0.19575668 | NA | II | 380411 | 381046 | 1 |
| YBR072W | 14.0240427 | 0.04166314 | 1.04169355 | 7.18903876 | 0.02747388 | 0.08319759 | HSP26 | II | 382030 | 382674 | 1 |
| YBR073W | 83.0644473 | 0.7800298 | 0.65626645 | 1.55134437 | 0.46039421 | 0.62448065 | RDH54 | II | 383112 | 385988 | 1 |
| YBR074W | 61.9368023 | -0.0369989 | 0.63078211 | 11.0109332 | 0.00406449 | 0.0194813 | PFF1 | II | 386286 | 389216 | 1 |
| YBR076W | 1522.50358 | 2.42783575 | 0.51563442 | 30.5600306 | 2.31E-07 | 4.96E-06 | ECM8 | II | 390227 | 391291 | 1 |
| YBR077C | 183.986512 | 2.36497044 | 0.53081495 | 30.0013666 | 3.06E-07 | 6.34E-06 | SLM4 | II | 391805 | 392293 | -1 |
| YBR078W | 103.239379 | 0.80221431 | 0.54578374 | 2.24629531 | 0.32525439 | 0.49807536 | ECM33 | II | 393123 | 394742 | 1 |
| YBR079C | 95.3169917 | -0.2929069 | 0.56662176 | 10.8314947 | 0.00444601 | 0.02079165 | RPG1 | II | 395383 | 398277 | -1 |

| | | | | | | | | | | | |
|---|---|---|---|---|---|---|---|---|---|---|---|
| YBR080C | 54.6286171 | -0.0843616 | 0.71600958 | 1.71443666 | 0.42434082 | 0.59489911 | SEC18 | II | 398614 | 400890 | -1 |
| YBR081C | 56.8902731 | 0.41090478 | 0.62067573 | 1.89907994 | 0.38691898 | 0.55936601 | SPT7 | II | 401253 | 405251 | -1 |
| YBR082C | 68.2219611 | 0.73953905 | 0.58030718 | 1.62729442 | 0.44323853 | 0.61162622 | UBC4 | II | 406628 | 407169 | -1 |
| YBR083W | 29.875278 | -0.2083403 | 0.74205063 | 2.52368894 | 0.28313132 | 0.45678444 | TEC1 | II | 409169 | 410629 | 1 |
| YBR084C-A | 115.153033 | -0.301433 | 0.60839799 | 15.5119297 | 0.00042818 | 0.00324208 | RPL19B | II | 414186 | 415261 | -1 |
| YBR084W | 52.8781684 | -0.5720431 | 0.73159684 | 0.71983765 | 0.69773296 | 0.81712659 | MIS1 | II | 411054 | 413981 | 1 |
| YBR085C-A | 29.4316524 | 1.71322176 | 0.83782571 | 11.7233328 | 0.0028465 | 0.01465951 | NA | II | 418907 | 419164 | -1 |
| YBR085W | 15.0200303 | -0.0028268 | 0.8413903 | 0.72071571 | 0.6974267 | 0.81693316 | AAC3 | II | 415983 | 416906 | 1 |
| YBR086C | 130.639245 | 1.00578494 | 0.55242662 | 8.19206232 | 0.01663858 | 0.05772162 | IST2 | II | 420201 | 423041 | -1 |
| YBR087W | 65.6285096 | 0.46295662 | 0.62698266 | 31.8148351 | 1.23E-07 | 2.89E-06 | RFC5 | II | 423765 | 424829 | 1 |
| YBR088C | 36.8058012 | -1.0934546 | 0.73696413 | 2.33798223 | 0.31068022 | 0.48342878 | POL30 | II | 424990 | 425766 | -1 |
| YBR089C-A | 5.36065785 | -0.2546261 | 1.28263385 | 0.0939936 | 0.95409045 | 0.96767513 | NHP6B | II | 426190 | 426489 | -1 |
| YBR091C | 152.571694 | 1.56865608 | 0.5991226 | 18.4444855 | 9.88E-05 | 0.00092581 | TIM12 | II | 427155 | 427484 | -1 |
| YBR092C | 611.598059 | 0.2642779 | 0.50418324 | 6.86955012 | 0.03223266 | 0.09435142 | PHO3 | II | 427698 | 429101 | -1 |
| YBR093C | 652.988375 | -2.3368558 | 0.50000926 | 48.8788694 | 2.43E-11 | 1.30E-09 | PHO5 | II | 429548 | 430951 | -1 |
| YBR094W | 22.9595185 | -0.4565949 | 0.72761836 | 1.51396105 | 0.46908067 | 0.63162258 | PBY1 | II | 432036 | 434297 | 1 |
| YBR095C | 22.3410063 | 0.24252367 | 0.71244012 | 2.28561671 | 0.31892212 | 0.49162914 | RXT2 | II | 434406 | 435698 | -1 |
| YBR096W | 26.172393 | 0.54710289 | 0.70311163 | 2.7227142 | 0.2563127 | 0.42842105 | NA | II | 436021 | 436713 | 1 |
| YBR097W | 46.4976753 | -1.2506455 | 0.64188269 | 6.42209377 | 0.04031439 | 0.11205967 | VPS15 | II | 436951 | 441315 | 1 |
| YBR098W | 816.189691 | -0.7515519 | 0.50815458 | 2.16230391 | 0.33920455 | 0.51239091 | MMS4 | II | 441515 | 443590 | 1 |
| YBR101C | 72.447252 | 0.46820346 | 0.57518041 | 1.13578945 | 0.56671728 | 0.71769314 | FES1 | II | 443821 | 444693 | -1 |
| YBR102C | 18.1135966 | -0.3909579 | 0.80305673 | 0.24829947 | 0.88324758 | 0.93100373 | EXO84 | II | 445062 | 447323 | -1 |
| YBR103W | 11.8892393 | -0.2862382 | 0.91981015 | 1.02518231 | 0.59894162 | 0.74609982 | SIF2 | II | 447709 | 449316 | 1 |
| YBR104W | 28.5549749 | -0.2519422 | 0.67270436 | 0.36024349 | 0.83516853 | 0.90300565 | YMC2 | II | 449667 | 450656 | 1 |
| YBR105C | 24.5390538 | -0.3727975 | 0.76295244 | 0.38022043 | 0.826868 | 0.89638002 | VID24 | II | 450881 | 451969 | -1 |
| YBR106W | 82.2178963 | -0.3066951 | 0.58416538 | 6.38335139 | 0.04110294 | 0.11365139 | PHO88 | II | 452658 | 453224 | 1 |
| YBR107C | 1644.84988 | 1.12973774 | 0.47937212 | 5.69620517 | 0.05795418 | 0.14626659 | IML3 | II | 453793 | 454530 | -1 |
| YBR108W | 114.444274 | 0.03133798 | 0.55429929 | 1.65352841 | 0.43746254 | 0.60627631 | AIM3 | II | 454822 | 457665 | 1 |
| YBR109C | 21.6162603 | -0.1679119 | 0.96789622 | 0.0762762 | 0.96258 | 0.97258112 | CMD1 | II | 457919 | 458362 | -1 |
| YBR110W | 18.064828 | 0.60316594 | 0.83109259 | 1.68083333 | 0.43153068 | 0.6001832 | ALG1 | II | 458872 | 460221 | 1 |
| YBR111C | 35.008564 | 0.10456079 | 0.63718656 | 0.11602606 | 0.94363765 | 0.96197675 | YSA1 | II | 461178 | 461873 | -1 |
| YBR111W-A | 13.233378 | 1.36672091 | 1.04164533 | 6.302141 | 0.04280628 | 0.11665747 | SUS1 | II | 462139 | 462579 | 1 |
| YBR112C | 259.797969 | 1.86389415 | 0.81221604 | 7.28367423 | 0.02620416 | 0.08040386 | CYC8 | II | 462870 | 465770 | -1 |

| | | | | | | | | | | | |
|---|---|---|---|---|---|---|---|---|---|---|---|
| YBR114W | 48.4270869 | 1.360518 | 0.80687312 | 3.5458067 | 0.16983917 | 0.32001317 | RAD16 | II | 467248 | 469620 | 1 |
| YBR115C | 50.7848437 | -0.024746 | 0.60171372 | 3.57788845 | 0.16713654 | 0.31645538 | LYS2 | II | 469748 | 473926 | -1 |
| YBR117C | 705.849896 | -1.3279552 | 0.48832387 | 11.0433267 | 0.00399919 | 0.01926399 | TKL2 | II | 474392 | 476437 | -1 |
| YBR118W | 147.181782 | -0.4527368 | 0.93279707 | 1.5433498 | 0.46223822 | 0.62560993 | TEF1 | II | 477671 | 479047 | 1 |
| YBR119W | 34.3293989 | -0.2943418 | 0.6399716 | 0.31353884 | 0.85490116 | 0.91395769 | MUD1 | II | 479338 | 480323 | 1 |
| YBR120C | 88.6386441 | 2.37154745 | 0.89420336 | 7.50661911 | 0.02344004 | 0.07454061 | CBP6 | II | 480435 | 480923 | -1 |
| YBR121C | 66.5495562 | -0.5615747 | 0.83449482 | 5.92500583 | 0.05168938 | 0.13420135 | NA | II | 481364 | 483367 | -1 |
| YBR122C | 32.8562004 | -1.4132661 | 0.67056163 | 9.28848634 | 0.00961681 | 0.03832161 | MRPL36 | II | 483970 | 484503 | -1 |
| YBR123C | 150.966778 | 1.11359391 | 0.60439215 | 59.7561633 | 1.06E-13 | 8.62E-12 | TFC1 | II | 484742 | 486691 | -1 |
| YBR125C | 36.2118893 | -0.627208 | 0.65828944 | 2.74278353 | 0.25375355 | 0.42586465 | PTC4 | II | 487199 | 488380 | -1 |
| YBR126C | 118.72519 | 0.41501882 | 0.60936674 | 0.56665 | 0.75327493 | 0.85085093 | TPS1 | II | 488905 | 490392 | -1 |
| YBR126W-A | 206.172159 | 0.54818816 | 0.4868285 | 1.53251012 | 0.46475027 | 0.62728439 | MEO1 | II | 490850 | 491056 | 1 |
| YBR127C | 132.563716 | -0.2275885 | 0.57688165 | 0.2060421 | 0.90210798 | 0.93824416 | VMA2 | II | 491269 | 492822 | -1 |
| YBR128C | 71.8252439 | -1.0901667 | 0.56028237 | 6.85301632 | 0.03250023 | 0.09503855 | ATG14 | II | 493081 | 494115 | -1 |
| YBR130C | 411.638871 | 2.01909141 | 0.49298474 | 44.937253 | 1.75E-10 | 7.66E-09 | SHE3 | II | 495592 | 496869 | -1 |
| YBR131W | 16.7561763 | 0.68747046 | 0.79367368 | 1.15706981 | 0.56071927 | 0.71212209 | CCZ1 | II | 497163 | 499277 | 1 |
| YBR132C | 46.1863543 | 0.00713913 | 0.59949314 | 1.78536444 | 0.40955576 | 0.5812078 | AGP2 | II | 499652 | 501442 | -1 |
| YBR133C | 33.8190352 | -0.090703 | 0.63764901 | 0.09986489 | 0.95129369 | 0.96614462 | HSL7 | II | 501804 | 504287 | -1 |
| YBR135W | 25.3770705 | 1.59510957 | 0.84021862 | 3.73155146 | 0.1547761 | 0.29951658 | CKS1 | II | 504854 | 505306 | 1 |
| YBR136W | 215.207331 | 0.26100221 | 0.53105219 | 12.8382891 | 0.00163005 | 0.00966238 | MEC1 | II | 505668 | 512774 | 1 |
| YBR137W | 16.0765811 | -0.8940254 | 0.84528819 | 1.21807896 | 0.54387302 | 0.69714961 | NA | II | 513044 | 513583 | 1 |
| YBR138C | 202.671335 | 1.4015573 | 0.55426394 | 31.4242992 | 1.50E-07 | 3.45E-06 | NA | II | 513762 | 515336 | -1 |
| YBR139W | 71.6788297 | 0.34707386 | 0.77419805 | 2.60379753 | 0.27201481 | 0.44540887 | NA | II | 515664 | 517190 | 1 |
| YBR140C | 301.086744 | 1.28314278 | 0.70233024 | 4.70408303 | 0.09517466 | 0.2111346 | IRA1 | II | 517350 | 526628 | -1 |
| YBR141C | 448.624387 | 1.09162924 | 0.53145487 | 4.70991657 | 0.09489747 | 0.21060035 | BMT2 | II | 527025 | 528038 | -1 |
| YBR142W | 87.434226 | 1.55994025 | 0.88680464 | 11.4256741 | 0.00330329 | 0.0164313 | MAK5 | II | 528317 | 530638 | 1 |
| YBR143C | 51.7063154 | -0.7099708 | 0.59226376 | 4.55268279 | 0.10265911 | 0.22312171 | SUP45 | II | 530869 | 532182 | -1 |
| YBR144C | 20.9465833 | -0.3801106 | 0.71697185 | 1.45443568 | 0.48325161 | 0.64559917 | NA | II | 533235 | 533549 | -1 |
| YBR145W | 70.140092 | -0.047445 | 0.65523288 | 0.10086927 | 0.95081608 | 0.96614462 | ADH5 | II | 533762 | 534817 | 1 |
| YBR146W | 29.9395107 | -0.3119094 | 0.66173282 | 4.86586832 | 0.0877789 | 0.20033103 | MRPS9 | II | 535260 | 536096 | 1 |
| YBR148W | 48.5252502 | -0.1096766 | 0.68656783 | 0.06059546 | 0.97015665 | 0.97724547 | YSW1 | II | 537876 | 539705 | 1 |
| YBR149W | 469.853298 | -3.0312916 | 0.52680529 | 45.7920648 | 1.14E-10 | 5.23E-09 | ARA1 | II | 539987 | 541021 | 1 |
| YBR150C | 444.125204 | 2.50453691 | 0.8932076 | 12.777176 | 0.00168063 | 0.00981378 | TBS1 | II | 541209 | 544493 | -1 |

| | | | | | | | | | | | |
|---|---|---|---|---|---|---|---|---|---|---|---|
| YBR151W | 53.8522041 | 0.53657353 | 0.7995373 | 1.90534234 | 0.38570935 | 0.55845627 | APD1 | II | 545028 | 545978 | 1 |
| YBR152W | 27.6191505 | -0.2620522 | 0.68377106 | 0.2097891 | 0.90041947 | 0.93824416 | SPP381 | II | 546376 | 547251 | 1 |
| YBR153W | 12.7059386 | -1.0942317 | 0.87214268 | 1.75606497 | 0.41559981 | 0.58733778 | RIB7 | II | 547460 | 548194 | 1 |
| YBR154C | 141.687499 | 0.37809443 | 0.61655318 | 9.27860018 | 0.00966446 | 0.03840187 | RPB5 | II | 548362 | 549009 | -1 |
| YBR155W | 64.2249131 | 1.81262634 | 1.07624881 | 3.39194056 | 0.18342117 | 0.33887818 | CNS1 | II | 549771 | 550928 | 1 |
| YBR156C | 48.4949478 | -1.0828184 | 0.67270831 | 5.69771997 | 0.0579103 | 0.14626659 | SLI15 | II | 551104 | 553200 | -1 |
| YBR157C | 260.555643 | -0.0851 | 0.50791212 | 0.24333189 | 0.8854441 | 0.931623 | ICS2 | II | 553543 | 554310 | -1 |
| YBR158W | 30.4405161 | -1.4932129 | 0.68308844 | 5.19704172 | 0.07438352 | 0.17629168 | AMN1 | II | 556549 | 558198 | 1 |
| YBR159W | 30.6797804 | 1.39848018 | 0.84067469 | 6.19831784 | 0.04508711 | 0.12108273 | IFA38 | II | 558685 | 559728 | 1 |
| YBR160W | 57.8786997 | 1.06546026 | 0.58960012 | 3.96930366 | 0.13742845 | 0.27509737 | CDC28 | II | 560078 | 560974 | 1 |
| YBR161W | 62.4827399 | 1.83593971 | 0.5941094 | 14.5529632 | 0.00069161 | 0.00483046 | CSH1 | II | 561634 | 562764 | 1 |
| YBR162C | 89.2506745 | -0.3261365 | 0.63324455 | 0.39218814 | 0.82193491 | 0.89337397 | TOS1 | II | 563203 | 564570 | -1 |
| YBR162W-A | 165.662514 | 1.30470595 | 0.59492436 | 5.54004469 | 0.0626606 | 0.15478025 | YSY6 | II | 565231 | 565428 | 1 |
| YBR163W | 66.2473236 | -0.9389129 | 0.62101381 | 3.21860746 | 0.20002684 | 0.35900663 | EXO5 | II | 565723 | 567480 | 1 |
| YBR164C | 10.2658955 | 0.09256317 | 0.95140959 | 0.55988069 | 0.75582883 | 0.85240532 | ARL1 | II | 567875 | 568426 | -1 |
| YBR165W | 14.719259 | 1.47266083 | 0.94944699 | 2.65144564 | 0.2656109 | 0.43764573 | UBS1 | II | 568852 | 569685 | 1 |
| YBR166C | 39.5956849 | 0.98237251 | 0.68268354 | 7.29781565 | 0.02601953 | 0.07997546 | TYR1 | II | 569842 | 571200 | -1 |
| YBR167C | 253.555078 | 3.67775619 | 0.57872408 | 64.833222 | 8.35E-15 | 8.48E-13 | POP7 | II | 571468 | 571890 | -1 |
| YBR168W | 13.026385 | 0.16736183 | 0.95937198 | 0.14459337 | 0.93025486 | 0.95618243 | PEX32 | II | 572371 | 573612 | 1 |
| YBR169C | 52.6384314 | 0.9949322 | 0.70624768 | 19.439104 | 6.01E-05 | 0.0006041 | SSE2 | II | 573915 | 575996 | -1 |
| YBR170C | 14.7348989 | 0.77412319 | 0.91932831 | 2.19018908 | 0.33450798 | 0.50802806 | NPL4 | II | 576344 | 578086 | -1 |
| YBR171W | 34.8072373 | 0.24039264 | 0.63613217 | 2.42024877 | 0.29816019 | 0.47161883 | SEC66 | II | 578364 | 578984 | 1 |
| YBR172C | 151.746889 | 1.53343893 | 0.68313595 | 5.44186166 | 0.06581346 | 0.16064998 | SMY2 | II | 579150 | 581372 | -1 |
| YBR173C | 113.904355 | -0.2206028 | 0.52492248 | 8.29133589 | 0.01583286 | 0.05593181 | UMP1 | II | 581726 | 582172 | -1 |
| YBR175W | 35.8064777 | -0.5530227 | 0.66155323 | 0.72627762 | 0.69548989 | 0.81611981 | SWD3 | II | 582408 | 583355 | 1 |
| YBR176W | 50.6200451 | 1.46351899 | 0.70457659 | 4.13653868 | 0.12640436 | 0.25966788 | ECM31 | II | 583720 | 584658 | 1 |
| YBR177C | 66.8611894 | 0.9441017 | 0.64528865 | 2.44414207 | 0.29461937 | 0.46889669 | EHT1 | II | 584807 | 586162 | -1 |
| YBR179C | 34.5451866 | -1.2343046 | 0.64431181 | 4.03387191 | 0.13306255 | 0.26909961 | FZO1 | II | 586547 | 589114 | -1 |
| YBR180W | 231.506979 | -0.9989336 | 0.50706238 | 7.49363034 | 0.02359277 | 0.07480948 | DTR1 | II | 589741 | 591459 | 1 |
| YBR181C | 38.048731 | -0.7200189 | 0.66462155 | 8.01524187 | 0.01817659 | 0.06161735 | RPS6A | II | 591712 | 592774 | -1 |
| YBR182C | 57.6368698 | -1.3271225 | 0.57308651 | 13.4732961 | 0.00118662 | 0.00756665 | SMP1 | II | 593506 | 594864 | -1 |
| YBR182C-A | 50.3317798 | 0.05781184 | 0.65538488 | 3.05578054 | 0.21699298 | 0.38107048 | NA | II | 595361 | 595555 | -1 |
| YBR183W | 14.2211668 | 0.70288337 | 0.85672548 | 3.27367866 | 0.19459412 | 0.35165417 | YPC1 | II | 596115 | 597065 | 1 |

| | | | | | | | | | | | |
|---|---|---|---|---|---|---|---|---|---|---|---|
| YBR184W | 52.9390505 | 0.44745626 | 0.60578491 | 3.76531306 | 0.15218528 | 0.29581776 | NA | II | 597363 | 598934 | 1 |
| YBR186W | 499.546392 | -1.3319804 | 0.47830135 | 32.3468233 | 9.46E-08 | 2.33E-06 | PCH2 | II | 600553 | 602360 | 1 |
| YBR187W | 256.798377 | 0.43362469 | 0.63007299 | 1.89547118 | 0.38761776 | 0.55995679 | GDT1 | II | 602634 | 603476 | 1 |
| YBR188C | 22.4129868 | -0.2033353 | 0.71334765 | 4.98154492 | 0.08284595 | 0.19095463 | NTC20 | II | 603685 | 604107 | -1 |
| YBR189W | 106.930075 | -0.3892945 | 0.90673031 | 10.2490286 | 0.00594911 | 0.02633435 | RPS9B | II | 604508 | 605508 | 1 |
| YBR191W | 65.5889934 | -0.5878425 | 0.69063006 | 14.019231 | 0.00090316 | 0.00605941 | RPL21A | II | 606270 | 607140 | 1 |
| YBR192W | 226.970427 | -0.2137001 | 0.50184778 | 5.66664818 | 0.05881702 | 0.14748831 | RIM2 | II | 607652 | 608785 | 1 |
| YBR193C | 26.1729235 | 0.01059455 | 0.70759205 | 0.09335679 | 0.95439429 | 0.96776584 | MED8 | II | 609082 | 609753 | -1 |
| YBR194W | 12.7124139 | -1.773904 | 0.91659768 | 3.93692561 | 0.13967139 | 0.27818967 | AIM4 | II | 610038 | 610409 | 1 |
| YBR195C | 246.347838 | 0.47900434 | 0.55284528 | 36.0642756 | 1.47E-08 | 4.25E-07 | MSI1 | II | 610614 | 611882 | -1 |
| YBR196C | 114.309634 | 0.89869642 | 0.84718847 | 1.27941806 | 0.52744587 | 0.68184633 | PGI1 | II | 612236 | 613900 | -1 |
| YBR196C-B | 62.4870326 | -0.6009848 | 0.57133733 | 13.3415387 | 0.00126742 | 0.00799388 | NA | II | 614526 | 614630 | -1 |
| YBR197C | 231.466843 | -0.1019978 | 0.55028031 | 0.39084997 | 0.82248504 | 0.89359303 | NA | II | 615203 | 615856 | -1 |
| YBR198C | 207.126918 | -0.1085349 | 0.51930175 | 3.56599695 | 0.16813325 | 0.31792668 | TAF5 | II | 616127 | 618523 | -1 |
| YBR199W | 40.0836265 | 0.98294067 | 0.75323387 | 2.12704444 | 0.34523766 | 0.51852261 | KTR4 | II | 618909 | 620303 | 1 |
| YBR200W | 29.2430876 | 0.91511681 | 0.72225418 | 2.42916241 | 0.2968343 | 0.47086866 | BEM1 | II | 620872 | 622527 | 1 |
| YBR200W-A | 226.766788 | 5.13266285 | 0.72749428 | 86.0814429 | 2.03E-19 | 4.90E-17 | NA | II | 622983 | 623147 | 1 |
| YBR201C-A | 33.4425987 | 0.54630404 | 0.66242392 | 2.19255469 | 0.33411256 | 0.5079834 | NA | II | 624493 | 624696 | -1 |
| YBR201W | 6.1737205 | -0.328413 | 1.18090331 | 1.18520531 | 0.55288644 | 0.70480239 | DER1 | II | 623576 | 624211 | 1 |
| YBR202W | 22.8064707 | -0.9092144 | 0.80934097 | 1.53320643 | 0.4645885 | 0.62728439 | MCM7 | II | 625772 | 628309 | 1 |
| YBR203W | 2248.36593 | 3.99449368 | 0.45516652 | 86.5598836 | 1.60E-19 | 4.02E-17 | COS111 | II | 629168 | 631942 | 1 |
| YBR204C | 25.5346675 | -0.8652185 | 0.71026419 | 1.55314847 | 0.45997909 | 0.62448065 | LDH1 | II | 632254 | 633381 | -1 |
| YBR205W | 30.5354967 | -0.486218 | 0.74446379 | 2.86717615 | 0.2384518 | 0.40666743 | KTR3 | II | 633622 | 634836 | 1 |
| YBR207W | 39.996762 | 0.91351998 | 0.72523451 | 1.80420851 | 0.40571503 | 0.57703022 | FTH1 | II | 635146 | 636543 | 1 |
| YBR208C | 74.2098808 | 0.58708334 | 0.57049043 | 15.8535928 | 0.00036094 | 0.00284348 | DUR1,2 | II | 636703 | 642210 | -1 |
| YBR209W | 350.089805 | 0.65812847 | 0.53630145 | 4.8251028 | 0.08958643 | 0.20301583 | NA | II | 642583 | 642900 | 1 |
| YBR210W | 9.39071885 | -0.0552134 | 0.94318564 | 1.95484544 | 0.37627963 | 0.54998972 | ERV15 | II | 645550 | 645978 | 1 |
| YBR211C | 39.6753003 | 2.21920725 | 0.85013105 | 9.78757365 | 0.00749299 | 0.03162131 | AME1 | II | 646158 | 647132 | -1 |
| YBR212W | 118.826427 | 1.89128669 | 0.58560993 | 31.3826065 | 1.53E-07 | 3.51E-06 | NGR1 | II | 647886 | 649904 | 1 |
| YBR213W | 16.338113 | 0.58601939 | 0.91984877 | 3.94428523 | 0.13915837 | 0.27774112 | MET8 | II | 650368 | 651192 | 1 |
| YBR214W | 345.850628 | -1.6814631 | 0.52609399 | 13.9170539 | 0.0009505 | 0.00631841 | SDS24 | II | 651415 | 652998 | 1 |
| YBR215W | 437.324909 | -0.4189725 | 0.51648528 | 0.7253552 | 0.69581073 | 0.81619613 | HPC2 | II | 653356 | 655317 | 1 |
| YBR216C | 53.5332662 | 0.33766507 | 0.62875936 | 5.49594147 | 0.06405772 | 0.15736105 | YBP1 | II | 655576 | 657600 | -1 |

| | | | | | | | | | | | |
|---|---|---|---|---|---|---|---|---|---|---|---|
| YBR217W | 12.8999714 | -1.3361787 | 0.92832795 | 2.49098003 | 0.28779985 | 0.46172289 | ATG12 | II | 657832 | 658392 | 1 |
| YBR218C | 93.8029793 | 0.17855056 | 0.53025817 | 6.10214652 | 0.04730812 | 0.12530377 | PYC2 | II | 658707 | 662249 | -1 |
| YBR220C | 44.5498562 | -0.5091876 | 0.66907733 | 4.99084963 | 0.08246141 | 0.19029557 | NA | II | 662995 | 664677 | -1 |
| YBR221C | 44.7315432 | 0.28925677 | 0.76788584 | 1.21744296 | 0.544046 | 0.69721698 | PDB1 | II | 665153 | 666253 | -1 |
| YBR221W-A | 1056.3207 | 2.5362485 | 0.70436178 | 13.9787369 | 0.00092163 | 0.00615027 | NA | II | 666538 | 666642 | 1 |
| YBR222C | 37.7833155 | 1.77557242 | 0.6883381 | 8.845311 | 0.01200232 | 0.04521368 | PCS60 | II | 666720 | 668351 | -1 |
| YBR223C | 8.79214549 | 0.71923258 | 1.04426495 | 1.37634122 | 0.50249449 | 0.66093664 | TDP1 | II | 668663 | 670297 | -1 |
| YBR225W | 24.6975037 | 0.09905591 | 0.74638547 | 1.146388 | 0.56372203 | 0.71483805 | NA | II | 670627 | 673329 | 1 |
| YBR227C | 213.57076 | 2.31695252 | 0.73328787 | 9.8661603 | 0.00720428 | 0.03068439 | MCX1 | II | 673572 | 675134 | -1 |
| YBR228W | 309.228892 | 0.13239443 | 0.67842912 | 5.50043539 | 0.06391395 | 0.15715409 | SLX1 | II | 675313 | 676227 | 1 |
| YBR229C | 115.033191 | -0.7081198 | 0.53388699 | 7.56352459 | 0.02278251 | 0.07300493 | ROT2 | II | 676357 | 679221 | -1 |
| YBR230C | 13.8569409 | 0.54570761 | 0.88433851 | 1.5918373 | 0.45116658 | 0.61755425 | OM14 | II | 679549 | 680050 | -1 |
| YBR230W-A | 42.025878 | 0.64037018 | 0.60750544 | 2.70078479 | 0.25913856 | 0.43082351 | NA | II | 680362 | 680562 | 1 |
| YBR231C | 19.1513397 | 0.64740632 | 0.82224403 | 1.23632984 | 0.53893252 | 0.69327244 | SWC5 | II | 682179 | 683090 | -1 |
| YBR233W | 27.4891304 | -0.9012907 | 0.74939512 | 1.8346541 | 0.39958569 | 0.57041448 | PBP2 | II | 683428 | 684669 | 1 |
| YBR234C | 88.6770024 | -0.4386941 | 0.53236244 | 7.89599379 | 0.01929331 | 0.0644223 | ARC40 | II | 685438 | 686592 | -1 |
| YBR235W | 287.738137 | -0.6240438 | 0.50542109 | 13.6383512 | 0.00109262 | 0.00709251 | VHC1 | II | 686901 | 690263 | 1 |
| YBR236C | 58.3708431 | -0.3208927 | 0.55580272 | 3.62240854 | 0.16345717 | 0.3120399 | ABD1 | II | 690383 | 691693 | -1 |
| YBR237W | 20.9281593 | -0.2062238 | 0.77237095 | 1.16181476 | 0.55939056 | 0.71059046 | PRP5 | II | 691969 | 694518 | 1 |
| YBR238C | 25.4665888 | 1.11117142 | 0.82895884 | 4.608339 | 0.09984168 | 0.21918634 | NA | II | 695107 | 697302 | -1 |
| YBR239C | 310.725895 | -0.4898759 | 0.53705752 | 1.4475218 | 0.48492507 | 0.64753601 | ERT1 | II | 698354 | 699943 | -1 |
| YBR240C | 12.730542 | -0.7442803 | 0.94026817 | 0.71768187 | 0.69848545 | 0.81736153 | THI2 | II | 700490 | 701842 | -1 |
| YBR241C | 247.186762 | 0.87613349 | 0.58344092 | 12.5517049 | 0.00188119 | 0.01071 | NA | II | 702589 | 704055 | -1 |
| YBR242W | 316.154853 | 0.82459539 | 0.58793058 | 5.63480427 | 0.05976099 | 0.14953161 | NA | II | 704670 | 705386 | 1 |
| YBR243C | 292.114403 | 0.01857196 | 0.63299623 | 1.62534504 | 0.44367076 | 0.61162622 | ALG7 | II | 705447 | 706793 | -1 |
| YBR244W | 31.8732958 | 1.3717812 | 0.79611563 | 13.6065575 | 0.00111013 | 0.00717372 | GPX2 | II | 707528 | 708016 | 1 |
| YBR245C | 58.233506 | 0.14079406 | 0.55624921 | 1.56823431 | 0.45652256 | 0.62207609 | ISW1 | II | 708150 | 711539 | -1 |
| YBR246W | 26.1981731 | 1.81111017 | 0.93891647 | 7.9062456 | 0.01919467 | 0.06416508 | RRT2 | II | 711591 | 712754 | 1 |
| YBR247C | 28.5119343 | -1.1434524 | 0.70427433 | 6.71432491 | 0.03483396 | 0.10013768 | ENP1 | II | 713004 | 714455 | -1 |
| YBR248C | 107.929456 | -1.3267683 | 0.52833462 | 18.3757855 | 0.00010227 | 0.00095353 | HIS7 | II | 714807 | 716465 | -1 |
| YBR249C | 39.1786025 | 0.43595355 | 0.67382182 | 4.3654089 | 0.11273623 | 0.23918753 | ARO4 | II | 716882 | 717994 | -1 |
| YBR251W | 12.8335635 | -0.6633579 | 0.84290386 | 1.79258047 | 0.40808074 | 0.5793986 | MRPS5 | II | 721390 | 722313 | 1 |
| YBR252W | 86.1719078 | -0.3696912 | 0.53472467 | 0.85720469 | 0.65141892 | 0.78236902 | DUT1 | II | 722611 | 723054 | 1 |

| | | | | | | | | | | | |
|---|---|---|---|---|---|---|---|---|---|---|---|
| YBR253W | 87.992407 | 1.73665967 | 0.87766751 | 7.34570743 | 0.02540387 | 0.07865691 | SRB6 | II | 723270 | 723635 | 1 |
| YBR254C | 9.19788425 | -1.2559409 | 1.03291906 | 3.01971313 | 0.22094167 | 0.38583539 | TRS20 | II | 723736 | 724263 | -1 |
| YBR255C-A | 239.809594 | 0.87877344 | 0.64590419 | 1.88011389 | 0.39060559 | 0.56175022 | RCF3 | II | 726618 | 727074 | -1 |
| YBR255W | 179.614665 | 1.84316917 | 0.66538176 | 8.37459505 | 0.01518727 | 0.05409075 | MTC4 | II | 724456 | 726540 | 1 |
| YBR256C | 33.4632869 | 0.85135835 | 0.68930659 | 3.20048553 | 0.20184751 | 0.36126649 | RIB5 | II | 727386 | 728102 | -1 |
| YBR257W | 240.368904 | 0.40931628 | 0.52808455 | 3.50396209 | 0.17343003 | 0.32507061 | POP4 | II | 728885 | 729724 | 1 |
| YBR258C | 126.136025 | 0.10574197 | 0.57935039 | 2.01682796 | 0.3647971 | 0.53799674 | SHG1 | II | 729734 | 730162 | -1 |
| YBR259W | 226.250356 | 2.72000026 | 0.64177614 | 40.4416519 | 1.65E-09 | 5.87E-08 | NA | II | 730387 | 732453 | 1 |
| YBR260C | 59.2857834 | -0.1432508 | 0.70263414 | 0.41317471 | 0.81335521 | 0.88771473 | RGD1 | II | 732639 | 734639 | -1 |
| YBR261C | 48.4329879 | -0.4132934 | 0.58952647 | 0.52061398 | 0.77081492 | 0.86042382 | TAE1 | II | 734832 | 735530 | -1 |
| YBR262C | 8.11284983 | -0.7822793 | 1.10663921 | 0.74452498 | 0.68917332 | 0.81068229 | MIC12 | II | 735720 | 736040 | -1 |
| YBR263W | 208.976017 | -0.4515224 | 0.54499905 | 13.744307 | 0.00103624 | 0.00679484 | SHM1 | II | 736264 | 737736 | 1 |
| YBR264C | 1214.10155 | 1.52562939 | 0.82905777 | 6.26259671 | 0.04366107 | 0.11846186 | YPT10 | II | 737770 | 738369 | -1 |
| YBR265W | 58.9057551 | -1.836019 | 0.61591181 | 10.2058203 | 0.00607903 | 0.02677727 | TSC10 | II | 738582 | 739544 | 1 |
| YBR266C | 3446.34823 | -1.2202793 | 0.52883166 | 20.6617721 | 3.26E-05 | 0.00036033 | NA | II | 739936 | 740388 | -1 |
| YBR267W | 36.9066703 | -1.0914908 | 0.69213334 | 5.72340266 | 0.05717141 | 0.14493103 | REI1 | II | 739841 | 741022 | 1 |
| YBR269C | 31.6057487 | 1.24133504 | 0.79234547 | 3.60312103 | 0.16504114 | 0.31392516 | SDH8 | II | 742160 | 742576 | -1 |
| YBR270C | 78.6176637 | 1.38780731 | 0.62209612 | 22.6892425 | 1.18E-05 | 0.00015327 | BIT2 | II | 742761 | 744398 | -1 |
| YBR271W | 19.2799028 | -0.4897182 | 0.73280969 | 0.73837259 | 0.69129661 | 0.81221741 | EFM2 | II | 744852 | 746111 | 1 |
| YBR272C | 13.8463636 | -0.1757817 | 0.83042861 | 0.26947329 | 0.87394604 | 0.92574965 | HSM3 | II | 746361 | 747803 | -1 |
| YBR273C | 68.696661 | 0.31025845 | 0.60827248 | 14.3504103 | 0.00076533 | 0.00525653 | UBX7 | II | 748061 | 749371 | -1 |
| YBR274W | 16.757086 | -0.1303489 | 0.78297692 | 0.50484125 | 0.77691788 | 0.86407117 | CHK1 | II | 749594 | 751177 | 1 |
| YBR275C | 190.420299 | -0.3012914 | 0.5140216 | 11.8970265 | 0.00260972 | 0.01368683 | RIF1 | II | 751356 | 757106 | -1 |
| YBR276C | 18.6933534 | 0.28051158 | 0.77582807 | 1.53621046 | 0.4638912 | 0.62711419 | PPS1 | II | 757620 | 760043 | -1 |
| YBR278W | 27.6081642 | -1.0733827 | 0.69883347 | 3.38904599 | 0.18368683 | 0.33924936 | DPB3 | II | 760294 | 760899 | 1 |
| YBR279W | 118.357209 | 1.69302309 | 0.65319051 | 7.42009695 | 0.02447634 | 0.07677031 | PAF1 | II | 761257 | 762594 | 1 |
| YBR280C | 91.3723663 | -0.4840144 | 0.55571246 | 3.55630353 | 0.16895012 | 0.31887463 | SAF1 | II | 762784 | 764697 | -1 |
| YBR281C | 17.4474364 | -1.120415 | 0.77643466 | 4.18786161 | 0.1232019 | 0.25458208 | DUG2 | II | 764970 | 767606 | -1 |
| YBR282W | 12.2737049 | 0.35132449 | 0.90781324 | 0.38386688 | 0.8253618 | 0.89541781 | MRPL27 | II | 768240 | 768680 | 1 |
| YBR283C | 7110.7187 | 6.00673841 | 0.5625591 | 90.4952702 | 2.23E-20 | 6.81E-18 | SSH1 | II | 768943 | 770415 | -1 |
| YBR284W | 22.2544745 | 2.5728956 | 0.87529959 | 12.8058449 | 0.00165671 | 0.00977813 | NA | II | 771239 | 773632 | 1 |
| YBR285W | 50.2825246 | -0.7066559 | 0.57952776 | 1.48370509 | 0.47623086 | 0.6387252 | NA | II | 773922 | 774356 | 1 |
| YBR286W | 144.470741 | 0.69686966 | 0.55632717 | 1.61853675 | 0.44518365 | 0.61297083 | APE3 | II | 774700 | 776313 | 1 |

| | | | | | | | | | | | |
|---|---|---|---|---|---|---|---|---|---|---|---|
| YBR287W | 126.330136 | -0.0301798 | 0.54534095 | 0.00492577 | 0.99754014 | 0.99771246 | NA | II | 776571 | 777854 | 1 |
| YBR288C | 2038.12009 | 6.04731525 | 0.50482357 | 142.746095 | 1.01E-31 | 5.83E-28 | APM3 | II | 778012 | 779463 | -1 |
| YBR289W | 55.4184523 | 2.20004103 | 0.83679775 | 6.55794807 | 0.03766688 | 0.1063341 | SNF5 | II | 779667 | 782384 | 1 |
| YBR290W | 86.8124611 | 1.46611336 | 0.69276692 | 4.31489224 | 0.11562002 | 0.242911 | BSD2 | II | 782591 | 783556 | 1 |
| YBR291C | 380.561205 | 2.14641758 | 0.62767781 | 10.9026714 | 0.00429057 | 0.02021351 | CTP1 | II | 783673 | 784572 | -1 |
| YBR292C | 57.6266749 | 1.25403282 | 0.75367785 | 7.24351027 | 0.02673571 | 0.08151646 | NA | II | 784702 | 785073 | -1 |
| YBR293W | 17.4933375 | 0.77982143 | 0.76677043 | 1.02037014 | 0.60038446 | 0.74682758 | VBA2 | II | 787006 | 788430 | 1 |
| YBR294W | 5455.36514 | 1.16471189 | 0.49963742 | 9.22697004 | 0.0099172 | 0.03919493 | SUL1 | II | 789235 | 791814 | 1 |
| YBR295W | 42.1290992 | -0.3683113 | 0.74044821 | 0.32388373 | 0.85049065 | 0.91140863 | PCA1 | II | 792849 | 796499 | 1 |
| YBR296C | 57.7098009 | 0.97976452 | 0.6254897 | 2.79186475 | 0.24760207 | 0.41832973 | PHO89 | II | 796798 | 798522 | -1 |
| YBR296C-A | 10.4001111 | -1.472036 | 0.92915194 | 5.83686869 | 0.0540182 | 0.13854305 | NA | II | 800123 | 800242 | -1 |
| YBR297W | 114.478453 | 0.6392798 | 0.65046842 | 58.3610555 | 2.12E-13 | 1.66E-11 | MAL33 | II | 800523 | 801929 | 1 |
| YBR298C | 21.3626856 | 0.55025113 | 0.87817968 | 6.8413479 | 0.0326904 | 0.09540191 | MAL31 | II | 802631 | 804475 | -1 |
| YBR299W | 55.577594 | -2.1591847 | 0.681675 | 14.0584599 | 0.00088561 | 0.00595552 | MAL32 | II | 805351 | 807105 | 1 |
| YBR302C | 8.53829875 | -0.8688969 | 1.07705907 | 3.75297392 | 0.1531271 | 0.29709307 | COS3 | II | 810340 | 811479 | -1 |
| YCL001W | 15.8275698 | -0.3481354 | 0.80103377 | 0.65967013 | 0.71904232 | 0.82938751 | RER1 | III | 111916 | 112482 | 1 |
| YCL001W-A | 106.218471 | 0.36101532 | 0.52161367 | 0.73062531 | 0.69397964 | 0.81470846 | NA | III | 113080 | 113541 | 1 |
| YCL001W-B | 168.048844 | 0.2211977 | 0.55349669 | 0.28441957 | 0.86743926 | 0.9222316 | NA | III | 113770 | 114024 | 1 |
| YCL002C | 601.128917 | 0.26843569 | 0.5087035 | 0.38399118 | 0.82531051 | 0.89541781 | NA | III | 110808 | 111675 | -1 |
| YCL004W | 341.831723 | -0.0736242 | 0.55278292 | 5.78563517 | 0.05541984 | 0.14137538 | PGS1 | III | 109105 | 110670 | 1 |
| YCL005W | 82.7667111 | -0.3433608 | 0.63262591 | 1.8475881 | 0.3970099 | 0.56785755 | LDB16 | III | 108021 | 108791 | 1 |
| YCL005W-A | 50.6195267 | -0.1619361 | 0.78436475 | 2.85276169 | 0.24017659 | 0.40828609 | VMA9 | III | 107023 | 107417 | 1 |
| YCL008C | 23.6180039 | -0.0887596 | 0.70914442 | 0.39072543 | 0.82253626 | 0.89359303 | STP22 | III | 105696 | 106853 | -1 |
| YCL009C | 247.105573 | -0.8564652 | 0.52465052 | 3.81600214 | 0.14837668 | 0.29072792 | ILV6 | III | 104619 | 105548 | -1 |
| YCL010C | 148.415847 | -1.7449467 | 0.56339483 | 21.1271333 | 2.58E-05 | 0.00029686 | SGF29 | III | 103571 | 104350 | -1 |
| YCL011C | 59.4034616 | -0.2374846 | 0.62921751 | 2.55271429 | 0.279052 | 0.45346929 | GBP2 | III | 102075 | 103358 | -1 |
| YCL012C | 62.5856088 | 2.04618965 | 0.74718023 | 7.35889409 | 0.02523693 | 0.07826556 | NA | III | 101317 | 101788 | -1 |
| YCL014W | 192.828109 | 0.82484049 | 0.59451962 | 1.94380222 | 0.37836304 | 0.55223646 | BUD3 | III | 96281 | 101191 | 1 |
| YCL016C | 10.5722747 | 0.14154639 | 0.94828311 | 0.25606986 | 0.87982265 | 0.92911891 | DCC1 | III | 94621 | 95763 | -1 |
| YCL017C | 253.865876 | -1.1022558 | 0.62250036 | 7.3695392 | 0.02510296 | 0.07810109 | NFS1 | III | 92777 | 94270 | -1 |
| YCL018W | 219.963722 | -0.9094024 | 0.50568787 | 12.2830307 | 0.00215166 | 0.01180864 | LEU2 | III | 91324 | 92418 | 1 |
| YCL021W-A | 22.0388119 | 2.15084377 | 0.78516363 | 7.99185174 | 0.01839041 | 0.06210569 | NA | III | 83620 | 83997 | 1 |
| YCL024W | 44.3843966 | 0.01260664 | 0.6072401 | 0.5023339 | 0.77789249 | 0.86498896 | KCC4 | III | 79162 | 82275 | 1 |

| | | | | | | | | | | | |
|---|---|---|---|---|---|---|---|---|---|---|---|
| YCL025C | 299.516182 | 0.68603252 | 0.50470219 | 13.5578054 | 0.00113752 | 0.00731806 | AGP1 | III | 76018 | 77919 | -1 |
| YCL026C-A | 27.3285305 | -2.5480697 | 0.75977892 | 17.8665274 | 0.00013193 | 0.00118796 | FRM2 | III | 74704 | 75285 | -1 |
| YCL026C-B | 11.3282441 | -0.4119627 | 0.97418786 | 3.92723822 | 0.14034956 | 0.27886889 | HBN1 | III | 73405 | 73986 | -1 |
| YCL027W | 77.7021553 | -0.9952344 | 0.59308074 | 9.92503456 | 0.0069953 | 0.02997984 | FUS1 | III | 71803 | 73341 | 1 |
| YCL028W | 126.493519 | 0.92103958 | 0.63303908 | 2.93032744 | 0.23104016 | 0.39792139 | RNQ1 | III | 70150 | 71367 | 1 |
| YCL029C | 26.6042279 | 0.70344566 | 0.73441377 | 0.9412615 | 0.62460817 | 0.76345394 | BIK1 | III | 68599 | 69921 | -1 |
| YCL030C | 63.7154504 | 1.30644051 | 0.624066 | 11.5910938 | 0.00304107 | 0.01538156 | HIS4 | III | 65934 | 68333 | -1 |
| YCL031C | 47.2983454 | 0.00019462 | 0.64439476 | 11.6534005 | 0.00294779 | 0.01503761 | RRP7 | III | 64675 | 65568 | -1 |
| YCL032W | 18.9617726 | 0.80103872 | 0.76601143 | 1.83051739 | 0.40041303 | 0.57131381 | STE50 | III | 63441 | 64481 | 1 |
| YCL033C | 107.919393 | 2.5568211 | 0.78114147 | 10.6337186 | 0.00490814 | 0.02256764 | MXR2 | III | 62776 | 63282 | -1 |
| YCL034W | 304.665661 | 2.60085374 | 0.81610982 | 9.21364627 | 0.00998348 | 0.03943 | LSB5 | III | 61658 | 62722 | 1 |
| YCL035C | 60.927361 | 0.18802941 | 0.55857594 | 0.35715833 | 0.83645784 | 0.90356173 | GRX1 | III | 60841 | 61173 | -1 |
| YCL036W | 150.413354 | 2.08008317 | 0.8223951 | 6.77059602 | 0.03386755 | 0.09820123 | GFD2 | III | 59026 | 60726 | 1 |
| YCL037C | 236.282174 | 2.91438616 | 0.58685522 | 29.1538847 | 4.67E-07 | 9.10E-06 | SRO9 | III | 57374 | 58678 | -1 |
| YCL038C | 162.787189 | 0.06681231 | 0.5540427 | 0.23491922 | 0.88917642 | 0.93304075 | ATG22 | III | 54941 | 56527 | -1 |
| YCL039W | 284.465486 | 0.24861322 | 0.77347935 | 1.71708384 | 0.42377954 | 0.59440008 | GID7 | III | 52645 | 54882 | 1 |
| YCL040W | 114.376159 | 2.08284531 | 0.56631711 | 15.127188 | 0.00051901 | 0.00378167 | GLK1 | III | 50838 | 52340 | 1 |
| YCL042W | 16.8862491 | 1.15129321 | 0.83485893 | 3.80165705 | 0.14944475 | 0.29203007 | NA | III | 50584 | 50943 | 1 |
| YCL043C | 294.126278 | 1.59285572 | 0.53737728 | 8.44110428 | 0.01469053 | 0.05261147 | PDI1 | III | 48653 | 50221 | -1 |
| YCL044C | 93.5532419 | -0.3444877 | 0.59098927 | 1.2357752 | 0.53908199 | 0.69331069 | MGR1 | III | 47111 | 48364 | -1 |
| YCL045C | 78.2732494 | -0.1263894 | 0.63972998 | 2.32244932 | 0.3131025 | 0.48516903 | EMC1 | III | 44623 | 46905 | -1 |
| YCL047C | 23.286016 | 0.11900208 | 0.76827804 | 0.57677301 | 0.74947186 | 0.8479727 | POF1 | III | 43661 | 44437 | -1 |
| YCL048W | 101.085519 | 2.22193574 | 0.72850037 | 9.46582969 | 0.00880078 | 0.03573389 | SPS22 | III | 42165 | 43556 | 1 |
| YCL048W-A | 37.7860573 | 2.61625181 | 0.89174642 | 11.2433619 | 0.00361855 | 0.01778559 | NA | III | 41488 | 41727 | 1 |
| YCL049C | 393.739341 | -0.8060418 | 0.51683095 | 2.54201377 | 0.280549 | 0.45466569 | NA | III | 39786 | 40724 | -1 |
| YCL050C | 51.8163578 | -0.3404369 | 0.59009177 | 0.43961437 | 0.80267355 | 0.881373 | APA1 | III | 37836 | 38801 | -1 |
| YCL051W | 149.050248 | 1.24913906 | 0.7207694 | 61.4237848 | 4.59E-14 | 4.03E-12 | LRE1 | III | 35865 | 37616 | 1 |
| YCL052C | 17.4005338 | 0.42060739 | 1.08265026 | 0.27399286 | 0.87197334 | 0.92527295 | PBN1 | III | 34143 | 35393 | -1 |
| YCL054W | 106.061843 | 1.06077341 | 0.86013861 | 3.51460045 | 0.17250997 | 0.32379018 | SPB1 | III | 31449 | 33974 | 1 |
| YCL055W | 7.10275474 | -1.2020633 | 1.04931894 | 2.2663983 | 0.32200148 | 0.49479526 | KAR4 | III | 27929 | 28936 | 1 |
| YCL056C | 794.042805 | 0.94079777 | 0.5149651 | 4.06638447 | 0.13091694 | 0.26590071 | PEX34 | III | 26925 | 27359 | -1 |
| YCL057C-A | 93.0929797 | -0.4203614 | 0.53038709 | 2.00439056 | 0.36707273 | 0.54025193 | MIC10 | III | 24032 | 24325 | -1 |
| YCL057W | 79.8728449 | 0.53037637 | 0.59573611 | 3.33785951 | 0.18844864 | 0.34420115 | PRD1 | III | 24768 | 26906 | 1 |

| | | | | | | | | | | | |
|---|---|---|---|---|---|---|---|---|---|---|---|
| YCL058C | 930.608544 | -1.4760959 | 0.50078424 | 28.4913856 | 6.50E-07 | 1.22E-05 | FYV5 | III | 23523 | 23981 | -1 |
| YCL058W-A | 723.627416 | 0.68507296 | 0.66343852 | 4.4659781 | 0.1072075 | 0.23024163 | ADF1 | III | 23584 | 23925 | 1 |
| YCL059C | 28.9969124 | -1.2328547 | 0.73162515 | 10.3439488 | 0.00567336 | 0.02535586 | KRR1 | III | 22429 | 23379 | -1 |
| YCL061C | 33.0284699 | -0.593654 | 0.68235389 | 2.39640059 | 0.30173676 | 0.47448396 | MRC1 | III | 18816 | 22106 | -1 |
| YCL063W | 436.06417 | -1.2865816 | 0.49301729 | 20.2877105 | 3.93E-05 | 0.0004228 | VAC17 | III | 17290 | 18561 | 1 |
| YCL064C | 123.37118 | -0.3237428 | 1.00647748 | 4.13595064 | 0.12644153 | 0.25966788 | CHA1 | III | 15798 | 16880 | -1 |
| YCL068C | 23.7561479 | 1.26585894 | 1.00405414 | 1.87739715 | 0.39113654 | 0.56237411 | NA | III | 11503 | 12285 | -1 |
| YCR001W | 336.656574 | 0.48154804 | 0.49659858 | 15.2329575 | 0.00049227 | 0.00363486 | NA | III | 115685 | 115999 | 1 |
| YCR002C | 15.7574711 | 0.86344092 | 0.93262284 | 1.01921125 | 0.60073245 | 0.74682758 | CDC10 | III | 117380 | 118348 | -1 |
| YCR003W | 18.3477302 | -1.2657777 | 0.78385289 | 2.89482966 | 0.23517748 | 0.40286319 | MRPL32 | III | 118620 | 119171 | 1 |
| YCR004C | 34.0747482 | 0.06080033 | 0.64345919 | 0.40343356 | 0.81732638 | 0.89020311 | YCP4 | III | 119575 | 120318 | -1 |
| YCR005C | 37.8609467 | 1.27181354 | 0.7733525 | 4.32972804 | 0.11476554 | 0.24172153 | CIT2 | III | 120946 | 122328 | -1 |
| YCR007C | 311.256928 | 0.97352214 | 0.56941396 | 4.06741363 | 0.13084959 | 0.26590071 | NA | III | 126011 | 126730 | -1 |
| YCR008W | 49.0829016 | -0.610598 | 0.86112271 | 0.54046554 | 0.76320182 | 0.85588583 | SAT4 | III | 128470 | 130281 | 1 |
| YCR009C | 67.6107321 | -0.0380109 | 0.65630495 | 3.94490681 | 0.13911513 | 0.27774112 | RVS161 | III | 130745 | 131542 | -1 |
| YCR010C | 170.08003 | 0.59228849 | 0.71539342 | 2.43569815 | 0.29586587 | 0.46995614 | ADY2 | III | 132273 | 133124 | -1 |
| YCR011C | 60.8555432 | 0.30600674 | 0.63651549 | 2.12237461 | 0.34604471 | 0.51947079 | ADP1 | III | 133724 | 136873 | -1 |
| YCR012W | 336.293428 | -0.3467679 | 0.70613247 | 4.66002125 | 0.09729471 | 0.21492841 | PGK1 | III | 137746 | 138996 | 1 |
| YCR014C | 44.2700354 | 0.48131492 | 0.60400384 | 0.8475861 | 0.65455933 | 0.78442961 | POL4 | III | 139185 | 140933 | -1 |
| YCR015C | 29.1308523 | -0.3813026 | 0.70928394 | 9.71719786 | 0.00776135 | 0.03237624 | CTO1 | III | 141217 | 142170 | -1 |
| YCR016W | 54.3935952 | -1.2916142 | 0.58153759 | 13.7483025 | 0.00103418 | 0.00679008 | NA | III | 143634 | 144506 | 1 |
| YCR017C | 34.9736628 | 0.02747737 | 0.89380125 | 0.06638544 | 0.96735211 | 0.97486964 | CWH43 | III | 144773 | 147634 | -1 |
| YCR018C | 229.212885 | -0.6833051 | 0.53856947 | 1.97104433 | 0.37324428 | 0.54640641 | SRD1 | III | 148238 | 148903 | -1 |
| YCR019W | 22.8264526 | 0.84517046 | 0.84058048 | 1.19996566 | 0.54882106 | 0.70069987 | MAK32 | III | 152837 | 153928 | 1 |
| YCR020C | 52.9609565 | 0.60092799 | 0.63744941 | 4.99462466 | 0.08230591 | 0.19005499 | PET18 | III | 154012 | 154659 | -1 |
| YCR020C-A | 10.0172298 | 0.5299868 | 0.99375154 | 1.37661848 | 0.50242483 | 0.66093664 | MAK31 | III | 154830 | 155096 | -1 |
| YCR020W-B | 11.8240907 | -0.6665777 | 1.10709502 | 6.91106627 | 0.03157047 | 0.09292985 | HTL1 | III | 155320 | 155556 | 1 |
| YCR021C | 334.952806 | 0.60472687 | 0.61373996 | 2.96486165 | 0.22708501 | 0.39307092 | HSP30 | III | 156109 | 157107 | -1 |
| YCR022C | 16.7210659 | -0.720714 | 0.80395192 | 1.12283461 | 0.57040006 | 0.72078052 | NA | III | 157421 | 157765 | -1 |
| YCR023C | 244.036595 | -0.9361765 | 0.51500776 | 3.55612731 | 0.168965 | 0.31887463 | NA | III | 158538 | 160373 | -1 |
| YCR024C | 100.911755 | -0.5796554 | 0.55494331 | 1.35888884 | 0.50689854 | 0.66464916 | SLM5 | III | 160744 | 162222 | -1 |
| YCR024C-A | 315.274774 | 2.00423432 | 0.63183717 | 18.7168757 | 8.62E-05 | 0.00082519 | PMP1 | III | 162945 | 163067 | -1 |
| YCR024C-B | 352.461296 | 2.88007272 | 0.64355735 | 26.159938 | 2.09E-06 | 3.35E-05 | NA | III | 162599 | 162865 | -1 |

| | | | | | | | | | | | |
|---|---|---|---|---|---|---|---|---|---|---|---|
| YCR025C | 290.595223 | 1.72696293 | 0.55675681 | 23.0053192 | 1.01E-05 | 0.00013356 | NA | III | 163446 | 163856 | -1 |
| YCR026C | 53.3210893 | -0.3487096 | 0.59451432 | 1.14726061 | 0.56347613 | 0.71468276 | NPP1 | III | 164111 | 166339 | -1 |
| YCR027C | 6.03799162 | -0.0435358 | 1.40130822 | 0.85716458 | 0.65143198 | 0.78236902 | RHB1 | III | 167370 | 167999 | -1 |
| YCR028C | 50.9138423 | -0.2647148 | 0.61055711 | 0.66741426 | 0.71626353 | 0.82794286 | FEN2 | III | 170886 | 172424 | -1 |
| YCR028C-A | 9.43023506 | -1.0798832 | 0.94635211 | 2.74098032 | 0.25398244 | 0.42600183 | RIM1 | III | 172950 | 173440 | -1 |
| YCR030C | 120.314438 | -1.2367976 | 0.51503887 | 8.85776568 | 0.01192781 | 0.04496224 | SYP1 | III | 173826 | 176438 | -1 |
| YCR031C | 73.7572058 | -0.868169 | 0.84142946 | 12.3206157 | 0.0021116 | 0.01164923 | RPS14A | III | 177500 | 178220 | -1 |
| YCR032W | 97.9721179 | 0.66362987 | 0.55940514 | 21.0206423 | 2.73E-05 | 0.00031124 | BPH1 | III | 179520 | 186023 | 1 |
| YCR033W | 98.9295984 | 1.05336683 | 0.57281475 | 6.31538478 | 0.04252376 | 0.1160873 | SNT1 | III | 186489 | 190169 | 1 |
| YCR034W | 52.9589909 | 0.22624481 | 0.58632787 | 1.20738814 | 0.54678803 | 0.6993379 | ELO2 | III | 190592 | 191635 | 1 |
| YCR035C | 94.8994726 | -0.0908735 | 0.52349824 | 12.6451571 | 0.00179531 | 0.01032258 | RRP43 | III | 191834 | 193018 | -1 |
| YCR036W | 70.474612 | 0.54307096 | 0.65034388 | 5.75136145 | 0.05637775 | 0.14342143 | RBK1 | III | 193297 | 194298 | 1 |
| YCR037C | 98.8286519 | 1.18426338 | 0.59930589 | 3.82132717 | 0.14798216 | 0.2902937 | PHO87 | III | 194414 | 197185 | -1 |
| YCR041W | 758.157492 | 1.0008431 | 0.59432752 | 4.61587992 | 0.09946594 | 0.21864382 | NA | III | 200911 | 201243 | 1 |
| YCR042C | 543.662284 | -0.7896277 | 0.59363246 | 10.2970603 | 0.00580794 | 0.02584777 | TAF2 | III | 201174 | 205397 | -1 |
| YCR043C | 7.23865365 | -1.6820868 | 1.24980574 | 4.91440003 | 0.0856745 | 0.19653541 | NA | III | 206261 | 206644 | -1 |
| YCR044C | 280.106854 | -1.8824147 | 0.51072444 | 26.2510662 | 1.99E-06 | 3.24E-05 | PER1 | III | 206877 | 207950 | -1 |
| YCR045C | 1395.06948 | -1.2294791 | 0.58343031 | 9.93040502 | 0.00697654 | 0.02994378 | RRT12 | III | 208135 | 209610 | -1 |
| YCR046C | 132.652168 | -0.4983028 | 0.52683455 | 7.95256421 | 0.01875524 | 0.06302544 | IMG1 | III | 209914 | 210423 | -1 |
| YCR047C | 63.2036226 | -0.2409422 | 0.66183292 | 0.68849199 | 0.70875456 | 0.82407349 | BUD23 | III | 210718 | 211545 | -1 |
| YCR048W | 21.713628 | 0.18839981 | 0.7567659 | 0.48086191 | 0.78628893 | 0.87021248 | ARE1 | III | 211929 | 213761 | 1 |
| YCR050C | 2212.76119 | -0.0165935 | 0.46897776 | 2.42760413 | 0.29706567 | 0.4709776 | NA | III | 213464 | 213772 | -1 |
| YCR051W | 146.173156 | 0.21957945 | 0.53007866 | 1.30930675 | 0.51962215 | 0.67578891 | NA | III | 214071 | 214739 | 1 |
| YCR052W | 36.2201719 | -0.199329 | 0.66231238 | 0.22871397 | 0.89193949 | 0.93370632 | RSC6 | III | 214994 | 216445 | 1 |
| YCR053W | 79.7704856 | -0.0223061 | 0.54893134 | 4.59895721 | 0.10031113 | 0.21944212 | THR4 | III | 216697 | 218241 | 1 |
| YCR054C | 34.9866406 | 0.61726686 | 0.81520913 | 4.96557022 | 0.08351032 | 0.19225635 | CTR86 | III | 218376 | 220067 | -1 |
| YCR057C | 27.9095756 | -1.0689797 | 0.69343099 | 6.44673536 | 0.03982073 | 0.11116077 | PWP2 | III | 220457 | 223228 | -1 |
| YCR059C | 148.648024 | -2.2085584 | 0.53774644 | 35.5672339 | 1.89E-08 | 5.26E-07 | YIH1 | III | 223454 | 224230 | -1 |
| YCR060W | 136.948626 | -0.6224786 | 0.59941993 | 14.5463924 | 0.00069389 | 0.00484051 | TAH1 | III | 224399 | 224734 | 1 |
| YCR061W | 1626.24945 | 1.50977555 | 0.51987048 | 53.7826072 | 2.10E-12 | 1.39E-10 | NA | III | 225563 | 227458 | 1 |
| YCR063W | 19.0498706 | -1.7904169 | 0.83695602 | 4.97666213 | 0.08304845 | 0.19134522 | BUD31 | III | 228318 | 228791 | 1 |
| YCR065W | 40.1115991 | -0.0206116 | 0.61122592 | 0.12171128 | 0.94095906 | 0.96087354 | HCM1 | III | 229310 | 231004 | 1 |
| YCR066W | 19.2850513 | -0.4218269 | 0.77563681 | 1.0925808 | 0.57909404 | 0.72795364 | RAD18 | III | 231500 | 232963 | 1 |

| | | | | | | | | | | | |
|---|---|---|---|---|---|---|---|---|---|---|---|
| YCR067C | 59.0079036 | 0.92739333 | 0.74649245 | 1.62788312 | 0.44310808 | 0.61162622 | SED4 | III | 233125 | 236322 | -1 |
| YCR068W | 17.8603771 | -0.4839523 | 0.85951193 | 0.71077454 | 0.70090195 | 0.81801018 | ATG15 | III | 237214 | 238776 | 1 |
| YCR069W | 26.2965861 | 0.25794316 | 0.71369047 | 1.05431342 | 0.59028092 | 0.73765326 | CPR4 | III | 239055 | 240011 | 1 |
| YCR071C | 608.495607 | 3.77353252 | 1.60699532 | 4.86100758 | 0.08799249 | 0.20050237 | IMG2 | III | 240103 | 240543 | -1 |
| YCR072C | 16.3417877 | -0.5345554 | 0.81524086 | 0.49751597 | 0.77976867 | 0.86640963 | RSA4 | III | 240805 | 242352 | -1 |
| YCR073C | 1117.23221 | -1.1008619 | 0.48449942 | 23.7522924 | 6.95E-06 | 9.52E-05 | SSK22 | III | 242588 | 246583 | -1 |
| YCR073W-A | 46.5795931 | 1.76003603 | 0.67882425 | 7.95875781 | 0.01869725 | 0.06286705 | SOL2 | III | 246963 | 247910 | 1 |
| YCR075C | 17.9377281 | 0.46821121 | 0.83598586 | 2.33093777 | 0.31177644 | 0.48409375 | ERS1 | III | 248033 | 248815 | -1 |
| YCR075W-A | 53.7519531 | 2.30870342 | 0.91327036 | 6.45881802 | 0.03958088 | 0.11076526 | EGO2 | III | 248975 | 249202 | 1 |
| YCR076C | 73.6517941 | 0.4047486 | 0.690451 | 0.54170441 | 0.76272921 | 0.85565759 | FUB1 | III | 249293 | 250045 | -1 |
| YCR077C | 19.1557254 | -0.0563713 | 0.83610034 | 2.28950833 | 0.31830216 | 0.49106568 | PAT1 | III | 250238 | 252628 | -1 |
| YCR079W | 99.654938 | -1.4496094 | 0.54859457 | 7.40262239 | 0.02469113 | 0.07709579 | PTC6 | III | 252845 | 254173 | 1 |
| YCR081W | 22.9218269 | -0.8730971 | 0.80170606 | 3.70444777 | 0.15688788 | 0.30265354 | SRB8 | III | 254371 | 258654 | 1 |
| YCR082W | 24.091331 | 0.99784265 | 0.82615437 | 1.48642994 | 0.47558247 | 0.63800336 | AHC2 | III | 258883 | 259269 | 1 |
| YCR083W | 26.5801793 | -0.3863683 | 0.74967245 | 1.3879461 | 0.49958724 | 0.65905903 | TRX3 | III | 259578 | 259961 | 1 |
| YCR084C | 113.174446 | 0.45337359 | 0.61983141 | 2.27780703 | 0.32016989 | 0.49276546 | TUP1 | III | 260311 | 262452 | -1 |
| YCR085W | 26.9306015 | -1.5092229 | 0.731732 | 18.5370214 | 9.43E-05 | 0.00088971 | NA | III | 262916 | 263269 | 1 |
| YCR086W | 118.76426 | 0.31262002 | 0.7625838 | 12.0848647 | 0.00237577 | 0.012725 | CSM1 | III | 263392 | 263964 | 1 |
| YCR087C-A | 57.1827257 | 0.1484146 | 0.73428038 | 3.91309119 | 0.14134584 | 0.28046348 | NA | III | 264006 | 264467 | -1 |
| YCR088W | 154.335812 | -0.1728734 | 0.55091599 | 0.47966806 | 0.78675843 | 0.8703346 | ABP1 | III | 265068 | 266846 | 1 |
| YCR089W | 179.551968 | 0.65684169 | 0.64734182 | 6.92840266 | 0.03129799 | 0.09240968 | FIG2 | III | 267434 | 272263 | 1 |
| YCR090C | 35.809202 | 2.82940861 | 1.03674744 | 9.40659015 | 0.00906536 | 0.03655182 | NA | III | 272315 | 272863 | -1 |
| YCR091W | 17.1212582 | 0.67413614 | 0.77988956 | 2.89869469 | 0.23472343 | 0.40225741 | KIN82 | III | 274404 | 276566 | 1 |
| YCR092C | 258.811154 | 0.99088054 | 0.66257298 | 2.98800691 | 0.22447219 | 0.38994719 | MSH3 | III | 276764 | 279820 | -1 |
| YCR093W | 66.601645 | 0.57911397 | 0.64395794 | 2.90854637 | 0.23357006 | 0.40094002 | CDC39 | III | 280117 | 286443 | 1 |
| YCR094W | 30.0038587 | -0.5722363 | 0.71314685 | 8.89042321 | 0.01173462 | 0.04440749 | CDC50 | III | 286762 | 287937 | 1 |
| YCR095C | 229.922913 | 1.41800789 | 0.72320982 | 7.67034847 | 0.02159758 | 0.07017394 | OCA4 | III | 288170 | 289258 | -1 |
| YCR095W-A | 16.1665781 | -1.2227639 | 0.83879035 | 2.19467386 | 0.33375872 | 0.50787464 | NA | III | 289639 | 289797 | 1 |
| YCR098C | 46.8822512 | 0.03588118 | 0.67718942 | 3.50643169 | 0.17321601 | 0.32488523 | GIT1 | III | 297049 | 298605 | -1 |
| YCR099C | 11.9341417 | 1.55594066 | 1.11412465 | 6.7134447 | 0.0348493 | 0.10013768 | NA | III | 300832 | 301299 | -1 |
| YCR100C | 6.73706912 | 1.3244941 | 1.52117984 | 7.46233843 | 0.0239648 | 0.07565017 | NA | III | 301271 | 302221 | -1 |
| YCR101C | 11.1913099 | 1.20257973 | 1.17005459 | 7.43089371 | 0.02434456 | 0.07668934 | NA | III | 302482 | 303030 | -1 |
| YCR102C | 26.2651998 | -0.1298151 | 0.69762022 | 1.55117522 | 0.46043315 | 0.62448065 | NA | III | 304361 | 305467 | -1 |

| | | | | | | | | | | | |
|---|---|---|---|---|---|---|---|---|---|---|---|
| YCR105W | 1389.81288 | 0.3090443 | 0.59138994 | 0.54766291 | 0.76046024 | 0.85479805 | ADH7 | III | 309070 | 310155 | 1 |
| YCR106W | 980.370505 | -1.7457206 | 0.52070062 | 39.2477513 | 3.00E-09 | 1.01E-07 | RDS1 | III | 310958 | 313456 | 1 |
| YCR107W | 338.190843 | -1.8251475 | 0.51427348 | 43.0456979 | 4.50E-10 | 1.77E-08 | AAD3 | III | 313890 | 314981 | 1 |
| YDL001W | 14.2188184 | -0.1927894 | 0.91470017 | 0.8191032 | 0.6639479 | 0.79099966 | RMD1 | IV | 447984 | 449276 | 1 |
| YDL002C | 29.0599948 | 0.1223469 | 0.67203508 | 0.62485025 | 0.73167041 | 0.83702784 | NHP10 | IV | 446967 | 447578 | -1 |
| YDL003W | 108.1267 | 0.19833644 | 0.56055169 | 12.9996249 | 0.00150372 | 0.00913878 | MCD1 | IV | 444683 | 446383 | 1 |
| YDL004W | 8.04822654 | 0.04187082 | 1.15684786 | 1.36411371 | 0.50557603 | 0.66363301 | ATP16 | IV | 443029 | 443511 | 1 |
| YDL005C | 278.661671 | 1.54918165 | 0.78906048 | 3.72930557 | 0.15495 | 0.29975293 | MED2 | IV | 441014 | 442309 | -1 |
| YDL006W | 76.5440348 | 0.27292716 | 0.70224413 | 1.34590438 | 0.51020014 | 0.66786132 | PTC1 | IV | 439909 | 440754 | 1 |
| YDL007W | 75.4203554 | 0.67295091 | 0.79787249 | 0.82702098 | 0.6613246 | 0.78897407 | RPT2 | IV | 438047 | 439360 | 1 |
| YDL008W | 207.013162 | -0.6020038 | 0.5412878 | 8.08143127 | 0.01758488 | 0.05999792 | APC11 | IV | 433497 | 433994 | 1 |
| YDL009C | 3549.89711 | 1.3827344 | 0.50766126 | 9.5357725 | 0.00849832 | 0.03482328 | NA | IV | 432925 | 433248 | -1 |
| YDL010W | 25.7091349 | 0.1222743 | 0.86472216 | 1.14244976 | 0.56483316 | 0.71577676 | GRX6 | IV | 432330 | 433025 | 1 |
| YDL012C | 50.3197723 | 0.63889658 | 0.68402231 | 1.76417171 | 0.41391864 | 0.58553357 | NA | IV | 431108 | 431517 | -1 |
| YDL013W | 93.2697411 | 0.80618033 | 0.57284573 | 2.0404844 | 0.36050761 | 0.53345156 | SLX5 | IV | 429067 | 430926 | 1 |
| YDL014W | 120.928798 | -1.1904262 | 0.81049832 | 8.80739519 | 0.01223203 | 0.04578115 | NOP1 | IV | 427364 | 428347 | 1 |
| YDL015C | 53.0464213 | -0.5999737 | 0.62576179 | 1.41110459 | 0.49383576 | 0.65469026 | TSC13 | IV | 426002 | 426934 | -1 |
| YDL017W | 8.0008098 | 0.23604652 | 1.08910656 | 0.62884198 | 0.73021155 | 0.83678999 | CDC7 | IV | 424209 | 425732 | 1 |
| YDL018C | 13.9249638 | -0.1092027 | 0.84954951 | 1.85479018 | 0.39558283 | 0.56679648 | ERP3 | IV | 422833 | 423510 | -1 |
| YDL019C | 44.3007987 | 0.36780437 | 0.6080791 | 0.85333012 | 0.65268213 | 0.78305626 | OSH2 | IV | 417663 | 421514 | -1 |
| YDL020C | 36.3343799 | 0.64492583 | 0.78045518 | 2.32167902 | 0.31322312 | 0.48516903 | RPN4 | IV | 415113 | 416708 | -1 |
| YDL021W | 41.7212904 | 0.35198657 | 0.67558281 | 5.04643784 | 0.08020103 | 0.18675877 | GPM2 | IV | 413953 | 414888 | 1 |
| YDL022W | 126.041901 | 2.08864074 | 0.66765142 | 11.932142 | 0.0025643 | 0.01350981 | GPD1 | IV | 411825 | 413000 | 1 |
| YDL024C | 10.0062406 | 0.2958574 | 1.00765481 | 0.1636896 | 0.92141495 | 0.95035295 | DIA3 | IV | 408451 | 409857 | -1 |
| YDL025C | 73.4476898 | -1.7485557 | 0.56929051 | 13.638276 | 0.00109266 | 0.00709251 | RTK1 | IV | 405344 | 407206 | -1 |
| YDL027C | 46.3521828 | 0.26988193 | 0.63969895 | 3.90945528 | 0.14160304 | 0.28078137 | MRX9 | IV | 403695 | 404957 | -1 |
| YDL028C | 19.6224839 | -0.2551699 | 0.75976339 | 0.13710473 | 0.93374456 | 0.95735444 | MPS1 | IV | 400997 | 403291 | -1 |
| YDL029W | 83.7846692 | 0.34270166 | 0.60735738 | 10.9232057 | 0.00424674 | 0.02008876 | ARP2 | IV | 399340 | 400638 | 1 |
| YDL030W | 74.7417788 | 0.43598567 | 0.57300905 | 6.73335497 | 0.03450409 | 0.09954094 | PRP9 | IV | 397537 | 399129 | 1 |
| YDL031W | 49.8269737 | -1.4988969 | 0.58436501 | 10.9613855 | 0.00416644 | 0.01978975 | DBP10 | IV | 394217 | 397204 | 1 |
| YDL033C | 145.008975 | 1.10205907 | 0.52563848 | 31.9266112 | 1.17E-07 | 2.76E-06 | SLM3 | IV | 392659 | 393912 | -1 |
| YDL035C | 154.996064 | -0.391995 | 0.55169323 | 1.28295167 | 0.5265148 | 0.68176637 | GPR1 | IV | 389172 | 392057 | -1 |
| YDL036C | 777.878867 | 0.04629096 | 0.49526735 | 4.14198718 | 0.12606047 | 0.25919393 | PUS9 | IV | 387513 | 388901 | -1 |

| | | | | | | | | | | | |
|---|---|---|---|---|---|---|---|---|---|---|---|
| YDL037C | 13.4495658 | -0.581446 | 1.01592133 | 6.44796779 | 0.0397962 | 0.11116077 | BSC1 | IV | 384601 | 385587 | -1 |
| YDL039C | 13.5411867 | 0.20874702 | 0.87465224 | 0.25322743 | 0.88107395 | 0.9300006 | PRM7 | IV | 381985 | 384081 | -1 |
| YDL040C | 43.2693753 | -0.0905017 | 0.62106317 | 4.07326588 | 0.13046726 | 0.26533384 | NAT1 | IV | 378874 | 381438 | -1 |
| YDL042C | 28.9576134 | -0.5324126 | 0.68967902 | 0.68489133 | 0.7100317 | 0.82438173 | SIR2 | IV | 376757 | 378445 | -1 |
| YDL043C | 9.54069089 | 0.790136 | 0.97738668 | 0.89476425 | 0.63929957 | 0.77519257 | PRP11 | IV | 375680 | 376480 | -1 |
| YDL044C | 41.8484206 | 0.57429385 | 0.6479122 | 9.00467586 | 0.01108305 | 0.04252544 | MTF2 | IV | 373967 | 375289 | -1 |
| YDL045C | 8.56391719 | 1.31346009 | 1.12285601 | 1.7253481 | 0.42203204 | 0.59293772 | FAD1 | IV | 372688 | 373608 | -1 |
| YDL045W-A | 15.1028616 | 1.00045223 | 0.90697007 | 3.11675128 | 0.21047769 | 0.37348017 | MRP10 | IV | 372248 | 372535 | 1 |
| YDL046W | 370.571418 | -0.1412141 | 0.49872018 | 5.13697657 | 0.07665133 | 0.18033776 | NPC2 | IV | 371240 | 371761 | 1 |
| YDL047W | 30.7840109 | 0.16911847 | 0.70543842 | 4.84979154 | 0.08848734 | 0.20139218 | SIT4 | IV | 369771 | 370706 | 1 |
| YDL048C | 209.675831 | 1.15944571 | 0.50929743 | 10.489619 | 0.00527483 | 0.0239164 | STP4 | IV | 366739 | 368211 | -1 |
| YDL049C | 72.0662531 | 0.50561637 | 0.55412602 | 1.47793519 | 0.47760674 | 0.64027392 | KNH1 | IV | 365068 | 365874 | -1 |
| YDL051W | 28.5085026 | -0.5415095 | 0.87603122 | 8.82074797 | 0.01215063 | 0.04558315 | LHP1 | IV | 363952 | 364779 | 1 |
| YDL052C | 27.3436663 | 0.14622947 | 0.74174913 | 0.08216472 | 0.95975008 | 0.97166515 | SLC1 | IV | 362672 | 363583 | -1 |
| YDL053C | 34.0407731 | -0.0084608 | 0.68059638 | 2.22954705 | 0.32798955 | 0.50041094 | PBP4 | IV | 361699 | 362256 | -1 |
| YDL054C | 45.1572706 | 1.17584833 | 0.68866716 | 6.98191132 | 0.03047174 | 0.09033864 | MCH1 | IV | 359825 | 361285 | -1 |
| YDL055C | 106.324777 | -0.0560703 | 0.87950853 | 2.19134155 | 0.33431528 | 0.50802806 | PSA1 | IV | 355674 | 356759 | -1 |
| YDL056W | 48.5245524 | 2.02832267 | 0.81467526 | 6.32169689 | 0.04238976 | 0.11597295 | MBP1 | IV | 352877 | 355378 | 1 |
| YDL057W | 419.537399 | 5.13975922 | 0.69652532 | 122.302566 | 2.77E-27 | 5.34E-24 | NA | IV | 351434 | 352420 | 1 |
| YDL058W | 53.1160216 | 0.27495208 | 0.61525025 | 0.23214308 | 0.89041152 | 0.93317728 | USO1 | IV | 345665 | 351037 | 1 |
| YDL059C | 7.69367683 | -1.246663 | 1.05068368 | 2.4185594 | 0.29841215 | 0.4716916 | RAD59 | IV | 344237 | 344953 | -1 |
| YDL060W | 72.6023375 | 0.02945308 | 0.61123739 | 0.38245425 | 0.82594497 | 0.89568423 | TSR1 | IV | 341619 | 343985 | 1 |
| YDL061C | 59.3751347 | -0.6946179 | 0.61049474 | 14.8634519 | 0.00059216 | 0.00424214 | RPS29B | IV | 340628 | 340798 | -1 |
| YDL063C | 89.6418855 | -0.0973456 | 0.61972999 | 40.5372613 | 1.58E-09 | 5.67E-08 | SYO1 | IV | 338272 | 340134 | -1 |
| YDL064W | 13.6504755 | -1.8893021 | 0.87356407 | 10.5790864 | 0.00504406 | 0.02308706 | UBC9 | IV | 337487 | 338070 | 1 |
| YDL065C | 124.326127 | 1.2425089 | 0.67590687 | 4.76981948 | 0.09209729 | 0.20646161 | PEX19 | IV | 336249 | 337277 | -1 |
| YDL066W | 70.9023283 | -0.1506063 | 0.551373 | 2.08351652 | 0.35283376 | 0.52643121 | IDP1 | IV | 334835 | 336121 | 1 |
| YDL067C | 46.1127844 | 0.19205189 | 0.71273187 | 0.68395646 | 0.71036367 | 0.82438173 | COX9 | IV | 334217 | 334396 | -1 |
| YDL069C | 463.082143 | 0.86199507 | 0.6420905 | 3.26777411 | 0.19516946 | 0.35245914 | CBS1 | IV | 333121 | 333810 | -1 |
| YDL070W | 38.0690166 | 0.94887169 | 0.62790737 | 2.64248231 | 0.26680395 | 0.43923653 | BDF2 | IV | 331024 | 332940 | 1 |
| YDL072C | 240.162161 | 0.27048407 | 0.52804235 | 12.5355052 | 0.00189649 | 0.01073943 | YET3 | IV | 329836 | 330447 | -1 |
| YDL073W | 363.718946 | 2.19606309 | 0.55858673 | 81.5952883 | 1.91E-18 | 3.96E-16 | AHK1 | IV | 326613 | 329567 | 1 |
| YDL074C | 187.067516 | -0.0079989 | 0.49595866 | 4.63766917 | 0.09838818 | 0.21668603 | BRE1 | IV | 324047 | 326149 | -1 |

| | | | | | | | | | | | |
|---|---|---|---|---|---|---|---|---|---|---|---|
| YDL075W | 141.390923 | -0.3104332 | 0.58063036 | 28.1198686 | 7.83E-07 | 1.44E-05 | RPL31A | IV | 322226 | 322988 | 1 |
| YDL076C | 118.624806 | 0.44416293 | 0.51692307 | 2.31033938 | 0.31500408 | 0.48701565 | RXT3 | IV | 320668 | 321552 | -1 |
| YDL077C | 26.0018394 | -0.9049982 | 0.73486526 | 2.12959832 | 0.3447971 | 0.51822506 | VAM6 | IV | 316971 | 320120 | -1 |
| YDL078C | 51.3206204 | -1.2716531 | 0.59430399 | 10.9322698 | 0.00422754 | 0.02004706 | MDH3 | IV | 315357 | 316388 | -1 |
| YDL079C | 642.937363 | -0.3030486 | 0.46000801 | 2.6253528 | 0.26909888 | 0.44226014 | MRK1 | IV | 312951 | 314748 | -1 |
| YDL080C | 40.0134938 | -0.0254488 | 0.66753253 | 0.6803479 | 0.71164652 | 0.82438173 | THI3 | IV | 310642 | 312471 | -1 |
| YDL081C | 118.463587 | -0.1106707 | 0.51632007 | 10.3211921 | 0.00573828 | 0.02555741 | RPP1A | IV | 309802 | 310122 | -1 |
| YDL082W | 39.1510609 | 0.37446306 | 0.60820153 | 9.63655342 | 0.0080807 | 0.03341947 | RPL13A | IV | 308424 | 309388 | 1 |
| YDL083C | 59.4584439 | -0.2854897 | 0.67711651 | 14.6682837 | 0.00065286 | 0.00461548 | RPS16A | IV | 306926 | 307789 | -1 |
| YDL084W | 81.459915 | 1.37926229 | 0.74971243 | 8.29735906 | 0.01578525 | 0.05583175 | SUB2 | IV | 305237 | 306577 | 1 |
| YDL085C-A | 44.4140201 | 0.65771382 | 0.59481458 | 1.588773 | 0.45185837 | 0.61806282 | NA | IV | 302464 | 302670 | -1 |
| YDL085W | 101.640533 | 0.70344042 | 0.53415965 | 3.08194967 | 0.21417222 | 0.37813088 | NDE2 | IV | 303211 | 304848 | 1 |
| YDL086W | 193.134209 | -0.5668468 | 0.51436301 | 6.55607059 | 0.03770226 | 0.1063821 | NA | IV | 301413 | 302234 | 1 |
| YDL088C | 32.763577 | -0.0947104 | 0.71949137 | 4.65739911 | 0.09742236 | 0.21492841 | ASM4 | IV | 298417 | 300003 | -1 |
| YDL089W | 144.625397 | 2.70482742 | 0.8610663 | 8.79104744 | 0.01233242 | 0.0461222 | NUR1 | IV | 296820 | 298274 | 1 |
| YDL090C | 44.4426734 | -0.2252525 | 0.63537559 | 0.86838091 | 0.64778887 | 0.77960872 | RAM1 | IV | 295034 | 296329 | -1 |
| YDL091C | 36.5121502 | 0.82336008 | 0.63558851 | 1.81368371 | 0.40379746 | 0.57458523 | UBX3 | IV | 293392 | 294759 | -1 |
| YDL092W | 47.2474625 | 1.06158634 | 0.65203263 | 4.79852059 | 0.09078508 | 0.20445182 | SRP14 | IV | 292781 | 293221 | 1 |
| YDL093W | 147.568304 | 0.69239641 | 0.57785122 | 11.6333555 | 0.00297748 | 0.01516237 | PMT5 | IV | 289909 | 292140 | 1 |
| YDL095W | 136.459951 | 0.84612213 | 0.53209212 | 3.11019474 | 0.21116882 | 0.37459175 | PMT1 | IV | 287059 | 289512 | 1 |
| YDL097C | 48.117063 | -0.1255066 | 0.630722 | 1.07658305 | 0.58374471 | 0.73199919 | RPN6 | IV | 285391 | 286695 | -1 |
| YDL099W | 99.8563851 | 1.09596964 | 0.6275921 | 3.31015777 | 0.19107698 | 0.34746725 | BUG1 | IV | 283419 | 284444 | 1 |
| YDL100C | 44.0981086 | 0.21779599 | 0.65088932 | 0.20613064 | 0.90206805 | 0.93824416 | GET3 | IV | 282112 | 283176 | -1 |
| YDL101C | 32.8447844 | 0.11944279 | 0.64318085 | 1.68615875 | 0.43038317 | 0.59945118 | DUN1 | IV | 280307 | 281848 | -1 |
| YDL102W | 89.7295819 | 0.7103385 | 0.55122193 | 13.3970943 | 0.0012327 | 0.00781955 | POL3 | IV | 276872 | 280165 | 1 |
| YDL103C | 11.8540505 | -0.7714907 | 0.88928332 | 0.77935549 | 0.6772751 | 0.8011991 | QRI1 | IV | 275148 | 276581 | -1 |
| YDL104C | 259.637937 | -0.6552464 | 0.60853468 | 4.32136721 | 0.11524631 | 0.24255767 | QRI7 | IV | 273653 | 274876 | -1 |
| YDL105W | 98.8256615 | -0.5714908 | 0.54282038 | 2.18043235 | 0.33614382 | 0.50936213 | NSE4 | IV | 272389 | 273597 | 1 |
| YDL106C | 27.4026596 | 0.86194082 | 0.82956383 | 1.21939774 | 0.54351451 | 0.69699868 | PHO2 | IV | 270222 | 271901 | -1 |
| YDL107W | 105.203975 | 0.15363025 | 0.51152031 | 0.09496677 | 0.95362632 | 0.96767513 | MSS2 | IV | 268921 | 269976 | 1 |
| YDL108W | 70.8727646 | 0.55497378 | 0.67634942 | 0.7987281 | 0.67074647 | 0.79566115 | KIN28 | IV | 267698 | 268699 | 1 |
| YDL109C | 46.5217634 | 0.620882 | 0.60404527 | 1.72003634 | 0.42315439 | 0.59395489 | NA | IV | 265258 | 267201 | -1 |
| YDL110C | 93.329144 | 1.57651887 | 0.54850081 | 10.0660704 | 0.00651899 | 0.02829458 | TMA17 | IV | 264512 | 264964 | -1 |

| | | | | | | | | | | | |
|---|---|---|---|---|---|---|---|---|---|---|---|
| YDL111C | 125.241481 | 0.01367178 | 0.70231002 | 10.3275598 | 0.00572004 | 0.02550402 | RRP42 | IV | 263313 | 264110 | -1 |
| YDL112W | 113.816495 | -0.1359589 | 0.53813171 | 0.08778042 | 0.95705903 | 0.96978855 | TRM3 | IV | 258915 | 263225 | 1 |
| YDL113C | 17.7860853 | 1.76384105 | 0.8415962 | 4.99417702 | 0.08232434 | 0.19005499 | ATG20 | IV | 256633 | 258555 | -1 |
| YDL114W | 33.5629784 | 2.4645067 | 1.03032904 | 5.51712028 | 0.06338297 | 0.15623132 | NA | IV | 255604 | 256530 | 1 |
| YDL115C | 258.465747 | -0.5900532 | 0.52086478 | 1.27382968 | 0.52892172 | 0.68319367 | IWR1 | IV | 253995 | 255126 | -1 |
| YDL116W | 7.35085685 | 0.4690546 | 1.19808652 | 0.33325447 | 0.8465151 | 0.90975468 | NUP84 | IV | 251566 | 253746 | 1 |
| YDL117W | 54.4486454 | -0.3022974 | 0.71463343 | 2.55132013 | 0.27924659 | 0.45351049 | CYK3 | IV | 248581 | 251238 | 1 |
| YDL119C | 36.8598278 | 0.86593024 | 0.67097797 | 1.62488726 | 0.44377233 | 0.61162622 | HEM25 | IV | 246689 | 247612 | -1 |
| YDL120W | 131.899966 | 1.46572409 | 0.64286035 | 10.8538365 | 0.00439662 | 0.02061251 | YFH1 | IV | 245923 | 246447 | 1 |
| YDL121C | 8.09890406 | -0.7953553 | 1.05445175 | 5.28454808 | 0.07119918 | 0.17091344 | NA | IV | 245133 | 245582 | -1 |
| YDL122W | 264.640069 | 0.43463801 | 0.60538219 | 8.63216016 | 0.01335212 | 0.04886775 | UBP1 | IV | 242552 | 244981 | 1 |
| YDL123W | 229.663303 | 0.91050899 | 0.49128232 | 6.17620823 | 0.0455883 | 0.12191042 | SNA4 | IV | 241418 | 241840 | 1 |
| YDL124W | 197.03977 | 1.18165958 | 0.65257212 | 3.23024937 | 0.19886587 | 0.35758801 | NA | IV | 240259 | 241197 | 1 |
| YDL125C | 57.8758117 | 0.30250441 | 0.57987689 | 0.30261696 | 0.85958249 | 0.91706593 | HNT1 | IV | 239019 | 239606 | -1 |
| YDL126C | 250.93767 | 0.3926452 | 0.50003269 | 0.62230746 | 0.73260124 | 0.83746519 | CDC48 | IV | 236157 | 238664 | -1 |
| YDL127W | 182.261519 | -0.0218191 | 0.55189071 | 2.79911403 | 0.24670623 | 0.41766412 | PCL2 | IV | 234927 | 235853 | 1 |
| YDL128W | 649.534225 | -0.2835701 | 0.47479697 | 0.64073737 | 0.72588137 | 0.83456178 | VCX1 | IV | 232652 | 233887 | 1 |
| YDL129W | 91.1220606 | 0.44015481 | 0.56997788 | 4.72833494 | 0.09402755 | 0.2093602 | NA | IV | 231024 | 231899 | 1 |
| YDL130W | 44.6251435 | -1.10207 | 0.68518611 | 17.5456645 | 0.00015488 | 0.00137233 | RPP1B | IV | 229906 | 230527 | 1 |
| YDL130W-A | 28.5033951 | 3.04989709 | 0.88922837 | 21.4473673 | 2.20E-05 | 0.00025693 | STF1 | IV | 229171 | 229431 | 1 |
| YDL131W | 31.4333264 | 0.51543143 | 0.67750096 | 0.67648802 | 0.71302128 | 0.8253485 | LYS21 | IV | 227393 | 228715 | 1 |
| YDL132W | 26.2996809 | 0.1086097 | 0.73069567 | 0.07724642 | 0.96211316 | 0.97258112 | CDC53 | IV | 224304 | 226751 | 1 |
| YDL133C-A | 637.842123 | -0.2904656 | 0.71870476 | 9.8441755 | 0.00728391 | 0.03098738 | RPL41A | IV | 221724 | 221801 | -1 |
| YDL133W | 1215.286 | 4.15358319 | 0.56145161 | 86.6757576 | 1.51E-19 | 3.97E-17 | SRF1 | IV | 222427 | 223740 | 1 |
| YDL134C | 92.1377053 | 0.53100708 | 0.54905796 | 8.60504444 | 0.01353438 | 0.04937874 | PPH21 | IV | 219662 | 220771 | -1 |
| YDL135C | 30.6744714 | -0.0241189 | 0.78571882 | 0.10881451 | 0.94704634 | 0.96419875 | RDI1 | IV | 218680 | 219288 | -1 |
| YDL136W | 104.392381 | 0.00116963 | 0.58120265 | 1.46624998 | 0.48040538 | 0.64328102 | RPL35A | IV | 217600 | 218367 | 1 |
| YDL137W | 94.4457558 | -1.6975623 | 0.6909308 | 15.7335741 | 0.00038326 | 0.00297068 | ARF2 | IV | 216529 | 217074 | 1 |
| YDL138W | 255.824156 | -0.8968796 | 0.49485545 | 12.0662313 | 0.00239801 | 0.01281215 | RGT2 | IV | 213351 | 215642 | 1 |
| YDL139C | 46.2250386 | -0.0201978 | 0.64933778 | 0.13125748 | 0.93647848 | 0.95873833 | SCM3 | IV | 211375 | 212046 | -1 |
| YDL140C | 163.073404 | 1.3517643 | 0.62025624 | 11.5974094 | 0.00303148 | 0.01538156 | RPO21 | IV | 205360 | 210561 | -1 |
| YDL141W | 45.9023388 | 1.0308596 | 0.73969682 | 26.8576949 | 1.47E-06 | 2.48E-05 | BPL1 | IV | 203039 | 205111 | 1 |
| YDL142C | 98.7539594 | -0.6421483 | 0.54341251 | 1.40310218 | 0.49581565 | 0.65587677 | CRD1 | IV | 201719 | 202570 | -1 |

| | | | | | | | | | | | |
|---|---|---|---|---|---|---|---|---|---|---|---|
| YDL143W | 52.2606569 | 0.4065214 | 0.64052006 | 4.05346448 | 0.1317654 | 0.26711622 | CCT4 | IV | 199996 | 201582 | 1 |
| YDL144C | 22.4116795 | 0.01313879 | 0.7013168 | 1.59605537 | 0.45021606 | 0.61669055 | NA | IV | 198662 | 199732 | -1 |
| YDL145C | 59.203475 | 0.0908464 | 0.67301773 | 0.35950928 | 0.83547518 | 0.90300565 | COP1 | IV | 194571 | 198176 | -1 |
| YDL146W | 26.9751983 | -0.0805928 | 0.76595917 | 0.21032935 | 0.90017627 | 0.93824416 | LDB17 | IV | 192750 | 194225 | 1 |
| YDL147W | 40.6151598 | 0.44400917 | 0.70277046 | 0.93724003 | 0.62586536 | 0.76430043 | RPN5 | IV | 190924 | 192261 | 1 |
| YDL148C | 48.9567783 | -1.142375 | 0.7392759 | 3.3127734 | 0.19082725 | 0.34723124 | NOP14 | IV | 188154 | 190586 | -1 |
| YDL149W | 172.034198 | -0.7867493 | 0.52397945 | 12.646335 | 0.00179425 | 0.01032258 | ATG9 | IV | 184925 | 187918 | 1 |
| YDL150W | 33.8252509 | -0.001852 | 0.67613349 | 9.97939021 | 0.00680774 | 0.02934982 | RPC53 | IV | 183343 | 184611 | 1 |
| YDL153C | 71.3586435 | 0.12633272 | 0.74654008 | 2.24200305 | 0.32595318 | 0.49874971 | SAS10 | IV | 181186 | 183018 | -1 |
| YDL154W | 10.132915 | -0.3896791 | 0.91510581 | 6.4451038 | 0.03985323 | 0.11116077 | MSH5 | IV | 178334 | 181039 | 1 |
| YDL155W | 12.2131574 | -1.8462188 | 0.89260494 | 5.57933389 | 0.06144167 | 0.15241958 | CLB3 | IV | 176773 | 178056 | 1 |
| YDL156W | 12.4497092 | -0.7317665 | 0.95448414 | 4.50709523 | 0.10502597 | 0.22698782 | CMR1 | IV | 174918 | 176486 | 1 |
| YDL157C | 11.6589645 | -0.1944718 | 0.91780803 | 4.69417081 | 0.09564753 | 0.21202113 | NA | IV | 174232 | 174588 | -1 |
| YDL159W | 207.118524 | 0.2897575 | 0.58880848 | 1.20995603 | 0.54608643 | 0.69889749 | STE7 | IV | 172481 | 174028 | 1 |
| YDL159W-A | 233.403292 | 0.39230571 | 0.74401227 | 0.92283845 | 0.63038835 | 0.76792521 | NA | IV | 172182 | 172313 | 1 |
| YDL160C | 9.3715743 | 0.50032213 | 1.02006622 | 0.35388042 | 0.83782988 | 0.90469306 | DHH1 | IV | 170410 | 171930 | -1 |
| YDL160C-A | 31.6922663 | 0.3067397 | 0.68475876 | 4.59484792 | 0.10051745 | 0.21962114 | MHF2 | IV | 169366 | 169608 | -1 |
| YDL161W | 276.349408 | 0.7388226 | 0.66339471 | 5.41739939 | 0.06662338 | 0.16194348 | ENT1 | IV | 167714 | 169078 | 1 |
| YDL164C | 51.6011219 | 0.44218102 | 0.63133924 | 2.77946262 | 0.24914224 | 0.42007384 | CDC9 | IV | 164987 | 167254 | -1 |
| YDL165W | 240.670259 | 0.24085532 | 0.50750527 | 2.9929934 | 0.22391322 | 0.38943778 | CDC36 | IV | 164290 | 164865 | 1 |
| YDL166C | 19.2616683 | -0.7203988 | 0.89083801 | 5.79116699 | 0.05526677 | 0.14109109 | FAP7 | IV | 163449 | 164042 | -1 |
| YDL167C | 69.6698499 | -0.7768346 | 0.67363441 | 18.4962238 | 9.63E-05 | 0.00090509 | NRP1 | IV | 160995 | 163154 | -1 |
| YDL168W | 21.2838689 | 0.7233355 | 0.79342881 | 0.95179722 | 0.62132648 | 0.76184702 | SFA1 | IV | 159604 | 160764 | 1 |
| YDL169C | 29.9602525 | 2.86911379 | 1.00120868 | 7.93295185 | 0.01894006 | 0.06346236 | UGX2 | IV | 158064 | 158735 | -1 |
| YDL170W | 11.4835194 | 0.41923839 | 1.12965401 | 2.49515839 | 0.28719921 | 0.4613994 | UGA3 | IV | 156318 | 157904 | 1 |
| YDL171C | 79.3804091 | 0.49294016 | 0.63811191 | 1.46345774 | 0.48107655 | 0.64358439 | GLT1 | IV | 149203 | 155640 | -1 |
| YDL173W | 302.118732 | 1.16317295 | 0.53906218 | 5.91939818 | 0.05183451 | 0.1343428 | PAR32 | IV | 148191 | 149078 | 1 |
| YDL174C | 122.630622 | 0.07807804 | 0.54775167 | 1.5126089 | 0.46939791 | 0.63190279 | DLD1 | IV | 145826 | 147589 | -1 |
| YDL175C | 63.8892846 | -0.7547058 | 0.64230891 | 5.44478283 | 0.06571741 | 0.16055012 | AIR2 | IV | 144484 | 145518 | -1 |
| YDL176W | 16.0379334 | 0.12406875 | 0.81879314 | 0.03191748 | 0.98416793 | 0.98751104 | NA | IV | 142097 | 144223 | 1 |
| YDL177C | 14.637641 | 1.14904407 | 0.93848008 | 3.07963423 | 0.21442031 | 0.37828653 | NA | IV | 141209 | 141721 | -1 |
| YDL178W | 43.223173 | 0.04381837 | 0.6982153 | 3.39643924 | 0.18300906 | 0.3384294 | DLD2 | IV | 139522 | 141114 | 1 |
| YDL179W | 29.6454138 | -0.1587756 | 0.77641947 | 0.07702981 | 0.96221736 | 0.97258112 | PCL9 | IV | 138291 | 139205 | 1 |

| | | | | | | | | | | | |
|---|---|---|---|---|---|---|---|---|---|---|---|
| YDL180W | 19.4840422 | 0.37415002 | 0.81864543 | 5.2458556 | 0.07259002 | 0.17317521 | NA | IV | 135896 | 137539 | 1 |
| YDL181W | 394.947947 | -0.2248459 | 0.55558537 | 4.65728522 | 0.09742791 | 0.21492841 | INH1 | IV | 135179 | 135436 | 1 |
| YDL182W | 37.1237245 | -1.7530977 | 0.85400825 | 10.2557953 | 0.00592901 | 0.02628559 | LYS20 | IV | 133437 | 134723 | 1 |
| YDL183C | 20.2589652 | -1.1924007 | 0.73856766 | 5.08404954 | 0.07870687 | 0.18416483 | NA | IV | 130871 | 131833 | -1 |
| YDL184C | 1517.23062 | -0.569431 | 0.69666155 | 14.9083378 | 0.00057902 | 0.00416465 | RPL41A | IV | 130407 | 130484 | -1 |
| YDL185W | 83.6142257 | -0.0121161 | 0.72485113 | 1.44399247 | 0.48578155 | 0.64773825 | VMA1 | IV | 126787 | 130002 | 1 |
| YDL186W | 270.9147 | -1.0559016 | 0.53475694 | 10.4250239 | 0.00544797 | 0.02452858 | NA | IV | 125616 | 126449 | 1 |
| YDL188C | 49.4125393 | -1.6130009 | 0.61895832 | 7.02408878 | 0.02983586 | 0.088635 | PPH22 | IV | 123865 | 124998 | -1 |
| YDL189W | 27.2519369 | -0.1766995 | 0.75042978 | 0.41392761 | 0.81304908 | 0.88754792 | RBS1 | IV | 122216 | 123589 | 1 |
| YDL190C | 38.8253492 | 0.08439069 | 0.68009908 | 2.05132377 | 0.35855906 | 0.53153496 | UFD2 | IV | 118707 | 121592 | -1 |
| YDL191W | 293.40126 | 1.84064558 | 0.81559232 | 6.40315128 | 0.04069803 | 0.11285424 | RPL35A | IV | 117664 | 118517 | 1 |
| YDL192W | 24.4474893 | -0.0069229 | 0.76121076 | 1.16740145 | 0.55783017 | 0.70907501 | ARF1 | IV | 116321 | 116866 | 1 |
| YDL193W | 26.5522463 | 0.52702643 | 0.72960512 | 0.96310294 | 0.61782411 | 0.75916842 | NUS1 | IV | 114672 | 115799 | 1 |
| YDL194W | 49.8079764 | 0.93700207 | 0.79858428 | 30.4410043 | 2.45E-07 | 5.22E-06 | SNF3 | IV | 111580 | 114234 | 1 |
| YDL195W | 98.5171519 | 0.65771552 | 0.61549759 | 2.95129395 | 0.22863076 | 0.39492008 | SEC31 | IV | 107208 | 111029 | 1 |
| YDL197C | 16.5109338 | -0.0989745 | 0.777133 | 0.0351144 | 0.98259603 | 0.98668592 | ASF2 | IV | 104917 | 106494 | -1 |
| YDL198C | 214.641586 | 0.2624321 | 0.54259646 | 1.13317405 | 0.56745886 | 0.71800411 | GGC1 | IV | 103649 | 104551 | -1 |
| YDL199C | 488.714912 | -0.1949034 | 0.47872707 | 2.35939186 | 0.30737219 | 0.47995819 | NA | IV | 101290 | 103353 | -1 |
| YDL200C | 115.299886 | 0.70249151 | 0.7357319 | 8.44801283 | 0.01463987 | 0.05258366 | MGT1 | IV | 100501 | 101067 | -1 |
| YDL201W | 130.609645 | -0.3655106 | 0.49990002 | 1.38978549 | 0.49912798 | 0.65860324 | TRM8 | IV | 99561 | 100421 | 1 |
| YDL202W | 10.9490693 | -0.1502894 | 0.98468905 | 4.54523371 | 0.10304218 | 0.22378628 | MRPL11 | IV | 98475 | 99224 | 1 |
| YDL203C | 42.9245349 | -0.0810769 | 0.79112414 | 20.2278926 | 4.05E-05 | 0.00043276 | ACK1 | IV | 96082 | 97953 | -1 |
| YDL204W | 17.5533068 | -0.0025402 | 0.78470559 | 0.47721025 | 0.78772587 | 0.87068194 | RTN2 | IV | 94605 | 95786 | 1 |
| YDL205C | 29.1659079 | 0.26882159 | 0.70582551 | 1.36158252 | 0.50621629 | 0.66417229 | HEM3 | IV | 92762 | 93745 | -1 |
| YDL206W | 18.7440155 | 0.12712248 | 0.76908532 | 0.10808811 | 0.94739037 | 0.96420992 | NA | IV | 90176 | 92464 | 1 |
| YDL207W | 298.142318 | 1.76378455 | 0.51927423 | 24.6734418 | 4.39E-06 | 6.35E-05 | GLE1 | IV | 88248 | 89864 | 1 |
| YDL208W | 110.659979 | -1.6238735 | 0.80559502 | 16.6436914 | 0.00024315 | 0.00202564 | NHP2 | IV | 87512 | 87982 | 1 |
| YDL209C | 8.43499462 | -0.714589 | 1.05533187 | 4.02644886 | 0.13355733 | 0.26953536 | CWC2 | IV | 86207 | 87226 | -1 |
| YDL210W | 162.418264 | 0.99215303 | 0.62545783 | 3.08160756 | 0.21420886 | 0.37813088 | UGA4 | IV | 84270 | 85985 | 1 |
| YDL211C | 143.359524 | 0.60903073 | 0.5983677 | 17.9419548 | 0.00012704 | 0.00114935 | NA | IV | 79294 | 80412 | -1 |
| YDL212W | 112.356912 | 0.18709522 | 0.59517825 | 8.2381671 | 0.01625941 | 0.05678441 | SHR3 | IV | 78426 | 79058 | 1 |
| YDL213C | 179.302254 | -0.5607293 | 0.49088355 | 12.4402594 | 0.00198899 | 0.01111606 | NOP6 | IV | 77289 | 77966 | -1 |
| YDL214C | 697.735197 | 1.37823579 | 0.47686283 | 18.1839788 | 0.00011256 | 0.00103124 | PRR2 | IV | 74446 | 76545 | -1 |

| | | | | | | | | | | | |
|---|---|---|---|---|---|---|---|---|---|---|---|
| YDL215C | 110.31734 | -1.3138636 | 0.57258136 | 5.35176388 | 0.06884608 | 0.16625162 | GDH2 | IV | 70640 | 73918 | -1 |
| YDL216C | 95.1245989 | -0.2951712 | 0.56434327 | 2.31900233 | 0.3136426 | 0.48568886 | RRI1 | IV | 68997 | 70319 | -1 |
| YDL217C | 12.2612598 | -0.4806805 | 1.04267012 | 0.28275529 | 0.86816139 | 0.92249119 | TIM22 | IV | 67983 | 68606 | -1 |
| YDL218W | 290.642644 | -0.5561988 | 0.53616447 | 1.61316481 | 0.44638101 | 0.61376064 | NA | IV | 66493 | 67446 | 1 |
| YDL219W | 27.8068376 | -0.5141678 | 0.69290515 | 14.5081095 | 0.0007073 | 0.00491629 | DTD1 | IV | 65242 | 65765 | 1 |
| YDL220C | 165.744042 | 1.62034006 | 0.50974158 | 24.0082962 | 6.12E-06 | 8.50E-05 | CDC13 | IV | 62244 | 65018 | -1 |
| YDL222C | 198.600521 | 2.72076649 | 0.62302788 | 64.428155 | 1.02E-14 | 9.87E-13 | FMP45 | IV | 60872 | 61801 | -1 |
| YDL223C | 883.363545 | -1.3684829 | 0.52008399 | 38.3153472 | 4.79E-09 | 1.54E-07 | HBT1 | IV | 57265 | 60405 | -1 |
| YDL224C | 115.662148 | 3.5888362 | 0.76963483 | 25.4245083 | 3.01E-06 | 4.63E-05 | WHI4 | IV | 54397 | 56346 | -1 |
| YDL225W | 63.9855735 | 0.10998081 | 0.54951004 | 0.0430486 | 0.9787057 | 0.98363235 | SHS1 | IV | 52445 | 54100 | 1 |
| YDL226C | 28.0553118 | -0.2790461 | 0.66682439 | 1.32317219 | 0.51603221 | 0.67293389 | GCS1 | IV | 51115 | 52173 | -1 |
| YDL227C | 69.7749212 | -0.5453876 | 0.54925488 | 3.58734675 | 0.16634799 | 0.31558154 | HO | IV | 46271 | 48031 | -1 |
| YDL229W | 116.760554 | -0.6291903 | 0.59043708 | 27.6421101 | 9.94E-07 | 1.77E-05 | SSB1 | IV | 44065 | 45906 | 1 |
| YDL230W | 37.4489022 | -0.3626055 | 0.63120851 | 0.47696037 | 0.7878243 | 0.87068194 | PTP1 | IV | 42700 | 43707 | 1 |
| YDL231C | 173.288301 | 1.90049883 | 0.94970284 | 6.17260021 | 0.04567062 | 0.12191465 | BRE4 | IV | 38867 | 42244 | -1 |
| YDL233W | 17.415683 | 1.58087022 | 0.83112856 | 4.76273971 | 0.09242388 | 0.20685781 | MFG1 | IV | 36797 | 38173 | 1 |
| YDL234C | 218.919544 | 0.05076393 | 0.52201834 | 0.07985731 | 0.96085799 | 0.97212597 | GYP7 | IV | 34237 | 36477 | -1 |
| YDL235C | 190.388624 | 2.18582968 | 0.60068336 | 15.321697 | 0.00047091 | 0.00350007 | YPD1 | IV | 33415 | 33918 | -1 |
| YDL236W | 149.9029 | 1.37906987 | 0.57056051 | 20.7880357 | 3.06E-05 | 0.0003442 | PHO13 | IV | 32296 | 33234 | 1 |
| YDL237W | 50.1960388 | 0.06861043 | 0.59891808 | 0.86085764 | 0.6502302 | 0.78140989 | AIM6 | IV | 30657 | 31829 | 1 |
| YDL238C | 52.1291465 | 0.21445661 | 0.71332679 | 27.2703173 | 1.20E-06 | 2.07E-05 | GUD1 | IV | 28985 | 30454 | -1 |
| YDL239C | 10.4984052 | 0.56158847 | 1.01912465 | 0.85026036 | 0.65368468 | 0.7839342 | ADY3 | IV | 26403 | 28775 | -1 |
| YDL241W | 11.8791569 | 0.37837627 | 1.00827341 | 1.72394927 | 0.42232732 | 0.59293772 | NA | IV | 20635 | 21006 | 1 |
| YDL242W | 446.063187 | 0.67424136 | 0.49758348 | 2.96807281 | 0.2267207 | 0.39279962 | NA | IV | 18959 | 19312 | 1 |
| YDL243C | 69.1838164 | -0.4620742 | 0.60961451 | 2.56008531 | 0.27802544 | 0.45261809 | AAD4 | IV | 17577 | 18566 | -1 |
| YDL244W | 8.08072935 | -1.5231496 | 1.06256157 | 3.19609806 | 0.2022908 | 0.36194799 | THI13 | IV | 16204 | 17226 | 1 |
| YDL245C | 539.502453 | 0.32789569 | 0.60528674 | 8.03794742 | 0.0179714 | 0.06102898 | HXT15 | IV | 11657 | 13360 | -1 |
| YDL247W | 34.7258308 | 1.65578485 | 0.67431139 | 6.41618691 | 0.04043363 | 0.11228331 | MPH2 | IV | 5985 | 7814 | 1 |
| YDR001C | 84.7276068 | 1.69715158 | 0.72497662 | 8.3862594 | 0.01509896 | 0.05383187 | NTH1 | IV | 450220 | 452475 | -1 |
| YDR002W | 133.215285 | -0.5936421 | 0.62656927 | 3.23693134 | 0.19820257 | 0.35661681 | YRB1 | IV | 453045 | 453650 | 1 |
| YDR003W | 25.8852123 | -1.1280258 | 0.69915821 | 2.65432879 | 0.26522828 | 0.43749737 | RCR2 | IV | 454122 | 454754 | 1 |
| YDR003W-A | 10.5238989 | -0.638819 | 0.9787101 | 7.2349064 | 0.02685097 | 0.08173877 | NA | IV | 454782 | 454904 | 1 |
| YDR004W | 788.407218 | 1.39326693 | 0.48606466 | 67.1264651 | 2.65E-15 | 2.95E-13 | RAD57 | IV | 455201 | 456583 | 1 |

| | | | | | | | | | | | |
|---|---|---|---|---|---|---|---|---|---|---|---|
| YDR005C | 658.39579 | 1.15916409 | 0.63151264 | 28.6399242 | 6.04E-07 | 1.15E-05 | MAF1 | IV | 456836 | 458103 | -1 |
| YDR006C | 70.0298313 | 0.57286747 | 0.56048033 | 1.91225473 | 0.38437856 | 0.5574307 | SOK1 | IV | 458542 | 461247 | -1 |
| YDR007W | 1359.77214 | -1.1068005 | 0.50219877 | 4.85640743 | 0.08819511 | 0.20080602 | TRP1 | IV | 461842 | 462516 | 1 |
| YDR009W | 96.0243104 | -0.1539775 | 0.58858296 | 2.6938992 | 0.26003225 | 0.43177137 | GAL3 | IV | 463434 | 464996 | 1 |
| YDR010C | 138.898764 | 0.01148613 | 0.59207125 | 0.39306048 | 0.82157649 | 0.89315206 | NA | IV | 465051 | 465383 | -1 |
| YDR011W | 89.3640715 | 0.58931622 | 0.78461091 | 4.92310885 | 0.08530225 | 0.19575903 | SNQ2 | IV | 465919 | 470424 | 1 |
| YDR012W | 72.1054899 | -0.3349137 | 0.6714902 | 4.73687178 | 0.09362705 | 0.20898252 | RPL4B | IV | 471853 | 472941 | 1 |
| YDR013W | 5.29562963 | 0.70119287 | 1.39880485 | 1.45944617 | 0.48204246 | 0.64436175 | PSF1 | IV | 473157 | 473783 | 1 |
| YDR014W | 37.7300773 | 0.34540672 | 0.7172986 | 3.87772588 | 0.14386744 | 0.28410385 | RAD61 | IV | 474046 | 475989 | 1 |
| YDR014W-A | 9.76288092 | 0.36783739 | 0.94232423 | 0.61219795 | 0.73631374 | 0.83938896 | HED1 | IV | 477797 | 478285 | 1 |
| YDR016C | 64.0435035 | 0.42455699 | 0.66558194 | 11.829213 | 0.00269972 | 0.01404438 | DAD1 | IV | 478474 | 478758 | -1 |
| YDR017C | 116.533952 | 1.16265969 | 0.54193212 | 28.1310724 | 7.79E-07 | 1.44E-05 | KCS1 | IV | 479115 | 482267 | -1 |
| YDR018C | 74.1186026 | -0.3498044 | 0.61916894 | 11.4420842 | 0.00327629 | 0.01636734 | NA | IV | 482670 | 483860 | -1 |
| YDR019C | 116.217421 | 1.80144506 | 0.63662684 | 52.3307152 | 4.33E-12 | 2.79E-10 | GCV1 | IV | 484163 | 485365 | -1 |
| YDR020C | 94.749602 | 1.37756694 | 0.62250036 | 27.8705059 | 8.87E-07 | 1.61E-05 | DAS2 | IV | 485746 | 486444 | -1 |
| YDR021W | 63.6009026 | 2.34814554 | 0.73635066 | 9.65211032 | 0.00801809 | 0.03327938 | FAL1 | IV | 486804 | 488003 | 1 |
| YDR022C | 30.1690674 | -0.8574752 | 0.68374095 | 2.58794545 | 0.27417938 | 0.44776846 | ATG31 | IV | 488072 | 488662 | -1 |
| YDR023W | 61.2264491 | -0.6308872 | 0.7872795 | 11.7073223 | 0.00286937 | 0.01473523 | SES1 | IV | 489508 | 490896 | 1 |
| YDR025W | 127.69053 | -0.2669716 | 0.50508261 | 29.6395354 | 3.66E-07 | 7.34E-06 | RPS11B | IV | 491515 | 492324 | 1 |
| YDR026C | 43.7395144 | 0.48641192 | 0.62646346 | 2.63305702 | 0.26806427 | 0.44106057 | NSI1 | IV | 492556 | 494268 | -1 |
| YDR027C | 117.341502 | 0.68571856 | 0.54279025 | 29.518613 | 3.89E-07 | 7.74E-06 | VPS54 | IV | 494649 | 497318 | -1 |
| YDR028C | 50.2984055 | 0.43368264 | 0.64063097 | 0.66745648 | 0.7162484 | 0.82794286 | REG1 | IV | 497835 | 500879 | -1 |
| YDR029W | 1893.6479 | 1.09642737 | 0.72803391 | 2.21866909 | 0.32977834 | 0.50287506 | NA | IV | 501100 | 501414 | 1 |
| YDR030C | 95.4792419 | -0.0775722 | 0.59246139 | 1.08193811 | 0.58218381 | 0.73072713 | RAD28 | IV | 501753 | 503273 | -1 |
| YDR031W | 12.5753819 | 1.97172407 | 1.16242729 | 3.00559681 | 0.22250662 | 0.38746266 | MIX14 | IV | 503498 | 503863 | 1 |
| YDR032C | 24.8086609 | 0.31347938 | 0.686456 | 0.29555329 | 0.86262376 | 0.91947562 | PST2 | IV | 504099 | 504695 | -1 |
| YDR033W | 51.7592807 | -1.1891328 | 0.85503901 | 11.8347448 | 0.00269227 | 0.01402028 | MRH1 | IV | 508147 | 509109 | 1 |
| YDR034C | 30.2079358 | -0.1218822 | 0.69881563 | 1.79269018 | 0.40805835 | 0.5793986 | LYS14 | IV | 509737 | 512109 | -1 |
| YDR034C-A | 16.4907013 | 0.33650066 | 0.78985632 | 0.29046346 | 0.86482186 | 0.92054956 | NA | IV | 520516 | 520692 | -1 |
| YDR034C-D | 18.4310403 | 0.64088352 | 0.80478664 | 0.73705841 | 0.69175101 | 0.81258639 | NA | IV | 514043 | 519356 | -1 |
| YDR034W-B | 109.111524 | 3.9721246 | 0.71424596 | 47.8919546 | 3.98E-11 | 2.01E-09 | NA | IV | 521314 | 521469 | 1 |
| YDR035W | 56.8585256 | 0.8303531 | 0.71376175 | 1.74830485 | 0.41721549 | 0.58892811 | ARO3 | IV | 521816 | 522928 | 1 |
| YDR036C | 65.5304343 | -0.0037498 | 0.63322455 | 7.32875046 | 0.02562017 | 0.07911509 | EHD3 | IV | 523211 | 524713 | -1 |

| | | | | | | | | | | | |
|---|---|---|---|---|---|---|---|---|---|---|---|
| YDR037W | 176.498124 | 0.26333092 | 0.51043538 | 5.72988709 | 0.05698635 | 0.14462713 | KRS1 | IV | 525440 | 527215 | 1 |
| YDR038C | 30.5074814 | 0.66993345 | 0.86188268 | 0.65978077 | 0.71900254 | 0.82938751 | ENA5 | IV | 527422 | 530697 | -1 |
| YDR040C | 159.590206 | -0.8333224 | 0.50168561 | 2.77316821 | 0.24992757 | 0.42103016 | ENA1 | IV | 535192 | 538467 | -1 |
| YDR041W | 6.63512044 | -0.9835751 | 1.18240153 | 0.81738544 | 0.66451839 | 0.79148192 | RSM10 | IV | 539803 | 540414 | 1 |
| YDR042C | 15.624754 | 0.09253719 | 0.80374582 | 5.96967539 | 0.05054771 | 0.13171522 | NA | IV | 540601 | 541203 | -1 |
| YDR043C | 113.452986 | -0.8947111 | 0.64540831 | 9.70558566 | 0.00780654 | 0.03254132 | NRG1 | IV | 542674 | 543369 | -1 |
| YDR044W | 7.78512596 | -0.9930003 | 1.14867794 | 1.36706083 | 0.50483158 | 0.66295642 | HEM13 | IV | 546642 | 547628 | 1 |
| YDR046C | 29.9593812 | -0.3705182 | 0.90261288 | 1.61080507 | 0.446908 | 0.61424863 | BAP3 | IV | 548762 | 550576 | -1 |
| YDR047W | 21.6630539 | 0.11071722 | 0.75240771 | 2.43986552 | 0.29525002 | 0.46951321 | HEM12 | IV | 551860 | 552948 | 1 |
| YDR049W | 39.6918006 | -0.4263926 | 0.65852029 | 1.93466611 | 0.38009538 | 0.55392707 | VMS1 | IV | 553254 | 555152 | 1 |
| YDR050C | 233.848319 | 0.41350947 | 0.53750356 | 0.84035497 | 0.65693022 | 0.78549013 | TPI1 | IV | 555726 | 556472 | -1 |
| YDR051C | 45.5405742 | 0.41774224 | 0.61215743 | 1.0189341 | 0.6008157 | 0.74682758 | DET1 | IV | 557056 | 558060 | -1 |
| YDR052C | 145.514307 | -1.0831283 | 0.52900203 | 11.804439 | 0.00273337 | 0.01414318 | DBF4 | IV | 558511 | 560625 | -1 |
| YDR054C | 61.2418144 | 0.75785033 | 0.673503 | 1.30469522 | 0.52082166 | 0.6765916 | CDC34 | IV | 561440 | 562327 | -1 |
| YDR055W | 181.28796 | 1.19194534 | 0.56228018 | 5.90697482 | 0.05215749 | 0.13493829 | PST1 | IV | 563527 | 564861 | 1 |
| YDR056C | 322.709443 | 3.66976559 | 0.79203093 | 17.9117435 | 0.00012898 | 0.00116502 | EMC10 | IV | 565023 | 565640 | -1 |
| YDR057W | 200.38693 | -0.3379962 | 0.5242721 | 8.4242075 | 0.01481517 | 0.05295051 | YOS9 | IV | 565927 | 567555 | 1 |
| YDR058C | 156.037859 | 0.08999928 | 0.51388286 | 7.56481642 | 0.0227678 | 0.07300493 | TGL2 | IV | 567723 | 568703 | -1 |
| YDR059C | 123.807223 | -0.7159313 | 0.60515303 | 9.53192392 | 0.00851469 | 0.03484104 | UBC5 | IV | 569234 | 569770 | -1 |
| YDR060W | 123.641223 | -0.8591899 | 0.52464634 | 20.7126491 | 3.18E-05 | 0.00035263 | MAK21 | IV | 570649 | 573726 | 1 |
| YDR061W | 507.113169 | 1.15735606 | 0.46677866 | 17.69195 | 0.00014396 | 0.0012883 | NA | IV | 574164 | 575783 | 1 |
| YDR062W | 45.0801462 | 0.38906515 | 0.62998463 | 0.88114944 | 0.64366639 | 0.77686719 | LCB2 | IV | 576474 | 578159 | 1 |
| YDR063W | 74.6079746 | -0.1995858 | 0.66388074 | 5.68905634 | 0.0581617 | 0.14654319 | AIM7 | IV | 578663 | 579112 | 1 |
| YDR064W | 84.047414 | -0.5685372 | 0.79835164 | 6.34397828 | 0.04192013 | 0.11497752 | RPS13 | IV | 579458 | 580452 | 1 |
| YDR065W | 115.324409 | -0.5411665 | 0.53869032 | 1.20953183 | 0.54620227 | 0.69889749 | RRG1 | IV | 580687 | 581784 | 1 |
| YDR066C | 485.520996 | -0.1865754 | 0.4575186 | 5.41797947 | 0.06660406 | 0.16194348 | RTR2 | IV | 581908 | 582498 | -1 |
| YDR067C | 130.671723 | -0.4000402 | 0.52419905 | 2.36834255 | 0.30599966 | 0.47867401 | OCA6 | IV | 582791 | 583465 | -1 |
| YDR068W | 429.516004 | 1.32744035 | 0.58515462 | 31.7210709 | 1.29E-07 | 3.01E-06 | DOS2 | IV | 583713 | 584645 | 1 |
| YDR069C | 1196.75898 | 0.22305133 | 0.50275616 | 88.2945526 | 6.72E-20 | 1.94E-17 | DOA4 | IV | 584939 | 587719 | -1 |
| YDR070C | 14.7327807 | 0.00637899 | 0.85536741 | 0.7436105 | 0.68948851 | 0.81075111 | FMP16 | IV | 588098 | 588379 | -1 |
| YDR071C | 17.7436025 | 0.63138062 | 0.98784029 | 0.46885132 | 0.79102504 | 0.87338577 | PAA1 | IV | 588827 | 589402 | -1 |
| YDR072C | 184.412977 | -0.8096575 | 0.52255208 | 5.88544646 | 0.05272196 | 0.13597334 | IPT1 | IV | 589761 | 591344 | -1 |
| YDR073W | 60.8939636 | -0.1774253 | 0.58872713 | 0.14914115 | 0.92814197 | 0.95498585 | SNF11 | IV | 592439 | 592948 | 1 |

| | | | | | | | | | | | |
|---|---|---|---|---|---|---|---|---|---|---|---|
| YDR074W | 54.6658773 | 0.77147564 | 0.59068212 | 1.85920006 | 0.39471155 | 0.56638907 | TPS2 | IV | 593893 | 596583 | 1 |
| YDR075W | 14.9661078 | -0.4747455 | 0.92671406 | 4.59002132 | 0.10076032 | 0.21998577 | PPH3 | IV | 597156 | 598082 | 1 |
| YDR076W | 383.57142 | 0.24442468 | 0.49618297 | 0.26580748 | 0.87554937 | 0.92710879 | RAD55 | IV | 598468 | 599688 | 1 |
| YDR077W | 4011.61506 | 3.68359124 | 0.69784346 | 23.9823169 | 6.20E-06 | 8.57E-05 | SED1 | IV | 600793 | 601809 | 1 |
| YDR078C | 375.786414 | 1.95883474 | 0.51608551 | 23.9184311 | 6.40E-06 | 8.82E-05 | SHU2 | IV | 602198 | 602869 | -1 |
| YDR079C-A | 86.9004218 | 1.10487878 | 0.66534974 | 4.33476063 | 0.11447712 | 0.24134059 | TFB5 | IV | 603593 | 603811 | -1 |
| YDR079W | 172.618252 | 0.84436928 | 0.7540805 | 2.94122851 | 0.2297843 | 0.3965577 | PET100 | IV | 603064 | 603399 | 1 |
| YDR080W | 165.199812 | 0.51190092 | 0.55734754 | 28.8028744 | 5.57E-07 | 1.06E-05 | VPS41 | IV | 604008 | 606986 | 1 |
| YDR081C | 95.6221685 | -0.2798103 | 0.51602905 | 3.78960978 | 0.15034767 | 0.2931317 | PDC2 | IV | 607304 | 610081 | -1 |
| YDR082W | 7.17947002 | -1.2647045 | 1.1606801 | 1.31622018 | 0.51782906 | 0.67436642 | STN1 | IV | 610441 | 611925 | 1 |
| YDR083W | 344.192033 | 1.95205056 | 0.86741816 | 10.8234009 | 0.00446404 | 0.02081063 | RRP8 | IV | 612073 | 613251 | 1 |
| YDR084C | 14.9287769 | -1.1756275 | 0.84718646 | 2.42850711 | 0.29693157 | 0.47089395 | TVP23 | IV | 613405 | 614004 | -1 |
| YDR085C | 120.213236 | -1.2015569 | 0.53291196 | 11.0084975 | 0.00406944 | 0.0194889 | AFR1 | IV | 614288 | 616150 | -1 |
| YDR086C | 52.1381611 | 0.34638736 | 0.64418374 | 3.40512837 | 0.18221569 | 0.33728544 | SSS1 | IV | 616928 | 617170 | -1 |
| YDR087C | 689.586689 | 0.41508155 | 0.46423686 | 2.524938 | 0.28295455 | 0.45678444 | RRP1 | IV | 617470 | 618306 | -1 |
| YDR088C | 18.1348426 | 0.48627065 | 0.7905715 | 0.51180296 | 0.77421824 | 0.86282905 | SLU7 | IV | 618497 | 619645 | -1 |
| YDR089W | 49.5810619 | 1.12265121 | 0.8343145 | 6.06972978 | 0.04808116 | 0.12677136 | VTC5 | IV | 622112 | 624721 | 1 |
| YDR090C | 16.4871873 | -0.9856895 | 0.80481996 | 1.62525609 | 0.44369049 | 0.61162622 | NA | IV | 625066 | 625998 | -1 |
| YDR091C | 21.1344131 | -0.875022 | 0.77343815 | 9.25412191 | 0.00978347 | 0.03871927 | RLI1 | IV | 626708 | 628534 | -1 |
| YDR092W | 62.8639181 | 0.63963658 | 0.59490047 | 1.46436763 | 0.48085774 | 0.64344033 | UBC13 | IV | 629876 | 630605 | 1 |
| YDR093W | 58.0365446 | 0.13058988 | 0.63194028 | 0.62453715 | 0.73178496 | 0.83702784 | DNF2 | IV | 631282 | 636120 | 1 |
| YDR096W | 11.5915247 | -0.5287972 | 0.93381661 | 2.16857805 | 0.33814211 | 0.51131962 | GIS1 | IV | 637139 | 639823 | 1 |
| YDR097C | 78.2613239 | -0.9205002 | 0.58533108 | 7.21747265 | 0.02708605 | 0.08236778 | MSH6 | IV | 640109 | 643837 | -1 |
| YDR098C | 51.4113358 | -0.1906346 | 0.59640217 | 3.12438324 | 0.20967604 | 0.37244958 | GRX3 | IV | 644178 | 644930 | -1 |
| YDR099W | 320.56756 | 0.70336148 | 0.50489856 | 3.09334732 | 0.21295516 | 0.37660671 | BMH2 | IV | 653607 | 654428 | 1 |
| YDR100W | 20.9524094 | 0.49932423 | 0.95970319 | 14.5726923 | 0.00068483 | 0.00479473 | TVP15 | IV | 655013 | 655444 | 1 |
| YDR101C | 153.510785 | 1.99239519 | 0.90755962 | 7.75346295 | 0.02071843 | 0.06786006 | ARX1 | IV | 655686 | 657467 | -1 |
| YDR102C | 81.189338 | -0.2495186 | 0.59077723 | 0.40768488 | 0.81559086 | 0.88902268 | NA | IV | 657648 | 657980 | -1 |
| YDR103W | 118.123307 | 0.97678219 | 0.56630508 | 4.08273379 | 0.1298511 | 0.26426638 | STE5 | IV | 658350 | 661103 | 1 |
| YDR104C | 88.2742595 | 1.00862347 | 0.62850854 | 8.08686749 | 0.01753715 | 0.05999792 | SPO71 | IV | 661173 | 664910 | -1 |
| YDR105C | 24.9634546 | 0.37948218 | 0.99591855 | 0.34255734 | 0.84258674 | 0.90781117 | TMS1 | IV | 665349 | 666770 | -1 |
| YDR106W | 240.63464 | 2.55847772 | 0.56194056 | 57.8340942 | 2.76E-13 | 2.12E-11 | ARP10 | IV | 667004 | 667858 | 1 |
| YDR107C | 1614.44327 | 0.02118906 | 0.51927188 | 0.80789236 | 0.66768005 | 0.79413877 | TMN2 | IV | 669016 | 671034 | -1 |

| | | | | | | | | | | | |
|---|---|---|---|---|---|---|---|---|---|---|---|
| YDR108W | 18.2758493 | 1.21207475 | 0.96676203 | 1.58190699 | 0.45341226 | 0.61945658 | TRS85 | IV | 671269 | 673365 | 1 |
| YDR109C | 30.5884662 | 0.8480797 | 0.66877087 | 2.65584878 | 0.26502678 | 0.43735364 | NA | IV | 673520 | 675667 | -1 |
| YDR110W | 140.486339 | -0.8669315 | 0.52009127 | 20.636973 | 3.30E-05 | 0.00036413 | FOB1 | IV | 676102 | 677802 | 1 |
| YDR111C | 15.9404174 | -0.4339186 | 0.94669783 | 4.11647104 | 0.12767906 | 0.26140797 | ALT2 | IV | 678241 | 679764 | -1 |
| YDR113C | 48.5576662 | -0.4251662 | 0.66643839 | 0.40934844 | 0.81491276 | 0.88874456 | PDS1 | IV | 680496 | 681617 | -1 |
| YDR114C | 855.255851 | -0.8351706 | 0.57908153 | 7.3808214 | 0.02496175 | 0.07778715 | NA | IV | 681924 | 682226 | -1 |
| YDR115W | 20.3368428 | -0.3923071 | 0.79856163 | 2.7956937 | 0.2471285 | 0.41801753 | MRX14 | IV | 682175 | 682492 | 1 |
| YDR116C | 35.9960642 | 1.11018231 | 0.77478315 | 3.09919217 | 0.21233372 | 0.37596705 | MRPL1 | IV | 682724 | 683581 | -1 |
| YDR117C | 71.0358177 | 0.84596871 | 0.64603325 | 1.6846065 | 0.43071733 | 0.59948398 | TMA64 | IV | 683946 | 685643 | -1 |
| YDR118W | 86.0469934 | -0.5620277 | 0.54135976 | 1.133303 | 0.56742228 | 0.71800411 | APC4 | IV | 685882 | 687840 | 1 |
| YDR119W | 67.7587896 | -0.2708878 | 0.56817401 | 2.40936271 | 0.29978751 | 0.47321965 | VBA4 | IV | 688227 | 690533 | 1 |
| YDR120C | 23.9164727 | -1.1314026 | 0.80317336 | 6.33132043 | 0.04218628 | 0.115598 | TRM1 | IV | 691549 | 693261 | -1 |
| YDR121W | 31.0915482 | -0.2494752 | 0.67561009 | 6.28278796 | 0.0432225 | 0.11765788 | DPB4 | IV | 693585 | 694175 | 1 |
| YDR122W | 66.2273539 | 1.23858124 | 0.64905053 | 3.61957179 | 0.16368918 | 0.3123054 | KIN1 | IV | 694700 | 697894 | 1 |
| YDR123C | 30.4482855 | -0.2385074 | 0.65829878 | 0.13697812 | 0.93380368 | 0.95735444 | INO2 | IV | 698554 | 699468 | -1 |
| YDR124W | 1222.34168 | 0.29641872 | 0.47759675 | 2.17510193 | 0.33704091 | 0.51030806 | NA | IV | 700315 | 701289 | 1 |
| YDR125C | 144.436664 | -0.4274359 | 0.5003458 | 0.76804855 | 0.68111489 | 0.80384598 | ECM18 | IV | 701397 | 702758 | -1 |
| YDR126W | 223.659643 | 1.83526607 | 0.56326592 | 20.1491385 | 4.21E-05 | 0.00044684 | SWF1 | IV | 703234 | 704244 | 1 |
| YDR127W | 46.7258556 | 0.28507978 | 0.67398058 | 0.3971078 | 0.81991558 | 0.89184881 | ARO1 | IV | 704484 | 709250 | 1 |
| YDR128W | 20.2424975 | -0.4061905 | 0.85985605 | 0.41565243 | 0.8123482 | 0.88731158 | MTC5 | IV | 709549 | 712995 | 1 |
| YDR129C | 56.579635 | 0.33253516 | 0.68954169 | 0.94728433 | 0.62273005 | 0.76281637 | SAC6 | IV | 713340 | 715379 | -1 |
| YDR130C | 13.5154257 | -0.0759429 | 0.8520431 | 1.00782825 | 0.60416126 | 0.7493774 | FIN1 | IV | 715747 | 716622 | -1 |
| YDR131C | 15.2432011 | -1.6241688 | 0.83840432 | 4.00122668 | 0.1352523 | 0.27200793 | NA | IV | 716790 | 718460 | -1 |
| YDR132C | 462.718116 | 0.15930491 | 0.5295089 | 11.1867927 | 0.00372236 | 0.01814182 | NA | IV | 718816 | 720303 | -1 |
| YDR134C | 174.515834 | -0.3105496 | 0.49969665 | 9.73740736 | 0.00768332 | 0.03213985 | NA | NA | NA | NA | NA |
| YDR135C | 163.041802 | -0.799235 | 0.58297765 | 2.30453266 | 0.31591998 | 0.4883013 | YCF1 | IV | 723004 | 727551 | -1 |
| YDR137W | 22.4751813 | -0.0354671 | 0.73971763 | 0.58242099 | 0.74735834 | 0.84631426 | RGP1 | IV | 728259 | 730250 | 1 |
| YDR138W | 82.0784747 | 0.87884285 | 0.62032218 | 22.3857789 | 1.38E-05 | 0.00017372 | HPR1 | IV | 730578 | 732836 | 1 |
| YDR139C | 75.7609835 | -0.3867234 | 0.57988601 | 0.46738339 | 0.79160584 | 0.87352731 | RUB1 | IV | 733618 | 733924 | -1 |
| YDR140W | 392.033508 | 0.56322751 | 0.52206721 | 6.8359705 | 0.03277841 | 0.09561057 | MTQ2 | IV | 734138 | 734803 | 1 |
| YDR141C | 206.551882 | -0.2432609 | 0.56038137 | 3.75248473 | 0.15316456 | 0.29709307 | DOP1 | IV | 734901 | 739997 | -1 |
| YDR142C | 31.6201435 | 0.81722766 | 0.7543179 | 17.2543314 | 0.00017917 | 0.00154376 | PEX7 | IV | 740473 | 741600 | -1 |
| YDR143C | 404.874966 | 1.59845569 | 0.51120498 | 59.3723446 | 1.28E-13 | 1.03E-11 | SAN1 | IV | 742042 | 743874 | -1 |

| | | | | | | | | | | | |
|---|---|---|---|---|---|---|---|---|---|---|---|
| YDR144C | 57.666145 | -0.1207011 | 0.63826399 | 9.72786244 | 0.00772007 | 0.03222728 | MKC7 | IV | 744311 | 746101 | -1 |
| YDR145W | 69.2029981 | 0.36695611 | 0.5808235 | 3.35538232 | 0.18680478 | 0.34260364 | TAF12 | IV | 746738 | 748357 | 1 |
| YDR146C | 18.4190428 | -0.68949 | 0.7431394 | 2.96741511 | 0.22679527 | 0.39280426 | SWI5 | IV | 748613 | 750742 | -1 |
| YDR147W | 11.458364 | 0.08409781 | 0.85694259 | 1.18799026 | 0.55211709 | 0.70397665 | EKI1 | IV | 751631 | 753235 | 1 |
| YDR148C | 32.3737561 | 0.77445339 | 0.66647124 | 1.35617647 | 0.50758645 | 0.66521629 | KGD2 | IV | 753675 | 755066 | -1 |
| YDR150W | 247.537976 | 0.44318691 | 0.48078814 | 9.81161368 | 0.00740347 | 0.03135777 | NUM1 | IV | 755628 | 763874 | 1 |
| YDR151C | 100.331106 | 0.25815628 | 0.53029451 | 8.90024554 | 0.01167713 | 0.04427675 | CTH1 | IV | 764178 | 765155 | -1 |
| YDR152W | 42.9765074 | -0.4516526 | 0.6135177 | 4.43382043 | 0.10894521 | 0.23299267 | GIR2 | IV | 765706 | 766503 | 1 |
| YDR153C | 39.2711666 | 1.0791364 | 0.68815932 | 3.93065961 | 0.14010967 | 0.27867477 | ENT5 | IV | 766736 | 767971 | -1 |
| YDR155C | 231.833547 | 0.94529323 | 0.61771791 | 2.41368688 | 0.29914004 | 0.47232639 | CPR1 | IV | 768512 | 769000 | -1 |
| YDR156W | 8.95649951 | -2.1532201 | 1.08031007 | 5.2708751 | 0.0716876 | 0.17187212 | RPA14 | IV | 769525 | 769938 | 1 |
| YDR157W | 52.3471625 | -2.0780119 | 0.60518615 | 56.9815644 | 4.23E-13 | 3.14E-11 | NA | IV | 769931 | 770332 | 1 |
| YDR158W | 39.4143757 | 0.0241278 | 0.67469878 | 4.83895454 | 0.08896811 | 0.20216851 | HOM2 | IV | 770357 | 771454 | 1 |
| YDR159W | 39.8862347 | -0.3433089 | 0.65037371 | 0.2941664 | 0.86322215 | 0.9197748 | SAC3 | IV | 771877 | 775782 | 1 |
| YDR160W | 89.7856824 | -0.5406428 | 0.63769013 | 28.0700886 | 8.03E-07 | 1.47E-05 | SSY1 | IV | 776163 | 778721 | 1 |
| YDR161W | 74.3538001 | 0.86591879 | 0.81474977 | 1.40147201 | 0.49621995 | 0.65603003 | ACL4 | IV | 779043 | 780206 | 1 |
| YDR162C | 41.7073421 | 0.19672751 | 0.65107731 | 0.64230884 | 0.72531124 | 0.83456178 | NBP2 | IV | 780390 | 781100 | -1 |
| YDR163W | 256.441841 | 1.1357679 | 0.62103869 | 6.74692686 | 0.03427074 | 0.09906519 | CWC15 | IV | 781423 | 781950 | 1 |
| YDR164C | 118.264862 | 2.28093508 | 0.82002516 | 9.08606531 | 0.01064109 | 0.04132253 | SEC1 | IV | 782041 | 784215 | -1 |
| YDR165W | 671.544708 | 0.66961997 | 0.51674587 | 2.96254207 | 0.22734854 | 0.39340945 | TRM82 | IV | 784871 | 786205 | 1 |
| YDR166C | 209.266491 | -0.0704631 | 0.50042396 | 6.86701921 | 0.03227347 | 0.09442315 | SEC5 | IV | 786306 | 789221 | -1 |
| YDR167W | 20.6193528 | 0.31553339 | 0.74574802 | 0.95944131 | 0.61895627 | 0.76021421 | TAF10 | IV | 789449 | 790069 | 1 |
| YDR168W | 19.2559238 | 0.8365566 | 0.77964511 | 1.19170856 | 0.55109158 | 0.70297868 | CDC37 | IV | 790328 | 791848 | 1 |
| YDR169C | 28.0530495 | 1.31403298 | 0.70913646 | 4.73924923 | 0.09351582 | 0.20881474 | STB3 | IV | 792348 | 793889 | -1 |
| YDR169C-A | 7.08680967 | -2.0237042 | 1.18083739 | 3.10862851 | 0.21133425 | 0.37477039 | NA | IV | 794574 | 794723 | -1 |
| YDR170C | 163.497298 | 1.17806328 | 0.64100494 | 7.85518129 | 0.01969106 | 0.06517227 | SEC7 | IV | 796193 | 802222 | -1 |
| YDR170W-A | 918.144124 | -0.2613708 | 0.48247159 | 1.47631709 | 0.47799331 | 0.64064381 | NA | IV | 803195 | 804517 | 1 |
| YDR171W | 169.589819 | -0.0799265 | 0.51760098 | 13.504056 | 0.00116851 | 0.00748414 | HSP42 | IV | 806621 | 807748 | 1 |
| YDR172W | 129.480321 | -0.2935794 | 0.56328405 | 13.1104212 | 0.00142268 | 0.00879118 | SUP35 | IV | 808324 | 810381 | 1 |
| YDR173C | 27.2184814 | 0.03907296 | 0.74956933 | 2.98985778 | 0.22426455 | 0.38976222 | ARG82 | IV | 810565 | 811632 | -1 |
| YDR174W | 51.3402329 | -0.3171442 | 0.58467679 | 7.96751118 | 0.0186156 | 0.0626889 | HMO1 | IV | 812110 | 812850 | 1 |
| YDR175C | 10.2734478 | 0.27797379 | 0.96497297 | 1.17703995 | 0.55514831 | 0.70690757 | RSM24 | IV | 813193 | 814152 | -1 |
| YDR176W | 73.8956684 | -0.8647591 | 0.55640508 | 2.95682207 | 0.22799968 | 0.39406513 | NGG1 | IV | 814452 | 816560 | 1 |

| | | | | | | | | | | | |
|---|---|---|---|---|---|---|---|---|---|---|---|
| YDR177W | 25.1545954 | 0.60811524 | 0.70171673 | 1.46972417 | 0.47957159 | 0.64246171 | UBC1 | IV | 816878 | 817525 | 1 |
| YDR178W | 23.9402945 | 1.63159696 | 0.86601583 | 7.52274145 | 0.02325185 | 0.07417531 | SDH4 | IV | 817950 | 818495 | 1 |
| YDR179C | 27.1226505 | 2.24716898 | 1.01642405 | 5.50451177 | 0.06378381 | 0.15701882 | CSN9 | IV | 818708 | 819196 | -1 |
| YDR179W-A | 133.515016 | -0.3426663 | 0.51809091 | 5.1566317 | 0.07590173 | 0.17886487 | NVJ3 | IV | 819433 | 820824 | 1 |
| YDR180W | 98.4599664 | 0.95191197 | 0.74365878 | 3.69197671 | 0.15786921 | 0.30407942 | SCC2 | IV | 821295 | 825776 | 1 |
| YDR181C | 11.7639461 | 0.46521634 | 0.89974595 | 0.48462398 | 0.78481129 | 0.86984253 | SAS4 | IV | 825910 | 827355 | -1 |
| YDR182W | 22.9288493 | 0.59911736 | 0.72077972 | 0.71080371 | 0.70089173 | 0.81801018 | CDC1 | IV | 827582 | 829057 | 1 |
| YDR182W-A | 47.4792837 | 0.8698077 | 0.64005517 | 4.60358955 | 0.10007906 | 0.21919967 | NA | IV | 829151 | 829354 | 1 |
| YDR183W | 11.1765531 | 0.38264985 | 0.89443534 | 0.27660793 | 0.87083395 | 0.92499148 | PLP1 | IV | 829585 | 830277 | 1 |
| YDR184C | 130.990693 | 2.55770507 | 0.694194 | 38.9758656 | 3.44E-09 | 1.14E-07 | ATC1 | IV | 830629 | 831513 | -1 |
| YDR185C | 415.503033 | -1.3445822 | 0.47348606 | 8.01402898 | 0.01818761 | 0.06161866 | UPS3 | IV | 831934 | 832473 | -1 |
| YDR186C | 1042.74668 | 3.78582198 | 0.51060098 | 78.6749297 | 8.24E-18 | 1.36E-15 | SND1 | IV | 832859 | 835492 | -1 |
| YDR188W | 33.2863256 | -0.2398796 | 0.84752049 | 1.05782517 | 0.58924538 | 0.73703407 | CCT6 | IV | 836421 | 838061 | 1 |
| YDR189W | 70.9667149 | 0.36191666 | 0.57127657 | 0.63990484 | 0.72618359 | 0.83474349 | SLY1 | IV | 838392 | 840392 | 1 |
| YDR190C | 23.6238727 | 0.02775181 | 0.82870485 | 1.20341148 | 0.5478763 | 0.70011119 | RVB1 | IV | 840604 | 841995 | -1 |
| YDR191W | 8.58172516 | -0.1209981 | 0.96056678 | 1.6251901 | 0.44370513 | 0.61162622 | HST4 | IV | 842337 | 843449 | 1 |
| YDR192C | 20.1133287 | 0.86230311 | 0.82878237 | 5.89828643 | 0.05238457 | 0.13534434 | NUP42 | IV | 843569 | 844861 | -1 |
| YDR194C | 26.2258674 | -0.2340256 | 0.71818722 | 1.90229913 | 0.38629669 | 0.55874541 | MSS116 | IV | 845952 | 847946 | -1 |
| YDR195W | 241.801929 | 1.47883069 | 0.61504662 | 6.08512538 | 0.04771246 | 0.12600836 | REF2 | IV | 848599 | 850200 | 1 |
| YDR196C | 70.5363179 | 2.15947155 | 0.90977322 | 6.69166631 | 0.03523085 | 0.10098348 | CAB5 | IV | 850274 | 850999 | -1 |
| YDR197W | 51.6223186 | -0.278927 | 0.65317797 | 3.98873528 | 0.13609969 | 0.27319883 | CBS2 | IV | 851228 | 852397 | 1 |
| YDR198C | 23.9307583 | -0.2686418 | 0.7078327 | 0.17749692 | 0.91507572 | 0.94797312 | RKM2 | IV | 852533 | 853972 | -1 |
| YDR200C | 32.5657853 | 0.78659685 | 0.72758788 | 1.7472069 | 0.4174446 | 0.58908219 | VPS64 | IV | 854279 | 856093 | -1 |
| YDR201W | 16.2378653 | 1.24259982 | 0.80892515 | 2.61987911 | 0.26983637 | 0.44309488 | SPC19 | IV | 856317 | 856814 | 1 |
| YDR202C | 76.4138293 | 0.15598989 | 0.62062302 | 7.0370411 | 0.02964326 | 0.0881533 | RAV2 | IV | 856898 | 857953 | -1 |
| YDR204W | 29.6644703 | -0.5664632 | 0.6614957 | 3.18359577 | 0.20355931 | 0.36388033 | COQ4 | IV | 858137 | 859144 | 1 |
| YDR205W | 14.746364 | -0.2112115 | 0.8080202 | 0.62506222 | 0.73159287 | 0.83702784 | MSC2 | IV | 859346 | 861520 | 1 |
| YDR206W | 22.5441119 | -0.8370414 | 0.85954308 | 2.17039316 | 0.33783537 | 0.51112275 | EBS1 | IV | 862054 | 864708 | 1 |
| YDR207C | 17.2944569 | -0.4804131 | 0.77052467 | 2.01599692 | 0.36494871 | 0.53808328 | UME6 | IV | 865012 | 867522 | -1 |
| YDR208W | 28.5280377 | 0.7534205 | 0.77960063 | 1.04368863 | 0.59342507 | 0.74066204 | MSS4 | IV | 868224 | 870563 | 1 |
| YDR209C | 21534.4233 | 1.99683097 | 0.45078237 | 19.9487156 | 4.66E-05 | 0.00048506 | NA | IV | 871036 | 871449 | -1 |
| YDR210W | 115.049412 | 2.00225304 | 0.61810617 | 10.0990169 | 0.00641248 | 0.02789503 | NA | IV | 871074 | 871301 | 1 |
| YDR211W | 75.9449653 | 0.43203411 | 0.65664141 | 2.20784697 | 0.33156763 | 0.50507144 | GCD6 | IV | 884727 | 886865 | 1 |

| | | | | | | | | | | | |
|---|---|---|---|---|---|---|---|---|---|---|---|
| YDR212W | 74.9226699 | 0.37647345 | 0.62201726 | 12.0888096 | 0.00237109 | 0.01271169 | TCP1 | IV | 887232 | 888911 | 1 |
| YDR213W | 37.1720412 | 0.66459405 | 0.65327017 | 1.24180852 | 0.53745822 | 0.69208525 | UPC2 | IV | 889751 | 892492 | 1 |
| YDR214W | 199.00934 | 0.73351465 | 0.50592215 | 2.1072109 | 0.34867834 | 0.52247608 | AHA1 | IV | 892875 | 893927 | 1 |
| YDR215C | 160.393793 | -1.8717939 | 0.61375365 | 21.4981921 | 2.15E-05 | 0.00025209 | NA | IV | 894089 | 894502 | -1 |
| YDR216W | 11.2443191 | 1.72969508 | 1.22873049 | 4.10254599 | 0.12857113 | 0.2622576 | ADR1 | IV | 895035 | 899006 | 1 |
| YDR217C | 143.026564 | 1.70311391 | 0.60827375 | 16.4448897 | 0.00026856 | 0.00221819 | RAD9 | IV | 899551 | 903480 | -1 |
| YDR218C | 402.426958 | 1.26745639 | 0.5140105 | 43.0114939 | 4.57E-10 | 1.79E-08 | SPR28 | IV | 903781 | 905052 | -1 |
| YDR219C | 9.9419742 | 0.44714268 | 0.98337852 | 0.31007274 | 0.85638403 | 0.91501449 | MFB1 | IV | 905455 | 906852 | -1 |
| YDR221W | 286.615605 | -0.6692328 | 0.47519099 | 2.87956643 | 0.23697913 | 0.40475196 | GTB1 | IV | 907330 | 909438 | 1 |
| YDR222W | 1023.55115 | 1.0102514 | 0.545819 | 14.5223682 | 0.00070228 | 0.00488723 | NA | IV | 910054 | 911301 | 1 |
| YDR223W | 4141.35204 | 1.66829336 | 0.46661203 | 27.7842189 | 9.26E-07 | 1.66E-05 | CRF1 | IV | 912099 | 913502 | 1 |
| YDR224C | 289.657422 | -0.708418 | 0.5832631 | 17.5440439 | 0.00015501 | 0.00137233 | HTB1 | IV | 914317 | 914712 | -1 |
| YDR225W | 20.8914668 | -1.5579168 | 0.80089371 | 12.1252492 | 0.00232828 | 0.01250534 | HTA1 | IV | 915530 | 915928 | 1 |
| YDR226W | 65.4540037 | -0.562202 | 0.80487947 | 0.8089624 | 0.66732293 | 0.79387708 | ADK1 | IV | 916486 | 917154 | 1 |
| YDR227W | 106.168682 | 0.9519208 | 0.59475158 | 15.2048472 | 0.00049924 | 0.00367131 | SIR4 | IV | 917571 | 921647 | 1 |
| YDR228C | 260.590408 | 0.67477949 | 0.53205001 | 74.4133837 | 6.94E-17 | 9.80E-15 | PCF11 | IV | 921926 | 923806 | -1 |
| YDR229W | 104.322787 | 2.22935058 | 0.68969682 | 9.76545018 | 0.00757634 | 0.03185694 | IVY1 | IV | 924785 | 926146 | 1 |
| YDR231C | 72.1422107 | -0.0412674 | 0.67193131 | 1.6103937 | 0.44699993 | 0.61424863 | COX20 | IV | 926293 | 926910 | -1 |
| YDR232W | 86.6431963 | 2.1938337 | 0.60253674 | 12.8501646 | 0.0016204 | 0.00963256 | HEM1 | IV | 927452 | 929098 | 1 |
| YDR233C | 82.8030985 | 0.30395654 | 0.56933925 | 0.7073423 | 0.70210582 | 0.81847842 | RTN1 | IV | 929470 | 930357 | -1 |
| YDR234W | 38.2969896 | -0.3989718 | 0.61506946 | 5.08415031 | 0.07870291 | 0.18416483 | LYS4 | IV | 931129 | 933210 | 1 |
| YDR235W | 174.212077 | 2.2997183 | 0.79746109 | 7.79600825 | 0.02028235 | 0.06680024 | PRP42 | IV | 933504 | 935138 | 1 |
| YDR236C | 41.4089579 | 0.45894525 | 0.71203495 | 1.56170451 | 0.4580155 | 0.62324553 | FMN1 | IV | 935236 | 935892 | -1 |
| YDR237W | 99.1319367 | -0.8255302 | 0.52648889 | 2.4551356 | 0.29300436 | 0.46722534 | MRPL7 | IV | 936615 | 937493 | 1 |
| YDR238C | 76.8303337 | 0.4483502 | 0.62646282 | 0.57299331 | 0.75088959 | 0.84898472 | SEC26 | IV | 937895 | 940816 | -1 |
| YDR239C | 18.3557136 | -1.2080241 | 0.89945295 | 7.10001726 | 0.02872439 | 0.08612855 | NA | IV | 941057 | 943420 | -1 |
| YDR240C | 48.9199332 | -1.2532737 | 0.60479865 | 4.34647725 | 0.11380844 | 0.24049301 | SNU56 | IV | 943674 | 945152 | -1 |
| YDR242W | 959.288404 | -1.2039074 | 0.49099391 | 15.605735 | 0.00040856 | 0.00312906 | AMD2 | IV | 946807 | 948456 | 1 |
| YDR243C | 327.330388 | 1.03440226 | 0.47943715 | 5.4187025 | 0.06657999 | 0.16194348 | PRP28 | IV | 948518 | 950284 | -1 |
| YDR244W | 21.1605285 | 0.39229051 | 0.78039053 | 0.96868254 | 0.61610291 | 0.75801867 | PEX5 | IV | 950563 | 952401 | 1 |
| YDR245W | 19.9441949 | -0.5845713 | 0.82385 | 3.84986723 | 0.14588544 | 0.28719849 | MNN10 | IV | 952800 | 953981 | 1 |
| YDR246W | 158.584625 | -0.4335809 | 0.51964322 | 1.9380379 | 0.37945512 | 0.55341188 | TRS23 | IV | 954288 | 954947 | 1 |
| YDR246W-A | 42.4636538 | 0.44903204 | 0.6728689 | 6.37546075 | 0.04126542 | 0.11399179 | NA | IV | 955133 | 955333 | 1 |

| | | | | | | | | | | | |
|---|---|---|---|---|---|---|---|---|---|---|---|
| YDR247W | 25.3283139 | 1.10090253 | 0.74673107 | 4.01151696 | 0.1345582 | 0.27098851 | VHS1 | IV | 956013 | 957398 | 1 |
| YDR248C | 632.775493 | 3.15062413 | 0.5635505 | 71.8640032 | 2.48E-16 | 3.19E-14 | NA | IV | 957758 | 958339 | -1 |
| YDR249C | 21.1576263 | 2.59178324 | 0.89706237 | 8.44341366 | 0.01467358 | 0.05261147 | NA | IV | 958681 | 959802 | -1 |
| YDR251W | 89.2560196 | -0.9802303 | 0.55572913 | 3.96691666 | 0.13759257 | 0.27528023 | PAM1 | IV | 960614 | 963106 | 1 |
| YDR253C | 933.304701 | -0.8131202 | 0.47624984 | 11.1885498 | 0.0037191 | 0.01814182 | MET32 | IV | 963990 | 964565 | -1 |
| YDR254W | 2047.39653 | 1.78703045 | 0.55150462 | 10.5536113 | 0.00510872 | 0.0233461 | CHL4 | IV | 965113 | 966489 | 1 |
| YDR255C | 495.95119 | -0.1364842 | 0.47780957 | 7.83678321 | 0.01987303 | 0.06563882 | RMD5 | IV | 966557 | 967822 | -1 |
| YDR256C | 1450.88045 | -1.0220513 | 0.4698057 | 6.52337551 | 0.03832366 | 0.10782022 | CTA1 | IV | 968133 | 969680 | -1 |
| YDR257C | 19.6778904 | -0.6472434 | 0.79030757 | 1.80145989 | 0.406273 | 0.57768189 | RKM4 | IV | 969990 | 971474 | -1 |
| YDR258C | 38.8444024 | 0.87296479 | 0.80709361 | 5.68712867 | 0.05821779 | 0.14655695 | HSP78 | IV | 971808 | 974243 | -1 |
| YDR259C | 69.999617 | 0.79166413 | 0.72081936 | 17.9600487 | 0.0001259 | 0.00114078 | YAP6 | IV | 974631 | 975782 | -1 |
| YDR260C | 236.933996 | 0.10671092 | 0.54829966 | 2.6438084 | 0.26662711 | 0.43907024 | SWM1 | IV | 976717 | 977229 | -1 |
| YDR261C | 199.924536 | -0.714417 | 0.54037822 | 1.74821757 | 0.4172337 | 0.58892811 | EXG2 | IV | 977521 | 979209 | -1 |
| YDR261C-D | 5.57146053 | 1.27019548 | 1.32754373 | 1.1200846 | 0.5711849 | 0.72145737 | NA | IV | 987534 | 992349 | -1 |
| YDR261W-B | 11.1163817 | -0.726581 | 0.88333494 | 3.01405755 | 0.22156733 | 0.38636302 | NA | IV | 981462 | 986775 | 1 |
| YDR262W | 26.7728231 | 0.34894959 | 0.75124937 | 0.23970833 | 0.88704979 | 0.93215081 | NA | IV | 993134 | 993952 | 1 |
| YDR263C | 93.7610117 | 0.64569126 | 0.58265711 | 6.31531479 | 0.04252524 | 0.1160873 | DIN7 | IV | 994242 | 995534 | -1 |
| YDR264C | 29.9666969 | 0.22298385 | 0.74108447 | 0.14694035 | 0.92916386 | 0.95541731 | AKR1 | IV | 996029 | 998323 | -1 |
| YDR265W | 148.685147 | -0.7490852 | 0.54492909 | 8.21883514 | 0.01641733 | 0.05715957 | PEX10 | IV | 998864 | 999877 | 1 |
| YDR266C | 87.192382 | -0.2491638 | 0.73223176 | 20.1454156 | 4.22E-05 | 0.00044686 | HEL2 | IV | 1000104 | 1E+06 | -1 |
| YDR267C | 31.7372522 | 0.23721247 | 0.68146056 | 0.18541495 | 0.91146009 | 0.94542349 | CIA1 | IV | 1002510 | 1E+06 | -1 |
| YDR268W | 73.7785833 | -2.0855892 | 0.6319511 | 25.1866777 | 3.39E-06 | 5.12E-05 | MSW1 | IV | 1004004 | 1E+06 | 1 |
| YDR270W | 67.523294 | 0.09272775 | 0.55927978 | 0.20890536 | 0.90081742 | 0.93824416 | CCC2 | IV | 1005675 | 1E+06 | 1 |
| YDR272W | 48.9182148 | 0.57098809 | 0.59882554 | 1.9173079 | 0.38340863 | 0.55665395 | GLO2 | IV | 1009010 | 1E+06 | 1 |
| YDR273W | 24.3426172 | 0.54808977 | 0.73591767 | 5.67940687 | 0.058443 | 0.14688185 | DON1 | IV | 1010176 | 1E+06 | 1 |
| YDR274C | 590.626191 | -0.1532842 | 0.65416514 | 1.42061931 | 0.49149198 | 0.65269233 | NA | IV | 1011589 | 1E+06 | -1 |
| YDR275W | 69.8160607 | 1.89458123 | 0.69231629 | 24.9829297 | 3.76E-06 | 5.61E-05 | BSC2 | IV | 1012252 | 1E+06 | 1 |
| YDR276C | 89.7201399 | 0.03328807 | 0.53164145 | 3.77533544 | 0.15142456 | 0.29470528 | PMP3 | IV | 1013476 | 1E+06 | -1 |
| YDR277C | 217.368528 | -0.1860825 | 0.52632037 | 16.1100726 | 0.0003175 | 0.0025639 | MTH1 | IV | 1014401 | 1E+06 | -1 |
| YDR279W | 10.4680648 | 0.11140672 | 0.94215855 | 0.02213677 | 0.98899265 | 0.99104663 | RNH202 | IV | 1019368 | 1E+06 | 1 |
| YDR280W | 71.9917212 | -0.257729 | 0.62034763 | 8.08347226 | 0.01756695 | 0.05999792 | RRP45 | IV | 1020747 | 1E+06 | 1 |
| YDR281C | 48.5108925 | -1.1740218 | 0.65730065 | 3.82689154 | 0.14757101 | 0.28982309 | PHM6 | IV | 1022007 | 1E+06 | -1 |
| YDR282C | 9.00741783 | 0.49943414 | 1.04814062 | 2.08729624 | 0.35216759 | 0.52579947 | MRX10 | IV | 1023511 | 1E+06 | -1 |

| | | | | | | | | | | | |
|---|---|---|---|---|---|---|---|---|---|---|---|
| YDR283C | 33.1679344 | 0.57740924 | 0.66248127 | 0.94303629 | 0.62405414 | 0.76326014 | GCN2 | IV | 1025070 | 1E+06 | -1 |
| YDR284C | 68.8294204 | -1.0114573 | 0.55430617 | 5.59257554 | 0.06103622 | 0.15186924 | DPP1 | IV | 1030550 | 1E+06 | -1 |
| YDR285W | 25.7633471 | -0.3907648 | 0.69584222 | 6.44687989 | 0.03981785 | 0.11116077 | ZIP1 | IV | 1032436 | 1E+06 | 1 |
| YDR286C | 12.6456038 | 1.33158598 | 0.98387546 | 1.84602896 | 0.39731952 | 0.56801976 | NA | IV | 1035231 | 1E+06 | -1 |
| YDR287W | 77.5386234 | 0.83223057 | 0.55351127 | 8.69255887 | 0.01295492 | 0.04783737 | INM2 | IV | 1035995 | 1E+06 | 1 |
| YDR288W | 29.5710637 | -0.220581 | 0.72566032 | 7.54586275 | 0.02298459 | 0.07344413 | NSE3 | IV | 1037195 | 1E+06 | 1 |
| YDR289C | 37.5243325 | 0.51149265 | 0.62555524 | 2.07091124 | 0.35506457 | 0.52857757 | RTT103 | IV | 1038280 | 1E+06 | -1 |
| YDR291W | 84.7972404 | -0.8881272 | 0.60472256 | 4.13706735 | 0.12637095 | 0.25966788 | HRQ1 | IV | 1039728 | 1E+06 | 1 |
| YDR292C | 137.725502 | 1.28656508 | 0.89459156 | 4.73185478 | 0.09386221 | 0.20926539 | SRP101 | IV | 1043146 | 1E+06 | -1 |
| YDR293C | 85.0855518 | 0.88374227 | 0.56951834 | 2.42132041 | 0.29800047 | 0.47161883 | SSD1 | IV | 1045640 | 1E+06 | -1 |
| YDR294C | 146.333189 | 2.04201206 | 0.51816527 | 22.9195427 | 1.05E-05 | 0.00013875 | DPL1 | IV | 1050459 | 1E+06 | -1 |
| YDR295C | 68.0682711 | 0.25205682 | 0.66336344 | 0.27781197 | 0.87030985 | 0.92460441 | HDA2 | IV | 1052623 | 1E+06 | -1 |
| YDR296W | 15.2667537 | -1.1531692 | 0.82257045 | 2.31332395 | 0.31453435 | 0.48654927 | MHR1 | IV | 1055212 | 1E+06 | 1 |
| YDR297W | 10.3935206 | 0.0371441 | 1.00423705 | 0.78371017 | 0.67580204 | 0.80051019 | SUR2 | IV | 1056551 | 1E+06 | 1 |
| YDR298C | 35.8648729 | 0.17832401 | 0.74719323 | 21.2178026 | 2.47E-05 | 0.0002854 | ATP5 | IV | 1058176 | 1E+06 | -1 |
| YDR299W | 32.2785168 | -1.1329101 | 0.69021768 | 15.6124455 | 0.00040719 | 0.00312271 | BFR2 | IV | 1059627 | 1E+06 | 1 |
| YDR300C | 15.6738021 | -0.989419 | 0.78993055 | 6.31541675 | 0.04252308 | 0.1160873 | PRO1 | IV | 1061505 | 1E+06 | -1 |
| YDR301W | 19.8011313 | -0.2407828 | 0.79255468 | 0.12217525 | 0.9407408 | 0.96082012 | CFT1 | IV | 1063352 | 1E+06 | 1 |
| YDR302W | 23.8244965 | -0.4105217 | 0.74561346 | 0.33218833 | 0.84696648 | 0.90980423 | GPI11 | IV | 1067731 | 1E+06 | 1 |
| YDR303C | 17.6984554 | -0.4084316 | 0.78509562 | 0.43486476 | 0.80458201 | 0.88279205 | RSC3 | IV | 1068729 | 1E+06 | -1 |
| YDR304C | 147.111184 | 0.23326199 | 0.54173894 | 5.49582086 | 0.06406158 | 0.15736105 | CPR5 | IV | 1071880 | 1E+06 | -1 |
| YDR305C | 16.0774104 | -0.1310572 | 0.8383952 | 0.1081083 | 0.9473808 | 0.96420992 | NA | IV | 1072746 | 1E+06 | -1 |
| YDR306C | 25.4172551 | -0.5024006 | 0.69757079 | 0.54910614 | 0.75991168 | 0.85451323 | NA | IV | 1073735 | 1E+06 | -1 |
| YDR307W | 160.954554 | -0.3336774 | 0.52540101 | 12.8552798 | 0.00161626 | 0.00961783 | PMT7 | IV | 1075865 | 1E+06 | 1 |
| YDR308C | 17.0599465 | -0.4234327 | 0.78542637 | 4.55951907 | 0.10230881 | 0.22277848 | SRB7 | IV | 1078027 | 1E+06 | -1 |
| YDR309C | 43.5667768 | 0.48142558 | 0.63914516 | 6.54633807 | 0.03788617 | 0.10674069 | GIC2 | IV | 1079048 | 1E+06 | -1 |
| YDR310C | 166.539173 | -0.0969027 | 0.63401155 | 1.18117381 | 0.55400204 | 0.70568075 | SUM1 | IV | 1081128 | 1E+06 | -1 |
| YDR311W | 16.5381371 | -0.2056253 | 0.925804 | 2.06736855 | 0.35569407 | 0.52929033 | TFB1 | IV | 1085065 | 1E+06 | 1 |
| YDR312W | 425.315029 | 1.90413459 | 0.55078313 | 70.2102156 | 5.68E-16 | 6.99E-14 | SSF2 | IV | 1087581 | 1E+06 | 1 |
| YDR313C | 8.27992837 | 0.5342854 | 1.20289912 | 3.14200064 | 0.20783717 | 0.3700422 | PIB1 | IV | 1089219 | 1E+06 | -1 |
| YDR314C | 22.4728227 | 0.7117818 | 0.9117983 | 7.9979285 | 0.01833462 | 0.06204409 | RAD34 | IV | 1090433 | 1E+06 | -1 |
| YDR315C | 23.085208 | -0.2426521 | 0.78678836 | 0.30585758 | 0.85819083 | 0.91626865 | IPK1 | IV | 1092741 | 1E+06 | -1 |
| YDR316W | 71.6735655 | 0.13890478 | 0.9281811 | 20.2951841 | 3.92E-05 | 0.00042234 | OMS1 | IV | 1093763 | 1E+06 | 1 |

| | | | | | | | | | | | |
|---|---|---|---|---|---|---|---|---|---|---|---|
| YDR317W | 622.743478 | -0.1069034 | 0.48544076 | 0.79984771 | 0.67037109 | 0.79537881 | HIM1 | IV | 1102184 | 1E+06 | 1 |
| YDR318W | 1175.04082 | 0.58771211 | 0.66401926 | 2.53070758 | 0.28213946 | 0.45618192 | MCM21 | IV | 1103758 | 1E+06 | 1 |
| YDR319C | 304.702776 | 1.7243617 | 0.76441475 | 6.6037691 | 0.03681372 | 0.10453726 | YFT2 | IV | 1105003 | 1E+06 | -1 |
| YDR320C | 46.5701599 | -0.5635177 | 0.65853318 | 3.36447636 | 0.1859573 | 0.34169876 | SWA2 | IV | 1106095 | 1E+06 | -1 |
| YDR320C-A | 260.239675 | 0.70767425 | 0.51054963 | 3.03846249 | 0.21888009 | 0.38305402 | DAD4 | IV | 1108280 | 1E+06 | -1 |
| YDR321W | 13.0422662 | -0.8485748 | 0.85969228 | 6.10073419 | 0.04734154 | 0.12533495 | ASP1 | IV | 1108702 | 1E+06 | 1 |
| YDR322C-A | 72.9965435 | 0.72805006 | 0.64747455 | 8.44078002 | 0.01469291 | 0.05261147 | TIM11 | IV | 1112003 | 1E+06 | -1 |
| YDR322W | 44.7391934 | 0.08189772 | 0.6798826 | 20.4624851 | 3.60E-05 | 0.00039327 | MRPL35 | IV | 1110589 | 1E+06 | 1 |
| YDR323C | 16.1746546 | -0.890005 | 0.9185744 | 1.73339751 | 0.4203369 | 0.59107763 | PEP7 | IV | 1112480 | 1E+06 | -1 |
| YDR324C | 96.842625 | 0.18419408 | 0.61512479 | 15.8337655 | 0.00036454 | 0.00285226 | UTP4 | IV | 1114433 | 1E+06 | -1 |
| YDR325W | 71.7712552 | 0.86507696 | 0.65702904 | 12.9312307 | 0.00155603 | 0.00937506 | YCG1 | IV | 1117126 | 1E+06 | 1 |
| YDR326C | 28.7770711 | -0.312322 | 0.73866385 | 0.54436511 | 0.76171519 | 0.85526961 | YSP2 | IV | 1120609 | 1E+06 | -1 |
| YDR328C | 54.3073416 | 0.44464864 | 0.61958734 | 1.01993936 | 0.60051379 | 0.74682758 | SKP1 | IV | 1125434 | 1E+06 | -1 |
| YDR329C | 23.2193538 | 0.14117322 | 0.83546491 | 0.64762183 | 0.72338701 | 0.83301726 | PEX3 | IV | 1126270 | 1E+06 | -1 |
| YDR330W | 16.0343649 | 0.37348739 | 0.8407344 | 0.81706133 | 0.66462609 | 0.79148192 | UBX5 | IV | 1127872 | 1E+06 | 1 |
| YDR331W | 71.6029717 | -0.5339242 | 0.59049525 | 8.95308469 | 0.01137267 | 0.04332089 | GPI8 | IV | 1129588 | 1E+06 | 1 |
| YDR332W | 177.183279 | 0.79416721 | 0.58269718 | 4.34006011 | 0.11417419 | 0.24102109 | IRC3 | IV | 1131001 | 1E+06 | 1 |
| YDR333C | 21.8051429 | -0.905168 | 0.76415174 | 3.15275487 | 0.20672261 | 0.36873811 | RQC1 | IV | 1133260 | 1E+06 | -1 |
| YDR334W | 37.3171476 | 0.03412193 | 0.65654857 | 0.71268263 | 0.70023358 | 0.81801018 | SWR1 | IV | 1135932 | 1E+06 | 1 |
| YDR335W | 81.608514 | 1.04386737 | 0.64764056 | 3.01503062 | 0.22145955 | 0.38636302 | MSN5 | IV | 1141168 | 1E+06 | 1 |
| YDR336W | 24.7301787 | 0.33184028 | 0.71547442 | 2.33350461 | 0.31137656 | 0.48399202 | MRX8 | IV | 1145092 | 1E+06 | 1 |
| YDR337W | 66.6952418 | 0.94634285 | 0.66499402 | 4.75374893 | 0.0928403 | 0.20754646 | MRPS28 | IV | 1146319 | 1E+06 | 1 |
| YDR338C | 83.9168802 | 1.53058199 | 0.61708072 | 16.7952551 | 0.0002254 | 0.00188867 | NA | IV | 1147379 | 1E+06 | -1 |
| YDR339C | 25.9608686 | 0.04052156 | 0.75313247 | 6.21810574 | 0.04464322 | 0.12011349 | FCF1 | IV | 1149951 | 1E+06 | -1 |
| YDR341C | 28.417631 | -0.5897957 | 0.75251813 | 8.15459284 | 0.01695324 | 0.05856757 | NA | IV | 1151803 | 1E+06 | -1 |
| YDR344C | 117.507842 | 0.5239739 | 0.52266405 | 8.27277046 | 0.01598051 | 0.05628173 | NA | IV | 1162006 | 1E+06 | -1 |
| YDR345C | 104.598713 | -0.277382 | 0.57734709 | 4.99851383 | 0.08214602 | 0.18987044 | HXT3 | IV | 1162957 | 1E+06 | -1 |
| YDR346C | 94.4122137 | 0.1161308 | 0.5376252 | 0.16838767 | 0.91925306 | 0.94974417 | SVF1 | IV | 1167214 | 1E+06 | -1 |
| YDR347W | 88.7493759 | -0.0448973 | 0.55535851 | 2.00642663 | 0.36669923 | 0.53983944 | MRP1 | IV | 1169178 | 1E+06 | 1 |
| YDR348C | 337.965777 | 1.55618962 | 0.67892532 | 49.6409204 | 1.66E-11 | 9.34E-10 | PAL1 | IV | 1170326 | 1E+06 | -1 |
| YDR349C | 62.3459624 | -1.0076808 | 0.57369423 | 4.09990969 | 0.12874072 | 0.26238612 | YPS7 | IV | 1172386 | 1E+06 | -1 |
| YDR350C | 13.2683833 | 0.88414458 | 0.92502554 | 1.01769388 | 0.60118839 | 0.74713045 | ATP22 | IV | 1176120 | 1E+06 | -1 |
| YDR351W | 21.6263266 | 0.58509771 | 0.89117387 | 1.2325776 | 0.53994457 | 0.69395761 | SBE2 | IV | 1178666 | 1E+06 | 1 |

| | | | | | | | | | | | |
|---|---|---|---|---|---|---|---|---|---|---|---|
| YDR352W | 98.8655909 | -0.3328459 | 0.5429022 | 0.46069587 | 0.7942572 | 0.87495229 | YPQ2 | IV | 1181801 | 1E+06 | 1 |
| YDR353W | 194.767144 | -0.3169773 | 0.59142925 | 0.7779117 | 0.67776419 | 0.8011991 | TRR1 | IV | 1183299 | 1E+06 | 1 |
| YDR354W | 17.2618509 | 0.05930016 | 0.94969836 | 2.37902354 | 0.30436983 | 0.47720463 | TRP4 | IV | 1184747 | 1E+06 | 1 |
| YDR356W | 26.3714575 | 0.7519519 | 0.71574971 | 3.80246279 | 0.14938455 | 0.29203007 | SPC110 | IV | 1186107 | 1E+06 | 1 |
| YDR357C | 176.623765 | 0.42852787 | 0.77700796 | 2.40839894 | 0.29993201 | 0.47325515 | CNL1 | IV | 1189199 | 1E+06 | -1 |
| YDR358W | 58.9108891 | -0.1372662 | 0.59065527 | 2.60241225 | 0.27220328 | 0.44559146 | GGA1 | IV | 1190059 | 1E+06 | 1 |
| YDR359C | 138.129953 | -0.4406206 | 0.50120214 | 7.59949791 | 0.02237639 | 0.07205745 | EAF1 | IV | 1191936 | 1E+06 | -1 |
| YDR361C | 47.7453056 | 0.35664294 | 0.63274887 | 0.3583629 | 0.8359542 | 0.9031862 | BCP1 | IV | 1195412 | 1E+06 | -1 |
| YDR362C | 247.672768 | 0.81718101 | 0.50842198 | 20.1652313 | 4.18E-05 | 0.00044467 | TFC6 | IV | 1196679 | 1E+06 | -1 |
| YDR363W | 27.0883058 | 0.24956611 | 0.71418501 | 4.03074544 | 0.13327072 | 0.26928639 | ESC2 | IV | 1199183 | 1E+06 | 1 |
| YDR363W-A | 90.7312727 | 0.56301307 | 0.59421058 | 3.60589087 | 0.16481273 | 0.31379996 | SEM1 | IV | 1202128 | 1E+06 | 1 |
| YDR364C | 87.6695912 | -0.1327207 | 0.66058506 | 28.7065655 | 5.84E-07 | 1.11E-05 | CDC40 | IV | 1202843 | 1E+06 | -1 |
| YDR365C | 93.357196 | -1.1338438 | 0.5801911 | 15.4053957 | 0.00045161 | 0.00338267 | ESF1 | IV | 1204497 | 1E+06 | -1 |
| YDR366C | 718.366383 | -1.5863933 | 0.56451893 | 22.6266855 | 1.22E-05 | 0.00015709 | NA | IV | 1212438 | 1E+06 | -1 |
| YDR367W | 23.6739416 | 0.3128271 | 0.74063338 | 2.1898692 | 0.33456149 | 0.50802806 | KEI1 | IV | 1212848 | 1E+06 | 1 |
| YDR368W | 175.516408 | -0.6963629 | 0.51950516 | 17.453997 | 0.00016215 | 0.00142248 | YPR1 | IV | 1213904 | 1E+06 | 1 |
| YDR369C | 57.4483607 | 0.64853911 | 0.60712353 | 4.02053127 | 0.13395309 | 0.27005166 | XRS2 | IV | 1215016 | 1E+06 | -1 |
| YDR370C | 9.78894465 | 0.11221529 | 0.93840449 | 0.12428858 | 0.93974728 | 0.96037943 | DXO1 | IV | 1217782 | 1E+06 | -1 |
| YDR371W | 69.1155655 | 1.94840128 | 0.78649303 | 7.76137061 | 0.02063668 | 0.06765932 | CTS2 | IV | 1219413 | 1E+06 | 1 |
| YDR372C | 32.7757912 | 0.33308061 | 0.69239979 | 0.24549892 | 0.88448523 | 0.9315066 | VPS74 | IV | 1221112 | 1E+06 | -1 |
| YDR373W | 209.21465 | 1.77486637 | 0.67254697 | 50.064797 | 1.34E-11 | 7.71E-10 | FRQ1 | IV | 1222759 | 1E+06 | 1 |
| YDR374C | 12.2377907 | -0.8546209 | 0.99433553 | 2.55520927 | 0.2787041 | 0.45315831 | PHO92 | IV | 1223474 | 1E+06 | -1 |
| YDR375C | 227.095816 | 2.01380794 | 0.64237404 | 9.17237015 | 0.01019166 | 0.04004855 | BCS1 | IV | 1225166 | 1E+06 | -1 |
| YDR376W | 99.3596946 | 0.58334435 | 0.58983311 | 27.863404 | 8.90E-07 | 1.61E-05 | ARH1 | IV | 1226822 | 1E+06 | 1 |
| YDR377W | 11.3062227 | -0.1953792 | 0.94203829 | 0.62274574 | 0.73244072 | 0.83744703 | ATP17 | IV | 1228611 | 1E+06 | 1 |
| YDR378C | 44.466696 | 0.31876112 | 0.85215767 | 7.15650087 | 0.02792451 | 0.08418952 | LSM6 | IV | 1229349 | 1E+06 | -1 |
| YDR379C-A | 36.1142658 | 3.23669087 | 0.97436244 | 10.9807539 | 0.00412629 | 0.01964738 | SDH6 | IV | 1233278 | 1E+06 | -1 |
| YDR379W | 615.473783 | 3.78242032 | 0.8756369 | 15.23665 | 0.00049136 | 0.00363486 | RGA2 | IV | 1230167 | 1E+06 | 1 |
| YDR380W | 1091.19385 | -0.978527 | 0.48295294 | 27.3759379 | 1.14E-06 | 1.98E-05 | ARO10 | IV | 1234218 | 1E+06 | 1 |
| YDR381C-A | 148.135608 | 1.35553166 | 0.58600888 | 10.1438389 | 0.00627037 | 0.02746253 | NA | IV | 1238312 | 1E+06 | -1 |
| YDR381W | 151.484618 | -0.0943591 | 0.54544837 | 6.08130329 | 0.04780373 | 0.12615478 | YRA1 | IV | 1236558 | 1E+06 | 1 |
| YDR382W | 127.817346 | -0.2798163 | 0.60659799 | 0.3206629 | 0.85186139 | 0.91195795 | RPP2B | IV | 1239492 | 1E+06 | 1 |
| YDR383C | 2525.48605 | 2.60684549 | 1.74030721 | 4.95912716 | 0.08377978 | 0.19280005 | NKP1 | IV | 1239960 | 1E+06 | -1 |

| | | | | | | | | | | | |
|---|---|---|---|---|---|---|---|---|---|---|---|
| YDR384C | 250.209484 | 1.33954943 | 0.52940747 | 40.4879245 | 1.61E-09 | 5.77E-08 | ATO3 | IV | 1241204 | 1E+06 | -1 |
| YDR385W | 31.6792435 | -0.1758799 | 0.84042226 | 2.28084828 | 0.3196834 | 0.49223979 | EFT1 | IV | 1243230 | 1E+06 | 1 |
| YDR386W | 35.3878877 | 0.58422444 | 0.71151612 | 5.99866446 | 0.04982033 | 0.13021013 | MUS81 | IV | 1246084 | 1E+06 | 1 |
| YDR387C | 31.542061 | -0.1658123 | 0.6976768 | 0.08681253 | 0.9575223 | 0.97008821 | CIN10 | IV | 1248154 | 1E+06 | -1 |
| YDR388W | 109.278628 | 1.10910102 | 0.6085678 | 3.29106625 | 0.19290969 | 0.34948282 | RVS167 | IV | 1250186 | 1E+06 | 1 |
| YDR389W | 42.3207474 | 0.58442265 | 0.62574269 | 2.14621837 | 0.3419437 | 0.51558698 | SAC7 | IV | 1252537 | 1E+06 | 1 |
| YDR390C | 63.2942293 | 1.82384053 | 0.74864978 | 6.02746503 | 0.04910804 | 0.12889191 | UBA2 | IV | 1254937 | 1E+06 | -1 |
| YDR391C | 12.2148509 | 1.12177824 | 1.00898405 | 5.03284152 | 0.08074811 | 0.18768829 | NA | IV | 1257358 | 1E+06 | -1 |
| YDR392W | 19.6990755 | 0.68032381 | 0.80024736 | 0.93696142 | 0.62595255 | 0.76430043 | SPT3 | IV | 1258696 | 1E+06 | 1 |
| YDR393W | 281.063151 | 0.17830342 | 0.47291623 | 1.97261606 | 0.37295108 | 0.5462653 | SHE9 | IV | 1259901 | 1E+06 | 1 |
| YDR394W | 29.8398429 | 0.31498158 | 0.82357322 | 0.20855247 | 0.90097638 | 0.93824416 | RPT3 | IV | 1261681 | 1E+06 | 1 |
| YDR395W | 64.1672337 | -0.3081477 | 0.60543434 | 0.29319573 | 0.86364121 | 0.91991868 | SXM1 | IV | 1263324 | 1E+06 | 1 |
| YDR397C | 47.8094006 | 0.28000663 | 0.59349993 | 1.77036372 | 0.41263912 | 0.58426143 | NCB2 | IV | 1266366 | 1E+06 | -1 |
| YDR398W | 691.820815 | 1.98666259 | 0.54319348 | 95.4774334 | 1.85E-21 | 6.72E-19 | UTP5 | IV | 1267471 | 1E+06 | 1 |
| YDR399W | 69.2561628 | 0.31610136 | 0.5593513 | 1.69027399 | 0.42949852 | 0.59890573 | HPT1 | IV | 1270068 | 1E+06 | 1 |
| YDR400W | 23.1586837 | 0.18635476 | 0.73694641 | 0.24247338 | 0.88582427 | 0.93185365 | URH1 | IV | 1271063 | 1E+06 | 1 |
| YDR402C | 500.253706 | 0.97610553 | 0.85984271 | 2.30359195 | 0.31606861 | 0.48830599 | DIT2 | IV | 1272234 | 1E+06 | -1 |
| YDR403W | 638.391017 | 0.99030549 | 0.53904314 | 22.466019 | 1.32E-05 | 0.00016799 | DIT1 | IV | 1274602 | 1E+06 | 1 |
| YDR404C | 76.3610355 | 0.41716031 | 0.60280294 | 11.9044314 | 0.00260007 | 0.013661 | RPB7 | IV | 1276654 | 1E+06 | -1 |
| YDR405W | 36.1791597 | 0.690201 | 0.66075398 | 6.05786584 | 0.04836722 | 0.12713301 | MRP20 | IV | 1277646 | 1E+06 | 1 |
| YDR406W | 342.054045 | -0.3474323 | 0.59867628 | 0.843519 | 0.65589176 | 0.78495521 | PDR15 | IV | 1279210 | 1E+06 | 1 |
| YDR407C | 30.1448295 | 0.01475979 | 0.68716138 | 0.22945807 | 0.89160771 | 0.93352778 | TRS120 | IV | 1284069 | 1E+06 | -1 |
| YDR408C | 200.102673 | -0.5282446 | 0.56198754 | 4.20193957 | 0.12233773 | 0.25315778 | ADE8 | IV | 1288215 | 1E+06 | -1 |
| YDR409W | 876.859297 | -0.9819321 | 0.46685778 | 8.11996024 | 0.01724936 | 0.05934272 | SIZ1 | IV | 1289406 | 1E+06 | 1 |
| YDR410C | 91.3475944 | 1.99657521 | 0.90971604 | 9.14967027 | 0.010308 | 0.04032978 | STE14 | IV | 1292372 | 1E+06 | -1 |
| YDR411C | 39.1475792 | 1.69979683 | 0.68711043 | 6.27341865 | 0.04342546 | 0.11809931 | DFM1 | IV | 1293369 | 1E+06 | -1 |
| YDR412W | 42.6492483 | -0.8012787 | 0.73246817 | 9.04180372 | 0.01087921 | 0.04193757 | RRP17 | IV | 1294693 | 1E+06 | 1 |
| YDR414C | 557.682694 | 0.56402556 | 0.49428993 | 1.45720079 | 0.48258394 | 0.644856 | ERD1 | IV | 1295598 | 1E+06 | -1 |
| YDR415C | 15.9850751 | 0.51699224 | 0.84482712 | 0.39589728 | 0.82041199 | 0.8920536 | NA | IV | 1297037 | 1E+06 | -1 |
| YDR416W | 59.2709766 | -0.6310513 | 0.5695688 | 6.32518604 | 0.04231587 | 0.11589825 | SYF1 | IV | 1298432 | 1E+06 | 1 |
| YDR418W | 179.122783 | -0.2200179 | 0.55720581 | 12.1886169 | 0.00225567 | 0.01226322 | RPL12B | IV | 1301616 | 1E+06 | 1 |
| YDR419W | 192.727971 | -0.7850748 | 0.50715349 | 2.37916907 | 0.30434768 | 0.47720463 | RAD30 | IV | 1303174 | 1E+06 | 1 |
| YDR420W | 23.6830874 | 0.23130029 | 0.74704328 | 0.63201853 | 0.7290527 | 0.83668919 | HKR1 | IV | 1306267 | 1E+06 | 1 |

| | | | | | | | | | | | |
|---|---|---|---|---|---|---|---|---|---|---|---|
| YDR421W | 186.883693 | -0.664351 | 0.58039959 | 7.16560454 | 0.02779769 | 0.08391483 | ARO80 | IV | 1312040 | 1E+06 | 1 |
| YDR422C | 64.3643335 | 0.3259017 | 0.60906128 | 24.4826429 | 4.83E-06 | 6.90E-05 | SIP1 | IV | 1315326 | 1E+06 | -1 |
| YDR423C | 24.5448454 | 1.11672639 | 0.77082235 | 3.05657473 | 0.21690683 | 0.38103476 | CAD1 | IV | 1318046 | 1E+06 | -1 |
| YDR424C | 65.1832492 | 0.1393592 | 0.57009158 | 0.06619872 | 0.96744243 | 0.97486964 | DYN2 | IV | 1319387 | 1E+06 | -1 |
| YDR425W | 30.3939495 | 1.26735277 | 0.683439 | 4.79693331 | 0.09085716 | 0.20453459 | SNX41 | IV | 1320064 | 1E+06 | 1 |
| YDR427W | 236.712341 | 0.17368787 | 0.58119501 | 11.8210096 | 0.00271082 | 0.01406419 | RPN9 | IV | 1322205 | 1E+06 | 1 |
| YDR428C | 362.801641 | 0.68097601 | 0.61444252 | 9.0751708 | 0.01069921 | 0.04143366 | BNA7 | IV | 1323454 | 1E+06 | -1 |
| YDR429C | 40.3697365 | -0.4798153 | 0.76025907 | 8.09997031 | 0.01742263 | 0.05974247 | TIF35 | IV | 1324477 | 1E+06 | -1 |
| YDR430C | 20.0190041 | -0.542741 | 0.72863431 | 0.87086933 | 0.64698339 | 0.77880121 | CYM1 | IV | 1325501 | 1E+06 | -1 |
| YDR432W | 183.436973 | 0.97717453 | 0.60305574 | 3.32465227 | 0.18969721 | 0.34571823 | NPL3 | IV | 1328783 | 1E+06 | 1 |
| YDR434W | 112.063018 | 2.12808773 | 0.69533742 | 12.3271562 | 0.00210471 | 0.01162812 | GPI17 | IV | 1331237 | 1E+06 | 1 |
| YDR435C | 14.4005583 | 1.67730808 | 0.84518822 | 4.1215517 | 0.12735512 | 0.26092929 | PPM1 | IV | 1332983 | 1E+06 | -1 |
| YDR436W | 84.9202402 | -0.5231372 | 0.60084425 | 11.8067806 | 0.00273017 | 0.01413927 | PPZ2 | IV | 1334821 | 1E+06 | 1 |
| YDR437W | 30.0280533 | -0.0767338 | 0.68416541 | 0.07652516 | 0.96246018 | 0.97258112 | GPI19 | IV | 1337352 | 1E+06 | 1 |
| YDR438W | 366.265586 | 0.57236093 | 0.57602241 | 4.01197125 | 0.13452764 | 0.27098851 | THI74 | IV | 1338274 | 1E+06 | 1 |
| YDR439W | 47.2765003 | 1.19817416 | 0.74843924 | 3.9902872 | 0.13599412 | 0.27312035 | LRS4 | IV | 1339676 | 1E+06 | 1 |
| YDR440W | 51.0372256 | 0.36132293 | 0.79086099 | 1.29007239 | 0.52464355 | 0.67987605 | DOT1 | IV | 1342493 | 1E+06 | 1 |
| YDR441C | 242.384671 | -0.0506471 | 0.51514625 | 1.62613244 | 0.44349612 | 0.61162622 | APT2 | IV | 1344517 | 1E+06 | -1 |
| YDR443C | 23.3769063 | 0.20682003 | 0.70077277 | 0.11556317 | 0.94385608 | 0.96197675 | SSN2 | IV | 1345676 | 1E+06 | -1 |
| YDR444W | 46.014738 | -0.0745526 | 0.70084289 | 16.841491 | 0.00022025 | 0.001854 | NA | IV | 1350290 | 1E+06 | 1 |
| YDR446W | 88.901944 | 0.59491808 | 0.74609974 | 22.0560907 | 1.62E-05 | 0.00019879 | ECM11 | IV | 1353725 | 1E+06 | 1 |
| YDR447C | 47.8813604 | -0.6954864 | 0.74978207 | 12.7268227 | 0.00172348 | 0.01001901 | RPS17B | IV | 1354829 | 1E+06 | -1 |
| YDR448W | 167.879642 | -1.5101074 | 0.50190547 | 41.1635014 | 1.15E-09 | 4.22E-08 | ADA2 | IV | 1356065 | 1E+06 | 1 |
| YDR449C | 18.2800135 | -1.7153619 | 0.85534985 | 9.68734821 | 0.00787806 | 0.03281579 | UTP6 | IV | 1357580 | 1E+06 | -1 |
| YDR450W | 79.4203755 | -0.6310366 | 1.97114788 | 2.85142558 | 0.24033709 | 0.40843903 | RPS18B | IV | 1359923 | 1E+06 | 1 |
| YDR451C | 166.235535 | -0.19864 | 0.50019693 | 0.55333371 | 0.75830708 | 0.85403579 | YHP1 | IV | 1361120 | 1E+06 | -1 |
| YDR452W | 1456.9095 | -1.1742992 | 0.4873224 | 23.6138779 | 7.45E-06 | 0.00010106 | PPN1 | IV | 1362878 | 1E+06 | 1 |
| YDR453C | 81.2840014 | 1.08028106 | 0.83307142 | 7.13122589 | 0.02827965 | 0.08501514 | TSA2 | IV | 1365072 | 1E+06 | -1 |
| YDR454C | 49.5704188 | -1.623779 | 0.61480869 | 7.24896761 | 0.02666286 | 0.08136057 | GUK1 | IV | 1366264 | 1E+06 | -1 |
| YDR456W | 127.309557 | 0.62597524 | 0.54321816 | 11.4615587 | 0.00324455 | 0.01625081 | NHX1 | IV | 1367485 | 1E+06 | 1 |
| YDR457W | 41.7733416 | -0.3628343 | 0.71416779 | 0.47002305 | 0.79056174 | 0.87307888 | TOM1 | IV | 1369790 | 1E+06 | 1 |
| YDR458C | 55.8958173 | -0.3476628 | 0.56720435 | 0.68490283 | 0.71002761 | 0.82438173 | HEH2 | IV | 1380055 | 1E+06 | -1 |
| YDR459C | 16.4619699 | 0.57126277 | 0.94298241 | 0.43635784 | 0.80398158 | 0.88247457 | PFA5 | IV | 1382319 | 1E+06 | -1 |

| | | | | | | | | | | | |
|---|---|---|---|---|---|---|---|---|---|---|---|
| YDR460W | 23.7977729 | 0.43045882 | 0.71293929 | 1.46571173 | 0.48053469 | 0.64330539 | TFB3 | IV | 1383811 | 1E+06 | 1 |
| YDR462W | 9.74320008 | -0.954169 | 0.94627319 | 2.02576046 | 0.36317145 | 0.53614552 | MRPL28 | IV | 1386073 | 1E+06 | 1 |
| YDR463W | 27.4529557 | 0.73082852 | 0.66579186 | 1.3836789 | 0.50065429 | 0.66001556 | STP1 | IV | 1386816 | 1E+06 | 1 |
| YDR464W | 87.7171636 | 0.1752369 | 0.59296562 | 6.08634473 | 0.04768338 | 0.12600836 | SPP41 | IV | 1388872 | 1E+06 | 1 |
| YDR465C | 91.4595215 | 0.28714065 | 0.61375359 | 11.9944255 | 0.00248567 | 0.01316746 | RMT2 | IV | 1393336 | 1E+06 | -1 |
| YDR466W | 278.549352 | -0.3330354 | 0.4716219 | 14.4537814 | 0.00072678 | 0.00503957 | PKH3 | IV | 1395121 | 1E+06 | 1 |
| YDR468C | 155.125067 | 1.30164537 | 0.78729741 | 6.6088863 | 0.03671965 | 0.10437251 | TLG1 | IV | 1398026 | 1E+06 | -1 |
| YDR469W | 85.0700435 | 1.61116452 | 0.74800608 | 6.10250765 | 0.04729958 | 0.12530377 | SDC1 | IV | 1399015 | 1E+06 | 1 |
| YDR470C | 165.299793 | -0.4574299 | 0.51463937 | 11.4437753 | 0.00327353 | 0.01636734 | UGO1 | IV | 1399706 | 1E+06 | -1 |
| YDR471W | 80.8401678 | -0.7084732 | 0.60107849 | 21.114373 | 2.60E-05 | 0.00029817 | RPL27B | IV | 1401770 | 1E+06 | 1 |
| YDR472W | 85.4094283 | -1.2240973 | 0.66931846 | 3.78215391 | 0.1509092 | 0.29389985 | TRS31 | IV | 1403322 | 1E+06 | 1 |
| YDR473C | 71.802308 | 0.30087893 | 0.5792382 | 0.39019657 | 0.82275379 | 0.89359303 | PRP3 | IV | 1404445 | 1E+06 | -1 |
| YDR475C | 412.200435 | -1.0940063 | 0.46769629 | 25.8250094 | 2.47E-06 | 3.86E-05 | JIP4 | IV | 1407464 | 1E+06 | -1 |
| YDR476C | 101.238069 | -1.1178514 | 0.56823677 | 9.78218448 | 0.00751321 | 0.03164938 | NA | IV | 1410453 | 1E+06 | -1 |
| YDR477W | 41.8219356 | 0.56824989 | 0.61260675 | 1.55286241 | 0.46004489 | 0.62448065 | SNF1 | IV | 1412373 | 1E+06 | 1 |
| YDR478W | 206.103414 | 2.1535785 | 0.70516068 | 9.04048462 | 0.01088639 | 0.04193757 | SNM1 | IV | 1414575 | 1E+06 | 1 |
| YDR479C | 679.294326 | -1.1555119 | 0.47497118 | 16.1744062 | 0.00030745 | 0.00249667 | PEX29 | IV | 1415210 | 1E+06 | -1 |
| YDR480W | 124.784615 | -0.7134046 | 0.54330479 | 4.81089492 | 0.09022511 | 0.20366605 | DIG2 | IV | 1417399 | 1E+06 | 1 |
| YDR481C | 198.427545 | -0.7717489 | 0.48329045 | 13.9147716 | 0.00095158 | 0.00631841 | PHO8 | IV | 1418550 | 1E+06 | -1 |
| YDR482C | 18.0635056 | 0.34564586 | 0.74570714 | 1.03212625 | 0.59686571 | 0.74415428 | CWC21 | IV | 1420431 | 1E+06 | -1 |
| YDR483W | 47.8625355 | 0.60835226 | 0.58598927 | 1.10063348 | 0.5767671 | 0.72593834 | KRE2 | IV | 1421157 | 1E+06 | 1 |
| YDR484W | 39.5615912 | 1.10963464 | 0.72367464 | 3.07895604 | 0.21449303 | 0.37828653 | VPS52 | IV | 1422763 | 1E+06 | 1 |
| YDR485C | 49.0156531 | 0.56037177 | 0.6195954 | 1.33696554 | 0.51248555 | 0.66951519 | VPS72 | IV | 1424820 | 1E+06 | -1 |
| YDR486C | 37.1347812 | 0.91240158 | 0.63171673 | 2.23479937 | 0.32712933 | 0.49988884 | VPS60 | IV | 1427431 | 1E+06 | -1 |
| YDR487C | 61.4657002 | 0.29365538 | 0.55689687 | 0.51236509 | 0.77400067 | 0.86281553 | RIB3 | IV | 1428354 | 1E+06 | -1 |
| YDR488C | 6.23681274 | -0.4350942 | 1.12908273 | 0.23681466 | 0.88833413 | 0.93279917 | PAC11 | IV | 1429188 | 1E+06 | -1 |
| YDR489W | 182.604605 | 0.46169848 | 0.52278412 | 2.45961915 | 0.29234824 | 0.46643602 | SLD5 | IV | 1431012 | 1E+06 | 1 |
| YDR490C | 655.772739 | -1.0923437 | 0.50100932 | 28.8493844 | 5.44E-07 | 1.04E-05 | PKH1 | IV | 1431968 | 1E+06 | -1 |
| YDR492W | 13.7360166 | -0.3616045 | 0.89424857 | 3.63770535 | 0.16221175 | 0.31078956 | IZH1 | IV | 1434924 | 1E+06 | 1 |
| YDR494W | 96.4198848 | 1.05111169 | 0.72450291 | 8.03396241 | 0.01800724 | 0.06111485 | RSM28 | IV | 1436930 | 1E+06 | 1 |
| YDR495C | 163.201288 | -1.7074984 | 0.51890553 | 16.6310425 | 0.00024469 | 0.00203556 | VPS3 | IV | 1438115 | 1E+06 | -1 |
| YDR496C | 32.104409 | -1.3335671 | 0.69300317 | 9.60218521 | 0.00822076 | 0.03390367 | PUF6 | IV | 1441433 | 1E+06 | -1 |
| YDR497C | 38.4561229 | 0.60717287 | 0.63023959 | 1.61683001 | 0.44556372 | 0.61307366 | ITR1 | IV | 1443713 | 1E+06 | -1 |

| | | | | | | | | | | | |
|---|---|---|---|---|---|---|---|---|---|---|---|
| YDR498C | 60.8878415 | -0.1719223 | 0.6078765 | 0.16575092 | 0.92046578 | 0.95024857 | SEC20 | IV | 1445843 | 1E+06 | -1 |
| YDR499W | 61.4240601 | -0.0225198 | 0.65549038 | 11.3796591 | 0.00338017 | 0.0167275 | LCD1 | IV | 1447830 | 1E+06 | 1 |
| YDR500C | 46.5357075 | -0.3582812 | 0.76063399 | 10.176656 | 0.00616832 | 0.02707703 | RPL37B | IV | 1450198 | 1E+06 | -1 |
| YDR501W | 290.791727 | 0.0014113 | 0.52359231 | 3.2096209 | 0.20092764 | 0.36006531 | PLM2 | IV | 1451353 | 1E+06 | 1 |
| YDR502C | 26.9680007 | -0.4662322 | 0.81644586 | 0.35316376 | 0.83813015 | 0.90469306 | SAM2 | IV | 1453310 | 1E+06 | -1 |
| YDR503C | 10.9543258 | -0.472545 | 1.00605796 | 1.74513064 | 0.41787818 | 0.58926319 | LPP1 | IV | 1455042 | 1E+06 | -1 |
| YDR504C | 213.079154 | -1.2444533 | 0.53396022 | 24.4794037 | 4.83E-06 | 6.90E-05 | SPG3 | IV | 1456311 | 1E+06 | -1 |
| YDR505C | 31.8644236 | 0.01827817 | 0.66878315 | 0.14029941 | 0.93225425 | 0.95731799 | PSP1 | IV | 1456695 | 1E+06 | -1 |
| YDR506C | 107.967187 | -0.6330901 | 0.51355834 | 1.50645495 | 0.47084446 | 0.63296713 | GMC1 | IV | 1459728 | 1E+06 | -1 |
| YDR507C | 782.888462 | 2.47855084 | 0.48762177 | 64.5900929 | 9.43E-15 | 9.25E-13 | GIN4 | IV | 1462358 | 1E+06 | -1 |
| YDR508C | 65.1517139 | 0.27165014 | 0.68319018 | 9.28198545 | 0.00964811 | 0.03839103 | GNP1 | IV | 1466453 | 1E+06 | -1 |
| YDR510W | 29.6747605 | 0.17942239 | 0.73789499 | 2.28047345 | 0.31974332 | 0.49223979 | SMT3 | IV | 1469400 | 1E+06 | 1 |
| YDR511W | 183.187563 | 1.70945414 | 0.77588075 | 6.59096553 | 0.03705015 | 0.10484868 | SDH7 | IV | 1470017 | 1E+06 | 1 |
| YDR512C | 2366.83895 | 0.15717343 | 0.47896273 | 0.61555873 | 0.73507748 | 0.83830975 | EMI1 | IV | 1470500 | 1E+06 | -1 |
| YDR513W | 11.4759735 | -0.5695252 | 0.99293916 | 0.33984426 | 0.84373052 | 0.90836736 | GRX2 | IV | 1471017 | 1E+06 | 1 |
| YDR514C | 97.1348473 | 2.42006393 | 0.97695163 | 8.17487422 | 0.01678219 | 0.05813116 | NA | IV | 1471548 | 1E+06 | -1 |
| YDR515W | 195.698393 | 1.46453589 | 0.6360444 | 67.8520408 | 1.85E-15 | 2.14E-13 | SLF1 | IV | 1473429 | 1E+06 | 1 |
| YDR516C | 144.010611 | -0.081935 | 0.52689403 | 16.0883409 | 0.00032097 | 0.0025883 | EMI2 | IV | 1474974 | 1E+06 | -1 |
| YDR517W | 46.7196339 | 0.37577674 | 0.6185595 | 3.71093067 | 0.15638016 | 0.30191434 | GRH1 | IV | 1477239 | 1E+06 | 1 |
| YDR518W | 335.848086 | 0.39351747 | 0.56183064 | 38.4724767 | 4.42E-09 | 1.44E-07 | EUG1 | IV | 1478608 | 1E+06 | 1 |
| YDR519W | 92.9916488 | 0.00897899 | 0.67717612 | 0.49458073 | 0.78091391 | 0.86705973 | FPR2 | IV | 1480425 | 1E+06 | 1 |
| YDR520C | 453.089386 | 1.73109959 | 0.65644776 | 7.87031169 | 0.01954265 | 0.06499882 | URC2 | IV | 1481085 | 1E+06 | -1 |
| YDR522C | 303.509566 | 0.06330602 | 0.55615434 | 0.20032064 | 0.90469237 | 0.94008773 | SPS2 | IV | 1483795 | 1E+06 | -1 |
| YDR523C | 11.5972407 | 0.52822037 | 1.13262438 | 2.47759374 | 0.28973259 | 0.46366825 | SPS1 | IV | 1485566 | 1E+06 | -1 |
| YDR524C | 113.452542 | 0.85307135 | 0.53622738 | 5.45765557 | 0.06529579 | 0.15958742 | AGE1 | IV | 1487542 | 1E+06 | -1 |
| YDR524C-B | 92.1323003 | -0.826486 | 0.70346007 | 4.07424882 | 0.13040316 | 0.26529666 | NA | IV | 1489598 | 1E+06 | -1 |
| YDR524W-C | 71688.527 | -0.3179883 | 0.50074983 | 15.5802559 | 0.0004138 | 0.00315666 | NA | IV | 1489403 | 1E+06 | 1 |
| YDR525W | 41.4819871 | -1.0390747 | 0.66602242 | 2.43512891 | 0.29595009 | 0.46995614 | NA | IV | 1489905 | 1E+06 | 1 |
| YDR525W-A | 13.3749558 | -1.1647483 | 0.90657693 | 2.11737096 | 0.34691153 | 0.5203673 | SNA2 | IV | 1490596 | 1E+06 | 1 |
| YDR527W | 82.6099973 | -0.5593787 | 0.615522 | 10.4841894 | 0.00528917 | 0.02396265 | RBA50 | IV | 1491094 | 1E+06 | 1 |
| YDR528W | 38.0072311 | -0.0124592 | 0.6827904 | 4.63316684 | 0.09860992 | 0.2170918 | HLR1 | IV | 1494586 | 1E+06 | 1 |
| YDR529C | 109.11031 | 0.28105789 | 0.60698048 | 2.23890501 | 0.32645848 | 0.49927259 | QCR7 | IV | 1496165 | 1E+06 | -1 |
| YDR530C | 11.5487157 | 1.02631212 | 0.88181288 | 1.4279184 | 0.48970153 | 0.65106127 | APA2 | IV | 1496791 | 1E+06 | -1 |

| | | | | | | | | | | | |
|---|---|---|---|---|---|---|---|---|---|---|---|
| YDR531W | 20.3198999 | 1.23213658 | 0.86264889 | 5.93695457 | 0.05138149 | 0.13358726 | CAB1 | IV | 1498232 | 1E+06 | 1 |
| YDR532C | 121.981467 | 2.47554004 | 0.93291134 | 10.5010286 | 0.00524482 | 0.02385369 | KRE28 | IV | 1499396 | 2E+06 | -1 |
| YDR533C | 101.205409 | -1.2457577 | 0.5733777 | 11.3444006 | 0.00344029 | 0.016967 | HSP31 | IV | 1501447 | 2E+06 | -1 |
| YDR534C | 7.19081197 | -0.0019335 | 1.3364136 | 6.43624071 | 0.04003023 | 0.1114839 | FIT1 | IV | 1503314 | 2E+06 | -1 |
| YDR536W | 734.822717 | -0.1983794 | 0.49535884 | 0.38546685 | 0.82470179 | 0.89537284 | STL1 | IV | 1508005 | 2E+06 | 1 |
| YDR539W | 11.3666958 | 0.00574335 | 0.97771436 | 0.06744877 | 0.96683794 | 0.97486964 | FDC1 | IV | 1512094 | 2E+06 | 1 |
| YDR540C | 84.7948889 | -0.8149704 | 0.56556764 | 3.40668037 | 0.18207434 | 0.33713158 | IRC4 | IV | 1517136 | 2E+06 | -1 |
| YDR541C | 103.76907 | 1.17592588 | 0.56427374 | 24.7214174 | 4.28E-06 | 6.26E-05 | NA | IV | 1519664 | 2E+06 | -1 |
| YEL001C | 334.341439 | 0.72033098 | 0.47402918 | 3.92603635 | 0.14043393 | 0.2789408 | IRC22 | V | 150301 | 150978 | -1 |
| YEL002C | 71.0830103 | 0.74625112 | 0.63627941 | 1.51800052 | 0.46813421 | 0.63108197 | WBP1 | V | 148722 | 150014 | -1 |
| YEL003W | 106.66296 | 2.38960248 | 1.00650938 | 8.3548487 | 0.01533796 | 0.05444929 | GIM4 | V | 148176 | 148599 | 1 |
| YEL004W | 199.797505 | -0.8776025 | 0.55247662 | 2.51689351 | 0.28409495 | 0.45768219 | YEA4 | V | 146951 | 147979 | 1 |
| YEL005C | 59.7027959 | -0.4679929 | 0.56700758 | 0.87373151 | 0.64605816 | 0.77803336 | VAB2 | V | 145907 | 146755 | -1 |
| YEL006W | 11.8299939 | 0.96073479 | 1.04760356 | 1.69060424 | 0.4294276 | 0.59890573 | YEA6 | V | 144327 | 145334 | 1 |
| YEL007W | 31.6733844 | -0.0504277 | 0.67304717 | 2.921963 | 0.23200845 | 0.39920621 | MIT1 | V | 141892 | 143892 | 1 |
| YEL008W | 257.313017 | -0.3318524 | 0.64090864 | 1.44430148 | 0.4857065 | 0.64773825 | NA | V | 140512 | 140892 | 1 |
| YEL009C | 58.5431425 | -0.1963314 | 0.72944029 | 1.15434386 | 0.56148404 | 0.71278066 | GCN4 | V | 138918 | 139763 | -1 |
| YEL009C-A | 8.80277354 | 0.01151063 | 1.01621905 | 1.41805766 | 0.4921219 | 0.6530873 | NA | V | 136371 | 136778 | -1 |
| YEL011W | 354.920885 | -1.4220173 | 0.52877252 | 19.9369042 | 4.69E-05 | 0.00048706 | GLC3 | V | 133120 | 135234 | 1 |
| YEL012W | 39.5710862 | 2.5588279 | 0.78735416 | 9.94607729 | 0.00692208 | 0.02979712 | UBC8 | V | 131772 | 132551 | 1 |
| YEL013W | 30.1812974 | 0.46355256 | 0.71717597 | 0.60028923 | 0.74071109 | 0.84208075 | VAC8 | V | 128825 | 130561 | 1 |
| YEL014C | 965.789855 | -0.0242205 | 0.51385215 | 6.40742318 | 0.04061119 | 0.11266833 | NA | V | 128303 | 128608 | -1 |
| YEL015W | 61.9439591 | 0.47723952 | 0.58174919 | 5.19886178 | 0.07431586 | 0.17629168 | EDC3 | V | 126629 | 128284 | 1 |
| YEL016C | 1157.15296 | 1.03483077 | 0.47207371 | 23.7339618 | 7.02E-06 | 9.56E-05 | NPP2 | V | 124737 | 126218 | -1 |
| YEL017C-A | 35.6311622 | -0.4774148 | 0.66608257 | 6.48389044 | 0.03908779 | 0.10976769 | PMP2 | V | 122798 | 122929 | -1 |
| YEL017W | 256.635516 | 0.85180977 | 0.86097019 | 4.72033582 | 0.09440437 | 0.20984715 | GTT3 | V | 123657 | 124670 | 1 |
| YEL018W | 41.3043116 | 0.3387303 | 0.67920985 | 4.56118436 | 0.10222365 | 0.22276062 | EAF5 | V | 121471 | 122310 | 1 |
| YEL020C | 24.9584881 | 1.68049032 | 0.76251349 | 4.81880731 | 0.08986887 | 0.20341703 | PXP1 | V | 118617 | 120299 | -1 |
| YEL020W-A | 18.7887843 | -0.3796957 | 0.8006153 | 2.98712733 | 0.22457093 | 0.39000171 | TIM9 | V | 117211 | 117474 | 1 |
| YEL021W | 131.258755 | 0.61632015 | 0.58294911 | 12.5204096 | 0.00191085 | 0.010794 | URA3 | V | 116167 | 116970 | 1 |
| YEL022W | 162.353839 | 0.7246735 | 0.5987863 | 43.0805948 | 4.42E-10 | 1.75E-08 | GEA2 | V | 111421 | 115800 | 1 |
| YEL023C | 67.0229087 | 1.85059541 | 0.9132584 | 9.33841952 | 0.00937968 | 0.03760965 | NA | V | 108504 | 110552 | -1 |
| YEL024W | 12.2085682 | -0.5406268 | 1.06137095 | 1.20767101 | 0.5467107 | 0.6993379 | RIP1 | V | 107260 | 107907 | 1 |

| | | | | | | | | | | | |
|---|---|---|---|---|---|---|---|---|---|---|---|
| YEL025C | 21.5367859 | -1.337618 | 0.81417979 | 3.6836856 | 0.15852503 | 0.3051396 | NA | V | 102581 | 106147 | -1 |
| YEL026W | 51.9787036 | 1.63066555 | 0.93672553 | 4.55563811 | 0.10250753 | 0.22304343 | SNU13 | V | 101943 | 102323 | 1 |
| YEL027W | 57.5917917 | 0.27286147 | 0.60523851 | 7.66035086 | 0.02170581 | 0.07037316 | VMA3 | V | 100769 | 101251 | 1 |
| YEL028W | 114.851982 | -0.5073736 | 0.60102346 | 1.88763953 | 0.38913857 | 0.56089429 | NA | V | 98668 | 99129 | 1 |
| YEL029C | 69.2822968 | -0.2189619 | 0.60915671 | 2.77678801 | 0.24947564 | 0.42048038 | BUD16 | V | 96858 | 97796 | -1 |
| YEL030W | 101.731766 | -0.7008114 | 0.56736835 | 1.85504264 | 0.39553289 | 0.56679648 | ECM10 | V | 94644 | 96578 | 1 |
| YEL031W | 39.0310599 | 0.38182968 | 0.84084416 | 0.94228775 | 0.62428775 | 0.7633846 | SPF1 | V | 90258 | 93905 | 1 |
| YEL032W | 22.3978604 | -1.1315161 | 0.76239302 | 7.36421218 | 0.02516991 | 0.07818335 | MCM3 | V | 86937 | 89852 | 1 |
| YEL033W | 118.715667 | -0.2539971 | 0.57173344 | 16.3591676 | 0.00028032 | 0.00230546 | MTC7 | V | 86179 | 86598 | 1 |
| YEL034W | 211.122941 | -0.1172794 | 0.53574386 | 14.152209 | 0.00084506 | 0.00572938 | HYP2 | V | 85676 | 86149 | 1 |
| YEL035C | 119.429613 | 0.1681509 | 0.73150029 | 0.23214863 | 0.89040905 | 0.93317728 | UTR5 | V | 85045 | 85545 | -1 |
| YEL036C | 131.007072 | 1.17837083 | 0.69084958 | 3.32320907 | 0.18983414 | 0.34583253 | ANP1 | V | 83050 | 84552 | -1 |
| YEL037C | 63.8755901 | -0.0113587 | 0.68154916 | 3.32868245 | 0.18931533 | 0.34544902 | RAD23 | V | 81407 | 82603 | -1 |
| YEL038W | 7.43708748 | 0.22375788 | 1.03388041 | 0.59686617 | 0.74197993 | 0.84243165 | UTR4 | V | 80462 | 81145 | 1 |
| YEL039C | 13.4990488 | 0.48055686 | 0.87736246 | 0.98374697 | 0.61147972 | 0.75489714 | CYC7 | V | 79636 | 79977 | -1 |
| YEL040W | 81.8173603 | -0.4182566 | 0.66363109 | 0.62749345 | 0.73070407 | 0.83678999 | UTR2 | V | 78053 | 79456 | 1 |
| YEL041W | 278.574714 | -0.3272461 | 0.48155143 | 6.48145759 | 0.03913536 | 0.10983701 | YEF1 | V | 75944 | 77431 | 1 |
| YEL042W | 57.1932105 | 0.02637386 | 0.73506748 | 9.33671129 | 0.00938769 | 0.03761574 | GDA1 | V | 73771 | 75327 | 1 |
| YEL043W | 37.2430899 | -0.3290135 | 0.65044456 | 1.67715946 | 0.4323241 | 0.60114231 | NA | V | 70478 | 73348 | 1 |
| YEL044W | 57.9697136 | -0.772731 | 0.56928105 | 3.8780161 | 0.14384657 | 0.28410385 | IES6 | V | 69757 | 70257 | 1 |
| YEL046C | 67.145133 | 0.29186415 | 0.58939368 | 2.68597711 | 0.2610643 | 0.43235173 | GLY1 | V | 67629 | 68792 | -1 |
| YEL047C | 24.178703 | -0.7964359 | 0.79223052 | 1.76763784 | 0.41320191 | 0.58466253 | FRD1 | V | 65385 | 66797 | -1 |
| YEL048C | 82.5617357 | 0.94355028 | 0.67974049 | 43.6547506 | 3.32E-10 | 1.36E-08 | TCA17 | V | 64709 | 65167 | -1 |
| YEL049W | 224.841917 | -1.1021305 | 0.49629721 | 8.91480643 | 0.01159243 | 0.0440421 | PAU2 | V | 63728 | 64090 | 1 |
| YEL050C | 45.2804867 | -0.5880879 | 0.60507875 | 2.06546128 | 0.35603343 | 0.52942744 | RML2 | V | 59670 | 60851 | -1 |
| YEL051W | 21.4255817 | -0.008103 | 0.88641832 | 2.60628356 | 0.2716769 | 0.44510732 | VMA8 | V | 58378 | 59148 | 1 |
| YEL052W | 18.7672271 | -0.7602938 | 0.85931522 | 1.07677398 | 0.58368899 | 0.73199919 | AFG1 | V | 56571 | 58100 | 1 |
| YEL053C | 24.1836648 | -1.1199156 | 0.76455951 | 3.50039479 | 0.17373964 | 0.32507061 | MAK10 | V | 53901 | 56102 | -1 |
| YEL054C | 109.073932 | -0.8991098 | 0.59806681 | 22.0167336 | 1.66E-05 | 0.00020089 | RPL12B | V | 52721 | 53218 | -1 |
| YEL055C | 695.87554 | 0.18354271 | 0.52732262 | 0.13110492 | 0.93654992 | 0.95873833 | POL5 | V | 48471 | 51539 | -1 |
| YEL056W | 270.431035 | -1.120164 | 0.48785349 | 37.3057663 | 7.93E-09 | 2.47E-07 | HAT2 | V | 47168 | 48373 | 1 |
| YEL057C | 248.856912 | 0.60797137 | 0.52114019 | 22.3024058 | 1.44E-05 | 0.00018033 | SDD1 | V | 45020 | 45721 | -1 |
| YEL058W | 117.167732 | 1.48371763 | 0.52081105 | 8.1443771 | 0.01704005 | 0.05876231 | PCM1 | V | 43252 | 44925 | 1 |

| | | | | | | | | | | | |
|---|---|---|---|---|---|---|---|---|---|---|---|
| YEL059C-A | 76.204023 | 1.44962088 | 0.56504812 | 8.87524218 | 0.01182403 | 0.04465829 | SOM1 | V | 42400 | 42624 | -1 |
| YEL059W | 982.020067 | 2.04900762 | 0.60216242 | 15.7357703 | 0.00038284 | 0.00297068 | NA | V | 42652 | 42960 | 1 |
| YEL060C | 603.363889 | 0.53354857 | 0.45678735 | 1.78442751 | 0.40974767 | 0.58133766 | PRB1 | V | 40046 | 41953 | -1 |
| YEL061C | 17.0691406 | -0.3501968 | 0.77682512 | 0.32559425 | 0.84976357 | 0.91079805 | CIN8 | V | 36535 | 39537 | -1 |
| YEL062W | 62.750231 | -0.5596021 | 0.62375153 | 15.8352929 | 0.00036426 | 0.00285226 | NPR2 | V | 34407 | 36254 | 1 |
| YEL063C | 327.659734 | 0.76019451 | 0.62030638 | 80.208346 | 3.83E-18 | 6.93E-16 | CAN1 | V | 31694 | 33466 | -1 |
| YEL064C | 79.2258103 | -0.7476384 | 0.58583976 | 5.66970174 | 0.05872728 | 0.14739097 | AVT2 | V | 29797 | 31239 | -1 |
| YEL065W | 25.2926885 | 0.98954308 | 0.93000125 | 5.16176646 | 0.07570711 | 0.17861909 | SIT1 | V | 27657 | 29543 | 1 |
| YEL066W | 35.9052292 | -1.8904252 | 0.68390652 | 20.5606113 | 3.43E-05 | 0.00037758 | NA | V | 26667 | 27206 | 1 |
| YEL067C | 532.526596 | -1.2462668 | 0.59459607 | 8.53164215 | 0.01404033 | 0.0509358 | NA | V | 26189 | 26776 | -1 |
| YEL069C | 211.190071 | -0.6637945 | 0.71277341 | 2.71627758 | 0.25713892 | 0.42955405 | HXT13 | V | 21537 | 23231 | -1 |
| YEL071W | 50.0681482 | 1.341585 | 0.74416877 | 3.29171546 | 0.19284708 | 0.34947875 | DLD3 | V | 16355 | 17845 | 1 |
| YEL073C | 12.6565869 | -1.4494793 | 0.90619084 | 2.62289299 | 0.26943004 | 0.44259056 | NA | V | 7230 | 7553 | -1 |
| YEL075C | 34.8718413 | -0.6624865 | 0.74676305 | 3.73898074 | 0.15420223 | 0.29860565 | NA | V | 5345 | 5713 | -1 |
| YEL077C | 42.2664748 | -1.0839142 | 0.65701458 | 2.90977851 | 0.23342621 | 0.40082886 | NA | V | 264 | 4097 | -1 |
| YER001W | 18.0438259 | -0.6380701 | 0.87515269 | 1.38156815 | 0.50118295 | 0.66037092 | MNN1 | V | 153520 | 155808 | 1 |
| YER002W | 107.938359 | 1.17002066 | 0.71648027 | 11.7127518 | 0.0028616 | 0.0147146 | NOP16 | V | 156803 | 157498 | 1 |
| YER003C | 32.2388659 | -0.0456161 | 0.68713498 | 1.59072015 | 0.45141866 | 0.61775326 | PMI40 | V | 157736 | 159118 | -1 |
| YER004W | 37.2066582 | -0.0465344 | 0.63852222 | 0.21368581 | 0.89866684 | 0.9375281 | FMP52 | V | 159580 | 160275 | 1 |
| YER005W | 781.007076 | 0.04054943 | 0.48400858 | 0.01066792 | 0.99468024 | 0.99571207 | YND1 | V | 160550 | 162442 | 1 |
| YER006W | 32.7792136 | -0.7600373 | 0.82136676 | 6.74073012 | 0.03437709 | 0.09926596 | NUG1 | V | 162723 | 164285 | 1 |
| YER007C-A | 53.6393664 | -0.3326378 | 0.5829597 | 0.71594565 | 0.69909207 | 0.81772588 | TMA20 | V | 166237 | 166885 | -1 |
| YER007W | 264.13632 | 0.11870452 | 0.57063482 | 4.35060011 | 0.11357407 | 0.24017307 | PAC2 | V | 164527 | 166083 | 1 |
| YER008C | 128.39489 | 0.54942237 | 0.59407865 | 1.1809538 | 0.55406299 | 0.70568075 | SEC3 | V | 167808 | 171818 | -1 |
| YER009W | 25.1105998 | -0.9569517 | 0.87184247 | 3.32683992 | 0.18948982 | 0.34544902 | NTF2 | V | 172115 | 172492 | 1 |
| YER010C | 2347.02392 | 3.4759501 | 1.94556775 | 4.60506215 | 0.1000054 | 0.21919967 | NA | V | 172634 | 173338 | -1 |
| YER011W | 21.8325162 | -0.6459731 | 0.76276825 | 2.05120952 | 0.35857955 | 0.53153496 | TIR1 | V | 175248 | 176012 | 1 |
| YER012W | 18.955544 | 0.01605877 | 0.82018319 | 0.77745964 | 0.67791741 | 0.80121286 | PRE1 | V | 177835 | 178431 | 1 |
| YER013W | 14.5510013 | -0.2711372 | 0.8639041 | 0.23945475 | 0.88716227 | 0.93215081 | PRP22 | V | 178841 | 182278 | 1 |
| YER014C-A | 1107.88932 | 0.61048808 | 0.48359528 | 2.10005343 | 0.3499284 | 0.52354248 | BUD25 | V | 183731 | 184700 | -1 |
| YER014W | 11.9309162 | -0.0197183 | 0.98097569 | 0.11470317 | 0.94426202 | 0.96220998 | HEM14 | V | 182600 | 184219 | 1 |
| YER015W | 570.477259 | -0.4788344 | 0.48424036 | 2.24390357 | 0.32564359 | 0.4985395 | FAA2 | V | 184541 | 186775 | 1 |
| YER016W | 52.078497 | -0.2567634 | 0.61689753 | 0.21980017 | 0.89592365 | 0.93584664 | BIM1 | V | 188277 | 189311 | 1 |

| | | | | | | | | | | | |
|---|---|---|---|---|---|---|---|---|---|---|---|
| YER017C | 38.1497047 | -0.7494789 | 0.79331101 | 1.84683475 | 0.39715948 | 0.56793119 | AFG3 | V | 189503 | 191788 | -1 |
| YER018C | 173.571655 | -0.3658535 | 0.57612509 | 0.52478845 | 0.76920772 | 0.85953436 | SPC25 | V | 191959 | 192624 | -1 |
| YER019C-A | 388.560547 | 1.43120502 | 0.62423795 | 7.25815164 | 0.0265407 | 0.08104993 | SBH2 | V | 194273 | 194539 | -1 |
| YER019W | 822.706836 | 1.17257166 | 0.78963735 | 2.18529322 | 0.33532784 | 0.50865815 | ISC1 | V | 192797 | 194230 | 1 |
| YER020W | 87.1020737 | 1.8562349 | 0.63573871 | 10.3431899 | 0.00567551 | 0.02535586 | GPA2 | V | 195168 | 196517 | 1 |
| YER021W | 31.1774024 | -0.2211667 | 0.75401606 | 1.07629292 | 0.5838294 | 0.73199919 | RPN3 | V | 196948 | 198519 | 1 |
| YER022W | 17.6630435 | 0.28673759 | 0.85729282 | 0.21813677 | 0.8966691 | 0.93645636 | SRB4 | V | 198812 | 200875 | 1 |
| YER023W | 29.8159364 | -0.274989 | 0.6504036 | 2.30018483 | 0.31660751 | 0.48897239 | PRO3 | V | 201076 | 201936 | 1 |
| YER024W | 276.166078 | -1.2532317 | 0.48018421 | 24.1703354 | 5.64E-06 | 7.89E-05 | YAT2 | V | 202192 | 204963 | 1 |
| YER025W | 81.5488209 | -0.2173183 | 0.56199641 | 0.56390223 | 0.75431056 | 0.85152235 | GCD11 | V | 205251 | 206834 | 1 |
| YER026C | 112.311171 | 0.34134386 | 0.51729216 | 4.58623426 | 0.10095129 | 0.22031963 | CHO1 | V | 207644 | 208474 | -1 |
| YER027C | 20.6536608 | 0.01254872 | 0.87983413 | 0.57716269 | 0.74932585 | 0.8479727 | GAL83 | V | 208979 | 210232 | -1 |
| YER028C | 108.335541 | -0.6938108 | 0.51833879 | 3.70133919 | 0.15713192 | 0.30296164 | MIG3 | V | 210692 | 211876 | -1 |
| YER029C | 41.6116818 | 0.00260327 | 0.66010685 | 3.79218763 | 0.150154 | 0.29297995 | SMB1 | V | 212587 | 213177 | -1 |
| YER030W | 128.507429 | 0.47803997 | 0.62107278 | 7.18437461 | 0.02753803 | 0.08334825 | CHZ1 | V | 213437 | 213898 | 1 |
| YER031C | 32.0906281 | 0.60370269 | 0.80532985 | 9.26302109 | 0.00974004 | 0.03862658 | YPT31 | V | 214076 | 214747 | -1 |
| YER032W | 54.1945932 | -0.188742 | 0.64462986 | 1.37946746 | 0.50170964 | 0.66050451 | FIR1 | V | 215063 | 217693 | 1 |
| YER033C | 127.501385 | 0.14934967 | 0.58102167 | 5.61787164 | 0.0602691 | 0.15054274 | ZRG8 | V | 218057 | 221287 | -1 |
| YER034W | 54.535686 | 2.32401779 | 0.79771062 | 9.38361722 | 0.00917009 | 0.03689701 | NA | V | 221846 | 222403 | 1 |
| YER035W | 55.6139693 | 1.48265185 | 0.64603856 | 6.91553631 | 0.03149999 | 0.09285753 | EDC2 | V | 222639 | 223076 | 1 |
| YER036C | 116.155692 | -0.0985923 | 0.51363291 | 11.8629207 | 0.0026546 | 0.01388984 | ARB1 | V | 223367 | 225199 | -1 |
| YER037W | 704.687182 | -0.7703229 | 0.45902914 | 21.630009 | 2.01E-05 | 0.00023746 | PHM8 | V | 225889 | 226854 | 1 |
| YER038C | 1951.91049 | 2.24414585 | 0.63898902 | 20.0285059 | 4.48E-05 | 0.00046777 | KRE29 | V | 226858 | 228252 | -1 |
| YER038W-A | 563.277799 | 1.88523251 | 0.50836484 | 24.7053771 | 4.32E-06 | 6.30E-05 | FMP49 | V | 228451 | 228831 | 1 |
| YER039C | 72.7048313 | -1.4466429 | 0.56583929 | 22.1641096 | 1.54E-05 | 0.00019051 | HVG1 | V | 228456 | 229205 | -1 |
| YER039C-A | 21.0623843 | -1.3221036 | 0.78927688 | 5.70673855 | 0.05764976 | 0.14582441 | NA | V | 229263 | 229481 | -1 |
| YER040W | 17.8587822 | 0.99707628 | 0.82586921 | 2.78166488 | 0.24886805 | 0.41973376 | GLN3 | V | 229795 | 231987 | 1 |
| YER041W | 13.2349626 | -0.0798888 | 0.90056077 | 0.66189522 | 0.7182428 | 0.82928083 | YEN1 | V | 232461 | 234740 | 1 |
| YER042W | 16.2822964 | 1.39763244 | 1.00612417 | 9.36558364 | 0.00925314 | 0.03719466 | MXR1 | V | 234937 | 235491 | 1 |
| YER043C | 80.0281878 | -0.660119 | 0.73504043 | 2.49111365 | 0.28778062 | 0.46172289 | SAH1 | V | 235770 | 237119 | -1 |
| YER044C | 8.4274271 | -0.1285231 | 1.24257737 | 0.20699369 | 0.90167887 | 0.93824416 | ERG28 | V | 237570 | 238016 | -1 |
| YER044C-A | 152.775915 | 0.46063427 | 0.50836746 | 4.22745953 | 0.12078662 | 0.25039546 | MEI4 | V | 238460 | 239774 | -1 |
| YER045C | 523.137559 | 0.19217041 | 0.50383086 | 0.95799194 | 0.61940498 | 0.76030111 | ACA1 | V | 240032 | 241501 | -1 |

| | | | | | | | | | | | |
|---|---|---|---|---|---|---|---|---|---|---|---|
| YER046W | 40.3832792 | 0.35040263 | 0.79040923 | 0.22601126 | 0.89314563 | 0.93429326 | SPO73 | V | 243180 | 243611 | 1 |
| YER047C | 73.5719866 | -0.5528821 | 0.60146666 | 2.73795128 | 0.25436739 | 0.42652395 | SAP1 | V | 243810 | 246503 | -1 |
| YER048C | 66.8559971 | -0.3762303 | 0.71434904 | 0.29645078 | 0.86223675 | 0.91923233 | CAJ1 | V | 246982 | 248157 | -1 |
| YER048W-A | 7.99170786 | 1.4112795 | 1.06248165 | 1.85695556 | 0.39515476 | 0.5664635 | ISD11 | V | 250718 | 251002 | 1 |
| YER049W | 35.2628584 | -0.8615542 | 0.73924714 | 2.79962569 | 0.24664312 | 0.41766412 | TPA1 | V | 251728 | 253662 | 1 |
| YER050C | 8.76293719 | 0.48015073 | 1.10893526 | 2.79949759 | 0.24665892 | 0.41766412 | RSM18 | V | 253971 | 254387 | -1 |
| YER051W | 61.2513181 | 0.24685488 | 0.64659393 | 12.7988505 | 0.00166251 | 0.00978886 | JHD1 | V | 254656 | 256134 | 1 |
| YER052C | 20.499073 | 0.9820755 | 0.74618201 | 2.27648392 | 0.32038177 | 0.49282956 | HOM3 | V | 256375 | 257958 | -1 |
| YER053C | 95.4406177 | 0.94842954 | 0.57324466 | 2.68618225 | 0.26103752 | 0.43235173 | PIC2 | V | 258737 | 259639 | -1 |
| YER054C | 109.06935 | -1.1404469 | 0.59851387 | 3.833026 | 0.14711907 | 0.28912102 | GIP2 | V | 262052 | 263698 | -1 |
| YER055C | 213.469682 | -0.9285909 | 0.54616705 | 4.06184963 | 0.13121412 | 0.26636668 | HIS1 | V | 264892 | 265785 | -1 |
| YER056C | 24.7771548 | -1.2464779 | 0.74094417 | 4.33990184 | 0.11418322 | 0.24102109 | FCY2 | V | 266512 | 268113 | -1 |
| YER056C-A | 80.3118705 | 0.85408518 | 0.65863257 | 20.5169036 | 3.51E-05 | 0.00038446 | RPL34A | V | 269423 | 270185 | -1 |
| YER057C | 223.640579 | -1.6164555 | 0.52175993 | 29.7611304 | 3.45E-07 | 6.95E-06 | HMF1 | V | 270737 | 271126 | -1 |
| YER058W | 16.830535 | -0.6252524 | 0.78063899 | 0.83628535 | 0.6582683 | 0.78633659 | PET117 | V | 271768 | 272091 | 1 |
| YER059W | 54.9343005 | -1.0567914 | 0.57825041 | 13.4945962 | 0.00117405 | 0.00750302 | PCL6 | V | 272624 | 273886 | 1 |
| YER060W | 891.417469 | -0.2875209 | 0.57934189 | 0.70152236 | 0.7041519 | 0.82015637 | FCY21 | V | 274567 | 276153 | 1 |
| YER060W-A | 831.668374 | 0.21007809 | 0.73643502 | 0.09831076 | 0.95203319 | 0.96672609 | FCY22 | V | 276572 | 278164 | 1 |
| YER061C | 29.8078867 | -1.1063534 | 0.73177683 | 6.12972977 | 0.04666015 | 0.12398451 | CEM1 | V | 278298 | 279626 | -1 |
| YER062C | 167.394336 | 0.35537785 | 0.50417596 | 12.4572526 | 0.00197216 | 0.01104333 | GPP2 | V | 279930 | 280682 | -1 |
| YER063W | 503.126894 | -0.6870559 | 0.52997976 | 19.1267114 | 7.03E-05 | 0.00069655 | THO1 | V | 281710 | 282366 | 1 |
| YER064C | 310.361037 | 3.22386427 | 0.56310599 | 34.0426745 | 4.05E-08 | 1.07E-06 | VHR2 | V | 282705 | 284222 | -1 |
| YER065C | 742.721524 | -2.1672812 | 0.4949509 | 46.3846954 | 8.47E-11 | 4.12E-09 | ICL1 | V | 285241 | 286914 | -1 |
| YER067W | 15.8475702 | -1.279801 | 0.97639324 | 3.16389159 | 0.2055747 | 0.36691662 | RGI1 | V | 292066 | 292551 | 1 |
| YER068W | 41.5307894 | 0.54953216 | 0.61032579 | 1.42234725 | 0.49106753 | 0.65227827 | MOT2 | V | 293050 | 294813 | 1 |
| YER069W | 106.025982 | 0.73571488 | 0.58508611 | 19.8211886 | 4.96E-05 | 0.00051177 | ARG5,6 | V | 295410 | 298001 | 1 |
| YER070W | 27.8771983 | -1.1409089 | 0.81679573 | 9.0516647 | 0.0108257 | 0.04184299 | RNR1 | V | 298950 | 301616 | 1 |
| YER071C | 53.9082163 | -0.3160092 | 0.58568202 | 4.15307598 | 0.12536347 | 0.25809632 | TDA2 | V | 301947 | 302327 | -1 |
| YER072W | 205.925721 | 1.79120888 | 0.48661317 | 12.8979994 | 0.0015821 | 0.00951154 | VTC1 | V | 302806 | 303195 | 1 |
| YER073W | 21.4982268 | -1.1917836 | 0.78822009 | 2.46359359 | 0.29176786 | 0.46615229 | ALD5 | V | 304030 | 305592 | 1 |
| YER074W | 125.098399 | -0.8674827 | 0.8064425 | 13.292862 | 0.00129865 | 0.00812884 | RPS24A | V | 306323 | 307196 | 1 |
| YER074W-A | 41.0455274 | 0.05651384 | 0.60653284 | 2.55028854 | 0.27939066 | 0.45351049 | YOS1 | V | 307653 | 308123 | 1 |
| YER075C | 193.969549 | 0.36833697 | 0.51926238 | 28.1164522 | 7.84E-07 | 1.44E-05 | PTP3 | V | 308413 | 311199 | -1 |

| | | | | | | | | | | | |
|---|---|---|---|---|---|---|---|---|---|---|---|
| YER076C | 25.9793048 | -0.6464067 | 0.7742741 | 3.22773716 | 0.19911583 | 0.35792631 | NA | V | 312590 | 313498 | -1 |
| YER077C | 40.994989 | 0.63320243 | 0.69221023 | 0.91648199 | 0.63239505 | 0.76955658 | MRX1 | V | 314534 | 316600 | -1 |
| YER078C | 124.420809 | 0.57144026 | 0.52720612 | 7.16108057 | 0.02786064 | 0.08406103 | ICP55 | V | 316807 | 318342 | -1 |
| YER079W | 71.9595585 | -0.6330431 | 0.6233496 | 1.7459641 | 0.41770408 | 0.58926319 | NA | V | 318920 | 319552 | 1 |
| YER080W | 32.1726407 | 0.31760196 | 0.64974584 | 1.22375928 | 0.54233052 | 0.69625138 | AIM9 | V | 319963 | 321846 | 1 |
| YER081W | 15.1841454 | 0.76176087 | 0.95978156 | 2.19351257 | 0.33395258 | 0.5079834 | SER3 | V | 322686 | 324095 | 1 |
| YER082C | 183.879808 | 2.34072353 | 0.78688374 | 9.23824059 | 0.00986147 | 0.0390013 | UTP7 | V | 324272 | 325936 | -1 |
| YER083C | 20.8384642 | -0.1129792 | 0.72366053 | 3.87157588 | 0.14431052 | 0.28488165 | GET2 | V | 326174 | 327031 | -1 |
| YER086W | 123.354459 | 0.73473904 | 0.58661365 | 18.2092969 | 0.00011115 | 0.0010215 | ILV1 | V | 328477 | 330207 | 1 |
| YER087C-B | 136.52163 | 0.58369291 | 0.55122989 | 5.03814171 | 0.0805344 | 0.18734197 | SBH1 | V | 332582 | 332830 | -1 |
| YER087W | 158.498974 | 1.45244722 | 0.5861062 | 12.8585911 | 0.00161359 | 0.00961783 | AIM10 | V | 330576 | 332306 | 1 |
| YER088C | 274.826881 | 0.71037763 | 0.51510004 | 8.43901666 | 0.01470587 | 0.05262485 | DOT6 | V | 333176 | 335188 | -1 |
| YER089C | 212.141068 | -0.5279044 | 0.49503284 | 3.93000409 | 0.1401556 | 0.27867477 | PTC2 | V | 335946 | 337340 | -1 |
| YER090W | 43.849504 | 0.33378173 | 0.59528088 | 0.44945763 | 0.79873279 | 0.87821171 | TRP2 | V | 337949 | 339472 | 1 |
| YER091C | 47.9214254 | -0.7432277 | 0.64840061 | 8.83884572 | 0.01204118 | 0.04533058 | MET6 | V | 339864 | 342167 | -1 |
| YER092W | 50.5999553 | 1.42821286 | 0.65354597 | 6.05766722 | 0.04837203 | 0.12713301 | IES5 | V | 342855 | 343232 | 1 |
| YER093C | 667.993694 | 0.4608331 | 0.47664403 | 3.90482227 | 0.14193144 | 0.28114371 | TSC11 | V | 343320 | 347612 | -1 |
| YER093C-A | 8.16772941 | -1.6231839 | 1.18744629 | 3.22437727 | 0.19945061 | 0.35830563 | AIM11 | V | 347912 | 348400 | -1 |
| YER094C | 31.0830692 | -0.1552034 | 0.65638811 | 0.31604408 | 0.85383096 | 0.91329785 | PUP3 | V | 348729 | 349346 | -1 |
| YER095W | 107.430999 | 1.10125333 | 0.57714796 | 14.9966055 | 0.00055402 | 0.00399975 | RAD51 | V | 349980 | 351182 | 1 |
| YER096W | 38.2973582 | 0.13265889 | 0.68105732 | 10.5622257 | 0.00508677 | 0.02326412 | SHC1 | V | 351698 | 353236 | 1 |
| YER098W | 281.276891 | -1.4851315 | 0.49704274 | 22.2014008 | 1.51E-05 | 0.00018804 | UBP9 | V | 355466 | 357730 | 1 |
| YER099C | 45.7971847 | 0.88750305 | 0.63912474 | 2.39625303 | 0.30175902 | 0.47448396 | PRS2 | V | 358105 | 359061 | -1 |
| YER100W | 15.2961059 | 0.69084828 | 0.83305126 | 0.77790226 | 0.67776739 | 0.8011991 | UBC6 | V | 359562 | 360314 | 1 |
| YER101C | 106.538811 | -0.2306519 | 0.57261063 | 9.18787421 | 0.01011296 | 0.03980562 | AST2 | V | 360502 | 361794 | -1 |
| YER102W | 76.8515991 | -0.6066468 | 0.59298869 | 25.2341191 | 3.31E-06 | 5.05E-05 | RPS8A | V | 363100 | 363702 | 1 |
| YER103W | 42.5748513 | 0.10070772 | 0.70272032 | 13.0051812 | 0.00149955 | 0.00912975 | SSA4 | V | 364589 | 366517 | 1 |
| YER104W | 205.489024 | -0.6519929 | 0.52428593 | 1.56093267 | 0.45819229 | 0.62328904 | RTT105 | V | 366802 | 367428 | 1 |
| YER105C | 204.95755 | -0.84155 | 0.516175 | 3.48502709 | 0.17507978 | 0.32688612 | NUP157 | V | 367838 | 372013 | -1 |
| YER107C | 17.0125847 | -0.2094056 | 0.79015352 | 0.50933162 | 0.77517551 | 0.86346022 | GLE2 | V | 373448 | 374545 | -1 |
| YER109C | 136.461112 | -0.6467822 | 0.58757768 | 10.7103193 | 0.00472372 | 0.02186276 | NA | NA | NA | NA | NA |
| YER110C | 67.4590919 | -0.7441571 | 0.5798044 | 3.1685125 | 0.20510028 | 0.36629569 | KAP123 | V | 378762 | 382103 | -1 |
| YER111C | 34.2479375 | 0.00615872 | 0.69929625 | 0.65914192 | 0.71923225 | 0.82938751 | SWI4 | V | 382595 | 385876 | -1 |

| | | | | | | | | | | | |
|---|---|---|---|---|---|---|---|---|---|---|---|
| YER112W | 81.5774478 | 1.34847372 | 0.91233609 | 8.1739764 | 0.01678972 | 0.05813116 | LSM4 | V | 387232 | 387795 | 1 |
| YER113C | 17.7845988 | -1.1180308 | 0.7802128 | 2.08539856 | 0.3525019 | 0.52616292 | TMN3 | V | 387932 | 390052 | -1 |
| YER114C | 47.2796174 | 0.47783268 | 0.75599796 | 6.68101773 | 0.03541893 | 0.10142216 | BOI2 | V | 390590 | 393712 | -1 |
| YER115C | 6.91550538 | -1.3793849 | 1.30900865 | 4.55305406 | 0.10264005 | 0.22312171 | SPR6 | V | 394292 | 394867 | -1 |
| YER116C | 35.8187016 | -1.2268758 | 0.71259402 | 7.40790697 | 0.02462598 | 0.07694787 | SLX8 | V | 395348 | 396172 | -1 |
| YER117W | 32.6679794 | -0.1050318 | 0.90761366 | 6.87410673 | 0.03215931 | 0.0942798 | RPL23A | V | 396769 | 397653 | 1 |
| YER118C | 23.2533793 | 0.55747681 | 0.86620293 | 3.67044552 | 0.15957795 | 0.3066566 | SHO1 | V | 397952 | 399055 | -1 |
| YER119C | 229.62588 | -0.8801272 | 0.52767703 | 2.75511064 | 0.25219433 | 0.42365049 | AVT6 | V | 399496 | 400842 | -1 |
| YER120W | 111.245703 | -0.3576227 | 0.51731467 | 2.26204284 | 0.32270347 | 0.4957424 | SCS2 | V | 401135 | 401869 | 1 |
| YER121W | 75.5519252 | 0.57142511 | 0.67049722 | 1.91331973 | 0.38417394 | 0.5573458 | NA | V | 402375 | 402719 | 1 |
| YER122C | 56.7675776 | 0.00340557 | 0.60126586 | 2.44684384 | 0.29422164 | 0.46865015 | GLO3 | V | 402871 | 404352 | -1 |
| YER123W | 11.0229101 | -0.534122 | 1.02158203 | 2.60966363 | 0.27121815 | 0.44467869 | YCK3 | V | 404813 | 406387 | 1 |
| YER124C | 180.569537 | -0.1287398 | 0.50306534 | 2.78209129 | 0.248815 | 0.41973376 | DSE1 | V | 407342 | 409063 | -1 |
| YER125W | 61.7124911 | 0.5114372 | 0.66132159 | 0.68278947 | 0.71077828 | 0.82438173 | RSP5 | V | 410189 | 412618 | 1 |
| YER126C | 45.4125309 | -0.7278403 | 0.65098758 | 1.32015586 | 0.51681106 | 0.67350835 | NSA2 | V | 413394 | 414179 | -1 |
| YER127W | 34.6723543 | -0.1045169 | 0.76499662 | 3.71782824 | 0.15584176 | 0.30107568 | LCP5 | V | 414481 | 415554 | 1 |
| YER128W | 17.8297179 | 0.7829734 | 0.85948837 | 0.84141393 | 0.65658247 | 0.78549013 | VFA1 | V | 415859 | 416470 | 1 |
| YER129W | 60.5964095 | 0.2910186 | 0.63313029 | 5.11238177 | 0.07759976 | 0.18227288 | SAK1 | V | 417281 | 420709 | 1 |
| YER130C | 27.5464413 | -0.7447183 | 0.84030463 | 1.34201726 | 0.51119271 | 0.66829159 | COM2 | V | 421115 | 422446 | -1 |
| YER131W | 164.32768 | -0.7602049 | 0.59543723 | 8.96290404 | 0.01131697 | 0.04316551 | RPS26B | V | 423952 | 424311 | 1 |
| YER132C | 113.425834 | 0.88172928 | 0.54686997 | 14.9883319 | 0.00055632 | 0.00401133 | PMD1 | V | 425188 | 430449 | -1 |
| YER133W | 87.5429717 | 0.492731 | 0.73398491 | 1.2013536 | 0.54844033 | 0.70063006 | GLC7 | V | 432495 | 433958 | 1 |
| YER134C | 27.567237 | 0.67159777 | 1.0240276 | 0.58460071 | 0.74654428 | 0.84572322 | NA | V | 437267 | 437803 | -1 |
| YER135C | 20.710723 | 0.46670439 | 0.80167285 | 6.66766967 | 0.03565611 | 0.10195005 | NA | V | 437952 | 438344 | -1 |
| YER136W | 25.5523655 | 0.71746654 | 0.81537126 | 0.88990421 | 0.64085497 | 0.77561669 | GDI1 | V | 439616 | 440971 | 1 |
| YER139C | 14.0625398 | -1.7367766 | 0.86130015 | 12.1770853 | 0.00226871 | 0.01229812 | RTR1 | V | 450563 | 451243 | -1 |
| YER140W | 19.9342804 | -0.3631177 | 0.73196895 | 1.00005253 | 0.60651473 | 0.75088539 | EMP65 | V | 451565 | 453235 | 1 |
| YER141W | 115.628197 | 0.18857219 | 0.52228305 | 0.35851966 | 0.83588868 | 0.9031862 | COX15 | V | 453459 | 454919 | 1 |
| YER142C | 365.812013 | 3.34135664 | 1.83512503 | 3.48636239 | 0.17496292 | 0.32678559 | MAG1 | V | 455146 | 456036 | -1 |
| YER143W | 237.726923 | 0.48132687 | 0.48149045 | 5.64877771 | 0.05934491 | 0.14861897 | DDI1 | V | 456319 | 457605 | 1 |
| YER144C | 74.2152784 | -0.7517417 | 0.5610175 | 2.00930101 | 0.36617259 | 0.53947564 | UBP5 | V | 457806 | 460223 | -1 |
| YER145C | 356.73164 | 0.99027701 | 0.60541909 | 3.31110018 | 0.19098697 | 0.34741267 | FTR1 | V | 460526 | 461740 | -1 |
| YER145C-A | 2194.74033 | -1.2192505 | 0.45563633 | 33.3456672 | 5.74E-08 | 1.48E-06 | NA | V | 462386 | 462823 | -1 |

| | | | | | | | | | | | |
|---|---|---|---|---|---|---|---|---|---|---|---|
| YER146W | 78.2436643 | 2.02329761 | 0.66949227 | 21.6783321 | 1.96E-05 | 0.00023274 | LSM5 | V | 462585 | 462866 | 1 |
| YER147C | 468.696204 | -0.2497646 | 0.50987806 | 7.6797415 | 0.02149638 | 0.06990254 | SCC4 | V | 462968 | 464842 | -1 |
| YER148W | 38.5568111 | -0.2204409 | 0.86720411 | 3.64959557 | 0.16125025 | 0.30925437 | SPT15 | V | 465303 | 466025 | 1 |
| YER149C | 169.133868 | 1.87858682 | 0.85203887 | 8.88051575 | 0.0117929 | 0.04456976 | PEA2 | V | 466208 | 467470 | -1 |
| YER150W | 132.564993 | 1.65550975 | 0.52318469 | 19.3215691 | 6.37E-05 | 0.00063406 | SPI1 | V | 468370 | 468816 | 1 |
| YER151C | 36.4183053 | 0.34232269 | 0.68682508 | 1.3093959 | 0.51959898 | 0.67578891 | UBP3 | V | 469686 | 472424 | -1 |
| YER152C | 47.1208303 | 1.7114096 | 0.67973568 | 20.0731907 | 4.38E-05 | 0.00045993 | NA | V | 472657 | 473988 | -1 |
| YER153C | 140.247832 | -0.0516868 | 0.6421335 | 7.22579712 | 0.02697355 | 0.08206876 | PET122 | V | 474041 | 474805 | -1 |
| YER154W | 451.471563 | 1.04262095 | 0.83355222 | 2.46084146 | 0.29216963 | 0.46640809 | OXA1 | V | 475020 | 476228 | 1 |
| YER155C | 91.0380692 | 0.50657364 | 0.64352756 | 3.00724011 | 0.22232388 | 0.38737744 | BEM2 | V | 476345 | 482848 | -1 |
| YER156C | 50.661402 | -1.5303807 | 0.63657284 | 16.2005893 | 0.00030345 | 0.00247461 | NA | V | 483325 | 484341 | -1 |
| YER157W | 7.65318783 | -0.5818501 | 1.11728851 | 0.33672242 | 0.84504854 | 0.90911019 | COG3 | V | 484788 | 487193 | 1 |
| YER158C | 611.580793 | -0.3463648 | 0.52696354 | 2.89843111 | 0.23475437 | 0.40225741 | NA | V | 488857 | 490578 | -1 |
| YER159C | 292.110878 | 1.99659855 | 0.58670509 | 18.6210311 | 9.05E-05 | 0.0008601 | BUR6 | V | 491530 | 491958 | -1 |
| YER161C | 41.5724261 | 0.67590291 | 0.63129322 | 1.20370361 | 0.54779628 | 0.70011119 | SPT2 | V | 499347 | 500348 | -1 |
| YER162C | 148.051859 | 0.04439649 | 0.53597474 | 30.097458 | 2.91E-07 | 6.11E-06 | RAD4 | V | 500630 | 502894 | -1 |
| YER163C | 36.44188 | -0.2307021 | 0.63720249 | 0.16822671 | 0.91932704 | 0.94974417 | GCG1 | V | 503084 | 503782 | -1 |
| YER164W | 57.415717 | -0.1292433 | 0.60933095 | 3.99531774 | 0.13565249 | 0.27253845 | CHD1 | V | 505392 | 509798 | 1 |
| YER165W | 149.69727 | 0.02444087 | 0.53269213 | 3.99524673 | 0.13565731 | 0.27253845 | PAB1 | V | 510373 | 512106 | 1 |
| YER166W | 178.286397 | 1.083911 | 0.54290047 | 3.94271701 | 0.13926753 | 0.2778632 | DNF1 | V | 512744 | 517459 | 1 |
| YER167W | 43.6177841 | 0.47201803 | 0.65532656 | 4.00449649 | 0.13503136 | 0.27175237 | BCK2 | V | 518216 | 520771 | 1 |
| YER168C | 14.1326525 | -0.2711217 | 0.8338498 | 0.11554003 | 0.943867 | 0.96197675 | CCA1 | V | 521029 | 522669 | -1 |
| YER169W | 33.2625146 | 0.20668975 | 0.65423644 | 0.16447046 | 0.92105527 | 0.95035295 | RPH1 | V | 523369 | 525759 | 1 |
| YER170W | 88.0582312 | -0.1275732 | 0.61096319 | 0.05609901 | 0.97234023 | 0.97893408 | ADK2 | V | 525974 | 526651 | 1 |
| YER171W | 164.200431 | 1.24043316 | 0.65158169 | 5.73204882 | 0.05692479 | 0.14462243 | RAD3 | V | 527082 | 529418 | 1 |
| YER172C | 515.31783 | 0.81235562 | 0.53722986 | 26.9157756 | 1.43E-06 | 2.43E-05 | BRR2 | V | 529530 | 536021 | -1 |
| YER173W | 13.9863034 | -0.8531487 | 0.83528539 | 3.25669735 | 0.19625339 | 0.35398975 | RAD24 | V | 536300 | 538279 | 1 |
| YER174C | 24.727032 | 2.40533221 | 1.05103689 | 6.60689919 | 0.03675615 | 0.10442499 | GRX4 | V | 538434 | 539168 | -1 |
| YER175C | 96.5024105 | -0.2231138 | 0.55248365 | 0.23381286 | 0.88966843 | 0.93317728 | TMT1 | V | 539464 | 540363 | -1 |
| YER175W-A | 257.230393 | -0.4632934 | 0.5860126 | 3.49977379 | 0.1737936 | 0.32507061 | NA | V | 540650 | 540814 | 1 |
| YER176W | 1271.50844 | 0.77770458 | 0.56209491 | 2.6927455 | 0.2601823 | 0.43177286 | ECM32 | V | 541690 | 545055 | 1 |
| YER177W | 284.608542 | 0.73647948 | 0.50115892 | 2.65562873 | 0.26505594 | 0.43735364 | BMH1 | V | 545611 | 546414 | 1 |
| YER178W | 39.7833935 | -0.6260943 | 0.8094442 | 0.90445319 | 0.63620999 | 0.77274089 | PDA1 | V | 546817 | 548079 | 1 |

| | | | | | | | | | | | |
|---|---|---|---|---|---|---|---|---|---|---|---|
| YER179W | 618.2578 | 1.05122779 | 0.5720739 | 6.87162126 | 0.0321993 | 0.09434916 | DMC1 | V | 548421 | 549517 | 1 |
| YER182W | 152.228797 | 3.20328726 | 1.92610959 | 2.75602617 | 0.25207892 | 0.42365049 | FMP10 | V | 552525 | 553259 | 1 |
| YER183C | 164.508521 | 2.92276272 | 0.9523469 | 8.33841023 | 0.01546455 | 0.05483143 | FAU1 | V | 553334 | 553969 | -1 |
| YER184C | 200.705848 | -0.1806311 | 0.53977699 | 0.42218647 | 0.80969857 | 0.8858947 | TOG1 | V | 556296 | 558680 | -1 |
| YER185W | 32.3488614 | 1.15614144 | 0.83904811 | 4.42845034 | 0.10923812 | 0.23339068 | PUG1 | V | 559454 | 560365 | 1 |
| YER186C | 9.61271573 | 0.44803004 | 1.00482001 | 2.46837647 | 0.29107095 | 0.46518644 | NA | V | 561705 | 562625 | -1 |
| YER187W | 92.0359168 | 1.03906116 | 0.68697858 | 35.7480425 | 1.73E-08 | 4.86E-07 | NA | V | 566230 | 566655 | 1 |
| YER188C-A | 70.8344263 | 4.02357425 | 1.11755916 | 21.9724132 | 1.69E-05 | 0.00020426 | NA | V | 569608 | 569907 | -1 |
| YER188W | 19.4880764 | -0.7048833 | 0.85682134 | 7.84583484 | 0.01978329 | 0.06537972 | NA | V | 568040 | 568759 | 1 |
| YER189W | 60.8917943 | -0.9612253 | 0.59192078 | 2.95583818 | 0.22811187 | 0.39414138 | NA | V | 571155 | 571523 | 1 |
| YER190W | 18.2747319 | -1.9565851 | 0.85727668 | 10.5119789 | 0.00521618 | 0.02376215 | YRF1-2 | V | 571480 | 576525 | 1 |
| YFL001W | 43.4772716 | 1.19181038 | 0.75992186 | 9.04810933 | 0.01084496 | 0.04188948 | DEG1 | VI | 147131 | 148459 | 1 |
| YFL002C | 33.4237746 | 0.195586 | 0.64915655 | 0.12949561 | 0.93730382 | 0.95917093 | SPB4 | VI | 145114 | 146934 | -1 |
| YFL003C | 93.51908 | -0.8071895 | 0.54290043 | 2.19722229 | 0.33333372 | 0.50736125 | MSH4 | VI | 134521 | 137157 | -1 |
| YFL004W | 43.2236981 | -0.0163214 | 0.61120721 | 2.6080258 | 0.27144034 | 0.44484562 | VTC2 | VI | 131810 | 134296 | 1 |
| YFL005W | 39.0120559 | 0.02446283 | 0.65712168 | 3.29223165 | 0.19279731 | 0.34947875 | SEC4 | VI | 130334 | 130981 | 1 |
| YFL007W | 200.668909 | 0.05828638 | 0.48324687 | 0.05284932 | 0.97392142 | 0.97984448 | BLM10 | VI | 123479 | 129910 | 1 |
| YFL008W | 51.0309369 | -0.1961687 | 0.64856362 | 4.53816053 | 0.10340724 | 0.22441077 | SMC1 | VI | 119429 | 123106 | 1 |
| YFL009W | 24.6214996 | 0.59929902 | 0.70455862 | 0.79267066 | 0.67278105 | 0.79774275 | CDC4 | VI | 116145 | 118484 | 1 |
| YFL010C | 163.385203 | 3.02913647 | 0.56874663 | 36.5929965 | 1.13E-08 | 3.34E-07 | WWM1 | VI | 115108 | 115743 | -1 |
| YFL010W-A | 6362.8586 | 0.67092227 | 0.46607826 | 2.08857228 | 0.35194297 | 0.52559964 | AUA1 | VI | 114990 | 115274 | 1 |
| YFL011W | 179.281868 | 3.53621707 | 0.66666104 | 31.1732447 | 1.70E-07 | 3.85E-06 | HXT10 | VI | 112345 | 113985 | 1 |
| YFL012W | 33.5108491 | -1.226 | 0.7052352 | 3.55543582 | 0.16902343 | 0.31888097 | NA | VI | 110647 | 111093 | 1 |
| YFL013C | 62.8349025 | 0.58558356 | 0.560508 | 1.34324403 | 0.51087925 | 0.66829159 | IES1 | VI | 107852 | 109930 | -1 |
| YFL014W | 83.9443555 | 1.73686106 | 0.59595184 | 22.1624431 | 1.54E-05 | 0.00019051 | HSP12 | VI | 107256 | 107585 | 1 |
| YFL015C | 260.605321 | 3.11539748 | 0.55465168 | 33.3831811 | 5.64E-08 | 1.46E-06 | NA | VI | 106469 | 106963 | -1 |
| YFL016C | 119.704688 | 0.9132156 | 0.5863676 | 6.04258366 | 0.04873822 | 0.12803733 | MDJ1 | VI | 104701 | 106236 | -1 |
| YFL017C | 180.579195 | 0.88329885 | 0.59475485 | 2.19044313 | 0.33446549 | 0.50802806 | GNA1 | VI | 103983 | 104462 | -1 |
| YFL017W-A | 207.19516 | 0.74771932 | 0.75788265 | 5.41375101 | 0.06674503 | 0.16211061 | SMX2 | VI | 103699 | 103932 | 1 |
| YFL018C | 170.835586 | 0.6458217 | 0.59366593 | 37.1241141 | 8.68E-09 | 2.65E-07 | LPD1 | VI | 101628 | 103127 | -1 |
| YFL019C | 731.05901 | 2.99662836 | 0.61009875 | 51.2746877 | 7.34E-12 | 4.52E-10 | NA | VI | 100252 | 100605 | -1 |
| YFL020C | 8.42144089 | 0.47045288 | 1.41462722 | 10.9818647 | 0.004124 | 0.01964738 | PAU5 | VI | 99231 | 99599 | -1 |
| YFL021W | 356.018105 | 0.10869964 | 0.60293925 | 21.3770886 | 2.28E-05 | 0.00026461 | GAT1 | VI | 95966 | 97498 | 1 |

| | | | | | | | | | | | |
|---|---|---|---|---|---|---|---|---|---|---|---|
| YFL022C | 69.1371007 | -0.1151518 | 0.61752439 | 0.64382985 | 0.72475985 | 0.8342663 | FRS2 | VI | 93499 | 95010 | -1 |
| YFL023W | 46.6981626 | -0.813234 | 0.82345381 | 3.42299985 | 0.18059471 | 0.33514211 | BUD27 | VI | 90986 | 93376 | 1 |
| YFL024C | 548.024339 | -1.1943999 | 0.47144434 | 20.2375107 | 4.03E-05 | 0.00043148 | EPL1 | VI | 87847 | 90345 | -1 |
| YFL025C | 23.9941525 | -0.8799397 | 0.78008095 | 4.28450187 | 0.11739031 | 0.24546419 | BST1 | VI | 84145 | 87234 | -1 |
| YFL026W | 139.186616 | 0.35637218 | 0.60369587 | 52.9500793 | 3.18E-12 | 2.07E-10 | STE2 | VI | 82580 | 83875 | 1 |
| YFL027C | 23.6084875 | -0.3120621 | 0.77803292 | 8.09744683 | 0.01744463 | 0.05976592 | GYP8 | VI | 80419 | 81912 | -1 |
| YFL028C | 12.4673956 | 1.44185581 | 0.89680217 | 3.34603657 | 0.18767974 | 0.34351531 | CAF16 | VI | 79344 | 80213 | -1 |
| YFL029C | 556.108056 | 1.35485095 | 0.57360382 | 11.2344192 | 0.00363477 | 0.01781991 | CAK1 | VI | 78055 | 79161 | -1 |
| YFL030W | 211.168915 | -0.1544502 | 0.53525847 | 4.22284135 | 0.12106585 | 0.25079473 | AGX1 | VI | 76831 | 77988 | 1 |
| YFL031W | 365.973031 | 1.66089002 | 0.59688814 | 7.99512566 | 0.01836033 | 0.06205856 | HAC1 | VI | 75179 | 76147 | 1 |
| YFL033C | 2432.97573 | 0.03028086 | 0.47880511 | 4.2401276 | 0.12002397 | 0.24929085 | RIM15 | VI | 69115 | 74427 | -1 |
| YFL034C-A | 56.2903263 | 1.28404588 | 0.64741698 | 13.3965604 | 0.00123303 | 0.00781955 | RPL22B | VI | 64243 | 64932 | -1 |
| YFL034C-B | 115.384806 | 1.71119474 | 0.68037716 | 14.5986965 | 0.00067598 | 0.00475567 | MOB2 | VI | 63016 | 63993 | -1 |
| YFL034W | 37.1725303 | 0.37530277 | 0.61640285 | 0.37283012 | 0.82992905 | 0.89868884 | MIL1 | VI | 65477 | 68698 | 1 |
| YFL036W | 15.4894288 | 0.69381282 | 0.79168022 | 2.48436374 | 0.28875351 | 0.46261284 | RPO41 | VI | 58782 | 62837 | 1 |
| YFL037W | 38.8773377 | -0.0148383 | 0.80080053 | 1.26392361 | 0.53154798 | 0.68590658 | TUB2 | VI | 56336 | 57709 | 1 |
| YFL038C | 102.743957 | 0.5920964 | 0.53448586 | 2.09160444 | 0.3514098 | 0.52507425 | YPT1 | VI | 55366 | 55986 | -1 |
| YFL039C | 332.45909 | 0.5265715 | 0.6938009 | 0.87367394 | 0.64607675 | 0.77803336 | ACT1 | VI | 53260 | 54696 | -1 |
| YFL040W | 1229.23072 | 2.06511155 | 0.50624139 | 26.1408681 | 2.11E-06 | 3.36E-05 | NA | VI | 51350 | 52972 | 1 |
| YFL041W | 12.6362744 | -0.6019589 | 0.96088205 | 0.58266132 | 0.74726855 | 0.84631426 | FET5 | VI | 49139 | 51007 | 1 |
| YFL042C | 25.6410458 | 0.97182076 | 0.78530193 | 4.8436064 | 0.08876142 | 0.20185727 | LAM5 | VI | 45720 | 47744 | -1 |
| YFL044C | 12.4536544 | -0.5493516 | 1.03024864 | 0.33096102 | 0.84748638 | 0.90980423 | OTU1 | VI | 44655 | 45560 | -1 |
| YFL045C | 51.8078165 | 0.00620163 | 0.74738767 | 2.55366053 | 0.27892 | 0.45338204 | SEC53 | VI | 43628 | 44392 | -1 |
| YFL046W | 26.2801263 | 0.42603441 | 0.6809379 | 4.11069311 | 0.12804845 | 0.26151694 | FMP32 | VI | 42815 | 43438 | 1 |
| YFL047W | 17.3574823 | 0.69961669 | 0.93305724 | 0.87858039 | 0.64449372 | 0.77725863 | RGD2 | VI | 40421 | 42565 | 1 |
| YFL048C | 29.8704495 | 0.01812334 | 0.65710922 | 0.43313363 | 0.80527873 | 0.88306133 | EMP47 | VI | 38843 | 40180 | -1 |
| YFL049W | 12.3242104 | 0.934164 | 0.94358558 | 1.60403604 | 0.44842312 | 0.61504128 | SWP82 | VI | 36803 | 38674 | 1 |
| YFL050C | 60.650699 | 1.06687948 | 0.65573676 | 35.0146898 | 2.49E-08 | 6.81E-07 | ALR2 | VI | 33272 | 35848 | -1 |
| YFL051C | 95.0709695 | -0.2937298 | 0.59189675 | 4.04865818 | 0.13208243 | 0.26749117 | NA | VI | 30058 | 30540 | -1 |
| YFL052W | 193.903239 | -1.2324212 | 0.48974484 | 20.9370553 | 2.84E-05 | 0.00032262 | ZNF1 | VI | 28232 | 29629 | 1 |
| YFL053W | 485.363874 | -0.5467753 | 0.52162874 | 1.25248107 | 0.53459783 | 0.68922766 | DAK2 | VI | 23423 | 25198 | 1 |
| YFL054C | 30.8251295 | -0.562421 | 0.81243275 | 19.0165331 | 7.42E-05 | 0.00072613 | AQY3 | VI | 20847 | 22787 | -1 |
| YFL055W | 148.302991 | 0.69899298 | 0.53203534 | 16.5789445 | 0.00025115 | 0.0020833 | AGP3 | VI | 17004 | 18680 | 1 |

| | | | | | | | | | | | |
|---|---|---|---|---|---|---|---|---|---|---|---|
| YFL056C | 11.7863485 | 0.3517259 | 1.04269612 | 2.09951305 | 0.35002296 | 0.52354248 | NA | VI | 14305 | 15431 | -1 |
| YFL059W | 35.7601537 | 1.0944733 | 0.69702952 | 7.63632836 | 0.02196809 | 0.07113829 | SNZ3 | VI | 11363 | 12259 | 1 |
| YFL060C | 7.65161202 | 0.38847291 | 1.05991173 | 0.62635656 | 0.73111956 | 0.83700349 | SNO3 | VI | 10301 | 10969 | -1 |
| YFL061W | 146.187721 | -1.0581517 | 0.58379259 | 3.54061024 | 0.17028102 | 0.32052333 | DDI3 | VI | 9545 | 10222 | 1 |
| YFL062W | 262.929821 | 4.69299085 | 0.61609316 | 53.4453943 | 2.48E-12 | 1.63E-10 | COS4 | VI | 6426 | 7565 | 1 |
| YFL064C | 16.5014881 | -2.3156543 | 0.92317326 | 7.78556552 | 0.02038853 | 0.06703554 | NA | VI | 3322 | 3846 | -1 |
| YFL066C | 5.50556082 | -0.8575012 | 1.30155207 | 0.88028052 | 0.64394609 | 0.77686719 | NA | VI | 1437 | 2615 | -1 |
| YFL067W | 1321.13539 | -0.2109055 | 0.56132016 | 0.14095644 | 0.93194804 | 0.95724306 | NA | VI | 836 | 1363 | 1 |
| YFR001W | 43.6172453 | -0.2852849 | 0.75931854 | 7.40190828 | 0.02469995 | 0.07709579 | LOC1 | VI | 149110 | 149724 | 1 |
| YFR002W | 280.452052 | 1.43284467 | 0.68942358 | 4.3607748 | 0.11299775 | 0.23947912 | NIC96 | VI | 150016 | 152535 | 1 |
| YFR003C | 33.523676 | 0.04054751 | 0.70095374 | 3.60812532 | 0.1646287 | 0.31355268 | YPI1 | VI | 152657 | 153124 | -1 |
| YFR004W | 53.4492989 | 0.69274751 | 0.64063177 | 2.3782414 | 0.30448888 | 0.47725789 | RPN11 | VI | 153393 | 154313 | 1 |
| YFR005C | 32.3799709 | -0.0971634 | 0.68561176 | 6.63792254 | 0.0361904 | 0.10317205 | SAD1 | VI | 154527 | 155873 | -1 |
| YFR006W | 36.3753694 | 0.32671936 | 0.66442557 | 0.6750165 | 0.71354609 | 0.82562587 | NA | VI | 156145 | 157752 | 1 |
| YFR007W | 148.949977 | -0.2993426 | 0.58990653 | 0.61107343 | 0.73672786 | 0.83969573 | YFH7 | VI | 159299 | 160360 | 1 |
| YFR009W | 17.1324887 | 0.32448755 | 0.92026727 | 2.67855719 | 0.26203463 | 0.43337041 | GCN20 | VI | 162488 | 164746 | 1 |
| YFR010W | 42.4636765 | 0.15194894 | 0.59747361 | 2.1792435 | 0.33634369 | 0.50953165 | UBP6 | VI | 165067 | 166566 | 1 |
| YFR011C | 11.5798321 | 1.00333135 | 0.92454411 | 2.28729336 | 0.31865487 | 0.49147887 | MIC19 | VI | 166746 | 167258 | -1 |
| YFR012W | 68.3639453 | 0.24383272 | 0.73721945 | 1.23293608 | 0.5398478 | 0.69395761 | DCV1 | VI | 167888 | 168496 | 1 |
| YFR013W | 22.6032938 | -0.6863392 | 0.72233602 | 2.1409614 | 0.34284367 | 0.51654043 | IOC3 | VI | 169922 | 172285 | 1 |
| YFR014C | 74.9792887 | -0.5854376 | 0.53365742 | 6.36455268 | 0.0414911 | 0.11428804 | CMK1 | VI | 172537 | 173877 | -1 |
| YFR015C | 290.139673 | 0.78191276 | 0.51678711 | 9.48074827 | 0.00873538 | 0.03551814 | GSY1 | VI | 174265 | 176391 | -1 |
| YFR016C | 88.920922 | 1.01798966 | 0.53865487 | 4.56014014 | 0.10227704 | 0.22277848 | NA | VI | 177042 | 180743 | -1 |
| YFR017C | 16.5912517 | 1.40466736 | 0.92758206 | 11.3944122 | 0.00335533 | 0.016633 | IGD1 | VI | 182274 | 182861 | -1 |
| YFR018C | 11.315605 | 0.09422818 | 0.94565698 | 1.12337264 | 0.57024663 | 0.72074394 | NA | VI | 183131 | 184222 | -1 |
| YFR019W | 43.8226135 | -0.0475053 | 0.61513038 | 0.93631049 | 0.62615631 | 0.76437804 | FAB1 | VI | 184502 | 191338 | 1 |
| YFR020W | 10.685797 | 0.42449196 | 1.02370433 | 5.87666346 | 0.052954 | 0.13651097 | CSS2 | VI | 192737 | 193435 | 1 |
| YFR021W | 102.74926 | -0.3518914 | 0.58576987 | 1.77326219 | 0.41204155 | 0.58387679 | ATG18 | VI | 194812 | 196314 | 1 |
| YFR022W | 81.4242109 | 1.31230598 | 0.57983735 | 10.2849226 | 0.00584329 | 0.02594528 | ROG3 | VI | 196833 | 199034 | 1 |
| YFR023W | 16.1848523 | 0.71992893 | 0.98098802 | 3.28170092 | 0.19381514 | 0.35057471 | PES4 | VI | 199874 | 201709 | 1 |
| YFR024C-A | 144.391475 | 0.84027313 | 0.60501289 | 41.0887951 | 1.20E-09 | 4.35E-08 | LSB3 | VI | 201960 | 203433 | -1 |
| YFR025C | 13.8681871 | 0.61155207 | 0.84711339 | 0.5493362 | 0.75982427 | 0.85451323 | HIS2 | VI | 203743 | 204750 | -1 |
| YFR026C | 158.938611 | 1.65872219 | 0.57120246 | 31.5971846 | 1.38E-07 | 3.19E-06 | ULI1 | VI | 205748 | 206257 | -1 |

| | | | | | | | | | | | |
|---|---|---|---|---|---|---|---|---|---|---|---|
| YFR027W | 30.6228025 | -0.1921309 | 0.68830219 | 0.320159 | 0.85207605 | 0.91195795 | ECO1 | VI | 207452 | 208297 | 1 |
| YFR028C | 38.5112836 | 0.58225518 | 0.69685193 | 0.93259971 | 0.62731915 | 0.76483005 | CDC14 | VI | 208413 | 210068 | -1 |
| YFR029W | 43.6420082 | 0.53351816 | 0.6638376 | 4.51904635 | 0.10440025 | 0.22597288 | PTR3 | VI | 210937 | 212973 | 1 |
| YFR030W | 101.603476 | -0.483335 | 0.7320371 | 20.4602889 | 3.61E-05 | 0.00039327 | MET10 | VI | 213312 | 216419 | 1 |
| YFR031C | 29.9127664 | 0.46819423 | 0.65152364 | 0.62775131 | 0.73060987 | 0.83678999 | SMC2 | VI | 216594 | 220106 | -1 |
| YFR031C-A | 204.524806 | -0.2857329 | 0.5646048 | 5.91696414 | 0.05189763 | 0.13440139 | RPL2A | VI | 220507 | 221418 | -1 |
| YFR032C-A | 367.372364 | -0.9930599 | 0.76411819 | 17.6463174 | 0.00014728 | 0.00131397 | RPL29 | VI | 223258 | 223437 | -1 |
| YFR032C-B | 18.5022029 | -1.062869 | 0.74881936 | 11.437847 | 0.00328324 | 0.01637994 | NA | VI | 223710 | 223973 | -1 |
| YFR033C | 743.646847 | 1.61407004 | 0.67853797 | 6.90276505 | 0.03170178 | 0.09322158 | QCR6 | VI | 224326 | 224769 | -1 |
| YFR034C | 55.6327479 | -1.4491696 | 0.58309305 | 10.9266742 | 0.00423938 | 0.02007035 | PHO4 | VI | 225020 | 225958 | -1 |
| YFR035C | 683.141128 | -0.8658733 | 0.47869239 | 20.7998264 | 3.04E-05 | 0.00034284 | NA | VI | 226121 | 226465 | -1 |
| YFR036W | 90.5633481 | 1.47299738 | 0.58245236 | 7.35274378 | 0.02531465 | 0.0784226 | CDC26 | VI | 226963 | 227337 | 1 |
| YFR037C | 106.730413 | -0.5093431 | 0.52172087 | 8.45930978 | 0.01455741 | 0.05238497 | RSC8 | VI | 227513 | 229186 | -1 |
| YFR038W | 29.5797463 | 0.07270446 | 0.67220577 | 0.48328369 | 0.7853374 | 0.87009253 | IRC5 | VI | 229380 | 231941 | 1 |
| YFR039C | 102.783252 | 1.33713869 | 0.72139116 | 5.69457641 | 0.0580014 | 0.14626659 | OSW7 | VI | 232012 | 233544 | -1 |
| YFR040W | 88.3240989 | 2.24254532 | 0.7636543 | 9.9615406 | 0.00686877 | 0.0295909 | SAP155 | VI | 234242 | 237250 | 1 |
| YFR041C | 55.9381398 | 0.89701444 | 0.61475208 | 4.72451935 | 0.09420711 | 0.20963072 | ERJ5 | VI | 237368 | 238255 | -1 |
| YFR042W | 26.1623037 | 2.55976475 | 0.98739665 | 7.13533258 | 0.02822164 | 0.08488482 | KEG1 | VI | 238458 | 239060 | 1 |
| YFR043C | 251.376006 | 1.02943786 | 0.60538402 | 8.14749518 | 0.01701351 | 0.05874074 | IRC6 | VI | 239114 | 239827 | -1 |
| YFR044C | 27.5080709 | 0.53230266 | 0.73815309 | 0.65389281 | 0.72112239 | 0.83085787 | DUG1 | VI | 239992 | 241437 | -1 |
| YFR045W | 145.598109 | 0.35127895 | 0.56633373 | 0.47830699 | 0.78729403 | 0.87059443 | NA | VI | 241998 | 242999 | 1 |
| YFR046C | 61.5709143 | 0.96990978 | 0.75165489 | 3.0382777 | 0.21890031 | 0.38305402 | CNN1 | VI | 243074 | 244159 | -1 |
| YFR047C | 16.0645305 | -0.0420678 | 0.89218291 | 0.2484314 | 0.88318932 | 0.93100373 | BNA6 | VI | 244279 | 245166 | -1 |
| YFR048W | 19.2180104 | 0.68560109 | 0.77433005 | 1.74447547 | 0.41801509 | 0.58931273 | RMD8 | VI | 246146 | 248134 | 1 |
| YFR050C | 25.2914112 | -0.1091348 | 0.74475999 | 0.84461982 | 0.65553085 | 0.78495521 | PRE4 | VI | 249066 | 249866 | -1 |
| YFR051C | 60.3495951 | 0.70853299 | 0.56778139 | 1.89991044 | 0.38675834 | 0.55927343 | RET2 | VI | 250163 | 251803 | -1 |
| YFR052W | 16.8714588 | 0.01119238 | 0.76987072 | 0.0017084 | 0.99914617 | 0.99914617 | RPN12 | VI | 252505 | 253329 | 1 |
| YFR053C | 13.0244428 | -0.7974828 | 0.88571578 | 0.84734054 | 0.6546397 | 0.78442961 | HXK1 | VI | 253592 | 255049 | -1 |
| YFR054C | 27.9680353 | -1.367148 | 0.70943549 | 8.30080078 | 0.01575811 | 0.05576982 | NA | VI | 258855 | 259433 | -1 |
| YFR055W | 35.243253 | -0.0166624 | 0.68557759 | 3.36776375 | 0.1856519 | 0.34157118 | IRC7 | VI | 264204 | 265226 | 1 |
| YFR057W | 5.74916313 | 2.85365544 | 1.49778443 | 4.60778321 | 0.09986943 | 0.21918634 | NA | VI | 269061 | 269516 | 1 |
| YGL001C | 18.2355552 | 0.48414876 | 0.88338769 | 0.49610238 | 0.78032 | 0.86685587 | ERG26 | VII | 495453 | 496502 | -1 |
| YGL002W | 94.1730664 | -0.0085874 | 0.58824091 | 5.36202146 | 0.06849389 | 0.16558648 | ERP6 | VII | 494517 | 495167 | 1 |

| | | | | | | | | | | | |
|---|---|---|---|---|---|---|---|---|---|---|---|
| YGL003C | 69.7960078 | -0.6511638 | 0.72718541 | 8.61358006 | 0.01347674 | 0.04923049 | CDH1 | VII | 492474 | 494174 | -1 |
| YGL004C | 99.2284674 | -0.0239036 | 0.52582526 | 0.15599634 | 0.92496612 | 0.95277599 | RPN14 | VII | 490703 | 491956 | -1 |
| YGL005C | 18.6169006 | -0.2747252 | 0.74945779 | 0.22357889 | 0.89423252 | 0.9350658 | COG7 | VII | 489708 | 490547 | -1 |
| YGL006W | 58.2576805 | 0.40120213 | 0.71304522 | 0.505838 | 0.77653078 | 0.8639961 | PMC1 | VII | 485921 | 489442 | 1 |
| YGL006W-A | 23.4880338 | 0.43675586 | 0.79594404 | 7.79330031 | 0.02030983 | 0.06685272 | NA | VII | 485423 | 485533 | 1 |
| YGL007C-A | 11.8198026 | -0.4783951 | 0.87607286 | 0.93358087 | 0.62701148 | 0.76477701 | NA | VII | 482941 | 483027 | -1 |
| YGL007W | 129.288991 | -0.1698473 | 0.56501805 | 0.8448618 | 0.65545154 | 0.78495521 | NA | VII | 483260 | 483637 | 1 |
| YGL008C | 258.808259 | -0.978778 | 0.71926685 | 10.092885 | 0.00643218 | 0.02793871 | PMA1 | VII | 479910 | 482666 | -1 |
| YGL009C | 15.3534269 | 1.37437516 | 0.91963846 | 2.45150306 | 0.29353702 | 0.46794585 | LEU1 | VII | 476313 | 478652 | -1 |
| YGL010W | 49.0758573 | -0.1431756 | 0.65344601 | 1.989782 | 0.36976375 | 0.54308878 | MPO1 | VII | 475545 | 476069 | 1 |
| YGL011C | 45.0174907 | 0.09013601 | 0.60955069 | 0.26225704 | 0.87710504 | 0.92756862 | SCL1 | VII | 474489 | 475247 | -1 |
| YGL012W | 271.898275 | 0.39476063 | 0.48919152 | 7.5756963 | 0.02264428 | 0.07271789 | ERG4 | VII | 472855 | 474276 | 1 |
| YGL013C | 65.29006 | -1.1017288 | 0.54801851 | 13.8375965 | 0.00098902 | 0.00652211 | PDR1 | VII | 469092 | 472298 | -1 |
| YGL014W | 26.4384243 | -0.3732575 | 0.70776378 | 0.62756808 | 0.73067681 | 0.83678999 | PUF4 | VII | 466141 | 468807 | 1 |
| YGL015C | 88.4477818 | 1.65062505 | 0.90091391 | 11.7064 | 0.0028707 | 0.01473523 | NA | VII | 465043 | 465435 | -1 |
| YGL016W | 17.819243 | 0.36334584 | 0.7943979 | 0.26987018 | 0.87377263 | 0.92573532 | KAP122 | VII | 461666 | 464911 | 1 |
| YGL017W | 18.9771416 | -1.0336743 | 0.81731904 | 2.4784871 | 0.28960321 | 0.46358932 | ATE1 | VII | 459854 | 461365 | 1 |
| YGL018C | 38.4878818 | 2.50979543 | 1.05373176 | 9.53281947 | 0.00851088 | 0.03484104 | JAC1 | VII | 459110 | 459664 | -1 |
| YGL019W | 238.104768 | -0.1925903 | 0.48876706 | 10.8936745 | 0.00430991 | 0.02028813 | CKB1 | VII | 458156 | 458992 | 1 |
| YGL020C | 64.0521661 | 1.34353966 | 0.95795385 | 6.9035237 | 0.03168975 | 0.09322158 | GET1 | VII | 457163 | 457870 | -1 |
| YGL021W | 79.4356268 | 0.66739982 | 0.56078061 | 1.41884388 | 0.49192848 | 0.6530873 | ALK1 | VII | 454785 | 457067 | 1 |
| YGL022W | 60.1428798 | 0.10257358 | 0.7321481 | 0.71040781 | 0.70103049 | 0.81801018 | STT3 | VII | 452404 | 454560 | 1 |
| YGL023C | 78.7708793 | 0.77858894 | 0.60067161 | 1.70660988 | 0.42600468 | 0.59578916 | PIB2 | VII | 450197 | 452104 | -1 |
| YGL025C | 49.0243762 | 0.00035121 | 0.62828057 | 4.43260204 | 0.1090116 | 0.23299267 | PGD1 | VII | 448764 | 449957 | -1 |
| YGL026C | 56.9355815 | 0.89066202 | 0.66603432 | 1.81844023 | 0.40283827 | 0.5737844 | TRP5 | VII | 446412 | 448535 | -1 |
| YGL027C | 97.5325178 | -1.210342 | 0.55006326 | 15.376968 | 0.00045807 | 0.00342224 | CWH41 | VII | 443642 | 446143 | -1 |
| YGL028C | 43.8483897 | -0.0286912 | 0.74659186 | 11.2718932 | 0.0035673 | 0.01754856 | SCW11 | VII | 441281 | 442909 | -1 |
| YGL029W | 53.0829909 | -1.4775676 | 0.58784773 | 23.7433666 | 6.99E-06 | 9.54E-05 | CGR1 | VII | 440063 | 440425 | 1 |
| YGL030W | 52.0978126 | -0.4038858 | 0.63972747 | 15.4363691 | 0.00044467 | 0.00334801 | RPL30 | VII | 439091 | 439638 | 1 |
| YGL031C | 101.672727 | -0.9511231 | 0.74445984 | 17.1266849 | 0.00019098 | 0.00163818 | RPL24A | VII | 437467 | 437934 | -1 |
| YGL034C | 50.7483735 | -1.122297 | 0.64413253 | 3.78630373 | 0.1505964 | 0.2934881 | NA | VII | 433214 | 433579 | -1 |
| YGL035C | 12.2809515 | -1.1267923 | 0.92369723 | 1.9154664 | 0.38376181 | 0.55688744 | MIG1 | VII | 431548 | 433062 | -1 |
| YGL036W | 91.723337 | 0.33692144 | 0.55403286 | 5.8488582 | 0.05369534 | 0.13805329 | NA | VII | 428601 | 431330 | 1 |

| | | | | | | | | | | | |
|---|---|---|---|---|---|---|---|---|---|---|---|
| YGL037C | 83.6235943 | 0.78959516 | 0.62999844 | 2.18087672 | 0.33606914 | 0.50936213 | PNC1 | VII | 427297 | 427947 | -1 |
| YGL038C | 33.3839526 | 0.51810963 | 0.83954914 | 0.52320405 | 0.76981733 | 0.85980755 | OCH1 | VII | 425362 | 426804 | -1 |
| YGL039W | 80.2761654 | -0.0971408 | 0.53685863 | 0.22960253 | 0.89154331 | 0.93352778 | NA | VII | 423961 | 425007 | 1 |
| YGL040C | 63.0124445 | -0.4310425 | 0.60726839 | 1.79565903 | 0.40745307 | 0.57879129 | HEM2 | VII | 419527 | 420555 | -1 |
| YGL041C-B | 573.0519 | -0.4775039 | 0.49238237 | 9.3532974 | 0.00931016 | 0.03735679 | NA | VII | 418704 | 418886 | -1 |
| YGL041W-A | 58.1610952 | 0.27946037 | 0.58490487 | 0.23502883 | 0.8891277 | 0.93304075 | NA | VII | 418825 | 419289 | 1 |
| YGL043W | 22.7465027 | 0.63078441 | 0.79319466 | 1.08635002 | 0.58090095 | 0.72940182 | DST1 | VII | 417483 | 418412 | 1 |
| YGL044C | 58.3989643 | 0.05789574 | 0.62168336 | 3.09542334 | 0.21273422 | 0.37639175 | RNA15 | VII | 416146 | 417036 | -1 |
| YGL045W | 42.1145364 | -0.1297672 | 0.86392685 | 12.0534229 | 0.00241342 | 0.01287898 | RIM8 | VII | 414102 | 415730 | 1 |
| YGL047W | 8.46088943 | -0.7889235 | 1.11791495 | 0.5982417 | 0.7414698 | 0.84243165 | ALG13 | VII | 411552 | 412160 | 1 |
| YGL048C | 59.4679122 | -0.151859 | 0.75531212 | 1.10716215 | 0.57488741 | 0.72422968 | RPT6 | VII | 410069 | 411286 | -1 |
| YGL049C | 105.279249 | 0.23694462 | 0.5184622 | 0.54807512 | 0.76030352 | 0.85478784 | TIF4632 | VII | 406860 | 409604 | -1 |
| YGL050W | 249.523882 | 0.20834918 | 0.50334367 | 1.03990744 | 0.59454806 | 0.74174387 | TYW3 | VII | 405776 | 406597 | 1 |
| YGL051W | 720.08248 | -0.533369 | 0.49620897 | 2.85800967 | 0.23954719 | 0.40781483 | MST27 | VII | 403687 | 404391 | 1 |
| YGL053W | 223.542802 | -0.925306 | 0.48219266 | 11.3534898 | 0.00342469 | 0.01690447 | PRM8 | VII | 402589 | 403302 | 1 |
| YGL054C | 9.12453776 | -0.5471389 | 0.98396238 | 1.27718469 | 0.52803519 | 0.68243834 | ERV14 | VII | 400871 | 401287 | -1 |
| YGL055W | 123.108241 | 0.34632446 | 0.55306625 | 1.89723417 | 0.38727622 | 0.55960303 | OLE1 | VII | 398628 | 400160 | 1 |
| YGL056C | 139.074079 | 1.5526011 | 0.66734435 | 24.8998263 | 3.92E-06 | 5.80E-05 | SDS23 | VII | 396035 | 397618 | -1 |
| YGL057C | 69.8702618 | -1.6393679 | 0.59311932 | 8.70440795 | 0.0128784 | 0.04764596 | GEP7 | VII | 394966 | 395829 | -1 |
| YGL058W | 43.4682959 | -0.5813494 | 0.7111576 | 1.85247259 | 0.39604149 | 0.56717295 | RAD6 | VII | 393986 | 394504 | 1 |
| YGL059W | 162.978222 | 0.84982878 | 0.5471653 | 23.3988694 | 8.30E-06 | 0.00011148 | PKP2 | VII | 392223 | 393698 | 1 |
| YGL060W | 16.4445429 | -0.6558402 | 0.84392325 | 0.68329442 | 0.71059885 | 0.82438173 | YBP2 | VII | 390065 | 391990 | 1 |
| YGL061C | 240.52493 | 2.10605025 | 0.49932617 | 19.1484129 | 6.95E-05 | 0.00069022 | DUO1 | VII | 388968 | 389711 | -1 |
| YGL062W | 91.211938 | 0.61389987 | 0.61433211 | 2.44491979 | 0.29450482 | 0.46884326 | PYC1 | VII | 385196 | 388732 | 1 |
| YGL063W | 139.128618 | -1.0064666 | 0.5362901 | 4.19468707 | 0.12278216 | 0.25398668 | PUS2 | VII | 383654 | 384766 | 1 |
| YGL064C | 106.605238 | -0.7716627 | 0.60553444 | 5.21162938 | 0.07384295 | 0.1754414 | MRH4 | VII | 381523 | 383208 | -1 |
| YGL065C | 375.732108 | 0.77913936 | 0.63378709 | 2.12925176 | 0.34485685 | 0.51822506 | ALG2 | VII | 379760 | 381271 | -1 |
| YGL066W | 120.231152 | 0.57878188 | 0.59116636 | 24.1824759 | 5.61E-06 | 7.86E-05 | SGF73 | VII | 377609 | 379582 | 1 |
| YGL067W | 20.0323337 | -0.6747354 | 0.78035742 | 6.16725724 | 0.04579279 | 0.12212817 | NPY1 | VII | 376101 | 377255 | 1 |
| YGL068W | 82.034876 | -0.2343832 | 0.59474347 | 5.49064764 | 0.0642275 | 0.15764189 | MNP1 | VII | 375087 | 375671 | 1 |
| YGL071W | 74.7011048 | 0.55493233 | 0.64250988 | 0.82832113 | 0.66089483 | 0.78866056 | AFT1 | VII | 372012 | 374084 | 1 |
| YGL073W | 1657.32762 | -0.2402677 | 0.45108674 | 3.9700182 | 0.13737936 | 0.27509737 | HSF1 | VII | 368753 | 371254 | 1 |
| YGL075C | 7.38898844 | -1.6986499 | 1.14743626 | 2.31797102 | 0.31380437 | 0.48580944 | MPS2 | VII | 366925 | 368088 | -1 |

| | | | | | | | | | | | |
|---|---|---|---|---|---|---|---|---|---|---|---|
| YGL076C | 53.0776461 | -0.8310598 | 0.80773186 | 9.09407969 | 0.01059853 | 0.04121256 | RPL7A | VII | 364335 | 365996 | -1 |
| YGL077C | 1267.9299 | 0.77675485 | 0.54866685 | 1.99113474 | 0.36951373 | 0.54287859 | HNM1 | VII | 362225 | 363916 | -1 |
| YGL078C | 229.554652 | -0.8843571 | 0.49054655 | 25.9773504 | 2.29E-06 | 3.61E-05 | DBP3 | VII | 360288 | 361859 | -1 |
| YGL079W | 379.220671 | -1.1015408 | 0.47933406 | 5.27642334 | 0.071489 | 0.1715269 | KXD1 | VII | 359444 | 360100 | 1 |
| YGL080W | 18.1243303 | 0.03228807 | 0.78642072 | 0.80562109 | 0.66843873 | 0.79450097 | MPC1 | VII | 358636 | 359028 | 1 |
| YGL081W | 43.5545693 | -0.2325397 | 0.61616753 | 2.76233337 | 0.25128521 | 0.42270232 | NA | VII | 357377 | 358339 | 1 |
| YGL082W | 17.5138914 | -0.0290429 | 0.81920343 | 1.60795944 | 0.44754431 | 0.61433892 | NA | VII | 355827 | 356972 | 1 |
| YGL083W | 576.818578 | 0.29710478 | 0.46179791 | 0.4148555 | 0.81267196 | 0.88731158 | SCY1 | VII | 353058 | 355472 | 1 |
| YGL084C | 49.2554351 | 1.3531577 | 0.72497956 | 4.47377235 | 0.10679051 | 0.22957814 | GUP1 | VII | 350616 | 352298 | -1 |
| YGL085W | 67.4886169 | 0.95936498 | 0.66855335 | 9.45513432 | 0.00884797 | 0.03585007 | LCL3 | VII | 349616 | 350440 | 1 |
| YGL086W | 32.7756323 | -0.9899602 | 0.64599416 | 2.73374587 | 0.25490281 | 0.42716573 | MAD1 | VII | 347119 | 349368 | 1 |
| YGL087C | 29.654439 | 0.88298197 | 0.70152432 | 2.03550281 | 0.36140669 | 0.53408966 | MMS2 | VII | 346406 | 346904 | -1 |
| YGL089C | 357.329818 | 0.28451359 | 0.46325566 | 0.82670201 | 0.66143008 | 0.78897407 | MF(ALPHA)2 | VII | 344791 | 345153 | -1 |
| YGL090W | 752.796528 | -3.3983663 | 0.62277664 | 50.9094892 | 8.81E-12 | 5.32E-10 | LIF1 | VII | 343319 | 344584 | 1 |
| YGL091C | 42.8939327 | -0.3312127 | 0.60886502 | 3.61095192 | 0.16439619 | 0.31321288 | NBP35 | VII | 342056 | 343042 | -1 |
| YGL092W | 48.8461922 | 1.39115295 | 0.75744326 | 6.6637846 | 0.03572544 | 0.10209787 | NUP145 | VII | 337906 | 341859 | 1 |
| YGL093W | 23.9984831 | -0.5038402 | 0.73232111 | 0.48067489 | 0.78636246 | 0.87021248 | SPC105 | VII | 334886 | 337639 | 1 |
| YGL094C | 19.6884986 | -0.4232479 | 0.75818373 | 0.32116502 | 0.85164755 | 0.91195795 | PAN2 | VII | 331118 | 334465 | -1 |
| YGL095C | 88.5586891 | 1.46551986 | 0.84735293 | 13.0755731 | 0.00144769 | 0.00890767 | VPS45 | VII | 328874 | 330607 | -1 |
| YGL096W | 86.1662258 | -0.0888968 | 0.55814873 | 2.38076522 | 0.30410489 | 0.47720463 | TOS8 | VII | 325331 | 326161 | 1 |
| YGL097W | 84.7872075 | -0.0991294 | 0.53767413 | 0.25745261 | 0.87921457 | 0.92911891 | SRM1 | VII | 321782 | 323230 | 1 |
| YGL098W | 13.5829956 | 0.6682381 | 1.31856219 | 0.45666729 | 0.79585868 | 0.8765497 | USE1 | VII | 317342 | 318079 | 1 |
| YGL099W | 19.1293074 | -1.2092124 | 0.75467548 | 2.68797851 | 0.26080318 | 0.43235173 | LSG1 | VII | 314631 | 316553 | 1 |
| YGL100W | 26.0745841 | -0.119697 | 0.67105025 | 1.7331452 | 0.42038993 | 0.59107763 | SEH1 | VII | 313234 | 314283 | 1 |
| YGL101W | 67.0938327 | -1.3698063 | 0.55350073 | 9.8284474 | 0.00734141 | 0.03116334 | NA | VII | 312193 | 312840 | 1 |
| YGL103W | 290.015156 | -0.7908004 | 0.71560188 | 7.97174877 | 0.01857619 | 0.06260545 | RPL28 | VII | 310967 | 311927 | 1 |
| YGL104C | 112.894762 | 0.08147563 | 0.51723714 | 0.05445076 | 0.97314189 | 0.97940058 | VPS73 | VII | 308713 | 310173 | -1 |
| YGL105W | 208.334285 | 0.23876124 | 0.50772453 | 13.2547619 | 0.00132363 | 0.00825839 | ARC1 | VII | 307437 | 308567 | 1 |
| YGL106W | 20.8700901 | -0.0715163 | 0.72708064 | 1.89002894 | 0.38867395 | 0.56060856 | MLC1 | VII | 306560 | 307009 | 1 |
| YGL107C | 24.3592471 | -0.9918567 | 0.77333836 | 3.14507885 | 0.20751754 | 0.36958676 | RMD9 | VII | 304333 | 306273 | -1 |
| YGL108C | 27.8800794 | 0.90412567 | 0.75554879 | 2.49713716 | 0.2869152 | 0.46119906 | NA | VII | 303649 | 304071 | -1 |
| YGL110C | 112.48558 | 0.65988297 | 0.71752953 | 0.97376909 | 0.61453798 | 0.75689745 | CUE3 | VII | 301537 | 303411 | -1 |
| YGL111W | 81.6624903 | -0.2557366 | 0.59744528 | 0.56129862 | 0.75529316 | 0.85213317 | NSA1 | VII | 299978 | 301369 | 1 |

| | | | | | | | | | | | |
|---|---|---|---|---|---|---|---|---|---|---|---|
| YGL112C | 50.7931724 | 1.01118574 | 0.75153747 | 1.92571434 | 0.38180046 | 0.55529381 | TAF6 | VII | 298178 | 299728 | -1 |
| YGL113W | 106.305412 | 1.52418235 | 0.60711899 | 32.1529831 | 1.04E-07 | 2.51E-06 | SLD3 | VII | 295932 | 297938 | 1 |
| YGL114W | 34.2179209 | -0.1330527 | 0.65211133 | 0.256511 | 0.8796286 | 0.92911891 | NA | VII | 293460 | 295637 | 1 |
| YGL115W | 63.4187715 | 0.39431397 | 0.58267595 | 1.30598405 | 0.52048614 | 0.6765916 | SNF4 | VII | 292033 | 293001 | 1 |
| YGL116W | 20.7691671 | -0.1518356 | 0.75645225 | 0.08268418 | 0.95950084 | 0.97158269 | CDC20 | VII | 289809 | 291641 | 1 |
| YGL117W | 24.8455311 | -1.6845295 | 0.72598862 | 14.756896 | 0.00062457 | 0.00444804 | NA | VII | 288512 | 289309 | 1 |
| YGL118C | 122.706991 | 1.30654241 | 0.54143534 | 5.81817472 | 0.05452547 | 0.13962957 | NA | VII | 288014 | 288451 | -1 |
| YGL119W | 16.1089108 | -0.4717637 | 0.85055506 | 1.55609708 | 0.45930145 | 0.62382251 | COQ8 | VII | 284442 | 285947 | 1 |
| YGL120C | 25.4212832 | -0.5288899 | 0.83304063 | 1.06837762 | 0.58614457 | 0.73378964 | PRP43 | VII | 281634 | 283937 | -1 |
| YGL121C | 59.453868 | -1.3035105 | 0.57248382 | 14.1887399 | 0.00082976 | 0.00565881 | GPG1 | VII | 280777 | 281157 | -1 |
| YGL122C | 305.348737 | 1.91121493 | 0.61033271 | 26.5898346 | 1.68E-06 | 2.80E-05 | NAB2 | VII | 278946 | 280523 | -1 |
| YGL123W | 928.086072 | -1.1240607 | 0.54656736 | 26.1287237 | 2.12E-06 | 3.36E-05 | RPS2 | VII | 277617 | 278381 | 1 |
| YGL124C | 349.881209 | 0.43683902 | 0.48222528 | 11.6836261 | 0.00290357 | 0.01483821 | MON1 | VII | 274780 | 276714 | -1 |
| YGL125W | 915.407718 | -0.4876613 | 0.52016251 | 0.95398435 | 0.62064739 | 0.76119942 | MET13 | VII | 272520 | 274322 | 1 |
| YGL126W | 169.798592 | 0.23542467 | 0.5346639 | 4.37547026 | 0.11217051 | 0.23842411 | SCS3 | VII | 270997 | 272139 | 1 |
| YGL127C | 45.7442302 | 1.44089411 | 0.67225418 | 12.8209779 | 0.00164422 | 0.00973419 | SOH1 | VII | 270392 | 270775 | -1 |
| YGL128C | 51.9198862 | -0.4329026 | 0.57456398 | 0.96740473 | 0.61649667 | 0.75817303 | CWC23 | VII | 269293 | 270144 | -1 |
| YGL129C | 30.2745529 | 0.01777655 | 0.65034015 | 0.30459998 | 0.85873063 | 0.91650698 | RSM23 | VII | 267723 | 269075 | -1 |
| YGL130W | 61.9859059 | 0.00626756 | 0.57921809 | 1.13517833 | 0.56689047 | 0.71775548 | CEG1 | VII | 266145 | 267524 | 1 |
| YGL131C | 57.2550088 | 0.02482853 | 0.58883811 | 3.94697428 | 0.1389714 | 0.27755929 | SNT2 | VII | 261648 | 265859 | -1 |
| YGL133W | 2247.16268 | 0.56886748 | 0.55255691 | 2.18875857 | 0.33474733 | 0.50817698 | ITC1 | VII | 257707 | 261501 | 1 |
| YGL134W | 93.712272 | -0.0891479 | 0.53639147 | 0.05702772 | 0.97188883 | 0.97864979 | PCL10 | VII | 255663 | 256964 | 1 |
| YGL135W | 27.5218963 | -0.9416045 | 0.91934389 | 8.64296652 | 0.01328017 | 0.04868773 | RPL1A | VII | 254641 | 255294 | 1 |
| YGL136C | 1250.81117 | 0.92772044 | 0.54696316 | 9.35390782 | 0.00930732 | 0.03735679 | MRM2 | VII | 252897 | 253859 | -1 |
| YGL137W | 275.033074 | 0.25484637 | 0.4677472 | 0.62784103 | 0.7305771 | 0.83678999 | SEC27 | VII | 249869 | 252738 | 1 |
| YGL139W | 16.1843891 | -0.7036646 | 0.79926803 | 1.78739463 | 0.40914023 | 0.58076047 | FLC3 | VII | 245716 | 248124 | 1 |
| YGL140C | 34.3666264 | 0.58261016 | 0.81930407 | 0.52772989 | 0.76807727 | 0.85918999 | NA | VII | 241353 | 245012 | -1 |
| YGL141W | 61.7083648 | 1.17525259 | 0.70701952 | 3.0627851 | 0.21623434 | 0.38031495 | HUL5 | VII | 238353 | 241085 | 1 |
| YGL142C | 32.3623598 | 0.25438034 | 0.6508483 | 0.91112345 | 0.63409168 | 0.77081479 | GPI10 | VII | 236269 | 238119 | -1 |
| YGL143C | 26.5822723 | 0.03363291 | 0.73781973 | 3.75622647 | 0.15287828 | 0.29674779 | MRF1 | VII | 234717 | 235958 | -1 |
| YGL144C | 101.266652 | 1.47233136 | 0.69632816 | 5.46410341 | 0.06508562 | 0.15927065 | ROG1 | VII | 232450 | 234507 | -1 |
| YGL145W | 11.9580561 | 0.20534172 | 0.8795636 | 0.20814208 | 0.90116128 | 0.93824416 | TIP20 | VII | 230243 | 232348 | 1 |
| YGL146C | 27.9101196 | -1.7979137 | 0.77425622 | 5.39942144 | 0.06722496 | 0.16286148 | RRT6 | VII | 228751 | 229686 | -1 |

| | | | | | | | | | | | |
|---|---|---|---|---|---|---|---|---|---|---|---|
| YGL147C | 152.941056 | -1.4315125 | 0.84812967 | 21.7143644 | 1.93E-05 | 0.00022952 | RPL9A | VII | 227754 | 228329 | -1 |
| YGL148W | 70.3199525 | -0.7311809 | 0.58649464 | 4.82926789 | 0.08940006 | 0.20267482 | ARO2 | VII | 226399 | 227529 | 1 |
| YGL149W | 509.369016 | 0.21426341 | 0.589943 | 2.39490932 | 0.30196183 | 0.47448396 | NA | VII | 225572 | 225877 | 1 |
| YGL150C | 155.229915 | 1.47800665 | 0.67802815 | 6.22626277 | 0.04446151 | 0.11984737 | INO80 | VII | 221104 | 225573 | -1 |
| YGL151W | 68.178915 | 0.72569544 | 0.64846948 | 2.90153422 | 0.23439042 | 0.40199067 | NUT1 | VII | 217524 | 220922 | 1 |
| YGL153W | 172.605315 | -0.6687335 | 0.59997198 | 8.10726793 | 0.01735918 | 0.05959198 | PEX14 | VII | 216273 | 217298 | 1 |
| YGL154C | 452.424049 | -0.6458488 | 0.4657183 | 18.9659443 | 7.61E-05 | 0.0007409 | LYS5 | VII | 215278 | 216096 | -1 |
| YGL155W | 38.8748404 | 0.27926345 | 0.79712982 | 0.99674467 | 0.60751869 | 0.7514491 | CDC43 | VII | 214081 | 215211 | 1 |
| YGL156W | 512.098438 | -2.4083679 | 0.53942448 | 48.6913961 | 2.67E-11 | 1.42E-09 | AMS1 | VII | 210416 | 213667 | 1 |
| YGL157W | 65.314318 | 1.49381619 | 0.71373841 | 9.55595803 | 0.00841298 | 0.03453249 | ARI1 | VII | 209006 | 210049 | 1 |
| YGL158W | 38.6731739 | 1.06087466 | 0.71353288 | 8.0713957 | 0.01767334 | 0.06020484 | RCK1 | VII | 207033 | 208571 | 1 |
| YGL159W | 63.0111839 | -1.688517 | 0.62001111 | 7.86625421 | 0.01958234 | 0.06501327 | NA | VII | 202721 | 203833 | 1 |
| YGL160W | 27.4260709 | -0.6216503 | 0.71999437 | 0.99096956 | 0.60927547 | 0.75306988 | AIM14 | VII | 200561 | 202273 | 1 |
| YGL161C | 62.3310456 | -0.1554981 | 0.62030343 | 0.14887382 | 0.92826604 | 0.95498585 | YIP5 | VII | 199210 | 200142 | -1 |
| YGL162W | 14.4182685 | -0.0943784 | 0.80023294 | 8.21551515 | 0.01644461 | 0.05722012 | SUT1 | VII | 198138 | 199037 | 1 |
| YGL163C | 27.6063779 | 0.75213391 | 0.73890453 | 2.35817081 | 0.3075599 | 0.48012182 | RAD54 | VII | 193707 | 196403 | -1 |
| YGL164C | 84.093683 | 1.09505976 | 0.57981985 | 3.81281166 | 0.14861357 | 0.29099512 | YRB30 | VII | 191978 | 193300 | -1 |
| YGL166W | 36.1747039 | 1.34904796 | 0.66388552 | 7.60943999 | 0.02226543 | 0.07177998 | CUP2 | VII | 191129 | 191806 | 1 |
| YGL167C | 48.8100926 | 0.18211069 | 0.66769553 | 1.10524503 | 0.57543873 | 0.72461728 | PMR1 | VII | 187616 | 190468 | -1 |
| YGL168W | 5495.69473 | 0.69373784 | 0.4473464 | 4.3167329 | 0.11551366 | 0.242911 | HUR1 | VII | 187464 | 187796 | 1 |
| YGL169W | 27.2474897 | 0.34424559 | 0.77065781 | 1.56723385 | 0.45675099 | 0.62210967 | SUA5 | VII | 186059 | 187339 | 1 |
| YGL170C | 286.923757 | -0.4035178 | 0.78550661 | 14.1949278 | 0.0008272 | 0.00564798 | SPO74 | VII | 184153 | 185394 | -1 |
| YGL171W | 20.8947615 | -0.8821786 | 0.7796987 | 1.8710628 | 0.3923773 | 0.56387803 | ROK1 | VII | 182390 | 184084 | 1 |
| YGL172W | 19.4032111 | -0.561158 | 0.79075031 | 1.39328788 | 0.49825467 | 0.6579007 | NUP49 | VII | 180700 | 182118 | 1 |
| YGL173C | 67.1401684 | 0.06466518 | 0.59514195 | 1.97875953 | 0.37180723 | 0.54514152 | XRN1 | VII | 175527 | 180113 | -1 |
| YGL174W | 87.4668753 | 0.56136347 | 0.7542819 | 6.86974545 | 0.03222951 | 0.09435142 | BUD13 | VII | 174545 | 175345 | 1 |
| YGL175C | 240.078984 | 0.27567654 | 0.54997632 | 24.1841826 | 5.60E-06 | 7.86E-05 | SAE2 | VII | 173285 | 174322 | -1 |
| YGL176C | 195.05709 | 0.3392696 | 0.56456485 | 39.8516156 | 2.22E-09 | 7.70E-08 | NA | VII | 171415 | 173079 | -1 |
| YGL178W | 122.179277 | 0.32175845 | 0.51848362 | 0.60525069 | 0.73887587 | 0.84040603 | MPT5 | VII | 167352 | 170571 | 1 |
| YGL179C | 26.1600606 | -1.6038581 | 0.7079441 | 5.44016662 | 0.06586927 | 0.16064998 | TOS3 | VII | 163409 | 165091 | -1 |
| YGL180W | 108.468344 | -2.0989196 | 0.57057186 | 20.0585478 | 4.41E-05 | 0.00046183 | ATG1 | VII | 160065 | 162758 | 1 |
| YGL181W | 42.6780784 | 1.68221148 | 0.67424451 | 6.44875142 | 0.03978061 | 0.11116077 | GTS1 | VII | 157906 | 159096 | 1 |
| YGL184C | 107.571463 | -0.0972469 | 0.54541032 | 4.65916543 | 0.09733636 | 0.21492841 | STR3 | VII | 154615 | 156012 | -1 |

| | | | | | | | | | | | |
|---|---|---|---|---|---|---|---|---|---|---|---|
| YGL185C | 23.512965 | -0.8656605 | 0.78262948 | 2.13886403 | 0.3432034 | 0.51667906 | NA | VII | 153167 | 154306 | -1 |
| YGL186C | 37.1146483 | -0.0719365 | 0.65999439 | 1.39176741 | 0.49863361 | 0.65825093 | TPN1 | VII | 151037 | 152776 | -1 |
| YGL187C | 58.3336215 | -0.686626 | 0.60566242 | 11.8597823 | 0.00265877 | 0.01389376 | COX4 | VII | 149704 | 150171 | -1 |
| YGL189C | 322.787215 | -0.4429115 | 0.61647587 | 20.1128683 | 4.29E-05 | 0.00045254 | RPS26A | VII | 148229 | 148588 | -1 |
| YGL190C | 52.8698749 | 0.72338648 | 0.63237095 | 3.38558598 | 0.18400488 | 0.3396201 | CDC55 | VII | 145809 | 147389 | -1 |
| YGL191W | 25.6908855 | 0.31704669 | 0.78720531 | 2.14296088 | 0.34250109 | 0.51615859 | COX13 | VII | 144808 | 145197 | 1 |
| YGL192W | 7094.27745 | 0.09480079 | 0.5235095 | 0.5639851 | 0.7542793 | 0.85152235 | IME4 | VII | 142246 | 144048 | 1 |
| YGL194C | 339.018628 | 0.97686692 | 0.48744961 | 17.1353681 | 0.00019015 | 0.00163351 | HOS2 | VII | 140368 | 141726 | -1 |
| YGL194C-A | 82.0249815 | 0.61818315 | 0.64302755 | 7.48454727 | 0.02370016 | 0.07502674 | NA | VII | 139719 | 139961 | -1 |
| YGL195W | 77.4018836 | 0.44749748 | 0.61694658 | 2.25111576 | 0.3244714 | 0.49763242 | GCN1 | VII | 131525 | 139543 | 1 |
| YGL196W | 38.8342515 | -1.2495281 | 0.70144995 | 8.2567839 | 0.01610876 | 0.05656139 | DSD1 | VII | 129883 | 131169 | 1 |
| YGL197W | 43.2867104 | 1.10675847 | 0.63739791 | 3.05997539 | 0.21653833 | 0.38073396 | MDS3 | VII | 124698 | 129161 | 1 |
| YGL198W | 53.1472986 | -0.2215971 | 0.57105252 | 1.40890851 | 0.4943783 | 0.65483292 | YIP4 | VII | 123591 | 124298 | 1 |
| YGL200C | 206.074287 | 0.12815354 | 0.75253237 | 0.68716697 | 0.70922427 | 0.82408359 | EMP24 | VII | 122694 | 123305 | -1 |
| YGL201C | 62.1026248 | 0.07600348 | 0.58788115 | 7.59498695 | 0.02242692 | 0.07215694 | MCM6 | VII | 117854 | 120907 | -1 |
| YGL202W | 92.047105 | 0.98726721 | 0.59878709 | 4.34554675 | 0.1138614 | 0.24051715 | ARO8 | VII | 116059 | 117561 | 1 |
| YGL203C | 157.096059 | 1.41266525 | 0.66894241 | 4.4757802 | 0.10668336 | 0.22954167 | KEX1 | VII | 112475 | 114664 | -1 |
| YGL204C | 22.2169155 | 0.89651428 | 0.91561072 | 13.0412406 | 0.00147276 | 0.00899499 | NA | VII | 111700 | 112005 | -1 |
| YGL205W | 668.500253 | -1.0037605 | 0.49121422 | 5.96624431 | 0.0506345 | 0.13188203 | POX1 | VII | 108158 | 110404 | 1 |
| YGL206C | 32.8787024 | 0.48448809 | 0.72161014 | 1.9207198 | 0.38275511 | 0.55598396 | CHC1 | VII | 102543 | 107504 | -1 |
| YGL207W | 306.944337 | 0.6447425 | 0.53665087 | 48.4563802 | 3.00E-11 | 1.55E-09 | SPT16 | VII | 98969 | 102076 | 1 |
| YGL208W | 90.7007233 | -1.3908859 | 0.60577674 | 5.69287621 | 0.05805072 | 0.14632725 | SIP2 | VII | 97338 | 98585 | 1 |
| YGL209W | 308.547492 | 1.35086688 | 0.51296868 | 10.7864489 | 0.00454729 | 0.02111371 | MIG2 | VII | 95858 | 97006 | 1 |
| YGL210W | 17.5816631 | 1.38781684 | 0.83605694 | 4.12269208 | 0.12728253 | 0.26090922 | YPT32 | VII | 93792 | 94460 | 1 |
| YGL211W | 250.322822 | 2.41839046 | 0.57691218 | 63.5058938 | 1.62E-14 | 1.51E-12 | NCS6 | VII | 92512 | 93591 | 1 |
| YGL212W | 8.57469329 | 0.26321357 | 1.01774404 | 0.49424235 | 0.78104604 | 0.86705973 | VAM7 | VII | 91432 | 92382 | 1 |
| YGL213C | 44.8018603 | -1.134944 | 0.6297554 | 5.7643959 | 0.05601152 | 0.14261508 | SKI8 | VII | 90054 | 91247 | -1 |
| YGL215W | 42.532926 | 0.47764008 | 0.67435596 | 0.6238721 | 0.73202834 | 0.83714084 | CLG1 | VII | 87981 | 89339 | 1 |
| YGL216W | 37.5954053 | 0.84921431 | 0.68719251 | 1.68895245 | 0.42978241 | 0.59904674 | KIP3 | VII | 84885 | 87302 | 1 |
| YGL219C | 1148.99621 | 2.85001089 | 0.96818156 | 10.1183028 | 0.00635095 | 0.02766891 | MDM34 | VII | 82878 | 84257 | -1 |
| YGL220W | 17.1760715 | -0.0756635 | 0.8857187 | 0.64160808 | 0.72556542 | 0.83456178 | BOL2 | VII | 82374 | 82736 | 1 |
| YGL221C | 54.772759 | -0.2612832 | 0.62571078 | 0.25621471 | 0.87975893 | 0.92911891 | NIF3 | VII | 81426 | 82292 | -1 |
| YGL222C | 34.4220691 | -0.5513597 | 0.69491868 | 0.97378765 | 0.61453228 | 0.75689745 | EDC1 | VII | 80659 | 81186 | -1 |

| | | | | | | | | | | | |
|---|---|---|---|---|---|---|---|---|---|---|---|
| YGL223C | 43.8260445 | 1.3126595 | 0.73738418 | 3.39187344 | 0.18342733 | 0.33887818 | COG1 | VII | 79112 | 80365 | -1 |
| YGL224C | 1907.65056 | 1.72002304 | 0.88026834 | 6.17543788 | 0.04560587 | 0.12191042 | SDT1 | VII | 78015 | 78857 | -1 |
| YGL225W | 28.9816834 | 0.51040836 | 0.71505453 | 1.05926894 | 0.58882016 | 0.73666135 | VRG4 | VII | 76894 | 77907 | 1 |
| YGL226C-A | 28.2472757 | 1.86647348 | 0.98911369 | 3.42675536 | 0.18025592 | 0.3349428 | OST5 | VII | 72749 | 73158 | -1 |
| YGL226W | 11.8306722 | 0.6192863 | 0.88287864 | 1.54840249 | 0.46107192 | 0.62486036 | MTC3 | VII | 73340 | 73711 | 1 |
| YGL227W | 182.366438 | -0.1803305 | 0.52769522 | 4.56130041 | 0.10221772 | 0.22276062 | VID30 | VII | 69671 | 72547 | 1 |
| YGL228W | 54.2472198 | 1.00444198 | 0.77063988 | 2.28322309 | 0.31930403 | 0.49195592 | SHE10 | VII | 67598 | 69331 | 1 |
| YGL229C | 42.9554226 | -0.5055014 | 0.68440352 | 0.57103677 | 0.75162452 | 0.84964974 | SAP4 | VII | 64503 | 66959 | -1 |
| YGL230C | 97.5783894 | 0.61069263 | 0.57460413 | 4.90512247 | 0.08607285 | 0.19729288 | NA | VII | 63772 | 64215 | -1 |
| YGL231C | 125.170884 | 1.63747051 | 0.7372802 | 11.8210464 | 0.00271077 | 0.01406419 | EMC4 | VII | 63048 | 63620 | -1 |
| YGL232W | 277.741902 | 3.38996087 | 0.96525201 | 12.7344281 | 0.00171694 | 0.00999815 | TAN1 | VII | 62075 | 63002 | 1 |
| YGL233W | 23.7761528 | -0.2697462 | 0.75693549 | 3.06668701 | 0.21581289 | 0.37968904 | SEC15 | VII | 59122 | 61854 | 1 |
| YGL234W | 99.6585088 | -0.0597464 | 0.61057511 | 12.2754569 | 0.00215982 | 0.01184098 | ADE5,7 | VII | 56482 | 58890 | 1 |
| YGL235W | 261.743606 | -1.6874775 | 0.55855459 | 30.3674698 | 2.55E-07 | 5.38E-06 | NA | VII | 55279 | 55815 | 1 |
| YGL236C | 42.1998781 | -0.4805363 | 0.67132579 | 6.41314335 | 0.04049521 | 0.11240041 | MTO1 | VII | 53787 | 55796 | -1 |
| YGL237C | 14.7157776 | -0.6981383 | 0.87693897 | 1.32814684 | 0.51475026 | 0.67171603 | HAP2 | VII | 52731 | 53528 | -1 |
| YGL238W | 27.4742485 | 0.89255322 | 0.74058165 | 1.75717594 | 0.41536901 | 0.58728945 | CSE1 | VII | 49552 | 52434 | 1 |
| YGL240W | 128.44674 | -0.4766937 | 0.56176626 | 1.00877477 | 0.60387541 | 0.74934389 | DOC1 | VII | 48613 | 49365 | 1 |
| YGL241W | 10.9410656 | 0.08892289 | 0.93083625 | 0.1070899 | 0.94786333 | 0.96435226 | KAP114 | VII | 45445 | 48459 | 1 |
| YGL242C | 51.305493 | 1.23206195 | 0.76890372 | 2.80556437 | 0.24591184 | 0.41681193 | NA | VII | 44652 | 45197 | -1 |
| YGL243W | 437.169956 | 2.7182982 | 0.83688729 | 15.4131084 | 0.00044987 | 0.00337792 | TAD1 | VII | 43307 | 44509 | 1 |
| YGL244W | 95.3922924 | 0.41361238 | 0.52561788 | 5.51731691 | 0.06337673 | 0.15623132 | RTF1 | VII | 41498 | 43174 | 1 |
| YGL245W | 93.1922743 | -0.2937438 | 0.78822347 | 4.4888891 | 0.10598639 | 0.22838155 | GUS1 | VII | 39023 | 41149 | 1 |
| YGL246C | 42.3951834 | 0.64697528 | 0.78721332 | 4.36117171 | 0.11297532 | 0.23947912 | RAI1 | VII | 37617 | 38780 | -1 |
| YGL247W | 382.397243 | 0.67647696 | 0.77015625 | 6.19027071 | 0.04526888 | 0.12151453 | BRR6 | VII | 36933 | 37526 | 1 |
| YGL248W | 55.3759054 | 0.40744961 | 0.64139592 | 0.43444614 | 0.80475043 | 0.88279205 | PDE1 | VII | 35653 | 36762 | 1 |
| YGL249W | 1329.84906 | 0.28493352 | 0.46961043 | 5.42301018 | 0.06643674 | 0.16169345 | ZIP2 | VII | 33098 | 35212 | 1 |
| YGL250W | 339.113139 | -0.0086136 | 0.53131514 | 1.38475171 | 0.50038581 | 0.65981185 | RMR1 | VII | 31910 | 32635 | 1 |
| YGL251C | 334.546996 | 0.30120841 | 0.4939532 | 15.8222579 | 0.00036664 | 0.00286098 | HFM1 | VII | 27921 | 31636 | -1 |
| YGL252C | 45.7510853 | 0.65255831 | 0.91554854 | 0.83939061 | 0.65724705 | 0.78549013 | RTG2 | VII | 25718 | 27484 | -1 |
| YGL253W | 101.465052 | 0.33901306 | 0.62702539 | 4.169413 | 0.12434361 | 0.25639227 | HXK2 | VII | 23935 | 25395 | 1 |
| YGL254W | 21.7250865 | -1.8004799 | 0.74950318 | 9.50702389 | 0.00862136 | 0.03516063 | FZF1 | VII | 22304 | 23203 | 1 |
| YGL255W | 50.6397061 | -0.8359127 | 0.62254984 | 5.11422386 | 0.07752832 | 0.18217898 | ZRT1 | VII | 20978 | 22108 | 1 |

| | | | | | | | | | | | |
|---|---|---|---|---|---|---|---|---|---|---|---|
| YGL256W | 7.57311263 | -0.3393839 | 1.21856253 | 0.94763173 | 0.62262189 | 0.76281637 | ADH4 | VII | 15159 | 16307 | 1 |
| YGL257C | 457.950788 | 0.54938978 | 0.61238925 | 4.82945096 | 0.08939188 | 0.20267482 | MNT2 | VII | 12481 | 14157 | -1 |
| YGL258W | 179.8594 | -0.2723605 | 0.50010042 | 0.47922189 | 0.78693396 | 0.87036249 | VEL1 | VII | 11110 | 11730 | 1 |
| YGL260W | 4.82872011 | 1.7645188 | 1.52354161 | 1.64734361 | 0.43881744 | 0.60769026 | NA | VII | 6860 | 7090 | 1 |
| YGR001C | 247.124933 | 1.69323043 | 0.50305509 | 10.8474836 | 0.00441061 | 0.02064466 | EFM5 | VII | 497133 | 498034 | -1 |
| YGR002C | 100.504987 | 0.69829438 | 0.57662 | 18.9589131 | 7.64E-05 | 0.00074226 | SWC4 | VII | 498476 | 499906 | -1 |
| YGR003W | 33.0487759 | -0.7310159 | 0.66704918 | 1.44201634 | 0.48626177 | 0.64797599 | CUL3 | VII | 500132 | 502366 | 1 |
| YGR004W | 33.0980316 | -0.4875936 | 0.68758219 | 5.08874167 | 0.07852244 | 0.18399228 | PEX31 | VII | 502938 | 504326 | 1 |
| YGR005C | 38.9371275 | 0.02166592 | 0.65837718 | 0.32955518 | 0.8480823 | 0.90980423 | TFG2 | VII | 504657 | 505859 | -1 |
| YGR006W | 15.5554726 | 0.78874886 | 0.82440527 | 2.24284756 | 0.32581557 | 0.49867094 | PRP18 | VII | 506070 | 506825 | 1 |
| YGR007W | 54.1340037 | 2.7644375 | 0.80083062 | 11.4304482 | 0.00329541 | 0.01642034 | ECT1 | VII | 506969 | 507940 | 1 |
| YGR008C | 187.811617 | 1.18583203 | 0.50304791 | 16.5285968 | 0.00025755 | 0.0021303 | STF2 | VII | 508110 | 508364 | -1 |
| YGR009C | 58.038851 | 0.74982142 | 0.58010594 | 2.19915814 | 0.33301123 | 0.50700368 | SEC9 | VII | 509047 | 511002 | -1 |
| YGR010W | 95.5238636 | -0.6030682 | 0.55264236 | 1.18104925 | 0.55403655 | 0.70568075 | NMA2 | VII | 511545 | 512732 | 1 |
| YGR012W | 75.6374511 | -1.4839613 | 0.64625534 | 5.19556447 | 0.07443848 | 0.17631337 | MCY1 | VII | 513159 | 514340 | 1 |
| YGR013W | 73.2854061 | 0.45485344 | 0.60508493 | 3.01368847 | 0.22160822 | 0.38636302 | SNU71 | VII | 514554 | 516416 | 1 |
| YGR014W | 43.4186146 | 0.6807214 | 0.64397079 | 3.94199242 | 0.139318 | 0.27786814 | MSB2 | VII | 516943 | 520863 | 1 |
| YGR015C | 15.2853535 | 0.06087504 | 0.84753074 | 1.42537534 | 0.49032459 | 0.65159041 | NA | VII | 521090 | 522076 | -1 |
| YGR016W | 26.8682014 | 0.1286234 | 0.75008564 | 7.94466286 | 0.01882948 | 0.06320157 | NA | VII | 522261 | 522833 | 1 |
| YGR017W | 71.4623018 | 0.32810951 | 0.56401608 | 1.7548088 | 0.41586092 | 0.58742004 | NA | VII | 523787 | 524680 | 1 |
| YGR018C | 2640.43518 | -0.2826772 | 0.46422244 | 7.5713538 | 0.0226935 | 0.07283556 | NA | VII | 524365 | 524694 | -1 |
| YGR019W | 66.4962043 | -0.3578323 | 0.54902291 | 2.58758219 | 0.27422918 | 0.44776846 | UGA1 | VII | 525229 | 526644 | 1 |
| YGR020C | 13.1517034 | -0.4504571 | 1.02800455 | 2.36957419 | 0.30581128 | 0.47867401 | VMA7 | VII | 526973 | 527329 | -1 |
| YGR021W | 29.8710104 | -0.0549819 | 0.71489973 | 4.06840579 | 0.13078469 | 0.26588601 | NA | VII | 527632 | 528504 | 1 |
| YGR023W | 33.6201665 | -0.7393417 | 0.78051542 | 1.03448333 | 0.5961627 | 0.74343787 | MTL1 | VII | 529264 | 530919 | 1 |
| YGR024C | 12.2876693 | 0.10339725 | 0.89773249 | 1.86944677 | 0.39269447 | 0.56412138 | THG1 | VII | 531883 | 532596 | -1 |
| YGR025W | 14.1128462 | 0.30731493 | 0.9400516 | 0.86173777 | 0.64994412 | 0.78139046 | NA | VII | 532639 | 532941 | 1 |
| YGR026W | 17.9142655 | 0.37208228 | 0.81226559 | 0.2931176 | 0.86367495 | 0.91991868 | NA | VII | 532985 | 533821 | 1 |
| YGR027C | 76.7771469 | 0.78228485 | 0.65769698 | 5.31459134 | 0.07013764 | 0.16864491 | RPS25A | VII | 534132 | 534458 | -1 |
| YGR028W | 25.8391292 | 0.96544034 | 0.75797489 | 6.28391878 | 0.04319807 | 0.11764668 | MSP1 | VII | 542203 | 543291 | 1 |
| YGR029W | 264.697207 | -0.1460101 | 0.52941273 | 4.76006172 | 0.09254772 | 0.20705228 | ERV1 | VII | 543553 | 544205 | 1 |
| YGR030C | 183.49793 | -1.0627159 | 0.51761564 | 11.0312802 | 0.00402335 | 0.01933212 | POP6 | VII | 545681 | 546157 | -1 |
| YGR031C-A | 861.159475 | -1.4312953 | 0.56715807 | 6.95698459 | 0.03085389 | 0.09137803 | NAG1 | VII | 546809 | 547300 | -1 |

| | | | | | | | | | | | |
|---|---|---|---|---|---|---|---|---|---|---|---|
| YGR031W | 66.1051386 | 1.54168352 | 0.61226797 | 8.71853978 | 0.01278772 | 0.04740135 | IMO32 | VII | 546441 | 547469 | 1 |
| YGR032W | 232.141312 | 2.65901199 | 0.67721461 | 15.0423525 | 0.0005415 | 0.00391418 | GSC2 | VII | 548264 | 553951 | 1 |
| YGR033C | 10.9864489 | 0.92912268 | 0.9728673 | 0.99995532 | 0.60654421 | 0.75088539 | TIM21 | VII | 554248 | 554967 | -1 |
| YGR034W | 64.4848146 | -0.4477074 | 0.7587891 | 14.7142652 | 0.00063803 | 0.00453272 | RPL26B | VII | 555812 | 556672 | 1 |
| YGR035C | 20.2639258 | 0.39381623 | 0.77954443 | 0.88826237 | 0.64138128 | 0.77591986 | NA | VII | 557072 | 557422 | -1 |
| YGR035W-A | 991.666309 | 0.2153537 | 0.54783153 | 6.10232792 | 0.04730383 | 0.12530377 | NA | VII | 557565 | 557786 | 1 |
| YGR036C | 17.2653411 | 1.60058051 | 0.84169612 | 3.65843135 | 0.16053943 | 0.30799315 | CAX4 | VII | 558149 | 558868 | -1 |
| YGR037C | 88.2969055 | 1.12944233 | 0.68266865 | 4.26627463 | 0.11846505 | 0.24726483 | ACB1 | VII | 559731 | 559994 | -1 |
| YGR038W | 517.495996 | 1.54138728 | 0.53056219 | 30.9743803 | 1.88E-07 | 4.16E-06 | ORM1 | VII | 560682 | 561350 | 1 |
| YGR039W | 48.4311911 | -0.0912365 | 0.58425669 | 0.8965754 | 0.6387209 | 0.77465312 | NA | VII | 574888 | 575199 | 1 |
| YGR040W | 46.3746771 | -0.613452 | 0.63276159 | 4.25982923 | 0.11884744 | 0.2479657 | KSS1 | VII | 575398 | 576504 | 1 |
| YGR041W | 1012.97484 | 0.74989054 | 0.48943203 | 13.386986 | 0.00123895 | 0.00784848 | BUD9 | VII | 577487 | 579130 | 1 |
| YGR042W | 139.532124 | 3.73926056 | 0.85109005 | 20.2386896 | 4.03E-05 | 0.00043148 | MTE1 | VII | 579476 | 580291 | 1 |
| YGR043C | 51.017049 | 2.92250319 | 0.85659276 | 10.4307477 | 0.0054324 | 0.02450414 | NQM1 | VII | 580435 | 581436 | -1 |
| YGR044C | 72.16624 | 1.92857762 | 0.63379485 | 13.8250147 | 0.00099526 | 0.0065558 | RME1 | VII | 582990 | 583892 | -1 |
| YGR045C | 53.4112199 | 1.81077892 | 0.81731664 | 7.4718398 | 0.02385122 | 0.07538131 | NA | VII | 583934 | 584296 | -1 |
| YGR046W | 11.1531913 | 0.50455309 | 1.12724003 | 0.52489261 | 0.76916766 | 0.85953436 | TAM41 | VII | 584895 | 586052 | 1 |
| YGR047C | 98.072144 | 3.26002393 | 0.9945089 | 10.9278082 | 0.00423698 | 0.02007035 | TFC4 | VII | 586392 | 589469 | -1 |
| YGR048W | 37.1231895 | 0.24497615 | 0.67082184 | 0.31877 | 0.85266802 | 0.91239103 | UFD1 | VII | 589826 | 590911 | 1 |
| YGR049W | 1268.70925 | 2.18655742 | 0.48484603 | 19.0291575 | 7.38E-05 | 0.00072393 | SCM4 | VII | 591314 | 591877 | 1 |
| YGR050C | 130.086023 | 0.36660609 | 0.57877582 | 0.76184801 | 0.68322981 | 0.80595449 | NA | VII | 592086 | 592442 | -1 |
| YGR052W | 39.0570244 | -1.1760417 | 0.66243987 | 5.14061931 | 0.07651185 | 0.18012252 | FMP48 | VII | 593596 | 594705 | 1 |
| YGR054W | 83.4827999 | -0.0796385 | 0.57218161 | 3.35684211 | 0.18666848 | 0.34246214 | NA | VII | 596693 | 598621 | 1 |
| YGR055W | 31.1267471 | 0.04667902 | 0.6847139 | 7.49650338 | 0.0235589 | 0.07474302 | MUP1 | VII | 599417 | 601141 | 1 |
| YGR056W | 164.24194 | 0.8583395 | 0.66716791 | 3.94980401 | 0.13877491 | 0.2772625 | RSC1 | VII | 601661 | 604447 | 1 |
| YGR057C | 165.194787 | 1.51755148 | 0.51682313 | 27.2355677 | 1.22E-06 | 2.09E-05 | LST7 | VII | 604773 | 605501 | -1 |
| YGR058W | 30.7612797 | -1.0672827 | 0.6767741 | 8.36925556 | 0.01522787 | 0.05412485 | PEF1 | VII | 606136 | 607143 | 1 |
| YGR059W | 15.9078194 | -0.2212959 | 0.95094604 | 2.23634571 | 0.3268765 | 0.49976629 | SPR3 | VII | 607563 | 609101 | 1 |
| YGR060W | 146.2102 | -0.0405417 | 0.50368058 | 1.23760358 | 0.5385894 | 0.69298502 | ERG25 | VII | 610564 | 611493 | 1 |
| YGR061C | 79.8559879 | -0.7695518 | 0.62494733 | 1.61809411 | 0.44528219 | 0.61297083 | ADE6 | VII | 611889 | 615965 | -1 |
| YGR062C | 29.5286947 | -2.0395852 | 0.79265598 | 9.91585575 | 0.00702747 | 0.03007323 | COX18 | VII | 616328 | 617278 | -1 |
| YGR063C | 95.4147023 | -0.7009589 | 0.5461535 | 1.64222066 | 0.4399429 | 0.60823052 | SPT4 | VII | 617516 | 617824 | -1 |
| YGR065C | 26.2186576 | -0.2599591 | 0.81766781 | 10.4321319 | 0.00542864 | 0.02450414 | VHT1 | VII | 618077 | 619858 | -1 |

| | | | | | | | | | | | |
|---|---|---|---|---|---|---|---|---|---|---|---|
| YGR066C | 197.19374 | -1.1668339 | 0.52990473 | 5.51890485 | 0.06332643 | 0.15622499 | NA | VII | 620921 | 621799 | -1 |
| YGR067C | 38.2686885 | 0.53330317 | 0.74134397 | 2.18759362 | 0.33494236 | 0.50833979 | NA | VII | 622372 | 624786 | -1 |
| YGR068C | 60.5496994 | 1.10258094 | 0.67017194 | 18.6474192 | 8.93E-05 | 0.00085024 | ART5 | VII | 625324 | 627084 | -1 |
| YGR070W | 34.5838589 | -1.1286937 | 0.66085646 | 2.93448194 | 0.23056073 | 0.39754218 | ROM1 | VII | 627806 | 631273 | 1 |
| YGR071C | 12.4031319 | 0.02126724 | 0.85799219 | 0.48581662 | 0.78434343 | 0.86949041 | ENV11 | VII | 631481 | 634063 | -1 |
| YGR072W | 66.8997026 | 0.41266936 | 0.62157025 | 4.17155478 | 0.12421052 | 0.25620909 | UPF3 | VII | 634304 | 635467 | 1 |
| YGR074W | 398.434296 | -0.293034 | 0.46932436 | 2.72274637 | 0.25630858 | 0.42842105 | SMD1 | VII | 635712 | 636152 | 1 |
| YGR075C | 1050.111 | -0.5102168 | 0.45573193 | 18.1109465 | 0.00011675 | 0.00106622 | PRP38 | VII | 636147 | 636875 | -1 |
| YGR076C | 7.86148688 | -1.9105432 | 1.09000301 | 3.10574833 | 0.21163881 | 0.37519557 | MRPL25 | VII | 637104 | 637577 | -1 |
| YGR077C | 61.8800081 | 0.15449164 | 0.63856558 | 4.25357892 | 0.11921944 | 0.24840182 | PEX8 | VII | 637744 | 639513 | -1 |
| YGR078C | 64.0004497 | -2.257267 | 0.60160076 | 26.2008014 | 2.04E-06 | 3.30E-05 | PAC10 | VII | 639772 | 640371 | -1 |
| YGR079W | 39.6895548 | -0.3019699 | 0.64664451 | 1.20667753 | 0.54698234 | 0.69943192 | NA | VII | 640720 | 641832 | 1 |
| YGR080W | 89.0060201 | -0.6136909 | 0.55985375 | 9.52838461 | 0.00852977 | 0.03487811 | TWF1 | VII | 642010 | 643008 | 1 |
| YGR081C | 22.4775208 | 1.51767405 | 0.99442193 | 7.86622965 | 0.01958258 | 0.06501327 | SLX9 | VII | 643148 | 643780 | -1 |
| YGR082W | 13.4625334 | -1.2705068 | 1.00954185 | 3.93599676 | 0.13973628 | 0.27822319 | TOM20 | VII | 644044 | 644595 | 1 |
| YGR083C | 115.275491 | 0.261501 | 0.55252929 | 3.91879482 | 0.14094333 | 0.2798566 | GCD2 | VII | 644860 | 646815 | -1 |
| YGR084C | 10.4695064 | -1.2608463 | 0.91826326 | 10.904648 | 0.00428633 | 0.02021351 | MRP13 | VII | 647123 | 648142 | -1 |
| YGR085C | 34.3943069 | -0.1655525 | 0.7344271 | 12.0656719 | 0.00239868 | 0.01281215 | RPL11B | VII | 648383 | 648907 | -1 |
| YGR086C | 257.48631 | 0.36699232 | 0.48406342 | 7.14561854 | 0.02807687 | 0.08453721 | PIL1 | VII | 649598 | 650617 | -1 |
| YGR087C | 28.1356343 | -0.2762299 | 0.87639235 | 6.03479692 | 0.04892834 | 0.1284785 | PDC6 | VII | 651290 | 652981 | -1 |
| YGR088W | 64.5413612 | -0.1253876 | 0.62638408 | 12.6641516 | 0.00177834 | 0.01025556 | CTT1 | VII | 654634 | 656322 | 1 |
| YGR089W | 40.3944955 | 0.04122048 | 0.64536337 | 2.42348479 | 0.29767815 | 0.47143231 | NNF2 | VII | 656960 | 659770 | 1 |
| YGR090W | 33.4237345 | -0.4795877 | 0.80162179 | 8.35546856 | 0.01533321 | 0.05444929 | UTP22 | VII | 662358 | 666071 | 1 |
| YGR091W | 23.2245783 | -0.1139759 | 0.75733363 | 6.09448158 | 0.04748978 | 0.12561252 | PRP31 | VII | 666341 | 667825 | 1 |
| YGR092W | 25.5711744 | 0.74713895 | 0.74532046 | 1.30468767 | 0.52082362 | 0.6765916 | DBF2 | VII | 668189 | 669907 | 1 |
| YGR093W | 36.067363 | -1.0026197 | 0.66312636 | 4.11169073 | 0.1279846 | 0.26147876 | DRN1 | VII | 670388 | 671911 | 1 |
| YGR094W | 135.375242 | 0.152255 | 0.53821149 | 0.241066 | 0.88644783 | 0.93215081 | VAS1 | VII | 672186 | 675500 | 1 |
| YGR095C | 9.97179505 | -0.4708467 | 0.90080153 | 2.48116905 | 0.28921512 | 0.46309611 | RRP46 | VII | 675671 | 676342 | -1 |
| YGR096W | 149.321279 | -0.806129 | 0.53342988 | 2.44785581 | 0.2940728 | 0.46854198 | TPC1 | VII | 676621 | 677565 | 1 |
| YGR097W | 66.6655705 | 0.28894859 | 0.5932843 | 0.53349272 | 0.7658673 | 0.85754625 | ASK10 | VII | 678695 | 682135 | 1 |
| YGR098C | 23.4857546 | -0.3487243 | 0.7620913 | 0.61849816 | 0.73399792 | 0.83775415 | ESP1 | VII | 682566 | 687458 | -1 |
| YGR099W | 27.104797 | -0.3503028 | 0.66479543 | 0.44344505 | 0.80113763 | 0.88018726 | TEL2 | VII | 687899 | 689965 | 1 |
| YGR100W | 49.2106248 | -0.2559545 | 0.71098376 | 0.14690461 | 0.92918046 | 0.95541731 | MDR1 | VII | 690245 | 693097 | 1 |

| | | | | | | | | | | | |
|---|---|---|---|---|---|---|---|---|---|---|---|
| YGR101W | 63.6824691 | 0.29621711 | 0.73565574 | 2.41653887 | 0.29871378 | 0.47191811 | PCP1 | VII | 693363 | 694403 | 1 |
| YGR102C | 32.7961106 | -0.0596566 | 0.67809808 | 0.1266866 | 0.93862119 | 0.95995205 | GTF1 | VII | 694584 | 695135 | -1 |
| YGR103W | 138.007939 | -1.0571732 | 0.56638639 | 32.5922975 | 8.37E-08 | 2.09E-06 | NOP7 | VII | 695417 | 697234 | 1 |
| YGR104C | 88.2012622 | -0.3227017 | 0.61839258 | 4.85961789 | 0.08805365 | 0.20056281 | SRB5 | VII | 697445 | 698368 | -1 |
| YGR105W | 13.3296522 | -0.0102106 | 0.90953549 | 6.54544557 | 0.03790308 | 0.10674069 | VMA21 | VII | 698599 | 698832 | 1 |
| YGR106C | 346.628425 | 2.15954351 | 1.60407891 | 4.17841322 | 0.12378531 | 0.25551406 | VOA1 | VII | 698989 | 699786 | -1 |
| YGR108W | 52.2261373 | -0.4274393 | 0.61989194 | 1.72404666 | 0.42230675 | 0.59293772 | CLB1 | VII | 703636 | 705051 | 1 |
| YGR109C | 184.179315 | -0.2430211 | 0.48561123 | 3.12411742 | 0.20970391 | 0.37244958 | CLB6 | VII | 705359 | 706501 | -1 |
| YGR109W-B | 291.950217 | -0.2384532 | 0.52203442 | 0.23841851 | 0.88762204 | 0.9322205 | NA | VII | 707610 | 712254 | 1 |
| YGR110W | 74.1632324 | -2.5969548 | 0.61691746 | 33.5928503 | 5.07E-08 | 1.33E-06 | CLD1 | VII | 713709 | 715046 | 1 |
| YGR111W | 38.055398 | -0.6481654 | 0.66489279 | 0.96700074 | 0.61662121 | 0.75817303 | NA | VII | 715828 | 717030 | 1 |
| YGR112W | 20.5537667 | -0.2053721 | 0.82809281 | 0.08851058 | 0.95670969 | 0.96960425 | SHY1 | VII | 717358 | 718527 | 1 |
| YGR113W | 73.5257086 | -0.9213305 | 0.59080714 | 3.34563767 | 0.18771718 | 0.34351531 | DAM1 | VII | 718893 | 719924 | 1 |
| YGR116W | 88.6879958 | 0.25946561 | 0.53928508 | 7.96076362 | 0.01867851 | 0.06284053 | SPT6 | VII | 720409 | 724764 | 1 |
| YGR117C | 48.7655781 | -0.1304096 | 0.60881644 | 0.19527579 | 0.90697727 | 0.94229291 | NA | VII | 725049 | 726479 | -1 |
| YGR118W | 120.90057 | -0.5546587 | 0.70383381 | 11.2352178 | 0.00363332 | 0.01781991 | RPS23B | VII | 726974 | 727730 | 1 |
| YGR119C | 48.6749629 | 0.26429053 | 0.66655745 | 0.27233374 | 0.87269699 | 0.92527295 | NUP57 | VII | 728046 | 729671 | -1 |
| YGR120C | 138.686245 | -0.0548112 | 0.54337593 | 2.12024834 | 0.34641279 | 0.51988856 | COG2 | VII | 730033 | 730821 | -1 |
| YGR121C | 1687.63223 | 0.67199778 | 0.56099004 | 2.96803705 | 0.22672476 | 0.39279962 | MEP1 | VII | 731449 | 732927 | -1 |
| YGR121W-A | 26.0304755 | -0.7570462 | 0.73821699 | 1.57585017 | 0.45478746 | 0.62066785 | NA | VII | 733412 | 733627 | 1 |
| YGR122W | 57.6364419 | -1.4897097 | 0.60447148 | 9.79066531 | 0.00748142 | 0.03161856 | NA | VII | 733935 | 735143 | 1 |
| YGR123C | 15.3340432 | -0.0347394 | 0.81351514 | 0.42009574 | 0.81054544 | 0.88665371 | PPT1 | VII | 736662 | 738203 | -1 |
| YGR124W | 70.08196 | 0.83883249 | 0.60844213 | 8.02269414 | 0.01810898 | 0.06142415 | ASN2 | VII | 739944 | 741662 | 1 |
| YGR125W | 71.7735703 | 0.51407965 | 0.60005759 | 6.56275786 | 0.03757641 | 0.10618223 | NA | VII | 742325 | 745435 | 1 |
| YGR126W | 40.2294814 | -0.2623121 | 0.65266861 | 10.4301303 | 0.00543408 | 0.02450414 | NA | VII | 745835 | 746527 | 1 |
| YGR127W | 27.6826594 | -0.3413894 | 0.7130662 | 2.51534257 | 0.28431535 | 0.45790983 | NA | VII | 746798 | 747736 | 1 |
| YGR128C | 31.1274652 | 1.05341601 | 0.92100964 | 2.52822455 | 0.28248996 | 0.45662112 | UTP8 | VII | 747950 | 750091 | -1 |
| YGR129W | 16.33605 | 0.43086635 | 0.88648227 | 0.31677161 | 0.85352043 | 0.91313438 | SYF2 | VII | 750400 | 751047 | 1 |
| YGR130C | 705.621384 | -1.4225552 | 0.4948477 | 11.2855965 | 0.00354294 | 0.01744356 | NA | VII | 751394 | 753844 | -1 |
| YGR131W | 7.60574127 | -0.7098021 | 1.08304406 | 0.63091881 | 0.72945368 | 0.83678999 | FHN1 | VII | 754726 | 755250 | 1 |
| YGR132C | 18.9169162 | -0.1418901 | 0.88248413 | 0.76049025 | 0.6836938 | 0.80606538 | PHB1 | VII | 755589 | 756452 | -1 |
| YGR133W | 475.228288 | 0.14759054 | 0.5450062 | 4.99481865 | 0.08229793 | 0.19005499 | PEX4 | VII | 756896 | 757447 | 1 |
| YGR134W | 80.7354192 | 0.07812995 | 0.64313407 | 8.37051387 | 0.0152183 | 0.05412404 | CAF130 | VII | 757770 | 761138 | 1 |

| | | | | | | | | | | | |
|---|---|---|---|---|---|---|---|---|---|---|---|
| YGR135W | 33.8072402 | 0.54554221 | 0.7492763 | 0.59073603 | 0.74425764 | 0.84379317 | PRE9 | VII | 761392 | 762168 | 1 |
| YGR136W | 48.4026226 | 1.69129337 | 0.69945954 | 6.39578997 | 0.0408481 | 0.11305473 | LSB1 | VII | 762428 | 763153 | 1 |
| YGR138C | 105.427501 | 0.01765498 | 0.52794178 | 2.50858948 | 0.28527697 | 0.45887782 | TPO2 | VII | 763762 | 765606 | -1 |
| YGR140W | 60.1304181 | -0.3398552 | 0.59370316 | 7.78591445 | 0.02038497 | 0.06703554 | CBF2 | VII | 767429 | 770299 | 1 |
| YGR141W | 604.27726 | -0.2218693 | 0.57360313 | 21.528309 | 2.11E-05 | 0.00024883 | VPS62 | VII | 770569 | 771972 | 1 |
| YGR142W | 118.993445 | 1.09152787 | 0.53906699 | 12.1728427 | 0.00227353 | 0.01230256 | BTN2 | VII | 772454 | 773686 | 1 |
| YGR143W | 88.5915044 | -1.5805958 | 0.54382487 | 9.39042732 | 0.00913891 | 0.03682277 | SKN1 | VII | 775193 | 777508 | 1 |
| YGR144W | 52.7161306 | 1.8666891 | 0.66809966 | 11.2168034 | 0.00366693 | 0.01794717 | THI4 | VII | 780399 | 781379 | 1 |
| YGR145W | 55.3857875 | -0.423775 | 0.64941929 | 0.84410948 | 0.65569814 | 0.78495521 | ENP2 | VII | 781767 | 783890 | 1 |
| YGR146C | 146.017916 | -1.7926513 | 0.52993547 | 15.8448366 | 0.00036252 | 0.0028442 | ECL1 | VII | 784223 | 784858 | -1 |
| YGR146C-A | 62.8746921 | -0.2942543 | 0.6643449 | 4.66063954 | 0.09726464 | 0.21492841 | NA | VII | 785276 | 785437 | -1 |
| YGR147C | 46.9472286 | 0.85296565 | 0.61767911 | 2.78725981 | 0.24817282 | 0.41904947 | NAT2 | VII | 786054 | 786920 | -1 |
| YGR148C | 115.845923 | -0.6741046 | 0.6461239 | 19.4452243 | 5.99E-05 | 0.0006041 | RPL24B | VII | 787312 | 787779 | -1 |
| YGR149W | 26.8801178 | 1.18352068 | 0.78197768 | 4.02653752 | 0.13355141 | 0.26953536 | GPC1 | VII | 789031 | 790329 | 1 |
| YGR150C | 461.861971 | 4.04762209 | 0.77798125 | 22.1901252 | 1.52E-05 | 0.0001887 | CCM1 | VII | 790459 | 793053 | -1 |
| YGR152C | 15.4282094 | -0.0288013 | 0.86967517 | 0.52663859 | 0.76849648 | 0.85932689 | RSR1 | VII | 794674 | 795492 | -1 |
| YGR153W | 443.091929 | 0.49768895 | 0.64996565 | 2.16441993 | 0.33884586 | 0.5121163 | NA | VII | 796092 | 796745 | 1 |
| YGR154C | 176.016378 | 0.87370743 | 0.50969199 | 7.50626736 | 0.02344416 | 0.07454061 | GTO1 | VII | 796798 | 797868 | -1 |
| YGR155W | 43.4153566 | -0.7516773 | 0.71172209 | 8.75240824 | 0.01257299 | 0.0468152 | CYS4 | VII | 798543 | 800066 | 1 |
| YGR156W | 348.763949 | -2.127852 | 0.49387179 | 44.5873923 | 2.08E-10 | 8.99E-09 | PTI1 | VII | 800546 | 801823 | 1 |
| YGR157W | 107.386806 | -0.4880291 | 0.57586069 | 10.6492989 | 0.00487006 | 0.02241465 | CHO2 | VII | 802440 | 805049 | 1 |
| YGR158C | 63.2640227 | -1.7379888 | 0.6069692 | 26.2067035 | 2.04E-06 | 3.30E-05 | MTR3 | VII | 805269 | 806021 | -1 |
| YGR159C | 105.827163 | -1.6830457 | 0.62809495 | 35.6141783 | 1.85E-08 | 5.17E-07 | NSR1 | VII | 806412 | 807656 | -1 |
| YGR161C | 218.737636 | -1.6377827 | 0.49602483 | 10.4898829 | 0.00527413 | 0.0239164 | RTS3 | VII | 808629 | 809420 | -1 |
| YGR161W-C | 7.76058394 | -0.4623459 | 1.017139 | 0.26381337 | 0.87642278 | 0.9271858 | NA | VII | 810227 | 810505 | 1 |
| YGR162W | 169.953386 | -0.6408071 | 0.53828677 | 17.5573328 | 0.00015398 | 0.00136743 | TIF4631 | VII | 824059 | 826917 | 1 |
| YGR163W | 429.819583 | 2.22132621 | 0.51789173 | 50.8334345 | 9.16E-12 | 5.46E-10 | GTR2 | VII | 827552 | 828577 | 1 |
| YGR164W | 231.776972 | -0.683567 | 0.5377553 | 35.1625593 | 2.31E-08 | 6.38E-07 | NA | VII | 828625 | 828960 | 1 |
| YGR165W | 23.8928623 | 0.56789906 | 0.78645712 | 0.65954003 | 0.7190891 | 0.82938751 | MRPS35 | VII | 829116 | 830153 | 1 |
| YGR166W | 184.91055 | -0.3123272 | 0.52505828 | 2.55661249 | 0.27850863 | 0.45301284 | TRS65 | VII | 830515 | 832197 | 1 |
| YGR167W | 73.2702201 | 0.86015553 | 0.64423805 | 1.79713679 | 0.40715212 | 0.57850572 | CLC1 | VII | 832456 | 833157 | 1 |
| YGR168C | 168.906165 | -0.6166757 | 0.5161109 | 9.60207823 | 0.0082212 | 0.03390367 | NA | VII | 833352 | 834482 | -1 |
| YGR169C | 11.6741497 | -0.5702934 | 0.97253255 | 1.3201147 | 0.51682169 | 0.67350835 | PUS6 | VII | 834689 | 835903 | -1 |

| | | | | | | | | | | | |
|---|---|---|---|---|---|---|---|---|---|---|---|
| YGR169C-A | 76.8577513 | 0.29164272 | 0.61236051 | 7.62628884 | 0.02207865 | 0.07131784 | LSO2 | VII | 836387 | 836665 | -1 |
| YGR170W | 494.170442 | -1.3259644 | 0.49394224 | 12.2370462 | 0.00220171 | 0.01203765 | PSD2 | VII | 837142 | 840558 | 1 |
| YGR171C | 119.285 | 1.35394235 | 0.76347604 | 4.47049971 | 0.1069654 | 0.22980693 | MSM1 | VII | 840824 | 842551 | -1 |
| YGR172C | 24.4945857 | -0.4433166 | 0.70596823 | 0.96537847 | 0.61712157 | 0.75846612 | YIP1 | VII | 842845 | 843591 | -1 |
| YGR173W | 37.654227 | -0.7959999 | 0.69788429 | 9.2062361 | 0.01002054 | 0.03951413 | RBG2 | VII | 843854 | 844960 | 1 |
| YGR174C | 1464.32768 | 3.61597046 | 0.59375776 | 49.0390478 | 2.25E-11 | 1.23E-09 | NA | VII | 845893 | 846405 | -1 |
| YGR174W-A | 43.214005 | 3.83925845 | 1.86744253 | 5.67922466 | 0.05844832 | 0.14688185 | NA | VII | 846655 | 846741 | 1 |
| YGR175C | 52.8898445 | 0.41448757 | 0.86209514 | 1.25566161 | 0.53374835 | 0.68828573 | ERG1 | VII | 846933 | 848423 | -1 |
| YGR177C | 127.964679 | 0.1620528 | 0.5331789 | 2.85359498 | 0.24007654 | 0.40823588 | ATF2 | VII | 848829 | 850436 | -1 |
| YGR178C | 258.261136 | -0.6619432 | 0.48197702 | 1.93114047 | 0.38076601 | 0.55448572 | PBP1 | VII | 851047 | 853215 | -1 |
| YGR180C | 28.4182515 | -0.2128609 | 0.67047685 | 2.71433615 | 0.25738865 | 0.42958807 | RNR4 | VII | 855264 | 856301 | -1 |
| YGR181W | 39.5652367 | -0.048399 | 0.76618282 | 2.98279326 | 0.22505811 | 0.3904964 | TIM13 | VII | 858287 | 858604 | 1 |
| YGR183C | 41.6803678 | -0.5107211 | 0.66207957 | 3.94035788 | 0.1394319 | 0.27799956 | QCR9 | VII | 859063 | 859476 | -1 |
| YGR184C | 53.7244738 | 1.4197098 | 0.6145856 | 5.5499553 | 0.06235087 | 0.15421253 | UBR1 | VII | 859901 | 865753 | -1 |
| YGR185C | 75.9300681 | 0.00858525 | 0.56986553 | 8.26704451 | 0.01602633 | 0.05640572 | TYS1 | VII | 866336 | 867520 | -1 |
| YGR186W | 49.484178 | 0.41039122 | 0.69661565 | 0.88991661 | 0.640851 | 0.77561669 | TFG1 | VII | 867774 | 869981 | 1 |
| YGR187C | 64.4279703 | -0.2830872 | 0.59755151 | 6.32106129 | 0.04240323 | 0.11597295 | HGH1 | VII | 870232 | 871416 | -1 |
| YGR188C | 40.9285061 | 0.05791385 | 0.67168617 | 12.6056569 | 0.00183112 | 0.01050761 | BUB1 | VII | 872044 | 875109 | -1 |
| YGR189C | 235.206657 | 3.21433035 | 0.58364089 | 27.1986945 | 1.24E-06 | 2.13E-05 | CRH1 | VII | 876669 | 878192 | -1 |
| YGR191W | 63.6451159 | -0.1012999 | 0.56857863 | 3.45152953 | 0.17803684 | 0.33145766 | HIP1 | VII | 880420 | 882231 | 1 |
| YGR192C | 842.365229 | -0.2965515 | 0.72917884 | 4.06058018 | 0.13129743 | 0.26636724 | TDH3 | VII | 882812 | 883810 | -1 |
| YGR193C | 62.439846 | 0.32963285 | 0.58423886 | 0.66522568 | 0.71704775 | 0.82851856 | PDX1 | VII | 884509 | 885741 | -1 |
| YGR194C | 69.7169034 | -0.7536203 | 0.5538014 | 3.40724241 | 0.18202319 | 0.33713158 | XKS1 | VII | 886073 | 887875 | -1 |
| YGR195W | 111.640546 | 0.93899596 | 0.79922789 | 5.78203506 | 0.05551969 | 0.14148724 | SKI6 | VII | 888882 | 889622 | 1 |
| YGR196C | 132.805055 | -0.7926557 | 0.53343191 | 5.42902475 | 0.06623724 | 0.16143398 | FYV8 | VII | 889733 | 892186 | -1 |
| YGR197C | 38.6617457 | -0.1787457 | 0.72656149 | 0.13949929 | 0.93262728 | 0.95731799 | SNG1 | VII | 892497 | 894140 | -1 |
| YGR198W | 41.3439075 | -0.6216504 | 0.70482838 | 1.10760826 | 0.57475919 | 0.72422968 | YPP1 | VII | 894693 | 897146 | 1 |
| YGR199W | 257.938692 | -1.0635419 | 0.5479185 | 6.91364023 | 0.03152986 | 0.09285753 | PMT6 | VII | 897502 | 899781 | 1 |
| YGR200C | 461.624139 | 0.71257419 | 0.64062373 | 2.74200019 | 0.25385296 | 0.42590803 | ELP2 | VII | 899904 | 902270 | -1 |
| YGR201C | 134.039587 | -1.5129635 | 0.56803723 | 14.0082707 | 0.00090812 | 0.00608566 | NA | VII | 902520 | 903197 | -1 |
| YGR202C | 196.54431 | 1.2438858 | 0.54587075 | 38.736485 | 3.88E-09 | 1.27E-07 | PCT1 | VII | 903474 | 904748 | -1 |
| YGR203W | 198.24288 | -0.625206 | 0.52899158 | 1.44519388 | 0.48548983 | 0.64773825 | YCH1 | VII | 905237 | 905683 | 1 |
| YGR204C-A | 235.032762 | -0.2116456 | 0.63628159 | 3.58534796 | 0.16651432 | 0.31564276 | NA | VII | 909061 | 909174 | -1 |

| | | | | | | | | | | | |
|---|---|---|---|---|---|---|---|---|---|---|---|
| YGR204W | 97.1955208 | -0.0909228 | 0.61159727 | 0.02888551 | 0.98566104 | 0.98832468 | ADE3 | VII | 905934 | 908774 | 1 |
| YGR205W | 95.4220637 | -0.7858133 | 0.58363084 | 5.48752119 | 0.06432798 | 0.15782161 | TDA10 | VII | 909213 | 910085 | 1 |
| YGR206W | 209.35029 | 1.9692572 | 0.94790865 | 7.69390051 | 0.02134473 | 0.06946937 | MVB12 | VII | 910432 | 910737 | 1 |
| YGR207C | 18.4804746 | 1.36225951 | 0.81282671 | 6.5936193 | 0.03700103 | 0.10476085 | CIR1 | VII | 910843 | 911628 | -1 |
| YGR208W | 19.8237596 | 0.33917391 | 0.76133373 | 4.52631097 | 0.10402173 | 0.22540636 | SER2 | VII | 911883 | 912812 | 1 |
| YGR209C | 34.2640386 | 0.50993887 | 0.8264845 | 3.55821065 | 0.16878909 | 0.3187519 | TRX2 | VII | 912913 | 913227 | -1 |
| YGR210C | 59.5265427 | -0.4516095 | 0.64295242 | 0.57633476 | 0.74963611 | 0.8479727 | NA | VII | 913503 | 914738 | -1 |
| YGR211W | 44.1599467 | -0.1393233 | 0.68082716 | 1.84772661 | 0.39698241 | 0.56785755 | ZPR1 | VII | 915241 | 916701 | 1 |
| YGR212W | 4335.86846 | 4.83100895 | 0.45223058 | 85.2404735 | 3.09E-19 | 7.16E-17 | SLI1 | VII | 917040 | 918446 | 1 |
| YGR213C | 418.861086 | -0.9014455 | 0.49641987 | 8.58854765 | 0.01364648 | 0.04972505 | RTA1 | VII | 918512 | 919465 | -1 |
| YGR214W | 163.065089 | -0.8025734 | 0.73141497 | 11.7007423 | 0.00287883 | 0.01475082 | RPS0A | VII | 920575 | 921788 | 1 |
| YGR215W | 92.3887364 | 2.1803467 | 0.98121088 | 9.74026212 | 0.00767236 | 0.03212072 | RSM27 | VII | 922175 | 922507 | 1 |
| YGR216C | 48.1609999 | -0.5530705 | 0.62914915 | 13.4742163 | 0.00118607 | 0.00756665 | GPI1 | VII | 922633 | 924462 | -1 |
| YGR217W | 44.9150882 | -0.3176927 | 0.60332354 | 0.28315875 | 0.86798628 | 0.9224744 | CCH1 | VII | 924696 | 930815 | 1 |
| YGR218W | 27.8598754 | 0.17894183 | 0.88450104 | 0.44726714 | 0.79960808 | 0.87884032 | CRM1 | VII | 932541 | 935795 | 1 |
| YGR220C | 12.989591 | 1.35491641 | 0.95272786 | 5.26301338 | 0.07196994 | 0.17226374 | MRPL9 | VII | 936074 | 936883 | -1 |
| YGR221C | 1295.30671 | 1.62881177 | 0.46832155 | 18.3410102 | 0.00010406 | 0.00096714 | TOS2 | VII | 937124 | 938992 | -1 |
| YGR222W | 48.5327019 | 1.81713302 | 0.89170223 | 4.31385617 | 0.11567993 | 0.24294045 | PET54 | VII | 939923 | 940804 | 1 |
| YGR223C | 109.292284 | 2.16895257 | 0.8445555 | 7.50521731 | 0.02345648 | 0.07454061 | HSV2 | VII | 940869 | 942215 | -1 |
| YGR224W | 1692.29434 | 0.71930533 | 0.5028689 | 7.27102708 | 0.02637039 | 0.08074276 | AZR1 | VII | 942806 | 944647 | 1 |
| YGR225W | 5.67939493 | -0.2142177 | 1.18011361 | 0.24394526 | 0.8851726 | 0.9315066 | AMA1 | VII | 945145 | 947019 | 1 |
| YGR227C-A | 319.653433 | 0.51050479 | 0.71915237 | 8.19299688 | 0.01663081 | 0.05772162 | NA | NA | NA | NA | NA |
| YGR227W | 19.2107175 | -0.3041445 | 0.80380173 | 0.24450795 | 0.88492359 | 0.9315066 | DIE2 | VII | 947420 | 948997 | 1 |
| YGR229C | 129.926616 | 0.16055463 | 0.59469763 | 3.15109206 | 0.20689455 | 0.36890584 | SMI1 | VII | 949382 | 950899 | -1 |
| YGR230W | 8.65971981 | 0.75732322 | 1.02849359 | 0.62486269 | 0.73166586 | 0.83702784 | BNS1 | VII | 951894 | 952307 | 1 |
| YGR231C | 32.684256 | -0.1540819 | 0.74425151 | 0.09112579 | 0.95545951 | 0.96850675 | PHB2 | VII | 952548 | 953480 | -1 |
| YGR232W | 128.454096 | 2.10443618 | 0.86285294 | 7.43968003 | 0.02423785 | 0.07639473 | NAS6 | VII | 953960 | 954646 | 1 |
| YGR233C | 176.424631 | 1.6299356 | 0.8222689 | 5.09092484 | 0.07843677 | 0.18386596 | PHO81 | VII | 954674 | 958210 | -1 |
| YGR234W | 107.342206 | -0.2285796 | 0.5663901 | 13.182497 | 0.00137233 | 0.00850724 | YHB1 | VII | 959904 | 961103 | 1 |
| YGR235C | 1621.45257 | 0.5865052 | 0.74014667 | 24.3122809 | 5.26E-06 | 7.46E-05 | MIC26 | VII | 961360 | 962061 | -1 |
| YGR237C | 174.551398 | 0.46726003 | 0.50421953 | 1.44367108 | 0.48585962 | 0.64773825 | NA | VII | 963298 | 965655 | -1 |
| YGR238C | 93.8336373 | -0.0315464 | 0.55874356 | 15.1310767 | 0.000518 | 0.00378167 | KEL2 | VII | 966039 | 968687 | -1 |
| YGR239C | 326.972644 | -1.463167 | 0.64799874 | 5.24021826 | 0.07279492 | 0.17359249 | PEX21 | VII | 969187 | 970053 | -1 |

| | | | | | | | | | | | |
|---|---|---|---|---|---|---|---|---|---|---|---|
| YGR240C | 109.310946 | 0.03874469 | 0.71413497 | 1.35878789 | 0.50692412 | 0.66464916 | PFK1 | VII | 970771 | 973734 | -1 |
| YGR240C-A | 804.206074 | -0.1986465 | 0.5410186 | 0.41157427 | 0.81400633 | 0.88825794 | NA | VII | 974577 | 974777 | -1 |
| YGR241C | 32.5535381 | 1.06569895 | 0.75671603 | 3.18882778 | 0.20302749 | 0.3631539 | YAP1802 | VII | 974875 | 976581 | -1 |
| YGR244C | 86.7203419 | -1.2053984 | 0.57361781 | 10.6000297 | 0.00499152 | 0.02286464 | LSC2 | VII | 978036 | 979319 | -1 |
| YGR245C | 68.6136283 | -1.6723924 | 0.56256443 | 18.6177839 | 9.06E-05 | 0.0008601 | SDA1 | VII | 979765 | 982068 | -1 |
| YGR246C | 13.1341319 | -0.1745741 | 0.93738999 | 0.8753967 | 0.64552047 | 0.77784881 | BRF1 | VII | 982482 | 984272 | -1 |
| YGR247W | 10.6784688 | -0.24768 | 0.96245937 | 0.88786883 | 0.64150749 | 0.77591986 | CPD1 | VII | 984966 | 985685 | 1 |
| YGR248W | 80.3010328 | 0.25596763 | 0.58435419 | 12.1812365 | 0.00226401 | 0.01228548 | SOL4 | VII | 985972 | 986739 | 1 |
| YGR249W | 83.3010538 | -1.1521015 | 0.58207284 | 7.28080831 | 0.02624174 | 0.08047651 | MGA1 | VII | 988049 | 989419 | 1 |
| YGR250C | 60.7895579 | 0.95539184 | 0.79143793 | 2.54629998 | 0.2799484 | 0.45403395 | RIE1 | VII | 991176 | 993521 | -1 |
| YGR251W | 20.5919623 | -0.4067766 | 0.78159816 | 1.53565129 | 0.46402092 | 0.62714312 | NOP19 | VII | 995639 | 996229 | 1 |
| YGR252W | 205.520044 | -0.0299349 | 0.54127386 | 2.70616602 | 0.25844225 | 0.43011803 | GCN5 | VII | 996869 | 998188 | 1 |
| YGR253C | 226.115219 | 0.74245833 | 0.67811931 | 5.09700013 | 0.07819887 | 0.18338253 | PUP2 | VII | 998358 | 999140 | -1 |
| YGR254W | 147.977753 | -0.0955679 | 0.5896986 | 1.95224343 | 0.37676949 | 0.55032677 | ENO1 | VII | 1000927 | 1E+06 | 1 |
| YGR255C | 17.171899 | -0.6909938 | 0.92022587 | 0.71211645 | 0.70043184 | 0.81801018 | COQ6 | VII | 1002523 | 1E+06 | -1 |
| YGR256W | 71.5619089 | -0.0954902 | 0.56842201 | 0.41732741 | 0.81166815 | 0.88731158 | GND2 | VII | 1004624 | 1E+06 | 1 |
| YGR257C | 57.6640789 | 1.81519427 | 0.80677492 | 4.81148772 | 0.09019838 | 0.20366605 | MTM1 | VII | 1006205 | 1E+06 | -1 |
| YGR258C | 53.0120634 | 0.56660592 | 0.62943702 | 4.49101939 | 0.10587356 | 0.22838155 | RAD2 | VII | 1007671 | 1E+06 | -1 |
| YGR260W | 215.488257 | 0.52316285 | 0.5135437 | 27.053088 | 1.34E-06 | 2.27E-05 | TNA1 | VII | 1012485 | 1E+06 | 1 |
| YGR261C | 33.9983192 | -0.2912603 | 0.76371946 | 0.49236824 | 0.78177827 | 0.86750947 | APL6 | VII | 1014321 | 1E+06 | -1 |
| YGR262C | 152.903638 | 0.34327387 | 0.67314403 | 60.6418372 | 6.79E-14 | 5.78E-12 | BUD32 | VII | 1016974 | 1E+06 | -1 |
| YGR263C | 89.1015434 | -0.571735 | 0.54733094 | 1.11381669 | 0.57297778 | 0.72277589 | SAY1 | VII | 1017967 | 1E+06 | -1 |
| YGR264C | 32.8954497 | -1.000309 | 0.79178788 | 4.80145616 | 0.09065193 | 0.20427392 | MES1 | VII | 1019598 | 1E+06 | -1 |
| YGR266W | 358.493564 | -0.4183003 | 0.50472529 | 11.8572797 | 0.0026621 | 0.01389861 | NA | VII | 1022656 | 1E+06 | 1 |
| YGR267C | 156.231856 | 2.83856562 | 0.92292387 | 10.5144304 | 0.00520979 | 0.02376215 | FOL2 | VII | 1025004 | 1E+06 | -1 |
| YGR268C | 52.5221446 | -0.670407 | 0.63553333 | 1.43737684 | 0.48739109 | 0.64903366 | HUA1 | VII | 1026057 | 1E+06 | -1 |
| YGR270W | 114.552403 | 0.94945799 | 0.57842038 | 2.89857517 | 0.23473746 | 0.40225741 | YTA7 | VII | 1027370 | 1E+06 | 1 |
| YGR271C-A | 121.595902 | 2.74511718 | 1.03930395 | 8.5264338 | 0.01407695 | 0.05103664 | EFG1 | VII | 1037800 | 1E+06 | -1 |
| YGR271W | 30.7590597 | -0.2241122 | 0.64317618 | 0.6394027 | 0.72636593 | 0.83478737 | SLH1 | VII | 1031791 | 1E+06 | 1 |
| YGR274C | 22.298224 | 0.06846247 | 0.78739078 | 2.52308617 | 0.28321666 | 0.45678444 | TAF1 | VII | 1039895 | 1E+06 | -1 |
| YGR275W | 59.4047774 | 2.78310637 | 0.90562925 | 11.6542639 | 0.00294652 | 0.01503761 | RTT102 | VII | 1043276 | 1E+06 | 1 |
| YGR276C | 234.277947 | 1.41689372 | 0.770092 | 8.06330574 | 0.01774498 | 0.06033083 | RNH70 | VII | 1043819 | 1E+06 | -1 |
| YGR277C | 23.1025815 | -0.0843437 | 0.73590809 | 0.07095063 | 0.96514656 | 0.97423267 | CAB4 | VII | 1045644 | 1E+06 | -1 |

| | | | | | | | | | | | |
|---|---|---|---|---|---|---|---|---|---|---|---|
| YGR278W | 44.9578612 | -0.0120991 | 0.67399062 | 1.96861682 | 0.37369758 | 0.54680541 | CWC22 | VII | 1046731 | 1E+06 | 1 |
| YGR279C | 58.2729932 | -0.2955607 | 0.58170087 | 0.71729601 | 0.69862022 | 0.81736153 | SCW4 | VII | 1048798 | 1E+06 | -1 |
| YGR280C | 186.52823 | -0.5801979 | 0.67275245 | 18.5528096 | 9.36E-05 | 0.00088415 | PXR1 | VII | 1050910 | 1E+06 | -1 |
| YGR281W | 83.255006 | 0.84232874 | 0.64012575 | 5.97226662 | 0.05048226 | 0.13160391 | YOR1 | VII | 1052824 | 1E+06 | 1 |
| YGR282C | 71.2078076 | 0.87710524 | 0.69806219 | 2.17235765 | 0.33750369 | 0.51075441 | BGL2 | VII | 1057783 | 1E+06 | -1 |
| YGR283C | 2033.14542 | 2.52744617 | 0.52793559 | 32.0801803 | 1.08E-07 | 2.58E-06 | NA | VII | 1059015 | 1E+06 | -1 |
| YGR284C | 60.2713665 | -0.7694854 | 0.60779924 | 8.66982276 | 0.01310304 | 0.04820096 | ERV29 | VII | 1060658 | 1E+06 | -1 |
| YGR285C | 135.562276 | 0.0305172 | 0.54078644 | 7.03063095 | 0.02973842 | 0.08839089 | ZUO1 | VII | 1061852 | 1E+06 | -1 |
| YGR286C | 9.19733501 | -0.3060338 | 0.97315637 | 0.15090866 | 0.92732208 | 0.95452353 | BIO2 | VII | 1063813 | 1E+06 | -1 |
| YGR288W | 59.4154946 | 0.28810165 | 0.63391448 | 23.3112355 | 8.67E-06 | 0.0001162 | MAL13 | VII | 1070293 | 1E+06 | 1 |
| YGR289C | 45.2200273 | 1.34338629 | 0.83049663 | 11.9246481 | 0.00257392 | 0.0135482 | MAL11 | VII | 1073963 | 1E+06 | -1 |
| YGR292W | 41.5154763 | -1.0332938 | 0.62210661 | 5.86858222 | 0.0531684 | 0.13694173 | MAL12 | VII | 1076599 | 1E+06 | 1 |
| YGR295C | 109.590786 | -1.7545405 | 0.52547449 | 18.7169411 | 8.62E-05 | 0.00082519 | COS6 | VII | 1081584 | 1E+06 | -1 |
| YGR296W | 11.8970325 | -1.0714083 | 0.97835656 | 2.1696518 | 0.33796062 | 0.51117868 | YRF1-7 | VII | 1084864 | 1E+06 | 1 |
| YHL001W | 25.5104093 | -0.4472894 | 0.7725923 | 3.54060489 | 0.17028148 | 0.32052333 | RPL14B | VIII | 104277 | 105091 | 1 |
| YHL002W | 135.650715 | -0.8426296 | 0.54248625 | 4.11195872 | 0.12796745 | 0.26147876 | HSE1 | VIII | 102612 | 103970 | 1 |
| YHL003C | 13.4117336 | 0.25424296 | 0.96373744 | 0.11822599 | 0.94260025 | 0.96176085 | LAG1 | VIII | 100648 | 101883 | -1 |
| YHL004W | 135.473134 | 2.19674701 | 0.86926235 | 11.169576 | 0.00375455 | 0.01825258 | MRP4 | VIII | 99219 | 100403 | 1 |
| YHL005C | 349.246839 | -0.79539 | 0.52591784 | 3.70775326 | 0.1566288 | 0.30229358 | NA | VIII | 98828 | 99220 | -1 |
| YHL006C | 66.2879648 | -1.7276309 | 0.68250528 | 21.0393022 | 2.70E-05 | 0.00030896 | SHU1 | VIII | 98343 | 98795 | -1 |
| YHL007C | 105.801095 | -0.2552421 | 0.52476039 | 1.68348061 | 0.43095987 | 0.5996774 | STE20 | VIII | 95118 | 97937 | -1 |
| YHL008C | 23.6077361 | 0.74844634 | 0.76538556 | 5.16764693 | 0.07548484 | 0.17824519 | NA | VIII | 92627 | 94510 | -1 |
| YHL009C | 32.4671778 | 0.93372544 | 0.7267411 | 10.5200798 | 0.0051951 | 0.02372209 | YAP3 | VIII | 84068 | 85060 | -1 |
| YHL010C | 90.8417042 | 1.40329965 | 0.63259511 | 25.6126808 | 2.74E-06 | 4.24E-05 | ETP1 | VIII | 81964 | 83721 | -1 |
| YHL011C | 22.6945235 | -0.3008801 | 0.81478779 | 1.20353933 | 0.54784128 | 0.70011119 | PRS3 | VIII | 80654 | 81616 | -1 |
| YHL012W | 1212.05141 | -0.9354687 | 0.47925541 | 29.9234555 | 3.18E-07 | 6.55E-06 | NA | VIII | 78936 | 80417 | 1 |
| YHL013C | 12.0078999 | -1.1260826 | 0.93268332 | 2.16368882 | 0.33896975 | 0.51216985 | OTU2 | VIII | 77430 | 78353 | -1 |
| YHL014C | 90.9708135 | 0.41463511 | 0.73933378 | 5.2757247 | 0.07151398 | 0.1715269 | YLF2 | VIII | 76097 | 77314 | -1 |
| YHL015W | 220.969664 | -0.8447542 | 0.68421683 | 18.2458983 | 0.00010913 | 0.00100778 | RPS20 | VIII | 75412 | 75777 | 1 |
| YHL015W-A | 454.204598 | 0.65393852 | 0.56889606 | 15.4717491 | 0.00043687 | 0.00329789 | NA | VIII | 74699 | 74782 | 1 |
| YHL016C | 128.610351 | -0.2397035 | 0.71234535 | 2.29334573 | 0.31769202 | 0.490255 | DUR3 | VIII | 72037 | 74244 | -1 |
| YHL017W | 18.406319 | 1.17993034 | 0.87868853 | 1.9109279 | 0.38463365 | 0.55747464 | NA | VIII | 70276 | 71874 | 1 |
| YHL019C | 84.9176728 | 0.40773403 | 0.5846745 | 0.50613866 | 0.77641405 | 0.8639961 | APM2 | VIII | 67731 | 69548 | -1 |

| | | | | | | | | | | | |
|---|---|---|---|---|---|---|---|---|---|---|---|
| YHL020C | 44.0676708 | 0.82291686 | 0.6406097 | 4.45912372 | 0.10757555 | 0.23077356 | OPI1 | VIII | 66242 | 67456 | -1 |
| YHL021C | 148.495758 | 0.30054169 | 0.52031264 | 0.40721634 | 0.81578196 | 0.88902268 | AIM17 | VIII | 64462 | 65859 | -1 |
| YHL022C | 111.502574 | -0.9189535 | 0.56548541 | 3.66490427 | 0.16002069 | 0.30728192 | SPO11 | VIII | 62961 | 64157 | -1 |
| YHL023C | 507.979672 | -0.3937896 | 0.50123886 | 4.81110709 | 0.09021554 | 0.20366605 | NPR3 | VIII | 59123 | 62563 | -1 |
| YHL024W | 2618.10836 | 0.3470225 | 0.4937902 | 5.61624845 | 0.06031803 | 0.1506 | RIM4 | VIII | 56649 | 58790 | 1 |
| YHL025W | 51.2035016 | -0.4853294 | 0.58858413 | 0.69937661 | 0.70490777 | 0.82055541 | SNF6 | VIII | 54851 | 55849 | 1 |
| YHL026C | 496.13448 | 1.10288741 | 0.50643618 | 14.7264565 | 0.00063415 | 0.00451071 | NA | VIII | 53219 | 54166 | -1 |
| YHL027W | 59.9396542 | 0.29619375 | 0.63215705 | 5.80929761 | 0.05476802 | 0.14006486 | RIM101 | VIII | 51111 | 52988 | 1 |
| YHL028W | 45.0019699 | -0.0422683 | 0.59786613 | 0.48033087 | 0.78649774 | 0.87021248 | WSC4 | VIII | 48763 | 50580 | 1 |
| YHL029C | 63.5717298 | -0.4857552 | 0.61039821 | 10.6325174 | 0.00491109 | 0.02256764 | OCA5 | VIII | 45929 | 47968 | -1 |
| YHL030W | 20.3851726 | -0.9761225 | 0.78572843 | 2.63177818 | 0.26823573 | 0.44109199 | ECM29 | VIII | 40084 | 45690 | 1 |
| YHL031C | 11.4256395 | -0.8476955 | 0.99382306 | 0.89127111 | 0.64041713 | 0.77561669 | GOS1 | VIII | 38815 | 39486 | -1 |
| YHL032C | 30.9581941 | 0.15001103 | 0.67825686 | 1.85925072 | 0.39470155 | 0.56638907 | GUT1 | VIII | 36379 | 38508 | -1 |
| YHL033C | 42.357202 | -0.2104878 | 0.78984947 | 8.43772017 | 0.01471541 | 0.05262645 | RPL8A | VIII | 35255 | 36025 | -1 |
| YHL034C | 76.8743048 | 0.6529281 | 0.76090067 | 1.55833874 | 0.45878693 | 0.62328904 | SBP1 | VIII | 33193 | 34077 | -1 |
| YHL035C | 86.8397376 | 1.8117196 | 0.61864524 | 8.64648396 | 0.01325684 | 0.048642 | VMR1 | VIII | 27978 | 32756 | -1 |
| YHL036W | 938.761105 | 1.59425407 | 0.71476111 | 5.89733458 | 0.05240951 | 0.13534837 | MUP3 | VIII | 26241 | 27881 | 1 |
| YHL037C | 21.3621546 | -0.9350348 | 0.82340004 | 1.65295951 | 0.43758699 | 0.60627631 | NA | VIII | 25778 | 26179 | -1 |
| YHL038C | 111.420251 | -0.6846805 | 0.6676824 | 4.36987831 | 0.11248458 | 0.23882864 | CBP2 | VIII | 23617 | 25509 | -1 |
| YHL039W | 43.6555215 | 1.1546499 | 0.72266447 | 4.69040295 | 0.09582789 | 0.21233965 | EFM1 | VIII | 21783 | 23540 | 1 |
| YHL040C | 294.889157 | 2.35051518 | 0.71309498 | 77.8938087 | 1.22E-17 | 1.96E-15 | ARN1 | VIII | 19088 | 20971 | -1 |
| YHL041W | 10.3974162 | -0.3727199 | 1.10002365 | 0.11886156 | 0.94230076 | 0.96176085 | NA | VIII | 17392 | 17841 | 1 |
| YHL042W | 257.902833 | 0.24681694 | 0.47814854 | 0.53734376 | 0.76439403 | 0.85689088 | NA | VIII | 15667 | 16119 | 1 |
| YHL043W | 798.884725 | 0.03044653 | 0.47600615 | 5.83645398 | 0.0540294 | 0.13854305 | ECM34 | VIII | 14901 | 15413 | 1 |
| YHL044W | 4.68901863 | 0.79560468 | 1.38431493 | 0.56904621 | 0.75237297 | 0.85032979 | NA | VIII | 13565 | 14272 | 1 |
| YHL046C | 27.4657212 | 0.16881095 | 0.78752062 | 3.86462526 | 0.14481291 | 0.28548409 | PAU13 | VIII | 11923 | 12285 | -1 |
| YHL047C | 12.8027714 | 0.30476869 | 0.83969693 | 0.36497645 | 0.83319445 | 0.90138189 | ARN2 | VIII | 8351 | 10213 | -1 |
| YHL048W | 31.8990859 | 0.60490825 | 0.66303719 | 1.60286997 | 0.44868465 | 0.61504428 | COS8 | VIII | 6401 | 7546 | 1 |
| YHL049C | 8.0658079 | -3.5430048 | 1.34598638 | 7.74867497 | 0.02076809 | 0.0679363 | NA | VIII | 3726 | 4541 | -1 |
| YHL050C | 18.9656218 | -0.9182844 | 0.77559412 | 1.90348314 | 0.38606807 | 0.55869386 | NA | VIII | 445 | 3311 | -1 |
| YHR001W | 23.5327889 | 0.17868047 | 0.73154101 | 1.28228242 | 0.52669102 | 0.68176637 | OSH7 | VIII | 106055 | 107368 | 1 |
| YHR001W-A | 10.0558087 | 1.66101513 | 1.06191755 | 4.3365499 | 0.11437475 | 0.24133739 | QCR10 | VIII | 107826 | 108122 | 1 |
| YHR002W | 238.564713 | -0.7791282 | 0.61366857 | 1.69984657 | 0.42744772 | 0.59709268 | LEU5 | VIII | 108812 | 109885 | 1 |

| | | | | | | | | | | | |
|---|---|---|---|---|---|---|---|---|---|---|---|
| YHR003C | 884.033303 | 1.45977957 | 0.49594636 | 81.6306568 | 1.88E-18 | 3.96E-16 | TCD1 | VIII | 110028 | 111317 | -1 |
| YHR004C | 82.9089337 | -0.9157691 | 0.6503216 | 2.34293524 | 0.30991177 | 0.48301189 | NEM1 | VIII | 111754 | 113094 | -1 |
| YHR005C | 28.4260432 | -0.0016622 | 0.65907916 | 0.17179222 | 0.91768956 | 0.94899492 | GPA1 | VIII | 113499 | 114917 | -1 |
| YHR005C-A | 53.1704556 | -1.1543746 | 0.59470358 | 16.8381131 | 0.00022062 | 0.001854 | TIM10 | VIII | 115620 | 115901 | -1 |
| YHR006W | 41.360877 | 0.99739693 | 0.68226198 | 6.65401974 | 0.03590029 | 0.10239541 | STP2 | VIII | 117814 | 119439 | 1 |
| YHR007C | 92.2979884 | -0.276219 | 0.73077256 | 0.99411042 | 0.6083194 | 0.75227879 | ERG11 | VIII | 120091 | 121683 | -1 |
| YHR007C-A | 242.988431 | 1.75041559 | 0.78683316 | 5.26184772 | 0.0720119 | 0.17229294 | NA | VIII | 122550 | 122765 | -1 |
| YHR008C | 59.9975433 | -0.0916109 | 0.65426972 | 0.33361048 | 0.84636443 | 0.90975468 | SOD2 | VIII | 122889 | 123590 | -1 |
| YHR009C | 53.6304476 | -0.0967548 | 0.64165211 | 13.741433 | 0.00103773 | 0.00679692 | TDA3 | VIII | 124109 | 125680 | -1 |
| YHR010W | 97.7733878 | -0.3948349 | 0.81542134 | 12.8378324 | 0.00163042 | 0.00966238 | RPL27A | VIII | 126521 | 127492 | 1 |
| YHR011W | 65.7708976 | -0.2704596 | 0.58552705 | 0.78254188 | 0.67619692 | 0.8007253 | DIA4 | VIII | 127780 | 129120 | 1 |
| YHR012W | 27.1982144 | 0.73842447 | 0.80082222 | 0.99741464 | 0.60731522 | 0.75135793 | VPS29 | VIII | 129481 | 130448 | 1 |
| YHR013C | 112.836371 | 1.9958297 | 0.7960571 | 8.86892633 | 0.01186143 | 0.04474117 | ARD1 | VIII | 130730 | 131446 | -1 |
| YHR014W | 260.514882 | -0.6761287 | 0.51194547 | 6.44225442 | 0.03991005 | 0.11125622 | SPO13 | VIII | 132047 | 132922 | 1 |
| YHR015W | 14.1099866 | 2.05596274 | 1.04638065 | 12.4340258 | 0.0019952 | 0.01112927 | MIP6 | VIII | 134554 | 136533 | 1 |
| YHR016C | 931.345573 | 2.31344226 | 0.48758382 | 87.1562835 | 1.19E-19 | 3.27E-17 | YSC84 | VIII | 136881 | 138455 | -1 |
| YHR017W | 248.308182 | 1.59625477 | 0.71840608 | 14.9393532 | 0.00057011 | 0.00410566 | YSC83 | VIII | 138694 | 139851 | 1 |
| YHR018C | 34.3587205 | 1.59068302 | 0.98070348 | 3.39474508 | 0.18316415 | 0.33860805 | ARG4 | VIII | 140011 | 141402 | -1 |
| YHR019C | 77.3617128 | -0.2500859 | 0.56048327 | 12.3708701 | 0.00205921 | 0.01140938 | DED81 | VIII | 141894 | 143558 | -1 |
| YHR020W | 56.6220245 | -0.6135674 | 0.61278482 | 6.79057721 | 0.03353088 | 0.09746173 | NA | VIII | 143996 | 146062 | 1 |
| YHR021C | 13.860296 | -0.109763 | 0.81658947 | 1.5443795 | 0.46200029 | 0.62543411 | RPS27B | VIII | 147871 | 148669 | -1 |
| YHR021W-A | 771.131606 | 2.71628743 | 0.50453841 | 37.0595574 | 8.97E-09 | 2.72E-07 | ECM12 | VIII | 149225 | 149680 | 1 |
| YHR022C-A | 31.7437473 | 0.07662243 | 0.64505375 | 0.88133781 | 0.64360576 | 0.77686719 | NA | VIII | 151217 | 151306 | -1 |
| YHR023W | 49.7546686 | 0.78849222 | 0.73516446 | 2.04283706 | 0.36008379 | 0.5332187 | MYO1 | VIII | 151666 | 157452 | 1 |
| YHR024C | 57.3428593 | -1.0148207 | 0.56707809 | 8.48435489 | 0.01437625 | 0.051862 | MAS2 | VIII | 157744 | 159192 | -1 |
| YHR025W | 63.4415567 | -0.1495625 | 0.59076785 | 1.62006059 | 0.44484459 | 0.61266655 | THR1 | VIII | 159438 | 160511 | 1 |
| YHR026W | 13.6975987 | -0.2840215 | 0.86729088 | 0.94405818 | 0.62373537 | 0.76319268 | VMA16 | VIII | 160844 | 161485 | 1 |
| YHR027C | 190.13927 | 0.59962835 | 0.57100905 | 2.53894377 | 0.28097997 | 0.45506966 | RPN1 | VIII | 161730 | 164711 | -1 |
| YHR028C | 462.172824 | -0.7555231 | 0.46844352 | 4.81475821 | 0.090051 | 0.20366605 | DAP2 | VIII | 164978 | 167434 | -1 |
| YHR029C | 89.9416328 | -1.4791347 | 0.57552483 | 13.057017 | 0.00146118 | 0.00894319 | YHI9 | VIII | 167677 | 168561 | -1 |
| YHR030C | 119.948735 | 2.74195288 | 0.81605239 | 10.2533014 | 0.00593641 | 0.02629825 | SLT2 | VIII | 168890 | 170344 | -1 |
| YHR031C | 107.537238 | 1.40053141 | 0.54780501 | 11.5871361 | 0.00304709 | 0.01538156 | RRM3 | VIII | 170799 | 172970 | -1 |
| YHR032W | 55.459718 | -0.3172684 | 0.64394982 | 0.89856543 | 0.63808568 | 0.77431376 | ERC1 | VIII | 173344 | 175089 | 1 |

| | | | | | | | | | | | |
|---|---|---|---|---|---|---|---|---|---|---|---|
| YHR033W | 62.1613368 | 0.30382785 | 0.7313232 | 45.0016384 | 1.69E-10 | 7.47E-09 | NA | VIII | 175548 | 176819 | 1 |
| YHR034C | 40.6566423 | -0.0213343 | 0.62548364 | 0.16527055 | 0.92068688 | 0.95024857 | PIH1 | VIII | 176965 | 177999 | -1 |
| YHR035W | 31.4699039 | -0.6173496 | 0.72486019 | 3.13888074 | 0.20816164 | 0.37050597 | NEL1 | VIII | 178219 | 180111 | 1 |
| YHR036W | 64.0043558 | -0.4309271 | 0.55790567 | 0.61947612 | 0.7336391 | 0.83775415 | BRL1 | VIII | 180345 | 181760 | 1 |
| YHR037W | 20.2326832 | -0.1882875 | 0.78963651 | 1.92355739 | 0.38221244 | 0.55547441 | PUT2 | VIII | 181977 | 183704 | 1 |
| YHR038W | 166.515375 | 2.10405405 | 0.85915455 | 9.54753533 | 0.00844849 | 0.03464359 | RRF1 | VIII | 184066 | 184758 | 1 |
| YHR039C | 48.7362825 | 0.09188546 | 0.58819003 | 1.45195961 | 0.48385026 | 0.64624982 | MSC7 | VIII | 184875 | 186809 | -1 |
| YHR039C-A | 175.816476 | -0.3575834 | 0.57389724 | 11.4860289 | 0.00320509 | 0.01608101 | VMA10 | VIII | 187173 | 187679 | -1 |
| YHR040W | 58.4036396 | 0.70453489 | 0.6857455 | 8.66498794 | 0.01313475 | 0.04828584 | BCD1 | VIII | 187924 | 189024 | 1 |
| YHR041C | 277.580421 | -0.704771 | 0.53903812 | 7.2859555 | 0.02617429 | 0.08039027 | SRB2 | VIII | 189131 | 189864 | -1 |
| YHR042W | 44.3959992 | -0.1500055 | 0.64559966 | 1.64244623 | 0.43989329 | 0.60823052 | NCP1 | VIII | 190543 | 192618 | 1 |
| YHR043C | 197.668394 | 1.13920599 | 0.81676093 | 6.2389066 | 0.04418132 | 0.11942569 | DOG2 | VIII | 192805 | 193545 | -1 |
| YHR044C | 5.47872682 | 1.55796639 | 1.52000185 | 3.30137081 | 0.19191832 | 0.34834078 | DOG1 | VIII | 194068 | 194808 | -1 |
| YHR045W | 51.1711908 | 0.54278006 | 0.72962499 | 11.1882176 | 0.00371971 | 0.01814182 | NA | VIII | 195551 | 197233 | 1 |
| YHR046C | 162.217745 | 0.06473508 | 0.62923002 | 3.25339991 | 0.19657722 | 0.35446344 | INM1 | VIII | 197398 | 198285 | -1 |
| YHR047C | 45.9527251 | 0.12291812 | 0.68230951 | 0.50849857 | 0.77549846 | 0.8634877 | AAP1 | VIII | 198740 | 201310 | -1 |
| YHR048W | 648.139114 | -0.9346678 | 0.47202129 | 10.2049947 | 0.00608154 | 0.02677727 | YHK8 | VIII | 204607 | 206151 | 1 |
| YHR049W | 129.754673 | -1.1341521 | 0.58706821 | 8.57752045 | 0.01372193 | 0.04993712 | FSH1 | VIII | 206462 | 207193 | 1 |
| YHR050W | 65.5978275 | 0.16544122 | 0.602967 | 15.3857596 | 0.00045606 | 0.00341163 | SMF2 | VIII | 207654 | 209303 | 1 |
| YHR050W-A | 9.73139916 | -0.7180936 | 1.0017901 | 0.50489694 | 0.77689625 | 0.86407117 | NA | VIII | 209475 | 209645 | 1 |
| YHR051W | 45.273619 | -1.7769807 | 0.63355194 | 9.74689304 | 0.00764696 | 0.03206077 | COX6 | VIII | 209705 | 210151 | 1 |
| YHR052W | 23.42874 | -0.6244142 | 0.87438167 | 3.53837657 | 0.17047131 | 0.32067215 | CIC1 | VIII | 210848 | 211978 | 1 |
| YHR054C | 140.921217 | -1.1396795 | 0.58537421 | 3.83727249 | 0.14680703 | 0.28872715 | NA | VIII | 213185 | 214249 | -1 |
| YHR056C | 46.1740423 | -0.9907611 | 0.6744259 | 3.03813105 | 0.21891636 | 0.38305402 | RSC30 | VIII | 215183 | 217834 | -1 |
| YHR057C | 103.123444 | 0.60387405 | 0.63175285 | 3.79179011 | 0.15018385 | 0.29297995 | CPR2 | VIII | 218226 | 218843 | -1 |
| YHR058C | 17.4532122 | -0.6912531 | 0.87063856 | 1.60866911 | 0.44738554 | 0.61428226 | MED6 | VIII | 218997 | 219884 | -1 |
| YHR060W | 51.8268212 | 1.06982337 | 0.87320695 | 4.91152974 | 0.08579755 | 0.19673972 | VMA22 | VIII | 220725 | 221270 | 1 |
| YHR061C | 517.483556 | -0.0011026 | 0.51894121 | 17.6621681 | 0.00014612 | 0.00130561 | GIC1 | VIII | 221534 | 222478 | -1 |
| YHR062C | 186.950339 | -0.882868 | 0.54041146 | 2.73016646 | 0.25535942 | 0.42756826 | RPP1 | VIII | 222877 | 223758 | -1 |
| YHR063C | 45.003073 | -1.8174718 | 0.64863089 | 17.4499943 | 0.00016247 | 0.00142318 | PAN5 | VIII | 224030 | 225169 | -1 |
| YHR064C | 36.4932875 | -0.6945853 | 0.75641995 | 5.24640806 | 0.07256997 | 0.17317521 | SSZ1 | VIII | 225525 | 227141 | -1 |
| YHR065C | 55.8662978 | 0.13950259 | 0.64480817 | 10.9191023 | 0.00425547 | 0.02011359 | RRP3 | VIII | 227532 | 229037 | -1 |
| YHR066W | 124.183948 | 0.03860733 | 0.55703177 | 16.0209989 | 0.00033196 | 0.00265843 | SSF1 | VIII | 229335 | 230696 | 1 |

| | | | | | | | | | | | |
|---|---|---|---|---|---|---|---|---|---|---|---|
| YHR067W | 12.730186 | -0.6317398 | 0.96072033 | 0.48249533 | 0.78564703 | 0.87017291 | HTD2 | VIII | 230970 | 231812 | 1 |
| YHR068W | 50.2707488 | -0.0966423 | 0.58095202 | 5.7016498 | 0.05779662 | 0.14608359 | DYS1 | VIII | 232133 | 233296 | 1 |
| YHR069C | 15.7826428 | -0.5585553 | 0.78132644 | 4.46423507 | 0.10730098 | 0.23035693 | RRP4 | VIII | 233579 | 234658 | -1 |
| YHR070W | 159.695575 | -0.1385214 | 0.59109198 | 0.85630553 | 0.65171185 | 0.78244649 | TRM5 | VIII | 234881 | 236380 | 1 |
| YHR071W | 95.7925513 | -0.461997 | 0.6528754 | 1.51482406 | 0.4688783 | 0.63149694 | PCL5 | VIII | 237004 | 237693 | 1 |
| YHR072W | 149.744652 | 0.40036219 | 0.52982379 | 4.67894765 | 0.09637834 | 0.21339601 | ERG7 | VIII | 239098 | 241293 | 1 |
| YHR072W-A | 14.1572986 | -0.6063791 | 0.83766466 | 4.71968636 | 0.09443503 | 0.20984715 | NOP10 | VIII | 241664 | 241840 | 1 |
| YHR073W | 36.6355245 | 0.23305623 | 0.70005935 | 0.88556997 | 0.64224529 | 0.77621415 | OSH3 | VIII | 242582 | 245572 | 1 |
| YHR074W | 61.9480824 | 0.36808653 | 0.84102858 | 9.58984488 | 0.00827164 | 0.03401477 | QNS1 | VIII | 246193 | 248337 | 1 |
| YHR075C | 271.643022 | 1.49866678 | 0.75912588 | 6.17732293 | 0.0455629 | 0.12191042 | PPE1 | VIII | 248439 | 249641 | -1 |
| YHR076W | 338.907306 | -1.049241 | 0.50670796 | 4.24856604 | 0.11951863 | 0.24874654 | PTC7 | VIII | 251101 | 252225 | 1 |
| YHR077C | 159.798725 | 0.74502764 | 0.57114306 | 1.69414055 | 0.42866898 | 0.59835906 | NMD2 | VIII | 252374 | 255756 | -1 |
| YHR078W | 42.0136743 | 1.65951968 | 0.67625132 | 12.6520106 | 0.00178917 | 0.01030774 | NA | VIII | 256360 | 258018 | 1 |
| YHR079C | 399.16565 | -1.1260176 | 0.51377473 | 16.1443729 | 0.0003121 | 0.00252383 | IRE1 | VIII | 258244 | 261591 | -1 |
| YHR079C-A | 48.8939776 | -0.2512006 | 0.69046753 | 20.6839844 | 3.23E-05 | 0.00035703 | SAE3 | VIII | 262192 | 262553 | -1 |
| YHR080C | 160.96475 | -0.2774437 | 0.57471341 | 25.1261764 | 3.50E-06 | 5.25E-05 | LAM4 | VIII | 262801 | 266838 | -1 |
| YHR081W | 21.7973695 | -0.965499 | 0.73690524 | 5.65101965 | 0.05927843 | 0.1485167 | LRP1 | VIII | 267538 | 268092 | 1 |
| YHR082C | 51.1855884 | 0.58158072 | 0.58935154 | 0.98257172 | 0.61183915 | 0.7550189 | KSP1 | VIII | 268459 | 271548 | -1 |
| YHR083W | 83.5435918 | -0.6550149 | 0.59622252 | 12.4742663 | 0.00195545 | 0.01099231 | SAM35 | VIII | 272627 | 273616 | 1 |
| YHR084W | 69.4974651 | -1.0066 | 0.7662046 | 7.17616542 | 0.0276513 | 0.08364733 | STE12 | VIII | 274174 | 276240 | 1 |
| YHR085W | 88.957631 | -0.5708001 | 0.59857971 | 9.49152615 | 0.00868843 | 0.03535208 | IPI1 | VIII | 276764 | 277768 | 1 |
| YHR086W | 70.2020459 | 1.01313593 | 0.7034425 | 7.98815743 | 0.01842441 | 0.06216629 | NAM8 | VIII | 278153 | 279724 | 1 |
| YHR086W-A | 44.4806486 | 0.17212549 | 0.74257112 | 1.67231529 | 0.4333725 | 0.60207676 | NA | VIII | 280231 | 280392 | 1 |
| YHR087W | 40.5228695 | 0.26511869 | 0.71459002 | 0.19098196 | 0.90892656 | 0.94414869 | RTC3 | VIII | 280820 | 281155 | 1 |
| YHR088W | 35.1185151 | -0.6946584 | 0.70180169 | 5.38939933 | 0.06756267 | 0.16360847 | RPF1 | VIII | 281495 | 282382 | 1 |
| YHR089C | 157.237438 | -1.0682174 | 0.61967166 | 5.80658544 | 0.05484234 | 0.14011689 | GAR1 | VIII | 282681 | 283298 | -1 |
| YHR090C | 13.0013571 | 1.44405178 | 0.89367687 | 2.67704249 | 0.26223316 | 0.43356082 | YNG2 | VIII | 283777 | 284625 | -1 |
| YHR091C | 207.309857 | -0.6815131 | 0.50050437 | 2.48515092 | 0.28863988 | 0.46255879 | MSR1 | VIII | 284839 | 286770 | -1 |
| YHR092C | 31.7026908 | 0.98293879 | 0.7950601 | 1.92662957 | 0.38162578 | 0.55529381 | HXT4 | VIII | 287081 | 288811 | -1 |
| YHR093W | 142.893984 | -1.3345548 | 0.56292452 | 5.56543226 | 0.06187023 | 0.15328568 | NA | VIII | 289142 | 289690 | 1 |
| YHR094C | 46.0186991 | 0.29022683 | 0.6425595 | 1.84321648 | 0.39787864 | 0.56853834 | HXT1 | VIII | 290913 | 292625 | -1 |
| YHR096C | 1386.792 | 1.43103902 | 0.64485962 | 6.98950247 | 0.0303563 | 0.09004251 | HXT5 | VIII | 294669 | 296447 | -1 |
| YHR097C | 208.99292 | 2.81531741 | 0.50935246 | 29.3797669 | 4.17E-07 | 8.21E-06 | NA | VIII | 297385 | 298609 | -1 |

| | | | | | | | | | | | |
|---|---|---|---|---|---|---|---|---|---|---|---|
| YHR098C | 77.118544 | 2.06237435 | 0.61311346 | 20.7356187 | 3.14E-05 | 0.00034994 | SFB3 | VIII | 299145 | 301934 | -1 |
| YHR099W | 72.6530054 | 0.02261799 | 0.56204726 | 0.21541138 | 0.89789182 | 0.93705725 | TRA1 | VIII | 302761 | 313995 | 1 |
| YHR100C | 513.667901 | -0.4316258 | 0.57967374 | 0.79591906 | 0.67168921 | 0.79661625 | GEP4 | VIII | 314116 | 314673 | -1 |
| YHR101C | 80.1368126 | -0.7344248 | 0.54820994 | 3.69953995 | 0.15727334 | 0.30313337 | BIG1 | VIII | 314874 | 315968 | -1 |
| YHR102W | 428.90623 | -0.1198036 | 0.5018278 | 0.07592391 | 0.96274957 | 0.97258112 | KIC1 | VIII | 316572 | 319814 | 1 |
| YHR103W | 16.1815043 | -0.5235324 | 0.8576208 | 0.99072727 | 0.60934929 | 0.75306988 | SBE22 | VIII | 320414 | 322972 | 1 |
| YHR104W | 22.3748001 | 0.39995456 | 0.79498259 | 3.69267236 | 0.15781431 | 0.30407483 | GRE3 | VIII | 323409 | 324392 | 1 |
| YHR105W | 14.3254284 | -0.3830568 | 0.88859769 | 4.81156735 | 0.09019478 | 0.20366605 | YPT35 | VIII | 324766 | 325410 | 1 |
| YHR106W | 35.1150828 | 0.36274106 | 0.71042847 | 1.85987572 | 0.39457823 | 0.56638907 | TRR2 | VIII | 325598 | 326626 | 1 |
| YHR107C | 31.8098845 | 0.27194648 | 0.64535045 | 1.14492049 | 0.56413582 | 0.71504956 | CDC12 | VIII | 326813 | 328036 | -1 |
| YHR108W | 41.6577891 | -0.1141627 | 0.61628038 | 0.33268049 | 0.84675808 | 0.90976606 | GGA2 | VIII | 328303 | 330060 | 1 |
| YHR109W | 20.840772 | -0.6277958 | 0.87759897 | 1.36165653 | 0.50619755 | 0.66417229 | CTM1 | VIII | 330310 | 332067 | 1 |
| YHR110W | 10.0573769 | 0.1314056 | 1.01421761 | 0.09233673 | 0.95488118 | 0.96809001 | ERP5 | VIII | 332282 | 332920 | 1 |
| YHR111W | 421.722519 | 0.16399209 | 0.46419794 | 0.22727835 | 0.89257996 | 0.93387025 | UBA4 | VIII | 333072 | 334394 | 1 |
| YHR112C | 493.886505 | -1.5233074 | 0.51258974 | 35.757229 | 1.72E-08 | 4.86E-07 | NA | VIII | 334527 | 335663 | -1 |
| YHR113W | 100.052273 | -1.0504311 | 0.58145578 | 3.6800863 | 0.15881057 | 0.30548612 | APE4 | VIII | 336337 | 337809 | 1 |
| YHR114W | 130.971018 | -1.2002141 | 0.52809222 | 15.8269778 | 0.00036578 | 0.00285809 | BZZ1 | VIII | 338083 | 339984 | 1 |
| YHR115C | 371.662103 | 1.1684528 | 0.6668981 | 9.91899309 | 0.00701646 | 0.0300483 | DMA1 | VIII | 340109 | 341359 | -1 |
| YHR116W | 21.7106812 | -2.0716875 | 0.82214278 | 9.02876601 | 0.01095036 | 0.04215597 | COX23 | VIII | 341665 | 342120 | 1 |
| YHR117W | 207.970067 | -0.5997989 | 0.54080465 | 11.4079538 | 0.00333269 | 0.01653492 | TOM71 | VIII | 342349 | 344268 | 1 |
| YHR118C | 120.598348 | 0.54794012 | 0.62547643 | 7.0720478 | 0.02912892 | 0.08689151 | ORC6 | VIII | 344321 | 345628 | -1 |
| YHR119W | 156.180524 | -0.9726949 | 0.50149453 | 7.50560235 | 0.02345196 | 0.07454061 | SET1 | VIII | 346043 | 349285 | 1 |
| YHR120W | 23.3135607 | -0.327977 | 0.75448117 | 0.22255234 | 0.89469163 | 0.9350658 | MSH1 | VIII | 349574 | 352453 | 1 |
| YHR121W | 29.2357214 | -0.9078551 | 0.72130341 | 4.04018594 | 0.13264313 | 0.26853278 | LSM12 | VIII | 352756 | 353319 | 1 |
| YHR122W | 26.6266893 | -0.2397149 | 0.69904498 | 0.44263869 | 0.8014607 | 0.88031084 | CIA2 | VIII | 353625 | 354320 | 1 |
| YHR123W | 35.5878082 | -0.0467285 | 0.70120883 | 10.3307812 | 0.00571083 | 0.025494 | EPT1 | VIII | 354815 | 356081 | 1 |
| YHR124W | 1493.9623 | -0.301385 | 0.4670483 | 0.41624796 | 0.81210634 | 0.88731158 | NDT80 | VIII | 356561 | 358444 | 1 |
| YHR125W | 9.13429095 | -0.2388863 | 1.1267974 | 1.72770977 | 0.42153398 | 0.59239848 | NA | VIII | 358858 | 359163 | 1 |
| YHR126C | 6.48106968 | 0.67186388 | 1.19748671 | 2.22956852 | 0.32798603 | 0.50041094 | ANS1 | VIII | 359702 | 360181 | -1 |
| YHR127W | 63.354218 | -2.9015999 | 0.62021588 | 43.2213978 | 4.12E-10 | 1.64E-08 | NA | VIII | 360913 | 361644 | 1 |
| YHR128W | 107.628796 | -0.1924313 | 0.52930035 | 0.40874108 | 0.81516027 | 0.88884707 | FUR1 | VIII | 362115 | 362765 | 1 |
| YHR129C | 44.9403704 | -0.2392655 | 0.66785432 | 5.04858957 | 0.08011479 | 0.18666585 | ARP1 | VIII | 362999 | 364153 | -1 |
| YHR130C | 32.001307 | 0.52958261 | 0.79836438 | 2.40754248 | 0.30006048 | 0.47326346 | NA | VIII | 364965 | 365300 | -1 |

| | | | | | | | | | | | |
|---|---|---|---|---|---|---|---|---|---|---|---|
| YHR131C | 48.9503403 | 0.17303213 | 0.63708829 | 0.11123634 | 0.94590024 | 0.96354018 | NA | VIII | 365340 | 367892 | -1 |
| YHR132C | 67.2291349 | 0.73918743 | 0.69908472 | 1.34151283 | 0.51132166 | 0.66829626 | ECM14 | VIII | 368502 | 369794 | -1 |
| YHR132W-A | 40.1234912 | 0.21131877 | 0.66731984 | 0.92617982 | 0.62933605 | 0.76680754 | IGO2 | VIII | 370054 | 370449 | 1 |
| YHR133C | 53.481965 | 0.59819609 | 0.67035216 | 1.04806012 | 0.59212942 | 0.73920425 | NSG1 | VIII | 370722 | 371597 | -1 |
| YHR134W | 1803.52476 | 2.86411218 | 0.47770331 | 46.2125226 | 9.23E-11 | 4.42E-09 | WSS1 | VIII | 371749 | 372558 | 1 |
| YHR135C | 1225.45617 | 1.03039136 | 0.45976419 | 6.56128726 | 0.03760405 | 0.1062085 | YCK1 | VIII | 372694 | 374310 | -1 |
| YHR136C | 6.11830014 | -0.5820841 | 1.24919737 | 1.7048867 | 0.42637188 | 0.59587092 | SPL2 | VIII | 374654 | 375100 | -1 |
| YHR137W | 248.108564 | 4.02708334 | 0.93092152 | 15.1956828 | 0.00050153 | 0.00368045 | ARO9 | VIII | 375709 | 377250 | 1 |
| YHR138C | 206.260095 | 1.95329683 | 0.81970463 | 5.47464679 | 0.06474341 | 0.15863916 | NA | VIII | 377355 | 377699 | -1 |
| YHR139C | 175.346183 | -1.2797356 | 0.54679866 | 8.39998243 | 0.01499571 | 0.05349671 | SPS100 | VIII | 378219 | 379199 | -1 |
| YHR139C-A | 105.269644 | -0.4465448 | 0.57379632 | 2.39477846 | 0.30198159 | 0.47448396 | NA | VIII | 380108 | 380419 | -1 |
| YHR140W | 36.947971 | 0.36127278 | 0.6556401 | 0.52159893 | 0.7704354 | 0.86016602 | NA | VIII | 380572 | 381291 | 1 |
| YHR141C | 113.083862 | -0.8170627 | 0.71464102 | 18.9800347 | 7.56E-05 | 0.00073818 | RPL42A | VIII | 381990 | 382751 | -1 |
| YHR142W | 510.825248 | -1.0331024 | 0.52851644 | 24.8806211 | 3.96E-06 | 5.84E-05 | CHS7 | VIII | 383538 | 384488 | 1 |
| YHR143W | 381.204132 | 0.26572935 | 0.49626228 | 4.42149471 | 0.10961869 | 0.23403106 | DSE2 | VIII | 385510 | 386487 | 1 |
| YHR143W-A | 15.3154485 | -0.1800203 | 0.82915124 | 0.1056259 | 0.94855742 | 0.96471939 | RPC10 | VIII | 387233 | 387445 | 1 |
| YHR144C | 24.7359693 | 0.77687516 | 0.73361861 | 4.86447683 | 0.08783999 | 0.20034795 | DCD1 | VIII | 387788 | 388726 | -1 |
| YHR146W | 53.3880307 | -0.2149592 | 0.6300153 | 0.71141646 | 0.70067703 | 0.81801018 | CRP1 | VIII | 390300 | 391697 | 1 |
| YHR147C | 182.239972 | -0.0046479 | 0.5323572 | 13.9779109 | 0.00092201 | 0.00615027 | MRPL6 | VIII | 392639 | 393283 | -1 |
| YHR148W | 7.76600939 | -0.2921335 | 1.00651858 | 6.34560741 | 0.041886 | 0.11493835 | IMP3 | VIII | 393534 | 394085 | 1 |
| YHR149C | 52.5911302 | -0.2214286 | 0.63724581 | 0.20328776 | 0.9033512 | 0.93919975 | SKG6 | VIII | 394455 | 396659 | -1 |
| YHR150W | 18.8336382 | -1.6407348 | 0.99330204 | 11.154229 | 0.00378347 | 0.01836234 | PEX28 | VIII | 397251 | 398990 | 1 |
| YHR151C | 12.6354726 | -0.7839975 | 0.93936335 | 3.8213306 | 0.1479819 | 0.2902937 | MTC6 | VIII | 399268 | 400848 | -1 |
| YHR152W | 75.0417121 | -1.2616895 | 0.64981914 | 18.2584721 | 0.00010845 | 0.00100306 | SPO12 | VIII | 401434 | 401955 | 1 |
| YHR153C | 8.8817812 | -0.8267238 | 0.94827185 | 6.68394776 | 0.03536708 | 0.10132379 | SPO16 | VIII | 402086 | 402682 | -1 |
| YHR154W | 118.307027 | -0.5018695 | 0.52328167 | 2.10338458 | 0.34934605 | 0.52293528 | RTT107 | VIII | 402966 | 406178 | 1 |
| YHR155W | 33.364221 | -0.4833745 | 0.64517034 | 0.68060863 | 0.71155375 | 0.82438173 | LAM1 | VIII | 407103 | 410789 | 1 |
| YHR156C | 271.65265 | -0.9142225 | 0.54241062 | 9.8950464 | 0.00710097 | 0.03034291 | LIN1 | VIII | 411384 | 412406 | -1 |
| YHR158C | 27.2764948 | -0.5850212 | 0.75409208 | 0.74758773 | 0.68811874 | 0.81029236 | KEL1 | VIII | 413685 | 417179 | -1 |
| YHR159W | 26.8608684 | 0.79444291 | 0.73582122 | 2.98374417 | 0.22495113 | 0.3904964 | TDA11 | VIII | 417549 | 419063 | 1 |
| YHR160C | 539.703211 | -0.0929086 | 0.4842328 | 4.3095163 | 0.11593123 | 0.24332354 | PEX18 | VIII | 419221 | 420072 | -1 |
| YHR161C | 73.3051109 | 0.9173793 | 0.58295999 | 2.86132498 | 0.23915044 | 0.40763705 | YAP1801 | VIII | 420373 | 422286 | -1 |
| YHR162W | 69.7417583 | 0.30120605 | 0.54758113 | 7.21043517 | 0.02718153 | 0.08256862 | MPC2 | VIII | 423072 | 423461 | 1 |

| | | | | | | | | | | | |
|---|---|---|---|---|---|---|---|---|---|---|---|
| YHR163W | 53.0931978 | 0.61161501 | 0.59141626 | 3.049795 | 0.21764336 | 0.38155469 | SOL3 | VIII | 423724 | 424473 | 1 |
| YHR164C | 172.361879 | 1.62918456 | 0.73239733 | 8.30136888 | 0.01575363 | 0.05576982 | DNA2 | VIII | 424610 | 429178 | -1 |
| YHR165C | 90.7947384 | 0.42890185 | 0.56504914 | 7.36069567 | 0.0252142 | 0.078237 | PRP8 | VIII | 429707 | 436948 | -1 |
| YHR166C | 16.9837717 | -0.3807454 | 0.83073097 | 1.97373835 | 0.37274186 | 0.546097 | CDC23 | VIII | 437170 | 439050 | -1 |
| YHR167W | 13.9078478 | -0.0726279 | 0.91970462 | 0.03124002 | 0.98450135 | 0.98751104 | THP2 | VIII | 439342 | 440127 | 1 |
| YHR168W | 22.9099055 | -1.3845291 | 0.70845513 | 12.0371563 | 0.00243313 | 0.01296026 | MTG2 | VIII | 440377 | 441933 | 1 |
| YHR169W | 39.9564375 | -0.0051247 | 0.76487608 | 0.32574404 | 0.84969992 | 0.91079805 | DBP8 | VIII | 442181 | 443476 | 1 |
| YHR170W | 34.377728 | -1.0139368 | 0.7195822 | 3.42110355 | 0.18076602 | 0.33535254 | NMD3 | VIII | 443828 | 445384 | 1 |
| YHR171W | 25.5372705 | -0.4983314 | 0.6990845 | 0.8982898 | 0.63817362 | 0.77431376 | ATG7 | VIII | 445713 | 447605 | 1 |
| YHR172W | 54.5244315 | 0.92161798 | 0.68987098 | 5.71589629 | 0.05738639 | 0.14528517 | SPC97 | VIII | 448335 | 450806 | 1 |
| YHR173C | 434.301909 | -0.9164039 | 0.46793172 | 10.1236357 | 0.00633403 | 0.02761601 | NA | VIII | 450815 | 451153 | -1 |
| YHR174W | 678.1549 | -0.2281321 | 0.63573518 | 6.25252025 | 0.0438816 | 0.11883745 | ENO2 | VIII | 451327 | 452640 | 1 |
| YHR175W | 27.6180233 | 0.52906961 | 0.7055308 | 0.9563941 | 0.61990003 | 0.76058936 | CTR2 | VIII | 452872 | 453441 | 1 |
| YHR176W | 31.0946548 | 0.88245138 | 0.69661583 | 1.60946785 | 0.4472069 | 0.61428226 | FMO1 | VIII | 454229 | 455527 | 1 |
| YHR177W | 1104.10105 | 0.84184831 | 0.52871471 | 10.4104042 | 0.00548794 | 0.02467017 | NA | VIII | 456591 | 457952 | 1 |
| YHR178W | 85.5455634 | -1.5504019 | 0.55357304 | 28.5393871 | 6.35E-07 | 1.20E-05 | STB5 | VIII | 459299 | 461530 | 1 |
| YHR179W | 74.2830905 | 1.43612137 | 0.67998274 | 4.54347725 | 0.10313271 | 0.22389892 | OYE2 | VIII | 462502 | 463704 | 1 |
| YHR180W | 730.352155 | 0.67040383 | 0.48692005 | 30.9721806 | 1.88E-07 | 4.16E-06 | NA | VIII | 465178 | 465669 | 1 |
| YHR181W | 20.1224725 | -1.0959501 | 0.75286418 | 2.07511295 | 0.35431941 | 0.52819501 | SVP26 | VIII | 467228 | 467914 | 1 |
| YHR182W | 377.380662 | 0.76672892 | 0.49181717 | 3.01103575 | 0.22190235 | 0.38675936 | NA | VIII | 468219 | 470576 | 1 |
| YHR183W | 118.404268 | 0.8764108 | 0.78465437 | 3.88458107 | 0.14337517 | 0.28351852 | GND1 | VIII | 470960 | 472429 | 1 |
| YHR186C | 61.8092633 | 0.85338241 | 0.68112033 | 14.0614319 | 0.0008843 | 0.00595552 | KOG1 | VIII | 475999 | 480672 | -1 |
| YHR187W | 170.953949 | 2.06269605 | 0.81835732 | 12.4000568 | 0.00202937 | 0.01126565 | IKI1 | VIII | 480990 | 481919 | 1 |
| YHR188C | 119.399674 | 1.0332263 | 0.7243454 | 2.55069991 | 0.2793332 | 0.45351049 | GPI16 | VIII | 482005 | 483837 | -1 |
| YHR189W | 202.737102 | -0.7748654 | 0.6204814 | 1.55002349 | 0.46069837 | 0.62469404 | PTH1 | VIII | 484028 | 484600 | 1 |
| YHR190W | 161.280792 | 0.75671939 | 0.59291601 | 5.5636494 | 0.06192541 | 0.15335677 | ERG9 | VIII | 484845 | 486179 | 1 |
| YHR191C | 1594.53219 | 1.60551111 | 0.63424534 | 9.63795636 | 0.00807503 | 0.03341947 | CTF8 | VIII | 486230 | 486631 | -1 |
| YHR192W | 1091.47309 | 2.72156999 | 1.60871088 | 5.24635764 | 0.0725718 | 0.17317521 | LNP1 | VIII | 486826 | 487662 | 1 |
| YHR193C | 530.214974 | 0.46167629 | 0.50297068 | 6.63062021 | 0.03632278 | 0.10344757 | EGD2 | VIII | 487712 | 488236 | -1 |
| YHR194W | 63.4478203 | -1.1913221 | 0.62399001 | 4.53058794 | 0.10379952 | 0.22509714 | MDM31 | VIII | 488657 | 490396 | 1 |
| YHR195W | 76.5297286 | -0.8316565 | 0.55534872 | 3.14527917 | 0.20749675 | 0.36958676 | NVJ1 | VIII | 490747 | 491712 | 1 |
| YHR196W | 27.5811457 | -1.3169285 | 0.93802225 | 2.68548535 | 0.26112849 | 0.43235173 | UTP9 | VIII | 491931 | 493658 | 1 |
| YHR197W | 73.9548284 | 0.1044126 | 0.69768586 | 2.7578336 | 0.25185121 | 0.42340839 | RIX1 | VIII | 493896 | 496187 | 1 |

| | | | | | | | | | | | |
|---|---|---|---|---|---|---|---|---|---|---|---|
| YHR198C | 129.36181 | -1.9657976 | 0.52835639 | 29.0096711 | 5.02E-07 | 9.75E-06 | AIM18 | VIII | 496315 | 497280 | -1 |
| YHR199C | 32.4552259 | -0.2350584 | 0.75973122 | 0.11876813 | 0.94234478 | 0.96176085 | AIM46 | VIII | 497490 | 498422 | -1 |
| YHR199C-A | 21.8386558 | -1.4188941 | 0.80749814 | 3.61694568 | 0.16390425 | 0.31248127 | NBL1 | VIII | 498571 | 498859 | -1 |
| YHR200W | 143.280763 | 1.59840351 | 0.7087275 | 9.42409569 | 0.00898636 | 0.0363345 | RPN10 | VIII | 499079 | 499885 | 1 |
| YHR201C | 159.190447 | 0.51210572 | 0.62169124 | 7.42667345 | 0.02439598 | 0.07671608 | PPX1 | VIII | 499950 | 501143 | -1 |
| YHR202W | 42.0338164 | -1.3867443 | 0.67313099 | 9.58999518 | 0.00827102 | 0.03401477 | NA | VIII | 502388 | 504196 | 1 |
| YHR203C | 75.3881069 | -0.8124078 | 0.72214222 | 13.8980416 | 0.00095957 | 0.00635691 | RPS4A | VIII | 504476 | 505530 | -1 |
| YHR204W | 171.634151 | -1.6118379 | 0.52066504 | 9.92617465 | 0.00699131 | 0.02997984 | MNL1 | VIII | 506319 | 508709 | 1 |
| YHR205W | 168.338849 | 0.80399176 | 0.57603449 | 8.26593772 | 0.0160352 | 0.05640572 | SCH9 | VIII | 509363 | 511837 | 1 |
| YHR206W | 31.2941275 | 0.13887098 | 0.66221612 | 0.28990772 | 0.86506221 | 0.92054956 | SKN7 | VIII | 512732 | 514600 | 1 |
| YHR207C | 49.4771919 | -0.5123349 | 0.6102889 | 0.72285294 | 0.69668182 | 0.81659461 | SET5 | VIII | 514905 | 516485 | -1 |
| YHR208W | 39.0195505 | 0.60444018 | 0.63636178 | 4.23387809 | 0.1203996 | 0.24977202 | BAT1 | VIII | 517532 | 518713 | 1 |
| YHR209W | 330.747873 | -0.4599696 | 0.48032053 | 5.0000732 | 0.08208199 | 0.18979822 | CRG1 | VIII | 519437 | 520312 | 1 |
| YHR210C | 11.024383 | 1.18221809 | 0.93476554 | 2.13406732 | 0.34402751 | 0.51765054 | NA | VIII | 520712 | 521737 | -1 |
| YHR211W | 154.662749 | -0.3431037 | 0.50494293 | 3.21752406 | 0.20013522 | 0.35908985 | FLO5 | VIII | 525392 | 528619 | 1 |
| YHR213W | 17.0950159 | 0.31943742 | 0.78921798 | 0.97052035 | 0.61553703 | 0.75780553 | NA | VIII | 539151 | 539747 | 1 |
| YHR213W-B | 151.170909 | 1.2144778 | 0.53684394 | 16.9576885 | 0.00020782 | 0.00176433 | NA | VIII | 540800 | 541099 | 1 |
| YHR214C-B | 8.68495834 | 0.3569992 | 0.98767411 | 2.3218241 | 0.3132004 | 0.48516903 | NA | VIII | 543964 | 549346 | -1 |
| YHR214C-E | 13.4042596 | -0.0944601 | 0.85618263 | 0.65206199 | 0.72178282 | 0.83133529 | NA | VIII | 551200 | 551499 | -1 |
| YHR214W | 172.718877 | -0.2965247 | 0.56289875 | 4.4084099 | 0.11033822 | 0.23522866 | NA | VIII | 541651 | 542262 | 1 |
| YHR216W | 31.8040126 | 0.02336211 | 0.76664703 | 3.53611718 | 0.170664 | 0.32088433 | IMD2 | VIII | 554396 | 555967 | 1 |
| YIL001W | 9.79863105 | -0.1158081 | 1.13725021 | 0.32851834 | 0.84852208 | 0.90980423 | NA | IX | 353940 | 355481 | 1 |
| YIL002C | 433.31064 | 1.97895715 | 0.71707697 | 8.05950837 | 0.0177787 | 0.06041002 | INP51 | IX | 350591 | 353431 | -1 |
| YIL002W-A | 86.236974 | 1.01239748 | 0.60909938 | 3.10045081 | 0.21220014 | 0.37592901 | NA | IX | 350301 | 350510 | 1 |
| YIL003W | 21.9248764 | 0.69979271 | 0.78344665 | 5.01877938 | 0.08131785 | 0.18855842 | CFD1 | IX | 349122 | 350003 | 1 |
| YIL004C | 398.232624 | 1.92020613 | 0.78687208 | 7.41543279 | 0.02453348 | 0.07690789 | BET1 | IX | 347946 | 348505 | -1 |
| YIL005W | 258.753057 | 0.72652119 | 0.63869718 | 1.6238387 | 0.44400505 | 0.61165577 | EPS1 | IX | 345692 | 347797 | 1 |
| YIL006W | 17.8462814 | -1.2107515 | 0.88224789 | 7.6572039 | 0.02173999 | 0.07043902 | YIA6 | IX | 344062 | 345183 | 1 |
| YIL007C | 13.924541 | 0.64197547 | 0.87025463 | 0.79186424 | 0.67305238 | 0.79774275 | NAS2 | IX | 342994 | 343656 | -1 |
| YIL008W | 106.066739 | 1.61449621 | 0.66961974 | 12.1858249 | 0.00225882 | 0.01226883 | URM1 | IX | 342536 | 342835 | 1 |
| YIL009C-A | 434.988398 | 2.3918449 | 0.60470714 | 14.5726399 | 0.00068484 | 0.00479473 | EST3 | IX | 335666 | 336212 | -1 |
| YIL009W | 20.8244296 | -0.6560087 | 0.82933057 | 0.6637434 | 0.71757938 | 0.8289674 | FAA3 | IX | 339344 | 341428 | 1 |
| YIL010W | 41.6564036 | 0.72087041 | 0.73374997 | 1.69489519 | 0.42850726 | 0.59827756 | DOT5 | IX | 334882 | 335529 | 1 |

| | | | | | | | | | | | |
|---|---|---|---|---|---|---|---|---|---|---|---|
| YIL011W | 109.11944 | 1.51840996 | 0.67200256 | 5.25178402 | 0.07237517 | 0.17294768 | TIR3 | IX | 333727 | 334536 | 1 |
| YIL012W | 11.9370033 | 1.67832013 | 0.96155604 | 2.9706482 | 0.22642894 | 0.39252203 | NA | IX | 333011 | 333352 | 1 |
| YIL013C | 1451.67203 | 0.05360371 | 0.63449981 | 2.0527081 | 0.35831097 | 0.53140894 | PDR11 | IX | 328207 | 332442 | -1 |
| YIL014C-A | 8.92910469 | 0.25579808 | 1.05486366 | 0.70114075 | 0.70428627 | 0.82015637 | NA | IX | 325212 | 325526 | -1 |
| YIL014W | 24.2396863 | 0.17647616 | 0.72489895 | 1.38095202 | 0.50133737 | 0.66037092 | MNT3 | IX | 326103 | 327995 | 1 |
| YIL015W | 245.773191 | -0.1172952 | 0.54017967 | 7.36568106 | 0.02515143 | 0.07816789 | BAR1 | IX | 322342 | 324105 | 1 |
| YIL016W | 71.1651908 | -0.622808 | 0.63682686 | 1.5008688 | 0.4721614 | 0.63393528 | SNL1 | IX | 321454 | 321933 | 1 |
| YIL017C | 59.2934768 | -0.0478827 | 0.57163973 | 1.50804469 | 0.47047035 | 0.63290505 | VID28 | IX | 318200 | 320965 | -1 |
| YIL018W | 142.084004 | -0.7019107 | 0.7869244 | 12.5000757 | 0.00193038 | 0.01089367 | RPL2A | IX | 316768 | 317932 | 1 |
| YIL019W | 34.3386162 | -2.2748424 | 0.86115083 | 23.5347838 | 7.75E-06 | 0.00010489 | FAF1 | IX | 315093 | 316133 | 1 |
| YIL020C | 153.413573 | -0.3153898 | 0.67379514 | 12.7816775 | 0.00167685 | 0.00980703 | HIS6 | IX | 314035 | 314820 | -1 |
| YIL021W | 20.9619285 | -0.3929936 | 0.74954217 | 6.42528192 | 0.04025017 | 0.11198871 | RPB3 | IX | 312905 | 313861 | 1 |
| YIL022W | 52.2732232 | -0.3025813 | 0.58643589 | 2.17711643 | 0.33670159 | 0.50994042 | TIM44 | IX | 311165 | 312460 | 1 |
| YIL023C | 144.538004 | 1.75885551 | 0.51953656 | 22.3764965 | 1.38E-05 | 0.00017415 | YKE4 | IX | 309386 | 310426 | -1 |
| YIL024C | 7.43760947 | 2.38206351 | 1.2086582 | 4.15260639 | 0.12539291 | 0.25809632 | NA | IX | 308602 | 309171 | -1 |
| YIL026C | 282.919182 | 0.66891401 | 0.49988664 | 2.7169607 | 0.25705111 | 0.42953129 | IRR1 | IX | 304477 | 307929 | -1 |
| YIL030C | 35.906532 | 0.03385792 | 0.63213392 | 2.58189031 | 0.27501073 | 0.44841232 | SSM4 | IX | 296050 | 300009 | -1 |
| YIL031W | 149.548759 | 1.21853846 | 0.51867178 | 17.296613 | 0.00017542 | 0.00151597 | ULP2 | IX | 292633 | 295737 | 1 |
| YIL033C | 231.459138 | -1.3609114 | 0.5305346 | 15.9438531 | 0.00034501 | 0.00274023 | BCY1 | IX | 290419 | 291669 | -1 |
| YIL034C | 519.175859 | -1.598775 | 0.48038918 | 22.5461931 | 1.27E-05 | 0.0001621 | CAP2 | IX | 289226 | 290089 | -1 |
| YIL035C | 24.5789834 | -0.0364635 | 0.78818528 | 3.12458943 | 0.20965442 | 0.37244958 | CKA1 | IX | 287790 | 288908 | -1 |
| YIL036W | 67.7484089 | -0.2529848 | 0.57975573 | 10.6786813 | 0.00479903 | 0.02212703 | CST6 | IX | 285666 | 287429 | 1 |
| YIL037C | 668.880673 | -0.3659737 | 0.51835749 | 0.59642313 | 0.74214431 | 0.84243165 | PRM2 | IX | 283029 | 284999 | -1 |
| YIL038C | 27.4108496 | 0.49722453 | 0.7187551 | 0.67961284 | 0.71190812 | 0.8243896 | NOT3 | IX | 280142 | 282652 | -1 |
| YIL039W | 17.373672 | -0.2288319 | 0.9880283 | 0.30274249 | 0.85952855 | 0.91706593 | TED1 | IX | 278426 | 279847 | 1 |
| YIL040W | 130.185184 | 0.12429184 | 0.53856236 | 4.95215402 | 0.08407239 | 0.19331976 | APQ12 | IX | 277723 | 278139 | 1 |
| YIL041W | 75.7198193 | 0.46920058 | 0.57806699 | 0.97660885 | 0.61366603 | 0.75646717 | GVP36 | IX | 276525 | 277505 | 1 |
| YIL042C | 7.5799814 | 1.44032126 | 1.19192019 | 1.69982488 | 0.42745236 | 0.59709268 | PKP1 | IX | 275108 | 276292 | -1 |
| YIL043C | 45.9601708 | -0.1970941 | 0.6982182 | 4.63113393 | 0.0987102 | 0.21722998 | CBR1 | IX | 274072 | 274926 | -1 |
| YIL044C | 21.1624471 | 0.22551931 | 0.81900847 | 2.69283232 | 0.260171 | 0.43177286 | AGE2 | IX | 272950 | 273846 | -1 |
| YIL045W | 142.626499 | 2.28948345 | 0.77882509 | 8.24925656 | 0.0161695 | 0.05670589 | PIG2 | IX | 271161 | 272777 | 1 |
| YIL046W | 40.9461182 | -0.0886852 | 0.63017385 | 1.74300542 | 0.41832246 | 0.589459 | MET30 | IX | 268651 | 270573 | 1 |
| YIL046W-A | 19.6365052 | 2.08117783 | 1.23820668 | 2.70983736 | 0.25796827 | 0.42992972 | NA | IX | 268309 | 268473 | 1 |

| | | | | | | | | | | | |
|---|---|---|---|---|---|---|---|---|---|---|---|
| YIL047C | 40.7818032 | 0.67398934 | 0.63631293 | 2.37373713 | 0.30517541 | 0.47807485 | SYG1 | IX | 265115 | 267823 | -1 |
| YIL048W | 85.473896 | -1.1059001 | 0.56294728 | 4.88293387 | 0.08703309 | 0.19878563 | NEO1 | IX | 261437 | 264892 | 1 |
| YIL049W | 86.7815832 | 0.6760185 | 0.56402278 | 1.95397334 | 0.37644374 | 0.54998972 | DFG10 | IX | 260158 | 260919 | 1 |
| YIL050W | 144.39429 | 0.15474774 | 0.5590967 | 3.61938917 | 0.16370413 | 0.3123054 | PCL7 | IX | 258913 | 259770 | 1 |
| YIL051C | 228.648128 | 0.45715867 | 0.51023142 | 0.83994247 | 0.65706572 | 0.78549013 | MMF1 | IX | 257843 | 258280 | -1 |
| YIL052C | 46.2448755 | -0.2519531 | 0.7394116 | 11.2269567 | 0.00364836 | 0.01787139 | RPL34B | IX | 256226 | 257063 | -1 |
| YIL053W | 67.5568594 | -0.2124879 | 0.58946234 | 12.5258667 | 0.00190565 | 0.0107751 | GPP1 | IX | 255115 | 255867 | 1 |
| YIL054W | 52.6158577 | 0.9913726 | 0.71718344 | 1.96527426 | 0.37432266 | 0.54758166 | NA | IX | 254543 | 254860 | 1 |
| YIL055C | 47.967356 | 1.89623758 | 0.75520758 | 6.36606337 | 0.04145977 | 0.11428804 | NA | IX | 252042 | 253925 | -1 |
| YIL056W | 79.5819769 | 2.27362372 | 0.84085562 | 6.77402217 | 0.03380958 | 0.09812404 | VHR1 | IX | 249991 | 251913 | 1 |
| YIL060W | 1319.77337 | 0.30828166 | 0.48922685 | 12.144879 | 0.00230554 | 0.01240622 | NA | IX | 246392 | 246826 | 1 |
| YIL061C | 13.1850525 | -0.77611 | 0.95963486 | 1.28021519 | 0.52723569 | 0.68184633 | SNP1 | IX | 244657 | 245559 | -1 |
| YIL062C | 216.89699 | 3.98539886 | 0.75975764 | 27.3701352 | 1.14E-06 | 1.98E-05 | ARC15 | IX | 243998 | 244462 | -1 |
| YIL063C | 1340.64674 | 1.53004765 | 0.4888984 | 42.2523172 | 6.68E-10 | 2.50E-08 | YRB2 | IX | 242761 | 243744 | -1 |
| YIL064W | 192.655507 | 0.15139892 | 0.52993663 | 6.3365012 | 0.04207714 | 0.11535353 | EFM4 | IX | 241943 | 242716 | 1 |
| YIL065C | 6.70773065 | 1.3831869 | 1.15150037 | 1.44378363 | 0.48583228 | 0.64773825 | FIS1 | IX | 241308 | 241775 | -1 |
| YIL066C | 30.1138084 | 0.64194036 | 0.70404017 | 7.48689938 | 0.0236723 | 0.07502059 | RNR3 | IX | 238099 | 240708 | -1 |
| YIL067C | 93.6303846 | 0.61573706 | 0.63869851 | 30.6728748 | 2.19E-07 | 4.70E-06 | NA | IX | 235724 | 237760 | -1 |
| YIL068C | 11.4114596 | 0.31971331 | 0.94611878 | 0.33576862 | 0.84545164 | 0.90923806 | SEC6 | IX | 233057 | 235474 | -1 |
| YIL069C | 47.4509569 | -0.6449726 | 0.94186036 | 10.1276335 | 0.00632139 | 0.02758163 | RPS24A | IX | 231553 | 232369 | -1 |
| YIL070C | 9.06439748 | -0.7054617 | 0.96912869 | 1.10112539 | 0.57662526 | 0.72593834 | MAM33 | IX | 230272 | 231072 | -1 |
| YIL071C | 151.270349 | 0.63536227 | 0.54229119 | 6.35872972 | 0.04161208 | 0.11445792 | PCI8 | IX | 228660 | 229994 | -1 |
| YIL072W | 683.280405 | 0.07585869 | 0.52025538 | 0.91267424 | 0.6336002 | 0.77037908 | HOP1 | IX | 226602 | 228419 | 1 |
| YIL073C | 1177.557 | 0.4932188 | 0.46361036 | 1.3807817 | 0.50138007 | 0.66037092 | SPO22 | IX | 222937 | 225954 | -1 |
| YIL074C | 83.3688747 | 1.21223354 | 0.66656439 | 3.36522895 | 0.18588734 | 0.34169876 | SER33 | IX | 221081 | 222490 | -1 |
| YIL075C | 1207.01814 | 1.90830916 | 0.73015843 | 6.71901579 | 0.03475236 | 0.10004686 | RPN2 | IX | 217863 | 220700 | -1 |
| YIL076W | 25.4445262 | 2.08734521 | 0.95548058 | 4.80809005 | 0.09035174 | 0.20387239 | SEC28 | IX | 216658 | 217548 | 1 |
| YIL077C | 68.1574905 | -1.6799818 | 0.59426073 | 7.98245852 | 0.01847699 | 0.06230737 | NA | IX | 214991 | 215953 | -1 |
| YIL078W | 72.5993962 | -0.1064818 | 0.69292302 | 4.64697453 | 0.09793148 | 0.2158444 | THS1 | IX | 212499 | 214703 | 1 |
| YIL079C | 81.9993244 | -0.3557994 | 0.58943146 | 0.36529811 | 0.83306046 | 0.90138189 | AIR1 | IX | 210923 | 212005 | -1 |
| YIL082W-A | 4094.14431 | -0.3830112 | 0.51396322 | 1.14174845 | 0.56503126 | 0.71587111 | NA | IX | 205635 | 210132 | 1 |
| YIL083C | 21.8435299 | -0.3460127 | 0.7168647 | 0.86461479 | 0.64900984 | 0.7805914 | CAB2 | IX | 203556 | 204653 | -1 |
| YIL084C | 101.777571 | -0.51816 | 0.52881631 | 1.09107942 | 0.57952892 | 0.72802613 | SDS3 | IX | 202276 | 203259 | -1 |

| | | | | | | | | | | | |
|---|---|---|---|---|---|---|---|---|---|---|---|
| YIL085C | 7.47599859 | 0.03597472 | 1.08482485 | 1.20980443 | 0.54612783 | 0.69889749 | KTR7 | IX | 200490 | 202043 | -1 |
| YIL086C | 44.3815795 | 0.7208624 | 0.77562023 | 11.0185513 | 0.00404904 | 0.01943942 | NA | IX | 200153 | 200461 | -1 |
| YIL087C | 14.9364468 | -0.7520486 | 0.80311859 | 0.98079082 | 0.6123842 | 0.7554524 | AIM19 | IX | 199646 | 200119 | -1 |
| YIL088C | 12.3482281 | 0.1004348 | 0.97582943 | 0.51251509 | 0.77394262 | 0.86281553 | AVT7 | IX | 197931 | 199403 | -1 |
| YIL089W | 353.360351 | 0.24086328 | 0.49503497 | 3.46959067 | 0.17643631 | 0.32890091 | NA | IX | 195599 | 196216 | 1 |
| YIL090W | 32.1425331 | 0.10104052 | 0.7850206 | 8.2380274 | 0.01626054 | 0.05678441 | ICE2 | IX | 193595 | 195070 | 1 |
| YIL091C | 140.303107 | 0.11085975 | 0.60646301 | 4.47783254 | 0.10657394 | 0.22939149 | UTP25 | IX | 191030 | 193195 | -1 |
| YIL092W | 107.933962 | -1.4327209 | 0.5400547 | 34.6504224 | 2.99E-08 | 8.09E-07 | NA | IX | 189066 | 190967 | 1 |
| YIL093C | 16.6518449 | -0.0934594 | 0.81825634 | 1.73739903 | 0.41949674 | 0.59032562 | RSM25 | IX | 187990 | 188784 | -1 |
| YIL094C | 25.8892758 | 0.12051262 | 0.76823301 | 4.35140314 | 0.11352848 | 0.24016437 | LYS12 | IX | 186517 | 187632 | -1 |
| YIL095W | 258.168863 | -0.2987496 | 0.65349171 | 17.9048959 | 0.00012942 | 0.0011672 | PRK1 | IX | 183937 | 186369 | 1 |
| YIL096C | 72.7100059 | 0.55043162 | 0.60839282 | 25.6982144 | 2.63E-06 | 4.09E-05 | BMT5 | IX | 182117 | 183127 | -1 |
| YIL097W | 46.3620208 | 1.1069053 | 0.75865575 | 3.61375143 | 0.16416624 | 0.31287772 | FYV10 | IX | 180427 | 181977 | 1 |
| YIL098C | 17.6222265 | 0.15296636 | 0.85666221 | 2.63218253 | 0.26818151 | 0.44109199 | FMC1 | IX | 179772 | 180239 | -1 |
| YIL099W | 34.1635665 | -1.0026656 | 0.66792471 | 11.777699 | 0.00277016 | 0.01429522 | SGA1 | IX | 178004 | 179653 | 1 |
| YIL101C | 526.385024 | -1.3310489 | 0.49947681 | 7.74628452 | 0.02079293 | 0.06797914 | XBP1 | IX | 175307 | 177250 | -1 |
| YIL102C | 59.17031 | -0.0744715 | 0.63482845 | 2.11932414 | 0.34657291 | 0.51999408 | NA | IX | 174582 | 174887 | -1 |
| YIL102C-A | 98.6561237 | 0.51416258 | 0.64616539 | 31.0571235 | 1.80E-07 | 4.03E-06 | NA | IX | 173365 | 173592 | -1 |
| YIL103W | 1753.47932 | 0.66487992 | 0.49181228 | 4.94522234 | 0.08436428 | 0.19386304 | DPH1 | IX | 171751 | 173028 | 1 |
| YIL104C | 10.4128709 | -0.9495529 | 0.94106029 | 1.40992465 | 0.49412719 | 0.65469026 | SHQ1 | IX | 169982 | 171505 | -1 |
| YIL105C | 20.421557 | -0.2108407 | 0.74420382 | 0.78194568 | 0.67639853 | 0.8007253 | SLM1 | IX | 167581 | 169641 | -1 |
| YIL105W-A | 1084.89459 | 0.67242953 | 0.47207768 | 2.43823851 | 0.2954903 | 0.46963735 | NA | IX | 167664 | 167804 | 1 |
| YIL106W | 158.090097 | 2.25977213 | 0.68783265 | 11.0648893 | 0.00395631 | 0.01908917 | MOB1 | IX | 166415 | 167444 | 1 |
| YIL107C | 959.758868 | 2.11825067 | 0.48658035 | 34.1638575 | 3.81E-08 | 1.01E-06 | PFK26 | IX | 163278 | 165761 | -1 |
| YIL108W | 92.060516 | 0.94216809 | 0.64917258 | 4.39800837 | 0.11091355 | 0.2359256 | NA | IX | 160887 | 162977 | 1 |
| YIL109C | 231.269482 | 1.58619811 | 0.64311756 | 7.07823004 | 0.02903901 | 0.08675743 | SEC24 | IX | 157385 | 160165 | -1 |
| YIL110W | 137.942905 | 1.95149549 | 0.56386378 | 27.4372308 | 1.10E-06 | 1.93E-05 | HPM1 | IX | 156045 | 157178 | 1 |
| YIL111W | 465.129622 | -0.4772054 | 0.47518204 | 7.75318179 | 0.02072135 | 0.06786006 | NA | IX | 155311 | 155765 | 1 |
| YIL112W | 1010.07033 | 1.04378719 | 0.54465629 | 16.5631939 | 0.00025313 | 0.00209676 | HOS4 | IX | 151595 | 154846 | 1 |
| YIL113W | 115.499867 | -1.2426718 | 0.54065163 | 9.15629591 | 0.01027391 | 0.04024758 | SDP1 | IX | 150559 | 151188 | 1 |
| YIL114C | 8.70018485 | -2.2655346 | 1.14749953 | 5.22372896 | 0.07339757 | 0.17466992 | POR2 | IX | 149143 | 149988 | -1 |
| YIL115C | 194.33159 | -1.4069023 | 0.4917481 | 8.20854394 | 0.01650203 | 0.05733167 | NUP159 | IX | 144327 | 148709 | -1 |
| YIL116W | 36.4921859 | 1.39774343 | 0.78480013 | 3.6281821 | 0.16298599 | 0.31175673 | HIS5 | IX | 142928 | 144085 | 1 |

| | | | | | | | | | | | |
|---|---|---|---|---|---|---|---|---|---|---|---|
| YIL117C | 61.8600166 | 2.26023251 | 0.6503446 | 11.5892287 | 0.0030439 | 0.01538156 | PRM5 | IX | 140613 | 141569 | -1 |
| YIL118W | 45.7768081 | 0.89115519 | 0.73712094 | 2.69773675 | 0.25953379 | 0.431191 | RHO3 | IX | 139752 | 140447 | 1 |
| YIL119C | 103.777618 | -2.0443781 | 0.56816307 | 44.8622051 | 1.81E-10 | 7.89E-09 | RPI1 | IX | 136654 | 137877 | -1 |
| YIL120W | 356.995797 | -0.1045944 | 0.57078661 | 0.41026827 | 0.81453805 | 0.88867068 | QDR1 | IX | 134417 | 136108 | 1 |
| YIL121W | 889.248043 | 0.03315014 | 0.5082555 | 0.29008937 | 0.86498364 | 0.92054956 | QDR2 | IX | 132244 | 133872 | 1 |
| YIL123W | 106.217052 | 1.44621873 | 0.62792786 | 5.24855978 | 0.07249194 | 0.17315525 | SIM1 | IX | 128151 | 129581 | 1 |
| YIL124W | 88.4615462 | -0.5654717 | 0.54989414 | 1.23451738 | 0.53942113 | 0.6935928 | AYR1 | IX | 126204 | 127097 | 1 |
| YIL125W | 48.1488981 | 0.41964824 | 0.65422867 | 2.1368831 | 0.3435435 | 0.51705663 | KGD1 | IX | 122689 | 125733 | 1 |
| YIL126W | 61.4675757 | 0.15195901 | 0.61916791 | 5.99806847 | 0.04983517 | 0.13021013 | STH1 | IX | 117992 | 122071 | 1 |
| YIL127C | 65.3755146 | 1.32815562 | 0.86167333 | 14.2155881 | 0.0008187 | 0.00559654 | RRT14 | IX | 117024 | 117644 | -1 |
| YIL128W | 318.993074 | -1.6953429 | 0.48909875 | 37.2261647 | 8.25E-09 | 2.53E-07 | MET18 | IX | 113806 | 116904 | 1 |
| YIL129C | 32.6923415 | -0.910623 | 0.74077876 | 2.4199603 | 0.2982032 | 0.47161883 | TAO3 | IX | 106107 | 113237 | -1 |
| YIL130W | 12.7734188 | -0.2707989 | 0.88682292 | 3.73992475 | 0.15412946 | 0.2985646 | ASG1 | IX | 102782 | 105676 | 1 |
| YIL131C | 31.9679858 | -0.3147411 | 0.67820452 | 4.334338 | 0.11450131 | 0.24134059 | FKH1 | IX | 100781 | 102235 | -1 |
| YIL132C | 17.529885 | -0.5623378 | 0.78563603 | 0.71724106 | 0.69863941 | 0.81736153 | CSM2 | IX | 99860 | 100501 | -1 |
| YIL133C | 40.4674163 | -0.4537303 | 0.73278429 | 10.4067899 | 0.00549787 | 0.02469562 | RPL16A | IX | 98527 | 99416 | -1 |
| YIL134W | 210.811528 | 0.66514206 | 0.58473248 | 1.31801991 | 0.51736329 | 0.67406244 | FLX1 | IX | 97395 | 98330 | 1 |
| YIL135C | 21.5076024 | 0.25168172 | 0.74624804 | 0.18274178 | 0.91267914 | 0.94617945 | VHS2 | IX | 95065 | 96375 | -1 |
| YIL136W | 80.5121934 | -0.3618443 | 0.5907115 | 1.297455 | 0.5227105 | 0.67858605 | OM45 | IX | 93619 | 94800 | 1 |
| YIL137C | 88.4740929 | 0.31594392 | 0.5891276 | 7.5809924 | 0.02258439 | 0.07260613 | TMA108 | IX | 89948 | 92788 | -1 |
| YIL138C | 37.5119893 | -0.1024088 | 0.60758082 | 1.99464457 | 0.36886584 | 0.54206426 | TPM2 | IX | 89230 | 89715 | -1 |
| YIL139C | 126.074958 | -0.0458354 | 0.55605064 | 6.30422625 | 0.04276167 | 0.11662274 | REV7 | IX | 87979 | 88716 | -1 |
| YIL140W | 93.5006198 | -0.0560556 | 0.59518219 | 3.81727352 | 0.14828239 | 0.29064152 | AXL2 | IX | 85366 | 87837 | 1 |
| YIL142W | 139.671254 | 0.93860013 | 0.51599226 | 5.8885574 | 0.05264002 | 0.1358225 | CCT2 | IX | 83302 | 84885 | 1 |
| YIL143C | 50.0357103 | 0.45698281 | 0.72592717 | 5.21948106 | 0.07355363 | 0.17489753 | SSL2 | IX | 80510 | 83041 | -1 |
| YIL144W | 19.4739474 | 1.57687914 | 0.81198098 | 4.49446626 | 0.10569125 | 0.22827341 | NDC80 | IX | 78074 | 80149 | 1 |
| YIL145C | 382.653333 | 1.43233561 | 0.50023969 | 28.1657094 | 7.65E-07 | 1.42E-05 | PAN6 | IX | 76354 | 77283 | -1 |
| YIL146C | 56.2396872 | -1.2579471 | 0.58874186 | 6.82579367 | 0.03294562 | 0.09600159 | ATG32 | IX | 74184 | 75773 | -1 |
| YIL147C | 58.7196971 | -0.6170636 | 0.55769212 | 2.39997139 | 0.30119852 | 0.47428323 | SLN1 | IX | 69791 | 73453 | -1 |
| YIL148W | 161.166612 | 1.34571156 | 0.72204848 | 12.6791619 | 0.00176504 | 0.01019919 | RPL40B | IX | 68708 | 69528 | 1 |
| YIL149C | 39.8050834 | -0.1175921 | 0.6235311 | 6.57964058 | 0.03726054 | 0.10539255 | MLP2 | IX | 63028 | 68067 | -1 |
| YIL150C | 31.8215211 | -0.5753348 | 0.71697219 | 10.0456487 | 0.0065859 | 0.02854218 | MCM10 | IX | 61013 | 62728 | -1 |
| YIL151C | 94.708406 | 0.88607991 | 0.61761014 | 2.54183916 | 0.28057349 | 0.45466569 | ESL1 | IX | 57338 | 60694 | -1 |

| | | | | | | | | | | | |
|---|---|---|---|---|---|---|---|---|---|---|---|
| YIL152W | 87.7985914 | -0.4167958 | 0.57591671 | 1.57249344 | 0.4555514 | 0.62135279 | NA | IX | 56545 | 57252 | 1 |
| YIL153W | 161.935523 | 0.73728313 | 0.51957886 | 15.7042613 | 0.00038892 | 0.00299848 | RRD1 | IX | 55198 | 56379 | 1 |
| YIL154C | 210.887786 | 0.63529146 | 0.56074371 | 1.5089576 | 0.47025565 | 0.63276324 | IMP2' | IX | 53981 | 55021 | -1 |
| YIL155C | 87.0775446 | -0.236325 | 0.5441153 | 1.29655388 | 0.52294607 | 0.67871828 | GUT2 | IX | 51759 | 53708 | -1 |
| YIL156W | 15.9377886 | 0.60403105 | 0.81560921 | 0.68003197 | 0.71175894 | 0.82438173 | UBP7 | IX | 48091 | 51306 | 1 |
| YIL157C | 23.8254105 | -0.8551458 | 0.72875934 | 2.52304756 | 0.28322213 | 0.45678444 | COA1 | IX | 46949 | 47542 | -1 |
| YIL158W | 438.192242 | 1.38467545 | 0.77472393 | 6.36634841 | 0.04145386 | 0.11428804 | AIM20 | IX | 46201 | 46815 | 1 |
| YIL159W | 162.441967 | 1.61345577 | 0.59684337 | 42.8138377 | 5.05E-10 | 1.95E-08 | BNR1 | IX | 41825 | 45952 | 1 |
| YIL160C | 509.511224 | -0.7168789 | 0.48932287 | 10.6030116 | 0.00498408 | 0.02284865 | POT1 | IX | 40191 | 41444 | -1 |
| YIL161W | 131.172184 | 0.40534964 | 0.51788582 | 2.33184935 | 0.31163437 | 0.48405923 | NA | IX | 39433 | 40140 | 1 |
| YIL162W | 7.85473547 | -0.982644 | 1.15375478 | 0.95394312 | 0.62066018 | 0.76119942 | SUC2 | IX | 37385 | 38983 | 1 |
| YIL163C | 303.459563 | -0.4350969 | 0.55617528 | 0.93293295 | 0.62721464 | 0.76483005 | NA | IX | 36899 | 37252 | -1 |
| YIL166C | 525.238228 | -0.0479287 | 0.54879399 | 0.20702628 | 0.90166418 | 0.93824416 | NA | IX | 30938 | 32566 | -1 |
| YIL167W | 26.0132288 | -0.8228878 | 0.79070525 | 12.1598884 | 0.0022883 | 0.01234789 | NA | NA | NA | NA | NA |
| YIL169C | 5707.65919 | -1.6166987 | 0.47718899 | 15.7156026 | 0.00038672 | 0.00298949 | CSS1 | IX | 23119 | 26106 | -1 |
| YIL170W | 586.665378 | 1.23076747 | 0.48308615 | 11.6957789 | 0.00288598 | 0.0147744 | NA | IX | 19847 | 21220 | 1 |
| YIL171W | 117.314393 | 1.06975298 | 0.58191811 | 5.72935287 | 0.05700157 | 0.14462713 | NA | IX | 19515 | 19844 | 1 |
| YIR001C | 66.2537475 | 0.52612322 | 0.6084754 | 1.68508122 | 0.43061511 | 0.59948398 | SGN1 | IX | 356143 | 356895 | -1 |
| YIR002C | 979.451431 | 3.60796824 | 0.48928489 | 75.3989585 | 4.24E-17 | 6.63E-15 | MPH1 | IX | 357415 | 360396 | -1 |
| YIR003W | 94.583085 | 0.21357027 | 0.53472298 | 0.16522649 | 0.92070717 | 0.95024857 | AIM21 | IX | 360885 | 362924 | 1 |
| YIR004W | 44.5477632 | 0.952722 | 0.73002462 | 2.59556725 | 0.2731365 | 0.44661404 | DJP1 | IX | 363221 | 364519 | 1 |
| YIR005W | 62.942193 | 0.63365991 | 0.61371296 | 1.1165271 | 0.5722018 | 0.72242661 | IST3 | IX | 364889 | 365335 | 1 |
| YIR006C | 58.2337649 | 0.70514587 | 0.59406452 | 1.52519407 | 0.46645346 | 0.62925571 | PAN1 | IX | 365466 | 369908 | -1 |
| YIR007W | 112.677182 | 0.61205197 | 0.68311181 | 4.05286323 | 0.13180501 | 0.26711622 | EGH1 | IX | 370704 | 372998 | 1 |
| YIR008C | 88.8782071 | 1.44776686 | 0.7498733 | 3.6727254 | 0.15939614 | 0.30640892 | PRI1 | IX | 373077 | 374306 | -1 |
| YIR009W | 29.8973564 | -1.1336816 | 0.67234082 | 10.9969008 | 0.00409311 | 0.0195435 | MSL1 | IX | 374525 | 374860 | 1 |
| YIR010W | 26.1677564 | -0.6194918 | 0.68832916 | 0.81335788 | 0.66585794 | 0.79245991 | DSN1 | IX | 375431 | 377161 | 1 |
| YIR011C | 35.2724218 | 0.88287512 | 0.71804521 | 4.63002577 | 0.09876491 | 0.21726779 | STS1 | IX | 377287 | 378246 | -1 |
| YIR012W | 37.9083094 | -0.4384884 | 0.6371006 | 4.56978747 | 0.10178488 | 0.22205518 | SQT1 | IX | 378486 | 379781 | 1 |
| YIR013C | 13.4716366 | -1.2730353 | 0.94194126 | 2.04907247 | 0.3589629 | 0.53196703 | GAT4 | IX | 380019 | 380384 | -1 |
| YIR014W | 21.0448467 | -1.0406761 | 0.73022527 | 4.37026477 | 0.11246284 | 0.23882864 | NA | IX | 381086 | 381814 | 1 |
| YIR015W | 16.2793808 | 0.60190088 | 0.79313069 | 0.71102279 | 0.70081496 | 0.81801018 | RPR2 | IX | 381948 | 382382 | 1 |
| YIR016W | 94.0653152 | 2.13207627 | 0.84789391 | 6.18391803 | 0.0454129 | 0.12167548 | NA | IX | 382628 | 383425 | 1 |

| | | | | | | | | | | | |
|---|---|---|---|---|---|---|---|---|---|---|---|
| YIR017C | 17.8101895 | 0.68885961 | 0.81457849 | 2.40652205 | 0.30021361 | 0.47337604 | MET28 | IX | 383556 | 384119 | -1 |
| YIR018C-A | 341.765649 | 0.00417653 | 0.58861926 | 0.88836351 | 0.64134884 | 0.77591986 | NA | IX | 385564 | 385701 | -1 |
| YIR018W | 23.1441236 | -0.0501035 | 0.72145462 | 1.46049248 | 0.48179034 | 0.64424159 | YAP5 | IX | 384609 | 385346 | 1 |
| YIR019C | 75.1340145 | -0.1176651 | 0.60824375 | 3.50139394 | 0.17365287 | 0.32507061 | FLO11 | IX | 389572 | 393675 | -1 |
| YIR021W | 21.1404179 | -0.1794129 | 0.84270591 | 9.86885262 | 0.00719459 | 0.03067501 | MRS1 | IX | 397294 | 398385 | 1 |
| YIR021W-A | 11.8945348 | -0.479646 | 0.91874038 | 1.06985021 | 0.58571316 | 0.73372548 | NA | IX | 398514 | 398726 | 1 |
| YIR022W | 47.5760305 | -1.3074174 | 0.76872105 | 2.91480899 | 0.23283983 | 0.40027153 | SEC11 | IX | 398733 | 399236 | 1 |
| YIR023W | 376.994198 | 0.55439518 | 0.62312923 | 49.9561545 | 1.42E-11 | 8.06E-10 | DAL81 | IX | 399777 | 402689 | 1 |
| YIR024C | 12.3374953 | -0.0501427 | 0.92103105 | 1.4876267 | 0.47529798 | 0.63776948 | INA22 | IX | 402841 | 403491 | -1 |
| YIR025W | 44.6138112 | -0.706183 | 0.72308042 | 5.92009829 | 0.05181637 | 0.1343428 | MND2 | IX | 403659 | 404765 | 1 |
| YIR026C | 169.640587 | -0.1593822 | 0.52262148 | 10.094684 | 0.00642639 | 0.02793454 | YVH1 | IX | 404873 | 405967 | -1 |
| YIR027C | 407.679798 | 0.55366869 | 0.5556207 | 11.0388309 | 0.00400819 | 0.01927527 | DAL1 | IX | 406260 | 407642 | -1 |
| YIR028W | 345.395778 | 0.51276422 | 0.5794256 | 0.78234082 | 0.67626491 | 0.8007253 | DAL4 | IX | 408468 | 410375 | 1 |
| YIR029W | 23.5433017 | -1.1544879 | 0.82059135 | 4.17747354 | 0.12384348 | 0.25554303 | DAL2 | IX | 410807 | 411838 | 1 |
| YIR030C | 176.302789 | -0.4337637 | 0.58300416 | 8.64207128 | 0.01328612 | 0.04868773 | DCG1 | IX | 412036 | 412770 | -1 |
| YIR031C | 15.3585968 | 0.54459095 | 0.89882285 | 0.48233367 | 0.78571053 | 0.87017291 | DAL7 | IX | 413015 | 414679 | -1 |
| YIR032C | 25.3673057 | -0.5947402 | 0.77630481 | 0.60606286 | 0.73857588 | 0.84040603 | DAL3 | IX | 415030 | 415617 | -1 |
| YIR033W | 58.2176227 | 0.41683293 | 0.60244207 | 1.37743943 | 0.50221864 | 0.66087407 | MGA2 | IX | 416124 | 419465 | 1 |
| YIR034C | 73.9418615 | -0.0946839 | 0.6267324 | 0.101853 | 0.95034852 | 0.96603194 | LYS1 | IX | 419615 | 420736 | -1 |
| YIR035C | 99.1665315 | -0.8834608 | 0.54638736 | 2.79191074 | 0.24759638 | 0.41832973 | NA | IX | 421026 | 421790 | -1 |
| YIR036C | 21.6222898 | -0.3913928 | 0.72999714 | 0.54938805 | 0.75980457 | 0.85451323 | IRC24 | IX | 422074 | 422865 | -1 |
| YIR037W | 18.8947794 | 1.15070322 | 0.85704721 | 2.1326766 | 0.34426681 | 0.51781024 | HYR1 | IX | 423128 | 423619 | 1 |
| YIR038C | 10.2064656 | 1.44439764 | 0.98028013 | 2.28099663 | 0.31965969 | 0.49223979 | GTT1 | IX | 423809 | 424513 | -1 |
| YIR039C | 14.0736163 | -0.5930128 | 0.8753992 | 0.54192172 | 0.76264635 | 0.85565759 | YPS6 | IX | 430498 | 432111 | -1 |
| YIR042C | 44.5687664 | -1.4947533 | 0.68156719 | 6.87430367 | 0.03215614 | 0.0942798 | NA | IX | 435273 | 435983 | -1 |
| YIR043C | 42.2360246 | -1.4922051 | 0.62928214 | 16.5892075 | 0.00024986 | 0.00207561 | NA | NA | NA | NA | NA |
| YJL001W | 115.938678 | 0.17956983 | 0.5709597 | 4.78764529 | 0.09128008 | 0.205327 | PRE3 | X | 435163 | 435926 | 1 |
| YJL002C | 46.018459 | 1.21599036 | 0.7240474 | 6.70389695 | 0.03501606 | 0.10046729 | OST1 | X | 433221 | 434651 | -1 |
| YJL003W | 37.4157626 | -0.2175729 | 0.6688344 | 2.65283363 | 0.26542663 | 0.43759117 | COX16 | X | 432644 | 433000 | 1 |
| YJL004C | 61.4410743 | 0.2957869 | 0.61508251 | 4.22768226 | 0.12077317 | 0.25039546 | SYS1 | X | 431589 | 432200 | -1 |
| YJL005W | 26.4296549 | 0.72727708 | 0.76948711 | 2.32575589 | 0.31258528 | 0.4849178 | CYR1 | X | 425157 | 431237 | 1 |
| YJL007C | 46.7306776 | -0.5648469 | 0.69491799 | 1.58783239 | 0.45207093 | 0.61810237 | NA | X | 422698 | 423012 | -1 |
| YJL008C | 36.1541747 | -0.5867267 | 0.8040247 | 6.3739102 | 0.04129743 | 0.1140258 | CCT8 | X | 419957 | 421663 | -1 |

| | | | | | | | | | | | |
|---|---|---|---|---|---|---|---|---|---|---|---|
| YJL010C | 21.8408143 | -0.9761559 | 0.80337783 | 8.95350578 | 0.01137027 | 0.04332089 | NOP9 | X | 417562 | 419562 | -1 |
| YJL012C | 20.0181818 | -0.5687322 | 0.77880695 | 0.93870687 | 0.6254065 | 0.76426839 | VTC4 | X | 411234 | 413399 | -1 |
| YJL013C | 39.5148822 | 0.19859761 | 0.70734196 | 7.88786662 | 0.01937187 | 0.06459254 | MAD3 | X | 409493 | 411040 | -1 |
| YJL014W | 137.611957 | -0.583682 | 0.79558824 | 5.17551942 | 0.0751883 | 0.17776245 | CCT3 | X | 407558 | 409162 | 1 |
| YJL016W | 250.047478 | 0.45801492 | 0.52938003 | 46.8523351 | 6.70E-11 | 3.29E-09 | TPH3 | X | 405588 | 407273 | 1 |
| YJL019W | 145.717939 | -0.7121947 | 0.58470326 | 2.30344593 | 0.31609169 | 0.48830599 | MPS3 | X | 402897 | 404945 | 1 |
| YJL020C | 143.271949 | 1.54477608 | 0.54194586 | 9.08180985 | 0.01066375 | 0.04137474 | BBC1 | X | 398937 | 402410 | -1 |
| YJL023C | 82.7049345 | -0.0977365 | 0.61966617 | 10.1916279 | 0.00612232 | 0.02692266 | PET130 | X | 397355 | 398398 | -1 |
| YJL024C | 144.642944 | -0.5732499 | 0.52812925 | 6.16563493 | 0.04582995 | 0.122171 | APS3 | X | 395931 | 396592 | -1 |
| YJL025W | 36.4432925 | -0.7778019 | 0.68256725 | 9.68434485 | 0.0078899 | 0.03284148 | RRN7 | X | 393967 | 395511 | 1 |
| YJL026W | 59.6352509 | -1.6339588 | 0.64690934 | 10.7025151 | 0.00474218 | 0.02193071 | RNR2 | X | 392404 | 393603 | 1 |
| YJL027C | 75.8158859 | -0.5139031 | 0.74326686 | 1.60377847 | 0.44848088 | 0.61504128 | NA | X | 391833 | 392249 | -1 |
| YJL028W | 8.78339306 | -1.0351773 | 0.98776842 | 3.83587822 | 0.14690941 | 0.28883039 | NA | X | 391311 | 391646 | 1 |
| YJL029C | 292.74313 | 0.3203092 | 0.4900642 | 2.18681187 | 0.33507331 | 0.50840526 | VPS53 | X | 388385 | 390853 | -1 |
| YJL030W | 24.1620609 | 0.36974609 | 0.86261643 | 4.95509791 | 0.08394874 | 0.19311211 | MAD2 | X | 387657 | 388247 | 1 |
| YJL031C | 16.0418051 | 0.74757803 | 0.86188276 | 0.84388666 | 0.6557712 | 0.78495521 | BET4 | X | 386368 | 387438 | -1 |
| YJL033W | 21.432247 | -0.9725134 | 0.74683854 | 4.78641623 | 0.0913362 | 0.20537343 | HCA4 | X | 383837 | 386149 | 1 |
| YJL034W | 321.984392 | 2.07707831 | 0.76090936 | 8.1617441 | 0.01689273 | 0.05839337 | KAR2 | X | 381327 | 383375 | 1 |
| YJL035C | 331.42487 | 2.44540086 | 0.49366273 | 30.043965 | 2.99E-07 | 6.26E-06 | TAD2 | X | 380249 | 381001 | -1 |
| YJL036W | 18.0479 | 1.96583964 | 0.96001305 | 4.24515929 | 0.11972239 | 0.2489022 | SNX4 | X | 378825 | 380096 | 1 |
| YJL037W | 48.9346594 | -0.3236525 | 0.71762413 | 10.8279454 | 0.00445391 | 0.02079165 | IRC18 | X | 376662 | 377336 | 1 |
| YJL038C | 664.986833 | 0.41972195 | 0.46129975 | 13.0953425 | 0.00143345 | 0.00883884 | LOH1 | X | 375115 | 375774 | -1 |
| YJL039C | 490.139535 | 1.08525673 | 0.59951828 | 97.3209389 | 7.36E-22 | 3.28E-19 | NUP192 | X | 368748 | 373799 | -1 |
| YJL041W | 506.202545 | 2.6051568 | 0.73358406 | 22.8255312 | 1.11E-05 | 0.00014382 | NA | X | 365903 | 368373 | 1 |
| YJL042W | 314.414421 | -1.2694239 | 0.49495528 | 7.6635744 | 0.02167085 | 0.07033309 | MHP1 | X | 361249 | 365445 | 1 |
| YJL043W | 16.761287 | -0.5824068 | 0.86211913 | 0.88660416 | 0.64191327 | 0.77608641 | NA | X | 360130 | 360903 | 1 |
| YJL044C | 33.7259217 | -1.4385321 | 0.65840473 | 8.92349507 | 0.01154218 | 0.04387997 | GYP6 | X | 358300 | 359676 | -1 |
| YJL045W | 152.280364 | -0.4569758 | 0.55191578 | 2.15983426 | 0.33962367 | 0.5126228 | NA | X | 356024 | 357928 | 1 |
| YJL046W | 8.34702949 | -0.8103002 | 1.02027044 | 0.67606504 | 0.7131721 | 0.82535807 | AIM22 | X | 352386 | 353615 | 1 |
| YJL047C | 49.1046436 | -0.5304823 | 0.59202974 | 0.90376144 | 0.63643008 | 0.77284609 | RTT101 | X | 349580 | 352108 | -1 |
| YJL048C | 66.387266 | 0.33445168 | 0.81436805 | 0.91886058 | 0.6316434 | 0.76880708 | UBX6 | X | 347447 | 348637 | -1 |
| YJL049W | 418.89263 | 3.64740184 | 1.00964178 | 10.67503 | 0.0048078 | 0.02214573 | CHM7 | X | 345973 | 347325 | 1 |
| YJL050W | 50.9237924 | -1.0040562 | 0.62983477 | 2.81448817 | 0.24481705 | 0.41519939 | MTR4 | X | 342522 | 345743 | 1 |

| | | | | | | | | | | | |
|---|---|---|---|---|---|---|---|---|---|---|---|
| YJL051W | 803.453439 | -1.2143727 | 0.50610059 | 10.1304875 | 0.00631237 | 0.02756307 | IRC8 | X | 339787 | 342255 | 1 |
| YJL052C-A | 19.011907 | 0.96822515 | 0.86667121 | 7.79624826 | 0.02027992 | 0.06680024 | NA | X | 337884 | 338003 | -1 |
| YJL052W | 12.2452933 | -1.4068831 | 1.07758471 | 3.66438097 | 0.16006257 | 0.30728192 | TDH1 | X | 338271 | 339269 | 1 |
| YJL053W | 52.5839467 | -0.2175897 | 0.59627096 | 7.40847655 | 0.02461896 | 0.07694787 | PEP8 | X | 335898 | 337037 | 1 |
| YJL054W | 42.9404315 | -0.0767907 | 0.60385041 | 1.09506594 | 0.57837492 | 0.72736551 | TIM54 | X | 334265 | 335701 | 1 |
| YJL055W | 48.9284572 | -0.0953203 | 0.59268511 | 6.22678324 | 0.04444994 | 0.11984737 | NA | X | 333357 | 334094 | 1 |
| YJL056C | 29.8260311 | -0.3137228 | 0.84426636 | 2.58457194 | 0.27464224 | 0.44819013 | ZAP1 | X | 330431 | 333073 | -1 |
| YJL057C | 120.154538 | 0.65606232 | 0.58258781 | 1.9319232 | 0.38061702 | 0.55440819 | IKS1 | X | 328118 | 330121 | -1 |
| YJL058C | 172.414194 | 0.94902181 | 0.49821717 | 12.319714 | 0.00211256 | 0.01164923 | BIT61 | X | 326242 | 327873 | -1 |
| YJL059W | 156.551187 | 0.40102005 | 0.53298289 | 0.62100822 | 0.73307731 | 0.83751334 | YHC3 | X | 324964 | 326190 | 1 |
| YJL060W | 128.837948 | -1.5120397 | 0.6406673 | 19.0622921 | 7.26E-05 | 0.00071568 | BNA3 | X | 323386 | 324720 | 1 |
| YJL061W | 25.7713721 | 0.52548796 | 0.70532078 | 0.68948069 | 0.70840427 | 0.82395756 | NUP82 | X | 320016 | 322157 | 1 |
| YJL062W | 238.909954 | -0.4899395 | 0.47919313 | 1.54088199 | 0.46280893 | 0.62608965 | LAS21 | X | 317284 | 319776 | 1 |
| YJL062W-A | 14.6114592 | 0.52152032 | 0.83046396 | 0.40759045 | 0.81562938 | 0.88902268 | COA3 | X | 316723 | 316980 | 1 |
| YJL063C | 36.4157979 | -2.1837364 | 0.69186972 | 19.7060503 | 5.26E-05 | 0.00053606 | MRPL8 | X | 315759 | 316475 | -1 |
| YJL065C | 11.7342153 | 1.17551474 | 0.93719317 | 1.86346629 | 0.39387048 | 0.5656027 | DLS1 | X | 315054 | 315557 | -1 |
| YJL066C | 13.2092808 | 0.60654443 | 0.86609012 | 0.5005859 | 0.77857267 | 0.86541289 | MPM1 | X | 314114 | 314872 | -1 |
| YJL068C | 42.5573331 | 0.18052326 | 0.59959812 | 0.17398411 | 0.91668438 | 0.94860224 | NA | X | 313016 | 313915 | -1 |
| YJL069C | 308.348865 | 1.62370502 | 0.55174247 | 25.9908689 | 2.27E-06 | 3.59E-05 | UTP18 | X | 310922 | 312706 | -1 |
| YJL070C | 84.6267711 | -0.0324895 | 0.53601672 | 2.23160323 | 0.32765252 | 0.50039829 | NA | X | 307971 | 310637 | -1 |
| YJL071W | 29.9092487 | 0.36795766 | 0.69054233 | 3.76511262 | 0.15220054 | 0.29581776 | ARG2 | X | 306132 | 307856 | 1 |
| YJL072C | 141.767339 | 2.29189467 | 0.74954368 | 9.88411235 | 0.0071399 | 0.03048675 | PSF2 | X | 305221 | 305862 | -1 |
| YJL073W | 466.03553 | -0.5204114 | 0.49480377 | 13.3646973 | 0.00125283 | 0.00791047 | JEM1 | X | 303181 | 305118 | 1 |
| YJL074C | 21.6632617 | 0.28862164 | 0.72181587 | 0.1679756 | 0.91944247 | 0.94974417 | SMC3 | X | 299157 | 302849 | -1 |
| YJL075C | 1961.2185 | -0.351742 | 0.45370634 | 4.52268889 | 0.10421029 | 0.22564606 | NA | X | 298460 | 298876 | -1 |
| YJL076W | 90.1237899 | -0.2459113 | 0.55643259 | 9.32619583 | 0.00943718 | 0.03773943 | NET1 | X | 295245 | 298814 | 1 |
| YJL077C | 1000.08439 | -0.1170711 | 0.4620126 | 0.61264547 | 0.736149 | 0.83936643 | ICS3 | X | 294666 | 295061 | -1 |
| YJL077W-A | 14.1315864 | -0.5523295 | 0.96423121 | 0.38895665 | 0.82326402 | 0.8939795 | NA | X | 294799 | 294885 | 1 |
| YJL077W-B | 105.047676 | -2.3317084 | 0.59147936 | 43.6618952 | 3.30E-10 | 1.36E-08 | NA | X | 294047 | 294145 | 1 |
| YJL078C | 3529.42512 | -0.5748277 | 0.49066596 | 3.25159832 | 0.19675437 | 0.35456204 | PRY3 | X | 291336 | 293981 | -1 |
| YJL079C | 126.311695 | -0.7188002 | 0.60661572 | 3.05461629 | 0.21711934 | 0.38117676 | PRY1 | X | 289875 | 290774 | -1 |
| YJL080C | 91.4973251 | -0.1994883 | 0.64874583 | 2.55648357 | 0.27852658 | 0.45301284 | SCP160 | X | 285558 | 289226 | -1 |
| YJL081C | 46.6140875 | 0.89175822 | 0.5990125 | 3.32806383 | 0.1893739 | 0.34544902 | ARP4 | X | 283802 | 285271 | -1 |

| | | | | | | | | | | | |
|---|---|---|---|---|---|---|---|---|---|---|---|
| YJL082W | 63.3023852 | 0.46238915 | 0.64513208 | 1.07443565 | 0.58437182 | 0.73236208 | IML2 | X | 281185 | 283380 | 1 |
| YJL083W | 134.580051 | -1.3782312 | 0.58544162 | 7.32610439 | 0.02565409 | 0.07917761 | TAX4 | X | 278841 | 280655 | 1 |
| YJL084C | 45.5815958 | 0.05139298 | 0.61828938 | 0.69936357 | 0.70491237 | 0.82055541 | ALY2 | X | 274862 | 278002 | -1 |
| YJL085W | 16.3586046 | 1.37704052 | 0.917202 | 7.30871628 | 0.0258781 | 0.07969905 | EXO70 | X | 272827 | 274698 | 1 |
| YJL087C | 19.4887545 | 0.11055167 | 0.79147993 | 3.4193827 | 0.18092163 | 0.33553116 | TRL1 | X | 270002 | 272485 | -1 |
| YJL088W | 29.5625519 | 3.31775409 | 1.13912514 | 10.9407011 | 0.00420976 | 0.01997909 | ARG3 | X | 268799 | 269815 | 1 |
| YJL089W | 140.498629 | 2.03549474 | 0.54143474 | 21.5596467 | 2.08E-05 | 0.00024546 | SIP4 | X | 265926 | 268415 | 1 |
| YJL090C | 49.9696471 | 0.31917017 | 0.60445439 | 1.54808644 | 0.46114478 | 0.62486036 | DPB11 | X | 262757 | 265051 | -1 |
| YJL091C | 125.97909 | -0.3497504 | 0.67415936 | 0.58884017 | 0.74496348 | 0.84442806 | GWT1 | X | 261080 | 262552 | -1 |
| YJL092W | 914.205947 | 0.92983148 | 0.47539218 | 6.4564698 | 0.03962738 | 0.11084181 | SRS2 | X | 257423 | 260947 | 1 |
| YJL093C | 27.5423638 | 0.34004891 | 0.77207678 | 6.3198085 | 0.0424298 | 0.11599082 | TOK1 | X | 254737 | 256812 | -1 |
| YJL094C | 170.346014 | 0.28365419 | 0.61058026 | 50.9243528 | 8.75E-12 | 5.32E-10 | KHA1 | X | 251821 | 254442 | -1 |
| YJL095W | 307.087897 | 1.22015327 | 0.62048482 | 4.10221692 | 0.12859229 | 0.2622576 | BCK1 | X | 247255 | 251691 | 1 |
| YJL096W | 6.42874269 | 1.02068068 | 1.20259353 | 1.19734257 | 0.54954134 | 0.7014648 | MRPL49 | X | 246490 | 246975 | 1 |
| YJL097W | 9.3053542 | -0.5225731 | 1.03189294 | 2.34047735 | 0.31029287 | 0.4830878 | PHS1 | X | 245591 | 246244 | 1 |
| YJL098W | 24.2571793 | -0.5180348 | 0.71598012 | 2.78498029 | 0.24845584 | 0.41928281 | SAP185 | X | 242082 | 245258 | 1 |
| YJL099W | 590.302935 | 2.70163427 | 0.51900633 | 64.2673809 | 1.11E-14 | 1.05E-12 | CHS6 | X | 239414 | 241654 | 1 |
| YJL100W | 1942.63559 | 3.4759764 | 0.46563053 | 66.1070287 | 4.42E-15 | 4.57E-13 | LSB6 | X | 237263 | 239086 | 1 |
| YJL101C | 196.975152 | 0.51162284 | 0.65566852 | 59.1352219 | 1.44E-13 | 1.14E-11 | GSH1 | X | 234320 | 236356 | -1 |
| YJL102W | 15.7476461 | 0.09601093 | 0.82126083 | 1.84875122 | 0.39677909 | 0.56785755 | MEF2 | X | 231301 | 233760 | 1 |
| YJL103C | 126.695169 | -0.8154054 | 0.52317762 | 2.43076461 | 0.2965966 | 0.47066158 | GSM1 | X | 229025 | 230881 | -1 |
| YJL104W | 16.9735945 | 0.42652371 | 0.8294764 | 0.69838158 | 0.70525856 | 0.82079338 | PAM16 | X | 227327 | 227776 | 1 |
| YJL105W | 796.982847 | -0.6450819 | 0.51598485 | 15.1261936 | 0.00051926 | 0.00378167 | SET4 | X | 225055 | 226737 | 1 |
| YJL106W | 44.1926295 | -2.503311 | 0.68164504 | 15.2307763 | 0.00049281 | 0.00363486 | IME2 | X | 221390 | 223327 | 1 |
| YJL107C | 50.6835132 | -0.8543559 | 0.62374495 | 8.24532493 | 0.01620132 | 0.05674873 | NA | X | 218853 | 220016 | -1 |
| YJL108C | 47.9525563 | -2.6734636 | 0.70802553 | 40.0268301 | 2.03E-09 | 7.09E-08 | PRM10 | X | 217705 | 218856 | -1 |
| YJL109C | 923.89909 | 3.30371085 | 0.52173784 | 62.2090364 | 3.10E-14 | 2.76E-12 | UTP10 | X | 212000 | 217309 | -1 |
| YJL110C | 8.33835741 | -0.0109694 | 1.19991687 | 3.58970704 | 0.16615179 | 0.31541602 | GZF3 | X | 209922 | 211577 | -1 |
| YJL111W | 35.6748238 | -0.6920727 | 0.8057462 | 0.85612498 | 0.65177068 | 0.78244649 | CCT7 | X | 207877 | 209529 | 1 |
| YJL112W | 27.4963476 | 0.10753466 | 0.70550398 | 3.91099448 | 0.1414941 | 0.28066148 | MDV1 | X | 205305 | 207449 | 1 |
| YJL114W | 12.6831236 | -0.6739548 | 0.98424088 | 4.42014546 | 0.10969267 | 0.23410268 | NA | X | 197915 | 198595 | 1 |
| YJL115W | 168.211743 | 0.61852184 | 0.64317956 | 1.33513183 | 0.51295564 | 0.66997815 | ASF1 | X | 196287 | 197126 | 1 |
| YJL116C | 358.539493 | -0.1201295 | 0.49216709 | 5.90330838 | 0.0522532 | 0.13512551 | NCA3 | X | 193860 | 194873 | -1 |

| | | | | | | | | | | | | |
|---|---|---|---|---|---|---|---|---|---|---|---|---|
| YJL117W | 154.771384 | 0.99736939 | 0.53479891 | 24.802001 | 4.11E-06 | 6.05E-05 | PHO86 | X | | 192531 | 193466 | 1 |
| YJL118W | 212.86262 | -0.0523432 | 0.57484064 | 0.24048444 | 0.88670563 | 0.93215081 | NA | X | | 191639 | 192298 | 1 |
| YJL121C | 74.8531822 | -0.897886 | 0.65307339 | 14.1297222 | 0.00085461 | 0.00578738 | RPE1 | X | | 190374 | 191090 | -1 |
| YJL122W | 115.710313 | -1.0082988 | 0.54389444 | 24.2329662 | 5.47E-06 | 7.70E-05 | ALB1 | X | | 189716 | 190243 | 1 |
| YJL123C | 263.422156 | 0.32967287 | 0.57918891 | 1.2165024 | 0.54430191 | 0.69728958 | MTC1 | X | | 188004 | 189440 | -1 |
| YJL124C | 107.656394 | -0.3135599 | 0.5447164 | 4.89231889 | 0.08662564 | 0.19825705 | LSM1 | X | | 187126 | 187644 | -1 |
| YJL125C | 185.626203 | 2.66971262 | 0.98607942 | 7.42584858 | 0.02440605 | 0.07671608 | GCD14 | X | | 185528 | 186679 | -1 |
| YJL126W | 13.3042666 | 0.1664878 | 0.88579179 | 0.08160692 | 0.96001779 | 0.97173192 | NIT2 | X | | 184501 | 185424 | 1 |
| YJL127C | 107.303658 | 1.34618084 | 0.57062872 | 10.8119676 | 0.00448964 | 0.02089629 | SPT10 | X | | 182298 | 184220 | -1 |
| YJL127C-B | 171.483754 | 3.38323807 | 0.60317332 | 47.0813539 | 5.98E-11 | 2.96E-09 | NA | X | | 181551 | 181709 | -1 |
| YJL128C | 56.8419471 | 0.72525684 | 0.58508022 | 2.53534907 | 0.28148545 | 0.45563342 | PBS2 | X | | 178097 | 180103 | -1 |
| YJL129C | 255.307575 | 2.43228599 | 0.57109687 | 20.7303881 | 3.15E-05 | 0.00035018 | TRK1 | X | | 173599 | 177306 | -1 |
| YJL130C | 98.7702461 | -0.2030645 | 0.57159045 | 2.10998122 | 0.3481957 | 0.52188793 | URA2 | X | | 165723 | 172367 | -1 |
| YJL131C | 269.370531 | 0.855529 | 0.63292475 | 1.83787645 | 0.3989424 | 0.56977714 | AIM23 | X | | 164278 | 165348 | -1 |
| YJL132W | 103.149947 | -0.0156241 | 0.53436179 | 0.20468453 | 0.90272053 | 0.93871262 | NA | X | | 161914 | 164166 | 1 |
| YJL133C-A | 5.93825843 | 0.38212416 | 1.3330064 | 3.37684409 | 0.18481092 | 0.34056499 | NA | X | | 159623 | 159847 | -1 |
| YJL133W | 2397.30506 | 4.93349101 | 0.50606777 | 92.006077 | 1.05E-20 | 3.38E-18 | MRS3 | X | | 160619 | 161563 | 1 |
| YJL134W | 60.6802689 | 1.05227278 | 0.73378301 | 17.532639 | 0.0001559 | 0.00137666 | LCB3 | X | | 158188 | 159417 | 1 |
| YJL136C | 57.4074546 | -0.4296148 | 0.68359217 | 12.4158378 | 0.00201342 | 0.01120935 | RPS21B | X | | 156550 | 157273 | -1 |
| YJL136W-A | 52.8886722 | -0.0840951 | 0.63329656 | 0.06654067 | 0.96727703 | 0.97486964 | NA | X | | 156169 | 156252 | 1 |
| YJL137C | 37.2184536 | -1.3335434 | 0.72226579 | 10.4222908 | 0.00545542 | 0.02454304 | GLG2 | X | | 154985 | 156127 | -1 |
| YJL138C | 34.3960905 | -0.6303201 | 0.65301804 | 6.17772353 | 0.04555378 | 0.12191042 | TIF2 | X | | 153504 | 154691 | -1 |
| YJL139C | 88.2817351 | 0.21384632 | 0.64501335 | 0.88509985 | 0.64239627 | 0.77621415 | YUR1 | X | | 151713 | 152999 | -1 |
| YJL140W | 1353.92133 | 0.5972943 | 0.47558616 | 2.35105226 | 0.30865654 | 0.48144434 | RPB4 | X | | 150961 | 151626 | 1 |
| YJL141C | 43.3554427 | 0.17789689 | 0.64564611 | 0.80501387 | 0.6686417 | 0.79450097 | YAK1 | X | | 147967 | 150390 | -1 |
| YJL142C | 556.20442 | 0.51805922 | 0.66178265 | 13.0598025 | 0.00145915 | 0.00894319 | NA | X | | 147819 | 148211 | -1 |
| YJL143W | 50.9498734 | 0.64833404 | 0.61458574 | 2.68628708 | 0.26102384 | 0.43235173 | TIM17 | X | | 147101 | 147577 | 1 |
| YJL144W | 103.596966 | 1.66036439 | 0.60462469 | 30.8212338 | 2.03E-07 | 4.42E-06 | NA | X | | 146359 | 146673 | 1 |
| YJL145W | 460.796424 | 2.88421452 | 0.54615709 | 98.8274313 | 3.47E-22 | 1.82E-19 | SFH5 | X | | 145160 | 146044 | 1 |
| YJL146W | 478.678088 | 0.43686467 | 0.50312913 | 2.23117927 | 0.32772199 | 0.50039829 | IDS2 | X | | 143292 | 144701 | 1 |
| YJL147C | 122.233543 | -1.6727277 | 0.52220007 | 12.4507016 | 0.00197863 | 0.01106886 | MRX5 | X | | 141419 | 142567 | -1 |
| YJL148W | 81.0747874 | -1.1765973 | 0.5750101 | 24.6798362 | 4.37E-06 | 6.35E-05 | RPA34 | X | | 140437 | 141138 | 1 |
| YJL149W | 44.5713871 | -0.6337369 | 0.62563814 | 1.29068429 | 0.52448306 | 0.67982022 | DAS1 | X | | 137379 | 139370 | 1 |

| | | | | | | | | | | | |
|---|---|---|---|---|---|---|---|---|---|---|---|
| YJL151C | 582.489954 | 2.42823972 | 0.50864675 | 39.3190518 | 2.90E-09 | 9.81E-08 | SNA3 | X | 136372 | 136773 | -1 |
| YJL153C | 62.6404634 | -0.876802 | 0.61744973 | 2.55641313 | 0.27853639 | 0.45301284 | INO1 | X | 134332 | 135933 | -1 |
| YJL154C | 415.148645 | 3.01336517 | 0.5249407 | 81.3829561 | 2.13E-18 | 4.25E-16 | VPS35 | X | 131101 | 133935 | -1 |
| YJL155C | 27.4101396 | 1.62925083 | 0.81751111 | 3.97400587 | 0.13710572 | 0.27478094 | FBP26 | X | 129285 | 130643 | -1 |
| YJL156C | 557.772413 | 0.00460549 | 0.52018832 | 0.00959614 | 0.99521342 | 0.99607359 | SSY5 | X | 126853 | 128952 | -1 |
| YJL157C | 6181.51688 | 1.24722589 | 0.4914152 | 7.85748842 | 0.01966836 | 0.06514862 | FAR1 | X | 123836 | 126328 | -1 |
| YJL158C | 69.2434908 | 1.35517212 | 0.63211013 | 4.55435286 | 0.10257342 | 0.22310297 | CIS3 | X | 122265 | 122948 | -1 |
| YJL159W | 309.371886 | 1.97897682 | 0.49842188 | 15.7570342 | 0.00037879 | 0.00294788 | HSP150 | X | 120449 | 121690 | 1 |
| YJL160C | 96.5299955 | 0.02193631 | 0.58111409 | 0.07406111 | 0.96364669 | 0.97305796 | PIR5 | X | 117962 | 118825 | -1 |
| YJL161W | 62.3408202 | -1.1973408 | 0.59694209 | 15.2757852 | 0.00048184 | 0.00357676 | FMP33 | X | 117245 | 117787 | 1 |
| YJL162C | 45.8846464 | 0.31429153 | 0.65182674 | 5.33734997 | 0.06934405 | 0.16715322 | JJJ2 | X | 113878 | 115629 | -1 |
| YJL163C | 266.428937 | -0.2248354 | 0.50468164 | 2.42431484 | 0.29755464 | 0.47136562 | NA | X | 111666 | 113333 | -1 |
| YJL164C | 99.2758131 | 0.00909072 | 0.56135213 | 0.46993561 | 0.7905963 | 0.87307888 | TPK1 | X | 109966 | 111159 | -1 |
| YJL165C | 120.058348 | 1.63660698 | 0.73572931 | 4.80691563 | 0.09040481 | 0.20391268 | HAL5 | X | 106894 | 109461 | -1 |
| YJL166W | 25.3914042 | -0.5091041 | 0.78240122 | 0.67856342 | 0.71228176 | 0.82465735 | QCR8 | X | 106434 | 106718 | 1 |
| YJL167W | 69.5022343 | 1.01433472 | 0.71406988 | 2.49327812 | 0.28746934 | 0.46170527 | ERG20 | X | 105014 | 106072 | 1 |
| YJL168C | 20.3496734 | -0.2634748 | 0.74842473 | 7.94157718 | 0.01885856 | 0.06326248 | SET2 | X | 102227 | 104428 | -1 |
| YJL170C | 70.6463743 | -0.3877849 | 0.59358559 | 1.21495354 | 0.5447236 | 0.69762213 | ASG7 | X | 101150 | 101779 | -1 |
| YJL171C | 75.1984857 | -0.2699082 | 0.9173252 | 0.70883965 | 0.70158037 | 0.81832198 | TOH1 | X | 99704 | 100894 | -1 |
| YJL172W | 88.8217577 | 1.83030825 | 0.56599545 | 15.6880163 | 0.00039209 | 0.00301892 | CPS1 | X | 97737 | 99467 | 1 |
| YJL173C | 281.600017 | 1.3873696 | 0.75557581 | 4.65700332 | 0.09744164 | 0.21492841 | RFA3 | X | 96161 | 96529 | -1 |
| YJL174W | 27.1285024 | 1.16535059 | 0.80223317 | 2.42028502 | 0.29815479 | 0.47161883 | KRE9 | X | 95092 | 95922 | 1 |
| YJL176C | 44.3130636 | -0.8333838 | 0.62315197 | 4.11276086 | 0.12791613 | 0.26147876 | SWI3 | X | 92053 | 94530 | -1 |
| YJL177W | 75.9171548 | -0.7722809 | 0.73308472 | 15.7106413 | 0.00038768 | 0.00299292 | RPL17B | X | 90786 | 91657 | 1 |
| YJL178C | 47.4464551 | 0.23954416 | 0.58874218 | 0.46096305 | 0.79415111 | 0.87495229 | ATG27 | X | 89282 | 90097 | -1 |
| YJL179W | 11.4726746 | 1.8862381 | 0.99828604 | 8.46387623 | 0.01452421 | 0.0523127 | PFD1 | X | 88787 | 89116 | 1 |
| YJL180C | 237.011716 | 1.05649662 | 0.49455968 | 33.6088349 | 5.03E-08 | 1.32E-06 | ATP12 | X | 87583 | 88560 | -1 |
| YJL181W | 243.379109 | 3.01040121 | 0.65128913 | 29.8273338 | 3.33E-07 | 6.80E-06 | RBH1 | X | 85660 | 87495 | 1 |
| YJL183W | 14.2238489 | -1.0339097 | 0.83293735 | 4.52877084 | 0.10389387 | 0.22521358 | MNN11 | X | 84068 | 85336 | 1 |
| YJL184W | 39.1326857 | -0.0759646 | 0.66118026 | 1.28602017 | 0.52570761 | 0.68095013 | GON7 | X | 83448 | 83819 | 1 |
| YJL185C | 171.215454 | -1.4477751 | 0.52164544 | 17.3915818 | 0.00016729 | 0.00145654 | ATG36 | X | 82095 | 82976 | -1 |
| YJL186W | 85.3785603 | -0.5472386 | 0.60203397 | 7.62468687 | 0.02209634 | 0.07131784 | MNN5 | X | 80155 | 81915 | 1 |
| YJL187C | 246.300995 | 0.62171903 | 0.53959331 | 1.32957135 | 0.51438376 | 0.67150244 | SWE1 | X | 76803 | 79262 | -1 |

| | | | | | | | | | | | |
|---|---|---|---|---|---|---|---|---|---|---|---|
| YJL189W | 408.889744 | -0.4619204 | 0.59224643 | 20.4221875 | 3.68E-05 | 0.00040008 | RPL39 | X | 75933 | 76474 | 1 |
| YJL190C | 57.1930306 | -0.8248182 | 0.74294698 | 19.0494741 | 7.30E-05 | 0.00071783 | RPS22A | X | 74910 | 75302 | -1 |
| YJL191W | 27.3181265 | 0.1160285 | 0.75552085 | 8.70903453 | 0.01284864 | 0.04759047 | RPS14B | X | 73787 | 74611 | 1 |
| YJL192C | 21.5020388 | 0.3993062 | 0.78103394 | 1.63672453 | 0.44115356 | 0.60960103 | SOP4 | X | 72710 | 73414 | -1 |
| YJL193W | 80.7304619 | 0.41798516 | 0.72858409 | 0.42941421 | 0.80677771 | 0.88420271 | NA | X | 71366 | 72574 | 1 |
| YJL194W | 28.877216 | -1.6405314 | 0.72750538 | 5.01345254 | 0.08153473 | 0.18875892 | CDC6 | X | 69338 | 70879 | 1 |
| YJL196C | 241.93037 | -0.8092824 | 0.49106629 | 3.41166879 | 0.18162078 | 0.33650698 | ELO1 | X | 67850 | 68782 | -1 |
| YJL197W | 50.3680959 | -0.4555986 | 0.61939222 | 3.27003262 | 0.19494919 | 0.3521859 | UBP12 | X | 63805 | 67569 | 1 |
| YJL198W | 12.6741171 | 0.24990947 | 0.89627659 | 3.16594119 | 0.20536414 | 0.36665383 | PHO90 | X | 60844 | 63489 | 1 |
| YJL199C | 10.7785091 | 0.61362828 | 1.20173419 | 2.06657827 | 0.35583465 | 0.52936346 | NA | X | 59856 | 60182 | -1 |
| YJL200C | 172.800056 | 0.81786138 | 0.61164631 | 27.796587 | 9.21E-07 | 1.66E-05 | ACO2 | X | 56444 | 58813 | -1 |
| YJL201W | 36.9538348 | -0.2244965 | 0.65005813 | 3.50498284 | 0.17334154 | 0.32501539 | ECM25 | X | 54379 | 56178 | 1 |
| YJL203W | 13.1056946 | 0.04853095 | 0.965468 | 0.20804341 | 0.90120574 | 0.93824416 | PRP21 | X | 53341 | 54183 | 1 |
| YJL204C | 18.2892024 | 0.04052132 | 0.76302823 | 0.55192272 | 0.75884225 | 0.85429037 | RCY1 | X | 50629 | 53151 | -1 |
| YJL205C | 76.5662057 | -0.4233492 | 0.55787903 | 3.95837753 | 0.13818129 | 0.27626715 | NCE101 | X | 50139 | 50443 | -1 |
| YJL206C | 30.1388211 | -1.4750725 | 0.68280165 | 11.4900971 | 0.00319858 | 0.01606225 | NA | X | 47659 | 49935 | -1 |
| YJL207C | 28.5499004 | 0.06838579 | 0.68008919 | 0.60901559 | 0.73748628 | 0.84022935 | LAA1 | X | 41389 | 47433 | -1 |
| YJL208C | 38.4753009 | 0.88059172 | 0.64116567 | 4.27579602 | 0.11790241 | 0.24644584 | NUC1 | X | 40194 | 41183 | -1 |
| YJL209W | 27.4203554 | -0.9983937 | 0.6806598 | 18.3843288 | 0.00010183 | 0.000951 | CBP1 | X | 38005 | 39969 | 1 |
| YJL210W | 70.791331 | 0.03562693 | 0.64673487 | 1.82846055 | 0.40082503 | 0.57147918 | PEX2 | X | 36919 | 37734 | 1 |
| YJL212C | 190.003944 | 0.05872218 | 0.5362383 | 6.82013011 | 0.03303905 | 0.0962254 | OPT1 | X | 33850 | 36249 | -1 |
| YJL213W | 27.9656877 | -0.9822078 | 0.70322774 | 2.07386057 | 0.35454135 | 0.52838982 | NA | X | 32163 | 33158 | 1 |
| YJL214W | 609.218516 | -0.040514 | 0.6361862 | 0.17415447 | 0.9166063 | 0.94860224 | HXT8 | X | 26887 | 28596 | 1 |
| YJL216C | 9.34708582 | -1.5333176 | 1.06303935 | 3.28562156 | 0.19343557 | 0.35020614 | IMA5 | X | 24341 | 26086 | -1 |
| YJL217W | 13.6900742 | -0.608233 | 0.87056601 | 5.06949516 | 0.07928173 | 0.18517193 | REE1 | X | 23133 | 23729 | 1 |
| YJL219W | 91.4101822 | -1.1258942 | 0.56857278 | 4.46284649 | 0.1073755 | 0.23043148 | HXT9 | X | 19497 | 21200 | 1 |
| YJL225C | 9.26512606 | -2.0382015 | 1.05010052 | 3.86164511 | 0.14502886 | 0.28581248 | NA | X | 466 | 6130 | -1 |
| YJR001W | 130.424412 | 0.04898992 | 0.53819709 | 1.53476619 | 0.46422632 | 0.62726488 | AVT1 | X | 436802 | 438610 | 1 |
| YJR002W | 115.246635 | -0.5874206 | 0.54576295 | 7.8872584 | 0.01937776 | 0.06459254 | MPP10 | X | 438865 | 440646 | 1 |
| YJR003C | 94.506714 | -0.8792995 | 0.62018358 | 25.638446 | 2.71E-06 | 4.19E-05 | MRX12 | X | 440994 | 442553 | -1 |
| YJR004C | 699.681641 | 0.42399551 | 0.48594876 | 12.7662857 | 0.0016898 | 0.00985293 | SAG1 | X | 442909 | 444861 | -1 |
| YJR005W | 14.6446637 | 1.42653204 | 1.04584779 | 2.03196518 | 0.36204651 | 0.53489393 | APL1 | X | 445923 | 448025 | 1 |
| YJR006W | 44.480877 | -1.5053371 | 0.72016127 | 10.1730025 | 0.0061796 | 0.02710599 | POL31 | X | 449202 | 450665 | 1 |

| | | | | | | | | | | | |
|---|---|---|---|---|---|---|---|---|---|---|---|
| YJR007W | 30.5629523 | -0.5746797 | 0.6711193 | 0.82240216 | 0.66285363 | 0.79018376 | SUI2 | X | 451020 | 451934 | 1 |
| YJR008W | 97.6650905 | -2.2884515 | 0.60711292 | 26.2023697 | 2.04E-06 | 3.30E-05 | MHO1 | X | 452430 | 453446 | 1 |
| YJR009C | 78.7815646 | -0.0719035 | 0.74062488 | 6.35438079 | 0.04170266 | 0.11459819 | TDH2 | X | 453683 | 454681 | -1 |
| YJR010C-A | 206.917255 | -0.6117531 | 0.5093075 | 2.96023281 | 0.22761119 | 0.39362867 | SPC1 | X | 458077 | 458361 | -1 |
| YJR010W | 58.4306194 | 0.34210016 | 0.60471458 | 6.09885672 | 0.047386 | 0.12539532 | MET3 | X | 456239 | 457774 | 1 |
| YJR011C | 14.0052138 | -0.211563 | 0.85992875 | 1.77780759 | 0.41110616 | 0.58269393 | NA | X | 458641 | 459426 | -1 |
| YJR012C | 12.8345677 | 0.70518479 | 0.91451065 | 3.63623943 | 0.16233069 | 0.31091456 | NA | X | 459795 | 460418 | -1 |
| YJR013W | 91.1343385 | -1.1984465 | 0.53062694 | 17.2664162 | 0.00017809 | 0.00153674 | GPI14 | X | 460382 | 461593 | 1 |
| YJR014W | 89.2123005 | 0.24549021 | 0.62054496 | 4.39297427 | 0.11119308 | 0.23643332 | TMA22 | X | 461829 | 462425 | 1 |
| YJR015W | 26.16382 | -0.3197563 | 0.7095088 | 2.50833947 | 0.28531263 | 0.45887782 | NA | X | 462721 | 464253 | 1 |
| YJR016C | 37.1563054 | -0.3748025 | 0.69359173 | 5.31280905 | 0.07020017 | 0.16872519 | ILV3 | X | 464451 | 466208 | -1 |
| YJR017C | 48.5289065 | -0.5256527 | 0.60176856 | 5.17701281 | 0.07513217 | 0.17770232 | ESS1 | X | 466521 | 467033 | -1 |
| YJR019C | 219.291914 | -0.1710468 | 0.5847772 | 1.51717305 | 0.46832793 | 0.63119616 | TES1 | X | 467232 | 468281 | -1 |
| YJR021C | 67.2569711 | -0.3618076 | 0.59942669 | 0.73086377 | 0.6938969 | 0.81470846 | REC107 | X | 468555 | 469579 | -1 |
| YJR022W | 25.5769993 | 2.59214807 | 1.26926416 | 5.32078152 | 0.06992089 | 0.16819359 | LSM8 | X | 469784 | 470113 | 1 |
| YJR024C | 30.6369115 | 1.46153703 | 0.82758976 | 7.32965814 | 0.02560855 | 0.07911509 | MDE1 | X | 470230 | 470964 | -1 |
| YJR025C | 63.8207648 | 0.57194196 | 0.75869028 | 1.65532277 | 0.43707023 | 0.60599536 | BNA1 | X | 471138 | 471671 | -1 |
| YJR030C | 267.553759 | -0.7331547 | 0.4838491 | 11.7274423 | 0.00284065 | 0.01464593 | RBH2 | X | 483959 | 486196 | -1 |
| YJR031C | 38.896291 | -0.0155544 | 0.74764552 | 7.4214932 | 0.02445926 | 0.07676865 | GEA1 | X | 486586 | 490812 | -1 |
| YJR032W | 136.464066 | 1.97258617 | 0.61700036 | 48.558231 | 2.86E-11 | 1.49E-09 | CPR7 | X | 491081 | 492262 | 1 |
| YJR033C | 147.525321 | 0.5336555 | 0.69982314 | 2.2277076 | 0.32829135 | 0.50073944 | RAV1 | X | 492378 | 496451 | -1 |
| YJR034W | 10.607743 | 0.17297992 | 1.05206685 | 4.1505244 | 0.12552351 | 0.25827332 | PET191 | X | 496683 | 497009 | 1 |
| YJR035W | 58.9579429 | 0.79845862 | 0.57817014 | 1.90741853 | 0.38530916 | 0.55815362 | RAD26 | X | 497355 | 500612 | 1 |
| YJR036C | 125.578664 | 0.32633654 | 0.6043794 | 2.07182544 | 0.35490231 | 0.52857757 | HUL4 | X | 500713 | 503391 | -1 |
| YJR039W | 58.7682428 | -1.7611134 | 0.58396355 | 15.0603514 | 0.00053664 | 0.00388561 | NA | X | 503936 | 507301 | 1 |
| YJR040W | 76.3046596 | 0.15189138 | 0.64394584 | 0.68799436 | 0.70893093 | 0.82407349 | GEF1 | X | 507746 | 510085 | 1 |
| YJR041C | 14.258325 | -0.3683375 | 0.85852071 | 0.94143264 | 0.62455473 | 0.76345394 | URB2 | X | 510239 | 513763 | -1 |
| YJR042W | 24.200017 | -0.1634625 | 0.71598127 | 0.94715896 | 0.62276908 | 0.76281637 | NUP85 | X | 514055 | 516289 | 1 |
| YJR043C | 17.9547947 | -0.3770784 | 0.78838424 | 5.4060733 | 0.06700174 | 0.16264366 | POL32 | X | 516461 | 517513 | -1 |
| YJR044C | 102.482711 | 1.61301016 | 0.53834348 | 12.6769725 | 0.00176697 | 0.01020018 | VPS55 | X | 518763 | 519185 | -1 |
| YJR045C | 122.322449 | -0.2967889 | 0.79241856 | 0.21370137 | 0.89865985 | 0.9375281 | SSC1 | X | 519638 | 521602 | -1 |
| YJR046W | 13.3295612 | -1.3523548 | 0.92978932 | 2.18320503 | 0.33567813 | 0.50893088 | TAH11 | X | 522048 | 523862 | 1 |
| YJR048W | 107.502032 | -0.8752803 | 0.62589241 | 11.4258067 | 0.00330307 | 0.0164313 | CYC1 | X | 526335 | 526664 | 1 |

| | | | | | | | | | | | |
|---|---|---|---|---|---|---|---|---|---|---|---|
| YJR049C | 53.3372832 | 0.26851501 | 0.67052371 | 0.80629162 | 0.66821466 | 0.79444823 | UTR1 | X | 526884 | 528476 | -1 |
| YJR050W | 14.8919863 | -0.5784763 | 0.87025415 | 5.04598524 | 0.08021918 | 0.18675877 | ISY1 | X | 528697 | 529404 | 1 |
| YJR051W | 51.3429471 | 0.30495882 | 0.58979107 | 1.89217391 | 0.38825732 | 0.56046121 | OSM1 | X | 529861 | 531366 | 1 |
| YJR052W | 35.1691426 | 0.39091999 | 0.6328088 | 0.58663954 | 0.74578362 | 0.84519224 | RAD7 | X | 532062 | 533759 | 1 |
| YJR053W | 68.7711988 | 1.01450307 | 0.63582319 | 3.38778747 | 0.18380245 | 0.33935465 | BFA1 | X | 534027 | 535751 | 1 |
| YJR054W | 81.3624894 | 0.96312815 | 0.54981098 | 7.70257323 | 0.02125238 | 0.06924663 | KCH1 | X | 536056 | 537549 | 1 |
| YJR055W | 6.42884824 | -0.2295124 | 1.08458902 | 0.46773885 | 0.79146515 | 0.87352731 | HIT1 | X | 538772 | 539266 | 1 |
| YJR056C | 16.6775826 | -0.4106178 | 0.80694875 | 1.37968087 | 0.50165611 | 0.66050451 | NA | X | 541792 | 542502 | -1 |
| YJR057W | 163.031073 | 0.60864 | 0.60909558 | 1.77026748 | 0.41265898 | 0.58426143 | CDC8 | X | 544062 | 544712 | 1 |
| YJR058C | 165.241021 | -0.0005036 | 0.52816436 | 9.08085919 | 0.01066882 | 0.04137474 | APS2 | X | 544732 | 545175 | -1 |
| YJR059W | 122.76682 | 0.10037198 | 0.51876781 | 2.4159619 | 0.29879996 | 0.47191811 | PTK2 | X | 545787 | 548243 | 1 |
| YJR060W | 42.5269261 | 0.20232499 | 0.68190973 | 0.72624006 | 0.69550295 | 0.81611981 | CBF1 | X | 548759 | 549814 | 1 |
| YJR061W | 1878.34739 | 0.04752311 | 0.47577334 | 1.09609868 | 0.57807634 | 0.727271 | NA | X | 550511 | 553318 | 1 |
| YJR062C | 8.50294115 | 0.44493727 | 1.03362086 | 0.41483735 | 0.81267933 | 0.88731158 | NTA1 | X | 553476 | 554849 | -1 |
| YJR063W | 6.67275698 | -0.9765223 | 1.18839433 | 2.44544435 | 0.29442759 | 0.46884326 | RPA12 | X | 555195 | 555572 | 1 |
| YJR064W | 61.9067673 | 1.06777442 | 0.63939616 | 18.6096648 | 9.10E-05 | 0.00086218 | CCT5 | X | 555914 | 557602 | 1 |
| YJR065C | 44.84875 | 0.87684147 | 0.72000743 | 1.51537577 | 0.46874898 | 0.63146966 | ARP3 | X | 557809 | 559158 | -1 |
| YJR066W | 333.281053 | -1.1747295 | 0.51659271 | 7.21374065 | 0.02713664 | 0.0824783 | TOR1 | X | 559416 | 566828 | 1 |
| YJR067C | 13.1671849 | 1.17219605 | 1.02808994 | 2.64601886 | 0.26633258 | 0.43871 | YAE1 | X | 567019 | 567444 | -1 |
| YJR068W | 489.633996 | 1.54988364 | 0.86726404 | 7.9366423 | 0.01890515 | 0.06338205 | RFC2 | X | 567643 | 568704 | 1 |
| YJR069C | 56.4966958 | -0.9115642 | 0.64222119 | 2.0150294 | 0.3651253 | 0.53820659 | HAM1 | X | 568806 | 569399 | -1 |
| YJR070C | 23.4144297 | -0.5584235 | 0.8906019 | 0.83926486 | 0.65728838 | 0.78549013 | LIA1 | X | 569621 | 570598 | -1 |
| YJR072C | 49.8537921 | -0.5154655 | 0.57629502 | 7.55798948 | 0.02284565 | 0.07312122 | NPA3 | X | 570967 | 572124 | -1 |
| YJR073C | 21.2435479 | -0.7583647 | 0.91062134 | 1.31187257 | 0.51895594 | 0.67522582 | OPI3 | X | 572315 | 572935 | -1 |
| YJR074W | 238.474681 | 0.88912794 | 0.65364806 | 5.12769041 | 0.07700806 | 0.18110344 | MOG1 | X | 573095 | 573751 | 1 |
| YJR075W | 61.6640786 | 0.04292606 | 0.60877012 | 4.3263156 | 0.11496152 | 0.24204626 | HOC1 | X | 573981 | 575171 | 1 |
| YJR076C | 156.896106 | 2.1919161 | 0.73958572 | 10.8123766 | 0.00448872 | 0.02089629 | CDC11 | X | 575354 | 576601 | -1 |
| YJR077C | 45.6762697 | 0.41406132 | 0.78498471 | 0.31132748 | 0.85584693 | 0.9146094 | MIR1 | X | 577255 | 578190 | -1 |
| YJR078W | 20.1889223 | 0.44322054 | 0.83237281 | 5.8133336 | 0.05465761 | 0.13990609 | BNA2 | X | 578860 | 580221 | 1 |
| YJR079W | 998.098346 | 0.09185872 | 0.46222708 | 3.24324596 | 0.19757777 | 0.35571372 | NA | X | 580205 | 581239 | 1 |
| YJR080C | 279.421625 | 1.4555714 | 0.50187376 | 20.7450632 | 3.13E-05 | 0.00034964 | AIM24 | X | 580432 | 581616 | -1 |
| YJR082C | 6.20026956 | 0.7526519 | 1.20155032 | 1.47483911 | 0.47834667 | 0.64096904 | EAF6 | X | 581914 | 582255 | -1 |
| YJR083C | 48.1456682 | 0.19822592 | 0.64818575 | 3.36242999 | 0.18614767 | 0.34183159 | ACF4 | X | 582608 | 583537 | -1 |

| | | | | | | | | | | | |
|---|---|---|---|---|---|---|---|---|---|---|---|
| YJR084W | 53.6844611 | 0.90797287 | 0.6378312 | 9.16983971 | 0.01020457 | 0.04005725 | NA | X | 583733 | 585004 | 1 |
| YJR085C | 42.9170799 | 1.76308924 | 0.81334924 | 6.0610479 | 0.04829033 | 0.12713301 | NA | X | 585120 | 585437 | -1 |
| YJR086W | 646.259016 | 1.14923572 | 0.64107637 | 5.12034559 | 0.07729138 | 0.18169594 | STE18 | X | 586068 | 586400 | 1 |
| YJR088C | 405.281537 | -0.8573026 | 0.51691562 | 20.1885205 | 4.13E-05 | 0.00044055 | EMC2 | X | 586495 | 587373 | -1 |
| YJR089W | 86.133817 | 0.64641545 | 0.57869176 | 25.9234627 | 2.35E-06 | 3.70E-05 | BIR1 | X | 587718 | 590582 | 1 |
| YJR090C | 27.1345701 | 0.55979897 | 0.77288952 | 2.5672124 | 0.27703645 | 0.45146103 | GRR1 | X | 590872 | 594327 | -1 |
| YJR091C | 106.55512 | 0.40028408 | 0.61170913 | 9.51720287 | 0.0085776 | 0.03502418 | JSN1 | X | 595061 | 598336 | -1 |
| YJR092W | 191.335585 | 0.17953903 | 0.50311477 | 0.30586355 | 0.85818827 | 0.91626865 | BUD4 | X | 598735 | 603078 | 1 |
| YJR093C | 20.4223302 | 0.00286415 | 0.74913724 | 1.53242476 | 0.46477011 | 0.62728439 | FIP1 | X | 603220 | 604203 | -1 |
| YJR094C | 8.95062326 | -0.4304843 | 1.00010531 | 0.23484644 | 0.88920878 | 0.93304075 | IME1 | X | 604569 | 605651 | -1 |
| YJR094W-A | 69.166065 | -0.6890249 | 0.78394245 | 11.0457843 | 0.00399428 | 0.01925635 | RPL43B | X | 608305 | 608858 | 1 |
| YJR095W | 60.022549 | 0.15490478 | 0.6628252 | 1.97779737 | 0.37198614 | 0.54526576 | SFC1 | X | 609773 | 610741 | 1 |
| YJR096W | 316.977027 | -0.1422335 | 0.53218592 | 0.13781763 | 0.93341179 | 0.95735444 | NA | X | 611195 | 612043 | 1 |
| YJR097W | 23.1712516 | 0.24569842 | 0.80526784 | 2.3955273 | 0.30186854 | 0.47448396 | JJJ3 | X | 612413 | 612931 | 1 |
| YJR098C | 16.5668403 | -0.5514996 | 0.77399993 | 1.39417308 | 0.4980342 | 0.65775958 | NA | X | 613186 | 615156 | -1 |
| YJR099W | 82.9140628 | 1.55580706 | 0.82316561 | 3.80221781 | 0.14940285 | 0.29203007 | YUH1 | X | 615576 | 616286 | 1 |
| YJR100C | 200.21551 | 0.80919987 | 0.67058944 | 3.85188572 | 0.14573828 | 0.28701519 | AIM25 | X | 616351 | 617334 | -1 |
| YJR101W | 52.446302 | 0.85697383 | 0.76054525 | 8.24978465 | 0.01616524 | 0.05670589 | RSM26 | X | 618234 | 619034 | 1 |
| YJR102C | 210.791026 | 1.0026122 | 0.75703305 | 4.48519734 | 0.10618221 | 0.22871838 | VPS25 | X | 619157 | 619765 | -1 |
| YJR103W | 1193.8047 | 1.9408935 | 0.94709797 | 6.36882746 | 0.04140251 | 0.11426146 | URA8 | X | 620754 | 622490 | 1 |
| YJR104C | 191.520352 | 0.57021783 | 0.59756528 | 1.88805007 | 0.3890587 | 0.56089429 | SOD1 | X | 622550 | 623014 | -1 |
| YJR105W | 927.15484 | 2.55746746 | 0.55700445 | 36.5408976 | 1.16E-08 | 3.42E-07 | ADO1 | X | 623581 | 624603 | 1 |
| YJR106W | 70.8389838 | -1.4204781 | 0.60185429 | 5.66749483 | 0.05879212 | 0.14748831 | ECM27 | X | 624838 | 627015 | 1 |
| YJR107C-A | 36.4917666 | 2.28539153 | 0.8537966 | 8.29214469 | 0.01582646 | 0.05593181 | NA | NA | NA | NA | NA |
| YJR107W | 626.375473 | -0.344998 | 0.53175783 | 2.71346853 | 0.25750033 | 0.42958807 | LIH1 | X | 627340 | 628326 | 1 |
| YJR108W | 56.8564524 | 0.30804299 | 0.6867756 | 0.76115738 | 0.68346578 | 0.80596066 | ABM1 | X | 628712 | 629083 | 1 |
| YJR109C | 202.446166 | -0.8295559 | 0.52271171 | 4.16272558 | 0.12476007 | 0.25697646 | CPA2 | X | 629585 | 632941 | -1 |
| YJR110W | 18.2875502 | 0.03597649 | 0.79930474 | 1.56866178 | 0.456425 | 0.62207609 | YMR1 | X | 633615 | 635681 | 1 |
| YJR111C | 29.7089708 | 1.36648356 | 0.81746535 | 3.32166669 | 0.1899806 | 0.34583253 | PXP2 | X | 635855 | 636706 | -1 |
| YJR112W | 16.6779089 | -1.9733959 | 0.87994423 | 9.49930821 | 0.00865469 | 0.03526435 | NNF1 | X | 637030 | 637635 | 1 |
| YJR112W-A | 238.546208 | 1.67139982 | 0.82148684 | 7.39521507 | 0.02478275 | 0.07731256 | NA | X | 637790 | 638168 | 1 |
| YJR113C | 110.813539 | 0.54992859 | 0.58512273 | 1.00660975 | 0.60452946 | 0.7496735 | RSM7 | X | 638232 | 638975 | -1 |
| YJR115W | 13.0246363 | 0.15498783 | 0.86690556 | 2.71323612 | 0.25753026 | 0.42958807 | NA | X | 639942 | 640451 | 1 |

| | | | | | | | | | | | |
|---|---|---|---|---|---|---|---|---|---|---|---|
| YJR116W | 194.925208 | -0.1390313 | 0.60545658 | 0.09638417 | 0.95295072 | 0.96731849 | TDA4 | X | 640825 | 641664 | 1 |
| YJR117W | 46.6450083 | 2.39472436 | 0.89255266 | 8.22917898 | 0.01633264 | 0.05693318 | STE24 | X | 642007 | 643368 | 1 |
| YJR118C | 358.282613 | 0.02795965 | 0.71150415 | 7.42878344 | 0.02437026 | 0.07671608 | ILM1 | X | 643490 | 644101 | -1 |
| YJR119C | 130.911035 | -0.7253691 | 0.54438575 | 8.69940929 | 0.01291063 | 0.04773469 | JHD2 | X | 644304 | 646490 | -1 |
| YJR120W | 29.9841622 | -0.2237105 | 0.86925084 | 11.1380611 | 0.00381418 | 0.0184804 | NA | X | 647126 | 647476 | 1 |
| YJR121W | 127.215234 | 0.18131485 | 0.75342018 | 1.55884525 | 0.45867076 | 0.62328904 | ATP2 | X | 647607 | 649142 | 1 |
| YJR122W | 67.5453114 | -0.864171 | 0.57553196 | 2.40020816 | 0.30116287 | 0.47428323 | IBA57 | X | 649776 | 651269 | 1 |
| YJR123W | 245.660746 | -0.6499467 | 0.68253993 | 20.5457397 | 3.46E-05 | 0.00037968 | RPS5 | X | 651901 | 652578 | 1 |
| YJR124C | 212.260736 | 1.20043059 | 0.508245 | 12.0219278 | 0.00245172 | 0.01302799 | NA | X | 652892 | 654238 | -1 |
| YJR125C | 52.2658751 | 0.7949455 | 0.65223221 | 2.14307455 | 0.34248162 | 0.51615859 | ENT3 | X | 654737 | 655963 | -1 |
| YJR126C | 30.3031533 | 1.08983179 | 0.83669787 | 1.73786191 | 0.41939967 | 0.59032562 | VPS70 | X | 656254 | 658689 | -1 |
| YJR127C | 175.78987 | 0.07087237 | 0.52654317 | 0.24558055 | 0.88444913 | 0.9315066 | RSF2 | X | 658917 | 663059 | -1 |
| YJR128W | 118.04878 | -1.0376599 | 0.65310112 | 5.07242535 | 0.07916566 | 0.18505013 | NA | X | 662921 | 663280 | 1 |
| YJR129C | 35.3831869 | -1.4299809 | 0.67926673 | 7.16846461 | 0.02775797 | 0.08383862 | EFM3 | X | 664000 | 665019 | -1 |
| YJR130C | 136.977473 | -0.6615688 | 0.51476316 | 1.95899121 | 0.37550045 | 0.54902718 | STR2 | X | 665218 | 667137 | -1 |
| YJR131W | 153.08395 | 0.85474032 | 0.54741472 | 16.7534563 | 0.00023016 | 0.00192578 | MNS1 | X | 667644 | 669293 | 1 |
| YJR132W | 86.3285409 | -1.2842938 | 0.64667055 | 9.32599758 | 0.00943812 | 0.03773943 | NMD5 | X | 669522 | 672668 | 1 |
| YJR133W | 14.057757 | 0.32891728 | 0.87750505 | 2.33425975 | 0.31125901 | 0.48393923 | XPT1 | X | 672991 | 673620 | 1 |
| YJR134C | 40.7222961 | 1.77639705 | 0.83300005 | 6.85023361 | 0.03254548 | 0.09507484 | SGM1 | X | 673729 | 675852 | -1 |
| YJR135C | 11.8983939 | -0.475339 | 0.94411586 | 0.4007655 | 0.81841744 | 0.8908887 | MCM22 | X | 676059 | 676778 | -1 |
| YJR135W-A | 6.73770371 | -1.5138371 | 1.18071185 | 2.48927476 | 0.28804534 | 0.46186168 | TIM8 | X | 676971 | 677234 | 1 |
| YJR136C | 33.3992789 | 0.16349281 | 0.67852426 | 0.17248593 | 0.91737131 | 0.94883528 | TTI2 | X | 677441 | 678706 | -1 |
| YJR137C | 33.0280823 | -0.4667803 | 0.65454706 | 0.90317324 | 0.63661728 | 0.77291131 | MET5 | X | 678957 | 683285 | -1 |
| YJR138W | 114.073731 | 1.40676055 | 0.66875758 | 5.36582491 | 0.06836376 | 0.16534092 | IML1 | X | 684567 | 689321 | 1 |
| YJR139C | 70.5652479 | 1.6342534 | 0.70202744 | 7.24839225 | 0.02667053 | 0.08136057 | HOM6 | X | 689445 | 690524 | -1 |
| YJR140C | 30.1369695 | -0.8086224 | 0.82786581 | 1.65617569 | 0.43688388 | 0.60588207 | HIR3 | X | 690750 | 695696 | -1 |
| YJR141W | 41.7615171 | 0.0506836 | 0.65381083 | 0.81538595 | 0.66518307 | 0.79181949 | IPA1 | X | 695906 | 696949 | 1 |
| YJR142W | 75.8418714 | 0.7304873 | 0.61420027 | 1.42963525 | 0.48928134 | 0.65095104 | NA | X | 697141 | 698169 | 1 |
| YJR143C | 51.7118877 | 0.21054228 | 0.59652623 | 0.80301426 | 0.66931055 | 0.79493499 | PMT4 | X | 698326 | 700614 | -1 |
| YJR144W | 16.3314162 | -0.3470942 | 0.84138141 | 2.36075005 | 0.30716352 | 0.4798912 | MGM101 | X | 700882 | 701691 | 1 |
| YJR145C | 103.381087 | -0.5411286 | 0.7858034 | 12.8956195 | 0.00158399 | 0.00951154 | RPS4A | X | 702027 | 703068 | -1 |
| YJR146W | 597.333776 | -0.7582372 | 0.64476126 | 3.28943858 | 0.19306675 | 0.34965796 | NA | X | 703885 | 704238 | 1 |
| YJR147W | 1543.38296 | -0.7540072 | 0.58348421 | 5.23885643 | 0.0728445 | 0.17363922 | HMS2 | X | 704196 | 705272 | 1 |

| | | | | | | | | | | | |
|---|---|---|---|---|---|---|---|---|---|---|---|
| YJR148W | 63.1141373 | 0.60114504 | 0.63577008 | 5.35150016 | 0.06885516 | 0.16625162 | BAT2 | X | 705744 | 706874 | 1 |
| YJR149W | 7.64958468 | 0.32316626 | 1.1342812 | 2.44279195 | 0.29481832 | 0.46908438 | NA | X | 707160 | 708374 | 1 |
| YJR150C | 20.8033201 | -1.6977054 | 0.89844344 | 4.59562132 | 0.10047858 | 0.2196191 | DAN1 | X | 708811 | 709707 | -1 |
| YJR151C | 85.5431181 | -0.8687516 | 0.56011739 | 4.12241331 | 0.12730027 | 0.26090922 | DAN4 | X | 712255 | 715740 | -1 |
| YJR151W-A | 36.0707463 | 0.6819757 | 0.77894523 | 7.11261061 | 0.02854409 | 0.08567667 | NA | X | 717580 | 717630 | 1 |
| YJR152W | 19.9763271 | 0.53365326 | 0.99516209 | 1.60495965 | 0.44821609 | 0.61496946 | DAL5 | X | 719666 | 721297 | 1 |
| YJR153W | 780.033524 | -0.0952598 | 0.54954506 | 0.15761856 | 0.92421617 | 0.95251186 | PGU1 | X | 722815 | 723900 | 1 |
| YJR154W | 12.8983937 | -0.250449 | 0.88382367 | 0.59410642 | 0.74300447 | 0.84319794 | NA | X | 725784 | 726824 | 1 |
| YJR155W | 29.1104515 | -0.6078841 | 0.68023152 | 1.39852758 | 0.49695103 | 0.65655356 | AAD10 | X | 727405 | 728271 | 1 |
| YJR158W | 14.3882458 | -0.7119975 | 0.91752086 | 3.13336633 | 0.20873638 | 0.37130066 | HXT16 | X | 732440 | 734143 | 1 |
| YJR159W | 226.440232 | 1.69848556 | 0.58740687 | 14.3995489 | 0.00074675 | 0.00515955 | SOR1 | X | 736044 | 737117 | 1 |
| YJR160C | 351.180581 | 1.10160489 | 0.62540676 | 8.81731858 | 0.01217149 | 0.04561353 | MPH3 | X | 738008 | 739816 | -1 |
| YJR161C | 23.499939 | -0.6500329 | 0.73762205 | 2.03991887 | 0.36060957 | 0.53345156 | COS5 | X | 742848 | 743999 | -1 |
| YKL001C | 93.3982281 | 4.33147752 | 1.05636533 | 17.4440476 | 0.00016296 | 0.00142526 | MET14 | XI | 438777 | 439385 | -1 |
| YKL002W | 100.06472 | -1.1009345 | 0.54453479 | 9.07399827 | 0.01070548 | 0.04143366 | DID4 | XI | 437778 | 438544 | 1 |
| YKL003C | 17.8148887 | -1.2106516 | 1.06559641 | 1.32655883 | 0.51515914 | 0.67209811 | MRP17 | XI | 437097 | 437492 | -1 |
| YKL004W | 254.887325 | 1.94423931 | 0.58805057 | 44.3485583 | 2.34E-10 | 1.01E-08 | AUR1 | XI | 435580 | 436785 | 1 |
| YKL005C | 43.3104528 | -0.3316392 | 0.59650588 | 0.59081701 | 0.7442275 | 0.84379317 | BYE1 | XI | 433093 | 434877 | -1 |
| YKL006C-A | 1270.7579 | 2.17493748 | 0.66654659 | 12.1708199 | 0.00227583 | 0.01230351 | SFT1 | XI | 430198 | 430632 | -1 |
| YKL006W | 51.0903532 | -0.5238468 | 0.74754 | 12.260768 | 0.00217575 | 0.01190696 | RPL14A | XI | 431906 | 432720 | 1 |
| YKL007W | 42.3518355 | 0.9024502 | 0.68000605 | 8.66977691 | 0.01310334 | 0.04820096 | CAP1 | XI | 429302 | 430108 | 1 |
| YKL008C | 139.71816 | -0.5772276 | 0.49800606 | 3.48841837 | 0.17478316 | 0.32676605 | LAC1 | XI | 427295 | 428551 | -1 |
| YKL009W | 44.513419 | -1.2855308 | 0.78996052 | 14.4091126 | 0.00074319 | 0.00514108 | MRT4 | XI | 426242 | 426952 | 1 |
| YKL010C | 86.4119865 | 1.0957629 | 0.60655919 | 4.6105826 | 0.09972974 | 0.2191405 | UFD4 | XI | 421424 | 425875 | -1 |
| YKL011C | 192.985953 | 1.35543657 | 0.56167694 | 5.67982383 | 0.05843081 | 0.14688185 | CCE1 | XI | 420155 | 421216 | -1 |
| YKL012W | 88.5743234 | 0.60079772 | 0.54957005 | 2.73320731 | 0.25497146 | 0.42716573 | PRP40 | XI | 418310 | 420061 | 1 |
| YKL013C | 33.7582319 | 0.85853757 | 0.73217682 | 5.58345857 | 0.06131509 | 0.15223601 | ARC19 | XI | 417508 | 418023 | -1 |
| YKL014C | 47.0224955 | 0.40069409 | 0.71000818 | 7.63129374 | 0.02202346 | 0.07124193 | URB1 | XI | 411619 | 416913 | -1 |
| YKL015W | 32.312395 | -0.7226536 | 0.70701449 | 4.30397286 | 0.116253 | 0.24370198 | PUT3 | XI | 408544 | 411483 | 1 |
| YKL016C | 17.2837261 | 0.19508068 | 0.78479744 | 0.47287353 | 0.7894358 | 0.87229643 | ATP7 | XI | 407465 | 407989 | -1 |
| YKL017C | 159.276209 | 0.9604224 | 0.70826613 | 6.77044552 | 0.0338701 | 0.09820123 | HCS1 | XI | 405190 | 407241 | -1 |
| YKL018C-A | 246.376603 | 2.17618763 | 0.85015325 | 8.1139634 | 0.01730116 | 0.05948558 | NA | XI | 403580 | 403879 | -1 |
| YKL018W | 65.0205263 | 0.52617761 | 0.69855104 | 1.46805352 | 0.47997236 | 0.64284986 | SWD2 | XI | 404102 | 405091 | 1 |

| | | | | | | | | | | | |
|---|---|---|---|---|---|---|---|---|---|---|---|
| YKL019W | 156.74458 | 1.72877944 | 0.71599739 | 12.8849968 | 0.00159242 | 0.00954465 | RAM2 | XI | 402568 | 403518 | 1 |
| YKL020C | 25.3439001 | 0.35868648 | 0.68979801 | 0.4302067 | 0.80645809 | 0.88416839 | SPT23 | XI | 398832 | 402080 | -1 |
| YKL021C | 29.6740585 | -0.8147109 | 0.69158671 | 5.84401993 | 0.05382539 | 0.13826487 | NA | XI | 397344 | 398750 | -1 |
| YKL022C | 496.310355 | 0.16907835 | 0.45644675 | 2.47572623 | 0.29000326 | 0.46384499 | CDC16 | XI | 394726 | 397248 | -1 |
| YKL023C-A | 100.395794 | 0.70986496 | 0.64601778 | 29.1644113 | 4.65E-07 | 9.09E-06 | NA | XI | 393146 | 393373 | -1 |
| YKL023W | 255.415489 | -0.0038317 | 0.53854229 | 2.34111563 | 0.31019386 | 0.4830878 | NA | XI | 393721 | 394554 | 1 |
| YKL024C | 5.35595216 | -0.38203 | 1.34542333 | 0.11572758 | 0.94377849 | 0.96197675 | URA6 | XI | 392526 | 393140 | -1 |
| YKL025C | 32.6859159 | 1.0937858 | 0.67612263 | 2.83602626 | 0.24219475 | 0.41135453 | PAN3 | XI | 390240 | 392279 | -1 |
| YKL026C | 10.257265 | -0.6597429 | 0.95722433 | 0.77295509 | 0.67944598 | 0.80269174 | GPX1 | XI | 389380 | 389883 | -1 |
| YKL027W | 1179.17709 | 1.54171687 | 0.51849158 | 107.482661 | 4.58E-24 | 2.94E-21 | TCD2 | XI | 387562 | 388905 | 1 |
| YKL028W | 54.9678836 | 0.25450274 | 0.60155677 | 1.54539571 | 0.46176561 | 0.62532091 | TFA1 | XI | 385758 | 387206 | 1 |
| YKL029C | 26.539856 | -0.8233775 | 0.75200461 | 1.50709423 | 0.47069398 | 0.63296713 | MAE1 | XI | 382716 | 384725 | -1 |
| YKL032C | 99.9374448 | -0.2846222 | 0.60551647 | 0.60545556 | 0.73880019 | 0.84040603 | IXR1 | XI | 380065 | 381858 | -1 |
| YKL033W | 118.584921 | -0.0871203 | 0.55272538 | 19.8550422 | 4.88E-05 | 0.00050469 | TTI1 | XI | 375456 | 378572 | 1 |
| YKL033W-A | 140.985854 | 0.72299115 | 0.51753908 | 12.1478111 | 0.00230216 | 0.01239956 | NA | XI | 374505 | 375215 | 1 |
| YKL034W | 50.3273129 | -0.3976651 | 0.62948959 | 4.03038954 | 0.13329444 | 0.26928639 | TUL1 | XI | 371829 | 374105 | 1 |
| YKL035W | 39.0289375 | -0.1059069 | 0.74378045 | 0.2276668 | 0.89240662 | 0.93387025 | UGP1 | XI | 369891 | 371390 | 1 |
| YKL037W | 195.938968 | 1.16628476 | 0.52295081 | 6.22827406 | 0.04441682 | 0.11983849 | AIM26 | XI | 369365 | 369721 | 1 |
| YKL038W | 24.448692 | 0.93770267 | 0.71389572 | 2.25198072 | 0.3243311 | 0.49763242 | RGT1 | XI | 365605 | 369117 | 1 |
| YKL039W | 39.6116109 | 1.23269139 | 0.7632471 | 2.92324815 | 0.23185941 | 0.39906837 | PTM1 | XI | 362622 | 364193 | 1 |
| YKL040C | 35.6615512 | 0.4153537 | 0.62553476 | 2.51887064 | 0.28381425 | 0.45748454 | NFU1 | XI | 361058 | 361828 | -1 |
| YKL041W | 552.459265 | 0.44387113 | 0.7371153 | 9.30548646 | 0.00953541 | 0.03806203 | VPS24 | XI | 360143 | 360817 | 1 |
| YKL042W | 10.4765366 | -0.8392209 | 1.00661416 | 3.97615931 | 0.13695818 | 0.27458028 | SPC42 | XI | 358475 | 359566 | 1 |
| YKL043W | 26.6077503 | 0.05996518 | 0.87274139 | 0.06617008 | 0.96745628 | 0.97486964 | PHD1 | XI | 356748 | 357848 | 1 |
| YKL044W | 3.79727751 | -0.554599 | 1.58999244 | 2.06405392 | 0.35628406 | 0.52948785 | MMO1 | XI | 356322 | 356642 | 1 |
| YKL045W | 234.737412 | -0.8209321 | 0.5593604 | 2.49101028 | 0.28779549 | 0.46172289 | PRI2 | XI | 353493 | 355079 | 1 |
| YKL046C | 52.1280153 | 0.97659548 | 0.87443643 | 3.32147462 | 0.18999884 | 0.34583253 | DCW1 | XI | 351275 | 352624 | -1 |
| YKL047W | 49.8150124 | -0.4727673 | 0.66002268 | 1.25051813 | 0.53512278 | 0.68975087 | ANR2 | XI | 349463 | 351013 | 1 |
| YKL048C | 24.9427364 | 0.2957791 | 0.73932474 | 2.99390703 | 0.22381096 | 0.38938265 | ELM1 | XI | 347214 | 349136 | -1 |
| YKL049C | 8.62966844 | 0.51688207 | 1.12956656 | 1.71569038 | 0.4240749 | 0.5946703 | CSE4 | XI | 346075 | 346764 | -1 |
| YKL050C | 42.0073784 | -0.038603 | 0.66145118 | 1.27540342 | 0.52850569 | 0.68289398 | NA | XI | 342852 | 345620 | -1 |
| YKL051W | 318.044893 | -0.1822028 | 0.50290415 | 2.91425693 | 0.23290411 | 0.40027153 | SFK1 | XI | 341312 | 342373 | 1 |
| YKL052C | 152.980799 | -0.4350128 | 0.50293881 | 1.62387919 | 0.44399606 | 0.61165577 | ASK1 | XI | 339668 | 340546 | -1 |

| | | | | | | | | | | | |
|---|---|---|---|---|---|---|---|---|---|---|---|
| YKL053C-A | 340.02936 | 2.45679635 | 0.49963124 | 39.2024545 | 3.07E-09 | 1.03E-07 | MDM35 | XI | 339182 | 339442 | -1 |
| YKL054C | 1654.29956 | 2.09564346 | 0.89091277 | 9.59198834 | 0.00826278 | 0.03401477 | DEF1 | XI | 336537 | 338753 | -1 |
| YKL055C | 48.2098984 | -1.0481196 | 0.61931105 | 8.99323396 | 0.01114664 | 0.04268456 | OAR1 | XI | 335321 | 336157 | -1 |
| YKL056C | 85.7976123 | -0.5513727 | 0.8487423 | 7.08402943 | 0.02895493 | 0.08664034 | TMA19 | XI | 334412 | 334915 | -1 |
| YKL057C | 43.0588934 | 1.13884263 | 0.76040486 | 8.68274432 | 0.01301865 | 0.04801146 | NUP120 | XI | 330856 | 333969 | -1 |
| YKL058W | 98.8202612 | -0.4952207 | 0.56520853 | 2.10582806 | 0.34891951 | 0.52261068 | TOA2 | XI | 330166 | 330534 | 1 |
| YKL059C | 17.2163274 | 0.28961368 | 0.81134456 | 0.15617945 | 0.92488144 | 0.95277599 | MPE1 | XI | 328118 | 329443 | -1 |
| YKL060C | 701.900665 | -0.1056788 | 0.7461676 | 6.52867875 | 0.03822218 | 0.10758697 | FBA1 | XI | 326408 | 327487 | -1 |
| YKL061W | 49.8142281 | 0.28734543 | 0.59449596 | 0.48659589 | 0.78403788 | 0.86949041 | BLI1 | XI | 325771 | 326112 | 1 |
| YKL062W | 35.7656423 | -0.664135 | 0.72119394 | 2.31432721 | 0.31437661 | 0.48643522 | MSN4 | XI | 323228 | 325120 | 1 |
| YKL063C | 67.9059538 | 0.00795111 | 0.57679695 | 13.2471959 | 0.00132864 | 0.00828077 | NA | XI | 321015 | 321518 | -1 |
| YKL064W | 162.276258 | 0.1997388 | 0.49621545 | 0.56482134 | 0.75396399 | 0.85146314 | MNR2 | XI | 317764 | 320673 | 1 |
| YKL065C | 91.5077303 | 1.241347 | 0.58082581 | 10.2210433 | 0.00603293 | 0.026624 | YET1 | XI | 316437 | 317057 | -1 |
| YKL065W-A | 75.5022367 | 2.30417988 | 0.78716588 | 8.07303161 | 0.01765889 | 0.06020484 | NA | XI | 315954 | 316175 | 1 |
| YKL067W | 89.1332694 | 0.53713923 | 0.64521271 | 11.8621522 | 0.00265562 | 0.01388984 | YNK1 | XI | 314812 | 315273 | 1 |
| YKL068W | 54.3222013 | -0.2896035 | 0.59782392 | 2.21142144 | 0.33097557 | 0.50430225 | NUP100 | XI | 310199 | 313078 | 1 |
| YKL068W-A | 201.976807 | 0.00309979 | 0.51651846 | 1.32014029 | 0.51681508 | 0.67350835 | NA | XI | 309206 | 309442 | 1 |
| YKL069W | 22.8156411 | -0.112831 | 0.73040707 | 0.48590662 | 0.78430813 | 0.86949041 | NA | XI | 307285 | 307827 | 1 |
| YKL070W | 1613.72921 | -0.6346684 | 0.48228991 | 2.31268792 | 0.3146344 | 0.48657403 | NA | XI | 306211 | 306720 | 1 |
| YKL071W | 1114.3498 | -1.0834233 | 0.48753728 | 18.7138072 | 8.64E-05 | 0.00082519 | NA | XI | 305114 | 305884 | 1 |
| YKL072W | 42.0245934 | 0.2846195 | 0.61182585 | 2.02110684 | 0.36401747 | 0.53712058 | STB6 | XI | 299582 | 301882 | 1 |
| YKL073W | 58.2412234 | 0.15447424 | 0.65364892 | 0.22264047 | 0.89465221 | 0.9350658 | LHS1 | XI | 296430 | 299075 | 1 |
| YKL074C | 247.868265 | 1.09047021 | 0.55289549 | 24.9607313 | 3.80E-06 | 5.66E-05 | MUD2 | XI | 294610 | 296193 | -1 |
| YKL075C | 16.0987689 | -0.1177426 | 0.80709287 | 0.21557346 | 0.89781906 | 0.93705725 | NA | XI | 292956 | 294308 | -1 |
| YKL077W | 62.034363 | -0.8121872 | 0.57431301 | 8.8087484 | 0.01222375 | 0.04577978 | NA | XI | 291453 | 292631 | 1 |
| YKL078W | 12.0498656 | 0.00037788 | 0.97322522 | 2.29938058 | 0.31673485 | 0.48903861 | DHR2 | XI | 288845 | 291052 | 1 |
| YKL079W | 39.8644475 | 1.71296394 | 0.77714074 | 9.10291801 | 0.0105518 | 0.04108602 | SMY1 | XI | 286603 | 288573 | 1 |
| YKL080W | 31.5482334 | 0.51307715 | 0.69719505 | 1.01699308 | 0.60139908 | 0.74720908 | VMA5 | XI | 285030 | 286208 | 1 |
| YKL081W | 259.591801 | -1.1907381 | 0.62456662 | 22.0607639 | 1.62E-05 | 0.00019875 | TEF4 | XI | 282891 | 284455 | 1 |
| YKL082C | 185.362814 | 0.4276611 | 0.77845226 | 4.64157702 | 0.09819613 | 0.21634535 | RRP14 | XI | 281025 | 282329 | -1 |
| YKL084W | 15.0554161 | -0.5256424 | 0.80528139 | 0.53650643 | 0.76471412 | 0.85692899 | HOT13 | XI | 280509 | 280859 | 1 |
| YKL085W | 39.0998012 | -0.6968833 | 0.68928449 | 6.42403399 | 0.0402753 | 0.11200479 | MDH1 | XI | 279123 | 280127 | 1 |
| YKL086W | 12.39688 | 0.33307726 | 1.64417285 | 30.8417139 | 2.01E-07 | 4.39E-06 | SRX1 | XI | 278281 | 278664 | 1 |

| | | | | | | | | | | | |
|---|---|---|---|---|---|---|---|---|---|---|---|
| YKL087C | 95.2933564 | 0.85771792 | 0.71585029 | 1.85314148 | 0.39590906 | 0.56712357 | CYT2 | XI | 277189 | 277863 | -1 |
| YKL088W | 133.476211 | 0.16086517 | 0.52966395 | 1.37101797 | 0.50383372 | 0.66239719 | CAB3 | XI | 275283 | 276998 | 1 |
| YKL089W | 16.8083708 | 0.12469314 | 0.78671719 | 0.02718395 | 0.98649998 | 0.98889108 | MIF2 | XI | 273394 | 275043 | 1 |
| YKL090W | 922.955746 | 0.33099367 | 0.52806327 | 42.6273681 | 5.54E-10 | 2.12E-08 | CUE2 | XI | 271878 | 273209 | 1 |
| YKL091C | 33.6886452 | 0.46906058 | 0.7400642 | 2.70059014 | 0.25916378 | 0.43082351 | NA | XI | 269718 | 270650 | -1 |
| YKL092C | 34.8173936 | -0.9603572 | 0.67552142 | 4.7665354 | 0.09224864 | 0.20670265 | BUD2 | XI | 266145 | 269459 | -1 |
| YKL093W | 978.232845 | -0.7592783 | 0.53079015 | 2.52476853 | 0.28297853 | 0.45678444 | MBR1 | XI | 264789 | 265808 | 1 |
| YKL094W | 33.2746764 | 0.98959485 | 0.6632341 | 2.20348232 | 0.33229201 | 0.50604175 | YJU3 | XI | 263349 | 264290 | 1 |
| YKL095W | 11.3325143 | 0.84417927 | 0.93468973 | 2.12937352 | 0.34483585 | 0.51822506 | YJU2 | XI | 262277 | 263113 | 1 |
| YKL096C-B | 7435.67403 | 0.4441725 | 0.50096661 | 0.84448905 | 0.65557372 | 0.78495521 | NA | XI | 259073 | 259222 | -1 |
| YKL096W | 43.3363155 | 0.40431953 | 0.68079622 | 7.99127368 | 0.01839573 | 0.06210569 | CWP1 | XI | 261132 | 261851 | 1 |
| YKL096W-A | 129.678295 | 1.08150348 | 0.53816175 | 4.49266867 | 0.10578629 | 0.22837533 | CWP2 | XI | 259253 | 259531 | 1 |
| YKL097C | 10.7542647 | -0.6953198 | 1.19228062 | 13.8668632 | 0.00097465 | 0.0064494 | NA | XI | 258503 | 258913 | -1 |
| YKL098W | 25.000075 | 0.2194221 | 0.7868256 | 10.499337 | 0.00524926 | 0.02385369 | MTC2 | XI | 256770 | 257843 | 1 |
| YKL099C | 10.9280332 | -1.5932467 | 0.94126245 | 9.36477008 | 0.00925691 | 0.03719466 | UTP11 | XI | 255720 | 256472 | -1 |
| YKL100C | 18.7510872 | 0.22191504 | 0.94021281 | 0.61821 | 0.73410369 | 0.83775415 | YPF1 | XI | 253697 | 255460 | -1 |
| YKL101W | 349.959848 | 1.21543719 | 0.61181434 | 5.86702695 | 0.05320976 | 0.13698733 | HSL1 | XI | 248920 | 253476 | 1 |
| YKL102C | 879.324165 | 2.95584168 | 0.4937543 | 33.2337381 | 6.07E-08 | 1.55E-06 | NA | XI | 248062 | 248367 | -1 |
| YKL103C | 2186.15313 | 2.32023537 | 0.49609379 | 20.0201564 | 4.49E-05 | 0.00046888 | APE1 | XI | 246138 | 247682 | -1 |
| YKL104C | 163.242181 | 1.5830906 | 0.58015178 | 8.45349495 | 0.0145998 | 0.05250487 | GFA1 | XI | 243220 | 245373 | -1 |
| YKL104W-A | 18547.0612 | 1.77425852 | 0.48474888 | 12.9412672 | 0.00154824 | 0.00935734 | NA | NA | NA | NA | NA |
| YKL105C | 1664.78033 | -0.5517128 | 0.57402255 | 1.58909246 | 0.4517862 | 0.61806282 | SEG2 | XI | 239185 | 242583 | -1 |
| YKL106C-A | 38.9752528 | 3.07301367 | 0.78247157 | 27.1865065 | 1.25E-06 | 2.13E-05 | NA | XI | 237147 | 237266 | -1 |
| YKL106W | 115.550022 | 1.09136217 | 0.77532481 | 15.101156 | 0.00052581 | 0.00381506 | AAT1 | XI | 237536 | 238891 | 1 |
| YKL107W | 167.171718 | 0.35683027 | 0.50078285 | 0.7493911 | 0.68749856 | 0.80989148 | NA | XI | 236141 | 237070 | 1 |
| YKL108W | 33.8109918 | 0.45150594 | 0.78439489 | 18.5539955 | 9.36E-05 | 0.00088415 | SLD2 | XI | 234426 | 235787 | 1 |
| YKL109W | 109.415335 | -1.9163485 | 0.63444273 | 10.6783112 | 0.00479992 | 0.02212703 | HAP4 | XI | 232227 | 233891 | 1 |
| YKL110C | 11.0140622 | -0.3096097 | 0.94578908 | 0.29790132 | 0.86161163 | 0.91873505 | KTI12 | XI | 228939 | 229880 | -1 |
| YKL112W | 59.0976138 | -0.1608765 | 0.57063687 | 0.25154817 | 0.88181404 | 0.9300006 | ABF1 | XI | 226570 | 228765 | 1 |
| YKL113C | 162.026644 | 0.80310194 | 0.51286574 | 3.27887222 | 0.19408946 | 0.35096126 | RAD27 | XI | 224727 | 225875 | -1 |
| YKL114C | 274.778156 | 0.22809841 | 0.55508636 | 15.345871 | 0.00046525 | 0.00346692 | APN1 | XI | 223352 | 224455 | -1 |
| YKL116C | 65.8837633 | 0.27330673 | 0.61495701 | 7.41292302 | 0.02456429 | 0.07694787 | PRR1 | XI | 221344 | 222900 | -1 |
| YKL117W | 86.3725139 | 0.2934003 | 0.54219335 | 0.8818227 | 0.64344975 | 0.77686719 | SBA1 | XI | 220324 | 220974 | 1 |

| | | | | | | | | | | | |
|---|---|---|---|---|---|---|---|---|---|---|---|
| YKL119C | 5.37406981 | -1.3489497 | 1.35218611 | 1.21415906 | 0.54494003 | 0.69774497 | VPH2 | XI | 218570 | 219217 | -1 |
| YKL120W | 13.0325104 | -0.6027169 | 0.8579927 | 0.5270828 | 0.76832581 | 0.859302 | OAC1 | XI | 217344 | 218318 | 1 |
| YKL121W | 87.3362291 | 1.39914113 | 0.54442093 | 6.67338925 | 0.03555428 | 0.10170914 | DGR2 | XI | 214142 | 216700 | 1 |
| YKL122C | 41.1769972 | 0.64417809 | 0.72221089 | 3.43914898 | 0.17914236 | 0.3333015 | SRP21 | XI | 212495 | 212998 | -1 |
| YKL124W | 106.862488 | -0.7128457 | 0.54710357 | 7.75203389 | 0.02073324 | 0.06786064 | SSH4 | XI | 210593 | 212332 | 1 |
| YKL125W | 50.6569691 | 0.25799307 | 0.6477876 | 3.62752415 | 0.16303961 | 0.31175673 | RRN3 | XI | 208247 | 210130 | 1 |
| YKL126W | 27.1494538 | 1.01403295 | 0.71676923 | 2.16047983 | 0.33951406 | 0.51259098 | YPK1 | XI | 205707 | 207749 | 1 |
| YKL127W | 252.717239 | -1.0816652 | 0.50507369 | 6.63239267 | 0.03629061 | 0.1034068 | PGM1 | XI | 203541 | 205253 | 1 |
| YKL128C | 684.588182 | 1.04444904 | 0.54640087 | 13.2969469 | 0.001296 | 0.00812103 | PMU1 | XI | 200884 | 201771 | -1 |
| YKL129C | 160.925232 | -1.2317823 | 0.50640032 | 6.76767824 | 0.03391699 | 0.09824187 | MYO3 | XI | 196349 | 200167 | -1 |
| YKL130C | 73.9642489 | -0.3197494 | 0.63435366 | 0.25436246 | 0.88057407 | 0.92970895 | SHE2 | XI | 195288 | 196028 | -1 |
| YKL132C | 22.3893949 | 0.42172582 | 0.7335826 | 2.56000038 | 0.27803725 | 0.45261809 | RMA1 | XI | 193574 | 194866 | -1 |
| YKL133C | 58.6644849 | -0.4300637 | 0.57331927 | 1.95435524 | 0.37637187 | 0.54998972 | NA | XI | 191679 | 193070 | -1 |
| YKL134C | 140.398003 | -0.7854375 | 0.50955285 | 3.41875732 | 0.18097821 | 0.33553116 | 1-Oct | XI | 189124 | 191442 | -1 |
| YKL135C | 153.693717 | 0.61845704 | 0.64154381 | 9.75272074 | 0.00762471 | 0.03199065 | APL2 | XI | 186460 | 188640 | -1 |
| YKL137W | 35.7339311 | -0.3929039 | 0.62361234 | 1.72400035 | 0.42231653 | 0.59293772 | CMC1 | XI | 185957 | 186292 | 1 |
| YKL138C | 17.3537142 | 1.01737223 | 0.9388051 | 1.71141747 | 0.42498188 | 0.59510585 | MRPL31 | XI | 185286 | 185681 | -1 |
| YKL138C-A | 29.1878324 | -0.4821472 | 0.65968909 | 0.80053615 | 0.67014038 | 0.79536351 | HSK3 | XI | 184803 | 185012 | -1 |
| YKL139W | 238.422738 | 2.40906376 | 0.84137321 | 8.44919768 | 0.0146312 | 0.05258366 | CTK1 | XI | 182958 | 184544 | 1 |
| YKL140W | 44.7127721 | 0.28840212 | 0.62776689 | 9.66820143 | 0.00795384 | 0.03308385 | TGL1 | XI | 180779 | 182425 | 1 |
| YKL141W | 32.827867 | 0.56492588 | 0.71811127 | 4.83015365 | 0.08936047 | 0.20267482 | SDH3 | XI | 179667 | 180263 | 1 |
| YKL142W | 24.4588257 | 1.44009817 | 0.77898593 | 3.5157943 | 0.17240703 | 0.32378745 | MRP8 | XI | 178515 | 179174 | 1 |
| YKL143W | 28.0490197 | -1.0373465 | 0.73235172 | 5.92419277 | 0.0517104 | 0.13420135 | LTV1 | XI | 176781 | 178172 | 1 |
| YKL144C | 277.927894 | -0.6184174 | 0.51031458 | 12.5760815 | 0.0018584 | 0.01061156 | RPC25 | XI | 175843 | 176481 | -1 |
| YKL145W | 114.279614 | 0.35873508 | 0.7676115 | 0.68126618 | 0.71131985 | 0.82438173 | RPT1 | XI | 174213 | 175616 | 1 |
| YKL146W | 69.8992525 | 0.42590699 | 0.65540709 | 0.4777698 | 0.78750552 | 0.87066201 | AVT3 | XI | 171783 | 173861 | 1 |
| YKL148C | 24.9651322 | 0.12065005 | 0.88609863 | 0.04482207 | 0.97783823 | 0.98293113 | SDH1 | XI | 169207 | 171129 | -1 |
| YKL149C | 50.9182633 | 1.92758385 | 0.73093141 | 22.698299 | 1.18E-05 | 0.00015292 | DBR1 | XI | 167612 | 168829 | -1 |
| YKL150W | 50.1128829 | 0.30972368 | 0.63012579 | 13.9835791 | 0.0009194 | 0.00615027 | MCR1 | XI | 166544 | 167452 | 1 |
| YKL151C | 616.280956 | -0.3654195 | 0.5565489 | 6.73998323 | 0.03438993 | 0.09926596 | NNR2 | XI | 164917 | 165930 | -1 |
| YKL152C | 433.030351 | 0.62509463 | 0.57410749 | 1.26494718 | 0.53127602 | 0.68570846 | GPM1 | XI | 163642 | 164385 | -1 |
| YKL154W | 12.9152514 | 1.81507239 | 1.19478529 | 8.63766939 | 0.01331539 | 0.04876415 | SRP102 | XI | 161601 | 162335 | 1 |
| YKL155C | 55.0812062 | -0.1505711 | 0.64699841 | 15.6030882 | 0.0004091 | 0.00312907 | RSM22 | XI | 159456 | 161342 | -1 |

| | | | | | | | | | | | |
|---|---|---|---|---|---|---|---|---|---|---|---|
| YKL156W | 12.8000601 | -0.7130301 | 0.84886322 | 5.01464042 | 0.08148631 | 0.1887223 | RPS27A | XI | 158613 | 159212 | 1 |
| YKL157W | 80.7514138 | -0.3459039 | 0.55964395 | 0.59869029 | 0.74130351 | 0.84242342 | APE2 | XI | 154991 | 158232 | 1 |
| YKL159C | 340.94678 | -0.678012 | 0.49982501 | 20.32271 | 3.86E-05 | 0.00041734 | RCN1 | XI | 153816 | 154451 | -1 |
| YKL160W | 182.560507 | 1.7038866 | 0.54832173 | 15.3679378 | 0.00046014 | 0.0034333 | ELF1 | XI | 153269 | 153706 | 1 |
| YKL161C | 39.3253805 | -0.2365532 | 0.69282828 | 0.54686612 | 0.76076326 | 0.85483951 | KDX1 | XI | 149386 | 150687 | -1 |
| YKL163W | 94.9762866 | 1.73104836 | 0.81655889 | 6.49172319 | 0.038935 | 0.10943382 | PIR3 | XI | 144401 | 145378 | 1 |
| YKL164C | 195.908905 | 1.25838298 | 0.49765305 | 6.91371965 | 0.03152861 | 0.09285753 | PIR1 | XI | 141794 | 142819 | -1 |
| YKL165C | 77.2701226 | 0.78474951 | 0.60640561 | 8.20801748 | 0.01650637 | 0.05733167 | MCD4 | XI | 137932 | 140691 | -1 |
| YKL166C | 17.8872574 | -0.4842131 | 0.79064786 | 0.39779448 | 0.81963412 | 0.89171017 | TPK3 | XI | 134509 | 135705 | -1 |
| YKL167C | 6.41677326 | 0.82966696 | 1.14664855 | 1.05628848 | 0.58969829 | 0.73728204 | MRP49 | XI | 133721 | 134134 | -1 |
| YKL168C | 22.093188 | 0.03520402 | 0.74223309 | 1.50462394 | 0.47127572 | 0.63325282 | KKQ8 | XI | 131288 | 133462 | -1 |
| YKL170W | 99.5446691 | 0.13737589 | 0.55394697 | 0.72140966 | 0.69718475 | 0.816815 | MRPL38 | XI | 130634 | 131050 | 1 |
| YKL171W | 413.976702 | -0.0424363 | 0.50498135 | 3.84703004 | 0.14609254 | 0.28741957 | NNK1 | XI | 127475 | 130261 | 1 |
| YKL172W | 95.8421433 | -0.7901311 | 0.55017078 | 23.1086106 | 9.59E-06 | 0.00012771 | EBP2 | XI | 125759 | 127042 | 1 |
| YKL173W | 21.8903198 | 0.39886376 | 0.71555535 | 0.62938175 | 0.73001451 | 0.83678999 | SNU114 | XI | 122517 | 125543 | 1 |
| YKL174C | 248.494257 | 1.96641247 | 0.67762065 | 8.54133698 | 0.01397244 | 0.05075309 | TPO5 | XI | 120380 | 122236 | -1 |
| YKL175W | 433.035587 | 3.01906068 | 0.91026361 | 10.1441256 | 0.00626947 | 0.02746253 | ZRT3 | XI | 118793 | 120304 | 1 |
| YKL176C | 183.431732 | -0.0703675 | 0.50130281 | 5.03721076 | 0.0805719 | 0.18735393 | LST4 | XI | 115500 | 117986 | -1 |
| YKL178C | 42.7980751 | -0.1266906 | 0.65143205 | 0.96181617 | 0.61822174 | 0.75949583 | STE3 | XI | 113215 | 114627 | -1 |
| YKL179C | 21.3681606 | 0.83102324 | 0.72514935 | 1.67666057 | 0.43243196 | 0.60114791 | COY1 | XI | 110464 | 112503 | -1 |
| YKL180W | 80.0674227 | -0.3604169 | 0.61647492 | 17.1991865 | 0.00018418 | 0.00158456 | RPL17A | XI | 109269 | 110129 | 1 |
| YKL181W | 56.6891656 | -0.3315467 | 0.60656445 | 0.86470051 | 0.64898203 | 0.7805914 | PRS1 | XI | 107316 | 108599 | 1 |
| YKL182W | 130.609291 | 0.16143017 | 0.7144547 | 0.08443682 | 0.95866038 | 0.97090145 | FAS1 | XI | 100671 | 106826 | 1 |
| YKL183C-A | 601.551244 | 0.30126858 | 0.63017761 | 0.88000448 | 0.64403498 | 0.77686719 | NA | XI | 98395 | 98607 | -1 |
| YKL183W | 21.8277522 | -1.2668836 | 0.78431287 | 3.45017362 | 0.17815759 | 0.33157583 | LOT5 | XI | 98721 | 99641 | 1 |
| YKL184W | 10.8236499 | -0.0083546 | 1.11527534 | 0.81052205 | 0.66680274 | 0.79342126 | SPE1 | XI | 96757 | 98157 | 1 |
| YKL185W | 55.8313792 | 0.46976382 | 0.66572701 | 0.59108314 | 0.74412848 | 0.84379317 | ASH1 | XI | 94499 | 96265 | 1 |
| YKL186C | 53.6163024 | 0.95014439 | 0.70650375 | 1.96330766 | 0.37469091 | 0.54798191 | MTR2 | XI | 92744 | 93298 | -1 |
| YKL187C | 238.343719 | 0.72042017 | 0.53539166 | 10.9729951 | 0.00414233 | 0.01969412 | FAT3 | XI | 89284 | 91536 | -1 |
| YKL188C | 994.381827 | 2.59322908 | 0.49761918 | 64.6574948 | 9.12E-15 | 9.10E-13 | PXA2 | XI | 86225 | 88786 | -1 |
| YKL189W | 19.7740721 | 1.82785357 | 0.88628491 | 14.2507426 | 0.00080443 | 0.00551204 | HYM1 | XI | 84704 | 85903 | 1 |
| YKL190W | 23.381838 | 0.18401457 | 0.93687298 | 0.18667932 | 0.91088406 | 0.94533405 | CNB1 | XI | 82947 | 83550 | 1 |
| YKL191W | 37.3742788 | -1.0013179 | 0.75324183 | 6.10332741 | 0.0472802 | 0.12530377 | DPH2 | XI | 81035 | 82639 | 1 |

| | | | | | | | | | | | |
|---|---|---|---|---|---|---|---|---|---|---|---|
| YKL192C | 301.428545 | 1.24570298 | 0.58599543 | 74.1717031 | 7.83E-17 | 1.08E-14 | ACP1 | XI | 80160 | 80537 | -1 |
| YKL193C | 12.2779565 | 1.32616568 | 0.94735827 | 4.25455789 | 0.1191611 | 0.24840182 | SDS22 | XI | 78866 | 79882 | -1 |
| YKL194C | 14.7588775 | 0.26861585 | 0.9044548 | 3.13452167 | 0.20861583 | 0.37120027 | MST1 | XI | 77258 | 78646 | -1 |
| YKL195W | 116.339296 | 0.29867312 | 0.50519226 | 0.67314143 | 0.71421538 | 0.82607012 | MIA40 | XI | 75821 | 77032 | 1 |
| YKL196C | 14.9451767 | -0.3745025 | 0.86074824 | 0.93693527 | 0.62596073 | 0.76430043 | YKT6 | XI | 74932 | 75534 | -1 |
| YKL197C | 58.6378411 | -0.0685295 | 0.59508259 | 5.0624278 | 0.07956238 | 0.18567762 | PEX1 | XI | 70734 | 73865 | -1 |
| YKL198C | 203.987887 | -1.0555522 | 0.50784521 | 14.2312941 | 0.00081229 | 0.00555932 | PTK1 | XI | 68232 | 70220 | -1 |
| YKL201C | 1426.57059 | 0.1795968 | 0.68485109 | 4.25277791 | 0.1192672 | 0.24840182 | MNN4 | XI | 63930 | 67466 | -1 |
| YKL203C | 31.4818697 | 0.68310637 | 0.64955873 | 3.36893663 | 0.18554306 | 0.34147943 | TOR2 | XI | 55935 | 63359 | -1 |
| YKL204W | 64.4175739 | 0.83652983 | 0.75082857 | 1.21817013 | 0.54384823 | 0.69714961 | EAP1 | XI | 53704 | 55602 | 1 |
| YKL205W | 23.5982772 | -0.6763028 | 0.71479522 | 2.24729226 | 0.3250923 | 0.49807536 | LOS1 | XI | 50051 | 53353 | 1 |
| YKL206C | 147.675493 | 2.37430862 | 0.93559948 | 9.40810165 | 0.00905851 | 0.03655068 | ADD66 | XI | 49006 | 49809 | -1 |
| YKL207W | 177.564364 | 2.46758085 | 0.79915227 | 13.3118183 | 0.0012864 | 0.00807836 | EMC3 | XI | 48194 | 48955 | 1 |
| YKL209C | 107.113163 | -0.9122254 | 0.53728755 | 3.72625902 | 0.15518621 | 0.30010961 | STE6 | XI | 42423 | 46295 | -1 |
| YKL210W | 52.9046751 | -0.0755835 | 0.68976961 | 0.3367829 | 0.84502299 | 0.90911019 | UBA1 | XI | 39163 | 42237 | 1 |
| YKL211C | 37.8750347 | -0.7786214 | 0.71896688 | 12.0328717 | 0.00243834 | 0.01297612 | TRP3 | XI | 36699 | 38153 | -1 |
| YKL212W | 61.7077875 | -0.9762473 | 0.57233433 | 6.80159174 | 0.03334672 | 0.09707265 | SAC1 | XI | 34543 | 36414 | 1 |
| YKL213C | 32.7376414 | 0.42853401 | 0.64410866 | 0.46282253 | 0.7934131 | 0.87452157 | DOA1 | XI | 31960 | 34107 | -1 |
| YKL214C | 32.5350753 | 0.10441976 | 0.65129125 | 0.14582891 | 0.92968035 | 0.95576159 | YRA2 | XI | 31082 | 31693 | -1 |
| YKL215C | 12.2519431 | 0.27141251 | 0.99466929 | 1.29445746 | 0.52349451 | 0.6789949 | OXP1 | XI | 26827 | 30687 | -1 |
| YKL216W | 19.8139908 | -0.437229 | 0.98058246 | 2.27725649 | 0.32025804 | 0.49277014 | URA1 | XI | 25215 | 26159 | 1 |
| YKL217W | 493.704277 | 0.40280756 | 0.55685641 | 7.63018919 | 0.02203563 | 0.07124193 | JEN1 | XI | 22234 | 24084 | 1 |
| YKL218C | 150.194662 | -0.2927653 | 0.51388262 | 1.6028215 | 0.44869552 | 0.61504428 | SRY1 | XI | 17359 | 18339 | -1 |
| YKL219W | 62.3996282 | -0.8679537 | 0.56939504 | 5.7305696 | 0.0569669 | 0.14462713 | COS9 | XI | 14485 | 15708 | 1 |
| YKL220C | 55.9706533 | -1.0023608 | 0.65188941 | 2.42160177 | 0.29795855 | 0.47161883 | FRE2 | XI | 9091 | 11226 | -1 |
| YKL221W | 193.874728 | -1.2660937 | 0.63876885 | 13.0590455 | 0.0014597 | 0.00894319 | MCH2 | XI | 6107 | 7528 | 1 |
| YKL222C | 44.8626822 | -0.0882095 | 0.69329445 | 6.25739567 | 0.04377476 | 0.11860359 | NA | XI | 3503 | 5620 | -1 |
| YKR001C | 68.4676191 | 0.7752188 | 0.58943367 | 2.15107892 | 0.34111369 | 0.51473762 | VPS1 | XI | 440608 | 442722 | -1 |
| YKR002W | 129.234309 | 1.10858676 | 0.58015809 | 34.2338539 | 3.68E-08 | 9.83E-07 | PAP1 | XI | 443232 | 444938 | 1 |
| YKR003W | 172.702423 | -0.6483055 | 0.48818739 | 3.02767896 | 0.22006342 | 0.38460377 | OSH6 | XI | 445381 | 446727 | 1 |
| YKR004C | 609.37646 | 2.70728522 | 0.92822499 | 12.3086527 | 0.00212427 | 0.01170082 | ECM9 | XI | 446799 | 448036 | -1 |
| YKR005C | 34.4496057 | 1.98949142 | 0.77453445 | 12.931382 | 0.00155592 | 0.00937506 | NA | XI | 448521 | 450167 | -1 |
| YKR007W | 31.9795813 | -0.5948725 | 0.64471077 | 4.50846439 | 0.1049541 | 0.22691719 | MEH1 | XI | 451434 | 451988 | 1 |

| | | | | | | | | | | | |
|---|---|---|---|---|---|---|---|---|---|---|---|
| YKR008W | 8.06518715 | 0.27914542 | 1.17192911 | 1.08146535 | 0.58232144 | 0.73074147 | RSC4 | XI | 452201 | 454078 | 1 |
| YKR009C | 1361.33197 | 0.55166485 | 0.51175693 | 10.7715671 | 0.00458125 | 0.02123734 | FOX2 | XI | 454352 | 457054 | -1 |
| YKR010C | 11.7927766 | -0.4232691 | 0.858278 | 0.4983563 | 0.7794411 | 0.8662119 | TOF2 | XI | 458924 | 461239 | -1 |
| YKR011C | 469.801923 | -1.2688504 | 0.48903899 | 27.5834387 | 1.02E-06 | 1.80E-05 | NA | XI | 461635 | 462696 | -1 |
| YKR013W | 392.440991 | 0.31091531 | 0.46786738 | 0.50113292 | 0.77835975 | 0.86534235 | PRY2 | XI | 463959 | 464948 | 1 |
| YKR014C | 46.2561887 | 0.80125654 | 0.64570795 | 1.64588164 | 0.43913833 | 0.60798922 | YPT52 | XI | 465367 | 466071 | -1 |
| YKR015C | 1633.46959 | 0.35877986 | 0.51945529 | 2.65739012 | 0.26482261 | 0.43721783 | NA | XI | 466961 | 468667 | -1 |
| YKR016W | 19.9458496 | 0.4080069 | 0.73890144 | 0.72167514 | 0.69709222 | 0.816815 | MIC60 | XI | 469717 | 471339 | 1 |
| YKR017C | 156.287335 | -0.2928154 | 0.5562529 | 2.59470582 | 0.27325417 | 0.44663164 | HEL1 | XI | 471694 | 473349 | -1 |
| YKR018C | 43.207686 | -0.3084675 | 0.66611621 | 0.36255822 | 0.83420249 | 0.90213531 | NA | XI | 473723 | 475900 | -1 |
| YKR019C | 76.1839461 | -0.4021313 | 0.57762604 | 0.55917136 | 0.75609694 | 0.85254164 | IRS4 | XI | 476216 | 478063 | -1 |
| YKR020W | 28.8894314 | 1.25368134 | 0.74608759 | 6.21627752 | 0.04468405 | 0.1201675 | VPS51 | XI | 478338 | 478832 | 1 |
| YKR021W | 75.7263776 | 0.20595058 | 0.69782402 | 10.0564336 | 0.00655048 | 0.02840995 | ALY1 | XI | 479234 | 481981 | 1 |
| YKR022C | 36.4194236 | -0.0735147 | 0.67891836 | 4.78989428 | 0.0911775 | 0.20517595 | NTR2 | XI | 482171 | 483139 | -1 |
| YKR023W | 46.8242679 | -0.9094837 | 0.63909643 | 4.19326351 | 0.12286959 | 0.25402775 | NA | XI | 483419 | 485011 | 1 |
| YKR024C | 13.6310549 | 0.1452199 | 0.90226108 | 0.1038283 | 0.94941037 | 0.96524777 | DBP7 | XI | 485144 | 487372 | -1 |
| YKR025W | 353.531283 | 0.7414352 | 0.63030708 | 9.10890053 | 0.01052028 | 0.04099087 | RPC37 | XI | 487770 | 488618 | 1 |
| YKR026C | 42.8567737 | 0.13360772 | 0.68222164 | 6.71724558 | 0.03478313 | 0.10004686 | GCN3 | XI | 488738 | 489655 | -1 |
| YKR027W | 16.4579408 | -0.1875219 | 0.82181102 | 0.10483132 | 0.94893435 | 0.96493324 | BCH2 | XI | 491364 | 493661 | 1 |
| YKR028W | 35.1784014 | 0.28632808 | 0.68744136 | 2.25072125 | 0.32453541 | 0.49763242 | SAP190 | XI | 494257 | 497358 | 1 |
| YKR029C | 31.2181708 | 0.10592578 | 0.69425682 | 4.884581 | 0.08696144 | 0.19877881 | SET3 | XI | 497578 | 499833 | -1 |
| YKR030W | 17.529784 | -0.6614919 | 0.84618466 | 5.59527788 | 0.06095381 | 0.15179465 | GMH1 | XI | 500282 | 501103 | 1 |
| YKR031C | 26.8621012 | -0.4638496 | 0.75273926 | 4.59590907 | 0.10046413 | 0.2196191 | SPO14 | XI | 501344 | 506395 | -1 |
| YKR034W | 6.55252156 | -1.6624937 | 1.13669922 | 2.70165182 | 0.25902624 | 0.43082351 | DAL80 | XI | 506898 | 507707 | 1 |
| YKR035W-A | 83.9636657 | -1.010746 | 0.53540681 | 17.5217136 | 0.00015675 | 0.00138141 | DID2 | XI | 507939 | 508553 | 1 |
| YKR036C | 34.7803364 | 0.07003389 | 0.65481711 | 3.29989171 | 0.19206031 | 0.34838007 | CAF4 | XI | 508702 | 510633 | -1 |
| YKR037C | 24.1793831 | -0.8055399 | 0.78868572 | 1.16894484 | 0.55739986 | 0.70899499 | SPC34 | XI | 510910 | 511797 | -1 |
| YKR038C | 24.0137057 | 0.30650175 | 0.7389588 | 0.71265746 | 0.70024239 | 0.81801018 | KAE1 | XI | 511999 | 513159 | -1 |
| YKR039W | 39.9995743 | -0.0630604 | 0.71611685 | 11.1498736 | 0.00379172 | 0.01838696 | GAP1 | XI | 515063 | 516871 | 1 |
| YKR041W | 8.16494176 | -0.4772339 | 1.05815964 | 1.24127539 | 0.5376015 | 0.69208525 | NA | XI | 518198 | 518950 | 1 |
| YKR042W | 112.956815 | 0.48181817 | 0.6774656 | 0.51132065 | 0.77440497 | 0.86282905 | UTH1 | XI | 519527 | 520624 | 1 |
| YKR043C | 140.486104 | -0.183088 | 0.50668696 | 1.32923371 | 0.51447061 | 0.67150244 | SHB17 | XI | 520897 | 521712 | -1 |
| YKR044W | 1150.46441 | -1.125083 | 0.47581462 | 15.865161 | 0.00035886 | 0.00283852 | UIP5 | XI | 522015 | 523346 | 1 |

| | | | | | | | | | | | |
|---|---|---|---|---|---|---|---|---|---|---|---|
| YKR045C | 79.4365272 | -0.5660148 | 0.54327559 | 13.4161528 | 0.00122101 | 0.00776032 | NA | XI | 523418 | 523969 | -1 |
| YKR046C | 1992.18604 | -1.8719371 | 0.47247595 | 45.8506971 | 1.11E-10 | 5.16E-09 | PET10 | XI | 524223 | 525074 | -1 |
| YKR048C | 200.983737 | 0.55235535 | 0.55600437 | 1.88459376 | 0.38973164 | 0.56101577 | NAP1 | XI | 525387 | 526640 | -1 |
| YKR049C | 391.686433 | 0.26541148 | 0.57054119 | 0.2988971 | 0.86118275 | 0.91844688 | FMP46 | XI | 526830 | 527231 | -1 |
| YKR050W | 25.897321 | 0.82262852 | 0.78667174 | 1.08600238 | 0.58100194 | 0.72940182 | TRK2 | XI | 527815 | 530484 | 1 |
| YKR051W | 152.527023 | 1.51101843 | 0.73999993 | 6.3635487 | 0.04151193 | 0.11429106 | NA | XI | 530946 | 532202 | 1 |
| YKR052C | 12.407362 | -0.120923 | 0.95599203 | 0.50853224 | 0.7754854 | 0.8634877 | MRS4 | XI | 532550 | 533464 | -1 |
| YKR053C | 482.630563 | -0.3768514 | 0.47863636 | 0.89051888 | 0.64065804 | 0.77561669 | YSR3 | XI | 534067 | 535281 | -1 |
| YKR054C | 98.5596393 | -0.4364778 | 0.52578478 | 2.22986439 | 0.32793751 | 0.50041094 | DYN1 | XI | 535647 | 547925 | -1 |
| YKR056W | 73.9645614 | 1.25397513 | 0.59506182 | 8.55653701 | 0.01386665 | 0.05043211 | TRM2 | XI | 549448 | 551367 | 1 |
| YKR057W | 27.6511602 | -1.4672326 | 0.76524115 | 18.8966499 | 7.88E-05 | 0.00076317 | RPS21A | XI | 551657 | 552242 | 1 |
| YKR058W | 26.732947 | -1.1050485 | 0.67313473 | 8.52011467 | 0.01412149 | 0.05110095 | GLG1 | XI | 552770 | 554620 | 1 |
| YKR059W | 19.4387559 | -0.3947605 | 0.7143342 | 8.75408646 | 0.01256245 | 0.04680603 | TIF2 | XI | 554987 | 556174 | 1 |
| YKR060W | 50.0613308 | 1.55854691 | 0.6213321 | 6.26887883 | 0.04352415 | 0.11820113 | UTP30 | XI | 556518 | 557342 | 1 |
| YKR061W | 126.33766 | -0.6621611 | 0.51695801 | 8.74983386 | 0.01258919 | 0.04684525 | KTR2 | XI | 557677 | 558954 | 1 |
| YKR062W | 182.607888 | -1.0770441 | 0.50152634 | 19.0588848 | 7.27E-05 | 0.00071568 | TFA2 | XI | 559666 | 560652 | 1 |
| YKR063C | 739.036832 | 0.15771452 | 0.56560537 | 4.28465863 | 0.11738111 | 0.24546419 | LAS1 | XI | 560789 | 562297 | -1 |
| YKR064W | 41.9874984 | 0.88920429 | 0.71928152 | 4.17870194 | 0.12376744 | 0.25551406 | OAF3 | XI | 562547 | 565138 | 1 |
| YKR065C | 15.8109404 | 1.56915031 | 0.92427121 | 4.3655456 | 0.11272852 | 0.23918753 | PAM17 | XI | 565299 | 565892 | -1 |
| YKR066C | 44.8801936 | 0.34622556 | 0.67807397 | 1.16783855 | 0.55770826 | 0.70907501 | CCP1 | XI | 566113 | 567198 | -1 |
| YKR067W | 146.561829 | 0.88132834 | 0.5756771 | 42.3726963 | 6.29E-10 | 2.37E-08 | GPT2 | XI | 567918 | 570149 | 1 |
| YKR068C | 653.86651 | 2.15731232 | 0.52035645 | 25.1865678 | 3.39E-06 | 5.12E-05 | BET3 | XI | 570328 | 570909 | -1 |
| YKR069W | 95.8746973 | 0.31543478 | 0.61561337 | 22.6389729 | 1.21E-05 | 0.00015675 | MET1 | XI | 571612 | 573393 | 1 |
| YKR070W | 65.1960973 | 0.05132493 | 0.63113493 | 1.81713027 | 0.40310221 | 0.57401913 | NA | XI | 573574 | 574632 | 1 |
| YKR071C | 65.1579686 | -0.2644986 | 0.59691939 | 5.54412962 | 0.06253275 | 0.15457697 | DRE2 | XI | 574934 | 575980 | -1 |
| YKR072C | 353.91002 | 1.14341825 | 0.57206059 | 4.8641233 | 0.08785552 | 0.20034795 | SIS2 | XI | 576435 | 578123 | -1 |
| YKR073C | 78.9742015 | 0.21104632 | 0.64434614 | 8.24321325 | 0.01621844 | 0.0567598 | NA | XI | 578187 | 578507 | -1 |
| YKR075C | 64.8983632 | -0.5423712 | 0.59040229 | 5.57715715 | 0.06150858 | 0.15245492 | NA | XI | 579827 | 580750 | -1 |
| YKR076W | 48.0971875 | -1.0424865 | 0.67576927 | 8.23088273 | 0.01631874 | 0.05693318 | ECM4 | XI | 582283 | 583395 | 1 |
| YKR077W | 57.3494246 | -1.2159136 | 0.60949771 | 9.09687109 | 0.01058375 | 0.04118273 | MSA2 | XI | 583653 | 584744 | 1 |
| YKR078W | 83.940083 | -0.6711321 | 0.61762847 | 10.3869777 | 0.0055526 | 0.02490283 | NA | XI | 584952 | 586709 | 1 |
| YKR079C | 160.534679 | -0.3716033 | 0.51277084 | 1.50650132 | 0.47083354 | 0.63296713 | TRZ1 | XI | 586789 | 589305 | -1 |
| YKR080W | 6.11961604 | -1.685434 | 1.21050421 | 4.0998286 | 0.12874594 | 0.26238612 | MTD1 | XI | 590395 | 591357 | 1 |

| | | | | | | | | | | | |
|---|---|---|---|---|---|---|---|---|---|---|---|
| YKR081C | 51.8453539 | -0.6578426 | 0.7276166 | 13.8131881 | 0.00100116 | 0.00658719 | RPF2 | XI | 591506 | 592540 | -1 |
| YKR082W | 57.7524104 | 1.30704543 | 0.71931417 | 8.72457462 | 0.01274919 | 0.04728881 | NUP133 | XI | 592825 | 596298 | 1 |
| YKR083C | 33.4870702 | -0.3265138 | 0.702078 | 7.42122376 | 0.02446255 | 0.07676865 | DAD2 | XI | 596421 | 596822 | -1 |
| YKR084C | 15.5883272 | -0.2214706 | 0.81118594 | 0.42703108 | 0.80773961 | 0.88441988 | HBS1 | XI | 597055 | 598890 | -1 |
| YKR085C | 7.43798492 | -1.0653482 | 1.19299616 | 1.06680159 | 0.58660665 | 0.73405068 | MRPL20 | XI | 599094 | 599681 | -1 |
| YKR086W | 34.6164534 | -0.3657786 | 0.65743206 | 0.94619231 | 0.62307016 | 0.76286238 | PRP16 | XI | 599857 | 603072 | 1 |
| YKR087C | 91.9285746 | -0.4866842 | 0.53722564 | 8.41248301 | 0.01490227 | 0.05319615 | OMA1 | XI | 603195 | 604232 | -1 |
| YKR088C | 34.2129609 | 0.49354871 | 0.79971611 | 0.6189664 | 0.7338261 | 0.83775415 | TVP38 | XI | 604405 | 605418 | -1 |
| YKR089C | 289.977292 | -0.4130218 | 0.48511031 | 1.10952374 | 0.57420898 | 0.7240135 | TGL4 | XI | 605633 | 608365 | -1 |
| YKR090W | 759.347307 | 1.29248643 | 0.48784601 | 13.2168547 | 0.00134895 | 0.00838929 | PXL1 | XI | 608946 | 611066 | 1 |
| YKR091W | 53.3464664 | 1.42933296 | 0.61650681 | 6.47956026 | 0.03917251 | 0.10988799 | SRL3 | XI | 611525 | 612265 | 1 |
| YKR092C | 44.7025307 | -0.7009325 | 0.76739068 | 0.97537035 | 0.61404616 | 0.75677464 | SRP40 | XI | 612665 | 613885 | -1 |
| YKR093W | 31.9544664 | -0.6570547 | 0.69639392 | 1.29351893 | 0.52374023 | 0.67900939 | PTR2 | XI | 615730 | 617535 | 1 |
| YKR094C | 60.8152594 | -0.6225913 | 0.67972163 | 16.2427813 | 0.00029712 | 0.0024298 | RPL40B | XI | 617996 | 618750 | -1 |
| YKR095W | 91.8030771 | 0.97477747 | 0.63241198 | 5.4003421 | 0.06719402 | 0.16286148 | MLP1 | XI | 619805 | 625432 | 1 |
| YKR095W-A | 35.1904138 | 0.41809143 | 0.85578134 | 1.82974576 | 0.40056754 | 0.57138844 | PCC1 | XI | 625864 | 626205 | 1 |
| YKR096W | 216.104016 | 1.58318342 | 0.55643665 | 46.024149 | 1.01E-10 | 4.77E-09 | ESL2 | XI | 626793 | 630380 | 1 |
| YKR097W | 86.6886956 | 0.23057426 | 0.5409683 | 5.91371151 | 0.0519821 | 0.13454465 | PCK1 | XI | 631152 | 632801 | 1 |
| YKR098C | 18.7655514 | 1.48420949 | 0.83955031 | 3.12617782 | 0.20948798 | 0.37244958 | UBP11 | XI | 633026 | 635179 | -1 |
| YKR099W | 19.6074458 | -0.6384947 | 0.72942331 | 1.43801012 | 0.48723679 | 0.64903366 | BAS1 | XI | 635851 | 638286 | 1 |
| YKR100C | 29.487636 | 0.34602986 | 0.69579258 | 1.59428335 | 0.45061513 | 0.61694528 | SKG1 | XI | 638901 | 639968 | -1 |
| YKR101W | 24.0404526 | 0.3291909 | 0.951896 | 20.7688628 | 3.09E-05 | 0.00034617 | SIR1 | XI | 640540 | 642504 | 1 |
| YKR102W | 378.193542 | 1.34025289 | 0.51306639 | 50.5184344 | 1.07E-11 | 6.20E-10 | FLO10 | XI | 646356 | 649865 | 1 |
| YKR103W | 28.4768296 | -0.1568473 | 0.75849516 | 10.2372454 | 0.00598426 | 0.02644951 | NFT1 | XI | 653080 | 656736 | 1 |
| YKR105C | 648.103478 | -0.5773444 | 0.48419387 | 1.70776282 | 0.42575918 | 0.5957336 | VBA5 | XI | 658716 | 660464 | -1 |
| YKR106W | 264.777876 | -0.0902633 | 0.51686007 | 0.1370999 | 0.93374682 | 0.95735444 | GEX2 | XI | 661442 | 663289 | 1 |
| YLL001W | 235.588272 | 1.11514631 | 0.57100091 | 47.385459 | 5.13E-11 | 2.56E-09 | DNM1 | XII | 147890 | 150163 | 1 |
| YLL002W | 33.1336444 | -1.055134 | 0.66992331 | 3.21921423 | 0.19996616 | 0.35900663 | RTT109 | XII | 146291 | 147601 | 1 |
| YLL003W | 20.5157482 | 0.20565834 | 0.74162227 | 0.77098275 | 0.68011636 | 0.80315597 | SFI1 | XII | 143201 | 146041 | 1 |
| YLL004W | 7.29859745 | -0.0977924 | 1.04451269 | 0.13328476 | 0.93552971 | 0.95803272 | ORC3 | XII | 141073 | 142923 | 1 |
| YLL005C | 2977.86199 | 0.23923549 | 0.53247943 | 0.62136767 | 0.73294557 | 0.83751334 | SPO75 | XII | 137940 | 140546 | -1 |
| YLL006W | 51.8320068 | 1.16917026 | 0.66002578 | 19.7395128 | 5.17E-05 | 0.0005281 | MMM1 | XII | 136586 | 137866 | 1 |
| YLL006W-A | 77.5372876 | -0.3929365 | 0.56036153 | 4.13549452 | 0.12647037 | 0.25966788 | NA | XII | 136345 | 136521 | 1 |

| | | | | | | | | | | | |
|---|---|---|---|---|---|---|---|---|---|---|---|
| YLL007C | 20.5227497 | -1.973483 | 0.77142441 | 6.59402781 | 0.03699347 | 0.10476085 | LMO1 | XII | 134302 | 136299 | -1 |
| YLL008W | 191.196088 | -0.8383702 | 0.5324096 | 22.953579 | 1.04E-05 | 0.00013674 | DRS1 | XII | 131729 | 133987 | 1 |
| YLL009C | 21.7580001 | 0.16399706 | 0.86239232 | 0.27433915 | 0.87182237 | 0.92527295 | COX17 | XII | 131205 | 131414 | -1 |
| YLL010C | 204.439929 | -0.5733526 | 0.55971541 | 1.76942677 | 0.41283248 | 0.58428259 | PSR1 | XII | 129330 | 130613 | -1 |
| YLL011W | 34.6834778 | -0.1240136 | 0.64086834 | 3.43185682 | 0.17979672 | 0.33430411 | SOF1 | XII | 127523 | 128992 | 1 |
| YLL012W | 104.168444 | 0.60400838 | 0.56497325 | 3.04899906 | 0.21773 | 0.38155469 | YEH1 | XII | 125534 | 127255 | 1 |
| YLL013C | 39.6325083 | 0.31964467 | 0.66122745 | 0.74664885 | 0.68844185 | 0.81049746 | PUF3 | XII | 122075 | 124714 | -1 |
| YLL014W | 6.91345042 | -0.8690912 | 1.24804307 | 0.93465171 | 0.62667585 | 0.76471951 | EMC6 | XII | 121322 | 121648 | 1 |
| YLL015W | 45.856788 | -0.2545029 | 0.72637792 | 0.23312044 | 0.8899765 | 0.93317728 | BPT1 | XII | 116432 | 121111 | 1 |
| YLL016W | 1162.55563 | -0.2951639 | 0.49048218 | 0.62154972 | 0.73287886 | 0.83751334 | NA | XII | 112847 | 115993 | 1 |
| YLL017W | 500.027079 | 0.21417438 | 0.51660148 | 0.17417096 | 0.91659874 | 0.94860224 | NA | XII | 112234 | 112545 | 1 |
| YLL018C | 83.8652628 | 0.24637402 | 0.62141747 | 3.56111382 | 0.16854426 | 0.31849584 | DPS1 | XII | 109902 | 111575 | -1 |
| YLL019C | 63.317529 | -1.4729376 | 0.59105477 | 10.4722581 | 0.00532081 | 0.02406837 | KNS1 | XII | 105686 | 107899 | -1 |
| YLL021W | 68.3782834 | -0.3739958 | 0.58385317 | 8.08198159 | 0.01758005 | 0.05999792 | SPA2 | XII | 100947 | 105347 | 1 |
| YLL022C | 68.8259455 | -1.1633378 | 0.58851226 | 3.89683419 | 0.14249946 | 0.28204189 | HIF1 | XII | 99044 | 100201 | -1 |
| YLL023C | 76.0218861 | -0.803486 | 0.57912206 | 4.71607539 | 0.09460569 | 0.21003333 | POM33 | XII | 97997 | 98836 | -1 |
| YLL024C | 151.286306 | -0.5898029 | 0.7710476 | 2.73167704 | 0.25516662 | 0.42736903 | SSA2 | XII | 95566 | 97485 | -1 |
| YLL025W | 35.0974549 | 0.05202145 | 0.68075853 | 1.45919589 | 0.48210278 | 0.64436175 | PAU17 | XII | 94747 | 95121 | 1 |
| YLL026W | 80.3172799 | 0.69478296 | 0.63181491 | 10.9727142 | 0.00414291 | 0.01969412 | HSP104 | XII | 88623 | 91349 | 1 |
| YLL027W | 22.3436809 | 1.03567723 | 0.71418296 | 2.08334809 | 0.35286348 | 0.52643121 | ISA1 | XII | 87403 | 88155 | 1 |
| YLL028W | 257.903926 | 0.73233983 | 0.48793399 | 15.2035807 | 0.00049956 | 0.00367131 | TPO1 | XII | 84804 | 86564 | 1 |
| YLL029W | 103.212649 | -0.0206358 | 0.53645352 | 0.3315859 | 0.84722163 | 0.90980423 | FRA1 | XII | 81461 | 83710 | 1 |
| YLL031C | 19.4109667 | -0.2690397 | 0.77774357 | 3.26661035 | 0.19528306 | 0.35245914 | GPI13 | XII | 77152 | 80205 | -1 |
| YLL032C | 282.078234 | -0.0887687 | 0.50028453 | 2.76159441 | 0.25137807 | 0.42273571 | NA | XII | 74270 | 76747 | -1 |
| YLL033W | 143.706275 | -0.9554198 | 0.52123248 | 7.13785792 | 0.02818603 | 0.08482177 | IRC19 | XII | 73409 | 74101 | 1 |
| YLL034C | 98.1423989 | 0.99255863 | 0.76555912 | 8.44076178 | 0.01469305 | 0.05261147 | RIX7 | XII | 70633 | 73146 | -1 |
| YLL035W | 27.1781537 | -1.0726413 | 0.67426729 | 6.0592691 | 0.0483333 | 0.12713301 | GRC3 | XII | 68580 | 70478 | 1 |
| YLL036C | 90.6045044 | -1.3946498 | 0.5462629 | 7.76553706 | 0.02059373 | 0.06755677 | PRP19 | XII | 66745 | 68256 | -1 |
| YLL038C | 75.6754422 | -0.0517455 | 0.58565877 | 12.7832308 | 0.00167555 | 0.00980703 | ENT4 | XII | 65775 | 66518 | -1 |
| YLL039C | 71.6765205 | 1.87722235 | 0.86016845 | 5.68729625 | 0.05821291 | 0.14655695 | UBI4 | XII | 64062 | 65207 | -1 |
| YLL040C | 35.1409347 | -0.0632536 | 0.94926962 | 0.32016111 | 0.85207515 | 0.91195795 | VPS13 | XII | 54211 | 63645 | -1 |
| YLL041C | 9.90484103 | 0.47218969 | 1.30329809 | 0.23413917 | 0.88952329 | 0.93317728 | SDH2 | XII | 53131 | 53931 | -1 |
| YLL042C | 62.4104949 | 2.27651902 | 0.82379481 | 9.76358512 | 0.00758341 | 0.03186352 | ATG10 | XII | 52087 | 52590 | -1 |

| | | | | | | | | | | | |
|---|---|---|---|---|---|---|---|---|---|---|---|
| YLL043W | 32.9942993 | 0.14678839 | 0.73568072 | 0.77879159 | 0.67746608 | 0.8011991 | FPS1 | XII | 49938 | 51947 | 1 |
| YLL045C | 135.942016 | -0.5855622 | 0.73156952 | 16.0338369 | 0.00032983 | 0.00264874 | RPL8B | XII | 47859 | 48629 | -1 |
| YLL048C | 87.2818636 | 0.43535301 | 0.62481662 | 5.58532203 | 0.06125799 | 0.15215948 | YBT1 | XII | 41280 | 46265 | -1 |
| YLL049W | 320.831728 | 0.08197559 | 0.62353261 | 15.9571191 | 0.00034273 | 0.00272586 | LDB18 | XII | 40666 | 41205 | 1 |
| YLL050C | 21.1108397 | 0.21078459 | 0.84016853 | 1.17536722 | 0.55561281 | 0.70703257 | COF1 | XII | 39804 | 40414 | -1 |
| YLL051C | 380.704381 | 1.92643368 | 0.50636422 | 13.8408822 | 0.00098739 | 0.00651883 | FRE6 | XII | 37333 | 39471 | -1 |
| YLL052C | 6.39995592 | 0.88049645 | 1.16804147 | 1.31553293 | 0.51800703 | 0.67444585 | AQY2 | XII | 35912 | 36361 | -1 |
| YLL053C | 9.11035867 | 0.56206165 | 1.10553987 | 0.73366584 | 0.69292541 | 0.81380084 | NA | XII | 35503 | 35961 | -1 |
| YLL054C | 39.1516127 | 0.0360408 | 0.65928204 | 12.5696514 | 0.00186438 | 0.01063524 | NA | XII | 32673 | 35204 | -1 |
| YLL055W | 36.903787 | -1.5801411 | 0.65789137 | 10.8724137 | 0.00435597 | 0.0204883 | YCT1 | XII | 30109 | 31704 | 1 |
| YLL056C | 17.3978481 | 1.64995868 | 0.95387135 | 9.59777662 | 0.0082389 | 0.03395248 | NA | XII | 27409 | 28305 | -1 |
| YLL057C | 557.469278 | -0.2774294 | 0.48851724 | 1.67125289 | 0.43360277 | 0.60207676 | JLP1 | XII | 25756 | 26994 | -1 |
| YLL058W | 22.7491351 | 0.26270005 | 0.98653587 | 0.17910121 | 0.91434199 | 0.94739444 | NA | XII | 23569 | 25296 | 1 |
| YLL060C | 691.743166 | 4.88041691 | 0.54719876 | 119.705901 | 1.01E-26 | 1.17E-23 | GTT2 | XII | 21138 | 21839 | -1 |
| YLL061W | 2093.38991 | 3.50062033 | 0.61486443 | 31.8303904 | 1.22E-07 | 2.88E-06 | MMP1 | XII | 17956 | 19707 | 1 |
| YLL062C | 46.5475082 | 0.6081161 | 0.62803656 | 11.962609 | 0.00252553 | 0.01335631 | MHT1 | XII | 16639 | 17613 | -1 |
| YLL063C | 6.02964683 | -0.7662054 | 1.30047681 | 2.34046867 | 0.31029422 | 0.4830878 | AYT1 | XII | 14648 | 16072 | -1 |
| YLR001C | 365.888899 | -0.5729902 | 0.49689516 | 1.58770031 | 0.45210078 | 0.61810237 | NA | XII | 151389 | 153977 | -1 |
| YLR002C | 53.8476042 | 0.30047912 | 0.61086835 | 10.4002427 | 0.00551589 | 0.02475739 | NOC3 | XII | 154343 | 156334 | -1 |
| YLR003C | 21.7499136 | -0.4965427 | 0.78295784 | 0.60486692 | 0.73901766 | 0.84040603 | CMS1 | XII | 156855 | 157730 | -1 |
| YLR004C | 385.713422 | 0.52167495 | 0.55208801 | 7.33870584 | 0.02549296 | 0.07880632 | THI73 | XII | 157934 | 159505 | -1 |
| YLR005W | 82.5953522 | 0.28513888 | 0.57266538 | 16.839874 | 0.00022043 | 0.001854 | SSL1 | XII | 160049 | 161434 | 1 |
| YLR006C | 228.440767 | -0.1209897 | 0.50043659 | 1.62603137 | 0.44351853 | 0.61162622 | SSK1 | XII | 161755 | 163893 | -1 |
| YLR007W | 561.424285 | 0.15831318 | 0.61905047 | 14.569631 | 0.00068587 | 0.00479615 | NSE1 | XII | 164392 | 165402 | 1 |
| YLR008C | 25.4241553 | -1.1193231 | 0.7318357 | 4.25917216 | 0.11888649 | 0.2479657 | PAM18 | XII | 165577 | 166083 | -1 |
| YLR009W | 69.0145571 | -0.1259047 | 0.63089117 | 18.0888671 | 0.00011805 | 0.00107636 | RLP24 | XII | 166537 | 167136 | 1 |
| YLR010C | 17.066754 | -1.7708791 | 0.81210969 | 6.223278 | 0.04452791 | 0.11991471 | TEN1 | XII | 167320 | 167802 | -1 |
| YLR011W | 21.1226735 | -0.0841406 | 0.78449414 | 0.15251612 | 0.92657706 | 0.95392624 | LOT6 | XII | 169103 | 169678 | 1 |
| YLR014C | 49.6776571 | -0.2837757 | 0.80377317 | 12.8066307 | 0.00165606 | 0.00977813 | PPR1 | XII | 172268 | 174982 | -1 |
| YLR015W | 155.647183 | -0.8384201 | 0.50027663 | 7.66019432 | 0.02170751 | 0.07037316 | BRE2 | XII | 175227 | 176744 | 1 |
| YLR016C | 172.079756 | -0.6089174 | 0.49554853 | 4.8621903 | 0.08794047 | 0.20046273 | PML1 | XII | 176802 | 177416 | -1 |
| YLR017W | 87.9483656 | 1.35232178 | 0.75712713 | 3.36367001 | 0.18603229 | 0.3417281 | MEU1 | XII | 177608 | 178621 | 1 |
| YLR018C | 92.9936797 | 1.68219011 | 0.81000827 | 5.42601793 | 0.0663369 | 0.16158631 | POM34 | XII | 178707 | 179606 | -1 |

| | | | | | | | | | | | |
|---|---|---|---|---|---|---|---|---|---|---|---|
| YLR019W | 91.9699081 | -0.0837268 | 0.68368593 | 0.20632417 | 0.90198076 | 0.93824416 | PSR2 | XII | 180288 | 181481 | 1 |
| YLR020C | 14.2542925 | -0.1742695 | 0.93732758 | 0.07730883 | 0.96208313 | 0.97258112 | YEH2 | XII | 181789 | 183405 | -1 |
| YLR021W | 256.618813 | -0.0376144 | 0.56810175 | 7.54917695 | 0.02294653 | 0.07336301 | IRC25 | XII | 183623 | 184162 | 1 |
| YLR022C | 147.332841 | 0.45156466 | 0.64814893 | 5.3264753 | 0.06972212 | 0.16778515 | SDO1 | XII | 184174 | 184926 | -1 |
| YLR023C | 7.17040836 | -0.3885961 | 1.0653026 | 0.51870938 | 0.77154932 | 0.86091165 | IZH3 | XII | 185498 | 187129 | -1 |
| YLR024C | 173.324556 | -0.1314754 | 0.49879917 | 2.28629215 | 0.31881443 | 0.49159402 | UBR2 | XII | 187663 | 193281 | -1 |
| YLR025W | 70.3988846 | -0.1212743 | 0.57871775 | 1.44438228 | 0.48568688 | 0.64773825 | SNF7 | XII | 194452 | 195174 | 1 |
| YLR026C | 52.0598605 | 0.23097489 | 0.58476018 | 0.53013068 | 0.76715582 | 0.85857988 | SED5 | XII | 195450 | 196472 | -1 |
| YLR027C | 73.5191573 | -0.2868597 | 0.58288152 | 7.40936581 | 0.02460802 | 0.07694787 | AAT2 | XII | 196829 | 198085 | -1 |
| YLR028C | 108.329668 | 0.68473556 | 0.63727654 | 1.60547976 | 0.44809954 | 0.61495528 | ADE16 | XII | 199540 | 201315 | -1 |
| YLR029C | 190.228142 | -0.6372701 | 0.59352047 | 15.9597593 | 0.00034228 | 0.00272586 | RPL15A | XII | 201976 | 202590 | -1 |
| YLR030W | 461.237851 | -1.3818754 | 0.52340912 | 12.4926628 | 0.00193755 | 0.01091285 | NA | XII | 203290 | 204081 | 1 |
| YLR031W | 131.582308 | -2.5274459 | 0.56234135 | 43.9760109 | 2.82E-10 | 1.18E-08 | NA | XII | 204225 | 204785 | 1 |
| YLR032W | 28.2710384 | -0.1303562 | 0.67840497 | 0.06727127 | 0.96692376 | 0.97486964 | RAD5 | XII | 204991 | 208500 | 1 |
| YLR033W | 72.88106 | 0.24256497 | 0.55200354 | 6.26145181 | 0.04368607 | 0.11847418 | RSC58 | XII | 208761 | 210269 | 1 |
| YLR034C | 46.565282 | -0.7691967 | 0.64814518 | 1.69306817 | 0.42889889 | 0.59853568 | SMF3 | XII | 210512 | 211933 | -1 |
| YLR035C | 78.8481265 | 0.03819841 | 0.54935137 | 0.93391144 | 0.62690785 | 0.76477701 | MLH2 | XII | 212369 | 214456 | -1 |
| YLR036C | 132.802365 | -0.3639979 | 0.6171247 | 1.10789983 | 0.5746754 | 0.72422968 | NA | XII | 221521 | 222132 | -1 |
| YLR038C | 493.818553 | 2.02808286 | 0.5573577 | 50.7077881 | 9.75E-12 | 5.76E-10 | COX12 | XII | 224921 | 225172 | -1 |
| YLR039C | 100.352284 | -0.4581832 | 0.56776413 | 26.2520918 | 1.99E-06 | 3.24E-05 | RIC1 | XII | 225426 | 228596 | -1 |
| YLR040C | 143.427358 | 1.14892131 | 0.53513057 | 4.60322323 | 0.1000974 | 0.21919967 | AFB1 | XII | 228908 | 229582 | -1 |
| YLR042C | 1874.38027 | 0.58041699 | 0.51990245 | 2.35671107 | 0.30778446 | 0.48023239 | NA | XII | 229966 | 230451 | -1 |
| YLR043C | 71.6066812 | -0.7499925 | 0.57383622 | 1.90420933 | 0.38592792 | 0.55863066 | TRX1 | XII | 231702 | 232013 | -1 |
| YLR044C | 688.9685 | -0.6620087 | 0.67325968 | 8.580594 | 0.01370086 | 0.04989179 | PDC1 | XII | 232390 | 234081 | -1 |
| YLR045C | 142.068932 | -0.6526632 | 0.52587445 | 9.50660719 | 0.00862316 | 0.03516063 | STU2 | XII | 235037 | 237703 | -1 |
| YLR046C | 21.5722861 | 0.04612443 | 0.76525335 | 0.12549684 | 0.93917972 | 0.96024203 | NA | XII | 238001 | 238813 | -1 |
| YLR047C | 394.564398 | 0.64019438 | 0.54863765 | 6.92326186 | 0.03137854 | 0.09258058 | FRE8 | XII | 239347 | 241407 | -1 |
| YLR048W | 168.218799 | -0.5697619 | 0.5762228 | 25.2795451 | 3.24E-06 | 4.95E-05 | RPS0B | XII | 242232 | 243349 | 1 |
| YLR049C | 30.3707644 | 0.60086497 | 0.71453907 | 2.59435911 | 0.27330154 | 0.44663164 | NA | XII | 243886 | 245172 | -1 |
| YLR050C | 473.770481 | -1.2440082 | 0.47616469 | 21.9267319 | 1.73E-05 | 0.00020811 | NA | XII | 245587 | 246072 | -1 |
| YLR051C | 13.9543765 | -0.2970509 | 0.85941341 | 5.52681326 | 0.06307652 | 0.15574118 | FCF2 | XII | 246324 | 246977 | -1 |
| YLR052W | 14.4461265 | 0.66738715 | 0.8601566 | 1.07291807 | 0.5848154 | 0.7327594 | IES3 | XII | 247201 | 247953 | 1 |
| YLR053C | 326.179925 | -0.8747436 | 0.57665669 | 10.1896076 | 0.00612851 | 0.02692266 | NA | XII | 248100 | 248426 | -1 |

| | | | | | | | | | | | |
|---|---|---|---|---|---|---|---|---|---|---|---|
| YLR054C | 392.045902 | 2.39490427 | 0.523281 | 51.2887422 | 7.29E-12 | 4.52E-10 | OSW2 | XII | 248717 | 250978 | -1 |
| YLR055C | 131.699232 | 0.54732446 | 0.53064211 | 3.82630148 | 0.14761456 | 0.28982309 | SPT8 | XII | 251272 | 253080 | -1 |
| YLR056W | 52.0152027 | -0.4391657 | 0.58364811 | 3.48285343 | 0.17527016 | 0.32699386 | ERG3 | XII | 253861 | 254958 | 1 |
| YLR057W | 143.094987 | 2.19440187 | 0.88150773 | 8.08486075 | 0.01755476 | 0.05999792 | MNL2 | XII | 255306 | 257855 | 1 |
| YLR058C | 32.9524941 | -0.2846999 | 0.66167095 | 1.01662504 | 0.60150976 | 0.74720908 | SHM2 | XII | 257992 | 259401 | -1 |
| YLR059C | 236.150597 | -1.0485557 | 0.50484708 | 7.66433135 | 0.02166265 | 0.07033309 | REX2 | XII | 259738 | 260547 | -1 |
| YLR060W | 118.650096 | 0.49412755 | 0.64709309 | 3.43689527 | 0.17934434 | 0.3335701 | FRS1 | XII | 260979 | 262766 | 1 |
| YLR061W | 115.776088 | -0.453572 | 0.54299714 | 32.5093148 | 8.72E-08 | 2.16E-06 | RPL22A | XII | 263194 | 263948 | 1 |
| YLR063W | 44.9216149 | -0.8507021 | 0.63856533 | 5.19515902 | 0.07445357 | 0.17631337 | BMT6 | XII | 264157 | 265254 | 1 |
| YLR064W | 163.842693 | 0.26179589 | 0.59310723 | 12.6003005 | 0.00183603 | 0.01052535 | PER33 | XII | 265456 | 266277 | 1 |
| YLR065C | 133.288523 | 0.57135991 | 0.64784392 | 5.89856594 | 0.05237725 | 0.13534434 | ENV10 | XII | 266373 | 266918 | -1 |
| YLR066W | 15.49876 | 1.50095635 | 0.85312198 | 3.39886488 | 0.18278724 | 0.33812719 | SPC3 | XII | 267169 | 267723 | 1 |
| YLR067C | 397.694641 | 2.91757669 | 0.9050722 | 13.8447177 | 0.0009855 | 0.00651377 | PET309 | XII | 267813 | 270710 | -1 |
| YLR068W | 59.4953894 | -0.6706827 | 0.64385785 | 18.5119312 | 9.55E-05 | 0.00089947 | FYV7 | XII | 271008 | 271463 | 1 |
| YLR069C | 46.5617288 | -0.4260118 | 0.61307805 | 1.88398601 | 0.38985009 | 0.56101577 | MEF1 | XII | 271630 | 273915 | -1 |
| YLR070C | 321.393428 | -0.2323963 | 0.46868227 | 1.38089626 | 0.50135135 | 0.66037092 | XYL2 | XII | 274141 | 275211 | -1 |
| YLR071C | 20.0380456 | -0.1943581 | 0.72499696 | 0.2213572 | 0.89522643 | 0.93531262 | RGR1 | XII | 275378 | 278626 | -1 |
| YLR072W | 54.6887811 | 0.93100538 | 0.69182813 | 6.82863601 | 0.03289884 | 0.09591352 | LAM6 | XII | 278862 | 280943 | 1 |
| YLR073C | 95.724713 | 0.11418613 | 0.68001311 | 6.87622048 | 0.03212534 | 0.0942798 | RFU1 | XII | 281019 | 281621 | -1 |
| YLR074C | 58.1086768 | -1.8512619 | 0.65260332 | 22.2849369 | 1.45E-05 | 0.00018113 | BUD20 | XII | 281956 | 282456 | -1 |
| YLR075W | 170.048347 | -0.606454 | 0.89521027 | 8.9965994 | 0.0111279 | 0.042641 | RPL10 | XII | 282927 | 283592 | 1 |
| YLR077W | 73.4438498 | 0.05983237 | 0.57469273 | 0.22997501 | 0.89137728 | 0.93352778 | FMP25 | XII | 283872 | 285623 | 1 |
| YLR078C | 132.619071 | 1.61222453 | 0.76610339 | 6.60052549 | 0.03687348 | 0.10460433 | BOS1 | XII | 285736 | 286559 | -1 |
| YLR079W | 62.6184079 | -0.2919819 | 0.62770703 | 0.23129441 | 0.89078944 | 0.93317728 | SIC1 | XII | 286820 | 287674 | 1 |
| YLR080W | 96.8951977 | 0.96074402 | 0.58759796 | 5.54352652 | 0.06255161 | 0.15457697 | EMP46 | XII | 287916 | 289250 | 1 |
| YLR081W | 308.082255 | -0.6004383 | 0.52376604 | 1.636287 | 0.44125008 | 0.60960103 | GAL2 | XII | 290212 | 291936 | 1 |
| YLR082C | 5.8500978 | -1.3798368 | 1.28163938 | 2.02428184 | 0.36344005 | 0.53640527 | SRL2 | XII | 292394 | 293572 | -1 |
| YLR083C | 26.0368959 | -1.2883191 | 0.78610009 | 3.04524808 | 0.21813873 | 0.38215529 | EMP70 | XII | 294091 | 296094 | -1 |
| YLR084C | 56.0018901 | 0.4625337 | 0.59783884 | 11.5940722 | 0.00303654 | 0.01538156 | RAX2 | XII | 296589 | 300251 | -1 |
| YLR085C | 23.255992 | -1.67732 | 0.79156434 | 7.53640785 | 0.0230935 | 0.0737108 | ARP6 | XII | 300673 | 301989 | -1 |
| YLR086W | 50.5364931 | -0.305411 | 0.57512578 | 0.68052752 | 0.71158261 | 0.82438173 | SMC4 | XII | 302243 | 306499 | 1 |
| YLR087C | 356.039151 | -0.9968568 | 0.50229592 | 9.64767894 | 0.00803587 | 0.03330545 | CSF1 | XII | 306855 | 315731 | -1 |
| YLR088W | 29.3705196 | 0.00227122 | 0.76411183 | 0.26990802 | 0.87375609 | 0.92573532 | GAA1 | XII | 316107 | 317951 | 1 |

| | | | | | | | | | | | |
|---|---|---|---|---|---|---|---|---|---|---|---|
| YLR089C | 103.938229 | -0.5739351 | 0.58321732 | 1.71355147 | 0.42452867 | 0.59492495 | ALT1 | XII | 318237 | 320015 | -1 |
| YLR090W | 18.1744279 | -0.751325 | 0.80214782 | 1.57793366 | 0.45431394 | 0.62039568 | XDJ1 | XII | 320701 | 322080 | 1 |
| YLR091W | 274.56831 | -0.5114773 | 0.59982086 | 1.50592922 | 0.47096824 | 0.63298657 | GEP5 | XII | 322297 | 323178 | 1 |
| YLR092W | 1297.45965 | 0.29028764 | 0.54560476 | 2.93805066 | 0.2301497 | 0.39706995 | SUL2 | XII | 323544 | 326225 | 1 |
| YLR093C | 30.5411433 | 0.73856947 | 0.72245667 | 3.12279929 | 0.20984216 | 0.37258084 | NYV1 | XII | 326513 | 327415 | -1 |
| YLR094C | 23.0068344 | -0.1616115 | 0.72881484 | 0.12243864 | 0.94061692 | 0.96082012 | GIS3 | XII | 327730 | 329238 | -1 |
| YLR095C | 263.740584 | -0.3885268 | 0.48917541 | 0.63329612 | 0.72858713 | 0.83634407 | IOC2 | XII | 329677 | 332115 | -1 |
| YLR096W | 38.4146317 | -0.0447759 | 0.62988958 | 2.96617067 | 0.22693643 | 0.3929312 | KIN2 | XII | 332590 | 336033 | 1 |
| YLR097C | 11.5237626 | -0.0079299 | 1.0241064 | 1.30534885 | 0.52065147 | 0.6765916 | HRT3 | XII | 336231 | 337265 | -1 |
| YLR098C | 62.7225267 | -1.2318472 | 0.63349858 | 3.97018058 | 0.13736821 | 0.27509737 | CHA4 | XII | 337527 | 339473 | -1 |
| YLR099C | 219.492179 | -1.7570349 | 0.53618101 | 10.4766619 | 0.00530911 | 0.02403421 | ICT1 | XII | 339744 | 340928 | -1 |
| YLR099W-A | 13.3433624 | -0.1723099 | 0.82022991 | 1.42719026 | 0.48987985 | 0.65114883 | MIM2 | XII | 341325 | 341588 | 1 |
| YLR100W | 414.85557 | 2.33090025 | 0.78103995 | 8.13501045 | 0.01712005 | 0.05900302 | ERG27 | XII | 341810 | 342853 | 1 |
| YLR102C | 153.33958 | 0.69070632 | 0.65078901 | 1.19119173 | 0.55123401 | 0.70300548 | APC9 | XII | 342970 | 343767 | -1 |
| YLR103C | 35.6019164 | -0.1156322 | 0.64002614 | 0.32244925 | 0.85110087 | 0.91173629 | CDC45 | XII | 343989 | 345941 | -1 |
| YLR104W | 157.46729 | 2.6428754 | 0.78795527 | 11.7220303 | 0.00284835 | 0.01465951 | LCL2 | XII | 346585 | 346980 | 1 |
| YLR105C | 62.2918952 | 2.11149931 | 0.8240243 | 6.16238562 | 0.04590447 | 0.12231333 | SEN2 | XII | 347047 | 348180 | -1 |
| YLR106C | 213.900546 | -0.0911903 | 0.56208396 | 1.34265316 | 0.5110302 | 0.66829159 | MDN1 | XII | 349006 | 363738 | -1 |
| YLR107W | 44.9989435 | 3.28179452 | 0.74181946 | 21.837798 | 1.81E-05 | 0.00021713 | REX3 | XII | 364116 | 365330 | 1 |
| YLR108C | 200.795583 | 1.55531087 | 0.54470868 | 32.1531342 | 1.04E-07 | 2.51E-06 | NA | XII | 366667 | 368124 | -1 |
| YLR109W | 92.449128 | -0.2322274 | 0.59382922 | 0.41621805 | 0.81211849 | 0.88731158 | AHP1 | XII | 368781 | 369311 | 1 |
| YLR110C | 292.164755 | 0.04576406 | 0.52819338 | 1.6153856 | 0.44588563 | 0.61337082 | CCW12 | XII | 369697 | 370098 | -1 |
| YLR111W | 136.924126 | 0.82324661 | 0.55694124 | 8.78554074 | 0.01236642 | 0.04619457 | NA | XII | 370391 | 370723 | 1 |
| YLR112W | 87.269329 | 1.69681441 | 0.63051832 | 15.5111567 | 0.00042835 | 0.00324208 | NA | XII | 370791 | 371210 | 1 |
| YLR113W | 276.476565 | 0.88566808 | 0.49676694 | 13.1040652 | 0.00142721 | 0.00880976 | HOG1 | XII | 371620 | 372927 | 1 |
| YLR114C | 88.2312069 | 0.15591843 | 0.62622019 | 0.74816734 | 0.68791935 | 0.81022234 | AVL9 | XII | 374944 | 377238 | -1 |
| YLR115W | 33.0740118 | -0.1028943 | 0.71127554 | 2.32539197 | 0.31264217 | 0.4849178 | CFT2 | XII | 377985 | 380564 | 1 |
| YLR116W | 14.3004611 | 0.10266472 | 0.82679537 | 0.83951872 | 0.65720495 | 0.78549013 | MSL5 | XII | 380822 | 382252 | 1 |
| YLR117C | 72.2386278 | -0.2544438 | 0.55199741 | 7.89299487 | 0.01932226 | 0.06448178 | CLF1 | XII | 382471 | 384534 | -1 |
| YLR118C | 10.6035215 | -0.3090327 | 0.94915156 | 0.41780203 | 0.81147556 | 0.88731158 | NA | XII | 384725 | 385408 | -1 |
| YLR119W | 685.564241 | 3.4918684 | 0.68524157 | 24.2829662 | 5.33E-06 | 7.55E-05 | SRN2 | XII | 385534 | 386175 | 1 |
| YLR120C | 113.47093 | -0.883239 | 0.6212103 | 3.63483103 | 0.16244505 | 0.31103069 | YPS1 | XII | 386511 | 388220 | -1 |
| YLR121C | 1135.88786 | -0.0504608 | 0.47699916 | 8.22628269 | 0.01635631 | 0.05698138 | YPS3 | XII | 388744 | 390270 | -1 |

| | | | | | | | | | | | |
|---|---|---|---|---|---|---|---|---|---|---|---|
| YLR122C | 61.6740397 | 2.22678134 | 0.6027451 | 14.653986 | 0.00065755 | 0.00463727 | NA | XII | 390954 | 391331 | -1 |
| YLR126C | 177.381549 | -0.8963155 | 0.55346775 | 11.3663124 | 0.0034028 | 0.01682512 | NA | XII | 394765 | 395520 | -1 |
| YLR127C | 322.108027 | -0.6982301 | 0.50702176 | 8.76274843 | 0.01250816 | 0.04666381 | APC2 | XII | 395758 | 398319 | -1 |
| YLR128W | 12.2940139 | 1.16297971 | 0.94158919 | 3.98333191 | 0.13646789 | 0.27378692 | DCN1 | XII | 398530 | 399433 | 1 |
| YLR129W | 67.6394381 | -0.5077004 | 0.5959707 | 3.01605084 | 0.22134661 | 0.38636302 | DIP2 | XII | 399657 | 402488 | 1 |
| YLR130C | 99.416519 | -0.8558759 | 0.60047725 | 11.4232031 | 0.00330737 | 0.01643749 | ZRT2 | XII | 402794 | 404062 | -1 |
| YLR131C | 276.934895 | 1.19183863 | 0.51520169 | 57.2043708 | 3.79E-13 | 2.85E-11 | ACE2 | XII | 404510 | 406822 | -1 |
| YLR132C | 30.5154113 | -0.8831433 | 0.8370407 | 8.89086521 | 0.01173203 | 0.04440749 | USB1 | XII | 407283 | 408155 | -1 |
| YLR133W | 18.6119003 | 0.29462643 | 0.78398873 | 0.3295122 | 0.84810053 | 0.90980423 | CKI1 | XII | 408445 | 410193 | 1 |
| YLR134W | 26.4535207 | -1.1910509 | 0.87584711 | 4.40601004 | 0.11047069 | 0.2353294 | PDC5 | XII | 410723 | 412414 | 1 |
| YLR135W | 173.742137 | -0.8590449 | 0.52087399 | 3.07908586 | 0.21447911 | 0.37828653 | SLX4 | XII | 413281 | 415527 | 1 |
| YLR136C | 61.8370891 | -0.6389997 | 0.62604004 | 3.91693547 | 0.14107442 | 0.28002087 | TIS11 | XII | 415801 | 416658 | -1 |
| YLR137W | 19.2836401 | 0.01745385 | 0.74556614 | 0.07200779 | 0.96463654 | 0.97388763 | RKM5 | XII | 417006 | 418109 | 1 |
| YLR138W | 213.049736 | 1.34522332 | 0.54489353 | 6.130323 | 0.04664631 | 0.12398451 | NHA1 | XII | 418437 | 421394 | 1 |
| YLR139C | 170.319651 | 1.78944653 | 0.73817622 | 8.67269485 | 0.01308423 | 0.04820096 | SLS1 | XII | 421542 | 423473 | -1 |
| YLR141W | 313.645258 | -0.1942449 | 0.49328146 | 0.2171266 | 0.89712211 | 0.9367605 | RRN5 | XII | 423683 | 424774 | 1 |
| YLR142W | 51.3655005 | -1.0408023 | 0.58211703 | 3.24476544 | 0.19742772 | 0.35555413 | PUT1 | XII | 425186 | 426616 | 1 |
| YLR143W | 38.7933049 | -0.5815718 | 0.65537999 | 0.79068995 | 0.67344768 | 0.7980479 | DPH6 | XII | 427329 | 429386 | 1 |
| YLR144C | 125.370496 | -0.2034054 | 0.52688984 | 1.71078614 | 0.42511606 | 0.59512137 | ACF2 | XII | 429677 | 432016 | -1 |
| YLR145W | 191.229209 | 1.65049461 | 0.84114657 | 5.47753579 | 0.06464995 | 0.15847724 | RMP1 | XII | 432168 | 432773 | 1 |
| YLR146C | 53.060647 | 2.14921175 | 0.96642486 | 4.72787077 | 0.09404937 | 0.2093602 | SPE4 | XII | 432823 | 433725 | -1 |
| YLR146W-A | 82.7316951 | 1.18696639 | 0.7788817 | 3.50904131 | 0.17299014 | 0.32456673 | NA | XII | 433870 | 434058 | 1 |
| YLR147C | 58.0255561 | 1.30038738 | 0.7053774 | 8.70801683 | 0.01285518 | 0.04759047 | SMD3 | XII | 434158 | 434463 | -1 |
| YLR148W | 12.8749628 | 0.03067402 | 0.93170657 | 0.46533851 | 0.79241562 | 0.87392122 | PEP3 | XII | 434641 | 437397 | 1 |
| YLR149C | 113.20265 | -0.3402881 | 0.53139635 | 13.7318785 | 0.0010427 | 0.00682175 | NA | XII | 437631 | 439823 | -1 |
| YLR150W | 356.670296 | -0.5811148 | 0.65761885 | 17.6137065 | 0.0001497 | 0.00133351 | STM1 | XII | 440467 | 441288 | 1 |
| YLR151C | 116.503544 | -0.4156482 | 0.55558524 | 8.82582583 | 0.01211982 | 0.04556739 | PCD1 | XII | 441715 | 442737 | -1 |
| YLR152C | 40.3578119 | -0.1515165 | 0.6688467 | 6.78951472 | 0.03354869 | 0.09746459 | NA | XII | 442958 | 444688 | -1 |
| YLR153C | 78.9372971 | -0.3002367 | 0.75567159 | 3.77282374 | 0.15161485 | 0.29497647 | ACS2 | XII | 445524 | 447575 | -1 |
| YLR154W-C | 1172.96589 | -1.1599808 | 0.64042736 | 5.46331853 | 0.06511116 | 0.15927065 | TAR1 | XII | 454696 | 455070 | 1 |
| YLR157W-E | 5.19751994 | -0.9228885 | 1.27984808 | 0.8485003 | 0.6542602 | 0.78442961 | NA | XII | 481873 | 482037 | 1 |
| YLR162W | 9.2268746 | 0.0352505 | 1.05618293 | 0.13494561 | 0.93475314 | 0.95774565 | NA | XII | 489573 | 489929 | 1 |
| YLR162W-A | 25.3462273 | -0.5285364 | 1.02262849 | 1.02714206 | 0.59835502 | 0.74552949 | RRT15 | XII | 490406 | 490594 | 1 |

| | | | | | | | | | | | |
|---|---|---|---|---|---|---|---|---|---|---|---|
| YLR163C | 137.951094 | -0.7338328 | 0.53868259 | 3.35176961 | 0.18714252 | 0.34300576 | MAS1 | XII | 491866 | 493254 | -1 |
| YLR164W | 106.040278 | 1.34306271 | 0.5312948 | 12.8568539 | 0.00161499 | 0.00961783 | SHH4 | XII | 493883 | 494389 | 1 |
| YLR165C | 44.551342 | -0.4465078 | 0.64121729 | 0.85715935 | 0.65143368 | 0.78236902 | PUS5 | XII | 494494 | 495258 | -1 |
| YLR166C | 33.1751793 | 0.55023255 | 0.64275961 | 1.24331741 | 0.53705289 | 0.69177669 | SEC10 | XII | 495429 | 498044 | -1 |
| YLR167W | 375.698986 | -0.6877741 | 0.67104251 | 20.8011501 | 3.04E-05 | 0.00034284 | RPS31 | XII | 498947 | 499405 | 1 |
| YLR168C | 10.7905189 | -0.0688894 | 0.90781322 | 3.06839262 | 0.21562892 | 0.37948068 | UPS2 | XII | 499578 | 500270 | -1 |
| YLR170C | 5.91356301 | -0.1439386 | 1.20769571 | 0.34033696 | 0.84352269 | 0.90831251 | APS1 | XII | 500579 | 501049 | -1 |
| YLR172C | 221.614274 | -0.1904113 | 0.50640743 | 0.1745274 | 0.9164354 | 0.94860224 | DPH5 | XII | 501260 | 502162 | -1 |
| YLR173W | 24.5198435 | -0.1971519 | 0.71596045 | 0.1264541 | 0.93873031 | 0.95995205 | NA | XII | 502421 | 504247 | 1 |
| YLR174W | 13.0328604 | -0.3987761 | 0.91600102 | 0.39818597 | 0.81947369 | 0.89171017 | IDP2 | XII | 504591 | 505829 | 1 |
| YLR175W | 671.100636 | -0.8824137 | 0.50409126 | 19.9113077 | 4.75E-05 | 0.00049245 | CBF5 | XII | 506134 | 507585 | 1 |
| YLR176C | 1277.10118 | 3.6976252 | 0.5764265 | 66.4598405 | 3.70E-15 | 4.02E-13 | RFX1 | XII | 507797 | 510232 | -1 |
| YLR177W | 20.2273986 | -0.3003914 | 0.7818575 | 1.73678239 | 0.4196261 | 0.59032562 | NA | XII | 511054 | 512940 | 1 |
| YLR178C | 23.4027875 | 0.45608736 | 0.77380096 | 2.40339551 | 0.30068329 | 0.47389383 | TFS1 | XII | 513162 | 513821 | -1 |
| YLR179C | 177.429136 | -0.0383792 | 0.51209782 | 3.15370603 | 0.20662432 | 0.36867637 | NA | XII | 514108 | 514713 | -1 |
| YLR180W | 51.5682664 | -1.3794445 | 0.67639988 | 25.6438004 | 2.70E-06 | 4.19E-05 | SAM1 | XII | 515262 | 516410 | 1 |
| YLR181C | 108.914756 | 0.71659653 | 0.77446046 | 9.40804597 | 0.00905876 | 0.03655068 | VTA1 | XII | 516678 | 517670 | -1 |
| YLR182W | 185.237221 | 0.4483024 | 0.62839986 | 74.465704 | 6.76E-17 | 9.79E-15 | SWI6 | XII | 517940 | 520351 | 1 |
| YLR183C | 16.345671 | 1.15791442 | 0.97930886 | 3.50022178 | 0.17375467 | 0.32507061 | TOS4 | XII | 520543 | 522012 | -1 |
| YLR185W | 42.4033398 | -0.7763017 | 0.74025631 | 12.5981378 | 0.00183802 | 0.01052632 | RPL37A | XII | 522663 | 523288 | 1 |
| YLR186W | 30.8537645 | -0.6729124 | 0.75759068 | 1.88094466 | 0.39044337 | 0.56165643 | EMG1 | XII | 523632 | 524390 | 1 |
| YLR187W | 29.1180825 | -0.3683057 | 0.66501327 | 0.59199366 | 0.74378979 | 0.8437584 | SKG3 | XII | 524865 | 527945 | 1 |
| YLR188W | 158.150221 | 0.91607313 | 0.54494418 | 8.17614244 | 0.01677155 | 0.05813116 | MDL1 | XII | 528300 | 530387 | 1 |
| YLR189C | 173.802973 | -0.9174616 | 0.5456169 | 3.28505471 | 0.19349041 | 0.35020614 | ATG26 | XII | 530797 | 534393 | -1 |
| YLR190W | 86.594231 | 0.24493305 | 0.60181371 | 17.482366 | 0.00015986 | 0.00140458 | MMR1 | XII | 535214 | 536689 | 1 |
| YLR191W | 48.2611282 | 0.44926166 | 0.62375388 | 12.1615797 | 0.00228637 | 0.01234789 | PEX13 | XII | 537272 | 538432 | 1 |
| YLR192C | 150.34374 | -0.0753687 | 0.54896395 | 1.98934581 | 0.3698444 | 0.54308878 | HCR1 | XII | 538792 | 539589 | -1 |
| YLR193C | 28.8743098 | -0.7233226 | 0.69170204 | 1.56213067 | 0.45791792 | 0.62324553 | UPS1 | XII | 540009 | 540536 | -1 |
| YLR194C | 648.95072 | 0.77215424 | 0.4617045 | 3.62938565 | 0.16288794 | 0.31167255 | NCW2 | XII | 540809 | 541573 | -1 |
| YLR195C | 1775.96751 | 4.34523398 | 0.47493353 | 80.5675993 | 3.20E-18 | 5.97E-16 | NMT1 | XII | 541937 | 543304 | -1 |
| YLR196W | 51.3886145 | -0.6435541 | 0.62821194 | 6.60130057 | 0.03685919 | 0.10460433 | PWP1 | XII | 543968 | 545698 | 1 |
| YLR197W | 496.519433 | -0.8220044 | 0.58257359 | 31.7971285 | 1.25E-07 | 2.91E-06 | NOP56 | XII | 546097 | 547611 | 1 |
| YLR199C | 50.2445665 | -0.2959156 | 0.59547966 | 3.03135183 | 0.21965966 | 0.3842385 | PBA1 | XII | 547853 | 548769 | -1 |

| | | | | | | | | | | | |
|---|---|---|---|---|---|---|---|---|---|---|---|
| YLR200W | 34.1807421 | 0.91395503 | 0.7217028 | 2.33914842 | 0.31049912 | 0.48327686 | YKE2 | XII | 549012 | 549356 | 1 |
| YLR201C | 151.013364 | 0.92384207 | 0.6586276 | 5.69481034 | 0.05799461 | 0.14626659 | COQ9 | XII | 549511 | 550293 | -1 |
| YLR203C | 27.4399533 | -0.9282357 | 0.7314347 | 3.60373648 | 0.16499036 | 0.31392516 | MSS51 | XII | 550650 | 551960 | -1 |
| YLR204W | 52.347963 | 0.67147962 | 0.66328776 | 3.47923706 | 0.17558737 | 0.32742379 | QRI5 | XII | 552270 | 552605 | 1 |
| YLR205C | 296.089123 | 1.94940191 | 0.84680596 | 5.47335179 | 0.06478534 | 0.15867476 | HMX1 | XII | 552724 | 553677 | -1 |
| YLR206W | 206.367522 | 2.99042005 | 0.79356041 | 12.7894361 | 0.00167036 | 0.00979022 | ENT2 | XII | 554578 | 556419 | 1 |
| YLR207W | 34.3457974 | 0.35893097 | 0.64820835 | 1.4099675 | 0.4941166 | 0.65469026 | HRD3 | XII | 556788 | 559289 | 1 |
| YLR208W | 20.6655441 | 0.15166027 | 0.80321885 | 0.70967669 | 0.7012868 | 0.81814438 | SEC13 | XII | 559551 | 560444 | 1 |
| YLR209C | 134.993546 | 1.90685811 | 0.85863814 | 12.856207 | 0.00161551 | 0.00961783 | PNP1 | XII | 560797 | 561732 | -1 |
| YLR210W | 7.50055348 | -0.6831216 | 1.041468 | 1.18144941 | 0.55392571 | 0.70568075 | CLB4 | XII | 562008 | 563390 | 1 |
| YLR212C | 23.0705508 | -0.3885177 | 0.71675401 | 2.24654081 | 0.32521447 | 0.49807536 | TUB4 | XII | 564860 | 566281 | -1 |
| YLR213C | 54.087663 | -1.0494082 | 0.61656396 | 4.29708861 | 0.11665385 | 0.24436533 | CRR1 | XII | 566654 | 567922 | -1 |
| YLR214W | 254.994056 | -0.5380784 | 0.74032169 | 6.41890117 | 0.04037879 | 0.11218484 | FRE1 | XII | 568567 | 570627 | 1 |
| YLR215C | 128.644294 | -0.0041978 | 0.61759895 | 6.22443179 | 0.04450223 | 0.11990132 | CDC123 | XII | 570776 | 571858 | -1 |
| YLR216C | 45.3891375 | 0.0156862 | 0.69015526 | 0.4533548 | 0.79717791 | 0.87733513 | CPR6 | XII | 572096 | 573211 | -1 |
| YLR218C | 109.388357 | 0.97625698 | 0.6640991 | 7.99647522 | 0.01834795 | 0.06205293 | NA | XII | 573466 | 573918 | -1 |
| YLR219W | 53.1535548 | 1.15916019 | 0.63858458 | 3.27483231 | 0.19448191 | 0.35156111 | MSC3 | XII | 574151 | 576337 | 1 |
| YLR220W | 68.1365476 | -0.205203 | 0.60777511 | 10.9078426 | 0.00427949 | 0.02021064 | CCC1 | XII | 576825 | 577793 | 1 |
| YLR221C | 25.7518269 | -1.1582914 | 0.70995836 | 11.0789508 | 0.00392859 | 0.01897124 | RSA3 | XII | 578362 | 579024 | -1 |
| YLR222C | 21.6798531 | -0.8636295 | 0.87565792 | 1.20005844 | 0.5487956 | 0.70069987 | UTP13 | XII | 579318 | 581771 | -1 |
| YLR223C | 117.493816 | 0.16896291 | 0.51581492 | 0.23523968 | 0.88903396 | 0.93304075 | IFH1 | XII | 582233 | 585490 | -1 |
| YLR224W | 311.442286 | 5.1308194 | 1.85985249 | 5.72046724 | 0.05725538 | 0.14501691 | UCC1 | XII | 586464 | 587573 | 1 |
| YLR225C | 75.1847186 | 2.88138943 | 0.76836764 | 14.0600057 | 0.00088493 | 0.00595552 | NA | XII | 587695 | 588918 | -1 |
| YLR226W | 61.8471132 | 0.40309519 | 0.56235605 | 1.98322148 | 0.37097866 | 0.5443402 | BUR2 | XII | 589354 | 590541 | 1 |
| YLR227C | 1287.92298 | -0.3713267 | 0.47008501 | 1.94243595 | 0.3786216 | 0.55247457 | ADY4 | XII | 590562 | 592043 | -1 |
| YLR227W-B | 5.67558581 | -0.4199066 | 1.24091059 | 1.01941435 | 0.60067144 | 0.74682758 | NA | XII | 593438 | 598706 | 1 |
| YLR228C | 20.9775114 | 0.15264411 | 0.72352519 | 0.37584617 | 0.82867844 | 0.89767038 | ECM22 | XII | 600019 | 602463 | -1 |
| YLR229C | 50.9993477 | 0.41936703 | 0.63388398 | 3.35044567 | 0.18726645 | 0.34312428 | CDC42 | XII | 604212 | 604787 | -1 |
| YLR231C | 35.9224423 | -0.5121275 | 0.64361977 | 0.65769419 | 0.71975306 | 0.82982282 | BNA5 | XII | 605758 | 607119 | -1 |
| YLR233C | 241.595612 | 0.27874575 | 0.50025182 | 10.0092512 | 0.00670685 | 0.02902292 | EST1 | XII | 607425 | 609524 | -1 |
| YLR234W | 170.08878 | -0.1482574 | 0.5642801 | 0.24656725 | 0.8840129 | 0.9315066 | TOP3 | XII | 609783 | 611753 | 1 |
| YLR236C | 189.380592 | -1.0072841 | 0.59579447 | 2.8003 | 0.24655998 | 0.41766412 | NA | XII | 611794 | 612117 | -1 |
| YLR237W | 210.595195 | 1.10485527 | 0.57095721 | 12.6274047 | 0.00181131 | 0.01040428 | THI7 | XII | 612367 | 614163 | 1 |

| | | | | | | | | | | | |
|---|---|---|---|---|---|---|---|---|---|---|---|
| YLR238W | 20.0029686 | -0.2190688 | 0.76991849 | 4.16531741 | 0.1245985 | 0.25673499 | FAR10 | XII | 614755 | 616191 | 1 |
| YLR239C | 6.29526663 | 0.76380437 | 1.24882125 | 1.23079571 | 0.54042584 | 0.69442202 | LIP2 | XII | 616332 | 617318 | -1 |
| YLR240W | 289.453827 | 0.86927086 | 0.52202265 | 16.0510516 | 0.00032701 | 0.00262969 | VPS34 | XII | 617533 | 620160 | 1 |
| YLR241W | 694.95083 | -0.3277576 | 0.50396069 | 67.7533773 | 1.94E-15 | 2.20E-13 | CSC1 | XII | 620473 | 622821 | 1 |
| YLR242C | 441.657544 | 3.51441367 | 0.88839437 | 13.4230038 | 0.00121684 | 0.0077508 | ARV1 | XII | 622918 | 623883 | -1 |
| YLR243W | 20.8778414 | -0.7703407 | 0.75876052 | 2.00287092 | 0.36735174 | 0.54052518 | GPN3 | XII | 624203 | 625021 | 1 |
| YLR244C | 110.156106 | 0.73897395 | 0.75169435 | 6.00150716 | 0.04974956 | 0.13016266 | MAP1 | XII | 625168 | 626331 | -1 |
| YLR245C | 89.7222558 | -1.8916864 | 0.60001209 | 27.6086917 | 1.01E-06 | 1.79E-05 | CDD1 | XII | 626502 | 626930 | -1 |
| YLR246W | 135.820924 | 2.86711411 | 0.66659612 | 70.5312315 | 4.83E-16 | 6.08E-14 | ERF2 | XII | 627118 | 628197 | 1 |
| YLR247C | 49.5707201 | 0.53955688 | 0.67374007 | 1.79899013 | 0.406775 | 0.57811175 | IRC20 | XII | 628684 | 633354 | -1 |
| YLR248W | 140.207065 | 1.36103368 | 0.5358592 | 30.2446738 | 2.71E-07 | 5.70E-06 | RCK2 | XII | 634252 | 636084 | 1 |
| YLR249W | 719.903702 | -0.4392935 | 0.60101004 | 15.1642588 | 0.00050948 | 0.0037336 | YEF3 | XII | 636780 | 639914 | 1 |
| YLR250W | 54.0795459 | 0.94704542 | 0.81517344 | 2.8800703 | 0.23691943 | 0.40475196 | SSP120 | XII | 640317 | 641021 | 1 |
| YLR251W | 154.186411 | -1.0134238 | 0.50782369 | 14.6725527 | 0.00065147 | 0.00461127 | SYM1 | XII | 641464 | 642057 | 1 |
| YLR253W | 138.062796 | 1.48024749 | 0.70067433 | 4.28450053 | 0.11739039 | 0.24546419 | MCP2 | XII | 642627 | 644336 | 1 |
| YLR254C | 74.8341915 | 1.05571837 | 0.71623586 | 4.23988559 | 0.1200385 | 0.24929085 | NDL1 | XII | 644404 | 644973 | -1 |
| YLR255C | 34.1392086 | 0.38314278 | 0.81067652 | 1.24903548 | 0.53551963 | 0.69010876 | NA | XII | 645600 | 645953 | -1 |
| YLR256W | 648.155449 | -0.7362087 | 0.54378521 | 7.079381 | 0.02902231 | 0.08675228 | HAP1 | XII | 646415 | 650923 | 1 |
| YLR256W-A | 231.759355 | 0.22112041 | 0.52454303 | 1.37478687 | 0.50288516 | 0.66130027 | NA | XII | 651117 | 652439 | 1 |
| YLR257W | 142.556692 | 1.09797556 | 0.57570436 | 3.832509 | 0.14715711 | 0.28912102 | NA | XII | 658826 | 659791 | 1 |
| YLR258W | 127.354862 | -0.7270829 | 0.5150197 | 3.74359195 | 0.15384711 | 0.29821719 | GSY2 | XII | 660716 | 662833 | 1 |
| YLR259C | 95.156923 | -0.2461283 | 0.71789432 | 0.24142623 | 0.88628818 | 0.93215081 | HSP60 | XII | 663284 | 665002 | -1 |
| YLR260W | 186.661515 | -0.5520474 | 0.54579648 | 11.181854 | 0.00373157 | 0.01817138 | LCB5 | XII | 665844 | 667907 | 1 |
| YLR262C-A | 36.9447575 | -1.0138005 | 0.79759401 | 2.55127162 | 0.27925336 | 0.45351049 | TMA7 | XII | 669468 | 669662 | -1 |
| YLR263W | 119.049387 | -1.1488538 | 0.50489297 | 15.8988759 | 0.00035286 | 0.00279489 | RED1 | XII | 670340 | 672823 | 1 |
| YLR264C-A | 19804.6944 | -0.3189184 | 0.52047871 | 36.9086183 | 9.67E-09 | 2.90E-07 | NA | XII | 673828 | 673944 | -1 |
| YLR264W | 16.3651907 | -1.3840731 | 0.97496652 | 5.22514296 | 0.07334569 | 0.17466992 | RPS28B | XII | 673131 | 673334 | 1 |
| YLR265C | 83.9125608 | -1.5428824 | 0.54987708 | 17.5316437 | 0.00015597 | 0.00137666 | NEJ1 | XII | 674427 | 675455 | -1 |
| YLR266C | 1349.55904 | -0.3442538 | 0.46623366 | 5.22442568 | 0.073372 | 0.17466992 | PDR8 | XII | 675619 | 677724 | -1 |
| YLR267W | 428.704063 | -0.9047594 | 0.47077814 | 16.2568327 | 0.00029504 | 0.0024162 | BOP2 | XII | 678212 | 679924 | 1 |
| YLR268W | 27.3274348 | -1.2550706 | 0.71505679 | 9.66550595 | 0.00796456 | 0.03310469 | SEC22 | XII | 680200 | 680844 | 1 |
| YLR270W | 12.2292995 | 0.78139129 | 1.04400489 | 1.65198617 | 0.4378 | 0.60642632 | DCS1 | XII | 681186 | 682238 | 1 |
| YLR271W | 87.8881142 | -1.0176252 | 0.54602602 | 16.8445262 | 0.00021992 | 0.001854 | CMG1 | XII | 682737 | 683561 | 1 |

| | | | | | | | | | | | |
|---|---|---|---|---|---|---|---|---|---|---|---|
| YLR272C | 235.979517 | -0.0874387 | 0.50611512 | 0.80441453 | 0.6688421 | 0.7945416 | YCS4 | XII | 683672 | 687202 | -1 |
| YLR273C | 20.7762509 | -1.0680131 | 0.83035786 | 9.06414335 | 0.01075837 | 0.04161051 | PIG1 | XII | 689083 | 691029 | -1 |
| YLR274W | 123.651581 | 0.37865339 | 0.66211204 | 69.7782342 | 7.04E-16 | 8.50E-14 | MCM5 | XII | 691555 | 693882 | 1 |
| YLR275W | 7.32997374 | -0.2160435 | 1.0407381 | 2.54019423 | 0.28080435 | 0.45491247 | SMD2 | XII | 694378 | 694800 | 1 |
| YLR276C | 93.884684 | 0.68136338 | 0.82419338 | 4.4051906 | 0.11051596 | 0.23533925 | DBP9 | XII | 695046 | 696830 | -1 |
| YLR277C | 11.8517903 | -0.9873614 | 1.04798939 | 1.01973503 | 0.60057514 | 0.74682758 | YSH1 | XII | 697156 | 699495 | -1 |
| YLR278C | 848.791947 | -0.5883047 | 0.4832132 | 2.04035597 | 0.36053077 | 0.53345156 | NA | XII | 699999 | 704024 | -1 |
| YLR281C | 28.0199004 | 0.15296588 | 0.68355328 | 1.22647853 | 0.54159366 | 0.69561386 | NA | XII | 704493 | 704960 | -1 |
| YLR283W | 17.8299394 | 0.57647317 | 0.826968 | 0.91892385 | 0.63162341 | 0.76880708 | NA | XII | 705186 | 706130 | 1 |
| YLR284C | 12.7107304 | 0.52358117 | 0.89172643 | 6.61237191 | 0.03665571 | 0.10424194 | ECI1 | XII | 706198 | 707040 | -1 |
| YLR285C-A | 675.63591 | 0.13642047 | 0.52929633 | 19.4412923 | 6.00E-05 | 0.0006041 | NA | XII | 708168 | 708338 | -1 |
| YLR285W | 125.000001 | -0.2318214 | 0.50833067 | 0.29494175 | 0.86288757 | 0.91958753 | NNT1 | XII | 707360 | 708145 | 1 |
| YLR286C | 162.40594 | -1.0009551 | 0.49690684 | 9.00077405 | 0.0111047 | 0.04258027 | CTS1 | XII | 708448 | 710136 | -1 |
| YLR287C | 40.8955548 | -0.778659 | 0.62707843 | 1.73856023 | 0.41925326 | 0.59032562 | NA | XII | 710991 | 712058 | -1 |
| YLR287C-A | 143.761962 | -0.6894414 | 0.77341983 | 7.77743391 | 0.0204716 | 0.06719418 | RPS30A | XII | 712537 | 713158 | -1 |
| YLR288C | 38.8439112 | -0.0836418 | 0.60636056 | 0.6809587 | 0.71142922 | 0.82438173 | MEC3 | XII | 713480 | 714904 | -1 |
| YLR289W | 29.2050879 | 0.7757766 | 0.75831752 | 1.88660334 | 0.38934024 | 0.56101577 | GUF1 | XII | 715089 | 717026 | 1 |
| YLR290C | 25.7809314 | 0.18325224 | 0.70973189 | 0.4022548 | 0.81780823 | 0.89056041 | COQ11 | XII | 717145 | 717978 | -1 |
| YLR291C | 15.0142801 | -0.2331676 | 0.9044881 | 0.32242465 | 0.85111134 | 0.91173629 | GCD7 | XII | 718317 | 719462 | -1 |
| YLR292C | 30.510899 | 0.53317234 | 0.88225502 | 0.56823596 | 0.75267784 | 0.85034238 | SEC72 | XII | 719789 | 720370 | -1 |
| YLR293C | 73.8217827 | -0.3360884 | 0.68170713 | 8.1311459 | 0.01715316 | 0.05904684 | GSP1 | XII | 720771 | 721430 | -1 |
| YLR295C | 18.9681436 | -1.4697465 | 0.77501447 | 12.4597866 | 0.00196966 | 0.01104002 | ATP14 | XII | 721999 | 722373 | -1 |
| YLR296W | 6.55619836 | -1.2134759 | 1.22214559 | 1.75547948 | 0.41572149 | 0.58736638 | NA | XII | 722978 | 723304 | 1 |
| YLR297W | 800.522384 | 1.33340329 | 0.53436657 | 43.3073909 | 3.94E-10 | 1.61E-08 | NA | XII | 724044 | 724433 | 1 |
| YLR298C | 154.999774 | 0.61657032 | 0.51708883 | 9.82476557 | 0.00735494 | 0.03119789 | YHC1 | XII | 724721 | 725416 | -1 |
| YLR299W | 255.76528 | 2.02185601 | 0.61255006 | 60.6769801 | 6.67E-14 | 5.76E-12 | ECM38 | XII | 726069 | 728051 | 1 |
| YLR300W | 201.328489 | 0.70335984 | 0.77061104 | 0.87717147 | 0.6449479 | 0.77748248 | EXG1 | XII | 728955 | 730301 | 1 |
| YLR301W | 29.7894719 | 0.03838667 | 0.6837825 | 0.73876911 | 0.69115957 | 0.81221741 | HRI1 | XII | 730825 | 731559 | 1 |
| YLR302C | 717.643346 | -0.8360056 | 0.71783394 | 5.93114773 | 0.05153089 | 0.13385547 | NA | XII | 732191 | 732553 | -1 |
| YLR303W | 46.0839993 | -0.5739489 | 0.59913325 | 1.23016112 | 0.54059734 | 0.69448827 | MET17 | XII | 732542 | 733876 | 1 |
| YLR304C | 46.5930886 | -0.4310137 | 0.65092355 | 6.67410748 | 0.03554152 | 0.10170914 | ACO1 | XII | 735212 | 737548 | -1 |
| YLR305C | 20.1229581 | -0.92954 | 0.75005248 | 4.45743308 | 0.10766653 | 0.2307994 | STT4 | XII | 738161 | 743863 | -1 |
| YLR306W | 11.3990355 | -0.7083163 | 0.94580353 | 0.62747756 | 0.73070988 | 0.83678999 | UBC12 | XII | 744151 | 744851 | 1 |

| | | | | | | | | | | | |
|---|---|---|---|---|---|---|---|---|---|---|---|
| YLR308W | 29.7025968 | 0.59514078 | 0.83865563 | 2.06522705 | 0.35607513 | 0.52942744 | CDA2 | XII | 747937 | 748875 | 1 |
| YLR309C | 110.525508 | -0.5957442 | 0.56731955 | 1.09177387 | 0.57932773 | 0.72802613 | IMH1 | XII | 749034 | 751769 | -1 |
| YLR310C | 43.069659 | -0.0352506 | 0.6590963 | 0.16101311 | 0.92264885 | 0.95140461 | CDC25 | XII | 752224 | 756993 | -1 |
| YLR312C | 26.4464701 | -1.1756494 | 0.69022455 | 2.88054341 | 0.23686339 | 0.40475196 | ATG39 | XII | 757637 | 758833 | -1 |
| YLR312W-A | 11.9168189 | -0.0876011 | 0.98714438 | 5.52056113 | 0.06327401 | 0.15616221 | MRPL15 | XII | 759480 | 760241 | 1 |
| YLR313C | 18.4485657 | -1.0054067 | 0.77438306 | 1.74047551 | 0.41885195 | 0.59006151 | NA | XII | 760750 | 762342 | -1 |
| YLR314C | 42.8119131 | 0.42394457 | 0.66393277 | 1.60087391 | 0.44913267 | 0.61535214 | CDC3 | XII | 762575 | 764137 | -1 |
| YLR315W | 68.8224074 | -0.0856987 | 0.56569464 | 7.06245681 | 0.02926894 | 0.08719023 | NKP2 | XII | 764808 | 765269 | 1 |
| YLR316C | 1328.8196 | -0.1498942 | 0.51463627 | 0.70666147 | 0.70234487 | 0.81847842 | TAD3 | XII | 765266 | 766358 | -1 |
| YLR318W | 45.2811153 | 1.13407124 | 0.90577416 | 3.72011618 | 0.15566359 | 0.30083183 | EST2 | XII | 766542 | 769196 | 1 |
| YLR319C | 26.0446135 | -0.0447493 | 0.94396981 | 1.54880253 | 0.4609797 | 0.62486036 | BUD6 | XII | 769318 | 771684 | -1 |
| YLR320W | 80.5201251 | 0.3582032 | 0.57920716 | 8.24626336 | 0.01619372 | 0.05674873 | MMS22 | XII | 771940 | 776304 | 1 |
| YLR321C | 109.026833 | -0.4630717 | 0.65027437 | 2.88141496 | 0.2367602 | 0.40473806 | SFH1 | XII | 776584 | 777864 | -1 |
| YLR323C | 15.6343226 | -1.6024263 | 0.95962066 | 2.95914199 | 0.22773537 | 0.39372582 | CWC24 | XII | 778173 | 778952 | -1 |
| YLR324W | 96.8263752 | -0.2570321 | 0.66970471 | 12.3402019 | 0.00209102 | 0.0115746 | PEX30 | XII | 779215 | 780786 | 1 |
| YLR325C | 82.938319 | -1.064147 | 0.66098628 | 12.98018 | 0.00151841 | 0.00921552 | RPL38 | XII | 781143 | 781379 | -1 |
| YLR326W | 20.5301731 | -0.3731071 | 0.79588351 | 3.81304897 | 0.14859594 | 0.29099512 | NA | XII | 782174 | 782896 | 1 |
| YLR327C | 18.2925609 | 1.27686445 | 1.01938867 | 11.5892557 | 0.00304386 | 0.01538156 | TMA10 | XII | 783127 | 783387 | -1 |
| YLR328W | 22.4730086 | -0.0924126 | 0.7275121 | 2.83726612 | 0.24204465 | 0.41122023 | NMA1 | XII | 784913 | 786118 | 1 |
| YLR329W | 153.122859 | -0.2301805 | 0.51836415 | 0.23169318 | 0.89061184 | 0.93317728 | REC102 | XII | 786442 | 787333 | 1 |
| YLR330W | 140.004968 | 0.84221529 | 0.5332075 | 2.94558577 | 0.22928423 | 0.39593071 | CHS5 | XII | 787664 | 789679 | 1 |
| YLR331C | 9223.46821 | 2.03670663 | 0.76246382 | 7.48541448 | 0.02368988 | 0.07502674 | NA | XII | 790669 | 791046 | -1 |
| YLR332W | 42.1062734 | 1.67560771 | 0.69433535 | 12.5874319 | 0.00184788 | 0.01056192 | MID2 | XII | 790676 | 791806 | 1 |
| YLR333C | 56.1160416 | -1.1852026 | 0.81225395 | 13.4171334 | 0.00122041 | 0.00776032 | RPS25B | XII | 795573 | 795899 | -1 |
| YLR335W | 124.261284 | 0.44806363 | 0.63559858 | 3.62251491 | 0.16344848 | 0.3120399 | NUP2 | XII | 797430 | 799592 | 1 |
| YLR336C | 63.9616569 | 0.53713886 | 0.67979811 | 0.6945959 | 0.70659477 | 0.82201802 | SGD1 | XII | 799697 | 802396 | -1 |
| YLR337C | 270.572216 | 1.66102661 | 0.55590187 | 28.2257013 | 7.43E-07 | 1.38E-05 | VRP1 | XII | 802653 | 805106 | -1 |
| YLR340W | 111.37672 | -0.3767988 | 0.59768985 | 13.2990529 | 0.00129464 | 0.00812103 | RPP0 | XII | 805887 | 806825 | 1 |
| YLR341W | 438.074077 | -0.7283969 | 0.48499484 | 13.6009588 | 0.00111324 | 0.00718581 | SPO77 | XII | 807385 | 808818 | 1 |
| YLR342W | 180.353166 | 0.70167583 | 0.55851061 | 6.23754315 | 0.04421145 | 0.11945136 | FKS1 | XII | 809997 | 815627 | 1 |
| YLR343W | 37.811294 | -1.0633423 | 0.68527978 | 2.41602404 | 0.29879068 | 0.47191811 | GAS2 | XII | 816094 | 817761 | 1 |
| YLR344W | 31.4887038 | -0.493569 | 0.63927924 | 9.17163074 | 0.01019543 | 0.04004855 | RPL26A | XII | 819312 | 820142 | 1 |
| YLR345W | 29.7648419 | 2.3986129 | 0.76025151 | 13.0132195 | 0.00149353 | 0.0091027 | NA | XII | 820511 | 822040 | 1 |

| | | | | | | | | | | | |
|---|---|---|---|---|---|---|---|---|---|---|---|
| YLR347C | 121.164709 | 1.24302714 | 0.5897991 | 6.92266851 | 0.03138785 | 0.09258058 | KAP95 | XII | 823828 | 826413 | -1 |
| YLR348C | 735.9817 | 4.49991925 | 0.60309877 | 50.5299894 | 1.07E-11 | 6.20E-10 | DIC1 | XII | 826977 | 827873 | -1 |
| YLR350W | 325.863296 | 2.24029777 | 0.75375479 | 9.16039721 | 0.01025286 | 0.04019232 | ORM2 | XII | 828730 | 829380 | 1 |
| YLR351C | 168.959922 | 2.11215278 | 0.69147096 | 12.3070704 | 0.00212595 | 0.01170082 | NIT3 | XII | 829489 | 830364 | -1 |
| YLR352W | 238.131289 | -0.555183 | 0.50041283 | 2.05913871 | 0.35716074 | 0.53038232 | NA | XII | 831116 | 833539 | 1 |
| YLR353W | 119.417136 | -0.1870396 | 0.52759406 | 1.83622676 | 0.3992716 | 0.57010668 | BUD8 | XII | 834352 | 836163 | 1 |
| YLR354C | 84.6699874 | 0.1277171 | 0.64078247 | 2.07790003 | 0.353826 | 0.5275953 | TAL1 | XII | 836350 | 837357 | -1 |
| YLR355C | 95.3119312 | -0.5744034 | 0.58462489 | 16.2353517 | 0.00029822 | 0.0024354 | ILV5 | XII | 838066 | 839253 | -1 |
| YLR356W | 39.8683312 | -0.3451662 | 0.64445367 | 2.09754991 | 0.3503667 | 0.523786 | ATG33 | XII | 840321 | 840914 | 1 |
| YLR357W | 19.0269109 | -0.1751026 | 0.76284209 | 0.70835916 | 0.70174894 | 0.81835374 | RSC2 | XII | 841331 | 844000 | 1 |
| YLR358C | 2052.82584 | -0.2213392 | 0.59800427 | 3.48250916 | 0.17530033 | 0.32699386 | NA | XII | 843487 | 844050 | -1 |
| YLR359W | 50.2970492 | 0.12828642 | 0.83370318 | 1.70585737 | 0.426165 | 0.59586944 | ADE13 | XII | 844282 | 845730 | 1 |
| YLR360W | 933.104889 | 0.16510761 | 0.46911179 | 1.00139145 | 0.60610883 | 0.75088539 | VPS38 | XII | 846103 | 847422 | 1 |
| YLR361C | 709.266517 | 1.00502556 | 0.65491309 | 2.46829378 | 0.29108298 | 0.46518644 | DCR2 | XII | 847388 | 849124 | -1 |
| YLR361C-A | 32.947318 | 0.5180561 | 0.6724747 | 0.70724354 | 0.70214049 | 0.81847842 | NA | XII | 849383 | 849679 | -1 |
| YLR362W | 25.2618034 | 1.02386306 | 0.80643841 | 2.04497741 | 0.35969864 | 0.53284736 | STE11 | XII | 849866 | 852019 | 1 |
| YLR363C | 13.557029 | 0.72710134 | 1.02189025 | 6.47615951 | 0.03923917 | 0.10991524 | NMD4 | XII | 852495 | 853151 | -1 |
| YLR363W-A | 170.483024 | 0.55772091 | 0.53286706 | 3.81781537 | 0.14824223 | 0.29064152 | NA | XII | 853461 | 853718 | 1 |
| YLR364W | 396.036105 | 0.64101135 | 0.49081374 | 5.91673649 | 0.05190354 | 0.13440139 | GRX8 | XII | 854062 | 854391 | 1 |
| YLR365W | 21.1043865 | -0.4813659 | 0.81875893 | 1.02382389 | 0.59934856 | 0.74628563 | NA | XII | 855199 | 855531 | 1 |
| YLR367W | 10.8332111 | -0.1458407 | 0.95551203 | 2.4927013 | 0.28755226 | 0.46171037 | RPS22B | XII | 856442 | 857317 | 1 |
| YLR368W | 16.7173321 | -0.1604826 | 1.0112039 | 6.10663417 | 0.04720209 | 0.12525211 | MDM30 | XII | 857540 | 859336 | 1 |
| YLR369W | 39.112502 | 2.03435561 | 0.80630239 | 9.40891199 | 0.00905484 | 0.03655068 | SSQ1 | XII | 859552 | 861525 | 1 |
| YLR370C | 15.6788637 | 0.2850948 | 0.81041005 | 0.16003411 | 0.9231006 | 0.95153151 | ARC18 | XII | 861718 | 862254 | -1 |
| YLR371W | 33.6482803 | -0.6286952 | 0.65452744 | 3.2160664 | 0.20028114 | 0.35924034 | ROM2 | XII | 862714 | 866784 | 1 |
| YLR372W | 212.946601 | 2.38663342 | 0.61335405 | 44.1592673 | 2.58E-10 | 1.09E-08 | ELO3 | XII | 867354 | 868391 | 1 |
| YLR373C | 57.9402541 | -0.1510449 | 0.64247793 | 3.10454301 | 0.2117664 | 0.37530684 | VID22 | XII | 868662 | 871367 | -1 |
| YLR375W | 86.0502495 | -0.535484 | 0.58186636 | 0.93457405 | 0.62670019 | 0.76471951 | STP3 | XII | 871697 | 872728 | 1 |
| YLR376C | 102.084059 | 3.35206265 | 1.68092634 | 3.67869753 | 0.15892089 | 0.30559679 | PSY3 | XII | 872826 | 873554 | -1 |
| YLR377C | 480.080057 | -0.4346305 | 0.49395025 | 2.03548504 | 0.3614099 | 0.53408966 | FBP1 | XII | 873746 | 874792 | -1 |
| YLR378C | 46.3427399 | 0.50445665 | 0.59960022 | 0.94323259 | 0.6239929 | 0.76326014 | SEC61 | XII | 875736 | 877178 | -1 |
| YLR380W | 31.1977536 | 0.15130539 | 0.66693409 | 2.04096518 | 0.36042096 | 0.53345156 | CSR1 | XII | 878282 | 879508 | 1 |
| YLR381W | 53.474129 | -0.1653339 | 0.61272827 | 2.69109941 | 0.26039653 | 0.43188081 | CTF3 | XII | 879723 | 881924 | 1 |

| | | | | | | | | | | | |
|---|---|---|---|---|---|---|---|---|---|---|---|
| YLR382C | 98.5483013 | -0.1883231 | 0.8178844 | 31.1424796 | 1.73E-07 | 3.88E-06 | NAM2 | XII | 882067 | 884751 | -1 |
| YLR383W | 18.8467481 | -0.802092 | 0.76334244 | 2.87563269 | 0.23744569 | 0.4053281 | SMC6 | XII | 885288 | 888632 | 1 |
| YLR384C | 26.8725254 | 0.11156858 | 0.76245366 | 2.26896898 | 0.32158786 | 0.49429087 | IKI3 | XII | 888851 | 892900 | -1 |
| YLR386W | 146.954453 | 0.91686941 | 0.7376517 | 12.5572496 | 0.00187598 | 0.01069086 | VAC14 | XII | 893628 | 896270 | 1 |
| YLR387C | 126.377764 | 0.13119533 | 0.52784676 | 6.40169714 | 0.04072763 | 0.11285424 | REH1 | XII | 896374 | 897672 | -1 |
| YLR388W | 71.5939671 | -0.4888119 | 0.5823232 | 10.4980046 | 0.00525276 | 0.02385369 | RPS29A | XII | 898651 | 898821 | 1 |
| YLR389C | 73.7614573 | -0.4491566 | 0.56911736 | 4.89220766 | 0.08663046 | 0.19825705 | STE23 | XII | 899577 | 902660 | -1 |
| YLR390W | 86.0331729 | 0.44179141 | 0.55701322 | 2.71129219 | 0.25778069 | 0.42975819 | ECM19 | XII | 903066 | 903404 | 1 |
| YLR390W-A | 50.3365091 | 1.12070344 | 0.61950376 | 3.60202665 | 0.16513147 | 0.31399383 | CCW14 | XII | 903724 | 904440 | 1 |
| YLR392C | 17.1737233 | -0.0847962 | 0.76744344 | 0.12271621 | 0.94048639 | 0.96082012 | ART10 | XII | 904749 | 906305 | -1 |
| YLR393W | 98.740742 | -0.484157 | 0.75433908 | 23.7027882 | 7.13E-06 | 9.69E-05 | ATP10 | XII | 907079 | 907918 | 1 |
| YLR394W | 132.497536 | 0.44537887 | 0.60118695 | 27.6279929 | 1.00E-06 | 1.77E-05 | CST9 | XII | 907950 | 909398 | 1 |
| YLR395C | 138.842959 | 0.39490095 | 0.60713292 | 8.52514377 | 0.01408603 | 0.05103761 | COX8 | XII | 909729 | 909965 | -1 |
| YLR396C | 397.83144 | -1.0562606 | 0.50448255 | 10.7949223 | 0.00452806 | 0.02104132 | VPS33 | XII | 910235 | 912310 | -1 |
| YLR397C | 88.9129803 | -0.5318902 | 0.60010815 | 7.85447528 | 0.01969801 | 0.06517227 | AFG2 | XII | 912550 | 914892 | -1 |
| YLR398C | 46.8528721 | -0.3177075 | 0.59454539 | 0.46299273 | 0.79334558 | 0.87452157 | SKI2 | XII | 915156 | 919019 | -1 |
| YLR399C | 75.0045811 | -0.3423153 | 0.5524405 | 0.83503994 | 0.65867834 | 0.78659971 | BDF1 | XII | 919536 | 921596 | -1 |
| YLR401C | 73.3995716 | -0.6977809 | 0.58630628 | 4.02838026 | 0.13342842 | 0.26946304 | DUS3 | XII | 922442 | 924448 | -1 |
| YLR403W | 20.7042522 | -0.1082425 | 0.7798059 | 0.13023441 | 0.93695764 | 0.95898617 | SFP1 | XII | 925568 | 927619 | 1 |
| YLR404W | 31.545634 | -0.5115558 | 0.66996365 | 0.63016281 | 0.72972947 | 0.83678999 | SEI1 | XII | 928743 | 929600 | 1 |
| YLR405W | 1654.88203 | 2.48368971 | 1.65852777 | 6.23293907 | 0.04431334 | 0.11967082 | DUS4 | XII | 929789 | 930892 | 1 |
| YLR406C | 64.5962666 | -0.6559046 | 0.62588889 | 11.8472367 | 0.0026755 | 0.013956 | RPL31B | XII | 931065 | 931755 | -1 |
| YLR407W | 66.6631141 | -0.1497237 | 0.59480956 | 3.32173737 | 0.18997388 | 0.34583253 | NA | XII | 932967 | 933653 | 1 |
| YLR408C | 66.1328068 | 1.2082589 | 0.60461938 | 15.522633 | 0.0004259 | 0.00323614 | BLS1 | XII | 933885 | 934253 | -1 |
| YLR409C | 36.3843517 | -0.9034421 | 0.70835308 | 7.92605436 | 0.01900549 | 0.06364477 | UTP21 | XII | 934414 | 937233 | -1 |
| YLR410W | 393.465136 | -0.7609321 | 0.48183051 | 6.39673558 | 0.04082879 | 0.11305473 | VIP1 | XII | 937541 | 940981 | 1 |
| YLR410W-B | 14.2480751 | 0.07901571 | 0.79942639 | 5.01622336 | 0.08142184 | 0.18867385 | NA | XII | 941483 | 946796 | 1 |
| YLR411W | 125.831203 | -0.4888614 | 0.54355426 | 2.50699167 | 0.28550497 | 0.45905965 | CTR3 | XII | 947253 | 947978 | 1 |
| YLR412W | 117.485249 | -1.2340067 | 0.54242437 | 10.2882482 | 0.00583358 | 0.02592206 | BER1 | XII | 948368 | 949192 | 1 |
| YLR413W | 528.507555 | 2.48899869 | 0.46176305 | 26.8421067 | 1.48E-06 | 2.50E-05 | INA1 | XII | 951156 | 953183 | 1 |
| YLR414C | 44.755932 | 0.70783383 | 0.70806515 | 1.42821436 | 0.48962907 | 0.65106127 | PUN1 | XII | 953353 | 954144 | -1 |
| YLR415C | 16.7057583 | -1.2102161 | 0.88720917 | 2.00738001 | 0.36652447 | 0.53971939 | NA | XII | 954259 | 954597 | -1 |
| YLR417W | 298.154617 | 0.59581921 | 0.52809638 | 4.00236118 | 0.1351756 | 0.27194814 | VPS36 | XII | 955010 | 956710 | 1 |

| | | | | | | | | | | | |
|---|---|---|---|---|---|---|---|---|---|---|---|
| YLR418C | 120.183553 | 1.55481862 | 0.61060195 | 39.8235649 | 2.25E-09 | 7.75E-08 | CDC73 | XII | 956914 | 958095 | -1 |
| YLR419W | 70.0461237 | -0.0818332 | 0.5408059 | 2.07174753 | 0.35491613 | 0.52857757 | NA | XII | 958428 | 962735 | 1 |
| YLR420W | 471.658182 | 1.65245356 | 0.69730057 | 6.73034408 | 0.03455607 | 0.09964126 | URA4 | XII | 963785 | 964879 | 1 |
| YLR421C | 42.0783491 | -0.1179658 | 0.75291307 | 0.06540246 | 0.96782768 | 0.97506912 | RPN13 | XII | 965090 | 965560 | -1 |
| YLR422W | 173.120198 | 1.66597663 | 0.62851184 | 13.6481553 | 0.00108728 | 0.00708137 | DCK1 | XII | 965897 | 971695 | 1 |
| YLR423C | 1821.66516 | 5.24893061 | 0.4906834 | 114.614454 | 1.29E-25 | 9.36E-23 | ATG17 | XII | 971917 | 973170 | -1 |
| YLR424W | 34.4625557 | 0.5842125 | 0.73770451 | 18.9698018 | 7.60E-05 | 0.00074072 | SPP382 | XII | 973395 | 975521 | 1 |
| YLR425W | 46.6984099 | -1.0676931 | 0.64015479 | 4.78090392 | 0.09158828 | 0.2057006 | TUS1 | XII | 982894 | 986817 | 1 |
| YLR426W | 40.0973189 | 1.6412429 | 0.68128826 | 8.37296192 | 0.01519968 | 0.05409105 | TDA5 | XII | 987062 | 988113 | 1 |
| YLR427W | 299.44852 | 2.43888323 | 0.51454426 | 21.6886717 | 1.95E-05 | 0.00023201 | MAG2 | XII | 988428 | 990440 | 1 |
| YLR429W | 222.879273 | 1.07358337 | 0.56835407 | 10.1902899 | 0.00612642 | 0.02692266 | CRN1 | XII | 990777 | 992732 | 1 |
| YLR430W | 108.245377 | 1.19490285 | 0.6447262 | 3.75615096 | 0.15288405 | 0.29674779 | SEN1 | XII | 993434 | 1E+06 | 1 |
| YLR431C | 80.8238326 | 0.98795368 | 0.60449058 | 31.3093365 | 1.59E-07 | 3.62E-06 | ATG23 | XII | 1000342 | 1E+06 | -1 |
| YLR432W | 152.716864 | 1.12680925 | 0.70529825 | 5.20919739 | 0.0739328 | 0.17558282 | IMD3 | XII | 1002557 | 1E+06 | 1 |
| YLR433C | 133.255444 | 0.76846285 | 0.62147945 | 6.77916082 | 0.03372282 | 0.09792134 | CNA1 | XII | 1004347 | 1E+06 | -1 |
| YLR435W | 34.2266832 | -1.4190372 | 0.69088993 | 14.5250107 | 0.00070135 | 0.00488665 | TSR2 | XII | 1006378 | 1E+06 | 1 |
| YLR436C | 218.530457 | 0.22647061 | 0.50878641 | 5.80590635 | 0.05486097 | 0.14011689 | ECM30 | XII | 1007421 | 1E+06 | -1 |
| YLR437C | 67.4003137 | -0.4519424 | 0.55411374 | 3.21057221 | 0.20083209 | 0.36006531 | DIF1 | XII | 1011622 | 1E+06 | -1 |
| YLR438C-A | 46.3295786 | -0.0709462 | 0.64647026 | 3.34165533 | 0.18809132 | 0.34376539 | LSM3 | XII | 1013909 | 1E+06 | -1 |
| YLR438W | 100.013468 | -1.2945613 | 0.53713548 | 7.26623887 | 0.0264336 | 0.08085078 | CAR2 | XII | 1012501 | 1E+06 | 1 |
| YLR439W | 130.660229 | 0.671385 | 0.55326407 | 4.43306447 | 0.10898639 | 0.23299267 | MRPL4 | XII | 1014491 | 1E+06 | 1 |
| YLR440C | 76.9080171 | 0.12519234 | 0.56577967 | 0.0948181 | 0.95369721 | 0.96767513 | SEC39 | XII | 1015568 | 1E+06 | -1 |
| YLR441C | 194.944006 | -0.5477081 | 0.69248605 | 18.0121966 | 0.00012266 | 0.00111316 | RPS1A | XII | 1018141 | 1E+06 | -1 |
| YLR442C | 18.4248633 | -0.0509899 | 0.94961849 | 0.51963827 | 0.77119105 | 0.86067776 | SIR3 | XII | 1019315 | 1E+06 | -1 |
| YLR443W | 32.1298008 | -1.1391972 | 0.68536888 | 3.20381388 | 0.20151188 | 0.36088889 | ECM7 | XII | 1022625 | 1E+06 | 1 |
| YLR445W | 31.6843425 | -1.6268432 | 0.6750403 | 7.34319792 | 0.02543577 | 0.07871357 | GMC2 | XII | 1024189 | 1E+06 | 1 |
| YLR446W | 1160.13324 | -1.2868682 | 0.45621558 | 36.3519797 | 1.28E-08 | 3.72E-07 | NA | XII | 1025214 | 1E+06 | 1 |
| YLR447C | 26.9764563 | -0.8358682 | 0.79209808 | 1.46445824 | 0.48083595 | 0.64344033 | VMA6 | XII | 1026856 | 1E+06 | -1 |
| YLR448W | 90.2431114 | -0.5220863 | 0.5905778 | 25.5618581 | 2.81E-06 | 4.33E-05 | RPL6B | XII | 1028854 | 1E+06 | 1 |
| YLR449W | 476.127598 | 0.59667547 | 0.59915718 | 8.94720999 | 0.01140612 | 0.04341976 | FPR4 | XII | 1030834 | 1E+06 | 1 |
| YLR450W | 352.791065 | -0.5744331 | 0.49877044 | 8.60506293 | 0.01353425 | 0.04937874 | HMG2 | XII | 1032627 | 1E+06 | 1 |
| YLR451W | 61.1017117 | -0.6238624 | 0.56972752 | 1.22273144 | 0.54260931 | 0.69630051 | LEU3 | XII | 1036093 | 1E+06 | 1 |
| YLR452C | 16.0225878 | -1.0406067 | 0.82717952 | 3.34730816 | 0.18756045 | 0.34351531 | SST2 | XII | 1039270 | 1E+06 | -1 |

| | | | | | | | | | | | |
|---|---|---|---|---|---|---|---|---|---|---|---|
| YLR453C | 143.578216 | -1.5443615 | 0.53211917 | 19.3763318 | 6.20E-05 | 0.00061906 | RIF2 | XII | 1041799 | 1E+06 | -1 |
| YLR454W | 54.2528009 | -0.1580567 | 0.650277 | 4.82365903 | 0.08965113 | 0.20308295 | FMP27 | XII | 1043998 | 1E+06 | 1 |
| YLR455W | 66.0789261 | 0.75805314 | 0.75790118 | 2.13241116 | 0.34431251 | 0.51781024 | PDP3 | XII | 1053629 | 1E+06 | 1 |
| YLR456W | 83.1167269 | 1.0009394 | 0.66858179 | 2.24989541 | 0.32466945 | 0.49770614 | NA | XII | 1055070 | 1E+06 | 1 |
| YLR457C | 13.366799 | 0.66477073 | 0.93763788 | 1.40502538 | 0.4953391 | 0.65554592 | NBP1 | XII | 1055812 | 1E+06 | -1 |
| YLR459W | 38.928909 | 1.47369675 | 0.84100969 | 5.57725217 | 0.06150566 | 0.15245492 | GAB1 | XII | 1057334 | 1E+06 | 1 |
| YLR460C | 25.3861504 | -1.7209221 | 0.75451184 | 8.89592136 | 0.01170241 | 0.04434355 | NA | XII | 1059757 | 1E+06 | -1 |
| YLR462W | 36.1186258 | -1.2406237 | 0.6875768 | 3.61739015 | 0.16386783 | 0.31248127 | NA | XII | 1065956 | 1E+06 | 1 |
| YLR466W | 7.9540639 | -1.6931054 | 1.06034906 | 6.14387703 | 0.04633125 | 0.12328031 | YRF1-4 | XII | 1067087 | 1E+06 | 1 |
| YML001W | 236.853208 | -1.4116366 | 0.55380535 | 8.90100562 | 0.0116727 | 0.04427675 | YPT7 | XIII | 267174 | 267800 | 1 |
| YML002W | 445.524361 | -0.1713607 | 0.49140624 | 0.13620961 | 0.93416256 | 0.95735444 | NA | XIII | 264541 | 266754 | 1 |
| YML003W | 9.24664629 | 0.36251492 | 1.07470896 | 2.33205427 | 0.31160244 | 0.48405923 | NA | XIII | 263483 | 264355 | 1 |
| YML004C | 56.7630591 | 0.23558228 | 0.70891451 | 0.34123925 | 0.84314222 | 0.9080717 | GLO1 | XIII | 261705 | 262685 | -1 |
| YML005W | 1877.57392 | 2.62530912 | 0.8232044 | 9.55537661 | 0.00841543 | 0.03453249 | TRM12 | XIII | 260221 | 261609 | 1 |
| YML006C | 65.3871588 | 0.2688657 | 0.59874031 | 4.35419834 | 0.11336992 | 0.24000433 | GIS4 | XIII | 256092 | 258416 | -1 |
| YML007C-A | 39.3994828 | -1.4388963 | 0.70887403 | 4.4136652 | 0.11004867 | 0.23477589 | NA | XIII | 253162 | 253272 | -1 |
| YML007W | 60.4121514 | 0.06761266 | 0.60001281 | 1.59551857 | 0.45033691 | 0.6167102 | YAP1 | XIII | 253848 | 255800 | 1 |
| YML008C | 56.9592058 | 0.45722965 | 0.68311364 | 4.31482422 | 0.11562396 | 0.242911 | ERG6 | XIII | 251839 | 252990 | -1 |
| YML009C | 10.7955892 | -0.571008 | 0.9679817 | 3.5144804 | 0.17252033 | 0.32379018 | MRPL39 | XIII | 251304 | 251516 | -1 |
| YML010W | 97.062783 | 0.19377017 | 0.57785373 | 1.73666295 | 0.41965116 | 0.59032562 | SPT5 | XIII | 247677 | 250868 | 1 |
| YML011C | 327.532665 | 0.20357141 | 0.57511913 | 12.2322452 | 0.002207 | 0.0120552 | RAD33 | XIII | 246895 | 247428 | -1 |
| YML012W | 55.6623557 | 0.77151879 | 0.58413618 | 3.07433513 | 0.21498918 | 0.37881539 | ERV25 | XIII | 246116 | 246751 | 1 |
| YML013W | 64.5221968 | 0.54085659 | 0.58825561 | 10.3269134 | 0.00572189 | 0.02550402 | UBX2 | XIII | 244149 | 245903 | 1 |
| YML014W | 13.0560467 | -0.8738256 | 0.86461008 | 1.50061398 | 0.47222156 | 0.63393528 | TRM9 | XIII | 243225 | 244064 | 1 |
| YML015C | 140.164885 | 1.91269133 | 0.55380119 | 12.4807064 | 0.00194917 | 0.01096762 | TAF11 | XIII | 241989 | 243029 | -1 |
| YML016C | 54.6503683 | 1.52144599 | 0.60473511 | 6.56435077 | 0.03754649 | 0.1061495 | PPZ1 | XIII | 239458 | 241536 | -1 |
| YML017W | 163.291274 | -0.3912457 | 0.5147088 | 7.19129488 | 0.02744291 | 0.08314728 | PSP2 | XIII | 236588 | 238731 | 1 |
| YML018C | 51.0715818 | -0.3391627 | 0.6617756 | 4.35512561 | 0.11331737 | 0.23998083 | NA | XIII | 234771 | 235952 | -1 |
| YML019W | 82.8842753 | -1.8549265 | 0.57504139 | 12.7891516 | 0.00167059 | 0.00979022 | OST6 | XIII | 233457 | 234455 | 1 |
| YML020W | 72.0218702 | 0.15969778 | 0.54676263 | 1.64472451 | 0.43939247 | 0.60805219 | NA | XIII | 231149 | 233143 | 1 |
| YML021C | 148.924859 | -0.8650821 | 0.61482953 | 22.0400599 | 1.64E-05 | 0.00019996 | UNG1 | XIII | 229734 | 230813 | -1 |
| YML022W | 35.7188785 | -0.4348635 | 0.66695133 | 0.44240539 | 0.80155419 | 0.88031084 | APT1 | XIII | 228937 | 229500 | 1 |
| YML023C | 453.966283 | 0.7051586 | 0.54816677 | 13.0884605 | 0.00143839 | 0.00885988 | NSE5 | XIII | 226994 | 228664 | -1 |

| | | | | | | | | | | | |
|---|---|---|---|---|---|---|---|---|---|---|---|
| YML024W | 67.3550106 | -0.6290155 | 0.70543999 | 10.6108569 | 0.00496457 | 0.02277723 | RPS17A | XIII | 225889 | 226697 | 1 |
| YML025C | 12.4565845 | -0.3995055 | 0.92079902 | 1.02257149 | 0.59972399 | 0.74659254 | YML6 | XIII | 224406 | 225365 | -1 |
| YML026C | 92.9752755 | -0.8968111 | 0.82496509 | 15.6790543 | 0.00039386 | 0.00302845 | RPS18B | XIII | 222987 | 223828 | -1 |
| YML027W | 190.49332 | 0.60992053 | 0.54986355 | 12.9323472 | 0.00155517 | 0.00937506 | YOX1 | XIII | 221406 | 222563 | 1 |
| YML028W | 153.334924 | 0.03980323 | 0.61655434 | 0.38214797 | 0.82607147 | 0.89568423 | TSA1 | XIII | 220138 | 220728 | 1 |
| YML029W | 275.17951 | -0.0049506 | 0.52743303 | 0.07573916 | 0.96283851 | 0.97258112 | USA1 | XIII | 217362 | 219878 | 1 |
| YML030W | 33.8801773 | 0.62181758 | 0.69610688 | 3.34206767 | 0.18805255 | 0.34376539 | RCF1 | XIII | 216435 | 216914 | 1 |
| YML031W | 47.3429009 | -0.1216764 | 0.62276291 | 2.79751881 | 0.24690308 | 0.41775828 | NDC1 | XIII | 214189 | 216156 | 1 |
| YML032C | 28.8329751 | -0.5422167 | 0.66097449 | 2.52669891 | 0.28270553 | 0.45678444 | RAD52 | XIII | 212515 | 213930 | -1 |
| YML034W | 35.4662262 | -0.3990998 | 0.65298252 | 1.67542986 | 0.43269814 | 0.60137356 | SRC1 | XIII | 209525 | 212155 | 1 |
| YML035C | 184.819314 | -0.2327145 | 0.52743896 | 0.48097787 | 0.78624334 | 0.87021248 | AMD1 | XIII | 206428 | 208860 | -1 |
| YML036W | 737.938283 | 0.56261211 | 0.57717816 | 2.34077371 | 0.3102469 | 0.4830878 | CGI121 | XIII | 205642 | 206293 | 1 |
| YML037C | 74.9023567 | 0.24286447 | 0.7793152 | 4.97092875 | 0.08328687 | 0.19181821 | NA | XIII | 204386 | 205408 | -1 |
| YML038C | 15.3053318 | 0.28798407 | 0.79509121 | 0.13344981 | 0.9354525 | 0.95803272 | YMD8 | XIII | 202775 | 204103 | -1 |
| YML041C | 50.5549655 | 2.2781981 | 0.94091366 | 5.58951753 | 0.06112962 | 0.1520363 | VPS71 | XIII | 194913 | 195755 | -1 |
| YML042W | 205.630072 | -0.2389277 | 0.53109722 | 0.26444191 | 0.87614739 | 0.92712351 | CAT2 | XIII | 192788 | 194800 | 1 |
| YML043C | 12.9528371 | -1.9707317 | 0.90024513 | 12.3273615 | 0.00210449 | 0.01162812 | RRN11 | XIII | 190244 | 191767 | -1 |
| YML046W | 10.1341204 | -1.6378606 | 0.93498923 | 5.44073594 | 0.06585052 | 0.16064998 | PRP39 | XIII | 181474 | 183363 | 1 |
| YML048W | 46.4032934 | -0.4183328 | 0.9962903 | 2.46015174 | 0.2922704 | 0.46643602 | GSF2 | XIII | 178426 | 179637 | 1 |
| YML049C | 48.0802931 | -0.2456707 | 0.59204261 | 0.17716428 | 0.91522793 | 0.94797312 | RSE1 | XIII | 174220 | 178305 | -1 |
| YML050W | 74.598849 | -0.3604474 | 0.54711563 | 0.46786483 | 0.7914153 | 0.87352731 | AIM32 | XIII | 173139 | 174074 | 1 |
| YML051W | 82.4893255 | -1.3529951 | 0.54074839 | 12.4643642 | 0.00196516 | 0.01103615 | GAL80 | XIII | 171594 | 172901 | 1 |
| YML052W | 187.630678 | -0.8441636 | 0.58713709 | 22.6354356 | 1.22E-05 | 0.00015675 | SUR7 | XIII | 170402 | 171310 | 1 |
| YML053C | 19.1901839 | -0.6692093 | 0.75068222 | 0.82081254 | 0.66338068 | 0.79064927 | NA | XIII | 169116 | 169754 | -1 |
| YML054C | 382.800786 | -0.9427681 | 0.50328607 | 20.364535 | 3.78E-05 | 0.00040962 | CYB2 | XIII | 165533 | 167308 | -1 |
| YML054C-A | 191.129122 | -0.5618499 | 0.55179668 | 1.04016847 | 0.59447047 | 0.74174387 | NA | XIII | 167623 | 167781 | -1 |
| YML055W | 30.5380914 | -0.0307607 | 0.65609244 | 0.0286761 | 0.98576425 | 0.98832468 | SPC2 | XIII | 164790 | 165326 | 1 |
| YML056C | 64.6264738 | 0.0193245 | 0.63922628 | 1.57827264 | 0.45423694 | 0.62039568 | IMD4 | XIII | 162194 | 164176 | -1 |
| YML057W | 149.624716 | 1.58070087 | 0.54967828 | 37.0185659 | 9.15E-09 | 2.76E-07 | CMP2 | XIII | 160180 | 161994 | 1 |
| YML058W | 186.328436 | 2.09820403 | 0.51854637 | 22.1348348 | 1.56E-05 | 0.00019234 | SML1 | XIII | 159383 | 159697 | 1 |
| YML058W-A | 25.9288503 | 0.85955122 | 0.70794703 | 2.28398552 | 0.31918233 | 0.49189931 | HUG1 | XIII | 158760 | 158966 | 1 |
| YML059C | 41.4330295 | 0.19751276 | 0.66220722 | 3.59140138 | 0.16601109 | 0.31525228 | NTE1 | XIII | 153219 | 158258 | -1 |
| YML060W | 7.15515563 | -0.815896 | 1.14090179 | 0.6849936 | 0.70999539 | 0.82438173 | OGG1 | XIII | 151871 | 153001 | 1 |

| | | | | | | | | | | | |
|---|---|---|---|---|---|---|---|---|---|---|---|
| YML061C | 31.1195425 | 0.48354861 | 0.64506952 | 3.46497779 | 0.17684372 | 0.32944824 | PIF1 | XIII | 148953 | 151532 | -1 |
| YML062C | 72.4756982 | -0.1111023 | 0.59568558 | 5.69663649 | 0.05794168 | 0.14626659 | MFT1 | XIII | 147505 | 148683 | -1 |
| YML063W | 158.93677 | -0.9519694 | 0.7164137 | 15.5747192 | 0.00041495 | 0.00316124 | RPS1B | XIII | 146482 | 147249 | 1 |
| YML064C | 17.0269822 | -0.1211971 | 0.80033295 | 5.08684574 | 0.07859691 | 0.18409228 | TEM1 | XIII | 145139 | 145876 | -1 |
| YML065W | 66.7441541 | 0.58647321 | 0.78804139 | 5.26666665 | 0.0718386 | 0.17209164 | ORC1 | XIII | 142210 | 144954 | 1 |
| YML066C | 994.627674 | 0.71621135 | 0.48291141 | 4.23724677 | 0.12019698 | 0.24944104 | SMA2 | XIII | 140424 | 141533 | -1 |
| YML067C | 399.78616 | 0.28011109 | 0.49306045 | 0.6427121 | 0.72516501 | 0.83456178 | ERV41 | XIII | 139063 | 140214 | -1 |
| YML068W | 188.4442 | 0.55702024 | 0.59910609 | 6.76661588 | 0.03393501 | 0.09824187 | ITT1 | XIII | 137550 | 138944 | 1 |
| YML069W | 44.6293787 | 0.28831178 | 0.59765015 | 0.99787162 | 0.60717647 | 0.75134682 | POB3 | XIII | 135500 | 137158 | 1 |
| YML070W | 55.4336425 | 0.03223246 | 0.65851243 | 1.98460515 | 0.37072209 | 0.54410163 | DAK1 | XIII | 133475 | 135229 | 1 |
| YML071C | 267.011606 | 2.47663326 | 0.71317094 | 11.1894268 | 0.00371746 | 0.01814182 | COG8 | XIII | 129749 | 131572 | -1 |
| YML072C | 262.166211 | 0.13097272 | 0.49867102 | 1.41290492 | 0.49339142 | 0.65446422 | TCB3 | XIII | 124730 | 129367 | -1 |
| YML073C | 186.67094 | -0.768851 | 0.61257892 | 22.5512317 | 1.27E-05 | 0.00016205 | RPL6A | XIII | 123227 | 124172 | -1 |
| YML074C | 680.951872 | 0.97813293 | 0.64402544 | 5.44109224 | 0.06583879 | 0.16064998 | FPR3 | XIII | 120089 | 121324 | -1 |
| YML075C | 75.2321225 | 0.2739121 | 0.5648475 | 0.24894529 | 0.88296241 | 0.93100373 | HMG1 | XIII | 115734 | 118898 | -1 |
| YML076C | 91.8079686 | 0.20459205 | 0.56452376 | 18.0516511 | 0.00012026 | 0.00109485 | WAR1 | XIII | 112513 | 115347 | -1 |
| YML077W | 48.2922499 | 0.58439581 | 0.59365793 | 1.66060621 | 0.43591714 | 0.60483111 | BET5 | XIII | 111865 | 112344 | 1 |
| YML078W | 106.227883 | 0.05429134 | 0.5634649 | 0.35122854 | 0.83894153 | 0.90540008 | CPR3 | XIII | 111002 | 111550 | 1 |
| YML079W | 24.0588246 | 0.17596697 | 0.73532814 | 0.13638016 | 0.93408291 | 0.95735444 | NA | XIII | 110247 | 110852 | 1 |
| YML080W | 16.1795302 | -0.6621442 | 0.95540052 | 1.0008501 | 0.60627291 | 0.75088539 | DUS1 | XIII | 108806 | 110077 | 1 |
| YML081C-A | 51.1614985 | -0.8753844 | 0.61530295 | 2.3316168 | 0.31167061 | 0.48405923 | ATP18 | XIII | 103983 | 104162 | -1 |
| YML081W | 44.2925372 | -0.8289917 | 0.69984981 | 4.10416492 | 0.1284671 | 0.26218699 | TDA9 | XIII | 104777 | 108532 | 1 |
| YML082W | 28.1193849 | -0.4567338 | 0.67198985 | 4.27259839 | 0.11809107 | 0.24673616 | NA | XIII | 101862 | 103811 | 1 |
| YML083C | 278.671987 | -0.4374675 | 0.48291959 | 0.82459892 | 0.66212597 | 0.78964147 | NA | XIII | 99794 | 101050 | -1 |
| YML085C | 37.3557016 | -0.9944395 | 0.72949039 | 2.59957376 | 0.27258988 | 0.44584616 | TUB1 | XIII | 97941 | 99400 | -1 |
| YML086C | 70.2729105 | 1.41874457 | 0.62743282 | 5.18786936 | 0.07472544 | 0.17681255 | ALO1 | XIII | 95791 | 97371 | -1 |
| YML087C | 1892.93277 | 2.29455322 | 0.47634501 | 35.9987802 | 1.52E-08 | 4.35E-07 | AIM33 | XIII | 94431 | 95369 | -1 |
| YML088W | 205.194481 | 2.48373741 | 0.51964116 | 26.4499081 | 1.80E-06 | 2.99E-05 | UFO1 | XIII | 92235 | 94241 | 1 |
| YML091C | 41.1568071 | -0.7194168 | 0.69985876 | 3.58330463 | 0.16668453 | 0.31570278 | RPM2 | XIII | 87123 | 90731 | -1 |
| YML092C | 36.4034058 | -0.7676798 | 0.83809901 | 0.8802231 | 0.64396458 | 0.77686719 | PRE8 | XIII | 85987 | 86739 | -1 |
| YML093W | 54.8671805 | -0.1822101 | 0.6567604 | 20.4773375 | 3.58E-05 | 0.0003914 | UTP14 | XIII | 83090 | 85789 | 1 |
| YML094W | 13.0292275 | -0.5946662 | 0.85292852 | 4.88886156 | 0.08677551 | 0.19851056 | GIM5 | XIII | 82275 | 82849 | 1 |
| YML095C | 185.069877 | -0.6494594 | 0.50367507 | 24.2487739 | 5.43E-06 | 7.66E-05 | RAD10 | XIII | 81481 | 82113 | -1 |

| | | | | | | | | | | | |
|---|---|---|---|---|---|---|---|---|---|---|---|
| YML096W | 869.395302 | 0.50775296 | 0.46138448 | 1.576424 | 0.454657 | 0.62066785 | NA | XIII | 79909 | 81486 | 1 |
| YML097C | 31.4509663 | -0.0224645 | 0.67005125 | 0.70572745 | 0.70267294 | 0.81860691 | VPS9 | XIII | 78335 | 79690 | -1 |
| YML098W | 35.8373151 | -0.0980784 | 0.65905781 | 0.23908611 | 0.8873258 | 0.93215081 | TAF13 | XIII | 77267 | 77770 | 1 |
| YML099C | 15.6251821 | 0.26908141 | 0.94805306 | 0.25261807 | 0.88134244 | 0.9300006 | ARG81 | XIII | 74398 | 77040 | -1 |
| YML100W | 126.786729 | 0.16819819 | 0.51844956 | 3.42928132 | 0.1800284 | 0.33462743 | TSL1 | XIII | 70624 | 73920 | 1 |
| YML100W-A | 129.566306 | -0.2020684 | 0.71444886 | 1.932583 | 0.38049148 | 0.55436479 | NA | XIII | 70138 | 70311 | 1 |
| YML101C | 158.046585 | 1.51063364 | 0.74745651 | 5.63283821 | 0.05981977 | 0.14961402 | CUE4 | XIII | 69735 | 70088 | -1 |
| YML102W | 433.992011 | 1.1610751 | 0.68535791 | 4.02216999 | 0.13384338 | 0.26992447 | CAC2 | XIII | 68294 | 69700 | 1 |
| YML103C | 11.3770154 | 0.22744756 | 1.10676907 | 0.5442316 | 0.76176604 | 0.85526961 | NUP188 | XIII | 62582 | 67549 | -1 |
| YML104C | 89.1669602 | 0.0176457 | 0.51926633 | 2.91446787 | 0.23287954 | 0.40027153 | MDM1 | XIII | 58939 | 62322 | -1 |
| YML105C | 66.6157127 | 0.81265621 | 0.72221549 | 13.6496431 | 0.00108647 | 0.00708137 | SEC65 | XIII | 57866 | 58687 | -1 |
| YML106W | 34.6021929 | -0.9392041 | 0.64074221 | 2.4959741 | 0.2870821 | 0.46133926 | URA5 | XIII | 56773 | 57453 | 1 |
| YML107C | 553.507954 | -0.3674036 | 0.57787423 | 5.19781115 | 0.07435491 | 0.17629168 | PML39 | XIII | 55265 | 56269 | -1 |
| YML108W | 182.735696 | 0.27543058 | 0.67015773 | 4.11450406 | 0.12780469 | 0.26147074 | NA | XIII | 54793 | 55110 | 1 |
| YML109W | 46.2871023 | 0.47169061 | 0.65307313 | 8.52022721 | 0.0141207 | 0.05110095 | ZDS2 | XIII | 51640 | 54468 | 1 |
| YML110C | 11.1069397 | -0.2778898 | 0.90076328 | 0.10004106 | 0.9512099 | 0.96614462 | COQ5 | XIII | 50031 | 50954 | -1 |
| YML111W | 87.8360469 | 0.69676921 | 0.60322689 | 1.3399216 | 0.51172864 | 0.66867723 | BUL2 | XIII | 46942 | 49704 | 1 |
| YML112W | 40.9382117 | 1.38399494 | 0.73599569 | 4.55162399 | 0.10271347 | 0.2231561 | CTK3 | XIII | 45063 | 45953 | 1 |
| YML113W | 46.447141 | 0.04522952 | 0.61598488 | 0.64075808 | 0.72587385 | 0.83456178 | DAT1 | XIII | 44045 | 44791 | 1 |
| YML114C | 48.9693991 | 0.58264159 | 0.63657255 | 2.92959932 | 0.23112429 | 0.39792139 | TAF8 | XIII | 42043 | 43575 | -1 |
| YML115C | 47.4230925 | -0.1246105 | 0.62875311 | 0.54926024 | 0.75985313 | 0.85451323 | VAN1 | XIII | 40187 | 41794 | -1 |
| YML116W | 244.627165 | 1.97428213 | 0.60454048 | 48.5814827 | 2.82E-11 | 1.49E-09 | ATR1 | XIII | 38196 | 39824 | 1 |
| YML117W | 169.692587 | 0.67842418 | 0.59945387 | 1.3118757 | 0.51895513 | 0.67522582 | NAB6 | XIII | 34243 | 37647 | 1 |
| YML118W | 183.049701 | 2.4580194 | 0.53103461 | 35.15155 | 2.33E-08 | 6.39E-07 | NGL3 | XIII | 32334 | 33851 | 1 |
| YML119W | 245.260817 | 2.20926212 | 0.5735312 | 29.9675796 | 3.11E-07 | 6.43E-06 | NA | XIII | 30611 | 31684 | 1 |
| YML120C | 24.57111 | 1.26781995 | 0.78144571 | 2.60001639 | 0.27252956 | 0.44584616 | NDI1 | XIII | 28266 | 29807 | -1 |
| YML121W | 13.6359108 | -0.5221142 | 0.82341736 | 3.33052931 | 0.18914059 | 0.34524718 | GTR1 | XIII | 26930 | 27862 | 1 |
| YML122C | 254.232888 | -0.0477746 | 0.50655233 | 2.23148788 | 0.32767142 | 0.50039829 | NA | XIII | 26038 | 26418 | -1 |
| YML123C | 1912.46145 | -0.3250802 | 0.47992791 | 0.5934124 | 0.74326235 | 0.8433253 | PHO84 | XIII | 24037 | 25800 | -1 |
| YML124C | 73.3289378 | 0.63308231 | 0.59365387 | 3.04936011 | 0.21769069 | 0.38155469 | TUB3 | XIII | 22048 | 23683 | -1 |
| YML125C | 526.521536 | -0.8187511 | 0.53882123 | 6.01353847 | 0.04945119 | 0.12955763 | PGA3 | XIII | 20761 | 21699 | -1 |
| YML126C | 113.072242 | 0.14216011 | 0.56029901 | 3.79773066 | 0.14973843 | 0.29250523 | ERG13 | XIII | 19060 | 20535 | -1 |
| YML127W | 12.3468812 | 0.02086511 | 0.91659834 | 0.06790372 | 0.96661804 | 0.97486964 | RSC9 | XIII | 17064 | 18809 | 1 |

| | | | | | | | | | | | |
|---|---|---|---|---|---|---|---|---|---|---|---|
| YML128C | 14.8391883 | 0.24909868 | 0.96141929 | 4.43364252 | 0.1089549 | 0.23299267 | MSC1 | XIII | 15135 | 16676 | -1 |
| YML129C | 6.88575157 | -0.5988021 | 1.20627935 | 0.66017479 | 0.71886091 | 0.82938751 | COX14 | XIII | 14541 | 14753 | -1 |
| YML130C | 88.1022797 | 1.89878902 | 0.61136636 | 10.4606207 | 0.00535186 | 0.02418992 | ERO1 | XIII | 11483 | 13174 | -1 |
| YML131W | 14.9524082 | 0.15786516 | 0.79636019 | 0.05551388 | 0.97262475 | 0.97905029 | NA | XIII | 10198 | 11295 | 1 |
| YML133C | 11.5094429 | -1.8283557 | 0.98856615 | 3.58466689 | 0.16657103 | 0.31564276 | NA | XIII | 461 | 4684 | -1 |
| YMR001C | 64.4256951 | -0.4612901 | 0.6326415 | 13.1517357 | 0.0013936 | 0.00862986 | CDC5 | XIII | 269019 | 271136 | -1 |
| YMR002W | 63.783678 | 0.15744458 | 0.58307806 | 0.18597073 | 0.91120684 | 0.94542349 | MIX17 | XIII | 272193 | 272663 | 1 |
| YMR003W | 46.5387994 | -0.5199356 | 0.62098019 | 2.4822426 | 0.28905991 | 0.46297563 | AIM34 | XIII | 273118 | 273714 | 1 |
| YMR004W | 48.8037284 | 0.06827196 | 0.60625078 | 5.02547894 | 0.08104591 | 0.1881539 | MVP1 | XIII | 274017 | 275552 | 1 |
| YMR005W | 236.768225 | 0.08060839 | 0.48661218 | 17.7652578 | 0.00013878 | 0.0012449 | TAF4 | XIII | 276045 | 277211 | 1 |
| YMR006C | 1733.72974 | 1.31268478 | 0.4872202 | 18.8439444 | 8.09E-05 | 0.00078224 | PLB2 | XIII | 277561 | 279681 | -1 |
| YMR007W | 2468.55699 | 2.30252282 | 0.55874505 | 18.8042747 | 8.25E-05 | 0.00079658 | NA | XIII | 279960 | 280340 | 1 |
| YMR008C | 112.197859 | 1.87239988 | 0.78307011 | 5.61451275 | 0.0603704 | 0.15063165 | PLB1 | XIII | 280590 | 282584 | -1 |
| YMR008C-A | 112.497476 | 1.46328708 | 0.5381687 | 10.1311087 | 0.00631041 | 0.02756307 | NA | NA | NA | NA | NA |
| YMR009W | 15.3253306 | 0.68399486 | 0.83906008 | 0.74594413 | 0.68868447 | 0.81049746 | ADI1 | XIII | 284102 | 284641 | 1 |
| YMR010W | 29.6458472 | -0.54911 | 0.69977745 | 1.05350261 | 0.59052028 | 0.73767258 | ANY1 | XIII | 285100 | 286317 | 1 |
| YMR011W | 171.711574 | -0.7268731 | 0.55479811 | 24.4765424 | 4.84E-06 | 6.90E-05 | HXT2 | XIII | 288079 | 289704 | 1 |
| YMR012W | 75.4907809 | -0.4850464 | 0.54049318 | 9.99047116 | 0.00677013 | 0.02923119 | CLU1 | XIII | 291134 | 294967 | 1 |
| YMR014W | 107.202825 | -0.1458057 | 0.62180552 | 6.62125991 | 0.03649318 | 0.10388176 | BUD22 | XIII | 298868 | 300427 | 1 |
| YMR015C | 25.1016391 | -0.20017 | 0.78506097 | 6.94376112 | 0.03105857 | 0.09182162 | ERG5 | XIII | 300869 | 302485 | -1 |
| YMR016C | 204.248051 | -0.4012668 | 0.50991587 | 3.0816558 | 0.21420369 | 0.37813088 | SOK2 | XIII | 303236 | 305593 | -1 |
| YMR017W | 36.7532861 | -1.4797481 | 0.63852327 | 9.14753313 | 0.01031902 | 0.04032978 | SPO20 | XIII | 307489 | 308682 | 1 |
| YMR018W | 59.5061239 | 2.16850314 | 0.9213148 | 6.00634065 | 0.04962948 | 0.12996593 | PEX9 | XIII | 310208 | 311752 | 1 |
| YMR019W | 246.029845 | 2.52169974 | 0.56510967 | 45.2526342 | 1.49E-10 | 6.80E-09 | STB4 | XIII | 312156 | 315005 | 1 |
| YMR020W | 268.222547 | 0.16126499 | 0.6478732 | 0.13938119 | 0.93268235 | 0.95731799 | FMS1 | XIII | 315377 | 316903 | 1 |
| YMR021C | 94.7806107 | -0.014692 | 0.55721549 | 2.42622122 | 0.29727115 | 0.47117436 | MAC1 | XIII | 317165 | 318418 | -1 |
| YMR022W | 11.0636959 | 0.89986519 | 0.98779072 | 1.36769023 | 0.50467273 | 0.66289817 | UBC7 | XIII | 318680 | 319177 | 1 |
| YMR023C | 110.571512 | 0.07684803 | 0.59593289 | 5.19168299 | 0.07458309 | 0.17654787 | MSS1 | XIII | 319437 | 321017 | -1 |
| YMR024W | 6.49353361 | -0.3813593 | 1.14237065 | 0.33730925 | 0.84480062 | 0.90911019 | MRPL3 | XIII | 321875 | 323047 | 1 |
| YMR025W | 188.326141 | 3.1048789 | 0.84350761 | 12.2737722 | 0.00216164 | 0.01184098 | CSI1 | XIII | 323300 | 324187 | 1 |
| YMR026C | 150.884966 | 0.09637025 | 0.51128361 | 1.88534121 | 0.38958601 | 0.56101577 | PEX12 | XIII | 324236 | 325435 | -1 |
| YMR027W | 26.2509223 | 0.7890558 | 0.96644455 | 0.70644384 | 0.70242129 | 0.81847842 | NA | XIII | 325877 | 327289 | 1 |
| YMR028W | 212.914341 | 2.63054036 | 0.93402169 | 8.69412505 | 0.01294478 | 0.04783043 | TAP42 | XIII | 327482 | 328582 | 1 |

| | | | | | | | | | | | |
|---|---|---|---|---|---|---|---|---|---|---|---|
| YMR029C | 123.952837 | 1.64302528 | 0.76075014 | 12.9990031 | 0.00150419 | 0.00913878 | FAR8 | XIII | 328660 | 330231 | -1 |
| YMR030W | 26.0597765 | 0.21907157 | 0.78363634 | 0.17369188 | 0.91681833 | 0.94860224 | RSF1 | XIII | 330793 | 331923 | 1 |
| YMR030W-A | 11.7322247 | 0.773577 | 1.01251329 | 2.98150187 | 0.22520348 | 0.39051457 | NA | XIII | 337313 | 337603 | 1 |
| YMR031C | 41.3945074 | 0.062789 | 0.66698178 | 0.4839551 | 0.7850738 | 0.86996695 | EIS1 | XIII | 332212 | 334743 | -1 |
| YMR032W | 15.4277085 | -0.8308562 | 0.97396405 | 1.27363345 | 0.52897362 | 0.68319367 | HOF1 | XIII | 335298 | 337307 | 1 |
| YMR033W | 103.899303 | 0.70987598 | 0.68250602 | 40.181675 | 1.88E-09 | 6.60E-08 | ARP9 | XIII | 337788 | 339277 | 1 |
| YMR034C | 115.266115 | -0.2958578 | 0.62533417 | 1.37049473 | 0.50396555 | 0.6624201 | RCH1 | XIII | 339418 | 340722 | -1 |
| YMR035W | 53.2344765 | 0.27705581 | 0.61610001 | 0.34738216 | 0.84055652 | 0.90645906 | IMP2 | XIII | 341142 | 341675 | 1 |
| YMR036C | 23.8936685 | 2.24973326 | 0.91210842 | 7.9667577 | 0.01862261 | 0.0626889 | MIH1 | XIII | 341856 | 343520 | -1 |
| YMR037C | 91.9025761 | 0.64016799 | 0.6048049 | 21.4440308 | 2.21E-05 | 0.00025693 | MSN2 | XIII | 344403 | 346517 | -1 |
| YMR038C | 537.351494 | -0.1723789 | 0.48585255 | 4.71678461 | 0.09457214 | 0.21003333 | CCS1 | XIII | 347511 | 348260 | -1 |
| YMR039C | 159.641959 | 0.49512675 | 0.50322407 | 3.84924875 | 0.14593056 | 0.28719849 | SUB1 | XIII | 348644 | 349522 | -1 |
| YMR040W | 139.229145 | 4.60072518 | 1.05085159 | 15.5023201 | 0.00043024 | 0.0032521 | YET2 | XIII | 350381 | 350863 | 1 |
| YMR041C | 132.286413 | 1.08350996 | 0.52989125 | 11.1546135 | 0.00378274 | 0.01836234 | ARA2 | XIII | 350966 | 351973 | -1 |
| YMR042W | 177.280757 | -0.5816983 | 0.50920945 | 2.77170394 | 0.25011062 | 0.42121597 | ARG80 | XIII | 352603 | 353136 | 1 |
| YMR043W | 234.040152 | 1.95020248 | 0.74677033 | 7.15218968 | 0.02798477 | 0.08430376 | MCM1 | XIII | 353871 | 354731 | 1 |
| YMR044W | 77.8223333 | 0.15212515 | 0.55380916 | 4.90355174 | 0.08614048 | 0.19736975 | IOC4 | XIII | 355384 | 356811 | 1 |
| YMR045C | 7.14014222 | -1.1771324 | 1.23488682 | 1.62796025 | 0.44309099 | 0.61162622 | NA | XIII | 357359 | 362627 | -1 |
| YMR047C | 48.5533361 | 0.11192623 | 0.61614224 | 3.07094727 | 0.21535367 | 0.37922681 | NUP116 | XIII | 363364 | 366705 | -1 |
| YMR048W | 47.6138775 | 0.36427955 | 0.78405717 | 17.3317136 | 0.00017237 | 0.00149406 | CSM3 | XIII | 366981 | 367934 | 1 |
| YMR049C | 47.4267326 | -1.1640769 | 0.62869721 | 17.4012326 | 0.00016648 | 0.00145171 | ERB1 | XIII | 368094 | 370517 | -1 |
| YMR052W | 63.2738524 | 2.62495366 | 0.65143067 | 16.3725746 | 0.00027845 | 0.00229332 | FAR3 | XIII | 379586 | 380200 | 1 |
| YMR053C | 395.186856 | 0.16664835 | 0.57404524 | 0.34738535 | 0.84055518 | 0.90645906 | STB2 | XIII | 380346 | 382898 | -1 |
| YMR054W | 323.372774 | -1.2150467 | 0.55142204 | 12.9031458 | 0.00157804 | 0.00949776 | STV1 | XIII | 383303 | 385975 | 1 |
| YMR055C | 90.8357744 | 0.24679575 | 0.5489972 | 0.55486397 | 0.7577271 | 0.85371471 | BUB2 | XIII | 386101 | 387021 | -1 |
| YMR056C | 62.5697369 | -1.6362099 | 0.70241947 | 13.2366635 | 0.00133566 | 0.00831554 | AAC1 | XIII | 387315 | 388244 | -1 |
| YMR058W | 427.046514 | 0.11585168 | 0.5827585 | 25.1894024 | 3.39E-06 | 5.12E-05 | FET3 | XIII | 388822 | 390732 | 1 |
| YMR060C | 183.718682 | -0.1050231 | 0.61921381 | 7.25902376 | 0.02652913 | 0.08104993 | SAM37 | XIII | 391532 | 392515 | -1 |
| YMR061W | 11.7091193 | -0.2649762 | 0.92678528 | 0.8017716 | 0.66972654 | 0.79510827 | RNA14 | XIII | 392755 | 394788 | 1 |
| YMR062C | 66.3844012 | -0.8657691 | 0.63882451 | 1.88965463 | 0.3887467 | 0.56060856 | ARG7 | XIII | 395054 | 396379 | -1 |
| YMR063W | 108.81423 | -0.153963 | 0.53187918 | 0.5111789 | 0.77445986 | 0.86282905 | RIM9 | XIII | 397077 | 397796 | 1 |
| YMR064W | 19.3955902 | -0.7389524 | 0.79036028 | 1.07913714 | 0.58299972 | 0.7314341 | AEP1 | XIII | 397903 | 399459 | 1 |
| YMR065W | 104.896978 | -0.4577996 | 0.54050552 | 7.55224637 | 0.02291134 | 0.07329098 | KAR5 | XIII | 399702 | 401216 | 1 |

| | | | | | | | | | | | |
|---|---|---|---|---|---|---|---|---|---|---|---|
| YMR066W | 52.0377484 | 1.86501844 | 0.93919014 | 4.33566817 | 0.11442518 | 0.24134059 | SOV1 | XIII | 401541 | 404237 | 1 |
| YMR067C | 21.7461241 | -0.0853778 | 0.74340581 | 0.6046635 | 0.73909283 | 0.84040603 | UBX4 | XIII | 404323 | 405573 | -1 |
| YMR068W | 17.0363636 | 0.01237999 | 0.80742418 | 1.64350459 | 0.43966056 | 0.60823052 | AVO2 | XIII | 406304 | 407584 | 1 |
| YMR069W | 25.0370625 | -0.0692079 | 0.82060168 | 0.94622382 | 0.62306034 | 0.76286238 | NAT4 | XIII | 407709 | 408566 | 1 |
| YMR070W | 635.192795 | 4.30711727 | 0.55940674 | 84.1103701 | 5.44E-19 | 1.21E-16 | MOT3 | XIII | 409154 | 410626 | 1 |
| YMR071C | 91.2768935 | 1.41594099 | 0.76280292 | 4.60431893 | 0.10004257 | 0.21919967 | TVP18 | XIII | 410762 | 411265 | -1 |
| YMR072W | 170.847745 | -0.1113183 | 0.50905387 | 2.14006927 | 0.34299664 | 0.51663645 | ABF2 | XIII | 411569 | 412120 | 1 |
| YMR073C | 24.9586269 | 0.92913118 | 0.78000652 | 3.05743229 | 0.21681385 | 0.380987 | IRC21 | XIII | 412268 | 412873 | -1 |
| YMR074C | 319.915176 | -0.6335602 | 0.52584601 | 1.50371684 | 0.47148951 | 0.6333931 | SDD2 | XIII | 413036 | 413473 | -1 |
| YMR075C-A | 1154.45491 | 0.53645139 | 0.61463397 | 1.17137007 | 0.55672436 | 0.70829137 | NA | XIII | 415688 | 416053 | -1 |
| YMR075W | 95.8032476 | -0.4668113 | 0.58756501 | 13.5332329 | 0.00115158 | 0.0073921 | RCO1 | XIII | 413982 | 416036 | 1 |
| YMR076C | 104.874823 | 0.01278877 | 0.51913991 | 3.05146422 | 0.21746179 | 0.38155469 | PDS5 | XIII | 416196 | 420029 | -1 |
| YMR077C | 18.688232 | -0.4549351 | 0.87020603 | 2.85500919 | 0.23990684 | 0.40806716 | VPS20 | XIII | 421484 | 422149 | -1 |
| YMR078C | 322.054751 | -0.7798436 | 0.49838433 | 8.87355536 | 0.01183401 | 0.04466683 | CTF18 | XIII | 422503 | 424728 | -1 |
| YMR079W | 24.2044627 | -0.3406593 | 0.84487538 | 0.44727155 | 0.79960632 | 0.87884032 | SEC14 | XIII | 424989 | 426059 | 1 |
| YMR080C | 20.4186915 | -0.7313743 | 0.77875744 | 0.94533827 | 0.62333628 | 0.76302686 | NAM7 | XIII | 426712 | 429627 | -1 |
| YMR081C | 455.523337 | -0.1964376 | 0.49388974 | 1.29572048 | 0.52316402 | 0.67871828 | ISF1 | XIII | 430079 | 431095 | -1 |
| YMR083W | 88.866678 | -0.4333067 | 1.69020309 | 0.11776601 | 0.94281707 | 0.96176085 | ADH3 | XIII | 434788 | 435915 | 1 |
| YMR085W | 10.9076129 | 0.42747351 | 1.12387513 | 1.00310118 | 0.60559091 | 0.75082898 | NA | XIII | 437491 | 438789 | 1 |
| YMR086W | 442.428445 | -0.8971309 | 0.58745318 | 5.92310874 | 0.05173843 | 0.13421395 | SEG1 | XIII | 439208 | 442090 | 1 |
| YMR087W | 417.279923 | 0.86876314 | 0.59874656 | 2.16077159 | 0.33946454 | 0.51259098 | NA | XIII | 442527 | 443381 | 1 |
| YMR088C | 633.536411 | 1.69578234 | 0.61847587 | 24.5185351 | 4.74E-06 | 6.83E-05 | VBA1 | XIII | 443414 | 445102 | -1 |
| YMR089C | 33.0238837 | 0.35859615 | 0.62920492 | 0.66270647 | 0.71795152 | 0.82923185 | YTA12 | XIII | 445609 | 448086 | -1 |
| YMR090W | 27.7869719 | 0.25564732 | 0.71488865 | 0.21321937 | 0.89887645 | 0.93754851 | NA | XIII | 449245 | 449928 | 1 |
| YMR091C | 75.6216463 | 0.59652262 | 0.63710961 | 0.92617228 | 0.62933842 | 0.76680754 | NPL6 | XIII | 450058 | 451365 | -1 |
| YMR092C | 23.8554022 | 0.76400057 | 0.87647027 | 0.81631602 | 0.66487381 | 0.7916141 | AIP1 | XIII | 451632 | 453479 | -1 |
| YMR093W | 9.83668325 | -0.1662379 | 1.22492648 | 3.53575451 | 0.17069495 | 0.32088433 | UTP15 | XIII | 454015 | 455556 | 1 |
| YMR094W | 600.557516 | 3.20310527 | 0.58067789 | 32.2747037 | 9.81E-08 | 2.40E-06 | CTF13 | XIII | 455825 | 457261 | 1 |
| YMR095C | 600.469589 | -1.1031766 | 0.48115426 | 7.46832016 | 0.02389323 | 0.07547289 | SNO1 | XIII | 457285 | 457959 | -1 |
| YMR096W | 44.5718488 | 2.94548181 | 0.73024222 | 17.4123505 | 0.00016556 | 0.00144584 | SNZ1 | XIII | 458408 | 459301 | 1 |
| YMR097C | 860.395237 | 3.87552864 | 0.7479702 | 22.4058367 | 1.36E-05 | 0.00017236 | MTG1 | XIII | 459424 | 460527 | -1 |
| YMR098C | 88.2131072 | -0.4889766 | 0.65155083 | 2.3461783 | 0.30940965 | 0.48239987 | ATP25 | XIII | 460771 | 462609 | -1 |
| YMR099C | 21.6920342 | 0.07204776 | 0.82022498 | 0.10920212 | 0.94686281 | 0.96418144 | NA | XIII | 463934 | 464827 | -1 |

| | | | | | | | | | | | |
|---|---|---|---|---|---|---|---|---|---|---|---|
| YMR100W | 38.3055422 | -1.3001991 | 0.78211342 | 6.70572308 | 0.0349841 | 0.10042536 | MUB1 | XIII | 466300 | 468162 | 1 |
| YMR101C | 11.2113408 | -1.1967656 | 0.96400402 | 3.42536117 | 0.18038162 | 0.33506648 | SRT1 | XIII | 468445 | 469476 | -1 |
| YMR102C | 63.8429626 | 1.71855806 | 0.62683342 | 23.9829022 | 6.20E-06 | 8.57E-05 | NA | XIII | 469848 | 472352 | -1 |
| YMR103C | 12.1600389 | 1.45196031 | 1.29831036 | 5.02167994 | 0.08120001 | 0.18843608 | NA | XIII | 472902 | 473264 | -1 |
| YMR104C | 83.8299788 | 1.66587401 | 0.66696265 | 7.06702577 | 0.02920215 | 0.08706512 | YPK2 | XIII | 473420 | 475453 | -1 |
| YMR105C | 23.3300621 | 1.05525842 | 0.81189028 | 10.3719875 | 0.00559437 | 0.02503202 | PGM2 | XIII | 475897 | 477606 | -1 |
| YMR105W-A | 276.814889 | -0.0207199 | 0.53343713 | 0.02646183 | 0.98685623 | 0.98907696 | NA | XIII | 478064 | 478258 | 1 |
| YMR106C | 21.2639694 | 1.04387228 | 0.79974583 | 2.39093065 | 0.30256313 | 0.47513982 | YKU80 | XIII | 478301 | 480190 | -1 |
| YMR108W | 61.6187542 | 0.2468038 | 0.58452931 | 6.39276525 | 0.04090992 | 0.11317174 | ILV2 | XIII | 484084 | 486147 | 1 |
| YMR109W | 36.0271123 | 1.74633723 | 0.83656225 | 5.14017788 | 0.07652874 | 0.18012252 | MYO5 | XIII | 486587 | 490246 | 1 |
| YMR110C | 44.2777111 | -0.5698198 | 0.63871536 | 0.87192461 | 0.6466421 | 0.77855225 | HFD1 | XIII | 490394 | 491992 | -1 |
| YMR111C | 23.6371965 | -0.4081437 | 0.71974593 | 0.34438912 | 0.84181537 | 0.90731776 | NA | XIII | 492404 | 493792 | -1 |
| YMR112C | 9.70930219 | 0.18996627 | 0.96002073 | 0.05840111 | 0.97122166 | 0.9781481 | NA | XIII | 494100 | 494495 | -1 |
| YMR113W | 129.627505 | 1.94389602 | 0.85063521 | 4.88649529 | 0.08687824 | 0.19866707 | FOL3 | XIII | 494999 | 496282 | 1 |
| YMR114C | 65.3201365 | 2.49764535 | 0.94398918 | 6.6577345 | 0.03583367 | 0.10225577 | NA | XIII | 496343 | 497449 | -1 |
| YMR115W | 13.0570643 | -0.1650992 | 0.86988472 | 3.00564288 | 0.2225015 | 0.38746266 | MGR3 | XIII | 497704 | 499209 | 1 |
| YMR116C | 56.3146113 | -0.7322728 | 0.67772011 | 15.4107974 | 0.00045039 | 0.00337792 | ASC1 | XIII | 499456 | 500688 | -1 |
| YMR117C | 33.5603311 | -1.4896692 | 0.63757875 | 12.5344517 | 0.00189749 | 0.01073943 | SPC24 | XIII | 501250 | 501891 | -1 |
| YMR118C | 557.329708 | -1.1135086 | 0.50389371 | 20.1290894 | 4.26E-05 | 0.0004497 | SHH3 | XIII | 502144 | 502734 | -1 |
| YMR119W | 20.0257178 | -0.1337689 | 0.84657682 | 0.06615971 | 0.9674613 | 0.97486964 | ASI1 | XIII | 505333 | 507207 | 1 |
| YMR120C | 556.624801 | 0.78994765 | 0.53551525 | 2.82769055 | 0.24320629 | 0.41283037 | ADE17 | XIII | 507502 | 509280 | -1 |
| YMR121C | 33.8845462 | 1.32940154 | 0.72281304 | 3.36556832 | 0.1858558 | 0.34169876 | RPL15B | XIII | 509734 | 510348 | -1 |
| YMR122C | 71.4408725 | -2.3000174 | 0.65872423 | 15.059504 | 0.00053687 | 0.00388561 | NA | XIII | 510701 | 511075 | -1 |
| YMR122W-A | 88.3203857 | 2.01045013 | 0.56454424 | 12.2081162 | 0.00223378 | 0.01216709 | NA | XIII | 511315 | 511569 | 1 |
| YMR123W | 94.8121014 | -0.0916902 | 0.51772901 | 0.05093939 | 0.97485192 | 0.98061025 | PKR1 | XIII | 513593 | 513961 | 1 |
| YMR124W | 92.996837 | 0.84941172 | 0.5879344 | 2.36798912 | 0.30605374 | 0.47867401 | EPO1 | XIII | 514456 | 517287 | 1 |
| YMR125W | 16.2205696 | -0.6531132 | 1.15512021 | 2.54785182 | 0.27973126 | 0.45380891 | STO1 | XIII | 517539 | 520446 | 1 |
| YMR126C | 18.8130016 | 0.20280833 | 0.88765532 | 2.7763635 | 0.2495286 | 0.42048038 | DLT1 | XIII | 520761 | 521789 | -1 |
| YMR127C | 992.383999 | 1.48117298 | 0.46383229 | 66.4345489 | 3.75E-15 | 4.02E-13 | SAS2 | XIII | 522329 | 523345 | -1 |
| YMR128W | 41.5662843 | -0.7531805 | 0.62983511 | 7.86122349 | 0.01963166 | 0.06510155 | ECM16 | XIII | 523696 | 527499 | 1 |
| YMR129W | 58.959327 | -0.2216299 | 0.66526361 | 6.87410442 | 0.03215934 | 0.0942798 | POM152 | XIII | 527804 | 531817 | 1 |
| YMR130W | 179.659093 | -0.2269524 | 0.60030217 | 0.3749221 | 0.82906141 | 0.89791724 | NA | XIII | 532119 | 533027 | 1 |
| YMR131C | 72.1682185 | -0.22323 | 0.60358475 | 8.51891203 | 0.01412999 | 0.05110095 | RRB1 | XIII | 533163 | 534698 | -1 |

| | | | | | | | | | | | |
|---|---|---|---|---|---|---|---|---|---|---|---|
| YMR132C | 58.6891595 | -1.6404416 | 0.57499912 | 28.4617005 | 6.60E-07 | 1.23E-05 | JLP2 | XIII | 534944 | 535570 | -1 |
| YMR133W | 888.151951 | 0.11142567 | 0.59341916 | 0.56043513 | 0.75561933 | 0.85233507 | REC114 | XIII | 536207 | 537609 | 1 |
| YMR134W | 27.4120029 | 0.24961046 | 0.71068919 | 0.13611666 | 0.93420598 | 0.95735444 | ERG29 | XIII | 537838 | 538551 | 1 |
| YMR135C | 372.328043 | 0.19241352 | 0.5115 | 4.33236104 | 0.11461455 | 0.24149136 | GID8 | XIII | 538689 | 540056 | -1 |
| YMR136W | 32.0518832 | -0.1146691 | 0.71438603 | 0.46459549 | 0.79271006 | 0.87407947 | GAT2 | XIII | 541199 | 542881 | 1 |
| YMR137C | 16.0910266 | -0.4902994 | 0.8329257 | 1.11004653 | 0.57405891 | 0.72398194 | PSO2 | XIII | 542978 | 544963 | -1 |
| YMR138W | 10.4156651 | 1.1569618 | 1.09244515 | 6.3229871 | 0.04236242 | 0.11597089 | CIN4 | XIII | 545155 | 545730 | 1 |
| YMR139W | 14.5939226 | 0.87473373 | 0.83326003 | 1.76998813 | 0.41271662 | 0.58426143 | RIM11 | XIII | 546125 | 547237 | 1 |
| YMR140W | 22.4743501 | 0.86877982 | 0.79155315 | 2.85577264 | 0.23981528 | 0.40803129 | SIP5 | XIII | 547714 | 549183 | 1 |
| YMR141C | 1612.15191 | 1.47001862 | 0.45991981 | 12.204249 | 0.00223811 | 0.01217918 | NA | XIII | 549736 | 550044 | -1 |
| YMR142C | 71.1469457 | -0.7243499 | 0.67516958 | 15.845667 | 0.00036237 | 0.0028442 | RPL13B | XIII | 550206 | 551207 | -1 |
| YMR143W | 30.3857902 | -0.705372 | 2.13760376 | 1.73008056 | 0.42103459 | 0.59184032 | RPS16A | XIII | 551928 | 552903 | 1 |
| YMR144W | 38.2350813 | 0.95506404 | 0.63144089 | 2.69944716 | 0.25931193 | 0.43094606 | FDO1 | XIII | 553362 | 554390 | 1 |
| YMR145C | 76.1691602 | 0.49732703 | 0.67193948 | 2.75494896 | 0.25221472 | 0.42365049 | NDE1 | XIII | 554793 | 556475 | -1 |
| YMR146C | 84.9429412 | -0.7226906 | 0.71268693 | 8.46331433 | 0.01452829 | 0.0523127 | TIF34 | XIII | 557481 | 558524 | -1 |
| YMR147W | 35.163466 | -0.969101 | 0.63309425 | 6.48368992 | 0.03909171 | 0.10976769 | NA | XIII | 559199 | 559870 | 1 |
| YMR149W | 74.0052761 | 1.75779064 | 0.7955937 | 8.44156905 | 0.01468712 | 0.05261147 | SWP1 | XIII | 560996 | 561856 | 1 |
| YMR150C | 131.440535 | 1.89051013 | 0.77480448 | 6.25911082 | 0.04373724 | 0.1185574 | IMP1 | XIII | 561956 | 562528 | -1 |
| YMR152W | 19.27489 | -0.8283613 | 0.7472036 | 3.34325046 | 0.18794137 | 0.34376539 | YIM1 | XIII | 563096 | 564193 | 1 |
| YMR153W | 689.723149 | -0.0547416 | 0.50996281 | 5.07005742 | 0.07925944 | 0.18517193 | NUP53 | XIII | 564435 | 565862 | 1 |
| YMR154C | 483.685533 | -0.3802482 | 0.45930178 | 8.88698187 | 0.01175483 | 0.04445491 | RIM13 | XIII | 565999 | 568182 | -1 |
| YMR155W | 50.6163872 | -0.041888 | 0.59732423 | 2.78594458 | 0.24833608 | 0.41920289 | NA | XIII | 568551 | 570194 | 1 |
| YMR156C | 725.556989 | 0.45877926 | 0.63107507 | 0.52992353 | 0.76723529 | 0.85857988 | TPP1 | XIII | 570300 | 571016 | -1 |
| YMR157C | 180.484619 | 0.33624154 | 0.61631726 | 0.76861878 | 0.68092072 | 0.80384598 | AIM36 | XIII | 571277 | 572044 | -1 |
| YMR158W | 31.1766209 | -1.5877103 | 0.7506974 | 7.3389446 | 0.02548992 | 0.07880632 | MRPS8 | XIII | 572248 | 572715 | 1 |
| YMR160W | 302.71924 | 0.14456055 | 0.51361973 | 20.782502 | 3.07E-05 | 0.00034448 | NA | XIII | 575066 | 577516 | 1 |
| YMR161W | 11.9788389 | -0.3381165 | 0.91810349 | 0.16990984 | 0.91855369 | 0.94954934 | HLJ1 | XIII | 577718 | 578392 | 1 |
| YMR162C | 40.4705831 | 0.1496792 | 0.64527624 | 1.51839345 | 0.46804224 | 0.63108197 | DNF3 | XIII | 578951 | 583921 | -1 |
| YMR163C | 46.2938112 | -2.1778581 | 0.63993675 | 19.8181619 | 4.97E-05 | 0.00051177 | INP2 | XIII | 584271 | 586388 | -1 |
| YMR164C | 102.876209 | 0.16316999 | 0.63573357 | 11.1748687 | 0.00374462 | 0.01821964 | MSS11 | XIII | 587274 | 589550 | -1 |
| YMR165C | 562.182382 | 0.228349 | 0.47068598 | 4.9449535 | 0.08437562 | 0.19386304 | PAH1 | XIII | 590040 | 592628 | -1 |
| YMR166C | 10.7988574 | -0.1857648 | 0.94799317 | 0.16993789 | 0.91854081 | 0.94954934 | MME1 | XIII | 593367 | 594473 | -1 |
| YMR167W | 852.509593 | 0.05207995 | 0.49749705 | 2.56054845 | 0.27796107 | 0.45261809 | MLH1 | XIII | 594886 | 597195 | 1 |

| | | | | | | | | | | | |
|---|---|---|---|---|---|---|---|---|---|---|---|
| YMR168C | 977.985175 | -0.3341945 | 0.47593874 | 2.43503816 | 0.29596352 | 0.46995614 | CEP3 | XIII | 597332 | 599158 | -1 |
| YMR169C | 35.1993292 | -0.8124473 | 0.63278658 | 5.19738861 | 0.07437062 | 0.17629168 | ALD3 | XIII | 599352 | 600872 | -1 |
| YMR170C | 166.388819 | 0.53264312 | 0.50000301 | 6.05900146 | 0.04833977 | 0.12713301 | ALD2 | XIII | 601562 | 603082 | -1 |
| YMR171C | 44.489071 | 0.55291988 | 0.7339755 | 28.9076869 | 5.28E-07 | 1.02E-05 | EAR1 | XIII | 603868 | 605520 | -1 |
| YMR172W | 118.53845 | -0.2827967 | 0.51455935 | 0.30868075 | 0.85698028 | 0.91531374 | HOT1 | XIII | 605981 | 608140 | 1 |
| YMR173W | 79.5212963 | -0.044985 | 0.54259284 | 1.43737404 | 0.48739177 | 0.64903366 | DDR48 | XIII | 608689 | 609981 | 1 |
| YMR174C | 135.488713 | 2.68847495 | 0.85049375 | 9.21220427 | 0.00999068 | 0.03943154 | PAI3 | XIII | 610159 | 610365 | -1 |
| YMR175W-A | 23.8353227 | 0.65942266 | 0.80756353 | 9.49180201 | 0.00868723 | 0.03535208 | NA | XIII | 611314 | 611508 | 1 |
| YMR176W | 47.8402548 | 0.16485219 | 0.60731807 | 3.26701851 | 0.19524321 | 0.35245914 | ECM5 | XIII | 611740 | 615975 | 1 |
| YMR177W | 42.6302027 | 1.24297627 | 0.78532374 | 34.2748927 | 3.61E-08 | 9.72E-07 | MMT1 | XIII | 616566 | 618098 | 1 |
| YMR178W | 316.059453 | -2.438919 | 0.50962311 | 48.4470682 | 3.02E-11 | 1.55E-09 | NA | XIII | 618479 | 619303 | 1 |
| YMR179W | 366.546287 | 2.05624834 | 0.56648276 | 12.5875969 | 0.00184773 | 0.01056192 | SPT21 | XIII | 619858 | 622134 | 1 |
| YMR180C | 349.881297 | 3.52449622 | 0.5056618 | 46.1211437 | 9.66E-11 | 4.58E-09 | CTL1 | XIII | 622251 | 623213 | -1 |
| YMR181C | 602.157426 | -0.8131022 | 0.46824133 | 14.4223485 | 0.00073829 | 0.00511327 | NA | XIII | 623616 | 624080 | -1 |
| YMR182C | 789.244588 | -0.2985402 | 0.48429627 | 0.83027263 | 0.66025027 | 0.78805382 | RGM1 | XIII | 624532 | 625167 | -1 |
| YMR182W-A | 30.6548282 | 1.0487834 | 0.77319327 | 11.5989766 | 0.0030291 | 0.01538156 | NA | XIII | 625811 | 625897 | 1 |
| YMR183C | 39.114781 | -0.2759312 | 0.74903556 | 5.05197647 | 0.07997924 | 0.18642503 | SSO2 | XIII | 626921 | 627808 | -1 |
| YMR184W | 21.6053344 | 2.08003252 | 0.96230554 | 5.15889535 | 0.07581587 | 0.17873529 | ADD37 | XIII | 628189 | 628785 | 1 |
| YMR185W | 12.4895121 | 0.23960185 | 0.9386584 | 0.08617217 | 0.95782893 | 0.9702291 | RTP1 | XIII | 629025 | 631970 | 1 |
| YMR186W | 1339.60841 | 0.27453599 | 0.52230592 | 2.79046993 | 0.24777481 | 0.41849946 | HSC82 | XIII | 632355 | 634472 | 1 |
| YMR187C | 184.922633 | 0.78169495 | 0.50513557 | 4.19293515 | 0.12288976 | 0.25402775 | NA | XIII | 634689 | 635984 | -1 |
| YMR188C | 17.9843346 | -0.841493 | 1.09081393 | 1.00083122 | 0.60627863 | 0.75088539 | MRPS17 | XIII | 636291 | 637004 | -1 |
| YMR189W | 21.7328492 | -0.1371873 | 0.73404247 | 0.42901673 | 0.80693806 | 0.88421109 | GCV2 | XIII | 637500 | 640604 | 1 |
| YMR190C | 21.2037388 | 0.55081815 | 0.73325691 | 1.69089565 | 0.42936504 | 0.59890573 | SGS1 | XIII | 640915 | 645258 | -1 |
| YMR191W | 78.868164 | 1.7968701 | 0.60289581 | 19.3870398 | 6.17E-05 | 0.00061682 | SPG5 | XIII | 645656 | 646777 | 1 |
| YMR192W | 86.7694842 | 1.14586034 | 0.60238852 | 32.0817225 | 1.08E-07 | 2.58E-06 | GYL1 | XIII | 647118 | 649280 | 1 |
| YMR193W | 44.0970783 | -0.7246202 | 0.59944149 | 5.58599298 | 0.06123744 | 0.15215948 | MRPL24 | XIII | 650036 | 650812 | 1 |
| YMR194C-B | 62.2148507 | -0.3539627 | 0.61491176 | 5.70144038 | 0.05780268 | 0.14608359 | CMC4 | XIII | 652594 | 652887 | -1 |
| YMR194W | 33.430147 | -0.9492853 | 0.67374085 | 14.1816655 | 0.0008327 | 0.00566552 | RPL36A | XIII | 651145 | 651910 | 1 |
| YMR195W | 42.1386467 | 2.61421638 | 1.07359236 | 5.67947817 | 0.05844091 | 0.14688185 | ICY1 | XIII | 654034 | 654417 | 1 |
| YMR196W | 512.657772 | -0.253323 | 0.48696483 | 12.6943449 | 0.00175169 | 0.01015246 | NA | XIII | 655076 | 658342 | 1 |
| YMR197C | 19.3812042 | 1.08445046 | 0.91258217 | 2.6521517 | 0.26551715 | 0.4376158 | VTI1 | XIII | 658545 | 659198 | -1 |
| YMR198W | 10.1150821 | -1.0446597 | 0.92958622 | 3.2304051 | 0.19885039 | 0.35758801 | CIK1 | XIII | 659745 | 661529 | 1 |

| | | | | | | | | | | | |
|---|---|---|---|---|---|---|---|---|---|---|---|
| YMR199W | 177.684583 | 1.31977851 | 0.57532342 | 22.5987904 | 1.24E-05 | 0.00015894 | CLN1 | XIII | 662644 | 664284 | 1 |
| YMR200W | 53.8132285 | 1.17186657 | 0.64058123 | 11.5874959 | 0.00304654 | 0.01538156 | ROT1 | XIII | 664752 | 665522 | 1 |
| YMR201C | 524.234841 | 2.7332871 | 0.88862414 | 9.63336205 | 0.00809361 | 0.03344895 | RAD14 | XIII | 665845 | 667044 | -1 |
| YMR202W | 53.4306376 | -0.3299651 | 0.67013222 | 0.74545519 | 0.68885285 | 0.81049746 | ERG2 | XIII | 667537 | 668205 | 1 |
| YMR203W | 27.731935 | 0.23507334 | 0.6555841 | 1.78056211 | 0.41054035 | 0.58217699 | TOM40 | XIII | 668492 | 669655 | 1 |
| YMR204C | 21.4404617 | -0.0342981 | 0.91413722 | 4.59874427 | 0.10032181 | 0.21944212 | INP1 | XIII | 670063 | 671325 | -1 |
| YMR205C | 70.8931425 | -0.1422174 | 0.8188046 | 5.22131703 | 0.07348614 | 0.17480884 | PFK2 | XIII | 671887 | 674766 | -1 |
| YMR206W | 182.174668 | 0.01987435 | 0.50170279 | 0.01498432 | 0.99253584 | 0.99425303 | NA | XIII | 675896 | 676837 | 1 |
| YMR207C | 49.0408939 | 0.19023975 | 0.61905359 | 0.68432214 | 0.7102338 | 0.82438173 | NA | XIII | 677193 | 683564 | -1 |
| YMR208W | 101.994348 | -0.2360809 | 0.5425788 | 0.55250415 | 0.75862168 | 0.85422394 | ERG12 | XIII | 684467 | 685798 | 1 |
| YMR209C | 24.8871767 | 0.76983614 | 0.72655017 | 6.36132442 | 0.04155813 | 0.11436385 | NA | XIII | 685911 | 687284 | -1 |
| YMR210W | 887.559967 | -0.8393933 | 0.50999797 | 2.69512177 | 0.25987335 | 0.43163129 | MGL2 | XIII | 687516 | 688865 | 1 |
| YMR211W | 80.7322697 | -1.0054826 | 0.56205279 | 24.4895548 | 4.81E-06 | 6.90E-05 | DML1 | XIII | 689083 | 690510 | 1 |
| YMR212C | 37.3184876 | -0.3084912 | 0.64767915 | 5.21754312 | 0.07362493 | 0.17499522 | EFR3 | XIII | 690695 | 693043 | -1 |
| YMR213W | 21.9636381 | 0.09428293 | 0.76842256 | 0.27038655 | 0.87354706 | 0.92573532 | CEF1 | XIII | 693381 | 695153 | 1 |
| YMR214W | 82.5827693 | 1.16848549 | 0.58216964 | 13.3792579 | 0.00124374 | 0.00786166 | SCJ1 | XIII | 695350 | 696483 | 1 |
| YMR215W | 144.308261 | -0.6540666 | 0.50468507 | 5.98227498 | 0.05023027 | 0.1311241 | GAS3 | XIII | 696796 | 698370 | 1 |
| YMR216C | 34.5712365 | 0.62365462 | 0.71206238 | 0.78061441 | 0.67684891 | 0.80093096 | SKY1 | XIII | 698811 | 701039 | -1 |
| YMR217W | 77.3674681 | -0.3650679 | 0.69725022 | 9.28234748 | 0.00964637 | 0.03839103 | GUA1 | XIII | 701790 | 703367 | 1 |
| YMR218C | 18.9631284 | 0.62278479 | 0.88432701 | 0.74246825 | 0.6898824 | 0.81104957 | TRS130 | XIII | 703579 | 706887 | -1 |
| YMR219W | 253.529671 | -0.6102729 | 0.48399323 | 3.6037226 | 0.1649915 | 0.31392516 | ESC1 | XIII | 707133 | 712109 | 1 |
| YMR220W | 416.984848 | 1.48626459 | 0.58314183 | 45.8103912 | 1.13E-10 | 5.23E-09 | ERG8 | XIII | 712316 | 713671 | 1 |
| YMR221C | 96.650884 | -0.0685309 | 0.58959593 | 5.62558327 | 0.06003716 | 0.15002811 | FMP42 | XIII | 713931 | 715445 | -1 |
| YMR222C | 15.9101257 | -0.8458096 | 0.80507004 | 3.41353387 | 0.18145149 | 0.33630094 | FSH2 | XIII | 715638 | 716309 | -1 |
| YMR223W | 326.772084 | 1.13144482 | 0.6243245 | 96.492647 | 1.11E-21 | 4.61E-19 | UBP8 | XIII | 716715 | 718130 | 1 |
| YMR224C | 53.2771094 | 1.53222731 | 0.72473888 | 12.5387154 | 0.00189344 | 0.01073788 | MRE11 | XIII | 718575 | 720653 | -1 |
| YMR225C | 65.9524926 | -0.0799439 | 0.6008165 | 9.83937275 | 0.00730142 | 0.03103908 | MRPL44 | XIII | 720960 | 721403 | -1 |
| YMR226C | 87.642058 | 0.53311582 | 0.60077634 | 1.05056873 | 0.59138717 | 0.73843686 | NA | XIII | 721593 | 722396 | -1 |
| YMR227C | 108.018012 | 0.9723077 | 0.60874904 | 2.61704585 | 0.2702189 | 0.44347149 | TAF7 | XIII | 722613 | 724385 | -1 |
| YMR228W | 21.6624428 | -0.2788599 | 0.73822404 | 0.44989323 | 0.79855885 | 0.87821055 | MTF1 | XIII | 724626 | 725651 | 1 |
| YMR229C | 58.2710905 | -0.5437429 | 0.68813306 | 0.67322275 | 0.71418634 | 0.82607012 | RRP5 | XIII | 725934 | 731123 | -1 |
| YMR230W | 62.8559353 | -0.3783352 | 0.64050325 | 18.1907132 | 0.00011219 | 0.0010294 | RPS10B | XIII | 732414 | 733141 | 1 |
| YMR231W | 12.0493044 | 0.89098422 | 0.92142243 | 1.23844777 | 0.53836211 | 0.69284654 | PEP5 | XIII | 733545 | 736634 | 1 |

| | | | | | | | | | | | |
|---|---|---|---|---|---|---|---|---|---|---|---|
| YMR232W | 16.9539949 | 0.55221333 | 0.8576627 | 2.04474196 | 0.35974099 | 0.53284736 | FUS2 | XIII | 736926 | 738959 | 1 |
| YMR233W | 24.966868 | 0.29971157 | 0.73027994 | 1.66247146 | 0.43551078 | 0.60441213 | TRI1 | XIII | 739185 | 739865 | 1 |
| YMR234W | 95.2734434 | 1.17088281 | 0.65693064 | 3.80217509 | 0.14940605 | 0.29203007 | RNH1 | XIII | 740266 | 741312 | 1 |
| YMR235C | 225.586251 | 0.27983251 | 0.57955922 | 38.3802886 | 4.63E-09 | 1.50E-07 | RNA1 | XIII | 741512 | 742735 | -1 |
| YMR236W | 9.33532621 | 0.27002193 | 1.06661532 | 0.89693168 | 0.63860713 | 0.77465312 | TAF9 | XIII | 742971 | 743444 | 1 |
| YMR237W | 167.31406 | 2.95546682 | 0.5668497 | 36.7564837 | 1.04E-08 | 3.11E-07 | BCH1 | XIII | 743749 | 745923 | 1 |
| YMR238W | 53.7888874 | 1.06024982 | 0.75411451 | 2.02996011 | 0.36240966 | 0.53529386 | DFG5 | XIII | 746353 | 747729 | 1 |
| YMR239C | 109.141091 | -0.3694195 | 0.55836697 | 0.72284097 | 0.69668599 | 0.81659461 | RNT1 | XIII | 748262 | 749677 | -1 |
| YMR240C | 16.1986197 | 0.32209845 | 0.90865754 | 0.27105016 | 0.87325726 | 0.92569746 | CUS1 | XIII | 749930 | 751240 | -1 |
| YMR241W | 54.6898996 | -0.7307502 | 0.6034633 | 6.89179255 | 0.03187618 | 0.09368684 | YHM2 | XIII | 751961 | 752905 | 1 |
| YMR242C | 61.3521873 | -0.5452305 | 0.55137597 | 19.4729207 | 5.91E-05 | 0.00059708 | NA | XIII | 753225 | 753742 | -1 |
| YMR242W-A | 324.962782 | -0.6538372 | 0.62176215 | 15.9625004 | 0.00034181 | 0.00272586 | NA | XIII | 754297 | 754386 | 1 |
| YMR243C | 11.7024977 | -1.1133594 | 1.06712773 | 6.16840794 | 0.04576645 | 0.12211417 | ZRC1 | XIII | 754838 | 756166 | -1 |
| YMR244C-A | 23.5356352 | -0.4190833 | 0.74085944 | 7.37666335 | 0.0250137 | 0.07790711 | COA6 | XIII | 758517 | 758831 | -1 |
| YMR244W | 16.0120832 | -1.7067901 | 0.86982485 | 5.59323216 | 0.06101619 | 0.15186924 | NA | XIII | 757250 | 758317 | 1 |
| YMR246W | 53.9384973 | -1.2675227 | 0.79238421 | 7.81021392 | 0.0201388 | 0.06640299 | FAA4 | XIII | 759807 | 761891 | 1 |
| YMR247C | 563.075547 | 0.42743033 | 0.48308622 | 1.92934131 | 0.3811087 | 0.55483451 | RKR1 | XIII | 763351 | 768039 | -1 |
| YMR247W-A | 182.409228 | 1.07516006 | 0.51955815 | 14.7870811 | 0.00061521 | 0.00439222 | NA | XIII | 769283 | 769426 | 1 |
| YMR250W | 33.4089057 | -0.2599038 | 0.69857112 | 1.34507349 | 0.51041215 | 0.66786132 | GAD1 | XIII | 770801 | 772558 | 1 |
| YMR251W | 522.415512 | 2.49010247 | 0.51004704 | 25.1356752 | 3.48E-06 | 5.24E-05 | GTO3 | XIII | 772915 | 774015 | 1 |
| YMR251W-A | 15.753937 | 1.96828927 | 0.87374598 | 5.17109151 | 0.07535494 | 0.17808372 | HOR7 | XIII | 774752 | 774931 | 1 |
| YMR252C | 21.1232859 | 0.63425045 | 0.73620014 | 1.36930801 | 0.50426467 | 0.66262751 | NA | XIII | 775315 | 775719 | -1 |
| YMR253C | 389.585196 | 0.32213752 | 0.50942499 | 4.81678072 | 0.08995998 | 0.20354369 | NA | XIII | 775946 | 777190 | -1 |
| YMR254C | 284.69552 | 0.57684977 | 0.73487856 | 2.53421914 | 0.28164452 | 0.45576349 | NA | XIII | 777615 | 777923 | -1 |
| YMR255W | 65.183087 | 0.39587375 | 0.66840952 | 0.42987045 | 0.80659369 | 0.88416839 | GFD1 | XIII | 778001 | 778567 | 1 |
| YMR257C | 20.0722321 | -0.8118836 | 0.78080825 | 1.27936691 | 0.52745936 | 0.68184633 | PET111 | XIII | 779629 | 782031 | -1 |
| YMR258C | 18.0929946 | 0.29096005 | 0.79490743 | 0.22346623 | 0.8942829 | 0.9350658 | ROY1 | XIII | 782620 | 784281 | -1 |
| YMR259C | 243.975792 | 1.46048869 | 0.51641393 | 10.8520785 | 0.00440049 | 0.02061395 | TRM732 | XIII | 784621 | 788883 | -1 |
| YMR260C | 68.4084866 | -0.914126 | 0.83474287 | 8.17285352 | 0.01679915 | 0.05813116 | TIF11 | XIII | 789378 | 789839 | -1 |
| YMR261C | 27.5113042 | -0.3322392 | 1.07646309 | 0.53625819 | 0.76480904 | 0.85692899 | TPS3 | XIII | 790205 | 793369 | -1 |
| YMR262W | 20.9902952 | 0.45257282 | 0.89255865 | 0.53609349 | 0.76487203 | 0.85692899 | NA | XIII | 793726 | 794667 | 1 |
| YMR263W | 55.8552871 | -0.7993285 | 0.56568699 | 16.0030104 | 0.00033496 | 0.00267504 | SAP30 | XIII | 794919 | 795524 | 1 |
| YMR264W | 185.539888 | -0.6675282 | 0.58746825 | 12.8650846 | 0.00160836 | 0.00961569 | CUE1 | XIII | 795805 | 796416 | 1 |

| | | | | | | | | | | | |
|---|---|---|---|---|---|---|---|---|---|---|---|
| YMR265C | 79.7643207 | -0.2592073 | 0.59507811 | 4.48897671 | 0.10598175 | 0.22838155 | NA | XIII | 796540 | 797925 | -1 |
| YMR266W | 46.0305882 | -0.2211678 | 0.59119051 | 0.80175799 | 0.6697311 | 0.79510827 | RSN1 | XIII | 798518 | 801379 | 1 |
| YMR267W | 60.5667982 | -0.2570502 | 0.60222376 | 1.17556657 | 0.55555743 | 0.70703257 | PPA2 | XIII | 801772 | 802704 | 1 |
| YMR268C | 20.5685782 | 0.07462515 | 0.80976372 | 5.95977042 | 0.05079866 | 0.13219068 | PRP24 | XIII | 802888 | 804222 | -1 |
| YMR269W | 46.7332342 | -0.4868838 | 0.6012178 | 2.98218304 | 0.22512679 | 0.39049854 | TMA23 | XIII | 804456 | 805091 | 1 |
| YMR270C | 8.39799049 | -0.0766159 | 1.13923476 | 0.16679576 | 0.91998503 | 0.95001129 | RRN9 | XIII | 805326 | 806423 | -1 |
| YMR271C | 47.632561 | -1.2620978 | 0.61022572 | 8.72677008 | 0.01273521 | 0.0472672 | URA10 | XIII | 806865 | 807548 | -1 |
| YMR272C | 9.94260701 | -0.0077804 | 1.02718886 | 0.09644403 | 0.9529222 | 0.96731849 | SCS7 | XIII | 809623 | 810777 | -1 |
| YMR272W-B | 164.572769 | 0.14756362 | 0.70265563 | 0.60835152 | 0.73773119 | 0.84034303 | NA | XIII | 811089 | 811196 | 1 |
| YMR273C | 130.163054 | 1.41659878 | 0.72654963 | 3.86973302 | 0.14444355 | 0.28495484 | ZDS1 | XIII | 811233 | 813980 | -1 |
| YMR274C | 16.6674444 | -0.1589551 | 0.97034307 | 0.10582163 | 0.9484646 | 0.96471939 | RCE1 | XIII | 814364 | 815311 | -1 |
| YMR275C | 71.9229385 | -0.5088519 | 0.55185778 | 0.98450411 | 0.61124828 | 0.75489714 | BUL1 | XIII | 815651 | 818581 | -1 |
| YMR276W | 84.895957 | -0.2024652 | 0.52124973 | 3.63405919 | 0.16250775 | 0.31104789 | DSK2 | XIII | 818827 | 819948 | 1 |
| YMR277W | 76.9720482 | -0.7122794 | 0.54331549 | 1.77860477 | 0.41094233 | 0.58260433 | FCP1 | XIII | 820256 | 822454 | 1 |
| YMR278W | 9.33355823 | -0.2204746 | 1.0483061 | 0.28667924 | 0.86645975 | 0.92150571 | PRM15 | XIII | 822763 | 824631 | 1 |
| YMR279C | 2973.57562 | 0.21552912 | 0.60143452 | 0.13834849 | 0.93316407 | 0.95735444 | NA | XIII | 824729 | 826351 | -1 |
| YMR280C | 1075.89824 | -0.691729 | 0.50867941 | 11.5294861 | 0.0031362 | 0.01579009 | CAT8 | XIII | 827028 | 831329 | -1 |
| YMR281W | 2186.48818 | 3.99099997 | 0.47526191 | 117.664425 | 2.82E-26 | 2.72E-23 | GPI12 | XIII | 832339 | 833253 | 1 |
| YMR282C | 45.6882923 | 1.08723967 | 0.63208868 | 6.11725021 | 0.04695221 | 0.12464616 | AEP2 | XIII | 833356 | 835098 | -1 |
| YMR283C | 17.457102 | -0.9643637 | 0.82519602 | 1.85120892 | 0.3962918 | 0.56739108 | RIT1 | XIII | 835326 | 836867 | -1 |
| YMR284W | 43.7424105 | 1.52291222 | 0.8670334 | 7.86803276 | 0.01956493 | 0.06501327 | YKU70 | XIII | 838187 | 839995 | 1 |
| YMR285C | 35.939145 | 0.34805678 | 0.64259495 | 0.48039831 | 0.78647122 | 0.87021248 | NGL2 | XIII | 840144 | 841691 | -1 |
| YMR286W | 13.4450905 | -0.2987823 | 0.83123743 | 0.20255045 | 0.90368428 | 0.93937738 | MRPL33 | XIII | 841942 | 842202 | 1 |
| YMR287C | 19.8916551 | -0.6422366 | 0.75681902 | 1.91201309 | 0.38442501 | 0.5574307 | DSS1 | XIII | 842436 | 845345 | -1 |
| YMR288W | 64.6967216 | -0.2701486 | 0.55442582 | 0.25178509 | 0.88170959 | 0.9300006 | HSH155 | XIII | 845571 | 848486 | 1 |
| YMR289W | 116.255289 | 1.31903538 | 0.59479707 | 44.1622714 | 2.57E-10 | 1.09E-08 | ABZ2 | XIII | 848685 | 849809 | 1 |
| YMR290C | 189.258027 | -0.0033211 | 0.49082816 | 0.42848721 | 0.80715174 | 0.88427787 | HAS1 | XIII | 850074 | 851591 | -1 |
| YMR291W | 38.8690368 | -1.4882332 | 0.66794667 | 9.01033048 | 0.01105176 | 0.04244332 | TDA1 | XIII | 852630 | 854390 | 1 |
| YMR292W | 249.383657 | 0.21140718 | 0.63485683 | 0.13707508 | 0.93375841 | 0.95735444 | GOT1 | XIII | 854795 | 855293 | 1 |
| YMR293C | 14.1865154 | -0.7081756 | 0.88807609 | 1.09576028 | 0.57817416 | 0.727271 | HER2 | XIII | 855399 | 856793 | -1 |
| YMR294W | 109.184096 | 3.36669208 | 1.05395221 | 10.7120277 | 0.00471968 | 0.02186157 | JNM1 | XIII | 856966 | 858087 | 1 |
| YMR295C | 136.859378 | 1.29340531 | 0.58155488 | 4.88314329 | 0.08702397 | 0.19878563 | NA | XIII | 858297 | 858890 | -1 |
| YMR296C | 253.459598 | 1.27213686 | 0.51193037 | 28.4700126 | 6.57E-07 | 1.23E-05 | LCB1 | XIII | 859215 | 860891 | -1 |

| | | | | | | | | | | | |
|---|---|---|---|---|---|---|---|---|---|---|---|
| YMR297W | 82.6784122 | 0.3343721 | 0.6231613 | 1.16533406 | 0.55840709 | 0.70949683 | PRC1 | XIII | 861922 | 863520 | 1 |
| YMR298W | 62.3430126 | -0.1703369 | 0.57046601 | 1.2801731 | 0.52724679 | 0.68184633 | LIP1 | XIII | 863819 | 864271 | 1 |
| YMR299C | 94.1038902 | -0.1857039 | 0.54622741 | 0.43412244 | 0.80488069 | 0.88279205 | DYN3 | XIII | 864412 | 865350 | -1 |
| YMR300C | 82.0272278 | 0.57060031 | 0.6460539 | 29.3881051 | 4.15E-07 | 8.21E-06 | ADE4 | XIII | 865559 | 867091 | -1 |
| YMR301C | 40.9017031 | -0.9429776 | 0.66383928 | 3.80344215 | 0.14931142 | 0.29203007 | ATM1 | XIII | 867555 | 869627 | -1 |
| YMR302C | 45.7471963 | 0.50849415 | 0.79960341 | 0.95139521 | 0.62145138 | 0.76184702 | YME2 | XIII | 870073 | 872625 | -1 |
| YMR303C | 589.12651 | -1.7574861 | 0.54739131 | 14.3897347 | 0.00075043 | 0.00517876 | ADH2 | XIII | 873291 | 874337 | -1 |
| YMR304W | 26.9568287 | 0.17680278 | 0.82260042 | 0.25634102 | 0.87970337 | 0.92911891 | UBP15 | XIII | 874987 | 878679 | 1 |
| YMR305C | 89.9559844 | 1.23295309 | 0.60340253 | 5.16232255 | 0.07568606 | 0.17861909 | SCW10 | XIII | 879063 | 880232 | -1 |
| YMR306W | 957.648805 | -0.4517422 | 0.53756358 | 3.5509367 | 0.16940409 | 0.31949501 | FKS3 | XIII | 881159 | 886516 | 1 |
| YMR307W | 84.8820185 | -0.2448293 | 0.72499495 | 3.95580616 | 0.13835906 | 0.27652708 | GAS1 | XIII | 887003 | 888682 | 1 |
| YMR308C | 32.0667848 | -1.014042 | 0.68746723 | 4.53055415 | 0.10380127 | 0.22509714 | PSE1 | XIII | 888952 | 892221 | -1 |
| YMR309C | 29.015788 | -0.2890901 | 0.92631798 | 5.83826413 | 0.05398052 | 0.13854043 | NIP1 | XIII | 892988 | 895426 | -1 |
| YMR310C | 20.7709717 | -1.439065 | 0.74066105 | 14.1577015 | 0.00084274 | 0.00572036 | NA | XIII | 895716 | 896669 | -1 |
| YMR311C | 112.785042 | 0.93322968 | 0.59102984 | 2.48822819 | 0.28819611 | 0.46197549 | GLC8 | XIII | 896914 | 897603 | -1 |
| YMR312W | 116.099661 | 1.52529045 | 0.69914839 | 8.49133962 | 0.01432613 | 0.05171342 | ELP6 | XIII | 898405 | 899226 | 1 |
| YMR313C | 42.9923358 | -1.1660656 | 0.61456617 | 5.89575703 | 0.05245086 | 0.13539478 | TGL3 | XIII | 899379 | 901307 | -1 |
| YMR314W | 51.4940978 | 1.47860104 | 0.73140856 | 4.12895091 | 0.12688483 | 0.2604265 | PRE5 | XIII | 901709 | 902413 | 1 |
| YMR315W | 3358.23331 | 4.3885451 | 0.4684489 | 73.9485637 | 8.76E-17 | 1.17E-14 | NA | XIII | 902800 | 903849 | 1 |
| YMR315W-A | 68.6713206 | -0.3031413 | 0.5806654 | 0.96542466 | 0.61710732 | 0.75846612 | NA | XIII | 904286 | 904393 | 1 |
| YMR316C-A | 7202.28902 | 4.33314153 | 0.56442287 | 45.0256027 | 1.67E-10 | 7.44E-09 | NA | XIII | 905662 | 905973 | -1 |
| YMR316W | 121.020751 | -0.1321419 | 0.53221838 | 4.2528651 | 0.119262 | 0.24840182 | DIA1 | XIII | 904825 | 905835 | 1 |
| YMR317W | 3148.13071 | -0.3097755 | 0.47045035 | 1.01956396 | 0.60062651 | 0.74682758 | NA | XIII | 907364 | 910786 | 1 |
| YMR318C | 200.206853 | 0.30278985 | 0.55895311 | 1.53240165 | 0.46477548 | 0.62728439 | ADH6 | XIII | 911061 | 912143 | -1 |
| YMR319C | 1223.15629 | 1.54131385 | 0.54871284 | 7.77854745 | 0.0204602 | 0.06719418 | FET4 | XIII | 912880 | 914538 | -1 |
| YNCA0001W | 790.335017 | -0.0417205 | 0.56389257 | 2.32191797 | 0.3131857 | 0.48516903 | NA | NA | NA | NA | NA |
| YNCA0003W | 10.0289279 | 0.16098181 | 1.37547776 | 12.7762643 | 0.00168139 | 0.00981378 | NA | NA | NA | NA | NA |
| YNCB0003W | 39.4998415 | 1.18305469 | 0.68192656 | 6.55496383 | 0.03772313 | 0.10638914 | NA | NA | NA | NA | NA |
| YNCB0006W | 5.1851498 | 1.9521709 | 1.58241226 | 7.90514899 | 0.01920519 | 0.06416508 | NA | NA | NA | NA | NA |
| YNCB0008W | 63.4237925 | -0.7836188 | 0.5931509 | 2.27159582 | 0.32116576 | 0.49377316 | NA | NA | NA | NA | NA |
| YNCB0009C | 33.4896672 | 0.02221662 | 0.63508646 | 0.16358129 | 0.92146485 | 0.95035295 | NA | NA | NA | NA | NA |
| YNCB0010W | 1454.72461 | 2.649427 | 0.89359877 | 7.85973526 | 0.01964627 | 0.06511272 | NA | NA | NA | NA | NA |
| YNCB0014W | 10538.2983 | -1.2626296 | 0.4973922 | 35.8151894 | 1.67E-08 | 4.74E-07 | NA | NA | NA | NA | NA |

| | | | | | | | | | | | |
|---|---|---|---|---|---|---|---|---|---|---|---|
| YNCB0016W | 15.3023853 | 1.79762005 | 1.19004648 | 17.3431793 | 0.00017139 | 0.00148775 | NA | NA | NA | NA | NA |
| YNCB0017C | 81.9735576 | 0.75852538 | 0.69102692 | 22.48513 | 1.31E-05 | 0.00016676 | NA | NA | NA | NA | NA |
| YNCB0019C | 308.399574 | 1.4203159 | 0.55754321 | 6.4987871 | 0.03879773 | 0.10910095 | NA | NA | NA | NA | NA |
| YNCC0002W | 27.8134993 | 0.29695369 | 0.66402255 | 0.86102168 | 0.65017687 | 0.78140989 | NA | NA | NA | NA | NA |
| YNCC0003C | 2675.13139 | 1.3709146 | 0.8836782 | 7.67922124 | 0.02150197 | 0.06990254 | NA | NA | NA | NA | NA |
| YNCC0011W | 60.0694742 | -0.3667701 | 0.61628878 | 0.59676395 | 0.74201785 | 0.84243165 | NA | NA | NA | NA | NA |
| YNCC0013W | 11.0760926 | -1.7588274 | 1.01702642 | 4.74396846 | 0.09329542 | 0.20840297 | NA | NA | NA | NA | NA |
| YNCD0002C | 28.3098886 | 0.53906843 | 0.76930694 | 7.27962801 | 0.02625723 | 0.08048139 | NA | NA | NA | NA | NA |
| YNCD0005C | 6.50321707 | -1.6219546 | 1.19105893 | 2.67849222 | 0.26204315 | 0.43337041 | NA | NA | NA | NA | NA |
| YNCD0008W | 9.86040035 | 3.46383677 | 1.5657668 | 13.5878559 | 0.00112056 | 0.00722498 | NA | NA | NA | NA | NA |
| YNCD0009W | 10.6730292 | 1.44684677 | 1.27769193 | 12.948177 | 0.0015429 | 0.00934751 | NA | NA | NA | NA | NA |
| YNCD0015C | 9.16206783 | -0.6275156 | 1.82990312 | 26.3314364 | 1.92E-06 | 3.15E-05 | NA | NA | NA | NA | NA |
| YNCD0016C | 11.6028954 | 1.48871018 | 1.02947163 | 5.26830682 | 0.07177971 | 0.17202174 | NA | NA | NA | NA | NA |
| YNCD0017W | 101.952971 | 4.64041037 | 0.77226737 | 55.6056023 | 8.42E-13 | 5.95E-11 | NA | NA | NA | NA | NA |
| YNCD0030W | 19.5957293 | -1.0830785 | 0.85605607 | 1.98159249 | 0.37128094 | 0.54464572 | NA | NA | NA | NA | NA |
| YNCD0032C | 279.60473 | -0.3765966 | 0.54907207 | 2.6919902 | 0.26028057 | 0.43181218 | NA | NA | NA | NA | NA |
| YNCE0003W | 13.6869103 | -0.7433273 | 0.8438748 | 11.1208852 | 0.00384707 | 0.01862421 | NA | NA | NA | NA | NA |
| YNCE0004C | 15.2555353 | -4.2291166 | 1.20940839 | 20.1625627 | 4.19E-05 | 0.00044467 | NA | NA | NA | NA | NA |
| YNCE0007C | 95.2876892 | 2.31119795 | 0.66862523 | 38.7697246 | 3.81E-09 | 1.26E-07 | NA | NA | NA | NA | NA |
| YNCE0011C | 94.2778616 | -0.5046072 | 0.55748102 | 0.88418798 | 0.64268923 | 0.77621415 | NA | NA | NA | NA | NA |
| YNCE0012W | 48.4629026 | -0.5272071 | 0.71069141 | 5.5145476 | 0.06346455 | 0.15629934 | NA | NA | NA | NA | NA |
| YNCE0013W | 13.2278782 | 0.35627153 | 1.0325343 | 3.30506668 | 0.191564 | 0.34813419 | NA | NA | NA | NA | NA |
| YNCE0014W | 9.31911418 | 1.26562004 | 1.37140494 | 8.8372941 | 0.01205052 | 0.04533628 | NA | NA | NA | NA | NA |
| YNCE0017W | 6.85800097 | 0.41705276 | 1.11568606 | 0.14003858 | 0.93237584 | 0.95731799 | NA | NA | NA | NA | NA |
| YNCE0018W | 17.9646158 | 2.37689392 | 1.09982751 | 23.2295086 | 9.03E-06 | 0.00012049 | NA | NA | NA | NA | NA |
| YNCE0019W | 831.715072 | 2.40832916 | 0.58614296 | 26.8880701 | 1.45E-06 | 2.45E-05 | NA | NA | NA | NA | NA |
| YNCE0020C | 23.2007434 | 0.77390945 | 0.79861307 | 8.24091578 | 0.01623708 | 0.05677094 | NA | NA | NA | NA | NA |
| YNCF0001C | 6782.19328 | 1.68995936 | 0.95712619 | 5.05997144 | 0.07966016 | 0.18583091 | NA | NA | NA | NA | NA |
| YNCF0003C | 1193.16544 | 1.04176276 | 0.59062358 | 5.67034957 | 0.05870826 | 0.14739097 | NA | NA | NA | NA | NA |
| YNCF0010C | 7.29313492 | -2.0365163 | 1.5292797 | 10.2276119 | 0.00601315 | 0.02655695 | NA | NA | NA | NA | NA |
| YNCF0013W | 9.59737856 | -2.2497014 | 1.09406273 | 4.48992906 | 0.1059313 | 0.22838155 | NA | NA | NA | NA | NA |
| YNCG0001C | 137.098384 | -0.0122385 | 0.57178691 | 1.55189179 | 0.46026821 | 0.62448065 | NA | NA | NA | NA | NA |
| YNCG0006C | 14.9671049 | 0.21907501 | 0.89926983 | 0.11768355 | 0.94285594 | 0.96176085 | NA | NA | NA | NA | NA |

| | | | | | | | | | | | |
|---|---|---|---|---|---|---|---|---|---|---|---|
| YNCG0008W | 21.6187333 | 3.08902813 | 0.8891826 | 12.404239 | 0.00202513 | 0.01126371 | NA | NA | NA | NA | NA |
| YNCG0010W | 39.4910692 | 2.12323354 | 1.03127211 | 14.173318 | 0.00083619 | 0.00568253 | NA | NA | NA | NA | NA |
| YNCG0015C | 25.5048131 | 0.25609863 | 0.76417125 | 0.89069905 | 0.64060033 | 0.77561669 | NA | NA | NA | NA | NA |
| YNCG0017W | 23.2246281 | -0.047388 | 0.79963202 | 1.5687622 | 0.45640208 | 0.62207609 | NA | NA | NA | NA | NA |
| YNCG0018C | 7.07812754 | 0.22848519 | 1.05955638 | 0.14385902 | 0.93059649 | 0.95636381 | NA | NA | NA | NA | NA |
| YNCG0024W | 9.72516321 | -1.2542969 | 0.93235837 | 11.5323541 | 0.00313171 | 0.01578119 | NA | NA | NA | NA | NA |
| YNCG0025C | 7.89994494 | 0.48926888 | 1.22163257 | 8.54605347 | 0.01393953 | 0.05066533 | NA | NA | NA | NA | NA |
| YNCG0026W | 45.9295051 | 1.23812735 | 0.77657868 | 2.63102091 | 0.26833731 | 0.44113375 | NA | NA | NA | NA | NA |
| YNCG0037W | 42.4356806 | -1.1064008 | 0.7135716 | 3.72032187 | 0.15564758 | 0.30083183 | NA | NA | NA | NA | NA |
| YNCG0038C | 8.39934761 | 3.07726015 | 1.33974309 | 7.73095988 | 0.02095286 | 0.06842475 | NA | NA | NA | NA | NA |
| YNCH0003C | 11.2755452 | -1.6614399 | 0.9254414 | 12.419671 | 0.00200957 | 0.01119865 | NA | NA | NA | NA | NA |
| YNCH0008W | 28.8518657 | -0.6049291 | 0.92017361 | 4.43930891 | 0.10864664 | 0.23264204 | NA | NA | NA | NA | NA |
| YNCH0009C | 10.9534641 | 1.1077484 | 0.93682691 | 2.68664398 | 0.26097726 | 0.43235173 | NA | NA | NA | NA | NA |
| YNCH0010C | 10.024353 | 0.57435781 | 0.98814826 | 0.53563979 | 0.76504556 | 0.85695759 | NA | NA | NA | NA | NA |
| YNCH0011W | 587.822998 | 2.54334933 | 0.48146475 | 37.2830838 | 8.02E-09 | 2.48E-07 | NA | NA | NA | NA | NA |
| YNCH0012W | 34.5506099 | 0.39536439 | 0.80585813 | 0.74418649 | 0.68928997 | 0.81068229 | NA | NA | NA | NA | NA |
| YNCH0014W | 111.011961 | 0.28374499 | 0.53420799 | 0.45521755 | 0.79643579 | 0.87685172 | NA | NA | NA | NA | NA |
| YNCI0001W | 49.6082987 | 0.488109 | 0.65134151 | 0.68154361 | 0.71122119 | 0.82438173 | NA | NA | NA | NA | NA |
| YNCI0003C | 5.97890307 | 0 | 2.29775058 | 24.0921757 | 5.87E-06 | 8.19E-05 | NA | NA | NA | NA | NA |
| YNCI0006W | 50.0742922 | 0.18554952 | 0.62171687 | 2.40270227 | 0.30078753 | 0.47389383 | NA | NA | NA | NA | NA |
| YNCI0012 | 18.7474564 | -0.6338105 | 0.77853978 | 0.68742949 | 0.70913118 | 0.82408359 | NA | NA | NA | NA | NA |
| YNCI0013W | 332.000789 | -0.4023651 | 0.57622155 | 2.79855419 | 0.24677529 | 0.41766412 | NA | NA | NA | NA | NA |
| YNCJ0004C | 10.4359128 | 1.49980346 | 1.04118506 | 4.12323797 | 0.12724779 | 0.26090922 | NA | NA | NA | NA | NA |
| YNCJ0006W | 6.73079234 | -0.2467131 | 1.2555705 | 2.99457915 | 0.22373576 | 0.38936881 | NA | NA | NA | NA | NA |
| YNCJ0007C | 27.55903 | -0.2570805 | 0.76104346 | 1.84760605 | 0.39700634 | 0.56785755 | NA | NA | NA | NA | NA |
| YNCJ0009C | 17.7194969 | 2.06951663 | 0.96288827 | 13.7479762 | 0.00103434 | 0.00679008 | NA | NA | NA | NA | NA |
| YNCJ0028C | 74.1698523 | -0.2384747 | 0.60525446 | 1.71289754 | 0.4246675 | 0.59492495 | NA | NA | NA | NA | NA |
| YNCJ0030W | 400.94239 | -0.4495183 | 0.4770603 | 0.94450726 | 0.62359533 | 0.76318262 | NA | NA | NA | NA | NA |
| YNCK0004W | 6.04427312 | -0.7684201 | 1.57842555 | 9.733855 | 0.00769698 | 0.03215404 | NA | NA | NA | NA | NA |
| YNCK0005C | 20.9076363 | 7.07865836 | 1.65750966 | 26.3837513 | 1.87E-06 | 3.09E-05 | NA | NA | NA | NA | NA |
| YNCK0010W | 35.6133643 | 5.23121245 | 1.63993317 | 60.1010224 | 8.90E-14 | 7.47E-12 | NA | NA | NA | NA | NA |
| YNCK0013W | 82.4758454 | 1.16736924 | 0.66066977 | 6.17336368 | 0.04565319 | 0.12191465 | NA | NA | NA | NA | NA |
| YNCK0015C | 134.642469 | -0.2652717 | 0.55152439 | 0.65360878 | 0.72122481 | 0.83085787 | NA | NA | NA | NA | NA |

| | | | | | | | | | | | |
|---|---|---|---|---|---|---|---|---|---|---|---|
| YNCK0017W | 7.78558252 | -1.2952063 | 1.0273367 | 3.42409234 | 0.18049609 | 0.33506648 | NA | NA | NA | NA | NA |
| YNCK0020C | 463.463479 | 4.32762839 | 0.59387326 | 42.4281448 | 6.12E-10 | 2.32E-08 | NA | NA | NA | NA | NA |
| YNCL0018W | 1392.084 | -2.7421074 | 0.64057029 | 29.5378476 | 3.85E-07 | 7.70E-06 | NA | NA | NA | NA | NA |
| YNCL0032C | 21.0259403 | 4.00338568 | 1.47786224 | 28.8698258 | 5.38E-07 | 1.04E-05 | NA | NA | NA | NA | NA |
| YNCL0036C | 16.7683428 | 0.51681838 | 1.01810033 | 0.40162603 | 0.81806538 | 0.89067292 | NA | NA | NA | NA | NA |
| YNCL0039W | 5.82013184 | -2.11768 | 1.56225445 | 6.94821515 | 0.03098948 | 0.09168578 | NA | NA | NA | NA | NA |
| YNCL0040W | 14.8728956 | -0.42737 | 0.89193169 | 0.7743864 | 0.67895991 | 0.8022812 | NA | NA | NA | NA | NA |
| YNCL0041C | 36.5714911 | 3.13593008 | 0.96088377 | 16.1881048 | 0.00030535 | 0.0024866 | NA | NA | NA | NA | NA |
| YNCL0044W | 8.36776444 | -0.7469352 | 1.45526661 | 16.2588412 | 0.00029474 | 0.0024162 | NA | NA | NA | NA | NA |
| YNCL0048W | 11.1850325 | 2.07279026 | 1.03520262 | 6.94321125 | 0.03106711 | 0.09182162 | NA | NA | NA | NA | NA |
| YNCM0001W | 1223.47622 | -0.362952 | 0.52054199 | 0.84789762 | 0.65445738 | 0.78442961 | NA | NA | NA | NA | NA |
| YNCM0002C | 32.190886 | -1.2897185 | 0.69227536 | 4.05612524 | 0.13159021 | 0.26686772 | NA | NA | NA | NA | NA |
| YNCM0003W | 11.6359297 | 2.31439827 | 1.06787758 | 7.20945311 | 0.02719488 | 0.08256862 | NA | NA | NA | NA | NA |
| YNCM0004C | 50.4566467 | 3.9131185 | 1.05433371 | 21.2235051 | 2.46E-05 | 0.00028516 | NA | NA | NA | NA | NA |
| YNCM0008C | 22.0944308 | 0.25421155 | 0.71239529 | 0.42442982 | 0.80879086 | 0.8854035 | NA | NA | NA | NA | NA |
| YNCM0011W | 10.5055136 | -1.9486408 | 1.03239072 | 3.90809521 | 0.14169937 | 0.28087618 | NA | NA | NA | NA | NA |
| YNCM0012W | 12.8602663 | 1.21515543 | 1.01850223 | 1.4140128 | 0.49311819 | 0.65425168 | NA | NA | NA | NA | NA |
| YNCM0019W | 24.9135411 | 0.64415449 | 0.77079422 | 2.82576531 | 0.24344052 | 0.41310685 | NA | NA | NA | NA | NA |
| YNCM0030W | 27.2305026 | 2.16801807 | 0.95574132 | 9.04398076 | 0.01086737 | 0.04192011 | NA | NA | NA | NA | NA |
| YNCM0031W | 260.606947 | 0.9554598 | 0.59615782 | 2.58324754 | 0.27482417 | 0.44836065 | NA | NA | NA | NA | NA |
| YNCM0032C | 80.8060757 | -0.4063029 | 0.58252176 | 3.88379059 | 0.14343185 | 0.28353376 | NA | NA | NA | NA | NA |
| YNCM0033C | 15.8194089 | 3.36651904 | 0.9802138 | 12.2111726 | 0.00223037 | 0.01215995 | NA | NA | NA | NA | NA |
| YNCM0034C | 124.942867 | -0.0925902 | 0.58748258 | 18.7538817 | 8.47E-05 | 0.00081555 | NA | NA | NA | NA | NA |
| YNCN0001W | 76.8726536 | 0.46831564 | 0.5726625 | 4.60859081 | 0.09982912 | 0.21918634 | NA | NA | NA | NA | NA |
| YNCN0002C | 14.0832359 | 1.34419378 | 1.01479461 | 4.98484958 | 0.08270917 | 0.19079127 | NA | NA | NA | NA | NA |
| YNCN0005C | 14.2686022 | -0.8059741 | 1.0063777 | 1.09306821 | 0.57895293 | 0.7279343 | NA | NA | NA | NA | NA |
| YNCN0008C | 26.7858364 | 0.08798381 | 0.84114948 | 0.20708138 | 0.90163933 | 0.93824416 | NA | NA | NA | NA | NA |
| YNCN0013W | 23.5620445 | -0.5712599 | 0.74680963 | 1.29609627 | 0.52306573 | 0.67871828 | NA | NA | NA | NA | NA |
| YNCN0018W | 14.9877506 | -0.4814482 | 0.81072914 | 4.3153395 | 0.11559417 | 0.242911 | NA | NA | NA | NA | NA |
| YNCN0019C | 15.8072341 | 1.50960579 | 1.41337149 | 30.5163193 | 2.36E-07 | 5.05E-06 | NA | NA | NA | NA | NA |
| YNCO0003C | 87.3822961 | -1.3758676 | 0.60173657 | 9.7825461 | 0.00751185 | 0.03164938 | NA | NA | NA | NA | NA |
| YNCO0007W | 10.639455 | -0.3750696 | 0.95819103 | 0.50578351 | 0.77655193 | 0.8639961 | NA | NA | NA | NA | NA |
| YNCO0014C | 727.000133 | 1.06521199 | 0.61915818 | 3.93131864 | 0.14006351 | 0.27867477 | NA | NA | NA | NA | NA |

| | | | | | | | | | | | |
|---|---|---|---|---|---|---|---|---|---|---|---|
| YNCO0015C | 172.896618 | 2.06935169 | 0.61414172 | 12.8639649 | 0.00160926 | 0.00961569 | NA | NA | NA | NA | NA |
| YNCO0016W | 32.4083216 | 0.32874001 | 0.67520265 | 0.967192 | 0.61656224 | 0.75817303 | NA | NA | NA | NA | NA |
| YNCO0022C | 76.6236034 | 0.20407419 | 0.56157401 | 4.77128852 | 0.09202967 | 0.20645168 | NA | NA | NA | NA | NA |
| YNCO0024C | 34.3450664 | 0.27442515 | 0.62413954 | 6.27077213 | 0.04348296 | 0.1181447 | NA | NA | NA | NA | NA |
| YNCO0025W | 126.379561 | 2.15508014 | 0.58152414 | 43.2647016 | 4.03E-10 | 1.62E-08 | NA | NA | NA | NA | NA |
| YNCO0026W | 24.7348683 | -1.1940816 | 0.77328762 | 2.44801694 | 0.29404911 | 0.46854198 | NA | NA | NA | NA | NA |
| YNCO0027C | 123.198071 | -0.7963247 | 0.53302188 | 8.90262769 | 0.01166323 | 0.04427675 | NA | NA | NA | NA | NA |
| YNCO0028W | 16.2343628 | -2.0994343 | 0.82567264 | 16.4191919 | 0.00027203 | 0.00224367 | NA | NA | NA | NA | NA |
| YNCO0029C | 350.558512 | 4.96472794 | 0.95266892 | 20.8070685 | 3.03E-05 | 0.00034284 | NA | NA | NA | NA | NA |
| YNCP0001C | 30.0954422 | -1.3932674 | 0.77767185 | 3.20985334 | 0.20090429 | 0.36006531 | NA | NA | NA | NA | NA |
| YNCP0002W | 1571.57567 | 1.01139678 | 0.52803348 | 9.29628236 | 0.00957939 | 0.03819881 | NA | NA | NA | NA | NA |
| YNCP0005C | 31.2905589 | -0.576296 | 0.7690153 | 7.60556033 | 0.02230866 | 0.07187933 | NA | NA | NA | NA | NA |
| YNCP0007C | 10.3773681 | 1.31494465 | 0.99812225 | 1.70517083 | 0.42631131 | 0.59587092 | NA | NA | NA | NA | NA |
| YNCP0013C | 10.4571387 | -0.0944452 | 1.05902413 | 5.6101777 | 0.0605014 | 0.15079771 | NA | NA | NA | NA | NA |
| YNCP0014C | 26.3667168 | 1.13108262 | 0.75893091 | 3.62566143 | 0.16319153 | 0.31184125 | NA | NA | NA | NA | NA |
| YNCP0015C | 22.1066181 | -0.6953917 | 0.75757466 | 1.19674074 | 0.54970673 | 0.70152126 | NA | NA | NA | NA | NA |
| YNCP0018C | 10.9465899 | -2.3046529 | 1.04031958 | 9.43599573 | 0.00893305 | 0.03616947 | NA | NA | NA | NA | NA |
| YNCP0021W | 30.116723 | -0.3849621 | 0.71656876 | 0.36732797 | 0.83221539 | 0.90069191 | NA | NA | NA | NA | NA |
| YNL001W | 7.25780802 | 0.42473882 | 1.04132221 | 0.37966906 | 0.82709598 | 0.89645933 | DOM34 | XIV | 627456 | 628616 | 1 |
| YNL002C | 45.455493 | -0.4403661 | 0.64815268 | 1.11485712 | 0.57267978 | 0.72277589 | RLP7 | XIV | 626174 | 627142 | -1 |
| YNL003C | 111.762781 | -0.7407743 | 0.55806961 | 1.77064369 | 0.41258136 | 0.58426143 | PET8 | XIV | 624975 | 625829 | -1 |
| YNL004W | 890.717428 | 1.61614562 | 0.66156368 | 8.6709681 | 0.01309553 | 0.04820096 | HRB1 | XIV | 622915 | 624621 | 1 |
| YNL005C | 136.723965 | 0.23041135 | 0.56632753 | 0.83735655 | 0.65791583 | 0.78607772 | MRP7 | XIV | 621313 | 622428 | -1 |
| YNL006W | 173.062834 | -1.0159996 | 0.55758387 | 4.61609088 | 0.09945545 | 0.21864382 | LST8 | XIV | 620067 | 620978 | 1 |
| YNL007C | 77.5664949 | -0.361186 | 0.66841707 | 0.61026608 | 0.73702531 | 0.83986943 | SIS1 | XIV | 618507 | 619565 | -1 |
| YNL008C | 683.128962 | -0.4348985 | 0.46927459 | 2.05971426 | 0.35705797 | 0.53036574 | ASI3 | XIV | 616211 | 618241 | -1 |
| YNL009W | 126.382162 | -0.3122195 | 0.52249515 | 0.76158009 | 0.68332134 | 0.80595449 | IDP3 | XIV | 614821 | 616083 | 1 |
| YNL010W | 10.9832782 | -0.2619962 | 0.99607003 | 0.32008919 | 0.85210579 | 0.91195795 | NA | XIV | 613635 | 614360 | 1 |
| YNL011C | 51.2603742 | 0.33258837 | 0.6179795 | 0.55338021 | 0.75828945 | 0.85403579 | NA | XIV | 611836 | 613170 | -1 |
| YNL012W | 1392.68163 | 0.20050059 | 0.47040643 | 7.47514509 | 0.02381184 | 0.07531616 | SPO1 | XIV | 609686 | 611665 | 1 |
| YNL014W | 66.9920647 | 0.59632587 | 0.58583791 | 2.10389503 | 0.3492569 | 0.52293528 | HEF3 | XIV | 606319 | 609453 | 1 |
| YNL015W | 28.5598836 | 1.0752978 | 0.69943541 | 3.56750512 | 0.16800651 | 0.31779081 | PBI2 | XIV | 605384 | 605611 | 1 |
| YNL016W | 158.658512 | 1.17818857 | 0.60950842 | 10.3765967 | 0.0055815 | 0.02501305 | PUB1 | XIV | 602907 | 604268 | 1 |

| | | | | | | | | | | | |
|---|---|---|---|---|---|---|---|---|---|---|---|
| YNL018C | 137.786783 | 1.09375145 | 0.56125766 | 4.49430725 | 0.10569966 | 0.22827341 | NA | XIV | 599936 | 601774 | -1 |
| YNL019C | 276.146147 | 1.56771885 | 0.53638856 | 17.3587795 | 0.00017005 | 0.0014784 | NA | XIV | 598376 | 599230 | -1 |
| YNL020C | 1420.64948 | -1.1190944 | 0.53519609 | 13.4946075 | 0.00117404 | 0.00750302 | ARK1 | XIV | 595623 | 597539 | -1 |
| YNL021W | 36.4262932 | 0.46879744 | 0.61659263 | 1.57264801 | 0.4555162 | 0.62135279 | HDA1 | XIV | 593227 | 595347 | 1 |
| YNL022C | 9.53352065 | -0.0344733 | 1.00492527 | 0.09991861 | 0.95126814 | 0.96614462 | RCM1 | XIV | 591427 | 592899 | -1 |
| YNL023C | 61.9186462 | -0.3249885 | 0.70733098 | 1.48944503 | 0.47486605 | 0.6373376 | FAP1 | XIV | 588263 | 591160 | -1 |
| YNL024C | 184.694133 | -0.7251412 | 0.60842657 | 7.69873909 | 0.02129316 | 0.06934048 | EFM6 | XIV | 587107 | 587847 | -1 |
| YNL024C-A | 14.3848552 | -1.2417878 | 0.87644105 | 3.89668496 | 0.14251009 | 0.28204189 | KSH1 | XIV | 586602 | 586820 | -1 |
| YNL025C | 8.12756857 | 0.42762616 | 1.06050062 | 0.41543972 | 0.8124346 | 0.88731158 | SSN8 | XIV | 584320 | 585291 | -1 |
| YNL026W | 91.4750315 | -0.6342384 | 0.56918329 | 1.3013873 | 0.52168378 | 0.67740505 | SAM50 | XIV | 581920 | 583374 | 1 |
| YNL027W | 15.7901544 | 0.73307403 | 0.85448601 | 1.33001602 | 0.51426941 | 0.67150244 | CRZ1 | XIV | 579580 | 581616 | 1 |
| YNL029C | 586.255355 | 2.48152703 | 1.80772378 | 3.86592186 | 0.14471906 | 0.28539624 | KTR5 | XIV | 577205 | 578773 | -1 |
| YNL030W | 132.707423 | -1.1941091 | 0.88161659 | 7.30360603 | 0.02594431 | 0.07981804 | HHF2 | XIV | 576727 | 577038 | 1 |
| YNL031C | 278.397464 | -0.9194468 | 0.49635305 | 15.3335618 | 0.00046812 | 0.00348384 | HHT2 | XIV | 575640 | 576050 | -1 |
| YNL032W | 28.5681931 | 0.6012924 | 0.72468955 | 6.9504792 | 0.03095442 | 0.09162887 | SIW14 | XIV | 574506 | 575351 | 1 |
| YNL033W | 92.8457098 | -0.2122462 | 0.59414102 | 3.69072033 | 0.15796842 | 0.30416931 | NA | XIV | 572999 | 573853 | 1 |
| YNL034W | 77.1518942 | 0.01134635 | 0.56013091 | 0.14256612 | 0.93119827 | 0.95681242 | NA | XIV | 570477 | 572315 | 1 |
| YNL035C | 48.1012349 | -0.8656653 | 0.61598991 | 2.25072135 | 0.32453539 | 0.49763242 | NA | XIV | 568522 | 569691 | -1 |
| YNL036W | 50.607255 | 2.54907541 | 0.75870306 | 12.8940185 | 0.00158526 | 0.00951154 | NCE103 | XIV | 559814 | 560479 | 1 |
| YNL037C | 67.2585874 | 0.73358689 | 0.60559912 | 15.5975706 | 0.00041023 | 0.00313357 | IDH1 | XIV | 557920 | 559002 | -1 |
| YNL038W | 44.6053403 | 3.02050309 | 0.94088599 | 15.2031916 | 0.00049965 | 0.00367131 | GPI15 | XIV | 557020 | 557783 | 1 |
| YNL039W | 51.8690704 | -0.3159261 | 0.57554354 | 0.89094646 | 0.64052109 | 0.77561669 | BDP1 | XIV | 555048 | 556832 | 1 |
| YNL040W | 13.6442109 | 0.17244956 | 0.90098315 | 0.74574462 | 0.68875318 | 0.81049746 | NA | XIV | 553380 | 554750 | 1 |
| YNL041C | 17.2744245 | 0.33257318 | 0.78855034 | 0.88663657 | 0.64190287 | 0.77608641 | COG6 | XIV | 549468 | 551987 | -1 |
| YNL042W | 125.767646 | 1.64490313 | 0.55217754 | 32.2712568 | 9.83E-08 | 2.40E-06 | BOP3 | XIV | 548100 | 549290 | 1 |
| YNL042W-B | 13.6064407 | -0.0160548 | 1.06993702 | 0.34814523 | 0.84023588 | 0.90645906 | NA | XIV | 547113 | 547370 | 1 |
| YNL044W | 77.7611086 | -0.8335007 | 0.6360344 | 2.88641632 | 0.23616887 | 0.40406514 | YIP3 | XIV | 545268 | 545877 | 1 |
| YNL045W | 715.326555 | 3.70709064 | 0.5194682 | 55.8888897 | 7.31E-13 | 5.29E-11 | LAP2 | XIV | 542963 | 544978 | 1 |
| YNL046W | 285.830051 | 0.71373608 | 0.49469767 | 3.54574225 | 0.16984464 | 0.32001317 | NA | XIV | 542304 | 542822 | 1 |
| YNL047C | 175.311761 | 1.53516427 | 0.72695242 | 4.45839767 | 0.10761461 | 0.23077356 | SLM2 | XIV | 539911 | 541881 | -1 |
| YNL048W | 33.54787 | 0.07351895 | 0.68727274 | 2.47501114 | 0.29010697 | 0.46388272 | ALG11 | XIV | 538173 | 539819 | 1 |
| YNL049C | 69.9526782 | -1.2044156 | 0.64898736 | 7.1193895 | 0.02844751 | 0.08543105 | SFB2 | XIV | 535281 | 537911 | -1 |
| YNL050C | 18.4094928 | 0.21382884 | 0.76767877 | 7.09599358 | 0.02878224 | 0.08625733 | NA | XIV | 534079 | 534982 | -1 |

| | | | | | | | | | | | |
|---|---|---|---|---|---|---|---|---|---|---|---|
| YNL051W | 26.1701317 | -0.3929661 | 0.76046896 | 6.43955116 | 0.03996403 | 0.11135309 | COG5 | XIV | 532659 | 533870 | 1 |
| YNL052W | 41.9136576 | -0.3613544 | 0.65502639 | 0.38349151 | 0.82551672 | 0.8954181 | COX5A | XIV | 531725 | 532186 | 1 |
| YNL053W | 615.515419 | -2.6374126 | 0.51303578 | 59.9851213 | 9.43E-14 | 7.80E-12 | MSG5 | XIV | 529942 | 531411 | 1 |
| YNL054W | 102.289492 | 0.78327965 | 0.54373684 | 4.83563205 | 0.08911603 | 0.2024252 | VAC7 | XIV | 526086 | 529583 | 1 |
| YNL054W-B | 18.947988 | -0.0014287 | 0.79450559 | 0.16378276 | 0.92137203 | 0.95035295 | NA | XIV | 519456 | 524706 | 1 |
| YNL055C | 261.220122 | 1.48191735 | 0.56738528 | 9.52578014 | 0.00854089 | 0.03489891 | POR1 | XIV | 517994 | 518845 | -1 |
| YNL056W | 36.6365605 | 0.29486178 | 0.61505431 | 1.56463602 | 0.45734465 | 0.62262533 | OCA2 | XIV | 517249 | 517842 | 1 |
| YNL058C | 355.288961 | 0.28191548 | 0.48144348 | 0.59907503 | 0.74116092 | 0.84242342 | NA | XIV | 515763 | 516713 | -1 |
| YNL059C | 98.7137308 | 0.41613651 | 0.55346073 | 7.06131431 | 0.02928566 | 0.08719023 | ARP5 | XIV | 512668 | 514935 | -1 |
| YNL061W | 35.3224471 | -0.6016532 | 0.65325385 | 12.5413696 | 0.00189093 | 0.01073788 | NOP2 | XIV | 510540 | 512396 | 1 |
| YNL062C | 65.9047539 | 0.82218395 | 0.64043708 | 1.63050581 | 0.4425274 | 0.61121985 | GCD10 | XIV | 508776 | 510212 | -1 |
| YNL063W | 270.248997 | -0.6183404 | 0.46714913 | 9.42689763 | 0.00897378 | 0.03630899 | MTQ1 | XIV | 507758 | 508702 | 1 |
| YNL064C | 103.463994 | -0.3004418 | 0.74003526 | 0.74075724 | 0.69047285 | 0.81157893 | YDJ1 | XIV | 505868 | 507097 | -1 |
| YNL065W | 49.0477664 | -0.6350086 | 0.58811058 | 1.61764083 | 0.44538312 | 0.61297083 | AQR1 | XIV | 503724 | 505484 | 1 |
| YNL066W | 212.457049 | 0.68297507 | 0.49877699 | 5.00540776 | 0.08186335 | 0.18936828 | SUN4 | XIV | 501515 | 502777 | 1 |
| YNL067W | 45.4721521 | 0.01919456 | 0.61433703 | 5.1068029 | 0.07781653 | 0.18263384 | RPL9B | XIV | 499681 | 500256 | 1 |
| YNL068C | 21.8112389 | -0.9876553 | 0.8901173 | 5.25470404 | 0.07226958 | 0.17276666 | FKH2 | XIV | 495701 | 498289 | -1 |
| YNL069C | 85.5661003 | -0.4995816 | 0.63664659 | 8.08302977 | 0.01757083 | 0.05999792 | RPL16B | XIV | 493956 | 495001 | -1 |
| YNL070W | 42.11087 | -1.4642431 | 0.68135896 | 13.2143386 | 0.00135065 | 0.00839084 | TOM7 | XIV | 493366 | 493548 | 1 |
| YNL071W | 40.26343 | -0.5552858 | 0.73175567 | 1.38553636 | 0.50018954 | 0.65970328 | LAT1 | XIV | 491523 | 492971 | 1 |
| YNL072W | 166.894019 | -0.7933347 | 0.62022268 | 1.97157013 | 0.37314617 | 0.54640641 | RNH201 | XIV | 490317 | 491240 | 1 |
| YNL073W | 150.43352 | -1.0984118 | 0.6126145 | 13.3338672 | 0.00127229 | 0.00800643 | MSK1 | XIV | 488386 | 490116 | 1 |
| YNL074C | 415.812097 | 1.94046422 | 0.48563026 | 42.5069732 | 5.88E-10 | 2.24E-08 | MLF3 | XIV | 486766 | 488124 | -1 |
| YNL075W | 59.7274771 | 0.44486463 | 0.71942661 | 1.55931797 | 0.45856236 | 0.62328904 | IMP4 | XIV | 485607 | 486479 | 1 |
| YNL076W | 47.2650051 | -0.9687505 | 0.58950802 | 8.64844587 | 0.01324384 | 0.04862512 | MKS1 | XIV | 483556 | 485310 | 1 |
| YNL077W | 92.9081468 | 1.61297561 | 0.55565265 | 11.8344444 | 0.00269267 | 0.01402028 | APJ1 | XIV | 481391 | 482977 | 1 |
| YNL078W | 68.5134103 | -0.7356663 | 0.61519183 | 1.84518856 | 0.39748651 | 0.56811822 | NIS1 | XIV | 479768 | 480991 | 1 |
| YNL079C | 495.02206 | 1.0885478 | 0.50311348 | 6.06067987 | 0.04829922 | 0.12713301 | TPM1 | XIV | 478566 | 479165 | -1 |
| YNL080C | 75.9093976 | 0.12211235 | 0.60300795 | 26.725215 | 1.57E-06 | 2.63E-05 | EOS1 | XIV | 476932 | 478032 | -1 |
| YNL081C | 13.6338551 | 0.31559827 | 0.88021917 | 0.37804202 | 0.82776912 | 0.897021 | SWS2 | XIV | 476188 | 476619 | -1 |
| YNL082W | 93.7699191 | 2.68632654 | 0.98181652 | 7.8187896 | 0.02005263 | 0.06615655 | PMS1 | XIV | 473391 | 476012 | 1 |
| YNL083W | 32.1224337 | 0.64902432 | 0.74327099 | 14.0779487 | 0.00087703 | 0.00593222 | NA | XIV | 471377 | 472861 | 1 |
| YNL084C | 39.2845386 | -0.4434959 | 0.77927668 | 4.84106223 | 0.0888744 | 0.20203486 | END3 | XIV | 470053 | 471102 | -1 |

| | | | | | | | | | | | |
|---|---|---|---|---|---|---|---|---|---|---|---|
| YNL085W | 59.9361002 | -0.1730113 | 0.91791494 | 0.14910096 | 0.92816062 | 0.95498585 | MKT1 | XIV | 467131 | 469623 | 1 |
| YNL086W | 106.565075 | -0.080746 | 0.58363953 | 2.93499614 | 0.23050146 | 0.39754218 | SNN1 | XIV | 466334 | 466642 | 1 |
| YNL087W | 431.10816 | 0.20796263 | 0.52138027 | 20.8314609 | 3.00E-05 | 0.00033944 | TCB2 | XIV | 462411 | 465947 | 1 |
| YNL088W | 54.1902413 | -0.8553041 | 0.57596742 | 2.32265586 | 0.31307017 | 0.48516903 | TOP2 | XIV | 457704 | 461990 | 1 |
| YNL090W | 12.2897595 | -0.1155155 | 0.94193426 | 2.39785612 | 0.30151725 | 0.47448396 | RHO2 | XIV | 456565 | 457143 | 1 |
| YNL091W | 394.688535 | 1.7339434 | 0.75729999 | 5.64251933 | 0.05953091 | 0.1490203 | NST1 | XIV | 452408 | 456130 | 1 |
| YNL092W | 101.437923 | 0.40097595 | 0.52931158 | 3.93884202 | 0.13953762 | 0.27801887 | NA | XIV | 450871 | 452073 | 1 |
| YNL093W | 42.2184248 | 1.33658667 | 0.7401361 | 10.9946479 | 0.00409772 | 0.0195435 | YPT53 | XIV | 449868 | 450530 | 1 |
| YNL094W | 44.4771965 | 0.37765463 | 0.62148695 | 2.18317352 | 0.33568342 | 0.50893088 | APP1 | XIV | 447611 | 449374 | 1 |
| YNL095C | 994.545445 | -0.0716929 | 0.50702695 | 0.89251773 | 0.64001807 | 0.77557652 | NA | XIV | 444912 | 446840 | -1 |
| YNL096C | 34.0681041 | -0.3767876 | 0.70165426 | 5.14292483 | 0.0764237 | 0.18002165 | RPS7B | XIV | 443398 | 444315 | -1 |
| YNL097C | 34.9026636 | 0.01506323 | 0.6776259 | 0.77193933 | 0.67979115 | 0.80293569 | PHO23 | XIV | 441366 | 442358 | -1 |
| YNL097C-B | 79.995857 | -0.4125764 | 0.60271781 | 11.7030494 | 0.00287551 | 0.01474687 | NA | XIV | 440797 | 440919 | -1 |
| YNL098C | 113.157832 | -0.4725782 | 0.54526587 | 8.10748173 | 0.01735732 | 0.05959198 | RAS2 | XIV | 439602 | 440570 | -1 |
| YNL099C | 65.1304785 | 0.30507836 | 0.58099166 | 0.91917889 | 0.63154288 | 0.76880708 | OCA1 | XIV | 438567 | 439283 | -1 |
| YNL100W | 93.552157 | -0.2143684 | 0.5987037 | 4.80174645 | 0.09063877 | 0.20427392 | MIC27 | XIV | 437613 | 438317 | 1 |
| YNL101W | 55.7589256 | -0.8435026 | 0.56827181 | 9.45783893 | 0.00883601 | 0.03582669 | AVT4 | XIV | 434999 | 437140 | 1 |
| YNL102W | 25.0197702 | -0.7419637 | 0.73817982 | 4.30465263 | 0.11621349 | 0.24370198 | POL1 | XIV | 430087 | 434493 | 1 |
| YNL103W | 55.7445089 | -0.1172325 | 0.59643343 | 2.87238664 | 0.23783139 | 0.40584843 | NA | XIV | 427735 | 429753 | 1 |
| YNL104C | 122.894442 | -0.7155811 | 0.65749873 | 1.34892316 | 0.50943063 | 0.66733108 | LEU4 | XIV | 424895 | 426754 | -1 |
| YNL106C | 114.247228 | 1.44909252 | 0.74472579 | 4.8440894 | 0.08873999 | 0.20185727 | INP52 | XIV | 420944 | 424495 | -1 |
| YNL107W | 55.3082166 | -0.2099796 | 0.57390986 | 2.49042978 | 0.28787904 | 0.46172289 | YAF9 | XIV | 420098 | 420778 | 1 |
| YNL108C | 76.4481733 | -0.0155879 | 0.54170197 | 0.00803877 | 0.99598868 | 0.99658361 | NA | XIV | 419014 | 419826 | -1 |
| YNL110C | 84.5231469 | -0.9774692 | 0.54264032 | 26.138301 | 2.11E-06 | 3.36E-05 | NOP15 | XIV | 417826 | 418488 | -1 |
| YNL111C | 6.86775019 | -0.2929716 | 1.1196967 | 4.14939552 | 0.12559438 | 0.25832734 | CYB5 | XIV | 416940 | 417302 | -1 |
| YNL112W | 122.087146 | -1.3598302 | 0.64960506 | 7.70595481 | 0.02121647 | 0.06916857 | DBP2 | XIV | 413639 | 416281 | 1 |
| YNL113W | 240.808963 | -0.0754731 | 0.53428064 | 3.28371509 | 0.19362005 | 0.35033128 | RPC19 | XIV | 412771 | 413199 | 1 |
| YNL115C | 74.0206021 | 0.93334143 | 0.68571445 | 6.00307398 | 0.04971061 | 0.13011953 | NA | XIV | 410119 | 412053 | -1 |
| YNL116W | 26.0244842 | 1.1436682 | 0.69726463 | 2.86005822 | 0.23930196 | 0.40763705 | DMA2 | XIV | 408341 | 409909 | 1 |
| YNL117W | 192.221024 | -0.5554801 | 0.52230313 | 1.36580812 | 0.50514788 | 0.66322137 | MLS1 | XIV | 406358 | 408022 | 1 |
| YNL118C | 98.9233882 | 1.03212495 | 0.68371582 | 3.85797586 | 0.14529517 | 0.28623989 | DCP2 | XIV | 402652 | 405564 | -1 |
| YNL119W | 189.391569 | 0.95672404 | 0.66867975 | 8.26445602 | 0.01604709 | 0.05641325 | NCS2 | XIV | 401040 | 402521 | 1 |
| YNL121C | 38.9229377 | -0.1826416 | 0.66686733 | 2.86008842 | 0.23929834 | 0.40763705 | TOM70 | XIV | 398684 | 400537 | -1 |

| | | | | | | | | | | | |
|---|---|---|---|---|---|---|---|---|---|---|---|
| YNL122C | 24.9515894 | 0.55169151 | 0.79283255 | 8.62409263 | 0.01340609 | 0.04903427 | NA | XIV | 398023 | 398370 | -1 |
| YNL123W | 91.5331075 | 1.24737321 | 0.62491125 | 14.6161484 | 0.00067011 | 0.00472009 | NMA111 | XIV | 394685 | 397678 | 1 |
| YNL124W | 213.548022 | 0.33897399 | 0.58418644 | 0.35323248 | 0.83810135 | 0.90469306 | NAF1 | XIV | 392892 | 394370 | 1 |
| YNL125C | 24.9550021 | 0.94363587 | 0.74400539 | 3.37805898 | 0.18469869 | 0.34046654 | ESBP6 | XIV | 390146 | 392167 | -1 |
| YNL126W | 33.32406 | 0.52444133 | 0.74512967 | 2.90270008 | 0.23425382 | 0.40187545 | SPC98 | XIV | 387227 | 389767 | 1 |
| YNL127W | 59.5018119 | -0.0438012 | 0.60169403 | 14.3716634 | 0.00075724 | 0.00521018 | FAR11 | XIV | 383987 | 386848 | 1 |
| YNL128W | 18.6545304 | -0.4578631 | 0.8000679 | 2.51141352 | 0.28487444 | 0.45855519 | TEP1 | XIV | 382359 | 383663 | 1 |
| YNL129W | 54.0429913 | -0.6477899 | 0.59096557 | 1.41803372 | 0.49212779 | 0.6530873 | NRK1 | XIV | 381480 | 382202 | 1 |
| YNL130C | 26.721403 | 0.29044756 | 0.77869565 | 3.06967394 | 0.21549082 | 0.37935295 | CPT1 | XIV | 379558 | 380831 | -1 |
| YNL130C-A | 43.966705 | -1.5406671 | 0.68930405 | 11.0028733 | 0.0040809 | 0.01952763 | DGR1 | XIV | 381245 | 381391 | -1 |
| YNL131W | 24.6389554 | -0.1888263 | 0.69062566 | 6.40125281 | 0.04073668 | 0.11285424 | TOM22 | XIV | 378767 | 379225 | 1 |
| YNL132W | 26.8956583 | -0.7124903 | 0.79153826 | 1.13161926 | 0.56790017 | 0.71824858 | KRE33 | XIV | 375321 | 378491 | 1 |
| YNL133C | 40.059583 | 0.39405589 | 0.61388541 | 1.22541186 | 0.54188259 | 0.6958306 | FYV6 | XIV | 374171 | 374692 | -1 |
| YNL134C | 583.416392 | -0.6169229 | 0.59163106 | 2.15014758 | 0.34127257 | 0.5148432 | NA | XIV | 372451 | 373581 | -1 |
| YNL136W | 696.624233 | 1.51740899 | 0.60874863 | 5.97756824 | 0.05034862 | 0.13131464 | EAF7 | XIV | 370368 | 371645 | 1 |
| YNL137C | 24.0473104 | -0.2856312 | 0.73613476 | 0.34163985 | 0.84297336 | 0.90805874 | NAM9 | XIV | 368595 | 370055 | -1 |
| YNL138W | 68.9820231 | 0.86868485 | 0.58831553 | 7.54001107 | 0.02305194 | 0.0736187 | SRV2 | XIV | 366741 | 368321 | 1 |
| YNL138W-A | 15.4470672 | 0.00901106 | 0.80649352 | 0.25226746 | 0.88149696 | 0.9300006 | YSF3 | XIV | 366033 | 366412 | 1 |
| YNL139C | 74.2041521 | -0.1514602 | 0.60896478 | 24.5971554 | 4.56E-06 | 6.58E-05 | THO2 | XIV | 360924 | 365717 | -1 |
| YNL141W | 795.159069 | 0.06179042 | 0.48588052 | 6.27371126 | 0.04341911 | 0.11809931 | AAH1 | XIV | 359596 | 360639 | 1 |
| YNL142W | 49.3757652 | -2.2525296 | 0.63853529 | 12.3879246 | 0.00204172 | 0.01132334 | MEP2 | XIV | 357453 | 358952 | 1 |
| YNL143C | 185.652425 | -0.2503614 | 0.54319331 | 0.49409776 | 0.78110251 | 0.86705973 | NA | XIV | 356794 | 357186 | -1 |
| YNL144C | 121.66511 | -0.4429299 | 0.54142646 | 2.36465471 | 0.30656442 | 0.47921383 | NA | XIV | 352820 | 355042 | -1 |
| YNL145W | 15.9440573 | -1.6358036 | 0.8099048 | 9.98230154 | 0.00679784 | 0.02932897 | MFA2 | XIV | 352414 | 352530 | 1 |
| YNL146C-A | 313.210433 | 0.39617404 | 0.66797688 | 8.7899673 | 0.01233908 | 0.0461222 | NA | XIV | 351386 | 351580 | -1 |
| YNL147W | 153.439426 | -0.1914806 | 0.59338254 | 0.71761253 | 0.69850966 | 0.81736153 | LSM7 | XIV | 350940 | 351383 | 1 |
| YNL148C | 23.6942946 | -0.6817985 | 0.69706976 | 1.28660785 | 0.52555316 | 0.6809024 | ALF1 | XIV | 349907 | 350671 | -1 |
| YNL149C | 40.3400702 | -0.2609315 | 0.73035596 | 1.00942572 | 0.60367889 | 0.74934389 | PGA2 | XIV | 349367 | 349756 | -1 |
| YNL151C | 32.2450808 | -0.3653682 | 0.68557365 | 1.91872314 | 0.38313741 | 0.55639971 | RPC31 | XIV | 347766 | 348521 | -1 |
| YNL152W | 14.6076563 | -0.7640563 | 0.913939 | 3.98062346 | 0.13665282 | 0.27406298 | INN1 | XIV | 346313 | 347542 | 1 |
| YNL153C | 56.715367 | -1.0331254 | 0.64360762 | 2.6110715 | 0.27102729 | 0.44454618 | GIM3 | XIV | 345669 | 346058 | -1 |
| YNL154C | 1031.05432 | 1.81091964 | 0.4603694 | 62.737617 | 2.38E-14 | 2.15E-12 | YCK2 | XIV | 343636 | 345276 | -1 |
| YNL155W | 15.9200379 | -0.0631319 | 0.83732715 | 0.97717188 | 0.6134933 | 0.75641529 | CUZ1 | XIV | 342516 | 343340 | 1 |

| | | | | | | | | | | | |
|---|---|---|---|---|---|---|---|---|---|---|---|
| YNL156C | 51.3730688 | 0.18465386 | 0.57676525 | 0.12465728 | 0.93957405 | 0.96037943 | NSG2 | XIV | 341069 | 341968 | -1 |
| YNL157W | 57.7125304 | 1.15304894 | 0.61427809 | 5.40464559 | 0.06704959 | 0.16264366 | IGO1 | XIV | 340352 | 340858 | 1 |
| YNL158W | 114.9382 | 1.32022563 | 0.59067149 | 33.5834895 | 5.10E-08 | 1.33E-06 | PGA1 | XIV | 339612 | 340208 | 1 |
| YNL159C | 80.0657402 | -1.5178971 | 0.57920035 | 12.8036988 | 0.00165849 | 0.00977866 | ASI2 | XIV | 338478 | 339347 | -1 |
| YNL160W | 233.930179 | -0.2714709 | 0.48229014 | 10.2800947 | 0.00585741 | 0.02598806 | YGP1 | XIV | 336545 | 337609 | 1 |
| YNL161W | 85.0185231 | 0.85633733 | 0.59742893 | 7.07495574 | 0.02908659 | 0.08680999 | CBK1 | XIV | 332597 | 334867 | 1 |
| YNL162W | 66.7535243 | -0.442074 | 0.71850589 | 14.0743575 | 0.0008786 | 0.00593594 | RPL42A | XIV | 331322 | 332154 | 1 |
| YNL162W-A | 78.7536025 | -0.0969409 | 0.55081568 | 2.36718795 | 0.30617637 | 0.47873648 | NA | XIV | 330329 | 330547 | 1 |
| YNL163C | 38.0530854 | -0.4888247 | 0.68215063 | 0.66930432 | 0.71558695 | 0.8274912 | RIA1 | XIV | 326741 | 330073 | -1 |
| YNL164C | 24.1225233 | 2.33506481 | 1.02310811 | 7.50755807 | 0.02342904 | 0.07454061 | IBD2 | XIV | 325265 | 326320 | -1 |
| YNL165W | 37.9920069 | -0.2844939 | 0.65118725 | 0.25599689 | 0.87985475 | 0.92911891 | NA | XIV | 323832 | 325052 | 1 |
| YNL166C | 151.849359 | 0.53688965 | 0.63400999 | 18.9433808 | 7.70E-05 | 0.00074679 | BNI5 | XIV | 322219 | 323565 | -1 |
| YNL167C | 155.266063 | 0.20964982 | 0.51874105 | 12.0193847 | 0.00245484 | 0.01302799 | SKO1 | XIV | 319416 | 321359 | -1 |
| YNL168C | 97.4701283 | -0.9187087 | 0.60274473 | 2.36258194 | 0.30688231 | 0.47958126 | FMP41 | XIV | 318030 | 318809 | -1 |
| YNL169C | 105.557281 | 0.64844924 | 0.68908593 | 6.66133962 | 0.03576914 | 0.10217233 | PSD1 | XIV | 316169 | 317671 | -1 |
| YNL172W | 1184.90243 | 3.50943636 | 0.47641275 | 78.9683086 | 7.12E-18 | 1.21E-15 | APC1 | XIV | 310636 | 315882 | 1 |
| YNL173C | 23.8126649 | -1.0651876 | 0.71183769 | 5.94080522 | 0.05128266 | 0.13339021 | MDG1 | XIV | 308957 | 310057 | -1 |
| YNL175C | 57.6833051 | -0.4885242 | 0.63857527 | 6.87576509 | 0.03213265 | 0.0942798 | NOP13 | XIV | 307401 | 308612 | -1 |
| YNL176C | 44.7625102 | -0.0137824 | 0.64415779 | 0.47108729 | 0.79014118 | 0.87290926 | TDA7 | XIV | 305070 | 306980 | -1 |
| YNL177C | 14.6632185 | -0.4780104 | 0.89945437 | 2.70472959 | 0.25862793 | 0.43030337 | MRPL22 | XIV | 303686 | 304615 | -1 |
| YNL178W | 100.034115 | -0.1800169 | 0.5481947 | 26.7607682 | 1.55E-06 | 2.59E-05 | RPS3 | XIV | 302680 | 303402 | 1 |
| YNL180C | 184.802711 | 1.86751957 | 0.82958869 | 5.11040941 | 0.07767633 | 0.18237873 | RHO5 | XIV | 299654 | 300649 | -1 |
| YNL181W | 30.7319179 | 0.58422427 | 0.88323245 | 4.2712764 | 0.11816915 | 0.24673616 | PBR1 | XIV | 298335 | 299558 | 1 |
| YNL182C | 13.6025685 | -0.6737423 | 0.81787116 | 4.28548586 | 0.11733257 | 0.24546419 | IPI3 | XIV | 295960 | 297627 | -1 |
| YNL183C | 23.6767746 | -0.5274185 | 0.71375727 | 2.74510867 | 0.25345871 | 0.42549317 | NPR1 | XIV | 293137 | 295509 | -1 |
| YNL185C | 6.2353873 | 0.07797008 | 1.20557911 | 0.42776017 | 0.8074452 | 0.88441988 | MRPL19 | XIV | 292193 | 292669 | -1 |
| YNL186W | 59.8978407 | -0.1829873 | 0.59444659 | 4.11558353 | 0.12773573 | 0.26143156 | UBP10 | XIV | 289498 | 291876 | 1 |
| YNL187W | 66.651575 | -1.9575616 | 0.662175 | 9.33293414 | 0.00940544 | 0.03766079 | SWT21 | XIV | 287994 | 289067 | 1 |
| YNL188W | 74.112174 | -0.2109042 | 0.61399811 | 19.0163206 | 7.42E-05 | 0.00072613 | KAR1 | XIV | 286308 | 287609 | 1 |
| YNL189W | 16.2443157 | -0.1855633 | 0.92022825 | 2.94342805 | 0.22953172 | 0.39623992 | SRP1 | XIV | 284260 | 285888 | 1 |
| YNL190W | 95.6737106 | 0.3930915 | 0.57658724 | 2.88704543 | 0.2360946 | 0.40406514 | NA | XIV | 282395 | 283009 | 1 |
| YNL191W | 32.5600283 | 0.08389253 | 0.73868895 | 16.832576 | 0.00022123 | 0.00185644 | DUG3 | XIV | 280432 | 281505 | 1 |
| YNL192W | 105.75236 | 2.39393743 | 0.68234351 | 12.5435889 | 0.00188884 | 0.01073788 | CHS1 | XIV | 276502 | 279897 | 1 |

| | | | | | | | | | | | |
|---|---|---|---|---|---|---|---|---|---|---|---|
| YNL193W | 200.047279 | 2.09378601 | 0.49258558 | 18.7244955 | 8.59E-05 | 0.00082519 | NA | XIV | 274368 | 276044 | 1 |
| YNL194C | 336.429552 | -1.8774241 | 0.49851554 | 15.413795 | 0.00044971 | 0.00337792 | NA | XIV | 272710 | 273615 | -1 |
| YNL195C | 83.5456041 | -2.1117713 | 0.57770018 | 16.9069798 | 0.00021316 | 0.00180434 | NA | XIV | 271523 | 272308 | -1 |
| YNL196C | 255.317109 | -0.7947412 | 0.56944755 | 2.2550751 | 0.32382969 | 0.49734056 | SLZ1 | XIV | 270275 | 271171 | -1 |
| YNL197C | 36.5653046 | -0.5180385 | 0.62895222 | 4.47807891 | 0.10656081 | 0.22939149 | WHI3 | XIV | 267608 | 269593 | -1 |
| YNL199C | 123.154241 | 1.12649976 | 0.61605989 | 32.6680712 | 8.06E-08 | 2.02E-06 | GCR2 | XIV | 264926 | 266530 | -1 |
| YNL200C | 54.468545 | -0.3116153 | 0.60160439 | 2.63670587 | 0.26757565 | 0.44038176 | NNR1 | XIV | 263713 | 264453 | -1 |
| YNL201C | 63.1730495 | 0.79855382 | 0.64225675 | 18.3623209 | 0.00010296 | 0.00095843 | PSY2 | XIV | 260627 | 263203 | -1 |
| YNL202W | 408.132515 | -0.906276 | 0.52900838 | 7.06082436 | 0.02929284 | 0.08719023 | SPS19 | XIV | 259578 | 260456 | 1 |
| YNL204C | 7.8099756 | -0.7204957 | 1.01806144 | 0.6316781 | 0.7291768 | 0.83668919 | SPS18 | XIV | 258375 | 259277 | -1 |
| YNL206C | 394.769881 | -1.1136446 | 0.46958645 | 5.50024007 | 0.06392019 | 0.15715409 | RTT106 | XIV | 256788 | 258155 | -1 |
| YNL207W | 206.635338 | 0.90909443 | 0.54581699 | 43.2619649 | 4.03E-10 | 1.62E-08 | RIO2 | XIV | 255353 | 256630 | 1 |
| YNL208W | 540.192045 | 0.71678206 | 0.5324164 | 1.8692073 | 0.3927415 | 0.56412138 | NA | XIV | 254418 | 255017 | 1 |
| YNL209W | 139.248198 | -0.6308692 | 0.59918973 | 10.6200701 | 0.00494175 | 0.02269053 | SSB2 | XIV | 252059 | 253900 | 1 |
| YNL210W | 540.133401 | -0.0497506 | 0.56953302 | 0.00785888 | 0.99607827 | 0.99658361 | MER1 | XIV | 250931 | 251743 | 1 |
| YNL212W | 39.2975461 | 0.50016459 | 0.78965184 | 1.13849288 | 0.56595176 | 0.71688048 | VID27 | XIV | 247461 | 249809 | 1 |
| YNL213C | 52.2152757 | 2.03813213 | 0.79737927 | 10.3743424 | 0.00558779 | 0.0250219 | RRG9 | XIV | 246460 | 247104 | -1 |
| YNL215W | 108.784861 | 0.44080264 | 0.67199051 | 1.24534432 | 0.53650888 | 0.69122974 | IES2 | XIV | 244468 | 245430 | 1 |
| YNL216W | 595.612072 | 0.93682673 | 0.49206359 | 30.6985151 | 2.16E-07 | 4.66E-06 | RAP1 | XIV | 241689 | 244172 | 1 |
| YNL217W | 34.9619619 | -0.0814539 | 0.63805606 | 0.03437917 | 0.98295731 | 0.98687755 | NA | XIV | 240331 | 241311 | 1 |
| YNL218W | 14.0170888 | -1.5262743 | 0.85470398 | 3.25183478 | 0.19673111 | 0.35456204 | MGS1 | XIV | 238238 | 240001 | 1 |
| YNL219C | 8.39230777 | -0.4582323 | 1.10161529 | 0.80029607 | 0.67022082 | 0.79536351 | ALG9 | XIV | 235996 | 237663 | -1 |
| YNL220W | 34.7808311 | -0.248068 | 0.67026008 | 1.05440221 | 0.59025472 | 0.73765326 | ADE12 | XIV | 234413 | 235714 | 1 |
| YNL221C | 55.2585846 | 0.34997233 | 0.64418324 | 18.3078154 | 0.00010581 | 0.00098018 | POP1 | XIV | 231068 | 233695 | -1 |
| YNL222W | 24.3789743 | 0.04112325 | 0.70470229 | 0.03108121 | 0.98457953 | 0.98751104 | SSU72 | XIV | 229094 | 229714 | 1 |
| YNL223W | 13.1989844 | 1.37363349 | 1.03352483 | 2.67878102 | 0.26200531 | 0.43337041 | ATG4 | XIV | 227370 | 228854 | 1 |
| YNL224C | 40.383097 | -1.3263997 | 0.64386105 | 4.20402501 | 0.12221023 | 0.25307484 | SQS1 | XIV | 224796 | 227099 | -1 |
| YNL225C | 29.6062206 | 0.06227951 | 0.76032805 | 20.7367585 | 3.14E-05 | 0.00034994 | CNM67 | XIV | 222724 | 224469 | -1 |
| YNL227C | 54.2701139 | 0.36806996 | 0.60369073 | 0.65577074 | 0.7204456 | 0.83045591 | JJJ1 | XIV | 220659 | 222431 | -1 |
| YNL229C | 49.939976 | 1.368312 | 0.66359573 | 4.65830808 | 0.09737809 | 0.21492841 | URE2 | XIV | 219137 | 220201 | -1 |
| YNL230C | 18.5218307 | 1.49887297 | 0.82067849 | 3.3324652 | 0.1889576 | 0.34502193 | ELA1 | XIV | 217523 | 218662 | -1 |
| YNL231C | 24.571091 | -0.1594709 | 0.82843144 | 2.05405247 | 0.3580702 | 0.53120548 | PDR16 | XIV | 215987 | 217042 | -1 |
| YNL232W | 30.6201998 | -0.2101942 | 0.79292787 | 0.10933652 | 0.94679919 | 0.96418144 | CSL4 | XIV | 214923 | 215801 | 1 |

| | | | | | | | | | | | |
|---|---|---|---|---|---|---|---|---|---|---|---|
| YNL233W | 36.0092696 | 0.47200227 | 0.7227391 | 3.7614841 | 0.15247692 | 0.29615611 | BNI4 | XIV | 211922 | 214600 | 1 |
| YNL234W | 10.1462143 | -1.4719307 | 1.08637876 | 2.74905005 | 0.25295972 | 0.42477864 | NA | XIV | 210233 | 211513 | 1 |
| YNL236W | 110.444384 | -0.0241571 | 0.62234744 | 8.28133921 | 0.01591219 | 0.05607523 | SIN4 | XIV | 206930 | 209854 | 1 |
| YNL237W | 1197.98362 | 0.19490954 | 0.54728912 | 0.15462871 | 0.92559884 | 0.95325814 | YTP1 | XIV | 205188 | 206567 | 1 |
| YNL238W | 25.2509688 | -0.5492508 | 0.76111487 | 1.1493231 | 0.56289535 | 0.71410256 | KEX2 | XIV | 202428 | 204872 | 1 |
| YNL239W | 344.374505 | 0.06531308 | 0.48094835 | 2.47665463 | 0.28986867 | 0.46375784 | NA | XIV | 200569 | 201933 | 1 |
| YNL240C | 155.487744 | -1.03722 | 0.56033262 | 15.2355786 | 0.00049163 | 0.00363486 | NAR1 | XIV | 198502 | 199977 | -1 |
| YNL241C | 166.074698 | 0.54133417 | 0.52942823 | 1.13174532 | 0.56786438 | 0.71824858 | ZWF1 | XIV | 196426 | 197943 | -1 |
| YNL242W | 195.193844 | -0.4924437 | 0.52909077 | 0.87441108 | 0.64583867 | 0.77803336 | ATG2 | XIV | 191324 | 196102 | 1 |
| YNL243W | 62.0550639 | 0.67841395 | 0.57778325 | 1.71330264 | 0.42458149 | 0.59492495 | SLA2 | XIV | 188051 | 190957 | 1 |
| YNL244C | 58.3218267 | 0.19430877 | 0.74567142 | 2.86024171 | 0.23928 | 0.40763705 | SUI1 | XIV | 187170 | 187496 | -1 |
| YNL245C | 487.202946 | -0.4891163 | 0.51064385 | 11.9622882 | 0.00252593 | 0.01335631 | CWC25 | XIV | 186346 | 186885 | -1 |
| YNL246W | 2197.14155 | 0.31537983 | 0.45826633 | 1.0868479 | 0.58075636 | 0.72940182 | VPS75 | XIV | 185460 | 186349 | 1 |
| YNL247W | 81.0404292 | -0.1219814 | 0.55881848 | 1.36896077 | 0.50435223 | 0.66262751 | NA | XIV | 182875 | 185178 | 1 |
| YNL248C | 21.8765074 | -0.8264411 | 0.73264889 | 1.35222039 | 0.50859147 | 0.66638258 | RPA49 | XIV | 181361 | 182608 | -1 |
| YNL249C | 385.060768 | -0.1432739 | 0.47129249 | 1.10887591 | 0.57439501 | 0.72409037 | MPA43 | XIV | 179394 | 181022 | -1 |
| YNL250W | 411.788885 | 0.2046572 | 0.46781921 | 4.08696602 | 0.12957661 | 0.26389327 | RAD50 | XIV | 175410 | 179348 | 1 |
| YNL251C | 96.0842937 | -0.427208 | 0.52779574 | 3.38324113 | 0.18422074 | 0.33979289 | NRD1 | XIV | 172588 | 174315 | -1 |
| YNL252C | 47.8379783 | 0.98516368 | 0.80435644 | 5.42874583 | 0.06624648 | 0.16143398 | MRPL17 | XIV | 171441 | 172286 | -1 |
| YNL253W | 32.7777418 | -1.1075874 | 0.7045635 | 3.17054814 | 0.20489163 | 0.36603596 | TEX1 | XIV | 170018 | 171286 | 1 |
| YNL254C | 166.907524 | -0.2724416 | 0.57732391 | 4.82924801 | 0.08940095 | 0.20267482 | RTC4 | XIV | 168043 | 169248 | -1 |
| YNL255C | 55.0902088 | -0.3824436 | 0.65965659 | 6.5969286 | 0.03693985 | 0.10474131 | GIS2 | XIV | 167329 | 167790 | -1 |
| YNL256W | 988.504187 | 0.74109137 | 0.69115182 | 1.28244423 | 0.5266484 | 0.68176637 | FOL1 | XIV | 164623 | 167097 | 1 |
| YNL257C | 74.709436 | -1.116008 | 0.57050063 | 4.40817738 | 0.11035104 | 0.23522866 | SIP3 | XIV | 160631 | 164320 | -1 |
| YNL258C | 76.3408158 | 0.11350617 | 0.615699 | 19.4116893 | 6.09E-05 | 0.00061032 | DSL1 | XIV | 158109 | 160373 | -1 |
| YNL260C | 107.750961 | -0.7832523 | 0.5768098 | 11.7871172 | 0.00275715 | 0.01424075 | LTO1 | XIV | 156859 | 157455 | -1 |
| YNL261W | 119.827604 | 0.86304552 | 0.61465924 | 2.51717331 | 0.28405521 | 0.45768219 | ORC5 | XIV | 155100 | 156539 | 1 |
| YNL262W | 74.9939808 | -0.3399659 | 0.56932611 | 1.21634957 | 0.54434351 | 0.69728958 | POL2 | XIV | 148212 | 154880 | 1 |
| YNL263C | 15.9385297 | -0.1914637 | 0.8498316 | 2.38288445 | 0.30378282 | 0.47692586 | YIF1 | XIV | 146896 | 147840 | -1 |
| YNL264C | 5.39301556 | 0.36607359 | 1.24885182 | 0.34706169 | 0.84069122 | 0.90645906 | PDR17 | XIV | 145563 | 146615 | -1 |
| YNL265C | 93.5840283 | 0.854359 | 0.66300168 | 5.01595415 | 0.08143281 | 0.18867385 | IST1 | XIV | 144280 | 145281 | -1 |
| YNL267W | 44.9170339 | 0.8753601 | 0.68232408 | 3.53054885 | 0.17113982 | 0.32161621 | PIK1 | XIV | 140878 | 144078 | 1 |
| YNL268W | 196.397832 | 0.64660881 | 0.53114775 | 7.19769378 | 0.02735525 | 0.08296851 | LYP1 | XIV | 138550 | 140385 | 1 |

| | | | | | | | | | | | |
|---|---|---|---|---|---|---|---|---|---|---|---|
| YNL269W | 56.4007457 | -0.1190054 | 0.74745881 | 1.88416498 | 0.3898152 | 0.56101577 | BSC4 | XIV | 137699 | 138094 | 1 |
| YNL270C | 73.1100454 | -0.1848346 | 0.69108421 | 21.8033093 | 1.84E-05 | 0.00021999 | ALP1 | XIV | 135940 | 137661 | -1 |
| YNL271C | 64.6654796 | 0.22388708 | 0.61006053 | 0.23893083 | 0.8873947 | 0.93215081 | BNI1 | XIV | 129522 | 135383 | -1 |
| YNL272C | 21.7670439 | -0.1134959 | 0.71598155 | 0.03189003 | 0.98418144 | 0.98751104 | SEC2 | XIV | 126805 | 129084 | -1 |
| YNL273W | 398.275851 | 0.20315775 | 0.49389651 | 2.39725054 | 0.30160856 | 0.47448396 | TOF1 | XIV | 122883 | 126599 | 1 |
| YNL274C | 20.7320622 | 0.32765732 | 0.83794843 | 0.27234951 | 0.87269011 | 0.92527295 | GOR1 | XIV | 121118 | 122170 | -1 |
| YNL275W | 36.2056233 | 0.53749872 | 0.78067587 | 0.48622922 | 0.78418163 | 0.86949041 | BOR1 | XIV | 119268 | 120998 | 1 |
| YNL277W | 2021.596 | -0.2533463 | 0.50426262 | 0.34589753 | 0.84118071 | 0.90680252 | MET2 | XIV | 117349 | 118809 | 1 |
| YNL278W | 48.6309448 | 0.06620646 | 0.63451486 | 0.1846427 | 0.91181209 | 0.9456192 | CAF120 | XIV | 113271 | 116453 | 1 |
| YNL279W | 891.428855 | -1.4692184 | 0.5040093 | 9.57327268 | 0.00834046 | 0.03427345 | PRM1 | XIV | 110917 | 112902 | 1 |
| YNL280C | 529.483013 | 0.56212724 | 0.5847879 | 1.53914283 | 0.46321155 | 0.62634163 | ERG24 | XIV | 109095 | 110411 | -1 |
| YNL281W | 70.7021229 | -0.8199125 | 0.55784659 | 2.79478717 | 0.24724054 | 0.4180849 | HCH1 | XIV | 108467 | 108928 | 1 |
| YNL282W | 24.0004457 | -0.9808994 | 0.7019784 | 2.57393812 | 0.27610638 | 0.45007205 | POP3 | XIV | 107687 | 108274 | 1 |
| YNL283C | 58.7146703 | 0.1730092 | 0.64514859 | 1.91634647 | 0.38359298 | 0.55678199 | WSC2 | XIV | 105184 | 106695 | -1 |
| YNL284C | 23.594413 | -0.0671405 | 0.71607248 | 0.04101004 | 0.97970378 | 0.98440542 | MRPL10 | XIV | 103134 | 104102 | -1 |
| YNL286W | 205.347349 | 1.04417447 | 0.49258756 | 22.0703246 | 1.61E-05 | 0.00019822 | CUS2 | XIV | 95223 | 96080 | 1 |
| YNL287W | 33.4801967 | 0.44343979 | 0.7513813 | 1.40676881 | 0.4949075 | 0.65527428 | SEC21 | XIV | 91994 | 94801 | 1 |
| YNL288W | 54.6704749 | 1.19007358 | 0.59344581 | 5.55559836 | 0.06217519 | 0.15384375 | CAF40 | XIV | 90303 | 91424 | 1 |
| YNL289W | 19.7573996 | -0.3379241 | 0.74445668 | 9.30483506 | 0.00953851 | 0.03806203 | PCL1 | XIV | 87897 | 88736 | 1 |
| YNL290W | 27.589636 | -0.8655836 | 0.73648989 | 5.34832286 | 0.06896464 | 0.16644654 | RFC3 | XIV | 86218 | 87240 | 1 |
| YNL291C | 22.0747864 | -0.1172418 | 0.85804596 | 6.65781712 | 0.03583219 | 0.10225577 | MID1 | XIV | 84169 | 85815 | -1 |
| YNL292W | 66.2504152 | -0.4221363 | 0.58903718 | 0.83479115 | 0.65876028 | 0.78659971 | PUS4 | XIV | 82806 | 84017 | 1 |
| YNL293W | 51.4356241 | 0.34201365 | 0.58929183 | 2.03625128 | 0.36127146 | 0.53408966 | MSB3 | XIV | 80640 | 82541 | 1 |
| YNL294C | 89.5947016 | 0.9972079 | 0.57016814 | 3.02363181 | 0.22050919 | 0.38525897 | RIM21 | XIV | 78659 | 80260 | -1 |
| YNL295W | 175.44022 | 2.06352075 | 0.63940528 | 11.5128383 | 0.00316242 | 0.01590824 | MRX6 | XIV | 76946 | 78520 | 1 |
| YNL297C | 23.5195825 | 0.03522364 | 0.70983879 | 0.59631688 | 0.74218374 | 0.84243165 | MON2 | XIV | 71673 | 76583 | -1 |
| YNL298W | 66.8407858 | 0.92444118 | 0.7118303 | 2.08926492 | 0.3518211 | 0.5255532 | CLA4 | XIV | 68915 | 71443 | 1 |
| YNL299W | 143.682919 | -0.026582 | 0.60692322 | 51.8950465 | 5.38E-12 | 3.43E-10 | TRF5 | XIV | 66517 | 68445 | 1 |
| YNL300W | 151.888724 | -0.8301029 | 0.69089326 | 1.43177719 | 0.48875761 | 0.65040372 | TOS6 | XIV | 65744 | 66052 | 1 |
| YNL301C | 44.2047003 | -0.57877 | 0.60161239 | 4.01033568 | 0.1346377 | 0.27105433 | RPL18B | XIV | 63570 | 64562 | -1 |
| YNL302C | 122.086514 | -0.6482663 | 0.87081028 | 9.20270099 | 0.01003827 | 0.03953849 | RPS19B | XIV | 61958 | 62943 | -1 |
| YNL304W | 42.5735855 | -0.8143697 | 0.72838598 | 4.24559713 | 0.11969618 | 0.2489022 | YPT11 | XIV | 60297 | 61550 | 1 |
| YNL305C | 108.733107 | 1.78168666 | 0.81484529 | 6.47651791 | 0.03923214 | 0.10991524 | BXI1 | XIV | 58898 | 59791 | -1 |

| | | | | | | | | | | | |
|---|---|---|---|---|---|---|---|---|---|---|---|
| YNL306W | 221.199628 | 3.26916704 | 0.90166027 | 11.4715595 | 0.00322836 | 0.01618375 | MRPS18 | XIV | 58155 | 58808 | 1 |
| YNL307C | 20.381995 | -0.3230681 | 0.792131 | 2.98955596 | 0.2242984 | 0.38976222 | MCK1 | XIV | 56446 | 57573 | -1 |
| YNL308C | 232.689932 | -0.3410861 | 0.53598884 | 12.0463478 | 0.00242197 | 0.01291271 | KRI1 | XIV | 54122 | 55897 | -1 |
| YNL309W | 17.3113154 | -0.6952988 | 0.80235396 | 0.9067155 | 0.63549075 | 0.77219127 | STB1 | XIV | 52662 | 53924 | 1 |
| YNL310C | 10.8437912 | -0.3346024 | 0.89188053 | 1.34525321 | 0.51036629 | 0.66786132 | ZIM17 | XIV | 51907 | 52431 | -1 |
| YNL311C | 37.3035245 | 1.07650145 | 0.77018698 | 2.80980887 | 0.24539051 | 0.41605008 | SKP2 | XIV | 49397 | 51688 | -1 |
| YNL312W | 53.128492 | -0.8611614 | 0.64263172 | 1.97643772 | 0.37223911 | 0.54549847 | RFA2 | XIV | 48287 | 49216 | 1 |
| YNL313C | 63.1044581 | 0.87155714 | 0.64553112 | 2.25286772 | 0.32418729 | 0.49763242 | EMW1 | XIV | 45308 | 48022 | -1 |
| YNL314W | 58.539092 | 0.77576132 | 0.89336572 | 4.76271407 | 0.09242507 | 0.20685781 | DAL82 | XIV | 44447 | 45214 | 1 |
| YNL315C | 16.2354101 | -0.2300227 | 0.77733859 | 0.14718791 | 0.92904885 | 0.95541731 | ATP11 | XIV | 43324 | 44280 | -1 |
| YNL316C | 306.475876 | 3.00315001 | 0.90322696 | 10.1383313 | 0.00628766 | 0.0275011 | PHA2 | XIV | 42071 | 43075 | -1 |
| YNL317W | 34.3889775 | 1.89362428 | 0.84721836 | 5.67297509 | 0.05863124 | 0.14727762 | PFS2 | XIV | 40619 | 42016 | 1 |
| YNL318C | 162.537016 | 2.23484772 | 0.84657828 | 8.47175688 | 0.0144671 | 0.05215721 | HXT14 | XIV | 38707 | 40329 | -1 |
| YNL320W | 27.6964423 | -0.4565102 | 0.8674418 | 2.12713689 | 0.3452217 | 0.51852261 | NA | XIV | 37700 | 38554 | 1 |
| YNL321W | 74.6297752 | -0.4505879 | 0.58292601 | 0.59722675 | 0.74184617 | 0.84243165 | VNX1 | XIV | 34696 | 37422 | 1 |
| YNL322C | 294.874699 | 2.41384967 | 0.82964762 | 9.11358629 | 0.01049566 | 0.04092248 | KRE1 | XIV | 33294 | 34235 | -1 |
| YNL323W | 305.087323 | 0.43981082 | 0.54107386 | 3.62638926 | 0.16313216 | 0.3118307 | LEM3 | XIV | 31944 | 33188 | 1 |
| YNL325C | 54.877303 | 0.15428747 | 0.58626648 | 0.32983346 | 0.84796431 | 0.90980423 | FIG4 | XIV | 28739 | 31378 | -1 |
| YNL326C | 8.18216707 | -0.3248403 | 1.16733992 | 0.18286926 | 0.91262097 | 0.94617945 | PFA3 | XIV | 27337 | 28347 | -1 |
| YNL327W | 72.2198397 | 0.51884999 | 0.69102571 | 1.11762453 | 0.57188791 | 0.72218779 | EGT2 | XIV | 24048 | 27173 | 1 |
| YNL328C | 80.1179336 | -0.230441 | 0.77090001 | 3.7147934 | 0.15607842 | 0.30143231 | MDJ2 | XIV | 22834 | 23274 | -1 |
| YNL329C | 38.7722231 | -0.6296915 | 0.62305407 | 2.99642148 | 0.22352975 | 0.38912726 | PEX6 | XIV | 19541 | 22633 | -1 |
| YNL330C | 13.4291546 | 0.19647176 | 0.95130913 | 0.17296047 | 0.91715368 | 0.94877967 | RPD3 | XIV | 18001 | 19302 | -1 |
| YNL331C | 348.213508 | -0.0997352 | 0.5064387 | 0.18561469 | 0.91136907 | 0.94542349 | AAD14 | XIV | 16118 | 17248 | -1 |
| YNL332W | 7.85211016 | -0.9148232 | 1.15068607 | 1.75671778 | 0.41546418 | 0.58728945 | THI12 | XIV | 14832 | 15854 | 1 |
| YNL333W | 47.5625948 | 0.816013 | 0.64047888 | 4.3583556 | 0.11313451 | 0.23968124 | SNZ2 | XIV | 13267 | 14163 | 1 |
| YNL334C | 6.48151049 | 0.05462895 | 1.17919757 | 1.06863326 | 0.58606966 | 0.73378964 | SNO2 | XIV | 12208 | 12876 | -1 |
| YNL335W | 134.095756 | -1.2279831 | 0.550335 | 5.9987314 | 0.04981866 | 0.13021013 | DDI3 | XIV | 11452 | 12129 | 1 |
| YNL336W | 229.973348 | 3.732618 | 0.60236041 | 36.6014371 | 1.13E-08 | 3.34E-07 | COS1 | XIV | 8330 | 9475 | 1 |
| YNR001C | 48.3517872 | 2.18066664 | 0.76386413 | 8.0055985 | 0.01826444 | 0.06184275 | CIT1 | XIV | 629622 | 631061 | -1 |
| YNR002C | 898.140829 | 0.71137732 | 0.50687475 | 4.66269792 | 0.09716459 | 0.21489634 | ATO2 | XIV | 633008 | 633856 | -1 |
| YNR004W | 681.992359 | 1.54322251 | 0.51613477 | 8.73556585 | 0.01267932 | 0.0471202 | SWM2 | XIV | 635943 | 636383 | 1 |
| YNR006W | 43.1013307 | 1.90503539 | 0.66971097 | 7.9242109 | 0.01902302 | 0.06366664 | VPS27 | XIV | 636986 | 638854 | 1 |

| | | | | | | | | | | | |
|---|---|---|---|---|---|---|---|---|---|---|---|
| YNR007C | 18.1083202 | 1.0833281 | 0.99341389 | 1.88372424 | 0.38990112 | 0.56101577 | ATG3 | XIV | 639182 | 640114 | -1 |
| YNR008W | 79.8538422 | 0.39773469 | 0.53850833 | 0.76233051 | 0.683065 | 0.80595449 | LRO1 | XIV | 640396 | 642381 | 1 |
| YNR009W | 195.101111 | 1.72172806 | 0.51614489 | 21.9352424 | 1.73E-05 | 0.00020766 | NRM1 | XIV | 642690 | 643439 | 1 |
| YNR010W | 61.9264056 | 1.10558622 | 0.58825983 | 7.17266937 | 0.02769967 | 0.08374992 | CSE2 | XIV | 643744 | 644193 | 1 |
| YNR011C | 464.512499 | 2.93897979 | 0.53705027 | 51.524773 | 6.48E-12 | 4.08E-10 | PRP2 | XIV | 644320 | 646950 | -1 |
| YNR012W | 546.595819 | 2.64605884 | 0.91889167 | 10.8591535 | 0.00438495 | 0.02059113 | URK1 | XIV | 647432 | 648937 | 1 |
| YNR013C | 37.0748061 | -0.4253459 | 0.61026089 | 2.19222677 | 0.33416734 | 0.5079834 | PHO91 | XIV | 649028 | 651712 | -1 |
| YNR014W | 212.26516 | -1.9511946 | 0.53854757 | 18.2394484 | 0.00010948 | 0.00100942 | NA | XIV | 652465 | 653103 | 1 |
| YNR015W | 79.8110266 | 1.31767176 | 0.88506781 | 4.5102024 | 0.10486293 | 0.22688953 | SMM1 | XIV | 653387 | 654541 | 1 |
| YNR016C | 85.0057386 | -0.6989543 | 0.63061483 | 2.5231267 | 0.28321092 | 0.45678444 | ACC1 | XIV | 654673 | 661374 | -1 |
| YNR017W | 19.5008405 | -0.7546098 | 0.90796737 | 3.81168304 | 0.14869746 | 0.29106095 | TIM23 | XIV | 662913 | 663581 | 1 |
| YNR018W | 112.511025 | -1.4203468 | 0.53118133 | 15.8553076 | 0.00036063 | 0.00284348 | RCF2 | XIV | 664270 | 664944 | 1 |
| YNR019W | 89.5544446 | 1.21061011 | 0.57371392 | 4.42674502 | 0.10933131 | 0.2335036 | ARE2 | XIV | 665339 | 667267 | 1 |
| YNR020C | 23.9278169 | 0.0134953 | 0.70177593 | 1.58437853 | 0.45285229 | 0.61883757 | NA | XIV | 667410 | 668222 | -1 |
| YNR021W | 62.4212231 | -0.5988176 | 0.56208736 | 6.47736626 | 0.0392155 | 0.10991524 | NA | XIV | 668377 | 669591 | 1 |
| YNR022C | 313.694246 | 1.12142698 | 0.84927395 | 11.4370966 | 0.00328448 | 0.01637994 | MRPL50 | XIV | 669774 | 670193 | -1 |
| YNR023W | 35.929788 | 0.23428453 | 0.76152034 | 9.93661107 | 0.00695492 | 0.02989533 | SNF12 | XIV | 670418 | 672118 | 1 |
| YNR024W | 309.290298 | 2.51204365 | 0.80606053 | 11.541699 | 0.00311711 | 0.0157213 | MPP6 | XIV | 672409 | 672969 | 1 |
| YNR026C | 387.891433 | 1.31880105 | 0.54810017 | 93.9485265 | 3.97E-21 | 1.35E-18 | SEC12 | XIV | 673274 | 674689 | -1 |
| YNR027W | 6.18698074 | 0.66928022 | 1.44257151 | 0.41736979 | 0.81165095 | 0.88731158 | BUD17 | XIV | 674923 | 675876 | 1 |
| YNR028W | 89.6730653 | 1.37541355 | 0.68664947 | 6.35292379 | 0.04173305 | 0.11462731 | CPR8 | XIV | 676177 | 677103 | 1 |
| YNR029C | 72.3947954 | 1.2425952 | 0.73663821 | 8.13159647 | 0.01714929 | 0.05904684 | NA | XIV | 677199 | 678488 | -1 |
| YNR030W | 11.0227003 | 0.19724309 | 1.04018543 | 0.32881706 | 0.84839536 | 0.90980423 | ALG12 | XIV | 678799 | 680454 | 1 |
| YNR031C | 12.0077344 | 0.33965433 | 0.87057726 | 0.75729564 | 0.68478674 | 0.80718958 | SSK2 | XIV | 680694 | 685433 | -1 |
| YNR032C-A | 111.418053 | 0.68159774 | 0.79234797 | 2.88645192 | 0.23616467 | 0.40406514 | HUB1 | XIV | 687243 | 687464 | -1 |
| YNR032W | 44.9744378 | -0.2362773 | 0.63810226 | 4.29069412 | 0.11702741 | 0.24505921 | PPG1 | XIV | 686010 | 687116 | 1 |
| YNR033W | 65.4243211 | 0.07307435 | 0.59064474 | 2.71143318 | 0.25776252 | 0.42975819 | ABZ1 | XIV | 687635 | 689998 | 1 |
| YNR034W | 17.4871431 | 1.08332023 | 0.87114973 | 2.85716543 | 0.23964833 | 0.40786709 | SOL1 | XIV | 690321 | 691286 | 1 |
| YNR034W-A | 38.4801718 | 0.76107856 | 0.63858211 | 1.53432954 | 0.46432768 | 0.62726488 | EGO4 | XIV | 692561 | 692857 | 1 |
| YNR035C | 49.9743691 | 0.92139844 | 0.60384163 | 2.36868452 | 0.30594735 | 0.47867401 | ARC35 | XIV | 693019 | 694047 | -1 |
| YNR036C | 24.030335 | 0.32329529 | 0.73971776 | 0.33557076 | 0.84553529 | 0.90923806 | MRPS12 | XIV | 694361 | 694822 | -1 |
| YNR037C | 8.00364484 | 0.82116281 | 1.17100814 | 0.56848459 | 0.75258428 | 0.85034238 | RSM19 | XIV | 695052 | 695327 | -1 |
| YNR038W | 27.2784937 | -0.2196433 | 0.68706507 | 5.58216062 | 0.0613549 | 0.15226954 | DBP6 | XIV | 695595 | 697484 | 1 |

| | | | | | | | | | | | |
|---|---|---|---|---|---|---|---|---|---|---|---|
| YNR039C | 80.2611807 | 0.8066872 | 0.69339006 | 10.9033002 | 0.00428922 | 0.02021351 | ZRG17 | XIV | 697614 | 699431 | -1 |
| YNR040W | 14.1816304 | -0.1716406 | 0.8637255 | 0.12095005 | 0.94131728 | 0.96106984 | NA | XIV | 699690 | 700460 | 1 |
| YNR041C | 56.2393097 | 0.31290194 | 0.58723827 | 0.31349152 | 0.85492139 | 0.91395769 | COQ2 | XIV | 700541 | 701659 | -1 |
| YNR043W | 45.6257504 | -0.2067852 | 0.64423142 | 0.71062027 | 0.70095602 | 0.81801018 | MVD1 | XIV | 701895 | 703085 | 1 |
| YNR044W | 20.5945621 | -0.7745835 | 0.78794474 | 0.95757607 | 0.61953379 | 0.76030111 | AGA1 | XIV | 703699 | 705876 | 1 |
| YNR045W | 58.6279175 | -1.534083 | 0.58997759 | 10.6812619 | 0.00479285 | 0.02212703 | PET494 | XIV | 706139 | 707608 | 1 |
| YNR046W | 14.5027222 | -1.3423174 | 0.83706584 | 3.08398878 | 0.21395397 | 0.37804141 | TRM112 | XIV | 707788 | 708195 | 1 |
| YNR047W | 53.1910999 | -1.2777724 | 0.61170463 | 12.1547266 | 0.00229422 | 0.01236827 | FPK1 | XIV | 708523 | 711204 | 1 |
| YNR048W | 15.6427688 | 0.17632601 | 0.84065123 | 3.48637707 | 0.17496164 | 0.32678559 | NA | XIV | 711630 | 712811 | 1 |
| YNR049C | 29.2830958 | -0.0710497 | 0.70470105 | 0.01717526 | 0.99144914 | 0.99333631 | MSO1 | XIV | 713023 | 713655 | -1 |
| YNR050C | 34.1369823 | 0.00775389 | 0.75522214 | 0.03251604 | 0.98387343 | 0.98751104 | LYS9 | XIV | 714048 | 715388 | -1 |
| YNR051C | 80.3609361 | 0.05825532 | 0.5281903 | 5.95981619 | 0.0507975 | 0.13219068 | BRE5 | XIV | 716780 | 718327 | -1 |
| YNR052C | 20.3978625 | 0.1061243 | 0.79414433 | 1.28012677 | 0.527259 | 0.68184633 | POP2 | XIV | 719346 | 720647 | -1 |
| YNR053C | 134.116212 | -0.5914729 | 0.65737641 | 1.83227671 | 0.40006096 | 0.57095216 | NOG2 | XIV | 721120 | 723112 | -1 |
| YNR054C | 37.3213495 | -0.8537576 | 0.70264973 | 11.8080925 | 0.00272838 | 0.01413927 | ESF2 | XIV | 723356 | 724306 | -1 |
| YNR055C | 71.0876633 | -0.0805183 | 0.56813339 | 5.60861605 | 0.06054865 | 0.15085056 | HOL1 | XIV | 728426 | 730186 | -1 |
| YNR056C | 2600.7659 | 0.67433358 | 0.61338375 | 2.35967645 | 0.30732845 | 0.47995819 | BIO5 | XIV | 731618 | 733303 | -1 |
| YNR057C | 70.6033261 | -1.0667826 | 0.61151636 | 4.72988211 | 0.09395484 | 0.2093602 | BIO4 | XIV | 733356 | 734069 | -1 |
| YNR058W | 755.807315 | 4.07965107 | 0.63118861 | 99.6467019 | 2.30E-22 | 1.33E-19 | BIO3 | XIV | 734291 | 735733 | 1 |
| YNR059W | 35.5637298 | -0.808591 | 0.73247688 | 3.18617346 | 0.20329712 | 0.36352388 | MNT4 | XIV | 736803 | 738545 | 1 |
| YNR060W | 601.466991 | 0.0614372 | 0.46573131 | 9.73618171 | 0.00768803 | 0.03213985 | FRE4 | XIV | 739951 | 742110 | 1 |
| YNR061C | 11.4951043 | -0.7420724 | 0.91916386 | 2.67459945 | 0.26255368 | 0.43396683 | NA | XIV | 742881 | 743540 | -1 |
| YNR062C | 965.310118 | 0.39128083 | 0.52651037 | 0.62617636 | 0.73118543 | 0.83700349 | NA | XIV | 744360 | 745343 | -1 |
| YNR063W | 719.023215 | 4.15948072 | 0.50512952 | 95.4702596 | 1.86E-21 | 6.72E-19 | NA | XIV | 746943 | 748766 | 1 |
| YNR064C | 413.403717 | -1.0690563 | 0.50130873 | 7.2612672 | 0.02649939 | 0.08100922 | NA | XIV | 749136 | 750008 | -1 |
| YNR065C | 82.7083224 | 2.56073188 | 0.64344574 | 17.1049429 | 0.00019307 | 0.0016512 | NA | XIV | 750350 | 753700 | -1 |
| YNR066C | 60.9852785 | -0.375728 | 0.61813445 | 2.29471987 | 0.31747382 | 0.49004889 | NA | XIV | 753725 | 755035 | -1 |
| YNR067C | 1658.65563 | 2.73907578 | 0.47057846 | 46.2706748 | 8.96E-11 | 4.32E-09 | DSE4 | XIV | 755746 | 759099 | -1 |
| YNR068C | 25.5915277 | -0.9609353 | 0.84110769 | 6.25006512 | 0.0439355 | 0.11887223 | NA | XIV | 760067 | 760885 | -1 |
| YNR069C | 53.7557522 | -1.0383908 | 0.68799514 | 2.76257547 | 0.25125479 | 0.42270232 | BSC5 | XIV | 761123 | 762592 | -1 |
| YNR070W | 23.6777157 | 0.22176899 | 0.89771413 | 3.35260168 | 0.18706468 | 0.34297166 | PDR18 | XIV | 765375 | 769376 | 1 |
| YNR072W | 12.4821222 | 0.57006386 | 0.9605388 | 3.09825428 | 0.21243332 | 0.3760284 | HXT17 | XIV | 772657 | 774351 | 1 |
| YNR075C-A | 679.999401 | 0.81388773 | 0.60115983 | 2.70838389 | 0.25815581 | 0.43001213 | NA | XIV | 781511 | 781603 | -1 |

| | | | | | | | | | | | |
|---|---|---|---|---|---|---|---|---|---|---|---|
| YNR075W | 102.688702 | 2.47736337 | 0.73007704 | 32.9628881 | 6.95E-08 | 1.75E-06 | COS10 | XIV | 779916 | 781040 | 1 |
| YOL001W | 63.4947046 | 1.29556843 | 0.67681123 | 21.9847393 | 1.68E-05 | 0.00020343 | PHO80 | XV | 325249 | 326130 | 1 |
| YOL002C | 111.771148 | 1.21269177 | 0.63283851 | 21.1478779 | 2.56E-05 | 0.00029438 | IZH2 | XV | 323411 | 324364 | -1 |
| YOL003C | 425.251481 | 0.95941467 | 0.47282754 | 9.8088094 | 0.00741386 | 0.03137882 | PFA4 | XV | 321858 | 322994 | -1 |
| YOL004W | 92.9126621 | 0.45428694 | 0.52354212 | 1.30229551 | 0.52144694 | 0.67724939 | SIN3 | XV | 316938 | 321548 | 1 |
| YOL005C | 35.2508043 | -0.2026452 | 0.71250482 | 0.08132799 | 0.96015169 | 0.97173192 | RPB11 | XV | 315813 | 316175 | -1 |
| YOL006C | 257.191758 | 0.37049276 | 0.51413748 | 0.60483753 | 0.73902852 | 0.84040603 | TOP1 | XV | 313078 | 315387 | -1 |
| YOL007C | 276.933098 | 0.18763745 | 0.5421995 | 20.9612818 | 2.81E-05 | 0.00031936 | CSI2 | XV | 311342 | 312367 | -1 |
| YOL008W | 73.3903805 | -0.3948906 | 0.60424187 | 0.83965085 | 0.65716153 | 0.78549013 | COQ10 | XV | 310312 | 310935 | 1 |
| YOL009C | 16.340064 | 1.27100139 | 1.00881869 | 7.29754111 | 0.0260231 | 0.07997546 | MDM12 | XV | 309324 | 310139 | -1 |
| YOL010W | 25.0995328 | 0.0583052 | 0.70762231 | 0.07981806 | 0.96087684 | 0.97212597 | RCL1 | XV | 307938 | 309041 | 1 |
| YOL011W | 22.9642181 | 0.19979962 | 0.72600453 | 0.49229392 | 0.78180732 | 0.86750947 | PLB3 | XV | 305349 | 307409 | 1 |
| YOL012C | 232.957717 | -0.1046833 | 0.53086336 | 0.35622844 | 0.83684683 | 0.90381331 | HTZ1 | XV | 303579 | 303983 | -1 |
| YOL013C | 44.7404919 | 0.29902203 | 0.6578006 | 1.05398654 | 0.59037741 | 0.73765326 | HRD1 | XV | 301380 | 303035 | -1 |
| YOL014W | 25.050078 | 0.56755307 | 0.7261663 | 2.53555322 | 0.28145671 | 0.45563342 | NA | XV | 299693 | 300067 | 1 |
| YOL015W | 39.6820177 | 0.73866674 | 0.6900798 | 1.149641 | 0.56280588 | 0.71410256 | IRC10 | XV | 297078 | 298838 | 1 |
| YOL016C | 98.538753 | 1.6672993 | 0.68017263 | 5.80695412 | 0.05483223 | 0.14011689 | CMK2 | XV | 294777 | 296120 | -1 |
| YOL017W | 306.619047 | 3.18372847 | 0.57785564 | 27.2520332 | 1.21E-06 | 2.08E-05 | ESC8 | XV | 292529 | 294673 | 1 |
| YOL018C | 27.0563243 | 0.66560442 | 0.70177431 | 1.47958926 | 0.47721191 | 0.63989277 | TLG2 | XV | 290881 | 292074 | -1 |
| YOL019W | 85.4745499 | 2.48378583 | 0.65582496 | 13.6217769 | 0.00110171 | 0.00712729 | NA | XV | 288898 | 290553 | 1 |
| YOL020W | 89.0652031 | 0.58479914 | 0.57114611 | 9.13180929 | 0.01040047 | 0.040606 | TAT2 | XV | 286172 | 287950 | 1 |
| YOL021C | 26.3072108 | -0.2166096 | 0.67591221 | 1.90264419 | 0.38623005 | 0.55874541 | DIS3 | XV | 282421 | 285426 | -1 |
| YOL022C | 41.3626991 | -0.2670767 | 0.64400999 | 4.67023911 | 0.09679891 | 0.21424529 | TSR4 | XV | 280272 | 281498 | -1 |
| YOL023W | 25.7067295 | 0.35869172 | 0.73384482 | 3.74389427 | 0.15382385 | 0.29821719 | IFM1 | XV | 278056 | 280086 | 1 |
| YOL025W | 26.809326 | 0.06577826 | 0.72969507 | 7.17104983 | 0.02772211 | 0.08377402 | LAG2 | XV | 274957 | 276939 | 1 |
| YOL026C | 14.8414966 | 0.64992195 | 0.8584411 | 0.78142093 | 0.67657602 | 0.8007717 | MIM1 | XV | 274012 | 274353 | -1 |
| YOL027C | 43.7781176 | 0.26970846 | 0.64499907 | 0.64571108 | 0.72407845 | 0.83364768 | MDM38 | XV | 272003 | 273724 | -1 |
| YOL028C | 25.6122913 | 0.65487314 | 0.75431481 | 0.95892285 | 0.61911674 | 0.76021421 | YAP7 | XV | 270633 | 271370 | -1 |
| YOL029C | 390.344253 | 3.07449306 | 1.5930328 | 4.06618957 | 0.13092969 | 0.26590071 | NA | XV | 269815 | 270420 | -1 |
| YOL030W | 38.6674049 | 0.28860805 | 0.65488651 | 0.20802644 | 0.90121338 | 0.93824416 | GAS5 | XV | 268187 | 269641 | 1 |
| YOL031C | 92.7343884 | -0.4008071 | 0.52829502 | 0.66179059 | 0.71828037 | 0.82928083 | SIL1 | XV | 266264 | 267529 | -1 |
| YOL032W | 179.817304 | 4.12503122 | 0.82712492 | 21.4512297 | 2.20E-05 | 0.00025693 | OPI10 | XV | 265429 | 266169 | 1 |
| YOL033W | 94.1959288 | -0.4840777 | 0.55500934 | 7.35746093 | 0.02525502 | 0.07827974 | MSE1 | XV | 263475 | 265085 | 1 |

| | | | | | | | | | | | |
|---|---|---|---|---|---|---|---|---|---|---|---|
| YOL034W | 19.3622168 | -0.3494194 | 0.80217594 | 0.63875136 | 0.72660253 | 0.83481707 | SMC5 | XV | 259923 | 263204 | 1 |
| YOL036W | 201.455001 | 2.73200019 | 0.57849353 | 45.0381059 | 1.66E-10 | 7.44E-09 | NA | XV | 256745 | 259030 | 1 |
| YOL038C-A | 31.5396252 | -0.7878372 | 0.65121554 | 4.34984823 | 0.11361678 | 0.24017566 | NA | XV | 254926 | 255021 | -1 |
| YOL038W | 82.1544553 | 0.70087202 | 0.60066459 | 1.97058222 | 0.37333053 | 0.54640641 | PRE6 | XV | 255337 | 256101 | 1 |
| YOL039W | 76.8831998 | -0.6593112 | 0.58019241 | 10.6974152 | 0.00475429 | 0.02196915 | RPP2A | XV | 254297 | 254617 | 1 |
| YOL040C | 165.496314 | -0.2818248 | 0.68562516 | 13.9818714 | 0.00092019 | 0.00615027 | RPS15 | XV | 253149 | 253577 | -1 |
| YOL041C | 60.9819771 | -0.0983528 | 0.62858762 | 3.30304434 | 0.1917578 | 0.34826777 | NOP12 | XV | 251267 | 252646 | -1 |
| YOL042W | 90.1501908 | 2.99912747 | 0.77336148 | 38.741722 | 3.87E-09 | 1.27E-07 | NGL1 | XV | 249825 | 250916 | 1 |
| YOL043C | 142.616706 | 0.55284675 | 0.61241542 | 26.3016207 | 1.94E-06 | 3.18E-05 | NTG2 | XV | 248392 | 249534 | -1 |
| YOL044W | 32.0489979 | 2.04659208 | 0.92562239 | 4.74562512 | 0.09321818 | 0.20831078 | PEX15 | XV | 247150 | 248301 | 1 |
| YOL045W | 31.7394034 | 0.74487735 | 0.77941152 | 12.020465 | 0.00245352 | 0.01302799 | PSK2 | XV | 243497 | 246802 | 1 |
| YOL047C | 406.592469 | 1.80724711 | 0.52116272 | 17.7635719 | 0.0001389 | 0.0012449 | LDS2 | XV | 241614 | 242747 | -1 |
| YOL048C | 156.235201 | 2.36383993 | 0.87339388 | 9.7957388 | 0.00746247 | 0.03156149 | RRT8 | XV | 240204 | 241310 | -1 |
| YOL049W | 125.820516 | 1.76888544 | 0.7005753 | 6.07735615 | 0.04789817 | 0.12634641 | GSH2 | XV | 238619 | 240094 | 1 |
| YOL051W | 83.5437118 | 0.48255447 | 0.59470034 | 2.53140434 | 0.28204119 | 0.45618192 | GAL11 | XV | 234940 | 238185 | 1 |
| YOL052C | 242.187212 | 1.34986138 | 0.50954986 | 32.0652819 | 1.09E-07 | 2.58E-06 | SPE2 | XV | 232446 | 233636 | -1 |
| YOL052C-A | 22.9689517 | 0.88594195 | 0.94820417 | 30.0239551 | 3.02E-07 | 6.30E-06 | DDR2 | XV | 231570 | 231755 | -1 |
| YOL053W | 138.382063 | 3.60785995 | 0.91626085 | 36.454432 | 1.21E-08 | 3.55E-07 | AIM39 | XV | 230085 | 231272 | 1 |
| YOL054W | 34.2377613 | 0.75974101 | 0.69976258 | 7.78070453 | 0.02043815 | 0.06716053 | PSH1 | XV | 228614 | 229834 | 1 |
| YOL055C | 114.80483 | -0.5355428 | 0.53392537 | 1.92400222 | 0.38212744 | 0.55547441 | THI20 | XV | 224420 | 226075 | -1 |
| YOL056W | 8.53704861 | -0.1996132 | 1.03068655 | 1.1683332 | 0.55757035 | 0.70905608 | GPM3 | XV | 223268 | 224179 | 1 |
| YOL057W | 31.3450712 | 0.11934482 | 0.73991053 | 0.55758391 | 0.75669731 | 0.8528864 | NA | XV | 220767 | 222902 | 1 |
| YOL058W | 1927.87114 | 0.79283394 | 0.47220734 | 7.46144673 | 0.02397549 | 0.07565017 | ARG1 | XV | 219211 | 220473 | 1 |
| YOL059W | 56.6939223 | 0.39724141 | 0.59040678 | 0.68813055 | 0.70888266 | 0.82407349 | GPD2 | XV | 217126 | 218448 | 1 |
| YOL060C | 193.292297 | 0.74394524 | 0.57938937 | 41.8093606 | 8.34E-10 | 3.08E-08 | MAM3 | XV | 214017 | 216137 | -1 |
| YOL061W | 85.6035009 | 0.93035048 | 0.72426095 | 1.85721957 | 0.39510261 | 0.5664635 | PRS5 | XV | 212244 | 213734 | 1 |
| YOL062C | 27.7127169 | 0.39017461 | 0.77951069 | 0.53434335 | 0.76554164 | 0.8573474 | APM4 | XV | 210520 | 211995 | -1 |
| YOL063C | 10.6249843 | -0.3746983 | 1.00595364 | 0.30923669 | 0.8567421 | 0.91522818 | CRT10 | XV | 207392 | 210265 | -1 |
| YOL064C | 8.13586319 | -0.2948855 | 1.08151987 | 0.75583236 | 0.68528794 | 0.80761595 | MET22 | XV | 206103 | 207176 | -1 |
| YOL066C | 78.0005346 | -0.5998995 | 0.71443948 | 8.72837003 | 0.01272502 | 0.0472597 | RIB2 | XV | 202696 | 204471 | -1 |
| YOL067C | 35.7252296 | -1.6524163 | 0.67949373 | 6.06460722 | 0.04820447 | 0.12703862 | RTG1 | XV | 201985 | 202518 | -1 |
| YOL068C | 181.704835 | -0.4468999 | 0.51454067 | 2.02583342 | 0.36315821 | 0.53614552 | HST1 | XV | 200368 | 201879 | -1 |
| YOL069W | 451.504409 | -1.0158517 | 0.51380113 | 5.50294205 | 0.06383389 | 0.15707532 | NUF2 | XV | 198942 | 200297 | 1 |

| | | | | | | | | | | | |
|---|---|---|---|---|---|---|---|---|---|---|---|
| YOL070C | 23.2665935 | -0.7514269 | 0.7383427 | 1.55827734 | 0.45880102 | 0.62328904 | NBA1 | XV | 197220 | 198725 | -1 |
| YOL071W | 56.9918828 | 0.91343396 | 0.59348827 | 3.20805348 | 0.20108517 | 0.36023611 | SDH5 | XV | 196507 | 196995 | 1 |
| YOL072W | 7.86451042 | 0.19241198 | 1.11125313 | 0.1134023 | 0.94487641 | 0.96266662 | THP1 | XV | 194970 | 196337 | 1 |
| YOL073C | 22.9062031 | -0.3384025 | 0.72916536 | 0.32963303 | 0.8480493 | 0.90980423 | DSC2 | XV | 193832 | 194800 | -1 |
| YOL075C | 473.147564 | 0.28820538 | 0.48300203 | 8.53421166 | 0.01402231 | 0.05090229 | NA | XV | 189658 | 193542 | -1 |
| YOL076W | 111.015564 | 0.39736974 | 0.71875961 | 5.36731415 | 0.06831287 | 0.16528689 | MDM20 | XV | 187024 | 189414 | 1 |
| YOL077C | 64.4396751 | 0.95282744 | 0.86647676 | 8.37420278 | 0.01519025 | 0.05409075 | BRX1 | XV | 185848 | 186723 | -1 |
| YOL077W-A | 46.5758531 | -0.1531891 | 0.66812705 | 1.35928264 | 0.50679874 | 0.66464916 | ATP19 | XV | 185438 | 185644 | 1 |
| YOL078W | 33.5714347 | -0.5723595 | 0.68158782 | 2.53105213 | 0.28209086 | 0.45618192 | AVO1 | XV | 181682 | 185212 | 1 |
| YOL080C | 39.9091585 | -0.6363535 | 0.64188775 | 18.1549589 | 0.00011421 | 0.00104466 | REX4 | XV | 180558 | 181427 | -1 |
| YOL081W | 59.2170702 | -0.6630118 | 0.59311913 | 2.60545949 | 0.27178887 | 0.44516479 | IRA2 | XV | 171070 | 180309 | 1 |
| YOL082W | 96.7384656 | -1.0705147 | 0.5550191 | 3.82102296 | 0.14800467 | 0.2902937 | ATG19 | XV | 168727 | 169974 | 1 |
| YOL083W | 29.9370306 | -0.407535 | 0.67244556 | 1.39956882 | 0.49669237 | 0.65643662 | ATG34 | XV | 165714 | 166952 | 1 |
| YOL084W | 14.3495785 | -0.0311644 | 0.86267576 | 1.13440139 | 0.56711074 | 0.71787738 | PHM7 | XV | 162356 | 165331 | 1 |
| YOL085C | 103.45469 | -0.083433 | 0.59516622 | 1.56036579 | 0.45832218 | 0.62328904 | NA | XV | 161673 | 162014 | -1 |
| YOL086C | 551.162842 | 0.01575104 | 0.65008277 | 8.96705991 | 0.01129348 | 0.04310431 | ADH1 | XV | 159548 | 160594 | -1 |
| YOL086W-A | 104.776284 | 2.24793035 | 0.7678309 | 16.1751178 | 0.00030734 | 0.00249667 | MHF1 | XV | 159173 | 159445 | 1 |
| YOL087C | 29.944492 | -0.1466463 | 0.8476911 | 1.40614747 | 0.49506128 | 0.65532803 | DUF1 | XV | 155287 | 158637 | -1 |
| YOL088C | 30.4572721 | -0.0662125 | 0.67403734 | 1.44595394 | 0.48530536 | 0.64773825 | MPD2 | XV | 153912 | 154745 | -1 |
| YOL089C | 55.6766668 | -0.8848227 | 0.58163675 | 2.60934262 | 0.27126168 | 0.44467869 | HAL9 | XV | 150398 | 153490 | -1 |
| YOL090W | 76.9468128 | 1.27888506 | 0.71626789 | 9.94467736 | 0.00692693 | 0.02979712 | MSH2 | XV | 147382 | 150276 | 1 |
| YOL091W | 57.2099947 | 1.38424662 | 0.62208921 | 17.8202421 | 0.00013502 | 0.00121388 | SPO21 | XV | 145334 | 147163 | 1 |
| YOL093W | 45.1446968 | -0.2634796 | 0.65671693 | 2.37900609 | 0.30437249 | 0.47720463 | TRM10 | XV | 142815 | 143696 | 1 |
| YOL094C | 41.0235307 | -1.8475566 | 0.85596486 | 4.87333134 | 0.08745196 | 0.19966358 | RFC4 | XV | 141584 | 142555 | -1 |
| YOL095C | 29.9833478 | 0.62785227 | 0.71977613 | 9.14680993 | 0.01032275 | 0.04032978 | HMI1 | XV | 139227 | 141347 | -1 |
| YOL096C | 8.97811843 | 1.29772204 | 1.07772382 | 2.05529474 | 0.35784785 | 0.53120548 | COQ3 | XV | 138107 | 139045 | -1 |
| YOL097C | 82.4596159 | -0.8991752 | 0.56975283 | 12.2166812 | 0.00222424 | 0.01213793 | WRS1 | XV | 136527 | 137825 | -1 |
| YOL097W-A | 123.63831 | 1.19606188 | 0.79552051 | 20.2855543 | 3.94E-05 | 0.0004228 | NA | XV | 136220 | 136405 | 1 |
| YOL098C | 91.9019144 | 0.15253621 | 0.5529951 | 4.40348551 | 0.11061022 | 0.23545338 | SDD3 | XV | 132725 | 135838 | -1 |
| YOL100W | 207.471582 | 0.83498042 | 0.56948184 | 2.42483306 | 0.29747755 | 0.47136562 | PKH2 | XV | 129237 | 132482 | 1 |
| YOL101C | 87.2250739 | 0.62722553 | 0.57046864 | 17.0003828 | 0.00020343 | 0.00173469 | IZH4 | XV | 126982 | 127920 | -1 |
| YOL102C | 39.0274342 | 0.03278388 | 0.63421048 | 3.52549223 | 0.17157306 | 0.32232576 | TPT1 | XV | 125997 | 126689 | -1 |
| YOL103W | 129.857466 | -0.1200339 | 0.539353 | 7.08455179 | 0.02894737 | 0.08664034 | ITR2 | XV | 124001 | 125830 | 1 |

| | | | | | | | | | | | |
|---|---|---|---|---|---|---|---|---|---|---|---|
| YOL103W-B | 7.44196863 | -1.1458228 | 1.06937452 | 2.54370611 | 0.28031171 | 0.45449588 | NA | XV | 118000 | 123268 | 1 |
| YOL104C | 11.586194 | -0.2196145 | 0.90165893 | 0.23943952 | 0.88716902 | 0.93215081 | NDJ1 | XV | 116396 | 117454 | -1 |
| YOL105C | 15.5387254 | -1.503679 | 0.8657547 | 2.88140581 | 0.23676128 | 0.40473806 | WSC3 | XV | 114138 | 115808 | -1 |
| YOL107W | 15.6360287 | -0.1063643 | 0.9908781 | 13.0502243 | 0.00146615 | 0.00896413 | NA | XV | 112102 | 113130 | 1 |
| YOL108C | 104.766931 | -0.2802573 | 0.63522367 | 23.8347354 | 6.67E-06 | 9.16E-05 | INO4 | XV | 111431 | 111886 | -1 |
| YOL109W | 421.347437 | -1.5090411 | 0.57220783 | 21.6306284 | 2.01E-05 | 0.00023746 | ZEO1 | XV | 110297 | 110638 | 1 |
| YOL110W | 16.267897 | 0.11143884 | 0.86059985 | 0.53929088 | 0.7636502 | 0.85622283 | SHR5 | XV | 109176 | 109889 | 1 |
| YOL111C | 175.71969 | 1.67447717 | 0.73321567 | 5.23010941 | 0.07316379 | 0.17432853 | MDY2 | XV | 108258 | 108896 | -1 |
| YOL112W | 111.054547 | 0.68497377 | 0.70430412 | 0.96904193 | 0.61599221 | 0.75801867 | MSB4 | XV | 106710 | 108188 | 1 |
| YOL113W | 43.1805564 | -0.0234872 | 0.65625023 | 0.55157088 | 0.75897576 | 0.85429037 | SKM1 | XV | 104326 | 106293 | 1 |
| YOL114C | 37.6135884 | 2.38196761 | 1.14318235 | 6.79925432 | 0.03338572 | 0.09713733 | PTH4 | XV | 103317 | 103925 | -1 |
| YOL115W | 26.0084112 | -1.3100687 | 0.68055455 | 6.71730722 | 0.03478206 | 0.10004686 | PAP2 | XV | 101475 | 103229 | 1 |
| YOL116W | 36.7293796 | -1.9851666 | 0.64647985 | 29.2027946 | 4.56E-07 | 8.94E-06 | MSN1 | XV | 99809 | 100957 | 1 |
| YOL117W | 11.2597536 | -0.3471424 | 0.97855957 | 1.54011979 | 0.46298534 | 0.62618199 | RRI2 | XV | 97551 | 99488 | 1 |
| YOL118C | 625.513705 | -0.1343777 | 0.51554486 | 0.4554138 | 0.79635764 | 0.87685172 | NA | XV | 96608 | 96916 | -1 |
| YOL119C | 22.9022304 | -0.0527509 | 0.73331156 | 0.05312988 | 0.9737848 | 0.97984448 | MCH4 | XV | 94856 | 96361 | -1 |
| YOL120C | 125.677943 | -0.5637006 | 0.77955845 | 14.3726163 | 0.00075688 | 0.00521018 | RPL18B | XV | 93395 | 94402 | -1 |
| YOL121C | 68.976464 | 0.44850238 | 0.6064908 | 12.8812236 | 0.00159543 | 0.00955278 | RPS19A | XV | 92026 | 92850 | -1 |
| YOL122C | 133.055772 | -0.8596091 | 0.55755245 | 15.7382166 | 0.00038238 | 0.00297068 | SMF1 | XV | 89692 | 91419 | -1 |
| YOL123W | 42.4202096 | -0.0023997 | 0.62977833 | 14.5792764 | 0.00068257 | 0.00479044 | HRP1 | XV | 87844 | 89448 | 1 |
| YOL124C | 104.269007 | 1.9255047 | 0.62035489 | 45.1895727 | 1.54E-10 | 6.96E-09 | TRM11 | XV | 85456 | 86757 | -1 |
| YOL125W | 256.89019 | 0.88214981 | 0.53895012 | 13.8991031 | 0.00095907 | 0.00635691 | TRM13 | XV | 83834 | 85264 | 1 |
| YOL126C | 859.195963 | 2.5387721 | 0.51455548 | 31.160816 | 1.71E-07 | 3.86E-06 | MDH2 | XV | 81787 | 82920 | -1 |
| YOL127W | 64.6320847 | -0.643315 | 0.83141908 | 11.6218065 | 0.00299472 | 0.01523678 | RPL25 | XV | 80348 | 81190 | 1 |
| YOL128C | 38.8809389 | -1.0316968 | 0.69824866 | 5.82127701 | 0.05444096 | 0.13947484 | YGK3 | XV | 78352 | 79479 | -1 |
| YOL129W | 65.2338704 | 0.17310924 | 0.56635864 | 0.51694053 | 0.77223199 | 0.86150115 | VPS68 | XV | 77560 | 78114 | 1 |
| YOL130W | 35.2755909 | -0.3828005 | 0.70612044 | 0.42238853 | 0.80961677 | 0.8858947 | ALR1 | XV | 74400 | 76979 | 1 |
| YOL131W | 10.9306547 | 0.20932425 | 0.89919816 | 0.68241579 | 0.7109111 | 0.82438173 | NA | XV | 73031 | 73357 | 1 |
| YOL132W | 181.722749 | -0.0851951 | 0.49578972 | 0.37656105 | 0.82838229 | 0.89751749 | GAS4 | XV | 71300 | 72715 | 1 |
| YOL133W | 14.8149122 | -0.1121799 | 0.81469257 | 1.05136591 | 0.5911515 | 0.7383018 | HRT1 | XV | 70325 | 70690 | 1 |
| YOL135C | 36.3126098 | -0.7738093 | 0.62326302 | 3.45746522 | 0.17750924 | 0.3305817 | MED7 | XV | 69376 | 70044 | -1 |
| YOL136C | 197.278376 | 2.67930072 | 0.58145713 | 63.3259808 | 1.77E-14 | 1.63E-12 | PFK27 | XV | 67561 | 68754 | -1 |
| YOL137W | 420.923363 | 2.31406167 | 0.65217449 | 39.5916206 | 2.53E-09 | 8.61E-08 | BSC6 | XV | 65621 | 67114 | 1 |

| | | | | | | | | | | | |
|---|---|---|---|---|---|---|---|---|---|---|---|
| YOL138C | 47.8426269 | -1.1307929 | 0.60143039 | 5.03116619 | 0.08081578 | 0.1877702 | RTC1 | XV | 61325 | 65350 | -1 |
| YOL139C | 26.0744634 | -1.3440788 | 0.82946092 | 14.0525421 | 0.00088824 | 0.00596624 | CDC33 | XV | 60383 | 61024 | -1 |
| YOL140W | 9.96772976 | -1.3671219 | 1.00705466 | 3.37211504 | 0.18524842 | 0.34115406 | ARG8 | XV | 58759 | 60030 | 1 |
| YOL141W | 9.70493127 | -0.354139 | 1.06694444 | 2.43926493 | 0.29533869 | 0.46952527 | PPM2 | XV | 56452 | 58539 | 1 |
| YOL142W | 12.8116719 | -0.8884353 | 0.85008256 | 3.32693721 | 0.18948061 | 0.34544902 | RRP40 | XV | 55558 | 56280 | 1 |
| YOL143C | 53.10384 | 0.11605121 | 0.59078736 | 0.04583216 | 0.9773445 | 0.98260543 | RIB4 | XV | 54595 | 55104 | -1 |
| YOL144W | 701.166036 | 1.45574705 | 0.66904117 | 5.49506186 | 0.0640859 | 0.15736105 | NOP8 | XV | 53098 | 54552 | 1 |
| YOL145C | 66.0989102 | 0.42777368 | 0.55682902 | 0.88434274 | 0.6426395 | 0.77621415 | CTR9 | XV | 49555 | 52788 | -1 |
| YOL146W | 118.11049 | 1.43438871 | 0.64444275 | 6.4323814 | 0.04010755 | 0.11164553 | PSF3 | XV | 48864 | 49448 | 1 |
| YOL147C | 26.9923362 | -1.0503875 | 0.71609813 | 3.02842966 | 0.21998084 | 0.38460377 | PEX11 | XV | 47933 | 48643 | -1 |
| YOL148C | 34.3962596 | 0.00859994 | 0.6499303 | 1.35626421 | 0.50756418 | 0.66521629 | SPT20 | XV | 45760 | 47574 | -1 |
| YOL149W | 14.7914644 | -0.3157754 | 0.89485233 | 6.11761034 | 0.04694375 | 0.12464616 | DCP1 | XV | 44938 | 45633 | 1 |
| YOL151W | 174.413073 | -1.8987128 | 0.53843926 | 29.7749926 | 3.42E-07 | 6.93E-06 | GRE2 | XV | 43694 | 44722 | 1 |
| YOL152W | 638.951866 | -0.444467 | 0.51238839 | 4.45516274 | 0.10778882 | 0.23089058 | FRE7 | XV | 40748 | 42610 | 1 |
| YOL153C | 1435.49136 | 0.58561515 | 0.4916054 | 1.43278151 | 0.48851224 | 0.65022664 | NA | NA | NA | NA | NA |
| YOL154W | 468.904414 | 0.79291158 | 0.47032898 | 12.8475359 | 0.00162253 | 0.00963534 | ZPS1 | XV | 34658 | 35407 | 1 |
| YOL155C | 6482.0637 | -1.2246618 | 0.46029039 | 8.42038218 | 0.01484353 | 0.05301915 | HPF1 | XV | 28703 | 31606 | -1 |
| YOL155W-A | 199.56082 | 1.97318959 | 0.70677811 | 7.32272561 | 0.02569747 | 0.07926923 | NA | XV | 27084 | 27218 | 1 |
| YOL156W | 526.968956 | 0.9547889 | 0.49406157 | 7.41193502 | 0.02457643 | 0.07694787 | HXT11 | XV | 25273 | 26976 | 1 |
| YOL158C | 146.132403 | 0.82058286 | 0.57033863 | 33.0677791 | 6.60E-08 | 1.67E-06 | ENB1 | XV | 19491 | 21311 | -1 |
| YOL159C | 27.3725643 | 1.41755897 | 0.8332289 | 8.24251641 | 0.01622409 | 0.0567598 | CSS3 | XV | 17281 | 17796 | -1 |
| YOL159C-A | 157.181904 | 2.09081768 | 0.5459643 | 32.1355047 | 1.05E-07 | 2.53E-06 | NA | XV | 15233 | 15505 | -1 |
| YOL160W | 24.9658078 | 0.2810797 | 0.73443161 | 0.26139268 | 0.87748419 | 0.92780012 | NA | XV | 14313 | 14654 | 1 |
| YOL163W | 47.6941897 | 0.35024018 | 0.79029713 | 19.7832217 | 5.06E-05 | 0.00051851 | NA | XV | 9597 | 10106 | 1 |
| YOL164W | 470.54277 | -0.9387353 | 0.59834362 | 3.7317351 | 0.15476189 | 0.29951658 | BDS1 | XV | 6175 | 8115 | 1 |
| YOL164W-A | 186.96233 | 0.22774517 | 0.48702324 | 2.51336859 | 0.2845961 | 0.45823454 | NA | XV | 4130 | 4312 | 1 |
| YOL165C | 13.6286884 | -2.2027997 | 1.00929779 | 6.18537427 | 0.04537985 | 0.12165304 | AAD15 | XV | 1647 | 2078 | -1 |
| YOR001W | 35.6340114 | 0.41746512 | 0.68240669 | 0.49433408 | 0.78101022 | 0.86705973 | RRP6 | XV | 326832 | 329033 | 1 |
| YOR002W | 53.8234793 | 0.64511267 | 0.75943907 | 1.74337839 | 0.41824446 | 0.589459 | ALG6 | XV | 329417 | 331051 | 1 |
| YOR003W | 363.902661 | 0.66846127 | 0.47563456 | 9.63692563 | 0.0080792 | 0.03341947 | YSP3 | XV | 331455 | 332891 | 1 |
| YOR004W | 16.1589526 | -1.46954 | 0.79627963 | 8.35341998 | 0.01534892 | 0.05445482 | UTP23 | XV | 333592 | 334356 | 1 |
| YOR005C | 998.780796 | 0.40860039 | 0.51210818 | 8.9770079 | 0.01123744 | 0.04297543 | DNL4 | XV | 334509 | 337343 | -1 |
| YOR006C | 794.243636 | 1.69182908 | 0.67313803 | 36.0213919 | 1.51E-08 | 4.32E-07 | TSR3 | XV | 337680 | 338621 | -1 |

| | | | | | | | | | | | |
|---|---|---|---|---|---|---|---|---|---|---|---|
| YOR007C | 81.4950711 | 0.21273368 | 0.65659798 | 0.11739765 | 0.94299073 | 0.96176085 | SGT2 | XV | 338938 | 339978 | -1 |
| YOR008C | 50.2356803 | -0.333648 | 0.68558957 | 1.00789072 | 0.60414239 | 0.7493774 | SLG1 | XV | 341278 | 342414 | -1 |
| YOR008C-A | 80.2524277 | -2.0235684 | 0.59393988 | 29.905815 | 3.21E-07 | 6.58E-06 | NA | XV | 342857 | 343081 | -1 |
| YOR009W | 16.7794487 | 3.32995925 | 1.14219894 | 19.3443547 | 6.30E-05 | 0.00062796 | TIR4 | XV | 344335 | 345798 | 1 |
| YOR010C | 103.83477 | 0.165026 | 0.58043989 | 7.30507212 | 0.0259253 | 0.07980195 | TIR2 | XV | 346195 | 346950 | -1 |
| YOR011W | 276.814243 | -0.0704349 | 0.52628426 | 0.20200154 | 0.90393234 | 0.93946657 | AUS1 | XV | 349679 | 353863 | 1 |
| YOR011W-A | 26.3636701 | -0.1111882 | 0.75285194 | 4.30925457 | 0.1159464 | 0.24332354 | NA | XV | 355652 | 355858 | 1 |
| YOR012W | 441.756498 | -0.7689351 | 0.5329613 | 8.97159336 | 0.01126791 | 0.04303508 | NA | XV | 356543 | 356956 | 1 |
| YOR014W | 152.183153 | 1.00040583 | 0.53352658 | 21.8281488 | 1.82E-05 | 0.00021773 | RTS1 | XV | 357674 | 359947 | 1 |
| YOR015W | 514.086934 | 1.50747893 | 0.80640957 | 4.35208785 | 0.11348962 | 0.24016437 | NA | XV | 359993 | 360352 | 1 |
| YOR016C | 109.53205 | 1.09397158 | 0.63762239 | 4.03179106 | 0.13320106 | 0.26928567 | ERP4 | XV | 360461 | 361084 | -1 |
| YOR017W | 1028.19863 | 3.08204125 | 0.48741553 | 98.2593317 | 4.61E-22 | 2.22E-19 | PET127 | XV | 361412 | 363814 | 1 |
| YOR018W | 74.8474412 | -0.2014755 | 0.54982831 | 1.81962936 | 0.40259883 | 0.57358445 | ROD1 | XV | 364369 | 366882 | 1 |
| YOR019W | 356.523054 | -1.0169531 | 0.47545819 | 8.59596015 | 0.01359599 | 0.0495723 | NA | XV | 368127 | 370319 | 1 |
| YOR020C | 56.4426165 | 0.91733163 | 0.59270355 | 2.37931841 | 0.30432496 | 0.47720463 | HSP10 | XV | 370524 | 370844 | -1 |
| YOR020W-A | 30.7133408 | -0.5927189 | 0.67892547 | 1.31621827 | 0.51782955 | 0.67436642 | NA | XV | 371685 | 371957 | 1 |
| YOR021C | 8.92841555 | -0.8621345 | 1.25313401 | 1.82927067 | 0.40066271 | 0.57138844 | SFM1 | XV | 372798 | 373439 | -1 |
| YOR022C | 158.926166 | -0.2444589 | 0.53722178 | 11.8975227 | 0.00260907 | 0.01368683 | DDL1 | XV | 373710 | 375857 | -1 |
| YOR023C | 36.3323203 | 1.07836517 | 0.69017929 | 10.2171509 | 0.00604469 | 0.02665555 | AHC1 | XV | 376012 | 377712 | -1 |
| YOR025W | 29.8815031 | -0.1929655 | 0.68658972 | 0.10005575 | 0.95120291 | 0.96614462 | HST3 | XV | 378219 | 379562 | 1 |
| YOR026W | 71.8919627 | -1.4535681 | 0.66005247 | 4.73268067 | 0.09382346 | 0.20925957 | BUB3 | XV | 379781 | 380806 | 1 |
| YOR027W | 126.038214 | 0.01755684 | 0.61249946 | 1.52445307 | 0.46662631 | 0.62934226 | STI1 | XV | 381053 | 382822 | 1 |
| YOR028C | 10.0234402 | -1.4970278 | 0.93543019 | 3.93495672 | 0.13980896 | 0.27827222 | CIN5 | XV | 383533 | 384420 | -1 |
| YOR029W | 96.3287753 | -1.6055756 | 0.56805883 | 22.9154154 | 1.06E-05 | 0.00013875 | NA | XV | 384600 | 384935 | 1 |
| YOR030W | 93.0652191 | 1.26039207 | 0.60516488 | 37.8747401 | 5.96E-09 | 1.89E-07 | DFG16 | XV | 386825 | 388684 | 1 |
| YOR031W | 23.6088454 | 2.14520962 | 0.97257218 | 22.2659689 | 1.46E-05 | 0.00018246 | NA | NA | NA | NA | NA |
| YOR032C | 543.844122 | -0.6971202 | 0.50941684 | 9.16132162 | 0.01024812 | 0.04019232 | HMS1 | XV | 389771 | 391075 | -1 |
| YOR032W-A | 100.805184 | 0.80180381 | 0.75452066 | 3.12443523 | 0.20967059 | 0.37244958 | NA | XV | 392176 | 392376 | 1 |
| YOR033C | 87.5243077 | -0.69361 | 0.59605276 | 11.8267849 | 0.002703 | 0.01404881 | EXO1 | XV | 392416 | 394524 | -1 |
| YOR034C | 57.1301806 | -2.1137762 | 0.61033209 | 11.881075 | 0.00263062 | 0.01378395 | AKR2 | XV | 394837 | 397086 | -1 |
| YOR034C-A | 11.0205544 | -1.3882348 | 1.15508252 | 7.71378356 | 0.02113359 | 0.06897602 | NA | XV | 397426 | 397668 | -1 |
| YOR035C | 44.7567629 | 1.51056987 | 0.65759936 | 15.1232077 | 0.00052004 | 0.00378167 | SHE4 | XV | 397735 | 400104 | -1 |
| YOR036W | 387.047932 | -0.7063837 | 0.49794929 | 3.15061344 | 0.20694407 | 0.36890584 | PEP12 | XV | 400348 | 401214 | 1 |

| | | | | | | | | | | | |
|---|---|---|---|---|---|---|---|---|---|---|---|
| YOR037W | 21.1699019 | 0.62307291 | 0.86944676 | 1.25976049 | 0.53265559 | 0.68718267 | CYC2 | XV | 401555 | 402655 | 1 |
| YOR038C | 41.3166425 | 0.46048846 | 0.8020181 | 5.16101052 | 0.07573573 | 0.17861909 | HIR2 | XV | 402761 | 405388 | -1 |
| YOR039W | 22.7492578 | -0.5710954 | 1.02874466 | 1.34197831 | 0.51120267 | 0.66829159 | CKB2 | XV | 405768 | 406544 | 1 |
| YOR040W | 172.030989 | 0.48508979 | 0.52195001 | 0.86181094 | 0.64992034 | 0.78139046 | GLO4 | XV | 407064 | 407921 | 1 |
| YOR042W | 149.347648 | -0.4445207 | 0.5234563 | 1.34635067 | 0.51008631 | 0.66786132 | CUE5 | XV | 408425 | 409660 | 1 |
| YOR043W | 47.0034561 | 0.10969012 | 0.74380429 | 0.9074327 | 0.6352629 | 0.77207645 | WHI2 | XV | 410870 | 412330 | 1 |
| YOR044W | 22.2652151 | 2.24582964 | 0.85932544 | 16.3364178 | 0.00028353 | 0.00232853 | IRC23 | XV | 413007 | 413480 | 1 |
| YOR045W | 41.9821184 | -1.0398464 | 0.71088677 | 4.20277091 | 0.12228689 | 0.25314304 | TOM6 | XV | 413852 | 414037 | 1 |
| YOR046C | 27.8629602 | -0.0373365 | 0.67054681 | 0.16056615 | 0.92285507 | 0.9514478 | DBP5 | XV | 414459 | 415907 | -1 |
| YOR047C | 16.6471161 | 0.47247084 | 0.8305616 | 2.27453001 | 0.32069492 | 0.49318024 | STD1 | XV | 416347 | 417681 | -1 |
| YOR048C | 62.0833798 | -1.0922414 | 0.56128237 | 3.78314289 | 0.15083459 | 0.2938534 | RAT1 | XV | 418630 | 421650 | -1 |
| YOR049C | 1879.43074 | 0.61558004 | 0.49551829 | 4.8955788 | 0.08648456 | 0.19807974 | NA | XV | 422668 | 423732 | -1 |
| YOR051C | 84.2332694 | 0.55962635 | 0.59467481 | 3.58844437 | 0.16625672 | 0.31551177 | ETT1 | XV | 424846 | 426084 | -1 |
| YOR052C | 59.8399416 | -1.0904264 | 0.61198535 | 13.6240747 | 0.00110045 | 0.00712707 | TMC1 | XV | 426772 | 427224 | -1 |
| YOR054C | 129.83011 | 1.9575319 | 0.62374447 | 9.28074133 | 0.00965412 | 0.03839103 | VHS3 | XV | 427833 | 429857 | -1 |
| YOR056C | 79.495714 | -0.4388844 | 0.56804964 | 0.64133662 | 0.72566391 | 0.83456178 | NOB1 | XV | 430247 | 431626 | -1 |
| YOR057W | 52.0609252 | 0.92501162 | 0.78372281 | 1.40382056 | 0.49563759 | 0.65579106 | SGT1 | XV | 432186 | 433373 | 1 |
| YOR058C | 782.967987 | 1.54746751 | 0.55656 | 22.8769491 | 1.08E-05 | 0.0001408 | ASE1 | XV | 433688 | 436345 | -1 |
| YOR059C | 23.6992291 | 0.81174623 | 0.8089103 | 5.32960872 | 0.06961297 | 0.16759214 | LPL1 | XV | 438906 | 440258 | -1 |
| YOR060C | 19.9926872 | -1.6808262 | 0.88333605 | 3.63826325 | 0.16216651 | 0.31078956 | SLD7 | XV | 440390 | 441163 | -1 |
| YOR061W | 30.0412746 | -0.0746737 | 0.71755708 | 6.14931215 | 0.04620552 | 0.12305885 | CKA2 | XV | 441534 | 442553 | 1 |
| YOR062C | 297.309682 | 0.91650315 | 0.6424643 | 5.78109215 | 0.05554587 | 0.14149168 | NA | XV | 442725 | 443531 | -1 |
| YOR063W | 220.199076 | -0.8157987 | 0.75109481 | 14.594028 | 0.00067756 | 0.004761 | RPL3 | XV | 444686 | 445849 | 1 |
| YOR064C | 178.261766 | -1.3979104 | 0.53035338 | 12.7903174 | 0.00166962 | 0.00979022 | YNG1 | XV | 446079 | 446738 | -1 |
| YOR065W | 184.811555 | -0.2826552 | 0.56959897 | 14.655679 | 0.00065699 | 0.00463727 | CYT1 | XV | 447439 | 448368 | 1 |
| YOR066W | 80.4108603 | 0.96403695 | 0.55842933 | 6.4449342 | 0.03985661 | 0.11116077 | MSA1 | XV | 449436 | 451325 | 1 |
| YOR067C | 29.0104386 | -0.1808569 | 0.72824988 | 7.83204282 | 0.01992019 | 0.06575707 | ALG8 | XV | 451729 | 453462 | -1 |
| YOR068C | 1152.90432 | 0.37503142 | 0.53478855 | 0.54484192 | 0.76153362 | 0.85526961 | VAM10 | XV | 453869 | 454213 | -1 |
| YOR069W | 74.5800439 | -0.8221863 | 0.6003676 | 11.0415674 | 0.00400271 | 0.01926491 | VPS5 | XV | 453768 | 455795 | 1 |
| YOR070C | 135.838085 | 2.40738193 | 0.83179227 | 9.66365492 | 0.00797194 | 0.03311157 | GYP1 | XV | 455907 | 457820 | -1 |
| YOR071C | 484.169463 | -0.3061838 | 0.51518218 | 12.5386544 | 0.0018935 | 0.01073788 | NRT1 | XV | 459480 | 461276 | -1 |
| YOR072W | 138.780857 | -0.056252 | 0.56708605 | 7.3742069 | 0.02504444 | 0.07796092 | NA | XV | 461502 | 461816 | 1 |
| YOR073W | 85.4348211 | -0.7470065 | 0.60566306 | 10.7734562 | 0.00457692 | 0.02123429 | SGO1 | XV | 464771 | 466543 | 1 |

| | | | | | | | | | | | |
|---|---|---|---|---|---|---|---|---|---|---|---|
| YOR074C | 17.9526262 | -1.2305164 | 0.86665778 | 5.78449733 | 0.05545138 | 0.14137538 | CDC21 | XV | 466675 | 467589 | -1 |
| YOR075W | 15.2867584 | 0.66221349 | 0.92067088 | 4.09162754 | 0.12927495 | 0.26337155 | UFE1 | XV | 468212 | 469252 | 1 |
| YOR076C | 9.57473995 | -1.0726186 | 0.99086007 | 2.24670402 | 0.32518793 | 0.49807536 | SKI7 | XV | 469377 | 471620 | -1 |
| YOR078W | 172.529836 | 1.92798 | 0.82853611 | 8.10682954 | 0.01736298 | 0.05959198 | BUD21 | XV | 472725 | 473369 | 1 |
| YOR079C | 41.4516387 | -1.1469131 | 0.61046998 | 12.8141145 | 0.00164987 | 0.00975767 | ATX2 | XV | 473475 | 474416 | -1 |
| YOR080W | 96.4419123 | -0.2440441 | 0.61755211 | 8.82153097 | 0.01214588 | 0.04558315 | DIA2 | XV | 474594 | 476792 | 1 |
| YOR081C | 257.22113 | 1.41488296 | 0.65796452 | 4.7018752 | 0.09527979 | 0.21128685 | TGL5 | XV | 476939 | 479188 | -1 |
| YOR083W | 14.9585073 | -0.1472676 | 0.80560685 | 4.55866132 | 0.10235269 | 0.22279026 | WHI5 | XV | 479533 | 480420 | 1 |
| YOR084W | 15.7058557 | -0.5844242 | 0.79712251 | 1.39006286 | 0.49905876 | 0.65860324 | LPX1 | XV | 480587 | 481750 | 1 |
| YOR085W | 58.8530464 | 0.89002045 | 0.60865245 | 2.37319715 | 0.30525781 | 0.47807485 | OST3 | XV | 482033 | 483085 | 1 |
| YOR086C | 116.515581 | 0.34780845 | 0.54025041 | 2.0105741 | 0.36593958 | 0.53926957 | TCB1 | XV | 483219 | 486779 | -1 |
| YOR087W | 26.0652891 | -0.0203193 | 0.78236128 | 4.80103965 | 0.09067081 | 0.20427392 | YVC1 | XV | 487707 | 489734 | 1 |
| YOR089C | 113.666329 | 0.41918919 | 0.55506099 | 1.43544608 | 0.48786184 | 0.64951024 | VPS21 | XV | 490196 | 490828 | -1 |
| YOR090C | 388.983381 | -0.8464925 | 0.50321748 | 5.72116153 | 0.05723551 | 0.14501691 | PTC5 | XV | 491123 | 492841 | -1 |
| YOR091W | 89.5624656 | -0.4650895 | 0.53532996 | 9.62742832 | 0.00811765 | 0.0335244 | TMA46 | XV | 493433 | 494470 | 1 |
| YOR092W | 146.795003 | -0.5714953 | 0.50063357 | 1.89400398 | 0.38790222 | 0.56008824 | ECM3 | XV | 495127 | 496968 | 1 |
| YOR093C | 43.6637967 | -0.3464497 | 0.62748434 | 2.82186295 | 0.24391598 | 0.41379241 | CMR2 | XV | 497506 | 502452 | -1 |
| YOR094W | 281.618257 | -1.4315225 | 0.58115274 | 37.3435535 | 7.78E-09 | 2.43E-07 | ARF3 | XV | 502795 | 503346 | 1 |
| YOR095C | 36.2023375 | -0.5008184 | 0.68487062 | 5.29342558 | 0.07088384 | 0.17022706 | RKI1 | XV | 503552 | 504328 | -1 |
| YOR096W | 58.791797 | 0.20282878 | 0.59593328 | 15.2394004 | 0.00049069 | 0.00363486 | RPS7A | XV | 505794 | 506767 | 1 |
| YOR097C | 353.146468 | 0.36879534 | 0.58368421 | 0.46534496 | 0.79241306 | 0.87392122 | NA | XV | 506978 | 507505 | -1 |
| YOR098C | 185.753282 | 2.33181001 | 0.57113894 | 22.5908543 | 1.24E-05 | 0.00015922 | NUP1 | XV | 507948 | 511178 | -1 |
| YOR099W | 37.9409054 | 0.66247684 | 0.80424413 | 4.73544232 | 0.093694 | 0.20905135 | KTR1 | XV | 511825 | 513006 | 1 |
| YOR101W | 27.7377992 | -0.7821562 | 0.72846871 | 1.1757923 | 0.55549473 | 0.70703257 | RAS1 | XV | 515244 | 516173 | 1 |
| YOR103C | 51.4427627 | -0.1328102 | 0.58045129 | 1.99739885 | 0.36835821 | 0.54145571 | OST2 | XV | 516449 | 516841 | -1 |
| YOR104W | 34.7964257 | 0.0229696 | 0.63715894 | 0.01335669 | 0.99334391 | 0.9947183 | PIN2 | XV | 517642 | 518490 | 1 |
| YOR106W | 25.70349 | -0.2711197 | 0.82619373 | 19.8764669 | 4.83E-05 | 0.0005002 | VAM3 | XV | 519121 | 519972 | 1 |
| YOR107W | 115.318134 | -0.9494407 | 0.52367936 | 25.3123462 | 3.19E-06 | 4.88E-05 | RGS2 | XV | 521353 | 522282 | 1 |
| YOR108W | 35.084505 | -0.635902 | 0.70120395 | 4.78351079 | 0.09146898 | 0.20551237 | LEU9 | XV | 523027 | 524841 | 1 |
| YOR109W | 60.4248504 | 0.39198858 | 0.62739785 | 0.72387426 | 0.69632614 | 0.81659461 | INP53 | XV | 525278 | 528601 | 1 |
| YOR110W | 49.9414387 | 0.58828493 | 0.59516889 | 2.65383197 | 0.26529417 | 0.43749737 | TFC7 | XV | 528941 | 530248 | 1 |
| YOR111W | 8.94155505 | 1.16145362 | 1.07965382 | 1.55959418 | 0.45849904 | 0.62328904 | NA | XV | 530429 | 531127 | 1 |
| YOR112W | 28.2133388 | -0.3471126 | 0.66351366 | 0.97191094 | 0.6151092 | 0.75743987 | CEX1 | XV | 531508 | 533793 | 1 |

| | | | | | | | | | | | |
|---|---|---|---|---|---|---|---|---|---|---|---|
| YOR113W | 47.5948239 | -1.1518255 | 0.70257376 | 5.62678845 | 0.06000099 | 0.15000247 | AZF1 | XV | 534075 | 536819 | 1 |
| YOR114W | 28.0008121 | -0.17609 | 0.68358531 | 0.07851177 | 0.96150464 | 0.97258112 | NA | XV | 537570 | 538454 | 1 |
| YOR115C | 252.673376 | 0.5742979 | 0.57518893 | 18.7190639 | 8.61E-05 | 0.00082519 | TRS33 | XV | 538659 | 539465 | -1 |
| YOR116C | 259.581436 | 0.93970085 | 0.56759781 | 54.2317599 | 1.67E-12 | 1.14E-10 | RPO31 | XV | 539763 | 544145 | -1 |
| YOR117W | 40.0537699 | -0.1109613 | 1.08816818 | 0.18811669 | 0.91022965 | 0.94499367 | RPT5 | XV | 545029 | 546333 | 1 |
| YOR118W | 1711.36309 | -1.02992 | 0.55995852 | 3.62447747 | 0.16328817 | 0.31192296 | RTC5 | XV | 546857 | 548560 | 1 |
| YOR119C | 202.664739 | -0.2327822 | 0.53757833 | 16.0287185 | 0.00033068 | 0.00265185 | RIO1 | XV | 548792 | 550246 | -1 |
| YOR120W | 122.857448 | 2.69396487 | 0.83356942 | 12.4948302 | 0.00193545 | 0.01091164 | GCY1 | XV | 551114 | 552052 | 1 |
| YOR122C | 26.8072216 | 0.31725405 | 0.80012227 | 0.2213026 | 0.89525087 | 0.93531262 | PFY1 | XV | 552298 | 552887 | -1 |
| YOR123C | 112.68325 | 0.87699685 | 0.57481725 | 3.57059574 | 0.16774709 | 0.3174038 | LEO1 | XV | 553176 | 554570 | -1 |
| YOR124C | 90.8470071 | -0.4951301 | 0.53245614 | 7.31979215 | 0.02573519 | 0.07934331 | UBP2 | XV | 554824 | 558642 | -1 |
| YOR125C | 71.2024743 | 1.53154073 | 0.62220813 | 27.4432826 | 1.10E-06 | 1.93E-05 | CAT5 | XV | 559030 | 559731 | -1 |
| YOR126C | 90.8834209 | -0.4208185 | 0.61027816 | 0.52385325 | 0.76956749 | 0.85969434 | IAH1 | XV | 559961 | 560677 | -1 |
| YOR127W | 29.9978847 | 0.38283085 | 0.6959285 | 0.79186421 | 0.67305239 | 0.79774275 | RGA1 | XV | 561170 | 564193 | 1 |
| YOR128C | 20.3164846 | 0.56190663 | 0.80135414 | 2.73527202 | 0.25470838 | 0.42697206 | ADE2 | XV | 564476 | 566191 | -1 |
| YOR129C | 80.6224739 | 0.26824524 | 0.60958868 | 2.72461242 | 0.25606954 | 0.42826189 | AFI1 | XV | 566877 | 569558 | -1 |
| YOR130C | 12.4647715 | -0.4834328 | 0.84590797 | 1.67138018 | 0.43357517 | 0.60207676 | ORT1 | XV | 569929 | 570807 | -1 |
| YOR131C | 27.9307282 | 0.89815054 | 0.68687749 | 3.37292189 | 0.18517371 | 0.34112496 | NA | XV | 572182 | 572838 | -1 |
| YOR132W | 21.9483489 | 0.72717249 | 0.75490555 | 1.40126455 | 0.49627142 | 0.65603003 | VPS17 | XV | 573175 | 574830 | 1 |
| YOR133W | 40.8188868 | 1.01047342 | 0.78180988 | 7.57766831 | 0.02262196 | 0.07268654 | EFT1 | XV | 575098 | 577626 | 1 |
| YOR134W | 280.410025 | -0.0703673 | 0.50460949 | 17.034807 | 0.00019996 | 0.00170761 | BAG7 | XV | 578564 | 579793 | 1 |
| YOR136W | 32.1745453 | 0.07267263 | 0.69594212 | 1.61015094 | 0.44705419 | 0.61424863 | IDH2 | XV | 580250 | 581359 | 1 |
| YOR137C | 32.9294103 | -0.075914 | 0.72146337 | 0.24417685 | 0.8850701 | 0.9315066 | SIA1 | XV | 581813 | 583681 | -1 |
| YOR138C | 56.4637835 | 0.36597644 | 0.64999415 | 4.77753625 | 0.09174263 | 0.20596736 | RUP1 | XV | 584309 | 586324 | -1 |
| YOR140W | 117.391406 | -0.5511754 | 0.51870317 | 9.27597543 | 0.00967715 | 0.0384035 | SFL1 | XV | 586981 | 589281 | 1 |
| YOR141C | 29.8101027 | 0.90493881 | 0.67044519 | 2.32843329 | 0.3121671 | 0.48457038 | ARP8 | XV | 589942 | 592587 | -1 |
| YOR142W | 32.1719218 | 0.50105679 | 0.64525863 | 0.9488558 | 0.62224094 | 0.76265348 | LSC1 | XV | 593057 | 594046 | 1 |
| YOR143C | 25.5927815 | -0.5106176 | 0.7895072 | 2.06479553 | 0.35615197 | 0.52942744 | THI80 | XV | 601383 | 602342 | -1 |
| YOR144C | 9.6568257 | 0.03250956 | 0.94081573 | 0.15607803 | 0.92492834 | 0.95277599 | ELG1 | XV | 602717 | 605092 | -1 |
| YOR145C | 75.9011814 | -1.1601153 | 0.72273511 | 5.38493965 | 0.06771349 | 0.16390515 | PNO1 | XV | 605347 | 606171 | -1 |
| YOR147W | 102.105408 | 1.79587985 | 0.61901953 | 34.6646705 | 2.97E-08 | 8.07E-07 | MDM32 | XV | 606607 | 608475 | 1 |
| YOR148C | 81.8643288 | 1.12588094 | 0.64278969 | 5.32977326 | 0.06960724 | 0.16759214 | SPP2 | XV | 608640 | 609197 | -1 |
| YOR149C | 127.652308 | 0.93554981 | 0.62660468 | 33.307188 | 5.85E-08 | 1.50E-06 | SMP3 | XV | 609838 | 611388 | -1 |

| | | | | | | | | | | | |
|---|---|---|---|---|---|---|---|---|---|---|---|
| YOR150W | 11.9670922 | 1.49050473 | 1.01112262 | 2.67988026 | 0.26186135 | 0.43337041 | MRPL23 | XV | 611999 | 612490 | 1 |
| YOR151C | 47.5063648 | -0.4695814 | 0.69754604 | 3.30203635 | 0.19185447 | 0.34833408 | RPB2 | XV | 612997 | 616671 | -1 |
| YOR152C | 49.7589791 | -0.1580567 | 0.60422561 | 2.3346658 | 0.31119583 | 0.48393923 | ATG40 | XV | 617518 | 618288 | -1 |
| YOR153W | 130.824396 | 2.43706792 | 0.65353045 | 18.4575234 | 9.82E-05 | 0.00092128 | PDR5 | XV | 619840 | 624375 | 1 |
| YOR154W | 42.3355903 | 0.51262784 | 0.65413402 | 15.161946 | 0.00051006 | 0.0037336 | SLP1 | XV | 624729 | 626492 | 1 |
| YOR155C | 104.02862 | 2.31184736 | 0.70475659 | 16.1575686 | 0.00031005 | 0.00251202 | ISN1 | XV | 626628 | 627980 | -1 |
| YOR156C | 36.2573285 | 0.55698998 | 0.66820356 | 9.07738187 | 0.01068739 | 0.04141899 | NFI1 | XV | 628360 | 630540 | -1 |
| YOR157C | 25.6050167 | 0.09135532 | 0.74313738 | 0.56332813 | 0.75452711 | 0.85160078 | PUP1 | XV | 630966 | 631751 | -1 |
| YOR158W | 308.492114 | -0.2283306 | 0.57880675 | 18.2100943 | 0.0001111 | 0.0010215 | PET123 | XV | 632164 | 633120 | 1 |
| YOR159C | 34.9980821 | -0.5995811 | 0.63009826 | 10.5043752 | 0.00523605 | 0.02383391 | SME1 | XV | 633282 | 633566 | -1 |
| YOR160W | 21.6087521 | 0.5769036 | 0.80357525 | 1.02425768 | 0.59921858 | 0.74628427 | MTR10 | XV | 633839 | 636757 | 1 |
| YOR161C | 146.625809 | 0.15978254 | 0.50561664 | 8.9855316 | 0.01118965 | 0.04282094 | PNS1 | XV | 636939 | 638558 | -1 |
| YOR161C-C | 21.9711705 | -0.7456616 | 0.71837034 | 2.40812248 | 0.29997347 | 0.47325515 | NA | XV | 639121 | 639267 | -1 |
| YOR162C | 209.988775 | -1.4717938 | 0.49115704 | 9.86529757 | 0.00720739 | 0.03068439 | YRR1 | XV | 639560 | 641992 | -1 |
| YOR163W | 374.975174 | 0.1326104 | 0.51903582 | 9.08936097 | 0.01062357 | 0.04128218 | DDP1 | XV | 642741 | 643307 | 1 |
| YOR164C | 63.7567297 | 0.03000967 | 0.56108077 | 2.07897527 | 0.35363583 | 0.52744756 | GET4 | XV | 643394 | 644332 | -1 |
| YOR165W | 14.8736446 | -0.10521 | 0.87223096 | 0.57713636 | 0.74933571 | 0.8479727 | SEY1 | XV | 644566 | 646896 | 1 |
| YOR166C | 77.9183439 | -0.7327522 | 0.57790288 | 4.83252469 | 0.0892546 | 0.20266044 | SWT1 | XV | 647126 | 648502 | -1 |
| YOR167C | 72.3060609 | -0.3952281 | 0.63780137 | 16.9620718 | 0.00020736 | 0.00176305 | RPS28A | XV | 648804 | 649007 | -1 |
| YOR168W | 43.6567533 | -0.2479203 | 0.80537079 | 7.42394163 | 0.02442933 | 0.0767476 | GLN4 | XV | 649303 | 651732 | 1 |
| YOR171C | 10.4305168 | 0.07777586 | 1.03998633 | 0.18820841 | 0.91018791 | 0.94499367 | LCB4 | XV | 652010 | 653884 | -1 |
| YOR172W | 26.8654167 | -0.2994006 | 0.70077503 | 0.96913523 | 0.61596347 | 0.75801867 | YRM1 | XV | 654210 | 656570 | 1 |
| YOR173W | 150.63964 | -0.0019722 | 0.51812353 | 18.3341759 | 0.00010442 | 0.0009689 | DCS2 | XV | 657264 | 658325 | 1 |
| YOR174W | 38.9371932 | -0.4518213 | 0.69910235 | 0.5510834 | 0.75916078 | 0.85433253 | MED4 | XV | 658747 | 659601 | 1 |
| YOR175C | 18.6225794 | 0.05003809 | 0.77459339 | 0.16771492 | 0.91956232 | 0.94974417 | ALE1 | XV | 659815 | 661674 | -1 |
| YOR176W | 75.7990219 | -0.4086946 | 0.69915812 | 1.68778762 | 0.43003279 | 0.59923342 | HEM15 | XV | 662401 | 663582 | 1 |
| YOR177C | 16.4096436 | -0.9594219 | 0.81629994 | 1.9248871 | 0.38195841 | 0.55538403 | MPC54 | XV | 665785 | 667179 | -1 |
| YOR178C | 1465.05242 | -0.8972357 | 0.45698606 | 11.6192485 | 0.00299856 | 0.01524288 | GAC1 | XV | 667860 | 670241 | -1 |
| YOR179C | 5.6168239 | -1.2991677 | 1.28930806 | 1.85715673 | 0.39511502 | 0.5664635 | SYC1 | XV | 671845 | 672411 | -1 |
| YOR180C | 11.1607266 | -0.4695561 | 0.93181229 | 3.30847263 | 0.19123805 | 0.34765095 | DCI1 | XV | 674352 | 675167 | -1 |
| YOR181W | 63.0818532 | 1.77124886 | 0.76746074 | 5.2636222 | 0.07194804 | 0.17226374 | LAS17 | XV | 675939 | 677840 | 1 |
| YOR182C | 38.9373858 | -0.8005363 | 0.68965117 | 11.9543693 | 0.00253596 | 0.01338485 | RPS30A | XV | 678191 | 678793 | -1 |
| YOR183W | 43.3308802 | 0.1517833 | 0.62578646 | 1.61472266 | 0.44603345 | 0.61342843 | FYV12 | XV | 678872 | 679261 | 1 |

| | | | | | | | | | | | |
|---|---|---|---|---|---|---|---|---|---|---|---|
| YOR184W | 317.66801 | 1.65581861 | 0.62110235 | 56.7473054 | 4.76E-13 | 3.49E-11 | SER1 | XV | 679357 | 680544 | 1 |
| YOR185C | 270.52564 | 1.88028703 | 0.49445188 | 16.1565449 | 0.00031021 | 0.00251202 | GSP2 | XV | 681444 | 682106 | -1 |
| YOR187W | 48.6657299 | -0.3226832 | 0.72713202 | 0.69744246 | 0.7055898 | 0.82101385 | TUF1 | XV | 684030 | 685343 | 1 |
| YOR188W | 46.7652818 | 0.96152071 | 0.66463313 | 2.25225488 | 0.32428665 | 0.49763242 | MSB1 | XV | 685767 | 689180 | 1 |
| YOR189W | 12.7717702 | 0.44866138 | 0.91835677 | 1.32579915 | 0.51535486 | 0.67220199 | IES4 | XV | 689624 | 689974 | 1 |
| YOR190W | 444.087938 | -0.0136377 | 0.51523586 | 3.77644821 | 0.15134033 | 0.2946404 | SPR1 | XV | 690695 | 692032 | 1 |
| YOR191W | 61.0258604 | -0.0823066 | 0.60804457 | 0.36367352 | 0.83373743 | 0.9018008 | ULS1 | XV | 692475 | 697334 | 1 |
| YOR192C | 1390.24366 | 1.24018291 | 0.58832978 | 13.2590477 | 0.00132079 | 0.00824961 | THI72 | XV | 698768 | 700567 | -1 |
| YOR192C-B | 7.39366793 | -0.0810637 | 1.10856146 | 0.21605929 | 0.89760098 | 0.93705725 | NA | XV | 704419 | 709732 | -1 |
| YOR192C-C | 154.024734 | -0.1710483 | 0.54379141 | 1.03532837 | 0.59591086 | 0.7432839 | NA | XV | 703988 | 704224 | -1 |
| YOR193W | 251.973422 | -0.3346548 | 0.48753488 | 8.81994614 | 0.01215551 | 0.04558315 | PEX27 | XV | 710446 | 711576 | 1 |
| YOR194C | 397.306355 | 1.93253831 | 0.70630369 | 7.15594249 | 0.02793231 | 0.08418952 | TOA1 | XV | 711683 | 712543 | -1 |
| YOR195W | 33.1060478 | -0.322978 | 0.70832608 | 0.26691332 | 0.87506539 | 0.9267658 | SLK19 | XV | 712866 | 715331 | 1 |
| YOR196C | 74.7984565 | -0.4419682 | 0.56039976 | 3.58433929 | 0.16659832 | 0.31564276 | LIP5 | XV | 715593 | 716837 | -1 |
| YOR197W | 101.853367 | 1.08779139 | 0.67633498 | 2.58196599 | 0.27500033 | 0.44841232 | MCA1 | XV | 717086 | 718384 | 1 |
| YOR198C | 157.167293 | 1.06856076 | 0.62013026 | 3.02763963 | 0.22006775 | 0.38460377 | BFR1 | XV | 718653 | 720065 | -1 |
| YOR201C | 130.22359 | 2.00730014 | 0.58586376 | 26.6366356 | 1.64E-06 | 2.74E-05 | MRM1 | XV | 720470 | 721708 | -1 |
| YOR202W | 228.175575 | -0.4728035 | 0.53924852 | 9.04501516 | 0.01086175 | 0.04192011 | HIS3 | XV | 721946 | 722608 | 1 |
| YOR204W | 43.5909786 | -0.6039623 | 0.69398545 | 5.72401536 | 0.0571539 | 0.14493103 | DED1 | XV | 722911 | 724725 | 1 |
| YOR205C | 16.3792919 | -0.3913026 | 0.80124865 | 0.62030254 | 0.73333602 | 0.83764363 | GEP3 | XV | 725564 | 727234 | -1 |
| YOR206W | 52.5328667 | -1.295916 | 0.70052646 | 20.0899528 | 4.34E-05 | 0.00045692 | NOC2 | XV | 727512 | 729644 | 1 |
| YOR207C | 56.3565503 | 0.66394598 | 0.61453764 | 1.82478297 | 0.40156274 | 0.57224915 | RET1 | XV | 730008 | 733457 | -1 |
| YOR208W | 75.5377254 | 1.95249402 | 0.64148775 | 20.0576829 | 4.41E-05 | 0.00046183 | PTP2 | XV | 733925 | 736177 | 1 |
| YOR209C | 35.15554 | 0.85813961 | 1.06407947 | 1.24109116 | 0.53765103 | 0.69208525 | NPT1 | XV | 736437 | 737726 | -1 |
| YOR211C | 18.9390104 | -0.2910114 | 0.78762088 | 0.27312282 | 0.87235275 | 0.92527295 | MGM1 | XV | 738924 | 741569 | -1 |
| YOR212W | 63.1175787 | -0.4047346 | 0.61803826 | 11.3374463 | 0.00345227 | 0.01701161 | STE4 | XV | 742910 | 744181 | 1 |
| YOR213C | 318.991893 | -1.5944289 | 0.4971684 | 12.0969448 | 0.00236147 | 0.01267182 | SAS5 | XV | 744535 | 745281 | -1 |
| YOR214C | 20.7181688 | -1.6503179 | 0.79819406 | 4.52314219 | 0.10418667 | 0.22564606 | SPR2 | XV | 745593 | 746303 | -1 |
| YOR215C | 11.3516725 | -0.0141937 | 0.96180269 | 0.03750069 | 0.98142435 | 0.98585131 | AIM41 | XV | 746725 | 747282 | -1 |
| YOR216C | 17.9759537 | 0.46853469 | 0.7605618 | 1.89040623 | 0.38860063 | 0.56060856 | RUD3 | XV | 747525 | 748979 | -1 |
| YOR217W | 31.7678766 | 0.59028241 | 0.68430612 | 1.71132191 | 0.42500219 | 0.59510585 | RFC1 | XV | 749301 | 751886 | 1 |
| YOR219C | 51.5677574 | -0.8316271 | 0.62824983 | 6.38027401 | 0.04116623 | 0.11377206 | STE13 | XV | 752214 | 755009 | -1 |
| YOR220W | 23.7660617 | 3.09674187 | 0.96320221 | 12.0795241 | 0.00238213 | 0.01274723 | RCN2 | XV | 755328 | 756125 | 1 |

| | | | | | | | | | | | |
|---|---|---|---|---|---|---|---|---|---|---|---|
| YOR221C | 26.1487751 | 0.97628567 | 0.85206073 | 5.33909458 | 0.06928358 | 0.16707703 | MCT1 | XV | 756476 | 757558 | -1 |
| YOR222W | 47.7326645 | 0.28150787 | 0.69007368 | 2.71464372 | 0.25734907 | 0.42958807 | ODC2 | XV | 758330 | 759253 | 1 |
| YOR223W | 170.241113 | -0.8135724 | 0.55648841 | 18.6655585 | 8.85E-05 | 0.00084395 | DSC3 | XV | 759782 | 760660 | 1 |
| YOR224C | 26.2863965 | 0.67281541 | 0.76525007 | 3.86969881 | 0.14444602 | 0.28495484 | RPB8 | XV | 760825 | 761265 | -1 |
| YOR226C | 25.5473901 | -0.4919446 | 0.72498383 | 0.6346967 | 0.72807708 | 0.83592431 | ISU2 | XV | 761614 | 762084 | -1 |
| YOR227W | 59.8373859 | 0.29606728 | 0.63916832 | 1.31443813 | 0.51829066 | 0.67466343 | HER1 | XV | 762825 | 766565 | 1 |
| YOR228C | 38.6345038 | -0.9892788 | 0.71254593 | 12.6880406 | 0.00175722 | 0.01017432 | MCP1 | XV | 766869 | 767777 | -1 |
| YOR229W | 10.7809174 | -0.4345685 | 0.93620962 | 1.73730867 | 0.4195157 | 0.59032562 | WTM2 | XV | 768409 | 769812 | 1 |
| YOR230W | 395.068118 | -1.3091478 | 0.52203364 | 7.36814153 | 0.02512051 | 0.07811371 | WTM1 | XV | 770800 | 772113 | 1 |
| YOR231W | 1030.94928 | 0.99701844 | 0.50885444 | 119.881288 | 9.29E-27 | 1.17E-23 | MKK1 | XV | 772601 | 774127 | 1 |
| YOR232W | 14.4879354 | -1.1389869 | 1.00283609 | 5.34224231 | 0.06917463 | 0.16688379 | MGE1 | XV | 774573 | 775259 | 1 |
| YOR233W | 80.167473 | 0.51798224 | 0.58895447 | 23.3042466 | 8.70E-06 | 0.00011634 | KIN4 | XV | 775846 | 778248 | 1 |
| YOR234C | 12.0094995 | -0.2914109 | 1.07778757 | 2.90523556 | 0.23395704 | 0.40148525 | RPL33B | XV | 778555 | 779405 | -1 |
| YOR235W | 11.4588188 | 2.58632959 | 1.29990373 | 5.61410393 | 0.06038274 | 0.15063165 | NA | XV | 779870 | 780184 | 1 |
| YOR236W | 196.569903 | 2.67304012 | 0.59653275 | 49.3317793 | 1.94E-11 | 1.07E-09 | DFR1 | XV | 780906 | 781541 | 1 |
| YOR237W | 18.218713 | 3.06910714 | 1.23429979 | 6.08453772 | 0.04772648 | 0.12600836 | HES1 | XV | 781994 | 783298 | 1 |
| YOR238W | 185.911328 | 2.53958689 | 0.55023889 | 27.8491641 | 8.97E-07 | 1.62E-05 | NA | XV | 783677 | 784588 | 1 |
| YOR239W | 87.8929727 | 0.32831964 | 0.71456988 | 0.51618439 | 0.772524 | 0.86150115 | ABP140 | XV | 784857 | 786744 | 1 |
| YOR241W | 21.9665429 | 0.22314545 | 0.79512905 | 3.38472097 | 0.18408448 | 0.33965875 | MET7 | XV | 786995 | 788641 | 1 |
| YOR242C | 381.65876 | 0.83649579 | 0.70332048 | 10.2464273 | 0.00595685 | 0.02634848 | SSP2 | XV | 788742 | 789857 | -1 |
| YOR243C | 58.4888433 | -0.9770208 | 0.63661302 | 4.40760236 | 0.11038278 | 0.23522866 | PUS7 | XV | 790211 | 792241 | -1 |
| YOR244W | 9.56326399 | -0.29124 | 0.93995374 | 0.24410041 | 0.88510393 | 0.9315066 | ESA1 | XV | 792531 | 793868 | 1 |
| YOR245C | 18.4755076 | -0.0215347 | 0.83550355 | 0.126555 | 0.93868295 | 0.95995205 | DGA1 | XV | 794076 | 795332 | -1 |
| YOR246C | 112.788849 | 0.54122776 | 0.55409056 | 14.859082 | 0.00059346 | 0.00424214 | ENV9 | XV | 795801 | 796793 | -1 |
| YOR247W | 250.324165 | 0.00064723 | 0.52457035 | 7.26854877 | 0.02640309 | 0.08080014 | SRL1 | XV | 797676 | 798308 | 1 |
| YOR249C | 71.9222831 | 0.30881315 | 0.6308434 | 9.83384892 | 0.00732161 | 0.03110209 | APC5 | XV | 798674 | 800731 | -1 |
| YOR250C | 28.1231595 | 0.33806601 | 0.78689396 | 1.00063012 | 0.6063396 | 0.75088539 | CLP1 | XV | 800970 | 802307 | -1 |
| YOR251C | 36.3483662 | 0.14984947 | 0.66624807 | 1.01300249 | 0.60260024 | 0.74824264 | TUM1 | XV | 802550 | 803464 | -1 |
| YOR252W | 14.9084759 | -1.2592807 | 0.8456855 | 7.94474565 | 0.0188287 | 0.06320157 | TMA16 | XV | 803667 | 804203 | 1 |
| YOR253W | 318.265424 | 0.32774667 | 0.47377164 | 0.99936574 | 0.60672304 | 0.75094621 | NAT5 | XV | 804377 | 804907 | 1 |
| YOR254C | 70.5239589 | 0.0292097 | 0.69847307 | 0.01092097 | 0.9945544 | 0.99571207 | SEC63 | XV | 805032 | 807023 | -1 |
| YOR256C | 16.3779748 | 1.14100802 | 0.8192524 | 2.40303043 | 0.30073818 | 0.47389383 | TRE2 | XV | 808254 | 810683 | -1 |
| YOR257W | 24.6494025 | -0.0237794 | 0.75615987 | 0.45136026 | 0.79797331 | 0.8780436 | CDC31 | XV | 811008 | 811493 | 1 |

| | | | | | | | | | | | |
|---|---|---|---|---|---|---|---|---|---|---|---|
| YOR258W | 188.973747 | 1.71085146 | 0.69637448 | 10.4267891 | 0.00544317 | 0.02452601 | HNT3 | XV | 811671 | 812324 | 1 |
| YOR259C | 247.807576 | 0.96781253 | 0.56520504 | 5.25878263 | 0.07212235 | 0.17248591 | RPT4 | XV | 812395 | 813708 | -1 |
| YOR260W | 87.3072357 | -0.2600923 | 0.59357956 | 5.17013394 | 0.07539103 | 0.17809631 | GCD1 | XV | 813984 | 815720 | 1 |
| YOR261C | 132.746662 | 1.28884378 | 0.76752964 | 13.548602 | 0.00114277 | 0.00734365 | RPN8 | XV | 815915 | 816931 | -1 |
| YOR262W | 197.811163 | 0.37639495 | 0.52522304 | 7.45080627 | 0.02410338 | 0.0760123 | GPN2 | XV | 817292 | 818335 | 1 |
| YOR264W | 12.7254128 | -0.1522612 | 0.94492555 | 1.89044291 | 0.38859351 | 0.56060856 | DSE3 | XV | 818866 | 820158 | 1 |
| YOR265W | 8.76054654 | -0.0812759 | 1.00639182 | 1.67117319 | 0.43362005 | 0.60207676 | RBL2 | XV | 820454 | 820774 | 1 |
| YOR266W | 138.966954 | 0.91349526 | 0.53100137 | 29.881675 | 3.25E-07 | 6.64E-06 | PNT1 | XV | 821023 | 822294 | 1 |
| YOR267C | 332.18335 | 2.06694953 | 0.75405607 | 7.08094987 | 0.02899955 | 0.08672903 | HRK1 | XV | 822588 | 824867 | -1 |
| YOR268C | 20.8965296 | -0.2836238 | 0.76556003 | 5.08041632 | 0.07884998 | 0.18438668 | NA | XV | 825534 | 825932 | -1 |
| YOR269W | 16.8205995 | -0.8993116 | 0.77670848 | 5.83397135 | 0.05409651 | 0.13865373 | PAC1 | XV | 826385 | 827869 | 1 |
| YOR270C | 39.2635651 | 0.07591808 | 0.8512206 | 2.37336834 | 0.30523168 | 0.47807485 | VPH1 | XV | 828052 | 830574 | -1 |
| YOR271C | 44.1304952 | -1.0995245 | 0.61976387 | 8.14484705 | 0.01703605 | 0.05876231 | FSF1 | XV | 831059 | 832042 | -1 |
| YOR272W | 77.8258738 | 1.91242482 | 0.94654009 | 8.33359184 | 0.01550185 | 0.05493006 | YTM1 | XV | 832813 | 834195 | 1 |
| YOR273C | 79.80623 | 1.04034966 | 0.65034313 | 11.2397567 | 0.00362508 | 0.01780257 | TPO4 | XV | 834452 | 836431 | -1 |
| YOR274W | 494.837565 | 1.59048322 | 0.54978659 | 9.64921365 | 0.00802971 | 0.03330374 | MOD5 | XV | 837674 | 838960 | 1 |
| YOR275C | 95.9573223 | -0.4176516 | 0.61230715 | 3.35720548 | 0.18663457 | 0.34246214 | RIM20 | XV | 839084 | 841069 | -1 |
| YOR276W | 42.3178799 | -0.7740133 | 0.62215842 | 10.9961238 | 0.0040947 | 0.0195435 | CAF20 | XV | 841333 | 841818 | 1 |
| YOR278W | 90.4495369 | 0.86804875 | 0.67656451 | 1.92133905 | 0.38263662 | 0.55595132 | HEM4 | XV | 842817 | 843644 | 1 |
| YOR279C | 73.7903509 | 1.32918328 | 0.75364289 | 3.96469784 | 0.1377453 | 0.27549061 | RFM1 | XV | 843698 | 844630 | -1 |
| YOR280C | 104.466537 | 0.93361514 | 0.60852383 | 9.00986737 | 0.01105432 | 0.04244332 | FSH3 | XV | 844992 | 845792 | -1 |
| YOR281C | 17.4444953 | 0.57072026 | 0.75309324 | 2.85831193 | 0.23951099 | 0.40781483 | PLP2 | XV | 846269 | 847129 | -1 |
| YOR283W | 11.3482217 | -0.0660761 | 1.1401569 | 0.50652753 | 0.7762631 | 0.8639961 | NA | XV | 847453 | 848145 | 1 |
| YOR284W | 22.5444158 | -0.2972053 | 0.71107233 | 6.18867342 | 0.04530505 | 0.12155526 | HUA2 | XV | 848478 | 849209 | 1 |
| YOR285W | 85.4736968 | 0.2372425 | 0.58281725 | 0.55762178 | 0.75668298 | 0.8528864 | RDL1 | XV | 849635 | 850054 | 1 |
| YOR286W | 6.96077264 | -1.6242386 | 1.16221941 | 3.40213661 | 0.18248847 | 0.3376824 | RDL2 | XV | 850280 | 850729 | 1 |
| YOR287C | 162.866501 | 0.59146931 | 0.75334394 | 13.331876 | 0.00127356 | 0.00800643 | RRP36 | XV | 850937 | 851839 | -1 |
| YOR288C | 29.9976978 | -0.8353601 | 0.75831674 | 4.24705879 | 0.11960874 | 0.24884462 | MPD1 | XV | 852121 | 853077 | -1 |
| YOR289W | 158.001314 | 0.86796246 | 0.5782749 | 4.76964316 | 0.09210541 | 0.20646161 | NA | XV | 853357 | 854112 | 1 |
| YOR290C | 52.2205755 | 0.54696569 | 0.60408436 | 1.19288758 | 0.5507668 | 0.70271921 | SNF2 | XV | 855147 | 860258 | -1 |
| YOR291W | 65.0876991 | 1.10922749 | 0.71649983 | 2.3345869 | 0.3112081 | 0.48393923 | YPK9 | XV | 861175 | 865593 | 1 |
| YOR292C | 115.549892 | 1.01561718 | 0.71931679 | 9.81284664 | 0.0073989 | 0.03135777 | NA | XV | 865653 | 866582 | -1 |
| YOR293C-A | 380.830429 | 1.53005477 | 0.98770787 | 15.8534856 | 0.00036096 | 0.00284348 | NA | XV | 867998 | 868147 | -1 |

| | | | | | | | | | | | |
|---|---|---|---|---|---|---|---|---|---|---|---|
| YOR293W | 239.156947 | -0.7073948 | 0.55077389 | 30.385615 | 2.52E-07 | 5.35E-06 | RPS10A | XV | 867098 | 867852 | 1 |
| YOR294W | 58.6664118 | -1.3480512 | 0.76745347 | 12.7955128 | 0.00166529 | 0.00978886 | RRS1 | XV | 868340 | 868951 | 1 |
| YOR295W | 78.7493891 | 0.09263049 | 0.57639419 | 3.34675255 | 0.18761256 | 0.34351531 | UAF30 | XV | 869208 | 869894 | 1 |
| YOR296W | 55.8722376 | -0.4664112 | 0.59348971 | 3.03843137 | 0.21888349 | 0.38305402 | NA | XV | 870202 | 874071 | 1 |
| YOR297C | 112.907707 | -0.3354744 | 0.51998363 | 0.71761044 | 0.69851039 | 0.81736153 | TIM18 | XV | 874743 | 875321 | -1 |
| YOR298C-A | 243.404967 | 0.16295061 | 0.50266397 | 0.43757009 | 0.80349442 | 0.88210707 | MBF1 | XV | 877230 | 877685 | -1 |
| YOR298W | 1488.23207 | 1.0436702 | 0.80245384 | 2.06340401 | 0.35639985 | 0.52952403 | MUM3 | XV | 875599 | 877038 | 1 |
| YOR299W | 48.6533455 | -1.0550379 | 0.67466352 | 9.75635815 | 0.00761086 | 0.03195568 | BUD7 | XV | 878435 | 880675 | 1 |
| YOR301W | 241.745519 | 0.13784026 | 0.58990041 | 4.23787242 | 0.12015938 | 0.24944104 | RAX1 | XV | 880965 | 882272 | 1 |
| YOR302W | 11.7856481 | 1.03322297 | 1.04846119 | 2.99242231 | 0.22397717 | 0.38943778 | NA | XV | 882765 | 882842 | 1 |
| YOR303W | 70.9760742 | 1.00303583 | 0.63114275 | 7.38583168 | 0.02489929 | 0.07763431 | CPA1 | XV | 882899 | 884134 | 1 |
| YOR304C-A | 19.6012972 | 0.33144401 | 0.77425139 | 1.30715607 | 0.52018122 | 0.67636408 | BIL1 | XV | 888520 | 888750 | -1 |
| YOR304W | 55.2867461 | 0.30384525 | 0.64418558 | 0.61607526 | 0.73488766 | 0.83825839 | ISW2 | XV | 884514 | 887876 | 1 |
| YOR305W | 86.2857957 | 2.44512043 | 0.64215959 | 19.7467908 | 5.15E-05 | 0.00052711 | RRG7 | XV | 889022 | 889750 | 1 |
| YOR306C | 1233.74031 | -0.6198623 | 0.52742276 | 11.6939112 | 0.00288868 | 0.01477514 | MCH5 | XV | 889867 | 891432 | -1 |
| YOR307C | 77.8204442 | 0.59460354 | 0.62758389 | 24.8414885 | 4.03E-06 | 5.94E-05 | SLY41 | XV | 892731 | 894092 | -1 |
| YOR308C | 30.5567725 | -0.8769353 | 0.71587534 | 3.76824547 | 0.15196231 | 0.29555317 | SNU66 | XV | 894624 | 896387 | -1 |
| YOR310C | 575.962285 | -0.1955808 | 0.69139329 | 14.7777229 | 0.0006181 | 0.00440738 | NOP58 | XV | 896825 | 898360 | -1 |
| YOR311C | 33.3328283 | -0.7265193 | 0.71676795 | 4.0341632 | 0.13304317 | 0.26909961 | DGK1 | XV | 899056 | 899928 | -1 |
| YOR312C | 50.7697352 | -0.4294011 | 0.64047337 | 20.3637945 | 3.78E-05 | 0.00040962 | NA | XV | 900250 | 900767 | -1 |
| YOR314W | 33.5880509 | -2.0410482 | 0.6884644 | 14.9020271 | 0.00058085 | 0.00417263 | NA | XV | 903043 | 903372 | 1 |
| YOR315W | 26.3145683 | -1.21114 | 0.71711298 | 3.36171565 | 0.18621417 | 0.34184529 | SFG1 | XV | 904760 | 905800 | 1 |
| YOR316C | 36.4264501 | -1.0358191 | 0.78103126 | 3.24616053 | 0.19729006 | 0.35541675 | COT1 | XV | 906236 | 907555 | -1 |
| YOR317W | 347.076466 | -2.2850751 | 0.50581067 | 26.1820149 | 2.06E-06 | 3.32E-05 | FAA1 | XV | 909343 | 911445 | 1 |
| YOR319W | 122.582243 | -0.4286285 | 0.5743135 | 21.1998998 | 2.49E-05 | 0.00028739 | HSH49 | XV | 912822 | 913463 | 1 |
| YOR320C | 318.903594 | -0.7103089 | 0.47463416 | 6.73988442 | 0.03439162 | 0.09926596 | GNT1 | XV | 913619 | 915094 | -1 |
| YOR321W | 107.901295 | -0.7408421 | 0.59343034 | 2.72812802 | 0.25561982 | 0.42788053 | PMT3 | XV | 916030 | 918291 | 1 |
| YOR322C | 233.869053 | 0.68645686 | 0.55154891 | 80.5985127 | 3.15E-18 | 5.97E-16 | LDB19 | XV | 918606 | 921062 | -1 |
| YOR323C | 86.0330806 | -0.1523485 | 0.74424163 | 0.38438885 | 0.82514642 | 0.89541781 | PRO2 | XV | 921535 | 922905 | -1 |
| YOR324C | 15.6933135 | -0.9613886 | 0.81148834 | 1.64170385 | 0.4400566 | 0.60824247 | FRT1 | XV | 923232 | 925040 | -1 |
| YOR326W | 110.890262 | 1.25354477 | 0.54166058 | 8.82108914 | 0.01214856 | 0.04558315 | MYO2 | XV | 925721 | 930445 | 1 |
| YOR327C | 123.911574 | 0.02460756 | 0.51784376 | 5.3993884 | 0.06722607 | 0.16286148 | SNC2 | XV | 930734 | 931081 | -1 |
| YOR328W | 108.330122 | -0.5865423 | 0.57306901 | 7.10075765 | 0.02871376 | 0.08612855 | PDR10 | XV | 931803 | 936497 | 1 |

| | | | | | | | | | | | |
|---|---|---|---|---|---|---|---|---|---|---|---|
| YOR329C | 104.770624 | 0.96948558 | 0.60512357 | 2.67378266 | 0.26266093 | 0.43402019 | SCD5 | XV | 936731 | 939349 | -1 |
| YOR330C | 294.861232 | 2.31058997 | 0.54402785 | 54.2419324 | 1.67E-12 | 1.14E-10 | MIP1 | XV | 939621 | 943385 | -1 |
| YOR332W | 151.369254 | 0.41418714 | 0.53916303 | 4.65524321 | 0.09752743 | 0.21503573 | VMA4 | XV | 943656 | 944357 | 1 |
| YOR334W | 40.8388899 | 0.05717001 | 0.64385477 | 2.12654714 | 0.34532352 | 0.51852261 | MRS2 | XV | 944596 | 946008 | 1 |
| YOR335C | 39.9192451 | -0.4124627 | 0.80798374 | 7.49727574 | 0.0235498 | 0.07474302 | NA | XV | 946233 | 949109 | -1 |
| YOR336W | 59.3633252 | -0.4434063 | 0.60096247 | 1.74555503 | 0.41778952 | 0.58926319 | KRE5 | XV | 949773 | 953870 | 1 |
| YOR337W | 20.7307779 | 0.23951247 | 0.894052 | 3.50139436 | 0.17365283 | 0.32507061 | TEA1 | XV | 954344 | 956623 | 1 |
| YOR338W | 18.4152278 | 0.44852333 | 0.84551001 | 2.52488256 | 0.28296239 | 0.45678444 | NA | XV | 956898 | 957989 | 1 |
| YOR339C | 178.820893 | 0.71497338 | 0.56452858 | 31.0223664 | 1.83E-07 | 4.09E-06 | UBC11 | XV | 958362 | 958832 | -1 |
| YOR340C | 57.8572901 | -0.9976574 | 0.58289773 | 9.02154112 | 0.01098999 | 0.04228042 | RPA43 | XV | 959202 | 960182 | -1 |
| YOR341W | 58.5309308 | -1.3194495 | 0.73292824 | 5.423599 | 0.06641718 | 0.16169345 | RPA190 | XV | 960987 | 965981 | 1 |
| YOR342C | 38.1283567 | 0.47193208 | 0.70716987 | 2.07057768 | 0.35512379 | 0.52857757 | NA | XV | 966666 | 967625 | -1 |
| YOR343C | 24.6685287 | -0.8940581 | 0.75343937 | 3.07749504 | 0.21464978 | 0.37844769 | NA | XV | 968148 | 968474 | -1 |
| YOR343W-B | 13.6096358 | 0.79347188 | 0.88509414 | 0.89944489 | 0.63780515 | 0.77419116 | NA | XV | 970578 | 975891 | 1 |
| YOR344C | 75.0057882 | 0.41826924 | 0.59609505 | 24.0597024 | 5.96E-06 | 8.30E-05 | TYE7 | XV | 977194 | 978069 | -1 |
| YOR346W | 1060.24576 | 1.01465341 | 0.47921194 | 7.24044987 | 0.02677665 | 0.0815554 | REV1 | XV | 981828 | 984785 | 1 |
| YOR347C | 33.9033217 | 0.14159099 | 0.69938888 | 2.39408135 | 0.30208686 | 0.4745206 | PYK2 | XV | 984942 | 986462 | -1 |
| YOR348C | 62.7778027 | 0.0703778 | 0.5638149 | 0.18056917 | 0.91367113 | 0.94703828 | PUT4 | XV | 986899 | 988782 | -1 |
| YOR349W | 758.857754 | -0.9258627 | 0.45457498 | 23.4208758 | 8.21E-06 | 0.00011052 | CIN1 | XV | 989789 | 992833 | 1 |
| YOR350C | 88.1724224 | -1.1073241 | 0.62444479 | 10.801742 | 0.00451265 | 0.02098654 | MNE1 | XV | 992864 | 994855 | -1 |
| YOR351C | 207.598597 | -0.832044 | 0.53954906 | 6.96713931 | 0.03069764 | 0.09096178 | MEK1 | XV | 995018 | 996511 | -1 |
| YOR352W | 13.9161607 | -0.2069165 | 0.82274476 | 0.09389455 | 0.9541377 | 0.96767513 | TFB6 | XV | 997213 | 998244 | 1 |
| YOR353C | 23.1342607 | 0.19108096 | 0.83203828 | 0.54282172 | 0.76230323 | 0.85554094 | SOG2 | XV | 998453 | 1E+06 | -1 |
| YOR354C | 7.27177081 | -0.993515 | 1.13487182 | 0.87780975 | 0.64474211 | 0.77739625 | MSC6 | XV | 1001147 | 1E+06 | -1 |
| YOR355W | 51.322971 | 0.89241355 | 0.60907321 | 8.09194629 | 0.01749267 | 0.05989508 | GDS1 | XV | 1005137 | 1E+06 | 1 |
| YOR356W | 116.45056 | 0.61901984 | 0.61210866 | 12.9454597 | 0.001545 | 0.00934751 | CIR2 | XV | 1007221 | 1E+06 | 1 |
| YOR357C | 294.924605 | 3.07708309 | 1.63355668 | 4.30015413 | 0.11647518 | 0.24407937 | SNX3 | XV | 1009224 | 1E+06 | -1 |
| YOR358W | 17.2642865 | -0.3808534 | 0.805891 | 0.24512999 | 0.88464841 | 0.9315066 | HAP5 | XV | 1010161 | 1E+06 | 1 |
| YOR359W | 222.713578 | 1.25937071 | 0.59275039 | 79.8053352 | 4.68E-18 | 8.22E-16 | VTS1 | XV | 1011189 | 1E+06 | 1 |
| YOR360C | 6.43066135 | -0.5387387 | 1.25634375 | 0.39793314 | 0.81957729 | 0.89171017 | PDE2 | XV | 1013241 | 1E+06 | -1 |
| YOR361C | 289.58826 | -0.2025423 | 0.65186193 | 37.2423066 | 8.18E-09 | 2.52E-07 | PRT1 | XV | 1015361 | 1E+06 | -1 |
| YOR362C | 40.3953689 | 0.13778988 | 0.63951805 | 0.31499744 | 0.85427791 | 0.91360714 | PRE10 | XV | 1017880 | 1E+06 | -1 |
| YOR363C | 29.6052992 | 0.12377325 | 0.68290098 | 3.59329389 | 0.16585407 | 0.31516084 | PIP2 | XV | 1020222 | 1E+06 | -1 |

| | | | | | | | | | | | |
|---|---|---|---|---|---|---|---|---|---|---|---|
| YOR365C | 802.435808 | 0.19646313 | 0.50405605 | 2.00765264 | 0.36647451 | 0.53971939 | NA | XV | 1023461 | 1E+06 | -1 |
| YOR367W | 32.7706104 | -1.0172419 | 0.652668 | 3.70404022 | 0.15691985 | 0.30265354 | SCP1 | XV | 1026007 | 1E+06 | 1 |
| YOR368W | 465.952288 | -0.9163273 | 0.47829506 | 23.839668 | 6.66E-06 | 9.16E-05 | RAD17 | XV | 1026843 | 1E+06 | 1 |
| YOR369C | 127.98923 | -0.8442817 | 0.83001719 | 13.0682237 | 0.00145302 | 0.00892151 | RPS12 | XV | 1028194 | 1E+06 | -1 |
| YOR370C | 22.1651221 | 0.23704804 | 0.79207166 | 0.34260887 | 0.84256503 | 0.90781117 | MRS6 | XV | 1029183 | 1E+06 | -1 |
| YOR371C | 128.10914 | -1.2187321 | 0.5276769 | 15.9403662 | 0.00034562 | 0.00274125 | GPB1 | XV | 1031489 | 1E+06 | -1 |
| YOR372C | 55.273807 | 0.3703246 | 0.60375699 | 5.86874346 | 0.05316411 | 0.13694173 | NDD1 | XV | 1034807 | 1E+06 | -1 |
| YOR373W | 51.6882843 | -0.3070703 | 0.61747959 | 0.51624752 | 0.77249962 | 0.86150115 | NUD1 | XV | 1036834 | 1E+06 | 1 |
| YOR374W | 84.471092 | -2.8042614 | 0.58297807 | 39.8132823 | 2.26E-09 | 7.75E-08 | ALD4 | XV | 1039840 | 1E+06 | 1 |
| YOR375C | 46.2371931 | -1.6572401 | 0.88082866 | 12.4598397 | 0.00196961 | 0.01104002 | GDH1 | XV | 1041678 | 1E+06 | -1 |
| YOR376W | 73.1900499 | -0.3642475 | 0.57038133 | 1.64300973 | 0.43976936 | 0.60823052 | NA | XV | 1043191 | 1E+06 | 1 |
| YOR376W-A | 35.65045 | -1.0461895 | 0.67798419 | 4.00025915 | 0.13531775 | 0.27204506 | NA | XV | 1045196 | 1E+06 | 1 |
| YOR377W | 108.561375 | -1.5158855 | 0.61314046 | 13.7262367 | 0.00104565 | 0.0068333 | ATF1 | XV | 1046226 | 1E+06 | 1 |
| YOR378W | 722.839546 | -0.3478675 | 0.52356432 | 3.37091733 | 0.18535939 | 0.34124989 | AMF1 | XV | 1049511 | 1E+06 | 1 |
| YOR380W | 93.8767858 | -1.1746115 | 0.551533 | 6.19883008 | 0.04507556 | 0.12108273 | RDR1 | XV | 1051290 | 1E+06 | 1 |
| YOR381W | 38.4545213 | -1.4724462 | 0.70638042 | 4.22509366 | 0.12092959 | 0.25060211 | FRE3 | XV | 1055545 | 1E+06 | 1 |
| YOR382W | 41.9240109 | 0.00735413 | 0.6287783 | 0.20619581 | 0.90203865 | 0.93824416 | FIT2 | XV | 1059531 | 1E+06 | 1 |
| YOR384W | 3179.00303 | 0.27069538 | 0.48215235 | 1.91085396 | 0.38464787 | 0.55747464 | FRE5 | XV | 1061564 | 1E+06 | 1 |
| YOR385W | 25.4800499 | 1.20870411 | 0.72801455 | 3.29559466 | 0.1924734 | 0.34902003 | NA | XV | 1065043 | 1E+06 | 1 |
| YOR386W | 13.2884771 | -0.5588153 | 0.84772388 | 0.63629785 | 0.72749444 | 0.83558675 | PHR1 | XV | 1066839 | 1E+06 | 1 |
| YOR387C | 376.852227 | 0.0752591 | 0.50831298 | 0.09395782 | 0.95410752 | 0.96767513 | NA | XV | 1069621 | 1E+06 | -1 |
| YOR388C | 41.9995532 | -0.2476943 | 0.75415063 | 0.45420016 | 0.79684103 | 0.8771311 | FDH1 | XV | 1071793 | 1E+06 | -1 |
| YOR389W | 259.975014 | 0.34695444 | 0.48612152 | 17.4915881 | 0.00015913 | 0.00140024 | NA | XV | 1074213 | 1E+06 | 1 |
| YOR390W | 20.3428834 | 0.25815139 | 0.86798219 | 3.98832565 | 0.13612757 | 0.27319883 | FEX1 | XV | 1076784 | 1E+06 | 1 |
| YOR391C | 7.16797393 | -0.3314869 | 1.16155618 | 1.06769645 | 0.58634424 | 0.73388092 | HSP33 | XV | 1078545 | 1E+06 | -1 |
| YPL001W | 39.8364639 | 1.47643356 | 0.71108518 | 10.8667777 | 0.00436827 | 0.02052944 | HAT1 | XVI | 554605 | 555729 | 1 |
| YPL002C | 480.214076 | 1.23759875 | 0.66892683 | 9.76857287 | 0.00756452 | 0.03183036 | SNF8 | XVI | 553627 | 554328 | -1 |
| YPL003W | 962.45856 | 0.84761901 | 0.56525144 | 2.21491676 | 0.33039764 | 0.50355418 | ULA1 | XVI | 552020 | 553408 | 1 |
| YPL004C | 63.0880644 | 0.42490554 | 0.59470614 | 2.96074017 | 0.22755346 | 0.39362867 | LSP1 | XVI | 550632 | 551657 | -1 |
| YPL005W | 40.8617127 | -0.4133822 | 0.64297676 | 1.93540351 | 0.37995527 | 0.55386228 | AEP3 | XVI | 548486 | 550306 | 1 |
| YPL006W | 623.098925 | 1.45982263 | 0.50534529 | 66.1472076 | 4.33E-15 | 4.56E-13 | NCR1 | XVI | 544631 | 548143 | 1 |
| YPL007C | 25.5416857 | -0.7619488 | 0.74613579 | 2.17462906 | 0.33712061 | 0.51030806 | TFC8 | XVI | 542082 | 543848 | -1 |
| YPL008W | 124.553814 | 0.213815 | 0.59091419 | 21.4140883 | 2.24E-05 | 0.00026028 | CHL1 | XVI | 539385 | 541970 | 1 |

| | | | | | | | | | | | |
|---|---|---|---|---|---|---|---|---|---|---|---|
| YPL009C | 272.474234 | 1.08622428 | 0.79488085 | 1.82564716 | 0.40138927 | 0.57214275 | RQC2 | XVI | 535820 | 538936 | -1 |
| YPL010W | 9.02302525 | -0.1960713 | 1.03082068 | 0.88497912 | 0.64243505 | 0.77621415 | RET3 | XVI | 535018 | 535587 | 1 |
| YPL011C | 13.1722737 | -0.7592974 | 0.9267414 | 1.11384272 | 0.57297032 | 0.72277589 | TAF3 | XVI | 533642 | 534703 | -1 |
| YPL012W | 34.8792974 | -1.3794723 | 0.72692433 | 7.40961616 | 0.02460494 | 0.07694787 | RRP12 | XVI | 529723 | 533409 | 1 |
| YPL013C | 19.0866289 | 0.02592636 | 0.79712325 | 0.52216856 | 0.770216 | 0.86008691 | MRPS16 | XVI | 528985 | 529350 | -1 |
| YPL014W | 78.8043128 | -0.7463983 | 0.54369308 | 8.17193965 | 0.01680683 | 0.05813116 | CIP1 | XVI | 527547 | 528692 | 1 |
| YPL015C | 69.1421886 | 0.33117341 | 0.56215408 | 5.9313571 | 0.05152549 | 0.13385547 | HST2 | XVI | 525810 | 526883 | -1 |
| YPL016W | 39.3259789 | -0.2693625 | 0.65191903 | 0.85209487 | 0.65308536 | 0.78337772 | SWI1 | XVI | 521014 | 524958 | 1 |
| YPL017C | 979.03059 | -0.2000145 | 0.47173639 | 0.26535334 | 0.8757482 | 0.92712351 | IRC15 | XVI | 518734 | 520233 | -1 |
| YPL018W | 1127.89283 | 1.23779329 | 0.55641629 | 28.8905979 | 5.33E-07 | 1.03E-05 | CTF19 | XVI | 517651 | 518760 | 1 |
| YPL019C | 244.541202 | 0.47866148 | 0.49045707 | 15.7207094 | 0.00038574 | 0.00298585 | VTC3 | XVI | 514511 | 517018 | -1 |
| YPL020C | 21.263298 | 0.37918853 | 0.73543324 | 0.61767272 | 0.73430092 | 0.83775415 | ULP1 | XVI | 512313 | 514178 | -1 |
| YPL021W | 8.32123492 | 4.6612654 | 1.75254212 | 19.101024 | 7.12E-05 | 0.00070435 | ECM23 | XVI | 511101 | 511664 | 1 |
| YPL022W | 27.3856249 | 0.01684061 | 0.7199661 | 0.2446333 | 0.88486813 | 0.9315066 | RAD1 | XVI | 506697 | 509999 | 1 |
| YPL023C | 127.830725 | 1.17315921 | 0.68438254 | 5.97956542 | 0.05029836 | 0.13124269 | MET12 | XVI | 504339 | 506312 | -1 |
| YPL024W | 315.363001 | 0.69738267 | 0.61494115 | 5.30424528 | 0.07050141 | 0.16937889 | RMI1 | XVI | 503517 | 504242 | 1 |
| YPL026C | 183.38368 | 1.3400576 | 0.81859805 | 2.66480448 | 0.26384269 | 0.43572423 | SKS1 | XVI | 500675 | 502183 | -1 |
| YPL027W | 34.8098475 | -1.6228 | 0.70919448 | 5.84537775 | 0.05378886 | 0.13823236 | SMA1 | XVI | 499665 | 500402 | 1 |
| YPL028W | 74.7152412 | 0.21126711 | 0.73679916 | 0.45082049 | 0.7981887 | 0.87811373 | ERG10 | XVI | 498096 | 499292 | 1 |
| YPL029W | 15.0239213 | -0.077087 | 0.85366885 | 3.17733354 | 0.20419767 | 0.3649088 | SUV3 | XVI | 495506 | 497719 | 1 |
| YPL030W | 8.09327774 | -0.2536316 | 1.01492858 | 0.27248868 | 0.87262939 | 0.92527295 | TRM44 | XVI | 493543 | 495246 | 1 |
| YPL031C | 244.18573 | -1.8446489 | 0.58494552 | 27.6885378 | 9.72E-07 | 1.73E-05 | PHO85 | XVI | 492018 | 493037 | -1 |
| YPL032C | 65.9095271 | 0.87942657 | 0.64165523 | 2.93150283 | 0.23090442 | 0.3978978 | SVL3 | XVI | 488887 | 491364 | -1 |
| YPL033C | 105.743499 | 1.57390957 | 0.70159883 | 33.2044107 | 6.16E-08 | 1.56E-06 | SRL4 | XVI | 487362 | 488207 | -1 |
| YPL034W | 17.1663606 | 0.93474452 | 0.88194935 | 10.1087207 | 0.00638145 | 0.02778089 | NA | XVI | 486712 | 487209 | 1 |
| YPL036W | 328.364258 | 0.08344552 | 0.60670995 | 4.24438087 | 0.119769 | 0.24890972 | PMA2 | XVI | 482843 | 485686 | 1 |
| YPL037C | 36.9572784 | -0.6603252 | 0.74552491 | 9.8701957 | 0.00718976 | 0.03067501 | EGD1 | XVI | 481428 | 481901 | -1 |
| YPL038W | 27.4989747 | -0.3534571 | 0.6941728 | 0.30469444 | 0.85869007 | 0.91650698 | MET31 | XVI | 480535 | 481068 | 1 |
| YPL039W | 31.9558132 | 0.30870193 | 0.7378283 | 0.41676505 | 0.81189641 | 0.88731158 | NA | XVI | 479225 | 480175 | 1 |
| YPL040C | 15.5684484 | 0.12537328 | 0.86071618 | 1.03072869 | 0.59728294 | 0.74451415 | ISM1 | XVI | 475993 | 479001 | -1 |
| YPL041C | 9.95484099 | -1.44398 | 1.06060173 | 3.10000623 | 0.21224731 | 0.37592901 | MRX11 | XVI | 475118 | 475741 | -1 |
| YPL042C | 22.1754736 | 0.55135567 | 0.74664976 | 3.3409666 | 0.18815611 | 0.34377528 | SSN3 | XVI | 473040 | 474707 | -1 |
| YPL043W | 257.359002 | -0.8292333 | 0.57569667 | 4.11387037 | 0.12784519 | 0.26147074 | NOP4 | XVI | 469939 | 471996 | 1 |

| | | | | | | | | | | | |
|---|---|---|---|---|---|---|---|---|---|---|---|
| YPL045W | 71.8395613 | -0.2652658 | 0.54260768 | 0.27294112 | 0.872432 | 0.92527295 | VPS16 | XVI | 467260 | 469656 | 1 |
| YPL046C | 83.8585488 | 0.51106699 | 0.68311815 | 5.08363203 | 0.07872331 | 0.18416483 | ELC1 | XVI | 466644 | 466943 | -1 |
| YPL047W | 10.2339223 | -0.7438724 | 0.92023516 | 4.12704761 | 0.12700564 | 0.26058208 | SGF11 | XVI | 465962 | 466261 | 1 |
| YPL048W | 50.3629168 | 0.02817545 | 0.72811733 | 0.23149807 | 0.89069873 | 0.93317728 | CAM1 | XVI | 464401 | 465648 | 1 |
| YPL049C | 126.681952 | 0.58370259 | 0.5633727 | 9.01596063 | 0.0110207 | 0.04237041 | DIG1 | XVI | 462481 | 463839 | -1 |
| YPL050C | 56.4463265 | -0.2026839 | 0.67892017 | 6.34622041 | 0.04187316 | 0.11493835 | MNN9 | XVI | 460779 | 461966 | -1 |
| YPL051W | 13.1748276 | -0.248968 | 0.95371512 | 0.18734868 | 0.91057926 | 0.94518714 | ARL3 | XVI | 459963 | 460559 | 1 |
| YPL052W | 137.485912 | 0.75429446 | 0.53742084 | 2.45570972 | 0.29292026 | 0.46721992 | OAZ1 | XVI | 458799 | 459678 | 1 |
| YPL053C | 27.9643379 | 1.34912523 | 0.82523698 | 2.83521528 | 0.24229298 | 0.41140068 | KTR6 | XVI | 457118 | 458458 | -1 |
| YPL054W | 189.300702 | 0.64027841 | 0.52588914 | 1.51560911 | 0.46869429 | 0.63146966 | LEE1 | XVI | 455759 | 456664 | 1 |
| YPL055C | 164.646292 | 1.39300478 | 0.49782271 | 7.60984637 | 0.02226091 | 0.07177998 | LGE1 | XVI | 453992 | 454990 | -1 |
| YPL056C | 313.006661 | 1.30726557 | 0.48008975 | 11.6482708 | 0.00295536 | 0.01506296 | LCL1 | XVI | 453430 | 453735 | -1 |
| YPL057C | 16.9818611 | -0.2520273 | 0.78706474 | 1.42335963 | 0.49081902 | 0.65209778 | SUR1 | XVI | 451909 | 453057 | -1 |
| YPL058C | 38.1879325 | 0.0201875 | 0.65715644 | 1.0089982 | 0.60380795 | 0.74934389 | PDR12 | XVI | 445842 | 450377 | -1 |
| YPL059W | 42.8501553 | -0.5032443 | 0.7322549 | 1.54195058 | 0.46256172 | 0.62590146 | GRX5 | XVI | 444579 | 445031 | 1 |
| YPL060C-A | 12.1185018 | -0.6643992 | 0.94595002 | 0.53144476 | 0.76665193 | 0.85825884 | NA | XVI | 437333 | 442742 | -1 |
| YPL060W | 18.9381274 | 0.42404362 | 0.77286469 | 0.57616003 | 0.7497016 | 0.8479727 | MFM1 | XVI | 434523 | 435764 | 1 |
| YPL061W | 71.694741 | 0.11072014 | 0.61787722 | 7.87011772 | 0.01954455 | 0.06499882 | ALD6 | XVI | 432588 | 434090 | 1 |
| YPL063W | 54.8978124 | 0.2164936 | 0.6084782 | 3.05873509 | 0.21667266 | 0.38085449 | TIM50 | XVI | 429939 | 431369 | 1 |
| YPL064C | 50.4793807 | 1.69439918 | 0.87555124 | 5.09780291 | 0.07816749 | 0.18338253 | CWC27 | XVI | 428711 | 429616 | -1 |
| YPL065W | 83.9920713 | 1.28638831 | 0.74872307 | 5.7609506 | 0.05610809 | 0.14279817 | VPS28 | XVI | 427927 | 428655 | 1 |
| YPL066W | 116.261524 | -0.426704 | 0.55592561 | 2.87125595 | 0.23796588 | 0.40595829 | RGL1 | XVI | 426233 | 427672 | 1 |
| YPL067C | 92.3994418 | 1.60684412 | 0.54114633 | 10.0257389 | 0.00665179 | 0.02880618 | NA | XVI | 425251 | 425847 | -1 |
| YPL068C | 13.7410094 | -1.2606106 | 1.01611879 | 5.19809214 | 0.07434446 | 0.17629168 | NA | XVI | 424215 | 425096 | -1 |
| YPL069C | 259.067028 | 1.83694223 | 0.53117414 | 12.0120734 | 0.00246383 | 0.01306373 | BTS1 | XVI | 422885 | 423892 | -1 |
| YPL070W | 122.815856 | 0.83642754 | 0.60689274 | 8.9265848 | 0.01152436 | 0.04384102 | MUK1 | XVI | 420948 | 422786 | 1 |
| YPL071C | 23.1309044 | 0.69074864 | 0.73149732 | 2.87554199 | 0.23745646 | 0.4053281 | NA | XVI | 420048 | 420518 | -1 |
| YPL072W | 216.458191 | 1.98048462 | 0.73368072 | 6.85153161 | 0.03252436 | 0.09506111 | UBP16 | XVI | 418512 | 420011 | 1 |
| YPL074W | 72.7913575 | -0.022749 | 0.58672589 | 7.47466104 | 0.0238176 | 0.07531616 | YTA6 | XVI | 415763 | 418027 | 1 |
| YPL075W | 58.3034661 | -0.5319007 | 0.66368578 | 11.2061682 | 0.00368648 | 0.01802762 | GCR1 | XVI | 412254 | 415362 | 1 |
| YPL076W | 150.082019 | 1.42449004 | 0.60759606 | 20.3754236 | 3.76E-05 | 0.00040878 | GPI2 | XVI | 410443 | 411285 | 1 |
| YPL077C | 26.1346564 | 0.3905696 | 0.76625962 | 4.36257592 | 0.11289603 | 0.23943884 | NA | XVI | 409368 | 410090 | -1 |
| YPL078C | 38.0277542 | 1.25012229 | 0.76032225 | 2.72622846 | 0.25586272 | 0.42803963 | ATP4 | XVI | 408010 | 408744 | -1 |

| | | | | | | | | | | | |
|---|---|---|---|---|---|---|---|---|---|---|---|
| YPL079W | 38.5673722 | -0.8383161 | 0.75403286 | 14.5001281 | 0.00071013 | 0.00493003 | RPL21B | XVI | 406636 | 407539 | 1 |
| YPL080C | 557.027376 | 0.87787711 | 0.61643332 | 28.1951653 | 7.54E-07 | 1.40E-05 | NA | XVI | 406172 | 406498 | -1 |
| YPL081W | 46.1188934 | -0.5232065 | 0.85707624 | 13.7141136 | 0.00105201 | 0.00686709 | RPS9A | XVI | 404950 | 406044 | 1 |
| YPL082C | 20.7410377 | -0.4656242 | 0.72940305 | 3.90539782 | 0.1418906 | 0.28114371 | MOT1 | XVI | 398480 | 404083 | -1 |
| YPL083C | 55.8367687 | 1.15115103 | 0.9079465 | 4.43851446 | 0.10868981 | 0.23264843 | SEN54 | XVI | 396702 | 398105 | -1 |
| YPL084W | 458.497616 | -1.2732776 | 0.49721435 | 31.5457724 | 1.41E-07 | 3.26E-06 | BRO1 | XVI | 394038 | 396572 | 1 |
| YPL085W | 79.9950468 | 0.36166116 | 0.5420442 | 1.5303399 | 0.46525485 | 0.62778504 | SEC16 | XVI | 387067 | 393654 | 1 |
| YPL086C | 19.0850517 | -1.2676758 | 0.82310414 | 6.46665889 | 0.03942601 | 0.11038521 | ELP3 | XVI | 384773 | 386446 | -1 |
| YPL087W | 14.6098457 | 0.808831 | 1.16364283 | 0.90610901 | 0.63568349 | 0.7722634 | YDC1 | XVI | 383455 | 384408 | 1 |
| YPL088W | 66.7766802 | 3.4249415 | 0.69672858 | 28.5904699 | 6.19E-07 | 1.17E-05 | NA | XVI | 381965 | 382993 | 1 |
| YPL089C | 596.910242 | -0.8934838 | 0.71121448 | 6.01490848 | 0.04941732 | 0.12952753 | RLM1 | XVI | 379120 | 381150 | -1 |
| YPL090C | 58.8007764 | -0.9684146 | 0.89180022 | 13.9270425 | 0.00094576 | 0.0062942 | RPS6A | XVI | 377291 | 378395 | -1 |
| YPL091W | 133.376952 | 0.39021281 | 0.53129698 | 0.54678951 | 0.7607924 | 0.85483951 | GLR1 | XVI | 375502 | 376953 | 1 |
| YPL092W | 19.5299865 | -0.9367373 | 0.79586009 | 2.32593153 | 0.31255783 | 0.4849178 | SSU1 | XVI | 373793 | 375169 | 1 |
| YPL093W | 48.3463212 | -1.389257 | 0.70839511 | 14.0664303 | 0.00088209 | 0.00595257 | NOG1 | XVI | 370978 | 372921 | 1 |
| YPL094C | 88.8940085 | 0.60135061 | 0.57524014 | 1.22294546 | 0.54255125 | 0.69630051 | SEC62 | XVI | 369839 | 370663 | -1 |
| YPL095C | 434.005486 | -0.5091597 | 0.50758889 | 2.21594712 | 0.33022747 | 0.50342734 | EEB1 | XVI | 368071 | 369441 | -1 |
| YPL096C-A | 31.2846611 | 5.50365456 | 2.31986995 | 4.68071113 | 0.09629339 | 0.2132895 | ERI1 | XVI | 366529 | 366735 | -1 |
| YPL096W | 71.0354076 | 0.75577962 | 0.81847858 | 0.87670551 | 0.64509818 | 0.77750176 | PNG1 | XVI | 366927 | 368018 | 1 |
| YPL097W | 371.247 | 2.72597646 | 0.97061191 | 7.59451493 | 0.02243221 | 0.07215694 | MSY1 | XVI | 364952 | 366430 | 1 |
| YPL098C | 13.9549594 | 0.00270168 | 0.82638587 | 0.60610456 | 0.73856048 | 0.84040603 | MGR2 | XVI | 364386 | 364727 | -1 |
| YPL099C | 159.247904 | -0.2369439 | 0.51855302 | 0.82303683 | 0.66264332 | 0.79009572 | INA17 | XVI | 363520 | 364068 | -1 |
| YPL100W | 157.301126 | 1.22726998 | 0.75814447 | 4.47472658 | 0.10673957 | 0.22957732 | ATG21 | XVI | 361872 | 363362 | 1 |
| YPL101W | 23.2508073 | -0.3186283 | 0.70696274 | 1.07561778 | 0.58402652 | 0.7320878 | ELP4 | XVI | 360209 | 361579 | 1 |
| YPL103C | 27.2967526 | -0.4188754 | 0.79321765 | 3.92893625 | 0.14023045 | 0.27872788 | FMP30 | XVI | 358000 | 359406 | -1 |
| YPL104W | 53.6088984 | -0.9984761 | 0.61183706 | 11.4473097 | 0.00326775 | 0.01635285 | MSD1 | XVI | 355700 | 357676 | 1 |
| YPL105C | 121.539729 | -0.1098981 | 0.56732651 | 0.06969875 | 0.96575087 | 0.97467275 | SYH1 | XVI | 352863 | 355412 | -1 |
| YPL106C | 185.594199 | -0.0683528 | 0.63901179 | 0.33532661 | 0.84563851 | 0.90923806 | SSE1 | XVI | 350194 | 352275 | -1 |
| YPL107W | 15.7866973 | 0.91288894 | 0.83491453 | 1.2221154 | 0.54277647 | 0.69636068 | NA | XVI | 349119 | 349865 | 1 |
| YPL108W | 46.960818 | -0.7494965 | 0.62705415 | 4.3043658 | 0.11623016 | 0.24370198 | NA | XVI | 348446 | 348952 | 1 |
| YPL109C | 604.813862 | 2.25704327 | 0.49425843 | 53.9545242 | 1.92E-12 | 1.29E-10 | NA | XVI | 345264 | 347389 | -1 |
| YPL110C | 175.582545 | 0.66004732 | 0.58394294 | 1.56864827 | 0.45642808 | 0.62207609 | GDE1 | XVI | 341068 | 344739 | -1 |
| YPL111W | 68.3600467 | 0.76035345 | 0.61529074 | 1.85854235 | 0.39484138 | 0.56643498 | CAR1 | XVI | 339944 | 340945 | 1 |

| | | | | | | | | | | | |
|---|---|---|---|---|---|---|---|---|---|---|---|
| YPL112C | 42.0847528 | 1.43913568 | 0.73993835 | 3.79529466 | 0.14992092 | 0.29275789 | PEX25 | XVI | 337436 | 338620 | -1 |
| YPL113C | 529.530621 | -0.4556267 | 0.48860993 | 2.09783434 | 0.35031688 | 0.523786 | NA | XVI | 335953 | 337143 | -1 |
| YPL115C | 151.681311 | 1.25903137 | 0.57500712 | 38.1615086 | 5.17E-09 | 1.64E-07 | BEM3 | XVI | 332100 | 335486 | -1 |
| YPL116W | 61.0156048 | -0.1964704 | 0.57964586 | 0.54361936 | 0.76199927 | 0.8553656 | HOS3 | XVI | 329606 | 331699 | 1 |
| YPL117C | 41.9580598 | 0.0782065 | 0.68181706 | 0.04078315 | 0.97981493 | 0.98440542 | IDI1 | XVI | 327864 | 328730 | -1 |
| YPL118W | 17.4520826 | -0.094059 | 0.82453168 | 0.33307667 | 0.84659036 | 0.90975468 | MRP51 | XVI | 326628 | 327662 | 1 |
| YPL119C | 285.037131 | -1.2783899 | 0.51573011 | 6.72521735 | 0.03464476 | 0.09984728 | DBP1 | XVI | 324411 | 326264 | -1 |
| YPL119C-A | 40.1708839 | -0.6257802 | 0.69126371 | 9.93144839 | 0.0069729 | 0.02994378 | NA | XVI | 324024 | 324287 | -1 |
| YPL120W | 13.0196837 | -0.3072617 | 0.90741662 | 0.22742625 | 0.89251396 | 0.93387025 | VPS30 | XVI | 322071 | 323744 | 1 |
| YPL121C | 30.1622048 | -1.8377797 | 0.82347357 | 5.41365784 | 0.06674814 | 0.16211061 | MEI5 | XVI | 320962 | 321630 | -1 |
| YPL122C | 25.2109492 | -0.124642 | 0.68129248 | 3.04954188 | 0.21767091 | 0.38155469 | TFB2 | XVI | 319226 | 320767 | -1 |
| YPL123C | 14.0181933 | 0.90454353 | 0.9046107 | 1.01481781 | 0.60205354 | 0.74772415 | RNY1 | XVI | 317641 | 318945 | -1 |
| YPL124W | 119.181686 | 3.2764927 | 0.94103808 | 10.8268375 | 0.00445638 | 0.02079165 | SPC29 | XVI | 316755 | 317516 | 1 |
| YPL125W | 10.9744936 | 0.11545323 | 0.90063952 | 0.3470276 | 0.84070555 | 0.90645906 | KAP120 | XVI | 313388 | 316486 | 1 |
| YPL126W | 26.3275124 | -0.8441742 | 0.74910911 | 1.40857376 | 0.49446106 | 0.65483292 | NAN1 | XVI | 310210 | 312900 | 1 |
| YPL127C | 63.9471792 | -1.1039381 | 0.56085328 | 12.715291 | 0.00173344 | 0.01006684 | HHO1 | XVI | 308828 | 309604 | -1 |
| YPL128C | 121.285395 | 0.98129906 | 0.55387535 | 11.7944723 | 0.00274703 | 0.01420115 | TBF1 | XVI | 306533 | 308221 | -1 |
| YPL129W | 74.3504342 | 0.01226547 | 0.57839789 | 1.28082869 | 0.52707399 | 0.68184633 | TAF14 | XVI | 305298 | 306137 | 1 |
| YPL131W | 298.94375 | -0.4992458 | 0.56367192 | 25.8485221 | 2.44E-06 | 3.83E-05 | RPL5 | XVI | 303121 | 304014 | 1 |
| YPL132W | 40.3992312 | 0.09413994 | 0.62317749 | 3.55820148 | 0.16878986 | 0.3187519 | COX11 | XVI | 301716 | 302618 | 1 |
| YPL133C | 24.2136169 | -0.255688 | 0.76060163 | 5.0200682 | 0.08126547 | 0.18851244 | RDS2 | XVI | 299888 | 301228 | -1 |
| YPL134C | 16.2367729 | 1.22656987 | 0.99640509 | 3.05020309 | 0.21759896 | 0.38155469 | ODC1 | XVI | 298571 | 299503 | -1 |
| YPL135W | 37.9698232 | 0.73252185 | 0.71327868 | 1.15002421 | 0.56269806 | 0.71410256 | ISU1 | XVI | 297553 | 298050 | 1 |
| YPL137C | 86.4841071 | 0.23493339 | 0.61837134 | 11.3604354 | 0.00341282 | 0.01686024 | GIP3 | XVI | 292817 | 296647 | -1 |
| YPL138C | 12.9784267 | 0.93505277 | 0.88087 | 1.65316125 | 0.43754285 | 0.60627631 | SPP1 | XVI | 291366 | 292427 | -1 |
| YPL139C | 34.9498958 | -1.286929 | 0.75004458 | 9.18527645 | 0.01012611 | 0.03983028 | UME1 | XVI | 289669 | 291051 | -1 |
| YPL140C | 129.469642 | 2.14914737 | 0.60249571 | 41.0624305 | 1.21E-09 | 4.38E-08 | MKK2 | XVI | 287514 | 289034 | -1 |
| YPL141C | 291.130431 | -0.3929812 | 0.48654979 | 0.68037438 | 0.7116371 | 0.82438173 | FRK1 | XVI | 283464 | 286061 | -1 |
| YPL143W | 117.819209 | -0.2278562 | 0.67004303 | 16.0649687 | 0.00032474 | 0.00261509 | RPL33A | XVI | 282122 | 282970 | 1 |
| YPL144W | 11.3359409 | -2.8217349 | 1.04566659 | 16.8569941 | 0.00021855 | 0.0018473 | POC4 | XVI | 280480 | 280926 | 1 |
| YPL145C | 55.834204 | -0.1265529 | 0.68199345 | 2.19214421 | 0.33418114 | 0.5079834 | KES1 | XVI | 278395 | 279699 | -1 |
| YPL146C | 54.8597733 | -1.3909479 | 0.72552329 | 13.2606858 | 0.00131971 | 0.00824961 | NOP53 | XVI | 276162 | 277529 | -1 |
| YPL147W | 33.9082953 | -0.1666803 | 0.6907269 | 1.70824284 | 0.425657 | 0.5957336 | PXA1 | XVI | 273255 | 275867 | 1 |

| | | | | | | | | | | | |
|---|---|---|---|---|---|---|---|---|---|---|---|
| YPL148C | 83.0580671 | 0.2808708 | 0.55285204 | 3.93887977 | 0.13953499 | 0.27801887 | PPT2 | XVI | 272294 | 272815 | -1 |
| YPL149W | 99.0589687 | -0.5732271 | 0.54057254 | 1.54592022 | 0.46164453 | 0.62532091 | ATG5 | XVI | 271310 | 272194 | 1 |
| YPL150W | 44.9007843 | 1.04090087 | 0.62785785 | 4.71938334 | 0.09444934 | 0.20984715 | NA | XVI | 268188 | 270893 | 1 |
| YPL151C | 160.083167 | 0.60388964 | 0.62383535 | 1.0913625 | 0.5794469 | 0.72802613 | PRP46 | XVI | 266180 | 267535 | -1 |
| YPL152W | 76.3505619 | 0.02142698 | 0.67903581 | 3.50053647 | 0.17372734 | 0.32507061 | RRD2 | XVI | 265028 | 266104 | 1 |
| YPL153C | 7.45896983 | -0.4878249 | 1.16907498 | 0.18897236 | 0.90984031 | 0.94492832 | RAD53 | XVI | 261727 | 264192 | -1 |
| YPL154C | 75.3691311 | -0.2899785 | 0.80724368 | 0.3672555 | 0.83224555 | 0.90069191 | PEP4 | XVI | 259714 | 260931 | -1 |
| YPL155C | 28.9025403 | 0.17635026 | 0.71340895 | 6.72112818 | 0.03471567 | 0.10000186 | KIP2 | XVI | 257216 | 259336 | -1 |
| YPL156C | 749.616911 | -0.7376254 | 0.50546234 | 2.44024757 | 0.29519362 | 0.46951321 | PRM4 | XVI | 255913 | 256767 | -1 |
| YPL157W | 116.662219 | 1.46213707 | 0.62272053 | 6.25140467 | 0.04390609 | 0.11884817 | TGS1 | XVI | 254814 | 255761 | 1 |
| YPL158C | 268.813706 | -0.2386423 | 0.49276526 | 1.71854031 | 0.42347104 | 0.59425528 | AIM44 | XVI | 252034 | 254310 | -1 |
| YPL159C | 119.904526 | 0.96529499 | 0.52942106 | 12.9468447 | 0.00154393 | 0.00934751 | PET20 | XVI | 250907 | 251668 | -1 |
| YPL160W | 52.3162552 | -0.8959518 | 0.66612962 | 9.30705206 | 0.00952795 | 0.03806203 | CDC60 | XVI | 246990 | 250262 | 1 |
| YPL161C | 71.2311932 | 0.31560022 | 0.59709447 | 6.75529053 | 0.03412772 | 0.09875038 | BEM4 | XVI | 244319 | 246220 | -1 |
| YPL162C | 1130.6657 | 1.01135323 | 0.49661976 | 6.17331954 | 0.0456542 | 0.12191465 | NA | XVI | 243206 | 244027 | -1 |
| YPL163C | 30.3554485 | 2.34723403 | 0.89236818 | 7.0600385 | 0.02930435 | 0.08719023 | SVS1 | XVI | 241918 | 242700 | -1 |
| YPL164C | 33.2342481 | -0.3482489 | 0.78605777 | 4.25102025 | 0.11937206 | 0.24853082 | MLH3 | XVI | 239350 | 241497 | -1 |
| YPL165C | 2534.39633 | 1.54632087 | 0.45759264 | 16.703864 | 0.00023594 | 0.00196844 | SET6 | XVI | 237956 | 239077 | -1 |
| YPL166W | 13.1536764 | -2.114515 | 0.99279862 | 5.02790264 | 0.08094776 | 0.18800141 | ATG29 | XVI | 237338 | 237979 | 1 |
| YPL167C | 41.7186997 | -0.3282523 | 0.62047573 | 0.57555708 | 0.74992765 | 0.84806272 | REV3 | XVI | 232593 | 237107 | -1 |
| YPL168W | 137.827041 | 1.11305341 | 0.57671208 | 4.31528333 | 0.11559742 | 0.242911 | MRX4 | XVI | 231217 | 232509 | 1 |
| YPL169C | 788.258884 | 2.54683507 | 0.53507413 | 134.021168 | 7.90E-30 | 2.29E-26 | MEX67 | XVI | 229039 | 230838 | -1 |
| YPL170W | 63.5742305 | 1.44726196 | 0.64774787 | 28.0599181 | 8.07E-07 | 1.47E-05 | DAP1 | XVI | 228314 | 228772 | 1 |
| YPL171C | 19.9927473 | 0.44422784 | 0.91882961 | 17.3191172 | 0.00017346 | 0.00150125 | OYE3 | XVI | 226169 | 227371 | -1 |
| YPL172C | 43.3699555 | 0.94664076 | 0.68109583 | 14.1836212 | 0.00083189 | 0.00566552 | COX10 | XVI | 224353 | 225741 | -1 |
| YPL173W | 49.524344 | 0.22965325 | 0.65492293 | 22.2894836 | 1.45E-05 | 0.00018111 | MRPL40 | XVI | 223143 | 224036 | 1 |
| YPL174C | 228.2472 | 0.94526067 | 0.55551689 | 3.09510017 | 0.2127686 | 0.37639175 | NIP100 | XVI | 220167 | 222773 | -1 |
| YPL175W | 138.123499 | -1.0291963 | 0.49613659 | 25.0485718 | 3.64E-06 | 5.44E-05 | SPT14 | XVI | 218630 | 220088 | 1 |
| YPL176C | 38.9679815 | 1.04588874 | 0.67199898 | 2.59979834 | 0.27255927 | 0.44584616 | TRE1 | XVI | 216012 | 218363 | -1 |
| YPL177C | 18.3234074 | 0.10170214 | 0.76204611 | 0.14158013 | 0.93165746 | 0.95711439 | CUP9 | XVI | 213042 | 213962 | -1 |
| YPL178W | 13.4110517 | 0.2845978 | 0.85386615 | 0.82035698 | 0.66353181 | 0.79066663 | CBC2 | XVI | 212158 | 212784 | 1 |
| YPL179W | 58.4547875 | 0.47392425 | 0.74027402 | 0.54452832 | 0.76165304 | 0.85526961 | PPQ1 | XVI | 208157 | 209806 | 1 |
| YPL180W | 37.5913026 | -0.3957189 | 0.64968972 | 1.44268397 | 0.48609948 | 0.64790884 | TCO89 | XVI | 205248 | 207647 | 1 |

| | | | | | | | | | | | |
|---|---|---|---|---|---|---|---|---|---|---|---|
| YPL181W | 90.0941349 | 0.66404909 | 0.55711272 | 1.56781234 | 0.45661889 | 0.62207609 | CTI6 | XVI | 203421 | 204941 | 1 |
| YPL183C | 252.884567 | 1.57473486 | 0.75510602 | 5.00767266 | 0.0817707 | 0.18922955 | RTT10 | XVI | 199495 | 202536 | -1 |
| YPL183W-A | 17.8925869 | -1.5802198 | 0.78997666 | 4.45569522 | 0.10776012 | 0.23089058 | RTC6 | XVI | 199095 | 199376 | 1 |
| YPL184C | 28.0105598 | 0.54786611 | 0.71512902 | 0.65402682 | 0.72107407 | 0.83085787 | MRN1 | XVI | 195950 | 197788 | -1 |
| YPL186C | 83.2509087 | -0.4602668 | 0.59284474 | 1.98715935 | 0.37024895 | 0.54354498 | UIP4 | XVI | 194512 | 195426 | -1 |
| YPL187W | 125.429403 | -1.7360646 | 0.61997854 | 44.0175255 | 2.77E-10 | 1.16E-08 | MF(ALPHA)1 | XVI | 193648 | 194145 | 1 |
| YPL188W | 571.553037 | -3.308356 | 0.48602233 | 116.064346 | 6.27E-26 | 5.18E-23 | POS5 | XVI | 191406 | 192650 | 1 |
| YPL189C-A | 20.4366754 | -1.3360872 | 0.75899485 | 3.08880672 | 0.21343918 | 0.37734744 | COA2 | XVI | 188307 | 188513 | -1 |
| YPL189W | 33.4723445 | -0.1958567 | 0.72990619 | 1.58561899 | 0.45257151 | 0.61859987 | GUP2 | XVI | 189154 | 190983 | 1 |
| YPL190C | 385.523114 | 0.97253118 | 0.63076914 | 3.29998135 | 0.1920517 | 0.34838007 | NAB3 | XVI | 185317 | 187725 | -1 |
| YPL191C | 14.5640653 | -0.2562668 | 0.8643261 | 0.54138636 | 0.76285052 | 0.85565759 | NA | XVI | 183597 | 184679 | -1 |
| YPL192C | 469.227848 | -1.2694518 | 0.46466697 | 25.2204051 | 3.34E-06 | 5.07E-05 | PRM3 | XVI | 182655 | 183056 | -1 |
| YPL193W | 15.4447164 | -0.3538322 | 0.83451818 | 0.40983528 | 0.81471441 | 0.88869564 | RSA1 | XVI | 181403 | 182548 | 1 |
| YPL194W | 9.94502394 | 0.88174104 | 0.97675799 | 0.93067435 | 0.62792335 | 0.76540551 | DDC1 | XVI | 179277 | 181115 | 1 |
| YPL195W | 33.4552174 | 0.03327415 | 0.6988778 | 0.42254769 | 0.80955235 | 0.8858947 | APL5 | XVI | 176223 | 179021 | 1 |
| YPL196W | 103.082693 | -0.109064 | 0.51950035 | 0.80492719 | 0.66867068 | 0.79450097 | OXR1 | XVI | 175043 | 175864 | 1 |
| YPL198W | 28.2920328 | -0.7566652 | 0.66578883 | 1.34514401 | 0.51039415 | 0.66786132 | RPL7B | XVI | 173152 | 174702 | 1 |
| YPL199C | 21.4068206 | -0.0455415 | 0.77266571 | 0.03101604 | 0.98461161 | 0.98751104 | NA | XVI | 172033 | 172755 | -1 |
| YPL200W | 44.6024688 | -0.5266971 | 0.73998339 | 12.401916 | 0.00202749 | 0.01126565 | CSM4 | XVI | 171484 | 171954 | 1 |
| YPL201C | 40.4085867 | 0.80493627 | 0.74704114 | 4.60713053 | 0.09990203 | 0.21918634 | YIG1 | XVI | 169770 | 171155 | -1 |
| YPL202C | 124.130738 | 0.81926392 | 0.63255414 | 2.13946059 | 0.34310104 | 0.5166593 | AFT2 | XVI | 168088 | 169338 | -1 |
| YPL203W | 14.5916833 | 0.95309578 | 1.04227185 | 1.57564185 | 0.45483484 | 0.62066785 | TPK2 | XVI | 166256 | 167398 | 1 |
| YPL204W | 47.8322832 | 1.11472478 | 0.61497023 | 3.59862128 | 0.16541288 | 0.31442566 | HRR25 | XVI | 164276 | 165760 | 1 |
| YPL206C | 69.1752654 | -0.5597991 | 0.5441637 | 1.13090904 | 0.56810187 | 0.71834677 | PGC1 | XVI | 162632 | 163597 | -1 |
| YPL207W | 74.6385325 | 0.13256478 | 0.61503646 | 2.61793289 | 0.27009907 | 0.44340052 | TYW1 | XVI | 159909 | 162341 | 1 |
| YPL208W | 10.3747291 | -1.4623722 | 0.96678136 | 3.07150475 | 0.21529365 | 0.37922681 | RKM1 | XVI | 157842 | 159593 | 1 |
| YPL209C | 6.39482843 | -0.5261362 | 1.24128286 | 0.38418133 | 0.82523204 | 0.89541781 | IPL1 | XVI | 156490 | 157593 | -1 |
| YPL210C | 33.9351836 | -0.452174 | 0.65925672 | 8.97387029 | 0.01125509 | 0.04301449 | SRP72 | XVI | 154290 | 156212 | -1 |
| YPL211W | 17.3917799 | -0.5930922 | 0.76782467 | 14.2862783 | 0.00079027 | 0.00542138 | NIP7 | XVI | 153495 | 154040 | 1 |
| YPL212C | 29.4611048 | -1.5679855 | 0.75511767 | 9.4584718 | 0.00883322 | 0.03582669 | PUS1 | XVI | 151515 | 153149 | -1 |
| YPL213W | 11.2003942 | -0.4220489 | 0.92580211 | 0.46695849 | 0.79177403 | 0.87354642 | LEA1 | XVI | 150614 | 151330 | 1 |
| YPL214C | 250.733721 | 1.26120216 | 0.77007962 | 4.47323406 | 0.10681926 | 0.22957814 | THI6 | XVI | 148569 | 150191 | -1 |
| YPL215W | 23.7834351 | 0.03458364 | 0.69802295 | 0.85571714 | 0.65190361 | 0.78244649 | CBP3 | XVI | 147416 | 148423 | 1 |

| | | | | | | | | | | | |
|---|---|---|---|---|---|---|---|---|---|---|---|
| YPL216W | 1410.0772 | -0.1574851 | 0.49934151 | 1.46072786 | 0.48173364 | 0.64424159 | NA | XVI | 143821 | 147129 | 1 |
| YPL217C | 191.233867 | -0.2656167 | 0.52878081 | 2.72674413 | 0.25579676 | 0.42803963 | BMS1 | XVI | 139620 | 143171 | -1 |
| YPL218W | 41.5656051 | 0.19698495 | 0.61761563 | 1.25800543 | 0.53312321 | 0.68763274 | SAR1 | XVI | 138698 | 139409 | 1 |
| YPL219W | 21.6007373 | -0.2970803 | 0.77508707 | 2.88995673 | 0.23575118 | 0.40372651 | PCL8 | XVI | 136750 | 138228 | 1 |
| YPL220W | 15.1209626 | -1.5532222 | 0.87982321 | 9.74501126 | 0.00765416 | 0.03206773 | RPL1A | XVI | 135790 | 136443 | 1 |
| YPL221W | 80.6127681 | 0.71778714 | 0.63496168 | 1.41028054 | 0.49403927 | 0.65469026 | FLC1 | XVI | 133043 | 135424 | 1 |
| YPL222W | 123.203915 | 1.34146012 | 0.52878202 | 19.662291 | 5.38E-05 | 0.00054696 | FMP40 | XVI | 130162 | 132228 | 1 |
| YPL223C | 107.83558 | 0.38047242 | 0.58627103 | 4.81142069 | 0.0902014 | 0.20366605 | GRE1 | XVI | 128632 | 129138 | -1 |
| YPL224C | 26.664884 | -0.9453098 | 0.70468693 | 1.90907768 | 0.38498964 | 0.55783034 | MMT2 | XVI | 126634 | 128088 | -1 |
| YPL225W | 84.5608594 | 1.30288696 | 0.6623047 | 5.40457715 | 0.06705188 | 0.16264366 | NA | XVI | 126006 | 126446 | 1 |
| YPL226W | 59.9064108 | -0.8421334 | 0.62438765 | 5.73912192 | 0.05672383 | 0.14417513 | NEW1 | XVI | 121767 | 125357 | 1 |
| YPL227C | 50.5108446 | 1.2694788 | 0.89373797 | 4.82284733 | 0.08968752 | 0.20308593 | ALG5 | XVI | 120163 | 121167 | -1 |
| YPL228W | 32.459821 | 0.73753404 | 0.69070652 | 2.70689917 | 0.25834753 | 0.43008401 | CET1 | XVI | 118382 | 120031 | 1 |
| YPL229W | 10.3792567 | -0.1585487 | 1.01010298 | 0.32881643 | 0.84839562 | 0.90980423 | NA | XVI | 117067 | 117687 | 1 |
| YPL230W | 162.210869 | -0.4887042 | 0.54948671 | 0.9805714 | 0.61245139 | 0.7554524 | USV1 | XVI | 115312 | 116487 | 1 |
| YPL231W | 177.939825 | 0.43512941 | 0.70171784 | 1.10029763 | 0.57686396 | 0.72593834 | FAS2 | XVI | 108652 | 114315 | 1 |
| YPL232W | 125.331993 | 0.85402408 | 0.52753006 | 2.98292368 | 0.22504344 | 0.3904964 | SSO1 | XVI | 107275 | 108147 | 1 |
| YPL233W | 9.23564103 | 1.43900477 | 1.08520129 | 2.93286724 | 0.23074695 | 0.39774482 | NSL1 | XVI | 106172 | 106822 | 1 |
| YPL234C | 169.026772 | -0.2819446 | 0.5770933 | 0.36794399 | 0.8319591 | 0.90069191 | VMA11 | XVI | 104946 | 105440 | -1 |
| YPL235W | 36.4765922 | 0.53122423 | 0.73784434 | 0.92241005 | 0.63052339 | 0.76792815 | RVB2 | XVI | 103232 | 104647 | 1 |
| YPL236C | 9.22046165 | 0.03638803 | 1.13938514 | 3.49946836 | 0.17382014 | 0.32507061 | ENV7 | XVI | 101608 | 102702 | -1 |
| YPL237W | 155.52167 | 0.22154836 | 0.4848821 | 3.48697122 | 0.17490967 | 0.32678559 | SUI3 | XVI | 100496 | 101353 | 1 |
| YPL239W | 47.430462 | -0.6966976 | 0.59250342 | 4.05184107 | 0.13187239 | 0.26715926 | YAR1 | XVI | 99484 | 100086 | 1 |
| YPL240C | 628.241469 | 0.86534002 | 0.49156098 | 31.2019876 | 1.68E-07 | 3.81E-06 | HSP82 | XVI | 96496 | 98625 | -1 |
| YPL241C | 480.450821 | -0.3123285 | 0.46993385 | 1.12115775 | 0.5708785 | 0.72122769 | CIN2 | XVI | 95372 | 96258 | -1 |
| YPL242C | 925.218825 | 3.97144452 | 0.48097318 | 69.2951204 | 8.97E-16 | 1.06E-13 | IQG1 | XVI | 90622 | 95109 | -1 |
| YPL243W | 351.574726 | 2.57519341 | 0.53683959 | 42.9669903 | 4.68E-10 | 1.82E-08 | SRP68 | XVI | 88517 | 90316 | 1 |
| YPL244C | 42.0135049 | -0.2623111 | 0.79593236 | 2.01797695 | 0.36458758 | 0.53782474 | HUT1 | XVI | 87014 | 88033 | -1 |
| YPL245W | 530.872484 | 0.98280079 | 0.61953172 | 6.79699875 | 0.03342339 | 0.0971981 | NA | XVI | 85586 | 86950 | 1 |
| YPL246C | 88.5673853 | -0.5103435 | 0.58445591 | 1.65825189 | 0.43643058 | 0.60539844 | RBD2 | XVI | 84509 | 85297 | -1 |
| YPL247C | 50.3800452 | 0.62395815 | 0.61646688 | 4.06128528 | 0.13125115 | 0.26636668 | NA | XVI | 82625 | 84196 | -1 |
| YPL248C | 1384.06982 | -0.722183 | 0.47925034 | 4.48985083 | 0.10593544 | 0.22838155 | GAL4 | XVI | 79711 | 82356 | -1 |
| YPL249C | 562.087254 | -1.5273138 | 0.51492428 | 23.0815831 | 9.73E-06 | 0.00012915 | GYP5 | XVI | 76669 | 79353 | -1 |

| | | | | | | | | | | | |
|---|---|---|---|---|---|---|---|---|---|---|---|
| YPL249C-A | 254.175645 | -0.45745 | 0.63010048 | 19.5120085 | 5.79E-05 | 0.00058655 | RPL36B | XVI | 75699 | 76239 | -1 |
| YPL250C | 10.6573536 | -0.0858823 | 0.99490233 | 0.12717452 | 0.93839223 | 0.95995205 | ATG41 | XVI | 74309 | 74719 | -1 |
| YPL252C | 27.4685969 | -0.4793387 | 0.73451823 | 8.28466083 | 0.01588579 | 0.05601627 | YAH1 | XVI | 73363 | 73881 | -1 |
| YPL253C | 580.071865 | -0.9761798 | 0.48812536 | 16.9365121 | 0.00021003 | 0.0017805 | VIK1 | XVI | 71063 | 73006 | -1 |
| YPL254W | 25.888218 | -0.3481674 | 0.7643588 | 0.39616487 | 0.82030223 | 0.8920536 | HFI1 | XVI | 69485 | 70951 | 1 |
| YPL255W | 14.2222383 | -1.2334769 | 0.84480674 | 2.10566082 | 0.34894868 | 0.52261068 | BBP1 | XVI | 67725 | 68882 | 1 |
| YPL256C | 160.696653 | -0.4809211 | 0.51022598 | 5.85472679 | 0.05353801 | 0.13770994 | CLN2 | XVI | 64977 | 66614 | -1 |
| YPL257W | 50.1080659 | -0.1515128 | 0.60979074 | 1.75778382 | 0.41524278 | 0.58726324 | NA | XVI | 63279 | 63860 | 1 |
| YPL258C | 475.460018 | 2.59442287 | 0.54355332 | 75.0910987 | 4.95E-17 | 7.53E-15 | THI21 | XVI | 53498 | 55153 | -1 |
| YPL259C | 23.3456495 | 0.9608742 | 0.72514904 | 2.3682758 | 0.30600988 | 0.47867401 | APM1 | XVI | 51244 | 52671 | -1 |
| YPL260W | 58.9210019 | 0.67338997 | 0.59798352 | 8.76993894 | 0.01246327 | 0.04652632 | CUB1 | XVI | 49303 | 50958 | 1 |
| YPL261C | 1331.73461 | 0.41051055 | 0.5538046 | 0.57814092 | 0.74895943 | 0.8479615 | NA | XVI | 48996 | 49304 | -1 |
| YPL262W | 23.214562 | 0.43107005 | 0.7957548 | 1.60226924 | 0.44881944 | 0.61506853 | FUM1 | XVI | 47336 | 48802 | 1 |
| YPL263C | 58.3671485 | -0.8236691 | 0.63396619 | 2.43059017 | 0.29662247 | 0.47066158 | KEL3 | XVI | 44551 | 46506 | -1 |
| YPL264C | 101.917013 | -0.2791038 | 0.57744178 | 1.93719688 | 0.37961472 | 0.55350522 | NA | XVI | 43283 | 44344 | -1 |
| YPL265W | 733.145839 | -0.1764079 | 0.5869188 | 1.90533383 | 0.38571099 | 0.55845627 | DIP5 | XVI | 41043 | 42869 | 1 |
| YPL266W | 38.6004515 | 1.06699038 | 0.71147978 | 6.76745468 | 0.03392078 | 0.09824187 | DIM1 | XVI | 39121 | 40077 | 1 |
| YPL267W | 5.88051903 | -1.67085 | 1.27089122 | 3.11964615 | 0.21017325 | 0.3730543 | ACM1 | XVI | 38169 | 38798 | 1 |
| YPL268W | 62.4517549 | -2.1536138 | 0.60191262 | 18.4407873 | 9.90E-05 | 0.00092602 | PLC1 | XVI | 35236 | 37845 | 1 |
| YPL269W | 25.2443604 | 0.66111405 | 0.72795638 | 2.59115026 | 0.27374038 | 0.44722258 | KAR9 | XVI | 33013 | 34947 | 1 |
| YPL270W | 26.0190313 | 0.17270302 | 0.69541745 | 1.05666483 | 0.58958734 | 0.73728204 | MDL2 | XVI | 30482 | 32803 | 1 |
| YPL271W | 16.7640561 | -0.3264892 | 0.79166434 | 0.16953419 | 0.91872623 | 0.94955817 | ATP15 | XVI | 30079 | 30267 | 1 |
| YPL272C | 8.06734631 | -1.3869183 | 1.04284245 | 1.81405111 | 0.40372329 | 0.57458523 | PBI1 | XVI | 26611 | 28164 | -1 |
| YPL273W | 11.4630495 | -0.4273131 | 0.85783319 | 4.93476028 | 0.08480675 | 0.19477631 | SAM4 | XVI | 25087 | 26064 | 1 |
| YPL274W | 131.473962 | -0.241903 | 0.53972731 | 3.59201425 | 0.16596023 | 0.31525228 | SAM3 | XVI | 22938 | 24701 | 1 |
| YPL275W | 158.924384 | -1.6730759 | 0.65045371 | 7.49910269 | 0.0235283 | 0.07472784 | NA | XVI | 18369 | 19079 | 1 |
| YPL276W | 158.569452 | -0.8282563 | 0.61669571 | 9.14736417 | 0.01031989 | 0.04032978 | NA | XVI | 17948 | 18385 | 1 |
| YPL277C | 25.7690514 | -0.0404137 | 0.86564035 | 13.6420655 | 0.00109059 | 0.00709251 | NA | XVI | 15405 | 16868 | -1 |
| YPL279C | 13.1864606 | 0.4491008 | 1.10870473 | 8.75707396 | 0.0125437 | 0.04676626 | FEX2 | XVI | 13228 | 14355 | -1 |
| YPL280W | 7.94479807 | -2.6184804 | 1.42989427 | 10.9948512 | 0.00409731 | 0.0195435 | HSP32 | XVI | 11887 | 12600 | 1 |
| YPL283C | 5.03946198 | -0.2671687 | 1.36842848 | 0.23248221 | 0.89026055 | 0.93317728 | YRF1-7 | XVI | 280 | 6007 | -1 |
| YPR001W | 283.284459 | -1.1619583 | 0.51151749 | 17.6048633 | 0.00015037 | 0.00133737 | CIT3 | XVI | 556377 | 557837 | 1 |
| YPR002W | 76.753294 | 0.13745843 | 0.58319764 | 4.27156744 | 0.11815196 | 0.24673616 | PDH1 | XVI | 558385 | 559935 | 1 |

| | | | | | | | | | | | |
|---|---|---|---|---|---|---|---|---|---|---|---|
| YPR003C | 27.2607812 | -0.749658 | 0.73610243 | 1.40216476 | 0.4960481 | 0.65603003 | NA | XVI | 561504 | 563768 | -1 |
| YPR004C | 302.593763 | 1.75181396 | 0.52881414 | 74.4700943 | 6.75E-17 | 9.79E-15 | AIM45 | XVI | 564007 | 565041 | -1 |
| YPR005C | 106.84206 | -0.3724553 | 0.555751 | 14.6889129 | 0.00064616 | 0.00457931 | HAL1 | XVI | 565787 | 566671 | -1 |
| YPR006C | 137.461258 | -0.5094904 | 0.51378993 | 0.98443897 | 0.61126819 | 0.75489714 | ICL2 | XVI | 567269 | 568996 | -1 |
| YPR007C | 24.881462 | -1.3365108 | 0.75707162 | 3.57238572 | 0.16759702 | 0.31722352 | REC8 | XVI | 569336 | 571378 | -1 |
| YPR008W | 149.498349 | 2.13648048 | 0.54849136 | 14.8607473 | 0.00059297 | 0.00424214 | HAA1 | XVI | 573018 | 575102 | 1 |
| YPR009W | 226.443113 | -1.4383206 | 0.50395413 | 35.1821864 | 2.29E-08 | 6.35E-07 | SUT2 | XVI | 576552 | 577358 | 1 |
| YPR010C | 44.5632116 | -1.3459983 | 0.7311906 | 15.1568779 | 0.00051136 | 0.00373834 | RPA135 | XVI | 577585 | 581196 | -1 |
| YPR010C-A | 12.556251 | -0.6675658 | 0.9008809 | 0.63499674 | 0.72796787 | 0.83592431 | NA | XVI | 582373 | 582734 | -1 |
| YPR011C | 226.558345 | 0.09712535 | 0.50278192 | 8.20805527 | 0.01650606 | 0.05733167 | NA | XVI | 583062 | 584042 | -1 |
| YPR013C | 35.7681176 | 0.2967457 | 0.68419983 | 3.56513586 | 0.16820565 | 0.31795975 | CMR3 | XVI | 584632 | 585585 | -1 |
| YPR014C | 136.241567 | 0.00951535 | 0.60882879 | 0.2129217 | 0.89901024 | 0.93754851 | NA | XVI | 587189 | 587518 | -1 |
| YPR015C | 57.2397923 | 0.37314217 | 0.75811281 | 24.679608 | 4.37E-06 | 6.35E-05 | NA | XVI | 590283 | 591026 | -1 |
| YPR016C | 21.0717727 | -0.7220837 | 0.88337369 | 5.05524392 | 0.07984868 | 0.18619567 | TIF6 | XVI | 592332 | 593069 | -1 |
| YPR017C | 42.0035366 | -0.8952481 | 0.72572271 | 3.0140964 | 0.22156302 | 0.38636302 | DSS4 | XVI | 593486 | 593917 | -1 |
| YPR018W | 78.6938773 | 0.48307887 | 0.63065416 | 2.07094979 | 0.35505772 | 0.52857757 | RLF2 | XVI | 594476 | 596296 | 1 |
| YPR019W | 19.6066079 | -0.4309397 | 0.86539235 | 0.29119275 | 0.86450657 | 0.9204658 | MCM4 | XVI | 596750 | 599551 | 1 |
| YPR020W | 8.86099299 | -0.2135934 | 1.27278496 | 0.32934998 | 0.84816932 | 0.90980423 | ATP20 | XVI | 599870 | 600217 | 1 |
| YPR021C | 54.3561844 | -0.5532677 | 0.66083385 | 22.8942104 | 1.07E-05 | 0.00013991 | AGC1 | XVI | 600649 | 603357 | -1 |
| YPR022C | 41.0964281 | -0.0479283 | 0.63623885 | 0.55653428 | 0.75709454 | 0.85316805 | SDD4 | XVI | 603911 | 607312 | -1 |
| YPR023C | 69.2204601 | 0.36492744 | 0.54483175 | 0.83405742 | 0.659002 | 0.7867261 | EAF3 | XVI | 608826 | 610031 | -1 |
| YPR024W | 84.2865756 | 0.5382629 | 0.59503966 | 19.5506316 | 5.68E-05 | 0.00057634 | YME1 | XVI | 610481 | 612724 | 1 |
| YPR025C | 52.1863232 | 0.94835265 | 0.71201514 | 1.99925069 | 0.36801729 | 0.5412294 | CCL1 | XVI | 613377 | 614558 | -1 |
| YPR026W | 83.3897458 | -1.0105128 | 0.58701768 | 3.14716305 | 0.2073014 | 0.36942908 | ATH1 | XVI | 615379 | 619014 | 1 |
| YPR027C | 839.67191 | -1.3640429 | 0.47066728 | 27.7898133 | 9.24E-07 | 1.66E-05 | NA | XVI | 620425 | 621258 | -1 |
| YPR028W | 23.3804826 | 0.38726838 | 0.93239224 | 0.86624248 | 0.64848187 | 0.78028055 | YOP1 | XVI | 623527 | 624202 | 1 |
| YPR029C | 44.4809343 | -0.3691785 | 0.71986342 | 1.12446081 | 0.56993646 | 0.72050919 | APL4 | XVI | 624469 | 626967 | -1 |
| YPR030W | 246.855467 | -1.4649451 | 0.51266248 | 34.2375443 | 3.68E-08 | 9.83E-07 | CSR2 | XVI | 627880 | 631245 | 1 |
| YPR031W | 18.606281 | -0.7468046 | 0.81161264 | 3.01943499 | 0.2209724 | 0.38583539 | NTO1 | XVI | 631515 | 633761 | 1 |
| YPR032W | 19.651125 | -0.0007358 | 0.76378185 | 0.35957453 | 0.83544792 | 0.90300565 | SRO7 | XVI | 634123 | 637224 | 1 |
| YPR033C | 31.5755907 | -0.9008304 | 0.72803149 | 4.08342841 | 0.12980601 | 0.26426638 | HTS1 | XVI | 637379 | 639019 | -1 |
| YPR034W | 38.4524955 | -0.174992 | 0.65306542 | 1.81567172 | 0.40339629 | 0.57429666 | ARP7 | XVI | 639525 | 640958 | 1 |
| YPR035W | 99.0273959 | 1.30281494 | 0.70347837 | 3.42419692 | 0.18048665 | 0.33506648 | GLN1 | XVI | 642208 | 643320 | 1 |

| | | | | | | | | | | | |
|---|---|---|---|---|---|---|---|---|---|---|---|
| YPR036W | 351.851788 | -0.0778537 | 0.56323148 | 7.36216253 | 0.02519572 | 0.07822156 | VMA13 | XVI | 643836 | 645272 | 1 |
| YPR036W-A | 100.69823 | 1.55124801 | 0.58839311 | 11.9087477 | 0.00259447 | 0.01364393 | SPO24 | XVI | 645950 | 646153 | 1 |
| YPR037C | 36.6514529 | 1.74622747 | 0.74893429 | 5.66510258 | 0.05886249 | 0.14753844 | ERV2 | XVI | 646448 | 647038 | -1 |
| YPR038W | 1219.38698 | -0.1921318 | 0.49065568 | 0.27284522 | 0.87247384 | 0.92527295 | NA | XVI | 646836 | 647195 | 1 |
| YPR040W | 99.1660519 | -0.7133325 | 0.5436846 | 2.09515949 | 0.35078571 | 0.52427705 | TIP41 | XVI | 647305 | 648375 | 1 |
| YPR041W | 36.8095756 | -0.6139285 | 0.84872002 | 6.8435885 | 0.03265379 | 0.09534315 | TIF5 | XVI | 648704 | 649921 | 1 |
| YPR042C | 18.9980674 | 1.08135599 | 0.96616468 | 1.29353474 | 0.52373609 | 0.67900939 | PUF2 | XVI | 650435 | 653662 | -1 |
| YPR043W | 70.3730866 | -0.8439295 | 0.71625809 | 13.9576547 | 0.00093139 | 0.00620573 | RPL43B | XVI | 654166 | 654847 | 1 |
| YPR045C | 48.7943914 | -0.2955576 | 0.58850505 | 2.62272405 | 0.2694528 | 0.44259056 | THP3 | XVI | 655140 | 656552 | -1 |
| YPR046W | 62.1102106 | -0.2370653 | 0.59718701 | 3.20132604 | 0.2017627 | 0.36122636 | MCM16 | XVI | 656799 | 657344 | 1 |
| YPR047W | 55.9062388 | -1.9108248 | 0.6151637 | 19.4157262 | 6.08E-05 | 0.00061014 | MSF1 | XVI | 657529 | 658938 | 1 |
| YPR048W | 46.1531846 | 1.07066566 | 0.87371418 | 1.7992975 | 0.40671249 | 0.57811175 | TAH18 | XVI | 659182 | 661053 | 1 |
| YPR049C | 24.7740599 | -0.5442728 | 0.70546015 | 0.71422503 | 0.69969377 | 0.81801018 | ATG11 | XVI | 661137 | 664673 | -1 |
| YPR051W | 48.2897862 | 1.41965989 | 0.74862442 | 8.74855413 | 0.01259725 | 0.04684525 | MAK3 | XVI | 664960 | 665490 | 1 |
| YPR053C | 24.0057033 | -2.0008308 | 0.75067362 | 15.84528 | 0.00036244 | 0.0028442 | NA | XVI | 665788 | 666243 | -1 |
| YPR054W | 1001.74801 | -1.167786 | 0.54883224 | 7.09211841 | 0.02883806 | 0.08637991 | SMK1 | XVI | 666280 | 667446 | 1 |
| YPR055W | 42.8673356 | 0.04631989 | 0.60842095 | 0.00753545 | 0.99623937 | 0.99658361 | SEC8 | XVI | 667676 | 670873 | 1 |
| YPR056W | 119.951034 | -0.492272 | 0.5723573 | 0.89354086 | 0.63969075 | 0.77534214 | TFB4 | XVI | 671126 | 672142 | 1 |
| YPR057W | 46.2769565 | -1.9192146 | 0.62146425 | 9.38821 | 0.00914905 | 0.03683798 | BRR1 | XVI | 672471 | 673496 | 1 |
| YPR058W | 85.2992813 | -1.1017713 | 0.53986975 | 27.422304 | 1.11E-06 | 1.94E-05 | YMC1 | XVI | 673751 | 674674 | 1 |
| YPR060C | 45.2808916 | -0.740462 | 0.58616544 | 10.5120197 | 0.00521608 | 0.02376215 | ARO7 | XVI | 674861 | 675631 | -1 |
| YPR061C | 36.4760078 | -2.2832808 | 0.67649256 | 11.5040644 | 0.00317632 | 0.01596431 | JID1 | XVI | 675977 | 676882 | -1 |
| YPR062W | 482.269379 | 0.89189071 | 0.78429463 | 10.2889377 | 0.00583157 | 0.02592206 | FCY1 | XVI | 677165 | 677641 | 1 |
| YPR063C | 22.2085036 | -0.1876918 | 0.72409644 | 0.66016039 | 0.71886608 | 0.82938751 | NA | XVI | 677812 | 678320 | -1 |
| YPR064W | 65.3512448 | 0.86207929 | 0.64827102 | 7.19936922 | 0.02733234 | 0.08294248 | NA | XVI | 678951 | 679370 | 1 |
| YPR065W | 114.260189 | 1.13091288 | 0.59710927 | 5.06598131 | 0.07942114 | 0.18542275 | ROX1 | XVI | 679693 | 680799 | 1 |
| YPR066W | 177.590596 | 0.08024181 | 0.49972179 | 1.16626555 | 0.55814708 | 0.70932212 | UBA3 | XVI | 681213 | 682112 | 1 |
| YPR067W | 18.7564939 | 0.84300519 | 0.79846793 | 3.76356093 | 0.15231867 | 0.29594801 | ISA2 | XVI | 682220 | 682777 | 1 |
| YPR068C | 132.828873 | 0.25730411 | 0.60906991 | 1.41148979 | 0.49374065 | 0.65469026 | HOS1 | XVI | 682941 | 684353 | -1 |
| YPR069C | 21.5254471 | -0.6591982 | 0.91240332 | 4.75842763 | 0.09262337 | 0.20714148 | SPE3 | XVI | 684556 | 685437 | -1 |
| YPR070W | 6.93047238 | -1.5740229 | 1.27346413 | 1.99854322 | 0.3681475 | 0.5412834 | MED1 | XVI | 685898 | 687598 | 1 |
| YPR071W | 6.27428739 | -0.8583919 | 1.27358948 | 0.91314566 | 0.63345087 | 0.77035929 | NA | XVI | 688172 | 688807 | 1 |
| YPR072W | 553.038133 | 1.0724688 | 0.67115959 | 5.61070849 | 0.06048534 | 0.15079771 | NOT5 | XVI | 690107 | 691789 | 1 |

| | | | | | | | | | | | |
|---|---|---|---|---|---|---|---|---|---|---|---|
| YPR073C | 39.5086949 | -0.4103083 | 0.66927303 | 2.41913912 | 0.29832566 | 0.47168367 | LTP1 | XVI | 691933 | 692418 | -1 |
| YPR074C | 111.917052 | 0.13978943 | 0.62272906 | 1.88514156 | 0.38962491 | 0.56101577 | TKL1 | XVI | 692796 | 694838 | -1 |
| YPR075C | 91.5168473 | 0.03621841 | 0.53453104 | 0.33791181 | 0.84454614 | 0.90907644 | OPY2 | XVI | 695737 | 696819 | -1 |
| YPR078C | 3551.47758 | 3.27789445 | 0.51437885 | 40.4220038 | 1.67E-09 | 5.89E-08 | NA | XVI | 697147 | 698265 | -1 |
| YPR079W | 15.5386709 | 0.80430603 | 0.88917412 | 1.39829974 | 0.49700765 | 0.65655356 | MRL1 | XVI | 698869 | 700014 | 1 |
| YPR080W | 108.552734 | -0.275229 | 0.7889275 | 5.84046541 | 0.05392114 | 0.1384494 | TEF1 | XVI | 700594 | 701970 | 1 |
| YPR081C | 300.970113 | 1.24762824 | 0.65718641 | 15.7651245 | 0.00037727 | 0.00293993 | GRS2 | XVI | 702114 | 703970 | -1 |
| YPR082C | 88.0293804 | 1.32873342 | 0.59780048 | 33.4497057 | 5.45E-08 | 1.42E-06 | DIB1 | XVI | 704225 | 704656 | -1 |
| YPR083W | 12.4207663 | -0.7375665 | 0.94293625 | 0.75416996 | 0.68585779 | 0.80812303 | MDM36 | XVI | 704854 | 706593 | 1 |
| YPR084W | 48.6250406 | 0.52413779 | 0.62215595 | 9.25738424 | 0.00976753 | 0.03870798 | NA | XVI | 706972 | 708342 | 1 |
| YPR085C | 281.912116 | -0.6352787 | 0.51348341 | 12.6850173 | 0.00175988 | 0.01017954 | ASA1 | XVI | 708497 | 709828 | -1 |
| YPR086W | 91.3255302 | 0.23109827 | 0.62149865 | 0.15321968 | 0.92625117 | 0.95376032 | SUA7 | XVI | 710101 | 711138 | 1 |
| YPR088C | 189.869246 | 2.20687008 | 0.80874073 | 11.1049768 | 0.0038778 | 0.0187416 | SRP54 | XVI | 711403 | 713028 | -1 |
| YPR089W | 151.229419 | 0.99905967 | 0.60586597 | 5.36116047 | 0.06852338 | 0.16558864 | NA | XVI | 713275 | 715941 | 1 |
| YPR091C | 71.4334244 | 0.7451343 | 0.65908155 | 2.88591515 | 0.23622806 | 0.40406514 | NVJ2 | XVI | 716156 | 718468 | -1 |
| YPR093C | 33.5784538 | -0.3644435 | 0.69440123 | 0.28636169 | 0.86659734 | 0.92150571 | ASR1 | XVI | 719558 | 720424 | -1 |
| YPR094W | 115.836428 | 1.86713942 | 0.79733943 | 8.66285808 | 0.01314874 | 0.04830662 | RDS3 | XVI | 720637 | 720960 | 1 |
| YPR095C | 25.6102537 | 0.80009868 | 0.70719317 | 1.74898461 | 0.41707371 | 0.58892811 | SYT1 | XVI | 721037 | 724717 | -1 |
| YPR096C | 13.1761713 | -0.5116302 | 0.89169942 | 1.14581722 | 0.56388293 | 0.71488553 | NA | XVI | 724842 | 725144 | -1 |
| YPR097W | 43.081126 | 0.63888805 | 0.68982749 | 1.93917433 | 0.37923957 | 0.55323686 | NA | XVI | 725394 | 728615 | 1 |
| YPR098C | 106.187201 | 0.76171464 | 0.57604569 | 22.0140575 | 1.66E-05 | 0.00020089 | NA | XVI | 728947 | 729528 | -1 |
| YPR099C | 2962.89302 | 0.05692622 | 0.56323418 | 3.36476788 | 0.1859302 | 0.34169876 | NA | XVI | 729757 | 730113 | -1 |
| YPR100W | 96.8956499 | -0.2475784 | 0.58097353 | 1.5008531 | 0.47216511 | 0.63393528 | MRPL51 | XVI | 729791 | 730213 | 1 |
| YPR101W | 79.8619494 | -0.6892082 | 0.52706657 | 4.98320405 | 0.08277725 | 0.19087227 | SNT309 | XVI | 730492 | 731019 | 1 |
| YPR102C | 105.88572 | 0.7329266 | 0.67980449 | 10.0014825 | 0.00673295 | 0.02911412 | RPL11A | XVI | 731224 | 731748 | -1 |
| YPR103W | 53.2917747 | 0.33990366 | 0.68203997 | 0.46231362 | 0.79361501 | 0.87457764 | PRE2 | XVI | 732349 | 733212 | 1 |
| YPR104C | 53.6749955 | -0.0327309 | 0.63140883 | 0.26431319 | 0.87620377 | 0.92712351 | FHL1 | XVI | 733623 | 736433 | -1 |
| YPR105C | 24.0145628 | -0.9415466 | 0.73532249 | 1.68736658 | 0.43012333 | 0.59923342 | COG4 | XVI | 736984 | 739569 | -1 |
| YPR106W | 188.100392 | 0.26414951 | 0.5057434 | 2.51056999 | 0.28499461 | 0.45862113 | ISR1 | XVI | 740061 | 741392 | 1 |
| YPR107C | 129.146495 | -0.4113954 | 0.58080276 | 3.79465473 | 0.1499689 | 0.29275789 | YTH1 | XVI | 741438 | 742064 | -1 |
| YPR108W | 49.8542291 | 0.44130582 | 0.70463032 | 0.42713621 | 0.80769715 | 0.88441988 | RPN7 | XVI | 742454 | 743743 | 1 |
| YPR109W | 22.6336722 | -0.4411758 | 0.70744922 | 7.42683642 | 0.024394 | 0.07671608 | NA | XVI | 744689 | 745573 | 1 |
| YPR110C | 42.6983544 | -0.6800938 | 0.84052969 | 1.60861814 | 0.44739694 | 0.61428226 | RPC40 | XVI | 745828 | 746835 | -1 |

| | | | | | | | | | | | |
|---|---|---|---|---|---|---|---|---|---|---|---|
| YPR111W | 14.7592286 | 0.23188162 | 0.85834758 | 2.09988779 | 0.34995738 | 0.52354248 | DBF20 | XVI | 747306 | 749000 | 1 |
| YPR112C | 56.5603051 | -0.9811805 | 0.61353983 | 13.5773329 | 0.00112647 | 0.00725502 | MRD1 | XVI | 749256 | 751919 | -1 |
| YPR113W | 14.006557 | -0.4287899 | 0.97540499 | 0.98375491 | 0.6114773 | 0.75489714 | PIS1 | XVI | 752257 | 752919 | 1 |
| YPR114W | 40.323588 | 1.03971406 | 0.68111066 | 2.39483099 | 0.30197366 | 0.47448396 | NA | XVI | 753301 | 754248 | 1 |
| YPR115W | 34.7210355 | -0.752229 | 0.74524948 | 1.20104716 | 0.54852436 | 0.70063006 | RGC1 | XVI | 754877 | 758128 | 1 |
| YPR116W | 12.3286806 | -1.5466358 | 0.92191225 | 3.08391093 | 0.2139623 | 0.37804141 | RRG8 | XVI | 758648 | 759481 | 1 |
| YPR117W | 56.5279885 | -0.397265 | 0.60079429 | 1.70689982 | 0.42594293 | 0.59578916 | NA | XVI | 760025 | 767494 | 1 |
| YPR118W | 65.464723 | 0.03462302 | 0.59042779 | 13.0142892 | 0.00149274 | 0.0091027 | MRI1 | XVI | 767752 | 768987 | 1 |
| YPR119W | 39.2633039 | -0.0047227 | 0.69413824 | 13.1900114 | 0.00136718 | 0.00848442 | CLB2 | XVI | 771653 | 773128 | 1 |
| YPR120C | 8.79983662 | -1.1097508 | 1.00188124 | 1.94417218 | 0.37829306 | 0.55223646 | CLB5 | XVI | 773875 | 775182 | -1 |
| YPR122W | 1392.99908 | -0.3364837 | 0.48619736 | 7.63006373 | 0.02203701 | 0.07124193 | AXL1 | XVI | 782045 | 785671 | 1 |
| YPR124W | 320.609035 | 0.05565263 | 0.4920583 | 11.1173075 | 0.00385396 | 0.01864197 | CTR1 | XVI | 786208 | 787428 | 1 |
| YPR125W | 21.0083741 | 0.12034168 | 0.80493038 | 0.29183376 | 0.86422954 | 0.92034008 | YLH47 | XVI | 787961 | 789325 | 1 |
| YPR127W | 261.990283 | 0.05510167 | 0.65155607 | 0.03594328 | 0.98218888 | 0.98644816 | NA | XVI | 790083 | 791120 | 1 |
| YPR128C | 27.7725382 | 0.42289689 | 0.67048498 | 2.52239651 | 0.28331434 | 0.45680591 | ANT1 | XVI | 791218 | 792204 | -1 |
| YPR129W | 346.507596 | 1.92725575 | 0.80112971 | 7.8518501 | 0.01972388 | 0.06522061 | SCD6 | XVI | 792687 | 793736 | 1 |
| YPR131C | 28.9309338 | 0.57662602 | 2.14583149 | 0.20658367 | 0.90186374 | 0.93824416 | NAT3 | XVI | 793911 | 794498 | -1 |
| YPR132W | 165.67757 | -0.7465485 | 0.70959498 | 15.1212461 | 0.00052055 | 0.00378167 | RPS23B | XVI | 794965 | 795767 | 1 |
| YPR133C | 82.2538506 | 0.29876878 | 0.59637696 | 1.54520921 | 0.46180867 | 0.62532091 | SPN1 | XVI | 795978 | 797210 | -1 |
| YPR133W-A | 18.2827308 | -1.8235475 | 0.9673153 | 7.01011563 | 0.03004504 | 0.08916492 | TOM5 | XVI | 797557 | 797709 | 1 |
| YPR134W | 167.716243 | -1.1367204 | 0.49279754 | 9.20529446 | 0.01002526 | 0.03951413 | MSS18 | XVI | 798051 | 798857 | 1 |
| YPR135W | 231.991345 | 0.17886336 | 0.50091074 | 5.78534621 | 0.05542785 | 0.14137538 | CTF4 | XVI | 799234 | 802017 | 1 |
| YPR137W | 42.3265204 | -0.8712846 | 0.65938577 | 3.54201691 | 0.1701613 | 0.32050551 | RRP9 | XVI | 802359 | 804080 | 1 |
| YPR138C | 2332.41412 | 1.10439856 | 0.7165968 | 6.22058673 | 0.04458787 | 0.12002035 | MEP3 | XVI | 810984 | 812453 | -1 |
| YPR139C | 38.116452 | 0.35707239 | 0.6564713 | 0.97459565 | 0.61428406 | 0.75689745 | LOA1 | XVI | 813156 | 814058 | -1 |
| YPR140W | 228.030671 | 0.50262923 | 0.61388887 | 73.9224345 | 8.87E-17 | 1.17E-14 | TAZ1 | XVI | 814391 | 815536 | 1 |
| YPR141C | 104.041312 | -0.3328774 | 0.57932259 | 13.1248784 | 0.00141244 | 0.00873719 | KAR3 | XVI | 815734 | 817923 | -1 |
| YPR143W | 47.2086741 | -0.9624217 | 0.68818528 | 10.1380085 | 0.00628868 | 0.0275011 | RRP15 | XVI | 818323 | 819075 | 1 |
| YPR144C | 96.6849167 | 0.26814984 | 0.56102898 | 23.4467957 | 8.10E-06 | 0.00010935 | NOC4 | XVI | 819765 | 821423 | -1 |
| YPR145C-A | 17.3632703 | 7.37023118 | 1.65239971 | 26.3143147 | 1.93E-06 | 3.17E-05 | NA | XVI | 824690 | 824926 | -1 |
| YPR145W | 65.2262466 | 0.14960672 | 0.63124259 | 7.56338517 | 0.02278409 | 0.07300493 | ASN1 | XVI | 822620 | 824338 | 1 |
| YPR147C | 17.2630117 | 0.9327983 | 0.85191251 | 2.76979909 | 0.25034895 | 0.42149474 | NA | XVI | 825645 | 826559 | -1 |
| YPR148C | 104.848504 | 1.32536571 | 0.60744847 | 4.78554442 | 0.09137602 | 0.20538321 | NA | XVI | 826833 | 828140 | -1 |

| | | | | | | | | | | | |
|---|---|---|---|---|---|---|---|---|---|---|---|
| YPR149W | 294.377135 | 0.9685012 | 0.47853737 | 6.59436646 | 0.03698721 | 0.10476085 | NCE102 | XVI | 829918 | 830439 | 1 |
| YPR151C | 76.4446537 | 1.75591493 | 0.69754583 | 8.28478312 | 0.01588482 | 0.05601627 | SUE1 | XVI | 831055 | 831675 | -1 |
| YPR152C | 47.3845413 | 0.30819619 | 0.66982541 | 13.6359354 | 0.00109394 | 0.00709286 | URN1 | XVI | 832061 | 833458 | -1 |
| YPR153W | 109.381204 | 1.77967859 | 0.64961599 | 49.5328177 | 1.75E-11 | 9.77E-10 | 24-May | XVI | 833689 | 834245 | 1 |
| YPR154W | 40.8861089 | 2.0871362 | 0.7549953 | 8.61677264 | 0.01345524 | 0.049183 | PIN3 | XVI | 834565 | 835212 | 1 |
| YPR155C | 55.1345794 | 0.9228178 | 0.61555554 | 6.30600471 | 0.04272366 | 0.11657399 | NCA2 | XVI | 835563 | 837413 | -1 |
| YPR156C | 49.1105503 | 1.06386665 | 0.86977133 | 13.0697735 | 0.00145189 | 0.00892151 | TPO3 | XVI | 837909 | 839777 | -1 |
| YPR157W | 29.4432781 | 1.32009489 | 0.80398876 | 24.7698062 | 4.18E-06 | 6.13E-05 | TDA6 | XVI | 841266 | 842669 | 1 |
| YPR158W | 21.5599466 | 1.05079689 | 0.83141233 | 5.86424349 | 0.05328386 | 0.13711714 | CUR1 | XVI | 843262 | 844020 | 1 |
| YPR159W | 75.5596377 | 1.76909897 | 0.65087397 | 8.07101308 | 0.01767672 | 0.06020484 | KRE6 | XVI | 857583 | 859745 | 1 |
| YPR160W | 250.266328 | -0.9518003 | 0.49886347 | 7.50605936 | 0.0234466 | 0.07454061 | GPH1 | XVI | 861306 | 864014 | 1 |
| YPR161C | 475.691265 | 0.55453647 | 0.53115399 | 30.9123856 | 1.94E-07 | 4.27E-06 | SGV1 | XVI | 864449 | 866422 | -1 |
| YPR162C | 122.055498 | 0.210508 | 0.53736035 | 14.8643199 | 0.00059191 | 0.00424214 | ORC4 | XVI | 866715 | 868304 | -1 |
| YPR163C | 90.8924626 | -0.4925791 | 0.61488156 | 9.78853969 | 0.00748938 | 0.03162131 | TIF3 | XVI | 868645 | 869955 | -1 |
| YPR164W | 35.2820128 | 0.79232925 | 0.68452462 | 6.61789132 | 0.03655469 | 0.10400574 | MMS1 | XVI | 870703 | 874926 | 1 |
| YPR165W | 102.945771 | 0.86066845 | 0.64085333 | 2.11010353 | 0.3481744 | 0.52188793 | RHO1 | XVI | 875368 | 875997 | 1 |
| YPR166C | 288.045616 | 2.67051979 | 0.49217502 | 30.8417151 | 2.01E-07 | 4.39E-06 | MRP2 | XVI | 876282 | 876629 | -1 |
| YPR167C | 96.1612302 | 1.80217137 | 0.56340146 | 12.1754355 | 0.00227058 | 0.01229812 | MET16 | XVI | 876847 | 877632 | -1 |
| YPR169W | 37.9062945 | -0.4020169 | 0.73355893 | 0.58581143 | 0.74609248 | 0.84537681 | JIP5 | XVI | 878690 | 880168 | 1 |
| YPR170W-B | 49.3001327 | 0.73623922 | 0.68592708 | 7.86136649 | 0.01963026 | 0.06510155 | NA | XVI | 883239 | 883595 | 1 |
| YPR171W | 224.073624 | -0.6191692 | 0.51349356 | 3.30318183 | 0.19174462 | 0.34826777 | BSP1 | XVI | 883828 | 885558 | 1 |
| YPR172W | 510.264979 | 0.04560079 | 0.60914873 | 3.54759931 | 0.16968701 | 0.31992439 | NA | XVI | 885796 | 886398 | 1 |
| YPR173C | 86.2053314 | 0.32736228 | 0.54697584 | 0.95865268 | 0.61920038 | 0.76021421 | VPS4 | XVI | 886524 | 887837 | -1 |
| YPR174C | 9.10928838 | 0.10797552 | 0.98578448 | 0.60505586 | 0.73894785 | 0.84040603 | NA | XVI | 888043 | 888708 | -1 |
| YPR175W | 730.649462 | 1.80864682 | 0.74569431 | 7.55983192 | 0.02282461 | 0.0730943 | DPB2 | XVI | 888974 | 891043 | 1 |
| YPR176C | 158.503041 | -0.5539253 | 0.55929722 | 11.0143749 | 0.0040575 | 0.01946391 | BET2 | XVI | 891101 | 892078 | -1 |
| YPR178W | 60.5813449 | -0.0218442 | 0.5877336 | 0.77860068 | 0.67753075 | 0.8011991 | PRP4 | XVI | 892332 | 893729 | 1 |
| YPR179C | 69.6040016 | 0.85290167 | 0.70633264 | 5.99677514 | 0.04986741 | 0.13023559 | HDA3 | XVI | 893797 | 895764 | -1 |
| YPR180W | 26.4959684 | 0.2365804 | 0.7340544 | 0.17658108 | 0.91549485 | 0.94807998 | AOS1 | XVI | 895961 | 897004 | 1 |
| YPR181C | 333.261303 | 1.60391267 | 0.55878987 | 57.8245057 | 2.78E-13 | 2.12E-11 | SEC23 | XVI | 897361 | 899667 | -1 |
| YPR182W | 17.8289841 | -0.8398528 | 0.89686991 | 2.46294141 | 0.29186302 | 0.46616698 | SMX3 | XVI | 900194 | 900454 | 1 |
| YPR183W | 71.2488422 | 0.58445966 | 0.61977423 | 0.9146666 | 0.63296933 | 0.76993539 | DPM1 | XVI | 900755 | 901558 | 1 |
| YPR184W | 29.7395926 | 0.18913201 | 0.86289968 | 2.92046462 | 0.23218233 | 0.39938672 | GDB1 | XVI | 902044 | 906654 | 1 |

| | | | | | | | | | | | |
|---|---|---|---|---|---|---|---|---|---|---|---|
| YPR185W | 32.4305316 | 1.11105063 | 0.70929498 | 3.5864862 | 0.16641958 | 0.31561394 | ATG13 | XVI | 907218 | 909434 | 1 |
| YPR186C | 57.2341255 | 1.5398481 | 0.68118628 | 8.06810298 | 0.01770246 | 0.06025706 | PZF1 | XVI | 909733 | 911022 | -1 |
| YPR187W | 61.8118719 | -0.8734363 | 0.67843203 | 15.5476264 | 0.00042061 | 0.00320015 | RPO26 | XVI | 911257 | 911800 | 1 |
| YPR188C | 164.438319 | -0.7457407 | 0.48795389 | 37.393422 | 7.59E-09 | 2.39E-07 | MLC2 | XVI | 911988 | 912479 | -1 |
| YPR189W | 73.763073 | -1.1090731 | 0.57361639 | 7.90556774 | 0.01920117 | 0.06416508 | SKI3 | XVI | 912664 | 916962 | 1 |
| YPR190C | 64.3540037 | 0.85367705 | 0.686639 | 6.14213605 | 0.0463716 | 0.123331 | RPC82 | XVI | 917077 | 919041 | -1 |
| YPR191W | 69.768793 | 0.90687284 | 0.89117533 | 1.68268579 | 0.43113117 | 0.59977162 | QCR2 | XVI | 919381 | 920487 | 1 |
| YPR192W | 1226.10006 | 1.25272614 | 0.52494332 | 22.1499196 | 1.55E-05 | 0.0001913 | AQY1 | XVI | 921860 | 922777 | 1 |
| YPR193C | 9.43111939 | 0.14537979 | 1.04011193 | 0.62824486 | 0.7304296 | 0.83678999 | HPA2 | XVI | 922909 | 923379 | -1 |
| YPR194C | 1639.24605 | -0.2376445 | 0.49267303 | 0.97986154 | 0.61266881 | 0.75555962 | OPT2 | XVI | 924304 | 926937 | -1 |
| YPR195C | 312.947221 | 0.33504698 | 0.5255681 | 0.98315896 | 0.61165953 | 0.75495815 | NA | XVI | 927965 | 928294 | -1 |
| YPR196W | 180.560311 | 0.27652588 | 0.53211318 | 24.389874 | 5.06E-06 | 7.19E-05 | NA | XVI | 931376 | 932788 | 1 |
| YPR198W | 52.9415312 | 0.34873823 | 0.59313076 | 0.80669905 | 0.66807855 | 0.79444823 | SGE1 | XVI | 934034 | 935665 | 1 |
| YPR199C | 558.329676 | 3.28396744 | 0.50557397 | 54.2806462 | 1.63E-12 | 1.14E-10 | ARR1 | XVI | 938148 | 939032 | -1 |
| YPR200C | 50.9232182 | 0.15572899 | 0.64777056 | 1.51007944 | 0.46999195 | 0.63255541 | ARR2 | XVI | 939279 | 939671 | -1 |
| YPR201W | 359.508137 | -0.2662636 | 0.50130383 | 0.40614167 | 0.81622042 | 0.88933313 | ARR3 | XVI | 939922 | 941136 | 1 |
| YPR202W | 21.6403239 | -1.5261144 | 0.76411174 | 6.71137743 | 0.03488534 | 0.10019152 | NA | XVI | 943032 | 943896 | 1 |

Table S6. Differential gene expression analysis for Myo3-AA treated with rapamycin for 5 hours.

| Row.names | baseMean | log2FoldCha | lfcSE | stat | pvalue | padj | genename | chrom | start_pos | end_pos | strand |
|---|---|---|---|---|---|---|---|---|---|---|---|
| Q0158 | 6.52489728 | -0.3800533 | 1.2165101 | 2.46242714 | 0.29193808 | 0.46616698 | NA | Mito | 58009 | 62447 | 1 |
| YAL001C | 152.194332 | 2.49367213 | 0.76566832 | 11.4156737 | 0.00331985 | 0.01648534 | TFC3 | I | 147594 | 151166 | -1 |
| YAL002W | 78.7260454 | -0.4489666 | 0.69556908 | 0.72275892 | 0.69671457 | 0.81659461 | VPS8 | I | 143707 | 147531 | 1 |
| YAL003W | 117.199149 | -0.9495519 | 0.59315419 | 2.55984277 | 0.27805916 | 0.45261809 | EFB1 | I | 142174 | 143160 | 1 |
| YAL005C | 85.6274446 | 1.44349859 | 0.63763535 | 6.74773434 | 0.0342569 | 0.09906519 | SSA1 | I | 139503 | 141431 | -1 |
| YAL007C | 171.012771 | 2.0395527 | 0.4923142 | 20.9886448 | 2.77E-05 | 0.00031564 | ERP2 | I | 137698 | 138345 | -1 |
| YAL008W | 16.4436684 | 1.22830781 | 0.81573601 | 2.61332338 | 0.2707223 | 0.44417176 | FUN14 | I | 136914 | 137510 | 1 |
| YAL009W | 16.1660335 | -0.1907233 | 0.75603377 | 0.07451714 | 0.96342699 | 0.9730058 | SPO7 | I | 135854 | 136633 | 1 |
| YAL010C | 74.0582927 | -1.3086421 | 0.55013551 | 6.09173003 | 0.04755516 | 0.12572802 | MDM10 | I | 134184 | 135665 | -1 |
| YAL011W | 219.581007 | -1.3253828 | 0.52196889 | 6.27233632 | 0.04344897 | 0.11810776 | SWC3 | I | 132199 | 134076 | 1 |
| YAL012W | 197.449702 | 0.95023848 | 0.48787994 | 4.18403485 | 0.12343786 | 0.25497866 | CYS3 | I | 130799 | 131983 | 1 |
| YAL013W | 173.375851 | -0.2419518 | 0.68869996 | 2.92971046 | 0.23111145 | 0.39792139 | DEP1 | I | 129270 | 130487 | 1 |
| YAL014C | 26.3703385 | -0.3169271 | 0.73582092 | 2.05406147 | 0.35806858 | 0.53120548 | SYN8 | I | 128252 | 129019 | -1 |
| YAL015C | 12.5995056 | 0.2544209 | 0.96476584 | 0.71046144 | 0.70101169 | 0.81801018 | NTG1 | I | 126903 | 128102 | -1 |
| YAL016W | 84.6687343 | 0.19549395 | 0.57183104 | 0.13350263 | 0.9354278 | 0.95803272 | TPD3 | I | 124879 | 126786 | 1 |
| YAL017W | 41.247613 | -0.1608243 | 0.66508234 | 0.10729795 | 0.94776474 | 0.96435226 | PSK1 | I | 120225 | 124295 | 1 |
| YAL018C | 8.76638298 | 0.09605213 | 0.9185981 | 1.06893526 | 0.58598117 | 0.73378964 | LDS1 | I | 118564 | 119541 | -1 |
| YAL019W | 33.9776591 | -1.5746316 | 0.70511753 | 6.17671038 | 0.04557686 | 0.12191042 | FUN30 | I | 114919 | 118314 | 1 |
| YAL020C | 39.0307728 | 0.14425252 | 0.66619878 | 1.00110037 | 0.60619705 | 0.75088539 | ATS1 | I | 113614 | 114615 | -1 |
| YAL021C | 79.1822695 | -0.3180904 | 0.61662532 | 0.43478161 | 0.80461546 | 0.88279205 | CCR4 | I | 110846 | 113359 | -1 |
| YAL022C | 119.31372 | 1.9709024 | 0.6085598 | 14.6928354 | 0.0006449 | 0.00457593 | FUN26 | I | 108877 | 110430 | -1 |
| YAL023C | 85.7619927 | 0.01105289 | 0.58999964 | 3.64225043 | 0.16184354 | 0.31028944 | PMT2 | I | 106272 | 108551 | -1 |
| YAL024C | 13.6197097 | 1.08276635 | 0.96238518 | 4.03902184 | 0.13272036 | 0.26859521 | LTE1 | I | 101565 | 105872 | -1 |
| YAL025C | 102.736369 | -3.5994047 | 0.60995535 | 30.7178657 | 2.14E-07 | 4.63E-06 | MAK16 | I | 100225 | 101145 | -1 |
| YAL026C | 26.8507426 | -0.0176943 | 0.8062049 | 0.04998457 | 0.97531744 | 0.9809081 | DRS2 | I | 95630 | 99697 | -1 |
| YAL027W | 15.4075861 | -0.5856643 | 0.81593611 | 2.31500854 | 0.31426953 | 0.48639952 | SAW1 | I | 94687 | 95472 | 1 |
| YAL028W | 22.7228518 | -1.8369321 | 0.733352 | 9.11580546 | 0.01048402 | 0.04090465 | FRT2 | I | 92900 | 94486 | 1 |
| YAL029C | 264.124837 | 0.65028733 | 0.90072359 | 7.8771885 | 0.01947557 | 0.06484392 | MYO4 | I | 87855 | 92270 | -1 |
| YAL030W | 174.597782 | -0.8821113 | 0.56173339 | 8.22967359 | 0.01632861 | 0.05693318 | SNC1 | I | 87286 | 87752 | 1 |
| YAL031C | 98.4896929 | 0.63579159 | 0.55663561 | 1.94896973 | 0.37738671 | 0.55108929 | GIP4 | I | 84749 | 87031 | -1 |
| YAL032C | 281.557756 | 4.0156194 | 0.54512161 | 48.3848365 | 3.11E-11 | 1.58E-09 | PRP45 | I | 83335 | 84474 | -1 |

| | | | | | | | | | | | |
|---|---|---|---|---|---|---|---|---|---|---|---|
| YAL033W | 52.7827913 | 0.86117669 | 0.65564554 | 2.54871713 | 0.27961026 | 0.45373975 | POP5 | I | 82706 | 83227 | 1 |
| YAL034C | 86.4634567 | 2.66974684 | 0.61258046 | 29.7533699 | 3.46E-07 | 6.96E-06 | FUN19 | I | 80710 | 81951 | -1 |
| YAL034W-A | 366.639954 | 3.91548884 | 0.62851362 | 32.5295341 | 8.64E-08 | 2.15E-06 | MTW1 | I | 79718 | 80587 | 1 |
| YAL035W | 690.188751 | -1.6699026 | 0.51955716 | 11.3865416 | 0.00336856 | 0.0166843 | FUN12 | I | 76427 | 79435 | 1 |
| YAL036C | 24.7767781 | -1.2817293 | 0.8974584 | 2.14681962 | 0.34184092 | 0.51556627 | RBG1 | I | 75043 | 76152 | -1 |
| YAL037C-A | 24876.3172 | 0.47639824 | 2.03425855 | 4.16720322 | 0.12448107 | 0.25658434 | NA | I | 73426 | 73518 | -1 |
| YAL037W | 182.809816 | 0.31735257 | 0.54651204 | 2.34600947 | 0.30943577 | 0.48239987 | NA | I | 74020 | 74823 | 1 |
| YAL038W | 1192.55953 | -0.8149701 | 0.47515188 | 8.09941568 | 0.01742747 | 0.05974247 | CDC19 | I | 71786 | 73288 | 1 |
| YAL039C | 185.960627 | -1.2339051 | 0.59344965 | 4.76423941 | 0.09235461 | 0.20685781 | CYC3 | I | 68716 | 69525 | -1 |
| YAL040C | 44.749424 | 1.08114333 | 0.59118371 | 4.66264229 | 0.09716729 | 0.21489634 | CLN3 | I | 65778 | 67520 | -1 |
| YAL041W | 143.741223 | 0.06584114 | 0.81917343 | 6.35754028 | 0.04163683 | 0.11447163 | CDC24 | I | 62840 | 65404 | 1 |
| YAL042W | 218.964554 | 2.66351594 | 0.58776986 | 25.776984 | 2.53E-06 | 3.94E-05 | ERV46 | I | 61316 | 62563 | 1 |
| YAL043C | 41.7538945 | -0.1240638 | 0.65235195 | 0.85382898 | 0.65251935 | 0.78302322 | PTA1 | I | 58695 | 61052 | -1 |
| YAL044C | 295.413083 | 0.63561544 | 0.88998946 | 9.89778291 | 0.00709127 | 0.0303238 | GCV3 | I | 57950 | 58462 | -1 |
| YAL044W-A | 15.1812068 | -0.976336 | 0.91100346 | 2.70746223 | 0.25827481 | 0.43008401 | BOL1 | I | 57518 | 57850 | 1 |
| YAL046C | 43.4614896 | -2.7608839 | 0.68234482 | 16.7369544 | 0.00023207 | 0.00193893 | BOL3 | I | 57029 | 57385 | -1 |
| YAL047C | 19.8213171 | 0.08760871 | 0.75668446 | 0.52608763 | 0.76870821 | 0.85939768 | SPC72 | I | 54989 | 56857 | -1 |
| YAL048C | 60.3846288 | -0.1520875 | 0.55433261 | 0.56227894 | 0.75492304 | 0.85188158 | GEM1 | I | 52801 | 54789 | -1 |
| YAL049C | 40.0566149 | 0.70343084 | 0.89012406 | 8.288864 | 0.01585244 | 0.05596684 | AIM2 | I | 51855 | 52595 | -1 |
| YAL051W | 25.0791615 | 1.60990335 | 0.73263889 | 5.04199906 | 0.08037923 | 0.18705615 | OAF1 | I | 48564 | 51707 | 1 |
| YAL053W | 134.787587 | 2.35278292 | 0.54481853 | 23.0246777 | 1.00E-05 | 0.00013257 | FLC2 | I | 45899 | 48250 | 1 |
| YAL054C | 497.265684 | -1.0704757 | 0.48929057 | 5.4794021 | 0.06458965 | 0.15839648 | ACS1 | I | 42881 | 45022 | -1 |
| YAL055W | 66.0301892 | 0.46990851 | 1.03651718 | 2.23566208 | 0.32698825 | 0.49980517 | PEX22 | I | 42177 | 42719 | 1 |
| YAL056W | 22.520543 | 0.33102243 | 0.78283042 | 0.25185743 | 0.88167769 | 0.9300006 | GPB2 | I | 39259 | 41901 | 1 |
| YAL058W | 33.6448257 | 0.14883299 | 0.63310163 | 0.26338621 | 0.87660998 | 0.92721443 | CNE1 | I | 37464 | 38972 | 1 |
| YAL059W | 17.4722431 | -2.2064412 | 0.82395877 | 8.26261199 | 0.01606189 | 0.05643103 | ECM1 | I | 36509 | 37147 | 1 |
| YAL060W | 49.8893578 | -1.2939656 | 0.63248257 | 5.70830547 | 0.05760461 | 0.1457739 | BDH1 | I | 35155 | 36303 | 1 |
| YAL061W | 827.748389 | -1.5883672 | 0.53860922 | 13.3813506 | 0.00124244 | 0.00786166 | BDH2 | I | 33448 | 34701 | 1 |
| YAL062W | 1323.60444 | -0.6995085 | 0.51123735 | 1.89413611 | 0.38787659 | 0.56008824 | GDH3 | I | 31567 | 32940 | 1 |
| YAL063C | 1429.86781 | -0.1079025 | 0.50700674 | 0.17975256 | 0.91404426 | 0.94725547 | FLO9 | I | 24000 | 27968 | -1 |
| YAL063C-A | 22.9971074 | -0.359072 | 0.71463457 | 0.71313174 | 0.70007636 | 0.81801018 | NA | I | 22395 | 22685 | -1 |
| YAL064C-A | 286.680621 | -1.1521046 | 0.5225971 | 8.06620496 | 0.01771927 | 0.06027884 | TDA8 | I | 13363 | 13743 | -1 |
| YAL064W-B | 34.2044202 | -1.1511513 | 0.65573958 | 3.25838733 | 0.19608762 | 0.35380098 | NA | I | 12046 | 12426 | 1 |

| | | | | | | | | | | | |
|---|---|---|---|---|---|---|---|---|---|---|---|
| YAL065C | 69.9340375 | -0.7029099 | 0.56580635 | 3.24044675 | 0.1978545 | 0.35610119 | NA | I | 11565 | 11951 | -1 |
| YAL067C | 148.676613 | 0.59593971 | 0.53809593 | 2.90969397 | 0.23343608 | 0.40082886 | SEO1 | I | 7235 | 9016 | -1 |
| YAR002C-A | 25.4323182 | -0.5092077 | 0.67283665 | 1.55151091 | 0.46035587 | 0.62448065 | ERP1 | I | 154065 | 154724 | -1 |
| YAR002W | 97.1288927 | -0.9004842 | 0.65621894 | 6.30174738 | 0.0428147 | 0.11665747 | NUP60 | I | 152257 | 153876 | 1 |
| YAR003W | 31.4873408 | 0.24868867 | 0.62857413 | 0.52461139 | 0.76927583 | 0.85953436 | SWD1 | I | 155005 | 156285 | 1 |
| YAR007C | 217.684773 | -0.2350494 | 0.69929867 | 6.20963477 | 0.0448327 | 0.12051131 | RFA1 | I | 156754 | 158619 | -1 |
| YAR008W | 9.97066093 | -0.4689096 | 0.89828081 | 1.64471825 | 0.43939385 | 0.60805219 | SEN34 | I | 158966 | 159793 | 1 |
| YAR010C | 8.12399364 | -0.7367097 | 0.97214054 | 1.21944634 | 0.54350131 | 0.69699868 | NA | I | 164544 | 165866 | -1 |
| YAR014C | 49.9580748 | -1.1517044 | 0.5964935 | 3.78940721 | 0.1503629 | 0.2931317 | BUD14 | I | 166742 | 168871 | -1 |
| YAR015W | 88.8911506 | -2.6503499 | 0.63353855 | 19.0755842 | 7.21E-05 | 0.00071215 | ADE1 | I | 169375 | 170295 | 1 |
| YAR018C | 163.651055 | 0.40687386 | 0.52954721 | 0.63853774 | 0.72668014 | 0.83481707 | KIN3 | I | 170396 | 171703 | -1 |
| YAR019C | 67.7457183 | -1.1550988 | 0.60331586 | 3.65945777 | 0.16045706 | 0.30793716 | CDC15 | I | 172211 | 175135 | -1 |
| YAR023C | 83.6213596 | 0.98943736 | 0.58414311 | 9.25607291 | 0.00977393 | 0.03870798 | NA | I | 179281 | 179820 | -1 |
| YAR027W | 69.3232713 | 0.64334524 | 0.58013463 | 1.26988498 | 0.52996597 | 0.68432269 | UIP3 | I | 183770 | 184477 | 1 |
| YAR028W | 33.6169365 | -1.434577 | 0.62452909 | 5.56038257 | 0.06202664 | 0.15354179 | NA | I | 184892 | 185596 | 1 |
| YAR029W | 35.7153511 | -1.7020277 | 0.72949379 | 6.18521251 | 0.04538352 | 0.12165304 | NA | I | 186321 | 186545 | 1 |
| YAR031W | 441.518308 | -1.2398681 | 0.53455252 | 6.26412197 | 0.04362779 | 0.11842705 | PRM9 | I | 186836 | 187732 | 1 |
| YAR033W | 291.320116 | -0.8733506 | 0.52910279 | 6.69695967 | 0.03513773 | 0.10076644 | MST28 | I | 188107 | 188811 | 1 |
| YAR035C-A | 55.3368512 | 1.03044514 | 0.72477094 | 2.03780421 | 0.36099105 | 0.53387949 | NA | I | 192337 | 192417 | -1 |
| YAR035W | 120.492423 | -0.6559016 | 0.72211306 | 1.97912543 | 0.37173921 | 0.54514152 | YAT1 | I | 190193 | 192256 | 1 |
| YAR042W | 51.8015762 | 0.32157167 | 0.60211288 | 0.30227492 | 0.85972951 | 0.91706593 | SWH1 | I | 192619 | 196185 | 1 |
| YAR050W | 305.750074 | 0.07173748 | 0.53715775 | 2.76248446 | 0.25126623 | 0.42270232 | FLO1 | I | 203403 | 208016 | 1 |
| YAR061W | 17.5073638 | -2.5910643 | 0.8224921 | 9.46594497 | 0.00880027 | 0.03573389 | NA | I | 218140 | 219145 | 1 |
| YAR064W | 147.531005 | 0.11212178 | 0.51627658 | 5.58603885 | 0.06123604 | 0.15215948 | NA | I | 220198 | 220497 | 1 |
| YAR066W | 8.62362385 | -0.9631414 | 1.05122343 | 6.14678384 | 0.04626396 | 0.12315786 | NA | I | 221049 | 221660 | 1 |
| YAR068W | 74.5682182 | 0.22748234 | 0.57874728 | 6.02355672 | 0.0492041 | 0.12908551 | NA | I | 222406 | 222891 | 1 |
| YBL001C | 53.1066746 | -1.2309543 | 0.64117194 | 3.89050831 | 0.14295089 | 0.2827761 | ECM15 | II | 237153 | 237467 | -1 |
| YBL002W | 56.0987531 | -3.23077 | 0.76473317 | 19.5508116 | 5.68E-05 | 0.00057634 | HTB2 | II | 236492 | 236887 | 1 |
| YBL003C | 9.30485265 | -2.874976 | 1.03438607 | 8.49506124 | 0.0142995 | 0.05164948 | HTA2 | II | 235394 | 235792 | -1 |
| YBL004W | 157.787747 | 0.09895928 | 0.58581553 | 1.9257807 | 0.38178779 | 0.55529381 | UTP20 | II | 227636 | 235117 | 1 |
| YBL005W | 3766.36863 | -1.2096766 | 0.48078042 | 7.24215194 | 0.02675387 | 0.08152891 | PDR3 | II | 217470 | 220400 | 1 |
| YBL005W-B | 8.43483097 | 0.39764042 | 0.95300614 | 0.2868475 | 0.86638686 | 0.92150571 | NA | II | 221330 | 226598 | 1 |
| YBL006C | 330.345265 | 2.33309413 | 0.5456018 | 19.7882846 | 5.05E-05 | 0.00051812 | LDB7 | II | 216587 | 217129 | -1 |

| | | | | | | | | | | | |
|---|---|---|---|---|---|---|---|---|---|---|---|
| YBL007C | 70.9587208 | -0.1445175 | 0.56717848 | 1.56572718 | 0.4570952 | 0.62243209 | SLA1 | II | 212632 | 216366 | -1 |
| YBL008W | 9.26943991 | 0.23481414 | 0.99902815 | 0.22319884 | 0.89440246 | 0.9350658 | HIR1 | II | 209653 | 212175 | 1 |
| YBL008W-A | 9.75644654 | 1.47944973 | 0.93678211 | 3.22021206 | 0.19986642 | 0.35894125 | NA | II | 209409 | 209648 | 1 |
| YBL009W | 81.3175795 | 0.39864787 | 0.55459332 | 0.52799488 | 0.7679755 | 0.85918999 | ALK2 | II | 207194 | 209224 | 1 |
| YBL010C | 11.6849997 | 0.2510452 | 0.84641624 | 0.25620379 | 0.87976373 | 0.92911891 | NA | II | 206107 | 206949 | -1 |
| YBL011W | 120.42145 | -1.3302361 | 0.50025605 | 7.62458665 | 0.02209744 | 0.07131784 | SCT1 | II | 203538 | 205817 | 1 |
| YBL013W | 45.950479 | -0.2317046 | 0.63209215 | 0.61788118 | 0.73422439 | 0.83775415 | FMT1 | II | 202056 | 203261 | 1 |
| YBL014C | 19.8043631 | -0.1569607 | 0.72407437 | 0.04845999 | 0.9760612 | 0.98148564 | RRN6 | II | 199064 | 201748 | -1 |
| YBL015W | 41.1753417 | 0.6303932 | 0.62151912 | 6.93193515 | 0.03124276 | 0.09229367 | ACH1 | II | 194122 | 195702 | 1 |
| YBL016W | 21.4586369 | 1.03468293 | 0.70820579 | 5.33290125 | 0.06949846 | 0.16745573 | FUS3 | II | 192451 | 193512 | 1 |
| YBL017C | 62.5472072 | 0.459 | 0.83914187 | 0.48857882 | 0.78326092 | 0.86895587 | PEP1 | II | 186844 | 191583 | -1 |
| YBL018C | 96.7110787 | 1.94485348 | 0.92740649 | 7.74443175 | 0.0208122 | 0.06800375 | POP8 | II | 185998 | 186474 | -1 |
| YBL019W | 428.27037 | 1.0638526 | 0.46579866 | 12.3005612 | 0.00213288 | 0.01172782 | APN2 | II | 184353 | 185915 | 1 |
| YBL020W | 32.5005079 | 0.05726825 | 0.62307182 | 0.91607105 | 0.632525 | 0.76955658 | RFT1 | II | 182401 | 184125 | 1 |
| YBL021C | 69.7223875 | -1.2230014 | 0.62594552 | 6.54887142 | 0.03783821 | 0.10666176 | HAP3 | II | 181660 | 182094 | -1 |
| YBL022C | 41.2835978 | -0.2107198 | 0.66123437 | 0.44984012 | 0.79858006 | 0.87821055 | PIM1 | II | 177874 | 181275 | -1 |
| YBL023C | 102.386519 | 1.04420656 | 0.51538733 | 4.40254901 | 0.11066203 | 0.23547709 | MCM2 | II | 174920 | 177526 | -1 |
| YBL024W | 43.4956088 | -2.2277141 | 0.88128145 | 6.22838802 | 0.04441429 | 0.11983849 | NCL1 | II | 172534 | 174588 | 1 |
| YBL025W | 112.266141 | 1.12139077 | 0.54701362 | 7.12594913 | 0.02835436 | 0.0851955 | RRN10 | II | 171481 | 171918 | 1 |
| YBL026W | 7.21219264 | 1.81934603 | 1.18054347 | 4.10672284 | 0.1283029 | 0.26194421 | LSM2 | II | 170623 | 171038 | 1 |
| YBL027W | 367.323685 | -1.648504 | 0.6147279 | 7.19141098 | 0.02744132 | 0.08314728 | RPL19B | II | 168423 | 169376 | 1 |
| YBL028C | 25.7178279 | -2.3769513 | 0.72412563 | 11.9589722 | 0.00253013 | 0.01336627 | NA | II | 167518 | 167838 | -1 |
| YBL029C-A | 47.5392462 | -1.9654564 | 0.61112193 | 9.99336414 | 0.00676034 | 0.02921072 | NA | II | 164488 | 164772 | -1 |
| YBL029W | 1712.60259 | 2.1590219 | 0.54740249 | 14.3704963 | 0.00075768 | 0.00521018 | NA | II | 166134 | 167264 | 1 |
| YBL030C | 95.1418224 | -0.3283582 | 0.64702623 | 0.276045 | 0.87107909 | 0.92508216 | PET9 | II | 163041 | 163997 | -1 |
| YBL032W | 89.9842815 | -1.3595954 | 0.51565418 | 8.69119495 | 0.01296376 | 0.0478395 | HEK2 | II | 160184 | 161329 | 1 |
| YBL033C | 26.6861463 | 1.34594124 | 0.76816987 | 3.38265696 | 0.18427456 | 0.33979289 | RIB1 | II | 158656 | 159693 | -1 |
| YBL034C | 109.306107 | 1.735026 | 0.55075977 | 22.4493111 | 1.33E-05 | 0.00016903 | STU1 | II | 153848 | 158389 | -1 |
| YBL035C | 317.806195 | 3.00617205 | 0.50060796 | 48.9089536 | 2.40E-11 | 1.30E-09 | POL12 | II | 151493 | 153610 | -1 |
| YBL036C | 32.439987 | 1.02006349 | 0.64637408 | 2.71430807 | 0.25739227 | 0.42958807 | NA | II | 150447 | 151220 | -1 |
| YBL037W | 21.1683494 | 0.98814659 | 0.77387332 | 1.78364015 | 0.40990901 | 0.58142409 | APL3 | II | 147209 | 150286 | 1 |
| YBL038W | 35.2141507 | -0.238712 | 0.67643159 | 2.70934266 | 0.25803209 | 0.42992972 | MRPL16 | II | 146187 | 146885 | 1 |
| YBL039C | 54.7332205 | -0.0618685 | 0.67076118 | 5.74223134 | 0.0566357 | 0.14401437 | URA7 | II | 143989 | 145728 | -1 |

| | | | | | | | | | | | |
|---|---|---|---|---|---|---|---|---|---|---|---|
| YBL039W-B | 15.1852469 | 0.31967608 | 0.83704569 | 0.40524203 | 0.81658766 | 0.88956586 | NA | II | 143393 | 143572 | 1 |
| YBL040C | 222.504594 | 0.09743968 | 0.70513606 | 15.5111039 | 0.00042836 | 0.00324208 | ERD2 | II | 142112 | 142868 | -1 |
| YBL041W | 52.1723582 | -0.6392016 | 0.63222261 | 9.2774313 | 0.00967011 | 0.03840187 | PRE7 | II | 141247 | 141972 | 1 |
| YBL042C | 53.4812589 | -0.5667978 | 0.57738333 | 2.79206681 | 0.24757706 | 0.41832973 | FUI1 | II | 138341 | 140260 | -1 |
| YBL045C | 27.9930743 | 0.11991212 | 0.82661413 | 0.06654808 | 0.96727345 | 0.97486964 | COR1 | II | 134143 | 135516 | -1 |
| YBL046W | 24.1545039 | 0.55447729 | 0.77588137 | 2.00228486 | 0.36745941 | 0.54054623 | PSY4 | II | 132424 | 133749 | 1 |
| YBL047C | 113.041398 | -0.447731 | 0.52243331 | 2.9115292 | 0.23322197 | 0.40069888 | EDE1 | II | 127895 | 132040 | -1 |
| YBL048W | 303.200081 | -2.4466031 | 0.7954696 | 12.2922292 | 0.00214179 | 0.01176561 | NA | II | 127299 | 127610 | 1 |
| YBL049W | 231.32859 | -3.3197212 | 0.57372133 | 29.8097622 | 3.36E-07 | 6.83E-06 | MOH1 | II | 126828 | 127244 | 1 |
| YBL050W | 36.1125016 | 0.47602245 | 0.76813697 | 0.72594747 | 0.6956047 | 0.81611981 | SEC17 | II | 125125 | 126119 | 1 |
| YBL051C | 57.4560575 | -0.4163463 | 0.5801615 | 1.11412119 | 0.57289055 | 0.72277589 | PIN4 | II | 122753 | 124759 | -1 |
| YBL052C | 222.49205 | 3.23834419 | 0.54283079 | 42.2045152 | 6.85E-10 | 2.54E-08 | SAS3 | II | 119379 | 121874 | -1 |
| YBL054W | 60.7158248 | 1.55868389 | 0.60737325 | 10.8555403 | 0.00439288 | 0.02061165 | TOD6 | II | 117589 | 119166 | 1 |
| YBL055C | 452.452705 | -1.8976917 | 0.53167704 | 12.7963205 | 0.00166462 | 0.00978886 | NA | II | 115573 | 116829 | -1 |
| YBL056W | 301.265659 | -1.0995365 | 0.47761563 | 7.31504693 | 0.02579632 | 0.07948946 | PTC3 | II | 113762 | 115168 | 1 |
| YBL057C | 297.034107 | 0.03735766 | 0.84942952 | 7.27661744 | 0.02629678 | 0.08055998 | PTH2 | II | 112801 | 113427 | -1 |
| YBL058W | 71.9131435 | 0.22079237 | 0.57449002 | 0.60468581 | 0.73908458 | 0.84040603 | SHP1 | II | 111437 | 112708 | 1 |
| YBL059C-A | 10.3655581 | -1.1430156 | 0.97229611 | 3.68273679 | 0.15860025 | 0.30518293 | CMC2 | II | 110125 | 110539 | -1 |
| YBL059W | 106.594204 | 0.4314422 | 0.59800927 | 2.23885061 | 0.32646736 | 0.49927259 | NA | II | 110594 | 111244 | 1 |
| YBL060W | 224.055767 | -1.5829778 | 0.62489574 | 7.07499372 | 0.02908604 | 0.08680999 | YEL1 | II | 107932 | 109995 | 1 |
| YBL061C | 68.3948846 | 0.37822884 | 0.71611981 | 4.3733762 | 0.11228802 | 0.23858629 | SKT5 | II | 105316 | 107406 | -1 |
| YBL063W | 114.113116 | -2.0337221 | 0.55723501 | 15.4428098 | 0.00044324 | 0.00334159 | KIP1 | II | 101886 | 105221 | 1 |
| YBL064C | 12.5639633 | 0.33169235 | 0.84336353 | 0.27244411 | 0.87264884 | 0.92527295 | PRX1 | II | 100371 | 101156 | -1 |
| YBL066C | 227.590091 | 2.20407694 | 0.54749858 | 22.8304011 | 1.10E-05 | 0.00014379 | SEF1 | II | 96669 | 100115 | -1 |
| YBL067C | 40.0367263 | -0.9733642 | 0.6078198 | 3.07495386 | 0.21492268 | 0.3788135 | UBP13 | II | 93639 | 95882 | -1 |
| YBL069W | 144.723039 | -1.241399 | 0.51729091 | 5.81016596 | 0.05474425 | 0.14006486 | AST1 | II | 90741 | 92030 | 1 |
| YBL071C | 1329.8121 | -1.0169414 | 0.50269559 | 4.1168539 | 0.12765462 | 0.26140797 | NA | II | 90223 | 90531 | -1 |
| YBL071W-A | 8.80941821 | -2.4218006 | 1.00388113 | 6.01916821 | 0.04931218 | 0.12931048 | KTI11 | II | 89978 | 90226 | 1 |
| YBL072C | 110.818765 | -2.6515705 | 0.61590257 | 16.9957962 | 0.0002039 | 0.00173612 | RPS8A | II | 88523 | 89125 | -1 |
| YBL075C | 32.4599416 | 0.2738383 | 0.73718991 | 1.69780288 | 0.42788473 | 0.59755248 | SSA3 | II | 84499 | 86448 | -1 |
| YBL076C | 102.332662 | -0.4232574 | 0.69668 | 0.44529037 | 0.80039879 | 0.87954242 | ILS1 | II | 81043 | 84261 | -1 |
| YBL078C | 9.26171442 | 5.25317282 | 1.54987744 | 13.335567 | 0.00127121 | 0.00800643 | ATG8 | II | 80378 | 80731 | -1 |
| YBL079W | 67.4164512 | 2.35638775 | 0.60147962 | 29.4295085 | 4.07E-07 | 8.07E-06 | NUP170 | II | 75259 | 79767 | 1 |

| | | | | | | | | | | | |
|---|---|---|---|---|---|---|---|---|---|---|---|
| YBL080C | 117.420865 | -1.6717309 | 0.86436942 | 7.71033728 | 0.02117003 | 0.06905605 | PET112 | II | 73070 | 74695 | -1 |
| YBL081W | 96.7393112 | -0.3571525 | 0.76253015 | 4.02228974 | 0.13383536 | 0.26992447 | NA | II | 71866 | 72972 | 1 |
| YBL082C | 12.0103079 | 0.21488496 | 0.84862816 | 0.31269134 | 0.8552635 | 0.91415464 | ALG3 | II | 69751 | 71127 | -1 |
| YBL084C | 8.01706199 | 0.39625883 | 1.00772544 | 3.46665665 | 0.17669533 | 0.32927775 | CDC27 | II | 67169 | 69445 | -1 |
| YBL085W | 24.9589338 | -1.2920986 | 0.73055058 | 3.66874054 | 0.15971405 | 0.3068163 | BOI1 | II | 63876 | 66818 | 1 |
| YBL086C | 23.4037512 | -0.6733468 | 0.70283542 | 1.10675015 | 0.57500584 | 0.72422968 | NA | II | 61202 | 62602 | -1 |
| YBL087C | 68.1187718 | 0.54886431 | 0.67761393 | 1.8757389 | 0.39146098 | 0.56270086 | RPL23A | II | 59822 | 60739 | -1 |
| YBL088C | 170.239302 | -1.1100148 | 0.57186421 | 4.600161 | 0.10025077 | 0.21944212 | TEL1 | II | 51019 | 59382 | -1 |
| YBL089W | 169.767331 | 0.12455455 | 0.68812281 | 3.34188847 | 0.1880694 | 0.34376539 | AVT5 | II | 49574 | 50953 | 1 |
| YBL090W | 21.3252378 | -0.9137396 | 0.82599641 | 2.11592003 | 0.3471633 | 0.52061007 | MRP21 | II | 48825 | 49358 | 1 |
| YBL091C | 86.2687368 | -0.8320937 | 0.59699666 | 3.74043188 | 0.15409038 | 0.2985646 | MAP2 | II | 47363 | 48628 | -1 |
| YBL091C-A | 207.301916 | -1.2034928 | 0.5108333 | 5.51527183 | 0.06344157 | 0.15629934 | SCS22 | II | 46565 | 47180 | -1 |
| YBL092W | 228.345945 | -2.9641643 | 0.59590594 | 22.0308759 | 1.64E-05 | 0.00020004 | RPL32 | II | 45978 | 46370 | 1 |
| YBL093C | 33.2344153 | 0.1333929 | 0.63503477 | 0.26435609 | 0.87618498 | 0.92712351 | ROX3 | II | 44256 | 44918 | -1 |
| YBL095W | 151.879964 | 0.91689149 | 0.55805736 | 3.88173361 | 0.14357944 | 0.28372866 | MRX3 | II | 43277 | 44089 | 1 |
| YBL097W | 27.4763209 | -0.2517011 | 0.7384052 | 0.28948797 | 0.86524378 | 0.92057359 | BRN1 | II | 40831 | 43095 | 1 |
| YBL098W | 15.5285733 | 0.80156055 | 0.92424482 | 1.02984144 | 0.59754797 | 0.74468418 | BNA4 | II | 39145 | 40527 | 1 |
| YBL099W | 55.9867717 | 0.85590214 | 0.62432375 | 1.8981586 | 0.38709726 | 0.55948406 | ATP1 | II | 37053 | 38690 | 1 |
| YBL100W-B | 14.6292161 | 1.23484352 | 0.8795254 | 3.96893687 | 0.13745366 | 0.27509737 | NA | II | 29935 | 35248 | 1 |
| YBL100W-C | 144.709954 | -0.338752 | 0.62566166 | 0.98468769 | 0.61119218 | 0.75489714 | NA | II | 28427 | 28546 | 1 |
| YBL101C | 72.7897334 | 0.44306903 | 0.61233344 | 4.60580758 | 0.09996814 | 0.21919967 | ECM21 | II | 24946 | 28299 | -1 |
| YBL102W | 14.1911084 | -2.4378104 | 0.89199048 | 7.5122462 | 0.02337418 | 0.07452452 | SFT2 | II | 24098 | 24745 | 1 |
| YBL103C | 22.2479841 | 1.33076478 | 0.75650393 | 4.56266755 | 0.10214787 | 0.22276062 | RTG3 | II | 22076 | 23536 | -1 |
| YBL104C | 31.1630661 | 1.02877137 | 0.68787949 | 3.89639134 | 0.14253101 | 0.28204189 | SEA4 | II | 18177 | 21293 | -1 |
| YBL105C | 31.6262671 | -0.0133712 | 0.71352308 | 2.35663212 | 0.30779661 | 0.48023239 | PKC1 | II | 14241 | 17696 | -1 |
| YBL106C | 1399.00522 | 1.71447745 | 0.47109353 | 12.7329993 | 0.00171816 | 0.00999815 | SRO77 | II | 10847 | 13879 | -1 |
| YBL107C | 18.6855308 | 0.17348357 | 0.75437987 | 0.39033667 | 0.82269616 | 0.89359303 | MIX23 | II | 9961 | 10551 | -1 |
| YBL111C | 14.091861 | -1.3547913 | 0.8083378 | 4.7218338 | 0.09433369 | 0.20983176 | NA | II | 2907 | 5009 | -1 |
| YBL113C | 30.8960372 | -2.0181334 | 0.65591439 | 12.133227 | 0.00231901 | 0.01246712 | NA | II | 280 | 2658 | -1 |
| YBR001C | 199.054503 | 0.64572333 | 0.49053461 | 1.71754101 | 0.42368268 | 0.59440008 | NTH2 | II | 238941 | 241283 | -1 |
| YBR002C | 129.37105 | 2.95437256 | 0.58653751 | 26.1409825 | 2.11E-06 | 3.36E-05 | RER2 | II | 241708 | 242568 | -1 |
| YBR003W | 13.5564005 | -0.0915764 | 0.92088914 | 1.80998806 | 0.4045443 | 0.57550651 | COQ1 | II | 242809 | 244230 | 1 |
| YBR004C | 48.6459951 | -0.2554511 | 0.75722344 | 4.72930219 | 0.09398209 | 0.2093602 | GPI18 | II | 244366 | 245667 | -1 |

| | | | | | | | | | | | |
|---|---|---|---|---|---|---|---|---|---|---|---|
| YBR005W | 69.3974136 | -1.2573297 | 0.54942282 | 7.28507241 | 0.02618585 | 0.08039027 | RCR1 | II | 245906 | 246547 | 1 |
| YBR006W | 197.431846 | -2.8805118 | 0.52933981 | 26.3373187 | 1.91E-06 | 3.15E-05 | UGA2 | II | 247010 | 248503 | 1 |
| YBR007C | 86.0059986 | -1.0302357 | 0.54673576 | 5.45980471 | 0.06522566 | 0.15948335 | DSF2 | II | 248806 | 251016 | -1 |
| YBR008C | 146.164609 | 0.70080562 | 0.59901193 | 1.67012904 | 0.43384649 | 0.60224674 | FLR1 | II | 252563 | 254209 | -1 |
| YBR009C | 118.531894 | -3.9868063 | 0.86267305 | 21.4571133 | 2.19E-05 | 0.0002568 | HHF2 | II | 255373 | 255684 | -1 |
| YBR010W | 97.8327738 | -2.6359798 | 0.60480211 | 22.0359264 | 1.64E-05 | 0.00019996 | HHT2 | II | 256331 | 256741 | 1 |
| YBR011C | 104.608298 | -0.8929075 | 0.63195334 | 2.66536803 | 0.26376835 | 0.43572423 | IPP1 | II | 257112 | 257975 | -1 |
| YBR014C | 13.4768211 | -1.2024262 | 0.83726956 | 2.43468659 | 0.29601555 | 0.46995614 | GRX7 | II | 266725 | 267336 | -1 |
| YBR015C | 28.9018869 | -1.2670355 | 0.70428961 | 4.59256369 | 0.10063232 | 0.21978918 | MNN2 | II | 267710 | 269503 | -1 |
| YBR016W | 120.602703 | -0.0076911 | 0.5708631 | 11.9442554 | 0.00254881 | 0.01344046 | NA | II | 270247 | 270633 | 1 |
| YBR017C | 66.2364598 | -0.4227725 | 0.62250929 | 4.50891672 | 0.10493036 | 0.22691719 | KAP104 | II | 270947 | 273703 | -1 |
| YBR019C | 1096.21125 | 0.05811228 | 0.56474581 | 0.01400937 | 0.99301979 | 0.99456575 | GAL10 | II | 276253 | 278352 | -1 |
| YBR021W | 124.651317 | 3.23423749 | 0.59188021 | 32.2013738 | 1.02E-07 | 2.48E-06 | FUR4 | II | 281443 | 283344 | 1 |
| YBR022W | 231.333759 | -0.1708925 | 0.58752188 | 6.24140653 | 0.04412613 | 0.11933221 | POA1 | II | 283738 | 284271 | 1 |
| YBR023C | 96.8530186 | 0.38776743 | 0.59369438 | 6.36520995 | 0.04147747 | 0.11428804 | CHS3 | II | 284428 | 287925 | -1 |
| YBR024W | 16.756137 | -1.697553 | 0.77307402 | 4.77563996 | 0.09182966 | 0.20608283 | SCO2 | II | 289445 | 290350 | 1 |
| YBR025C | 116.807621 | -0.5310771 | 0.57427853 | 0.93796399 | 0.62563885 | 0.76430043 | OLA1 | II | 290681 | 291865 | -1 |
| YBR026C | 20.2491715 | 0.66344179 | 0.79703188 | 0.76804189 | 0.68111716 | 0.80384598 | ETR1 | II | 292877 | 294019 | -1 |
| YBR028C | 22.6467773 | -0.6326354 | 0.68748292 | 1.83877443 | 0.39876332 | 0.56966189 | YPK3 | II | 294425 | 296002 | -1 |
| YBR029C | 28.5505022 | -1.1896845 | 0.6626652 | 3.53928631 | 0.17039378 | 0.32063048 | CDS1 | II | 296369 | 297742 | -1 |
| YBR030W | 42.0143239 | -1.8494546 | 0.63240864 | 9.78143041 | 0.00751605 | 0.03164938 | RKM3 | II | 298292 | 299950 | 1 |
| YBR031W | 53.6380664 | -2.0189779 | 0.72026534 | 7.88036144 | 0.0194447 | 0.06477837 | RPL4A | II | 300166 | 301254 | 1 |
| YBR032W | 298.776708 | 0.6556942 | 0.55899941 | 1.37841317 | 0.50197418 | 0.66070255 | NA | II | 301519 | 301821 | 1 |
| YBR033W | 1841.15713 | -0.0641037 | 0.49700313 | 1.92887702 | 0.38119718 | 0.55483451 | EDS1 | II | 301944 | 304703 | 1 |
| YBR034C | 29.3050338 | 0.71961222 | 0.6714735 | 1.15630228 | 0.5609345 | 0.7122392 | HMT1 | II | 304930 | 305976 | -1 |
| YBR035C | 94.6979886 | -0.2920806 | 0.57517622 | 0.99138978 | 0.60914747 | 0.75306988 | PDX3 | II | 306269 | 306955 | -1 |
| YBR036C | 30.0687325 | 0.8574857 | 0.70869837 | 2.16639785 | 0.33851092 | 0.51174366 | CSG2 | II | 309081 | 310313 | -1 |
| YBR038W | 36.0318055 | -0.6354386 | 0.66922598 | 0.89362721 | 0.63966313 | 0.77534214 | CHS2 | II | 311898 | 314789 | 1 |
| YBR039W | 18.382251 | -1.2476905 | 0.88322546 | 2.05398635 | 0.35808203 | 0.53120548 | ATP3 | II | 315575 | 316510 | 1 |
| YBR041W | 83.6906172 | 2.25675206 | 0.56533037 | 15.6761362 | 0.00039443 | 0.00302885 | FAT1 | II | 318266 | 320275 | 1 |
| YBR042C | 240.702337 | 2.83007551 | 0.7655415 | 12.4347806 | 0.00199444 | 0.01112927 | CST26 | II | 320416 | 321609 | -1 |
| YBR043C | 60.1466405 | 0.94473227 | 0.57132733 | 5.46987457 | 0.06489808 | 0.15888366 | QDR3 | II | 321876 | 323945 | -1 |
| YBR044C | 49.1064942 | 0.37345907 | 0.59265942 | 1.42840495 | 0.48958241 | 0.65106127 | TCM62 | II | 324341 | 326059 | -1 |

| | | | | | | | | | | | |
|---|---|---|---|---|---|---|---|---|---|---|---|
| YBR045C | 664.991476 | 3.93042412 | 0.48241858 | 55.8194913 | 7.57E-13 | 5.41E-11 | GIP1 | II | 328173 | 330092 | -1 |
| YBR046C | 197.333702 | 3.85071328 | 0.58951404 | 38.1700814 | 5.15E-09 | 1.64E-07 | ZTA1 | II | 330507 | 331511 | -1 |
| YBR048W | 146.668327 | -3.1170967 | 0.69668113 | 18.0211897 | 0.00012211 | 0.00110991 | RPS11B | II | 332831 | 333812 | 1 |
| YBR049C | 115.166347 | 1.1653751 | 0.57017113 | 10.8300046 | 0.00444933 | 0.02079165 | REB1 | II | 334386 | 336818 | -1 |
| YBR050C | 80.620294 | 2.50087629 | 0.56916435 | 24.9539852 | 3.81E-06 | 5.66E-05 | REG2 | II | 337183 | 338199 | -1 |
| YBR052C | 13.7935772 | 0.14176316 | 0.82339315 | 0.12415145 | 0.93981171 | 0.96037943 | RFS1 | II | 338720 | 339352 | -1 |
| YBR053C | 80.3652142 | 2.43675082 | 0.62309947 | 17.1158279 | 0.00019202 | 0.00164466 | NA | II | 339675 | 340751 | -1 |
| YBR054W | 73.2061936 | -0.691138 | 0.60785405 | 4.01230815 | 0.13450498 | 0.27098851 | YRO2 | II | 343101 | 344135 | 1 |
| YBR055C | 77.4420046 | 0.84616709 | 1.01237531 | 3.48445728 | 0.17512966 | 0.32688612 | PRP6 | II | 344602 | 347301 | -1 |
| YBR056W | 65.3642353 | 1.64469713 | 0.61040544 | 12.7116323 | 0.00173662 | 0.01007516 | NA | II | 347879 | 349384 | 1 |
| YBR056W-A | 82.332978 | 2.42633242 | 0.58086913 | 19.8164904 | 4.98E-05 | 0.00051177 | NA | II | 351255 | 351455 | 1 |
| YBR057C | 132.032365 | 3.18180183 | 0.53536595 | 36.3115969 | 1.30E-08 | 3.77E-07 | MUM2 | II | 352193 | 353293 | -1 |
| YBR058C | 28.4624625 | 0.18167265 | 0.74741372 | 0.0761986 | 0.96261735 | 0.97258112 | UBP14 | II | 353672 | 356017 | -1 |
| YBR058C-A | 145.863237 | -0.3659777 | 0.49260601 | 0.78822588 | 0.6742779 | 0.79886823 | TSC3 | II | 356324 | 356566 | -1 |
| YBR059C | 117.715 | 0.03299143 | 0.53320797 | 0.28396682 | 0.86763565 | 0.92227105 | AKL1 | II | 356861 | 360187 | -1 |
| YBR060C | 53.1128642 | -0.801903 | 0.57721053 | 2.58611939 | 0.27442983 | 0.44796975 | ORC2 | II | 360652 | 362514 | -1 |
| YBR061C | 20.8055825 | 1.11621886 | 0.74761794 | 3.22679308 | 0.19920984 | 0.35798416 | TRM7 | II | 364787 | 365719 | -1 |
| YBR062C | 85.6958833 | -1.9930164 | 0.5230724 | 16.0155032 | 0.00033287 | 0.00266206 | NA | II | 365976 | 366600 | -1 |
| YBR063C | 27.4086118 | 0.6799913 | 0.7429286 | 1.26810472 | 0.53043791 | 0.68477938 | NA | II | 366970 | 368184 | -1 |
| YBR065C | 24.1402283 | 1.24308172 | 0.70606747 | 7.01824775 | 0.02992312 | 0.08884865 | ECM2 | II | 368584 | 369678 | -1 |
| YBR066C | 273.271372 | -0.4514174 | 0.63250827 | 0.66597171 | 0.71678033 | 0.82837488 | NRG2 | II | 370037 | 370699 | -1 |
| YBR067C | 1218.53451 | 1.81713188 | 0.58707892 | 13.5142455 | 0.00116257 | 0.00745435 | TIP1 | II | 372103 | 372735 | -1 |
| YBR068C | 147.634629 | -0.6530138 | 0.5069421 | 1.68482801 | 0.43066963 | 0.59948398 | BAP2 | II | 373861 | 375690 | -1 |
| YBR069C | 544.691312 | -1.3765633 | 0.46917665 | 8.5079435 | 0.01420769 | 0.0513499 | TAT1 | II | 376574 | 378433 | -1 |
| YBR070C | 34.4064612 | -0.8368611 | 0.64036165 | 1.68990479 | 0.42957781 | 0.59890573 | ALG14 | II | 379221 | 379934 | -1 |
| YBR071W | 140.944152 | 1.14435965 | 0.51861789 | 4.92392579 | 0.08526742 | 0.19575668 | NA | II | 380411 | 381046 | 1 |
| YBR072W | 14.0240427 | 2.06331046 | 0.96018109 | 7.18903876 | 0.02747388 | 0.08319759 | HSP26 | II | 382030 | 382674 | 1 |
| YBR073W | 83.0644473 | 0.23618514 | 0.65450397 | 1.55134437 | 0.46039421 | 0.62448065 | RDH54 | II | 383112 | 385988 | 1 |
| YBR074W | 61.9368023 | 1.64253424 | 0.60235204 | 11.0109332 | 0.00406449 | 0.0194813 | PFF1 | II | 386286 | 389216 | 1 |
| YBR076W | 1522.50358 | 3.05776033 | 0.51502557 | 30.5600306 | 2.31E-07 | 4.96E-06 | ECM8 | II | 390227 | 391291 | 1 |
| YBR077C | 183.986512 | 3.04366208 | 0.52582962 | 30.0013666 | 3.06E-07 | 6.34E-06 | SLM4 | II | 391805 | 392293 | -1 |
| YBR078W | 103.239379 | 0.31513293 | 0.54371467 | 2.24629531 | 0.32525439 | 0.49807536 | ECM33 | II | 393123 | 394742 | 1 |
| YBR079C | 95.3169917 | -1.8784652 | 0.57508845 | 10.8314947 | 0.00444601 | 0.02079165 | RPG1 | II | 395383 | 398277 | -1 |

| | | | | | | | | | | | |
|---|---|---|---|---|---|---|---|---|---|---|---|
| YBR080C | 54.6286171 | -0.8657812 | 0.71575329 | 1.71443666 | 0.42434082 | 0.59489911 | SEC18 | II | 398614 | 400890 | -1 |
| YBR081C | 56.8902731 | -0.4347973 | 0.62113451 | 1.89907994 | 0.38691898 | 0.55936601 | SPT7 | II | 401253 | 405251 | -1 |
| YBR082C | 68.2219611 | 0.41498607 | 0.57590534 | 1.62729442 | 0.44323853 | 0.61162622 | UBC4 | II | 406628 | 407169 | -1 |
| YBR083W | 29.875278 | 0.83963173 | 0.70675066 | 2.52368894 | 0.28313132 | 0.45678444 | TEC1 | II | 409169 | 410629 | 1 |
| YBR084C-A | 115.153033 | -2.4688797 | 0.62222896 | 15.5119297 | 0.00042818 | 0.00324208 | RPL19B | II | 414186 | 415261 | -1 |
| YBR084W | 52.8781684 | -0.4352806 | 0.7203065 | 0.71983765 | 0.69773296 | 0.81712659 | MIS1 | II | 411054 | 413981 | 1 |
| YBR085C-A | 29.4316524 | 2.7845475 | 0.81288838 | 11.7233328 | 0.0028465 | 0.01465951 | NA | II | 418907 | 419164 | -1 |
| YBR085W | 15.0200303 | -0.6161297 | 0.83429691 | 0.72071571 | 0.6974267 | 0.81693316 | AAC3 | II | 415983 | 416906 | 1 |
| YBR086C | 130.639245 | 1.58950062 | 0.54553229 | 8.19206232 | 0.01663858 | 0.05772162 | IST2 | II | 420201 | 423041 | -1 |
| YBR087W | 65.6285096 | 2.89247638 | 0.58831003 | 31.8148351 | 1.23E-07 | 2.89E-06 | RFC5 | II | 423765 | 424829 | 1 |
| YBR088C | 36.8058012 | -0.2534763 | 0.70666352 | 2.33798223 | 0.31068022 | 0.48342878 | POL30 | II | 424990 | 425766 | -1 |
| YBR089C-A | 5.36065785 | -0.097596 | 1.23192418 | 0.0939936 | 0.95409045 | 0.96767513 | NHP6B | II | 426190 | 426489 | -1 |
| YBR091C | 152.571694 | 2.64402805 | 0.59163225 | 18.4444855 | 9.88E-05 | 0.00092581 | TIM12 | II | 427155 | 427484 | -1 |
| YBR092C | 611.598059 | 1.21868683 | 0.50180791 | 6.86955012 | 0.03223266 | 0.09435142 | PHO3 | II | 427698 | 429101 | -1 |
| YBR093C | 652.988375 | -3.4195742 | 0.5011956 | 48.8788694 | 2.43E-11 | 1.30E-09 | PHO5 | II | 429548 | 430951 | -1 |
| YBR094W | 22.9595185 | 0.41022149 | 0.68657749 | 1.51396105 | 0.46908067 | 0.63162258 | PBY1 | II | 432036 | 434297 | 1 |
| YBR095C | 22.3410063 | -0.8036968 | 0.71793479 | 2.28561671 | 0.31892212 | 0.49162914 | RXT2 | II | 434406 | 435698 | -1 |
| YBR096W | 26.172393 | 1.11802663 | 0.6769263 | 2.7227142 | 0.2563127 | 0.42842105 | NA | II | 436021 | 436713 | 1 |
| YBR097W | 46.4976753 | -1.438816 | 0.62948411 | 6.42209377 | 0.04031439 | 0.11205967 | VPS15 | II | 436951 | 441315 | 1 |
| YBR098W | 816.189691 | -0.327108 | 0.50662137 | 2.16230391 | 0.33920455 | 0.51239091 | MMS4 | II | 441515 | 443590 | 1 |
| YBR101C | 72.447252 | -0.0878051 | 0.57285344 | 1.13578945 | 0.56671728 | 0.71769314 | FES1 | II | 443821 | 444693 | -1 |
| YBR102C | 18.1135966 | -0.1775475 | 0.77459216 | 0.24829947 | 0.88324758 | 0.93100373 | EXO84 | II | 445062 | 447323 | -1 |
| YBR103W | 11.8892393 | -0.8971205 | 0.91162792 | 1.02518231 | 0.59894162 | 0.74609982 | SIF2 | II | 447709 | 449316 | 1 |
| YBR104W | 28.5549749 | 0.14239594 | 0.6473838 | 0.36024349 | 0.83516853 | 0.90300565 | YMC2 | II | 449667 | 450656 | 1 |
| YBR105C | 24.5390538 | 0.05604167 | 0.73590864 | 0.38022043 | 0.826868 | 0.89638002 | VID24 | II | 450881 | 451969 | -1 |
| YBR106W | 82.2178963 | -1.4707397 | 0.58812136 | 6.38335139 | 0.04110294 | 0.11365139 | PHO88 | II | 452658 | 453224 | 1 |
| YBR107C | 1644.84988 | 0.88889673 | 0.47911449 | 5.69620517 | 0.05795418 | 0.14626659 | IML3 | II | 453793 | 454530 | -1 |
| YBR108W | 114.444274 | -0.6141051 | 0.55318901 | 1.65352841 | 0.43746254 | 0.60627631 | AIM3 | II | 454822 | 457665 | 1 |
| YBR109C | 21.6162603 | 0.06267016 | 0.94786384 | 0.0762762 | 0.96258 | 0.97258112 | CMD1 | II | 457919 | 458362 | -1 |
| YBR110W | 18.064828 | 1.02540169 | 0.80392255 | 1.68083333 | 0.43153068 | 0.6001832 | ALG1 | II | 458872 | 460221 | 1 |
| YBR111C | 35.008564 | -0.108147 | 0.62666877 | 0.11602606 | 0.94363765 | 0.96197675 | YSA1 | II | 461178 | 461873 | -1 |
| YBR111W-A | 13.233378 | 2.50295405 | 0.99402419 | 6.302141 | 0.04280628 | 0.11665747 | SUS1 | II | 462139 | 462579 | 1 |
| YBR112C | 259.797969 | 0.07891232 | 0.81425765 | 7.28367423 | 0.02620416 | 0.08040386 | CYC8 | II | 462870 | 465770 | -1 |

| | | | | | | | | | | | |
|---|---|---|---|---|---|---|---|---|---|---|---|
| YBR114W | 48.4270869 | 0.18475045 | 0.81061556 | 3.5458067 | 0.16983917 | 0.32001317 | RAD16 | II | 467248 | 469620 | 1 |
| YBR115C | 50.7848437 | -1.0324511 | 0.60460633 | 3.57788845 | 0.16713654 | 0.31645538 | LYS2 | II | 469748 | 473926 | -1 |
| YBR117C | 705.849896 | -1.3800777 | 0.48689015 | 11.0433267 | 0.00399919 | 0.01926399 | TKL2 | II | 474392 | 476437 | -1 |
| YBR118W | 147.181782 | -1.168683 | 0.93249443 | 1.5433498 | 0.46223822 | 0.62560993 | TEF1 | II | 477671 | 479047 | 1 |
| YBR119W | 34.3293989 | 0.03135421 | 0.61860675 | 0.31353884 | 0.85490116 | 0.91395769 | MUD1 | II | 479338 | 480323 | 1 |
| YBR120C | 88.6386441 | 0.79734047 | 0.8980134 | 7.50661911 | 0.02344004 | 0.07454061 | CBP6 | II | 480435 | 480923 | -1 |
| YBR121C | 66.5495562 | -2.094145 | 0.84304889 | 5.92500583 | 0.05168938 | 0.13420135 | NA | II | 481364 | 483367 | -1 |
| YBR122C | 32.8562004 | -1.8602612 | 0.6599485 | 9.28848634 | 0.00961681 | 0.03832161 | MRPL36 | II | 483970 | 484503 | -1 |
| YBR123C | 150.966778 | 4.13195615 | 0.57892896 | 59.7561633 | 1.06E-13 | 8.62E-12 | TFC1 | II | 484742 | 486691 | -1 |
| YBR125C | 36.2118893 | 0.42890701 | 0.62416417 | 2.74278353 | 0.25375355 | 0.42586465 | PTC4 | II | 487199 | 488380 | -1 |
| YBR126C | 118.72519 | 0.38436505 | 0.60522549 | 0.56665 | 0.75327493 | 0.85085093 | TPS1 | II | 488905 | 490392 | -1 |
| YBR126W-A | 206.172159 | 0.51406694 | 0.48389818 | 1.53251012 | 0.46475027 | 0.62728439 | MEO1 | II | 490850 | 491056 | 1 |
| YBR127C | 132.563716 | -0.2149288 | 0.57214613 | 0.2060421 | 0.90210798 | 0.93824416 | VMA2 | II | 491269 | 492822 | -1 |
| YBR128C | 71.8252439 | -1.3231407 | 0.55208433 | 6.85301632 | 0.03250023 | 0.09503855 | ATG14 | II | 493081 | 494115 | -1 |
| YBR130C | 411.638871 | 3.47376678 | 0.48871047 | 44.937253 | 1.75E-10 | 7.66E-09 | SHE3 | II | 495592 | 496869 | -1 |
| YBR131W | 16.7561763 | 0.80155966 | 0.77041916 | 1.15706981 | 0.56071927 | 0.71212209 | CCZ1 | II | 497163 | 499277 | 1 |
| YBR132C | 46.1863543 | -0.702723 | 0.59785427 | 1.78536444 | 0.40955576 | 0.5812078 | AGP2 | II | 499652 | 501442 | -1 |
| YBR133C | 33.8190352 | 0.10385564 | 0.619436 | 0.09986489 | 0.95129369 | 0.96614462 | HSL7 | II | 501804 | 504287 | -1 |
| YBR135W | 25.3770705 | 1.2086277 | 0.83387205 | 3.73155146 | 0.1547761 | 0.29951658 | CKS1 | II | 504854 | 505306 | 1 |
| YBR136W | 215.207331 | 1.67761009 | 0.52266689 | 12.8382891 | 0.00163005 | 0.00966238 | MEC1 | II | 505668 | 512774 | 1 |
| YBR137W | 16.0765811 | -0.4863274 | 0.80556244 | 1.21807896 | 0.54387302 | 0.69714961 | NA | II | 513044 | 513583 | 1 |
| YBR138C | 202.671335 | 3.08702866 | 0.54516286 | 31.4242992 | 1.50E-07 | 3.45E-06 | NA | II | 513762 | 515336 | -1 |
| YBR139W | 71.6788297 | 1.16538274 | 0.76280162 | 2.60379753 | 0.27201481 | 0.44540887 | NA | II | 515664 | 517190 | 1 |
| YBR140C | 301.086744 | 0.01387416 | 0.70327307 | 4.70408303 | 0.09517466 | 0.2111346 | IRA1 | II | 517350 | 526628 | -1 |
| YBR141C | 448.624387 | 0.99408459 | 0.53038786 | 4.70991657 | 0.09489747 | 0.21060035 | BMT2 | II | 527025 | 528038 | -1 |
| YBR142W | 87.434226 | -1.5946203 | 0.90711969 | 11.4256741 | 0.00330329 | 0.0164313 | MAK5 | II | 528317 | 530638 | 1 |
| YBR143C | 51.7063154 | -1.2466276 | 0.58763132 | 4.55268279 | 0.10265911 | 0.22312171 | SUP45 | II | 530869 | 532182 | -1 |
| YBR144C | 20.9465833 | -0.850261 | 0.70676524 | 1.45443568 | 0.48325161 | 0.64559917 | NA | II | 533235 | 533549 | -1 |
| YBR145W | 70.140092 | -0.1894073 | 0.64897755 | 0.10086927 | 0.95081608 | 0.96614462 | ADH5 | II | 533762 | 534817 | 1 |
| YBR146W | 29.9395107 | -1.4474783 | 0.66908756 | 4.86586832 | 0.0877789 | 0.20033103 | MRPS9 | II | 535260 | 536096 | 1 |
| YBR148W | 48.5252502 | -0.1315185 | 0.67642588 | 0.06059546 | 0.97015665 | 0.97724547 | YSW1 | II | 537876 | 539705 | 1 |
| YBR149W | 469.853298 | -2.8885097 | 0.52148915 | 45.7920648 | 1.14E-10 | 5.23E-09 | ARA1 | II | 539987 | 541021 | 1 |
| YBR150C | 444.125204 | -0.317028 | 0.89606004 | 12.777176 | 0.00168063 | 0.00981378 | TBS1 | II | 541209 | 544493 | -1 |

| | | | | | | | | | | | |
|---|---|---|---|---|---|---|---|---|---|---|---|
| YBR151W | 53.8522041 | -0.5553359 | 0.80243252 | 1.90534234 | 0.38570935 | 0.55845627 | APD1 | II | 545028 | 545978 | 1 |
| YBR152W | 27.6191505 | -0.0018561 | 0.6613115 | 0.2097891 | 0.90041947 | 0.93824416 | SPP381 | II | 546376 | 547251 | 1 |
| YBR153W | 12.7059386 | -0.7659661 | 0.82304041 | 1.75606497 | 0.41559981 | 0.58733778 | RIB7 | II | 547460 | 548194 | 1 |
| YBR154C | 141.687499 | -1.5512372 | 0.62390292 | 9.27860018 | 0.00966446 | 0.03840187 | RPB5 | II | 548362 | 549009 | -1 |
| YBR155W | 64.2249131 | 0.35577744 | 1.08008474 | 3.39194056 | 0.18342117 | 0.33887818 | CNS1 | II | 549771 | 550928 | 1 |
| YBR156C | 48.4949478 | -1.5233705 | 0.66605868 | 5.69771997 | 0.0579103 | 0.14626659 | SLI15 | II | 551104 | 553200 | -1 |
| YBR157C | 260.555643 | 0.15882958 | 0.50449029 | 0.24333189 | 0.8854441 | 0.931623 | ICS2 | II | 553543 | 554310 | -1 |
| YBR158W | 30.4405161 | -0.9532938 | 0.64653202 | 5.19704172 | 0.07438352 | 0.17629168 | AMN1 | II | 556549 | 558198 | 1 |
| YBR159W | 30.6797804 | 2.05766498 | 0.82157137 | 6.19831784 | 0.04508711 | 0.12108273 | IFA38 | II | 558685 | 559728 | 1 |
| YBR160W | 57.8786997 | 1.03473866 | 0.58193388 | 3.96930366 | 0.13742845 | 0.27509737 | CDC28 | II | 560078 | 560974 | 1 |
| YBR161W | 62.4827399 | 2.25093011 | 0.58370465 | 14.5529632 | 0.00069161 | 0.00483046 | CSH1 | II | 561634 | 562764 | 1 |
| YBR162C | 89.2506745 | -0.3463546 | 0.62697812 | 0.39218814 | 0.82193491 | 0.89337397 | TOS1 | II | 563203 | 564570 | -1 |
| YBR162W-A | 165.662514 | 0.30362962 | 0.59573665 | 5.54004469 | 0.0626606 | 0.15478025 | YSY6 | II | 565231 | 565428 | 1 |
| YBR163W | 66.2473236 | 0.12300524 | 0.59803286 | 3.21860746 | 0.20002684 | 0.35900663 | EXO5 | II | 565723 | 567480 | 1 |
| YBR164C | 10.2658955 | -0.5697004 | 0.94484141 | 0.55988069 | 0.75582883 | 0.85240532 | ARL1 | II | 567875 | 568426 | -1 |
| YBR165W | 14.719259 | 1.24143098 | 0.93649399 | 2.65144564 | 0.2656109 | 0.43764573 | UBS1 | II | 568852 | 569685 | 1 |
| YBR166C | 39.5956849 | 1.78199736 | 0.66251512 | 7.29781565 | 0.02601953 | 0.07997546 | TYR1 | II | 569842 | 571200 | -1 |
| YBR167C | 253.555078 | 5.02069025 | 0.57352794 | 64.833222 | 8.35E-15 | 8.48E-13 | POP7 | II | 571468 | 571890 | -1 |
| YBR168W | 13.026385 | 0.30142625 | 0.932633 | 0.14459337 | 0.93025486 | 0.95618243 | PEX32 | II | 572371 | 573612 | 1 |
| YBR169C | 52.6384314 | 2.82088591 | 0.67731689 | 19.439104 | 6.01E-05 | 0.0006041 | SSE2 | II | 573915 | 575996 | -1 |
| YBR170C | 14.7348989 | 1.27562149 | 0.88843807 | 2.19018908 | 0.33450798 | 0.50802806 | NPL4 | II | 576344 | 578086 | -1 |
| YBR171W | 34.8072373 | -0.7236652 | 0.63880418 | 2.42024877 | 0.29816019 | 0.47161883 | SEC66 | II | 578364 | 578984 | 1 |
| YBR172C | 151.746889 | 1.39489972 | 0.68095408 | 5.44186166 | 0.06581346 | 0.16064998 | SMY2 | II | 579150 | 581372 | -1 |
| YBR173C | 113.904355 | -1.4844837 | 0.5290704 | 8.29133589 | 0.01583286 | 0.05593181 | UMP1 | II | 581726 | 582172 | -1 |
| YBR175W | 35.8064777 | -0.331188 | 0.64230899 | 0.72627762 | 0.69548989 | 0.81611981 | SWD3 | II | 582408 | 583355 | 1 |
| YBR176W | 50.6200451 | 0.90611696 | 0.70219887 | 4.13653868 | 0.12640436 | 0.25966788 | ECM31 | II | 583720 | 584658 | 1 |
| YBR177C | 66.8611894 | 0.83615076 | 0.63967681 | 2.44414207 | 0.29461937 | 0.46889669 | EHT1 | II | 584807 | 586162 | -1 |
| YBR179C | 34.5451866 | -0.8496581 | 0.61581217 | 4.03387191 | 0.13306255 | 0.26909961 | FZO1 | II | 586547 | 589114 | -1 |
| YBR180W | 231.506979 | -1.2881036 | 0.50447515 | 7.49363034 | 0.02359277 | 0.07480948 | DTR1 | II | 589741 | 591459 | 1 |
| YBR181C | 38.048731 | -1.9544453 | 0.67388105 | 8.01524187 | 0.01817659 | 0.06161735 | RPS6A | II | 591712 | 592774 | -1 |
| YBR182C | 57.6368698 | -2.0257551 | 0.57063842 | 13.4732961 | 0.00118662 | 0.00756665 | SMP1 | II | 593506 | 594864 | -1 |
| YBR182C-A | 50.3317798 | 0.95815059 | 0.63472087 | 3.05578054 | 0.21699298 | 0.38107048 | NA | II | 595361 | 595555 | -1 |
| YBR183W | 14.2211668 | 1.4497973 | 0.81458938 | 3.27367866 | 0.19459412 | 0.35165417 | YPC1 | II | 596115 | 597065 | 1 |

| | | | | | | | | | | | |
|---|---|---|---|---|---|---|---|---|---|---|---|
| YBR184W | 52.9390505 | -0.743312 | 0.61094536 | 3.76531306 | 0.15218528 | 0.29581776 | NA | II | 597363 | 598934 | 1 |
| YBR186W | 499.546392 | -2.846458 | 0.48110747 | 32.3468233 | 9.46E-08 | 2.33E-06 | PCH2 | II | 600553 | 602360 | 1 |
| YBR187W | 256.798377 | 0.86727934 | 0.62704511 | 1.89547118 | 0.38761776 | 0.55995679 | GDT1 | II | 602634 | 603476 | 1 |
| YBR188C | 22.4129868 | -1.5703408 | 0.72943674 | 4.98154492 | 0.08284595 | 0.19095463 | NTC20 | II | 603685 | 604107 | -1 |
| YBR189W | 106.930075 | -3.0871634 | 0.92468208 | 10.2490286 | 0.00594911 | 0.02633435 | RPS9B | II | 604508 | 605508 | 1 |
| YBR191W | 65.5889934 | -2.7795553 | 0.71361681 | 14.019231 | 0.00090316 | 0.00605941 | RPL21A | II | 606270 | 607140 | 1 |
| YBR192W | 226.970427 | -1.1710935 | 0.50259442 | 5.66664818 | 0.05881702 | 0.14748831 | RIM2 | II | 607652 | 608785 | 1 |
| YBR193C | 26.1729235 | -0.1559318 | 0.69356747 | 0.09335679 | 0.95439429 | 0.96776584 | MED8 | II | 609082 | 609753 | -1 |
| YBR194W | 12.7124139 | -0.9057026 | 0.83403964 | 3.93692561 | 0.13967139 | 0.27818967 | AIM4 | II | 610038 | 610409 | 1 |
| YBR195C | 246.347838 | 2.90972548 | 0.54022955 | 36.0642756 | 1.47E-08 | 4.25E-07 | MSI1 | II | 610614 | 611882 | -1 |
| YBR196C | 114.309634 | 0.22843367 | 0.84669759 | 1.27941806 | 0.52744587 | 0.68184633 | PGI1 | II | 612236 | 613900 | -1 |
| YBR196C-B | 62.4870326 | -2.2056275 | 0.58507442 | 13.3415387 | 0.00126742 | 0.00799388 | NA | II | 614526 | 614630 | -1 |
| YBR197C | 231.466843 | 0.22859511 | 0.54643347 | 0.39084997 | 0.82248504 | 0.89359303 | NA | II | 615203 | 615856 | -1 |
| YBR198C | 207.126918 | -0.9296652 | 0.5194189 | 3.56599695 | 0.16813325 | 0.31792668 | TAF5 | II | 616127 | 618523 | -1 |
| YBR199W | 40.0836265 | 0.94969933 | 0.74459064 | 2.12704444 | 0.34523766 | 0.51852261 | KTR4 | II | 618909 | 620303 | 1 |
| YBR200W | 29.2430876 | 1.06836679 | 0.70665276 | 2.42916241 | 0.2968343 | 0.47086866 | BEM1 | II | 620872 | 622527 | 1 |
| YBR200W-A | 226.766788 | 6.87952819 | 0.72150808 | 86.0814429 | 2.03E-19 | 4.90E-17 | NA | II | 622983 | 623147 | 1 |
| YBR201C-A | 33.4425987 | -0.4178182 | 0.66505287 | 2.19255469 | 0.33411256 | 0.5079834 | NA | II | 624493 | 624696 | -1 |
| YBR201W | 6.1737205 | 0.79104881 | 1.08044369 | 1.18520531 | 0.55288644 | 0.70480239 | DER1 | II | 623576 | 624211 | 1 |
| YBR202W | 22.8064707 | -0.7065604 | 0.78346179 | 1.53320643 | 0.4645885 | 0.62728439 | MCM7 | II | 625772 | 628309 | 1 |
| YBR203W | 2248.36593 | 4.91454396 | 0.45460673 | 86.5598836 | 1.60E-19 | 4.02E-17 | COS111 | II | 629168 | 631942 | 1 |
| YBR204C | 25.5346675 | -0.2203281 | 0.67352193 | 1.55314847 | 0.45997909 | 0.62448065 | LDH1 | II | 632254 | 633381 | -1 |
| YBR205W | 30.5354967 | 0.69829734 | 0.70671571 | 2.86717615 | 0.2384518 | 0.40666743 | KTR3 | II | 633622 | 634836 | 1 |
| YBR207W | 39.996762 | 0.76992946 | 0.71748705 | 1.80420851 | 0.40571503 | 0.57703022 | FTH1 | II | 635146 | 636543 | 1 |
| YBR208C | 74.2098808 | 2.04674363 | 0.54912883 | 15.8535928 | 0.00036094 | 0.00284348 | DUR1,2 | II | 636703 | 642210 | -1 |
| YBR209W | 350.089805 | 1.18650133 | 0.53359694 | 4.8251028 | 0.08958643 | 0.20301583 | NA | II | 642583 | 642900 | 1 |
| YBR210W | 9.39071885 | -1.2056513 | 0.95728996 | 1.95484544 | 0.37627963 | 0.54998972 | ERV15 | II | 645550 | 645978 | 1 |
| YBR211C | 39.6753003 | 0.0481797 | 0.86762144 | 9.78757365 | 0.00749299 | 0.03162131 | AME1 | II | 646158 | 647132 | -1 |
| YBR212W | 118.826427 | 3.29830902 | 0.57383735 | 31.3826065 | 1.53E-07 | 3.51E-06 | NGR1 | II | 647886 | 649904 | 1 |
| YBR213W | 16.338113 | 1.62271696 | 0.87760266 | 3.94428523 | 0.13915837 | 0.27774112 | MET8 | II | 650368 | 651192 | 1 |
| YBR214W | 345.850628 | -1.5802187 | 0.52244522 | 13.9170539 | 0.0009505 | 0.00631841 | SDS24 | II | 651415 | 652998 | 1 |
| YBR215W | 437.324909 | -0.3134542 | 0.51455492 | 0.7253552 | 0.69581073 | 0.81619613 | HPC2 | II | 653356 | 655317 | 1 |
| YBR216C | 53.5332662 | -1.142309 | 0.63819277 | 5.49594147 | 0.06405772 | 0.15736105 | YBP1 | II | 655576 | 657600 | -1 |

| | | | | | | | | | | | |
|---|---|---|---|---|---|---|---|---|---|---|---|
| YBR217W | 12.8999714 | -0.1191513 | 0.84608304 | 2.49098003 | 0.28779985 | 0.46172289 | ATG12 | II | 657832 | 658392 | 1 |
| YBR218C | 93.8029793 | 1.14557897 | 0.51601027 | 6.10214652 | 0.04730812 | 0.12530377 | PYC2 | II | 658707 | 662249 | -1 |
| YBR220C | 44.5498562 | 0.88850921 | 0.63545983 | 4.99084963 | 0.08246141 | 0.19029557 | NA | II | 662995 | 664677 | -1 |
| YBR221C | 44.7315432 | 0.80141202 | 0.75340839 | 1.21744296 | 0.544046 | 0.69721698 | PDB1 | II | 665153 | 666253 | -1 |
| YBR221W-A | 1056.3207 | 2.67529076 | 0.70395614 | 13.9787369 | 0.00092163 | 0.00615027 | NA | II | 666538 | 666642 | 1 |
| YBR222C | 37.7833155 | 1.92414408 | 0.67716254 | 8.845311 | 0.01200232 | 0.04521368 | PCS60 | II | 666720 | 668351 | -1 |
| YBR223C | 8.79214549 | 1.16035113 | 1.00092411 | 1.37634122 | 0.50249449 | 0.66093664 | TDP1 | II | 668663 | 670297 | -1 |
| YBR225W | 24.6975037 | 0.69359291 | 0.71711582 | 1.146388 | 0.56372203 | 0.71483805 | NA | II | 670627 | 673329 | 1 |
| YBR227C | 213.57076 | 2.10151557 | 0.73208322 | 9.8661603 | 0.00720428 | 0.03068439 | MCX1 | II | 673572 | 675134 | -1 |
| YBR228W | 309.228892 | -1.4114231 | 0.68045789 | 5.50043539 | 0.06391395 | 0.15715409 | SLX1 | II | 675313 | 676227 | 1 |
| YBR229C | 115.033191 | -1.4826164 | 0.5336163 | 7.56352459 | 0.02278251 | 0.07300493 | ROT2 | II | 676357 | 679221 | -1 |
| YBR230C | 13.8569409 | 1.05010355 | 0.84897437 | 1.5918373 | 0.45116658 | 0.61755425 | OM14 | II | 679549 | 680050 | -1 |
| YBR230W-A | 42.025878 | -0.3232807 | 0.60978343 | 2.70078479 | 0.25913856 | 0.43082351 | NA | II | 680362 | 680562 | 1 |
| YBR231C | 19.1513397 | 0.86975087 | 0.79970361 | 1.23632984 | 0.53893252 | 0.69327244 | SWC5 | II | 682179 | 683090 | -1 |
| YBR233W | 27.4891304 | 0.00336398 | 0.70949119 | 1.8346541 | 0.39958569 | 0.57041448 | PBP2 | II | 683428 | 684669 | 1 |
| YBR234C | 88.6770024 | -1.5217886 | 0.5355294 | 7.89599379 | 0.01929331 | 0.0644223 | ARC40 | II | 685438 | 686592 | -1 |
| YBR235W | 287.738137 | -1.9359491 | 0.50775895 | 13.6383512 | 0.00109262 | 0.00709251 | VHC1 | II | 686901 | 690263 | 1 |
| YBR236C | 58.3708431 | -1.0551903 | 0.55463767 | 3.62240854 | 0.16345717 | 0.3120399 | ABD1 | II | 690383 | 691693 | -1 |
| YBR237W | 20.9281593 | 0.55391336 | 0.73320006 | 1.16181476 | 0.55939056 | 0.71059046 | PRP5 | II | 691969 | 694518 | 1 |
| YBR238C | 25.4665888 | -0.6341812 | 0.84781878 | 4.608339 | 0.09984168 | 0.21918634 | NA | II | 695107 | 697302 | -1 |
| YBR239C | 310.725895 | 0.1321956 | 0.53302638 | 1.4475218 | 0.48492507 | 0.64753601 | ERT1 | II | 698354 | 699943 | -1 |
| YBR240C | 12.730542 | -0.3870227 | 0.89922972 | 0.71768187 | 0.69848545 | 0.81736153 | THI2 | II | 700490 | 701842 | -1 |
| YBR241C | 247.186762 | 2.04250232 | 0.5780952 | 12.5517049 | 0.00188119 | 0.01071 | NA | II | 702589 | 704055 | -1 |
| YBR242W | 316.154853 | -0.5563833 | 0.58938522 | 5.63480427 | 0.05976099 | 0.14953161 | NA | II | 704670 | 705386 | 1 |
| YBR243C | 292.114403 | -0.7118037 | 0.63283523 | 1.62534504 | 0.44367076 | 0.61162622 | ALG7 | II | 705447 | 706793 | -1 |
| YBR244W | 31.8732958 | -1.8198479 | 0.85297614 | 13.6065575 | 0.00111013 | 0.00717372 | GPX2 | II | 707528 | 708016 | 1 |
| YBR245C | 58.233506 | -0.5275938 | 0.55441683 | 1.56823431 | 0.45652256 | 0.62207609 | ISW1 | II | 708150 | 711539 | -1 |
| YBR246W | 26.1981731 | -0.7170022 | 0.97293502 | 7.9062456 | 0.01919467 | 0.06416508 | RRT2 | II | 711591 | 712754 | 1 |
| YBR247C | 28.5119343 | -1.7283057 | 0.69749312 | 6.71432491 | 0.03483396 | 0.10013768 | ENP1 | II | 713004 | 714455 | -1 |
| YBR248C | 107.929456 | -2.2543371 | 0.52981696 | 18.3757855 | 0.00010227 | 0.00095353 | HIS7 | II | 714807 | 716465 | -1 |
| YBR249C | 39.1786025 | -0.9801888 | 0.68374109 | 4.3654089 | 0.11273623 | 0.23918753 | ARO4 | II | 716882 | 717994 | -1 |
| YBR251W | 12.8335635 | -1.0959372 | 0.82621926 | 1.79258047 | 0.40808074 | 0.5793986 | MRPS5 | II | 721390 | 722313 | 1 |
| YBR252W | 86.1719078 | 0.10310285 | 0.52260245 | 0.85720469 | 0.65141892 | 0.78236902 | DUT1 | II | 722611 | 723054 | 1 |

| | | | | | | | | | | | |
|---|---|---|---|---|---|---|---|---|---|---|---|
| YBR253W | 87.992407 | -0.4606799 | 0.88644164 | 7.34570743 | 0.02540387 | 0.07865691 | SRB6 | II | 723270 | 723635 | 1 |
| YBR254C | 9.19788425 | -1.5751113 | 1.00478074 | 3.01971313 | 0.22094167 | 0.38583539 | TRS20 | II | 723736 | 724263 | -1 |
| YBR255C-A | 239.809594 | 0.64482238 | 0.6445313 | 1.88011389 | 0.39060559 | 0.56175022 | RCF3 | II | 726618 | 727074 | -1 |
| YBR255W | 179.614665 | 0.5637791 | 0.66694364 | 8.37459505 | 0.01518727 | 0.05409075 | MTC4 | II | 724456 | 726540 | 1 |
| YBR256C | 33.4632869 | -0.2836988 | 0.69454805 | 3.20048553 | 0.20184751 | 0.36126649 | RIB5 | II | 727386 | 728102 | -1 |
| YBR257W | 240.368904 | -0.5891979 | 0.52882006 | 3.50396209 | 0.17343003 | 0.32507061 | POP4 | II | 728885 | 729724 | 1 |
| YBR258C | 126.136025 | -0.6758089 | 0.57924762 | 2.01682796 | 0.3647971 | 0.53799674 | SHG1 | II | 729734 | 730162 | -1 |
| YBR259W | 226.250356 | 4.35737286 | 0.63520167 | 40.4416519 | 1.65E-09 | 5.87E-08 | NA | II | 730387 | 732453 | 1 |
| YBR260C | 59.2857834 | -0.4397718 | 0.69712253 | 0.41317471 | 0.81335521 | 0.88771473 | RGD1 | II | 732639 | 734639 | -1 |
| YBR261C | 48.4329879 | -0.2785275 | 0.57517831 | 0.52061398 | 0.77081492 | 0.86042382 | TAE1 | II | 734832 | 735530 | -1 |
| YBR262C | 8.11284983 | -0.7575715 | 1.06586917 | 0.74452498 | 0.68917332 | 0.81068229 | MIC12 | II | 735720 | 736040 | -1 |
| YBR263W | 208.976017 | -2.0913906 | 0.54997262 | 13.744307 | 0.00103624 | 0.00679484 | SHM1 | II | 736264 | 737736 | 1 |
| YBR264C | 1214.10155 | -0.3750137 | 0.8295787 | 6.26259671 | 0.04366107 | 0.11846186 | YPT10 | II | 737770 | 738369 | -1 |
| YBR265W | 58.9057551 | -1.3002175 | 0.59125213 | 10.2058203 | 0.00607903 | 0.02677727 | TSC10 | II | 738582 | 739544 | 1 |
| YBR266C | 3446.34823 | -2.4790351 | 0.52905241 | 20.6617721 | 3.26E-05 | 0.00036033 | NA | II | 739936 | 740388 | -1 |
| YBR267W | 36.9066703 | -1.5509415 | 0.68426971 | 5.72340266 | 0.05717141 | 0.14493103 | REI1 | II | 739841 | 741022 | 1 |
| YBR269C | 31.6057487 | 1.42040645 | 0.77972119 | 3.60312103 | 0.16504114 | 0.31392516 | SDH8 | II | 742160 | 742576 | -1 |
| YBR270C | 78.6176637 | 2.8605949 | 0.60457665 | 22.6892425 | 1.18E-05 | 0.00015327 | BIT2 | II | 742761 | 744398 | -1 |
| YBR271W | 19.2799028 | -0.5560657 | 0.71042703 | 0.73837259 | 0.69129661 | 0.81221741 | EFM2 | II | 744852 | 746111 | 1 |
| YBR272C | 13.8463636 | 0.23515034 | 0.79026719 | 0.26947329 | 0.87394604 | 0.92574965 | HSM3 | II | 746361 | 747803 | -1 |
| YBR273C | 68.696661 | 1.95489711 | 0.58359606 | 14.3504103 | 0.00076533 | 0.00525653 | UBX7 | II | 748061 | 749371 | -1 |
| YBR274W | 16.757086 | -0.5202485 | 0.770053 | 0.50484125 | 0.77691788 | 0.86407117 | CHK1 | II | 749594 | 751177 | 1 |
| YBR275C | 190.420299 | -1.7913098 | 0.51831364 | 11.8970265 | 0.00260972 | 0.01368683 | RIF1 | II | 751356 | 757106 | -1 |
| YBR276C | 18.6933534 | -0.6619133 | 0.77891586 | 1.53621046 | 0.4638912 | 0.62711419 | PPS1 | II | 757620 | 760043 | -1 |
| YBR278W | 27.6081642 | -1.0680456 | 0.67662899 | 3.38904599 | 0.18368683 | 0.33924936 | DPB3 | II | 760294 | 760899 | 1 |
| YBR279W | 118.357209 | 0.49809777 | 0.65514857 | 7.42009695 | 0.02447634 | 0.07677031 | PAF1 | II | 761257 | 762594 | 1 |
| YBR280C | 91.3723663 | -1.0550109 | 0.55337394 | 3.55630353 | 0.16895012 | 0.31887463 | SAF1 | II | 762784 | 764697 | -1 |
| YBR281C | 17.4474364 | -1.4283086 | 0.75664031 | 4.18786161 | 0.1232019 | 0.25458208 | DUG2 | II | 764970 | 767606 | -1 |
| YBR282W | 12.2737049 | -0.1884874 | 0.89862851 | 0.38386688 | 0.8253618 | 0.89541781 | MRPL27 | II | 768240 | 768680 | 1 |
| YBR283C | 7110.7187 | 6.78555977 | 0.5624324 | 90.4952702 | 2.23E-20 | 6.81E-18 | SSH1 | II | 768943 | 770415 | -1 |
| YBR284W | 22.2544745 | 2.99984225 | 0.85721625 | 12.8058449 | 0.00165671 | 0.00977813 | NA | II | 771239 | 773632 | 1 |
| YBR285W | 50.2825246 | -0.3102432 | 0.56052061 | 1.48370509 | 0.47623086 | 0.6387252 | NA | II | 773922 | 774356 | 1 |
| YBR286W | 144.470741 | 0.28629883 | 0.55448362 | 1.61853675 | 0.44518365 | 0.61297083 | APE3 | II | 774700 | 776313 | 1 |

| | | | | | | | | | | | |
|---|---|---|---|---|---|---|---|---|---|---|---|
| YBR287W | 126.330136 | -0.0047437 | 0.54020864 | 0.00492577 | 0.99754014 | 0.99771246 | NA | II | 776571 | 777854 | 1 |
| YBR288C | 2038.12009 | 7.33414616 | 0.50410851 | 142.746095 | 1.01E-31 | 5.83E-28 | APM3 | II | 778012 | 779463 | -1 |
| YBR289W | 55.4184523 | 1.50903777 | 0.83604159 | 6.55794807 | 0.03766688 | 0.1063341 | SNF5 | II | 779667 | 782384 | 1 |
| YBR290W | 86.8124611 | 0.869714 | 0.6915851 | 4.31489224 | 0.11562002 | 0.242911 | BSD2 | II | 782591 | 783556 | 1 |
| YBR291C | 380.561205 | 1.61641705 | 0.62730914 | 10.9026714 | 0.00429057 | 0.02021351 | CTP1 | II | 783673 | 784572 | -1 |
| YBR292C | 57.6266749 | 2.04628708 | 0.7405481 | 7.24351027 | 0.02673571 | 0.08151646 | NA | II | 784702 | 785073 | -1 |
| YBR293W | 17.4933375 | 0.4279155 | 0.75507894 | 1.02037014 | 0.60038446 | 0.74682758 | VBA2 | II | 787006 | 788430 | 1 |
| YBR294W | 5455.36514 | 1.5324949 | 0.49948247 | 9.22697004 | 0.0099172 | 0.03919493 | SUL1 | II | 789235 | 791814 | 1 |
| YBR295W | 42.1290992 | -0.0271375 | 0.72473675 | 0.32388373 | 0.85049065 | 0.91140863 | PCA1 | II | 792849 | 796499 | 1 |
| YBR296C | 57.7098009 | 0.87084885 | 0.61877309 | 2.79186475 | 0.24760207 | 0.41832973 | PHO89 | II | 796798 | 798522 | -1 |
| YBR296C-A | 10.4001111 | -2.1105382 | 0.91511827 | 5.83686869 | 0.0540182 | 0.13854305 | NA | II | 800123 | 800242 | -1 |
| YBR297W | 114.478453 | 4.09626825 | 0.60926502 | 58.3610555 | 2.12E-13 | 1.66E-11 | MAL33 | II | 800523 | 801929 | 1 |
| YBR298C | 21.3626856 | 2.00263737 | 0.8314384 | 6.8413479 | 0.0326904 | 0.09540191 | MAL31 | II | 802631 | 804475 | -1 |
| YBR299W | 55.577594 | -1.9878604 | 0.66071921 | 14.0584599 | 0.00088561 | 0.00595552 | MAL32 | II | 805351 | 807105 | 1 |
| YBR302C | 8.53829875 | -2.1050686 | 1.09969787 | 3.75297392 | 0.1531271 | 0.29709307 | COS3 | II | 810340 | 811479 | -1 |
| YCL001W | 15.8275698 | -0.6287386 | 0.7828806 | 0.65967013 | 0.71904232 | 0.82938751 | RER1 | III | 111916 | 112482 | 1 |
| YCL001W-A | 106.218471 | -0.0353032 | 0.5186754 | 0.73062531 | 0.69397964 | 0.81470846 | NA | III | 113080 | 113541 | 1 |
| YCL001W-B | 168.048844 | 0.28183684 | 0.54973188 | 0.28441957 | 0.86743926 | 0.9222316 | NA | III | 113770 | 114024 | 1 |
| YCL002C | 601.128917 | -0.0038524 | 0.50797602 | 0.38399118 | 0.82531051 | 0.89541781 | NA | III | 110808 | 111675 | -1 |
| YCL004W | 341.831723 | -1.2547821 | 0.55390051 | 5.78563517 | 0.05541984 | 0.14137538 | PGS1 | III | 109105 | 110670 | 1 |
| YCL005W | 82.7667111 | 0.49171242 | 0.6187047 | 1.8475881 | 0.3970099 | 0.56785755 | LDB16 | III | 108021 | 108791 | 1 |
| YCL005W-A | 50.6195267 | 1.00450349 | 0.76233271 | 2.85276169 | 0.24017659 | 0.40828609 | VMA9 | III | 107023 | 107417 | 1 |
| YCL008C | 23.6180039 | -0.4135681 | 0.69720515 | 0.39072543 | 0.82253626 | 0.89359303 | STP22 | III | 105696 | 106853 | -1 |
| YCL009C | 247.105573 | -0.8667108 | 0.5213305 | 3.81600214 | 0.14837668 | 0.29072792 | ILV6 | III | 104619 | 105548 | -1 |
| YCL010C | 148.415847 | -2.4740725 | 0.56261448 | 21.1271333 | 2.58E-05 | 0.00029686 | SGF29 | III | 103571 | 104350 | -1 |
| YCL011C | 59.4034616 | -0.9808565 | 0.62842141 | 2.55271429 | 0.279052 | 0.45346929 | GBP2 | III | 102075 | 103358 | -1 |
| YCL012C | 62.5856088 | 0.97646853 | 0.74923881 | 7.35889409 | 0.02523693 | 0.07826556 | NA | III | 101317 | 101788 | -1 |
| YCL014W | 192.828109 | 0.58875604 | 0.59266448 | 1.94380222 | 0.37836304 | 0.55223646 | BUD3 | III | 96281 | 101191 | 1 |
| YCL016C | 10.5722747 | 0.43620871 | 0.90763399 | 0.25606986 | 0.87982265 | 0.92911891 | DCC1 | III | 94621 | 95763 | -1 |
| YCL017C | 253.865876 | -1.6224051 | 0.62136404 | 7.3695392 | 0.02510296 | 0.07810109 | NFS1 | III | 92777 | 94270 | -1 |
| YCL018W | 219.963722 | -1.7978013 | 0.50621553 | 12.2830307 | 0.00215166 | 0.01180864 | LEU2 | III | 91324 | 92418 | 1 |
| YCL021W-A | 22.0388119 | 1.88613306 | 0.77595209 | 7.99185174 | 0.01839041 | 0.06210569 | NA | III | 83620 | 83997 | 1 |
| YCL024W | 44.3843966 | 0.35864338 | 0.59070741 | 0.5023339 | 0.77789249 | 0.86498896 | KCC4 | III | 79162 | 82275 | 1 |

| | | | | | | | | | | | |
|---|---|---|---|---|---|---|---|---|---|---|---|
| YCL025C | 299.516182 | 1.80528436 | 0.49964164 | 13.5578054 | 0.00113752 | 0.00731806 | AGP1 | III | 76018 | 77919 | -1 |
| YCL026C-A | 27.3285305 | -2.5676512 | 0.72379564 | 17.8665274 | 0.00013193 | 0.00118796 | FRM2 | III | 74704 | 75285 | -1 |
| YCL026C-B | 11.3282441 | -1.9429748 | 1.00681397 | 3.92723822 | 0.14034956 | 0.27886889 | HBN1 | III | 73405 | 73986 | -1 |
| YCL027W | 77.7021553 | 0.84895208 | 0.56224113 | 9.92503456 | 0.0069953 | 0.02997984 | FUS1 | III | 71803 | 73341 | 1 |
| YCL028W | 126.493519 | 0.0145428 | 0.63359873 | 2.93032744 | 0.23104016 | 0.39792139 | RNQ1 | III | 70150 | 71367 | 1 |
| YCL029C | 26.6042279 | 0.28117791 | 0.72722326 | 0.9412615 | 0.62460817 | 0.76345394 | BIK1 | III | 68599 | 69921 | -1 |
| YCL030C | 63.7154504 | 2.11448732 | 0.61010213 | 11.5910938 | 0.00304107 | 0.01538156 | HIS4 | III | 65934 | 68333 | -1 |
| YCL031C | 47.2983454 | -2.1260472 | 0.66939264 | 11.6534005 | 0.00294779 | 0.01503761 | RRP7 | III | 64675 | 65568 | -1 |
| YCL032W | 18.9617726 | -0.1029092 | 0.76791869 | 1.83051739 | 0.40041303 | 0.57131381 | STE50 | III | 63441 | 64481 | 1 |
| YCL033C | 107.919393 | 1.11256836 | 0.78399672 | 10.6337186 | 0.00490814 | 0.02256764 | MXR2 | III | 62776 | 63282 | -1 |
| YCL034W | 304.665661 | 1.55221814 | 0.81647317 | 9.21364627 | 0.00998348 | 0.03943 | LSB5 | III | 61658 | 62722 | 1 |
| YCL035C | 60.927361 | -0.1410534 | 0.5529608 | 0.35715833 | 0.83645784 | 0.90356173 | GRX1 | III | 60841 | 61173 | -1 |
| YCL036W | 150.413354 | 0.71978407 | 0.82415185 | 6.77059602 | 0.03386755 | 0.09820123 | GFD2 | III | 59026 | 60726 | 1 |
| YCL037C | 236.282174 | 3.30752686 | 0.58415252 | 29.1538847 | 4.67E-07 | 9.10E-06 | SRO9 | III | 57374 | 58678 | -1 |
| YCL038C | 162.787189 | -0.1911628 | 0.5514313 | 0.23491922 | 0.88917642 | 0.93304075 | ATG22 | III | 54941 | 56527 | -1 |
| YCL039W | 284.465486 | -0.7577777 | 0.77395534 | 1.71708384 | 0.42377954 | 0.59440008 | GID7 | III | 52645 | 54882 | 1 |
| YCL040W | 114.376159 | 2.0639506 | 0.56249889 | 15.127188 | 0.00051901 | 0.00378167 | GLK1 | III | 50838 | 52340 | 1 |
| YCL042W | 16.8862491 | 1.58879783 | 0.80699558 | 3.80165705 | 0.14944475 | 0.29203007 | NA | III | 50584 | 50943 | 1 |
| YCL043C | 294.126278 | 0.97501271 | 0.53698686 | 8.44110428 | 0.01469053 | 0.05261147 | PDI1 | III | 48653 | 50221 | -1 |
| YCL044C | 93.5532419 | -0.6514168 | 0.58671585 | 1.2357752 | 0.53908199 | 0.69331069 | MGR1 | III | 47111 | 48364 | -1 |
| YCL045C | 78.2732494 | 0.73103425 | 0.6258291 | 2.32244932 | 0.3131025 | 0.48516903 | EMC1 | III | 44623 | 46905 | -1 |
| YCL047C | 23.286016 | 0.52697513 | 0.74226683 | 0.57677301 | 0.74947186 | 0.8479727 | POF1 | III | 43661 | 44437 | -1 |
| YCL048W | 101.085519 | 2.04035775 | 0.72579296 | 9.46582969 | 0.00880078 | 0.03573389 | SPS22 | III | 42165 | 43556 | 1 |
| YCL048W-A | 37.7860573 | 2.974992 | 0.88096071 | 11.2433619 | 0.00361855 | 0.01778559 | NA | III | 41488 | 41727 | 1 |
| YCL049C | 393.739341 | -0.1880208 | 0.51323187 | 2.54201377 | 0.280549 | 0.45466569 | NA | III | 39786 | 40724 | -1 |
| YCL050C | 51.8163578 | -0.3171443 | 0.57840543 | 0.43961437 | 0.80267355 | 0.881373 | APA1 | III | 37836 | 38801 | -1 |
| YCL051W | 149.050248 | 4.95896609 | 0.68727057 | 61.4237848 | 4.59E-14 | 4.03E-12 | LRE1 | III | 35865 | 37616 | 1 |
| YCL052C | 17.4005338 | -0.0500373 | 1.07604121 | 0.27399286 | 0.87197334 | 0.92527295 | PBN1 | III | 34143 | 35393 | -1 |
| YCL054W | 106.061843 | -0.4973111 | 0.86394356 | 3.51460045 | 0.17250997 | 0.32379018 | SPB1 | III | 31449 | 33974 | 1 |
| YCL055W | 7.10275474 | -1.362173 | 1.00317394 | 2.2663983 | 0.32200148 | 0.49479526 | KAR4 | III | 27929 | 28936 | 1 |
| YCL056C | 794.042805 | 0.14313949 | 0.51497275 | 4.06638447 | 0.13091694 | 0.26590071 | PEX34 | III | 26925 | 27359 | -1 |
| YCL057C-A | 93.0929797 | -0.740053 | 0.52572824 | 2.00439056 | 0.36707273 | 0.54025193 | MIC10 | III | 24032 | 24325 | -1 |
| YCL057W | 79.8728449 | -0.566015 | 0.59842207 | 3.33785951 | 0.18844864 | 0.34420115 | PRD1 | III | 24768 | 26906 | 1 |

| | | | | | | | | | | | |
|---|---|---|---|---|---|---|---|---|---|---|---|
| YCL058C | 930.608544 | -2.7353308 | 0.50173224 | 28.4913856 | 6.50E-07 | 1.22E-05 | FYV5 | III | 23523 | 23981 | -1 |
| YCL058W-A | 723.627416 | -0.7358046 | 0.66407409 | 4.4659781 | 0.1072075 | 0.23024163 | ADF1 | III | 23584 | 23925 | 1 |
| YCL059C | 28.9969124 | -2.3492915 | 0.74025031 | 10.3439488 | 0.00567336 | 0.02535586 | KRR1 | III | 22429 | 23379 | -1 |
| YCL061C | 33.0284699 | -1.0265691 | 0.67434929 | 2.39640059 | 0.30173676 | 0.47448396 | MRC1 | III | 18816 | 22106 | -1 |
| YCL063W | 436.06417 | -2.2226806 | 0.49353163 | 20.2877105 | 3.93E-05 | 0.0004228 | VAC17 | III | 17290 | 18561 | 1 |
| YCL064C | 123.37118 | 1.50412677 | 0.99581862 | 4.13595064 | 0.12644153 | 0.25966788 | CHA1 | III | 15798 | 16880 | -1 |
| YCL068C | 23.7561479 | 1.18903655 | 0.99362278 | 1.87739715 | 0.39113654 | 0.56237411 | NA | III | 11503 | 12285 | -1 |
| YCR001W | 336.656574 | 1.80207951 | 0.49129245 | 15.2329575 | 0.00049227 | 0.00363486 | NA | III | 115685 | 115999 | 1 |
| YCR002C | 15.7574711 | 0.70374809 | 0.91835999 | 1.01921125 | 0.60073245 | 0.74682758 | CDC10 | III | 117380 | 118348 | -1 |
| YCR003W | 18.3477302 | -0.8457499 | 0.74058239 | 2.89482966 | 0.23517748 | 0.40286319 | MRPL32 | III | 118620 | 119171 | 1 |
| YCR004C | 34.0747482 | -0.3148929 | 0.63553272 | 0.40343356 | 0.81732638 | 0.89020311 | YCP4 | III | 119575 | 120318 | -1 |
| YCR005C | 37.8609467 | 1.55004266 | 0.7613633 | 4.32972804 | 0.11476554 | 0.24172153 | CIT2 | III | 120946 | 122328 | -1 |
| YCR007C | 311.256928 | 1.09385233 | 0.56751417 | 4.06741363 | 0.13084959 | 0.26590071 | NA | III | 126011 | 126730 | -1 |
| YCR008W | 49.0829016 | -0.3594953 | 0.84941157 | 0.54046554 | 0.76320182 | 0.85588583 | SAT4 | III | 128470 | 130281 | 1 |
| YCR009C | 67.6107321 | -1.1970223 | 0.66024594 | 3.94490681 | 0.13911513 | 0.27774112 | RVS161 | III | 130745 | 131542 | -1 |
| YCR010C | 170.08003 | 1.12328174 | 0.71114163 | 2.43569815 | 0.29586587 | 0.46995614 | ADY2 | III | 132273 | 133124 | -1 |
| YCR011C | 60.8555432 | 0.87577457 | 0.62310785 | 2.12237461 | 0.34604471 | 0.51947079 | ADP1 | III | 133724 | 136873 | -1 |
| YCR012W | 336.293428 | -1.5319183 | 0.70711574 | 4.66002125 | 0.09729471 | 0.21492841 | PGK1 | III | 137746 | 138996 | 1 |
| YCR014C | 44.2700354 | 0.49158184 | 0.59285508 | 0.8475861 | 0.65455933 | 0.78442961 | POL4 | III | 139185 | 140933 | -1 |
| YCR015C | 29.1308523 | 1.48240874 | 0.65113542 | 9.71719786 | 0.00776135 | 0.03237624 | CTO1 | III | 141217 | 142170 | -1 |
| YCR016W | 54.3935952 | -2.1077394 | 0.58137235 | 13.7483025 | 0.00103418 | 0.00679008 | NA | III | 143634 | 144506 | 1 |
| YCR017C | 34.9736628 | -0.1550111 | 0.88571013 | 0.06638544 | 0.96735211 | 0.97486964 | CWH43 | III | 144773 | 147634 | -1 |
| YCR018C | 229.212885 | -0.0271799 | 0.53275899 | 1.97104433 | 0.37324428 | 0.54640641 | SRD1 | III | 148238 | 148903 | -1 |
| YCR019W | 22.8264526 | 0.1567952 | 0.83817957 | 1.19996566 | 0.54882106 | 0.70069987 | MAK32 | III | 152837 | 153928 | 1 |
| YCR020C | 52.9609565 | -0.8554121 | 0.64604678 | 4.99462466 | 0.08230591 | 0.19005499 | PET18 | III | 154012 | 154659 | -1 |
| YCR020C-A | 10.0172298 | 1.09816139 | 0.9464159 | 1.37661848 | 0.50242483 | 0.66093664 | MAK31 | III | 154830 | 155096 | -1 |
| YCR020W-B | 11.8240907 | 1.81190912 | 0.97496121 | 6.91106627 | 0.03157047 | 0.09292985 | HTL1 | III | 155320 | 155556 | 1 |
| YCR021C | 334.952806 | -0.4459499 | 0.61430696 | 2.96486165 | 0.22708501 | 0.39307092 | HSP30 | III | 156109 | 157107 | -1 |
| YCR022C | 16.7210659 | -0.6964612 | 0.7748446 | 1.12283461 | 0.57040006 | 0.72078052 | NA | III | 157421 | 157765 | -1 |
| YCR023C | 244.036595 | -0.1459145 | 0.50831803 | 3.55612731 | 0.168965 | 0.31887463 | NA | III | 158538 | 160373 | -1 |
| YCR024C | 100.911755 | -0.0147921 | 0.54350648 | 1.35888884 | 0.50689854 | 0.66464916 | SLM5 | III | 160744 | 162222 | -1 |
| YCR024C-A | 315.274774 | 2.89209972 | 0.62886747 | 18.7168757 | 8.62E-05 | 0.00082519 | PMP1 | III | 162945 | 163067 | -1 |
| YCR024C-B | 352.461296 | 3.56311166 | 0.64140633 | 26.159938 | 2.09E-06 | 3.35E-05 | NA | III | 162599 | 162865 | -1 |

| | | | | | | | | | | | |
|---|---|---|---|---|---|---|---|---|---|---|---|
| YCR025C | 290.595223 | 2.78627046 | 0.55260568 | 23.0053192 | 1.01E-05 | 0.00013356 | NA | III | 163446 | 163856 | -1 |
| YCR026C | 53.3210893 | 0.27324324 | 0.57552065 | 1.14726061 | 0.56347613 | 0.71468276 | NPP1 | III | 164111 | 166339 | -1 |
| YCR027C | 6.03799162 | 0.96585532 | 1.32450022 | 0.85716458 | 0.65143198 | 0.78236902 | RHB1 | III | 167370 | 167999 | -1 |
| YCR028C | 50.9138423 | 0.22252229 | 0.59322036 | 0.66741426 | 0.71626353 | 0.82794286 | FEN2 | III | 170886 | 172424 | -1 |
| YCR028C-A | 9.43023506 | -1.4323457 | 0.91922868 | 2.74098032 | 0.25398244 | 0.42600183 | RIM1 | III | 172950 | 173440 | -1 |
| YCR030C | 120.314438 | -1.3092737 | 0.50791194 | 8.85776568 | 0.01192781 | 0.04496224 | SYP1 | III | 173826 | 176438 | -1 |
| YCR031C | 73.7572058 | -3.2075717 | 0.86432071 | 12.3206157 | 0.0021116 | 0.01164923 | RPS14A | III | 177500 | 178220 | -1 |
| YCR032W | 97.9721179 | 2.32066632 | 0.54067442 | 21.0206423 | 2.73E-05 | 0.00031124 | BPH1 | III | 179520 | 186023 | 1 |
| YCR033W | 98.9295984 | 1.43566086 | 0.5654308 | 6.31538478 | 0.04252376 | 0.1160873 | SNT1 | III | 186489 | 190169 | 1 |
| YCR034W | 52.9589909 | -0.4102904 | 0.58401498 | 1.20738814 | 0.54678803 | 0.6993379 | ELO2 | III | 190592 | 191635 | 1 |
| YCR035C | 94.8994726 | -1.8048179 | 0.53380014 | 12.6451571 | 0.00179531 | 0.01032258 | RRP43 | III | 191834 | 193018 | -1 |
| YCR036W | 70.474612 | 1.47384709 | 0.63610011 | 5.75136145 | 0.05637775 | 0.14342143 | RBK1 | III | 193297 | 194298 | 1 |
| YCR037C | 98.8286519 | 0.61638589 | 0.59788991 | 3.82132717 | 0.14798216 | 0.2902937 | PHO87 | III | 194414 | 197185 | -1 |
| YCR041W | 758.157492 | -0.134903 | 0.59465501 | 4.61587992 | 0.09946594 | 0.21864382 | NA | III | 200911 | 201243 | 1 |
| YCR042C | 543.662284 | -1.9719713 | 0.59447601 | 10.2970603 | 0.00580794 | 0.02584777 | TAF2 | III | 201174 | 205397 | -1 |
| YCR043C | 7.23865365 | -2.6084322 | 1.25491064 | 4.91440003 | 0.0856745 | 0.19653541 | NA | III | 206261 | 206644 | -1 |
| YCR044C | 280.106854 | -2.4102794 | 0.50902082 | 26.2510662 | 1.99E-06 | 3.24E-05 | PER1 | III | 206877 | 207950 | -1 |
| YCR045C | 1395.06948 | -1.7577407 | 0.5832017 | 9.93040502 | 0.00697654 | 0.02994378 | RRT12 | III | 208135 | 209610 | -1 |
| YCR046C | 132.652168 | -1.5166041 | 0.52859001 | 7.95256421 | 0.01875524 | 0.06302544 | IMG1 | III | 209914 | 210423 | -1 |
| YCR047C | 63.2036226 | 0.29968003 | 0.64799618 | 0.68849199 | 0.70875456 | 0.82407349 | BUD23 | III | 210718 | 211545 | -1 |
| YCR048W | 21.713628 | -0.3091182 | 0.7494559 | 0.48086191 | 0.78628893 | 0.87021248 | ARE1 | III | 211929 | 213761 | 1 |
| YCR050C | 2212.76119 | 0.61061157 | 0.46837969 | 2.42760413 | 0.29706567 | 0.4709776 | NA | III | 213464 | 213772 | -1 |
| YCR051W | 146.173156 | 0.58829186 | 0.52405807 | 1.30930675 | 0.51962215 | 0.67578891 | NA | III | 214071 | 214739 | 1 |
| YCR052W | 36.2201719 | -0.301243 | 0.65014526 | 0.22871397 | 0.89193949 | 0.93370632 | RSC6 | III | 214994 | 216445 | 1 |
| YCR053W | 79.7704856 | -1.0814074 | 0.55169733 | 4.59895721 | 0.10031113 | 0.21944212 | THR4 | III | 216697 | 218241 | 1 |
| YCR054C | 34.9866406 | -1.2634049 | 0.83394996 | 4.96557022 | 0.08351032 | 0.19225635 | CTR86 | III | 218376 | 220067 | -1 |
| YCR057C | 27.9095756 | -1.6992008 | 0.68772995 | 6.44673536 | 0.03982073 | 0.11116077 | PWP2 | III | 220457 | 223228 | -1 |
| YCR059C | 148.648024 | -3.0295441 | 0.53787624 | 35.5672339 | 1.89E-08 | 5.26E-07 | YIH1 | III | 223454 | 224230 | -1 |
| YCR060W | 136.948626 | 1.48779418 | 0.58028498 | 14.5463924 | 0.00069389 | 0.00484051 | TAH1 | III | 224399 | 224734 | 1 |
| YCR061W | 1626.24945 | 3.86175663 | 0.51798243 | 53.7826072 | 2.10E-12 | 1.39E-10 | NA | III | 225563 | 227458 | 1 |
| YCR063W | 19.0498706 | -1.0075649 | 0.77804543 | 4.97666213 | 0.08304845 | 0.19134522 | BUD31 | III | 228318 | 228791 | 1 |
| YCR065W | 40.1115991 | -0.1911583 | 0.60071082 | 0.12171128 | 0.94095906 | 0.96087354 | HCM1 | III | 229310 | 231004 | 1 |
| YCR066W | 19.2850513 | -0.8026771 | 0.76281917 | 1.0925808 | 0.57909404 | 0.72795364 | RAD18 | III | 231500 | 232963 | 1 |

| | | | | | | | | | | | |
|---|---|---|---|---|---|---|---|---|---|---|---|
| YCR067C | 59.0079036 | 0.7530949 | 0.74129609 | 1.62788312 | 0.44310808 | 0.61162622 | SED4 | III | 233125 | 236322 | -1 |
| YCR068W | 17.8603771 | 0.19263067 | 0.8199776 | 0.71077454 | 0.70090195 | 0.81801018 | ATG15 | III | 237214 | 238776 | 1 |
| YCR069W | 26.2965861 | -0.4578155 | 0.71156031 | 1.05431342 | 0.59028092 | 0.73765326 | CPR4 | III | 239055 | 240011 | 1 |
| YCR071C | 608.495607 | 1.91997974 | 1.60741742 | 4.86100758 | 0.08799249 | 0.20050237 | IMG2 | III | 240103 | 240543 | -1 |
| YCR072C | 16.3417877 | -0.3512783 | 0.78458368 | 0.49751597 | 0.77976867 | 0.86640963 | RSA4 | III | 240805 | 242352 | -1 |
| YCR073C | 1117.23221 | -2.4591718 | 0.48536768 | 23.7522924 | 6.95E-06 | 9.52E-05 | SSK22 | III | 242588 | 246583 | -1 |
| YCR073W-A | 46.5795931 | 0.42578221 | 0.68481201 | 7.95875781 | 0.01869725 | 0.06286705 | SOL2 | III | 246963 | 247910 | 1 |
| YCR075C | 17.9377281 | -0.8274937 | 0.84930297 | 2.33093777 | 0.31177644 | 0.48409375 | ERS1 | III | 248033 | 248815 | -1 |
| YCR075W-A | 53.7519531 | 0.94053188 | 0.91758589 | 6.45881802 | 0.03958088 | 0.11076526 | EGO2 | III | 248975 | 249202 | 1 |
| YCR076C | 73.6517941 | -0.0507196 | 0.68766093 | 0.54170441 | 0.76272921 | 0.85565759 | FUB1 | III | 249293 | 250045 | -1 |
| YCR077C | 19.1557254 | -1.159975 | 0.84372802 | 2.28950833 | 0.31830216 | 0.49106568 | PAT1 | III | 250238 | 252628 | -1 |
| YCR079W | 99.654938 | -0.9179305 | 0.53389677 | 7.40262239 | 0.02469113 | 0.07709579 | PTC6 | III | 252845 | 254173 | 1 |
| YCR081W | 22.9218269 | 0.61211432 | 0.7429282 | 3.70444777 | 0.15688788 | 0.30265354 | SRB8 | III | 254371 | 258654 | 1 |
| YCR082W | 24.091331 | 0.4271743 | 0.82178948 | 1.48642994 | 0.47558247 | 0.63800336 | AHC2 | III | 258883 | 259269 | 1 |
| YCR083W | 26.5801793 | 0.45589854 | 0.71528411 | 1.3879461 | 0.49958724 | 0.65905903 | TRX3 | III | 259578 | 259961 | 1 |
| YCR084C | 113.174446 | -0.4891341 | 0.62074402 | 2.27780703 | 0.32016989 | 0.49276546 | TUP1 | III | 260311 | 262452 | -1 |
| YCR085W | 26.9306015 | -3.4330534 | 0.77924845 | 18.5370214 | 9.43E-05 | 0.00088971 | NA | III | 262916 | 263269 | 1 |
| YCR086W | 118.76426 | -2.4914852 | 0.77961294 | 12.0848647 | 0.00237577 | 0.012725 | CSM1 | III | 263392 | 263964 | 1 |
| YCR087C-A | 57.1827257 | -1.2717672 | 0.74183057 | 3.91309119 | 0.14134584 | 0.28046348 | NA | III | 264006 | 264467 | -1 |
| YCR088W | 154.335812 | -0.3794648 | 0.54775743 | 0.47966806 | 0.78675843 | 0.8703346 | ABP1 | III | 265068 | 266846 | 1 |
| YCR089W | 179.551968 | -1.1001355 | 0.65161727 | 6.92840266 | 0.03129799 | 0.09240968 | FIG2 | III | 267434 | 272263 | 1 |
| YCR090C | 35.809202 | 0.47087907 | 1.05526624 | 9.40659015 | 0.00906536 | 0.03655182 | NA | III | 272315 | 272863 | -1 |
| YCR091W | 17.1212582 | -0.6603543 | 0.79447072 | 2.89869469 | 0.23472343 | 0.40225741 | KIN82 | III | 274404 | 276566 | 1 |
| YCR092C | 258.811154 | 1.07858206 | 0.66067807 | 2.98800691 | 0.22447219 | 0.38994719 | MSH3 | III | 276764 | 279820 | -1 |
| YCR093W | 66.601645 | -0.5248713 | 0.64697937 | 2.90854637 | 0.23357006 | 0.40094002 | CDC39 | III | 280117 | 286443 | 1 |
| YCR094W | 30.0038587 | 1.30491049 | 0.65476492 | 8.89042321 | 0.01173462 | 0.04440749 | CDC50 | III | 286762 | 287937 | 1 |
| YCR095C | 229.922913 | -0.4567993 | 0.72626327 | 7.67034847 | 0.02159758 | 0.07017394 | OCA4 | III | 288170 | 289258 | -1 |
| YCR095W-A | 16.1665781 | -0.3061485 | 0.77537736 | 2.19467386 | 0.33375872 | 0.50787464 | NA | III | 289639 | 289797 | 1 |
| YCR098C | 46.8822512 | 1.04942211 | 0.65361854 | 3.50643169 | 0.17321601 | 0.32488523 | GIT1 | III | 297049 | 298605 | -1 |
| YCR099C | 11.9341417 | 2.67165519 | 1.06905992 | 6.7134447 | 0.0348493 | 0.10013768 | NA | III | 300832 | 301299 | -1 |
| YCR100C | 6.73706912 | 3.41555709 | 1.41663619 | 7.46233843 | 0.0239648 | 0.07565017 | NA | III | 301271 | 302221 | -1 |
| YCR101C | 11.1913099 | 2.82636506 | 1.10514342 | 7.43089371 | 0.02434456 | 0.07668934 | NA | III | 302482 | 303030 | -1 |
| YCR102C | 26.2651998 | 0.63117702 | 0.66403577 | 1.55117522 | 0.46043315 | 0.62448065 | NA | III | 304361 | 305467 | -1 |

| | | | | | | | | | | | |
|---|---|---|---|---|---|---|---|---|---|---|---|
| YCR105W | 1389.81288 | 0.428754 | 0.59093014 | 0.54766291 | 0.76046024 | 0.85479805 | ADH7 | III | 309070 | 310155 | 1 |
| YCR106W | 980.370505 | -3.4286329 | 0.52283677 | 39.2477513 | 3.00E-09 | 1.01E-07 | RDS1 | III | 310958 | 313456 | 1 |
| YCR107W | 338.190843 | -3.555267 | 0.52083763 | 43.0456979 | 4.50E-10 | 1.77E-08 | AAD3 | III | 313890 | 314981 | 1 |
| YDL001W | 14.2188184 | -0.773376 | 0.90718467 | 0.8191032 | 0.6639479 | 0.79099966 | RMD1 | IV | 447984 | 449276 | 1 |
| YDL002C | 29.0599948 | -0.3865354 | 0.66580061 | 0.62485025 | 0.73167041 | 0.83702784 | NHP10 | IV | 446967 | 447578 | -1 |
| YDL003W | 108.1267 | 1.71100662 | 0.54424016 | 12.9996249 | 0.00150372 | 0.00913878 | MCD1 | IV | 444683 | 446383 | 1 |
| YDL004W | 8.04822654 | -1.1478497 | 1.17224941 | 1.36411371 | 0.50557603 | 0.66363301 | ATP16 | IV | 443029 | 443511 | 1 |
| YDL005C | 278.661671 | 0.82074913 | 0.78895734 | 3.72930557 | 0.15495 | 0.29975293 | MED2 | IV | 441014 | 442309 | -1 |
| YDL006W | 76.5440348 | 0.78479947 | 0.69242474 | 1.34590438 | 0.51020014 | 0.66786132 | PTC1 | IV | 439909 | 440754 | 1 |
| YDL007W | 75.4203554 | 0.58555331 | 0.79348308 | 0.82702098 | 0.6613246 | 0.78897407 | RPT2 | IV | 438047 | 439360 | 1 |
| YDL008W | 207.013162 | -1.5721461 | 0.54220755 | 8.08143127 | 0.01758488 | 0.05999792 | APC11 | IV | 433497 | 433994 | 1 |
| YDL009C | 3549.89711 | 1.5143564 | 0.50747973 | 9.5357725 | 0.00849832 | 0.03482328 | NA | IV | 432925 | 433248 | -1 |
| YDL010W | 25.7091349 | 0.7936977 | 0.8395554 | 1.14244976 | 0.56483316 | 0.71577676 | GRX6 | IV | 432330 | 433025 | 1 |
| YDL012C | 50.3197723 | -0.2222063 | 0.68468857 | 1.76417171 | 0.41391864 | 0.58553357 | NA | IV | 431108 | 431517 | -1 |
| YDL013W | 93.2697411 | 0.59345949 | 0.56880478 | 2.0404844 | 0.36050761 | 0.53345156 | SLX5 | IV | 429067 | 430926 | 1 |
| YDL014W | 120.928798 | -2.4799806 | 0.81453416 | 8.80739519 | 0.01223203 | 0.04578115 | NOP1 | IV | 427364 | 428347 | 1 |
| YDL015C | 53.0464213 | -0.6529404 | 0.61494699 | 1.41110459 | 0.49383576 | 0.65469026 | TSC13 | IV | 426002 | 426934 | -1 |
| YDL017W | 8.0008098 | -0.5819688 | 1.08829601 | 0.62884198 | 0.73021155 | 0.83678999 | CDC7 | IV | 424209 | 425732 | 1 |
| YDL018C | 13.9249638 | 0.84550471 | 0.793877 | 1.85479018 | 0.39558283 | 0.56679648 | ERP3 | IV | 422833 | 423510 | -1 |
| YDL019C | 44.3007987 | -0.1710004 | 0.60419113 | 0.85333012 | 0.65268213 | 0.78305626 | OSH2 | IV | 417663 | 421514 | -1 |
| YDL020C | 36.3343799 | 1.15996854 | 0.76399298 | 2.32167902 | 0.31322312 | 0.48516903 | RPN4 | IV | 415113 | 416708 | -1 |
| YDL021W | 41.7212904 | 1.35150526 | 0.651618 | 5.04643784 | 0.08020103 | 0.18675877 | GPM2 | IV | 413953 | 414888 | 1 |
| YDL022W | 126.041901 | 2.27031291 | 0.66391602 | 11.932142 | 0.0025643 | 0.01350981 | GPD1 | IV | 411825 | 413000 | 1 |
| YDL024C | 10.0062406 | 0.38552517 | 0.97624299 | 0.1636896 | 0.92141495 | 0.95035295 | DIA3 | IV | 408451 | 409857 | -1 |
| YDL025C | 73.4476898 | -1.705386 | 0.55501976 | 13.638276 | 0.00109266 | 0.00709251 | RTK1 | IV | 405344 | 407206 | -1 |
| YDL027C | 46.3521828 | 1.13075745 | 0.61830821 | 3.90945528 | 0.14160304 | 0.28078137 | MRX9 | IV | 403695 | 404957 | -1 |
| YDL028C | 19.6224839 | -0.0218613 | 0.7315906 | 0.13710473 | 0.93374456 | 0.95735444 | MPS1 | IV | 400997 | 403291 | -1 |
| YDL029W | 83.7846692 | 1.75904197 | 0.58945156 | 10.9232057 | 0.00424674 | 0.02008876 | ARP2 | IV | 399340 | 400638 | 1 |
| YDL030W | 74.7417788 | 1.37917918 | 0.55747155 | 6.73335497 | 0.03450409 | 0.09954094 | PRP9 | IV | 397537 | 399129 | 1 |
| YDL031W | 49.8269737 | -1.6629045 | 0.57030481 | 10.9613855 | 0.00416644 | 0.01978975 | DBP10 | IV | 394217 | 397204 | 1 |
| YDL033C | 145.008975 | 2.82160441 | 0.51194452 | 31.9266112 | 1.17E-07 | 2.76E-06 | SLM3 | IV | 392659 | 393912 | -1 |
| YDL035C | 154.996064 | -0.6089312 | 0.5483895 | 1.28295167 | 0.5265148 | 0.68176637 | GPR1 | IV | 389172 | 392057 | -1 |
| YDL036C | 777.878867 | 0.86915552 | 0.4934554 | 4.14198718 | 0.12606047 | 0.25919393 | PUS9 | IV | 387513 | 388901 | -1 |

| | | | | | | | | | | | |
|---|---|---|---|---|---|---|---|---|---|---|---|
| YDL037C | 13.4495658 | 1.58252783 | 0.91357858 | 6.44796779 | 0.0397962 | 0.11116077 | BSC1 | IV | 384601 | 385587 | -1 |
| YDL039C | 13.5411867 | 0.38641328 | 0.8458995 | 0.25322743 | 0.88107395 | 0.9300006 | PRM7 | IV | 381985 | 384081 | -1 |
| YDL040C | 43.2693753 | -1.1706169 | 0.62561958 | 4.07326588 | 0.13046726 | 0.26533384 | NAT1 | IV | 378874 | 381438 | -1 |
| YDL042C | 28.9576134 | -0.4044118 | 0.66931066 | 0.68489133 | 0.7100317 | 0.82438173 | SIR2 | IV | 376757 | 378445 | -1 |
| YDL043C | 9.54069089 | 0.7818644 | 0.95068568 | 0.89476425 | 0.63929957 | 0.77519257 | PRP11 | IV | 375680 | 376480 | -1 |
| YDL044C | 41.8484206 | 1.76474914 | 0.6202295 | 9.00467586 | 0.01108305 | 0.04252544 | MTF2 | IV | 373967 | 375289 | -1 |
| YDL045C | 8.56391719 | 1.2727879 | 1.09944106 | 1.7253481 | 0.42203204 | 0.59293772 | FAD1 | IV | 372688 | 373608 | -1 |
| YDL045W-A | 15.1028616 | 1.52007657 | 0.87680727 | 3.11675128 | 0.21047769 | 0.37348017 | MRP10 | IV | 372248 | 372535 | 1 |
| YDL046W | 370.571418 | 0.86640902 | 0.49437048 | 5.13697657 | 0.07665133 | 0.18033776 | NPC2 | IV | 371240 | 371761 | 1 |
| YDL047W | 30.7840109 | 1.30432438 | 0.67058265 | 4.84979154 | 0.08848734 | 0.20139218 | SIT4 | IV | 369771 | 370706 | 1 |
| YDL048C | 209.675831 | 1.66960978 | 0.50493128 | 10.489619 | 0.00527483 | 0.0239164 | STP4 | IV | 366739 | 368211 | -1 |
| YDL049C | 72.0662531 | 0.64439428 | 0.54537179 | 1.47793519 | 0.47760674 | 0.64027392 | KNH1 | IV | 365068 | 365874 | -1 |
| YDL051W | 28.5085026 | -2.7419459 | 0.91429342 | 8.82074797 | 0.01215063 | 0.04558315 | LHP1 | IV | 363952 | 364779 | 1 |
| YDL052C | 27.3436663 | -0.0110998 | 0.72891609 | 0.08216472 | 0.95975008 | 0.97166515 | SLC1 | IV | 362672 | 363583 | -1 |
| YDL053C | 34.0407731 | 0.81718927 | 0.65286521 | 2.22954705 | 0.32798955 | 0.50041094 | PBP4 | IV | 361699 | 362256 | -1 |
| YDL054C | 45.1572706 | 1.81996482 | 0.67251011 | 6.98191132 | 0.03047174 | 0.09033864 | MCH1 | IV | 359825 | 361285 | -1 |
| YDL055C | 106.324777 | -1.2098104 | 0.88153077 | 2.19134155 | 0.33431528 | 0.50802806 | PSA1 | IV | 355674 | 356759 | -1 |
| YDL056W | 48.5245524 | 1.71361093 | 0.81071821 | 6.32169689 | 0.04238976 | 0.11597295 | MBP1 | IV | 352877 | 355378 | 1 |
| YDL057W | 419.537399 | 7.76351461 | 0.69044033 | 122.302566 | 2.77E-27 | 5.34E-24 | NA | IV | 351434 | 352420 | 1 |
| YDL058W | 53.1160216 | 0.23448165 | 0.60622833 | 0.23214308 | 0.89041152 | 0.93317728 | USO1 | IV | 345665 | 351037 | 1 |
| YDL059C | 7.69367683 | -1.3606263 | 1.0069307 | 2.4185594 | 0.29841215 | 0.4716916 | RAD59 | IV | 344237 | 344953 | -1 |
| YDL060W | 72.6023375 | -0.313243 | 0.60678344 | 0.38245425 | 0.82594497 | 0.89568423 | TSR1 | IV | 341619 | 343985 | 1 |
| YDL061C | 59.3751347 | -2.4852119 | 0.62842393 | 14.8634519 | 0.00059216 | 0.00424214 | RPS29B | IV | 340628 | 340798 | -1 |
| YDL063C | 89.6418855 | 2.966615 | 0.57541237 | 40.5372613 | 1.58E-09 | 5.67E-08 | SYO1 | IV | 338272 | 340134 | -1 |
| YDL064W | 13.6504755 | -2.6173328 | 0.86505635 | 10.5790864 | 0.00504406 | 0.02308706 | UBC9 | IV | 337487 | 338070 | 1 |
| YDL065C | 124.326127 | 0.00931495 | 0.67803723 | 4.76981948 | 0.09209729 | 0.20646161 | PEX19 | IV | 336249 | 337277 | -1 |
| YDL066W | 70.9023283 | -0.7619156 | 0.54909578 | 2.08351652 | 0.35283376 | 0.52643121 | IDP1 | IV | 334835 | 336121 | 1 |
| YDL067C | 46.1127844 | -0.3784891 | 0.70981418 | 0.68395646 | 0.71036367 | 0.82438173 | COX9 | IV | 334217 | 334396 | -1 |
| YDL069C | 463.082143 | -0.2072559 | 0.64249457 | 3.26777411 | 0.19516946 | 0.35245914 | CBS1 | IV | 333121 | 333810 | -1 |
| YDL070W | 38.0690166 | 0.8631335 | 0.61778856 | 2.64248231 | 0.26680395 | 0.43923653 | BDF2 | IV | 331024 | 332940 | 1 |
| YDL072C | 240.162161 | 1.6593821 | 0.52057793 | 12.5355052 | 0.00189649 | 0.01073943 | YET3 | IV | 329836 | 330447 | -1 |
| YDL073W | 363.718946 | 5.03586678 | 0.54827987 | 81.5952883 | 1.91E-18 | 3.96E-16 | AHK1 | IV | 326613 | 329567 | 1 |
| YDL074C | 187.067516 | -0.9690926 | 0.49682787 | 4.63766917 | 0.09838818 | 0.21668603 | BRE1 | IV | 324047 | 326149 | -1 |

| | | | | | | | | | | | |
|---|---|---|---|---|---|---|---|---|---|---|---|
| YDL075W | 141.390923 | -3.3199318 | 0.60585891 | 28.1198686 | 7.83E-07 | 1.44E-05 | RPL31A | IV | 322226 | 322988 | 1 |
| YDL076C | 118.624806 | 0.77763681 | 0.50987153 | 2.31033938 | 0.31500408 | 0.48701565 | RXT3 | IV | 320668 | 321552 | -1 |
| YDL077C | 26.0018394 | -0.8456411 | 0.7131383 | 2.12959832 | 0.3447971 | 0.51822506 | VAM6 | IV | 316971 | 320120 | -1 |
| YDL078C | 51.3206204 | -1.8765227 | 0.58988171 | 10.9322698 | 0.00422754 | 0.02004706 | MDH3 | IV | 315357 | 316388 | -1 |
| YDL079C | 642.937363 | -0.7477696 | 0.45941771 | 2.6253528 | 0.26909888 | 0.44226014 | MRK1 | IV | 312951 | 314748 | -1 |
| YDL080C | 40.0134938 | -0.4865839 | 0.66209949 | 0.6803479 | 0.71164652 | 0.82438173 | THI3 | IV | 310642 | 312471 | -1 |
| YDL081C | 118.463587 | -1.6038494 | 0.52251204 | 10.3211921 | 0.00573828 | 0.02555741 | RPP1A | IV | 309802 | 310122 | -1 |
| YDL082W | 39.1510609 | -1.5525754 | 0.63048608 | 9.63655342 | 0.0080807 | 0.03341947 | RPL13A | IV | 308424 | 309388 | 1 |
| YDL083C | 59.4584439 | -2.7025378 | 0.70551354 | 14.6682837 | 0.00065286 | 0.00461548 | RPS16A | IV | 306926 | 307789 | -1 |
| YDL084W | 81.459915 | -0.736738 | 0.76023438 | 8.29735906 | 0.01578525 | 0.05583175 | SUB2 | IV | 305237 | 306577 | 1 |
| YDL085C-A | 44.4140201 | 0.05052094 | 0.59194201 | 1.588773 | 0.45185837 | 0.61806282 | NA | IV | 302464 | 302670 | -1 |
| YDL085W | 101.640533 | 0.91473683 | 0.52733872 | 3.08194967 | 0.21417222 | 0.37813088 | NDE2 | IV | 303211 | 304848 | 1 |
| YDL086W | 193.134209 | -1.3352961 | 0.51419871 | 6.55607059 | 0.03770226 | 0.1063821 | NA | IV | 301413 | 302234 | 1 |
| YDL088C | 32.763577 | 1.16830326 | 0.68402788 | 4.65739911 | 0.09742236 | 0.21492841 | ASM4 | IV | 298417 | 300003 | -1 |
| YDL089W | 144.625397 | 1.85854379 | 0.86121535 | 8.79104744 | 0.01233242 | 0.0461222 | NUR1 | IV | 296820 | 298274 | 1 |
| YDL090C | 44.4426734 | 0.34492121 | 0.61509133 | 0.86838091 | 0.64778887 | 0.77960872 | RAM1 | IV | 295034 | 296329 | -1 |
| YDL091C | 36.5121502 | 0.6622965 | 0.62599138 | 1.81368371 | 0.40379746 | 0.57458523 | UBX3 | IV | 293392 | 294759 | -1 |
| YDL092W | 47.2474625 | 1.38288703 | 0.63971721 | 4.79852059 | 0.09078508 | 0.20445182 | SRP14 | IV | 292781 | 293221 | 1 |
| YDL093W | 147.568304 | 1.8929886 | 0.56848842 | 11.6333555 | 0.00297748 | 0.01516237 | PMT5 | IV | 289909 | 292140 | 1 |
| YDL095W | 136.459951 | 0.12897428 | 0.53166682 | 3.11019474 | 0.21116882 | 0.37459175 | PMT1 | IV | 287059 | 289512 | 1 |
| YDL097C | 48.117063 | -0.6189261 | 0.62624536 | 1.07658305 | 0.58374471 | 0.73199919 | RPN6 | IV | 285391 | 286695 | -1 |
| YDL099W | 99.8563851 | 0.92089989 | 0.62406894 | 3.31015777 | 0.19107698 | 0.34746725 | BUG1 | IV | 283419 | 284444 | 1 |
| YDL100C | 44.0981086 | 0.27158482 | 0.63950287 | 0.20613064 | 0.90206805 | 0.93824416 | GET3 | IV | 282112 | 283176 | -1 |
| YDL101C | 32.8447844 | -0.6681523 | 0.64253671 | 1.68615875 | 0.43038317 | 0.59945118 | DUN1 | IV | 280307 | 281848 | -1 |
| YDL102W | 89.7295819 | 1.90139692 | 0.53573566 | 13.3970943 | 0.0012327 | 0.00781955 | POL3 | IV | 276872 | 280165 | 1 |
| YDL103C | 11.8540505 | -0.3132142 | 0.83828545 | 0.77935549 | 0.6772751 | 0.8011991 | QRI1 | IV | 275148 | 276581 | -1 |
| YDL104C | 259.637937 | -1.2728441 | 0.60789313 | 4.32136721 | 0.11524631 | 0.24255767 | QRI7 | IV | 273653 | 274876 | -1 |
| YDL105W | 98.8256615 | -0.7505746 | 0.53711535 | 2.18043235 | 0.33614382 | 0.50936213 | NSE4 | IV | 272389 | 273597 | 1 |
| YDL106C | 27.4026596 | 0.20825131 | 0.82704108 | 1.21939774 | 0.54351451 | 0.69699868 | PHO2 | IV | 270222 | 271901 | -1 |
| YDL107W | 105.203975 | 0.10825879 | 0.50581806 | 0.09496677 | 0.95362632 | 0.96767513 | MSS2 | IV | 268921 | 269976 | 1 |
| YDL108W | 70.8727646 | 0.5084336 | 0.67029375 | 0.7987281 | 0.67074647 | 0.79566115 | KIN28 | IV | 267698 | 268699 | 1 |
| YDL109C | 46.5217634 | -0.0868762 | 0.60263919 | 1.72003634 | 0.42315439 | 0.59395489 | NA | IV | 265258 | 267201 | -1 |
| YDL110C | 93.329144 | 1.60930225 | 0.54319117 | 10.0660704 | 0.00651899 | 0.02829458 | TMA17 | IV | 264512 | 264964 | -1 |

| | | | | | | | | | | | |
|---|---|---|---|---|---|---|---|---|---|---|---|
| YDL111C | 125.241481 | -2.2039575 | 0.71342774 | 10.3275598 | 0.00572004 | 0.02550402 | RRP42 | IV | 263313 | 264110 | -1 |
| YDL112W | 113.816495 | 0.00209097 | 0.53148912 | 0.08778042 | 0.95705903 | 0.96978855 | TRM3 | IV | 258915 | 263225 | 1 |
| YDL113C | 17.7860853 | 1.58427871 | 0.82934891 | 4.99417702 | 0.08232434 | 0.19005499 | ATG20 | IV | 256633 | 258555 | -1 |
| YDL114W | 33.5629784 | 1.20968641 | 1.03504185 | 5.51712028 | 0.06338297 | 0.15623132 | NA | IV | 255604 | 256530 | 1 |
| YDL115C | 258.465747 | -0.2648933 | 0.51675879 | 1.27382968 | 0.52892172 | 0.68319367 | IWR1 | IV | 253995 | 255126 | -1 |
| YDL116W | 7.35085685 | -0.0801884 | 1.18780859 | 0.33325447 | 0.8465151 | 0.90975468 | NUP84 | IV | 251566 | 253746 | 1 |
| YDL117W | 54.4486454 | 0.74862732 | 0.69417641 | 2.55132013 | 0.27924659 | 0.45351049 | CYK3 | IV | 248581 | 251238 | 1 |
| YDL119C | 36.8598278 | 0.44836869 | 0.66536436 | 1.62488726 | 0.44377233 | 0.61162622 | HEM25 | IV | 246689 | 247612 | -1 |
| YDL120W | 131.899966 | 2.17044527 | 0.63670077 | 10.8538365 | 0.00439662 | 0.02061251 | YFH1 | IV | 245923 | 246447 | 1 |
| YDL121C | 8.09890406 | -2.5179861 | 1.11051547 | 5.28454808 | 0.07119918 | 0.17091344 | NA | IV | 245133 | 245582 | -1 |
| YDL122W | 264.640069 | -1.3880079 | 0.60896433 | 8.63216016 | 0.01335212 | 0.04886775 | UBP1 | IV | 242552 | 244981 | 1 |
| YDL123W | 229.663303 | 1.21094991 | 0.48774097 | 6.17620823 | 0.0455883 | 0.12191042 | SNA4 | IV | 241418 | 241840 | 1 |
| YDL124W | 197.03977 | 0.64264516 | 0.65184572 | 3.23024937 | 0.19886587 | 0.35758801 | NA | IV | 240259 | 241197 | 1 |
| YDL125C | 57.8758117 | 0.08215954 | 0.57319007 | 0.30261696 | 0.85958249 | 0.91706593 | HNT1 | IV | 239019 | 239606 | -1 |
| YDL126C | 250.93767 | 0.18033487 | 0.49815194 | 0.62230746 | 0.73260124 | 0.83746519 | CDC48 | IV | 236157 | 238664 | -1 |
| YDL127W | 182.261519 | 0.75787123 | 0.54523536 | 2.79911403 | 0.24670623 | 0.41766412 | PCL2 | IV | 234927 | 235853 | 1 |
| YDL128W | 649.534225 | -0.3557929 | 0.47368293 | 0.64073737 | 0.72588137 | 0.83456178 | VCX1 | IV | 232652 | 233887 | 1 |
| YDL129W | 91.1220606 | 1.18506294 | 0.55861419 | 4.72833494 | 0.09402755 | 0.2093602 | NA | IV | 231024 | 231899 | 1 |
| YDL130W | 44.6251435 | -3.0990665 | 0.71575135 | 17.5456645 | 0.00015488 | 0.00137233 | RPP1B | IV | 229906 | 230527 | 1 |
| YDL130W-A | 28.5033951 | 4.02339053 | 0.86769061 | 21.4473673 | 2.20E-05 | 0.00025693 | STF1 | IV | 229171 | 229431 | 1 |
| YDL131W | 31.4333264 | 0.45889632 | 0.66490432 | 0.67648802 | 0.71302128 | 0.8253485 | LYS21 | IV | 227393 | 228715 | 1 |
| YDL132W | 26.2996809 | 0.15536297 | 0.71417575 | 0.07724642 | 0.96211316 | 0.97258112 | CDC53 | IV | 224304 | 226751 | 1 |
| YDL133C-A | 637.842123 | -2.2999266 | 0.72068105 | 9.8441755 | 0.00728391 | 0.03098738 | RPL41A | IV | 221724 | 221801 | -1 |
| YDL133W | 1215.286 | 6.11049344 | 0.55973409 | 86.6757576 | 1.51E-19 | 3.97E-17 | SRF1 | IV | 222427 | 223740 | 1 |
| YDL134C | 92.1377053 | 1.51314145 | 0.53559933 | 8.60504444 | 0.01353438 | 0.04937874 | PPH21 | IV | 219662 | 220771 | -1 |
| YDL135C | 30.6744714 | -0.2104959 | 0.77540394 | 0.10881451 | 0.94704634 | 0.96419875 | RDI1 | IV | 218680 | 219288 | -1 |
| YDL136W | 104.392381 | -0.6231264 | 0.5798723 | 1.46624998 | 0.48040538 | 0.64328102 | RPL35A | IV | 217600 | 218367 | 1 |
| YDL137W | 94.4457558 | -2.689188 | 0.69345235 | 15.7335741 | 0.00038326 | 0.00297068 | ARF2 | IV | 216529 | 217074 | 1 |
| YDL138W | 255.824156 | -1.7386062 | 0.49508288 | 12.0662313 | 0.00239801 | 0.01281215 | RGT2 | IV | 213351 | 215642 | 1 |
| YDL139C | 46.2250386 | -0.2098726 | 0.64089739 | 0.13125748 | 0.93647848 | 0.95873833 | SCM3 | IV | 211375 | 212046 | -1 |
| YDL140C | 163.073404 | -0.7036723 | 0.62630725 | 11.5974094 | 0.00303148 | 0.01538156 | RPO21 | IV | 205360 | 210561 | -1 |
| YDL141W | 45.9023388 | 3.31078238 | 0.69898484 | 26.8576949 | 1.47E-06 | 2.48E-05 | BPL1 | IV | 203039 | 205111 | 1 |
| YDL142C | 98.7539594 | -0.361124 | 0.53368505 | 1.40310218 | 0.49581565 | 0.65587677 | CRD1 | IV | 201719 | 202570 | -1 |

| | | | | | | | | | | | |
|---|---|---|---|---|---|---|---|---|---|---|---|
| YDL143W | 52.2606569 | -0.8931302 | 0.64699096 | 4.05346448 | 0.1317654 | 0.26711622 | CCT4 | IV | 199996 | 201582 | 1 |
| YDL144C | 22.4116795 | -0.7727061 | 0.7002259 | 1.59605537 | 0.45021606 | 0.61669055 | NA | IV | 198662 | 199732 | -1 |
| YDL145C | 59.203475 | -0.2872747 | 0.66860448 | 0.35950928 | 0.83547518 | 0.90300565 | COP1 | IV | 194571 | 198176 | -1 |
| YDL146W | 26.9751983 | 0.22876403 | 0.74499197 | 0.21032935 | 0.90017627 | 0.93824416 | LDB17 | IV | 192750 | 194225 | 1 |
| YDL147W | 40.6151598 | -0.197512 | 0.70053977 | 0.93724003 | 0.62586536 | 0.76430043 | RPN5 | IV | 190924 | 192261 | 1 |
| YDL148C | 48.9567783 | -1.065644 | 0.72585029 | 3.3127734 | 0.19082725 | 0.34723124 | NOP14 | IV | 188154 | 190586 | -1 |
| YDL149W | 172.034198 | -1.9170173 | 0.52635482 | 12.646335 | 0.00179425 | 0.01032258 | ATG9 | IV | 184925 | 187918 | 1 |
| YDL150W | 33.8252509 | -2.1064244 | 0.70780056 | 9.97939021 | 0.00680774 | 0.02934982 | RPC53 | IV | 183343 | 184611 | 1 |
| YDL153C | 71.3586435 | -0.9466073 | 0.74903314 | 2.24200305 | 0.32595318 | 0.49874971 | SAS10 | IV | 181186 | 183018 | -1 |
| YDL154W | 10.132915 | -2.4291578 | 0.98332312 | 6.4451038 | 0.03985323 | 0.11116077 | MSH5 | IV | 178334 | 181039 | 1 |
| YDL155W | 12.2131574 | -1.4969846 | 0.82695638 | 5.57933389 | 0.06144167 | 0.15241958 | CLB3 | IV | 176773 | 178056 | 1 |
| YDL156W | 12.4497092 | 1.08139351 | 0.85512485 | 4.50709523 | 0.10502597 | 0.22698782 | CMR1 | IV | 174918 | 176486 | 1 |
| YDL157C | 11.6589645 | -1.9508555 | 0.95838271 | 4.69417081 | 0.09564753 | 0.21202113 | NA | IV | 174232 | 174588 | -1 |
| YDL159W | 207.118524 | -0.3604027 | 0.58828716 | 1.20995603 | 0.54608643 | 0.69889749 | STE7 | IV | 172481 | 174028 | 1 |
| YDL159W-A | 233.403292 | -0.3211979 | 0.74381373 | 0.92283845 | 0.63038835 | 0.76792521 | NA | IV | 172182 | 172313 | 1 |
| YDL160C | 9.3715743 | 0.06460691 | 1.0067027 | 0.35388042 | 0.83782988 | 0.90469306 | DHH1 | IV | 170410 | 171930 | -1 |
| YDL160C-A | 31.6922663 | 1.29822577 | 0.65377769 | 4.59484792 | 0.10051745 | 0.21962114 | MHF2 | IV | 169366 | 169608 | -1 |
| YDL161W | 276.349408 | -0.8347384 | 0.66548756 | 5.41739939 | 0.06662338 | 0.16194348 | ENT1 | IV | 167714 | 169078 | 1 |
| YDL164C | 51.6011219 | 1.00480178 | 0.61605883 | 2.77946262 | 0.24914224 | 0.42007384 | CDC9 | IV | 164987 | 167254 | -1 |
| YDL165W | 240.670259 | -0.6292795 | 0.50779109 | 2.9929934 | 0.22391322 | 0.38943778 | CDC36 | IV | 164290 | 164865 | 1 |
| YDL166C | 19.2616683 | -2.1944215 | 0.91243622 | 5.79116699 | 0.05526677 | 0.14109109 | FAP7 | IV | 163449 | 164042 | -1 |
| YDL167C | 69.6698499 | 1.81998006 | 0.63110105 | 18.4962238 | 9.63E-05 | 0.00090509 | NRP1 | IV | 160995 | 163154 | -1 |
| YDL168W | 21.2838689 | 0.55721972 | 0.78076138 | 0.95179722 | 0.62132648 | 0.76184702 | SFA1 | IV | 159604 | 160764 | 1 |
| YDL169C | 29.9602525 | 1.44768851 | 1.00848358 | 7.93295185 | 0.01894006 | 0.06346236 | UGX2 | IV | 158064 | 158735 | -1 |
| YDL170W | 11.4835194 | 1.53410394 | 1.07793743 | 2.49515839 | 0.28719921 | 0.4613994 | UGA3 | IV | 156318 | 157904 | 1 |
| YDL171C | 79.3804091 | -0.2598095 | 0.6377261 | 1.46345774 | 0.48107655 | 0.64358439 | GLT1 | IV | 149203 | 155640 | -1 |
| YDL173W | 302.118732 | 1.23051408 | 0.5371642 | 5.91939818 | 0.05183451 | 0.1343428 | PAR32 | IV | 148191 | 149078 | 1 |
| YDL174C | 122.630622 | 0.59847276 | 0.53971769 | 1.5126089 | 0.46939791 | 0.63190279 | DLD1 | IV | 145826 | 147589 | -1 |
| YDL175C | 63.8892846 | 0.70613028 | 0.61553597 | 5.44478283 | 0.06571741 | 0.16055012 | AIR2 | IV | 144484 | 145518 | -1 |
| YDL176W | 16.0379334 | -0.0022964 | 0.79923305 | 0.03191748 | 0.98416793 | 0.98751104 | NA | IV | 142097 | 144223 | 1 |
| YDL177C | 14.637641 | -0.4014105 | 0.95930092 | 3.07963423 | 0.21442031 | 0.37828653 | NA | IV | 141209 | 141721 | -1 |
| YDL178W | 43.223173 | 1.06516099 | 0.67367498 | 3.39643924 | 0.18300906 | 0.3384294 | DLD2 | IV | 139522 | 141114 | 1 |
| YDL179W | 29.6454138 | -0.003337 | 0.75968342 | 0.07702981 | 0.96221736 | 0.97258112 | PCL9 | IV | 138291 | 139205 | 1 |

| | | | | | | | | | | | |
|---|---|---|---|---|---|---|---|---|---|---|---|
| YDL180W | 19.4840422 | 1.59732193 | 0.77272552 | 5.2458556 | 0.07259002 | 0.17317521 | NA | IV | 135896 | 137539 | 1 |
| YDL181W | 394.947947 | 0.87479302 | 0.55163092 | 4.65728522 | 0.09742791 | 0.21492841 | INH1 | IV | 135179 | 135436 | 1 |
| YDL182W | 37.1237245 | -2.6217533 | 0.85532747 | 10.2557953 | 0.00592901 | 0.02628559 | LYS20 | IV | 133437 | 134723 | 1 |
| YDL183C | 20.2589652 | -1.5066648 | 0.72001514 | 5.08404954 | 0.07870687 | 0.18416483 | NA | IV | 130871 | 131833 | -1 |
| YDL184C | 1517.23062 | -2.8693123 | 0.69796747 | 14.9083378 | 0.00057902 | 0.00416465 | RPL41A | IV | 130407 | 130484 | -1 |
| YDL185W | 83.6142257 | -0.7767328 | 0.72459896 | 1.44399247 | 0.48578155 | 0.64773825 | VMA1 | IV | 126787 | 130002 | 1 |
| YDL186W | 270.9147 | -1.6907125 | 0.53403031 | 10.4250239 | 0.00544797 | 0.02452858 | NA | IV | 125616 | 126449 | 1 |
| YDL188C | 49.4125393 | -0.9514376 | 0.58983024 | 7.02408878 | 0.02983586 | 0.088635 | PPH22 | IV | 123865 | 124998 | -1 |
| YDL189W | 27.2519369 | -0.4544382 | 0.73976821 | 0.41392761 | 0.81304908 | 0.88754792 | RBS1 | IV | 122216 | 123589 | 1 |
| YDL190C | 38.8253492 | -0.814104 | 0.68152139 | 2.05132377 | 0.35855906 | 0.53153496 | UFD2 | IV | 118707 | 121592 | -1 |
| YDL191W | 293.40126 | 0.25651525 | 0.81696851 | 6.40315128 | 0.04069803 | 0.11285424 | RPL35A | IV | 117664 | 118517 | 1 |
| YDL192W | 24.4474893 | -0.7202958 | 0.75887984 | 1.16740145 | 0.55783017 | 0.70907501 | ARF1 | IV | 116321 | 116866 | 1 |
| YDL193W | 26.5522463 | -0.1135201 | 0.7263373 | 0.96310294 | 0.61782411 | 0.75916842 | NUS1 | IV | 114672 | 115799 | 1 |
| YDL194W | 49.8079764 | 3.70169147 | 0.75045363 | 30.4410043 | 2.45E-07 | 5.22E-06 | SNF3 | IV | 111580 | 114234 | 1 |
| YDL195W | 98.5171519 | -0.370686 | 0.61709194 | 2.95129395 | 0.22863076 | 0.39492008 | SEC31 | IV | 107208 | 111029 | 1 |
| YDL197C | 16.5109338 | 0.03470725 | 0.74886287 | 0.0351144 | 0.98259603 | 0.98668592 | ASF2 | IV | 104917 | 106494 | -1 |
| YDL198C | 214.641586 | 0.57310107 | 0.5387355 | 1.13317405 | 0.56745886 | 0.71800411 | GGC1 | IV | 103649 | 104551 | -1 |
| YDL199C | 488.714912 | -0.7244539 | 0.47817268 | 2.35939186 | 0.30737219 | 0.47995819 | NA | IV | 101290 | 103353 | -1 |
| YDL200C | 115.299886 | -1.5522199 | 0.74576318 | 8.44801283 | 0.01463987 | 0.05258366 | MGT1 | IV | 100501 | 101067 | -1 |
| YDL201W | 130.609645 | -0.5744051 | 0.49562697 | 1.38978549 | 0.49912798 | 0.65860324 | TRM8 | IV | 99561 | 100421 | 1 |
| YDL202W | 10.9490693 | -2.0408403 | 1.0329996 | 4.54523371 | 0.10304218 | 0.22378628 | MRPL11 | IV | 98475 | 99224 | 1 |
| YDL203C | 42.9245349 | 2.6595345 | 0.73213681 | 20.2278926 | 4.05E-05 | 0.00043276 | ACK1 | IV | 96082 | 97953 | -1 |
| YDL204W | 17.5533068 | 0.44150771 | 0.7506693 | 0.47721025 | 0.78772587 | 0.87068194 | RTN2 | IV | 94605 | 95786 | 1 |
| YDL205C | 29.1659079 | 0.7616765 | 0.68311941 | 1.36158252 | 0.50621629 | 0.66417229 | HEM3 | IV | 92762 | 93745 | -1 |
| YDL206W | 18.7440155 | -0.0995799 | 0.75407546 | 0.10808811 | 0.94739037 | 0.96420992 | NA | IV | 90176 | 92464 | 1 |
| YDL207W | 298.142318 | 2.69580768 | 0.5153051 | 24.6734418 | 4.39E-06 | 6.35E-05 | GLE1 | IV | 88248 | 89864 | 1 |
| YDL208W | 110.659979 | -3.4906994 | 0.81913052 | 16.6436914 | 0.00024315 | 0.00202564 | NHP2 | IV | 87512 | 87982 | 1 |
| YDL209C | 8.43499462 | -2.1582233 | 1.08980279 | 4.02644886 | 0.13355733 | 0.26953536 | CWC2 | IV | 86207 | 87226 | -1 |
| YDL210W | 162.418264 | 0.9955573 | 0.62257841 | 3.08160756 | 0.21420886 | 0.37813088 | UGA4 | IV | 84270 | 85985 | 1 |
| YDL211C | 143.359524 | 2.3244932 | 0.58548293 | 17.9419548 | 0.00012704 | 0.00114935 | NA | IV | 79294 | 80412 | -1 |
| YDL212W | 112.356912 | 1.47604188 | 0.58202943 | 8.2381671 | 0.01625941 | 0.05678441 | SHR3 | IV | 78426 | 79058 | 1 |
| YDL213C | 179.302254 | -1.7815324 | 0.49379095 | 12.4402594 | 0.00198899 | 0.01111606 | NOP6 | IV | 77289 | 77966 | -1 |
| YDL214C | 697.735197 | 2.10484672 | 0.47521458 | 18.1839788 | 0.00011256 | 0.00103124 | PRR2 | IV | 74446 | 76545 | -1 |

| | | | | | | | | | | | |
|---|---|---|---|---|---|---|---|---|---|---|---|
| YDL215C | 110.31734 | -0.7175153 | 0.55981477 | 5.35176388 | 0.06884608 | 0.16625162 | GDH2 | IV | 70640 | 73918 | -1 |
| YDL216C | 95.1245989 | 0.52980914 | 0.55103559 | 2.31900233 | 0.3136426 | 0.48568886 | RRI1 | IV | 68997 | 70319 | -1 |
| YDL217C | 12.2612598 | -0.2841955 | 1.01089418 | 0.28275529 | 0.86816139 | 0.92249119 | TIM22 | IV | 67983 | 68606 | -1 |
| YDL218W | 290.642644 | -0.5979913 | 0.53371753 | 1.61316481 | 0.44638101 | 0.61376064 | NA | IV | 66493 | 67446 | 1 |
| YDL219W | 27.8068376 | -2.8917954 | 0.74720721 | 14.5081095 | 0.0007073 | 0.00491629 | DTD1 | IV | 65242 | 65765 | 1 |
| YDL220C | 165.744042 | 2.55555529 | 0.50253293 | 24.0082962 | 6.12E-06 | 8.50E-05 | CDC13 | IV | 62244 | 65018 | -1 |
| YDL222C | 198.600521 | 5.07724099 | 0.61086527 | 64.428155 | 1.02E-14 | 9.87E-13 | FMP45 | IV | 60872 | 61801 | -1 |
| YDL223C | 883.363545 | -3.5004726 | 0.52353756 | 38.3153472 | 4.79E-09 | 1.54E-07 | HBT1 | IV | 57265 | 60405 | -1 |
| YDL224C | 115.662148 | 4.08477524 | 0.76516242 | 25.4245083 | 3.01E-06 | 4.63E-05 | WHI4 | IV | 54397 | 56346 | -1 |
| YDL225W | 63.9855735 | 0.03058527 | 0.54129333 | 0.0430486 | 0.9787057 | 0.98363235 | SHS1 | IV | 52445 | 54100 | 1 |
| YDL226C | 28.0553118 | -0.7546651 | 0.65898433 | 1.32317219 | 0.51603221 | 0.67293389 | GCS1 | IV | 51115 | 52173 | -1 |
| YDL227C | 69.7749212 | -1.0360631 | 0.54512826 | 3.58734675 | 0.16634799 | 0.31558154 | HO | IV | 46271 | 48031 | -1 |
| YDL229W | 116.760554 | -3.4310254 | 0.61802315 | 27.6421101 | 9.94E-07 | 1.77E-05 | SSB1 | IV | 44065 | 45906 | 1 |
| YDL230W | 37.4489022 | -0.3779212 | 0.61674834 | 0.47696037 | 0.7878243 | 0.87068194 | PTP1 | IV | 42700 | 43707 | 1 |
| YDL231C | 173.288301 | -0.1286586 | 0.95316705 | 6.17260021 | 0.04567062 | 0.12191465 | BRE4 | IV | 38867 | 42244 | -1 |
| YDL233W | 17.415683 | 1.71828851 | 0.81101369 | 4.76273971 | 0.09242388 | 0.20685781 | MFG1 | IV | 36797 | 38173 | 1 |
| YDL234C | 218.919544 | 0.14374625 | 0.51871719 | 0.07985731 | 0.96085799 | 0.97212597 | GYP7 | IV | 34237 | 36477 | -1 |
| YDL235C | 190.388624 | 2.30593061 | 0.5980812 | 15.321697 | 0.00047091 | 0.00350007 | YPD1 | IV | 33415 | 33918 | -1 |
| YDL236W | 149.9029 | 2.62485629 | 0.56153489 | 20.7880357 | 3.06E-05 | 0.0003442 | PHO13 | IV | 32296 | 33234 | 1 |
| YDL237W | 50.1960388 | 0.49123533 | 0.58296741 | 0.86085764 | 0.6502302 | 0.78140989 | AIM6 | IV | 30657 | 31829 | 1 |
| YDL238C | 52.1291465 | 2.91688558 | 0.66191594 | 27.2703173 | 1.20E-06 | 2.07E-05 | GUD1 | IV | 28985 | 30454 | -1 |
| YDL239C | 10.4984052 | -0.3248552 | 1.0211598 | 0.85026036 | 0.65368468 | 0.7839342 | ADY3 | IV | 26403 | 28775 | -1 |
| YDL241W | 11.8791569 | 1.15606396 | 0.96385171 | 1.72394927 | 0.42232732 | 0.59293772 | NA | IV | 20635 | 21006 | 1 |
| YDL242W | 446.063187 | 0.82883087 | 0.49594252 | 2.96807281 | 0.2267207 | 0.39279962 | NA | IV | 18959 | 19312 | 1 |
| YDL243C | 69.1838164 | 0.49497785 | 0.59086772 | 2.56008531 | 0.27802544 | 0.45261809 | AAD4 | IV | 17577 | 18566 | -1 |
| YDL244W | 8.08072935 | -1.5733082 | 1.00997343 | 3.19609806 | 0.2022908 | 0.36194799 | THI13 | IV | 16204 | 17226 | 1 |
| YDL245C | 539.502453 | 1.57405671 | 0.60260012 | 8.03794742 | 0.0179714 | 0.06102898 | HXT15 | IV | 11657 | 13360 | -1 |
| YDL247W | 34.7258308 | 1.40407065 | 0.666997 | 6.41618691 | 0.04043363 | 0.11228331 | MPH2 | IV | 5985 | 7814 | 1 |
| YDR001C | 84.7276068 | 2.09902532 | 0.71852884 | 8.3862594 | 0.01509896 | 0.05383187 | NTH1 | IV | 450220 | 452475 | -1 |
| YDR002W | 133.215285 | -1.1202491 | 0.62483512 | 3.23693134 | 0.19820257 | 0.35661681 | YRB1 | IV | 453045 | 453650 | 1 |
| YDR003W | 25.8852123 | -0.6159225 | 0.6626202 | 2.65432879 | 0.26522828 | 0.43749737 | RCR2 | IV | 454122 | 454754 | 1 |
| YDR003W-A | 10.5238989 | -2.8718667 | 1.06596501 | 7.2349064 | 0.02685097 | 0.08173877 | NA | IV | 454782 | 454904 | 1 |
| YDR004W | 788.407218 | 3.92206601 | 0.48142762 | 67.1264651 | 2.65E-15 | 2.95E-13 | RAD57 | IV | 455201 | 456583 | 1 |

| | | | | | | | | | | | |
|---|---|---|---|---|---|---|---|---|---|---|---|
| YDR005C | 658.39579 | 3.31392219 | 0.6280887 | 28.6399242 | 6.04E-07 | 1.15E-05 | MAF1 | IV | 456836 | 458103 | -1 |
| YDR006C | 70.0298313 | -0.1456395 | 0.55960939 | 1.91225473 | 0.38437856 | 0.5574307 | SOK1 | IV | 458542 | 461247 | -1 |
| YDR007W | 1359.77214 | -0.3058722 | 0.50086555 | 4.85640743 | 0.08819511 | 0.20080602 | TRP1 | IV | 461842 | 462516 | 1 |
| YDR009W | 96.0243104 | -0.9281769 | 0.58835469 | 2.6938992 | 0.26003225 | 0.43177137 | GAL3 | IV | 463434 | 464996 | 1 |
| YDR010C | 138.898764 | -0.3176682 | 0.58953064 | 0.39306048 | 0.82157649 | 0.89315206 | NA | IV | 465051 | 465383 | -1 |
| YDR011W | 89.3640715 | -1.1884232 | 0.79173514 | 4.92310885 | 0.08530225 | 0.19575903 | SNQ2 | IV | 465919 | 470424 | 1 |
| YDR012W | 72.1054899 | -1.4581134 | 0.67502313 | 4.73687178 | 0.09362705 | 0.20898252 | RPL4B | IV | 471853 | 472941 | 1 |
| YDR013W | 5.29562963 | -0.9651616 | 1.44119924 | 1.45944617 | 0.48204246 | 0.64436175 | PSF1 | IV | 473157 | 473783 | 1 |
| YDR014W | 37.7300773 | 1.2816582 | 0.69249395 | 3.87772588 | 0.14386744 | 0.28410385 | RAD61 | IV | 474046 | 475989 | 1 |
| YDR014W-A | 9.76288092 | -0.3391365 | 0.93702292 | 0.61219795 | 0.73631374 | 0.83938896 | HED1 | IV | 477797 | 478285 | 1 |
| YDR016C | 64.0435035 | 2.01975813 | 0.6415796 | 11.829213 | 0.00269972 | 0.01404438 | DAD1 | IV | 478474 | 478758 | -1 |
| YDR017C | 116.533952 | 2.75006237 | 0.52687692 | 28.1310724 | 7.79E-07 | 1.44E-05 | KCS1 | IV | 479115 | 482267 | -1 |
| YDR018C | 74.1186026 | 1.47326592 | 0.59193576 | 11.4420842 | 0.00327629 | 0.01636734 | NA | IV | 482670 | 483860 | -1 |
| YDR019C | 116.217421 | 4.31276731 | 0.6144194 | 52.3307152 | 4.33E-12 | 2.79E-10 | GCV1 | IV | 484163 | 485365 | -1 |
| YDR020C | 94.749602 | 3.16004 | 0.6043603 | 27.8705059 | 8.87E-07 | 1.61E-05 | DAS2 | IV | 485746 | 486444 | -1 |
| YDR021W | 63.6009026 | 1.3264004 | 0.73798474 | 9.65211032 | 0.00801809 | 0.03327938 | FAL1 | IV | 486804 | 488003 | 1 |
| YDR022C | 30.1690674 | 0.20857681 | 0.64147567 | 2.58794545 | 0.27417938 | 0.44776846 | ATG31 | IV | 488072 | 488662 | -1 |
| YDR023W | 61.2264491 | -2.8698694 | 0.81082727 | 11.7073223 | 0.00286937 | 0.01473523 | SES1 | IV | 489508 | 490896 | 1 |
| YDR025W | 127.69053 | -2.9229708 | 0.52772018 | 29.6395354 | 3.66E-07 | 7.34E-06 | RPS11B | IV | 491515 | 492324 | 1 |
| YDR026C | 43.7395144 | 0.98319541 | 0.60938546 | 2.63305702 | 0.26806427 | 0.44106057 | NSI1 | IV | 492556 | 494268 | -1 |
| YDR027C | 117.341502 | 2.62997967 | 0.52360886 | 29.518613 | 3.89E-07 | 7.74E-06 | VPS54 | IV | 494649 | 497318 | -1 |
| YDR028C | 50.2984055 | 0.46222639 | 0.63115635 | 0.66745648 | 0.7162484 | 0.82794286 | REG1 | IV | 497835 | 500879 | -1 |
| YDR029W | 1893.6479 | 0.69138448 | 0.72792851 | 2.21866909 | 0.32977834 | 0.50287506 | NA | IV | 501100 | 501414 | 1 |
| YDR030C | 95.4792419 | -0.575168 | 0.5900035 | 1.08193811 | 0.58218381 | 0.73072713 | RAD28 | IV | 501753 | 503273 | -1 |
| YDR031W | 12.5753819 | 0.90868667 | 1.16820344 | 3.00559681 | 0.22250662 | 0.38746266 | MIX14 | IV | 503498 | 503863 | 1 |
| YDR032C | 24.8086609 | -0.0031989 | 0.67573983 | 0.29555329 | 0.86262376 | 0.91947562 | PST2 | IV | 504099 | 504695 | -1 |
| YDR033W | 51.7592807 | -3.1331424 | 0.87804356 | 11.8347448 | 0.00269227 | 0.01402028 | MRH1 | IV | 508147 | 509109 | 1 |
| YDR034C | 30.2079358 | 0.67735372 | 0.66929346 | 1.79269018 | 0.40805835 | 0.5793986 | LYS14 | IV | 509737 | 512109 | -1 |
| YDR034C-A | 16.4907013 | -0.0294036 | 0.77693005 | 0.29046346 | 0.86482186 | 0.92054956 | NA | IV | 520516 | 520692 | -1 |
| YDR034C-D | 18.4310403 | 0.20226169 | 0.79616341 | 0.73705841 | 0.69175101 | 0.81258639 | NA | IV | 514043 | 519356 | -1 |
| YDR034W-B | 109.111524 | 5.06854589 | 0.70617743 | 47.8919546 | 3.98E-11 | 2.01E-09 | NA | IV | 521314 | 521469 | 1 |
| YDR035W | 56.8585256 | 0.10061368 | 0.71311052 | 1.74830485 | 0.41721549 | 0.58892811 | ARO3 | IV | 521816 | 522928 | 1 |
| YDR036C | 65.5304343 | 1.37460192 | 0.61075863 | 7.32875046 | 0.02562017 | 0.07911509 | EHD3 | IV | 523211 | 524713 | -1 |

| | | | | | | | | | | | |
|---|---|---|---|---|---|---|---|---|---|---|---|
| YDR037W | 176.498124 | 1.12063798 | 0.50303206 | 5.72988709 | 0.05698635 | 0.14462713 | KRS1 | IV | 525440 | 527215 | 1 |
| YDR038C | 30.5074814 | 0.53377268 | 0.8522918 | 0.65978077 | 0.71900254 | 0.82938751 | ENA5 | IV | 527422 | 530697 | -1 |
| YDR040C | 159.590206 | -0.4478044 | 0.49409488 | 2.77316821 | 0.24992757 | 0.42103016 | ENA1 | IV | 535192 | 538467 | -1 |
| YDR041W | 6.63512044 | -0.6721389 | 1.11960488 | 0.81738544 | 0.66451839 | 0.79148192 | RSM10 | IV | 539803 | 540414 | 1 |
| YDR042C | 15.624754 | -1.8848662 | 0.84876696 | 5.96967539 | 0.05054771 | 0.13171522 | NA | IV | 540601 | 541203 | -1 |
| YDR043C | 113.452986 | -2.0679095 | 0.64886529 | 9.70558566 | 0.00780654 | 0.03254132 | NRG1 | IV | 542674 | 543369 | -1 |
| YDR044W | 7.78512596 | -1.1811818 | 1.11506492 | 1.36706083 | 0.50483158 | 0.66295642 | HEM13 | IV | 546642 | 547628 | 1 |
| YDR046C | 29.9593812 | -1.1329685 | 0.90182385 | 1.61080507 | 0.446908 | 0.61424863 | BAP3 | IV | 548762 | 550576 | -1 |
| YDR047W | 21.6630539 | -0.9910707 | 0.75965637 | 2.43986552 | 0.29525002 | 0.46951321 | HEM12 | IV | 551860 | 552948 | 1 |
| YDR049W | 39.6918006 | 0.4594605 | 0.63121482 | 1.93466611 | 0.38009538 | 0.55392707 | VMS1 | IV | 553254 | 555152 | 1 |
| YDR050C | 233.848319 | -0.0098328 | 0.53628596 | 0.84035497 | 0.65693022 | 0.78549013 | TPI1 | IV | 555726 | 556472 | -1 |
| YDR051C | 45.5405742 | 0.59809172 | 0.59912325 | 1.0189341 | 0.6008157 | 0.74682758 | DET1 | IV | 557056 | 558060 | -1 |
| YDR052C | 145.514307 | 0.74429583 | 0.50987924 | 11.804439 | 0.00273337 | 0.01414318 | DBF4 | IV | 558511 | 560625 | -1 |
| YDR054C | 61.2418144 | 0.47537455 | 0.66895628 | 1.30469522 | 0.52082166 | 0.6765916 | CDC34 | IV | 561440 | 562327 | -1 |
| YDR055W | 181.28796 | 1.29535891 | 0.55917244 | 5.90697482 | 0.05215749 | 0.13493829 | PST1 | IV | 563527 | 564861 | 1 |
| YDR056C | 322.709443 | 2.49777503 | 0.79253465 | 17.9117435 | 0.00012898 | 0.00116502 | EMC10 | IV | 565023 | 565640 | -1 |
| YDR057W | 200.38693 | 1.06195926 | 0.51440038 | 8.4242075 | 0.01481517 | 0.05295051 | YOS9 | IV | 565927 | 567555 | 1 |
| YDR058C | 156.037859 | 1.20277287 | 0.50380188 | 7.56481642 | 0.0227678 | 0.07300493 | TGL2 | IV | 567723 | 568703 | -1 |
| YDR059C | 123.807223 | 1.08619508 | 0.58700835 | 9.53192392 | 0.00851469 | 0.03484104 | UBC5 | IV | 569234 | 569770 | -1 |
| YDR060W | 123.641223 | -2.53233 | 0.53449231 | 20.7126491 | 3.18E-05 | 0.00035263 | MAK21 | IV | 570649 | 573726 | 1 |
| YDR061W | 507.113169 | 2.00948863 | 0.46419794 | 17.69195 | 0.00014396 | 0.0012883 | NA | IV | 574164 | 575783 | 1 |
| YDR062W | 45.0801462 | -0.1708259 | 0.62662117 | 0.88114944 | 0.64366639 | 0.77686719 | LCB2 | IV | 576474 | 578159 | 1 |
| YDR063W | 74.6079746 | -1.5487066 | 0.66987018 | 5.68905634 | 0.0581617 | 0.14654319 | AIM7 | IV | 578663 | 579112 | 1 |
| YDR064W | 84.047414 | -2.0801628 | 0.80532113 | 6.34397828 | 0.04192013 | 0.11497752 | RPS13 | IV | 579458 | 580452 | 1 |
| YDR065W | 115.324409 | -0.4578215 | 0.53183498 | 1.20953183 | 0.54620227 | 0.69889749 | RRG1 | IV | 580687 | 581784 | 1 |
| YDR066C | 485.520996 | -1.036918 | 0.4576541 | 5.41797947 | 0.06660406 | 0.16194348 | RTR2 | IV | 581908 | 582498 | -1 |
| YDR067C | 130.671723 | -0.8055092 | 0.52136701 | 2.36834255 | 0.30599966 | 0.47867401 | OCA6 | IV | 582791 | 583465 | -1 |
| YDR068W | 429.516004 | 3.27787296 | 0.58022802 | 31.7210709 | 1.29E-07 | 3.01E-06 | DOS2 | IV | 583713 | 584645 | 1 |
| YDR069C | 1196.75898 | 4.07788546 | 0.49557998 | 88.2945526 | 6.72E-20 | 1.94E-17 | DOA4 | IV | 584939 | 587719 | -1 |
| YDR070C | 14.7327807 | 0.5994147 | 0.81318442 | 0.7436105 | 0.68948851 | 0.81075111 | FMP16 | IV | 588098 | 588379 | -1 |
| YDR071C | 17.7436025 | 0.45324817 | 0.9754325 | 0.46885132 | 0.79102504 | 0.87338577 | PAA1 | IV | 588827 | 589402 | -1 |
| YDR072C | 184.412977 | 0.46351993 | 0.51155946 | 5.88544646 | 0.05272196 | 0.13597334 | IPT1 | IV | 589761 | 591344 | -1 |
| YDR073W | 60.8939636 | -0.1981738 | 0.57953675 | 0.14914115 | 0.92814197 | 0.95498585 | SNF11 | IV | 592439 | 592948 | 1 |

| | | | | | | | | | | | |
|---|---|---|---|---|---|---|---|---|---|---|---|
| YDR074W | 54.6658773 | 0.62614582 | 0.5834597 | 1.85920006 | 0.39471155 | 0.56638907 | TPS2 | IV | 593893 | 596583 | 1 |
| YDR075W | 14.9661078 | 1.2436311 | 0.84857865 | 4.59002132 | 0.10076032 | 0.21998577 | PPH3 | IV | 597156 | 598082 | 1 |
| YDR076W | 383.57142 | 0.19344703 | 0.49455018 | 0.26580748 | 0.87554937 | 0.92710879 | RAD55 | IV | 598468 | 599688 | 1 |
| YDR077W | 4011.61506 | 2.04978669 | 0.69795499 | 23.9823169 | 6.20E-06 | 8.57E-05 | SED1 | IV | 600793 | 601809 | 1 |
| YDR078C | 375.786414 | 2.64715402 | 0.51346666 | 23.9184311 | 6.40E-06 | 8.82E-05 | SHU2 | IV | 602198 | 602869 | -1 |
| YDR079C-A | 86.9004218 | -0.1124573 | 0.66832166 | 4.33476063 | 0.11447712 | 0.24134059 | TFB5 | IV | 603593 | 603811 | -1 |
| YDR079W | 172.618252 | -0.4087009 | 0.75565752 | 2.94122851 | 0.2297843 | 0.3965577 | PET100 | IV | 603064 | 603399 | 1 |
| YDR080W | 165.199812 | 2.63265778 | 0.54204875 | 28.8028744 | 5.57E-07 | 1.06E-05 | VPS41 | IV | 604008 | 606986 | 1 |
| YDR081C | 95.6221685 | -0.9964473 | 0.51515341 | 3.78960978 | 0.15034767 | 0.2931317 | PDC2 | IV | 607304 | 610081 | -1 |
| YDR082W | 7.17947002 | -0.7458281 | 1.08501825 | 1.31622018 | 0.51782906 | 0.67436642 | STN1 | IV | 610441 | 611925 | 1 |
| YDR083W | 344.192033 | -0.7812577 | 0.8711367 | 10.8234009 | 0.00446404 | 0.02081063 | RRP8 | IV | 612073 | 613251 | 1 |
| YDR084C | 14.9287769 | -0.0171031 | 0.77313487 | 2.42850711 | 0.29693157 | 0.47089395 | TVP23 | IV | 613405 | 614004 | -1 |
| YDR085C | 120.213236 | -1.6712418 | 0.52955913 | 11.0084975 | 0.00406944 | 0.0194889 | AFR1 | IV | 614288 | 616150 | -1 |
| YDR086C | 52.1381611 | -0.8328918 | 0.64899505 | 3.40512837 | 0.18221569 | 0.33728544 | SSS1 | IV | 616928 | 617170 | -1 |
| YDR087C | 689.586689 | 0.73963826 | 0.46283801 | 2.524938 | 0.28295455 | 0.45678444 | RRP1 | IV | 617470 | 618306 | -1 |
| YDR088C | 18.1348426 | 0.46284838 | 0.77199918 | 0.51180296 | 0.77421824 | 0.86282905 | SLU7 | IV | 618497 | 619645 | -1 |
| YDR089W | 49.5810619 | -0.9757999 | 0.85001355 | 6.06972978 | 0.04808116 | 0.12677136 | VTC5 | IV | 622112 | 624721 | 1 |
| YDR090C | 16.4871873 | -0.1772817 | 0.74908975 | 1.62525609 | 0.44369049 | 0.61162622 | NA | IV | 625066 | 625998 | -1 |
| YDR091C | 21.1344131 | -2.4699125 | 0.80115646 | 9.25412191 | 0.00978347 | 0.03871927 | RLI1 | IV | 626708 | 628534 | -1 |
| YDR092W | 62.8639181 | 0.07227897 | 0.59250223 | 1.46436763 | 0.48085774 | 0.64344033 | UBC13 | IV | 629876 | 630605 | 1 |
| YDR093W | 58.0365446 | -0.3468097 | 0.62827046 | 0.62453715 | 0.73178496 | 0.83702784 | DNF2 | IV | 631282 | 636120 | 1 |
| YDR096W | 11.5915247 | 0.74948374 | 0.85449143 | 2.16857805 | 0.33814211 | 0.51131962 | GIS1 | IV | 637139 | 639823 | 1 |
| YDR097C | 78.2613239 | 0.64221753 | 0.55883479 | 7.21747265 | 0.02708605 | 0.08236778 | MSH6 | IV | 640109 | 643837 | -1 |
| YDR098C | 51.4113358 | -1.0181287 | 0.59661563 | 3.12438324 | 0.20967604 | 0.37244958 | GRX3 | IV | 644178 | 644930 | -1 |
| YDR099W | 320.56756 | -0.0891586 | 0.50489847 | 3.09334732 | 0.21295516 | 0.37660671 | BMH2 | IV | 653607 | 654428 | 1 |
| YDR100W | 20.9524094 | 2.91506597 | 0.88482074 | 14.5726923 | 0.00068483 | 0.00479473 | TVP15 | IV | 655013 | 655444 | 1 |
| YDR101C | 153.510785 | -0.2126681 | 0.91238329 | 7.75346295 | 0.02071843 | 0.06786006 | ARX1 | IV | 655686 | 657467 | -1 |
| YDR102C | 81.189338 | 0.11945361 | 0.5800961 | 0.40768488 | 0.81559086 | 0.88902268 | NA | IV | 657648 | 657980 | -1 |
| YDR103W | 118.123307 | 1.07455068 | 0.56149743 | 4.08273379 | 0.1298511 | 0.26426638 | STE5 | IV | 658350 | 661103 | 1 |
| YDR104C | 88.2742595 | -0.7796883 | 0.63680757 | 8.08686749 | 0.01753715 | 0.05999792 | SPO71 | IV | 661173 | 664910 | -1 |
| YDR105C | 24.9634546 | 0.53738566 | 0.98179652 | 0.34255734 | 0.84258674 | 0.90781117 | TMS1 | IV | 665349 | 666770 | -1 |
| YDR106W | 240.63464 | 4.44591266 | 0.55366499 | 57.8340942 | 2.76E-13 | 2.12E-11 | ARP10 | IV | 667004 | 667858 | 1 |
| YDR107C | 1614.44327 | 0.40842934 | 0.5186603 | 0.80789236 | 0.66768005 | 0.79413877 | TMN2 | IV | 669016 | 671034 | -1 |

| | | | | | | | | | | | |
|---|---|---|---|---|---|---|---|---|---|---|---|
| YDR108W | 18.2758493 | 0.85305324 | 0.95794104 | 1.58190699 | 0.45341226 | 0.61945658 | TRS85 | IV | 671269 | 673365 | 1 |
| YDR109C | 30.5884662 | 1.02752073 | 0.65280863 | 2.65584878 | 0.26502678 | 0.43735364 | NA | IV | 673520 | 675667 | -1 |
| YDR110W | 140.486339 | -2.5048195 | 0.5285574 | 20.636973 | 3.30E-05 | 0.00036413 | FOB1 | IV | 676102 | 677802 | 1 |
| YDR111C | 15.9404174 | 1.23322937 | 0.87538193 | 4.11647104 | 0.12767906 | 0.26140797 | ALT2 | IV | 678241 | 679764 | -1 |
| YDR113C | 48.5576662 | -0.2267688 | 0.65239239 | 0.40934844 | 0.81491276 | 0.88874456 | PDS1 | IV | 680496 | 681617 | -1 |
| YDR114C | 855.255851 | -1.5827183 | 0.57902652 | 7.3808214 | 0.02496175 | 0.07778715 | NA | IV | 681924 | 682226 | -1 |
| YDR115W | 20.3368428 | 0.81127039 | 0.7476266 | 2.7956937 | 0.2471285 | 0.41801753 | MRX14 | IV | 682175 | 682492 | 1 |
| YDR116C | 35.9960642 | -0.0689029 | 0.78000936 | 3.09919217 | 0.21233372 | 0.37596705 | MRPL1 | IV | 682724 | 683581 | -1 |
| YDR117C | 71.0358177 | 0.42916809 | 0.64289262 | 1.6846065 | 0.43071733 | 0.59948398 | TMA64 | IV | 683946 | 685643 | -1 |
| YDR118W | 86.0469934 | -0.1506562 | 0.52939652 | 1.133303 | 0.56742228 | 0.71800411 | APC4 | IV | 685882 | 687840 | 1 |
| YDR119W | 67.7587896 | 0.55778216 | 0.55021333 | 2.40936271 | 0.29978751 | 0.47321965 | VBA4 | IV | 688227 | 690533 | 1 |
| YDR120C | 23.9164727 | -1.9763244 | 0.80369285 | 6.33132043 | 0.04218628 | 0.115598 | TRM1 | IV | 691549 | 693261 | -1 |
| YDR121W | 31.0915482 | -1.680666 | 0.69001063 | 6.28278796 | 0.0432225 | 0.11765788 | DPB4 | IV | 693585 | 694175 | 1 |
| YDR122W | 66.2273539 | 0.93859441 | 0.64496596 | 3.61957179 | 0.16368918 | 0.3123054 | KIN1 | IV | 694700 | 697894 | 1 |
| YDR123C | 30.4482855 | -0.0740286 | 0.63846251 | 0.13697812 | 0.93380368 | 0.95735444 | INO2 | IV | 698554 | 699468 | -1 |
| YDR124W | 1222.34168 | 0.69804521 | 0.47675631 | 2.17510193 | 0.33704091 | 0.51030806 | NA | IV | 700315 | 701289 | 1 |
| YDR125C | 144.436664 | -0.1094051 | 0.49322965 | 0.76804855 | 0.68111489 | 0.80384598 | ECM18 | IV | 701397 | 702758 | -1 |
| YDR126W | 223.659643 | 2.63277737 | 0.55890109 | 20.1491385 | 4.21E-05 | 0.00044684 | SWF1 | IV | 703234 | 704244 | 1 |
| YDR127W | 46.7258556 | 0.41027564 | 0.66280648 | 0.3971078 | 0.81991558 | 0.89184881 | ARO1 | IV | 704484 | 709250 | 1 |
| YDR128W | 20.2424975 | 0.08971776 | 0.82932593 | 0.41565243 | 0.8123482 | 0.88731158 | MTC5 | IV | 709549 | 712995 | 1 |
| YDR129C | 56.579635 | -0.3252642 | 0.688033 | 0.94728433 | 0.62273005 | 0.76281637 | SAC6 | IV | 713340 | 715379 | -1 |
| YDR130C | 13.5154257 | -0.778985 | 0.84742837 | 1.00782825 | 0.60416126 | 0.7493774 | FIN1 | IV | 715747 | 716622 | -1 |
| YDR131C | 15.2432011 | -0.9830714 | 0.7739093 | 4.00122668 | 0.1352523 | 0.27200793 | NA | IV | 716790 | 718460 | -1 |
| YDR132C | 462.718116 | 1.54089236 | 0.52556501 | 11.1867927 | 0.00372236 | 0.01814182 | NA | IV | 718816 | 720303 | -1 |
| YDR134C | 174.515834 | 1.1031284 | 0.48789598 | 9.73740736 | 0.00768332 | 0.03213985 | NA | NA | NA | NA | NA |
| YDR135C | 163.041802 | -0.6889363 | 0.57790164 | 2.30453266 | 0.31591998 | 0.4883013 | YCF1 | IV | 723004 | 727551 | -1 |
| YDR137W | 22.4751813 | 0.43200602 | 0.7111561 | 0.58242099 | 0.74735834 | 0.84631426 | RGP1 | IV | 728259 | 730250 | 1 |
| YDR138W | 82.0784747 | 2.67033728 | 0.59903799 | 22.3857789 | 1.38E-05 | 0.00017372 | HPR1 | IV | 730578 | 732836 | 1 |
| YDR139C | 75.7609835 | -0.1254464 | 0.56922014 | 0.46738339 | 0.79160584 | 0.87352731 | RUB1 | IV | 733618 | 733924 | -1 |
| YDR140W | 392.033508 | -0.8275934 | 0.52348644 | 6.8359705 | 0.03277841 | 0.09561057 | MTQ2 | IV | 734138 | 734803 | 1 |
| YDR141C | 206.551882 | -1.0721921 | 0.56053425 | 3.75248473 | 0.15316456 | 0.29709307 | DOP1 | IV | 734901 | 739997 | -1 |
| YDR142C | 31.6201435 | 2.7162236 | 0.7077876 | 17.2543314 | 0.00017917 | 0.00154376 | PEX7 | IV | 740473 | 741600 | -1 |
| YDR143C | 404.874966 | 3.90912844 | 0.50384139 | 59.3723446 | 1.28E-13 | 1.03E-11 | SAN1 | IV | 742042 | 743874 | -1 |

| | | | | | | | | | | | |
|---|---|---|---|---|---|---|---|---|---|---|---|
| YDR144C | 57.666145 | 1.50967217 | 0.60911886 | 9.72786244 | 0.00772007 | 0.03222728 | MKC7 | IV | 744311 | 746101 | -1 |
| YDR145W | 69.2029981 | -0.7009041 | 0.58366423 | 3.35538232 | 0.18680478 | 0.34260364 | TAF12 | IV | 746738 | 748357 | 1 |
| YDR146C | 18.4190428 | -1.2501886 | 0.7337503 | 2.96741511 | 0.22679527 | 0.39280426 | SWI5 | IV | 748613 | 750742 | -1 |
| YDR147W | 11.458364 | -0.7786021 | 0.85764323 | 1.18799026 | 0.55211709 | 0.70397665 | EKI1 | IV | 751631 | 753235 | 1 |
| YDR148C | 32.3737561 | 0.41576656 | 0.6591507 | 1.35617647 | 0.50758645 | 0.66521629 | KGD2 | IV | 753675 | 755066 | -1 |
| YDR150W | 247.537976 | 1.42346615 | 0.4748002 | 9.81161368 | 0.00740347 | 0.03135777 | NUM1 | IV | 755628 | 763874 | 1 |
| YDR151C | 100.331106 | 1.39351458 | 0.51564939 | 8.90024554 | 0.01167713 | 0.04427675 | CTH1 | IV | 764178 | 765155 | -1 |
| YDR152W | 42.9765074 | -1.2962137 | 0.61399265 | 4.43382043 | 0.10894521 | 0.23299267 | GIR2 | IV | 765706 | 766503 | 1 |
| YDR153C | 39.2711666 | -0.116451 | 0.6936044 | 3.93065961 | 0.14010967 | 0.27867477 | ENT5 | IV | 766736 | 767971 | -1 |
| YDR155C | 231.833547 | 0.38805875 | 0.61708914 | 2.41368688 | 0.29914004 | 0.47232639 | CPR1 | IV | 768512 | 769000 | -1 |
| YDR156W | 8.95649951 | -1.6776109 | 0.99154162 | 5.2708751 | 0.0716876 | 0.17187212 | RPA14 | IV | 769525 | 769938 | 1 |
| YDR157W | 52.3471625 | -5.6022437 | 0.75206008 | 56.9815644 | 4.23E-13 | 3.14E-11 | NA | IV | 769931 | 770332 | 1 |
| YDR158W | 39.4143757 | -1.3470659 | 0.68465266 | 4.83895454 | 0.08896811 | 0.20216851 | HOM2 | IV | 770357 | 771454 | 1 |
| YDR159W | 39.8862347 | -0.1713565 | 0.63454611 | 0.2941664 | 0.86322215 | 0.9197748 | SAC3 | IV | 771877 | 775782 | 1 |
| YDR160W | 89.7856824 | 2.33323749 | 0.5976941 | 28.0700886 | 8.03E-07 | 1.47E-05 | SSY1 | IV | 776163 | 778721 | 1 |
| YDR161W | 74.3538001 | 0.10573022 | 0.81451863 | 1.40147201 | 0.49621995 | 0.65603003 | ACL4 | IV | 779043 | 780206 | 1 |
| YDR162C | 41.7073421 | -0.3133586 | 0.64668029 | 0.64230884 | 0.72531124 | 0.83456178 | NBP2 | IV | 780390 | 781100 | -1 |
| YDR163W | 256.441841 | -0.3792458 | 0.62306479 | 6.74692686 | 0.03427074 | 0.09906519 | CWC15 | IV | 781423 | 781950 | 1 |
| YDR164C | 118.264862 | 0.50310649 | 0.82417613 | 9.08606531 | 0.01064109 | 0.04132253 | SEC1 | IV | 782041 | 784215 | -1 |
| YDR165W | 671.544708 | 0.8729288 | 0.51563707 | 2.96254207 | 0.22734854 | 0.39340945 | TRM82 | IV | 784871 | 786205 | 1 |
| YDR166C | 209.266491 | 1.04420796 | 0.49246177 | 6.86701921 | 0.03227347 | 0.09442315 | SEC5 | IV | 786306 | 789221 | -1 |
| YDR167W | 20.6193528 | -0.410403 | 0.74334658 | 0.95944131 | 0.61895627 | 0.76021421 | TAF10 | IV | 789449 | 790069 | 1 |
| YDR168W | 19.2559238 | 0.48755887 | 0.7694852 | 1.19170856 | 0.55109158 | 0.70297868 | CDC37 | IV | 790328 | 791848 | 1 |
| YDR169C | 28.0530495 | 1.42473994 | 0.69458974 | 4.73924923 | 0.09351582 | 0.20881474 | STB3 | IV | 792348 | 793889 | -1 |
| YDR169C-A | 7.08680967 | -0.7623129 | 1.03449572 | 3.10862851 | 0.21133425 | 0.37477039 | NA | IV | 794574 | 794723 | -1 |
| YDR170C | 163.497298 | -0.5524323 | 0.64510963 | 7.85518129 | 0.01969106 | 0.06517227 | SEC7 | IV | 796193 | 802222 | -1 |
| YDR170W-A | 918.144124 | 0.32115753 | 0.48105067 | 1.47631709 | 0.47799331 | 0.64064381 | NA | IV | 803195 | 804517 | 1 |
| YDR171W | 169.589819 | 1.50108583 | 0.50520461 | 13.504056 | 0.00116851 | 0.00748414 | HSP42 | IV | 806621 | 807748 | 1 |
| YDR172W | 129.480321 | -2.0673843 | 0.57174493 | 13.1104212 | 0.00142268 | 0.00879118 | SUP35 | IV | 808324 | 810381 | 1 |
| YDR173C | 27.2184814 | 1.05836055 | 0.71379044 | 2.98985778 | 0.22426455 | 0.38976222 | ARG82 | IV | 810565 | 811632 | -1 |
| YDR174W | 51.3402329 | -1.6523233 | 0.59370774 | 7.96751118 | 0.0186156 | 0.0626889 | HMO1 | IV | 812110 | 812850 | 1 |
| YDR175C | 10.2734478 | -0.7193958 | 0.97105719 | 1.17703995 | 0.55514831 | 0.70690757 | RSM24 | IV | 813193 | 814152 | -1 |
| YDR176W | 73.8956684 | -0.7260458 | 0.5448116 | 2.95682207 | 0.22799968 | 0.39406513 | NGG1 | IV | 814452 | 816560 | 1 |

| | | | | | | | | | | | |
|---|---|---|---|---|---|---|---|---|---|---|---|
| YDR177W | 25.1545954 | -0.1856559 | 0.70116534 | 1.46972417 | 0.47957159 | 0.64246171 | UBC1 | IV | 816878 | 817525 | 1 |
| YDR178W | 23.9402945 | 2.40890826 | 0.84061027 | 7.52274145 | 0.02325185 | 0.07417531 | SDH4 | IV | 817950 | 818495 | 1 |
| YDR179C | 27.1226505 | 0.59740234 | 1.02832361 | 5.50451177 | 0.06378381 | 0.15701882 | CSN9 | IV | 818708 | 819196 | -1 |
| YDR179W-A | 133.515016 | 0.76651371 | 0.50560822 | 5.1566317 | 0.07590173 | 0.17886487 | NVJ3 | IV | 819433 | 820824 | 1 |
| YDR180W | 98.4599664 | -0.4244822 | 0.74716036 | 3.69197671 | 0.15786921 | 0.30407942 | SCC2 | IV | 821295 | 825776 | 1 |
| YDR181C | 11.7639461 | -0.1016009 | 0.89126403 | 0.48462398 | 0.78481129 | 0.86984253 | SAS4 | IV | 825910 | 827355 | -1 |
| YDR182W | 22.9288493 | 0.26747858 | 0.71068799 | 0.71080371 | 0.70089173 | 0.81801018 | CDC1 | IV | 827582 | 829057 | 1 |
| YDR182W-A | 47.4792837 | 1.34308691 | 0.62553184 | 4.60358955 | 0.10007906 | 0.21919967 | NA | IV | 829151 | 829354 | 1 |
| YDR183W | 11.1765531 | -0.015434 | 0.87903204 | 0.27660793 | 0.87083395 | 0.92499148 | PLP1 | IV | 829585 | 830277 | 1 |
| YDR184C | 130.990693 | 4.41763582 | 0.68220401 | 38.9758656 | 3.44E-09 | 1.14E-07 | ATC1 | IV | 830629 | 831513 | -1 |
| YDR185C | 415.503033 | -0.3503665 | 0.46804262 | 8.01402898 | 0.01818761 | 0.06161866 | UPS3 | IV | 831934 | 832473 | -1 |
| YDR186C | 1042.74668 | 5.19648028 | 0.50907483 | 78.6749297 | 8.24E-18 | 1.36E-15 | SND1 | IV | 832859 | 835492 | -1 |
| YDR188W | 33.2863256 | -0.8437664 | 0.84427239 | 1.05782517 | 0.58924538 | 0.73703407 | CCT6 | IV | 836421 | 838061 | 1 |
| YDR189W | 70.9667149 | -0.0498949 | 0.56741506 | 0.63990484 | 0.72618359 | 0.83474349 | SLY1 | IV | 838392 | 840392 | 1 |
| YDR190C | 23.6238727 | -0.7780343 | 0.82853305 | 1.20341148 | 0.5478763 | 0.70011119 | RVB1 | IV | 840604 | 841995 | -1 |
| YDR191W | 8.58172516 | -1.1656976 | 0.97012701 | 1.6251901 | 0.44370513 | 0.61162622 | HST4 | IV | 842337 | 843449 | 1 |
| YDR192C | 20.1133287 | 1.85819573 | 0.7920978 | 5.89828643 | 0.05238457 | 0.13534434 | NUP42 | IV | 843569 | 844861 | -1 |
| YDR194C | 26.2258674 | -0.955672 | 0.71585683 | 1.90229913 | 0.38629669 | 0.55874541 | MSS116 | IV | 845952 | 847946 | -1 |
| YDR195W | 241.801929 | 1.29326598 | 0.61362051 | 6.08512538 | 0.04771246 | 0.12600836 | REF2 | IV | 848599 | 850200 | 1 |
| YDR196C | 70.5363179 | 0.44591454 | 0.91557967 | 6.69166631 | 0.03523085 | 0.10098348 | CAB5 | IV | 850274 | 850999 | -1 |
| YDR197W | 51.6223186 | -1.3005948 | 0.65638346 | 3.98873528 | 0.13609969 | 0.27319883 | CBS2 | IV | 851228 | 852397 | 1 |
| YDR198C | 23.9307583 | -0.2377448 | 0.68769417 | 0.17749692 | 0.91507572 | 0.94797312 | RKM2 | IV | 852533 | 853972 | -1 |
| YDR200C | 32.5657853 | 0.8968976 | 0.71390454 | 1.7472069 | 0.4174446 | 0.58908219 | VPS64 | IV | 854279 | 856093 | -1 |
| YDR201W | 16.2378653 | 0.36414204 | 0.81023242 | 2.61987911 | 0.26983637 | 0.44309488 | SPC19 | IV | 856317 | 856814 | 1 |
| YDR202C | 76.4138293 | -1.4622913 | 0.62976016 | 7.0370411 | 0.02964326 | 0.0881533 | RAV2 | IV | 856898 | 857953 | -1 |
| YDR204W | 29.6644703 | -1.1760841 | 0.65622855 | 3.18359577 | 0.20355931 | 0.36388033 | COQ4 | IV | 858137 | 859144 | 1 |
| YDR205W | 14.746364 | -0.6205642 | 0.79382095 | 0.62506222 | 0.73159287 | 0.83702784 | MSC2 | IV | 859346 | 861520 | 1 |
| YDR206W | 22.5441119 | -1.1842528 | 0.84737051 | 2.17039316 | 0.33783537 | 0.51112275 | EBS1 | IV | 862054 | 864708 | 1 |
| YDR207C | 17.2944569 | -1.0756605 | 0.76269415 | 2.01599692 | 0.36494871 | 0.53808328 | UME6 | IV | 865012 | 867522 | -1 |
| YDR208W | 28.5280377 | 0.56940268 | 0.77002691 | 1.04368863 | 0.59342507 | 0.74066204 | MSS4 | IV | 868224 | 870563 | 1 |
| YDR209C | 21534.4233 | 0.79689977 | 0.45079842 | 19.9487156 | 4.66E-05 | 0.00048506 | NA | IV | 871036 | 871449 | -1 |
| YDR210W | 115.049412 | 1.31903414 | 0.61758261 | 10.0990169 | 0.00641248 | 0.02789503 | NA | IV | 871074 | 871301 | 1 |
| YDR211W | 75.9449653 | 0.94697332 | 0.64670141 | 2.20784697 | 0.33156763 | 0.50507144 | GCD6 | IV | 884727 | 886865 | 1 |

| | | | | | | | | | | | |
|---|---|---|---|---|---|---|---|---|---|---|---|
| YDR212W | 74.9226699 | 1.89561494 | 0.60115414 | 12.0888096 | 0.00237109 | 0.01271169 | TCP1 | IV | 887232 | 888911 | 1 |
| YDR213W | 37.1720412 | 0.10446081 | 0.64943515 | 1.24180852 | 0.53745822 | 0.69208525 | UPC2 | IV | 889751 | 892492 | 1 |
| YDR214W | 199.00934 | 0.35526857 | 0.50433126 | 2.1072109 | 0.34867834 | 0.52247608 | AHA1 | IV | 892875 | 893927 | 1 |
| YDR215C | 160.393793 | -2.7622574 | 0.61462184 | 21.4981921 | 2.15E-05 | 0.00025209 | NA | IV | 894089 | 894502 | -1 |
| YDR216W | 11.2443191 | 2.40182724 | 1.196625 | 4.10254599 | 0.12857113 | 0.2622576 | ADR1 | IV | 895035 | 899006 | 1 |
| YDR217C | 143.026564 | 2.54047368 | 0.60170597 | 16.4448897 | 0.00026856 | 0.00221819 | RAD9 | IV | 899551 | 903480 | -1 |
| YDR218C | 402.426958 | 3.30689002 | 0.50769396 | 43.0114939 | 4.57E-10 | 1.79E-08 | SPR28 | IV | 903781 | 905052 | -1 |
| YDR219C | 9.9419742 | -0.0391922 | 0.97117376 | 0.31007274 | 0.85638403 | 0.91501449 | MFB1 | IV | 905455 | 906852 | -1 |
| YDR221W | 286.615605 | 0.08798164 | 0.46957153 | 2.87956643 | 0.23697913 | 0.40475196 | GTB1 | IV | 907330 | 909438 | 1 |
| YDR222W | 1023.55115 | 2.1068133 | 0.54449482 | 14.5223682 | 0.00070228 | 0.00488723 | NA | IV | 910054 | 911301 | 1 |
| YDR223W | 4141.35204 | 2.59932304 | 0.4662873 | 27.7842189 | 9.26E-07 | 1.66E-05 | CRF1 | IV | 912099 | 913502 | 1 |
| YDR224C | 289.657422 | -2.6010927 | 0.58851866 | 17.5440439 | 0.00015501 | 0.00137233 | HTB1 | IV | 914317 | 914712 | -1 |
| YDR225W | 20.8914668 | -2.7564523 | 0.81575242 | 12.1252492 | 0.00232828 | 0.01250534 | HTA1 | IV | 915530 | 915928 | 1 |
| YDR226W | 65.4540037 | 0.11696258 | 0.79183478 | 0.8089624 | 0.66732293 | 0.79387708 | ADK1 | IV | 916486 | 917154 | 1 |
| YDR227W | 106.168682 | 2.23045019 | 0.58217613 | 15.2048472 | 0.00049924 | 0.00367131 | SIR4 | IV | 917571 | 921647 | 1 |
| YDR228C | 260.590408 | 3.96787166 | 0.51144361 | 74.4133837 | 6.94E-17 | 9.80E-15 | PCF11 | IV | 921926 | 923806 | -1 |
| YDR229W | 104.322787 | 1.41120813 | 0.68980884 | 9.76545018 | 0.00757634 | 0.03185694 | IVY1 | IV | 924785 | 926146 | 1 |
| YDR231C | 72.1422107 | 0.68312833 | 0.65890674 | 1.6103937 | 0.44699993 | 0.61424863 | COX20 | IV | 926293 | 926910 | -1 |
| YDR232W | 86.6431963 | 1.30139519 | 0.60313022 | 12.8501646 | 0.0016204 | 0.00963256 | HEM1 | IV | 927452 | 929098 | 1 |
| YDR233C | 82.8030985 | -0.1581633 | 0.56643306 | 0.7073423 | 0.70210582 | 0.81847842 | RTN1 | IV | 929470 | 930357 | -1 |
| YDR234W | 38.2969896 | -1.3928023 | 0.61861246 | 5.08415031 | 0.07870291 | 0.18416483 | LYS4 | IV | 931129 | 933210 | 1 |
| YDR235W | 174.212077 | 1.26329927 | 0.7980876 | 7.79600825 | 0.02028235 | 0.06680024 | PRP42 | IV | 933504 | 935138 | 1 |
| YDR236C | 41.4089579 | -0.4125148 | 0.71291761 | 1.56170451 | 0.4580155 | 0.62324553 | FMN1 | IV | 935236 | 935892 | -1 |
| YDR237W | 99.1319367 | -0.4139763 | 0.51491386 | 2.4551356 | 0.29300436 | 0.46722534 | MRPL7 | IV | 936615 | 937493 | 1 |
| YDR238C | 76.8303337 | 0.36006145 | 0.62083035 | 0.57299331 | 0.75088959 | 0.84898472 | SEC26 | IV | 937895 | 940816 | -1 |
| YDR239C | 18.3557136 | 1.03740064 | 0.79932796 | 7.10001726 | 0.02872439 | 0.08612855 | NA | IV | 941057 | 943420 | -1 |
| YDR240C | 48.9199332 | -0.6489736 | 0.5793638 | 4.34647725 | 0.11380844 | 0.24049301 | SNU56 | IV | 943674 | 945152 | -1 |
| YDR242W | 959.288404 | -1.9009603 | 0.49084917 | 15.605735 | 0.00040856 | 0.00312906 | AMD2 | IV | 946807 | 948456 | 1 |
| YDR243C | 327.330388 | 0.98349206 | 0.47771132 | 5.4187025 | 0.06657999 | 0.16194348 | PRP28 | IV | 948518 | 950284 | -1 |
| YDR244W | 21.1605285 | 0.72264089 | 0.75654648 | 0.96868254 | 0.61610291 | 0.75801867 | PEX5 | IV | 950563 | 952401 | 1 |
| YDR245W | 19.9441949 | -1.628992 | 0.83040323 | 3.84986723 | 0.14588544 | 0.28719849 | MNN10 | IV | 952800 | 953981 | 1 |
| YDR246W | 158.584625 | 0.28806842 | 0.51114978 | 1.9380379 | 0.37945512 | 0.55341188 | TRS23 | IV | 954288 | 954947 | 1 |
| YDR246W-A | 42.4636538 | 1.54032598 | 0.64740084 | 6.37546075 | 0.04126542 | 0.11399179 | NA | IV | 955133 | 955333 | 1 |

| | | | | | | | | | | | |
|---|---|---|---|---|---|---|---|---|---|---|---|
| YDR247W | 25.3283139 | 1.45025892 | 0.72719625 | 4.01151696 | 0.1345582 | 0.27098851 | VHS1 | IV | 956013 | 957398 | 1 |
| YDR248C | 632.775493 | 5.27999563 | 0.55984523 | 71.8640032 | 2.48E-16 | 3.19E-14 | NA | IV | 957758 | 958339 | -1 |
| YDR249C | 21.1576263 | 2.04315854 | 0.89295003 | 8.44341366 | 0.01467358 | 0.05261147 | NA | IV | 958681 | 959802 | -1 |
| YDR251W | 89.2560196 | -0.8678009 | 0.54584633 | 3.96691666 | 0.13759257 | 0.27528023 | PAM1 | IV | 960614 | 963106 | 1 |
| YDR253C | 933.304701 | -1.6114072 | 0.47626062 | 11.1885498 | 0.0037191 | 0.01814182 | MET32 | IV | 963990 | 964565 | -1 |
| YDR254W | 2047.39653 | 1.5355098 | 0.55134057 | 10.5536113 | 0.00510872 | 0.0233461 | CHL4 | IV | 965113 | 966489 | 1 |
| YDR255C | 495.95119 | -1.2855067 | 0.47863444 | 7.83678321 | 0.01987303 | 0.06563882 | RMD5 | IV | 966557 | 967822 | -1 |
| YDR256C | 1450.88045 | -0.9939298 | 0.46909083 | 6.52337551 | 0.03832366 | 0.10782022 | CTA1 | IV | 968133 | 969680 | -1 |
| YDR257C | 19.6778904 | -0.9986234 | 0.77669778 | 1.80145989 | 0.406273 | 0.57768189 | RKM4 | IV | 969990 | 971474 | -1 |
| YDR258C | 38.8444024 | 1.84233258 | 0.7869422 | 5.68712867 | 0.05821779 | 0.14655695 | HSP78 | IV | 971808 | 974243 | -1 |
| YDR259C | 69.999617 | 2.78207899 | 0.69564789 | 17.9600487 | 0.0001259 | 0.00114078 | YAP6 | IV | 974631 | 975782 | -1 |
| YDR260C | 236.933996 | 0.79683392 | 0.54349107 | 2.6438084 | 0.26662711 | 0.43907024 | SWM1 | IV | 976717 | 977229 | -1 |
| YDR261C | 199.924536 | -0.2463865 | 0.53448531 | 1.74821757 | 0.4172337 | 0.58892811 | EXG2 | IV | 977521 | 979209 | -1 |
| YDR261C-D | 5.57146053 | 0.49238387 | 1.32570779 | 1.1200846 | 0.5711849 | 0.72145737 | NA | IV | 987534 | 992349 | -1 |
| YDR261W-B | 11.1163817 | -1.5203072 | 0.88047328 | 3.01405755 | 0.22156733 | 0.38636302 | NA | IV | 981462 | 986775 | 1 |
| YDR262W | 26.7728231 | 0.11243952 | 0.74077516 | 0.23970833 | 0.88704979 | 0.93215081 | NA | IV | 993134 | 993952 | 1 |
| YDR263C | 93.7610117 | 1.43581452 | 0.57162638 | 6.31531479 | 0.04252524 | 0.1160873 | DIN7 | IV | 994242 | 995534 | -1 |
| YDR264C | 29.9666969 | 0.21418783 | 0.72779244 | 0.14694035 | 0.92916386 | 0.95541731 | AKR1 | IV | 996029 | 998323 | -1 |
| YDR265W | 148.685147 | -1.5849464 | 0.54519239 | 8.21883514 | 0.01641733 | 0.05715957 | PEX10 | IV | 998864 | 999877 | 1 |
| YDR266C | 87.192382 | 2.45566317 | 0.69996618 | 20.1454156 | 4.22E-05 | 0.00044686 | HEL2 | IV | 1000104 | 1E+06 | -1 |
| YDR267C | 31.7372522 | -0.0180533 | 0.67149742 | 0.18541495 | 0.91146009 | 0.94542349 | CIA1 | IV | 1002510 | 1E+06 | -1 |
| YDR268W | 73.7785833 | -3.0456712 | 0.63488682 | 25.1866777 | 3.39E-06 | 5.12E-05 | MSW1 | IV | 1004004 | 1E+06 | 1 |
| YDR270W | 67.523294 | 0.24724988 | 0.54922833 | 0.20890536 | 0.90081742 | 0.93824416 | CCC2 | IV | 1005675 | 1E+06 | 1 |
| YDR272W | 48.9182148 | 0.8120432 | 0.58595715 | 1.9173079 | 0.38340863 | 0.55665395 | GLO2 | IV | 1009010 | 1E+06 | 1 |
| YDR273W | 24.3426172 | 1.58511289 | 0.6990331 | 5.67940687 | 0.058443 | 0.14688185 | DON1 | IV | 1010176 | 1E+06 | 1 |
| YDR274C | 590.626191 | 0.57260119 | 0.65239873 | 1.42061931 | 0.49149198 | 0.65269233 | NA | IV | 1011589 | 1E+06 | -1 |
| YDR275W | 69.8160607 | 3.39367746 | 0.67476651 | 24.9829297 | 3.76E-06 | 5.61E-05 | BSC2 | IV | 1012252 | 1E+06 | 1 |
| YDR276C | 89.7201399 | 0.86213932 | 0.51774678 | 3.77533544 | 0.15142456 | 0.29470528 | PMP3 | IV | 1013476 | 1E+06 | -1 |
| YDR277C | 217.368528 | 1.6221249 | 0.51520059 | 16.1100726 | 0.0003175 | 0.0025639 | MTH1 | IV | 1014401 | 1E+06 | -1 |
| YDR279W | 10.4680648 | 0.05724244 | 0.91466326 | 0.02213677 | 0.98899265 | 0.99104663 | RNH202 | IV | 1019368 | 1E+06 | 1 |
| YDR280W | 71.9917212 | 1.27703172 | 0.59663994 | 8.08347226 | 0.01756695 | 0.05999792 | RRP45 | IV | 1020747 | 1E+06 | 1 |
| YDR281C | 48.5108925 | -0.0225951 | 0.62506205 | 3.82689154 | 0.14757101 | 0.28982309 | PHM6 | IV | 1022007 | 1E+06 | -1 |
| YDR282C | 9.00741783 | 1.34778584 | 0.98928558 | 2.08729624 | 0.35216759 | 0.52579947 | MRX10 | IV | 1023511 | 1E+06 | -1 |

| | | | | | | | | | | | |
|---|---|---|---|---|---|---|---|---|---|---|---|
| YDR283C | 33.1679344 | 0.54326708 | 0.65008516 | 0.94303629 | 0.62405414 | 0.76326014 | GCN2 | IV | 1025070 | 1E+06 | -1 |
| YDR284C | 68.8294204 | -1.1582553 | 0.54486127 | 5.59257554 | 0.06103622 | 0.15186924 | DPP1 | IV | 1030550 | 1E+06 | -1 |
| YDR285W | 25.7633471 | -1.7827476 | 0.71163871 | 6.44687989 | 0.03981785 | 0.11116077 | ZIP1 | IV | 1032436 | 1E+06 | 1 |
| YDR286C | 12.6456038 | 0.72805611 | 0.97811554 | 1.84602896 | 0.39731952 | 0.56801976 | NA | IV | 1035231 | 1E+06 | -1 |
| YDR287W | 77.5386234 | 1.59754665 | 0.54044706 | 8.69255887 | 0.01295492 | 0.04783737 | INM2 | IV | 1035995 | 1E+06 | 1 |
| YDR288W | 29.5710637 | -1.9978413 | 0.75050487 | 7.54586275 | 0.02298459 | 0.07344413 | NSE3 | IV | 1037195 | 1E+06 | 1 |
| YDR289C | 37.5243325 | -0.3778093 | 0.62677104 | 2.07091124 | 0.35506457 | 0.52857757 | RTT103 | IV | 1038280 | 1E+06 | -1 |
| YDR291W | 84.7972404 | -1.1429232 | 0.59875995 | 4.13706735 | 0.12637095 | 0.25966788 | HRQ1 | IV | 1039728 | 1E+06 | 1 |
| YDR292C | 137.725502 | -0.5926142 | 0.89882431 | 4.73185478 | 0.09386221 | 0.20926539 | SRP101 | IV | 1043146 | 1E+06 | -1 |
| YDR293C | 85.0855518 | 0.56576093 | 0.56593888 | 2.42132041 | 0.29800047 | 0.47161883 | SSD1 | IV | 1045640 | 1E+06 | -1 |
| YDR294C | 146.333189 | 2.5246217 | 0.51265939 | 22.9195427 | 1.05E-05 | 0.00013875 | DPL1 | IV | 1050459 | 1E+06 | -1 |
| YDR295C | 68.0682711 | -0.0724683 | 0.65897312 | 0.27781197 | 0.87030985 | 0.92460441 | HDA2 | IV | 1052623 | 1E+06 | -1 |
| YDR296W | 15.2667537 | -0.8672942 | 0.77924689 | 2.31332395 | 0.31453435 | 0.48654927 | MHR1 | IV | 1055212 | 1E+06 | 1 |
| YDR297W | 10.3935206 | -0.7298245 | 1.00199981 | 0.78371017 | 0.67580204 | 0.80051019 | SUR2 | IV | 1056551 | 1E+06 | 1 |
| YDR298C | 35.8648729 | 2.62826934 | 0.68819979 | 21.2178026 | 2.47E-05 | 0.0002854 | ATP5 | IV | 1058176 | 1E+06 | -1 |
| YDR299W | 32.2785168 | -2.9647073 | 0.72291272 | 15.6124455 | 0.00040719 | 0.00312271 | BFR2 | IV | 1059627 | 1E+06 | 1 |
| YDR300C | 15.6738021 | -1.9850234 | 0.79606249 | 6.31541675 | 0.04252308 | 0.1160873 | PRO1 | IV | 1061505 | 1E+06 | -1 |
| YDR301W | 19.8011313 | -0.050098 | 0.76758092 | 0.12217525 | 0.9407408 | 0.96082012 | CFT1 | IV | 1063352 | 1E+06 | 1 |
| YDR302W | 23.8244965 | -0.2426127 | 0.72291172 | 0.33218833 | 0.84696648 | 0.90980423 | GPI11 | IV | 1067731 | 1E+06 | 1 |
| YDR303C | 17.6984554 | 0.07309302 | 0.74701314 | 0.43486476 | 0.80458201 | 0.88279205 | RSC3 | IV | 1068729 | 1E+06 | -1 |
| YDR304C | 147.111184 | 1.14715861 | 0.53303223 | 5.49582086 | 0.06406158 | 0.15736105 | CPR5 | IV | 1071880 | 1E+06 | -1 |
| YDR305C | 16.0774104 | 0.13639632 | 0.80736348 | 0.1081083 | 0.9473808 | 0.96420992 | NA | IV | 1072746 | 1E+06 | -1 |
| YDR306C | 25.4172551 | -0.3141979 | 0.6737071 | 0.54910614 | 0.75991168 | 0.85451323 | NA | IV | 1073735 | 1E+06 | -1 |
| YDR307W | 160.954554 | -1.913715 | 0.53113953 | 12.8552798 | 0.00161626 | 0.00961783 | PMT7 | IV | 1075865 | 1E+06 | 1 |
| YDR308C | 17.0599465 | -1.6971201 | 0.80141905 | 4.55951907 | 0.10230881 | 0.22277848 | SRB7 | IV | 1078027 | 1E+06 | -1 |
| YDR309C | 43.5667768 | 1.50134076 | 0.61459336 | 6.54633807 | 0.03788617 | 0.10674069 | GIC2 | IV | 1079048 | 1E+06 | -1 |
| YDR310C | 166.539173 | -0.649084 | 0.63292977 | 1.18117381 | 0.55400204 | 0.70568075 | SUM1 | IV | 1081128 | 1E+06 | -1 |
| YDR311W | 16.5381371 | 0.94097735 | 0.87513207 | 2.06736855 | 0.35569407 | 0.52929033 | TFB1 | IV | 1085065 | 1E+06 | 1 |
| YDR312W | 425.315029 | 4.62012008 | 0.54244438 | 70.2102156 | 5.68E-16 | 6.99E-14 | SSF2 | IV | 1087581 | 1E+06 | 1 |
| YDR313C | 8.27992837 | 1.81849219 | 1.13122229 | 3.14200064 | 0.20783717 | 0.3700422 | PIB1 | IV | 1089219 | 1E+06 | -1 |
| YDR314C | 22.4728227 | 2.30642746 | 0.86430313 | 7.9979285 | 0.01833462 | 0.06204409 | RAD34 | IV | 1090433 | 1E+06 | -1 |
| YDR315C | 23.085208 | 0.18496639 | 0.75871215 | 0.30585758 | 0.85819083 | 0.91626865 | IPK1 | IV | 1092741 | 1E+06 | -1 |
| YDR316W | 71.6735655 | 3.36963511 | 0.88615403 | 20.2951841 | 3.92E-05 | 0.00042234 | OMS1 | IV | 1093763 | 1E+06 | 1 |

| | | | | | | | | | | | |
|---|---|---|---|---|---|---|---|---|---|---|---|
| YDR317W | 622.743478 | 0.30622248 | 0.48367355 | 0.79984771 | 0.67037109 | 0.79537881 | HIM1 | IV | 1102184 | 1E+06 | 1 |
| YDR318W | 1175.04082 | -0.4653627 | 0.66417361 | 2.53070758 | 0.28213946 | 0.45618192 | MCM21 | IV | 1103758 | 1E+06 | 1 |
| YDR319C | 304.702776 | 1.95822472 | 0.76290596 | 6.6037691 | 0.03681372 | 0.10453726 | YFT2 | IV | 1105003 | 1E+06 | -1 |
| YDR320C | 46.5701599 | 0.59772629 | 0.62999971 | 3.36447636 | 0.1859573 | 0.34169876 | SWA2 | IV | 1106095 | 1E+06 | -1 |
| YDR320C-A | 260.239675 | 0.85842046 | 0.50785386 | 3.03846249 | 0.21888009 | 0.38305402 | DAD4 | IV | 1108280 | 1E+06 | -1 |
| YDR321W | 13.0422662 | -2.1830693 | 0.88245504 | 6.10073419 | 0.04734154 | 0.12533495 | ASP1 | IV | 1108702 | 1E+06 | 1 |
| YDR322C-A | 72.9965435 | 1.80712084 | 0.632178 | 8.44078002 | 0.01469291 | 0.05261147 | TIM11 | IV | 1112003 | 1E+06 | -1 |
| YDR322W | 44.7391934 | 2.37385037 | 0.63040251 | 20.4624851 | 3.60E-05 | 0.00039327 | MRPL35 | IV | 1110589 | 1E+06 | 1 |
| YDR323C | 16.1746546 | -1.0566931 | 0.89776474 | 1.73339751 | 0.4203369 | 0.59107763 | PEP7 | IV | 1112480 | 1E+06 | -1 |
| YDR324C | 96.842625 | 2.03534403 | 0.59477872 | 15.8337655 | 0.00036454 | 0.00285226 | UTP4 | IV | 1114433 | 1E+06 | -1 |
| YDR325W | 71.7712552 | 2.21825303 | 0.6395243 | 12.9312307 | 0.00155603 | 0.00937506 | YCG1 | IV | 1117126 | 1E+06 | 1 |
| YDR326C | 28.7770711 | -0.5145297 | 0.72661372 | 0.54436511 | 0.76171519 | 0.85526961 | YSP2 | IV | 1120609 | 1E+06 | -1 |
| YDR328C | 54.3073416 | -0.1363268 | 0.61700142 | 1.01993936 | 0.60051379 | 0.74682758 | SKP1 | IV | 1125434 | 1E+06 | -1 |
| YDR329C | 23.2193538 | -0.4801288 | 0.83158337 | 0.64762183 | 0.72338701 | 0.83301726 | PEX3 | IV | 1126270 | 1E+06 | -1 |
| YDR330W | 16.0343649 | 0.72815691 | 0.80992073 | 0.81706133 | 0.66462609 | 0.79148192 | UBX5 | IV | 1127872 | 1E+06 | 1 |
| YDR331W | 71.6029717 | 1.10203873 | 0.56293205 | 8.95308469 | 0.01137267 | 0.04332089 | GPI8 | IV | 1129588 | 1E+06 | 1 |
| YDR332W | 177.183279 | -0.3789719 | 0.5842918 | 4.34006011 | 0.11417419 | 0.24102109 | IRC3 | IV | 1131001 | 1E+06 | 1 |
| YDR333C | 21.8051429 | -1.2566754 | 0.75017114 | 3.15275487 | 0.20672261 | 0.36873811 | RQC1 | IV | 1133260 | 1E+06 | -1 |
| YDR334W | 37.3171476 | 0.46392625 | 0.63736867 | 0.71268263 | 0.70023358 | 0.81801018 | SWR1 | IV | 1135932 | 1E+06 | 1 |
| YDR335W | 81.608514 | 0.22421104 | 0.64779506 | 3.01503062 | 0.22145955 | 0.38636302 | MSN5 | IV | 1141168 | 1E+06 | 1 |
| YDR336W | 24.7301787 | 1.00691431 | 0.68518216 | 2.33350461 | 0.31137656 | 0.48399202 | MRX8 | IV | 1145092 | 1E+06 | 1 |
| YDR337W | 66.6952418 | 1.44877039 | 0.65443031 | 4.75374893 | 0.0928403 | 0.20754646 | MRPS28 | IV | 1146319 | 1E+06 | 1 |
| YDR338C | 83.9168802 | 2.55486904 | 0.60469055 | 16.7952551 | 0.0002254 | 0.00188867 | NA | IV | 1147379 | 1E+06 | -1 |
| YDR339C | 25.9608686 | -1.748921 | 0.77788515 | 6.21810574 | 0.04464322 | 0.12011349 | FCF1 | IV | 1149951 | 1E+06 | -1 |
| YDR341C | 28.417631 | -2.2191301 | 0.77435183 | 8.15459284 | 0.01695324 | 0.05856757 | NA | IV | 1151803 | 1E+06 | -1 |
| YDR344C | 117.507842 | 1.43245287 | 0.51195054 | 8.27277046 | 0.01598051 | 0.05628173 | NA | IV | 1162006 | 1E+06 | -1 |
| YDR345C | 104.598713 | 0.9055094 | 0.56258711 | 4.99851383 | 0.08214602 | 0.18987044 | HXT3 | IV | 1162957 | 1E+06 | -1 |
| YDR346C | 94.4122137 | 0.21755209 | 0.53043068 | 0.16838767 | 0.91925306 | 0.94974417 | SVF1 | IV | 1167214 | 1E+06 | -1 |
| YDR347W | 88.7493759 | 0.6280482 | 0.54298342 | 2.00642663 | 0.36669923 | 0.53983944 | MRP1 | IV | 1169178 | 1E+06 | 1 |
| YDR348C | 337.965777 | 4.64042671 | 0.6680015 | 49.6409204 | 1.66E-11 | 9.34E-10 | PAL1 | IV | 1170326 | 1E+06 | -1 |
| YDR349C | 62.3459624 | 0.06947791 | 0.54743847 | 4.09990969 | 0.12874072 | 0.26238612 | YPS7 | IV | 1172386 | 1E+06 | -1 |
| YDR350C | 13.2683833 | 0.59447313 | 0.9115494 | 1.01769388 | 0.60118839 | 0.74713045 | ATP22 | IV | 1176120 | 1E+06 | -1 |
| YDR351W | 21.6263266 | -0.3554997 | 0.89372982 | 1.2325776 | 0.53994457 | 0.69395761 | SBE2 | IV | 1178666 | 1E+06 | 1 |

| | | | | | | | | | | | |
|---|---|---|---|---|---|---|---|---|---|---|---|
| YDR352W | 98.8655909 | -0.2933969 | 0.53578687 | 0.46069587 | 0.7942572 | 0.87495229 | YPQ2 | IV | 1181801 | 1E+06 | 1 |
| YDR353W | 194.767144 | -0.5111163 | 0.58894283 | 0.7779117 | 0.67776419 | 0.8011991 | TRR1 | IV | 1183299 | 1E+06 | 1 |
| YDR354W | 17.2618509 | 1.1573334 | 0.90631826 | 2.37902354 | 0.30436983 | 0.47720463 | TRP4 | IV | 1184747 | 1E+06 | 1 |
| YDR356W | 26.3714575 | 1.35795299 | 0.68992186 | 3.80246279 | 0.14938455 | 0.29203007 | SPC110 | IV | 1186107 | 1E+06 | 1 |
| YDR357C | 176.623765 | -0.7972225 | 0.77853801 | 2.40839894 | 0.29993201 | 0.47325515 | CNL1 | IV | 1189199 | 1E+06 | -1 |
| YDR358W | 58.9108891 | -0.9023669 | 0.59007666 | 2.60241225 | 0.27220328 | 0.44559146 | GGA1 | IV | 1190059 | 1E+06 | 1 |
| YDR359C | 138.129953 | -1.3968775 | 0.50241072 | 7.59949791 | 0.02237639 | 0.07205745 | EAF1 | IV | 1191936 | 1E+06 | -1 |
| YDR361C | 47.7453056 | 0.07871246 | 0.6260491 | 0.3583629 | 0.8359542 | 0.9031862 | BCP1 | IV | 1195412 | 1E+06 | -1 |
| YDR362C | 247.672768 | 2.2000536 | 0.50139667 | 20.1652313 | 4.18E-05 | 0.00044467 | TFC6 | IV | 1196679 | 1E+06 | -1 |
| YDR363W | 27.0883058 | 1.23623937 | 0.67967454 | 4.03074544 | 0.13327072 | 0.26928639 | ESC2 | IV | 1199183 | 1E+06 | 1 |
| YDR363W-A | 90.7312727 | -0.5748142 | 0.59694993 | 3.60589087 | 0.16481273 | 0.31379996 | SEM1 | IV | 1202128 | 1E+06 | 1 |
| YDR364C | 87.6695912 | 2.68501126 | 0.62320215 | 28.7065655 | 5.84E-07 | 1.11E-05 | CDC40 | IV | 1202843 | 1E+06 | -1 |
| YDR365C | 93.357196 | -2.3415973 | 0.58547198 | 15.4053957 | 0.00045161 | 0.00338267 | ESF1 | IV | 1204497 | 1E+06 | -1 |
| YDR366C | 718.366383 | -2.7003025 | 0.56527487 | 22.6266855 | 1.22E-05 | 0.00015709 | NA | IV | 1212438 | 1E+06 | -1 |
| YDR367W | 23.6739416 | 1.00132016 | 0.71010789 | 2.1898692 | 0.33456149 | 0.50802806 | KEI1 | IV | 1212848 | 1E+06 | 1 |
| YDR368W | 175.516408 | -2.2876073 | 0.52560776 | 17.453997 | 0.00016215 | 0.00142248 | YPR1 | IV | 1213904 | 1E+06 | 1 |
| YDR369C | 57.4483607 | 1.18566625 | 0.59342042 | 4.02053127 | 0.13395309 | 0.27005166 | XRS2 | IV | 1215016 | 1E+06 | -1 |
| YDR370C | 9.78894465 | 0.30175129 | 0.90021182 | 0.12428858 | 0.93974728 | 0.96037943 | DXO1 | IV | 1217782 | 1E+06 | -1 |
| YDR371W | 69.1155655 | 0.27361332 | 0.79301708 | 7.76137061 | 0.02063668 | 0.06765932 | CTS2 | IV | 1219413 | 1E+06 | 1 |
| YDR372C | 32.7757912 | 0.15664237 | 0.68212299 | 0.24549892 | 0.88448523 | 0.9315066 | VPS74 | IV | 1221112 | 1E+06 | -1 |
| YDR373W | 209.21465 | 4.63939087 | 0.65728236 | 50.064797 | 1.34E-11 | 7.71E-10 | FRQ1 | IV | 1222759 | 1E+06 | 1 |
| YDR374C | 12.2377907 | 0.66935853 | 0.90352049 | 2.55520927 | 0.2787041 | 0.45315831 | PHO92 | IV | 1223474 | 1E+06 | -1 |
| YDR375C | 227.095816 | 1.28719588 | 0.6422196 | 9.17237015 | 0.01019166 | 0.04004855 | BCS1 | IV | 1225166 | 1E+06 | -1 |
| YDR376W | 99.3596946 | 2.71173256 | 0.56646289 | 27.863404 | 8.90E-07 | 1.61E-05 | ARH1 | IV | 1226822 | 1E+06 | 1 |
| YDR377W | 11.3062227 | -0.7012937 | 0.92995102 | 0.62274574 | 0.73244072 | 0.83744703 | ATP17 | IV | 1228611 | 1E+06 | 1 |
| YDR378C | 44.466696 | -2.0768788 | 0.87904306 | 7.15650087 | 0.02792451 | 0.08418952 | LSM6 | IV | 1229349 | 1E+06 | -1 |
| YDR379C-A | 36.1142658 | 1.53064526 | 0.98376748 | 10.9807539 | 0.00412629 | 0.01964738 | SDH6 | IV | 1233278 | 1E+06 | -1 |
| YDR379W | 615.473783 | 2.58996036 | 0.8758898 | 15.23665 | 0.00049136 | 0.00363486 | RGA2 | IV | 1230167 | 1E+06 | 1 |
| YDR380W | 1091.19385 | -2.6827457 | 0.48446357 | 27.3759379 | 1.14E-06 | 1.98E-05 | ARO10 | IV | 1234218 | 1E+06 | 1 |
| YDR381C-A | 148.135608 | 1.90196352 | 0.58056715 | 10.1438389 | 0.00627037 | 0.02746253 | NA | IV | 1238312 | 1E+06 | -1 |
| YDR381W | 151.484618 | -1.2740559 | 0.54785725 | 6.08130329 | 0.04780373 | 0.12615478 | YRA1 | IV | 1236558 | 1E+06 | 1 |
| YDR382W | 127.817346 | -0.3010275 | 0.60203991 | 0.3206629 | 0.85186139 | 0.91195795 | RPP2B | IV | 1239492 | 1E+06 | 1 |
| YDR383C | 2525.48605 | -1.2344861 | 1.74088088 | 4.95912716 | 0.08377978 | 0.19280005 | NKP1 | IV | 1239960 | 1E+06 | -1 |

| | | | | | | | | | | | |
|---|---|---|---|---|---|---|---|---|---|---|---|
| YDR384C | 250.209484 | 3.29141079 | 0.5202011 | 40.4879245 | 1.61E-09 | 5.77E-08 | ATO3 | IV | 1241204 | 1E+06 | -1 |
| YDR385W | 31.6792435 | -1.2156568 | 0.84456356 | 2.28084828 | 0.3196834 | 0.49223979 | EFT1 | IV | 1243230 | 1E+06 | 1 |
| YDR386W | 35.3878877 | 1.59984929 | 0.68531948 | 5.99866446 | 0.04982033 | 0.13021013 | MUS81 | IV | 1246084 | 1E+06 | 1 |
| YDR387C | 31.542061 | -0.1842166 | 0.68299596 | 0.08681253 | 0.9575223 | 0.97008821 | CIN10 | IV | 1248154 | 1E+06 | -1 |
| YDR388W | 109.278628 | 0.63038325 | 0.60684698 | 3.29106625 | 0.19290969 | 0.34948282 | RVS167 | IV | 1250186 | 1E+06 | 1 |
| YDR389W | 42.3207474 | 0.88838214 | 0.61108927 | 2.14621837 | 0.3419437 | 0.51558698 | SAC7 | IV | 1252537 | 1E+06 | 1 |
| YDR390C | 63.2942293 | 0.79636941 | 0.75038302 | 6.02746503 | 0.04910804 | 0.12889191 | UBA2 | IV | 1254937 | 1E+06 | -1 |
| YDR391C | 12.2148509 | 2.09217642 | 0.96372323 | 5.03284152 | 0.08074811 | 0.18768829 | NA | IV | 1257358 | 1E+06 | -1 |
| YDR392W | 19.6990755 | 0.0625084 | 0.79580121 | 0.93696142 | 0.62595255 | 0.76430043 | SPT3 | IV | 1258696 | 1E+06 | 1 |
| YDR393W | 281.063151 | 0.63055408 | 0.4690588 | 1.97261606 | 0.37295108 | 0.5462653 | SHE9 | IV | 1259901 | 1E+06 | 1 |
| YDR394W | 29.8398429 | 0.28472394 | 0.81192631 | 0.20855247 | 0.90097638 | 0.93824416 | RPT3 | IV | 1261681 | 1E+06 | 1 |
| YDR395W | 64.1672337 | -0.227234 | 0.59553123 | 0.29319573 | 0.86364121 | 0.91991868 | SXM1 | IV | 1263324 | 1E+06 | 1 |
| YDR397C | 47.8094006 | -0.5049998 | 0.59309626 | 1.77036372 | 0.41263912 | 0.58426143 | NCB2 | IV | 1266366 | 1E+06 | -1 |
| YDR398W | 691.820815 | 5.30652983 | 0.53546752 | 95.4774334 | 1.85E-21 | 6.72E-19 | UTP5 | IV | 1267471 | 1E+06 | 1 |
| YDR399W | 69.2561628 | 0.7078395 | 0.5476298 | 1.69027399 | 0.42949852 | 0.59890573 | HPT1 | IV | 1270068 | 1E+06 | 1 |
| YDR400W | 23.1586837 | 0.34042053 | 0.71650748 | 0.24247338 | 0.88582427 | 0.93185365 | URH1 | IV | 1271063 | 1E+06 | 1 |
| YDR402C | 500.253706 | -0.2186161 | 0.86025245 | 2.30359195 | 0.31606861 | 0.48830599 | DIT2 | IV | 1272234 | 1E+06 | -1 |
| YDR403W | 638.391017 | 2.52338454 | 0.53620089 | 22.466019 | 1.32E-05 | 0.00016799 | DIT1 | IV | 1274602 | 1E+06 | 1 |
| YDR404C | 76.3610355 | 1.84272515 | 0.58308241 | 11.9044314 | 0.00260007 | 0.013661 | RPB7 | IV | 1276654 | 1E+06 | -1 |
| YDR405W | 36.1791597 | 1.53314302 | 0.63663251 | 6.05786584 | 0.04836722 | 0.12713301 | MRP20 | IV | 1277646 | 1E+06 | 1 |
| YDR406W | 342.054045 | 0.19794655 | 0.59564947 | 0.843519 | 0.65589176 | 0.78495521 | PDR15 | IV | 1279210 | 1E+06 | 1 |
| YDR407C | 30.1448295 | -0.2729607 | 0.67703093 | 0.22945807 | 0.89160771 | 0.93352778 | TRS120 | IV | 1284069 | 1E+06 | -1 |
| YDR408C | 200.102673 | 0.60828049 | 0.55377542 | 4.20193957 | 0.12233773 | 0.25315778 | ADE8 | IV | 1288215 | 1E+06 | -1 |
| YDR409W | 876.859297 | -1.2222307 | 0.46603948 | 8.11996024 | 0.01724936 | 0.05934272 | SIZ1 | IV | 1289406 | 1E+06 | 1 |
| YDR410C | 91.3475944 | -0.5592149 | 0.9207881 | 9.14967027 | 0.010308 | 0.04032978 | STE14 | IV | 1292372 | 1E+06 | -1 |
| YDR411C | 39.1475792 | 1.33069433 | 0.68200791 | 6.27341865 | 0.04342546 | 0.11809931 | DFM1 | IV | 1293369 | 1E+06 | -1 |
| YDR412W | 42.6492483 | -2.2777336 | 0.74602345 | 9.04180372 | 0.01087921 | 0.04193757 | RRP17 | IV | 1294693 | 1E+06 | 1 |
| YDR414C | 557.682694 | 0.13464012 | 0.49375161 | 1.45720079 | 0.48258394 | 0.644856 | ERD1 | IV | 1295598 | 1E+06 | -1 |
| YDR415C | 15.9850751 | 0.36986007 | 0.82766621 | 0.39589728 | 0.82041199 | 0.8920536 | NA | IV | 1297037 | 1E+06 | -1 |
| YDR416W | 59.2709766 | -1.4539718 | 0.56968414 | 6.32518604 | 0.04231587 | 0.11589825 | SYF1 | IV | 1298432 | 1E+06 | 1 |
| YDR418W | 179.122783 | -1.9380619 | 0.56297718 | 12.1886169 | 0.00225567 | 0.01226322 | RPL12B | IV | 1301616 | 1E+06 | 1 |
| YDR419W | 192.727971 | -0.373949 | 0.50084324 | 2.37916907 | 0.30434768 | 0.47720463 | RAD30 | IV | 1303174 | 1E+06 | 1 |
| YDR420W | 23.6830874 | 0.55936581 | 0.72395587 | 0.63201853 | 0.7290527 | 0.83668919 | HKR1 | IV | 1306267 | 1E+06 | 1 |

| | | | | | | | | | | | |
|---|---|---|---|---|---|---|---|---|---|---|---|
| YDR421W | 186.883693 | 0.8592839 | 0.56955176 | 7.16560454 | 0.02779769 | 0.08391483 | ARO80 | IV | 1312040 | 1E+06 | 1 |
| YDR422C | 64.3643335 | 2.46766106 | 0.57420313 | 24.4826429 | 4.83E-06 | 6.90E-05 | SIP1 | IV | 1315326 | 1E+06 | -1 |
| YDR423C | 24.5448454 | 1.26934322 | 0.75398667 | 3.05657473 | 0.21690683 | 0.38103476 | CAD1 | IV | 1318046 | 1E+06 | -1 |
| YDR424C | 65.1832492 | 0.03397156 | 0.56254577 | 0.06619872 | 0.96744243 | 0.97486964 | DYN2 | IV | 1319387 | 1E+06 | -1 |
| YDR425W | 30.3939495 | 0.05264981 | 0.69030402 | 4.79693331 | 0.09085716 | 0.20453459 | SNX41 | IV | 1320064 | 1E+06 | 1 |
| YDR427W | 236.712341 | -1.8178222 | 0.58660216 | 11.8210096 | 0.00271082 | 0.01406419 | RPN9 | IV | 1322205 | 1E+06 | 1 |
| YDR428C | 362.801641 | -1.2383478 | 0.61720294 | 9.0751708 | 0.01069921 | 0.04143366 | BNA7 | IV | 1323454 | 1E+06 | -1 |
| YDR429C | 40.3697365 | -2.2378326 | 0.77922003 | 8.09997031 | 0.01742263 | 0.05974247 | TIF35 | IV | 1324477 | 1E+06 | -1 |
| YDR430C | 20.0190041 | -0.5897113 | 0.70603984 | 0.87086933 | 0.64698339 | 0.77880121 | CYM1 | IV | 1325501 | 1E+06 | -1 |
| YDR432W | 183.436973 | 0.10832694 | 0.60332842 | 3.32465227 | 0.18969721 | 0.34571823 | NPL3 | IV | 1328783 | 1E+06 | 1 |
| YDR434W | 112.063018 | 0.22267807 | 0.70133566 | 12.3271562 | 0.00210471 | 0.01162812 | GPI17 | IV | 1331237 | 1E+06 | 1 |
| YDR435C | 14.4005583 | 0.71008432 | 0.8487587 | 4.1215517 | 0.12735512 | 0.26092929 | PPM1 | IV | 1332983 | 1E+06 | -1 |
| YDR436W | 84.9202402 | 1.35027668 | 0.57481686 | 11.8067806 | 0.00273017 | 0.01413927 | PPZ2 | IV | 1334821 | 1E+06 | 1 |
| YDR437W | 30.0280533 | -0.1761044 | 0.67048746 | 0.07652516 | 0.96246018 | 0.97258112 | GPI19 | IV | 1337352 | 1E+06 | 1 |
| YDR438W | 366.265586 | 1.15578174 | 0.5734618 | 4.01197125 | 0.13452764 | 0.27098851 | THI74 | IV | 1338274 | 1E+06 | 1 |
| YDR439W | 47.2765003 | -0.1085857 | 0.75417322 | 3.9902872 | 0.13599412 | 0.27312035 | LRS4 | IV | 1339676 | 1E+06 | 1 |
| YDR440W | 51.0372256 | -0.542723 | 0.79200129 | 1.29007239 | 0.52464355 | 0.67987605 | DOT1 | IV | 1342493 | 1E+06 | 1 |
| YDR441C | 242.384671 | 0.52671288 | 0.51041591 | 1.62613244 | 0.44349612 | 0.61162622 | APT2 | IV | 1344517 | 1E+06 | -1 |
| YDR443C | 23.3769063 | 0.01772407 | 0.68678096 | 0.11556317 | 0.94385608 | 0.96197675 | SSN2 | IV | 1345676 | 1E+06 | -1 |
| YDR444W | 46.014738 | 2.1763315 | 0.65396623 | 16.841491 | 0.00022025 | 0.001854 | NA | IV | 1350290 | 1E+06 | 1 |
| YDR446W | 88.901944 | -3.467771 | 0.79864532 | 22.0560907 | 1.62E-05 | 0.00019879 | ECM11 | IV | 1353725 | 1E+06 | 1 |
| YDR447C | 47.8813604 | -2.8712677 | 0.77842667 | 12.7268227 | 0.00172348 | 0.01001901 | RPS17B | IV | 1354829 | 1E+06 | -1 |
| YDR448W | 167.879642 | -3.4741296 | 0.51635915 | 41.1635014 | 1.15E-09 | 4.22E-08 | ADA2 | IV | 1356065 | 1E+06 | 1 |
| YDR449C | 18.2800135 | -2.4685141 | 0.85120326 | 9.68734821 | 0.00787806 | 0.03281579 | UTP6 | IV | 1357580 | 1E+06 | -1 |
| YDR450W | 79.4203755 | -3.683719 | 1.98870706 | 2.85142558 | 0.24033709 | 0.40843903 | RPS18B | IV | 1359923 | 1E+06 | 1 |
| YDR451C | 166.235535 | -0.3691217 | 0.49675404 | 0.55333371 | 0.75830708 | 0.85403579 | YHP1 | IV | 1361120 | 1E+06 | -1 |
| YDR452W | 1456.9095 | -2.4483969 | 0.48789284 | 23.6138779 | 7.45E-06 | 0.00010106 | PPN1 | IV | 1362878 | 1E+06 | 1 |
| YDR453C | 81.2840014 | -1.2406691 | 0.84547071 | 7.13122589 | 0.02827965 | 0.08501514 | TSA2 | IV | 1365072 | 1E+06 | -1 |
| YDR454C | 49.5704188 | -0.9023323 | 0.58491525 | 7.24896761 | 0.02666286 | 0.08136057 | GUK1 | IV | 1366264 | 1E+06 | -1 |
| YDR456W | 127.309557 | 1.74677357 | 0.53231152 | 11.4615587 | 0.00324455 | 0.01625081 | NHX1 | IV | 1367485 | 1E+06 | 1 |
| YDR457W | 41.7733416 | -0.4410301 | 0.70349221 | 0.47002305 | 0.79056174 | 0.87307888 | TOM1 | IV | 1369790 | 1E+06 | 1 |
| YDR458C | 55.8958173 | 0.09977717 | 0.55045712 | 0.68490283 | 0.71002761 | 0.82438173 | HEH2 | IV | 1380055 | 1E+06 | -1 |
| YDR459C | 16.4619699 | 0.41705181 | 0.92855411 | 0.43635784 | 0.80398158 | 0.88247457 | PFA5 | IV | 1382319 | 1E+06 | -1 |

| | | | | | | | | | | | |
|---|---|---|---|---|---|---|---|---|---|---|---|
| YDR460W | 23.7977729 | -0.4191905 | 0.71353115 | 1.46571173 | 0.48053469 | 0.64330539 | TFB3 | IV | 1383811 | 1E+06 | 1 |
| YDR462W | 9.74320008 | -1.235833 | 0.91730439 | 2.02576046 | 0.36317145 | 0.53614552 | MRPL28 | IV | 1386073 | 1E+06 | 1 |
| YDR463W | 27.4529557 | 0.16516232 | 0.66095915 | 1.3836789 | 0.50065429 | 0.66001556 | STP1 | IV | 1386816 | 1E+06 | 1 |
| YDR464W | 87.7171636 | -1.2487317 | 0.59899363 | 6.08634473 | 0.04768338 | 0.12600836 | SPP41 | IV | 1388872 | 1E+06 | 1 |
| YDR465C | 91.4595215 | -1.8902801 | 0.62836325 | 11.9944255 | 0.00248567 | 0.01316746 | RMT2 | IV | 1393336 | 1E+06 | -1 |
| YDR466W | 278.549352 | -1.8127024 | 0.47482709 | 14.4537814 | 0.00072678 | 0.00503957 | PKH3 | IV | 1395121 | 1E+06 | 1 |
| YDR468C | 155.125067 | -0.6720922 | 0.79202872 | 6.6088863 | 0.03671965 | 0.10437251 | TLG1 | IV | 1398026 | 1E+06 | -1 |
| YDR469W | 85.0700435 | 0.14107985 | 0.75230027 | 6.10250765 | 0.04729958 | 0.12530377 | SDC1 | IV | 1399015 | 1E+06 | 1 |
| YDR470C | 165.299793 | -1.7878547 | 0.51842685 | 11.4437753 | 0.00327353 | 0.01636734 | UGO1 | IV | 1399706 | 1E+06 | -1 |
| YDR471W | 80.8401678 | -3.0037223 | 0.62552835 | 21.114373 | 2.60E-05 | 0.00029817 | RPL27B | IV | 1401770 | 1E+06 | 1 |
| YDR472W | 85.4094283 | -0.1084278 | 0.65070966 | 3.78215391 | 0.1509092 | 0.29389985 | TRS31 | IV | 1403322 | 1E+06 | 1 |
| YDR473C | 71.802308 | 0.32577438 | 0.57147365 | 0.39019657 | 0.82275379 | 0.89359303 | PRP3 | IV | 1404445 | 1E+06 | -1 |
| YDR475C | 412.200435 | -2.4807835 | 0.47016893 | 25.8250094 | 2.47E-06 | 3.86E-05 | JIP4 | IV | 1407464 | 1E+06 | -1 |
| YDR476C | 101.238069 | -1.7195679 | 0.56600572 | 9.78218448 | 0.00751321 | 0.03164938 | NA | IV | 1410453 | 1E+06 | -1 |
| YDR477W | 41.8219356 | -0.1348855 | 0.61099043 | 1.55286241 | 0.46004489 | 0.62448065 | SNF1 | IV | 1412373 | 1E+06 | 1 |
| YDR478W | 206.103414 | 1.03726521 | 0.70597252 | 9.04048462 | 0.01088639 | 0.04193757 | SNM1 | IV | 1414575 | 1E+06 | 1 |
| YDR479C | 679.294326 | -1.8843899 | 0.4748315 | 16.1744062 | 0.00030745 | 0.00249667 | PEX29 | IV | 1415210 | 1E+06 | -1 |
| YDR480W | 124.784615 | -1.1724055 | 0.54054115 | 4.81089492 | 0.09022511 | 0.20366605 | DIG2 | IV | 1417399 | 1E+06 | 1 |
| YDR481C | 198.427545 | -1.8540943 | 0.48516149 | 13.9147716 | 0.00095158 | 0.00631841 | PHO8 | IV | 1418550 | 1E+06 | -1 |
| YDR482C | 18.0635056 | -0.4032713 | 0.74350218 | 1.03212625 | 0.59686571 | 0.74415428 | CWC21 | IV | 1420431 | 1E+06 | -1 |
| YDR483W | 47.8625355 | 0.40249116 | 0.57830477 | 1.10063348 | 0.5767671 | 0.72593834 | KRE2 | IV | 1421157 | 1E+06 | 1 |
| YDR484W | 39.5615912 | 0.08735568 | 0.72654941 | 3.07895604 | 0.21449303 | 0.37828653 | VPS52 | IV | 1422763 | 1E+06 | 1 |
| YDR485C | 49.0156531 | -0.0856033 | 0.6175466 | 1.33696554 | 0.51248555 | 0.66951519 | VPS72 | IV | 1424820 | 1E+06 | -1 |
| YDR486C | 37.1347812 | 0.72851787 | 0.62265454 | 2.23479937 | 0.32712933 | 0.49988884 | VPS60 | IV | 1427431 | 1E+06 | -1 |
| YDR487C | 61.4657002 | 0.37824769 | 0.5470657 | 0.51236509 | 0.77400067 | 0.86281553 | RIB3 | IV | 1428354 | 1E+06 | -1 |
| YDR488C | 6.23681274 | -0.4770414 | 1.08464316 | 0.23681466 | 0.88833413 | 0.93279917 | PAC11 | IV | 1429188 | 1E+06 | -1 |
| YDR489W | 182.604605 | -0.3524759 | 0.52286782 | 2.45961915 | 0.29234824 | 0.46643602 | SLD5 | IV | 1431012 | 1E+06 | 1 |
| YDR490C | 655.772739 | -2.8750932 | 0.50380311 | 28.8493844 | 5.44E-07 | 1.04E-05 | PKH1 | IV | 1431968 | 1E+06 | -1 |
| YDR492W | 13.7360166 | 1.1024412 | 0.81837908 | 3.63770535 | 0.16221175 | 0.31078956 | IZH1 | IV | 1434924 | 1E+06 | 1 |
| YDR494W | 96.4198848 | -1.062342 | 0.73423341 | 8.03396241 | 0.01800724 | 0.06111485 | RSM28 | IV | 1436930 | 1E+06 | 1 |
| YDR495C | 163.201288 | -1.7924374 | 0.51261828 | 16.6310425 | 0.00024469 | 0.00203556 | VPS3 | IV | 1438115 | 1E+06 | -1 |
| YDR496C | 32.104409 | -2.0784766 | 0.6905685 | 9.60218521 | 0.00822076 | 0.03390367 | PUF6 | IV | 1441433 | 1E+06 | -1 |
| YDR497C | 38.4561229 | -0.1122415 | 0.62881583 | 1.61683001 | 0.44556372 | 0.61307366 | ITR1 | IV | 1443713 | 1E+06 | -1 |

| | | | | | | | | | | | |
|---|---|---|---|---|---|---|---|---|---|---|---|
| YDR498C | 60.8878415 | -0.2332577 | 0.59935207 | 0.16575092 | 0.92046578 | 0.95024857 | SEC20 | IV | 1445843 | 1E+06 | -1 |
| YDR499W | 61.4240601 | -2.1486213 | 0.67577829 | 11.3796591 | 0.00338017 | 0.0167275 | LCD1 | IV | 1447830 | 1E+06 | 1 |
| YDR500C | 46.5357075 | -2.4999081 | 0.78590541 | 10.176656 | 0.00616832 | 0.02707703 | RPL37B | IV | 1450198 | 1E+06 | -1 |
| YDR501W | 290.791727 | 0.78472292 | 0.51915789 | 3.2096209 | 0.20092764 | 0.36006531 | PLM2 | IV | 1451353 | 1E+06 | 1 |
| YDR502C | 26.9680007 | -0.1872249 | 0.79531875 | 0.35316376 | 0.83813015 | 0.90469306 | SAM2 | IV | 1453310 | 1E+06 | -1 |
| YDR503C | 10.9543258 | -1.3072716 | 1.00615785 | 1.74513064 | 0.41787818 | 0.58926319 | LPP1 | IV | 1455042 | 1E+06 | -1 |
| YDR504C | 213.079154 | -2.7798801 | 0.53959961 | 24.4794037 | 4.83E-06 | 6.90E-05 | SPG3 | IV | 1456311 | 1E+06 | -1 |
| YDR505C | 31.8644236 | 0.21462491 | 0.65034249 | 0.14029941 | 0.93225425 | 0.95731799 | PSP1 | IV | 1456695 | 1E+06 | -1 |
| YDR506C | 107.967187 | -0.2679055 | 0.50353698 | 1.50645495 | 0.47084446 | 0.63296713 | GMC1 | IV | 1459728 | 1E+06 | -1 |
| YDR507C | 782.888462 | 4.23201982 | 0.48488568 | 64.5900929 | 9.43E-15 | 9.25E-13 | GIN4 | IV | 1462358 | 1E+06 | -1 |
| YDR508C | 65.1517139 | -1.83974 | 0.69991254 | 9.28198545 | 0.00964811 | 0.03839103 | GNP1 | IV | 1466453 | 1E+06 | -1 |
| YDR510W | 29.6747605 | -0.8804345 | 0.74264059 | 2.28047345 | 0.31974332 | 0.49223979 | SMT3 | IV | 1469400 | 1E+06 | 1 |
| YDR511W | 183.187563 | 0.09930669 | 0.77830576 | 6.59096553 | 0.03705015 | 0.10484868 | SDH7 | IV | 1470017 | 1E+06 | 1 |
| YDR512C | 2366.83895 | -0.2183805 | 0.47879899 | 0.61555873 | 0.73507748 | 0.83830975 | EMI1 | IV | 1470500 | 1E+06 | -1 |
| YDR513W | 11.4759735 | -0.2705871 | 0.9518224 | 0.33984426 | 0.84373052 | 0.90836736 | GRX2 | IV | 1471017 | 1E+06 | 1 |
| YDR514C | 97.1348473 | 0.20125435 | 0.98379795 | 8.17487422 | 0.01678219 | 0.05813116 | NA | IV | 1471548 | 1E+06 | -1 |
| YDR515W | 195.698393 | 4.7935684 | 0.61320677 | 67.8520408 | 1.85E-15 | 2.14E-13 | SLF1 | IV | 1473429 | 1E+06 | 1 |
| YDR516C | 144.010611 | 1.6588298 | 0.51128238 | 16.0883409 | 0.00032097 | 0.0025883 | EMI2 | IV | 1474974 | 1E+06 | -1 |
| YDR517W | 46.7196339 | 1.10898977 | 0.59888492 | 3.71093067 | 0.15638016 | 0.30191434 | GRH1 | IV | 1477239 | 1E+06 | 1 |
| YDR518W | 335.848086 | 3.02467059 | 0.55142681 | 38.4724767 | 4.42E-09 | 1.44E-07 | EUG1 | IV | 1478608 | 1E+06 | 1 |
| YDR519W | 92.9916488 | 0.40189971 | 0.6693495 | 0.49458073 | 0.78091391 | 0.86705973 | FPR2 | IV | 1480425 | 1E+06 | 1 |
| YDR520C | 453.089386 | 0.4446416 | 0.65710516 | 7.87031169 | 0.01954265 | 0.06499882 | URC2 | IV | 1481085 | 1E+06 | -1 |
| YDR522C | 303.509566 | 0.23784207 | 0.55369204 | 0.20032064 | 0.90469237 | 0.94008773 | SPS2 | IV | 1483795 | 1E+06 | -1 |
| YDR523C | 11.5972407 | 1.62292381 | 1.07674276 | 2.47759374 | 0.28973259 | 0.46366825 | SPS1 | IV | 1485566 | 1E+06 | -1 |
| YDR524C | 113.452542 | 1.25154829 | 0.52910536 | 5.45765557 | 0.06529579 | 0.15958742 | AGE1 | IV | 1487542 | 1E+06 | -1 |
| YDR524C-B | 92.1323003 | -1.3951954 | 0.7014117 | 4.07424882 | 0.13040316 | 0.26529666 | NA | IV | 1489598 | 1E+06 | -1 |
| YDR524W-C | 71688.527 | -2.0038159 | 0.50076705 | 15.5802559 | 0.0004138 | 0.00315666 | NA | IV | 1489403 | 1E+06 | 1 |
| YDR525W | 41.4819871 | -0.4692339 | 0.64084818 | 2.43512891 | 0.29595009 | 0.46995614 | NA | IV | 1489905 | 1E+06 | 1 |
| YDR525W-A | 13.3749558 | -1.015938 | 0.86442593 | 2.11737096 | 0.34691153 | 0.5203673 | SNA2 | IV | 1490596 | 1E+06 | 1 |
| YDR527W | 82.6099973 | -2.0779852 | 0.62448654 | 10.4841894 | 0.00528917 | 0.02396265 | RBA50 | IV | 1491094 | 1E+06 | 1 |
| YDR528W | 38.0072311 | 1.17068035 | 0.6511866 | 4.63316684 | 0.09860992 | 0.2170918 | HLR1 | IV | 1494586 | 1E+06 | 1 |
| YDR529C | 109.11031 | -0.6275246 | 0.60773584 | 2.23890501 | 0.32645848 | 0.49927259 | QCR7 | IV | 1496165 | 1E+06 | -1 |
| YDR530C | 11.5487157 | 0.38579213 | 0.87582785 | 1.4279184 | 0.48970153 | 0.65106127 | APA2 | IV | 1496791 | 1E+06 | -1 |

| | | | | | | | | | | | |
|---|---|---|---|---|---|---|---|---|---|---|---|
| YDR531W | 20.3198999 | -0.8667104 | 0.89527275 | 5.93695457 | 0.05138149 | 0.13358726 | CAB1 | IV | 1498232 | 1E+06 | 1 |
| YDR532C | 121.981467 | -0.082898 | 0.94071055 | 10.5010286 | 0.00524482 | 0.02385369 | KRE28 | IV | 1499396 | 2E+06 | -1 |
| YDR533C | 101.205409 | -1.8599825 | 0.57117574 | 11.3444006 | 0.00344029 | 0.016967 | HSP31 | IV | 1501447 | 2E+06 | -1 |
| YDR534C | 7.19081197 | 2.40381684 | 1.18185038 | 6.43624071 | 0.04003023 | 0.1114839 | FIT1 | IV | 1503314 | 2E+06 | -1 |
| YDR536W | 734.822717 | 0.10583426 | 0.4939878 | 0.38546685 | 0.82470179 | 0.89537284 | STL1 | IV | 1508005 | 2E+06 | 1 |
| YDR539W | 11.3666958 | 0.1420316 | 0.94702219 | 0.06744877 | 0.96683794 | 0.97486964 | FDC1 | IV | 1512094 | 2E+06 | 1 |
| YDR540C | 84.7948889 | 0.19078152 | 0.54711717 | 3.40668037 | 0.18207434 | 0.33713158 | IRC4 | IV | 1517136 | 2E+06 | -1 |
| YDR541C | 103.76907 | 2.70208341 | 0.54867118 | 24.7214174 | 4.28E-06 | 6.26E-05 | NA | IV | 1519664 | 2E+06 | -1 |
| YEL001C | 334.341439 | 0.91673679 | 0.47166694 | 3.92603635 | 0.14043393 | 0.2789408 | IRC22 | V | 150301 | 150978 | -1 |
| YEL002C | 71.0830103 | 0.19141503 | 0.63418847 | 1.51800052 | 0.46813421 | 0.63108197 | WBP1 | V | 148722 | 150014 | -1 |
| YEL003W | 106.66296 | -0.0675431 | 1.01413727 | 8.3548487 | 0.01533796 | 0.05444929 | GIM4 | V | 148176 | 148599 | 1 |
| YEL004W | 199.797505 | -0.4381803 | 0.54658527 | 2.51689351 | 0.28409495 | 0.45768219 | YEA4 | V | 146951 | 147979 | 1 |
| YEL005C | 59.7027959 | -0.0031585 | 0.55049 | 0.87373151 | 0.64605816 | 0.77803336 | VAB2 | V | 145907 | 146755 | -1 |
| YEL006W | 11.8299939 | 1.27805504 | 1.01879341 | 1.69060424 | 0.4294276 | 0.59890573 | YEA6 | V | 144327 | 145334 | 1 |
| YEL007W | 31.6733844 | -1.0604665 | 0.67705643 | 2.921963 | 0.23200845 | 0.39920621 | MIT1 | V | 141892 | 143892 | 1 |
| YEL008W | 257.313017 | -0.7747325 | 0.639832 | 1.44430148 | 0.4857065 | 0.64773825 | NA | V | 140512 | 140892 | 1 |
| YEL009C | 58.5431425 | -0.7570371 | 0.72684503 | 1.15434386 | 0.56148404 | 0.71278066 | GCN4 | V | 138918 | 139763 | -1 |
| YEL009C-A | 8.80277354 | 0.94396802 | 0.94587823 | 1.41805766 | 0.4921219 | 0.6530873 | NA | V | 136371 | 136778 | -1 |
| YEL011W | 354.920885 | -2.3399972 | 0.52930844 | 19.9369042 | 4.69E-05 | 0.00048706 | GLC3 | V | 133120 | 135234 | 1 |
| YEL012W | 39.5710862 | 2.11611362 | 0.78373551 | 9.94607729 | 0.00692208 | 0.02979712 | UBC8 | V | 131772 | 132551 | 1 |
| YEL013W | 30.1812974 | 0.4850625 | 0.70356095 | 0.60028923 | 0.74071109 | 0.84208075 | VAC8 | V | 128825 | 130561 | 1 |
| YEL014C | 965.789855 | -1.2035448 | 0.51427801 | 6.40742318 | 0.04061119 | 0.11266833 | NA | V | 128303 | 128608 | -1 |
| YEL015W | 61.9439591 | 1.25322606 | 0.56553191 | 5.19886178 | 0.07431586 | 0.17629168 | EDC3 | V | 126629 | 128284 | 1 |
| YEL016C | 1157.15296 | 2.31322394 | 0.47055442 | 23.7339618 | 7.02E-06 | 9.56E-05 | NPP2 | V | 124737 | 126218 | -1 |
| YEL017C-A | 35.6311622 | -1.7228205 | 0.67555321 | 6.48389044 | 0.03908779 | 0.10976769 | PMP2 | V | 122798 | 122929 | -1 |
| YEL017W | 256.635516 | -1.077856 | 0.8637091 | 4.72033582 | 0.09440437 | 0.20984715 | GTT3 | V | 123657 | 124670 | 1 |
| YEL018W | 41.3043116 | -1.1158788 | 0.68967249 | 4.56118436 | 0.10222365 | 0.22276062 | EAF5 | V | 121471 | 122310 | 1 |
| YEL020C | 24.9584881 | 1.20717432 | 0.75688126 | 4.81880731 | 0.08986887 | 0.20341703 | PXP1 | V | 118617 | 120299 | -1 |
| YEL020W-A | 18.7887843 | -1.3619833 | 0.80523703 | 2.98712733 | 0.22457093 | 0.39000171 | TIM9 | V | 117211 | 117474 | 1 |
| YEL021W | 131.258755 | 1.93245907 | 0.5717396 | 12.5204096 | 0.00191085 | 0.010794 | URA3 | V | 116167 | 116970 | 1 |
| YEL022W | 162.353839 | 3.44067272 | 0.57851954 | 43.0805948 | 4.42E-10 | 1.75E-08 | GEA2 | V | 111421 | 115800 | 1 |
| YEL023C | 67.0229087 | -0.8723724 | 0.93063524 | 9.33841952 | 0.00937968 | 0.03760965 | NA | V | 108504 | 110552 | -1 |
| YEL024W | 12.2085682 | -1.1410154 | 1.05361527 | 1.20767101 | 0.5467107 | 0.6993379 | RIP1 | V | 107260 | 107907 | 1 |

| | | | | | | | | | | | |
|---|---|---|---|---|---|---|---|---|---|---|---|
| YEL025C | 21.5367859 | 0.07567012 | 0.74998977 | 3.6836856 | 0.15852503 | 0.3051396 | NA | V | 102581 | 106147 | -1 |
| YEL026W | 51.9787036 | -0.076142 | 0.94467245 | 4.55563811 | 0.10250753 | 0.22304343 | SNU13 | V | 101943 | 102323 | 1 |
| YEL027W | 57.5917917 | -1.3912408 | 0.6173133 | 7.66035086 | 0.02170581 | 0.07037316 | VMA3 | V | 100769 | 101251 | 1 |
| YEL028W | 114.851982 | 0.30758258 | 0.5902473 | 1.88763953 | 0.38913857 | 0.56089429 | NA | V | 98668 | 99129 | 1 |
| YEL029C | 69.2822968 | -0.9923414 | 0.60878748 | 2.77678801 | 0.24947564 | 0.42048038 | BUD16 | V | 96858 | 97796 | -1 |
| YEL030W | 101.731766 | -0.5993802 | 0.55953794 | 1.85504264 | 0.39553289 | 0.56679648 | ECM10 | V | 94644 | 96578 | 1 |
| YEL031W | 39.0310599 | -0.4157451 | 0.84079232 | 0.94228775 | 0.62428775 | 0.7633846 | SPF1 | V | 90258 | 93905 | 1 |
| YEL032W | 22.3978604 | -2.0441145 | 0.76494957 | 7.36421218 | 0.02516991 | 0.07818335 | MCM3 | V | 86937 | 89852 | 1 |
| YEL033W | 118.715667 | -2.356933 | 0.58479634 | 16.3591676 | 0.00028032 | 0.00230546 | MTC7 | V | 86179 | 86598 | 1 |
| YEL034W | 211.122941 | -1.9792512 | 0.54172244 | 14.152209 | 0.00084506 | 0.00572938 | HYP2 | V | 85676 | 86149 | 1 |
| YEL035C | 119.429613 | -0.1821269 | 0.72927136 | 0.23214863 | 0.89040905 | 0.93317728 | UTR5 | V | 85045 | 85545 | -1 |
| YEL036C | 131.007072 | 0.27561513 | 0.69132005 | 3.32320907 | 0.18983414 | 0.34583253 | ANP1 | V | 83050 | 84552 | -1 |
| YEL037C | 63.8755901 | -1.1287313 | 0.6850338 | 3.32868245 | 0.18931533 | 0.34544902 | RAD23 | V | 81407 | 82603 | -1 |
| YEL038W | 7.43708748 | -0.5417097 | 1.02995273 | 0.59686617 | 0.74197993 | 0.84243165 | UTR4 | V | 80462 | 81145 | 1 |
| YEL039C | 13.4990488 | -0.3692706 | 0.87795522 | 0.98374697 | 0.61147972 | 0.75489714 | CYC7 | V | 79636 | 79977 | -1 |
| YEL040W | 81.8173603 | -0.4623171 | 0.65716288 | 0.62749345 | 0.73070407 | 0.83678999 | UTR2 | V | 78053 | 79456 | 1 |
| YEL041W | 278.574714 | -1.2283329 | 0.48197772 | 6.48145759 | 0.03913536 | 0.10983701 | YEF1 | V | 75944 | 77431 | 1 |
| YEL042W | 57.1932105 | 1.78847342 | 0.70777435 | 9.33671129 | 0.00938769 | 0.03761574 | GDA1 | V | 73771 | 75327 | 1 |
| YEL043W | 37.2430899 | 0.48155594 | 0.62240618 | 1.67715946 | 0.4323241 | 0.60114231 | NA | V | 70478 | 73348 | 1 |
| YEL044W | 57.9697136 | -1.0544899 | 0.56107812 | 3.8780161 | 0.14384657 | 0.28410385 | IES6 | V | 69757 | 70257 | 1 |
| YEL046C | 67.145133 | 0.90620975 | 0.5757625 | 2.68597711 | 0.2610643 | 0.43235173 | GLY1 | V | 67629 | 68792 | -1 |
| YEL047C | 24.178703 | -0.9315566 | 0.77531675 | 1.76763784 | 0.41320191 | 0.58466253 | FRD1 | V | 65385 | 66797 | -1 |
| YEL048C | 82.5617357 | 3.82816172 | 0.64235357 | 43.6547506 | 3.32E-10 | 1.36E-08 | TCA17 | V | 64709 | 65167 | -1 |
| YEL049W | 224.841917 | 0.38088793 | 0.48495354 | 8.91480643 | 0.01159243 | 0.0440421 | PAU2 | V | 63728 | 64090 | 1 |
| YEL050C | 45.2804867 | 0.26516444 | 0.57831226 | 2.06546128 | 0.35603343 | 0.52942744 | RML2 | V | 59670 | 60851 | -1 |
| YEL051W | 21.4255817 | -1.2865262 | 0.89755876 | 2.60628356 | 0.2716769 | 0.44510732 | VMA8 | V | 58378 | 59148 | 1 |
| YEL052W | 18.7672271 | -0.0094679 | 0.81763023 | 1.07677398 | 0.58368899 | 0.73199919 | AFG1 | V | 56571 | 58100 | 1 |
| YEL053C | 24.1836648 | -1.254691 | 0.74434881 | 3.50039479 | 0.17373964 | 0.32507061 | MAK10 | V | 53901 | 56102 | -1 |
| YEL054C | 109.073932 | -3.0364682 | 0.615519 | 22.0167336 | 1.66E-05 | 0.00020089 | RPL12B | V | 52721 | 53218 | -1 |
| YEL055C | 695.87554 | 0.04873859 | 0.52655069 | 0.13110492 | 0.93654992 | 0.95873833 | POL5 | V | 48471 | 51539 | -1 |
| YEL056W | 270.431035 | -3.2526399 | 0.49787075 | 37.3057663 | 7.93E-09 | 2.47E-07 | HAT2 | V | 47168 | 48373 | 1 |
| YEL057C | 248.856912 | -2.0102409 | 0.53051286 | 22.3024058 | 1.44E-05 | 0.00018033 | SDD1 | V | 45020 | 45721 | -1 |
| YEL058W | 117.167732 | 0.71573645 | 0.52063586 | 8.1443771 | 0.01704005 | 0.05876231 | PCM1 | V | 43252 | 44925 | 1 |

| | | | | | | | | | | | |
|---|---|---|---|---|---|---|---|---|---|---|---|
| YEL059C-A | 76.204023 | 1.61005888 | 0.55770788 | 8.87524218 | 0.01182403 | 0.04465829 | SOM1 | V | 42400 | 42624 | -1 |
| YEL059W | 982.020067 | 2.48383445 | 0.60146152 | 15.7357703 | 0.00038284 | 0.00297068 | NA | V | 42652 | 42960 | 1 |
| YEL060C | 603.363889 | 0.5466567 | 0.45563008 | 1.78442751 | 0.40974767 | 0.58133766 | PRB1 | V | 40046 | 41953 | -1 |
| YEL061C | 17.0691406 | 0.04863111 | 0.74117867 | 0.32559425 | 0.84976357 | 0.91079805 | CIN8 | V | 36535 | 39537 | -1 |
| YEL062W | 62.750231 | 1.64158912 | 0.58339297 | 15.8352929 | 0.00036426 | 0.00285226 | NPR2 | V | 34407 | 36254 | 1 |
| YEL063C | 327.659734 | 4.84841116 | 0.59674354 | 80.208346 | 3.83E-18 | 6.93E-16 | CAN1 | V | 31694 | 33466 | -1 |
| YEL064C | 79.2258103 | 0.63351696 | 0.56274427 | 5.66970174 | 0.05872728 | 0.14739097 | AVT2 | V | 29797 | 31239 | -1 |
| YEL065W | 25.2926885 | -1.2236707 | 0.96003925 | 5.16176646 | 0.07570711 | 0.17861909 | SIT1 | V | 27657 | 29543 | 1 |
| YEL066W | 35.9052292 | -3.062593 | 0.69540974 | 20.5606113 | 3.43E-05 | 0.00037758 | NA | V | 26667 | 27206 | 1 |
| YEL067C | 532.526596 | -1.6091061 | 0.59366937 | 8.53164215 | 0.01404033 | 0.0509358 | NA | V | 26189 | 26776 | -1 |
| YEL069C | 211.190071 | -1.1704095 | 0.71169152 | 2.71627758 | 0.25713892 | 0.42955405 | HXT13 | V | 21537 | 23231 | -1 |
| YEL071W | 50.0681482 | 0.61363924 | 0.74349154 | 3.29171546 | 0.19284708 | 0.34947875 | DLD3 | V | 16355 | 17845 | 1 |
| YEL073C | 12.6565869 | -0.7712003 | 0.83733972 | 2.62289299 | 0.26943004 | 0.44259056 | NA | V | 7230 | 7553 | -1 |
| YEL075C | 34.8718413 | 0.72963093 | 0.70712315 | 3.73898074 | 0.15420223 | 0.29860565 | NA | V | 5345 | 5713 | -1 |
| YEL077C | 42.2664748 | -0.722251 | 0.6348115 | 2.90977851 | 0.23342621 | 0.40082886 | NA | V | 264 | 4097 | -1 |
| YER001W | 18.0438259 | -0.9507835 | 0.86071131 | 1.38156815 | 0.50118295 | 0.66037092 | MNN1 | V | 153520 | 155808 | 1 |
| YER002W | 107.938359 | -1.3967565 | 0.72968311 | 11.7127518 | 0.0028616 | 0.0147146 | NOP16 | V | 156803 | 157498 | 1 |
| YER003C | 32.2388659 | -0.7824261 | 0.68559296 | 1.59072015 | 0.45141866 | 0.61775326 | PMI40 | V | 157736 | 159118 | -1 |
| YER004W | 37.2066582 | -0.2737438 | 0.62849761 | 0.21368581 | 0.89866684 | 0.9375281 | FMP52 | V | 159580 | 160275 | 1 |
| YER005W | 781.007076 | 0.04567896 | 0.48308326 | 0.01066792 | 0.99468024 | 0.99571207 | YND1 | V | 160550 | 162442 | 1 |
| YER006W | 32.7792136 | -2.1926191 | 0.83564747 | 6.74073012 | 0.03437709 | 0.09926596 | NUG1 | V | 162723 | 164285 | 1 |
| YER007C-A | 53.6393664 | 0.14806355 | 0.56545605 | 0.71594565 | 0.69909207 | 0.81772588 | TMA20 | V | 166237 | 166885 | -1 |
| YER007W | 264.13632 | -1.0192175 | 0.57179754 | 4.35060011 | 0.11357407 | 0.24017307 | PAC2 | V | 164527 | 166083 | 1 |
| YER008C | 128.39489 | -0.0009174 | 0.59276489 | 1.1809538 | 0.55406299 | 0.70568075 | SEC3 | V | 167808 | 171818 | -1 |
| YER009W | 25.1105998 | -1.5374148 | 0.86592584 | 3.32683992 | 0.18948982 | 0.34544902 | NTF2 | V | 172115 | 172492 | 1 |
| YER010C | 2347.02392 | -0.1377615 | 1.94600512 | 4.60506215 | 0.1000054 | 0.21919967 | NA | V | 172634 | 173338 | -1 |
| YER011W | 21.8325162 | 0.43157042 | 0.71332143 | 2.05120952 | 0.35857955 | 0.53153496 | TIR1 | V | 175248 | 176012 | 1 |
| YER012W | 18.955544 | -0.608835 | 0.81520534 | 0.77745964 | 0.67791741 | 0.80121286 | PRE1 | V | 177835 | 178431 | 1 |
| YER013W | 14.5510013 | 0.09082118 | 0.82857634 | 0.23945475 | 0.88716227 | 0.93215081 | PRP22 | V | 178841 | 182278 | 1 |
| YER014C-A | 1107.88932 | 0.63487722 | 0.48299623 | 2.10005343 | 0.3499284 | 0.52354248 | BUD25 | V | 183731 | 184700 | -1 |
| YER014W | 11.9309162 | 0.19417992 | 0.94940557 | 0.11470317 | 0.94426202 | 0.96220998 | HEM14 | V | 182600 | 184219 | 1 |
| YER015W | 570.477259 | 0.24294301 | 0.48161578 | 2.24390357 | 0.32564359 | 0.4985395 | FAA2 | V | 184541 | 186775 | 1 |
| YER016W | 52.078497 | -0.0280609 | 0.60348493 | 0.21980017 | 0.89592365 | 0.93584664 | BIM1 | V | 188277 | 189311 | 1 |

| | | | | | | | | | | | |
|---|---|---|---|---|---|---|---|---|---|---|---|
| YER017C | 38.1497047 | 0.28448692 | 0.76511731 | 1.84683475 | 0.39715948 | 0.56793119 | AFG3 | V | 189503 | 191788 | -1 |
| YER018C | 173.571655 | -0.3448986 | 0.57228728 | 0.52478845 | 0.76920772 | 0.85953436 | SPC25 | V | 191959 | 192624 | -1 |
| YER019C-A | 388.560547 | 0.05311032 | 0.6252457 | 7.25815164 | 0.0265407 | 0.08104993 | SBH2 | V | 194273 | 194539 | -1 |
| YER019W | 822.706836 | 0.56275206 | 0.78953179 | 2.18529322 | 0.33532784 | 0.50865815 | ISC1 | V | 192797 | 194230 | 1 |
| YER020W | 87.1020737 | 1.96071927 | 0.63040286 | 10.3431899 | 0.00567551 | 0.02535586 | GPA2 | V | 195168 | 196517 | 1 |
| YER021W | 31.1774024 | -0.7503303 | 0.74881824 | 1.07629292 | 0.5838294 | 0.73199919 | RPN3 | V | 196948 | 198519 | 1 |
| YER022W | 17.6630435 | 0.31832933 | 0.83781707 | 0.21813677 | 0.8966691 | 0.93645636 | SRB4 | V | 198812 | 200875 | 1 |
| YER023W | 29.8159364 | -0.9683886 | 0.6474442 | 2.30018483 | 0.31660751 | 0.48897239 | PRO3 | V | 201076 | 201936 | 1 |
| YER024W | 276.166078 | -2.4123307 | 0.48234345 | 24.1703354 | 5.64E-06 | 7.89E-05 | YAT2 | V | 202192 | 204963 | 1 |
| YER025W | 81.5488209 | -0.4167908 | 0.55616507 | 0.56390223 | 0.75431056 | 0.85152235 | GCD11 | V | 205251 | 206834 | 1 |
| YER026C | 112.311171 | 1.04575869 | 0.50724082 | 4.58623426 | 0.10095129 | 0.22031963 | CHO1 | V | 207644 | 208474 | -1 |
| YER027C | 20.6536608 | -0.5547806 | 0.87440551 | 0.57716269 | 0.74932585 | 0.8479727 | GAL83 | V | 208979 | 210232 | -1 |
| YER028C | 108.335541 | -0.9398463 | 0.51322188 | 3.70133919 | 0.15713192 | 0.30296164 | MIG3 | V | 210692 | 211876 | -1 |
| YER029C | 41.6116818 | -1.1666283 | 0.66620709 | 3.79218763 | 0.150154 | 0.29297995 | SMB1 | V | 212587 | 213177 | -1 |
| YER030W | 128.507429 | -1.229411 | 0.62700653 | 7.18437461 | 0.02753803 | 0.08334825 | CHZ1 | V | 213437 | 213898 | 1 |
| YER031C | 32.0906281 | -2.0082722 | 0.84574339 | 9.26302109 | 0.00974004 | 0.03862658 | YPT31 | V | 214076 | 214747 | -1 |
| YER032W | 54.1945932 | -0.7323082 | 0.6412619 | 1.37946746 | 0.50170964 | 0.66050451 | FIR1 | V | 215063 | 217693 | 1 |
| YER033C | 127.501385 | 1.20423083 | 0.57050607 | 5.61787164 | 0.0602691 | 0.15054274 | ZRG8 | V | 218057 | 221287 | -1 |
| YER034W | 54.535686 | 2.28745447 | 0.79216526 | 9.38361722 | 0.00917009 | 0.03689701 | NA | V | 221846 | 222403 | 1 |
| YER035W | 55.6139693 | 1.62905157 | 0.63729825 | 6.91553631 | 0.03149999 | 0.09285753 | EDC2 | V | 222639 | 223076 | 1 |
| YER036C | 116.155692 | -1.7133243 | 0.52128135 | 11.8629207 | 0.0026546 | 0.01388984 | ARB1 | V | 223367 | 225199 | -1 |
| YER037W | 704.687182 | -2.2353485 | 0.46056787 | 21.630009 | 2.01E-05 | 0.00023746 | PHM8 | V | 225889 | 226854 | 1 |
| YER038C | 1951.91049 | 3.09055038 | 0.63853118 | 20.0285059 | 4.48E-05 | 0.00046777 | KRE29 | V | 226858 | 228252 | -1 |
| YER038W-A | 563.277799 | 2.66329867 | 0.50644526 | 24.7053771 | 4.32E-06 | 6.30E-05 | FMP49 | V | 228451 | 228831 | 1 |
| YER039C | 72.7048313 | -2.7308386 | 0.5747558 | 22.1641096 | 1.54E-05 | 0.00019051 | HVG1 | V | 228456 | 229205 | -1 |
| YER039C-A | 21.0623843 | -1.7491033 | 0.77437651 | 5.70673855 | 0.05764976 | 0.14582441 | NA | V | 229263 | 229481 | -1 |
| YER040W | 17.8587822 | -0.2784537 | 0.83741655 | 2.78166488 | 0.24886805 | 0.41973376 | GLN3 | V | 229795 | 231987 | 1 |
| YER041W | 13.2349626 | 0.51834028 | 0.85777983 | 0.66189522 | 0.7182428 | 0.82928083 | YEN1 | V | 232461 | 234740 | 1 |
| YER042W | 16.2822964 | -2.001099 | 1.10411868 | 9.36558364 | 0.00925314 | 0.03719466 | MXR1 | V | 234937 | 235491 | 1 |
| YER043C | 80.0281878 | -1.1415383 | 0.7321566 | 2.49111365 | 0.28778062 | 0.46172289 | SAH1 | V | 235770 | 237119 | -1 |
| YER044C | 8.4274271 | -0.4386623 | 1.22432527 | 0.20699369 | 0.90167887 | 0.93824416 | ERG28 | V | 237570 | 238016 | -1 |
| YER044C-A | 152.775915 | 1.02484275 | 0.50162914 | 4.22745953 | 0.12078662 | 0.25039546 | MEI4 | V | 238460 | 239774 | -1 |
| YER045C | 523.137559 | -0.3002231 | 0.5033236 | 0.95799194 | 0.61940498 | 0.76030111 | ACA1 | V | 240032 | 241501 | -1 |

| | | | | | | | | | | | |
|---|---|---|---|---|---|---|---|---|---|---|---|
| YER046W | 40.3832792 | 0.09429601 | 0.78349183 | 0.22601126 | 0.89314563 | 0.93429326 | SPO73 | V | 243180 | 243611 | 1 |
| YER047C | 73.5719866 | 0.42525344 | 0.58311941 | 2.73795128 | 0.25436739 | 0.42652395 | SAP1 | V | 243810 | 246503 | -1 |
| YER048C | 66.8559971 | -0.2192223 | 0.705376 | 0.29645078 | 0.86223675 | 0.91923233 | CAJ1 | V | 246982 | 248157 | -1 |
| YER048W-A | 7.99170786 | 0.92997514 | 1.05071623 | 1.85695556 | 0.39515476 | 0.5664635 | ISD11 | V | 250718 | 251002 | 1 |
| YER049W | 35.2628584 | -1.1360154 | 0.72877491 | 2.79962569 | 0.24664312 | 0.41766412 | TPA1 | V | 251728 | 253662 | 1 |
| YER050C | 8.76293719 | -1.3930299 | 1.15592152 | 2.79949759 | 0.24665892 | 0.41766412 | RSM18 | V | 253971 | 254387 | -1 |
| YER051W | 61.2513181 | 1.94774187 | 0.61901801 | 12.7988505 | 0.00166251 | 0.00978886 | JHD1 | V | 254656 | 256134 | 1 |
| YER052C | 20.499073 | 0.11987872 | 0.74706507 | 2.27648392 | 0.32038177 | 0.49282956 | HOM3 | V | 256375 | 257958 | -1 |
| YER053C | 95.4406177 | 0.53341548 | 0.57065487 | 2.68618225 | 0.26103752 | 0.43235173 | PIC2 | V | 258737 | 259639 | -1 |
| YER054C | 109.06935 | -0.2032888 | 0.58396312 | 3.833026 | 0.14711907 | 0.28912102 | GIP2 | V | 262052 | 263698 | -1 |
| YER055C | 213.469682 | 0.11248 | 0.53787192 | 4.06184963 | 0.13121412 | 0.26636668 | HIS1 | V | 264892 | 265785 | -1 |
| YER056C | 24.7771548 | -1.2664038 | 0.71755093 | 4.33990184 | 0.11418322 | 0.24102109 | FCY2 | V | 266512 | 268113 | -1 |
| YER056C-A | 80.3118705 | -2.4969911 | 0.69491264 | 20.5169036 | 3.51E-05 | 0.00038446 | RPL34A | V | 269423 | 270185 | -1 |
| YER057C | 223.640579 | -2.8997918 | 0.5256874 | 29.7611304 | 3.45E-07 | 6.95E-06 | HMF1 | V | 270737 | 271126 | -1 |
| YER058W | 16.830535 | 0.00838378 | 0.73433076 | 0.83628535 | 0.6582683 | 0.78633659 | PET117 | V | 271768 | 272091 | 1 |
| YER059W | 54.9343005 | -2.1752323 | 0.58419736 | 13.4945962 | 0.00117405 | 0.00750302 | PCL6 | V | 272624 | 273886 | 1 |
| YER060W | 891.417469 | -0.4800311 | 0.57877577 | 0.70152236 | 0.7041519 | 0.82015637 | FCY21 | V | 274567 | 276153 | 1 |
| YER060W-A | 831.668374 | 0.02517218 | 0.73600646 | 0.09831076 | 0.95203319 | 0.96672609 | FCY22 | V | 276572 | 278164 | 1 |
| YER061C | 29.8078867 | -1.7732533 | 0.72737273 | 6.12972977 | 0.04666015 | 0.12398451 | CEM1 | V | 278298 | 279626 | -1 |
| YER062C | 167.394336 | 1.60537974 | 0.49416826 | 12.4572526 | 0.00197216 | 0.01104333 | GPP2 | V | 279930 | 280682 | -1 |
| YER063W | 503.126894 | -2.4491521 | 0.53282229 | 19.1267114 | 7.03E-05 | 0.00069655 | THO1 | V | 281710 | 282366 | 1 |
| YER064C | 310.361037 | 3.40790065 | 0.56137036 | 34.0426745 | 4.05E-08 | 1.07E-06 | VHR2 | V | 282705 | 284222 | -1 |
| YER065C | 742.721524 | -3.3448286 | 0.49628097 | 46.3846954 | 8.47E-11 | 4.12E-09 | ICL1 | V | 285241 | 286914 | -1 |
| YER067W | 15.8475702 | 0.37007504 | 0.89396851 | 3.16389159 | 0.2055747 | 0.36691662 | RGI1 | V | 292066 | 292551 | 1 |
| YER068W | 41.5307894 | -0.1112505 | 0.6080617 | 1.42234725 | 0.49106753 | 0.65227827 | MOT2 | V | 293050 | 294813 | 1 |
| YER069W | 106.025982 | 2.39804041 | 0.56831822 | 19.8211886 | 4.96E-05 | 0.00051177 | ARG5,6 | V | 295410 | 298001 | 1 |
| YER070W | 27.8771983 | -2.5244684 | 0.83332937 | 9.0516647 | 0.0108257 | 0.04184299 | RNR1 | V | 298950 | 301616 | 1 |
| YER071C | 53.9082163 | -1.1802149 | 0.58644817 | 4.15307598 | 0.12536347 | 0.25809632 | TDA2 | V | 301947 | 302327 | -1 |
| YER072W | 205.925721 | 1.14987857 | 0.48608753 | 12.8979994 | 0.0015821 | 0.00951154 | VTC1 | V | 302806 | 303195 | 1 |
| YER073W | 21.4982268 | -0.2636534 | 0.73785383 | 2.46359359 | 0.29176786 | 0.46615229 | ALD5 | V | 304030 | 305592 | 1 |
| YER074W | 125.098399 | -3.1899815 | 0.820965 | 13.292862 | 0.00129865 | 0.00812884 | RPS24A | V | 306323 | 307196 | 1 |
| YER074W-A | 41.0455274 | -0.8313197 | 0.60773829 | 2.55028854 | 0.27939066 | 0.45351049 | YOS1 | V | 307653 | 308123 | 1 |
| YER075C | 193.969549 | 2.38563176 | 0.50596943 | 28.1164522 | 7.84E-07 | 1.44E-05 | PTP3 | V | 308413 | 311199 | -1 |

| | | | | | | | | | | | |
|---|---|---|---|---|---|---|---|---|---|---|---|
| YER076C | 25.9793048 | -1.3827981 | 0.77219136 | 3.22773716 | 0.19911583 | 0.35792631 | NA | V | 312590 | 313498 | -1 |
| YER077C | 40.994989 | 0.47927185 | 0.68405992 | 0.91648199 | 0.63239505 | 0.76955658 | MRX1 | V | 314534 | 316600 | -1 |
| YER078C | 124.420809 | 1.37084463 | 0.51787857 | 7.16108057 | 0.02786064 | 0.08406103 | ICP55 | V | 316807 | 318342 | -1 |
| YER079W | 71.9595585 | -0.7392324 | 0.61574412 | 1.7459641 | 0.41770408 | 0.58926319 | NA | V | 318920 | 319552 | 1 |
| YER080W | 32.1726407 | -0.3904958 | 0.6477292 | 1.22375928 | 0.54233052 | 0.69625138 | AIM9 | V | 319963 | 321846 | 1 |
| YER081W | 15.1841454 | -0.6034194 | 0.97413617 | 2.19351257 | 0.33395258 | 0.5079834 | SER3 | V | 322686 | 324095 | 1 |
| YER082C | 183.879808 | 0.8503541 | 0.78873345 | 9.23824059 | 0.00986147 | 0.0390013 | UTP7 | V | 324272 | 325936 | -1 |
| YER083C | 20.8384642 | -1.345596 | 0.7352913 | 3.87157588 | 0.14431052 | 0.28488165 | GET2 | V | 326174 | 327031 | -1 |
| YER086W | 123.354459 | 2.33238957 | 0.57267416 | 18.2092969 | 0.00011115 | 0.0010215 | ILV1 | V | 328477 | 330207 | 1 |
| YER087C-B | 136.52163 | 1.21899204 | 0.54403894 | 5.03814171 | 0.0805344 | 0.18734197 | SBH1 | V | 332582 | 332830 | -1 |
| YER087W | 158.498974 | 2.15897495 | 0.58038811 | 12.8585911 | 0.00161359 | 0.00961783 | AIM10 | V | 330576 | 332306 | 1 |
| YER088C | 274.826881 | -0.8119102 | 0.51756918 | 8.43901666 | 0.01470587 | 0.05262485 | DOT6 | V | 333176 | 335188 | -1 |
| YER089C | 212.141068 | 0.45262297 | 0.48699582 | 3.93000409 | 0.1401556 | 0.27867477 | PTC2 | V | 335946 | 337340 | -1 |
| YER090W | 43.849504 | 0.35659494 | 0.58352126 | 0.44945763 | 0.79873279 | 0.87821171 | TRP2 | V | 337949 | 339472 | 1 |
| YER091C | 47.9214254 | 1.06101553 | 0.6074115 | 8.83884572 | 0.01204118 | 0.04533058 | MET6 | V | 339864 | 342167 | -1 |
| YER092W | 50.5999553 | 0.19473083 | 0.65836898 | 6.05766722 | 0.04837203 | 0.12713301 | IES5 | V | 342855 | 343232 | 1 |
| YER093C | 667.993694 | -0.4859563 | 0.47686671 | 3.90482227 | 0.14193144 | 0.28114371 | TSC11 | V | 343320 | 347612 | -1 |
| YER093C-A | 8.16772941 | 0.3404541 | 1.0369404 | 3.22437727 | 0.19945061 | 0.35830563 | AIM11 | V | 347912 | 348400 | -1 |
| YER094C | 31.0830692 | 0.19841635 | 0.63410692 | 0.31604408 | 0.85383096 | 0.91329785 | PUP3 | V | 348729 | 349346 | -1 |
| YER095W | 107.430999 | 2.21079615 | 0.56569165 | 14.9966055 | 0.00055402 | 0.00399975 | RAD51 | V | 349980 | 351182 | 1 |
| YER096W | 38.2973582 | 1.7765485 | 0.64182643 | 10.5622257 | 0.00508677 | 0.02326412 | SHC1 | V | 351698 | 353236 | 1 |
| YER098W | 281.276891 | -2.2858244 | 0.4970575 | 22.2014008 | 1.51E-05 | 0.00018804 | UBP9 | V | 355466 | 357730 | 1 |
| YER099C | 45.7971847 | 0.11770544 | 0.63864208 | 2.39625303 | 0.30175902 | 0.47448396 | PRS2 | V | 358105 | 359061 | -1 |
| YER100W | 15.2961059 | 0.54784323 | 0.81585966 | 0.77790226 | 0.67776739 | 0.8011991 | UBC6 | V | 359562 | 360314 | 1 |
| YER101C | 106.538811 | 1.28534914 | 0.55507661 | 9.18787421 | 0.01011296 | 0.03980562 | AST2 | V | 360502 | 361794 | -1 |
| YER102W | 76.8515991 | -3.2856234 | 0.62786158 | 25.2341191 | 3.31E-06 | 5.05E-05 | RPS8A | V | 363100 | 363702 | 1 |
| YER103W | 42.5748513 | 2.01051762 | 0.66197086 | 13.0051812 | 0.00149955 | 0.00912975 | SSA4 | V | 364589 | 366517 | 1 |
| YER104W | 205.489024 | -0.3505704 | 0.51922802 | 1.56093267 | 0.45819229 | 0.62328904 | RTT105 | V | 366802 | 367428 | 1 |
| YER105C | 204.95755 | -0.7793207 | 0.51181157 | 3.48502709 | 0.17507978 | 0.32688612 | NUP157 | V | 367838 | 372013 | -1 |
| YER107C | 17.0125847 | 0.31048258 | 0.75289576 | 0.50933162 | 0.77517551 | 0.86346022 | GLE2 | V | 373448 | 374545 | -1 |
| YER109C | 136.461112 | 1.17685853 | 0.57064285 | 10.7103193 | 0.00472372 | 0.02186276 | NA | NA | NA | NA | NA |
| YER110C | 67.4590919 | 0.26641096 | 0.55803549 | 3.1685125 | 0.20510028 | 0.36629569 | KAP123 | V | 378762 | 382103 | -1 |
| YER111C | 34.2479375 | 0.47141937 | 0.67847694 | 0.65914192 | 0.71923225 | 0.82938751 | SWI4 | V | 382595 | 385876 | -1 |

| | | | | | | | | | | | |
|---|---|---|---|---|---|---|---|---|---|---|---|
| YER112W | 81.5774478 | -1.3802905 | 0.9279584 | 8.1739764 | 0.01678972 | 0.05813116 | LSM4 | V | 387232 | 387795 | 1 |
| YER113C | 17.7845988 | -0.4793481 | 0.73071396 | 2.08539856 | 0.3525019 | 0.52616292 | TMN3 | V | 387932 | 390052 | -1 |
| YER114C | 47.2796174 | 1.74940458 | 0.73353092 | 6.68101773 | 0.03541893 | 0.10142216 | BOI2 | V | 390590 | 393712 | -1 |
| YER115C | 6.91550538 | 1.19284239 | 1.08587817 | 4.55305406 | 0.10264005 | 0.22312171 | SPR6 | V | 394292 | 394867 | -1 |
| YER116C | 35.8187016 | -1.8532246 | 0.70788998 | 7.40790697 | 0.02462598 | 0.07694787 | SLX8 | V | 395348 | 396172 | -1 |
| YER117W | 32.6679794 | -2.3506443 | 0.93836685 | 6.87410673 | 0.03215931 | 0.0942798 | RPL23A | V | 396769 | 397653 | 1 |
| YER118C | 23.2533793 | -1.1201905 | 0.88558149 | 3.67044552 | 0.15957795 | 0.3066566 | SHO1 | V | 397952 | 399055 | -1 |
| YER119C | 229.62588 | -0.333169 | 0.52185057 | 2.75511064 | 0.25219433 | 0.42365049 | AVT6 | V | 399496 | 400842 | -1 |
| YER120W | 111.245703 | -0.7785739 | 0.51413305 | 2.26204284 | 0.32270347 | 0.4957424 | SCS2 | V | 401135 | 401869 | 1 |
| YER121W | 75.5519252 | -0.339705 | 0.67143511 | 1.91331973 | 0.38417394 | 0.5573458 | NA | V | 402375 | 402719 | 1 |
| YER122C | 56.7675776 | -0.8315028 | 0.60160299 | 2.44684384 | 0.29422164 | 0.46865015 | GLO3 | V | 402871 | 404352 | -1 |
| YER123W | 11.0229101 | -1.6281264 | 1.03241603 | 2.60966363 | 0.27121815 | 0.44467869 | YCK3 | V | 404813 | 406387 | 1 |
| YER124C | 180.569537 | 0.62934535 | 0.49566387 | 2.78209129 | 0.248815 | 0.41973376 | DSE1 | V | 407342 | 409063 | -1 |
| YER125W | 61.7124911 | 0.13253112 | 0.6573781 | 0.68278947 | 0.71077828 | 0.82438173 | RSP5 | V | 410189 | 412618 | 1 |
| YER126C | 45.4125309 | -0.1876059 | 0.6299829 | 1.32015586 | 0.51681106 | 0.67350835 | NSA2 | V | 413394 | 414179 | -1 |
| YER127W | 34.6723543 | -1.4285793 | 0.77481737 | 3.71782824 | 0.15584176 | 0.30107568 | LCP5 | V | 414481 | 415554 | 1 |
| YER128W | 17.8297179 | 0.38860277 | 0.8497658 | 0.84141393 | 0.65658247 | 0.78549013 | VFA1 | V | 415859 | 416470 | 1 |
| YER129W | 60.5964095 | 1.2809785 | 0.61483632 | 5.11238177 | 0.07759976 | 0.18227288 | SAK1 | V | 417281 | 420709 | 1 |
| YER130C | 27.5464413 | -0.8647892 | 0.82590208 | 1.34201726 | 0.51119271 | 0.66829159 | COM2 | V | 421115 | 422446 | -1 |
| YER131W | 164.32768 | -1.8296104 | 0.59720557 | 8.96290404 | 0.01131697 | 0.04316551 | RPS26B | V | 423952 | 424311 | 1 |
| YER132C | 113.425834 | 2.04629497 | 0.53481234 | 14.9883319 | 0.00055632 | 0.00401133 | PMD1 | V | 425188 | 430449 | -1 |
| YER133W | 87.5429717 | -0.2865171 | 0.73387669 | 1.2013536 | 0.54844033 | 0.70063006 | GLC7 | V | 432495 | 433958 | 1 |
| YER134C | 27.567237 | 0.05050931 | 1.02163145 | 0.58460071 | 0.74654428 | 0.84572322 | NA | V | 437267 | 437803 | -1 |
| YER135C | 20.710723 | 1.78417672 | 0.75468197 | 6.66766967 | 0.03565611 | 0.10195005 | NA | V | 437952 | 438344 | -1 |
| YER136W | 25.5523655 | 0.55517123 | 0.80484194 | 0.88990421 | 0.64085497 | 0.77561669 | GDI1 | V | 439616 | 440971 | 1 |
| YER139C | 14.0625398 | -2.956378 | 0.882364 | 12.1770853 | 0.00226871 | 0.01229812 | RTR1 | V | 450563 | 451243 | -1 |
| YER140W | 19.9342804 | -0.7118155 | 0.71841373 | 1.00005253 | 0.60651473 | 0.75088539 | EMP65 | V | 451565 | 453235 | 1 |
| YER141W | 115.628197 | 0.30785508 | 0.51615223 | 0.35851966 | 0.83588868 | 0.9031862 | COX15 | V | 453459 | 454919 | 1 |
| YER142C | 365.812013 | 1.03418089 | 1.83613623 | 3.48636239 | 0.17496292 | 0.32678559 | MAG1 | V | 455146 | 456036 | -1 |
| YER143W | 237.726923 | -0.6761389 | 0.48294212 | 5.64877771 | 0.05934491 | 0.14861897 | DDI1 | V | 456319 | 457605 | 1 |
| YER144C | 74.2152784 | -0.558551 | 0.549314 | 2.00930101 | 0.36617259 | 0.53947564 | UBP5 | V | 457806 | 460223 | -1 |
| YER145C | 356.73164 | 1.00256604 | 0.60404122 | 3.31110018 | 0.19098697 | 0.34741267 | FTR1 | V | 460526 | 461740 | -1 |
| YER145C-A | 2194.74033 | -2.7659307 | 0.45633807 | 33.3456672 | 5.74E-08 | 1.48E-06 | NA | V | 462386 | 462823 | -1 |

| | | | | | | | | | | | |
|---|---|---|---|---|---|---|---|---|---|---|---|
| YER146W | 78.2436643 | 3.16022262 | 0.65650058 | 21.6783321 | 1.96E-05 | 0.00023274 | LSM5 | V | 462585 | 462866 | 1 |
| YER147C | 468.696204 | -1.4004164 | 0.51073237 | 7.6797415 | 0.02149638 | 0.06990254 | SCC4 | V | 462968 | 464842 | -1 |
| YER148W | 38.5568111 | -1.6097232 | 0.87658514 | 3.64959557 | 0.16125025 | 0.30925437 | SPT15 | V | 465303 | 466025 | 1 |
| YER149C | 169.133868 | -0.4441112 | 0.85734778 | 8.88051575 | 0.0117929 | 0.04456976 | PEA2 | V | 466208 | 467470 | -1 |
| YER150W | 132.564993 | 2.34396222 | 0.5159032 | 19.3215691 | 6.37E-05 | 0.00063406 | SPI1 | V | 468370 | 468816 | 1 |
| YER151C | 36.4183053 | -0.4427234 | 0.68640018 | 1.3093959 | 0.51959898 | 0.67578891 | UBP3 | V | 469686 | 472424 | -1 |
| YER152C | 47.1208303 | 2.94751831 | 0.65783366 | 20.0731907 | 4.38E-05 | 0.00045993 | NA | V | 472657 | 473988 | -1 |
| YER153C | 140.247832 | 1.3850972 | 0.63096155 | 7.22579712 | 0.02697355 | 0.08206876 | PET122 | V | 474041 | 474805 | -1 |
| YER154W | 451.471563 | -0.1067052 | 0.83395899 | 2.46084146 | 0.29216963 | 0.46640809 | OXA1 | V | 475020 | 476228 | 1 |
| YER155C | 91.0380692 | -0.6211933 | 0.6460467 | 3.00724011 | 0.22232388 | 0.38737744 | BEM2 | V | 476345 | 482848 | -1 |
| YER156C | 50.661402 | -2.509529 | 0.64012647 | 16.2005893 | 0.00030345 | 0.00247461 | NA | V | 483325 | 484341 | -1 |
| YER157W | 7.65318783 | -0.2611725 | 1.06528716 | 0.33672242 | 0.84504854 | 0.90911019 | COG3 | V | 484788 | 487193 | 1 |
| YER158C | 611.580793 | 0.53549179 | 0.52453351 | 2.89843111 | 0.23475437 | 0.40225741 | NA | V | 488857 | 490578 | -1 |
| YER159C | 292.110878 | 2.65444009 | 0.58380671 | 18.6210311 | 9.05E-05 | 0.0008601 | BUR6 | V | 491530 | 491958 | -1 |
| YER161C | 41.5724261 | 0.51836078 | 0.6223923 | 1.20370361 | 0.54779628 | 0.70011119 | SPT2 | V | 499347 | 500348 | -1 |
| YER162C | 148.051859 | 2.35576066 | 0.51560004 | 30.097458 | 2.91E-07 | 6.11E-06 | RAD4 | V | 500630 | 502894 | -1 |
| YER163C | 36.44188 | -0.2132527 | 0.62254514 | 0.16822671 | 0.91932704 | 0.94974417 | GCG1 | V | 503084 | 503782 | -1 |
| YER164W | 57.415717 | -1.1526619 | 0.61224405 | 3.99531774 | 0.13565249 | 0.27253845 | CHD1 | V | 505392 | 509798 | 1 |
| YER165W | 149.69727 | -0.9446401 | 0.5337269 | 3.99524673 | 0.13565731 | 0.27253845 | PAB1 | V | 510373 | 512106 | 1 |
| YER166W | 178.286397 | 0.53273703 | 0.54195953 | 3.94271701 | 0.13926753 | 0.2778632 | DNF1 | V | 512744 | 517459 | 1 |
| YER167W | 43.6177841 | 1.23303754 | 0.63534575 | 4.00449649 | 0.13503136 | 0.27175237 | BCK2 | V | 518216 | 520771 | 1 |
| YER168C | 14.1326525 | -0.1738312 | 0.80384112 | 0.11554003 | 0.943867 | 0.96197675 | CCA1 | V | 521029 | 522669 | -1 |
| YER169W | 33.2625146 | -0.0300071 | 0.64402967 | 0.16447046 | 0.92105527 | 0.95035295 | RPH1 | V | 523369 | 525759 | 1 |
| YER170W | 88.0582312 | -0.119411 | 0.60448383 | 0.05609901 | 0.97234023 | 0.97893408 | ADK2 | V | 525974 | 526651 | 1 |
| YER171W | 164.200431 | 1.55007506 | 0.64791784 | 5.73204882 | 0.05692479 | 0.14462243 | RAD3 | V | 527082 | 529418 | 1 |
| YER172C | 515.31783 | 2.65421479 | 0.532904 | 26.9157756 | 1.43E-06 | 2.43E-05 | BRR2 | V | 529530 | 536021 | -1 |
| YER173W | 13.9863034 | -1.4965558 | 0.8266393 | 3.25669735 | 0.19625339 | 0.35398975 | RAD24 | V | 536300 | 538279 | 1 |
| YER174C | 24.727032 | 0.34995187 | 1.07117679 | 6.60689919 | 0.03675615 | 0.10442499 | GRX4 | V | 538434 | 539168 | -1 |
| YER175C | 96.5024105 | 0.01656768 | 0.5440217 | 0.23381286 | 0.88966843 | 0.93317728 | TMT1 | V | 539464 | 540363 | -1 |
| YER175W-A | 257.230393 | -1.1101148 | 0.58548339 | 3.49977379 | 0.1737936 | 0.32507061 | NA | V | 540650 | 540814 | 1 |
| YER176W | 1271.50844 | 0.86942381 | 0.5616218 | 2.6927455 | 0.2601823 | 0.43177286 | ECM32 | V | 541690 | 545055 | 1 |
| YER177W | 284.608542 | 0.10295504 | 0.50071116 | 2.65562873 | 0.26505594 | 0.43735364 | BMH1 | V | 545611 | 546414 | 1 |
| YER178W | 39.7833935 | -0.6562918 | 0.79802394 | 0.90445319 | 0.63620999 | 0.77274089 | PDA1 | V | 546817 | 548079 | 1 |

| | | | | | | | | | | | |
|---|---|---|---|---|---|---|---|---|---|---|---|
| YER179W | 618.2578 | 1.52607365 | 0.57075271 | 6.87162126 | 0.0321993 | 0.09434916 | DMC1 | V | 548421 | 549517 | 1 |
| YER182W | 152.228797 | 1.21783631 | 1.9277869 | 2.75602617 | 0.25207892 | 0.42365049 | FMP10 | V | 552525 | 553259 | 1 |
| YER183C | 164.508521 | 1.81737488 | 0.95304066 | 8.33841023 | 0.01546455 | 0.05483143 | FAU1 | V | 553334 | 553969 | -1 |
| YER184C | 200.705848 | -0.349155 | 0.53710449 | 0.42218647 | 0.80969857 | 0.8858947 | TOG1 | V | 556296 | 558680 | -1 |
| YER185W | 32.3488614 | 1.7615769 | 0.82081446 | 4.42845034 | 0.10923812 | 0.23339068 | PUG1 | V | 559454 | 560365 | 1 |
| YER186C | 9.61271573 | 1.39397622 | 0.94190983 | 2.46837647 | 0.29107095 | 0.46518644 | NA | V | 561705 | 562625 | -1 |
| YER187W | 92.0359168 | 3.66292506 | 0.65821381 | 35.7480425 | 1.73E-08 | 4.86E-07 | NA | V | 566230 | 566655 | 1 |
| YER188C-A | 70.8344263 | 5.52661462 | 1.1067816 | 21.9724132 | 1.69E-05 | 0.00020426 | NA | V | 569608 | 569907 | -1 |
| YER188W | 19.4880764 | 1.42304972 | 0.77053831 | 7.84583484 | 0.01978329 | 0.06537972 | NA | V | 568040 | 568759 | 1 |
| YER189W | 60.8917943 | -0.7007088 | 0.57675118 | 2.95583818 | 0.22811187 | 0.39414138 | NA | V | 571155 | 571523 | 1 |
| YER190W | 18.2747319 | -2.4790541 | 0.84180803 | 10.5119789 | 0.00521618 | 0.02376215 | YRF1-2 | V | 571480 | 576525 | 1 |
| YFL001W | 43.4772716 | 2.24458589 | 0.74018712 | 9.04810933 | 0.01084496 | 0.04188948 | DEG1 | VI | 147131 | 148459 | 1 |
| YFL002C | 33.4237746 | 0.00603807 | 0.63805035 | 0.12949561 | 0.93730382 | 0.95917093 | SPB4 | VI | 145114 | 146934 | -1 |
| YFL003C | 93.51908 | -0.4215974 | 0.53122571 | 2.19722229 | 0.33333372 | 0.50736125 | MSH4 | VI | 134521 | 137157 | -1 |
| YFL004W | 43.2236981 | -0.8837993 | 0.61206297 | 2.6080258 | 0.27144034 | 0.44484562 | VTC2 | VI | 131810 | 134296 | 1 |
| YFL005W | 39.0120559 | -1.0552003 | 0.66176705 | 3.29223165 | 0.19279731 | 0.34947875 | SEC4 | VI | 130334 | 130981 | 1 |
| YFL007W | 200.668909 | -0.0518949 | 0.48021483 | 0.05284932 | 0.97392142 | 0.97984448 | BLM10 | VI | 123479 | 129910 | 1 |
| YFL008W | 51.0309369 | 1.00386333 | 0.62291816 | 4.53816053 | 0.10340724 | 0.22441077 | SMC1 | VI | 119429 | 123106 | 1 |
| YFL009W | 24.6214996 | 0.18330969 | 0.69647296 | 0.79267066 | 0.67278105 | 0.79774275 | CDC4 | VI | 116145 | 118484 | 1 |
| YFL010C | 163.385203 | 3.59764719 | 0.56411723 | 36.5929965 | 1.13E-08 | 3.34E-07 | WWM1 | VI | 115108 | 115743 | -1 |
| YFL010W-A | 6362.8586 | 0.44927285 | 0.46600203 | 2.08857228 | 0.35194297 | 0.52559964 | AUA1 | VI | 114990 | 115274 | 1 |
| YFL011W | 179.281868 | 3.98032171 | 0.66340272 | 31.1732447 | 1.70E-07 | 3.85E-06 | HXT10 | VI | 112345 | 113985 | 1 |
| YFL012W | 33.5108491 | -0.0766876 | 0.66276338 | 3.55543582 | 0.16902343 | 0.31888097 | NA | VI | 110647 | 111093 | 1 |
| YFL013C | 62.8349025 | 0.55376083 | 0.55253796 | 1.34324403 | 0.51087925 | 0.66829159 | IES1 | VI | 107852 | 109930 | -1 |
| YFL014W | 83.9443555 | 2.82402224 | 0.58284444 | 22.1624431 | 1.54E-05 | 0.00019051 | HSP12 | VI | 107256 | 107585 | 1 |
| YFL015C | 260.605321 | 3.32132185 | 0.55248869 | 33.3831811 | 5.64E-08 | 1.46E-06 | NA | VI | 106469 | 106963 | -1 |
| YFL016C | 119.704688 | 1.44770765 | 0.57943358 | 6.04258366 | 0.04873822 | 0.12803733 | MDJ1 | VI | 104701 | 106236 | -1 |
| YFL017C | 180.579195 | 0.42749267 | 0.5935474 | 2.19044313 | 0.33446549 | 0.50802806 | GNA1 | VI | 103983 | 104462 | -1 |
| YFL017W-A | 207.19516 | -1.0729698 | 0.76130614 | 5.41375101 | 0.06674503 | 0.16211061 | SMX2 | VI | 103699 | 103932 | 1 |
| YFL018C | 170.835586 | 3.17466348 | 0.57618281 | 37.1241141 | 8.68E-09 | 2.65E-07 | LPD1 | VI | 101628 | 103127 | -1 |
| YFL019C | 731.05901 | 4.86615698 | 0.6075863 | 51.2746877 | 7.34E-12 | 4.52E-10 | NA | VI | 100252 | 100605 | -1 |
| YFL020C | 8.42144089 | 3.44523026 | 1.23892588 | 10.9818647 | 0.004124 | 0.01964738 | PAU5 | VI | 99231 | 99599 | -1 |
| YFL021W | 356.018105 | 2.34270112 | 0.59546757 | 21.3770886 | 2.28E-05 | 0.00026461 | GAT1 | VI | 95966 | 97498 | 1 |

| | | | | | | | | | | | |
|---|---|---|---|---|---|---|---|---|---|---|---|
| YFL022C | 69.1371007 | 0.34011691 | 0.60542598 | 0.64382985 | 0.72475985 | 0.8342663 | FRS2 | VI | 93499 | 95010 | -1 |
| YFL023W | 46.6981626 | -1.5013792 | 0.82174765 | 3.42299985 | 0.18059471 | 0.33514211 | BUD27 | VI | 90986 | 93376 | 1 |
| YFL024C | 548.024339 | -2.1325511 | 0.47186759 | 20.2375107 | 4.03E-05 | 0.00043148 | EPL1 | VI | 87847 | 90345 | -1 |
| YFL025C | 23.9941525 | 0.66627355 | 0.72105897 | 4.28450187 | 0.11739031 | 0.24546419 | BST1 | VI | 84145 | 87234 | -1 |
| YFL026W | 139.186616 | 3.59661767 | 0.57097913 | 52.9500793 | 3.18E-12 | 2.07E-10 | STE2 | VI | 82580 | 83875 | 1 |
| YFL027C | 23.6084875 | 1.53038333 | 0.71352878 | 8.09744683 | 0.01744463 | 0.05976592 | GYP8 | VI | 80419 | 81912 | -1 |
| YFL028C | 12.4673956 | 0.20860859 | 0.90854874 | 3.34603657 | 0.18767974 | 0.34351531 | CAF16 | VI | 79344 | 80213 | -1 |
| YFL029C | 556.108056 | 1.9848101 | 0.57199707 | 11.2344192 | 0.00363477 | 0.01781991 | CAK1 | VI | 78055 | 79161 | -1 |
| YFL030W | 211.168915 | 0.83131125 | 0.5282682 | 4.22284135 | 0.12106585 | 0.25079473 | AGX1 | VI | 76831 | 77988 | 1 |
| YFL031W | 365.973031 | 1.44315493 | 0.59598768 | 7.99512566 | 0.01836033 | 0.06205856 | HAC1 | VI | 75179 | 76147 | 1 |
| YFL033C | 2432.97573 | -0.8728428 | 0.47885462 | 4.2401276 | 0.12002397 | 0.24929085 | RIM15 | VI | 69115 | 74427 | -1 |
| YFL034C-A | 56.2903263 | 2.32204486 | 0.62977578 | 13.3965604 | 0.00123303 | 0.00781955 | RPL22B | VI | 64243 | 64932 | -1 |
| YFL034C-B | 115.384806 | 2.6838749 | 0.67232357 | 14.5986965 | 0.00067598 | 0.00475567 | MOB2 | VI | 63016 | 63993 | -1 |
| YFL034W | 37.1725303 | 0.17593241 | 0.60658022 | 0.37283012 | 0.82992905 | 0.89868884 | MIL1 | VI | 65477 | 68698 | 1 |
| YFL036W | 15.4894288 | -0.5408966 | 0.8037805 | 2.48436374 | 0.28875351 | 0.46261284 | RPO41 | VI | 58782 | 62837 | 1 |
| YFL037W | 38.8773377 | -0.7939913 | 0.80043815 | 1.26392361 | 0.53154798 | 0.68590658 | TUB2 | VI | 56336 | 57709 | 1 |
| YFL038C | 102.743957 | -0.1074815 | 0.53375621 | 2.09160444 | 0.3514098 | 0.52507425 | YPT1 | VI | 55366 | 55986 | -1 |
| YFL039C | 332.45909 | -0.0436334 | 0.69340027 | 0.87367394 | 0.64607675 | 0.77803336 | ACT1 | VI | 53260 | 54696 | -1 |
| YFL040W | 1229.23072 | 2.74708682 | 0.50542596 | 26.1408681 | 2.11E-06 | 3.36E-05 | NA | VI | 51350 | 52972 | 1 |
| YFL041W | 12.6362744 | -0.0050265 | 0.9140627 | 0.58266132 | 0.74726855 | 0.84631426 | FET5 | VI | 49139 | 51007 | 1 |
| YFL042C | 25.6410458 | 1.6600962 | 0.76090915 | 4.8436064 | 0.08876142 | 0.20185727 | LAM5 | VI | 45720 | 47744 | -1 |
| YFL044C | 12.4536544 | -0.3068933 | 0.99626612 | 0.33096102 | 0.84748638 | 0.90980423 | OTU1 | VI | 44655 | 45560 | -1 |
| YFL045C | 51.8078165 | -1.0684285 | 0.75071177 | 2.55366053 | 0.27892 | 0.45338204 | SEC53 | VI | 43628 | 44392 | -1 |
| YFL046W | 26.2801263 | -0.9643548 | 0.69404678 | 4.11069311 | 0.12804845 | 0.26151694 | FMP32 | VI | 42815 | 43438 | 1 |
| YFL047W | 17.3574823 | -0.0183257 | 0.93101621 | 0.87858039 | 0.64449372 | 0.77725863 | RGD2 | VI | 40421 | 42565 | 1 |
| YFL048C | 29.8704495 | -0.3620741 | 0.6482836 | 0.43313363 | 0.80527873 | 0.88306133 | EMP47 | VI | 38843 | 40180 | -1 |
| YFL049W | 12.3242104 | 1.07958372 | 0.91830734 | 1.60403604 | 0.44842312 | 0.61504128 | SWP82 | VI | 36803 | 38674 | 1 |
| YFL050C | 60.650699 | 3.38534354 | 0.61960905 | 35.0146898 | 2.49E-08 | 6.81E-07 | ALR2 | VI | 33272 | 35848 | -1 |
| YFL051C | 95.0709695 | 0.80984106 | 0.57677514 | 4.04865818 | 0.13208243 | 0.26749117 | NA | VI | 30058 | 30540 | -1 |
| YFL052W | 193.903239 | -2.2678338 | 0.49158047 | 20.9370553 | 2.84E-05 | 0.00032262 | ZNF1 | VI | 28232 | 29629 | 1 |
| YFL053W | 485.363874 | -0.0677387 | 0.51916509 | 1.25248107 | 0.53459783 | 0.68922766 | DAK2 | VI | 23423 | 25198 | 1 |
| YFL054C | 30.8251295 | 2.30436544 | 0.72737833 | 19.0165331 | 7.42E-05 | 0.00072613 | AQY3 | VI | 20847 | 22787 | -1 |
| YFL055W | 148.302991 | 2.04681393 | 0.52104579 | 16.5789445 | 0.00025115 | 0.0020833 | AGP3 | VI | 17004 | 18680 | 1 |

| | | | | | | | | | | | |
|---|---|---|---|---|---|---|---|---|---|---|---|
| YFL056C | 11.7863485 | 1.29094307 | 0.99411072 | 2.09951305 | 0.35002296 | 0.52354248 | NA | VI | 14305 | 15431 | -1 |
| YFL059W | 35.7601537 | 1.88691408 | 0.67510912 | 7.63632836 | 0.02196809 | 0.07113829 | SNZ3 | VI | 11363 | 12259 | 1 |
| YFL060C | 7.65161202 | 0.80002292 | 1.00812846 | 0.62635656 | 0.73111956 | 0.83700349 | SNO3 | VI | 10301 | 10969 | -1 |
| YFL061W | 146.187721 | -0.1684889 | 0.57311804 | 3.54061024 | 0.17028102 | 0.32052333 | DDI3 | VI | 9545 | 10222 | 1 |
| YFL062W | 262.929821 | 4.72745841 | 0.61455827 | 53.4453943 | 2.48E-12 | 1.63E-10 | COS4 | VI | 6426 | 7565 | 1 |
| YFL064C | 16.5014881 | -1.661812 | 0.85276217 | 7.78556552 | 0.02038853 | 0.06703554 | NA | VI | 3322 | 3846 | -1 |
| YFL066C | 5.50556082 | -1.0534808 | 1.26337321 | 0.88028052 | 0.64394609 | 0.77686719 | NA | VI | 1437 | 2615 | -1 |
| YFL067W | 1321.13539 | -0.1016207 | 0.56075549 | 0.14095644 | 0.93194804 | 0.95724306 | NA | VI | 836 | 1363 | 1 |
| YFR001W | 43.6172453 | -2.069874 | 0.77687154 | 7.40190828 | 0.02469995 | 0.07709579 | LOC1 | VI | 149110 | 149724 | 1 |
| YFR002W | 280.452052 | 0.61002935 | 0.68948621 | 4.3607748 | 0.11299775 | 0.23947912 | NIC96 | VI | 150016 | 152535 | 1 |
| YFR003C | 33.523676 | -1.1861225 | 0.70915274 | 3.60812532 | 0.1646287 | 0.31355268 | YPI1 | VI | 152657 | 153124 | -1 |
| YFR004W | 53.4492989 | -0.2434866 | 0.64215568 | 2.3782414 | 0.30448888 | 0.47725789 | RPN11 | VI | 153393 | 154313 | 1 |
| YFR005C | 32.3799709 | 1.32315897 | 0.64476973 | 6.63792254 | 0.0361904 | 0.10317205 | SAD1 | VI | 154527 | 155873 | -1 |
| YFR006W | 36.3753694 | -0.1961506 | 0.65986498 | 0.6750165 | 0.71354609 | 0.82562587 | NA | VI | 156145 | 157752 | 1 |
| YFR007W | 148.949977 | 0.15320487 | 0.58352983 | 0.61107343 | 0.73672786 | 0.83969573 | YFH7 | VI | 159299 | 160360 | 1 |
| YFR009W | 17.1324887 | -1.1446447 | 0.93779937 | 2.67855719 | 0.26203463 | 0.43337041 | GCN20 | VI | 162488 | 164746 | 1 |
| YFR010W | 42.4636765 | -0.6901253 | 0.59789387 | 2.1792435 | 0.33634369 | 0.50953165 | UBP6 | VI | 165067 | 166566 | 1 |
| YFR011C | 11.5798321 | -0.2930862 | 0.93992599 | 2.28729336 | 0.31865487 | 0.49147887 | MIC19 | VI | 166746 | 167258 | -1 |
| YFR012W | 68.3639453 | 0.78109009 | 0.72647518 | 1.23293608 | 0.5398478 | 0.69395761 | DCV1 | VI | 167888 | 168496 | 1 |
| YFR013W | 22.6032938 | -0.9983632 | 0.70768273 | 2.1409614 | 0.34284367 | 0.51654043 | IOC3 | VI | 169922 | 172285 | 1 |
| YFR014C | 74.9792887 | -1.3619924 | 0.53319576 | 6.36455268 | 0.0414911 | 0.11428804 | CMK1 | VI | 172537 | 173877 | -1 |
| YFR015C | 290.139673 | 1.59249293 | 0.51268078 | 9.48074827 | 0.00873538 | 0.03551814 | GSY1 | VI | 174265 | 176391 | -1 |
| YFR016C | 88.920922 | 1.04530331 | 0.53263425 | 4.56014014 | 0.10227704 | 0.22277848 | NA | VI | 177042 | 180743 | -1 |
| YFR017C | 16.5912517 | 2.84047898 | 0.87705881 | 11.3944122 | 0.00335533 | 0.016633 | IGD1 | VI | 182274 | 182861 | -1 |
| YFR018C | 11.315605 | 0.80819067 | 0.89630855 | 1.12337264 | 0.57024663 | 0.72074394 | NA | VI | 183131 | 184222 | -1 |
| YFR019W | 43.8226135 | 0.46520836 | 0.59584562 | 0.93631049 | 0.62615631 | 0.76437804 | FAB1 | VI | 184502 | 191338 | 1 |
| YFR020W | 10.685797 | 2.00497215 | 0.94440781 | 5.87666346 | 0.052954 | 0.13651097 | CSS2 | VI | 192737 | 193435 | 1 |
| YFR021W | 102.74926 | 0.41109309 | 0.57425813 | 1.77326219 | 0.41204155 | 0.58387679 | ATG18 | VI | 194812 | 196314 | 1 |
| YFR022W | 81.4242109 | 1.8716888 | 0.57005912 | 10.2849226 | 0.00584329 | 0.02594528 | ROG3 | VI | 196833 | 199034 | 1 |
| YFR023W | 16.1848523 | -1.1454924 | 1.01168794 | 3.28170092 | 0.19381514 | 0.35057471 | PES4 | VI | 199874 | 201709 | 1 |
| YFR024C-A | 144.391475 | 3.44041258 | 0.58410485 | 41.0887951 | 1.20E-09 | 4.35E-08 | LSB3 | VI | 201960 | 203433 | -1 |
| YFR025C | 13.8681871 | 0.31356294 | 0.83238893 | 0.5493362 | 0.75982427 | 0.85451323 | HIS2 | VI | 203743 | 204750 | -1 |
| YFR026C | 158.938611 | 3.23230547 | 0.5609149 | 31.5971846 | 1.38E-07 | 3.19E-06 | ULI1 | VI | 205748 | 206257 | -1 |

| | | | | | | | | | | | |
|---|---|---|---|---|---|---|---|---|---|---|---|
| YFR027W | 30.6228025 | 0.18720616 | 0.66543084 | 0.320159 | 0.85207605 | 0.91195795 | ECO1 | VI | 207452 | 208297 | 1 |
| YFR028C | 38.5112836 | 0.02391461 | 0.69326685 | 0.93259971 | 0.62731915 | 0.76483005 | CDC14 | VI | 208413 | 210068 | -1 |
| YFR029W | 43.6420082 | 1.33701772 | 0.64326905 | 4.51904635 | 0.10440025 | 0.22597288 | PTR3 | VI | 210937 | 212973 | 1 |
| YFR030W | 101.603476 | 2.36774896 | 0.70091744 | 20.4602889 | 3.61E-05 | 0.00039327 | MET10 | VI | 213312 | 216419 | 1 |
| YFR031C | 29.9127664 | 0.06830465 | 0.64382632 | 0.62775131 | 0.73060987 | 0.83678999 | SMC2 | VI | 216594 | 220106 | -1 |
| YFR031C-A | 204.524806 | -1.3681034 | 0.56597569 | 5.91696414 | 0.05189763 | 0.13440139 | RPL2A | VI | 220507 | 221418 | -1 |
| YFR032C-A | 367.372364 | -3.544774 | 0.7713608 | 17.6463174 | 0.00014728 | 0.00131397 | RPL29 | VI | 223258 | 223437 | -1 |
| YFR032C-B | 18.5022029 | -2.695835 | 0.7841113 | 11.437847 | 0.00328324 | 0.01637994 | NA | VI | 223710 | 223973 | -1 |
| YFR033C | 743.646847 | 0.27478984 | 0.67897951 | 6.90276505 | 0.03170178 | 0.09322158 | QCR6 | VI | 224326 | 224769 | -1 |
| YFR034C | 55.6327479 | -1.6999717 | 0.57209515 | 10.9266742 | 0.00423938 | 0.02007035 | PHO4 | VI | 225020 | 225958 | -1 |
| YFR035C | 683.141128 | -2.2839463 | 0.48014314 | 20.7998264 | 3.04E-05 | 0.00034284 | NA | VI | 226121 | 226465 | -1 |
| YFR036W | 90.5633481 | 1.44705079 | 0.57751225 | 7.35274378 | 0.02531465 | 0.0784226 | CDC26 | VI | 226963 | 227337 | 1 |
| YFR037C | 106.730413 | -1.5426701 | 0.52398066 | 8.45930978 | 0.01455741 | 0.05238497 | RSC8 | VI | 227513 | 229186 | -1 |
| YFR038W | 29.5797463 | -0.3633066 | 0.66464393 | 0.48328369 | 0.7853374 | 0.87009253 | IRC5 | VI | 229380 | 231941 | 1 |
| YFR039C | 102.783252 | -0.1947786 | 0.72563488 | 5.69457641 | 0.0580014 | 0.14626659 | OSW7 | VI | 232012 | 233544 | -1 |
| YFR040W | 88.3240989 | 0.55028156 | 0.76892597 | 9.9615406 | 0.00686877 | 0.0295909 | SAP155 | VI | 234242 | 237250 | 1 |
| YFR041C | 55.9381398 | 1.31032411 | 0.60261045 | 4.72451935 | 0.09420711 | 0.20963072 | ERJ5 | VI | 237368 | 238255 | -1 |
| YFR042W | 26.1623037 | 0.94192026 | 0.99904412 | 7.13533258 | 0.02822164 | 0.08488482 | KEG1 | VI | 238458 | 239060 | 1 |
| YFR043C | 251.376006 | -0.6812425 | 0.60825625 | 8.14749518 | 0.01701351 | 0.05874074 | IRC6 | VI | 239114 | 239827 | -1 |
| YFR044C | 27.5080709 | 0.09126488 | 0.73163018 | 0.65389281 | 0.72112239 | 0.83085787 | DUG1 | VI | 239992 | 241437 | -1 |
| YFR045W | 145.598109 | 0.33398144 | 0.56252973 | 0.47830699 | 0.78729403 | 0.87059443 | NA | VI | 241998 | 242999 | 1 |
| YFR046C | 61.5709143 | -0.2410472 | 0.75537436 | 3.0382777 | 0.21890031 | 0.38305402 | CNN1 | VI | 243074 | 244159 | -1 |
| YFR047C | 16.0645305 | 0.29400766 | 0.86297785 | 0.2484314 | 0.88318932 | 0.93100373 | BNA6 | VI | 244279 | 245166 | -1 |
| YFR048W | 19.2180104 | -0.2888649 | 0.77802187 | 1.74447547 | 0.41801509 | 0.58931273 | RMD8 | VI | 246146 | 248134 | 1 |
| YFR050C | 25.2914112 | -0.6331017 | 0.73836709 | 0.84461982 | 0.65553085 | 0.78495521 | PRE4 | VI | 249066 | 249866 | -1 |
| YFR051C | 60.3495951 | 0.10081638 | 0.56559482 | 1.89991044 | 0.38675834 | 0.55927343 | RET2 | VI | 250163 | 251803 | -1 |
| YFR052W | 16.8714588 | 0.02083157 | 0.74632052 | 0.0017084 | 0.99914617 | 0.99914617 | RPN12 | VI | 252505 | 253329 | 1 |
| YFR053C | 13.0244428 | -0.3394556 | 0.83878588 | 0.84734054 | 0.6546397 | 0.78442961 | HXK1 | VI | 253592 | 255049 | -1 |
| YFR054C | 27.9680353 | -1.9265408 | 0.70087432 | 8.30080078 | 0.01575811 | 0.05576982 | NA | VI | 258855 | 259433 | -1 |
| YFR055W | 35.243253 | -1.1434487 | 0.69140495 | 3.36776375 | 0.1856519 | 0.34157118 | IRC7 | VI | 264204 | 265226 | 1 |
| YFR057W | 5.74916313 | 3.09563101 | 1.46526149 | 4.60778321 | 0.09986943 | 0.21918634 | NA | VI | 269061 | 269516 | 1 |
| YGL001C | 18.2355552 | 0.52949487 | 0.86527956 | 0.49610238 | 0.78032 | 0.86685587 | ERG26 | VII | 495453 | 496502 | -1 |
| YGL002W | 94.1730664 | 1.11161085 | 0.57343681 | 5.36202146 | 0.06849389 | 0.16558648 | ERP6 | VII | 494517 | 495167 | 1 |

| | | | | | | | | | | | |
|---|---|---|---|---|---|---|---|---|---|---|---|
| YGL003C | 69.7960078 | 1.34309622 | 0.69803021 | 8.61358006 | 0.01347674 | 0.04923049 | CDH1 | VII | 492474 | 494174 | -1 |
| YGL004C | 99.2284674 | -0.1892413 | 0.52062474 | 0.15599634 | 0.92496612 | 0.95277599 | RPN14 | VII | 490703 | 491956 | -1 |
| YGL005C | 18.6169006 | -0.316709 | 0.72740104 | 0.22357889 | 0.89423252 | 0.9350658 | COG7 | VII | 489708 | 490547 | -1 |
| YGL006W | 58.2576805 | -0.0435843 | 0.70967012 | 0.505838 | 0.77653078 | 0.8639961 | PMC1 | VII | 485921 | 489442 | 1 |
| YGL006W-A | 23.4880338 | 1.88853481 | 0.75048765 | 7.79330031 | 0.02030983 | 0.06685272 | NA | VII | 485423 | 485533 | 1 |
| YGL007C-A | 11.8198026 | -0.8127139 | 0.85501405 | 0.93358087 | 0.62701148 | 0.76477701 | NA | VII | 482941 | 483027 | -1 |
| YGL007W | 129.288991 | -0.5129117 | 0.56212967 | 0.8448618 | 0.65545154 | 0.78495521 | NA | VII | 483260 | 483637 | 1 |
| YGL008C | 258.808259 | -2.3811791 | 0.72182349 | 10.092885 | 0.00643218 | 0.02793871 | PMA1 | VII | 479910 | 482666 | -1 |
| YGL009C | 15.3534269 | 0.46929002 | 0.92168455 | 2.45150306 | 0.29353702 | 0.46794585 | LEU1 | VII | 476313 | 478652 | -1 |
| YGL010W | 49.0758573 | 0.6790302 | 0.63214106 | 1.989782 | 0.36976375 | 0.54308878 | MPO1 | VII | 475545 | 476069 | 1 |
| YGL011C | 45.0174907 | 0.29240401 | 0.59536853 | 0.26225704 | 0.87710504 | 0.92756862 | SCL1 | VII | 474489 | 475247 | -1 |
| YGL012W | 271.898275 | 1.2761538 | 0.48412547 | 7.5756963 | 0.02264428 | 0.07271789 | ERG4 | VII | 472855 | 474276 | 1 |
| YGL013C | 65.29006 | -2.045156 | 0.55018655 | 13.8375965 | 0.00098902 | 0.00652211 | PDR1 | VII | 469092 | 472298 | -1 |
| YGL014W | 26.4384243 | 0.16587006 | 0.67842841 | 0.62756808 | 0.73067681 | 0.83678999 | PUF4 | VII | 466141 | 468807 | 1 |
| YGL015C | 88.4477818 | -1.5926076 | 0.92187588 | 11.7064 | 0.0028707 | 0.01473523 | NA | VII | 465043 | 465435 | -1 |
| YGL016W | 17.819243 | 0.35268397 | 0.77415564 | 0.26987018 | 0.87377263 | 0.92573532 | KAP122 | VII | 461666 | 464911 | 1 |
| YGL017W | 18.9771416 | -1.0838503 | 0.79273654 | 2.4784871 | 0.28960321 | 0.46358932 | ATE1 | VII | 459854 | 461365 | 1 |
| YGL018C | 38.4878818 | -0.4016786 | 1.0818765 | 9.53281947 | 0.00851088 | 0.03484104 | JAC1 | VII | 459110 | 459664 | -1 |
| YGL019W | 238.104768 | -1.5870234 | 0.4917316 | 10.8936745 | 0.00430991 | 0.02028813 | CKB1 | VII | 458156 | 458992 | 1 |
| YGL020C | 64.0521661 | -1.2697825 | 0.97504371 | 6.9035237 | 0.03168975 | 0.09322158 | GET1 | VII | 457163 | 457870 | -1 |
| YGL021W | 79.4356268 | 0.42175521 | 0.55614355 | 1.41884388 | 0.49192848 | 0.6530873 | ALK1 | VII | 454785 | 457067 | 1 |
| YGL022W | 60.1428798 | -0.4745654 | 0.72996929 | 0.71040781 | 0.70103049 | 0.81801018 | STT3 | VII | 452404 | 454560 | 1 |
| YGL023C | 78.7708793 | 0.53934626 | 0.59628151 | 1.70660988 | 0.42600468 | 0.59578916 | PIB2 | VII | 450197 | 452104 | -1 |
| YGL025C | 49.0243762 | -1.2231964 | 0.63473875 | 4.43260204 | 0.1090116 | 0.23299267 | PGD1 | VII | 448764 | 449957 | -1 |
| YGL026C | 56.9355815 | 0.5560115 | 0.66167521 | 1.81844023 | 0.40283827 | 0.5737844 | TRP5 | VII | 446412 | 448535 | -1 |
| YGL027C | 97.5325178 | -2.1572225 | 0.55182268 | 15.376968 | 0.00045807 | 0.00342224 | CWH41 | VII | 443642 | 446143 | -1 |
| YGL028C | 43.8483897 | 1.93847948 | 0.70746963 | 11.2718932 | 0.0035673 | 0.01754856 | SCW11 | VII | 441281 | 442909 | -1 |
| YGL029W | 53.0829909 | -2.980005 | 0.605023 | 23.7433666 | 6.99E-06 | 9.54E-05 | CGR1 | VII | 440063 | 440425 | 1 |
| YGL030W | 52.0978126 | -2.6455712 | 0.66869291 | 15.4363691 | 0.00044467 | 0.00334801 | RPL30 | VII | 439091 | 439638 | 1 |
| YGL031C | 101.672727 | -3.38676 | 0.76588695 | 17.1266849 | 0.00019098 | 0.00163818 | RPL24A | VII | 437467 | 437934 | -1 |
| YGL034C | 50.7483735 | -0.9544954 | 0.62792486 | 3.78630373 | 0.1505964 | 0.2934881 | NA | VII | 433214 | 433579 | -1 |
| YGL035C | 12.2809515 | -0.8957591 | 0.8799481 | 1.9154664 | 0.38376181 | 0.55688744 | MIG1 | VII | 431548 | 433062 | -1 |
| YGL036W | 91.723337 | 1.22932591 | 0.5411042 | 5.8488582 | 0.05369534 | 0.13805329 | NA | VII | 428601 | 431330 | 1 |

| | | | | | | | | | | | |
|---|---|---|---|---|---|---|---|---|---|---|---|
| YGL037C | 83.6235943 | 0.85795672 | 0.62407436 | 2.18087672 | 0.33606914 | 0.50936213 | PNC1 | VII | 427297 | 427947 | -1 |
| YGL038C | 33.3839526 | 0.5349616 | 0.8288764 | 0.52320405 | 0.76981733 | 0.85980755 | OCH1 | VII | 425362 | 426804 | -1 |
| YGL039W | 80.2761654 | -0.2520374 | 0.53040766 | 0.22960253 | 0.89154331 | 0.93352778 | NA | VII | 423961 | 425007 | 1 |
| YGL040C | 63.0124445 | 0.37099184 | 0.58868531 | 1.79565903 | 0.40745307 | 0.57879129 | HEM2 | VII | 419527 | 420555 | -1 |
| YGL041C-B | 573.0519 | -1.5370098 | 0.4929525 | 9.3532974 | 0.00931016 | 0.03735679 | NA | VII | 418704 | 418886 | -1 |
| YGL041W-A | 58.1610952 | 0.17555231 | 0.57685363 | 0.23502883 | 0.8891277 | 0.93304075 | NA | VII | 418825 | 419289 | 1 |
| YGL043W | 22.7465027 | -0.0961709 | 0.79145982 | 1.08635002 | 0.58090095 | 0.72940182 | DST1 | VII | 417483 | 418412 | 1 |
| YGL044C | 58.3989643 | 0.92096658 | 0.60324809 | 3.09542334 | 0.21273422 | 0.37639175 | RNA15 | VII | 416146 | 417036 | -1 |
| YGL045W | 42.1145364 | 2.30470982 | 0.81527074 | 12.0534229 | 0.00241342 | 0.01287898 | RIM8 | VII | 414102 | 415730 | 1 |
| YGL047W | 8.46088943 | -0.2174435 | 1.05735031 | 0.5982417 | 0.7414698 | 0.84243165 | ALG13 | VII | 411552 | 412160 | 1 |
| YGL048C | 59.4679122 | 0.56705113 | 0.74114896 | 1.10716215 | 0.57488741 | 0.72422968 | RPT6 | VII | 410069 | 411286 | -1 |
| YGL049C | 105.279249 | -0.1377327 | 0.51525116 | 0.54807512 | 0.76030352 | 0.85478784 | TIF4632 | VII | 406860 | 409604 | -1 |
| YGL050W | 249.523882 | -0.3044504 | 0.50235976 | 1.03990744 | 0.59454806 | 0.74174387 | TYW3 | VII | 405776 | 406597 | 1 |
| YGL051W | 720.08248 | 0.3068256 | 0.49398399 | 2.85800967 | 0.23954719 | 0.40781483 | MST27 | VII | 403687 | 404391 | 1 |
| YGL053W | 223.542802 | -1.6162605 | 0.4815705 | 11.3534898 | 0.00342469 | 0.01690447 | PRM8 | VII | 402589 | 403302 | 1 |
| YGL054C | 9.12453776 | -1.0698608 | 0.96925789 | 1.27718469 | 0.52803519 | 0.68243834 | ERV14 | VII | 400871 | 401287 | -1 |
| YGL055W | 123.108241 | -0.4128228 | 0.55282114 | 1.89723417 | 0.38727622 | 0.55960303 | OLE1 | VII | 398628 | 400160 | 1 |
| YGL056C | 139.074079 | 3.29890412 | 0.65598715 | 24.8998263 | 3.92E-06 | 5.80E-05 | SDS23 | VII | 396035 | 397618 | -1 |
| YGL057C | 69.8702618 | -0.0546116 | 0.55797945 | 8.70440795 | 0.0128784 | 0.04764596 | GEP7 | VII | 394966 | 395829 | -1 |
| YGL058W | 43.4682959 | 0.36234409 | 0.68594945 | 1.85247259 | 0.39604149 | 0.56717295 | RAD6 | VII | 393986 | 394504 | 1 |
| YGL059W | 162.978222 | 2.49867425 | 0.53555668 | 23.3988694 | 8.30E-06 | 0.00011148 | PKP2 | VII | 392223 | 393698 | 1 |
| YGL060W | 16.4445429 | -0.1270511 | 0.80298232 | 0.68329442 | 0.71059885 | 0.82438173 | YBP2 | VII | 390065 | 391990 | 1 |
| YGL061C | 240.52493 | 2.05789951 | 0.49730596 | 19.1484129 | 6.95E-05 | 0.00069022 | DUO1 | VII | 388968 | 389711 | -1 |
| YGL062W | 91.211938 | 0.94518831 | 0.60691381 | 2.44491979 | 0.29450482 | 0.46884326 | PYC1 | VII | 385196 | 388732 | 1 |
| YGL063W | 139.128618 | -0.8177781 | 0.52899092 | 4.19468707 | 0.12278216 | 0.25398668 | PUS2 | VII | 383654 | 384766 | 1 |
| YGL064C | 106.605238 | 0.59903759 | 0.58878875 | 5.21162938 | 0.07384295 | 0.1754414 | MRH4 | VII | 381523 | 383208 | -1 |
| YGL065C | 375.732108 | -0.0061378 | 0.6337778 | 2.12925176 | 0.34485685 | 0.51822506 | ALG2 | VII | 379760 | 381271 | -1 |
| YGL066W | 120.231152 | 2.57992198 | 0.57291589 | 24.1824759 | 5.61E-06 | 7.86E-05 | SGF73 | VII | 377609 | 379582 | 1 |
| YGL067W | 20.0323337 | -1.9742674 | 0.79609218 | 6.16725724 | 0.04579279 | 0.12212817 | NPY1 | VII | 376101 | 377255 | 1 |
| YGL068W | 82.034876 | 1.003601 | 0.57641671 | 5.49064764 | 0.0642275 | 0.15764189 | MNP1 | VII | 375087 | 375671 | 1 |
| YGL071W | 74.7011048 | 0.13351633 | 0.63939827 | 0.82832113 | 0.66089483 | 0.78866056 | AFT1 | VII | 372012 | 374084 | 1 |
| YGL073W | 1657.32762 | -0.8910517 | 0.45099052 | 3.9700182 | 0.13737936 | 0.27509737 | HSF1 | VII | 368753 | 371254 | 1 |
| YGL075C | 7.38898844 | -0.6932774 | 1.03317416 | 2.31797102 | 0.31380437 | 0.48580944 | MPS2 | VII | 366925 | 368088 | -1 |

| | | | | | | | | | | | |
|---|---|---|---|---|---|---|---|---|---|---|---|
| YGL076C | 53.0776461 | -2.5624469 | 0.82341984 | 9.09407969 | 0.01059853 | 0.04121256 | RPL7A | VII | 364335 | 365996 | -1 |
| YGL077C | 1267.9299 | 0.48950136 | 0.54838349 | 1.99113474 | 0.36951373 | 0.54287859 | HNM1 | VII | 362225 | 363916 | -1 |
| YGL078C | 229.554652 | -2.6748413 | 0.49763492 | 25.9773504 | 2.29E-06 | 3.61E-05 | DBP3 | VII | 360288 | 361859 | -1 |
| YGL079W | 379.220671 | -0.320513 | 0.47449409 | 5.27642334 | 0.071489 | 0.1715269 | KXD1 | VII | 359444 | 360100 | 1 |
| YGL080W | 18.1243303 | -0.5959413 | 0.78095987 | 0.80562109 | 0.66843873 | 0.79450097 | MPC1 | VII | 358636 | 359028 | 1 |
| YGL081W | 43.5545693 | 0.69565414 | 0.58945409 | 2.76233337 | 0.25128521 | 0.42270232 | NA | VII | 357377 | 358339 | 1 |
| YGL082W | 17.5138914 | 0.79523896 | 0.77727159 | 1.60795944 | 0.44754431 | 0.61433892 | NA | VII | 355827 | 356972 | 1 |
| YGL083W | 576.818578 | 0.17256955 | 0.46075123 | 0.4148555 | 0.81267196 | 0.88731158 | SCY1 | VII | 353058 | 355472 | 1 |
| YGL084C | 49.2554351 | 1.40841058 | 0.71693531 | 4.47377235 | 0.10679051 | 0.22957814 | GUP1 | VII | 350616 | 352298 | -1 |
| YGL085W | 67.4886169 | -1.1693503 | 0.68349608 | 9.45513432 | 0.00884797 | 0.03585007 | LCL3 | VII | 349616 | 350440 | 1 |
| YGL086W | 32.7756323 | -0.7682577 | 0.62174093 | 2.73374587 | 0.25490281 | 0.42716573 | MAD1 | VII | 347119 | 349368 | 1 |
| YGL087C | 29.654439 | 0.08440069 | 0.70123626 | 2.03550281 | 0.36140669 | 0.53408966 | MMS2 | VII | 346406 | 346904 | -1 |
| YGL089C | 357.329818 | 0.41355892 | 0.46098237 | 0.82670201 | 0.66143008 | 0.78897407 | MF(ALPHA)2 | VII | 344791 | 345153 | -1 |
| YGL090W | 752.796528 | -4.1434217 | 0.62253983 | 50.9094892 | 8.81E-12 | 5.32E-10 | LIF1 | VII | 343319 | 344584 | 1 |
| YGL091C | 42.8939327 | -1.1477538 | 0.6088271 | 3.61095192 | 0.16439619 | 0.31321288 | NBP35 | VII | 342056 | 343042 | -1 |
| YGL092W | 48.8461922 | -0.4325289 | 0.76944609 | 6.6637846 | 0.03572544 | 0.10209787 | NUP145 | VII | 337906 | 341859 | 1 |
| YGL093W | 23.9984831 | -0.2723248 | 0.70711219 | 0.48067489 | 0.78636246 | 0.87021248 | SPC105 | VII | 334886 | 337639 | 1 |
| YGL094C | 19.6884986 | -0.1608679 | 0.72896936 | 0.32116502 | 0.85164755 | 0.91195795 | PAN2 | VII | 331118 | 334465 | -1 |
| YGL095C | 88.5586891 | -1.8140799 | 0.87070701 | 13.0755731 | 0.00144769 | 0.00890767 | VPS45 | VII | 328874 | 330607 | -1 |
| YGL096W | 86.1662258 | 0.6609538 | 0.54478254 | 2.38076522 | 0.30410489 | 0.47720463 | TOS8 | VII | 325331 | 326161 | 1 |
| YGL097W | 84.7872075 | -0.2662081 | 0.53164716 | 0.25745261 | 0.87921457 | 0.92911891 | SRM1 | VII | 321782 | 323230 | 1 |
| YGL098W | 13.5829956 | 0.79456491 | 1.30082525 | 0.45666729 | 0.79585868 | 0.8765497 | USE1 | VII | 317342 | 318079 | 1 |
| YGL099W | 19.1293074 | -0.6967235 | 0.70951031 | 2.68797851 | 0.26080318 | 0.43235173 | LSG1 | VII | 314631 | 316553 | 1 |
| YGL100W | 26.0745841 | -0.8332909 | 0.6683051 | 1.7331452 | 0.42038993 | 0.59107763 | SEH1 | VII | 313234 | 314283 | 1 |
| YGL101W | 67.0938327 | -1.4834421 | 0.5420726 | 9.8284474 | 0.00734141 | 0.03116334 | NA | VII | 312193 | 312840 | 1 |
| YGL103W | 290.015156 | -2.0991372 | 0.71736953 | 7.97174877 | 0.01857619 | 0.06260545 | RPL28 | VII | 310967 | 311927 | 1 |
| YGL104C | 112.894762 | -0.0337996 | 0.51233603 | 0.05445076 | 0.97314189 | 0.97940058 | VPS73 | VII | 308713 | 310173 | -1 |
| YGL105W | 208.334285 | -1.6124118 | 0.51343044 | 13.2547619 | 0.00132363 | 0.00825839 | ARC1 | VII | 307437 | 308567 | 1 |
| YGL106W | 20.8700901 | -0.912983 | 0.7273956 | 1.89002894 | 0.38867395 | 0.56060856 | MLC1 | VII | 306560 | 307009 | 1 |
| YGL107C | 24.3592471 | 0.31783289 | 0.72023434 | 3.14507885 | 0.20751754 | 0.36958676 | RMD9 | VII | 304333 | 306273 | -1 |
| YGL108C | 27.8800794 | -0.1764919 | 0.76042693 | 2.49713716 | 0.2869152 | 0.46119906 | NA | VII | 303649 | 304071 | -1 |
| YGL110C | 112.48558 | 0.59373116 | 0.71403732 | 0.97376909 | 0.61453798 | 0.75689745 | CUE3 | VII | 301537 | 303411 | -1 |
| YGL111W | 81.6624903 | -0.4358658 | 0.59176029 | 0.56129862 | 0.75529316 | 0.85213317 | NSA1 | VII | 299978 | 301369 | 1 |

| | | | | | | | | | | | |
|---|---|---|---|---|---|---|---|---|---|---|---|
| YGL112C | 50.7931724 | 0.856694 | 0.74541741 | 1.92571434 | 0.38180046 | 0.55529381 | TAF6 | VII | 298178 | 299728 | -1 |
| YGL113W | 106.305412 | 3.32259481 | 0.5907221 | 32.1529831 | 1.04E-07 | 2.51E-06 | SLD3 | VII | 295932 | 297938 | 1 |
| YGL114W | 34.2179209 | 0.18501754 | 0.63205744 | 0.256511 | 0.8796286 | 0.92911891 | NA | VII | 293460 | 295637 | 1 |
| YGL115W | 63.4187715 | 0.65138703 | 0.57194819 | 1.30598405 | 0.52048614 | 0.6765916 | SNF4 | VII | 292033 | 293001 | 1 |
| YGL116W | 20.7691671 | 0.04607964 | 0.73086994 | 0.08268418 | 0.95950084 | 0.97158269 | CDC20 | VII | 289809 | 291641 | 1 |
| YGL117W | 24.8455311 | -2.7167683 | 0.73329157 | 14.756896 | 0.00062457 | 0.00444804 | NA | VII | 288512 | 289309 | 1 |
| YGL118C | 122.706991 | 0.96427976 | 0.53905915 | 5.81817472 | 0.05452547 | 0.13962957 | NA | VII | 288014 | 288451 | -1 |
| YGL119W | 16.1089108 | -1.0267449 | 0.84199303 | 1.55609708 | 0.45930145 | 0.62382251 | COQ8 | VII | 284442 | 285947 | 1 |
| YGL120C | 25.4212832 | 0.28738121 | 0.80072612 | 1.06837762 | 0.58614457 | 0.73378964 | PRP43 | VII | 281634 | 283937 | -1 |
| YGL121C | 59.453868 | -2.1090547 | 0.57215702 | 14.1887399 | 0.00082976 | 0.00565881 | GPG1 | VII | 280777 | 281157 | -1 |
| YGL122C | 305.348737 | 3.29689973 | 0.60585426 | 26.5898346 | 1.68E-06 | 2.80E-05 | NAB2 | VII | 278946 | 280523 | -1 |
| YGL123W | 928.086072 | -2.9933799 | 0.5486431 | 26.1287237 | 2.12E-06 | 3.36E-05 | RPS2 | VII | 277617 | 278381 | 1 |
| YGL124C | 349.881209 | 1.54721584 | 0.47757879 | 11.6836261 | 0.00290357 | 0.01483821 | MON1 | VII | 274780 | 276714 | -1 |
| YGL125W | 915.407718 | -0.0986562 | 0.51895419 | 0.95398435 | 0.62064739 | 0.76119942 | MET13 | VII | 272520 | 274322 | 1 |
| YGL126W | 169.798592 | 1.02998094 | 0.52753061 | 4.37547026 | 0.11217051 | 0.23842411 | SCS3 | VII | 270997 | 272139 | 1 |
| YGL127C | 45.7442302 | 2.38754295 | 0.65268605 | 12.8209779 | 0.00164422 | 0.00973419 | SOH1 | VII | 270392 | 270775 | -1 |
| YGL128C | 51.9198862 | -0.512975 | 0.56360769 | 0.96740473 | 0.61649667 | 0.75817303 | CWC23 | VII | 269293 | 270144 | -1 |
| YGL129C | 30.2745529 | -0.2984882 | 0.64030432 | 0.30459998 | 0.85873063 | 0.91650698 | RSM23 | VII | 267723 | 269075 | -1 |
| YGL130W | 61.9859059 | -0.5353264 | 0.57599984 | 1.13517833 | 0.56689047 | 0.71775548 | CEG1 | VII | 266145 | 267524 | 1 |
| YGL131C | 57.2550088 | 0.95583902 | 0.56839603 | 3.94697428 | 0.1389714 | 0.27755929 | SNT2 | VII | 261648 | 265859 | -1 |
| YGL133W | 2247.16268 | 0.81322883 | 0.55222764 | 2.18875857 | 0.33474733 | 0.50817698 | ITC1 | VII | 257707 | 261501 | 1 |
| YGL134W | 93.712272 | -0.1194634 | 0.52981518 | 0.05702772 | 0.97188883 | 0.97864979 | PCL10 | VII | 255663 | 256964 | 1 |
| YGL135W | 27.5218963 | -2.8755135 | 0.95232268 | 8.64296652 | 0.01328017 | 0.04868773 | RPL1A | VII | 254641 | 255294 | 1 |
| YGL136C | 1250.81117 | -0.7492225 | 0.54759926 | 9.35390782 | 0.00930732 | 0.03735679 | MRM2 | VII | 252897 | 253859 | -1 |
| YGL137W | 275.033074 | -0.1010127 | 0.46630096 | 0.62784103 | 0.7305771 | 0.83678999 | SEC27 | VII | 249869 | 252738 | 1 |
| YGL139W | 16.1843891 | -1.0126768 | 0.78089261 | 1.78739463 | 0.40914023 | 0.58076047 | FLC3 | VII | 245716 | 248124 | 1 |
| YGL140C | 34.3666264 | 0.35582225 | 0.81181141 | 0.52772989 | 0.76807727 | 0.85918999 | NA | VII | 241353 | 245012 | -1 |
| YGL141W | 61.7083648 | 0.31715817 | 0.7075276 | 3.0627851 | 0.21623434 | 0.38031495 | HUL5 | VII | 238353 | 241085 | 1 |
| YGL142C | 32.3623598 | 0.59451041 | 0.63081344 | 0.91112345 | 0.63409168 | 0.77081479 | GPI10 | VII | 236269 | 238119 | -1 |
| YGL143C | 26.5822723 | 1.13030185 | 0.7009432 | 3.75622647 | 0.15287828 | 0.29674779 | MRF1 | VII | 234717 | 235958 | -1 |
| YGL144C | 101.266652 | 0.24601709 | 0.69872125 | 5.46410341 | 0.06508562 | 0.15927065 | ROG1 | VII | 232450 | 234507 | -1 |
| YGL145W | 11.9580561 | -0.1632565 | 0.86428606 | 0.20814208 | 0.90116128 | 0.93824416 | TIP20 | VII | 230243 | 232348 | 1 |
| YGL146C | 27.9101196 | -0.6933698 | 0.72020366 | 5.39942144 | 0.06722496 | 0.16286148 | RRT6 | VII | 228751 | 229686 | -1 |

| | | | | | | | | | | | |
|---|---|---|---|---|---|---|---|---|---|---|---|
| YGL147C | 152.941056 | -4.5314822 | 0.87699746 | 21.7143644 | 1.93E-05 | 0.00022952 | RPL9A | VII | 227754 | 228329 | -1 |
| YGL148W | 70.3199525 | -1.2686375 | 0.5830236 | 4.82926789 | 0.08940006 | 0.20267482 | ARO2 | VII | 226399 | 227529 | 1 |
| YGL149W | 509.369016 | -0.6855558 | 0.59011717 | 2.39490932 | 0.30196183 | 0.47448396 | NA | VII | 225572 | 225877 | 1 |
| YGL150C | 155.229915 | 0.1373269 | 0.6801308 | 6.22626277 | 0.04446151 | 0.11984737 | INO80 | VII | 221104 | 225573 | -1 |
| YGL151W | 68.178915 | -0.3436563 | 0.65099516 | 2.90153422 | 0.23439042 | 0.40199067 | NUT1 | VII | 217524 | 220922 | 1 |
| YGL153W | 172.605315 | 0.99897106 | 0.5875951 | 8.10726793 | 0.01735918 | 0.05959198 | PEX14 | VII | 216273 | 217298 | 1 |
| YGL154C | 452.424049 | -2.1134361 | 0.46794687 | 18.9659443 | 7.61E-05 | 0.0007409 | LYS5 | VII | 215278 | 216096 | -1 |
| YGL155W | 38.8748404 | -0.5135097 | 0.79695936 | 0.99674467 | 0.60751869 | 0.7514491 | CDC43 | VII | 214081 | 215211 | 1 |
| YGL156W | 512.098438 | -3.7814117 | 0.54258637 | 48.6913961 | 2.67E-11 | 1.42E-09 | AMS1 | VII | 210416 | 213667 | 1 |
| YGL157W | 65.314318 | 2.22912848 | 0.70261904 | 9.55595803 | 0.00841298 | 0.03453249 | ARI1 | VII | 209006 | 210049 | 1 |
| YGL158W | 38.6731739 | 1.99369481 | 0.69154612 | 8.0713957 | 0.01767334 | 0.06020484 | RCK1 | VII | 207033 | 208571 | 1 |
| YGL159W | 63.0111839 | -0.2076198 | 0.58395565 | 7.86625421 | 0.01958234 | 0.06501327 | NA | VII | 202721 | 203833 | 1 |
| YGL160W | 27.4260709 | -0.0070217 | 0.68904592 | 0.99096956 | 0.60927547 | 0.75306988 | AIM14 | VII | 200561 | 202273 | 1 |
| YGL161C | 62.3310456 | 0.06950311 | 0.60926452 | 0.14887382 | 0.92826604 | 0.95498585 | YIP5 | VII | 199210 | 200142 | -1 |
| YGL162W | 14.4182685 | -2.3632188 | 0.86597549 | 8.21551515 | 0.01644461 | 0.05722012 | SUT1 | VII | 198138 | 199037 | 1 |
| YGL163C | 27.6063779 | -0.3300016 | 0.74388682 | 2.35817081 | 0.3075599 | 0.48012182 | RAD54 | VII | 193707 | 196403 | -1 |
| YGL164C | 84.093683 | 0.36157668 | 0.57931316 | 3.81281166 | 0.14861357 | 0.29099512 | YRB30 | VII | 191978 | 193300 | -1 |
| YGL166W | 36.1747039 | 1.78168548 | 0.64746015 | 7.60943999 | 0.02226543 | 0.07177998 | CUP2 | VII | 191129 | 191806 | 1 |
| YGL167C | 48.8100926 | -0.4943429 | 0.66604359 | 1.10524503 | 0.57543873 | 0.72461728 | PMR1 | VII | 187616 | 190468 | -1 |
| YGL168W | 5495.69473 | -0.1634311 | 0.44735839 | 4.3167329 | 0.11551366 | 0.242911 | HUR1 | VII | 187464 | 187796 | 1 |
| YGL169W | 27.2474897 | 0.91195796 | 0.74658335 | 1.56723385 | 0.45675099 | 0.62210967 | SUA5 | VII | 186059 | 187339 | 1 |
| YGL170C | 286.923757 | -3.1966401 | 0.79440238 | 14.1949278 | 0.0008272 | 0.00564798 | SPO74 | VII | 184153 | 185394 | -1 |
| YGL171W | 20.8947615 | -0.8868367 | 0.75551168 | 1.8710628 | 0.3923773 | 0.56387803 | ROK1 | VII | 182390 | 184084 | 1 |
| YGL172W | 19.4032111 | 0.35167732 | 0.74352778 | 1.39328788 | 0.49825467 | 0.6579007 | NUP49 | VII | 180700 | 182118 | 1 |
| YGL173C | 67.1401684 | -0.702865 | 0.5947261 | 1.97875953 | 0.37180723 | 0.54514152 | XRN1 | VII | 175527 | 180113 | -1 |
| YGL174W | 87.4668753 | -1.5134772 | 0.76523205 | 6.86974545 | 0.03222951 | 0.09435142 | BUD13 | VII | 174545 | 175345 | 1 |
| YGL175C | 240.078984 | 2.32859895 | 0.53938851 | 24.1841826 | 5.60E-06 | 7.86E-05 | SAE2 | VII | 173285 | 174322 | -1 |
| YGL176C | 195.05709 | 3.01268661 | 0.5464919 | 39.8516156 | 2.22E-09 | 7.70E-08 | NA | VII | 171415 | 173079 | -1 |
| YGL178W | 122.179277 | 0.37456556 | 0.51319662 | 0.60525069 | 0.73887587 | 0.84040603 | MPT5 | VII | 167352 | 170571 | 1 |
| YGL179C | 26.1600606 | -0.9325284 | 0.66234956 | 5.44016662 | 0.06586927 | 0.16064998 | TOS3 | VII | 163409 | 165091 | -1 |
| YGL180W | 108.468344 | -2.0942633 | 0.55951761 | 20.0585478 | 4.41E-05 | 0.00046183 | ATG1 | VII | 160065 | 162758 | 1 |
| YGL181W | 42.6780784 | 1.32025517 | 0.6693488 | 6.44875142 | 0.03978061 | 0.11116077 | GTS1 | VII | 157906 | 159096 | 1 |
| YGL184C | 107.571463 | 0.91257132 | 0.53231215 | 4.65916543 | 0.09733636 | 0.21492841 | STR3 | VII | 154615 | 156012 | -1 |

| | | | | | | | | | | | |
|---|---|---|---|---|---|---|---|---|---|---|---|
| YGL185C | 23.512965 | 0.20385309 | 0.73650519 | 2.13886403 | 0.3432034 | 0.51667906 | NA | VII | 153167 | 154306 | -1 |
| YGL186C | 37.1146483 | 0.59066605 | 0.63668366 | 1.39176741 | 0.49863361 | 0.65825093 | TPN1 | VII | 151037 | 152776 | -1 |
| YGL187C | 58.3336215 | 1.23494818 | 0.56734856 | 11.8597823 | 0.00265877 | 0.01389376 | COX4 | VII | 149704 | 150171 | -1 |
| YGL189C | 322.787215 | -2.9395154 | 0.6241294 | 20.1128683 | 4.29E-05 | 0.00045254 | RPS26A | VII | 148229 | 148588 | -1 |
| YGL190C | 52.8698749 | -0.4031074 | 0.63619611 | 3.38558598 | 0.18400488 | 0.3396201 | CDC55 | VII | 145809 | 147389 | -1 |
| YGL191W | 25.6908855 | 1.03550509 | 0.76027472 | 2.14296088 | 0.34250109 | 0.51615859 | COX13 | VII | 144808 | 145197 | 1 |
| YGL192W | 7094.27745 | 0.37471391 | 0.52338524 | 0.5639851 | 0.7542793 | 0.85152235 | IME4 | VII | 142246 | 144048 | 1 |
| YGL194C | 339.018628 | 2.02311507 | 0.4831684 | 17.1353681 | 0.00019015 | 0.00163351 | HOS2 | VII | 140368 | 141726 | -1 |
| YGL194C-A | 82.0249815 | 1.67403056 | 0.62952444 | 7.48454727 | 0.02370016 | 0.07502674 | NA | VII | 139719 | 139961 | -1 |
| YGL195W | 77.4018836 | -0.4858088 | 0.61813528 | 2.25111576 | 0.3244714 | 0.49763242 | GCN1 | VII | 131525 | 139543 | 1 |
| YGL196W | 38.8342515 | -1.9497215 | 0.69864521 | 8.2567839 | 0.01610876 | 0.05656139 | DSD1 | VII | 129883 | 131169 | 1 |
| YGL197W | 43.2867104 | 0.71126509 | 0.63238938 | 3.05997539 | 0.21653833 | 0.38073396 | MDS3 | VII | 124698 | 129161 | 1 |
| YGL198W | 53.1472986 | -0.6674677 | 0.56575388 | 1.40890851 | 0.4943783 | 0.65483292 | YIP4 | VII | 123591 | 124298 | 1 |
| YGL200C | 206.074287 | -0.472145 | 0.75196934 | 0.68716697 | 0.70922427 | 0.82408359 | EMP24 | VII | 122694 | 123305 | -1 |
| YGL201C | 62.1026248 | -1.4787594 | 0.59836765 | 7.59498695 | 0.02242692 | 0.07215694 | MCM6 | VII | 117854 | 120907 | -1 |
| YGL202W | 92.047105 | 1.19886612 | 0.5923861 | 4.34554675 | 0.1138614 | 0.24051715 | ARO8 | VII | 116059 | 117561 | 1 |
| YGL203C | 157.096059 | 0.62609058 | 0.66892046 | 4.4757802 | 0.10668336 | 0.22954167 | KEX1 | VII | 112475 | 114664 | -1 |
| YGL204C | 22.2169155 | 2.86061616 | 0.85825213 | 13.0412406 | 0.00147276 | 0.00899499 | NA | VII | 111700 | 112005 | -1 |
| YGL205W | 668.500253 | -1.0121634 | 0.48980609 | 5.96624431 | 0.0506345 | 0.13188203 | POX1 | VII | 108158 | 110404 | 1 |
| YGL206C | 32.8787024 | -0.4981537 | 0.72447502 | 1.9207198 | 0.38275511 | 0.55598396 | CHC1 | VII | 102543 | 107504 | -1 |
| YGL207W | 306.944337 | 3.3202976 | 0.52465906 | 48.4563802 | 3.00E-11 | 1.55E-09 | SPT16 | VII | 98969 | 102076 | 1 |
| YGL208W | 90.7007233 | -0.1869651 | 0.58478767 | 5.69287621 | 0.05805072 | 0.14632725 | SIP2 | VII | 97338 | 98585 | 1 |
| YGL209W | 308.547492 | 1.69066524 | 0.51045387 | 10.7864489 | 0.00454729 | 0.02111371 | MIG2 | VII | 95858 | 97006 | 1 |
| YGL210W | 17.5816631 | -0.0341979 | 0.85079446 | 4.12269208 | 0.12728253 | 0.26090922 | YPT32 | VII | 93792 | 94460 | 1 |
| YGL211W | 250.322822 | 4.68562051 | 0.56702176 | 63.5058938 | 1.62E-14 | 1.51E-12 | NCS6 | VII | 92512 | 93591 | 1 |
| YGL212W | 8.57469329 | 0.65344693 | 0.97175934 | 0.49424235 | 0.78104604 | 0.86705973 | VAM7 | VII | 91432 | 92382 | 1 |
| YGL213C | 44.8018603 | -1.3586916 | 0.61789429 | 5.7643959 | 0.05601152 | 0.14261508 | SKI8 | VII | 90054 | 91247 | -1 |
| YGL215W | 42.532926 | 0.04986224 | 0.66926347 | 0.6238721 | 0.73202834 | 0.83714084 | CLG1 | VII | 87981 | 89339 | 1 |
| YGL216W | 37.5954053 | 0.68359159 | 0.67867233 | 1.68895245 | 0.42978241 | 0.59904674 | KIP3 | VII | 84885 | 87302 | 1 |
| YGL219C | 1148.99621 | 0.59792105 | 0.96877753 | 10.1183028 | 0.00635095 | 0.02766891 | MDM34 | VII | 82878 | 84257 | -1 |
| YGL220W | 17.1760715 | -0.6238425 | 0.87867513 | 0.64160808 | 0.72556542 | 0.83456178 | BOL2 | VII | 82374 | 82736 | 1 |
| YGL221C | 54.772759 | -0.2761103 | 0.61578393 | 0.25621471 | 0.87975893 | 0.92911891 | NIF3 | VII | 81426 | 82292 | -1 |
| YGL222C | 34.4220691 | -0.6001171 | 0.68004473 | 0.97378765 | 0.61453228 | 0.75689745 | EDC1 | VII | 80659 | 81186 | -1 |

| | | | | | | | | | | | |
|---|---|---|---|---|---|---|---|---|---|---|---|
| YGL223C | 43.8260445 | 0.4722487 | 0.73779805 | 3.39187344 | 0.18342733 | 0.33887818 | COG1 | VII | 79112 | 80365 | -1 |
| YGL224C | 1907.65056 | -0.196149 | 0.88057807 | 6.17543788 | 0.04560587 | 0.12191042 | SDT1 | VII | 78015 | 78857 | -1 |
| YGL225W | 28.9816834 | 0.70620841 | 0.69782637 | 1.05926894 | 0.58882016 | 0.73666135 | VRG4 | VII | 76894 | 77907 | 1 |
| YGL226C-A | 28.2472757 | 1.04037726 | 0.98943308 | 3.42675536 | 0.18025592 | 0.3349428 | OST5 | VII | 72749 | 73158 | -1 |
| YGL226W | 11.8306722 | -0.4605837 | 0.89140342 | 1.54840249 | 0.46107192 | 0.62486036 | MTC3 | VII | 73340 | 73711 | 1 |
| YGL227W | 182.366438 | -1.0904804 | 0.52829723 | 4.56130041 | 0.10221772 | 0.22276062 | VID30 | VII | 69671 | 72547 | 1 |
| YGL228W | 54.2472198 | 0.08471435 | 0.77173233 | 2.28322309 | 0.31930403 | 0.49195592 | SHE10 | VII | 67598 | 69331 | 1 |
| YGL229C | 42.9554226 | -0.1358691 | 0.66641356 | 0.57103677 | 0.75162452 | 0.84964974 | SAP4 | VII | 64503 | 66959 | -1 |
| YGL230C | 97.5783894 | 1.25160875 | 0.56505391 | 4.90512247 | 0.08607285 | 0.19729288 | NA | VII | 63772 | 64215 | -1 |
| YGL231C | 125.170884 | -0.8390862 | 0.74692421 | 11.8210464 | 0.00271077 | 0.01406419 | EMC4 | VII | 63048 | 63620 | -1 |
| YGL232W | 277.741902 | 1.15057855 | 0.96760142 | 12.7344281 | 0.00171694 | 0.00999815 | TAN1 | VII | 62075 | 63002 | 1 |
| YGL233W | 23.7761528 | 0.89315963 | 0.71208869 | 3.06668701 | 0.21581289 | 0.37968904 | SEC15 | VII | 59122 | 61854 | 1 |
| YGL234W | 99.6585088 | 1.6785856 | 0.59145192 | 12.2754569 | 0.00215982 | 0.01184098 | ADE5,7 | VII | 56482 | 58890 | 1 |
| YGL235W | 261.743606 | -3.19459 | 0.56372185 | 30.3674698 | 2.55E-07 | 5.38E-06 | NA | VII | 55279 | 55815 | 1 |
| YGL236C | 42.1998781 | 1.07173674 | 0.63302524 | 6.41314335 | 0.04049521 | 0.11240041 | MTO1 | VII | 53787 | 55796 | -1 |
| YGL237C | 14.7157776 | 0.26341287 | 0.81787026 | 1.32814684 | 0.51475026 | 0.67171603 | HAP2 | VII | 52731 | 53528 | -1 |
| YGL238W | 27.4742485 | 0.15066429 | 0.73934898 | 1.75717594 | 0.41536901 | 0.58728945 | CSE1 | VII | 49552 | 52434 | 1 |
| YGL240W | 128.44674 | -0.4834865 | 0.55648052 | 1.00877477 | 0.60387541 | 0.74934389 | DOC1 | VII | 48613 | 49365 | 1 |
| YGL241W | 10.9410656 | -0.1783322 | 0.91124209 | 0.1070899 | 0.94786333 | 0.96435226 | KAP114 | VII | 45445 | 48459 | 1 |
| YGL242C | 51.305493 | 0.42651949 | 0.76896273 | 2.80556437 | 0.24591184 | 0.41681193 | NA | VII | 44652 | 45197 | -1 |
| YGL243W | 437.169956 | -0.0546341 | 0.83979801 | 15.4131084 | 0.00044987 | 0.00337792 | TAD1 | VII | 43307 | 44509 | 1 |
| YGL244W | 95.3922924 | -0.834378 | 0.52960547 | 5.51731691 | 0.06337673 | 0.15623132 | RTF1 | VII | 41498 | 43174 | 1 |
| YGL245W | 93.1922743 | -1.6617112 | 0.79275348 | 4.4888891 | 0.10598639 | 0.22838155 | GUS1 | VII | 39023 | 41149 | 1 |
| YGL246C | 42.3951834 | -1.0627685 | 0.80034916 | 4.36117171 | 0.11297532 | 0.23947912 | RAI1 | VII | 37617 | 38780 | -1 |
| YGL247W | 382.397243 | -1.3163173 | 0.77245542 | 6.19027071 | 0.04526888 | 0.12151453 | BRR6 | VII | 36933 | 37526 | 1 |
| YGL248W | 55.3759054 | 0.29308403 | 0.6340998 | 0.43444614 | 0.80475043 | 0.88279205 | PDE1 | VII | 35653 | 36762 | 1 |
| YGL249W | 1329.84906 | 1.03705886 | 0.46858398 | 5.42301018 | 0.06643674 | 0.16169345 | ZIP2 | VII | 33098 | 35212 | 1 |
| YGL250W | 339.113139 | 0.52403583 | 0.5281484 | 1.38475171 | 0.50038581 | 0.65981185 | RMR1 | VII | 31910 | 32635 | 1 |
| YGL251C | 334.546996 | 1.75276273 | 0.48800901 | 15.8222579 | 0.00036664 | 0.00286098 | HFM1 | VII | 27921 | 31636 | -1 |
| YGL252C | 45.7510853 | -0.0816554 | 0.91495445 | 0.83939061 | 0.65724705 | 0.78549013 | RTG2 | VII | 25718 | 27484 | -1 |
| YGL253W | 101.465052 | -0.9364143 | 0.63055445 | 4.169413 | 0.12434361 | 0.25639227 | HXK2 | VII | 23935 | 25395 | 1 |
| YGL254W | 21.7250865 | -1.9556535 | 0.72120773 | 9.50702389 | 0.00862136 | 0.03516063 | FZF1 | VII | 22304 | 23203 | 1 |
| YGL255W | 50.6397061 | -1.3836368 | 0.61791778 | 5.11422386 | 0.07752832 | 0.18217898 | ZRT1 | VII | 20978 | 22108 | 1 |

| | | | | | | | | | | | |
|---|---|---|---|---|---|---|---|---|---|---|---|
| YGL256W | 7.57311263 | -1.131645 | 1.21688752 | 0.94763173 | 0.62262189 | 0.76281637 | ADH4 | VII | 15159 | 16307 | 1 |
| YGL257C | 457.950788 | 1.32956765 | 0.61015076 | 4.82945096 | 0.08939188 | 0.20267482 | MNT2 | VII | 12481 | 14157 | -1 |
| YGL258W | 179.8594 | -0.3141156 | 0.49625645 | 0.47922189 | 0.78693396 | 0.87036249 | VEL1 | VII | 11110 | 11730 | 1 |
| YGL260W | 4.82872011 | 1.55100008 | 1.50141131 | 1.64734361 | 0.43881744 | 0.60769026 | NA | VII | 6860 | 7090 | 1 |
| YGR001C | 247.124933 | 1.19764564 | 0.50222711 | 10.8474836 | 0.00441061 | 0.02064466 | EFM5 | VII | 497133 | 498034 | -1 |
| YGR002C | 100.504987 | 2.28973945 | 0.55970777 | 18.9589131 | 7.64E-05 | 0.00074226 | SWC4 | VII | 498476 | 499906 | -1 |
| YGR003W | 33.0487759 | -0.5995104 | 0.64732396 | 1.44201634 | 0.48626177 | 0.64797599 | CUL3 | VII | 500132 | 502366 | 1 |
| YGR004W | 33.0980316 | 0.93044311 | 0.64478827 | 5.08874167 | 0.07852244 | 0.18399228 | PEX31 | VII | 502938 | 504326 | 1 |
| YGR005C | 38.9371275 | -0.3064121 | 0.65084322 | 0.32955518 | 0.8480823 | 0.90980423 | TFG2 | VII | 504657 | 505859 | -1 |
| YGR006W | 15.5554726 | 1.18099089 | 0.79404902 | 2.24284756 | 0.32581557 | 0.49867094 | PRP18 | VII | 506070 | 506825 | 1 |
| YGR007W | 54.1340037 | 2.29257011 | 0.79847291 | 11.4304482 | 0.00329541 | 0.01642034 | ECT1 | VII | 506969 | 507940 | 1 |
| YGR008C | 187.811617 | 2.06899481 | 0.49658158 | 16.5285968 | 0.00025755 | 0.0021303 | STF2 | VII | 508110 | 508364 | -1 |
| YGR009C | 58.038851 | 0.77361536 | 0.57141333 | 2.19915814 | 0.33301123 | 0.50700368 | SEC9 | VII | 509047 | 511002 | -1 |
| YGR010W | 95.5238636 | -0.2349337 | 0.54210569 | 1.18104925 | 0.55403655 | 0.70568075 | NMA2 | VII | 511545 | 512732 | 1 |
| YGR012W | 75.6374511 | -0.5863696 | 0.62559536 | 5.19556447 | 0.07443848 | 0.17631337 | MCY1 | VII | 513159 | 514340 | 1 |
| YGR013W | 73.2854061 | 1.01388781 | 0.59366392 | 3.01368847 | 0.22160822 | 0.38636302 | SNU71 | VII | 514554 | 516416 | 1 |
| YGR014W | 43.4186146 | -0.594405 | 0.65085426 | 3.94199242 | 0.139318 | 0.27786814 | MSB2 | VII | 516943 | 520863 | 1 |
| YGR015C | 15.2853535 | 0.82612369 | 0.80369352 | 1.42537534 | 0.49032459 | 0.65159041 | NA | VII | 521090 | 522076 | -1 |
| YGR016W | 26.8682014 | -2.035969 | 0.78694194 | 7.94466286 | 0.01882948 | 0.06320157 | NA | VII | 522261 | 522833 | 1 |
| YGR017W | 71.4623018 | -0.41648 | 0.56339296 | 1.7548088 | 0.41586092 | 0.58742004 | NA | VII | 523787 | 524680 | 1 |
| YGR018C | 2640.43518 | -1.2718444 | 0.46431447 | 7.5713538 | 0.0226935 | 0.07283556 | NA | VII | 524365 | 524694 | -1 |
| YGR019W | 66.4962043 | -0.8798614 | 0.54533637 | 2.58758219 | 0.27422918 | 0.44776846 | UGA1 | VII | 525229 | 526644 | 1 |
| YGR020C | 13.1517034 | -1.5500973 | 1.03767773 | 2.36957419 | 0.30581128 | 0.47867401 | VMA7 | VII | 526973 | 527329 | -1 |
| YGR021W | 29.8710104 | 1.11462172 | 0.67794012 | 4.06840579 | 0.13078469 | 0.26588601 | NA | VII | 527632 | 528504 | 1 |
| YGR023W | 33.6201665 | -0.0722458 | 0.75344863 | 1.03448333 | 0.5961627 | 0.74343787 | MTL1 | VII | 529264 | 530919 | 1 |
| YGR024C | 12.2876693 | -1.0350834 | 0.90890372 | 1.86944677 | 0.39269447 | 0.56412138 | THG1 | VII | 531883 | 532596 | -1 |
| YGR025W | 14.1128462 | 0.82798762 | 0.90415038 | 0.86173777 | 0.64994412 | 0.78139046 | NA | VII | 532639 | 532941 | 1 |
| YGR026W | 17.9142655 | 0.32574163 | 0.7941588 | 0.2931176 | 0.86367495 | 0.91991868 | NA | VII | 532985 | 533821 | 1 |
| YGR027C | 76.7771469 | -0.7537293 | 0.66429711 | 5.31459134 | 0.07013764 | 0.16864491 | RPS25A | VII | 534132 | 534458 | -1 |
| YGR028W | 25.8391292 | 1.80873932 | 0.73008995 | 6.28391878 | 0.04319807 | 0.11764668 | MSP1 | VII | 542203 | 543291 | 1 |
| YGR029W | 264.697207 | 0.88194029 | 0.52362385 | 4.76006172 | 0.09254772 | 0.20705228 | ERV1 | VII | 543553 | 544205 | 1 |
| YGR030C | 183.49793 | -1.6750319 | 0.51635043 | 11.0312802 | 0.00402335 | 0.01933212 | POP6 | VII | 545681 | 546157 | -1 |
| YGR031C-A | 861.159475 | -0.9951592 | 0.56551748 | 6.95698459 | 0.03085389 | 0.09137803 | NAG1 | VII | 546809 | 547300 | -1 |

| | | | | | | | | | | | |
|---|---|---|---|---|---|---|---|---|---|---|---|
| YGR031W | 66.1051386 | 1.73563387 | 0.60432943 | 8.71853978 | 0.01278772 | 0.04740135 | IMO32 | VII | 546441 | 547469 | 1 |
| YGR032W | 232.141312 | 2.47821323 | 0.67598259 | 15.0423525 | 0.0005415 | 0.00391418 | GSC2 | VII | 548264 | 553951 | 1 |
| YGR033C | 10.9864489 | 0.48978472 | 0.96177193 | 0.99995532 | 0.60654421 | 0.75088539 | TIM21 | VII | 554248 | 554967 | -1 |
| YGR034W | 64.4848146 | -3.1202023 | 0.79098974 | 14.7142652 | 0.00063803 | 0.00453272 | RPL26B | VII | 555812 | 556672 | 1 |
| YGR035C | 20.2639258 | 0.71327805 | 0.75353174 | 0.88826237 | 0.64138128 | 0.77591986 | NA | VII | 557072 | 557422 | -1 |
| YGR035W-A | 991.666309 | -1.114873 | 0.5483569 | 6.10232792 | 0.04730383 | 0.12530377 | NA | VII | 557565 | 557786 | 1 |
| YGR036C | 17.2653411 | 1.25735942 | 0.83146583 | 3.65843135 | 0.16053943 | 0.30799315 | CAX4 | VII | 558149 | 558868 | -1 |
| YGR037C | 88.2969055 | 1.37253805 | 0.67669085 | 4.26627463 | 0.11846505 | 0.24726483 | ACB1 | VII | 559731 | 559994 | -1 |
| YGR038W | 517.495996 | 3.04451806 | 0.52720491 | 30.9743803 | 1.88E-07 | 4.16E-06 | ORM1 | VII | 560682 | 561350 | 1 |
| YGR039W | 48.4311911 | -0.5189999 | 0.57844808 | 0.8965754 | 0.6387209 | 0.77465312 | NA | VII | 574888 | 575199 | 1 |
| YGR040W | 46.3746771 | -1.3044537 | 0.63062059 | 4.25982923 | 0.11884744 | 0.2479657 | KSS1 | VII | 575398 | 576504 | 1 |
| YGR041W | 1012.97484 | 1.77787033 | 0.48797008 | 13.386986 | 0.00123895 | 0.00784848 | BUD9 | VII | 577487 | 579130 | 1 |
| YGR042W | 139.532124 | 4.05237466 | 0.84819535 | 20.2386896 | 4.03E-05 | 0.00043148 | MTE1 | VII | 579476 | 580291 | 1 |
| YGR043C | 51.017049 | 1.86743151 | 0.85855791 | 10.4307477 | 0.0054324 | 0.02450414 | NQM1 | VII | 580435 | 581436 | -1 |
| YGR044C | 72.16624 | 2.37655097 | 0.62505848 | 13.8250147 | 0.00099526 | 0.0065558 | RME1 | VII | 582990 | 583892 | -1 |
| YGR045C | 53.4112199 | 2.30275158 | 0.80732944 | 7.4718398 | 0.02385122 | 0.07538131 | NA | VII | 583934 | 584296 | -1 |
| YGR046W | 11.1531913 | 0.7245081 | 1.10006097 | 0.52489261 | 0.76916766 | 0.85953436 | TAM41 | VII | 584895 | 586052 | 1 |
| YGR047C | 98.072144 | 1.25034351 | 0.99957382 | 10.9278082 | 0.00423698 | 0.02007035 | TFC4 | VII | 586392 | 589469 | -1 |
| YGR048W | 37.1231895 | -0.1087423 | 0.66350426 | 0.31877 | 0.85266802 | 0.91239103 | UFD1 | VII | 589826 | 590911 | 1 |
| YGR049W | 1268.70925 | 1.70879377 | 0.48467542 | 19.0291575 | 7.38E-05 | 0.00072393 | SCM4 | VII | 591314 | 591877 | 1 |
| YGR050C | 130.086023 | -0.109216 | 0.57699713 | 0.76184801 | 0.68322981 | 0.80595449 | NA | VII | 592086 | 592442 | -1 |
| YGR052W | 39.0570244 | 0.26450433 | 0.61854449 | 5.14061931 | 0.07651185 | 0.18012252 | FMP48 | VII | 593596 | 594705 | 1 |
| YGR054W | 83.4827999 | -0.9767316 | 0.57310139 | 3.35684211 | 0.18666848 | 0.34246214 | NA | VII | 596693 | 598621 | 1 |
| YGR055W | 31.1267471 | -1.7392554 | 0.70745854 | 7.49650338 | 0.0235589 | 0.07474302 | MUP1 | VII | 599417 | 601141 | 1 |
| YGR056W | 164.24194 | -0.4284212 | 0.66916498 | 3.94980401 | 0.13877491 | 0.2772625 | RSC1 | VII | 601661 | 604447 | 1 |
| YGR057C | 165.194787 | 2.72923998 | 0.50812823 | 27.2355677 | 1.22E-06 | 2.09E-05 | LST7 | VII | 604773 | 605501 | -1 |
| YGR058W | 30.7612797 | -1.9533309 | 0.6782612 | 8.36925556 | 0.01522787 | 0.05412485 | PEF1 | VII | 606136 | 607143 | 1 |
| YGR059W | 15.9078194 | -1.4242426 | 0.96351568 | 2.23634571 | 0.3268765 | 0.49976629 | SPR3 | VII | 607563 | 609101 | 1 |
| YGR060W | 146.2102 | 0.44868283 | 0.49635587 | 1.23760358 | 0.5385894 | 0.69298502 | ERG25 | VII | 610564 | 611493 | 1 |
| YGR061C | 79.8559879 | -0.4825364 | 0.6142329 | 1.61809411 | 0.44528219 | 0.61297083 | ADE6 | VII | 611889 | 615965 | -1 |
| YGR062C | 29.5286947 | -1.9816963 | 0.76487229 | 9.91585575 | 0.00702747 | 0.03007323 | COX18 | VII | 616328 | 617278 | -1 |
| YGR063C | 95.4147023 | -0.3135532 | 0.53515235 | 1.64222066 | 0.4399429 | 0.60823052 | SPT4 | VII | 617516 | 617824 | -1 |
| YGR065C | 26.2186576 | 1.8751352 | 0.75251965 | 10.4321319 | 0.00542864 | 0.02450414 | VHT1 | VII | 618077 | 619858 | -1 |

| | | | | | | | | | | | |
|---|---|---|---|---|---|---|---|---|---|---|---|
| YGR066C | 197.19374 | -0.8825039 | 0.52384461 | 5.51890485 | 0.06332643 | 0.15622499 | NA | VII | 620921 | 621799 | -1 |
| YGR067C | 38.2686885 | 1.07268386 | 0.72360735 | 2.18759362 | 0.33494236 | 0.50833979 | NA | VII | 622372 | 624786 | -1 |
| YGR068C | 60.5496994 | 2.71179315 | 0.64654614 | 18.6474192 | 8.93E-05 | 0.00085024 | ART5 | VII | 625324 | 627084 | -1 |
| YGR070W | 34.5838589 | -0.5643546 | 0.63032586 | 2.93448194 | 0.23056073 | 0.39754218 | ROM1 | VII | 627806 | 631273 | 1 |
| YGR071C | 12.4031319 | -0.4990457 | 0.84693923 | 0.48581662 | 0.78434343 | 0.86949041 | ENV11 | VII | 631481 | 634063 | -1 |
| YGR072W | 66.8997026 | -0.8727996 | 0.62685191 | 4.17155478 | 0.12421052 | 0.25620909 | UPF3 | VII | 634304 | 635467 | 1 |
| YGR074W | 398.434296 | -0.7752377 | 0.4684932 | 2.72274637 | 0.25630858 | 0.42842105 | SMD1 | VII | 635712 | 636152 | 1 |
| YGR075C | 1050.111 | -2.0030022 | 0.45673526 | 18.1109465 | 0.00011675 | 0.00106622 | PRP38 | VII | 636147 | 636875 | -1 |
| YGR076C | 7.86148688 | -0.7021519 | 0.95403454 | 3.10574833 | 0.21163881 | 0.37519557 | MRPL25 | VII | 637104 | 637577 | -1 |
| YGR077C | 61.8800081 | 1.14049299 | 0.62026386 | 4.25357892 | 0.11921944 | 0.24840182 | PEX8 | VII | 637744 | 639513 | -1 |
| YGR078C | 64.0004497 | -2.7295675 | 0.59301516 | 26.2008014 | 2.04E-06 | 3.30E-05 | PAC10 | VII | 639772 | 640371 | -1 |
| YGR079W | 39.6895548 | 0.38663858 | 0.62259073 | 1.20667753 | 0.54698234 | 0.69943192 | NA | VII | 640720 | 641832 | 1 |
| YGR080W | 89.0060201 | -1.7802221 | 0.56412178 | 9.52838461 | 0.00852977 | 0.03487811 | TWF1 | VII | 642010 | 643008 | 1 |
| YGR081C | 22.4775208 | -1.4535168 | 1.04920642 | 7.86622965 | 0.01958258 | 0.06501327 | SLX9 | VII | 643148 | 643780 | -1 |
| YGR082W | 13.4625334 | -1.8777109 | 1.00014977 | 3.93599676 | 0.13973628 | 0.27822319 | TOM20 | VII | 644044 | 644595 | 1 |
| YGR083C | 115.275491 | -0.8242083 | 0.55460869 | 3.91879482 | 0.14094333 | 0.2798566 | GCD2 | VII | 644860 | 646815 | -1 |
| YGR084C | 10.4695064 | -3.3298784 | 1.0054892 | 10.904648 | 0.00428633 | 0.02021351 | MRP13 | VII | 647123 | 648142 | -1 |
| YGR085C | 34.3943069 | -2.5944047 | 0.77566466 | 12.0656719 | 0.00239868 | 0.01281215 | RPL11B | VII | 648383 | 648907 | -1 |
| YGR086C | 257.48631 | -0.9343486 | 0.48603565 | 7.14561854 | 0.02807687 | 0.08453721 | PIL1 | VII | 649598 | 650617 | -1 |
| YGR087C | 28.1356343 | 1.56447343 | 0.82479349 | 6.03479692 | 0.04892834 | 0.1284785 | PDC6 | VII | 651290 | 652981 | -1 |
| YGR088W | 64.5413612 | -2.2433562 | 0.64703301 | 12.6641516 | 0.00177834 | 0.01025556 | CTT1 | VII | 654634 | 656322 | 1 |
| YGR089W | 40.3944955 | 0.83057863 | 0.62163148 | 2.42348479 | 0.29767815 | 0.47143231 | NNF2 | VII | 656960 | 659770 | 1 |
| YGR090W | 33.4237345 | -2.3942895 | 0.82749596 | 8.35546856 | 0.01533321 | 0.05444929 | UTP22 | VII | 662358 | 666071 | 1 |
| YGR091W | 23.2245783 | 1.37322905 | 0.70450965 | 6.09448158 | 0.04748978 | 0.12561252 | PRP31 | VII | 666341 | 667825 | 1 |
| YGR092W | 25.5711744 | 0.76812068 | 0.7301829 | 1.30468767 | 0.52082362 | 0.6765916 | DBF2 | VII | 668189 | 669907 | 1 |
| YGR093W | 36.067363 | 0.3056824 | 0.61973477 | 4.11169073 | 0.1279846 | 0.26147876 | DRN1 | VII | 670388 | 671911 | 1 |
| YGR094W | 135.375242 | -0.1073465 | 0.53511823 | 0.241066 | 0.88644783 | 0.93215081 | VAS1 | VII | 672186 | 675500 | 1 |
| YGR095C | 9.97179505 | -1.4192229 | 0.90532896 | 2.48116905 | 0.28921512 | 0.46309611 | RRP46 | VII | 675671 | 676342 | -1 |
| YGR096W | 149.321279 | -0.5531787 | 0.52659511 | 2.44785581 | 0.2940728 | 0.46854198 | TPC1 | VII | 676621 | 677565 | 1 |
| YGR097W | 66.6655705 | -0.1268012 | 0.58929307 | 0.53349272 | 0.7658673 | 0.85754625 | ASK10 | VII | 678695 | 682135 | 1 |
| YGR098C | 23.4857546 | -0.5670688 | 0.74807882 | 0.61849816 | 0.73399792 | 0.83775415 | ESP1 | VII | 682566 | 687458 | -1 |
| YGR099W | 27.104797 | -0.3926804 | 0.64683795 | 0.44344505 | 0.80113763 | 0.88018726 | TEL2 | VII | 687899 | 689965 | 1 |
| YGR100W | 49.2106248 | -0.1309575 | 0.69971678 | 0.14690461 | 0.92918046 | 0.95541731 | MDR1 | VII | 690245 | 693097 | 1 |

| | | | | | | | | | | | |
|---|---|---|---|---|---|---|---|---|---|---|---|
| YGR101W | 63.6824691 | -0.8482426 | 0.73912699 | 2.41653887 | 0.29871378 | 0.47191811 | PCP1 | VII | 693363 | 694403 | 1 |
| YGR102C | 32.7961106 | 0.15998009 | 0.66010094 | 0.1266866 | 0.93862119 | 0.95995205 | GTF1 | VII | 694584 | 695135 | -1 |
| YGR103W | 138.007939 | -3.6155288 | 0.59000521 | 32.5922975 | 8.37E-08 | 2.09E-06 | NOP7 | VII | 695417 | 697234 | 1 |
| YGR104C | 88.2012622 | 0.93700436 | 0.60112319 | 4.85961789 | 0.08805365 | 0.20056281 | SRB5 | VII | 697445 | 698368 | -1 |
| YGR105W | 13.3296522 | -2.4283692 | 0.9845941 | 6.54544557 | 0.03790308 | 0.10674069 | VMA21 | VII | 698599 | 698832 | 1 |
| YGR106C | 346.628425 | -1.063501 | 1.60701977 | 4.17841322 | 0.12378531 | 0.25551406 | VOA1 | VII | 698989 | 699786 | -1 |
| YGR108W | 52.2261373 | 0.37042975 | 0.59837733 | 1.72404666 | 0.42230675 | 0.59293772 | CLB1 | VII | 703636 | 705051 | 1 |
| YGR109C | 184.179315 | 0.56931148 | 0.47768021 | 3.12411742 | 0.20970391 | 0.37244958 | CLB6 | VII | 705359 | 706501 | -1 |
| YGR109W-B | 291.950217 | -0.1922922 | 0.51949704 | 0.23841851 | 0.88762204 | 0.9322205 | NA | VII | 707610 | 712254 | 1 |
| YGR110W | 74.1632324 | -3.2588598 | 0.61258206 | 33.5928503 | 5.07E-08 | 1.33E-06 | CLD1 | VII | 713709 | 715046 | 1 |
| YGR111W | 38.055398 | -0.2495015 | 0.64351184 | 0.96700074 | 0.61662121 | 0.75817303 | NA | VII | 715828 | 717030 | 1 |
| YGR112W | 20.5537667 | -0.1427206 | 0.80779086 | 0.08851058 | 0.95670969 | 0.96960425 | SHY1 | VII | 717358 | 718527 | 1 |
| YGR113W | 73.5257086 | 0.07106274 | 0.5703577 | 3.34563767 | 0.18771718 | 0.34351531 | DAM1 | VII | 718893 | 719924 | 1 |
| YGR116W | 88.6879958 | -1.2564003 | 0.54664571 | 7.96076362 | 0.01867851 | 0.06284053 | SPT6 | VII | 720409 | 724764 | 1 |
| YGR117C | 48.7655781 | -0.2580078 | 0.59907679 | 0.19527579 | 0.90697727 | 0.94229291 | NA | VII | 725049 | 726479 | -1 |
| YGR118W | 120.90057 | -2.4709783 | 0.71353529 | 11.2352178 | 0.00363332 | 0.01781991 | RPS23B | VII | 726974 | 727730 | 1 |
| YGR119C | 48.6749629 | -0.0542447 | 0.66051552 | 0.27233374 | 0.87269699 | 0.92527295 | NUP57 | VII | 728046 | 729671 | -1 |
| YGR120C | 138.686245 | 0.62917071 | 0.53508433 | 2.12024834 | 0.34641279 | 0.51988856 | COG2 | VII | 730033 | 730821 | -1 |
| YGR121C | 1687.63223 | 0.96643691 | 0.56054502 | 2.96803705 | 0.22672476 | 0.39279962 | MEP1 | VII | 731449 | 732927 | -1 |
| YGR121W-A | 26.0304755 | 0.10879305 | 0.69844992 | 1.57585017 | 0.45478746 | 0.62066785 | NA | VII | 733412 | 733627 | 1 |
| YGR122W | 57.6364419 | -1.6135323 | 0.59191736 | 9.79066531 | 0.00748142 | 0.03161856 | NA | VII | 733935 | 735143 | 1 |
| YGR123C | 15.3340432 | -0.4667924 | 0.80137975 | 0.42009574 | 0.81054544 | 0.88665371 | PPT1 | VII | 736662 | 738203 | -1 |
| YGR124W | 70.08196 | -0.9008533 | 0.61835164 | 8.02269414 | 0.01810898 | 0.06142415 | ASN2 | VII | 739944 | 741662 | 1 |
| YGR125W | 71.7735703 | 1.44141322 | 0.58487183 | 6.56275786 | 0.03757641 | 0.10618223 | NA | VII | 742325 | 745435 | 1 |
| YGR126W | 40.2294814 | 1.49225566 | 0.60942062 | 10.4301303 | 0.00543408 | 0.02450414 | NA | VII | 745835 | 746527 | 1 |
| YGR127W | 27.6826594 | 0.69600635 | 0.67448227 | 2.51534257 | 0.28431535 | 0.45790983 | NA | VII | 746798 | 747736 | 1 |
| YGR128C | 31.1274652 | -0.3243424 | 0.92941013 | 2.52822455 | 0.28248996 | 0.45662112 | UTP8 | VII | 747950 | 750091 | -1 |
| YGR129W | 16.33605 | 0.02974836 | 0.87570383 | 0.31677161 | 0.85352043 | 0.91313438 | SYF2 | VII | 750400 | 751047 | 1 |
| YGR130C | 705.621384 | -1.3490752 | 0.49316418 | 11.2855965 | 0.00354294 | 0.01744356 | NA | VII | 751394 | 753844 | -1 |
| YGR131W | 7.60574127 | 0.07655919 | 0.99977331 | 0.63091881 | 0.72945368 | 0.83678999 | FHN1 | VII | 754726 | 755250 | 1 |
| YGR132C | 18.9169162 | -0.71842 | 0.87645845 | 0.76049025 | 0.6836938 | 0.80606538 | PHB1 | VII | 755589 | 756452 | -1 |
| YGR133W | 475.228288 | 1.08493699 | 0.54216543 | 4.99481865 | 0.08229793 | 0.19005499 | PEX4 | VII | 756896 | 757447 | 1 |
| YGR134W | 80.7354192 | 1.54613126 | 0.6239 | 8.37051387 | 0.0152183 | 0.05412404 | CAF130 | VII | 757770 | 761138 | 1 |

| | | | | | | | | | | | |
|---|---|---|---|---|---|---|---|---|---|---|---|
| YGR135W | 33.8072402 | 0.40156976 | 0.7399383 | 0.59073603 | 0.74425764 | 0.84379317 | PRE9 | VII | 761392 | 762168 | 1 |
| YGR136W | 48.4026226 | 1.49837814 | 0.69377011 | 6.39578997 | 0.0408481 | 0.11305473 | LSB1 | VII | 762428 | 763153 | 1 |
| YGR138C | 105.427501 | -0.7351005 | 0.52753214 | 2.50858948 | 0.28527697 | 0.45887782 | TPO2 | VII | 763762 | 765606 | -1 |
| YGR140W | 60.1304181 | 1.13535557 | 0.56499111 | 7.78591445 | 0.02038497 | 0.06703554 | CBF2 | VII | 767429 | 770299 | 1 |
| YGR141W | 604.27726 | 2.08300698 | 0.56858409 | 21.528309 | 2.11E-05 | 0.00024883 | VPS62 | VII | 770569 | 771972 | 1 |
| YGR142W | 118.993445 | 1.88614036 | 0.53011788 | 12.1728427 | 0.00227353 | 0.01230256 | BTN2 | VII | 772454 | 773686 | 1 |
| YGR143W | 88.5915044 | -1.1030765 | 0.52711016 | 9.39042732 | 0.00913891 | 0.03682277 | SKN1 | VII | 775193 | 777508 | 1 |
| YGR144W | 52.7161306 | 2.19891362 | 0.65777116 | 11.2168034 | 0.00366693 | 0.01794717 | THI4 | VII | 780399 | 781379 | 1 |
| YGR145W | 55.3857875 | 0.16152324 | 0.63213927 | 0.84410948 | 0.65569814 | 0.78495521 | ENP2 | VII | 781767 | 783890 | 1 |
| YGR146C | 146.017916 | -1.6869092 | 0.52125663 | 15.8448366 | 0.00036252 | 0.0028442 | ECL1 | VII | 784223 | 784858 | -1 |
| YGR146C-A | 62.8746921 | 1.00687445 | 0.64154589 | 4.66063954 | 0.09726464 | 0.21492841 | NA | VII | 785276 | 785437 | -1 |
| YGR147C | 46.9472286 | 0.9587042 | 0.60684838 | 2.78725981 | 0.24817282 | 0.41904947 | NAT2 | VII | 786054 | 786920 | -1 |
| YGR148C | 115.845923 | -3.0899082 | 0.66502 | 19.4452243 | 5.99E-05 | 0.0006041 | RPL24B | VII | 787312 | 787779 | -1 |
| YGR149W | 26.8801178 | 1.49383973 | 0.76475489 | 4.02653752 | 0.13355141 | 0.26953536 | GPC1 | VII | 789031 | 790329 | 1 |
| YGR150C | 461.861971 | 2.6295724 | 0.77858943 | 22.1901252 | 1.52E-05 | 0.0001887 | CCM1 | VII | 790459 | 793053 | -1 |
| YGR152C | 15.4282094 | -0.5447597 | 0.86080684 | 0.52663859 | 0.76849648 | 0.85932689 | RSR1 | VII | 794674 | 795492 | -1 |
| YGR153W | 443.091929 | 0.96057058 | 0.6482324 | 2.16441993 | 0.33884586 | 0.5121163 | NA | VII | 796092 | 796745 | 1 |
| YGR154C | 176.016378 | 1.40396285 | 0.50426321 | 7.50626736 | 0.02344416 | 0.07454061 | GTO1 | VII | 796798 | 797868 | -1 |
| YGR155W | 43.4153566 | -2.1794327 | 0.72401816 | 8.75240824 | 0.01257299 | 0.0468152 | CYS4 | VII | 798543 | 800066 | 1 |
| YGR156W | 348.763949 | -3.257873 | 0.49614747 | 44.5873923 | 2.08E-10 | 8.99E-09 | PTI1 | VII | 800546 | 801823 | 1 |
| YGR157W | 107.386806 | -1.9404901 | 0.58240741 | 10.6492989 | 0.00487006 | 0.02241465 | CHO2 | VII | 802440 | 805049 | 1 |
| YGR158C | 63.2640227 | -3.1957991 | 0.6221004 | 26.2067035 | 2.04E-06 | 3.30E-05 | MTR3 | VII | 805269 | 806021 | -1 |
| YGR159C | 105.827163 | -4.2060032 | 0.66200763 | 35.6141783 | 1.85E-08 | 5.17E-07 | NSR1 | VII | 806412 | 807656 | -1 |
| YGR161C | 218.737636 | -0.7077329 | 0.48569143 | 10.4898829 | 0.00527413 | 0.0239164 | RTS3 | VII | 808629 | 809420 | -1 |
| YGR161W-C | 7.76058394 | -0.3849992 | 0.97280138 | 0.26381337 | 0.87642278 | 0.9271858 | NA | VII | 810227 | 810505 | 1 |
| YGR162W | 169.953386 | -2.3698603 | 0.54553497 | 17.5573328 | 0.00015398 | 0.00136743 | TIF4631 | VII | 824059 | 826917 | 1 |
| YGR163W | 429.819583 | 3.90478448 | 0.51339495 | 50.8334345 | 9.16E-12 | 5.46E-10 | GTR2 | VII | 827552 | 828577 | 1 |
| YGR164W | 231.776972 | -3.5613919 | 0.55560701 | 35.1625593 | 2.31E-08 | 6.38E-07 | NA | VII | 828625 | 828960 | 1 |
| YGR165W | 23.8928623 | 0.06886738 | 0.78047516 | 0.65954003 | 0.7190891 | 0.82938751 | MRPS35 | VII | 829116 | 830153 | 1 |
| YGR166W | 184.91055 | 0.50396159 | 0.51756134 | 2.55661249 | 0.27850863 | 0.45301284 | TRS65 | VII | 830515 | 832197 | 1 |
| YGR167W | 73.2702201 | 0.44686478 | 0.64125553 | 1.79713679 | 0.40715212 | 0.57850572 | CLC1 | VII | 832456 | 833157 | 1 |
| YGR168C | 168.906165 | -1.6352308 | 0.51760366 | 9.60207823 | 0.0082212 | 0.03390367 | NA | VII | 833352 | 834482 | -1 |
| YGR169C | 11.6741497 | -1.085761 | 0.96045919 | 1.3201147 | 0.51682169 | 0.67350835 | PUS6 | VII | 834689 | 835903 | -1 |

| | | | | | | | | | | | |
|---|---|---|---|---|---|---|---|---|---|---|---|
| YGR169C-A | 76.8577513 | -1.4126292 | 0.62230648 | 7.62628884 | 0.02207865 | 0.07131784 | LSO2 | VII | 836387 | 836665 | -1 |
| YGR170W | 494.170442 | -1.5413419 | 0.49232701 | 12.2370462 | 0.00220171 | 0.01203765 | PSD2 | VII | 837142 | 840558 | 1 |
| YGR171C | 119.285 | 0.00439794 | 0.76600146 | 4.47049971 | 0.1069654 | 0.22980693 | MSM1 | VII | 840824 | 842551 | -1 |
| YGR172C | 24.4945857 | -0.6590532 | 0.69055148 | 0.96537847 | 0.61712157 | 0.75846612 | YIP1 | VII | 842845 | 843591 | -1 |
| YGR173W | 37.654227 | 1.19030595 | 0.64411656 | 9.2062361 | 0.01002054 | 0.03951413 | RBG2 | VII | 843854 | 844960 | 1 |
| YGR174C | 1464.32768 | 4.80868509 | 0.59294903 | 49.0390478 | 2.25E-11 | 1.23E-09 | NA | VII | 845893 | 846405 | -1 |
| YGR174W-A | 43.214005 | 0.16384791 | 1.89209326 | 5.67922466 | 0.05844832 | 0.14688185 | NA | VII | 846655 | 846741 | 1 |
| YGR175C | 52.8898445 | -0.5422179 | 0.86359255 | 1.25566161 | 0.53374835 | 0.68828573 | ERG1 | VII | 846933 | 848423 | -1 |
| YGR177C | 127.964679 | 0.81789923 | 0.52458919 | 2.85359498 | 0.24007654 | 0.40823588 | ATF2 | VII | 848829 | 850436 | -1 |
| YGR178C | 258.261136 | -0.1993587 | 0.4769835 | 1.93114047 | 0.38076601 | 0.55448572 | PBP1 | VII | 851047 | 853215 | -1 |
| YGR180C | 28.4182515 | -1.0646979 | 0.6710868 | 2.71433615 | 0.25738865 | 0.42958807 | RNR4 | VII | 855264 | 856301 | -1 |
| YGR181W | 39.5652367 | -1.223016 | 0.77217849 | 2.98279326 | 0.22505811 | 0.3904964 | TIM13 | VII | 858287 | 858604 | 1 |
| YGR183C | 41.6803678 | 0.74011734 | 0.62892653 | 3.94035788 | 0.1394319 | 0.27799956 | QCR9 | VII | 859063 | 859476 | -1 |
| YGR184C | 53.7244738 | 0.61724493 | 0.61455994 | 5.5499553 | 0.06235087 | 0.15421253 | UBR1 | VII | 859901 | 865753 | -1 |
| YGR185C | 75.9300681 | -1.5166025 | 0.57847062 | 8.26704451 | 0.01602633 | 0.05640572 | TYS1 | VII | 866336 | 867520 | -1 |
| YGR186W | 49.484178 | -0.2193461 | 0.69462439 | 0.88991661 | 0.640851 | 0.77561669 | TFG1 | VII | 867774 | 869981 | 1 |
| YGR187C | 64.4279703 | -1.5025323 | 0.60317329 | 6.32106129 | 0.04240323 | 0.11597295 | HGH1 | VII | 870232 | 871416 | -1 |
| YGR188C | 40.9285061 | 1.85994596 | 0.63082159 | 12.6056569 | 0.00183112 | 0.01050761 | BUB1 | VII | 872044 | 875109 | -1 |
| YGR189C | 235.206657 | 2.03486657 | 0.58458968 | 27.1986945 | 1.24E-06 | 2.13E-05 | CRH1 | VII | 876669 | 878192 | -1 |
| YGR191W | 63.6451159 | 0.80700886 | 0.54910328 | 3.45152953 | 0.17803684 | 0.33145766 | HIP1 | VII | 880420 | 882231 | 1 |
| YGR192C | 842.365229 | -1.4667266 | 0.72954401 | 4.06058018 | 0.13129743 | 0.26636724 | TDH3 | VII | 882812 | 883810 | -1 |
| YGR193C | 62.439846 | 0.46209728 | 0.57447373 | 0.66522568 | 0.71704775 | 0.82851856 | PDX1 | VII | 884509 | 885741 | -1 |
| YGR194C | 69.7169034 | -0.9340431 | 0.54563106 | 3.40724241 | 0.18202319 | 0.33713158 | XKS1 | VII | 886073 | 887875 | -1 |
| YGR195W | 111.640546 | -1.0423644 | 0.80613233 | 5.78203506 | 0.05551969 | 0.14148724 | SKI6 | VII | 888882 | 889622 | 1 |
| YGR196C | 132.805055 | -1.2002612 | 0.53036831 | 5.42902475 | 0.06623724 | 0.16143398 | FYV8 | VII | 889733 | 892186 | -1 |
| YGR197C | 38.6617457 | 0.08135837 | 0.71084312 | 0.13949929 | 0.93262728 | 0.95731799 | SNG1 | VII | 892497 | 894140 | -1 |
| YGR198W | 41.3439075 | -0.6203303 | 0.69191099 | 1.10760826 | 0.57475919 | 0.72422968 | YPP1 | VII | 894693 | 897146 | 1 |
| YGR199W | 257.938692 | -1.3218275 | 0.54557427 | 6.91364023 | 0.03152986 | 0.09285753 | PMT6 | VII | 897502 | 899781 | 1 |
| YGR200C | 461.624139 | -0.3061582 | 0.64096301 | 2.74200019 | 0.25385296 | 0.42590803 | ELP2 | VII | 899904 | 902270 | -1 |
| YGR201C | 134.039587 | -1.9696589 | 0.56469056 | 14.0082707 | 0.00090812 | 0.00608566 | NA | VII | 902520 | 903197 | -1 |
| YGR202C | 196.54431 | 3.26089777 | 0.53404387 | 38.736485 | 3.88E-09 | 1.27E-07 | PCT1 | VII | 903474 | 904748 | -1 |
| YGR203W | 198.24288 | -0.1778782 | 0.52320498 | 1.44519388 | 0.48548983 | 0.64773825 | YCH1 | VII | 905237 | 905683 | 1 |
| YGR204C-A | 235.032762 | -1.1845298 | 0.63693089 | 3.58534796 | 0.16651432 | 0.31564276 | NA | VII | 909061 | 909174 | -1 |

| | | | | | | | | | | | |
|---|---|---|---|---|---|---|---|---|---|---|---|
| YGR204W | 97.1955208 | -0.062647 | 0.60566107 | 0.02888551 | 0.98566104 | 0.98832468 | ADE3 | VII | 905934 | 908774 | 1 |
| YGR205W | 95.4220637 | -1.3515434 | 0.58124621 | 5.48752119 | 0.06432798 | 0.15782161 | TDA10 | VII | 909213 | 910085 | 1 |
| YGR206W | 209.35029 | -0.4203969 | 0.95200694 | 7.69390051 | 0.02134473 | 0.06946937 | MVB12 | VII | 910432 | 910737 | 1 |
| YGR207C | 18.4804746 | -0.6324082 | 0.84421568 | 6.5936193 | 0.03700103 | 0.10476085 | CIR1 | VII | 910843 | 911628 | -1 |
| YGR208W | 19.8237596 | -1.3142873 | 0.78581185 | 4.52631097 | 0.10402173 | 0.22540636 | SER2 | VII | 911883 | 912812 | 1 |
| YGR209C | 34.2640386 | -1.0973338 | 0.84016145 | 3.55821065 | 0.16878909 | 0.3187519 | TRX2 | VII | 912913 | 913227 | -1 |
| YGR210C | 59.5265427 | -0.3556859 | 0.63231504 | 0.57633476 | 0.74963611 | 0.8479727 | NA | VII | 913503 | 914738 | -1 |
| YGR211W | 44.1599467 | -0.8730994 | 0.67970755 | 1.84772661 | 0.39698241 | 0.56785755 | ZPR1 | VII | 915241 | 916701 | 1 |
| YGR212W | 4335.86846 | 4.29700751 | 0.45219073 | 85.2404735 | 3.09E-19 | 7.16E-17 | SLI1 | VII | 917040 | 918446 | 1 |
| YGR213C | 418.861086 | -1.4226146 | 0.49561468 | 8.58854765 | 0.01364648 | 0.04972505 | RTA1 | VII | 918512 | 919465 | -1 |
| YGR214W | 163.065089 | -2.6541656 | 0.73861415 | 11.7007423 | 0.00287883 | 0.01475082 | RPS0A | VII | 920575 | 921788 | 1 |
| YGR215W | 92.3887364 | -0.7106258 | 0.9944898 | 9.74026212 | 0.00767236 | 0.03212072 | RSM27 | VII | 922175 | 922507 | 1 |
| YGR216C | 48.1609999 | -2.4295255 | 0.65173734 | 13.4742163 | 0.00118607 | 0.00756665 | GPI1 | VII | 922633 | 924462 | -1 |
| YGR217W | 44.9150882 | -0.1035599 | 0.58738148 | 0.28315875 | 0.86798628 | 0.9224744 | CCH1 | VII | 924696 | 930815 | 1 |
| YGR218W | 27.8598754 | -0.3778559 | 0.88045818 | 0.44726714 | 0.79960808 | 0.87884032 | CRM1 | VII | 932541 | 935795 | 1 |
| YGR220C | 12.989591 | 2.06261482 | 0.91579303 | 5.26301338 | 0.07196994 | 0.17226374 | MRPL9 | VII | 936074 | 936883 | -1 |
| YGR221C | 1295.30671 | 2.05469394 | 0.46762361 | 18.3410102 | 0.00010406 | 0.00096714 | TOS2 | VII | 937124 | 938992 | -1 |
| YGR222W | 48.5327019 | 0.68227606 | 0.89458765 | 4.31385617 | 0.11567993 | 0.24294045 | PET54 | VII | 939923 | 940804 | 1 |
| YGR223C | 109.292284 | 0.55517877 | 0.84809506 | 7.50521731 | 0.02345648 | 0.07454061 | HSV2 | VII | 940869 | 942215 | -1 |
| YGR224W | 1692.29434 | 1.37113117 | 0.50220996 | 7.27102708 | 0.02637039 | 0.08074276 | AZR1 | VII | 942806 | 944647 | 1 |
| YGR225W | 5.67939493 | 0.31456998 | 1.10841384 | 0.24394526 | 0.8851726 | 0.9315066 | AMA1 | VII | 945145 | 947019 | 1 |
| YGR227C-A | 319.653433 | -1.6310426 | 0.72273206 | 8.19299688 | 0.01663081 | 0.05772162 | NA | NA | NA | NA | NA |
| YGR227W | 19.2107175 | -0.3558434 | 0.78370677 | 0.24450795 | 0.88492359 | 0.9315066 | DIE2 | VII | 947420 | 948997 | 1 |
| YGR229C | 129.926616 | -0.8612159 | 0.59608966 | 3.15109206 | 0.20689455 | 0.36890584 | SMI1 | VII | 949382 | 950899 | -1 |
| YGR230W | 8.65971981 | 0.63176452 | 1.00344592 | 0.62486269 | 0.73166586 | 0.83702784 | BNS1 | VII | 951894 | 952307 | 1 |
| YGR231C | 32.684256 | 0.01907553 | 0.72815295 | 0.09112579 | 0.95545951 | 0.96850675 | PHB2 | VII | 952548 | 953480 | -1 |
| YGR232W | 128.454096 | 0.30928335 | 0.86665625 | 7.43968003 | 0.02423785 | 0.07639473 | NAS6 | VII | 953960 | 954646 | 1 |
| YGR233C | 176.424631 | 0.16869785 | 0.8241732 | 5.09092484 | 0.07843677 | 0.18386596 | PHO81 | VII | 954674 | 958210 | -1 |
| YGR234W | 107.342206 | -2.076718 | 0.57710354 | 13.182497 | 0.00137233 | 0.00850724 | YHB1 | VII | 959904 | 961103 | 1 |
| YGR235C | 1621.45257 | -3.5459673 | 0.74378134 | 24.3122809 | 5.26E-06 | 7.46E-05 | MIC26 | VII | 961360 | 962061 | -1 |
| YGR237C | 174.551398 | 0.58026896 | 0.50019177 | 1.44367108 | 0.48585962 | 0.64773825 | NA | VII | 963298 | 965655 | -1 |
| YGR238C | 93.8336373 | 1.71641478 | 0.53647393 | 15.1310767 | 0.000518 | 0.00378167 | KEL2 | VII | 966039 | 968687 | -1 |
| YGR239C | 326.972644 | -0.8669481 | 0.64378817 | 5.24021826 | 0.07279492 | 0.17359249 | PEX21 | VII | 969187 | 970053 | -1 |

| | | | | | | | | | | | |
|---|---|---|---|---|---|---|---|---|---|---|---|
| YGR240C | 109.310946 | -0.7165178 | 0.71389555 | 1.35878789 | 0.50692412 | 0.66464916 | PFK1 | VII | 970771 | 973734 | -1 |
| YGR240C-A | 804.206074 | -0.3446753 | 0.54031617 | 0.41157427 | 0.81400633 | 0.88825794 | NA | VII | 974577 | 974777 | -1 |
| YGR241C | 32.5535381 | -0.1296853 | 0.76278446 | 3.18882778 | 0.20302749 | 0.3631539 | YAP1802 | VII | 974875 | 976581 | -1 |
| YGR244C | 86.7203419 | -1.7887201 | 0.57073113 | 10.6000297 | 0.00499152 | 0.02286464 | LSC2 | VII | 978036 | 979319 | -1 |
| YGR245C | 68.6136283 | -2.2553351 | 0.55799729 | 18.6177839 | 9.06E-05 | 0.0008601 | SDA1 | VII | 979765 | 982068 | -1 |
| YGR246C | 13.1341319 | -0.8055887 | 0.93109698 | 0.8753967 | 0.64552047 | 0.77784881 | BRF1 | VII | 982482 | 984272 | -1 |
| YGR247W | 10.6784688 | 0.57640911 | 0.90200696 | 0.88786883 | 0.64150749 | 0.77591986 | CPD1 | VII | 984966 | 985685 | 1 |
| YGR248W | 80.3010328 | 1.74746552 | 0.56375833 | 12.1812365 | 0.00226401 | 0.01228548 | SOL4 | VII | 985972 | 986739 | 1 |
| YGR249W | 83.3010538 | -1.4361716 | 0.57555953 | 7.28080831 | 0.02624174 | 0.08047651 | MGA1 | VII | 988049 | 989419 | 1 |
| YGR250C | 60.7895579 | 1.23114958 | 0.78360612 | 2.54629998 | 0.2799484 | 0.45403395 | RIE1 | VII | 991176 | 993521 | -1 |
| YGR251W | 20.5919623 | -0.9513898 | 0.77373878 | 1.53565129 | 0.46402092 | 0.62714312 | NOP19 | VII | 995639 | 996229 | 1 |
| YGR252W | 205.520044 | -0.8125221 | 0.54121909 | 2.70616602 | 0.25844225 | 0.43011803 | GCN5 | VII | 996869 | 998188 | 1 |
| YGR253C | 226.115219 | -0.8172226 | 0.68055837 | 5.09700013 | 0.07819887 | 0.18338253 | PUP2 | VII | 998358 | 999140 | -1 |
| YGR254W | 147.977753 | 0.63453423 | 0.58219407 | 1.95224343 | 0.37676949 | 0.55032677 | ENO1 | VII | 1000927 | 1E+06 | 1 |
| YGR255C | 17.171899 | -0.0421349 | 0.88013121 | 0.71211645 | 0.70043184 | 0.81801018 | COQ6 | VII | 1002523 | 1E+06 | -1 |
| YGR256W | 71.5619089 | 0.25094026 | 0.55666287 | 0.41732741 | 0.81166815 | 0.88731158 | GND2 | VII | 1004624 | 1E+06 | 1 |
| YGR257C | 57.6640789 | 1.17664233 | 0.80560546 | 4.81148772 | 0.09019838 | 0.20366605 | MTM1 | VII | 1006205 | 1E+06 | -1 |
| YGR258C | 53.0120634 | 1.29351435 | 0.6124003 | 4.49101939 | 0.10587356 | 0.22838155 | RAD2 | VII | 1007671 | 1E+06 | -1 |
| YGR260W | 215.488257 | 2.39764521 | 0.50250329 | 27.053088 | 1.34E-06 | 2.27E-05 | TNA1 | VII | 1012485 | 1E+06 | 1 |
| YGR261C | 33.9983192 | -0.5234263 | 0.75406934 | 0.49236824 | 0.78177827 | 0.86750947 | APL6 | VII | 1014321 | 1E+06 | -1 |
| YGR262C | 152.903638 | 4.23536154 | 0.63319399 | 60.6418372 | 6.79E-14 | 5.78E-12 | BUD32 | VII | 1016974 | 1E+06 | -1 |
| YGR263C | 89.1015434 | -0.1830487 | 0.5359846 | 1.11381669 | 0.57297778 | 0.72277589 | SAY1 | VII | 1017967 | 1E+06 | -1 |
| YGR264C | 32.8954497 | -1.6777548 | 0.78879435 | 4.80145616 | 0.09065193 | 0.20427392 | MES1 | VII | 1019598 | 1E+06 | -1 |
| YGR266W | 358.493564 | -1.7760848 | 0.50667888 | 11.8572797 | 0.0026621 | 0.01389861 | NA | VII | 1022656 | 1E+06 | 1 |
| YGR267C | 156.231856 | 0.78312045 | 0.92663752 | 10.5144304 | 0.00520979 | 0.02376215 | FOL2 | VII | 1025004 | 1E+06 | -1 |
| YGR268C | 52.5221446 | 0.01863132 | 0.61455063 | 1.43737684 | 0.48739109 | 0.64903366 | HUA1 | VII | 1026057 | 1E+06 | -1 |
| YGR270W | 114.552403 | 0.32027275 | 0.57747165 | 2.89857517 | 0.23473746 | 0.40225741 | YTA7 | VII | 1027370 | 1E+06 | 1 |
| YGR271C-A | 121.595902 | 0.47404469 | 1.04460481 | 8.5264338 | 0.01407695 | 0.05103664 | EFG1 | VII | 1037800 | 1E+06 | -1 |
| YGR271W | 30.7590597 | -0.503524 | 0.63192569 | 0.6394027 | 0.72636593 | 0.83478737 | SLH1 | VII | 1031791 | 1E+06 | 1 |
| YGR274C | 22.298224 | 1.03227674 | 0.74857917 | 2.52308617 | 0.28321666 | 0.45678444 | TAF1 | VII | 1039895 | 1E+06 | -1 |
| YGR275W | 59.4047774 | 0.51010786 | 0.9176289 | 11.6542639 | 0.00294652 | 0.01503761 | RTT102 | VII | 1043276 | 1E+06 | 1 |
| YGR276C | 234.277947 | -0.7087749 | 0.77382024 | 8.06330574 | 0.01774498 | 0.06033083 | RNH70 | VII | 1043819 | 1E+06 | -1 |
| YGR277C | 23.1025815 | -0.1744874 | 0.71944107 | 0.07095063 | 0.96514656 | 0.97423267 | CAB4 | VII | 1045644 | 1E+06 | -1 |

| | | | | | | | | | | | |
|---|---|---|---|---|---|---|---|---|---|---|---|
| YGR278W | 44.9578612 | -0.8445091 | 0.67433085 | 1.96861682 | 0.37369758 | 0.54680541 | CWC22 | VII | 1046731 | 1E+06 | 1 |
| YGR279C | 58.2729932 | -0.4779222 | 0.57356259 | 0.71729601 | 0.69862022 | 0.81736153 | SCW4 | VII | 1048798 | 1E+06 | -1 |
| YGR280C | 186.52823 | -3.1677545 | 0.68639144 | 18.5528096 | 9.36E-05 | 0.00088415 | PXR1 | VII | 1050910 | 1E+06 | -1 |
| YGR281W | 83.255006 | 1.54451085 | 0.62997885 | 5.97226662 | 0.05048226 | 0.13160391 | YOR1 | VII | 1052824 | 1E+06 | 1 |
| YGR282C | 71.2078076 | 0.95005782 | 0.69183807 | 2.17235765 | 0.33750369 | 0.51075441 | BGL2 | VII | 1057783 | 1E+06 | -1 |
| YGR283C | 2033.14542 | 3.2417331 | 0.52746166 | 32.0801803 | 1.08E-07 | 2.58E-06 | NA | VII | 1059015 | 1E+06 | -1 |
| YGR284C | 60.2713665 | -1.8335774 | 0.61180507 | 8.66982276 | 0.01310304 | 0.04820096 | ERV29 | VII | 1060658 | 1E+06 | -1 |
| YGR285C | 135.562276 | -1.3019743 | 0.54455202 | 7.03063095 | 0.02973842 | 0.08839089 | ZUO1 | VII | 1061852 | 1E+06 | -1 |
| YGR286C | 9.19733501 | -0.270477 | 0.93717373 | 0.15090866 | 0.92732208 | 0.95452353 | BIO2 | VII | 1063813 | 1E+06 | -1 |
| YGR288W | 59.4154946 | 2.48279957 | 0.59641801 | 23.3112355 | 8.67E-06 | 0.0001162 | MAL13 | VII | 1070293 | 1E+06 | 1 |
| YGR289C | 45.2200273 | 2.80919123 | 0.80687749 | 11.9246481 | 0.00257392 | 0.0135482 | MAL11 | VII | 1073963 | 1E+06 | -1 |
| YGR292W | 41.5154763 | -1.4067951 | 0.61270763 | 5.86858222 | 0.0531684 | 0.13694173 | MAL12 | VII | 1076599 | 1E+06 | 1 |
| YGR295C | 109.590786 | -1.9736638 | 0.51782521 | 18.7169411 | 8.62E-05 | 0.00082519 | COS6 | VII | 1081584 | 1E+06 | -1 |
| YGR296W | 11.8970325 | -1.2865253 | 0.95186631 | 2.1696518 | 0.33796062 | 0.51117868 | YRF1-7 | VII | 1084864 | 1E+06 | 1 |
| YHL001W | 25.5104093 | -1.4593219 | 0.77729881 | 3.54060489 | 0.17028148 | 0.32052333 | RPL14B | VIII | 104277 | 105091 | 1 |
| YHL002W | 135.650715 | -0.9893187 | 0.53767574 | 4.11195872 | 0.12796745 | 0.26147876 | HSE1 | VIII | 102612 | 103970 | 1 |
| YHL003C | 13.4117336 | 0.25713349 | 0.94142692 | 0.11822599 | 0.94260025 | 0.96176085 | LAG1 | VIII | 100648 | 101883 | -1 |
| YHL004W | 135.473134 | -0.4253294 | 0.8774363 | 11.169576 | 0.00375455 | 0.01825258 | MRP4 | VIII | 99219 | 100403 | 1 |
| YHL005C | 349.246839 | -0.902283 | 0.52385756 | 3.70775326 | 0.1566288 | 0.30229358 | NA | VIII | 98828 | 99220 | -1 |
| YHL006C | 66.2879648 | -3.2220307 | 0.69696507 | 21.0393022 | 2.70E-05 | 0.00030896 | SHU1 | VIII | 98343 | 98795 | -1 |
| YHL007C | 105.801095 | 0.40463207 | 0.51340155 | 1.68348061 | 0.43095987 | 0.5996774 | STE20 | VIII | 95118 | 97937 | -1 |
| YHL008C | 23.6077361 | 1.63187955 | 0.73301123 | 5.16764693 | 0.07548484 | 0.17824519 | NA | VIII | 92627 | 94510 | -1 |
| YHL009C | 32.4671778 | 2.20647163 | 0.69426397 | 10.5200798 | 0.0051951 | 0.02372209 | YAP3 | VIII | 84068 | 85060 | -1 |
| YHL010C | 90.8417042 | 3.09762581 | 0.61501207 | 25.6126808 | 2.74E-06 | 4.24E-05 | ETP1 | VIII | 81964 | 83721 | -1 |
| YHL011C | 22.6945235 | -0.8637643 | 0.80886489 | 1.20353933 | 0.54784128 | 0.70011119 | PRS3 | VIII | 80654 | 81616 | -1 |
| YHL012W | 1212.05141 | -2.8050332 | 0.4809347 | 29.9234555 | 3.18E-07 | 6.55E-06 | NA | VIII | 78936 | 80417 | 1 |
| YHL013C | 12.0078999 | -1.0858008 | 0.89558151 | 2.16368882 | 0.33896975 | 0.51216985 | OTU2 | VIII | 77430 | 78353 | -1 |
| YHL014C | 90.9708135 | -1.3222483 | 0.7467931 | 5.2757247 | 0.07151398 | 0.1715269 | YLF2 | VIII | 76097 | 77314 | -1 |
| YHL015W | 220.969664 | -3.1777199 | 0.69411472 | 18.2458983 | 0.00010913 | 0.00100778 | RPS20 | VIII | 75412 | 75777 | 1 |
| YHL015W-A | 454.204598 | 2.12938993 | 0.56515126 | 15.4717491 | 0.00043687 | 0.00329789 | NA | VIII | 74699 | 74782 | 1 |
| YHL016C | 128.610351 | 0.76545822 | 0.70338573 | 2.29334573 | 0.31769202 | 0.490255 | DUR3 | VIII | 72037 | 74244 | -1 |
| YHL017W | 18.406319 | 0.48468983 | 0.87629814 | 1.9109279 | 0.38463365 | 0.55747464 | NA | VIII | 70276 | 71874 | 1 |
| YHL019C | 84.9176728 | 0.29339434 | 0.57924112 | 0.50613866 | 0.77641405 | 0.8639961 | APM2 | VIII | 67731 | 69548 | -1 |

| | | | | | | | | | | | |
|---|---|---|---|---|---|---|---|---|---|---|---|
| YHL020C | 44.0676708 | 1.33358605 | 0.62455422 | 4.45912372 | 0.10757555 | 0.23077356 | OPI1 | VIII | 66242 | 67456 | -1 |
| YHL021C | 148.495758 | 0.03841923 | 0.51749328 | 0.40721634 | 0.81578196 | 0.88902268 | AIM17 | VIII | 64462 | 65859 | -1 |
| YHL022C | 111.502574 | -0.8997381 | 0.5584403 | 3.66490427 | 0.16002069 | 0.30728192 | SPO11 | VIII | 62961 | 64157 | -1 |
| YHL023C | 507.979672 | 0.67716507 | 0.49776681 | 4.81110709 | 0.09021554 | 0.20366605 | NPR3 | VIII | 59123 | 62563 | -1 |
| YHL024W | 2618.10836 | 1.12378976 | 0.49328913 | 5.61624845 | 0.06031803 | 0.1506 | RIM4 | VIII | 56649 | 58790 | 1 |
| YHL025W | 51.2035016 | -0.2961973 | 0.57385696 | 0.69937661 | 0.70490777 | 0.82055541 | SNF6 | VIII | 54851 | 55849 | 1 |
| YHL026C | 496.13448 | 1.98636826 | 0.50393081 | 14.7264565 | 0.00063415 | 0.00451071 | NA | VIII | 53219 | 54166 | -1 |
| YHL027W | 59.9396542 | 1.35698177 | 0.61287098 | 5.80929761 | 0.05476802 | 0.14006486 | RIM101 | VIII | 51111 | 52988 | 1 |
| YHL028W | 45.0019699 | 0.32036365 | 0.58078944 | 0.48033087 | 0.78649774 | 0.87021248 | WSC4 | VIII | 48763 | 50580 | 1 |
| YHL029C | 63.5717298 | 1.30233124 | 0.57862856 | 10.6325174 | 0.00491109 | 0.02256764 | OCA5 | VIII | 45929 | 47968 | -1 |
| YHL030W | 20.3851726 | -1.0924036 | 0.76443525 | 2.63177818 | 0.26823573 | 0.44109199 | ECM29 | VIII | 40084 | 45690 | 1 |
| YHL031C | 11.4256395 | -0.6497493 | 0.95450006 | 0.89127111 | 0.64041713 | 0.77561669 | GOS1 | VIII | 38815 | 39486 | -1 |
| YHL032C | 30.9581941 | -0.725574 | 0.67937862 | 1.85925072 | 0.39470155 | 0.56638907 | GUT1 | VIII | 36379 | 38508 | -1 |
| YHL033C | 42.357202 | -2.3147717 | 0.81434408 | 8.43772017 | 0.01471541 | 0.05262645 | RPL8A | VIII | 35255 | 36025 | -1 |
| YHL034C | 76.8743048 | -0.2366418 | 0.76155385 | 1.55833874 | 0.45878693 | 0.62328904 | SBP1 | VIII | 33193 | 34077 | -1 |
| YHL035C | 86.8397376 | 1.46927642 | 0.61584601 | 8.64648396 | 0.01325684 | 0.048642 | VMR1 | VIII | 27978 | 32756 | -1 |
| YHL036W | 938.761105 | 0.32031348 | 0.71505135 | 5.89733458 | 0.05240951 | 0.13534837 | MUP3 | VIII | 26241 | 27881 | 1 |
| YHL037C | 21.3621546 | -0.8354878 | 0.79695775 | 1.65295951 | 0.43758699 | 0.60627631 | NA | VIII | 25778 | 26179 | -1 |
| YHL038C | 111.420251 | -1.4137826 | 0.66714065 | 4.36987831 | 0.11248458 | 0.23882864 | CBP2 | VIII | 23617 | 25509 | -1 |
| YHL039W | 43.6555215 | -0.284955 | 0.73084461 | 4.69040295 | 0.09582789 | 0.21233965 | EFM1 | VIII | 21783 | 23540 | 1 |
| YHL040C | 294.889157 | 6.10986759 | 0.69498693 | 77.8938087 | 1.22E-17 | 1.96E-15 | ARN1 | VIII | 19088 | 20971 | -1 |
| YHL041W | 10.3974162 | -0.2273581 | 1.06456654 | 0.11886156 | 0.94230076 | 0.96176085 | NA | VIII | 17392 | 17841 | 1 |
| YHL042W | 257.902833 | 0.34107083 | 0.47519581 | 0.53734376 | 0.76439403 | 0.85689088 | NA | VIII | 15667 | 16119 | 1 |
| YHL043W | 798.884725 | 0.97790107 | 0.47401177 | 5.83645398 | 0.0540294 | 0.13854305 | ECM34 | VIII | 14901 | 15413 | 1 |
| YHL044W | 4.68901863 | 0.87117018 | 1.34380598 | 0.56904621 | 0.75237297 | 0.85032979 | NA | VIII | 13565 | 14272 | 1 |
| YHL046C | 27.4657212 | 1.31764305 | 0.75131623 | 3.86462526 | 0.14481291 | 0.28548409 | PAU13 | VIII | 11923 | 12285 | -1 |
| YHL047C | 12.8027714 | -0.1845435 | 0.82823927 | 0.36497645 | 0.83319445 | 0.90138189 | ARN2 | VIII | 8351 | 10213 | -1 |
| YHL048W | 31.8990859 | 0.79599336 | 0.64681602 | 1.60286997 | 0.44868465 | 0.61504428 | COS8 | VIII | 6401 | 7546 | 1 |
| YHL049C | 8.0658079 | -1.3175281 | 1.02910104 | 7.74867497 | 0.02076809 | 0.0679363 | NA | VIII | 3726 | 4541 | -1 |
| YHL050C | 18.9656218 | 0.06773762 | 0.71931087 | 1.90348314 | 0.38606807 | 0.55869386 | NA | VIII | 445 | 3311 | -1 |
| YHR001W | 23.5327889 | 0.7484792 | 0.70174391 | 1.28228242 | 0.52669102 | 0.68176637 | OSH7 | VIII | 106055 | 107368 | 1 |
| YHR001W-A | 10.0558087 | 2.16237154 | 1.02504671 | 4.3365499 | 0.11437475 | 0.24133739 | QCR10 | VIII | 107826 | 108122 | 1 |
| YHR002W | 238.564713 | -0.1597082 | 0.60868171 | 1.69984657 | 0.42744772 | 0.59709268 | LEU5 | VIII | 108812 | 109885 | 1 |

| | | | | | | | | | | | |
|---|---|---|---|---|---|---|---|---|---|---|---|
| YHR003C | 884.033303 | 4.40215295 | 0.49067479 | 81.6306568 | 1.88E-18 | 3.96E-16 | TCD1 | VIII | 110028 | 111317 | -1 |
| YHR004C | 82.9089337 | -0.7286758 | 0.64070103 | 2.34293524 | 0.30991177 | 0.48301189 | NEM1 | VIII | 111754 | 113094 | -1 |
| YHR005C | 28.4260432 | -0.2313857 | 0.64694672 | 0.17179222 | 0.91768956 | 0.94899492 | GPA1 | VIII | 113499 | 114917 | -1 |
| YHR005C-A | 53.1704556 | -2.5631769 | 0.60782853 | 16.8381131 | 0.00022062 | 0.001854 | TIM10 | VIII | 115620 | 115901 | -1 |
| YHR006W | 41.360877 | 1.71060664 | 0.66403722 | 6.65401974 | 0.03590029 | 0.10239541 | STP2 | VIII | 117814 | 119439 | 1 |
| YHR007C | 92.2979884 | -0.7227993 | 0.7282901 | 0.99411042 | 0.6083194 | 0.75227879 | ERG11 | VIII | 120091 | 121683 | -1 |
| YHR007C-A | 242.988431 | 1.63405134 | 0.78562481 | 5.26184772 | 0.0720119 | 0.17229294 | NA | VIII | 122550 | 122765 | -1 |
| YHR008C | 59.9975433 | 0.25630832 | 0.64229021 | 0.33361048 | 0.84636443 | 0.90975468 | SOD2 | VIII | 122889 | 123590 | -1 |
| YHR009C | 53.6304476 | 1.79610649 | 0.60612424 | 13.741433 | 0.00103773 | 0.00679692 | TDA3 | VIII | 124109 | 125680 | -1 |
| YHR010W | 97.7733878 | -3.1157296 | 0.83716413 | 12.8378324 | 0.00163042 | 0.00966238 | RPL27A | VIII | 126521 | 127492 | 1 |
| YHR011W | 65.7708976 | -0.513325 | 0.57892434 | 0.78254188 | 0.67619692 | 0.8007253 | DIA4 | VIII | 127780 | 129120 | 1 |
| YHR012W | 27.1982144 | 0.61259473 | 0.7901673 | 0.99741464 | 0.60731522 | 0.75135793 | VPS29 | VIII | 129481 | 130448 | 1 |
| YHR013C | 112.836371 | 0.04849899 | 0.80168207 | 8.86892633 | 0.01186143 | 0.04474117 | ARD1 | VIII | 130730 | 131446 | -1 |
| YHR014W | 260.514882 | -1.3051621 | 0.51122914 | 6.44225442 | 0.03991005 | 0.11125622 | SPO13 | VIII | 132047 | 132922 | 1 |
| YHR015W | 14.1099866 | 3.37322906 | 1.00011237 | 12.4340258 | 0.0019952 | 0.01112927 | MIP6 | VIII | 134554 | 136533 | 1 |
| YHR016C | 931.345573 | 4.8016634 | 0.4838565 | 87.1562835 | 1.19E-19 | 3.27E-17 | YSC84 | VIII | 136881 | 138455 | -1 |
| YHR017W | 248.308182 | -1.235456 | 0.72540241 | 14.9393532 | 0.00057011 | 0.00410566 | YSC83 | VIII | 138694 | 139851 | 1 |
| YHR018C | 34.3587205 | 0.26100042 | 0.98653424 | 3.39474508 | 0.18316415 | 0.33860805 | ARG4 | VIII | 140011 | 141402 | -1 |
| YHR019C | 77.3617128 | -1.979666 | 0.57286023 | 12.3708701 | 0.00205921 | 0.01140938 | DED81 | VIII | 141894 | 143558 | -1 |
| YHR020W | 56.6220245 | -1.6167335 | 0.61576707 | 6.79057721 | 0.03353088 | 0.09746173 | NA | VIII | 143996 | 146062 | 1 |
| YHR021C | 13.860296 | -0.9504667 | 0.81660559 | 1.5443795 | 0.46200029 | 0.62543411 | RPS27B | VIII | 147871 | 148669 | -1 |
| YHR021W-A | 771.131606 | 0.33848879 | 0.50645246 | 37.0595574 | 8.97E-09 | 2.72E-07 | ECM12 | VIII | 149225 | 149680 | 1 |
| YHR022C-A | 31.7437473 | -0.4835327 | 0.64001435 | 0.88133781 | 0.64360576 | 0.77686719 | NA | VIII | 151217 | 151306 | -1 |
| YHR023W | 49.7546686 | -0.1799189 | 0.73699549 | 2.04283706 | 0.36008379 | 0.5332187 | MYO1 | VIII | 151666 | 157452 | 1 |
| YHR024C | 57.3428593 | -1.607555 | 0.56308022 | 8.48435489 | 0.01437625 | 0.051862 | MAS2 | VIII | 157744 | 159192 | -1 |
| YHR025W | 63.4415567 | -0.7176297 | 0.58792549 | 1.62006059 | 0.44484459 | 0.61266655 | THR1 | VIII | 159438 | 160511 | 1 |
| YHR026W | 13.6975987 | 0.49538383 | 0.81609729 | 0.94405818 | 0.62373537 | 0.76319268 | VMA16 | VIII | 160844 | 161485 | 1 |
| YHR027C | 190.13927 | -0.2741668 | 0.57131998 | 2.53894377 | 0.28097997 | 0.45506966 | RPN1 | VIII | 161730 | 164711 | -1 |
| YHR028C | 462.172824 | -0.9503955 | 0.46691771 | 4.81475821 | 0.090051 | 0.20366605 | DAP2 | VIII | 164978 | 167434 | -1 |
| YHR029C | 89.9416328 | -1.9218579 | 0.57054688 | 13.057017 | 0.00146118 | 0.00894319 | YHI9 | VIII | 167677 | 168561 | -1 |
| YHR030C | 119.948735 | 1.84628453 | 0.81640141 | 10.2533014 | 0.00593641 | 0.02629825 | SLT2 | VIII | 168890 | 170344 | -1 |
| YHR031C | 107.537238 | 1.8706216 | 0.54045995 | 11.5871361 | 0.00304709 | 0.01538156 | RRM3 | VIII | 170799 | 172970 | -1 |
| YHR032W | 55.459718 | -0.5992908 | 0.63737977 | 0.89856543 | 0.63808568 | 0.77431376 | ERC1 | VIII | 173344 | 175089 | 1 |

| | | | | | | | | | | | |
|---|---|---|---|---|---|---|---|---|---|---|---|
| YHR033W | 62.1613368 | 3.73802575 | 0.66729884 | 45.0016384 | 1.69E-10 | 7.47E-09 | NA | VIII | 175548 | 176819 | 1 |
| YHR034C | 40.6566423 | -0.2191205 | 0.61555901 | 0.16527055 | 0.92068688 | 0.95024857 | PIH1 | VIII | 176965 | 177999 | -1 |
| YHR035W | 31.4699039 | -1.2788138 | 0.72149892 | 3.13888074 | 0.20816164 | 0.37050597 | NEL1 | VIII | 178219 | 180111 | 1 |
| YHR036W | 64.0043558 | -0.124623 | 0.5442382 | 0.61947612 | 0.7336391 | 0.83775415 | BRL1 | VIII | 180345 | 181760 | 1 |
| YHR037W | 20.2326832 | 0.75530329 | 0.74747789 | 1.92355739 | 0.38221244 | 0.55547441 | PUT2 | VIII | 181977 | 183704 | 1 |
| YHR038W | 166.515375 | -0.2068738 | 0.86430679 | 9.54753533 | 0.00844849 | 0.03464359 | RRF1 | VIII | 184066 | 184758 | 1 |
| YHR039C | 48.7362825 | 0.62189227 | 0.57027288 | 1.45195961 | 0.48385026 | 0.64624982 | MSC7 | VIII | 184875 | 186809 | -1 |
| YHR039C-A | 175.816476 | -1.9777026 | 0.57910238 | 11.4860289 | 0.00320509 | 0.01608101 | VMA10 | VIII | 187173 | 187679 | -1 |
| YHR040W | 58.4036396 | -1.4062654 | 0.70277094 | 8.66498794 | 0.01313475 | 0.04828584 | BCD1 | VIII | 187924 | 189024 | 1 |
| YHR041C | 277.580421 | 0.74826606 | 0.53130662 | 7.2859555 | 0.02617429 | 0.08039027 | SRB2 | VIII | 189131 | 189864 | -1 |
| YHR042W | 44.3959992 | -0.7849334 | 0.6429094 | 1.64244623 | 0.43989329 | 0.60823052 | NCP1 | VIII | 190543 | 192618 | 1 |
| YHR043C | 197.668394 | -0.9225285 | 0.82084346 | 6.2389066 | 0.04418132 | 0.11942569 | DOG2 | VIII | 192805 | 193545 | -1 |
| YHR044C | 5.47872682 | 2.52829241 | 1.45800181 | 3.30137081 | 0.19191832 | 0.34834078 | DOG1 | VIII | 194068 | 194808 | -1 |
| YHR045W | 51.1711908 | -2.0750445 | 0.76008473 | 11.1882176 | 0.00371971 | 0.01814182 | NA | VIII | 195551 | 197233 | 1 |
| YHR046C | 162.217745 | -0.9983248 | 0.63055408 | 3.25339991 | 0.19657722 | 0.35446344 | INM1 | VIII | 197398 | 198285 | -1 |
| YHR047C | 45.9527251 | 0.44392324 | 0.66850085 | 0.50849857 | 0.77549846 | 0.8634877 | AAP1 | VIII | 198740 | 201310 | -1 |
| YHR048W | 648.139114 | -1.4739513 | 0.47150235 | 10.2049947 | 0.00608154 | 0.02677727 | YHK8 | VIII | 204607 | 206151 | 1 |
| YHR049W | 129.754673 | -1.6415756 | 0.58461259 | 8.57752045 | 0.01372193 | 0.04993712 | FSH1 | VIII | 206462 | 207193 | 1 |
| YHR050W | 65.5978275 | 1.92619517 | 0.57477965 | 15.3857596 | 0.00045606 | 0.00341163 | SMF2 | VIII | 207654 | 209303 | 1 |
| YHR050W-A | 9.73139916 | -0.3651808 | 0.94891178 | 0.50489694 | 0.77689625 | 0.86407117 | NA | VIII | 209475 | 209645 | 1 |
| YHR051W | 45.273619 | -1.4074795 | 0.60692375 | 9.74689304 | 0.00764696 | 0.03206077 | COX6 | VIII | 209705 | 210151 | 1 |
| YHR052W | 23.42874 | -1.6430233 | 0.87949875 | 3.53837657 | 0.17047131 | 0.32067215 | CIC1 | VIII | 210848 | 211978 | 1 |
| YHR054C | 140.921217 | -0.6395672 | 0.57648364 | 3.83727249 | 0.14680703 | 0.28872715 | NA | VIII | 213185 | 214249 | -1 |
| YHR056C | 46.1740423 | -0.979568 | 0.66026617 | 3.03813105 | 0.21891636 | 0.38305402 | RSC30 | VIII | 215183 | 217834 | -1 |
| YHR057C | 103.123444 | 1.2212529 | 0.62343236 | 3.79179011 | 0.15018385 | 0.29297995 | CPR2 | VIII | 218226 | 218843 | -1 |
| YHR058C | 17.4532122 | 0.39390658 | 0.81493615 | 1.60866911 | 0.44738554 | 0.61428226 | MED6 | VIII | 218997 | 219884 | -1 |
| YHR060W | 51.8268212 | -0.9028621 | 0.88595364 | 4.91152974 | 0.08579755 | 0.19673972 | VMA22 | VIII | 220725 | 221270 | 1 |
| YHR061C | 517.483556 | 1.79635622 | 0.51425164 | 17.6621681 | 0.00014612 | 0.00130561 | GIC1 | VIII | 221534 | 222478 | -1 |
| YHR062C | 186.950339 | -0.2416229 | 0.53302969 | 2.73016646 | 0.25535942 | 0.42756826 | RPP1 | VIII | 222877 | 223758 | -1 |
| YHR063C | 45.003073 | -2.5306128 | 0.64545506 | 17.4499943 | 0.00016247 | 0.00142318 | PAN5 | VIII | 224030 | 225169 | -1 |
| YHR064C | 36.4932875 | -1.7469524 | 0.76121722 | 5.24640806 | 0.07256997 | 0.17317521 | SSZ1 | VIII | 225525 | 227141 | -1 |
| YHR065C | 55.8662978 | -2.0023425 | 0.66640343 | 10.9191023 | 0.00425547 | 0.02011359 | RRP3 | VIII | 227532 | 229037 | -1 |
| YHR066W | 124.183948 | 1.80238741 | 0.54007015 | 16.0209989 | 0.00033196 | 0.00265843 | SSF1 | VIII | 229335 | 230696 | 1 |

| | | | | | | | | | | | |
|---|---|---|---|---|---|---|---|---|---|---|---|
| YHR067W | 12.730186 | -0.2136192 | 0.91906869 | 0.48249533 | 0.78564703 | 0.87017291 | HTD2 | VIII | 230970 | 231812 | 1 |
| YHR068W | 50.2707488 | -1.3178505 | 0.58756124 | 5.7016498 | 0.05779662 | 0.14608359 | DYS1 | VIII | 232133 | 233296 | 1 |
| YHR069C | 15.7826428 | -1.6986225 | 0.79276504 | 4.46423507 | 0.10730098 | 0.23035693 | RRP4 | VIII | 233579 | 234658 | -1 |
| YHR070W | 159.695575 | -0.5334335 | 0.58911544 | 0.85630553 | 0.65171185 | 0.78244649 | TRM5 | VIII | 234881 | 236380 | 1 |
| YHR071W | 95.7925513 | 0.33004558 | 0.64128867 | 1.51482406 | 0.4688783 | 0.63149694 | PCL5 | VIII | 237004 | 237693 | 1 |
| YHR072W | 149.744652 | 1.1012055 | 0.52241089 | 4.67894765 | 0.09637834 | 0.21339601 | ERG7 | VIII | 239098 | 241293 | 1 |
| YHR072W-A | 14.1572986 | -1.8410521 | 0.85379256 | 4.71968636 | 0.09443503 | 0.20984715 | NOP10 | VIII | 241664 | 241840 | 1 |
| YHR073W | 36.6355245 | -0.4054412 | 0.69740781 | 0.88556997 | 0.64224529 | 0.77621415 | OSH3 | VIII | 242582 | 245572 | 1 |
| YHR074W | 61.9480824 | 2.2417647 | 0.81797875 | 9.58984488 | 0.00827164 | 0.03401477 | QNS1 | VIII | 246193 | 248337 | 1 |
| YHR075C | 271.643022 | -0.1475224 | 0.76094794 | 6.17732293 | 0.0455629 | 0.12191042 | PPE1 | VIII | 248439 | 249641 | -1 |
| YHR076W | 338.907306 | -0.3341633 | 0.50184386 | 4.24856604 | 0.11951863 | 0.24874654 | PTC7 | VIII | 251101 | 252225 | 1 |
| YHR077C | 159.798725 | 0.37927085 | 0.56932927 | 1.69414055 | 0.42866898 | 0.59835906 | NMD2 | VIII | 252374 | 255756 | -1 |
| YHR078W | 42.0136743 | 2.38595096 | 0.65862438 | 12.6520106 | 0.00178917 | 0.01030774 | NA | VIII | 256360 | 258018 | 1 |
| YHR079C | 399.16565 | -2.0882425 | 0.51437922 | 16.1443729 | 0.0003121 | 0.00252383 | IRE1 | VIII | 258244 | 261591 | -1 |
| YHR079C-A | 48.8939776 | 2.25959494 | 0.6383042 | 20.6839844 | 3.23E-05 | 0.00035703 | SAE3 | VIII | 262192 | 262553 | -1 |
| YHR080C | 160.96475 | 2.16611556 | 0.55499493 | 25.1261764 | 3.50E-06 | 5.25E-05 | LAM4 | VIII | 262801 | 266838 | -1 |
| YHR081W | 21.7973695 | -1.7091097 | 0.73382405 | 5.65101965 | 0.05927843 | 0.1485167 | LRP1 | VIII | 267538 | 268092 | 1 |
| YHR082C | 51.1855884 | 0.37407909 | 0.58210771 | 0.98257172 | 0.61183915 | 0.7550189 | KSP1 | VIII | 268459 | 271548 | -1 |
| YHR083W | 83.5435918 | -2.2270595 | 0.6064176 | 12.4742663 | 0.00195545 | 0.01099231 | SAM35 | VIII | 272627 | 273616 | 1 |
| YHR084W | 69.4974651 | 1.00511273 | 0.73673712 | 7.17616542 | 0.0276513 | 0.08364733 | STE12 | VIII | 274174 | 276240 | 1 |
| YHR085W | 88.957631 | 1.14593717 | 0.57534237 | 9.49152615 | 0.00868843 | 0.03535208 | IPI1 | VIII | 276764 | 277768 | 1 |
| YHR086W | 70.2020459 | 1.96042797 | 0.69068375 | 7.98815743 | 0.01842441 | 0.06216629 | NAM8 | VIII | 278153 | 279724 | 1 |
| YHR086W-A | 44.4806486 | -0.7653271 | 0.74438891 | 1.67231529 | 0.4333725 | 0.60207676 | NA | VIII | 280231 | 280392 | 1 |
| YHR087W | 40.5228695 | 0.03059894 | 0.7070389 | 0.19098196 | 0.90892656 | 0.94414869 | RTC3 | VIII | 280820 | 281155 | 1 |
| YHR088W | 35.1185151 | -1.6672564 | 0.70517831 | 5.38939933 | 0.06756267 | 0.16360847 | RPF1 | VIII | 281495 | 282382 | 1 |
| YHR089C | 157.237438 | -1.3779284 | 0.61662512 | 5.80658544 | 0.05484234 | 0.14011689 | GAR1 | VIII | 282681 | 283298 | -1 |
| YHR090C | 13.0013571 | 0.73898038 | 0.89057056 | 2.67704249 | 0.26223316 | 0.43356082 | YNG2 | VIII | 283777 | 284625 | -1 |
| YHR091C | 207.309857 | 0.03707731 | 0.49334552 | 2.48515092 | 0.28863988 | 0.46255879 | MSR1 | VIII | 284839 | 286770 | -1 |
| YHR092C | 31.7026908 | 0.11492351 | 0.79596993 | 1.92662957 | 0.38162578 | 0.55529381 | HXT4 | VIII | 287081 | 288811 | -1 |
| YHR093W | 142.893984 | -0.4074144 | 0.55044883 | 5.56543226 | 0.06187023 | 0.15328568 | NA | VIII | 289142 | 289690 | 1 |
| YHR094C | 46.0186991 | 0.82889205 | 0.62541633 | 1.84321648 | 0.39787864 | 0.56853834 | HXT1 | VIII | 290913 | 292625 | -1 |
| YHR096C | 1386.792 | 1.70217327 | 0.64443999 | 6.98950247 | 0.0303563 | 0.09004251 | HXT5 | VIII | 294669 | 296447 | -1 |
| YHR097C | 208.99292 | 2.49959349 | 0.50795008 | 29.3797669 | 4.17E-07 | 8.21E-06 | NA | VIII | 297385 | 298609 | -1 |

| | | | | | | | | | | | |
|---|---|---|---|---|---|---|---|---|---|---|---|
| YHR098C | 77.118544 | 2.81604014 | 0.6024199 | 20.7356187 | 3.14E-05 | 0.00034994 | SFB3 | VIII | 299145 | 301934 | -1 |
| YHR099W | 72.6530054 | 0.23016454 | 0.5520393 | 0.21541138 | 0.89789182 | 0.93705725 | TRA1 | VIII | 302761 | 313995 | 1 |
| YHR100C | 513.667901 | 0.04629299 | 0.57763332 | 0.79591906 | 0.67168921 | 0.79661625 | GEP4 | VIII | 314116 | 314673 | -1 |
| YHR101C | 80.1368126 | 0.29861307 | 0.52851311 | 3.69953995 | 0.15727334 | 0.30313337 | BIG1 | VIII | 314874 | 315968 | -1 |
| YHR102W | 428.90623 | -0.1185339 | 0.50015933 | 0.07592391 | 0.96274957 | 0.97258112 | KIC1 | VIII | 316572 | 319814 | 1 |
| YHR103W | 16.1815043 | 0.27111046 | 0.8112158 | 0.99072727 | 0.60934929 | 0.75306988 | SBE22 | VIII | 320414 | 322972 | 1 |
| YHR104W | 22.3748001 | 1.35204524 | 0.75999169 | 3.69267236 | 0.15781431 | 0.30407483 | GRE3 | VIII | 323409 | 324392 | 1 |
| YHR105W | 14.3254284 | 1.26896909 | 0.80754305 | 4.81156735 | 0.09019478 | 0.20366605 | YPT35 | VIII | 324766 | 325410 | 1 |
| YHR106W | 35.1150828 | 0.91527762 | 0.69056852 | 1.85987572 | 0.39457823 | 0.56638907 | TRR2 | VIII | 325598 | 326626 | 1 |
| YHR107C | 31.8098845 | -0.4090498 | 0.64273463 | 1.14492049 | 0.56413582 | 0.71504956 | CDC12 | VIII | 326813 | 328036 | -1 |
| YHR108W | 41.6577891 | -0.3439053 | 0.6069305 | 0.33268049 | 0.84675808 | 0.90976606 | GGA2 | VIII | 328303 | 330060 | 1 |
| YHR109W | 20.840772 | 0.34434487 | 0.83599183 | 1.36165653 | 0.50619755 | 0.66417229 | CTM1 | VIII | 330310 | 332067 | 1 |
| YHR110W | 10.0573769 | -0.052605 | 0.99249267 | 0.09233673 | 0.95488118 | 0.96809001 | ERP5 | VIII | 332282 | 332920 | 1 |
| YHR111W | 421.722519 | -0.0444979 | 0.46291215 | 0.22727835 | 0.89257996 | 0.93387025 | UBA4 | VIII | 333072 | 334394 | 1 |
| YHR112C | 493.886505 | -3.2426145 | 0.51654773 | 35.757229 | 1.72E-08 | 4.86E-07 | NA | VIII | 334527 | 335663 | -1 |
| YHR113W | 100.052273 | -0.7758785 | 0.57133076 | 3.6800863 | 0.15881057 | 0.30548612 | APE4 | VIII | 336337 | 337809 | 1 |
| YHR114W | 130.971018 | -2.0928627 | 0.52897256 | 15.8269778 | 0.00036578 | 0.00285809 | BZZ1 | VIII | 338083 | 339984 | 1 |
| YHR115C | 371.662103 | 2.1512459 | 0.66418013 | 9.91899309 | 0.00701646 | 0.0300483 | DMA1 | VIII | 340109 | 341359 | -1 |
| YHR116W | 21.7106812 | -1.8522204 | 0.78123111 | 9.02876601 | 0.01095036 | 0.04215597 | COX23 | VIII | 341665 | 342120 | 1 |
| YHR117W | 207.970067 | -1.8947994 | 0.54365955 | 11.4079538 | 0.00333269 | 0.01653492 | TOM71 | VIII | 342349 | 344268 | 1 |
| YHR118C | 120.598348 | -1.1659695 | 0.63168341 | 7.0720478 | 0.02912892 | 0.08689151 | ORC6 | VIII | 344321 | 345628 | -1 |
| YHR119W | 156.180524 | -1.2815846 | 0.49783975 | 7.50560235 | 0.02345196 | 0.07454061 | SET1 | VIII | 346043 | 349285 | 1 |
| YHR120W | 23.3135607 | -0.2618795 | 0.73401927 | 0.22255234 | 0.89469163 | 0.9350658 | MSH1 | VIII | 349574 | 352453 | 1 |
| YHR121W | 29.2357214 | -1.3724077 | 0.7124893 | 4.04018594 | 0.13264313 | 0.26853278 | LSM12 | VIII | 352756 | 353319 | 1 |
| YHR122W | 26.6266893 | -0.4562537 | 0.68577537 | 0.44263869 | 0.8014607 | 0.88031084 | CIA2 | VIII | 353625 | 354320 | 1 |
| YHR123W | 35.5878082 | 1.7203632 | 0.65618961 | 10.3307812 | 0.00571083 | 0.025494 | EPT1 | VIII | 354815 | 356081 | 1 |
| YHR124W | 1493.9623 | -0.1526129 | 0.4664107 | 0.41624796 | 0.81210634 | 0.88731158 | NDT80 | VIII | 356561 | 358444 | 1 |
| YHR125W | 9.13429095 | 1.01076391 | 1.04830089 | 1.72770977 | 0.42153398 | 0.59239848 | NA | VIII | 358858 | 359163 | 1 |
| YHR126C | 6.48106968 | 1.61921513 | 1.12263211 | 2.22956852 | 0.32798603 | 0.50041094 | ANS1 | VIII | 359702 | 360181 | -1 |
| YHR127W | 63.354218 | -3.6979242 | 0.6184724 | 43.2213978 | 4.12E-10 | 1.64E-08 | NA | VIII | 360913 | 361644 | 1 |
| YHR128W | 107.628796 | -0.3337312 | 0.52414127 | 0.40874108 | 0.81516027 | 0.88884707 | FUR1 | VIII | 362115 | 362765 | 1 |
| YHR129C | 44.9403704 | -1.4863813 | 0.67542869 | 5.04858957 | 0.08011479 | 0.18666585 | ARP1 | VIII | 362999 | 364153 | -1 |
| YHR130C | 32.001307 | -0.72467 | 0.80604495 | 2.40754248 | 0.30006048 | 0.47326346 | NA | VIII | 364965 | 365300 | -1 |

| | | | | | | | | | | | |
|---|---|---|---|---|---|---|---|---|---|---|---|
| YHR131C | 48.9503403 | 0.18089431 | 0.62701854 | 0.11123634 | 0.94590024 | 0.96354018 | NA | VIII | 365340 | 367892 | -1 |
| YHR132C | 67.2291349 | 0.67288362 | 0.69342569 | 1.34151283 | 0.51132166 | 0.66829626 | ECM14 | VIII | 368502 | 369794 | -1 |
| YHR132W-A | 40.1234912 | 0.61260359 | 0.65006165 | 0.92617982 | 0.62933605 | 0.76680754 | IGO2 | VIII | 370054 | 370449 | 1 |
| YHR133C | 53.481965 | 0.61600188 | 0.66175596 | 1.04806012 | 0.59212942 | 0.73920425 | NSG1 | VIII | 370722 | 371597 | -1 |
| YHR134W | 1803.52476 | 3.56310028 | 0.47712734 | 46.2125226 | 9.23E-11 | 4.42E-09 | WSS1 | VIII | 371749 | 372558 | 1 |
| YHR135C | 1225.45617 | 1.10512031 | 0.45919761 | 6.56128726 | 0.03760405 | 0.1062085 | YCK1 | VIII | 372694 | 374310 | -1 |
| YHR136C | 6.11830014 | 0.91845689 | 1.12126316 | 1.7048867 | 0.42637188 | 0.59587092 | SPL2 | VIII | 374654 | 375100 | -1 |
| YHR137W | 248.108564 | 2.9141192 | 0.93138366 | 15.1956828 | 0.00050153 | 0.00368045 | ARO9 | VIII | 375709 | 377250 | 1 |
| YHR138C | 206.260095 | 0.97354098 | 0.82011205 | 5.47464679 | 0.06474341 | 0.15863916 | NA | VIII | 377355 | 377699 | -1 |
| YHR139C | 175.346183 | -1.3629501 | 0.54203917 | 8.39998243 | 0.01499571 | 0.05349671 | SPS100 | VIII | 378219 | 379199 | -1 |
| YHR139C-A | 105.269644 | 0.42673719 | 0.56129218 | 2.39477846 | 0.30198159 | 0.47448396 | NA | VIII | 380108 | 380419 | -1 |
| YHR140W | 36.947971 | 0.44671399 | 0.64182265 | 0.52159893 | 0.7704354 | 0.86016602 | NA | VIII | 380572 | 381291 | 1 |
| YHR141C | 113.083862 | -3.4256875 | 0.73701981 | 18.9800347 | 7.56E-05 | 0.00073818 | RPL42A | VIII | 381990 | 382751 | -1 |
| YHR142W | 510.825248 | -2.8172103 | 0.53182444 | 24.8806211 | 3.96E-06 | 5.84E-05 | CHS7 | VIII | 383538 | 384488 | 1 |
| YHR143W | 381.204132 | 0.98355024 | 0.49297812 | 4.42149471 | 0.10961869 | 0.23403106 | DSE2 | VIII | 385510 | 386487 | 1 |
| YHR143W-A | 15.3154485 | -0.2453761 | 0.80631125 | 0.1056259 | 0.94855742 | 0.96471939 | RPC10 | VIII | 387233 | 387445 | 1 |
| YHR144C | 24.7359693 | 1.53527989 | 0.70437773 | 4.86447683 | 0.08783999 | 0.20034795 | DCD1 | VIII | 387788 | 388726 | -1 |
| YHR146W | 53.3880307 | -0.5188675 | 0.62351732 | 0.71141646 | 0.70067703 | 0.81801018 | CRP1 | VIII | 390300 | 391697 | 1 |
| YHR147C | 182.239972 | 1.61711161 | 0.52088589 | 13.9779109 | 0.00092201 | 0.00615027 | MRPL6 | VIII | 392639 | 393283 | -1 |
| YHR148W | 7.76600939 | -2.7081164 | 1.11308516 | 6.34560741 | 0.041886 | 0.11493835 | IMP3 | VIII | 393534 | 394085 | 1 |
| YHR149C | 52.5911302 | 0.04921122 | 0.6236858 | 0.20328776 | 0.9033512 | 0.93919975 | SKG6 | VIII | 394455 | 396659 | -1 |
| YHR150W | 18.8336382 | 1.43932825 | 0.84829779 | 11.154229 | 0.00378347 | 0.01836234 | PEX28 | VIII | 397251 | 398990 | 1 |
| YHR151C | 12.6354726 | 0.89936941 | 0.84586116 | 3.8213306 | 0.1479819 | 0.2902937 | MTC6 | VIII | 399268 | 400848 | -1 |
| YHR152W | 75.0417121 | -2.9512543 | 0.66530959 | 18.2584721 | 0.00010845 | 0.00100306 | SPO12 | VIII | 401434 | 401955 | 1 |
| YHR153C | 8.8817812 | -2.6730215 | 1.01556446 | 6.68394776 | 0.03536708 | 0.10132379 | SPO16 | VIII | 402086 | 402682 | -1 |
| YHR154W | 118.307027 | 0.24779541 | 0.51176905 | 2.10338458 | 0.34934605 | 0.52293528 | RTT107 | VIII | 402966 | 406178 | 1 |
| YHR155W | 33.364221 | -0.4136357 | 0.62729256 | 0.68060863 | 0.71155375 | 0.82438173 | LAM1 | VIII | 407103 | 410789 | 1 |
| YHR156C | 271.65265 | -1.718315 | 0.54245041 | 9.8950464 | 0.00710097 | 0.03034291 | LIN1 | VIII | 411384 | 412406 | -1 |
| YHR158C | 27.2764948 | -0.5115266 | 0.73475291 | 0.74758773 | 0.68811874 | 0.81029236 | KEL1 | VIII | 413685 | 417179 | -1 |
| YHR159W | 26.8608684 | 1.23814275 | 0.71403174 | 2.98374417 | 0.22495113 | 0.3904964 | TDA11 | VIII | 417549 | 419063 | 1 |
| YHR160C | 539.703211 | -0.9483033 | 0.48435747 | 4.3095163 | 0.11593123 | 0.24332354 | PEX18 | VIII | 419221 | 420072 | -1 |
| YHR161C | 73.3051109 | 0.84501698 | 0.57687707 | 2.86132498 | 0.23915044 | 0.40763705 | YAP1801 | VIII | 420373 | 422286 | -1 |
| YHR162W | 69.7417583 | -1.1719887 | 0.55584259 | 7.21043517 | 0.02718153 | 0.08256862 | MPC2 | VIII | 423072 | 423461 | 1 |

| | | | | | | | | | | | |
|---|---|---|---|---|---|---|---|---|---|---|---|
| YHR163W | 53.0931978 | -0.4053998 | 0.59408762 | 3.049795 | 0.21764336 | 0.38155469 | SOL3 | VIII | 423724 | 424473 | 1 |
| YHR164C | 172.361879 | -0.246448 | 0.73627903 | 8.30136888 | 0.01575363 | 0.05576982 | DNA2 | VIII | 424610 | 429178 | -1 |
| YHR165C | 90.7947384 | 1.41556131 | 0.55154284 | 7.36069567 | 0.0252142 | 0.078237 | PRP8 | VIII | 429707 | 436948 | -1 |
| YHR166C | 16.9837717 | -1.1406452 | 0.82860192 | 1.97373835 | 0.37274186 | 0.546097 | CDC23 | VIII | 437170 | 439050 | -1 |
| YHR167W | 13.9078478 | -0.0019735 | 0.89201848 | 0.03124002 | 0.98450135 | 0.98751104 | THP2 | VIII | 439342 | 440127 | 1 |
| YHR168W | 22.9099055 | -2.4346361 | 0.7162136 | 12.0371563 | 0.00243313 | 0.01296026 | MTG2 | VIII | 440377 | 441933 | 1 |
| YHR169W | 39.9564375 | -0.3745834 | 0.75864231 | 0.32574404 | 0.84969992 | 0.91079805 | DBP8 | VIII | 442181 | 443476 | 1 |
| YHR170W | 34.377728 | -1.1625846 | 0.70537964 | 3.42110355 | 0.18076602 | 0.33535254 | NMD3 | VIII | 443828 | 445384 | 1 |
| YHR171W | 25.5372705 | -0.5989805 | 0.68145383 | 0.8982898 | 0.63817362 | 0.77431376 | ATG7 | VIII | 445713 | 447605 | 1 |
| YHR172W | 54.5244315 | 1.61259492 | 0.67601009 | 5.71589629 | 0.05738639 | 0.14528517 | SPC97 | VIII | 448335 | 450806 | 1 |
| YHR173C | 434.301909 | -1.4574384 | 0.46716249 | 10.1236357 | 0.00633403 | 0.02761601 | NA | VIII | 450815 | 451153 | -1 |
| YHR174W | 678.1549 | -1.5700255 | 0.63650177 | 6.25252025 | 0.0438816 | 0.11883745 | ENO2 | VIII | 451327 | 452640 | 1 |
| YHR175W | 27.6180233 | -0.0764632 | 0.70165122 | 0.9563941 | 0.61990003 | 0.76058936 | CTR2 | VIII | 452872 | 453441 | 1 |
| YHR176W | 31.0946548 | 0.54550416 | 0.68910228 | 1.60946785 | 0.4472069 | 0.61428226 | FMO1 | VIII | 454229 | 455527 | 1 |
| YHR177W | 1104.10105 | 1.72266323 | 0.52760118 | 10.4104042 | 0.00548794 | 0.02467017 | NA | VIII | 456591 | 457952 | 1 |
| YHR178W | 85.5455634 | -3.1065996 | 0.56756522 | 28.5393871 | 6.35E-07 | 1.20E-05 | STB5 | VIII | 459299 | 461530 | 1 |
| YHR179W | 74.2830905 | 1.07064944 | 0.67708971 | 4.54347725 | 0.10313271 | 0.22389892 | OYE2 | VIII | 462502 | 463704 | 1 |
| YHR180W | 730.352155 | 2.53600934 | 0.4834632 | 30.9721806 | 1.88E-07 | 4.16E-06 | NA | VIII | 465178 | 465669 | 1 |
| YHR181W | 20.1224725 | -0.3994571 | 0.70449513 | 2.07511295 | 0.35431941 | 0.52819501 | SVP26 | VIII | 467228 | 467914 | 1 |
| YHR182W | 377.380662 | 0.10009645 | 0.49154464 | 3.01103575 | 0.22190235 | 0.38675936 | NA | VIII | 468219 | 470576 | 1 |
| YHR183W | 118.404268 | -0.6514414 | 0.78826987 | 3.88458107 | 0.14337517 | 0.28351852 | GND1 | VIII | 470960 | 472429 | 1 |
| YHR186C | 61.8092633 | 2.35876389 | 0.65950109 | 14.0614319 | 0.0008843 | 0.00595552 | KOG1 | VIII | 475999 | 480672 | -1 |
| YHR187W | 170.953949 | -0.6342138 | 0.82582943 | 12.4000568 | 0.00202937 | 0.01126565 | IKI1 | VIII | 480990 | 481919 | 1 |
| YHR188C | 119.399674 | 0.12132126 | 0.72489343 | 2.55069991 | 0.2793332 | 0.45351049 | GPI16 | VIII | 482005 | 483837 | -1 |
| YHR189W | 202.737102 | -0.3230262 | 0.6153729 | 1.55002349 | 0.46069837 | 0.62469404 | PTH1 | VIII | 484028 | 484600 | 1 |
| YHR190W | 161.280792 | -0.6473539 | 0.59583683 | 5.5636494 | 0.06192541 | 0.15335677 | ERG9 | VIII | 484845 | 486179 | 1 |
| YHR191C | 1594.53219 | -0.0704999 | 0.63463053 | 9.63795636 | 0.00807503 | 0.03341947 | CTF8 | VIII | 486230 | 486631 | -1 |
| YHR192W | 1091.47309 | -0.7210336 | 1.60975986 | 5.24635764 | 0.0725718 | 0.17317521 | LNP1 | VIII | 486826 | 487662 | 1 |
| YHR193C | 530.214974 | -0.8545601 | 0.50393475 | 6.63062021 | 0.03632278 | 0.10344757 | EGD2 | VIII | 487712 | 488236 | -1 |
| YHR194W | 63.4478203 | 0.00702536 | 0.59699831 | 4.53058794 | 0.10379952 | 0.22509714 | MDM31 | VIII | 488657 | 490396 | 1 |
| YHR195W | 76.5297286 | -0.8213807 | 0.54549878 | 3.14527917 | 0.20749675 | 0.36958676 | NVJ1 | VIII | 490747 | 491712 | 1 |
| YHR196W | 27.5811457 | -1.1834865 | 0.91722635 | 2.68548535 | 0.26112849 | 0.43235173 | UTP9 | VIII | 491931 | 493658 | 1 |
| YHR197W | 73.9548284 | -0.9967432 | 0.70052898 | 2.7578336 | 0.25185121 | 0.42340839 | RIX1 | VIII | 493896 | 496187 | 1 |

| | | | | | | | | | | | |
|---|---|---|---|---|---|---|---|---|---|---|---|
| YHR198C | 129.36181 | -2.6607986 | 0.52679471 | 29.0096711 | 5.02E-07 | 9.75E-06 | AIM18 | VIII | 496315 | 497280 | -1 |
| YHR199C | 32.4552259 | -0.050617 | 0.74320087 | 0.11876813 | 0.94234478 | 0.96176085 | AIM46 | VIII | 497490 | 498422 | -1 |
| YHR199C-A | 21.8386558 | -1.0378011 | 0.77002512 | 3.61694568 | 0.16390425 | 0.31248127 | NBL1 | VIII | 498571 | 498859 | -1 |
| YHR200W | 143.280763 | -0.3958281 | 0.71421968 | 9.42409569 | 0.00898636 | 0.0363345 | RPN10 | VIII | 499079 | 499885 | 1 |
| YHR201C | 159.190447 | -1.2347788 | 0.62676688 | 7.42667345 | 0.02439598 | 0.07671608 | PPX1 | VIII | 499950 | 501143 | -1 |
| YHR202W | 42.0338164 | -1.9778204 | 0.667821 | 9.58999518 | 0.00827102 | 0.03401477 | NA | VIII | 502388 | 504196 | 1 |
| YHR203C | 75.3881069 | -2.8881401 | 0.74109679 | 13.8980416 | 0.00095957 | 0.00635691 | RPS4A | VIII | 504476 | 505530 | -1 |
| YHR204W | 171.634151 | -0.96166 | 0.51013796 | 9.92617465 | 0.00699131 | 0.02997984 | MNL1 | VIII | 506319 | 508709 | 1 |
| YHR205W | 168.338849 | 1.64372513 | 0.56968996 | 8.26593772 | 0.0160352 | 0.05640572 | SCH9 | VIII | 509363 | 511837 | 1 |
| YHR206W | 31.2941275 | -0.2122216 | 0.6534566 | 0.28990772 | 0.86506221 | 0.92054956 | SKN7 | VIII | 512732 | 514600 | 1 |
| YHR207C | 49.4771919 | -0.1483934 | 0.59285583 | 0.72285294 | 0.69668182 | 0.81659461 | SET5 | VIII | 514905 | 516485 | -1 |
| YHR208W | 39.0195505 | 1.25374169 | 0.61585823 | 4.23387809 | 0.1203996 | 0.24977202 | BAT1 | VIII | 517532 | 518713 | 1 |
| YHR209W | 330.747873 | -1.0847666 | 0.47974758 | 5.0000732 | 0.08208199 | 0.18979822 | CRG1 | VIII | 519437 | 520312 | 1 |
| YHR210C | 11.024383 | 1.21515702 | 0.91015495 | 2.13406732 | 0.34402751 | 0.51765054 | NA | VIII | 520712 | 521737 | -1 |
| YHR211W | 154.662749 | 0.53596342 | 0.4952895 | 3.21752406 | 0.20013522 | 0.35908985 | FLO5 | VIII | 525392 | 528619 | 1 |
| YHR213W | 17.0950159 | -0.4411597 | 0.78721758 | 0.97052035 | 0.61553703 | 0.75780553 | NA | VIII | 539151 | 539747 | 1 |
| YHR213W-B | 151.170909 | 2.22211206 | 0.52867772 | 16.9576885 | 0.00020782 | 0.00176433 | NA | VIII | 540800 | 541099 | 1 |
| YHR214C-B | 8.68495834 | -1.1673346 | 1.01997214 | 2.3218241 | 0.3132004 | 0.48516903 | NA | VIII | 543964 | 549346 | -1 |
| YHR214C-E | 13.4042596 | -0.6228131 | 0.84525636 | 0.65206199 | 0.72178282 | 0.83133529 | NA | VIII | 551200 | 551499 | -1 |
| YHR214W | 172.718877 | 0.80926606 | 0.55393896 | 4.4084099 | 0.11033822 | 0.23522866 | NA | VIII | 541651 | 542262 | 1 |
| YHR216W | 31.8040126 | 1.15054596 | 0.7354704 | 3.53611718 | 0.170664 | 0.32088433 | IMD2 | VIII | 554396 | 555967 | 1 |
| YIL001W | 9.79863105 | 0.40049574 | 1.09414673 | 0.32851834 | 0.84852208 | 0.90980423 | NA | IX | 353940 | 355481 | 1 |
| YIL002C | 433.31064 | 0.68005723 | 0.71770651 | 8.05950837 | 0.0177787 | 0.06041002 | INP51 | IX | 350591 | 353431 | -1 |
| YIL002W-A | 86.236974 | 0.26069847 | 0.60875854 | 3.10045081 | 0.21220014 | 0.37592901 | NA | IX | 350301 | 350510 | 1 |
| YIL003W | 21.9248764 | 1.6387039 | 0.74770539 | 5.01877938 | 0.08131785 | 0.18855842 | CFD1 | IX | 349122 | 350003 | 1 |
| YIL004C | 398.232624 | 0.28878233 | 0.78798962 | 7.41543279 | 0.02453348 | 0.07690789 | BET1 | IX | 347946 | 348505 | -1 |
| YIL005W | 258.753057 | 0.07708956 | 0.63834562 | 1.6238387 | 0.44400505 | 0.61165577 | EPS1 | IX | 345692 | 347797 | 1 |
| YIL006W | 17.8462814 | 1.06909401 | 0.77581195 | 7.6572039 | 0.02173999 | 0.07043902 | YIA6 | IX | 344062 | 345183 | 1 |
| YIL007C | 13.924541 | 0.7032839 | 0.84605808 | 0.79186424 | 0.67305238 | 0.79774275 | NAS2 | IX | 342994 | 343656 | -1 |
| YIL008W | 106.066739 | 2.39968671 | 0.66181276 | 12.1858249 | 0.00225882 | 0.01226883 | URM1 | IX | 342536 | 342835 | 1 |
| YIL009C-A | 434.988398 | 1.34615241 | 0.60504966 | 14.5726399 | 0.00068484 | 0.00479473 | EST3 | IX | 335666 | 336212 | -1 |
| YIL009W | 20.8244296 | -0.1253524 | 0.79487173 | 0.6637434 | 0.71757938 | 0.8289674 | FAA3 | IX | 339344 | 341428 | 1 |
| YIL010W | 41.6564036 | -0.1578436 | 0.73470941 | 1.69489519 | 0.42850726 | 0.59827756 | DOT5 | IX | 334882 | 335529 | 1 |

| | | | | | | | | | | | |
|---|---|---|---|---|---|---|---|---|---|---|---|
| YIL011W | 109.11944 | 1.28096673 | 0.66937168 | 5.25178402 | 0.07237517 | 0.17294768 | TIR3 | IX | 333727 | 334536 | 1 |
| YIL012W | 11.9370033 | 1.01893741 | 0.95698853 | 2.9706482 | 0.22642894 | 0.39252203 | NA | IX | 333011 | 333352 | 1 |
| YIL013C | 1451.67203 | -0.7880779 | 0.63452821 | 2.0527081 | 0.35831097 | 0.53140894 | PDR11 | IX | 328207 | 332442 | -1 |
| YIL014C-A | 8.92910469 | 0.75484385 | 1.00880877 | 0.70114075 | 0.70428627 | 0.82015637 | NA | IX | 325212 | 325526 | -1 |
| YIL014W | 24.2396863 | 0.74983776 | 0.696592 | 1.38095202 | 0.50133737 | 0.66037092 | MNT3 | IX | 326103 | 327995 | 1 |
| YIL015W | 245.773191 | 1.1496704 | 0.53316916 | 7.36568106 | 0.02515143 | 0.07816789 | BAR1 | IX | 322342 | 324105 | 1 |
| YIL016W | 71.1651908 | 0.11490633 | 0.62082442 | 1.5008688 | 0.4721614 | 0.63393528 | SNL1 | IX | 321454 | 321933 | 1 |
| YIL017C | 59.2934768 | -0.6393193 | 0.56881056 | 1.50804469 | 0.47047035 | 0.63290505 | VID28 | IX | 318200 | 320965 | -1 |
| YIL018W | 142.084004 | -2.9963905 | 0.7989682 | 12.5000757 | 0.00193038 | 0.01089367 | RPL2A | IX | 316768 | 317932 | 1 |
| YIL019W | 34.3386162 | -4.4146105 | 0.92115131 | 23.5347838 | 7.75E-06 | 0.00010489 | FAF1 | IX | 315093 | 316133 | 1 |
| YIL020C | 153.413573 | -2.5106061 | 0.68411219 | 12.7816775 | 0.00167685 | 0.00980703 | HIS6 | IX | 314035 | 314820 | -1 |
| YIL021W | 20.9619285 | -1.9216697 | 0.77187237 | 6.42528192 | 0.04025017 | 0.11198871 | RPB3 | IX | 312905 | 313861 | 1 |
| YIL022W | 52.2732232 | -0.8567692 | 0.58262111 | 2.17711643 | 0.33670159 | 0.50994042 | TIM44 | IX | 311165 | 312460 | 1 |
| YIL023C | 144.538004 | 2.51513838 | 0.51250324 | 22.3764965 | 1.38E-05 | 0.00017415 | YKE4 | IX | 309386 | 310426 | -1 |
| YIL024C | 7.43760947 | 0.97811532 | 1.22832736 | 4.15260639 | 0.12539291 | 0.25809632 | NA | IX | 308602 | 309171 | -1 |
| YIL026C | 282.919182 | 0.78390116 | 0.49743804 | 2.7169607 | 0.25705111 | 0.42953129 | IRR1 | IX | 304477 | 307929 | -1 |
| YIL030C | 35.906532 | -0.8838362 | 0.63398077 | 2.58189031 | 0.27501073 | 0.44841232 | SSM4 | IX | 296050 | 300009 | -1 |
| YIL031W | 149.548759 | 2.17012739 | 0.51051088 | 17.296613 | 0.00017542 | 0.00151597 | ULP2 | IX | 292633 | 295737 | 1 |
| YIL033C | 231.459138 | -2.0482009 | 0.52986756 | 15.9438531 | 0.00034501 | 0.00274023 | BCY1 | IX | 290419 | 291669 | -1 |
| YIL034C | 519.175859 | -2.1351274 | 0.47954892 | 22.5461931 | 1.27E-05 | 0.0001621 | CAP2 | IX | 289226 | 290089 | -1 |
| YIL035C | 24.5789834 | -1.2691251 | 0.79788885 | 3.12458943 | 0.20965442 | 0.37244958 | CKA1 | IX | 287790 | 288908 | -1 |
| YIL036W | 67.7484089 | 1.3673087 | 0.55191632 | 10.6786813 | 0.00479903 | 0.02212703 | CST6 | IX | 285666 | 287429 | 1 |
| YIL037C | 668.880673 | -0.3132825 | 0.51718238 | 0.59642313 | 0.74214431 | 0.84243165 | PRM2 | IX | 283029 | 284999 | -1 |
| YIL038C | 27.4108496 | -0.0137098 | 0.71304532 | 0.67961284 | 0.71190812 | 0.8243896 | NOT3 | IX | 280142 | 282652 | -1 |
| YIL039W | 17.373672 | -0.5040392 | 0.97518196 | 0.30274249 | 0.85952855 | 0.91706593 | TED1 | IX | 278426 | 279847 | 1 |
| YIL040W | 130.185184 | 1.04197516 | 0.52853694 | 4.95215402 | 0.08407239 | 0.19331976 | APQ12 | IX | 277723 | 278139 | 1 |
| YIL041W | 75.7198193 | -0.0231855 | 0.5752997 | 0.97660885 | 0.61366603 | 0.75646717 | GVP36 | IX | 276525 | 277505 | 1 |
| YIL042C | 7.5799814 | 1.20306412 | 1.17373283 | 1.69982488 | 0.42745236 | 0.59709268 | PKP1 | IX | 275108 | 276292 | -1 |
| YIL043C | 45.9601708 | -1.4530535 | 0.70525495 | 4.63113393 | 0.0987102 | 0.21722998 | CBR1 | IX | 274072 | 274926 | -1 |
| YIL044C | 21.1624471 | -1.0758255 | 0.83100875 | 2.69283232 | 0.260171 | 0.43177286 | AGE2 | IX | 272950 | 273846 | -1 |
| YIL045W | 142.626499 | 1.15530905 | 0.77993095 | 8.24925656 | 0.0161695 | 0.05670589 | PIG2 | IX | 271161 | 272777 | 1 |
| YIL046W | 40.9461182 | 0.63208477 | 0.60652247 | 1.74300542 | 0.41832246 | 0.589459 | MET30 | IX | 268651 | 270573 | 1 |
| YIL046W-A | 19.6365052 | 1.68516019 | 1.23267409 | 2.70983736 | 0.25796827 | 0.42992972 | NA | IX | 268309 | 268473 | 1 |

| | | | | | | | | | | | |
|---|---|---|---|---|---|---|---|---|---|---|---|
| YIL047C | 40.7818032 | -0.2533671 | 0.6380203 | 2.37373713 | 0.30517541 | 0.47807485 | SYG1 | IX | 265115 | 267823 | -1 |
| YIL048W | 85.473896 | -0.960644 | 0.55202577 | 4.88293387 | 0.08703309 | 0.19878563 | NEO1 | IX | 261437 | 264892 | 1 |
| YIL049W | 86.7815832 | 0.71141841 | 0.55781766 | 1.95397334 | 0.37644374 | 0.54998972 | DFG10 | IX | 260158 | 260919 | 1 |
| YIL050W | 144.39429 | 0.9507844 | 0.5510184 | 3.61938917 | 0.16370413 | 0.3123054 | PCL7 | IX | 258913 | 259770 | 1 |
| YIL051C | 228.648128 | 0.33550676 | 0.50792481 | 0.83994247 | 0.65706572 | 0.78549013 | MMF1 | IX | 257843 | 258280 | -1 |
| YIL052C | 46.2448755 | -2.5323392 | 0.76818905 | 11.2269567 | 0.00364836 | 0.01787139 | RPL34B | IX | 256226 | 257063 | -1 |
| YIL053W | 67.5568594 | -2.1035188 | 0.60564775 | 12.5258667 | 0.00190565 | 0.0107751 | GPP1 | IX | 255115 | 255867 | 1 |
| YIL054W | 52.6158577 | 0.75895208 | 0.7117358 | 1.96527426 | 0.37432266 | 0.54758166 | NA | IX | 254543 | 254860 | 1 |
| YIL055C | 47.967356 | 0.83715322 | 0.75772136 | 6.36606337 | 0.04145977 | 0.11428804 | NA | IX | 252042 | 253925 | -1 |
| YIL056W | 79.5819769 | 1.69672278 | 0.83977677 | 6.77402217 | 0.03380958 | 0.09812404 | VHR1 | IX | 249991 | 251913 | 1 |
| YIL060W | 1319.77337 | 1.55872735 | 0.4878589 | 12.144879 | 0.00230554 | 0.01240622 | NA | IX | 246392 | 246826 | 1 |
| YIL061C | 13.1850525 | 0.23521536 | 0.89745521 | 1.28021519 | 0.52723569 | 0.68184633 | SNP1 | IX | 244657 | 245559 | -1 |
| YIL062C | 216.89699 | 4.25505202 | 0.75776677 | 27.3701352 | 1.14E-06 | 1.98E-05 | ARC15 | IX | 243998 | 244462 | -1 |
| YIL063C | 1340.64674 | 3.27173149 | 0.48724873 | 42.2523172 | 6.68E-10 | 2.50E-08 | YRB2 | IX | 242761 | 243744 | -1 |
| YIL064W | 192.655507 | 1.17334118 | 0.52244496 | 6.3365012 | 0.04207714 | 0.11535353 | EFM4 | IX | 241943 | 242716 | 1 |
| YIL065C | 6.70773065 | 0.8053394 | 1.14194704 | 1.44378363 | 0.48583228 | 0.64773825 | FIS1 | IX | 241308 | 241775 | -1 |
| YIL066C | 30.1138084 | 1.7656377 | 0.67098177 | 7.48689938 | 0.0236723 | 0.07502059 | RNR3 | IX | 238099 | 240708 | -1 |
| YIL067C | 93.6303846 | 3.04310931 | 0.61144533 | 30.6728748 | 2.19E-07 | 4.70E-06 | NA | IX | 235724 | 237760 | -1 |
| YIL068C | 11.4114596 | -0.17339 | 0.9354627 | 0.33576862 | 0.84545164 | 0.90923806 | SEC6 | IX | 233057 | 235474 | -1 |
| YIL069C | 47.4509569 | -3.2428672 | 0.97770401 | 10.1276335 | 0.00632139 | 0.02758163 | RPS24A | IX | 231553 | 232369 | -1 |
| YIL070C | 9.06439748 | -0.9212739 | 0.93926652 | 1.10112539 | 0.57662526 | 0.72593834 | MAM33 | IX | 230272 | 231072 | -1 |
| YIL071C | 151.270349 | 1.35238066 | 0.53523266 | 6.35872972 | 0.04161208 | 0.11445792 | PCI8 | IX | 228660 | 229994 | -1 |
| YIL072W | 683.280405 | -0.393816 | 0.51984075 | 0.91267424 | 0.6336002 | 0.77037908 | HOP1 | IX | 226602 | 228419 | 1 |
| YIL073C | 1177.557 | 0.46842796 | 0.46304536 | 1.3807817 | 0.50138007 | 0.66037092 | SPO22 | IX | 222937 | 225954 | -1 |
| YIL074C | 83.3688747 | 0.94585405 | 0.66316401 | 3.36522895 | 0.18588734 | 0.34169876 | SER33 | IX | 221081 | 222490 | -1 |
| YIL075C | 1207.01814 | 0.88016095 | 0.73025899 | 6.71901579 | 0.03475236 | 0.10004686 | RPN2 | IX | 217863 | 220700 | -1 |
| YIL076W | 25.4445262 | 1.35547042 | 0.95458938 | 4.80809005 | 0.09035174 | 0.20387239 | SEC28 | IX | 216658 | 217548 | 1 |
| YIL077C | 68.1574905 | -0.3998607 | 0.56271899 | 7.98245852 | 0.01847699 | 0.06230737 | NA | IX | 214991 | 215953 | -1 |
| YIL078W | 72.5993962 | -1.4218429 | 0.69827561 | 4.64697453 | 0.09793148 | 0.2158444 | THS1 | IX | 212499 | 214703 | 1 |
| YIL079C | 81.9993244 | -0.1509226 | 0.5800306 | 0.36529811 | 0.83306046 | 0.90138189 | AIR1 | IX | 210923 | 212005 | -1 |
| YIL082W-A | 4094.14431 | -0.5249382 | 0.5138091 | 1.14174845 | 0.56503126 | 0.71587111 | NA | IX | 205635 | 210132 | 1 |
| YIL083C | 21.8435299 | -0.6455157 | 0.70301605 | 0.86461479 | 0.64900984 | 0.7805914 | CAB2 | IX | 203556 | 204653 | -1 |
| YIL084C | 101.777571 | -0.0696548 | 0.51812774 | 1.09107942 | 0.57952892 | 0.72802613 | SDS3 | IX | 202276 | 203259 | -1 |

| | | | | | | | | | | | |
|---|---|---|---|---|---|---|---|---|---|---|---|
| YIL085C | 7.47599859 | -1.0157327 | 1.09476927 | 1.20980443 | 0.54612783 | 0.69889749 | KTR7 | IX | 200490 | 202043 | -1 |
| YIL086C | 44.3815795 | 2.31573643 | 0.74787514 | 11.0185513 | 0.00404904 | 0.01943942 | NA | IX | 200153 | 200461 | -1 |
| YIL087C | 14.9364468 | -0.5461697 | 0.76562501 | 0.98079082 | 0.6123842 | 0.7554524 | AIM19 | IX | 199646 | 200119 | -1 |
| YIL088C | 12.3482281 | -0.5365786 | 0.96987186 | 0.51251509 | 0.77394262 | 0.86281553 | AVT7 | IX | 197931 | 199403 | -1 |
| YIL089W | 353.360351 | 0.87151871 | 0.49169961 | 3.46959067 | 0.17643631 | 0.32890091 | NA | IX | 195599 | 196216 | 1 |
| YIL090W | 32.1425331 | 1.79994729 | 0.74287376 | 8.2380274 | 0.01626054 | 0.05678441 | ICE2 | IX | 193595 | 195070 | 1 |
| YIL091C | 140.303107 | -1.1169027 | 0.60904779 | 4.47783254 | 0.10657394 | 0.22939149 | UTP25 | IX | 191030 | 193195 | -1 |
| YIL092W | 107.933962 | -3.4373511 | 0.56014002 | 34.6504224 | 2.99E-08 | 8.09E-07 | NA | IX | 189066 | 190967 | 1 |
| YIL093C | 16.6518449 | -1.0190597 | 0.82120852 | 1.73739903 | 0.41949674 | 0.59032562 | RSM25 | IX | 187990 | 188784 | -1 |
| YIL094C | 25.8892758 | -1.397963 | 0.78452256 | 4.35140314 | 0.11352848 | 0.24016437 | LYS12 | IX | 186517 | 187632 | -1 |
| YIL095W | 258.168863 | -2.9260846 | 0.66318876 | 17.9048959 | 0.00012942 | 0.0011672 | PRK1 | IX | 183937 | 186369 | 1 |
| YIL096C | 72.7100059 | 2.63414911 | 0.57874098 | 25.6982144 | 2.63E-06 | 4.09E-05 | BMT5 | IX | 182117 | 183127 | -1 |
| YIL097W | 46.3620208 | 1.41490258 | 0.74759827 | 3.61375143 | 0.16416624 | 0.31287772 | FYV10 | IX | 180427 | 181977 | 1 |
| YIL098C | 17.6222265 | -1.1806729 | 0.87156915 | 2.63218253 | 0.26818151 | 0.44109199 | FMC1 | IX | 179772 | 180239 | -1 |
| YIL099W | 34.1635665 | -2.3639744 | 0.68261007 | 11.777699 | 0.00277016 | 0.01429522 | SGA1 | IX | 178004 | 179653 | 1 |
| YIL101C | 526.385024 | -0.1260988 | 0.49491407 | 7.74628452 | 0.02079293 | 0.06797914 | XBP1 | IX | 175307 | 177250 | -1 |
| YIL102C | 59.17031 | 0.72315668 | 0.61702117 | 2.11932414 | 0.34657291 | 0.51999408 | NA | IX | 174582 | 174887 | -1 |
| YIL102C-A | 98.6561237 | 3.05985982 | 0.61818697 | 31.0571235 | 1.80E-07 | 4.03E-06 | NA | IX | 173365 | 173592 | -1 |
| YIL103W | 1753.47932 | 1.10571356 | 0.49125719 | 4.94522234 | 0.08436428 | 0.19386304 | DPH1 | IX | 171751 | 173028 | 1 |
| YIL104C | 10.4128709 | -0.850153 | 0.89892185 | 1.40992465 | 0.49412719 | 0.65469026 | SHQ1 | IX | 169982 | 171505 | -1 |
| YIL105C | 20.421557 | -0.6350633 | 0.73324278 | 0.78194568 | 0.67639853 | 0.8007253 | SLM1 | IX | 167581 | 169641 | -1 |
| YIL105W-A | 1084.89459 | 0.10456689 | 0.47189939 | 2.43823851 | 0.2954903 | 0.46963735 | NA | IX | 167664 | 167804 | 1 |
| YIL106W | 158.090097 | 0.89992354 | 0.68977473 | 11.0648893 | 0.00395631 | 0.01908917 | MOB1 | IX | 166415 | 167444 | 1 |
| YIL107C | 959.758868 | 3.04537161 | 0.48528298 | 34.1638575 | 3.81E-08 | 1.01E-06 | PFK26 | IX | 163278 | 165761 | -1 |
| YIL108W | 92.060516 | 1.34814901 | 0.6420462 | 4.39800837 | 0.11091355 | 0.2359256 | NA | IX | 160887 | 162977 | 1 |
| YIL109C | 231.269482 | 1.59011111 | 0.6412936 | 7.07823004 | 0.02903901 | 0.08675743 | SEC24 | IX | 157385 | 160165 | -1 |
| YIL110W | 137.942905 | 3.04293305 | 0.55526624 | 27.4372308 | 1.10E-06 | 1.93E-05 | HPM1 | IX | 156045 | 157178 | 1 |
| YIL111W | 465.129622 | -1.345255 | 0.47537662 | 7.75318179 | 0.02072135 | 0.06786006 | NA | IX | 155311 | 155765 | 1 |
| YIL112W | 1010.07033 | 2.24099704 | 0.54322385 | 16.5631939 | 0.00025313 | 0.00209676 | HOS4 | IX | 151595 | 154846 | 1 |
| YIL113W | 115.499867 | -1.4610737 | 0.53482251 | 9.15629591 | 0.01027391 | 0.04024758 | SDP1 | IX | 150559 | 151188 | 1 |
| YIL114C | 8.70018485 | -1.8524369 | 1.05714961 | 5.22372896 | 0.07339757 | 0.17466992 | POR2 | IX | 149143 | 149988 | -1 |
| YIL115C | 194.33159 | -0.3283737 | 0.48009181 | 8.20854394 | 0.01650203 | 0.05733167 | NUP159 | IX | 144327 | 148709 | -1 |
| YIL116W | 36.4921859 | 0.37384411 | 0.78754925 | 3.6281821 | 0.16298599 | 0.31175673 | HIS5 | IX | 142928 | 144085 | 1 |

| | | | | | | | | | | | |
|---|---|---|---|---|---|---|---|---|---|---|---|
| YIL117C | 61.8600166 | 1.72534353 | 0.64827173 | 11.5892287 | 0.0030439 | 0.01538156 | PRM5 | IX | 140613 | 141569 | -1 |
| YIL118W | 45.7768081 | 1.16503824 | 0.72597974 | 2.69773675 | 0.25953379 | 0.431191 | RHO3 | IX | 139752 | 140447 | 1 |
| YIL119C | 103.777618 | -4.0720435 | 0.59477699 | 44.8622051 | 1.81E-10 | 7.89E-09 | RPI1 | IX | 136654 | 137877 | -1 |
| YIL120W | 356.995797 | 0.24788827 | 0.56829625 | 0.41026827 | 0.81453805 | 0.88867068 | QDR1 | IX | 134417 | 136108 | 1 |
| YIL121W | 889.248043 | 0.24960713 | 0.50728501 | 0.29008937 | 0.86498364 | 0.92054956 | QDR2 | IX | 132244 | 133872 | 1 |
| YIL123W | 106.217052 | 1.04789644 | 0.6258813 | 5.24855978 | 0.07249194 | 0.17315525 | SIM1 | IX | 128151 | 129581 | 1 |
| YIL124W | 88.4615462 | -0.4572324 | 0.54104348 | 1.23451738 | 0.53942113 | 0.6935928 | AYR1 | IX | 126204 | 127097 | 1 |
| YIL125W | 48.1488981 | 0.91670336 | 0.63919922 | 2.1368831 | 0.3435435 | 0.51705663 | KGD1 | IX | 122689 | 125733 | 1 |
| YIL126W | 61.4675757 | -1.3065686 | 0.62753689 | 5.99806847 | 0.04983517 | 0.13021013 | STH1 | IX | 117992 | 122071 | 1 |
| YIL127C | 65.3755146 | -2.3005364 | 0.90151027 | 14.2155881 | 0.0008187 | 0.00559654 | RRT14 | IX | 117024 | 117644 | -1 |
| YIL128W | 318.993074 | -3.0597981 | 0.49281771 | 37.2261647 | 8.25E-09 | 2.53E-07 | MET18 | IX | 113806 | 116904 | 1 |
| YIL129C | 32.6923415 | -1.0093796 | 0.72573305 | 2.4199603 | 0.2982032 | 0.47161883 | TAO3 | IX | 106107 | 113237 | -1 |
| YIL130W | 12.7734188 | -1.7238375 | 0.9130945 | 3.73992475 | 0.15412946 | 0.2985646 | ASG1 | IX | 102782 | 105676 | 1 |
| YIL131C | 31.9679858 | 0.93911938 | 0.63822788 | 4.334338 | 0.11450131 | 0.24134059 | FKH1 | IX | 100781 | 102235 | -1 |
| YIL132C | 17.529885 | 0.04045978 | 0.74250455 | 0.71724106 | 0.69863941 | 0.81736153 | CSM2 | IX | 99860 | 100501 | -1 |
| YIL133C | 40.4674163 | -2.4640983 | 0.75875824 | 10.4067899 | 0.00549787 | 0.02469562 | RPL16A | IX | 98527 | 99416 | -1 |
| YIL134W | 210.811528 | 0.47300265 | 0.58282723 | 1.31801991 | 0.51736329 | 0.67406244 | FLX1 | IX | 97395 | 98330 | 1 |
| YIL135C | 21.5076024 | -0.0336607 | 0.73389727 | 0.18274178 | 0.91267914 | 0.94617945 | VHS2 | IX | 95065 | 96375 | -1 |
| YIL136W | 80.5121934 | 0.29891166 | 0.57726631 | 1.297455 | 0.5227105 | 0.67858605 | OM45 | IX | 93619 | 94800 | 1 |
| YIL137C | 88.4740929 | 1.4465346 | 0.57439035 | 7.5809924 | 0.02258439 | 0.07260613 | TMA108 | IX | 89948 | 92788 | -1 |
| YIL138C | 37.5119893 | -0.8011916 | 0.6053115 | 1.99464457 | 0.36886584 | 0.54206426 | TPM2 | IX | 89230 | 89715 | -1 |
| YIL139C | 126.074958 | -1.3105347 | 0.5595476 | 6.30422625 | 0.04276167 | 0.11662274 | REV7 | IX | 87979 | 88716 | -1 |
| YIL140W | 93.5006198 | -1.0732875 | 0.59707447 | 3.81727352 | 0.14828239 | 0.29064152 | AXL2 | IX | 85366 | 87837 | 1 |
| YIL142W | 139.671254 | 1.23686577 | 0.51055344 | 5.8885574 | 0.05264002 | 0.1358225 | CCT2 | IX | 83302 | 84885 | 1 |
| YIL143C | 50.0357103 | 1.51713557 | 0.70644522 | 5.21948106 | 0.07355363 | 0.17489753 | SSL2 | IX | 80510 | 83041 | -1 |
| YIL144W | 19.4739474 | 1.50229056 | 0.79771844 | 4.49446626 | 0.10569125 | 0.22827341 | NDC80 | IX | 78074 | 80149 | 1 |
| YIL145C | 382.653333 | 2.71874139 | 0.49605123 | 28.1657094 | 7.65E-07 | 1.42E-05 | PAN6 | IX | 76354 | 77283 | -1 |
| YIL146C | 56.2396872 | -1.2820581 | 0.57512294 | 6.82579367 | 0.03294562 | 0.09600159 | ATG32 | IX | 74184 | 75773 | -1 |
| YIL147C | 58.7196971 | -0.8037475 | 0.54858433 | 2.39997139 | 0.30119852 | 0.47428323 | SLN1 | IX | 69791 | 73453 | -1 |
| YIL148W | 161.166612 | -1.321213 | 0.73159626 | 12.6791619 | 0.00176504 | 0.01019919 | RPL40B | IX | 68708 | 69528 | 1 |
| YIL149C | 39.8050834 | -1.5326909 | 0.63524064 | 6.57964058 | 0.03726054 | 0.10539255 | MLP2 | IX | 63028 | 68067 | -1 |
| YIL150C | 31.8215211 | 1.4132146 | 0.65910246 | 10.0456487 | 0.0065859 | 0.02854218 | MCM10 | IX | 61013 | 62728 | -1 |
| YIL151C | 94.708406 | 0.88097477 | 0.61265514 | 2.54183916 | 0.28057349 | 0.45466569 | ESL1 | IX | 57338 | 60694 | -1 |

| | | | | | | | | | | | |
|---|---|---|---|---|---|---|---|---|---|---|---|
| YIL152W | 87.7985914 | -0.711384 | 0.57106017 | 1.57249344 | 0.4555514 | 0.62135279 | NA | IX | 56545 | 57252 | 1 |
| YIL153W | 161.935523 | 1.97282428 | 0.51001092 | 15.7042613 | 0.00038892 | 0.00299848 | RRD1 | IX | 55198 | 56379 | 1 |
| YIL154C | 210.887786 | 0.58704054 | 0.55829832 | 1.5089576 | 0.47025565 | 0.63276324 | IMP2' | IX | 53981 | 55021 | -1 |
| YIL155C | 87.0775446 | 0.36089438 | 0.53161456 | 1.29655388 | 0.52294607 | 0.67871828 | GUT2 | IX | 51759 | 53708 | -1 |
| YIL156W | 15.9377886 | 0.54061168 | 0.79651156 | 0.68003197 | 0.71175894 | 0.82438173 | UBP7 | IX | 48091 | 51306 | 1 |
| YIL157C | 23.8254105 | -1.0487743 | 0.71123974 | 2.52304756 | 0.28322213 | 0.45678444 | COA1 | IX | 46949 | 47542 | -1 |
| YIL158W | 438.192242 | -0.4432874 | 0.77616089 | 6.36634841 | 0.04145386 | 0.11428804 | AIM20 | IX | 46201 | 46815 | 1 |
| YIL159W | 162.441967 | 3.79960735 | 0.58267206 | 42.8138377 | 5.05E-10 | 1.95E-08 | BNR1 | IX | 41825 | 45952 | 1 |
| YIL160C | 509.511224 | -1.6279883 | 0.48962138 | 10.6030116 | 0.00498408 | 0.02284865 | POT1 | IX | 40191 | 41444 | -1 |
| YIL161W | 131.172184 | 0.78438842 | 0.51117079 | 2.33184935 | 0.31163437 | 0.48405923 | NA | IX | 39433 | 40140 | 1 |
| YIL162W | 7.85473547 | -0.0491973 | 1.06969078 | 0.95394312 | 0.62066018 | 0.76119942 | SUC2 | IX | 37385 | 38983 | 1 |
| YIL163C | 303.459563 | -0.4765462 | 0.55399197 | 0.93293295 | 0.62721464 | 0.76483005 | NA | IX | 36899 | 37252 | -1 |
| YIL166C | 525.238228 | 0.18635152 | 0.54722227 | 0.20702628 | 0.90166418 | 0.93824416 | NA | IX | 30938 | 32566 | -1 |
| YIL167W | 26.0132288 | 1.61163204 | 0.70617353 | 12.1598884 | 0.0022883 | 0.01234789 | NA | NA | NA | NA | NA |
| YIL169C | 5707.65919 | -1.53634 | 0.47695098 | 15.7156026 | 0.00038672 | 0.00298949 | CSS1 | IX | 23119 | 26106 | -1 |
| YIL170W | 586.665378 | 1.67765807 | 0.48151492 | 11.6957789 | 0.00288598 | 0.0147744 | NA | IX | 19847 | 21220 | 1 |
| YIL171W | 117.314393 | 1.37083053 | 0.57628564 | 5.72935287 | 0.05700157 | 0.14462713 | NA | IX | 19515 | 19844 | 1 |
| YIR001C | 66.2537475 | 0.77819879 | 0.59881777 | 1.68508122 | 0.43061511 | 0.59948398 | SGN1 | IX | 356143 | 356895 | -1 |
| YIR002C | 979.451431 | 4.84327657 | 0.48777915 | 75.3989585 | 4.24E-17 | 6.63E-15 | MPH1 | IX | 357415 | 360396 | -1 |
| YIR003W | 94.583085 | 0.14437337 | 0.52889503 | 0.16522649 | 0.92070717 | 0.95024857 | AIM21 | IX | 360885 | 362924 | 1 |
| YIR004W | 44.5477632 | -0.0712119 | 0.73265416 | 2.59556725 | 0.2731365 | 0.44661404 | DJP1 | IX | 363221 | 364519 | 1 |
| YIR005W | 62.942193 | 0.47511657 | 0.6074793 | 1.1165271 | 0.5722018 | 0.72242661 | IST3 | IX | 364889 | 365335 | 1 |
| YIR006C | 58.2337649 | 0.22394621 | 0.59061509 | 1.52519407 | 0.46645346 | 0.62925571 | PAN1 | IX | 365466 | 369908 | -1 |
| YIR007W | 112.677182 | -0.7907854 | 0.68684031 | 4.05286323 | 0.13180501 | 0.26711622 | EGH1 | IX | 370704 | 372998 | 1 |
| YIR008C | 88.8782071 | 0.69891183 | 0.74962172 | 3.6727254 | 0.15939614 | 0.30640892 | PRI1 | IX | 373077 | 374306 | -1 |
| YIR009W | 29.8973564 | -2.2856283 | 0.68214381 | 10.9969008 | 0.00409311 | 0.0195435 | MSL1 | IX | 374525 | 374860 | 1 |
| YIR010W | 26.1677564 | -0.2294756 | 0.65956985 | 0.81335788 | 0.66585794 | 0.79245991 | DSN1 | IX | 375431 | 377161 | 1 |
| YIR011C | 35.2724218 | -0.6587861 | 0.73026723 | 4.63002577 | 0.09876491 | 0.21726779 | STS1 | IX | 377287 | 378246 | -1 |
| YIR012W | 37.9083094 | -1.3699319 | 0.6393243 | 4.56978747 | 0.10178488 | 0.22205518 | SQT1 | IX | 378486 | 379781 | 1 |
| YIR013C | 13.4716366 | -0.1553894 | 0.86403019 | 2.04907247 | 0.3589629 | 0.53196703 | GAT4 | IX | 380019 | 380384 | -1 |
| YIR014W | 21.0448467 | -1.4245792 | 0.71478413 | 4.37026477 | 0.11246284 | 0.23882864 | NA | IX | 381086 | 381814 | 1 |
| YIR015W | 16.2793808 | 0.57682681 | 0.77253856 | 0.71102279 | 0.70081496 | 0.81801018 | RPR2 | IX | 381948 | 382382 | 1 |
| YIR016W | 94.0653152 | 1.79643634 | 0.84599113 | 6.18391803 | 0.0454129 | 0.12167548 | NA | IX | 382628 | 383425 | 1 |

| | | | | | | | | | | | |
|---|---|---|---|---|---|---|---|---|---|---|---|
| YIR017C | 17.8101895 | 1.205743 | 0.78416905 | 2.40652205 | 0.30021361 | 0.47337604 | MET28 | IX | 383556 | 384119 | -1 |
| YIR018C-A | 341.765649 | -0.4881503 | 0.58792276 | 0.88836351 | 0.64134884 | 0.77591986 | NA | IX | 385564 | 385701 | -1 |
| YIR018W | 23.1441236 | 0.66592496 | 0.68723471 | 1.46049248 | 0.48179034 | 0.64424159 | YAP5 | IX | 384609 | 385346 | 1 |
| YIR019C | 75.1340145 | 0.87364512 | 0.59146743 | 3.50139394 | 0.17365287 | 0.32507061 | FLO11 | IX | 389572 | 393675 | -1 |
| YIR021W | 21.1404179 | 1.90613298 | 0.76698197 | 9.86885262 | 0.00719459 | 0.03067501 | MRS1 | IX | 397294 | 398385 | 1 |
| YIR021W-A | 11.8945348 | 0.41544023 | 0.85678408 | 1.06985021 | 0.58571316 | 0.73372548 | NA | IX | 398514 | 398726 | 1 |
| YIR022W | 47.5760305 | -0.4246864 | 0.74305604 | 2.91480899 | 0.23283983 | 0.40027153 | SEC11 | IX | 398733 | 399236 | 1 |
| YIR023W | 376.994198 | 3.89169877 | 0.61019634 | 49.9561545 | 1.42E-11 | 8.06E-10 | DAL81 | IX | 399777 | 402689 | 1 |
| YIR024C | 12.3374953 | -1.0001468 | 0.92519408 | 1.4876267 | 0.47529798 | 0.63776948 | INA22 | IX | 402841 | 403491 | -1 |
| YIR025W | 44.6138112 | -1.7829524 | 0.72781709 | 5.92009829 | 0.05181637 | 0.1343428 | MND2 | IX | 403659 | 404765 | 1 |
| YIR026C | 169.640587 | -1.6269876 | 0.52696605 | 10.094684 | 0.00642639 | 0.02793454 | YVH1 | IX | 404873 | 405967 | -1 |
| YIR027C | 407.679798 | 1.76126904 | 0.55196485 | 11.0388309 | 0.00400819 | 0.01927527 | DAL1 | IX | 406260 | 407642 | -1 |
| YIR028W | 345.395778 | 0.31660796 | 0.57822158 | 0.78234082 | 0.67626491 | 0.8007253 | DAL4 | IX | 408468 | 410375 | 1 |
| YIR029W | 23.5433017 | 0.46997771 | 0.75595771 | 4.17747354 | 0.12384348 | 0.25554303 | DAL2 | IX | 410807 | 411838 | 1 |
| YIR030C | 176.302789 | 1.16392899 | 0.57148083 | 8.64207128 | 0.01328612 | 0.04868773 | DCG1 | IX | 412036 | 412770 | -1 |
| YIR031C | 15.3585968 | 0.50800461 | 0.87823866 | 0.48233367 | 0.78571053 | 0.87017291 | DAL7 | IX | 413015 | 414679 | -1 |
| YIR032C | 25.3673057 | -0.1999751 | 0.74915901 | 0.60606286 | 0.73857588 | 0.84040603 | DAL3 | IX | 415030 | 415617 | -1 |
| YIR033W | 58.2176227 | -0.2783821 | 0.60115346 | 1.37743943 | 0.50221864 | 0.66087407 | MGA2 | IX | 416124 | 419465 | 1 |
| YIR034C | 73.9418615 | 0.09263053 | 0.6178709 | 0.101853 | 0.95034852 | 0.96603194 | LYS1 | IX | 419615 | 420736 | -1 |
| YIR035C | 99.1665315 | -0.5989723 | 0.53601323 | 2.79191074 | 0.24759638 | 0.41832973 | NA | IX | 421026 | 421790 | -1 |
| YIR036C | 21.6222898 | -0.5012406 | 0.71131665 | 0.54938805 | 0.75980457 | 0.85451323 | IRC24 | IX | 422074 | 422865 | -1 |
| YIR037W | 18.8947794 | 0.28972243 | 0.85801058 | 2.1326766 | 0.34426681 | 0.51781024 | HYR1 | IX | 423128 | 423619 | 1 |
| YIR038C | 10.2064656 | 0.62847702 | 0.9798937 | 2.28099663 | 0.31965969 | 0.49223979 | GTT1 | IX | 423809 | 424513 | -1 |
| YIR039C | 14.0736163 | -0.447681 | 0.8429238 | 0.54192172 | 0.76264635 | 0.85565759 | YPS6 | IX | 430498 | 432111 | -1 |
| YIR042C | 44.5687664 | -1.4612322 | 0.66403534 | 6.87430367 | 0.03215614 | 0.0942798 | NA | IX | 435273 | 435983 | -1 |
| YIR043C | 42.2360246 | -2.554334 | 0.63548774 | 16.5892075 | 0.00024986 | 0.00207561 | NA | NA | NA | NA | NA |
| YJL001W | 115.938678 | 1.11153356 | 0.56021537 | 4.78764529 | 0.09128008 | 0.205327 | PRE3 | X | 435163 | 435926 | 1 |
| YJL002C | 46.018459 | -0.5948178 | 0.73711498 | 6.70389695 | 0.03501606 | 0.10046729 | OST1 | X | 433221 | 434651 | -1 |
| YJL003W | 37.4157626 | 0.75685845 | 0.63946803 | 2.65283363 | 0.26542663 | 0.43759117 | COX16 | X | 432644 | 433000 | 1 |
| YJL004C | 61.4410743 | 1.15595817 | 0.59798371 | 4.22768226 | 0.12077317 | 0.25039546 | SYS1 | X | 431589 | 432200 | -1 |
| YJL005W | 26.4296549 | 1.12405312 | 0.74953116 | 2.32575589 | 0.31258528 | 0.4849178 | CYR1 | X | 425157 | 431237 | 1 |
| YJL007C | 46.7306776 | -0.8472691 | 0.68691832 | 1.58783239 | 0.45207093 | 0.61810237 | NA | X | 422698 | 423012 | -1 |
| YJL008C | 36.1541747 | -2.0795822 | 0.81800201 | 6.3739102 | 0.04129743 | 0.1140258 | CCT8 | X | 419957 | 421663 | -1 |

| | | | | | | | | | | | |
|---|---|---|---|---|---|---|---|---|---|---|---|
| YJL010C | 21.8408143 | -2.5287948 | 0.82914503 | 8.95350578 | 0.01137027 | 0.04332089 | NOP9 | X | 417562 | 419562 | -1 |
| YJL012C | 20.0181818 | -0.6707293 | 0.75893736 | 0.93870687 | 0.6254065 | 0.76426839 | VTC4 | X | 411234 | 413399 | -1 |
| YJL013C | 39.5148822 | 1.63846493 | 0.67514482 | 7.88786662 | 0.01937187 | 0.06459254 | MAD3 | X | 409493 | 411040 | -1 |
| YJL014W | 137.611957 | -1.8590524 | 0.79839851 | 5.17551942 | 0.0751883 | 0.17776245 | CCT3 | X | 407558 | 409162 | 1 |
| YJL016W | 250.047478 | 3.12483838 | 0.51449465 | 46.8523351 | 6.70E-11 | 3.29E-09 | TPH3 | X | 405588 | 407273 | 1 |
| YJL019W | 145.717939 | -0.7808363 | 0.58024597 | 2.30344593 | 0.31609169 | 0.48830599 | MPS3 | X | 402897 | 404945 | 1 |
| YJL020C | 143.271949 | 1.45443835 | 0.53888106 | 9.08180985 | 0.01066375 | 0.04137474 | BBC1 | X | 398937 | 402410 | -1 |
| YJL023C | 82.7049345 | 1.54025601 | 0.59789415 | 10.1916279 | 0.00612232 | 0.02692266 | PET130 | X | 397355 | 398398 | -1 |
| YJL024C | 144.642944 | 0.7159203 | 0.51493398 | 6.16563493 | 0.04582995 | 0.122171 | APS3 | X | 395931 | 396592 | -1 |
| YJL025W | 36.4432925 | 1.17591284 | 0.62814472 | 9.68434485 | 0.0078899 | 0.03284148 | RRN7 | X | 393967 | 395511 | 1 |
| YJL026W | 59.6352509 | -1.8197634 | 0.63580308 | 10.7025151 | 0.00474218 | 0.02193071 | RNR2 | X | 392404 | 393603 | 1 |
| YJL027C | 75.8158859 | -0.9410581 | 0.73983415 | 1.60377847 | 0.44848088 | 0.61504128 | NA | X | 391833 | 392249 | -1 |
| YJL028W | 8.78339306 | -1.9354942 | 0.9898227 | 3.83587822 | 0.14690941 | 0.28883039 | NA | X | 391311 | 391646 | 1 |
| YJL029C | 292.74313 | -0.4059308 | 0.48985611 | 2.18681187 | 0.33507331 | 0.50840526 | VPS53 | X | 388385 | 390853 | -1 |
| YJL030W | 24.1620609 | 1.65271802 | 0.82503215 | 4.95509791 | 0.08394874 | 0.19311211 | MAD2 | X | 387657 | 388247 | 1 |
| YJL031C | 16.0418051 | 0.30882364 | 0.85287562 | 0.84388666 | 0.6557712 | 0.78495521 | BET4 | X | 386368 | 387438 | -1 |
| YJL033W | 21.432247 | -1.567326 | 0.73906046 | 4.78641623 | 0.0913362 | 0.20537343 | HCA4 | X | 383837 | 386149 | 1 |
| YJL034W | 321.984392 | 2.07595237 | 0.7598459 | 8.1617441 | 0.01689273 | 0.05839337 | KAR2 | X | 381327 | 383375 | 1 |
| YJL035C | 331.42487 | 2.7878342 | 0.49141507 | 30.043965 | 2.99E-07 | 6.26E-06 | TAD2 | X | 380249 | 381001 | -1 |
| YJL036W | 18.0479 | 1.24672719 | 0.95840344 | 4.24515929 | 0.11972239 | 0.2489022 | SNX4 | X | 378825 | 380096 | 1 |
| YJL037W | 48.9346594 | 1.68797842 | 0.67806348 | 10.8279454 | 0.00445391 | 0.02079165 | IRC18 | X | 376662 | 377336 | 1 |
| YJL038C | 664.986833 | 1.56560473 | 0.45865966 | 13.0953425 | 0.00143345 | 0.00883884 | LOH1 | X | 375115 | 375774 | -1 |
| YJL039C | 490.139535 | 5.36000661 | 0.5810507 | 97.3209389 | 7.36E-22 | 3.28E-19 | NUP192 | X | 368748 | 373799 | -1 |
| YJL041W | 506.202545 | 3.85828106 | 0.73156436 | 22.8255312 | 1.11E-05 | 0.00014382 | NA | X | 365903 | 368373 | 1 |
| YJL042W | 314.414421 | -0.9869351 | 0.49072143 | 7.6635744 | 0.02167085 | 0.07033309 | MHP1 | X | 361249 | 365445 | 1 |
| YJL043W | 16.761287 | 0.1850866 | 0.81628548 | 0.88660416 | 0.64191327 | 0.77608641 | NA | X | 360130 | 360903 | 1 |
| YJL044C | 33.7259217 | -1.7594691 | 0.64444034 | 8.92349507 | 0.01154218 | 0.04387997 | GYP6 | X | 358300 | 359676 | -1 |
| YJL045W | 152.280364 | 0.35361931 | 0.54300647 | 2.15983426 | 0.33962367 | 0.5126228 | NA | X | 356024 | 357928 | 1 |
| YJL046W | 8.34702949 | -0.4832102 | 0.96336957 | 0.67606504 | 0.7131721 | 0.82535807 | AIM22 | X | 352386 | 353615 | 1 |
| YJL047C | 49.1046436 | -0.4030268 | 0.57761459 | 0.90376144 | 0.63643008 | 0.77284609 | RTT101 | X | 349580 | 352108 | -1 |
| YJL048C | 66.387266 | 0.758309 | 0.80562124 | 0.91886058 | 0.6316434 | 0.76880708 | UBX6 | X | 347447 | 348637 | -1 |
| YJL049W | 418.89263 | 2.66418679 | 1.00979296 | 10.67503 | 0.0048078 | 0.02214573 | CHM7 | X | 345973 | 347325 | 1 |
| YJL050W | 50.9237924 | -0.7069591 | 0.6121594 | 2.81448817 | 0.24481705 | 0.41519939 | MTR4 | X | 342522 | 345743 | 1 |

| | | | | | | | | | | | |
|---|---|---|---|---|---|---|---|---|---|---|---|
| YJL051W | 803.453439 | -1.4555276 | 0.50520446 | 10.1304875 | 0.00631237 | 0.02756307 | IRC8 | X | 339787 | 342255 | 1 |
| YJL052C-A | 19.011907 | 2.20376299 | 0.8245754 | 7.79624826 | 0.02027992 | 0.06680024 | NA | X | 337884 | 338003 | -1 |
| YJL052W | 12.2452933 | 0.53707053 | 0.968086 | 3.66438097 | 0.16006257 | 0.30728192 | TDH1 | X | 338271 | 339269 | 1 |
| YJL053W | 52.5839467 | 1.16770818 | 0.56652725 | 7.40847655 | 0.02461896 | 0.07694787 | PEP8 | X | 335898 | 337037 | 1 |
| YJL054W | 42.9404315 | -0.5827428 | 0.59883222 | 1.09506594 | 0.57837492 | 0.72736551 | TIM54 | X | 334265 | 335701 | 1 |
| YJL055W | 48.9284572 | -1.4089649 | 0.60116518 | 6.22678324 | 0.04444994 | 0.11984737 | NA | X | 333357 | 334094 | 1 |
| YJL056C | 29.8260311 | 0.89971557 | 0.8102151 | 2.58457194 | 0.27464224 | 0.44819013 | ZAP1 | X | 330431 | 333073 | -1 |
| YJL057C | 120.154538 | -0.057833 | 0.58210422 | 1.9319232 | 0.38061702 | 0.55440819 | IKS1 | X | 328118 | 330121 | -1 |
| YJL058C | 172.414194 | 1.75074071 | 0.49135742 | 12.319714 | 0.00211256 | 0.01164923 | BIT61 | X | 326242 | 327873 | -1 |
| YJL059W | 156.551187 | 0.10217336 | 0.53055568 | 0.62100822 | 0.73307731 | 0.83751334 | YHC3 | X | 324964 | 326190 | 1 |
| YJL060W | 128.837948 | 1.30102173 | 0.6090524 | 19.0622921 | 7.26E-05 | 0.00071568 | BNA3 | X | 323386 | 324720 | 1 |
| YJL061W | 25.7713721 | 0.05914636 | 0.69830432 | 0.68948069 | 0.70840427 | 0.82395756 | NUP82 | X | 320016 | 322157 | 1 |
| YJL062W | 238.909954 | 0.06418139 | 0.47364177 | 1.54088199 | 0.46280893 | 0.62608965 | LAS21 | X | 317284 | 319776 | 1 |
| YJL062W-A | 14.6114592 | 0.20939353 | 0.81610414 | 0.40759045 | 0.81562938 | 0.88902268 | COA3 | X | 316723 | 316980 | 1 |
| YJL063C | 36.4157979 | -2.7965475 | 0.68375117 | 19.7060503 | 5.26E-05 | 0.00053606 | MRPL8 | X | 315759 | 316475 | -1 |
| YJL065C | 11.7342153 | 0.30614666 | 0.9384626 | 1.86346629 | 0.39387048 | 0.5656027 | DLS1 | X | 315054 | 315557 | -1 |
| YJL066C | 13.2092808 | 0.34615488 | 0.8498494 | 0.5005859 | 0.77857267 | 0.86541289 | MPM1 | X | 314114 | 314872 | -1 |
| YJL068C | 42.5573331 | 0.23656175 | 0.58678125 | 0.17398411 | 0.91668438 | 0.94860224 | NA | X | 313016 | 313915 | -1 |
| YJL069C | 308.348865 | 2.89978893 | 0.54713192 | 25.9908689 | 2.27E-06 | 3.59E-05 | UTP18 | X | 310922 | 312706 | -1 |
| YJL070C | 84.6267711 | 0.64351682 | 0.52273132 | 2.23160323 | 0.32765252 | 0.50039829 | NA | X | 307971 | 310637 | -1 |
| YJL071W | 29.9092487 | 1.20463233 | 0.66201501 | 3.76511262 | 0.15220054 | 0.29581776 | ARG2 | X | 306132 | 307856 | 1 |
| YJL072C | 141.767339 | 0.79089247 | 0.75210468 | 9.88411235 | 0.0071399 | 0.03048675 | PSF2 | X | 305221 | 305862 | -1 |
| YJL073W | 466.03553 | -1.8647636 | 0.49636053 | 13.3646973 | 0.00125283 | 0.00791047 | JEM1 | X | 303181 | 305118 | 1 |
| YJL074C | 21.6632617 | 0.15835234 | 0.70618353 | 0.1679756 | 0.91944247 | 0.94974417 | SMC3 | X | 299157 | 302849 | -1 |
| YJL075C | 1961.2185 | -0.9715556 | 0.45360469 | 4.52268889 | 0.10421029 | 0.22564606 | NA | X | 298460 | 298876 | -1 |
| YJL076W | 90.1237899 | 1.2335176 | 0.5360097 | 9.32619583 | 0.00943718 | 0.03773943 | NET1 | X | 295245 | 298814 | 1 |
| YJL077C | 1000.08439 | 0.23517944 | 0.46090708 | 0.61264547 | 0.736149 | 0.83936643 | ICS3 | X | 294666 | 295061 | -1 |
| YJL077W-A | 14.1315864 | -0.1263668 | 0.92672846 | 0.38895665 | 0.82326402 | 0.8939795 | NA | X | 294799 | 294885 | 1 |
| YJL077W-B | 105.047676 | -4.0832803 | 0.61239734 | 43.6618952 | 3.30E-10 | 1.36E-08 | NA | X | 294047 | 294145 | 1 |
| YJL078C | 3529.42512 | 0.31327751 | 0.49018446 | 3.25159832 | 0.19675437 | 0.35456204 | PRY3 | X | 291336 | 293981 | -1 |
| YJL079C | 126.311695 | -1.0141165 | 0.60304839 | 3.05461629 | 0.21711934 | 0.38117676 | PRY1 | X | 289875 | 290774 | -1 |
| YJL080C | 91.4973251 | 0.74006962 | 0.63613707 | 2.55648357 | 0.27852658 | 0.45301284 | SCP160 | X | 285558 | 289226 | -1 |
| YJL081C | 46.6140875 | -0.0438027 | 0.60066421 | 3.32806383 | 0.1893739 | 0.34544902 | ARP4 | X | 283802 | 285271 | -1 |

| | | | | | | | | | | | |
|---|---|---|---|---|---|---|---|---|---|---|---|
| YJL082W | 63.3023852 | 0.64965534 | 0.63612358 | 1.07443565 | 0.58437182 | 0.73236208 | IML2 | X | 281185 | 283380 | 1 |
| YJL083W | 134.580051 | -1.2494767 | 0.5779982 | 7.32610439 | 0.02565409 | 0.07917761 | TAX4 | X | 278841 | 280655 | 1 |
| YJL084C | 45.5815958 | -0.417456 | 0.61318111 | 0.69936357 | 0.70491237 | 0.82055541 | ALY2 | X | 274862 | 278002 | -1 |
| YJL085W | 16.3586046 | -1.1103126 | 0.96742901 | 7.30871628 | 0.0258781 | 0.07969905 | EXO70 | X | 272827 | 274698 | 1 |
| YJL087C | 19.4887545 | -1.2776832 | 0.80767717 | 3.4193827 | 0.18092163 | 0.33553116 | TRL1 | X | 270002 | 272485 | -1 |
| YJL088W | 29.5625519 | 0.45129844 | 1.16977203 | 10.9407011 | 0.00420976 | 0.01997909 | ARG3 | X | 268799 | 269815 | 1 |
| YJL089W | 140.498629 | 2.58053482 | 0.53561585 | 21.5596467 | 2.08E-05 | 0.00024546 | SIP4 | X | 265926 | 268415 | 1 |
| YJL090C | 49.9696471 | -0.4274314 | 0.60360517 | 1.54808644 | 0.46114478 | 0.62486036 | DPB11 | X | 262757 | 265051 | -1 |
| YJL091C | 125.97909 | -0.4969558 | 0.67046878 | 0.58884017 | 0.74496348 | 0.84442806 | GWT1 | X | 261080 | 262552 | -1 |
| YJL092W | 914.205947 | -0.1576186 | 0.47568399 | 6.4564698 | 0.03962738 | 0.11084181 | SRS2 | X | 257423 | 260947 | 1 |
| YJL093C | 27.5423638 | 1.67252477 | 0.73316236 | 6.3198085 | 0.0424298 | 0.11599082 | TOK1 | X | 254737 | 256812 | -1 |
| YJL094C | 170.346014 | 3.57829089 | 0.58287011 | 50.9243528 | 8.75E-12 | 5.32E-10 | KHA1 | X | 251821 | 254442 | -1 |
| YJL095W | 307.087897 | 1.04666662 | 0.61931021 | 4.10221692 | 0.12859229 | 0.2622576 | BCK1 | X | 247255 | 251691 | 1 |
| YJL096W | 6.42874269 | -0.0939667 | 1.21447121 | 1.19734257 | 0.54954134 | 0.7014648 | MRPL49 | X | 246490 | 246975 | 1 |
| YJL097W | 9.3053542 | -1.606218 | 1.04425869 | 2.34047735 | 0.31029287 | 0.4830878 | PHS1 | X | 245591 | 246244 | 1 |
| YJL098W | 24.2571793 | -1.1803017 | 0.71160885 | 2.78498029 | 0.24845584 | 0.41928281 | SAP185 | X | 242082 | 245258 | 1 |
| YJL099W | 590.302935 | 4.52299187 | 0.51546574 | 64.2673809 | 1.11E-14 | 1.05E-12 | CHS6 | X | 239414 | 241654 | 1 |
| YJL100W | 1942.63559 | 4.27793266 | 0.46504639 | 66.1070287 | 4.42E-15 | 4.57E-13 | LSB6 | X | 237263 | 239086 | 1 |
| YJL101C | 196.975152 | 4.2557533 | 0.62610543 | 59.1352219 | 1.44E-13 | 1.14E-11 | GSH1 | X | 234320 | 236356 | -1 |
| YJL102W | 15.7476461 | 0.92447783 | 0.77479965 | 1.84875122 | 0.39677909 | 0.56785755 | MEF2 | X | 231301 | 233760 | 1 |
| YJL103C | 126.695169 | -0.4159187 | 0.51405495 | 2.43076461 | 0.2965966 | 0.47066158 | GSM1 | X | 229025 | 230881 | -1 |
| YJL104W | 16.9735945 | -0.251837 | 0.8256067 | 0.69838158 | 0.70525856 | 0.82079338 | PAM16 | X | 227327 | 227776 | 1 |
| YJL105W | 796.982847 | -2.090714 | 0.51711335 | 15.1261936 | 0.00051926 | 0.00378167 | SET4 | X | 225055 | 226737 | 1 |
| YJL106W | 44.1926295 | -1.6525145 | 0.63601355 | 15.2307763 | 0.00049281 | 0.00363486 | IME2 | X | 221390 | 223327 | 1 |
| YJL107C | 50.6835132 | -1.8315394 | 0.62672635 | 8.24532493 | 0.01620132 | 0.05674873 | NA | X | 218853 | 220016 | -1 |
| YJL108C | 47.9525563 | -4.6851008 | 0.76248364 | 40.0268301 | 2.03E-09 | 7.09E-08 | PRM10 | X | 217705 | 218856 | -1 |
| YJL109C | 923.89909 | 4.65676029 | 0.5201032 | 62.2090364 | 3.10E-14 | 2.76E-12 | UTP10 | X | 212000 | 217309 | -1 |
| YJL110C | 8.33835741 | 1.67554471 | 1.10346548 | 3.58970704 | 0.16615179 | 0.31541602 | GZF3 | X | 209922 | 211577 | -1 |
| YJL111W | 35.6748238 | -0.5236314 | 0.78976027 | 0.85612498 | 0.65177068 | 0.78244649 | CCT7 | X | 207877 | 209529 | 1 |
| YJL112W | 27.4963476 | 1.14504192 | 0.66976278 | 3.91099448 | 0.1414941 | 0.28066148 | MDV1 | X | 205305 | 207449 | 1 |
| YJL114W | 12.6831236 | 1.15730203 | 0.89101328 | 4.42014546 | 0.10969267 | 0.23410268 | NA | X | 197915 | 198595 | 1 |
| YJL115W | 168.211743 | -0.0253791 | 0.64260948 | 1.33513183 | 0.51295564 | 0.66997815 | ASF1 | X | 196287 | 197126 | 1 |
| YJL116C | 358.539493 | 0.93172289 | 0.48750517 | 5.90330838 | 0.0522532 | 0.13512551 | NCA3 | X | 193860 | 194873 | -1 |

| | | | | | | | | | | | |
|---|---|---|---|---|---|---|---|---|---|---|---|
| YJL117W | 154.771384 | 2.5545997 | 0.52318591 | 24.802001 | 4.11E-06 | 6.05E-05 | PHO86 | X | 192531 | 193466 | 1 |
| YJL118W | 212.86262 | 0.21046275 | 0.57107478 | 0.24048444 | 0.88670563 | 0.93215081 | NA | X | 191639 | 192298 | 1 |
| YJL121C | 74.8531822 | -2.622877 | 0.66732936 | 14.1297222 | 0.00085461 | 0.00578738 | RPE1 | X | 190374 | 191090 | -1 |
| YJL122W | 115.710313 | -2.9045057 | 0.55824256 | 24.2329662 | 5.47E-06 | 7.70E-05 | ALB1 | X | 189716 | 190243 | 1 |
| YJL123C | 263.422156 | -0.3097871 | 0.57874679 | 1.2165024 | 0.54430191 | 0.69728958 | MTC1 | X | 188004 | 189440 | -1 |
| YJL124C | 107.656394 | 0.80212194 | 0.53025959 | 4.89231889 | 0.08662564 | 0.19825705 | LSM1 | X | 187126 | 187644 | -1 |
| YJL125C | 185.626203 | 1.03791741 | 0.987862 | 7.42584858 | 0.02440605 | 0.07671608 | GCD14 | X | 185528 | 186679 | -1 |
| YJL126W | 13.3042666 | 0.00441645 | 0.8644823 | 0.08160692 | 0.96001779 | 0.97173192 | NIT2 | X | 184501 | 185424 | 1 |
| YJL127C | 107.303658 | 1.89661251 | 0.56299081 | 10.8119676 | 0.00448964 | 0.02089629 | SPT10 | X | 182298 | 184220 | -1 |
| YJL127C-B | 171.483754 | 4.43126461 | 0.5971953 | 47.0813539 | 5.98E-11 | 2.96E-09 | NA | X | 181551 | 181709 | -1 |
| YJL128C | 56.8419471 | -0.104035 | 0.58533105 | 2.53534907 | 0.28148545 | 0.45563342 | PBS2 | X | 178097 | 180103 | -1 |
| YJL129C | 255.307575 | 2.6029972 | 0.56896811 | 20.7303881 | 3.15E-05 | 0.00035018 | TRK1 | X | 173599 | 177306 | -1 |
| YJL130C | 98.7702461 | -0.8097825 | 0.56997076 | 2.10998122 | 0.3481957 | 0.52188793 | URA2 | X | 165723 | 172367 | -1 |
| YJL131C | 269.370531 | 0.59929151 | 0.63172116 | 1.83787645 | 0.3989424 | 0.56977714 | AIM23 | X | 164278 | 165348 | -1 |
| YJL132W | 103.149947 | 0.19592391 | 0.52669199 | 0.20468453 | 0.90272053 | 0.93871262 | NA | X | 161914 | 164166 | 1 |
| YJL133C-A | 5.93825843 | 1.95818746 | 1.2258759 | 3.37684409 | 0.18481092 | 0.34056499 | NA | X | 159623 | 159847 | -1 |
| YJL133W | 2397.30506 | 5.8346762 | 0.50560683 | 92.006077 | 1.05E-20 | 3.38E-18 | MRS3 | X | 160619 | 161563 | 1 |
| YJL134W | 60.6802689 | 2.85502778 | 0.70915806 | 17.532639 | 0.0001559 | 0.00137666 | LCB3 | X | 158188 | 159417 | 1 |
| YJL136C | 57.4074546 | -2.5103288 | 0.70550559 | 12.4158378 | 0.00201342 | 0.01120935 | RPS21B | X | 156550 | 157273 | -1 |
| YJL136W-A | 52.8886722 | 0.07578247 | 0.62139552 | 0.06654067 | 0.96727703 | 0.97486964 | NA | X | 156169 | 156252 | 1 |
| YJL137C | 37.2184536 | -2.311243 | 0.72621168 | 10.4222908 | 0.00545542 | 0.02454304 | GLG2 | X | 154985 | 156127 | -1 |
| YJL138C | 34.3960905 | -1.6686219 | 0.65796854 | 6.17772353 | 0.04555378 | 0.12191042 | TIF2 | X | 153504 | 154691 | -1 |
| YJL139C | 88.2817351 | -0.3885964 | 0.64350364 | 0.88509985 | 0.64239627 | 0.77621415 | YUR1 | X | 151713 | 152999 | -1 |
| YJL140W | 1353.92133 | -0.0410383 | 0.47548702 | 2.35105226 | 0.30865654 | 0.48144434 | RPB4 | X | 150961 | 151626 | 1 |
| YJL141C | 43.3554427 | 0.5406688 | 0.62983489 | 0.80501387 | 0.6686417 | 0.79450097 | YAK1 | X | 147967 | 150390 | -1 |
| YJL142C | 556.20442 | -2.0005101 | 0.66502838 | 13.0598025 | 0.00145915 | 0.00894319 | NA | X | 147819 | 148211 | -1 |
| YJL143W | 50.9498734 | 0.99272475 | 0.60148272 | 2.68628708 | 0.26102384 | 0.43235173 | TIM17 | X | 147101 | 147577 | 1 |
| YJL144W | 103.596966 | 3.30920966 | 0.58923671 | 30.8212338 | 2.03E-07 | 4.42E-06 | NA | X | 146359 | 146673 | 1 |
| YJL145W | 460.796424 | 5.62459003 | 0.5384117 | 98.8274313 | 3.47E-22 | 1.82E-19 | SFH5 | X | 145160 | 146044 | 1 |
| YJL146W | 478.678088 | 0.75335986 | 0.50129074 | 2.23117927 | 0.32772199 | 0.50039829 | IDS2 | X | 143292 | 144701 | 1 |
| YJL147C | 122.233543 | -1.3366919 | 0.51022362 | 12.4507016 | 0.00197863 | 0.01106886 | MRX5 | X | 141419 | 142567 | -1 |
| YJL148W | 81.0747874 | -3.0659599 | 0.59441351 | 24.6798362 | 4.37E-06 | 6.35E-05 | RPA34 | X | 140437 | 141138 | 1 |
| YJL149W | 44.5713871 | -0.5501996 | 0.6109952 | 1.29068429 | 0.52448306 | 0.67982022 | DAS1 | X | 137379 | 139370 | 1 |

| | | | | | | | | | | | |
|---|---|---|---|---|---|---|---|---|---|---|---|
| YJL151C | 582.489954 | 3.44750057 | 0.50649714 | 39.3190518 | 2.90E-09 | 9.81E-08 | SNA3 | X | 136372 | 136773 | -1 |
| YJL153C | 62.6404634 | -0.7700662 | 0.60544273 | 2.55641313 | 0.27853639 | 0.45301284 | INO1 | X | 134332 | 135933 | -1 |
| YJL154C | 415.148645 | 5.05737686 | 0.5192689 | 81.3829561 | 2.13E-18 | 4.25E-16 | VPS35 | X | 131101 | 133935 | -1 |
| YJL155C | 27.4101396 | 1.01509021 | 0.81479392 | 3.97400587 | 0.13710572 | 0.27478094 | FBP26 | X | 129285 | 130643 | -1 |
| YJL156C | 557.772413 | 0.04553395 | 0.51892158 | 0.00959614 | 0.99521342 | 0.99607359 | SSY5 | X | 126853 | 128952 | -1 |
| YJL157C | 6181.51688 | 1.28399108 | 0.49131695 | 7.85748842 | 0.01966836 | 0.06514862 | FAR1 | X | 123836 | 126328 | -1 |
| YJL158C | 69.2434908 | 0.78864042 | 0.63027849 | 4.55435286 | 0.10257342 | 0.22310297 | CIS3 | X | 122265 | 122948 | -1 |
| YJL159W | 309.371886 | 0.89269668 | 0.49911218 | 15.7570342 | 0.00037879 | 0.00294788 | HSP150 | X | 120449 | 121690 | 1 |
| YJL160C | 96.5299955 | -0.1184936 | 0.57606813 | 0.07406111 | 0.96364669 | 0.97305796 | PIR5 | X | 117962 | 118825 | -1 |
| YJL161W | 62.3408202 | -2.3856768 | 0.60394181 | 15.2757852 | 0.00048184 | 0.00357676 | FMP33 | X | 117245 | 117787 | 1 |
| YJL162C | 45.8846464 | 1.33630737 | 0.62866002 | 5.33734997 | 0.06934405 | 0.16715322 | JJJ2 | X | 113878 | 115629 | -1 |
| YJL163C | 266.428937 | 0.52463135 | 0.49954819 | 2.42431484 | 0.29755464 | 0.47136562 | NA | X | 111666 | 113333 | -1 |
| YJL164C | 99.2758131 | 0.32916205 | 0.55302503 | 0.46993561 | 0.7905963 | 0.87307888 | TPK1 | X | 109966 | 111159 | -1 |
| YJL165C | 120.058348 | 0.80861857 | 0.73587341 | 4.80691563 | 0.09040481 | 0.20391268 | HAL5 | X | 106894 | 109461 | -1 |
| YJL166W | 25.3914042 | -0.5546688 | 0.76560054 | 0.67856342 | 0.71228176 | 0.82465735 | QCR8 | X | 106434 | 106718 | 1 |
| YJL167W | 69.5022343 | 0.17697189 | 0.71437032 | 2.49327812 | 0.28746934 | 0.46170527 | ERG20 | X | 105014 | 106072 | 1 |
| YJL168C | 20.3496734 | -2.1378556 | 0.78411259 | 7.94157718 | 0.01885856 | 0.06326248 | SET2 | X | 102227 | 104428 | -1 |
| YJL170C | 70.6463743 | -0.6404201 | 0.5875049 | 1.21495354 | 0.5447236 | 0.69762213 | ASG7 | X | 101150 | 101779 | -1 |
| YJL171C | 75.1984857 | -0.7662438 | 0.91521783 | 0.70883965 | 0.70158037 | 0.81832198 | TOH1 | X | 99704 | 100894 | -1 |
| YJL172W | 88.8217577 | -0.039169 | 0.57475537 | 15.6880163 | 0.00039209 | 0.00301892 | CPS1 | X | 97737 | 99467 | 1 |
| YJL173C | 281.600017 | 0.04681649 | 0.75665274 | 4.65700332 | 0.09744164 | 0.21492841 | RFA3 | X | 96161 | 96529 | -1 |
| YJL174W | 27.1285024 | 1.0074183 | 0.79267296 | 2.42028502 | 0.29815479 | 0.47161883 | KRE9 | X | 95092 | 95922 | 1 |
| YJL176C | 44.3130636 | -1.1998483 | 0.61479505 | 4.11276086 | 0.12791613 | 0.26147876 | SWI3 | X | 92053 | 94530 | -1 |
| YJL177W | 75.9171548 | -3.1567222 | 0.75796794 | 15.7106413 | 0.00038768 | 0.00299292 | RPL17B | X | 90786 | 91657 | 1 |
| YJL178C | 47.4464551 | -0.1503798 | 0.58285814 | 0.46096305 | 0.79415111 | 0.87495229 | ATG27 | X | 89282 | 90097 | -1 |
| YJL179W | 11.4726746 | -0.9477747 | 1.07688824 | 8.46387623 | 0.01452421 | 0.0523127 | PFD1 | X | 88787 | 89116 | 1 |
| YJL180C | 237.011716 | 2.75951043 | 0.48556043 | 33.6088349 | 5.03E-08 | 1.32E-06 | ATP12 | X | 87583 | 88560 | -1 |
| YJL181W | 243.379109 | 3.86411517 | 0.64780517 | 29.8273338 | 3.33E-07 | 6.80E-06 | RBH1 | X | 85660 | 87495 | 1 |
| YJL183W | 14.2238489 | -1.6977176 | 0.82507825 | 4.52877084 | 0.10389387 | 0.22521358 | MNN11 | X | 84068 | 85336 | 1 |
| YJL184W | 39.1326857 | -0.6862731 | 0.6578431 | 1.28602017 | 0.52570761 | 0.68095013 | GON7 | X | 83448 | 83819 | 1 |
| YJL185C | 171.215454 | -2.073261 | 0.5201809 | 17.3915818 | 0.00016729 | 0.00145654 | ATG36 | X | 82095 | 82976 | -1 |
| YJL186W | 85.3785603 | -1.7136066 | 0.60617369 | 7.62468687 | 0.02209634 | 0.07131784 | MNN5 | X | 80155 | 81915 | 1 |
| YJL187C | 246.300995 | 0.40332231 | 0.53789288 | 1.32957135 | 0.51438376 | 0.67150244 | SWE1 | X | 76803 | 79262 | -1 |

| | | | | | | | | | | | |
|---|---|---|---|---|---|---|---|---|---|---|---|
| YJL189W | 408.889744 | -2.8365708 | 0.5979331 | 20.4221875 | 3.68E-05 | 0.00040008 | RPL39 | X | 75933 | 76474 | 1 |
| YJL190C | 57.1930306 | -3.6068352 | 0.78772616 | 19.0494741 | 7.30E-05 | 0.00071783 | RPS22A | X | 74910 | 75302 | -1 |
| YJL191W | 27.3181265 | -2.0951114 | 0.79200397 | 8.70903453 | 0.01284864 | 0.04759047 | RPS14B | X | 73787 | 74611 | 1 |
| YJL192C | 21.5020388 | -0.5946483 | 0.78504762 | 1.63672453 | 0.44115356 | 0.60960103 | SOP4 | X | 72710 | 73414 | -1 |
| YJL193W | 80.7304619 | 0.41920947 | 0.72316731 | 0.42941421 | 0.80677771 | 0.88420271 | NA | X | 71366 | 72574 | 1 |
| YJL194W | 28.877216 | -0.5837644 | 0.67621322 | 5.01345254 | 0.08153473 | 0.18875892 | CDC6 | X | 69338 | 70879 | 1 |
| YJL196C | 241.93037 | -0.7146594 | 0.48707092 | 3.41166879 | 0.18162078 | 0.33650698 | ELO1 | X | 67850 | 68782 | -1 |
| YJL197W | 50.3680959 | -1.1197668 | 0.61708866 | 3.27003262 | 0.19494919 | 0.3521859 | UBP12 | X | 63805 | 67569 | 1 |
| YJL198W | 12.6741171 | -1.3648146 | 0.92614013 | 3.16594119 | 0.20536414 | 0.36665383 | PHO90 | X | 60844 | 63489 | 1 |
| YJL199C | 10.7785091 | 1.55493317 | 1.15601622 | 2.06657827 | 0.35583465 | 0.52936346 | NA | X | 59856 | 60182 | -1 |
| YJL200C | 172.800056 | 2.95221376 | 0.59850416 | 27.796587 | 9.21E-07 | 1.66E-05 | ACO2 | X | 56444 | 58813 | -1 |
| YJL201W | 36.9538348 | 0.85205969 | 0.6175669 | 3.50498284 | 0.17334154 | 0.32501539 | ECM25 | X | 54379 | 56178 | 1 |
| YJL203W | 13.1056946 | -0.3401697 | 0.95251185 | 0.20804341 | 0.90120574 | 0.93824416 | PRP21 | X | 53341 | 54183 | 1 |
| YJL204C | 18.2892024 | -0.4639601 | 0.75392935 | 0.55192272 | 0.75884225 | 0.85429037 | RCY1 | X | 50629 | 53151 | -1 |
| YJL205C | 76.5662057 | 0.6395929 | 0.538515 | 3.95837753 | 0.13818129 | 0.27626715 | NCE101 | X | 50139 | 50443 | -1 |
| YJL206C | 30.1388211 | -2.2118637 | 0.67952727 | 11.4900971 | 0.00319858 | 0.01606225 | NA | X | 47659 | 49935 | -1 |
| YJL207C | 28.5499004 | 0.45819989 | 0.65719209 | 0.60901559 | 0.73748628 | 0.84022935 | LAA1 | X | 41389 | 47433 | -1 |
| YJL208C | 38.4753009 | 1.30499603 | 0.62409238 | 4.27579602 | 0.11790241 | 0.24644584 | NUC1 | X | 40194 | 41183 | -1 |
| YJL209W | 27.4203554 | -3.2317063 | 0.73577152 | 18.3843288 | 0.00010183 | 0.000951 | CBP1 | X | 38005 | 39969 | 1 |
| YJL210W | 70.791331 | 0.74267746 | 0.63322378 | 1.82846055 | 0.40082503 | 0.57147918 | PEX2 | X | 36919 | 37734 | 1 |
| YJL212C | 190.003944 | 1.18385352 | 0.52813196 | 6.82013011 | 0.03303905 | 0.0962254 | OPT1 | X | 33850 | 36249 | -1 |
| YJL213W | 27.9656877 | -0.6356568 | 0.67436414 | 2.07386057 | 0.35454135 | 0.52838982 | NA | X | 32163 | 33158 | 1 |
| YJL214W | 609.218516 | -0.2492542 | 0.63549658 | 0.17415447 | 0.9166063 | 0.94860224 | HXT8 | X | 26887 | 28596 | 1 |
| YJL216C | 9.34708582 | -1.5627264 | 1.01803786 | 3.28562156 | 0.19343557 | 0.35020614 | IMA5 | X | 24341 | 26086 | -1 |
| YJL217W | 13.6900742 | -1.992338 | 0.8937694 | 5.06949516 | 0.07928173 | 0.18517193 | REE1 | X | 23133 | 23729 | 1 |
| YJL219W | 91.4101822 | -0.0937412 | 0.54977998 | 4.46284649 | 0.1073755 | 0.23043148 | HXT9 | X | 19497 | 21200 | 1 |
| YJL225C | 9.26512606 | -0.941884 | 0.92728951 | 3.86164511 | 0.14502886 | 0.28581248 | NA | X | 466 | 6130 | -1 |
| YJR001W | 130.424412 | 0.57880756 | 0.53044377 | 1.53476619 | 0.46422632 | 0.62726488 | AVT1 | X | 436802 | 438610 | 1 |
| YJR002W | 115.246635 | 0.88512767 | 0.52836878 | 7.8872584 | 0.01937776 | 0.06459254 | MPP10 | X | 438865 | 440646 | 1 |
| YJR003C | 94.506714 | -3.5321541 | 0.65145093 | 25.638446 | 2.71E-06 | 4.19E-05 | MRX12 | X | 440994 | 442553 | -1 |
| YJR004C | 699.681641 | 1.62512472 | 0.4834784 | 12.7662857 | 0.0016898 | 0.00985293 | SAG1 | X | 442909 | 444861 | -1 |
| YJR005W | 14.6446637 | 0.66778692 | 1.04481443 | 2.03196518 | 0.36204651 | 0.53489393 | APL1 | X | 445923 | 448025 | 1 |
| YJR006W | 44.480877 | -2.1656345 | 0.71672784 | 10.1730025 | 0.0061796 | 0.02710599 | POL31 | X | 449202 | 450665 | 1 |

| | | | | | | | | | | | |
|---|---|---|---|---|---|---|---|---|---|---|---|
| YJR007W | 30.5629523 | -0.4272853 | 0.65031902 | 0.82240216 | 0.66285363 | 0.79018376 | SUI2 | X | 451020 | 451934 | 1 |
| YJR008W | 97.6650905 | -2.7782887 | 0.60161829 | 26.2023697 | 2.04E-06 | 3.30E-05 | MHO1 | X | 452430 | 453446 | 1 |
| YJR009C | 78.7815646 | -1.7868813 | 0.74977724 | 6.35438079 | 0.04170266 | 0.11459819 | TDH2 | X | 453683 | 454681 | -1 |
| YJR010C-A | 206.917255 | -0.8300462 | 0.50645508 | 2.96023281 | 0.22761119 | 0.39362867 | SPC1 | X | 458077 | 458361 | -1 |
| YJR010W | 58.4306194 | 1.3464151 | 0.58506953 | 6.09885672 | 0.047386 | 0.12539532 | MET3 | X | 456239 | 457774 | 1 |
| YJR011C | 14.0052138 | -1.1001218 | 0.86177931 | 1.77780759 | 0.41110616 | 0.58269393 | NA | X | 458641 | 459426 | -1 |
| YJR012C | 12.8345677 | -1.063827 | 0.94834171 | 3.63623943 | 0.16233069 | 0.31091456 | NA | X | 459795 | 460418 | -1 |
| YJR013W | 91.1343385 | -2.2335106 | 0.53380342 | 17.2664162 | 0.00017809 | 0.00153674 | GPI14 | X | 460382 | 461593 | 1 |
| YJR014W | 89.2123005 | -1.0415499 | 0.6248337 | 4.39297427 | 0.11119308 | 0.23643332 | TMA22 | X | 461829 | 462425 | 1 |
| YJR015W | 26.16382 | 0.70002193 | 0.6698306 | 2.50833947 | 0.28531263 | 0.45887782 | NA | X | 462721 | 464253 | 1 |
| YJR016C | 37.1563054 | -1.5963844 | 0.70161245 | 5.31280905 | 0.07020017 | 0.16872519 | ILV3 | X | 464451 | 466208 | -1 |
| YJR017C | 48.5289065 | -1.3769509 | 0.60236221 | 5.17701281 | 0.07513217 | 0.17770232 | ESS1 | X | 466521 | 467033 | -1 |
| YJR019C | 219.291914 | -0.7028307 | 0.58378974 | 1.51717305 | 0.46832793 | 0.63119616 | TES1 | X | 467232 | 468281 | -1 |
| YJR021C | 67.2569711 | -0.4813049 | 0.59178483 | 0.73086377 | 0.6938969 | 0.81470846 | REC107 | X | 468555 | 469579 | -1 |
| YJR022W | 25.5769993 | 0.35075989 | 1.28887664 | 5.32078152 | 0.06992089 | 0.16819359 | LSM8 | X | 469784 | 470113 | 1 |
| YJR024C | 30.6369115 | -0.7105255 | 0.85178952 | 7.32965814 | 0.02560855 | 0.07911509 | MDE1 | X | 470230 | 470964 | -1 |
| YJR025C | 63.8207648 | -0.376345 | 0.75998277 | 1.65532277 | 0.43707023 | 0.60599536 | BNA1 | X | 471138 | 471671 | -1 |
| YJR030C | 267.553759 | -1.693425 | 0.48463382 | 11.7274423 | 0.00284065 | 0.01464593 | RBH2 | X | 483959 | 486196 | -1 |
| YJR031C | 38.896291 | 1.59306207 | 0.71205952 | 7.4214932 | 0.02445926 | 0.07676865 | GEA1 | X | 486586 | 490812 | -1 |
| YJR032W | 136.464066 | 4.19379604 | 0.60053165 | 48.558231 | 2.86E-11 | 1.49E-09 | CPR7 | X | 491081 | 492262 | 1 |
| YJR033C | 147.525321 | -0.5183181 | 0.70095486 | 2.2277076 | 0.32829135 | 0.50073944 | RAV1 | X | 492378 | 496451 | -1 |
| YJR034W | 10.607743 | 1.69817661 | 0.97251805 | 4.1505244 | 0.12552351 | 0.25827332 | PET191 | X | 496683 | 497009 | 1 |
| YJR035W | 58.9579429 | 0.53075899 | 0.57252341 | 1.90741853 | 0.38530916 | 0.55815362 | RAD26 | X | 497355 | 500612 | 1 |
| YJR036C | 125.578664 | -0.548204 | 0.60484087 | 2.07182544 | 0.35490231 | 0.52857757 | HUL4 | X | 500713 | 503391 | -1 |
| YJR039W | 58.7682428 | -1.936337 | 0.57073491 | 15.0603514 | 0.00053664 | 0.00388561 | NA | X | 503936 | 507301 | 1 |
| YJR040W | 76.3046596 | -0.3693087 | 0.64147238 | 0.68799436 | 0.70893093 | 0.82407349 | GEF1 | X | 507746 | 510085 | 1 |
| YJR041C | 14.258325 | -0.8066644 | 0.84519316 | 0.94143264 | 0.62455473 | 0.76345394 | URB2 | X | 510239 | 513763 | -1 |
| YJR042W | 24.200017 | -0.6632093 | 0.70829743 | 0.94715896 | 0.62276908 | 0.76281637 | NUP85 | X | 514055 | 516289 | 1 |
| YJR043C | 17.9547947 | -1.8413041 | 0.81063307 | 5.4060733 | 0.06700174 | 0.16264366 | POL32 | X | 516461 | 517513 | -1 |
| YJR044C | 102.482711 | 1.8879466 | 0.53192044 | 12.6769725 | 0.00176697 | 0.01020018 | VPS55 | X | 518763 | 519185 | -1 |
| YJR045C | 122.322449 | -0.3207741 | 0.78872737 | 0.21370137 | 0.89865985 | 0.9375281 | SSC1 | X | 519638 | 521602 | -1 |
| YJR046W | 13.3295612 | -0.6900202 | 0.86929434 | 2.18320503 | 0.33567813 | 0.50893088 | TAH11 | X | 522048 | 523862 | 1 |
| YJR048W | 107.502032 | -2.1975915 | 0.63120946 | 11.4258067 | 0.00330307 | 0.0164313 | CYC1 | X | 526335 | 526664 | 1 |

| | | | | | | | | | | | |
|---|---|---|---|---|---|---|---|---|---|---|---|
| YJR049C | 53.3372832 | -0.3306266 | 0.66810234 | 0.80629162 | 0.66821466 | 0.79444823 | UTR1 | X | 526884 | 528476 | -1 |
| YJR050W | 14.8919863 | -1.9879902 | 0.89347835 | 5.04598524 | 0.08021918 | 0.18675877 | ISY1 | X | 528697 | 529404 | 1 |
| YJR051W | 51.3429471 | 0.77550844 | 0.57403605 | 1.89217391 | 0.38825732 | 0.56046121 | OSM1 | X | 529861 | 531366 | 1 |
| YJR052W | 35.1691426 | 0.44551199 | 0.6186482 | 0.58663954 | 0.74578362 | 0.84519224 | RAD7 | X | 532062 | 533759 | 1 |
| YJR053W | 68.7711988 | 0.08968533 | 0.63689379 | 3.38778747 | 0.18380245 | 0.33935465 | BFA1 | X | 534027 | 535751 | 1 |
| YJR054W | 81.3624894 | 1.52010487 | 0.5391444 | 7.70257323 | 0.02125238 | 0.06924663 | KCH1 | X | 536056 | 537549 | 1 |
| YJR055W | 6.42884824 | -0.712102 | 1.06473277 | 0.46773885 | 0.79146515 | 0.87352731 | HIT1 | X | 538772 | 539266 | 1 |
| YJR056C | 16.6775826 | -0.938239 | 0.79695323 | 1.37968087 | 0.50165611 | 0.66050451 | NA | X | 541792 | 542502 | -1 |
| YJR057W | 163.031073 | -0.1406152 | 0.608906 | 1.77026748 | 0.41265898 | 0.58426143 | CDC8 | X | 544062 | 544712 | 1 |
| YJR058C | 165.241021 | -1.4821441 | 0.53248227 | 9.08085919 | 0.01066882 | 0.04137474 | APS2 | X | 544732 | 545175 | -1 |
| YJR059W | 122.76682 | 0.71585496 | 0.50972347 | 2.4159619 | 0.29879996 | 0.47191811 | PTK2 | X | 545787 | 548243 | 1 |
| YJR060W | 42.5269261 | -0.3717462 | 0.67859822 | 0.72624006 | 0.69550295 | 0.81611981 | CBF1 | X | 548759 | 549814 | 1 |
| YJR061W | 1878.34739 | -0.4131 | 0.47560208 | 1.09609868 | 0.57807634 | 0.727271 | NA | X | 550511 | 553318 | 1 |
| YJR062C | 8.50294115 | 0.61957471 | 0.99416135 | 0.41483735 | 0.81267933 | 0.88731158 | NTA1 | X | 553476 | 554849 | -1 |
| YJR063W | 6.67275698 | -1.8672038 | 1.19087994 | 2.44544435 | 0.29442759 | 0.46884326 | RPA12 | X | 555195 | 555572 | 1 |
| YJR064W | 61.9067673 | -1.9192478 | 0.67321038 | 18.6096648 | 9.10E-05 | 0.00086218 | CCT5 | X | 555914 | 557602 | 1 |
| YJR065C | 44.84875 | 0.51503432 | 0.71520705 | 1.51537577 | 0.46874898 | 0.63146966 | ARP3 | X | 557809 | 559158 | -1 |
| YJR066W | 333.281053 | -1.1472668 | 0.51365727 | 7.21374065 | 0.02713664 | 0.0824783 | TOR1 | X | 559416 | 566828 | 1 |
| YJR067C | 13.1671849 | -0.3815748 | 1.04853009 | 2.64601886 | 0.26633258 | 0.43871 | YAE1 | X | 567019 | 567444 | -1 |
| YJR068W | 489.633996 | -0.8470095 | 0.86931833 | 7.9366423 | 0.01890515 | 0.06338205 | RFC2 | X | 567643 | 568704 | 1 |
| YJR069C | 56.4966958 | -0.4658294 | 0.62502541 | 2.0150294 | 0.3651253 | 0.53820659 | HAM1 | X | 568806 | 569399 | -1 |
| YJR070C | 23.4144297 | -0.7371382 | 0.87694638 | 0.83926486 | 0.65728838 | 0.78549013 | LIA1 | X | 569621 | 570598 | -1 |
| YJR072C | 49.8537921 | -1.6166938 | 0.58142075 | 7.55798948 | 0.02284565 | 0.07312122 | NPA3 | X | 570967 | 572124 | -1 |
| YJR073C | 21.2435479 | -0.9359008 | 0.89491315 | 1.31187257 | 0.51895594 | 0.67522582 | OPI3 | X | 572315 | 572935 | -1 |
| YJR074W | 238.474681 | 1.50199764 | 0.65028038 | 5.12769041 | 0.07700806 | 0.18110344 | MOG1 | X | 573095 | 573751 | 1 |
| YJR075W | 61.6640786 | 1.0404099 | 0.5895599 | 4.3263156 | 0.11496152 | 0.24204626 | HOC1 | X | 573981 | 575171 | 1 |
| YJR076C | 156.896106 | 0.37364331 | 0.74326347 | 10.8123766 | 0.00448872 | 0.02089629 | CDC11 | X | 575354 | 576601 | -1 |
| YJR077C | 45.6762697 | 0.12594685 | 0.77945438 | 0.31132748 | 0.85584693 | 0.9146094 | MIR1 | X | 577255 | 578190 | -1 |
| YJR078W | 20.1889223 | 1.73907802 | 0.78618357 | 5.8133336 | 0.05465761 | 0.13990609 | BNA2 | X | 578860 | 580221 | 1 |
| YJR079W | 998.098346 | -0.6930058 | 0.46222149 | 3.24324596 | 0.19757777 | 0.35571372 | NA | X | 580205 | 581239 | 1 |
| YJR080C | 279.421625 | 2.35138049 | 0.49753822 | 20.7450632 | 3.13E-05 | 0.00034964 | AIM24 | X | 580432 | 581616 | -1 |
| YJR082C | 6.20026956 | 1.34867942 | 1.14405486 | 1.47483911 | 0.47834667 | 0.64096904 | EAF6 | X | 581914 | 582255 | -1 |
| YJR083C | 48.1456682 | 1.04732172 | 0.62783168 | 3.36242999 | 0.18614767 | 0.34183159 | ACF4 | X | 582608 | 583537 | -1 |

| | | | | | | | | | | | |
|---|---|---|---|---|---|---|---|---|---|---|---|
| YJR084W | 53.6844611 | 1.8790453 | 0.61892273 | 9.16983971 | 0.01020457 | 0.04005725 | NA | X | 583733 | 585004 | 1 |
| YJR085C | 42.9170799 | 0.21075157 | 0.82177996 | 6.0610479 | 0.04829033 | 0.12713301 | NA | X | 585120 | 585437 | -1 |
| YJR086W | 646.259016 | -0.1266492 | 0.64157847 | 5.12034559 | 0.07729138 | 0.18169594 | STE18 | X | 586068 | 586400 | 1 |
| YJR088C | 405.281537 | -2.4550825 | 0.51993616 | 20.1885205 | 4.13E-05 | 0.00044055 | EMC2 | X | 586495 | 587373 | -1 |
| YJR089W | 86.133817 | 2.58956156 | 0.55447139 | 25.9234627 | 2.35E-06 | 3.70E-05 | BIR1 | X | 587718 | 590582 | 1 |
| YJR090C | 27.1345701 | 1.1737564 | 0.74939334 | 2.5672124 | 0.27703645 | 0.45146103 | GRR1 | X | 590872 | 594327 | -1 |
| YJR091C | 106.55512 | 1.71019258 | 0.5982583 | 9.51720287 | 0.0085776 | 0.03502418 | JSN1 | X | 595061 | 598336 | -1 |
| YJR092W | 191.335585 | 0.27385104 | 0.4992959 | 0.30586355 | 0.85818827 | 0.91626865 | BUD4 | X | 598735 | 603078 | 1 |
| YJR093C | 20.4223302 | -0.8093778 | 0.7487529 | 1.53242476 | 0.46477011 | 0.62728439 | FIP1 | X | 603220 | 604203 | -1 |
| YJR094C | 8.95062326 | -0.2113615 | 0.9561256 | 0.23484644 | 0.88920878 | 0.93304075 | IME1 | X | 604569 | 605651 | -1 |
| YJR094W-A | 69.166065 | -2.7716893 | 0.8024101 | 11.0457843 | 0.00399428 | 0.01925635 | RPL43B | X | 608305 | 608858 | 1 |
| YJR095W | 60.022549 | 0.83912601 | 0.64778764 | 1.97779737 | 0.37198614 | 0.54526576 | SFC1 | X | 609773 | 610741 | 1 |
| YJR096W | 316.977027 | 0.04790873 | 0.52958237 | 0.13781763 | 0.93341179 | 0.95735444 | NA | X | 611195 | 612043 | 1 |
| YJR097W | 23.1712516 | 1.1067117 | 0.77098756 | 2.3955273 | 0.30186854 | 0.47448396 | JJJ3 | X | 612413 | 612931 | 1 |
| YJR098C | 16.5668403 | -0.8830054 | 0.75696982 | 1.39417308 | 0.4980342 | 0.65775958 | NA | X | 613186 | 615156 | -1 |
| YJR099W | 82.9140628 | 0.51467156 | 0.82453928 | 3.80221781 | 0.14940285 | 0.29203007 | YUH1 | X | 615576 | 616286 | 1 |
| YJR100C | 200.21551 | -0.4849982 | 0.67227427 | 3.85188572 | 0.14573828 | 0.28701519 | AIM25 | X | 616351 | 617334 | -1 |
| YJR101W | 52.446302 | -1.4550781 | 0.78123951 | 8.24978465 | 0.01616524 | 0.05670589 | RSM26 | X | 618234 | 619034 | 1 |
| YJR102C | 210.791026 | -0.5680454 | 0.75930727 | 4.48519734 | 0.10618221 | 0.22871838 | VPS25 | X | 619157 | 619765 | -1 |
| YJR103W | 1193.8047 | -0.0969853 | 0.94761037 | 6.36882746 | 0.04140251 | 0.11426146 | URA8 | X | 620754 | 622490 | 1 |
| YJR104C | 191.520352 | -0.2131566 | 0.59752752 | 1.88805007 | 0.3890587 | 0.56089429 | SOD1 | X | 622550 | 623014 | -1 |
| YJR105W | 927.15484 | 3.6947444 | 0.55566606 | 36.5408976 | 1.16E-08 | 3.42E-07 | ADO1 | X | 623581 | 624603 | 1 |
| YJR106W | 70.8389838 | -0.7836639 | 0.58209682 | 5.66749483 | 0.05879212 | 0.14748831 | ECM27 | X | 624838 | 627015 | 1 |
| YJR107C-A | 36.4917666 | 0.64211619 | 0.86382329 | 8.29214469 | 0.01582646 | 0.05593181 | NA | NA | NA | NA | NA |
| YJR107W | 626.375473 | -0.8816528 | 0.53136033 | 2.71346853 | 0.25750033 | 0.42958807 | LIH1 | X | 627340 | 628326 | 1 |
| YJR108W | 56.8564524 | 0.59319111 | 0.67582216 | 0.76115738 | 0.68346578 | 0.80596066 | ABM1 | X | 628712 | 629083 | 1 |
| YJR109C | 202.446166 | -0.9543435 | 0.51922608 | 4.16272558 | 0.12476007 | 0.25697646 | CPA2 | X | 629585 | 632941 | -1 |
| YJR110W | 18.2875502 | 0.81130187 | 0.75887738 | 1.56866178 | 0.456425 | 0.62207609 | YMR1 | X | 633615 | 635681 | 1 |
| YJR111C | 29.7089708 | 1.3296564 | 0.80706352 | 3.32166669 | 0.1899806 | 0.34583253 | PXP2 | X | 635855 | 636706 | -1 |
| YJR112W | 16.6779089 | -2.3651434 | 0.8572247 | 9.49930821 | 0.00865469 | 0.03526435 | NNF1 | X | 637030 | 637635 | 1 |
| YJR112W-A | 238.546208 | -0.3561173 | 0.82447419 | 7.39521507 | 0.02478275 | 0.07731256 | NA | X | 637790 | 638168 | 1 |
| YJR113C | 110.813539 | 0.47457286 | 0.58083186 | 1.00660975 | 0.60452946 | 0.7496735 | RSM7 | X | 638232 | 638975 | -1 |
| YJR115W | 13.0246363 | -1.1995892 | 0.88546643 | 2.71323612 | 0.25753026 | 0.42958807 | NA | X | 639942 | 640451 | 1 |

| | | | | | | | | | | | |
|---|---|---|---|---|---|---|---|---|---|---|---|
| YJR116W | 194.925208 | -0.1753285 | 0.60253547 | 0.09638417 | 0.95295072 | 0.96731849 | TDA4 | X | 640825 | 641664 | 1 |
| YJR117W | 46.6450083 | 0.66882049 | 0.90112236 | 8.22917898 | 0.01633264 | 0.05693318 | STE24 | X | 642007 | 643368 | 1 |
| YJR118C | 358.282613 | -1.8335422 | 0.71418909 | 7.42878344 | 0.02437026 | 0.07671608 | ILM1 | X | 643490 | 644101 | -1 |
| YJR119C | 130.911035 | -1.6381881 | 0.54531914 | 8.69940929 | 0.01291063 | 0.04773469 | JHD2 | X | 644304 | 646490 | -1 |
| YJR120W | 29.9841622 | 2.12474444 | 0.80660149 | 11.1380611 | 0.00381418 | 0.0184804 | NA | X | 647126 | 647476 | 1 |
| YJR121W | 127.215234 | -0.7286098 | 0.7539873 | 1.55884525 | 0.45867076 | 0.62328904 | ATP2 | X | 647607 | 649142 | 1 |
| YJR122W | 67.5453114 | -0.1633493 | 0.55653714 | 2.40020816 | 0.30116287 | 0.47428323 | IBA57 | X | 649776 | 651269 | 1 |
| YJR123W | 245.660746 | -3.3855645 | 0.69461131 | 20.5457397 | 3.46E-05 | 0.00037968 | RPS5 | X | 651901 | 652578 | 1 |
| YJR124C | 212.260736 | 1.79149284 | 0.50365347 | 12.0219278 | 0.00245172 | 0.01302799 | NA | X | 652892 | 654238 | -1 |
| YJR125C | 52.2658751 | -0.002943 | 0.65216158 | 2.14307455 | 0.34248162 | 0.51615859 | ENT3 | X | 654737 | 655963 | -1 |
| YJR126C | 30.3031533 | 0.63710066 | 0.83201765 | 1.73786191 | 0.41939967 | 0.59032562 | VPS70 | X | 656254 | 658689 | -1 |
| YJR127C | 175.78987 | 0.24976584 | 0.5221134 | 0.24558055 | 0.88444913 | 0.9315066 | RSF2 | X | 658917 | 663059 | -1 |
| YJR128W | 118.04878 | -1.3812328 | 0.64948995 | 5.07242535 | 0.07916566 | 0.18505013 | NA | X | 662921 | 663280 | 1 |
| YJR129C | 35.3831869 | -1.5522008 | 0.66183243 | 7.16846461 | 0.02775797 | 0.08383862 | EFM3 | X | 664000 | 665019 | -1 |
| YJR130C | 136.977473 | -0.5452029 | 0.50829553 | 1.95899121 | 0.37550045 | 0.54902718 | STR2 | X | 665218 | 667137 | -1 |
| YJR131W | 153.08395 | 2.166263 | 0.53742143 | 16.7534563 | 0.00023016 | 0.00192578 | MNS1 | X | 667644 | 669293 | 1 |
| YJR132W | 86.3285409 | 0.69712272 | 0.61731325 | 9.32599758 | 0.00943812 | 0.03773943 | NMD5 | X | 669522 | 672668 | 1 |
| YJR133W | 14.057757 | 1.17634622 | 0.83027614 | 2.33425975 | 0.31125901 | 0.48393923 | XPT1 | X | 672991 | 673620 | 1 |
| YJR134C | 40.7222961 | -0.064394 | 0.84573439 | 6.85023361 | 0.03254548 | 0.09507484 | SGM1 | X | 673729 | 675852 | -1 |
| YJR135C | 11.8983939 | -0.5107942 | 0.91562534 | 0.4007655 | 0.81841744 | 0.8908887 | MCM22 | X | 676059 | 676778 | -1 |
| YJR135W-A | 6.73770371 | -1.4812478 | 1.12217745 | 2.48927476 | 0.28804534 | 0.46186168 | TIM8 | X | 676971 | 677234 | 1 |
| YJR136C | 33.3992789 | 0.26864995 | 0.66335258 | 0.17248593 | 0.91737131 | 0.94883528 | TTI2 | X | 677441 | 678706 | -1 |
| YJR137C | 33.0280823 | -0.563225 | 0.64000707 | 0.90317324 | 0.63661728 | 0.77291131 | MET5 | X | 678957 | 683285 | -1 |
| YJR138W | 114.073731 | 1.44657193 | 0.664968 | 5.36582491 | 0.06836376 | 0.16534092 | IML1 | X | 684567 | 689321 | 1 |
| YJR139C | 70.5652479 | 0.13293963 | 0.70765401 | 7.24839225 | 0.02667053 | 0.08136057 | HOM6 | X | 689445 | 690524 | -1 |
| YJR140C | 30.1369695 | 0.19140717 | 0.79419807 | 1.65617569 | 0.43688388 | 0.60588207 | HIR3 | X | 690750 | 695696 | -1 |
| YJR141W | 41.7615171 | 0.51140545 | 0.63538528 | 0.81538595 | 0.66518307 | 0.79181949 | IPA1 | X | 695906 | 696949 | 1 |
| YJR142W | 75.8418714 | 0.46288534 | 0.61000003 | 1.42963525 | 0.48928134 | 0.65095104 | NA | X | 697141 | 698169 | 1 |
| YJR143C | 51.7118877 | -0.3184062 | 0.59286261 | 0.80301426 | 0.66931055 | 0.79493499 | PMT4 | X | 698326 | 700614 | -1 |
| YJR144W | 16.3314162 | -1.2640454 | 0.84416267 | 2.36075005 | 0.30716352 | 0.4798912 | MGM101 | X | 700882 | 701691 | 1 |
| YJR145C | 103.381087 | -3.0275876 | 0.80402369 | 12.8956195 | 0.00158399 | 0.00951154 | RPS4A | X | 702027 | 703068 | -1 |
| YJR146W | 597.333776 | -1.1306647 | 0.64413518 | 3.28943858 | 0.19306675 | 0.34965796 | NA | X | 703885 | 704238 | 1 |
| YJR147W | 1543.38296 | -1.3296934 | 0.58334125 | 5.23885643 | 0.0728445 | 0.17363922 | HMS2 | X | 704196 | 705272 | 1 |

| | | | | | | | | | | | |
|---|---|---|---|---|---|---|---|---|---|---|---|
| YJR148W | 63.1141373 | 1.40326002 | 0.62111289 | 5.35150016 | 0.06885516 | 0.16625162 | BAT2 | X | 705744 | 706874 | 1 |
| YJR149W | 7.64958468 | 1.46770369 | 1.05788194 | 2.44279195 | 0.29481832 | 0.46908438 | NA | X | 707160 | 708374 | 1 |
| YJR150C | 20.8033201 | 0.02572509 | 0.81999559 | 4.59562132 | 0.10047858 | 0.2196191 | DAN1 | X | 708811 | 709707 | -1 |
| YJR151C | 85.5431181 | -1.0209293 | 0.55282439 | 4.12241331 | 0.12730027 | 0.26090922 | DAN4 | X | 712255 | 715740 | -1 |
| YJR151W-A | 36.0707463 | 1.94153906 | 0.75016678 | 7.11261061 | 0.02854409 | 0.08567667 | NA | X | 717580 | 717630 | 1 |
| YJR152W | 19.9763271 | -0.7041449 | 1.00397915 | 1.60495965 | 0.44821609 | 0.61496946 | DAL5 | X | 719666 | 721297 | 1 |
| YJR153W | 780.033524 | 0.12188105 | 0.54849094 | 0.15761856 | 0.92421617 | 0.95251186 | PGU1 | X | 722815 | 723900 | 1 |
| YJR154W | 12.8983937 | 0.38737881 | 0.83463487 | 0.59410642 | 0.74300447 | 0.84319794 | NA | X | 725784 | 726824 | 1 |
| YJR155W | 29.1104515 | 0.14686038 | 0.64658372 | 1.39852758 | 0.49695103 | 0.65655356 | AAD10 | X | 727405 | 728271 | 1 |
| YJR158W | 14.3882458 | 0.84402426 | 0.83491922 | 3.13336633 | 0.20873638 | 0.37130066 | HXT16 | X | 732440 | 734143 | 1 |
| YJR159W | 226.440232 | 2.31359501 | 0.58373747 | 14.3995489 | 0.00074675 | 0.00515955 | SOR1 | X | 736044 | 737117 | 1 |
| YJR160C | 351.180581 | 1.90139697 | 0.62270998 | 8.81731858 | 0.01217149 | 0.04561353 | MPH3 | X | 738008 | 739816 | -1 |
| YJR161C | 23.499939 | -1.0151213 | 0.72514751 | 2.03991887 | 0.36060957 | 0.53345156 | COS5 | X | 742848 | 743999 | -1 |
| YKL001C | 93.3982281 | 1.4837251 | 1.06674079 | 17.4440476 | 0.00016296 | 0.00142526 | MET14 | XI | 438777 | 439385 | -1 |
| YKL002W | 100.06472 | -1.5590003 | 0.54062 | 9.07399827 | 0.01070548 | 0.04143366 | DID4 | XI | 437778 | 438544 | 1 |
| YKL003C | 17.8148887 | -0.4480232 | 1.02307226 | 1.32655883 | 0.51515914 | 0.67209811 | MRP17 | XI | 437097 | 437492 | -1 |
| YKL004W | 254.887325 | 3.98022389 | 0.57969169 | 44.3485583 | 2.34E-10 | 1.01E-08 | AUR1 | XI | 435580 | 436785 | 1 |
| YKL005C | 43.3104528 | -0.4247251 | 0.58459656 | 0.59081701 | 0.7442275 | 0.84379317 | BYE1 | XI | 433093 | 434877 | -1 |
| YKL006C-A | 1270.7579 | 0.53345982 | 0.6669563 | 12.1708199 | 0.00227583 | 0.01230351 | SFT1 | XI | 430198 | 430632 | -1 |
| YKL006W | 51.0903532 | -2.7732816 | 0.77523962 | 12.260768 | 0.00217575 | 0.01190696 | RPL14A | XI | 431906 | 432720 | 1 |
| YKL007W | 42.3518355 | -1.1595501 | 0.70071643 | 8.66977691 | 0.01310334 | 0.04820096 | CAP1 | XI | 429302 | 430108 | 1 |
| YKL008C | 139.71816 | -0.9054977 | 0.49453997 | 3.48841837 | 0.17478316 | 0.32676605 | LAC1 | XI | 427295 | 428551 | -1 |
| YKL009W | 44.513419 | -3.2063908 | 0.81742499 | 14.4091126 | 0.00074319 | 0.00514108 | MRT4 | XI | 426242 | 426952 | 1 |
| YKL010C | 86.4119865 | 1.22781537 | 0.60045137 | 4.6105826 | 0.09972974 | 0.2191405 | UFD4 | XI | 421424 | 425875 | -1 |
| YKL011C | 192.985953 | 0.73150142 | 0.56110651 | 5.67982383 | 0.05843081 | 0.14688185 | CCE1 | XI | 420155 | 421216 | -1 |
| YKL012W | 88.5743234 | 0.89489331 | 0.54126861 | 2.73320731 | 0.25497146 | 0.42716573 | PRP40 | XI | 418310 | 420061 | 1 |
| YKL013C | 33.7582319 | 1.67103922 | 0.70963388 | 5.58345857 | 0.06131509 | 0.15223601 | ARC19 | XI | 417508 | 418023 | -1 |
| YKL014C | 47.0224955 | -1.608989 | 0.72906814 | 7.63129374 | 0.02202346 | 0.07124193 | URB1 | XI | 411619 | 416913 | -1 |
| YKL015W | 32.312395 | -1.4583287 | 0.70511927 | 4.30397286 | 0.116253 | 0.24370198 | PUT3 | XI | 408544 | 411483 | 1 |
| YKL016C | 17.2837261 | 0.49845396 | 0.7559199 | 0.47287353 | 0.7894358 | 0.87229643 | ATP7 | XI | 407465 | 407989 | -1 |
| YKL017C | 159.276209 | -0.9174009 | 0.71309736 | 6.77044552 | 0.0338701 | 0.09820123 | HCS1 | XI | 405190 | 407241 | -1 |
| YKL018C-A | 246.376603 | 0.34364627 | 0.85226363 | 8.1139634 | 0.01730116 | 0.05948558 | NA | XI | 403580 | 403879 | -1 |
| YKL018W | 65.0205263 | -0.3046822 | 0.69883515 | 1.46805352 | 0.47997236 | 0.64284986 | SWD2 | XI | 404102 | 405091 | 1 |

| | | | | | | | | | | | |
|---|---|---|---|---|---|---|---|---|---|---|---|
| YKL019W | 156.74458 | -0.7390018 | 0.72382697 | 12.8849968 | 0.00159242 | 0.00954465 | RAM2 | XI | 402568 | 403518 | 1 |
| YKL020C | 25.3439001 | -0.0445515 | 0.68115993 | 0.4302067 | 0.80645809 | 0.88416839 | SPT23 | XI | 398832 | 402080 | -1 |
| YKL021C | 29.6740585 | -1.6789592 | 0.69252806 | 5.84401993 | 0.05382539 | 0.13826487 | NA | XI | 397344 | 398750 | -1 |
| YKL022C | 496.310355 | -0.5324842 | 0.45626256 | 2.47572623 | 0.29000326 | 0.46384499 | CDC16 | XI | 394726 | 397248 | -1 |
| YKL023C-A | 100.395794 | 3.07794721 | 0.62137016 | 29.1644113 | 4.65E-07 | 9.09E-06 | NA | XI | 393146 | 393373 | -1 |
| YKL023W | 255.415489 | -0.7380714 | 0.53833355 | 2.34111563 | 0.31019386 | 0.4830878 | NA | XI | 393721 | 394554 | 1 |
| YKL024C | 5.35595216 | -0.2964598 | 1.2973416 | 0.11572758 | 0.94377849 | 0.96197675 | URA6 | XI | 392526 | 393140 | -1 |
| YKL025C | 32.6859159 | 0.90029582 | 0.66687743 | 2.83602626 | 0.24219475 | 0.41135453 | PAN3 | XI | 390240 | 392279 | -1 |
| YKL026C | 10.257265 | -0.6832835 | 0.92342225 | 0.77295509 | 0.67944598 | 0.80269174 | GPX1 | XI | 389380 | 389883 | -1 |
| YKL027W | 1179.17709 | 5.27661855 | 0.51216133 | 107.482661 | 4.58E-24 | 2.94E-21 | TCD2 | XI | 387562 | 388905 | 1 |
| YKL028W | 54.9678836 | -0.4814044 | 0.60066345 | 1.54539571 | 0.46176561 | 0.62532091 | TFA1 | XI | 385758 | 387206 | 1 |
| YKL029C | 26.539856 | 0.00647123 | 0.71443361 | 1.50709423 | 0.47069398 | 0.63296713 | MAE1 | XI | 382716 | 384725 | -1 |
| YKL032C | 99.9374448 | 0.18351077 | 0.59616806 | 0.60545556 | 0.73880019 | 0.84040603 | IXR1 | XI | 380065 | 381858 | -1 |
| YKL033W | 118.584921 | 1.91448696 | 0.53195897 | 19.8550422 | 4.88E-05 | 0.00050469 | TTI1 | XI | 375456 | 378572 | 1 |
| YKL033W-A | 140.985854 | 1.74773422 | 0.50793729 | 12.1478111 | 0.00230216 | 0.01239956 | NA | XI | 374505 | 375215 | 1 |
| YKL034W | 50.3273129 | -1.2623212 | 0.63031039 | 4.03038954 | 0.13329444 | 0.26928639 | TUL1 | XI | 371829 | 374105 | 1 |
| YKL035W | 39.0289375 | -0.3303182 | 0.73544834 | 0.2276668 | 0.89240662 | 0.93387025 | UGP1 | XI | 369891 | 371390 | 1 |
| YKL037W | 195.938968 | 1.21277849 | 0.5200483 | 6.22827406 | 0.04441682 | 0.11983849 | AIM26 | XI | 369365 | 369721 | 1 |
| YKL038W | 24.448692 | 0.94770041 | 0.69862573 | 2.25198072 | 0.3243311 | 0.49763242 | RGT1 | XI | 365605 | 369117 | 1 |
| YKL039W | 39.6116109 | 0.38444319 | 0.76377519 | 2.92324815 | 0.23185941 | 0.39906837 | PTM1 | XI | 362622 | 364193 | 1 |
| YKL040C | 35.6615512 | -0.5799505 | 0.62871289 | 2.51887064 | 0.28381425 | 0.45748454 | NFU1 | XI | 361058 | 361828 | -1 |
| YKL041W | 552.459265 | -1.8957471 | 0.73964992 | 9.30548646 | 0.00953541 | 0.03806203 | VPS24 | XI | 360143 | 360817 | 1 |
| YKL042W | 10.4765366 | 0.99381726 | 0.89155316 | 3.97615931 | 0.13695818 | 0.27458028 | SPC42 | XI | 358475 | 359566 | 1 |
| YKL043W | 26.6077503 | 0.18681502 | 0.85626876 | 0.06617008 | 0.96745628 | 0.97486964 | PHD1 | XI | 356748 | 357848 | 1 |
| YKL044W | 3.79727751 | 1.29375589 | 1.41302516 | 2.06405392 | 0.35628406 | 0.52948785 | MMO1 | XI | 356322 | 356642 | 1 |
| YKL045W | 234.737412 | -0.646219 | 0.55542612 | 2.49101028 | 0.28779549 | 0.46172289 | PRI2 | XI | 353493 | 355079 | 1 |
| YKL046C | 52.1280153 | 1.58630582 | 0.86353464 | 3.32147462 | 0.18999884 | 0.34583253 | DCW1 | XI | 351275 | 352624 | -1 |
| YKL047W | 49.8150124 | 0.24693798 | 0.63957687 | 1.25051813 | 0.53512278 | 0.68975087 | ANR2 | XI | 349463 | 351013 | 1 |
| YKL048C | 24.9427364 | 1.14993737 | 0.70571818 | 2.99390703 | 0.22381096 | 0.38938265 | ELM1 | XI | 347214 | 349136 | -1 |
| YKL049C | 8.62966844 | 1.31147932 | 1.0756362 | 1.71569038 | 0.4240749 | 0.5946703 | CSE4 | XI | 346075 | 346764 | -1 |
| YKL050C | 42.0073784 | -0.6750486 | 0.65864983 | 1.27540342 | 0.52850569 | 0.68289398 | NA | XI | 342852 | 345620 | -1 |
| YKL051W | 318.044893 | 0.61612852 | 0.49846701 | 2.91425693 | 0.23290411 | 0.40027153 | SFK1 | XI | 341312 | 342373 | 1 |
| YKL052C | 152.980799 | -0.6131083 | 0.4990294 | 1.62387919 | 0.44399606 | 0.61165577 | ASK1 | XI | 339668 | 340546 | -1 |

| | | | | | | | | | | | |
|---|---|---|---|---|---|---|---|---|---|---|---|
| YKL053C-A | 340.02936 | 3.33479493 | 0.49627637 | 39.2024545 | 3.07E-09 | 1.03E-07 | MDM35 | XI | 339182 | 339442 | -1 |
| YKL054C | 1654.29956 | -0.3872587 | 0.89151222 | 9.59198834 | 0.00826278 | 0.03401477 | DEF1 | XI | 336537 | 338753 | -1 |
| YKL055C | 48.2098984 | -1.8414429 | 0.61875386 | 8.99323396 | 0.01114664 | 0.04268456 | OAR1 | XI | 335321 | 336157 | -1 |
| YKL056C | 85.7976123 | -2.3489459 | 0.85852719 | 7.08402943 | 0.02895493 | 0.08664034 | TMA19 | XI | 334412 | 334915 | -1 |
| YKL057C | 43.0588934 | -1.1879835 | 0.78420288 | 8.68274432 | 0.01301865 | 0.04801146 | NUP120 | XI | 330856 | 333969 | -1 |
| YKL058W | 98.8202612 | 0.31345936 | 0.55213362 | 2.10582806 | 0.34891951 | 0.52261068 | TOA2 | XI | 330166 | 330534 | 1 |
| YKL059C | 17.2163274 | 0.11280012 | 0.79519395 | 0.15617945 | 0.92488144 | 0.95277599 | MPE1 | XI | 328118 | 329443 | -1 |
| YKL060C | 701.900665 | -1.853402 | 0.74735143 | 6.52867875 | 0.03822218 | 0.10758697 | FBA1 | XI | 326408 | 327487 | -1 |
| YKL061W | 49.8142281 | 0.3998912 | 0.58283114 | 0.48659589 | 0.78403788 | 0.86949041 | BLI1 | XI | 325771 | 326112 | 1 |
| YKL062W | 35.7656423 | -1.0663253 | 0.7133534 | 2.31432721 | 0.31437661 | 0.48643522 | MSN4 | XI | 323228 | 325120 | 1 |
| YKL063C | 67.9059538 | -2.0227193 | 0.59507527 | 13.2471959 | 0.00132864 | 0.00828077 | NA | XI | 321015 | 321518 | -1 |
| YKL064W | 162.276258 | -0.1699399 | 0.4939715 | 0.56482134 | 0.75396399 | 0.85146314 | MNR2 | XI | 317764 | 320673 | 1 |
| YKL065C | 91.5077303 | 1.8659989 | 0.57160256 | 10.2210433 | 0.00603293 | 0.026624 | YET1 | XI | 316437 | 317057 | -1 |
| YKL065W-A | 75.5022367 | 1.75204861 | 0.78582094 | 8.07303161 | 0.01765889 | 0.06020484 | NA | XI | 315954 | 316175 | 1 |
| YKL067W | 89.1332694 | 2.01724016 | 0.62892366 | 11.8621522 | 0.00265562 | 0.01388984 | YNK1 | XI | 314812 | 315273 | 1 |
| YKL068W | 54.3222013 | 0.55138281 | 0.57629771 | 2.21142144 | 0.33097557 | 0.50430225 | NUP100 | XI | 310199 | 313078 | 1 |
| YKL068W-A | 201.976807 | 0.50120486 | 0.5112695 | 1.32014029 | 0.51681508 | 0.67350835 | NA | XI | 309206 | 309442 | 1 |
| YKL069W | 22.8156411 | -0.4689323 | 0.7187945 | 0.48590662 | 0.78430813 | 0.86949041 | NA | XI | 307285 | 307827 | 1 |
| YKL070W | 1613.72921 | -0.6089079 | 0.48173827 | 2.31268792 | 0.3146344 | 0.48657403 | NA | XI | 306211 | 306720 | 1 |
| YKL071W | 1114.3498 | -2.1545418 | 0.4879296 | 18.7138072 | 8.64E-05 | 0.00082519 | NA | XI | 305114 | 305884 | 1 |
| YKL072W | 42.0245934 | 0.81709374 | 0.5924804 | 2.02110684 | 0.36401747 | 0.53712058 | STB6 | XI | 299582 | 301882 | 1 |
| YKL073W | 58.2412234 | -0.1467466 | 0.64828296 | 0.22264047 | 0.89465221 | 0.9350658 | LHS1 | XI | 296430 | 299075 | 1 |
| YKL074C | 247.868265 | 2.70873523 | 0.54551649 | 24.9607313 | 3.80E-06 | 5.66E-05 | MUD2 | XI | 294610 | 296193 | -1 |
| YKL075C | 16.0987689 | 0.212382 | 0.7746799 | 0.21557346 | 0.89781906 | 0.93705725 | NA | XI | 292956 | 294308 | -1 |
| YKL077W | 62.034363 | -1.7315569 | 0.57598331 | 8.8087484 | 0.01222375 | 0.04577978 | NA | XI | 291453 | 292631 | 1 |
| YKL078W | 12.0498656 | 1.15392964 | 0.90860915 | 2.29938058 | 0.31673485 | 0.48903861 | DHR2 | XI | 288845 | 291052 | 1 |
| YKL079W | 39.8644475 | 2.33260329 | 0.76198138 | 9.10291801 | 0.0105518 | 0.04108602 | SMY1 | XI | 286603 | 288573 | 1 |
| YKL080W | 31.5482334 | -0.140014 | 0.69447275 | 1.01699308 | 0.60139908 | 0.74720908 | VMA5 | XI | 285030 | 286208 | 1 |
| YKL081W | 259.591801 | -3.1572318 | 0.63175083 | 22.0607639 | 1.62E-05 | 0.00019875 | TEF4 | XI | 282891 | 284455 | 1 |
| YKL082C | 185.362814 | -1.2902615 | 0.78195269 | 4.64157702 | 0.09819613 | 0.21634535 | RRP14 | XI | 281025 | 282329 | -1 |
| YKL084W | 15.0554161 | -0.4640808 | 0.77472614 | 0.53650643 | 0.76471412 | 0.85692899 | HOT13 | XI | 280509 | 280859 | 1 |
| YKL085W | 39.0998012 | 0.94351961 | 0.64666145 | 6.42403399 | 0.0402753 | 0.11200479 | MDH1 | XI | 279123 | 280127 | 1 |
| YKL086W | 12.39688 | 5.31521739 | 1.31857037 | 30.8417139 | 2.01E-07 | 4.39E-06 | SRX1 | XI | 278281 | 278664 | 1 |

| | | | | | | | | | | | |
|---|---|---|---|---|---|---|---|---|---|---|---|
| YKL087C | 95.2933564 | 0.06746097 | 0.71582017 | 1.85314148 | 0.39590906 | 0.56712357 | CYT2 | XI | 277189 | 277863 | -1 |
| YKL088W | 133.476211 | -0.4436682 | 0.52846388 | 1.37101797 | 0.50383372 | 0.66239719 | CAB3 | XI | 275283 | 276998 | 1 |
| YKL089W | 16.8083708 | 0.07589384 | 0.76556768 | 0.02718395 | 0.98649998 | 0.98889108 | MIF2 | XI | 273394 | 275043 | 1 |
| YKL090W | 922.955746 | 3.01475184 | 0.52381371 | 42.6273681 | 5.54E-10 | 2.12E-08 | CUE2 | XI | 271878 | 273209 | 1 |
| YKL091C | 33.6886452 | 1.1525804 | 0.71859436 | 2.70059014 | 0.25916378 | 0.43082351 | NA | XI | 269718 | 270650 | -1 |
| YKL092C | 34.8173936 | -1.3943666 | 0.66687542 | 4.7665354 | 0.09224864 | 0.20670265 | BUD2 | XI | 266145 | 269459 | -1 |
| YKL093W | 978.232845 | -0.6588901 | 0.52986327 | 2.52476853 | 0.28297853 | 0.45678444 | MBR1 | XI | 264789 | 265808 | 1 |
| YKL094W | 33.2746764 | 0.61601147 | 0.65654017 | 2.20348232 | 0.33229201 | 0.50604175 | YJU3 | XI | 263349 | 264290 | 1 |
| YKL095W | 11.3325143 | 1.30552937 | 0.89610729 | 2.12937352 | 0.34483585 | 0.51822506 | YJU2 | XI | 262277 | 263113 | 1 |
| YKL096C-B | 7435.67403 | 0.34634618 | 0.50089028 | 0.84448905 | 0.65557372 | 0.78495521 | NA | XI | 259073 | 259222 | -1 |
| YKL096W | 43.3363155 | -1.5535754 | 0.70037952 | 7.99127368 | 0.01839573 | 0.06210569 | CWP1 | XI | 261132 | 261851 | 1 |
| YKL096W-A | 129.678295 | 0.29168638 | 0.53812129 | 4.49266867 | 0.10578629 | 0.22837533 | CWP2 | XI | 259253 | 259531 | 1 |
| YKL097C | 10.7542647 | 2.62689555 | 0.98819124 | 13.8668632 | 0.00097465 | 0.0064494 | NA | XI | 258503 | 258913 | -1 |
| YKL098W | 25.000075 | 2.01954072 | 0.73275267 | 10.499337 | 0.00524926 | 0.02385369 | MTC2 | XI | 256770 | 257843 | 1 |
| YKL099C | 10.9280332 | -2.847155 | 0.96617446 | 9.36477008 | 0.00925691 | 0.03719466 | UTP11 | XI | 255720 | 256472 | -1 |
| YKL100C | 18.7510872 | -0.4787696 | 0.93773187 | 0.61821 | 0.73410369 | 0.83775415 | YPF1 | XI | 253697 | 255460 | -1 |
| YKL101W | 349.959848 | -0.0508017 | 0.6127447 | 5.86702695 | 0.05320976 | 0.13698733 | HSL1 | XI | 248920 | 253476 | 1 |
| YKL102C | 879.324165 | 1.50769906 | 0.4943079 | 33.2337381 | 6.07E-08 | 1.55E-06 | NA | XI | 248062 | 248367 | -1 |
| YKL103C | 2186.15313 | 1.43500361 | 0.49612474 | 20.0201564 | 4.49E-05 | 0.00046888 | APE1 | XI | 246138 | 247682 | -1 |
| YKL104C | 163.242181 | 1.52745996 | 0.57752904 | 8.45349495 | 0.0145998 | 0.05250487 | GFA1 | XI | 243220 | 245373 | -1 |
| YKL104W-A | 18547.0612 | 1.37647288 | 0.48473372 | 12.9412672 | 0.00154824 | 0.00935734 | NA | NA | NA | NA | NA |
| YKL105C | 1664.78033 | 0.15174695 | 0.57325108 | 1.58909246 | 0.4517862 | 0.61806282 | SEG2 | XI | 239185 | 242583 | -1 |
| YKL106C-A | 38.9752528 | 3.99345295 | 0.76485688 | 27.1865065 | 1.25E-06 | 2.13E-05 | NA | XI | 237147 | 237266 | -1 |
| YKL106W | 115.550022 | 2.91062016 | 0.76255795 | 15.101156 | 0.00052581 | 0.00381506 | AAT1 | XI | 237536 | 238891 | 1 |
| YKL107W | 167.171718 | -0.0249726 | 0.49874069 | 0.7493911 | 0.68749856 | 0.80989148 | NA | XI | 236141 | 237070 | 1 |
| YKL108W | 33.8109918 | 2.74024062 | 0.72964753 | 18.5539955 | 9.36E-05 | 0.00088415 | SLD2 | XI | 234426 | 235787 | 1 |
| YKL109W | 109.415335 | 0.02293213 | 0.6071386 | 10.6783112 | 0.00479992 | 0.02212703 | HAP4 | XI | 232227 | 233891 | 1 |
| YKL110C | 11.0140622 | 0.1757251 | 0.89853214 | 0.29790132 | 0.86161163 | 0.91873505 | KTI12 | XI | 228939 | 229880 | -1 |
| YKL112W | 59.0976138 | -0.2795121 | 0.56193711 | 0.25154817 | 0.88181404 | 0.9300006 | ABF1 | XI | 226570 | 228765 | 1 |
| YKL113C | 162.026644 | 0.84942161 | 0.50913251 | 3.27887222 | 0.19408946 | 0.35096126 | RAD27 | XI | 224727 | 225875 | -1 |
| YKL114C | 274.778156 | 1.891959 | 0.54769229 | 15.345871 | 0.00046525 | 0.00346692 | APN1 | XI | 223352 | 224455 | -1 |
| YKL116C | 65.8837633 | 1.46954493 | 0.59497696 | 7.41292302 | 0.02456429 | 0.07694787 | PRR1 | XI | 221344 | 222900 | -1 |
| YKL117W | 86.3725139 | -0.2058203 | 0.53956523 | 0.8818227 | 0.64344975 | 0.77686719 | SBA1 | XI | 220324 | 220974 | 1 |

| | | | | | | | | | | | | |
|---|---|---|---|---|---|---|---|---|---|---|---|---|
| YKL119C | 5.37406981 | -0.8334013 | 1.26519397 | 1.21415906 | 0.54494003 | 0.69774497 | VPH2 | XI | | 218570 | 219217 | -1 |
| YKL120W | 13.0325104 | -0.1617477 | 0.81210379 | 0.5270828 | 0.76832581 | 0.859302 | OAC1 | XI | | 217344 | 218318 | 1 |
| YKL121W | 87.3362291 | 1.09083135 | 0.54091956 | 6.67338925 | 0.03555428 | 0.10170914 | DGR2 | XI | | 214142 | 216700 | 1 |
| YKL122C | 41.1769972 | -0.7155833 | 0.73034501 | 3.43914898 | 0.17914236 | 0.3333015 | SRP21 | XI | | 212495 | 212998 | -1 |
| YKL124W | 106.862488 | 0.77647978 | 0.52781982 | 7.75203389 | 0.02073324 | 0.06786064 | SSH4 | XI | | 210593 | 212332 | 1 |
| YKL125W | 50.6569691 | 1.10288094 | 0.628652 | 3.62752415 | 0.16303961 | 0.31175673 | RRN3 | XI | | 208247 | 210130 | 1 |
| YKL126W | 27.1494538 | 0.86440712 | 0.70552405 | 2.16047983 | 0.33951406 | 0.51259098 | YPK1 | XI | | 205707 | 207749 | 1 |
| YKL127W | 252.717239 | 0.17096866 | 0.49625914 | 6.63239267 | 0.03629061 | 0.1034068 | PGM1 | XI | | 203541 | 205253 | 1 |
| YKL128C | 684.588182 | 2.02843994 | 0.54458118 | 13.2969469 | 0.001296 | 0.00812103 | PMU1 | XI | | 200884 | 201771 | -1 |
| YKL129C | 160.925232 | -0.9233198 | 0.49844978 | 6.76767824 | 0.03391699 | 0.09824187 | MYO3 | XI | | 196349 | 200167 | -1 |
| YKL130C | 73.9642489 | -0.1403795 | 0.62509572 | 0.25436246 | 0.88057407 | 0.92970895 | SHE2 | XI | | 195288 | 196028 | -1 |
| YKL132C | 22.3893949 | 1.09663087 | 0.70155846 | 2.56000038 | 0.27803725 | 0.45261809 | RMA1 | XI | | 193574 | 194866 | -1 |
| YKL133C | 58.6644849 | 0.35405508 | 0.55301071 | 1.95435524 | 0.37637187 | 0.54998972 | NA | XI | | 191679 | 193070 | -1 |
| YKL134C | 140.398003 | -0.7999708 | 0.50382377 | 3.41875732 | 0.18097821 | 0.33553116 | 1-Oct | XI | | 189124 | 191442 | -1 |
| YKL135C | 153.693717 | -1.4762473 | 0.64901327 | 9.75272074 | 0.00762471 | 0.03199065 | APL2 | XI | | 186460 | 188640 | -1 |
| YKL137W | 35.7339311 | -0.8108808 | 0.61552174 | 1.72400035 | 0.42231653 | 0.59293772 | CMC1 | XI | | 185957 | 186292 | 1 |
| YKL138C | 17.3537142 | -0.0290038 | 0.9441784 | 1.71141747 | 0.42498188 | 0.59510585 | MRPL31 | XI | | 185286 | 185681 | -1 |
| YKL138C-A | 29.1878324 | 0.06139908 | 0.63032549 | 0.80053615 | 0.67014038 | 0.79536351 | HSK3 | XI | | 184803 | 185012 | -1 |
| YKL139W | 238.422738 | 0.89831395 | 0.84275502 | 8.44919768 | 0.0146312 | 0.05258366 | CTK1 | XI | | 182958 | 184544 | 1 |
| YKL140W | 44.7127721 | 1.65462531 | 0.59739644 | 9.66820143 | 0.00795384 | 0.03308385 | TGL1 | XI | | 180779 | 182425 | 1 |
| YKL141W | 32.827867 | 1.48372187 | 0.69090293 | 4.83015365 | 0.08936047 | 0.20267482 | SDH3 | XI | | 179667 | 180263 | 1 |
| YKL142W | 24.4588257 | 1.00526762 | 0.77286219 | 3.5157943 | 0.17240703 | 0.32378745 | MRP8 | XI | | 178515 | 179174 | 1 |
| YKL143W | 28.0490197 | -1.7221304 | 0.72852787 | 5.92419277 | 0.0517104 | 0.13420135 | LTV1 | XI | | 176781 | 178172 | 1 |
| YKL144C | 277.927894 | -1.8718063 | 0.51239506 | 12.5760815 | 0.0018584 | 0.01061156 | RPC25 | XI | | 175843 | 176481 | -1 |
| YKL145W | 114.279614 | -0.2632042 | 0.76677704 | 0.68126618 | 0.71131985 | 0.82438173 | RPT1 | XI | | 174213 | 175616 | 1 |
| YKL146W | 69.8992525 | 0.36242678 | 0.64908796 | 0.4777698 | 0.78750552 | 0.87066201 | AVT3 | XI | | 171783 | 173861 | 1 |
| YKL148C | 24.9651322 | 0.08699504 | 0.87274948 | 0.04482207 | 0.97783823 | 0.98293113 | SDH1 | XI | | 169207 | 171129 | -1 |
| YKL149C | 50.9182633 | 3.38554767 | 0.7087941 | 22.698299 | 1.18E-05 | 0.00015292 | DBR1 | XI | | 167612 | 168829 | -1 |
| YKL150W | 50.1128829 | 1.968685 | 0.59775478 | 13.9835791 | 0.0009194 | 0.00615027 | MCR1 | XI | | 166544 | 167452 | 1 |
| YKL151C | 616.280956 | -1.4589898 | 0.55706799 | 6.73998323 | 0.03438993 | 0.09926596 | NNR2 | XI | | 164917 | 165930 | -1 |
| YKL152C | 433.030351 | 0.48908031 | 0.57307172 | 1.26494718 | 0.53127602 | 0.68570846 | GPM1 | XI | | 163642 | 164385 | -1 |
| YKL154W | 12.9152514 | -1.9244343 | 1.31862265 | 8.63766939 | 0.01331539 | 0.04876415 | SRP102 | XI | | 161601 | 162335 | 1 |
| YKL155C | 55.0812062 | 1.90499605 | 0.60875634 | 15.6030882 | 0.0004091 | 0.00312907 | RSM22 | XI | | 159456 | 161342 | -1 |

| | | | | | | | | | | | |
|---|---|---|---|---|---|---|---|---|---|---|---|
| YKL156W | 12.8000601 | -1.9681781 | 0.86771873 | 5.01464042 | 0.08148631 | 0.1887223 | RPS27A | XI | 158613 | 159212 | 1 |
| YKL157W | 80.7514138 | 0.05157454 | 0.54812856 | 0.59869029 | 0.74130351 | 0.84242342 | APE2 | XI | 154991 | 158232 | 1 |
| YKL159C | 340.94678 | -2.3732644 | 0.50376829 | 20.32271 | 3.86E-05 | 0.00041734 | RCN1 | XI | 153816 | 154451 | -1 |
| YKL160W | 182.560507 | -0.1586666 | 0.55298332 | 15.3679378 | 0.00046014 | 0.0034333 | ELF1 | XI | 153269 | 153706 | 1 |
| YKL161C | 39.3253805 | 0.2536799 | 0.67341283 | 0.54686612 | 0.76076326 | 0.85483951 | KDX1 | XI | 149386 | 150687 | -1 |
| YKL163W | 94.9762866 | 2.08622535 | 0.81162953 | 6.49172319 | 0.038935 | 0.10943382 | PIR3 | XI | 144401 | 145378 | 1 |
| YKL164C | 195.908905 | 1.10870381 | 0.49532294 | 6.91371965 | 0.03152861 | 0.09285753 | PIR1 | XI | 141794 | 142819 | -1 |
| YKL165C | 77.2701226 | 1.69062576 | 0.59304745 | 8.20801748 | 0.01650637 | 0.05733167 | MCD4 | XI | 137932 | 140691 | -1 |
| YKL166C | 17.8872574 | -0.1895277 | 0.75876021 | 0.39779448 | 0.81963412 | 0.89171017 | TPK3 | XI | 134509 | 135705 | -1 |
| YKL167C | 6.41677326 | 1.08363489 | 1.10312426 | 1.05628848 | 0.58969829 | 0.73728204 | MRP49 | XI | 133721 | 134134 | -1 |
| YKL168C | 22.093188 | 0.74496121 | 0.70750268 | 1.50462394 | 0.47127572 | 0.63325282 | KKQ8 | XI | 131288 | 133462 | -1 |
| YKL170W | 99.5446691 | -0.3223522 | 0.55128472 | 0.72140966 | 0.69718475 | 0.816815 | MRPL38 | XI | 130634 | 131050 | 1 |
| YKL171W | 413.976702 | 0.80747179 | 0.50155753 | 3.84703004 | 0.14609254 | 0.28741957 | NNK1 | XI | 127475 | 130261 | 1 |
| YKL172W | 95.8421433 | -2.8510738 | 0.56859361 | 23.1086106 | 9.59E-06 | 0.00012771 | EBP2 | XI | 125759 | 127042 | 1 |
| YKL173W | 21.8903198 | 0.53944968 | 0.69442947 | 0.62938175 | 0.73001451 | 0.83678999 | SNU114 | XI | 122517 | 125543 | 1 |
| YKL174C | 248.494257 | 0.80608301 | 0.67843735 | 8.54133698 | 0.01397244 | 0.05075309 | TPO5 | XI | 120380 | 122236 | -1 |
| YKL175W | 433.035587 | 1.53002733 | 0.91091828 | 10.1441256 | 0.00626947 | 0.02746253 | ZRT3 | XI | 118793 | 120304 | 1 |
| YKL176C | 183.431732 | -1.0510121 | 0.5023019 | 5.03721076 | 0.0805719 | 0.18735393 | LST4 | XI | 115500 | 117986 | -1 |
| YKL178C | 42.7980751 | 0.45637982 | 0.63131257 | 0.96181617 | 0.61822174 | 0.75949583 | STE3 | XI | 113215 | 114627 | -1 |
| YKL179C | 21.3681606 | 0.0886729 | 0.72332775 | 1.67666057 | 0.43243196 | 0.60114791 | COY1 | XI | 110464 | 112503 | -1 |
| YKL180W | 80.0674227 | -2.6978293 | 0.63899594 | 17.1991865 | 0.00018418 | 0.00158456 | RPL17A | XI | 109269 | 110129 | 1 |
| YKL181W | 56.6891656 | 0.2170949 | 0.58991981 | 0.86470051 | 0.64898203 | 0.7805914 | PRS1 | XI | 107316 | 108599 | 1 |
| YKL182W | 130.609291 | -0.0227126 | 0.71169764 | 0.08443682 | 0.95866038 | 0.97090145 | FAS1 | XI | 100671 | 106826 | 1 |
| YKL183C-A | 601.551244 | -0.2907983 | 0.62994463 | 0.88000448 | 0.64403498 | 0.77686719 | NA | XI | 98395 | 98607 | -1 |
| YKL183W | 21.8277522 | -1.1013932 | 0.75422779 | 3.45017362 | 0.17815759 | 0.33157583 | LOT5 | XI | 98721 | 99641 | 1 |
| YKL184W | 10.8236499 | -0.8464545 | 1.11590566 | 0.81052205 | 0.66680274 | 0.79342126 | SPE1 | XI | 96757 | 98157 | 1 |
| YKL185W | 55.8313792 | 0.08144438 | 0.66143396 | 0.59108314 | 0.74412848 | 0.84379317 | ASH1 | XI | 94499 | 96265 | 1 |
| YKL186C | 53.6163024 | 0.79608682 | 0.70035723 | 1.96330766 | 0.37469091 | 0.54798191 | MTR2 | XI | 92744 | 93298 | -1 |
| YKL187C | 238.343719 | 1.7363063 | 0.52985714 | 10.9729951 | 0.00414233 | 0.01969412 | FAT3 | XI | 89284 | 91536 | -1 |
| YKL188C | 994.381827 | 4.37946605 | 0.49545944 | 64.6574948 | 9.12E-15 | 9.10E-13 | PXA2 | XI | 86225 | 88786 | -1 |
| YKL189W | 19.7740721 | 3.11434001 | 0.84756422 | 14.2507426 | 0.00080443 | 0.00551204 | HYM1 | XI | 84704 | 85903 | 1 |
| YKL190W | 23.381838 | -0.173951 | 0.92896854 | 0.18667932 | 0.91088406 | 0.94533405 | CNB1 | XI | 82947 | 83550 | 1 |
| YKL191W | 37.3742788 | 0.79994817 | 0.70633108 | 6.10332741 | 0.0472802 | 0.12530377 | DPH2 | XI | 81035 | 82639 | 1 |

| | | | | | | | | | | | |
|---|---|---|---|---|---|---|---|---|---|---|---|
| YKL192C | 301.428545 | 4.62897917 | 0.56900799 | 74.1717031 | 7.83E-17 | 1.08E-14 | ACP1 | XI | 80160 | 80537 | -1 |
| YKL193C | 12.2779565 | 1.90343966 | 0.91044664 | 4.25455789 | 0.1191611 | 0.24840182 | SDS22 | XI | 78866 | 79882 | -1 |
| YKL194C | 14.7588775 | -1.2979752 | 0.92889711 | 3.13452167 | 0.20861583 | 0.37120027 | MST1 | XI | 77258 | 78646 | -1 |
| YKL195W | 116.339296 | -0.0924351 | 0.50235375 | 0.67314143 | 0.71421538 | 0.82607012 | MIA40 | XI | 75821 | 77032 | 1 |
| YKL196C | 14.9451767 | -0.7964908 | 0.84788396 | 0.93693527 | 0.62596073 | 0.76430043 | YKT6 | XI | 74932 | 75534 | -1 |
| YKL197C | 58.6378411 | 1.03798136 | 0.57270163 | 5.0624278 | 0.07956238 | 0.18567762 | PEX1 | XI | 70734 | 73865 | -1 |
| YKL198C | 203.987887 | -1.921439 | 0.50827885 | 14.2312941 | 0.00081229 | 0.00555932 | PTK1 | XI | 68232 | 70220 | -1 |
| YKL201C | 1426.57059 | -1.1976291 | 0.68517744 | 4.25277791 | 0.1192672 | 0.24840182 | MNN4 | XI | 63930 | 67466 | -1 |
| YKL203C | 31.4818697 | -0.4934623 | 0.65629218 | 3.36893663 | 0.18554306 | 0.34147943 | TOR2 | XI | 55935 | 63359 | -1 |
| YKL204W | 64.4175739 | 0.45908321 | 0.74752118 | 1.21817013 | 0.54384823 | 0.69714961 | EAP1 | XI | 53704 | 55602 | 1 |
| YKL205W | 23.5982772 | -1.023127 | 0.70177061 | 2.24729226 | 0.3250923 | 0.49807536 | LOS1 | XI | 50051 | 53353 | 1 |
| YKL206C | 147.675493 | -0.0317656 | 0.94128474 | 9.40810165 | 0.00905851 | 0.03655068 | ADD66 | XI | 49006 | 49809 | -1 |
| YKL207W | 177.564364 | 0.08677182 | 0.80448945 | 13.3118183 | 0.0012864 | 0.00807836 | EMC3 | XI | 48194 | 48955 | 1 |
| YKL209C | 107.113163 | -0.8270125 | 0.52908732 | 3.72625902 | 0.15518621 | 0.30010961 | STE6 | XI | 42423 | 46295 | -1 |
| YKL210W | 52.9046751 | -0.3713256 | 0.68386019 | 0.3367829 | 0.84502299 | 0.90911019 | UBA1 | XI | 39163 | 42237 | 1 |
| YKL211C | 37.8750347 | -2.6678716 | 0.74585812 | 12.0328717 | 0.00243834 | 0.01297612 | TRP3 | XI | 36699 | 38153 | -1 |
| YKL212W | 61.7077875 | -1.4163701 | 0.56644607 | 6.80159174 | 0.03334672 | 0.09707265 | SAC1 | XI | 34543 | 36414 | 1 |
| YKL213C | 32.7376414 | 0.14897071 | 0.63483526 | 0.46282253 | 0.7934131 | 0.87452157 | DOA1 | XI | 31960 | 34107 | -1 |
| YKL214C | 32.5350753 | -0.1318707 | 0.64054892 | 0.14582891 | 0.92968035 | 0.95576159 | YRA2 | XI | 31082 | 31693 | -1 |
| YKL215C | 12.2519431 | 0.98012632 | 0.95210799 | 1.29445746 | 0.52349451 | 0.6789949 | OXP1 | XI | 26827 | 30687 | -1 |
| YKL216W | 19.8139908 | -1.4573637 | 0.98584425 | 2.27725649 | 0.32025804 | 0.49277014 | URA1 | XI | 25215 | 26159 | 1 |
| YKL217W | 493.704277 | 1.45141314 | 0.55407359 | 7.63018919 | 0.02203563 | 0.07124193 | JEN1 | XI | 22234 | 24084 | 1 |
| YKL218C | 150.194662 | 0.34814219 | 0.50561741 | 1.6028215 | 0.44869552 | 0.61504428 | SRY1 | XI | 17359 | 18339 | -1 |
| YKL219W | 62.3996282 | -1.3074508 | 0.56376227 | 5.7305696 | 0.0569669 | 0.14462713 | COS9 | XI | 14485 | 15708 | 1 |
| YKL220C | 55.9706533 | -0.290419 | 0.63080181 | 2.42160177 | 0.29795855 | 0.47161883 | FRE2 | XI | 9091 | 11226 | -1 |
| YKL221W | 193.874728 | 1.08823575 | 0.6222083 | 13.0590455 | 0.0014597 | 0.00894319 | MCH2 | XI | 6107 | 7528 | 1 |
| YKL222C | 44.8626822 | 1.32689522 | 0.66324365 | 6.25739567 | 0.04377476 | 0.11860359 | NA | XI | 3503 | 5620 | -1 |
| YKR001C | 68.4676191 | 0.09686589 | 0.58825689 | 2.15107892 | 0.34111369 | 0.51473762 | VPS1 | XI | 440608 | 442722 | -1 |
| YKR002W | 129.234309 | 3.16950467 | 0.56290281 | 34.2338539 | 3.68E-08 | 9.83E-07 | PAP1 | XI | 443232 | 444938 | 1 |
| YKR003W | 172.702423 | -0.7717474 | 0.48409055 | 3.02767896 | 0.22006342 | 0.38460377 | OSH6 | XI | 445381 | 446727 | 1 |
| YKR004C | 609.37646 | -0.0295252 | 0.93006026 | 12.3086527 | 0.00212427 | 0.01170082 | ECM9 | XI | 446799 | 448036 | -1 |
| YKR005C | 34.4496057 | 2.7915417 | 0.75481545 | 12.931382 | 0.00155592 | 0.00937506 | NA | XI | 448521 | 450167 | -1 |
| YKR007W | 31.9795813 | -1.3894819 | 0.64389598 | 4.50846439 | 0.1049541 | 0.22691719 | MEH1 | XI | 451434 | 451988 | 1 |

| | | | | | | | | | | | |
|---|---|---|---|---|---|---|---|---|---|---|---|
| YKR008W | 8.06518715 | -0.8719139 | 1.1850358 | 1.08146535 | 0.58232144 | 0.73074147 | RSC4 | XI | 452201 | 454078 | 1 |
| YKR009C | 1361.33197 | 1.62545967 | 0.51065694 | 10.7715671 | 0.00458125 | 0.02123734 | FOX2 | XI | 454352 | 457054 | -1 |
| YKR010C | 11.7927766 | -0.562924 | 0.83056637 | 0.4983563 | 0.7794411 | 0.8662119 | TOF2 | XI | 458924 | 461239 | -1 |
| YKR011C | 469.801923 | -2.6735952 | 0.49137698 | 27.5834387 | 1.02E-06 | 1.80E-05 | NA | XI | 461635 | 462696 | -1 |
| YKR013W | 392.440991 | 0.06391095 | 0.46662214 | 0.50113292 | 0.77835975 | 0.86534235 | PRY2 | XI | 463959 | 464948 | 1 |
| YKR014C | 46.2561887 | 0.28102349 | 0.64227139 | 1.64588164 | 0.43913833 | 0.60798922 | YPT52 | XI | 465367 | 466071 | -1 |
| YKR015C | 1633.46959 | 0.84014492 | 0.5188432 | 2.65739012 | 0.26482261 | 0.43721783 | NA | XI | 466961 | 468667 | -1 |
| YKR016W | 19.9458496 | 0.60579084 | 0.71509047 | 0.72167514 | 0.69709222 | 0.816815 | MIC60 | XI | 469717 | 471339 | 1 |
| YKR017C | 156.287335 | 0.56988849 | 0.54765922 | 2.59470582 | 0.27325417 | 0.44663164 | HEL1 | XI | 471694 | 473349 | -1 |
| YKR018C | 43.207686 | 0.06305871 | 0.64888116 | 0.36255822 | 0.83420249 | 0.90213531 | NA | XI | 473723 | 475900 | -1 |
| YKR019C | 76.1839461 | -0.0572902 | 0.56608523 | 0.55917136 | 0.75609694 | 0.85254164 | IRS4 | XI | 476216 | 478063 | -1 |
| YKR020W | 28.8894314 | 1.8105451 | 0.72574273 | 6.21627752 | 0.04468405 | 0.1201675 | VPS51 | XI | 478338 | 478832 | 1 |
| YKR021W | 75.7263776 | 1.86592941 | 0.67742389 | 10.0564336 | 0.00655048 | 0.02840995 | ALY1 | XI | 479234 | 481981 | 1 |
| YKR022C | 36.4194236 | -1.4113348 | 0.68937984 | 4.78989428 | 0.0911775 | 0.20517595 | NTR2 | XI | 482171 | 483139 | -1 |
| YKR023W | 46.8242679 | -1.2139654 | 0.63017148 | 4.19326351 | 0.12286959 | 0.25402775 | NA | XI | 483419 | 485011 | 1 |
| YKR024C | 13.6310549 | -0.1425081 | 0.88616944 | 0.1038283 | 0.94941037 | 0.96524777 | DBP7 | XI | 485144 | 487372 | -1 |
| YKR025W | 353.531283 | -1.2335055 | 0.63321228 | 9.10890053 | 0.01052028 | 0.04099087 | RPC37 | XI | 487770 | 488618 | 1 |
| YKR026C | 42.8567737 | -1.5724552 | 0.69774046 | 6.71724558 | 0.03478313 | 0.10004686 | GCN3 | XI | 488738 | 489655 | -1 |
| YKR027W | 16.4579408 | 0.00832036 | 0.7936586 | 0.10483132 | 0.94893435 | 0.96493324 | BCH2 | XI | 491364 | 493661 | 1 |
| YKR028W | 35.1784014 | -0.7232704 | 0.6907811 | 2.25072125 | 0.32453541 | 0.49763242 | SAP190 | XI | 494257 | 497358 | 1 |
| YKR029C | 31.2181708 | 1.26119633 | 0.658932 | 4.884581 | 0.08696144 | 0.19877881 | SET3 | XI | 497578 | 499833 | -1 |
| YKR030W | 17.529784 | -2.0814117 | 0.86746255 | 5.59527788 | 0.06095381 | 0.15179465 | GMH1 | XI | 500282 | 501103 | 1 |
| YKR031C | 26.8621012 | -1.6168012 | 0.76092614 | 4.59590907 | 0.10046413 | 0.2196191 | SPO14 | XI | 501344 | 506395 | -1 |
| YKR034W | 6.55252156 | -1.2525356 | 1.04810714 | 2.70165182 | 0.25902624 | 0.43082351 | DAL80 | XI | 506898 | 507707 | 1 |
| YKR035W-A | 83.9636657 | -2.339573 | 0.54322898 | 17.5217136 | 0.00015675 | 0.00138141 | DID2 | XI | 507939 | 508553 | 1 |
| YKR036C | 34.7803364 | 0.97820316 | 0.62605168 | 3.29989171 | 0.19206031 | 0.34838007 | CAF4 | XI | 508702 | 510633 | -1 |
| YKR037C | 24.1793831 | -0.1744146 | 0.75478588 | 1.16894484 | 0.55739986 | 0.70899499 | SPC34 | XI | 510910 | 511797 | -1 |
| YKR038C | 24.0137057 | -0.2988517 | 0.7344921 | 0.71265746 | 0.70024239 | 0.81801018 | KAE1 | XI | 511999 | 513159 | -1 |
| YKR039W | 39.9995743 | 1.81298929 | 0.67448419 | 11.1498736 | 0.00379172 | 0.01838696 | GAP1 | XI | 515063 | 516871 | 1 |
| YKR041W | 8.16494176 | -1.1610032 | 1.05034649 | 1.24127539 | 0.5376015 | 0.69208525 | NA | XI | 518198 | 518950 | 1 |
| YKR042W | 112.956815 | 0.24490589 | 0.67458636 | 0.51132065 | 0.77440497 | 0.86282905 | UTH1 | XI | 519527 | 520624 | 1 |
| YKR043C | 140.486104 | -0.5752429 | 0.50405066 | 1.32923371 | 0.51447061 | 0.67150244 | SHB17 | XI | 520897 | 521712 | -1 |
| YKR044W | 1150.46441 | -1.8785971 | 0.47576622 | 15.865161 | 0.00035886 | 0.00283852 | UIP5 | XI | 522015 | 523346 | 1 |

| | | | | | | | | | | | |
|---|---|---|---|---|---|---|---|---|---|---|---|
| YKR045C | 79.4365272 | -2.0634418 | 0.55297226 | 13.4161528 | 0.00122101 | 0.00776032 | NA | XI | 523418 | 523969 | -1 |
| YKR046C | 1992.18604 | -3.2720025 | 0.47325367 | 45.8506971 | 1.11E-10 | 5.16E-09 | PET10 | XI | 524223 | 525074 | -1 |
| YKR048C | 200.983737 | -0.159332 | 0.55569769 | 1.88459376 | 0.38973164 | 0.56101577 | NAP1 | XI | 525387 | 526640 | -1 |
| YKR049C | 391.686433 | 0.28000434 | 0.56903905 | 0.2988971 | 0.86118275 | 0.91844688 | FMP46 | XI | 526830 | 527231 | -1 |
| YKR050W | 25.897321 | 0.53140905 | 0.7778896 | 1.08600238 | 0.58100194 | 0.72940182 | TRK2 | XI | 527815 | 530484 | 1 |
| YKR051W | 152.527023 | -0.0933983 | 0.74308765 | 6.3635487 | 0.04151193 | 0.11429106 | NA | XI | 530946 | 532202 | 1 |
| YKR052C | 12.407362 | 0.46671149 | 0.91216606 | 0.50853224 | 0.7754854 | 0.8634877 | MRS4 | XI | 532550 | 533464 | -1 |
| YKR053C | 482.630563 | 0.03775397 | 0.4761828 | 0.89051888 | 0.64065804 | 0.77561669 | YSR3 | XI | 534067 | 535281 | -1 |
| YKR054C | 98.5596393 | 0.34025835 | 0.51217666 | 2.22986439 | 0.32793751 | 0.50041094 | DYN1 | XI | 535647 | 547925 | -1 |
| YKR056W | 73.9645614 | 1.73667863 | 0.58519137 | 8.55653701 | 0.01386665 | 0.05043211 | TRM2 | XI | 549448 | 551367 | 1 |
| YKR057W | 27.6511602 | -3.6482084 | 0.8220939 | 18.8966499 | 7.88E-05 | 0.00076317 | RPS21A | XI | 551657 | 552242 | 1 |
| YKR058W | 26.732947 | -1.9451978 | 0.6731097 | 8.52011467 | 0.01412149 | 0.05110095 | GLG1 | XI | 552770 | 554620 | 1 |
| YKR059W | 19.4387559 | -2.2064203 | 0.75111106 | 8.75408646 | 0.01256245 | 0.04680603 | TIF2 | XI | 554987 | 556174 | 1 |
| YKR060W | 50.0613308 | 0.87854457 | 0.62000842 | 6.26887883 | 0.04352415 | 0.11820113 | UTP30 | XI | 556518 | 557342 | 1 |
| YKR061W | 126.33766 | -1.5562391 | 0.51775735 | 8.74983386 | 0.01258919 | 0.04684525 | KTR2 | XI | 557677 | 558954 | 1 |
| YKR062W | 182.607888 | -2.257951 | 0.50453931 | 19.0588848 | 7.27E-05 | 0.00071568 | TFA2 | XI | 559666 | 560652 | 1 |
| YKR063C | 739.036832 | -0.974259 | 0.56602338 | 4.28465863 | 0.11738111 | 0.24546419 | LAS1 | XI | 560789 | 562297 | -1 |
| YKR064W | 41.9874984 | -0.5614606 | 0.72832988 | 4.17870194 | 0.12376744 | 0.25551406 | OAF3 | XI | 562547 | 565138 | 1 |
| YKR065C | 15.8109404 | 1.81252943 | 0.90196009 | 4.3655456 | 0.11272852 | 0.23918753 | PAM17 | XI | 565299 | 565892 | -1 |
| YKR066C | 44.8801936 | 0.71045059 | 0.66386035 | 1.16783855 | 0.55770826 | 0.70907501 | CCP1 | XI | 566113 | 567198 | -1 |
| YKR067W | 146.561829 | 3.35272548 | 0.55572317 | 42.3726963 | 6.29E-10 | 2.37E-08 | GPT2 | XI | 567918 | 570149 | 1 |
| YKR068C | 653.86651 | 2.75362892 | 0.51897482 | 25.1865678 | 3.39E-06 | 5.12E-05 | BET3 | XI | 570328 | 570909 | -1 |
| YKR069W | 95.8746973 | 2.47074371 | 0.59116043 | 22.6389729 | 1.21E-05 | 0.00015675 | MET1 | XI | 571612 | 573393 | 1 |
| YKR070W | 65.1960973 | 0.72080684 | 0.6168731 | 1.81713027 | 0.40310221 | 0.57401913 | NA | XI | 573574 | 574632 | 1 |
| YKR071C | 65.1579686 | -1.3865053 | 0.6010535 | 5.54412962 | 0.06253275 | 0.15457697 | DRE2 | XI | 574934 | 575980 | -1 |
| YKR072C | 353.91002 | 0.18410522 | 0.57239216 | 4.8641233 | 0.08785552 | 0.20034795 | SIS2 | XI | 576435 | 578123 | -1 |
| YKR073C | 78.9742015 | 1.59819799 | 0.62598749 | 8.24321325 | 0.01621844 | 0.0567598 | NA | XI | 578187 | 578507 | -1 |
| YKR075C | 64.8983632 | -1.4186934 | 0.59131659 | 5.57715715 | 0.06150858 | 0.15245492 | NA | XI | 579827 | 580750 | -1 |
| YKR076W | 48.0971875 | -1.9414072 | 0.67735857 | 8.23088273 | 0.01631874 | 0.05693318 | ECM4 | XI | 582283 | 583395 | 1 |
| YKR077W | 57.3494246 | -1.7573297 | 0.60456224 | 9.09687109 | 0.01058375 | 0.04118273 | MSA2 | XI | 583653 | 584744 | 1 |
| YKR078W | 83.940083 | -2.0859187 | 0.62528119 | 10.3869777 | 0.0055526 | 0.02490283 | NA | XI | 584952 | 586709 | 1 |
| YKR079C | 160.534679 | -0.6197511 | 0.50954367 | 1.50650132 | 0.47083354 | 0.63296713 | TRZ1 | XI | 586789 | 589305 | -1 |
| YKR080W | 6.11961604 | -2.1761578 | 1.18057327 | 4.0998286 | 0.12874594 | 0.26238612 | MTD1 | XI | 590395 | 591357 | 1 |

| | | | | | | | | | | | |
|---|---|---|---|---|---|---|---|---|---|---|---|
| YKR081C | 51.8453539 | -2.9577401 | 0.75882849 | 13.8131881 | 0.00100116 | 0.00658719 | RPF2 | XI | 591506 | 592540 | -1 |
| YKR082W | 57.7524104 | -0.786325 | 0.73400104 | 8.72457462 | 0.01274919 | 0.04728881 | NUP133 | XI | 592825 | 596298 | 1 |
| YKR083C | 33.4870702 | -1.9685185 | 0.72196438 | 7.42122376 | 0.02446255 | 0.07676865 | DAD2 | XI | 596421 | 596822 | -1 |
| YKR084C | 15.5883272 | -0.5127393 | 0.79435985 | 0.42703108 | 0.80773961 | 0.88441988 | HBS1 | XI | 597055 | 598890 | -1 |
| YKR085C | 7.43798492 | -0.8199882 | 1.13904577 | 1.06680159 | 0.58660665 | 0.73405068 | MRPL20 | XI | 599094 | 599681 | -1 |
| YKR086W | 34.6164534 | 0.25724062 | 0.63127539 | 0.94619231 | 0.62307016 | 0.76286238 | PRP16 | XI | 599857 | 603072 | 1 |
| YKR087C | 91.9285746 | -1.5877271 | 0.54053617 | 8.41248301 | 0.01490227 | 0.05319615 | OMA1 | XI | 603195 | 604232 | -1 |
| YKR088C | 34.2129609 | -0.0421519 | 0.79601419 | 0.6189664 | 0.7338261 | 0.83775415 | TVP38 | XI | 604405 | 605418 | -1 |
| YKR089C | 289.977292 | -0.4550489 | 0.48251549 | 1.10952374 | 0.57420898 | 0.7240135 | TGL4 | XI | 605633 | 608365 | -1 |
| YKR090W | 759.347307 | 1.81807388 | 0.48656772 | 13.2168547 | 0.00134895 | 0.00838929 | PXL1 | XI | 608946 | 611066 | 1 |
| YKR091W | 53.3464664 | 1.43810696 | 0.60852123 | 6.47956026 | 0.03917251 | 0.10988799 | SRL3 | XI | 611525 | 612265 | 1 |
| YKR092C | 44.7025307 | -0.1022598 | 0.74850019 | 0.97537035 | 0.61404616 | 0.75677464 | SRP40 | XI | 612665 | 613885 | -1 |
| YKR093W | 31.9544664 | -0.6535923 | 0.67981003 | 1.29351893 | 0.52374023 | 0.67900939 | PTR2 | XI | 615730 | 617535 | 1 |
| YKR094C | 60.8152594 | -2.9468235 | 0.70813466 | 16.2427813 | 0.00029712 | 0.0024298 | RPL40B | XI | 617996 | 618750 | -1 |
| YKR095W | 91.8030771 | -0.4343404 | 0.63690828 | 5.4003421 | 0.06719402 | 0.16286148 | MLP1 | XI | 619805 | 625432 | 1 |
| YKR095W-A | 35.1904138 | 1.11237289 | 0.83637263 | 1.82974576 | 0.40056754 | 0.57138844 | PCC1 | XI | 625864 | 626205 | 1 |
| YKR096W | 216.104016 | 3.70317982 | 0.54532389 | 46.024149 | 1.01E-10 | 4.77E-09 | ESL2 | XI | 626793 | 630380 | 1 |
| YKR097W | 86.6886956 | 1.16958006 | 0.52630688 | 5.91371151 | 0.0519821 | 0.13454465 | PCK1 | XI | 631152 | 632801 | 1 |
| YKR098C | 18.7655514 | 0.73739975 | 0.83805995 | 3.12617782 | 0.20948798 | 0.37244958 | UBP11 | XI | 633026 | 635179 | -1 |
| YKR099W | 19.6074458 | -0.7983923 | 0.70877423 | 1.43801012 | 0.48723679 | 0.64903366 | BAS1 | XI | 635851 | 638286 | 1 |
| YKR100C | 29.487636 | 0.8249459 | 0.6735555 | 1.59428335 | 0.45061513 | 0.61694528 | SKG1 | XI | 638901 | 639968 | -1 |
| YKR101W | 24.0404526 | 3.29699156 | 0.8566949 | 20.7688628 | 3.09E-05 | 0.00034617 | SIR1 | XI | 640540 | 642504 | 1 |
| YKR102W | 378.193542 | 3.56268639 | 0.50557049 | 50.5184344 | 1.07E-11 | 6.20E-10 | FLO10 | XI | 646356 | 649865 | 1 |
| YKR103W | 28.4768296 | 1.77920908 | 0.70065855 | 10.2372454 | 0.00598426 | 0.02644951 | NFT1 | XI | 653080 | 656736 | 1 |
| YKR105C | 648.103478 | -0.0345325 | 0.48209374 | 1.70776282 | 0.42575918 | 0.5957336 | VBA5 | XI | 658716 | 660464 | -1 |
| YKR106W | 264.777876 | -0.1904735 | 0.51457666 | 0.1370999 | 0.93374682 | 0.95735444 | GEX2 | XI | 661442 | 663289 | 1 |
| YLL001W | 235.588272 | 3.65643609 | 0.55787651 | 47.385459 | 5.13E-11 | 2.56E-09 | DNM1 | XII | 147890 | 150163 | 1 |
| YLL002W | 33.1336444 | 0.05079121 | 0.62740123 | 3.21921423 | 0.19996616 | 0.35900663 | RTT109 | XII | 146291 | 147601 | 1 |
| YLL003W | 20.5157482 | 0.60599851 | 0.71270499 | 0.77098275 | 0.68011636 | 0.80315597 | SFI1 | XII | 143201 | 146041 | 1 |
| YLL004W | 7.29859745 | -0.3552659 | 1.01684913 | 0.13328476 | 0.93552971 | 0.95803272 | ORC3 | XII | 141073 | 142923 | 1 |
| YLL005C | 2977.86199 | -0.1779117 | 0.53237544 | 0.62136767 | 0.73294557 | 0.83751334 | SPO75 | XII | 137940 | 140546 | -1 |
| YLL006W | 51.8320068 | 2.74216392 | 0.63348986 | 19.7395128 | 5.17E-05 | 0.0005281 | MMM1 | XII | 136586 | 137866 | 1 |
| YLL006W-A | 77.5372876 | -1.1481994 | 0.55976186 | 4.13549452 | 0.12647037 | 0.25966788 | NA | XII | 136345 | 136521 | 1 |

| | | | | | | | | | | | |
|---|---|---|---|---|---|---|---|---|---|---|---|
| YLL007C | 20.5227497 | -0.8497561 | 0.69661206 | 6.59402781 | 0.03699347 | 0.10476085 | LMO1 | XII | 134302 | 136299 | -1 |
| YLL008W | 191.196088 | -2.7263493 | 0.54108916 | 22.953579 | 1.04E-05 | 0.00013674 | DRS1 | XII | 131729 | 133987 | 1 |
| YLL009C | 21.7580001 | -0.2854361 | 0.85466312 | 0.27433915 | 0.87182237 | 0.92527295 | COX17 | XII | 131205 | 131414 | -1 |
| YLL010C | 204.439929 | 0.14762331 | 0.55330426 | 1.76942677 | 0.41283248 | 0.58428259 | PSR1 | XII | 129330 | 130613 | -1 |
| YLL011W | 34.6834778 | -1.1328679 | 0.64471541 | 3.43185682 | 0.17979672 | 0.33430411 | SOF1 | XII | 127523 | 128992 | 1 |
| YLL012W | 104.168444 | 0.97903689 | 0.55758885 | 3.04899906 | 0.21773 | 0.38155469 | YEH1 | XII | 125534 | 127255 | 1 |
| YLL013C | 39.6325083 | -0.243992 | 0.65747535 | 0.74664885 | 0.68844185 | 0.81049746 | PUF3 | XII | 122075 | 124714 | -1 |
| YLL014W | 6.91345042 | -1.0894049 | 1.21583154 | 0.93465171 | 0.62667585 | 0.76471951 | EMC6 | XII | 121322 | 121648 | 1 |
| YLL015W | 45.856788 | -0.3259645 | 0.71680841 | 0.23312044 | 0.8899765 | 0.93317728 | BPT1 | XII | 116432 | 121111 | 1 |
| YLL016W | 1162.55563 | -0.358366 | 0.48986778 | 0.62154972 | 0.73287886 | 0.83751334 | NA | XII | 112847 | 115993 | 1 |
| YLL017W | 500.027079 | 0.0881335 | 0.51550154 | 0.17417096 | 0.91659874 | 0.94860224 | NA | XII | 112234 | 112545 | 1 |
| YLL018C | 83.8652628 | -0.9019776 | 0.6244898 | 3.56111382 | 0.16854426 | 0.31849584 | DPS1 | XII | 109902 | 111575 | -1 |
| YLL019C | 63.317529 | -1.6510354 | 0.58026713 | 10.4722581 | 0.00532081 | 0.02406837 | KNS1 | XII | 105686 | 107899 | -1 |
| YLL021W | 68.3782834 | -1.673631 | 0.59060849 | 8.08198159 | 0.01758005 | 0.05999792 | SPA2 | XII | 100947 | 105347 | 1 |
| YLL022C | 68.8259455 | -0.5128866 | 0.56943262 | 3.89683419 | 0.14249946 | 0.28204189 | HIF1 | XII | 99044 | 100201 | -1 |
| YLL023C | 76.0218861 | -1.2149611 | 0.57422464 | 4.71607539 | 0.09460569 | 0.21003333 | POM33 | XII | 97997 | 98836 | -1 |
| YLL024C | 151.286306 | -1.2826544 | 0.77056134 | 2.73167704 | 0.25516662 | 0.42736903 | SSA2 | XII | 95566 | 97485 | -1 |
| YLL025W | 35.0974549 | 0.69350996 | 0.65747453 | 1.45919589 | 0.48210278 | 0.64436175 | PAU17 | XII | 94747 | 95121 | 1 |
| YLL026W | 80.3172799 | 1.95883222 | 0.61600633 | 10.9727142 | 0.00414291 | 0.01969412 | HSP104 | XII | 88623 | 91349 | 1 |
| YLL027W | 22.3436809 | 0.53454768 | 0.70759976 | 2.08334809 | 0.35286348 | 0.52643121 | ISA1 | XII | 87403 | 88155 | 1 |
| YLL028W | 257.903926 | 1.84978332 | 0.48193859 | 15.2035807 | 0.00049956 | 0.00367131 | TPO1 | XII | 84804 | 86564 | 1 |
| YLL029W | 103.212649 | 0.24802441 | 0.52849474 | 0.3315859 | 0.84722163 | 0.90980423 | FRA1 | XII | 81461 | 83710 | 1 |
| YLL031C | 19.4109667 | -1.3780148 | 0.78627863 | 3.26661035 | 0.19528306 | 0.35245914 | GPI13 | XII | 77152 | 80205 | -1 |
| YLL032C | 282.078234 | -0.7803209 | 0.49993184 | 2.76159441 | 0.25137807 | 0.42273571 | NA | XII | 74270 | 76747 | -1 |
| YLL033W | 143.706275 | -1.3164978 | 0.51780652 | 7.13785792 | 0.02818603 | 0.08482177 | IRC19 | XII | 73409 | 74101 | 1 |
| YLL034C | 98.1423989 | -1.3432691 | 0.77705032 | 8.44076178 | 0.01469305 | 0.05261147 | RIX7 | XII | 70633 | 73146 | -1 |
| YLL035W | 27.1781537 | -1.5604982 | 0.66426669 | 6.0592691 | 0.0483333 | 0.12713301 | GRC3 | XII | 68580 | 70478 | 1 |
| YLL036C | 90.6045044 | -1.095715 | 0.53289213 | 7.76553706 | 0.02059373 | 0.06755677 | PRP19 | XII | 66745 | 68256 | -1 |
| YLL038C | 75.6754422 | 1.62863957 | 0.56078741 | 12.7832308 | 0.00167555 | 0.00980703 | ENT4 | XII | 65775 | 66518 | -1 |
| YLL039C | 71.6765205 | 1.9627816 | 0.85554106 | 5.68729625 | 0.05821291 | 0.14655695 | UBI4 | XII | 64062 | 65207 | -1 |
| YLL040C | 35.1409347 | -0.4871505 | 0.94435204 | 0.32016111 | 0.85207515 | 0.91195795 | VPS13 | XII | 54211 | 63645 | -1 |
| YLL041C | 9.90484103 | 0.52953591 | 1.28022028 | 0.23413917 | 0.88952329 | 0.93317728 | SDH2 | XII | 53131 | 53931 | -1 |
| YLL042C | 62.4104949 | 0.31211433 | 0.83329289 | 9.76358512 | 0.00758341 | 0.03186352 | ATG10 | XII | 52087 | 52590 | -1 |

| | | | | | | | | | | | |
|---|---|---|---|---|---|---|---|---|---|---|---|
| YLL043W | 32.9942993 | -0.4679855 | 0.73242108 | 0.77879159 | 0.67746608 | 0.8011991 | FPS1 | XII | 49938 | 51947 | 1 |
| YLL045C | 135.942016 | -3.1627948 | 0.74824449 | 16.0338369 | 0.00032983 | 0.00264874 | RPL8B | XII | 47859 | 48629 | -1 |
| YLL048C | 87.2818636 | -1.0663834 | 0.63106502 | 5.58532203 | 0.06125799 | 0.15215948 | YBT1 | XII | 41280 | 46265 | -1 |
| YLL049W | 320.831728 | -2.4248973 | 0.63007223 | 15.9571191 | 0.00034273 | 0.00272586 | LDB18 | XII | 40666 | 41205 | 1 |
| YLL050C | 21.1108397 | -0.654934 | 0.84129273 | 1.17536722 | 0.55561281 | 0.70703257 | COF1 | XII | 39804 | 40414 | -1 |
| YLL051C | 380.704381 | 1.46030009 | 0.50579186 | 13.8408822 | 0.00098739 | 0.00651883 | FRE6 | XII | 37333 | 39471 | -1 |
| YLL052C | 6.39995592 | 1.29524608 | 1.11532654 | 1.31553293 | 0.51800703 | 0.67444585 | AQY2 | XII | 35912 | 36361 | -1 |
| YLL053C | 9.11035867 | -0.3812774 | 1.10989216 | 0.73366584 | 0.69292541 | 0.81380084 | NA | XII | 35503 | 35961 | -1 |
| YLL054C | 39.1516127 | 1.81344542 | 0.61624094 | 12.5696514 | 0.00186438 | 0.01063524 | NA | XII | 32673 | 35204 | -1 |
| YLL055W | 36.903787 | -1.9534247 | 0.64551266 | 10.8724137 | 0.00435597 | 0.0204883 | YCT1 | XII | 30109 | 31704 | 1 |
| YLL056C | 17.3978481 | 2.77553507 | 0.91604614 | 9.59777662 | 0.0082389 | 0.03395248 | NA | XII | 27409 | 28305 | -1 |
| YLL057C | 557.469278 | -0.6331854 | 0.48772652 | 1.67125289 | 0.43360277 | 0.60207676 | JLP1 | XII | 25756 | 26994 | -1 |
| YLL058W | 22.7491351 | 0.35661425 | 0.97174557 | 0.17910121 | 0.91434199 | 0.94739444 | NA | XII | 23569 | 25296 | 1 |
| YLL060C | 691.743166 | 6.72295114 | 0.54436613 | 119.705901 | 1.01E-26 | 1.17E-23 | GTT2 | XII | 21138 | 21839 | -1 |
| YLL061W | 2093.38991 | 3.77113537 | 0.61460489 | 31.8303904 | 1.22E-07 | 2.88E-06 | MMP1 | XII | 17956 | 19707 | 1 |
| YLL062C | 46.5475082 | 1.95422251 | 0.59996869 | 11.962609 | 0.00252553 | 0.01335631 | MHT1 | XII | 16639 | 17613 | -1 |
| YLL063C | 6.02964683 | 1.03827812 | 1.15279192 | 2.34046867 | 0.31029422 | 0.4830878 | AYT1 | XII | 14648 | 16072 | -1 |
| YLR001C | 365.888899 | -0.4805298 | 0.49444685 | 1.58770031 | 0.45210078 | 0.61810237 | NA | XII | 151389 | 153977 | -1 |
| YLR002C | 53.8476042 | 1.68889089 | 0.58400843 | 10.4002427 | 0.00551589 | 0.02475739 | NOC3 | XII | 154343 | 156334 | -1 |
| YLR003C | 21.7499136 | -0.5214471 | 0.76227465 | 0.60486692 | 0.73901766 | 0.84040603 | CMS1 | XII | 156855 | 157730 | -1 |
| YLR004C | 385.713422 | 1.45020303 | 0.54883036 | 7.33870584 | 0.02549296 | 0.07880632 | THI73 | XII | 157934 | 159505 | -1 |
| YLR005W | 82.5953522 | 1.98849545 | 0.54977316 | 16.839874 | 0.00022043 | 0.001854 | SSL1 | XII | 160049 | 161434 | 1 |
| YLR006C | 228.440767 | 0.46867546 | 0.49516683 | 1.62603137 | 0.44351853 | 0.61162622 | SSK1 | XII | 161755 | 163893 | -1 |
| YLR007W | 561.424285 | -2.2261748 | 0.62238268 | 14.569631 | 0.00068587 | 0.00479615 | NSE1 | XII | 164392 | 165402 | 1 |
| YLR008C | 25.4241553 | 0.35262859 | 0.67053473 | 4.25917216 | 0.11888649 | 0.2479657 | PAM18 | XII | 165577 | 166083 | -1 |
| YLR009W | 69.0145571 | -2.7880742 | 0.66257471 | 18.0888671 | 0.00011805 | 0.00107636 | RLP24 | XII | 166537 | 167136 | 1 |
| YLR010C | 17.066754 | -1.4799522 | 0.76320355 | 6.223278 | 0.04452791 | 0.11991471 | TEN1 | XII | 167320 | 167802 | -1 |
| YLR011W | 21.1226735 | 0.19077464 | 0.75834548 | 0.15251612 | 0.92657706 | 0.95392624 | LOT6 | XII | 169103 | 169678 | 1 |
| YLR014C | 49.6776571 | 2.13123107 | 0.75938545 | 12.8066307 | 0.00165606 | 0.00977813 | PPR1 | XII | 172268 | 174982 | -1 |
| YLR015W | 155.647183 | -1.3573445 | 0.49819145 | 7.66019432 | 0.02170751 | 0.07037316 | BRE2 | XII | 175227 | 176744 | 1 |
| YLR016C | 172.079756 | -1.083772 | 0.49351961 | 4.8621903 | 0.08794047 | 0.20046273 | PML1 | XII | 176802 | 177416 | -1 |
| YLR017W | 87.9483656 | 0.47244573 | 0.75760967 | 3.36367001 | 0.18603229 | 0.3417281 | MEU1 | XII | 177608 | 178621 | 1 |
| YLR018C | 92.9936797 | 0.23259768 | 0.81350821 | 5.42601793 | 0.0663369 | 0.16158631 | POM34 | XII | 178707 | 179606 | -1 |

| | | | | | | | | | | | |
|---|---|---|---|---|---|---|---|---|---|---|---|
| YLR019W | 91.9699081 | -0.2960112 | 0.67960319 | 0.20632417 | 0.90198076 | 0.93824416 | PSR2 | XII | 180288 | 181481 | 1 |
| YLR020C | 14.2542925 | -0.1575171 | 0.91353778 | 0.07730883 | 0.96208313 | 0.97258112 | YEH2 | XII | 181789 | 183405 | -1 |
| YLR021W | 256.618813 | -1.4705776 | 0.57058995 | 7.54917695 | 0.02294653 | 0.07336301 | IRC25 | XII | 183623 | 184162 | 1 |
| YLR022C | 147.332841 | -1.0788355 | 0.65205305 | 5.3264753 | 0.06972212 | 0.16778515 | SDO1 | XII | 184174 | 184926 | -1 |
| YLR023C | 7.17040836 | -0.7189557 | 1.03973958 | 0.51870938 | 0.77154932 | 0.86091165 | IZH3 | XII | 185498 | 187129 | -1 |
| YLR024C | 173.324556 | 0.5571363 | 0.49138883 | 2.28629215 | 0.31881443 | 0.49159402 | UBR2 | XII | 187663 | 193281 | -1 |
| YLR025W | 70.3988846 | -0.6584108 | 0.57577175 | 1.44438228 | 0.48568688 | 0.64773825 | SNF7 | XII | 194452 | 195174 | 1 |
| YLR026C | 52.0598605 | 0.41893196 | 0.5723367 | 0.53013068 | 0.76715582 | 0.85857988 | SED5 | XII | 195450 | 196472 | -1 |
| YLR027C | 73.5191573 | -1.5789216 | 0.58894899 | 7.40936581 | 0.02460802 | 0.07694787 | AAT2 | XII | 196829 | 198085 | -1 |
| YLR028C | 108.329668 | 0.01399124 | 0.63655758 | 1.60547976 | 0.44809954 | 0.61495528 | ADE16 | XII | 199540 | 201315 | -1 |
| YLR029C | 190.228142 | -2.5032673 | 0.60069191 | 15.9597593 | 0.00034228 | 0.00272586 | RPL15A | XII | 201976 | 202590 | -1 |
| YLR030W | 461.237851 | -1.6773237 | 0.52194601 | 12.4926628 | 0.00193755 | 0.01091285 | NA | XII | 203290 | 204081 | 1 |
| YLR031W | 131.582308 | -3.5886473 | 0.56663023 | 43.9760109 | 2.82E-10 | 1.18E-08 | NA | XII | 204225 | 204785 | 1 |
| YLR032W | 28.2710384 | -0.1580546 | 0.66219605 | 0.06727127 | 0.96692376 | 0.97486964 | RAD5 | XII | 204991 | 208500 | 1 |
| YLR033W | 72.88106 | -1.1263144 | 0.55863114 | 6.26145181 | 0.04368607 | 0.11847418 | RSC58 | XII | 208761 | 210269 | 1 |
| YLR034C | 46.565282 | -0.0610736 | 0.624815 | 1.69306817 | 0.42889889 | 0.59853568 | SMF3 | XII | 210512 | 211933 | -1 |
| YLR035C | 78.8481265 | 0.46048164 | 0.53793108 | 0.93391144 | 0.62690785 | 0.76477701 | MLH2 | XII | 212369 | 214456 | -1 |
| YLR036C | 132.802365 | 0.28620684 | 0.60902303 | 1.10789983 | 0.5746754 | 0.72422968 | NA | XII | 221521 | 222132 | -1 |
| YLR038C | 493.818553 | 4.13302813 | 0.55254404 | 50.7077881 | 9.75E-12 | 5.76E-10 | COX12 | XII | 224921 | 225172 | -1 |
| YLR039C | 100.352284 | 2.03641813 | 0.53528331 | 26.2520918 | 1.99E-06 | 3.24E-05 | RIC1 | XII | 225426 | 228596 | -1 |
| YLR040C | 143.427358 | 0.54827543 | 0.53419228 | 4.60322323 | 0.1000974 | 0.21919967 | AFB1 | XII | 228908 | 229582 | -1 |
| YLR042C | 1874.38027 | -0.1670365 | 0.51988399 | 2.35671107 | 0.30778446 | 0.48023239 | NA | XII | 229966 | 230451 | -1 |
| YLR043C | 71.6066812 | -0.1223439 | 0.55738498 | 1.90420933 | 0.38592792 | 0.55863066 | TRX1 | XII | 231702 | 232013 | -1 |
| YLR044C | 688.9685 | -2.0476094 | 0.67416242 | 8.580594 | 0.01370086 | 0.04989179 | PDC1 | XII | 232390 | 234081 | -1 |
| YLR045C | 142.068932 | -1.6546579 | 0.52746407 | 9.50660719 | 0.00862316 | 0.03516063 | STU2 | XII | 235037 | 237703 | -1 |
| YLR046C | 21.5722861 | 0.23559556 | 0.74271041 | 0.12549684 | 0.93917972 | 0.96024203 | NA | XII | 238001 | 238813 | -1 |
| YLR047C | 394.564398 | -0.8266089 | 0.5501488 | 6.92326186 | 0.03137854 | 0.09258058 | FRE8 | XII | 239347 | 241407 | -1 |
| YLR048W | 168.218799 | -3.1414531 | 0.59274408 | 25.2795451 | 3.24E-06 | 4.95E-05 | RPS0B | XII | 242232 | 243349 | 1 |
| YLR049C | 30.3707644 | 1.10070162 | 0.69390728 | 2.59435911 | 0.27330154 | 0.44663164 | NA | XII | 243886 | 245172 | -1 |
| YLR050C | 473.770481 | -2.2496057 | 0.47689568 | 21.9267319 | 1.73E-05 | 0.00020811 | NA | XII | 245587 | 246072 | -1 |
| YLR051C | 13.9543765 | -2.0234676 | 0.89682039 | 5.52681326 | 0.06307652 | 0.15574118 | FCF2 | XII | 246324 | 246977 | -1 |
| YLR052W | 14.4461265 | -0.1381327 | 0.85948029 | 1.07291807 | 0.5848154 | 0.7327594 | IES3 | XII | 247201 | 247953 | 1 |
| YLR053C | 326.179925 | -1.8828214 | 0.57742711 | 10.1896076 | 0.00612851 | 0.02692266 | NA | XII | 248100 | 248426 | -1 |

| | | | | | | | | | | | |
|---|---|---|---|---|---|---|---|---|---|---|---|
| YLR054C | 392.045902 | 4.00093091 | 0.51866818 | 51.2887422 | 7.29E-12 | 4.52E-10 | OSW2 | XII | 248717 | 250978 | -1 |
| YLR055C | 131.699232 | 1.03188023 | 0.52366555 | 3.82630148 | 0.14761456 | 0.28982309 | SPT8 | XII | 251272 | 253080 | -1 |
| YLR056W | 52.0152027 | -1.0907861 | 0.58108228 | 3.48285343 | 0.17527016 | 0.32699386 | ERG3 | XII | 253861 | 254958 | 1 |
| YLR057W | 143.094987 | 0.22689044 | 0.88557104 | 8.08486075 | 0.01755476 | 0.05999792 | MNL2 | XII | 255306 | 257855 | 1 |
| YLR058C | 32.9524941 | -0.6509595 | 0.65291265 | 1.01662504 | 0.60150976 | 0.74720908 | SHM2 | XII | 257992 | 259401 | -1 |
| YLR059C | 236.150597 | -1.2659052 | 0.5019322 | 7.66433135 | 0.02166265 | 0.07033309 | REX2 | XII | 259738 | 260547 | -1 |
| YLR060W | 118.650096 | 1.16645905 | 0.63974819 | 3.43689527 | 0.17934434 | 0.3335701 | FRS1 | XII | 260979 | 262766 | 1 |
| YLR061W | 115.776088 | -3.4126505 | 0.57456156 | 32.5093148 | 8.72E-08 | 2.16E-06 | RPL22A | XII | 263194 | 263948 | 1 |
| YLR063W | 44.9216149 | 0.57038881 | 0.60176102 | 5.19515902 | 0.07445357 | 0.17631337 | BMT6 | XII | 264157 | 265254 | 1 |
| YLR064W | 163.842693 | -1.8827895 | 0.60176804 | 12.6003005 | 0.00183603 | 0.01052535 | PER33 | XII | 265456 | 266277 | 1 |
| YLR065C | 133.288523 | -1.0493347 | 0.65265391 | 5.89856594 | 0.05237725 | 0.13534434 | ENV10 | XII | 266373 | 266918 | -1 |
| YLR066W | 15.49876 | 0.51483595 | 0.85709936 | 3.39886488 | 0.18278724 | 0.33812719 | SPC3 | XII | 267169 | 267723 | 1 |
| YLR067C | 397.694641 | 0.20842396 | 0.90783508 | 13.8447177 | 0.0009855 | 0.00651377 | PET309 | XII | 267813 | 270710 | -1 |
| YLR068W | 59.4953894 | -3.0355617 | 0.67602182 | 18.5119312 | 9.55E-05 | 0.00089947 | FYV7 | XII | 271008 | 271463 | 1 |
| YLR069C | 46.5617288 | 0.39801496 | 0.58860057 | 1.88398601 | 0.38985009 | 0.56101577 | MEF1 | XII | 271630 | 273915 | -1 |
| YLR070C | 321.393428 | -0.5502513 | 0.46717102 | 1.38089626 | 0.50135135 | 0.66037092 | XYL2 | XII | 274141 | 275211 | -1 |
| YLR071C | 20.0380456 | -0.3288692 | 0.70645328 | 0.2213572 | 0.89522643 | 0.93531262 | RGR1 | XII | 275378 | 278626 | -1 |
| YLR072W | 54.6887811 | -0.9159809 | 0.70472833 | 6.82863601 | 0.03289884 | 0.09591352 | LAM6 | XII | 278862 | 280943 | 1 |
| YLR073C | 95.724713 | -1.6292678 | 0.68834401 | 6.87622048 | 0.03212534 | 0.0942798 | RFU1 | XII | 281019 | 281621 | -1 |
| YLR074C | 58.1086768 | -3.0449128 | 0.66143583 | 22.2849369 | 1.45E-05 | 0.00018113 | BUD20 | XII | 281956 | 282456 | -1 |
| YLR075W | 170.048347 | -2.8622334 | 0.90363979 | 8.9965994 | 0.0111279 | 0.042641 | RPL10 | XII | 282927 | 283592 | 1 |
| YLR077W | 73.4438498 | -0.1983871 | 0.56924112 | 0.22997501 | 0.89137728 | 0.93352778 | FMP25 | XII | 283872 | 285623 | 1 |
| YLR078C | 132.619071 | -0.0580353 | 0.76980776 | 6.60052549 | 0.03687348 | 0.10460433 | BOS1 | XII | 285736 | 286559 | -1 |
| YLR079W | 62.6184079 | -0.1223795 | 0.61708047 | 0.23129441 | 0.89078944 | 0.93317728 | SIC1 | XII | 286820 | 287674 | 1 |
| YLR080W | 96.8951977 | -0.3482959 | 0.59137677 | 5.54352652 | 0.06255161 | 0.15457697 | EMP46 | XII | 287916 | 289250 | 1 |
| YLR081W | 308.082255 | -0.5327681 | 0.52104827 | 1.636287 | 0.44125008 | 0.60960103 | GAL2 | XII | 290212 | 291936 | 1 |
| YLR082C | 5.8500978 | -1.4768725 | 1.23217372 | 2.02428184 | 0.36344005 | 0.53640527 | SRL2 | XII | 292394 | 293572 | -1 |
| YLR083C | 26.0368959 | -0.9059536 | 0.75489202 | 3.04524808 | 0.21813873 | 0.38215529 | EMP70 | XII | 294091 | 296094 | -1 |
| YLR084C | 56.0018901 | 1.80025088 | 0.57295102 | 11.5940722 | 0.00303654 | 0.01538156 | RAX2 | XII | 296589 | 300251 | -1 |
| YLR085C | 23.255992 | -1.8423138 | 0.76845862 | 7.53640785 | 0.0230935 | 0.0737108 | ARP6 | XII | 300673 | 301989 | -1 |
| YLR086W | 50.5364931 | -0.4566207 | 0.56524302 | 0.68052752 | 0.71158261 | 0.82438173 | SMC4 | XII | 302243 | 306499 | 1 |
| YLR087C | 356.039151 | -1.5131418 | 0.50131235 | 9.64767894 | 0.00803587 | 0.03330545 | CSF1 | XII | 306855 | 315731 | -1 |
| YLR088W | 29.3705196 | -0.3275953 | 0.75554216 | 0.26990802 | 0.87375609 | 0.92573532 | GAA1 | XII | 316107 | 317951 | 1 |

| | | | | | | | | | | | |
|---|---|---|---|---|---|---|---|---|---|---|---|
| YLR089C | 103.938229 | 0.16486752 | 0.5712589 | 1.71355147 | 0.42452867 | 0.59492495 | ALT1 | XII | 318237 | 320015 | -1 |
| YLR090W | 18.1744279 | -0.8883186 | 0.78071064 | 1.57793366 | 0.45431394 | 0.62039568 | XDJ1 | XII | 320701 | 322080 | 1 |
| YLR091W | 274.56831 | -0.6996087 | 0.59794784 | 1.50592922 | 0.47096824 | 0.63298657 | GEP5 | XII | 322297 | 323178 | 1 |
| YLR092W | 1297.45965 | 0.9031099 | 0.54478416 | 2.93805066 | 0.2301497 | 0.39706995 | SUL2 | XII | 323544 | 326225 | 1 |
| YLR093C | 30.5411433 | 1.22953566 | 0.70266062 | 3.12279929 | 0.20984216 | 0.37258084 | NYV1 | XII | 326513 | 327415 | -1 |
| YLR094C | 23.0068344 | -0.2465322 | 0.71147315 | 0.12243864 | 0.94061692 | 0.96082012 | GIS3 | XII | 327730 | 329238 | -1 |
| YLR095C | 263.740584 | -0.2119899 | 0.48562465 | 0.63329612 | 0.72858713 | 0.83634407 | IOC2 | XII | 329677 | 332115 | -1 |
| YLR096W | 38.4146317 | -0.9887171 | 0.63217392 | 2.96617067 | 0.22693643 | 0.3929312 | KIN2 | XII | 332590 | 336033 | 1 |
| YLR097C | 11.5237626 | 0.87901839 | 0.97255369 | 1.30534885 | 0.52065147 | 0.6765916 | HRT3 | XII | 336231 | 337265 | -1 |
| YLR098C | 62.7225267 | -0.7349558 | 0.61554076 | 3.97018058 | 0.13736821 | 0.27509737 | CHA4 | XII | 337527 | 339473 | -1 |
| YLR099C | 219.492179 | -0.4521853 | 0.52425578 | 10.4766619 | 0.00530911 | 0.02403421 | ICT1 | XII | 339744 | 340928 | -1 |
| YLR099W-A | 13.3433624 | -0.9349818 | 0.81717379 | 1.42719026 | 0.48987985 | 0.65114883 | MIM2 | XII | 341325 | 341588 | 1 |
| YLR100W | 414.85557 | 1.50243776 | 0.78108149 | 8.13501045 | 0.01712005 | 0.05900302 | ERG27 | XII | 341810 | 342853 | 1 |
| YLR102C | 153.33958 | 0.22301035 | 0.64948433 | 1.19119173 | 0.55123401 | 0.70300548 | APC9 | XII | 342970 | 343767 | -1 |
| YLR103C | 35.6019164 | -0.3502335 | 0.62964831 | 0.32244925 | 0.85110087 | 0.91173629 | CDC45 | XII | 343989 | 345941 | -1 |
| YLR104W | 157.46729 | 2.68521634 | 0.78579954 | 11.7220303 | 0.00284835 | 0.01465951 | LCL2 | XII | 346585 | 346980 | 1 |
| YLR105C | 62.2918952 | 1.40423635 | 0.82344116 | 6.16238562 | 0.04590447 | 0.12231333 | SEN2 | XII | 347047 | 348180 | -1 |
| YLR106C | 213.900546 | -0.6149107 | 0.56101864 | 1.34265316 | 0.5110302 | 0.66829159 | MDN1 | XII | 349006 | 363738 | -1 |
| YLR107W | 44.9989435 | 3.20812987 | 0.73554206 | 21.837798 | 1.81E-05 | 0.00021713 | REX3 | XII | 364116 | 365330 | 1 |
| YLR108C | 200.795583 | 3.11359891 | 0.53617138 | 32.1531342 | 1.04E-07 | 2.51E-06 | NA | XII | 366667 | 368124 | -1 |
| YLR109W | 92.449128 | 0.14032288 | 0.58465839 | 0.41621805 | 0.81211849 | 0.88731158 | AHP1 | XII | 368781 | 369311 | 1 |
| YLR110C | 292.164755 | -0.5698005 | 0.52765901 | 1.6153856 | 0.44588563 | 0.61337082 | CCW12 | XII | 369697 | 370098 | -1 |
| YLR111W | 136.924126 | 1.64200804 | 0.54902293 | 8.78554074 | 0.01236642 | 0.04619457 | NA | XII | 370391 | 370723 | 1 |
| YLR112W | 87.269329 | 2.53415481 | 0.62023743 | 15.5111567 | 0.00042835 | 0.00324208 | NA | XII | 370791 | 371210 | 1 |
| YLR113W | 276.476565 | 1.79859352 | 0.49204535 | 13.1040652 | 0.00142721 | 0.00880976 | HOG1 | XII | 371620 | 372927 | 1 |
| YLR114C | 88.2312069 | -0.3697599 | 0.62411155 | 0.74816734 | 0.68791935 | 0.81022234 | AVL9 | XII | 374944 | 377238 | -1 |
| YLR115W | 33.0740118 | 0.80634563 | 0.68276224 | 2.32539197 | 0.31264217 | 0.4849178 | CFT2 | XII | 377985 | 380564 | 1 |
| YLR116W | 14.3004611 | -0.6067284 | 0.82276724 | 0.83951872 | 0.65720495 | 0.78549013 | MSL5 | XII | 380822 | 382252 | 1 |
| YLR117C | 72.2386278 | -1.5391146 | 0.55827901 | 7.89299487 | 0.01932226 | 0.06448178 | CLF1 | XII | 382471 | 384534 | -1 |
| YLR118C | 10.6035215 | 0.27569054 | 0.89556141 | 0.41780203 | 0.81147556 | 0.88731158 | NA | XII | 384725 | 385408 | -1 |
| YLR119W | 685.564241 | 1.5943492 | 0.68617247 | 24.2829662 | 5.33E-06 | 7.55E-05 | SRN2 | XII | 385534 | 386175 | 1 |
| YLR120C | 113.47093 | -1.0792655 | 0.61646731 | 3.63483103 | 0.16244505 | 0.31103069 | YPS1 | XII | 386511 | 388220 | -1 |
| YLR121C | 1135.88786 | -1.2811664 | 0.47744887 | 8.22628269 | 0.01635631 | 0.05698138 | YPS3 | XII | 388744 | 390270 | -1 |

| | | | | | | | | | | | |
|---|---|---|---|---|---|---|---|---|---|---|---|
| YLR122C | 61.6740397 | 2.04522899 | 0.59765763 | 14.653986 | 0.00065755 | 0.00463727 | NA | XII | 390954 | 391331 | -1 |
| YLR126C | 177.381549 | -1.913476 | 0.55495071 | 11.3663124 | 0.0034028 | 0.01682512 | NA | XII | 394765 | 395520 | -1 |
| YLR127C | 322.108027 | -1.5249862 | 0.50714155 | 8.76274843 | 0.01250816 | 0.04666381 | APC2 | XII | 395758 | 398319 | -1 |
| YLR128W | 12.2940139 | 1.78945443 | 0.90384447 | 3.98333191 | 0.13646789 | 0.27378692 | DCN1 | XII | 398530 | 399433 | 1 |
| YLR129W | 67.6394381 | 0.51174675 | 0.57556871 | 3.01605084 | 0.22134661 | 0.38636302 | DIP2 | XII | 399657 | 402488 | 1 |
| YLR130C | 99.416519 | 1.1049883 | 0.57554794 | 11.4232031 | 0.00330737 | 0.01643749 | ZRT2 | XII | 402794 | 404062 | -1 |
| YLR131C | 276.934895 | 3.66756542 | 0.50332105 | 57.2043708 | 3.79E-13 | 2.85E-11 | ACE2 | XII | 404510 | 406822 | -1 |
| YLR132C | 30.5154113 | 1.40143229 | 0.77131136 | 8.89086521 | 0.01173203 | 0.04440749 | USB1 | XII | 407283 | 408155 | -1 |
| YLR133W | 18.6119003 | 0.40910245 | 0.76188397 | 0.3295122 | 0.84810053 | 0.90980423 | CKI1 | XII | 408445 | 410193 | 1 |
| YLR134W | 26.4535207 | -1.7470037 | 0.86912436 | 4.40601004 | 0.11047069 | 0.2353294 | PDC5 | XII | 410723 | 412414 | 1 |
| YLR135W | 173.742137 | -0.6487813 | 0.51499892 | 3.07908586 | 0.21447911 | 0.37828653 | SLX4 | XII | 413281 | 415527 | 1 |
| YLR136C | 61.8370891 | -1.2320516 | 0.62313488 | 3.91693547 | 0.14107442 | 0.28002087 | TIS11 | XII | 415801 | 416658 | -1 |
| YLR137W | 19.2836401 | 0.17087421 | 0.72021614 | 0.07200779 | 0.96463654 | 0.97388763 | RKM5 | XII | 417006 | 418109 | 1 |
| YLR138W | 213.049736 | 0.59925366 | 0.54474257 | 6.130323 | 0.04664631 | 0.12398451 | NHA1 | XII | 418437 | 421394 | 1 |
| YLR139C | 170.319651 | 2.2248305 | 0.73483787 | 8.67269485 | 0.01308423 | 0.04820096 | SLS1 | XII | 421542 | 423473 | -1 |
| YLR141W | 313.645258 | 0.01151763 | 0.49037079 | 0.2171266 | 0.89712211 | 0.9367605 | RRN5 | XII | 423683 | 424774 | 1 |
| YLR142W | 51.3655005 | -0.2854554 | 0.55619983 | 3.24476544 | 0.19742772 | 0.35555413 | PUT1 | XII | 425186 | 426616 | 1 |
| YLR143W | 38.7933049 | -0.30979 | 0.63609135 | 0.79068995 | 0.67344768 | 0.7980479 | DPH6 | XII | 427329 | 429386 | 1 |
| YLR144C | 125.370496 | 0.45211549 | 0.51734992 | 1.71078614 | 0.42511606 | 0.59512137 | ACF2 | XII | 429677 | 432016 | -1 |
| YLR145W | 191.229209 | 0.01653331 | 0.84338949 | 5.47753579 | 0.06464995 | 0.15847724 | RMP1 | XII | 432168 | 432773 | 1 |
| YLR146C | 53.060647 | 1.65583352 | 0.96446752 | 4.72787077 | 0.09404937 | 0.2093602 | SPE4 | XII | 432823 | 433725 | -1 |
| YLR146W-A | 82.7316951 | -0.0753401 | 0.78187517 | 3.50904131 | 0.17299014 | 0.32456673 | NA | XII | 433870 | 434058 | 1 |
| YLR147C | 58.0255561 | 2.08267778 | 0.69179741 | 8.70801683 | 0.01285518 | 0.04759047 | SMD3 | XII | 434158 | 434463 | -1 |
| YLR148W | 12.8749628 | -0.515596 | 0.92291854 | 0.46533851 | 0.79241562 | 0.87392122 | PEP3 | XII | 434641 | 437397 | 1 |
| YLR149C | 113.20265 | -2.0066471 | 0.5402387 | 13.7318785 | 0.0010427 | 0.00682175 | NA | XII | 437631 | 439823 | -1 |
| YLR150W | 356.670296 | -2.9563066 | 0.66376915 | 17.6137065 | 0.0001497 | 0.00133351 | STM1 | XII | 440467 | 441288 | 1 |
| YLR151C | 116.503544 | -1.6758678 | 0.55969213 | 8.82582583 | 0.01211982 | 0.04556739 | PCD1 | XII | 441715 | 442737 | -1 |
| YLR152C | 40.3578119 | 1.30352579 | 0.63272043 | 6.78951472 | 0.03354869 | 0.09746459 | NA | XII | 442958 | 444688 | -1 |
| YLR153C | 78.9372971 | -1.456348 | 0.75893949 | 3.77282374 | 0.15161485 | 0.29497647 | ACS2 | XII | 445524 | 447575 | -1 |
| YLR154W-C | 1172.96589 | -1.3337863 | 0.63989666 | 5.46331853 | 0.06511116 | 0.15927065 | TAR1 | XII | 454696 | 455070 | 1 |
| YLR157W-E | 5.19751994 | 0.12498721 | 1.16438797 | 0.8485003 | 0.6542602 | 0.78442961 | NA | XII | 481873 | 482037 | 1 |
| YLR162W | 9.2268746 | 0.29739794 | 1.01733077 | 0.13494561 | 0.93475314 | 0.95774565 | NA | XII | 489573 | 489929 | 1 |
| YLR162W-A | 25.3462273 | -1.0086774 | 1.01657436 | 1.02714206 | 0.59835502 | 0.74552949 | RRT15 | XII | 490406 | 490594 | 1 |

| | | | | | | | | | | | |
|---|---|---|---|---|---|---|---|---|---|---|---|
| YLR163C | 137.951094 | -0.9070275 | 0.53421596 | 3.35176961 | 0.18714252 | 0.34300576 | MAS1 | XII | 491866 | 493254 | -1 |
| YLR164W | 106.040278 | 1.91847709 | 0.52295067 | 12.8568539 | 0.00161499 | 0.00961783 | SHH4 | XII | 493883 | 494389 | 1 |
| YLR165C | 44.551342 | 0.12046368 | 0.62015889 | 0.85715935 | 0.65143368 | 0.78236902 | PUS5 | XII | 494494 | 495258 | -1 |
| YLR166C | 33.1751793 | 0.67569523 | 0.6274005 | 1.24331741 | 0.53705289 | 0.69177669 | SEC10 | XII | 495429 | 498044 | -1 |
| YLR167W | 375.698986 | -3.3483911 | 0.67883827 | 20.8011501 | 3.04E-05 | 0.00034284 | RPS31 | XII | 498947 | 499405 | 1 |
| YLR168C | 10.7905189 | -1.5078717 | 0.93395718 | 3.06839262 | 0.21562892 | 0.37948068 | UPS2 | XII | 499578 | 500270 | -1 |
| YLR170C | 5.91356301 | -0.6511782 | 1.19033342 | 0.34033696 | 0.84352269 | 0.90831251 | APS1 | XII | 500579 | 501049 | -1 |
| YLR172C | 221.614274 | -0.0134422 | 0.50254152 | 0.1745274 | 0.9164354 | 0.94860224 | DPH5 | XII | 501260 | 502162 | -1 |
| YLR173W | 24.5198435 | 0.01701836 | 0.69320071 | 0.1264541 | 0.93873031 | 0.95995205 | NA | XII | 502421 | 504247 | 1 |
| YLR174W | 13.0328604 | 0.10318904 | 0.87359339 | 0.39818597 | 0.81947369 | 0.89171017 | IDP2 | XII | 504591 | 505829 | 1 |
| YLR175W | 671.100636 | -2.360692 | 0.50567113 | 19.9113077 | 4.75E-05 | 0.00049245 | CBF5 | XII | 506134 | 507585 | 1 |
| YLR176C | 1277.10118 | 5.45523021 | 0.57502477 | 66.4598405 | 3.70E-15 | 4.02E-13 | RFX1 | XII | 507797 | 510232 | -1 |
| YLR177W | 20.2273986 | 0.63142086 | 0.73867501 | 1.73678239 | 0.4196261 | 0.59032562 | NA | XII | 511054 | 512940 | 1 |
| YLR178C | 23.4027875 | 1.11589515 | 0.74514217 | 2.40339551 | 0.30068329 | 0.47389383 | TFS1 | XII | 513162 | 513821 | -1 |
| YLR179C | 177.429136 | 0.73785906 | 0.50473335 | 3.15370603 | 0.20662432 | 0.36867637 | NA | XII | 514108 | 514713 | -1 |
| YLR180W | 51.5682664 | -3.7984444 | 0.7225971 | 25.6438004 | 2.70E-06 | 4.19E-05 | SAM1 | XII | 515262 | 516410 | 1 |
| YLR181C | 108.914756 | -1.804002 | 0.78737665 | 9.40804597 | 0.00905876 | 0.03655068 | VTA1 | XII | 516678 | 517670 | -1 |
| YLR182W | 185.237221 | 4.41873707 | 0.59132799 | 74.465704 | 6.76E-17 | 9.79E-15 | SWI6 | XII | 517940 | 520351 | 1 |
| YLR183C | 16.345671 | -0.6456389 | 1.00588875 | 3.50022178 | 0.17375467 | 0.32507061 | TOS4 | XII | 520543 | 522012 | -1 |
| YLR185W | 42.4033398 | -2.8375844 | 0.76937932 | 12.5981378 | 0.00183802 | 0.01052632 | RPL37A | XII | 522663 | 523288 | 1 |
| YLR186W | 30.8537645 | -0.9786333 | 0.74726648 | 1.88094466 | 0.39044337 | 0.56165643 | EMG1 | XII | 523632 | 524390 | 1 |
| YLR187W | 29.1180825 | 0.11629793 | 0.63786481 | 0.59199366 | 0.74378979 | 0.8437584 | SKG3 | XII | 524865 | 527945 | 1 |
| YLR188W | 158.150221 | 1.57403743 | 0.53873094 | 8.17614244 | 0.01677155 | 0.05813116 | MDL1 | XII | 528300 | 530387 | 1 |
| YLR189C | 173.802973 | -0.7150256 | 0.53994676 | 3.28505471 | 0.19349041 | 0.35020614 | ATG26 | XII | 530797 | 534393 | -1 |
| YLR190W | 86.594231 | 2.09667187 | 0.57908061 | 17.482366 | 0.00015986 | 0.00140458 | MMR1 | XII | 535214 | 536689 | 1 |
| YLR191W | 48.2611282 | 1.89183858 | 0.59435588 | 12.1615797 | 0.00228637 | 0.01234789 | PEX13 | XII | 537272 | 538432 | 1 |
| YLR192C | 150.34374 | 0.60407031 | 0.54138345 | 1.98934581 | 0.3698444 | 0.54308878 | HCR1 | XII | 538792 | 539589 | -1 |
| YLR193C | 28.8743098 | -0.7264785 | 0.67230939 | 1.56213067 | 0.45791792 | 0.62324553 | UPS1 | XII | 540009 | 540536 | -1 |
| YLR194C | 648.95072 | 0.05967544 | 0.46159745 | 3.62938565 | 0.16288794 | 0.31167255 | NCW2 | XII | 540809 | 541573 | -1 |
| YLR195C | 1775.96751 | 4.89492571 | 0.47443069 | 80.5675993 | 3.20E-18 | 5.97E-16 | NMT1 | XII | 541937 | 543304 | -1 |
| YLR196W | 51.3886145 | -1.6407156 | 0.63129152 | 6.60130057 | 0.03685919 | 0.10460433 | PWP1 | XII | 543968 | 545698 | 1 |
| YLR197W | 496.519433 | -3.6686036 | 0.59104199 | 31.7971285 | 1.25E-07 | 2.91E-06 | NOP56 | XII | 546097 | 547611 | 1 |
| YLR199C | 50.2445665 | 0.66733358 | 0.57051186 | 3.03135183 | 0.21965966 | 0.3842385 | PBA1 | XII | 547853 | 548769 | -1 |

| | | | | | | | | | | | |
|---|---|---|---|---|---|---|---|---|---|---|---|
| YLR200W | 34.1807421 | -0.0310192 | 0.7237764 | 2.33914842 | 0.31049912 | 0.48327686 | YKE2 | XII | 549012 | 549356 | 1 |
| YLR201C | 151.013364 | -0.6360065 | 0.66219888 | 5.69481034 | 0.05799461 | 0.14626659 | COQ9 | XII | 549511 | 550293 | -1 |
| YLR203C | 27.4399533 | 0.40781224 | 0.68168729 | 3.60373648 | 0.16499036 | 0.31392516 | MSS51 | XII | 550650 | 551960 | -1 |
| YLR204W | 52.347963 | -0.5579363 | 0.66837653 | 3.47923706 | 0.17558737 | 0.32742379 | QRI5 | XII | 552270 | 552605 | 1 |
| YLR205C | 296.089123 | 0.73229862 | 0.84745353 | 5.47335179 | 0.06478534 | 0.15867476 | HMX1 | XII | 552724 | 553677 | -1 |
| YLR206W | 206.367522 | 1.74143257 | 0.79454086 | 12.7894361 | 0.00167036 | 0.00979022 | ENT2 | XII | 554578 | 556419 | 1 |
| YLR207W | 34.3457974 | 0.73966713 | 0.62911339 | 1.4099675 | 0.4941166 | 0.65469026 | HRD3 | XII | 556788 | 559289 | 1 |
| YLR208W | 20.6655441 | -0.4794595 | 0.79888838 | 0.70967669 | 0.7012868 | 0.81814438 | SEC13 | XII | 559551 | 560444 | 1 |
| YLR209C | 134.993546 | -1.1881772 | 0.87128962 | 12.856207 | 0.00161551 | 0.00961783 | PNP1 | XII | 560797 | 561732 | -1 |
| YLR210W | 7.50055348 | -1.0764566 | 1.01502117 | 1.18144941 | 0.55392571 | 0.70568075 | CLB4 | XII | 562008 | 563390 | 1 |
| YLR212C | 23.0705508 | -1.0770369 | 0.71295766 | 2.24654081 | 0.32521447 | 0.49807536 | TUB4 | XII | 564860 | 566281 | -1 |
| YLR213C | 54.087663 | -1.0584069 | 0.60384339 | 4.29708861 | 0.11665385 | 0.24436533 | CRR1 | XII | 566654 | 567922 | -1 |
| YLR214W | 254.994056 | 1.23778578 | 0.73322505 | 6.41890117 | 0.04037879 | 0.11218484 | FRE1 | XII | 568567 | 570627 | 1 |
| YLR215C | 128.644294 | -1.4373756 | 0.62200136 | 6.22443179 | 0.04450223 | 0.11990132 | CDC123 | XII | 570776 | 571858 | -1 |
| YLR216C | 45.3891375 | 0.38235664 | 0.67553729 | 0.4533548 | 0.79717791 | 0.87733513 | CPR6 | XII | 572096 | 573211 | -1 |
| YLR218C | 109.388357 | 1.88794404 | 0.65525644 | 7.99647522 | 0.01834795 | 0.06205293 | NA | XII | 573466 | 573918 | -1 |
| YLR219W | 53.1535548 | 0.56833011 | 0.63635 | 3.27483231 | 0.19448191 | 0.35156111 | MSC3 | XII | 574151 | 576337 | 1 |
| YLR220W | 68.1365476 | 1.48168465 | 0.58053803 | 10.9078426 | 0.00427949 | 0.02021064 | CCC1 | XII | 576825 | 577793 | 1 |
| YLR221C | 25.7518269 | -2.4222163 | 0.72441147 | 11.0789508 | 0.00392859 | 0.01897124 | RSA3 | XII | 578362 | 579024 | -1 |
| YLR222C | 21.6798531 | -0.1005458 | 0.83831148 | 1.20005844 | 0.5487956 | 0.70069987 | UTP13 | XII | 579318 | 581771 | -1 |
| YLR223C | 117.493816 | -0.0723984 | 0.51198608 | 0.23523968 | 0.88903396 | 0.93304075 | IFH1 | XII | 582233 | 585490 | -1 |
| YLR224W | 311.442286 | 3.27238761 | 1.86054521 | 5.72046724 | 0.05725538 | 0.14501691 | UCC1 | XII | 586464 | 587573 | 1 |
| YLR225C | 75.1847186 | 1.29213215 | 0.77337906 | 14.0600057 | 0.00088493 | 0.00595552 | NA | XII | 587695 | 588918 | -1 |
| YLR226W | 61.8471132 | -0.3856456 | 0.56212599 | 1.98322148 | 0.37097866 | 0.5443402 | BUR2 | XII | 589354 | 590541 | 1 |
| YLR227C | 1287.92298 | -0.6503736 | 0.46965844 | 1.94243595 | 0.3786216 | 0.55247457 | ADY4 | XII | 590562 | 592043 | -1 |
| YLR227W-B | 5.67558581 | -1.2087323 | 1.2376494 | 1.01941435 | 0.60067144 | 0.74682758 | NA | XII | 593438 | 598706 | 1 |
| YLR228C | 20.9775114 | 0.41835302 | 0.697702 | 0.37584617 | 0.82867844 | 0.89767038 | ECM22 | XII | 600019 | 602463 | -1 |
| YLR229C | 50.9993477 | -0.740821 | 0.63854961 | 3.35044567 | 0.18726645 | 0.34312428 | CDC42 | XII | 604212 | 604787 | -1 |
| YLR231C | 35.9224423 | -0.1402451 | 0.62123451 | 0.65769419 | 0.71975306 | 0.82982282 | BNA5 | XII | 605758 | 607119 | -1 |
| YLR233C | 241.595612 | 1.42880569 | 0.49350415 | 10.0092512 | 0.00670685 | 0.02902292 | EST1 | XII | 607425 | 609524 | -1 |
| YLR234W | 170.08878 | -0.2783007 | 0.56114996 | 0.24656725 | 0.8840129 | 0.9315066 | TOP3 | XII | 609783 | 611753 | 1 |
| YLR236C | 189.380592 | -0.369626 | 0.58885242 | 2.8003 | 0.24655998 | 0.41766412 | NA | XII | 611794 | 612117 | -1 |
| YLR237W | 210.595195 | 2.05820128 | 0.56549842 | 12.6274047 | 0.00181131 | 0.01040428 | THI7 | XII | 612367 | 614163 | 1 |

| | | | | | | | | | | | |
|---|---|---|---|---|---|---|---|---|---|---|---|
| YLR238W | 20.0029686 | -1.5215306 | 0.7840715 | 4.16531741 | 0.1245985 | 0.25673499 | FAR10 | XII | 614755 | 616191 | 1 |
| YLR239C | 6.29526663 | -0.5676672 | 1.27123586 | 1.23079571 | 0.54042584 | 0.69442202 | LIP2 | XII | 616332 | 617318 | -1 |
| YLR240W | 289.453827 | 2.05981404 | 0.51683861 | 16.0510516 | 0.00032701 | 0.00262969 | VPS34 | XII | 617533 | 620160 | 1 |
| YLR241W | 694.95083 | 3.25669509 | 0.49338173 | 67.7533773 | 1.94E-15 | 2.20E-13 | CSC1 | XII | 620473 | 622821 | 1 |
| YLR242C | 441.657544 | 2.09302294 | 0.88896657 | 13.4230038 | 0.00121684 | 0.0077508 | ARV1 | XII | 622918 | 623883 | -1 |
| YLR243W | 20.8778414 | 0.27760199 | 0.70603933 | 2.00287092 | 0.36735174 | 0.54052518 | GPN3 | XII | 624203 | 625021 | 1 |
| YLR244C | 110.156106 | -1.1760271 | 0.7588112 | 6.00150716 | 0.04974956 | 0.13016266 | MAP1 | XII | 625168 | 626331 | -1 |
| YLR245C | 89.7222558 | -3.1856505 | 0.60887173 | 27.6086917 | 1.01E-06 | 1.79E-05 | CDD1 | XII | 626502 | 626930 | -1 |
| YLR246W | 135.820924 | 5.40231635 | 0.64839132 | 70.5312315 | 4.83E-16 | 6.08E-14 | ERF2 | XII | 627118 | 628197 | 1 |
| YLR247C | 49.5707201 | 0.87658075 | 0.66148977 | 1.79899013 | 0.406775 | 0.57811175 | IRC20 | XII | 628684 | 633354 | -1 |
| YLR248W | 140.207065 | 2.8898281 | 0.52372714 | 30.2446738 | 2.71E-07 | 5.70E-06 | RCK2 | XII | 634252 | 636084 | 1 |
| YLR249W | 719.903702 | -2.4359706 | 0.60317387 | 15.1642588 | 0.00050948 | 0.0037336 | YEF3 | XII | 636780 | 639914 | 1 |
| YLR250W | 54.0795459 | 1.36270757 | 0.80555972 | 2.8800703 | 0.23691943 | 0.40475196 | SSP120 | XII | 640317 | 641021 | 1 |
| YLR251W | 154.186411 | -1.9767693 | 0.50919388 | 14.6725527 | 0.00065147 | 0.00461127 | SYM1 | XII | 641464 | 642057 | 1 |
| YLR253W | 138.062796 | 0.92722552 | 0.69978544 | 4.28450053 | 0.11739039 | 0.24546419 | MCP2 | XII | 642627 | 644336 | 1 |
| YLR254C | 74.8341915 | -0.3258566 | 0.7208692 | 4.23988559 | 0.1200385 | 0.24929085 | NDL1 | XII | 644404 | 644973 | -1 |
| YLR255C | 34.1392086 | -0.5312557 | 0.81236237 | 1.24903548 | 0.53551963 | 0.69010876 | NA | XII | 645600 | 645953 | -1 |
| YLR256W | 648.155449 | -1.4586866 | 0.54366663 | 7.079381 | 0.02902231 | 0.08675228 | HAP1 | XII | 646415 | 650923 | 1 |
| YLR256W-A | 231.759355 | -0.3920769 | 0.52388022 | 1.37478687 | 0.50288516 | 0.66130027 | NA | XII | 651117 | 652439 | 1 |
| YLR257W | 142.556692 | 0.88001168 | 0.57319741 | 3.832509 | 0.14715711 | 0.28912102 | NA | XII | 658826 | 659791 | 1 |
| YLR258W | 127.354862 | 0.24870844 | 0.50190055 | 3.74359195 | 0.15384711 | 0.29821719 | GSY2 | XII | 660716 | 662833 | 1 |
| YLR259C | 95.156923 | -0.33088 | 0.71316989 | 0.24142623 | 0.88628818 | 0.93215081 | HSP60 | XII | 663284 | 665002 | -1 |
| YLR260W | 186.661515 | 1.16912567 | 0.5332499 | 11.181854 | 0.00373157 | 0.01817138 | LCB5 | XII | 665844 | 667907 | 1 |
| YLR262C-A | 36.9447575 | -1.0790805 | 0.78427332 | 2.55127162 | 0.27925336 | 0.45351049 | TMA7 | XII | 669468 | 669662 | -1 |
| YLR263W | 119.049387 | -2.0016138 | 0.50538327 | 15.8988759 | 0.00035286 | 0.00279489 | RED1 | XII | 670340 | 672823 | 1 |
| YLR264C-A | 19804.6944 | -3.3993751 | 0.52072892 | 36.9086183 | 9.67E-09 | 2.90E-07 | NA | XII | 673828 | 673944 | -1 |
| YLR264W | 16.3651907 | -2.1288155 | 0.97160029 | 5.22514296 | 0.07334569 | 0.17466992 | RPS28B | XII | 673131 | 673334 | 1 |
| YLR265C | 83.9125608 | -2.176807 | 0.54702779 | 17.5316437 | 0.00015597 | 0.00137666 | NEJ1 | XII | 674427 | 675455 | -1 |
| YLR266C | 1349.55904 | -1.0717111 | 0.46617993 | 5.22442568 | 0.073372 | 0.17466992 | PDR8 | XII | 675619 | 677724 | -1 |
| YLR267W | 428.704063 | -1.9459333 | 0.47161896 | 16.2568327 | 0.00029504 | 0.0024162 | BOP2 | XII | 678212 | 679924 | 1 |
| YLR268W | 27.3274348 | -2.1935943 | 0.71828627 | 9.66550595 | 0.00796456 | 0.03310469 | SEC22 | XII | 680200 | 680844 | 1 |
| YLR270W | 12.2292995 | 1.25778311 | 1.01203819 | 1.65198617 | 0.4378 | 0.60642632 | DCS1 | XII | 681186 | 682238 | 1 |
| YLR271W | 87.8881142 | -2.3305718 | 0.55326422 | 16.8445262 | 0.00021992 | 0.001854 | CMG1 | XII | 682737 | 683561 | 1 |

| | | | | | | | | | | | |
|---|---|---|---|---|---|---|---|---|---|---|---|
| YLR272C | 235.979517 | -0.433383 | 0.50438634 | 0.80441453 | 0.6688421 | 0.7945416 | YCS4 | XII | 683672 | 687202 | -1 |
| YLR273C | 20.7762509 | 1.2496919 | 0.73139286 | 9.06414335 | 0.01075837 | 0.04161051 | PIG1 | XII | 689083 | 691029 | -1 |
| YLR274W | 123.651581 | 4.33062393 | 0.6113937 | 69.7782342 | 7.04E-16 | 8.50E-14 | MCM5 | XII | 691555 | 693882 | 1 |
| YLR275W | 7.32997374 | -1.6347844 | 1.07367824 | 2.54019423 | 0.28080435 | 0.45491247 | SMD2 | XII | 694378 | 694800 | 1 |
| YLR276C | 93.884684 | -1.111438 | 0.83083514 | 4.4051906 | 0.11051596 | 0.23533925 | DBP9 | XII | 695046 | 696830 | -1 |
| YLR277C | 11.8517903 | -0.5897156 | 1.00371077 | 1.01973503 | 0.60057514 | 0.74682758 | YSH1 | XII | 697156 | 699495 | -1 |
| YLR278C | 848.791947 | -0.5827474 | 0.48220955 | 2.04035597 | 0.36053077 | 0.53345156 | NA | XII | 699999 | 704024 | -1 |
| YLR281C | 28.0199004 | -0.5640015 | 0.68143988 | 1.22647853 | 0.54159366 | 0.69561386 | NA | XII | 704493 | 704960 | -1 |
| YLR283W | 17.8299394 | -0.1374641 | 0.82415742 | 0.91892385 | 0.63162341 | 0.76880708 | NA | XII | 705186 | 706130 | 1 |
| YLR284C | 12.7107304 | -1.925307 | 0.96326156 | 6.61237191 | 0.03665571 | 0.10424194 | ECI1 | XII | 706198 | 707040 | -1 |
| YLR285C-A | 675.63591 | -2.20454 | 0.53240949 | 19.4412923 | 6.00E-05 | 0.0006041 | NA | XII | 708168 | 708338 | -1 |
| YLR285W | 125.000001 | -0.2401202 | 0.5027965 | 0.29494175 | 0.86288757 | 0.91958753 | NNT1 | XII | 707360 | 708145 | 1 |
| YLR286C | 162.40594 | -1.4139325 | 0.49404648 | 9.00077405 | 0.0111047 | 0.04258027 | CTS1 | XII | 708448 | 710136 | -1 |
| YLR287C | 40.8955548 | -0.5703691 | 0.60831603 | 1.73856023 | 0.41925326 | 0.59032562 | NA | XII | 710991 | 712058 | -1 |
| YLR287C-A | 143.761962 | -2.2485796 | 0.77833929 | 7.77743391 | 0.0204716 | 0.06719418 | RPS30A | XII | 712537 | 713158 | -1 |
| YLR288C | 38.8439112 | -0.4680007 | 0.5988193 | 0.6809587 | 0.71142922 | 0.82438173 | MEC3 | XII | 713480 | 714904 | -1 |
| YLR289W | 29.2050879 | -0.184346 | 0.76092169 | 1.88660334 | 0.38934024 | 0.56101577 | GUF1 | XII | 715089 | 717026 | 1 |
| YLR290C | 25.7809314 | 0.43359521 | 0.68815604 | 0.4022548 | 0.81780823 | 0.89056041 | COQ11 | XII | 717145 | 717978 | -1 |
| YLR291C | 15.0142801 | 0.21441357 | 0.86951944 | 0.32242465 | 0.85111134 | 0.91173629 | GCD7 | XII | 718317 | 719462 | -1 |
| YLR292C | 30.510899 | 0.58828626 | 0.87083305 | 0.56823596 | 0.75267784 | 0.85034238 | SEC72 | XII | 719789 | 720370 | -1 |
| YLR293C | 73.8217827 | -1.9608168 | 0.69154656 | 8.1311459 | 0.01715316 | 0.05904684 | GSP1 | XII | 720771 | 721430 | -1 |
| YLR295C | 18.9681436 | -2.7640312 | 0.7953732 | 12.4597866 | 0.00196966 | 0.01104002 | ATP14 | XII | 721999 | 722373 | -1 |
| YLR296W | 6.55619836 | 0.28062743 | 1.08661726 | 1.75547948 | 0.41572149 | 0.58736638 | NA | XII | 722978 | 723304 | 1 |
| YLR297W | 800.522384 | 3.50160031 | 0.53103187 | 43.3073909 | 3.94E-10 | 1.61E-08 | NA | XII | 724044 | 724433 | 1 |
| YLR298C | 154.999774 | 1.56593464 | 0.50862768 | 9.82476557 | 0.00735494 | 0.03119789 | YHC1 | XII | 724721 | 725416 | -1 |
| YLR299W | 255.76528 | 4.71743022 | 0.60036425 | 60.6769801 | 6.67E-14 | 5.76E-12 | ECM38 | XII | 726069 | 728051 | 1 |
| YLR300W | 201.328489 | 0.25313705 | 0.76973937 | 0.87717147 | 0.6449479 | 0.77748248 | EXG1 | XII | 728955 | 730301 | 1 |
| YLR301W | 29.7894719 | 0.48863995 | 0.66073373 | 0.73876911 | 0.69115957 | 0.81221741 | HRI1 | XII | 730825 | 731559 | 1 |
| YLR302C | 717.643346 | -1.7832658 | 0.71803901 | 5.93114773 | 0.05153089 | 0.13385547 | NA | XII | 732191 | 732553 | -1 |
| YLR303W | 46.0839993 | -0.5401579 | 0.58534391 | 1.23016112 | 0.54059734 | 0.69448827 | MET17 | XII | 732542 | 733876 | 1 |
| YLR304C | 46.5930886 | -1.6951487 | 0.65884236 | 6.67410748 | 0.03554152 | 0.10170914 | ACO1 | XII | 735212 | 737548 | -1 |
| YLR305C | 20.1229581 | -1.5234154 | 0.74199754 | 4.45743308 | 0.10766653 | 0.2307994 | STT4 | XII | 738161 | 743863 | -1 |
| YLR306W | 11.3990355 | -0.4369855 | 0.90365579 | 0.62747756 | 0.73070988 | 0.83678999 | UBC12 | XII | 744151 | 744851 | 1 |

| | | | | | | | | | | | |
|---|---|---|---|---|---|---|---|---|---|---|---|
| YLR308W | 29.7025968 | -0.626029 | 0.84585023 | 2.06522705 | 0.35607513 | 0.52942744 | CDA2 | XII | 747937 | 748875 | 1 |
| YLR309C | 110.525508 | -0.2289404 | 0.55836308 | 1.09177387 | 0.57932773 | 0.72802613 | IMH1 | XII | 749034 | 751769 | -1 |
| YLR310C | 43.069659 | -0.2307838 | 0.65037265 | 0.16101311 | 0.92264885 | 0.95140461 | CDC25 | XII | 752224 | 756993 | -1 |
| YLR312C | 26.4464701 | -0.4159295 | 0.64744686 | 2.88054341 | 0.23686339 | 0.40475196 | ATG39 | XII | 757637 | 758833 | -1 |
| YLR312W-A | 11.9168189 | -2.277132 | 1.0510183 | 5.52056113 | 0.06327401 | 0.15616221 | MRPL15 | XII | 759480 | 760241 | 1 |
| YLR313C | 18.4485657 | -0.5634544 | 0.73284528 | 1.74047551 | 0.41885195 | 0.59006151 | NA | XII | 760750 | 762342 | -1 |
| YLR314C | 42.8119131 | -0.4008067 | 0.66415656 | 1.60087391 | 0.44913267 | 0.61535214 | CDC3 | XII | 762575 | 764137 | -1 |
| YLR315W | 68.8224074 | -1.4278065 | 0.57266988 | 7.06245681 | 0.02926894 | 0.08719023 | NKP2 | XII | 764808 | 765269 | 1 |
| YLR316C | 1328.8196 | 0.2737608 | 0.51383592 | 0.70666147 | 0.70234487 | 0.81847842 | TAD3 | XII | 765266 | 766358 | -1 |
| YLR318W | 45.2811153 | -0.5727535 | 0.91581305 | 3.72011618 | 0.15566359 | 0.30083183 | EST2 | XII | 766542 | 769196 | 1 |
| YLR319C | 26.0446135 | -1.0586137 | 0.94782296 | 1.54880253 | 0.4609797 | 0.62486036 | BUD6 | XII | 769318 | 771684 | -1 |
| YLR320W | 80.5201251 | -1.3265172 | 0.58871612 | 8.24626336 | 0.01619372 | 0.05674873 | MMS22 | XII | 771940 | 776304 | 1 |
| YLR321C | 109.026833 | 0.6114214 | 0.63807933 | 2.88141496 | 0.2367602 | 0.40473806 | SFH1 | XII | 776584 | 777864 | -1 |
| YLR323C | 15.6343226 | -0.8280564 | 0.90024966 | 2.95914199 | 0.22773537 | 0.39372582 | CWC24 | XII | 778173 | 778952 | -1 |
| YLR324W | 96.8263752 | -2.4303565 | 0.68488715 | 12.3402019 | 0.00209102 | 0.0115746 | PEX30 | XII | 779215 | 780786 | 1 |
| YLR325C | 82.938319 | -2.4781936 | 0.66918208 | 12.98018 | 0.00151841 | 0.00921552 | RPL38 | XII | 781143 | 781379 | -1 |
| YLR326W | 20.5301731 | 1.00962399 | 0.73835097 | 3.81304897 | 0.14859594 | 0.29099512 | NA | XII | 782174 | 782896 | 1 |
| YLR327C | 18.2925609 | 3.09453055 | 0.96386442 | 11.5892557 | 0.00304386 | 0.01538156 | TMA10 | XII | 783127 | 783387 | -1 |
| YLR328W | 22.4730086 | -1.1391405 | 0.73325813 | 2.83726612 | 0.24204465 | 0.41122023 | NMA1 | XII | 784913 | 786118 | 1 |
| YLR329W | 153.122859 | -0.1934602 | 0.51365279 | 0.23169318 | 0.89061184 | 0.93317728 | REC102 | XII | 786442 | 787333 | 1 |
| YLR330W | 140.004968 | 0.16487546 | 0.53259031 | 2.94558577 | 0.22928423 | 0.39593071 | CHS5 | XII | 787664 | 789679 | 1 |
| YLR331C | 9223.46821 | 0.71688287 | 0.76249298 | 7.48541448 | 0.02368988 | 0.07502674 | NA | XII | 790669 | 791046 | -1 |
| YLR332W | 42.1062734 | 2.45435487 | 0.67648367 | 12.5874319 | 0.00184788 | 0.01056192 | MID2 | XII | 790676 | 791806 | 1 |
| YLR333C | 56.1160416 | -3.1899886 | 0.83595132 | 13.4171334 | 0.00122041 | 0.00776032 | RPS25B | XII | 795573 | 795899 | -1 |
| YLR335W | 124.261284 | 1.16979581 | 0.62802713 | 3.62251491 | 0.16344848 | 0.3120399 | NUP2 | XII | 797430 | 799592 | 1 |
| YLR336C | 63.9616569 | 0.44443088 | 0.67353899 | 0.6945959 | 0.70659477 | 0.82201802 | SGD1 | XII | 799697 | 802396 | -1 |
| YLR337C | 270.572216 | 3.04290877 | 0.55031463 | 28.2257013 | 7.43E-07 | 1.38E-05 | VRP1 | XII | 802653 | 805106 | -1 |
| YLR340W | 111.37672 | -2.2425857 | 0.60828368 | 13.2990529 | 0.00129464 | 0.00812103 | RPP0 | XII | 805887 | 806825 | 1 |
| YLR341W | 438.074077 | -1.8432294 | 0.48598805 | 13.6009588 | 0.00111324 | 0.00718581 | SPO77 | XII | 807385 | 808818 | 1 |
| YLR342W | 180.353166 | -0.7000468 | 0.56126614 | 6.23754315 | 0.04421145 | 0.11945136 | FKS1 | XII | 809997 | 815627 | 1 |
| YLR343W | 37.811294 | -0.5143905 | 0.65872909 | 2.41602404 | 0.29879068 | 0.47191811 | GAS2 | XII | 816094 | 817761 | 1 |
| YLR344W | 31.4887038 | -2.0243087 | 0.65825592 | 9.17163074 | 0.01019543 | 0.04004855 | RPL26A | XII | 819312 | 820142 | 1 |
| YLR345W | 29.7648419 | 2.60987963 | 0.74699671 | 13.0132195 | 0.00149353 | 0.0091027 | NA | XII | 820511 | 822040 | 1 |

| | | | | | | | | | | | |
|---|---|---|---|---|---|---|---|---|---|---|---|
| YLR347C | 121.164709 | 1.51493957 | 0.58466209 | 6.92266851 | 0.03138785 | 0.09258058 | KAP95 | XII | 823828 | 826413 | -1 |
| YLR348C | 735.9817 | 4.82348951 | 0.60232238 | 50.5299894 | 1.07E-11 | 6.20E-10 | DIC1 | XII | 826977 | 827873 | -1 |
| YLR350W | 325.863296 | 0.82404157 | 0.75473116 | 9.16039721 | 0.01025286 | 0.04019232 | ORM2 | XII | 828730 | 829380 | 1 |
| YLR351C | 168.959922 | 0.20450271 | 0.69555757 | 12.3070704 | 0.00212595 | 0.01170082 | NIT3 | XII | 829489 | 830364 | -1 |
| YLR352W | 238.131289 | -0.6526521 | 0.49744883 | 2.05913871 | 0.35716074 | 0.53038232 | NA | XII | 831116 | 833539 | 1 |
| YLR353W | 119.417136 | -0.6994893 | 0.52544229 | 1.83622676 | 0.3992716 | 0.57010668 | BUD8 | XII | 834352 | 836163 | 1 |
| YLR354C | 84.6699874 | -0.7457776 | 0.64139808 | 2.07790003 | 0.353826 | 0.5275953 | TAL1 | XII | 836350 | 837357 | -1 |
| YLR355C | 95.3119312 | -2.4765578 | 0.59837654 | 16.2353517 | 0.00029822 | 0.0024354 | ILV5 | XII | 838066 | 839253 | -1 |
| YLR356W | 39.8683312 | 0.5450966 | 0.61638656 | 2.09754991 | 0.3503667 | 0.523786 | ATG33 | XII | 840321 | 840914 | 1 |
| YLR357W | 19.0269109 | -0.6133489 | 0.75209946 | 0.70835916 | 0.70174894 | 0.81835374 | RSC2 | XII | 841331 | 844000 | 1 |
| YLR358C | 2052.82584 | -1.097738 | 0.5980428 | 3.48250916 | 0.17530033 | 0.32699386 | NA | XII | 843487 | 844050 | -1 |
| YLR359W | 50.2970492 | -0.895259 | 0.83616386 | 1.70585737 | 0.426165 | 0.59586944 | ADE13 | XII | 844282 | 845730 | 1 |
| YLR360W | 933.104889 | -0.3010725 | 0.4687786 | 1.00139145 | 0.60610883 | 0.75088539 | VPS38 | XII | 846103 | 847422 | 1 |
| YLR361C | 709.266517 | 0.35703455 | 0.65479321 | 2.46829378 | 0.29108298 | 0.46518644 | DCR2 | XII | 847388 | 849124 | -1 |
| YLR361C-A | 32.947318 | 0.10274958 | 0.66602374 | 0.70724354 | 0.70214049 | 0.81847842 | NA | XII | 849383 | 849679 | -1 |
| YLR362W | 25.2618034 | 0.97645506 | 0.79436306 | 2.04497741 | 0.35969864 | 0.53284736 | STE11 | XII | 849866 | 852019 | 1 |
| YLR363C | 13.557029 | 2.23730272 | 0.96148776 | 6.47615951 | 0.03923917 | 0.10991524 | NMD4 | XII | 852495 | 853151 | -1 |
| YLR363W-A | 170.483024 | 1.04121176 | 0.52745761 | 3.81781537 | 0.14824223 | 0.29064152 | NA | XII | 853461 | 853718 | 1 |
| YLR364W | 396.036105 | 1.19992695 | 0.48812598 | 5.91673649 | 0.05190354 | 0.13440139 | GRX8 | XII | 854062 | 854391 | 1 |
| YLR365W | 21.1043865 | 0.32323369 | 0.77929395 | 1.02382389 | 0.59934856 | 0.74628563 | NA | XII | 855199 | 855531 | 1 |
| YLR367W | 10.8332111 | 1.07332585 | 0.88122295 | 2.4927013 | 0.28755226 | 0.46171037 | RPS22B | XII | 856442 | 857317 | 1 |
| YLR368W | 16.7173321 | 1.78514057 | 0.94105351 | 6.10663417 | 0.04720209 | 0.12525211 | MDM30 | XII | 857540 | 859336 | 1 |
| YLR369W | 39.112502 | 0.00661708 | 0.82233042 | 9.40891199 | 0.00905484 | 0.03655068 | SSQ1 | XII | 859552 | 861525 | 1 |
| YLR370C | 15.6788637 | 0.02175274 | 0.79457626 | 0.16003411 | 0.9231006 | 0.95153151 | ARC18 | XII | 861718 | 862254 | -1 |
| YLR371W | 33.6482803 | -1.161698 | 0.64823311 | 3.2160664 | 0.20028114 | 0.35924034 | ROM2 | XII | 862714 | 866784 | 1 |
| YLR372W | 212.946601 | 4.22262519 | 0.60501922 | 44.1592673 | 2.58E-10 | 1.09E-08 | ELO3 | XII | 867354 | 868391 | 1 |
| YLR373C | 57.9402541 | -1.0849361 | 0.64414103 | 3.10454301 | 0.2117664 | 0.37530684 | VID22 | XII | 868662 | 871367 | -1 |
| YLR375W | 86.0502495 | -0.3947113 | 0.5729973 | 0.93457405 | 0.62670019 | 0.76471951 | STP3 | XII | 871697 | 872728 | 1 |
| YLR376C | 102.084059 | 1.60294827 | 1.68307174 | 3.67869753 | 0.15892089 | 0.30559679 | PSY3 | XII | 872826 | 873554 | -1 |
| YLR377C | 480.080057 | 0.26909312 | 0.49096657 | 2.03548504 | 0.3614099 | 0.53408966 | FBP1 | XII | 873746 | 874792 | -1 |
| YLR378C | 46.3427399 | 0.02876135 | 0.5951365 | 0.94323259 | 0.6239929 | 0.76326014 | SEC61 | XII | 875736 | 877178 | -1 |
| YLR380W | 31.1977536 | -0.7594174 | 0.66875206 | 2.04096518 | 0.36042096 | 0.53345156 | CSR1 | XII | 878282 | 879508 | 1 |
| YLR381W | 53.474129 | -0.9702214 | 0.61264218 | 2.69109941 | 0.26039653 | 0.43188081 | CTF3 | XII | 879723 | 881924 | 1 |

| | | | | | | | | | | | |
|---|---|---|---|---|---|---|---|---|---|---|---|
| YLR382C | 98.5483013 | 3.48647601 | 0.77200978 | 31.1424796 | 1.73E-07 | 3.88E-06 | NAM2 | XII | 882067 | 884751 | -1 |
| YLR383W | 18.8467481 | -1.2284919 | 0.75012824 | 2.87563269 | 0.23744569 | 0.4053281 | SMC6 | XII | 885288 | 888632 | 1 |
| YLR384C | 26.8725254 | -0.9545262 | 0.7676409 | 2.26896898 | 0.32158786 | 0.49429087 | IKI3 | XII | 888851 | 892900 | -1 |
| YLR386W | 146.954453 | -1.9041622 | 0.7502917 | 12.5572496 | 0.00187598 | 0.01069086 | VAC14 | XII | 893628 | 896270 | 1 |
| YLR387C | 126.377764 | -1.1495923 | 0.5314388 | 6.40169714 | 0.04072763 | 0.11285424 | REH1 | XII | 896374 | 897672 | -1 |
| YLR388W | 71.5939671 | -1.9352787 | 0.59125491 | 10.4980046 | 0.00525276 | 0.02385369 | RPS29A | XII | 898651 | 898821 | 1 |
| YLR389C | 73.7614573 | 0.74366768 | 0.54791329 | 4.89220766 | 0.08663046 | 0.19825705 | STE23 | XII | 899577 | 902660 | -1 |
| YLR390W | 86.0331729 | 0.90338227 | 0.54706768 | 2.71129219 | 0.25778069 | 0.42975819 | ECM19 | XII | 903066 | 903404 | 1 |
| YLR390W-A | 50.3365091 | 0.95725207 | 0.61256334 | 3.60202665 | 0.16513147 | 0.31399383 | CCW14 | XII | 903724 | 904440 | 1 |
| YLR392C | 17.1737233 | -0.2542461 | 0.74861995 | 0.12271621 | 0.94048639 | 0.96082012 | ART10 | XII | 904749 | 906305 | -1 |
| YLR393W | 98.740742 | 2.63884263 | 0.71816271 | 23.7027882 | 7.13E-06 | 9.69E-05 | ATP10 | XII | 907079 | 907918 | 1 |
| YLR394W | 132.497536 | 2.73374756 | 0.58156413 | 27.6279929 | 1.00E-06 | 1.77E-05 | CST9 | XII | 907950 | 909398 | 1 |
| YLR395C | 138.842959 | 1.62633905 | 0.59710817 | 8.52514377 | 0.01408603 | 0.05103761 | COX8 | XII | 909729 | 909965 | -1 |
| YLR396C | 397.83144 | -1.6090367 | 0.50369671 | 10.7949223 | 0.00452806 | 0.02104132 | VPS33 | XII | 910235 | 912310 | -1 |
| YLR397C | 88.9129803 | 1.04991836 | 0.57840047 | 7.85447528 | 0.01969801 | 0.06517227 | AFG2 | XII | 912550 | 914892 | -1 |
| YLR398C | 46.8528721 | 0.05940285 | 0.57680284 | 0.46299273 | 0.79334558 | 0.87452157 | SKI2 | XII | 915156 | 919019 | -1 |
| YLR399C | 75.0045811 | -0.4791318 | 0.5451914 | 0.83503994 | 0.65867834 | 0.78659971 | BDF1 | XII | 919536 | 921596 | -1 |
| YLR401C | 73.3995716 | 0.47375106 | 0.5644183 | 4.02838026 | 0.13342842 | 0.26946304 | DUS3 | XII | 922442 | 924448 | -1 |
| YLR403W | 20.7042522 | 0.12567124 | 0.75541289 | 0.13023441 | 0.93695764 | 0.95898617 | SFP1 | XII | 925568 | 927619 | 1 |
| YLR404W | 31.545634 | -0.0986829 | 0.64490669 | 0.63016281 | 0.72972947 | 0.83678999 | SEI1 | XII | 928743 | 929600 | 1 |
| YLR405W | 1654.88203 | -1.8722549 | 1.65988547 | 6.23293907 | 0.04431334 | 0.11967082 | DUS4 | XII | 929789 | 930892 | 1 |
| YLR406C | 64.5962666 | -2.2614613 | 0.63850649 | 11.8472367 | 0.0026755 | 0.013956 | RPL31B | XII | 931065 | 931755 | -1 |
| YLR407W | 66.6631141 | 0.803092 | 0.57622006 | 3.32173737 | 0.18997388 | 0.34583253 | NA | XII | 932967 | 933653 | 1 |
| YLR408C | 66.1328068 | 2.31948585 | 0.58744583 | 15.522633 | 0.0004259 | 0.00323614 | BLS1 | XII | 933885 | 934253 | -1 |
| YLR409C | 36.3843517 | -2.0422537 | 0.71570305 | 7.92605436 | 0.01900549 | 0.06364477 | UTP21 | XII | 934414 | 937233 | -1 |
| YLR410W | 393.465136 | -1.1889319 | 0.48072441 | 6.39673558 | 0.04082879 | 0.11305473 | VIP1 | XII | 937541 | 940981 | 1 |
| YLR410W-B | 14.2480751 | -1.6636493 | 0.83447881 | 5.01622336 | 0.08142184 | 0.18867385 | NA | XII | 941483 | 946796 | 1 |
| YLR411W | 125.831203 | 0.36847809 | 0.53235667 | 2.50699167 | 0.28550497 | 0.45905965 | CTR3 | XII | 947253 | 947978 | 1 |
| YLR412W | 117.485249 | -1.6160073 | 0.53816723 | 10.2882482 | 0.00583358 | 0.02592206 | BER1 | XII | 948368 | 949192 | 1 |
| YLR413W | 528.507555 | 1.96865129 | 0.4614001 | 26.8421067 | 1.48E-06 | 2.50E-05 | INA1 | XII | 951156 | 953183 | 1 |
| YLR414C | 44.755932 | 0.00016305 | 0.70690781 | 1.42821436 | 0.48962907 | 0.65106127 | PUN1 | XII | 953353 | 954144 | -1 |
| YLR415C | 16.7057583 | -0.8115956 | 0.84315664 | 2.00738001 | 0.36652447 | 0.53971939 | NA | XII | 954259 | 954597 | -1 |
| YLR417W | 298.154617 | -0.4553247 | 0.52882759 | 4.00236118 | 0.1351756 | 0.27194814 | VPS36 | XII | 955010 | 956710 | 1 |

| | | | | | | | | | | | |
|---|---|---|---|---|---|---|---|---|---|---|---|
| YLR418C | 120.183553 | 3.67037295 | 0.59281384 | 39.8235649 | 2.25E-09 | 7.75E-08 | CDC73 | XII | 956914 | 958095 | -1 |
| YLR419W | 70.0461237 | -0.7244295 | 0.53886047 | 2.07174753 | 0.35491613 | 0.52857757 | NA | XII | 958428 | 962735 | 1 |
| YLR420W | 471.658182 | 0.31383295 | 0.6979719 | 6.73034408 | 0.03455607 | 0.09964126 | URA4 | XII | 963785 | 964879 | 1 |
| YLR421C | 42.0783491 | -0.1518743 | 0.74287962 | 0.06540246 | 0.96782768 | 0.97506912 | RPN13 | XII | 965090 | 965560 | -1 |
| YLR422W | 173.120198 | 2.39565512 | 0.62361553 | 13.6481553 | 0.00108728 | 0.00708137 | DCK1 | XII | 965897 | 971695 | 1 |
| YLR423C | 1821.66516 | 6.32412016 | 0.48997361 | 114.614454 | 1.29E-25 | 9.36E-23 | ATG17 | XII | 971917 | 973170 | -1 |
| YLR424W | 34.4625557 | 2.65878958 | 0.68904497 | 18.9698018 | 7.60E-05 | 0.00074072 | SPP382 | XII | 973395 | 975521 | 1 |
| YLR425W | 46.6984099 | 0.2817014 | 0.60394769 | 4.78090392 | 0.09158828 | 0.2057006 | TUS1 | XII | 982894 | 986817 | 1 |
| YLR426W | 40.0973189 | 1.88383314 | 0.66918042 | 8.37296192 | 0.01519968 | 0.05409105 | TDA5 | XII | 987062 | 988113 | 1 |
| YLR427W | 299.44852 | 2.14800822 | 0.51349541 | 21.6886717 | 1.95E-05 | 0.00023201 | MAG2 | XII | 988428 | 990440 | 1 |
| YLR429W | 222.879273 | 1.8405596 | 0.56383832 | 10.1902899 | 0.00612642 | 0.02692266 | CRN1 | XII | 990777 | 992732 | 1 |
| YLR430W | 108.245377 | 0.3510585 | 0.64498889 | 3.75615096 | 0.15288405 | 0.29674779 | SEN1 | XII | 993434 | 1E+06 | 1 |
| YLR431C | 80.8238326 | 3.04288297 | 0.57866439 | 31.3093365 | 1.59E-07 | 3.62E-06 | ATG23 | XII | 1000342 | 1E+06 | -1 |
| YLR432W | 152.716864 | -0.3920937 | 0.70829729 | 5.20919739 | 0.0739328 | 0.17558282 | IMD3 | XII | 1002557 | 1E+06 | 1 |
| YLR433C | 133.255444 | 1.60417855 | 0.61395471 | 6.77916082 | 0.03372282 | 0.09792134 | CNA1 | XII | 1004347 | 1E+06 | -1 |
| YLR435W | 34.2266832 | -2.6422105 | 0.70260712 | 14.5250107 | 0.00070135 | 0.00488665 | TSR2 | XII | 1006378 | 1E+06 | 1 |
| YLR436C | 218.530457 | 1.11406072 | 0.50258224 | 5.80590635 | 0.05486097 | 0.14011689 | ECM30 | XII | 1007421 | 1E+06 | -1 |
| YLR437C | 67.4003137 | -0.992955 | 0.55061831 | 3.21057221 | 0.20083209 | 0.36006531 | DIF1 | XII | 1011622 | 1E+06 | -1 |
| YLR438C-A | 46.3295786 | -1.1058695 | 0.65001461 | 3.34165533 | 0.18809132 | 0.34376539 | LSM3 | XII | 1013909 | 1E+06 | -1 |
| YLR438W | 100.013468 | -1.1000334 | 0.52614734 | 7.26623887 | 0.0264336 | 0.08085078 | CAR2 | XII | 1012501 | 1E+06 | 1 |
| YLR439W | 130.660229 | 1.16426818 | 0.546554 | 4.43306447 | 0.10898639 | 0.23299267 | MRPL4 | XII | 1014491 | 1E+06 | 1 |
| YLR440C | 76.9080171 | 0.16530318 | 0.55801443 | 0.0948181 | 0.95369721 | 0.96767513 | SEC39 | XII | 1015568 | 1E+06 | -1 |
| YLR441C | 194.944006 | -3.1773156 | 0.70544887 | 18.0121966 | 0.00012266 | 0.00111316 | RPS1A | XII | 1018141 | 1E+06 | -1 |
| YLR442C | 18.4248633 | -0.5924743 | 0.94334999 | 0.51963827 | 0.77119105 | 0.86067776 | SIR3 | XII | 1019315 | 1E+06 | -1 |
| YLR443W | 32.1298008 | -0.8641284 | 0.6596582 | 3.20381388 | 0.20151188 | 0.36088889 | ECM7 | XII | 1022625 | 1E+06 | 1 |
| YLR445W | 31.6843425 | -1.3260739 | 0.64384578 | 7.34319792 | 0.02543577 | 0.07871357 | GMC2 | XII | 1024189 | 1E+06 | 1 |
| YLR446W | 1160.13324 | -2.9005788 | 0.45771484 | 36.3519797 | 1.28E-08 | 3.72E-07 | NA | XII | 1025214 | 1E+06 | 1 |
| YLR447C | 26.9764563 | -0.7139183 | 0.77157565 | 1.46445824 | 0.48083595 | 0.64344033 | VMA6 | XII | 1026856 | 1E+06 | -1 |
| YLR448W | 90.2431114 | -3.268324 | 0.62212108 | 25.5618581 | 2.81E-06 | 4.33E-05 | RPL6B | XII | 1028854 | 1E+06 | 1 |
| YLR449W | 476.127598 | -1.2564157 | 0.60119545 | 8.94720999 | 0.01140612 | 0.04341976 | FPR4 | XII | 1030834 | 1E+06 | 1 |
| YLR450W | 352.791065 | -1.4955628 | 0.49920013 | 8.60506293 | 0.01353425 | 0.04937874 | HMG2 | XII | 1032627 | 1E+06 | 1 |
| YLR451W | 61.1017117 | -0.3440754 | 0.55537941 | 1.22273144 | 0.54260931 | 0.69630051 | LEU3 | XII | 1036093 | 1E+06 | 1 |
| YLR452C | 16.0225878 | 0.42048393 | 0.747546 | 3.34730816 | 0.18756045 | 0.34351531 | SST2 | XII | 1039270 | 1E+06 | -1 |

| | | | | | | | | | | | |
|---|---|---|---|---|---|---|---|---|---|---|---|
| YLR453C | 143.578216 | -2.2506807 | 0.53110641 | 19.3763318 | 6.20E-05 | 0.00061906 | RIF2 | XII | 1041799 | 1E+06 | -1 |
| YLR454W | 54.2528009 | -1.3723255 | 0.65608397 | 4.82365903 | 0.08965113 | 0.20308295 | FMP27 | XII | 1043998 | 1E+06 | 1 |
| YLR455W | 66.0789261 | -0.3027077 | 0.76012555 | 2.13241116 | 0.34431251 | 0.51781024 | PDP3 | XII | 1053629 | 1E+06 | 1 |
| YLR456W | 83.1167269 | 0.44026799 | 0.66697822 | 2.24989541 | 0.32466945 | 0.49770614 | NA | XII | 1055070 | 1E+06 | 1 |
| YLR457C | 13.366799 | 1.03443207 | 0.90740235 | 1.40502538 | 0.4953391 | 0.65554592 | NBP1 | XII | 1055812 | 1E+06 | -1 |
| YLR459W | 38.928909 | 2.00108567 | 0.82706895 | 5.57725217 | 0.06150566 | 0.15245492 | GAB1 | XII | 1057334 | 1E+06 | 1 |
| YLR460C | 25.3861504 | -1.9618136 | 0.73336052 | 8.89592136 | 0.01170241 | 0.04434355 | NA | XII | 1059757 | 1E+06 | -1 |
| YLR462W | 36.1186258 | -0.8650777 | 0.66174875 | 3.61739015 | 0.16386783 | 0.31248127 | NA | XII | 1065956 | 1E+06 | 1 |
| YLR466W | 7.9540639 | -2.5026844 | 1.05486184 | 6.14387703 | 0.04633125 | 0.12328031 | YRF1-4 | XII | 1067087 | 1E+06 | 1 |
| YML001W | 236.853208 | -1.3350675 | 0.54942266 | 8.90100562 | 0.0116727 | 0.04427675 | YPT7 | XIII | 267174 | 267800 | 1 |
| YML002W | 445.524361 | -0.1347009 | 0.4896731 | 0.13620961 | 0.93416256 | 0.95735444 | NA | XIII | 264541 | 266754 | 1 |
| YML003W | 9.24664629 | 1.4376255 | 1.00463251 | 2.33205427 | 0.31160244 | 0.48405923 | NA | XIII | 263483 | 264355 | 1 |
| YML004C | 56.7630591 | 0.4045266 | 0.69952525 | 0.34123925 | 0.84314222 | 0.9080717 | GLO1 | XIII | 261705 | 262685 | -1 |
| YML005W | 1877.57392 | 1.30986084 | 0.82332925 | 9.55537661 | 0.00841543 | 0.03453249 | TRM12 | XIII | 260221 | 261609 | 1 |
| YML006C | 65.3871588 | 1.13153251 | 0.58209289 | 4.35419834 | 0.11336992 | 0.24000433 | GIS4 | XIII | 256092 | 258416 | -1 |
| YML007C-A | 39.3994828 | -0.8956708 | 0.68120887 | 4.4136652 | 0.11004867 | 0.23477589 | NA | XIII | 253162 | 253272 | -1 |
| YML007W | 60.4121514 | 0.65043225 | 0.58506861 | 1.59551857 | 0.45033691 | 0.6167102 | YAP1 | XIII | 253848 | 255800 | 1 |
| YML008C | 56.9592058 | -0.9724746 | 0.69043839 | 4.31482422 | 0.11562396 | 0.242911 | ERG6 | XIII | 251839 | 252990 | -1 |
| YML009C | 10.7955892 | -1.8261383 | 0.98677922 | 3.5144804 | 0.17252033 | 0.32379018 | MRPL39 | XIII | 251304 | 251516 | -1 |
| YML010W | 97.062783 | -0.5500844 | 0.57741443 | 1.73666295 | 0.41965116 | 0.59032562 | SPT5 | XIII | 247677 | 250868 | 1 |
| YML011C | 327.532665 | -1.8101395 | 0.57918599 | 12.2322452 | 0.002207 | 0.0120552 | RAD33 | XIII | 246895 | 247428 | -1 |
| YML012W | 55.6623557 | -0.1586466 | 0.58557836 | 3.07433513 | 0.21498918 | 0.37881539 | ERV25 | XIII | 246116 | 246751 | 1 |
| YML013W | 64.5221968 | 1.73144371 | 0.56810085 | 10.3269134 | 0.00572189 | 0.02550402 | UBX2 | XIII | 244149 | 245903 | 1 |
| YML014W | 13.0560467 | -0.8620194 | 0.83087449 | 1.50061398 | 0.47222156 | 0.63393528 | TRM9 | XIII | 243225 | 244064 | 1 |
| YML015C | 140.164885 | 1.70553191 | 0.55133647 | 12.4807064 | 0.00194917 | 0.01096762 | TAF11 | XIII | 241989 | 243029 | -1 |
| YML016C | 54.6503683 | 1.24047127 | 0.59961372 | 6.56435077 | 0.03754649 | 0.1061495 | PPZ1 | XIII | 239458 | 241536 | -1 |
| YML017W | 163.291274 | 0.90734743 | 0.50306779 | 7.19129488 | 0.02744291 | 0.08314728 | PSP2 | XIII | 236588 | 238731 | 1 |
| YML018C | 51.0715818 | 0.92968785 | 0.63429165 | 4.35512561 | 0.11331737 | 0.23998083 | NA | XIII | 234771 | 235952 | -1 |
| YML019W | 82.8842753 | -1.4784649 | 0.55765907 | 12.7891516 | 0.00167059 | 0.00979022 | OST6 | XIII | 233457 | 234455 | 1 |
| YML020W | 72.0218702 | -0.5180271 | 0.54536462 | 1.64472451 | 0.43939247 | 0.60805219 | NA | XIII | 231149 | 233143 | 1 |
| YML021C | 148.924859 | -3.1816748 | 0.63045712 | 22.0400599 | 1.64E-05 | 0.00019996 | UNG1 | XIII | 229734 | 230813 | -1 |
| YML022W | 35.7188785 | -0.262387 | 0.64925261 | 0.44240539 | 0.80155419 | 0.88031084 | APT1 | XIII | 228937 | 229500 | 1 |
| YML023C | 453.966283 | 1.929758 | 0.54485316 | 13.0884605 | 0.00143839 | 0.00885988 | NSE5 | XIII | 226994 | 228664 | -1 |

| | | | | | | | | | | | |
|---|---|---|---|---|---|---|---|---|---|---|---|
| YML024W | 67.3550106 | -2.424738 | 0.7198364 | 10.6108569 | 0.00496457 | 0.02277723 | RPS17A | XIII | 225889 | 226697 | 1 |
| YML025C | 12.4565845 | -0.8982456 | 0.90892296 | 1.02257149 | 0.59972399 | 0.74659254 | YML6 | XIII | 224406 | 225365 | -1 |
| YML026C | 92.9752755 | -3.6290182 | 0.85216257 | 15.6790543 | 0.00039386 | 0.00302845 | RPS18B | XIII | 222987 | 223828 | -1 |
| YML027W | 190.49332 | 1.87498074 | 0.54182368 | 12.9323472 | 0.00155517 | 0.00937506 | YOX1 | XIII | 221406 | 222563 | 1 |
| YML028W | 153.334924 | -0.308797 | 0.61447252 | 0.38214797 | 0.82607147 | 0.89568423 | TSA1 | XIII | 220138 | 220728 | 1 |
| YML029W | 275.17951 | -0.1275947 | 0.52539659 | 0.07573916 | 0.96283851 | 0.97258112 | USA1 | XIII | 217362 | 219878 | 1 |
| YML030W | 33.8801773 | -0.6574007 | 0.70414523 | 3.34206767 | 0.18805255 | 0.34376539 | RCF1 | XIII | 216435 | 216914 | 1 |
| YML031W | 47.3429009 | 0.77777553 | 0.59917519 | 2.79751881 | 0.24690308 | 0.41775828 | NDC1 | XIII | 214189 | 216156 | 1 |
| YML032C | 28.8329751 | -1.0348071 | 0.65287259 | 2.52669891 | 0.28270553 | 0.45678444 | RAD52 | XIII | 212515 | 213930 | -1 |
| YML034W | 35.4662262 | 0.41899817 | 0.62354644 | 1.67542986 | 0.43269814 | 0.60137356 | SRC1 | XIII | 209525 | 212155 | 1 |
| YML035C | 184.819314 | 0.12900666 | 0.5221295 | 0.48097787 | 0.78624334 | 0.87021248 | AMD1 | XIII | 206428 | 208860 | -1 |
| YML036W | 737.938283 | -0.298308 | 0.57725168 | 2.34077371 | 0.3102469 | 0.4830878 | CGI121 | XIII | 205642 | 206293 | 1 |
| YML037C | 74.9023567 | 1.5385447 | 0.7638975 | 4.97092875 | 0.08328687 | 0.19181821 | NA | XIII | 204386 | 205408 | -1 |
| YML038C | 15.3053318 | 0.13258388 | 0.77582887 | 0.13344981 | 0.9354525 | 0.95803272 | YMD8 | XIII | 202775 | 204103 | -1 |
| YML041C | 50.5549655 | 1.36719272 | 0.94174458 | 5.58951753 | 0.06112962 | 0.1520363 | VPS71 | XIII | 194913 | 195755 | -1 |
| YML042W | 205.630072 | -0.0033642 | 0.52686436 | 0.26444191 | 0.87614739 | 0.92712351 | CAT2 | XIII | 192788 | 194800 | 1 |
| YML043C | 12.9528371 | -3.0429628 | 0.91314429 | 12.3273615 | 0.00210449 | 0.01162812 | RRN11 | XIII | 190244 | 191767 | -1 |
| YML046W | 10.1341204 | -1.8385039 | 0.89507187 | 5.44073594 | 0.06585052 | 0.16064998 | PRP39 | XIII | 181474 | 183363 | 1 |
| YML048W | 46.4032934 | -1.5661748 | 1.00053993 | 2.46015174 | 0.2922704 | 0.46643602 | GSF2 | XIII | 178426 | 179637 | 1 |
| YML049C | 48.0802931 | -0.0839992 | 0.57780003 | 0.17716428 | 0.91522793 | 0.94797312 | RSE1 | XIII | 174220 | 178305 | -1 |
| YML050W | 74.598849 | -0.2552267 | 0.53719263 | 0.46786483 | 0.7914153 | 0.87352731 | AIM32 | XIII | 173139 | 174074 | 1 |
| YML051W | 82.4893255 | -1.7592046 | 0.53485957 | 12.4643642 | 0.00196516 | 0.01103615 | GAL80 | XIII | 171594 | 172901 | 1 |
| YML052W | 187.630678 | 1.78892862 | 0.56726473 | 22.6354356 | 1.22E-05 | 0.00015675 | SUR7 | XIII | 170402 | 171310 | 1 |
| YML053C | 19.1901839 | -0.4156329 | 0.71815173 | 0.82081254 | 0.66338068 | 0.79064927 | NA | XIII | 169116 | 169754 | -1 |
| YML054C | 382.800786 | -2.3819287 | 0.50587725 | 20.364535 | 3.78E-05 | 0.00040962 | CYB2 | XIII | 165533 | 167308 | -1 |
| YML054C-A | 191.129122 | -0.2049935 | 0.5464794 | 1.04016847 | 0.59447047 | 0.74174387 | NA | XIII | 167623 | 167781 | -1 |
| YML055W | 30.5380914 | 0.0734585 | 0.63840933 | 0.0286761 | 0.98576425 | 0.98832468 | SPC2 | XIII | 164790 | 165326 | 1 |
| YML056C | 64.6264738 | 0.67018348 | 0.62505389 | 1.57827264 | 0.45423694 | 0.62039568 | IMD4 | XIII | 162194 | 164176 | -1 |
| YML057W | 149.624716 | 3.29140304 | 0.53737292 | 37.0185659 | 9.15E-09 | 2.76E-07 | CMP2 | XIII | 160180 | 161994 | 1 |
| YML058W | 186.328436 | 2.47724656 | 0.51457117 | 22.1348348 | 1.56E-05 | 0.00019234 | SML1 | XIII | 159383 | 159697 | 1 |
| YML058W-A | 25.9288503 | 1.01675788 | 0.69018383 | 2.28398552 | 0.31918233 | 0.49189931 | HUG1 | XIII | 158760 | 158966 | 1 |
| YML059C | 41.4330295 | 1.08763544 | 0.63887575 | 3.59140138 | 0.16601109 | 0.31525228 | NTE1 | XIII | 153219 | 158258 | -1 |
| YML060W | 7.15515563 | -0.0020899 | 1.05737719 | 0.6849936 | 0.70999539 | 0.82438173 | OGG1 | XIII | 151871 | 153001 | 1 |

| | | | | | | | | | | | |
|---|---|---|---|---|---|---|---|---|---|---|---|
| YML061C | 31.1195425 | -0.7256747 | 0.65288812 | 3.46497779 | 0.17684372 | 0.32944824 | PIF1 | XIII | 148953 | 151532 | -1 |
| YML062C | 72.4756982 | -1.3503372 | 0.60078099 | 5.69663649 | 0.05794168 | 0.14626659 | MFT1 | XIII | 147505 | 148683 | -1 |
| YML063W | 158.93677 | -3.0538291 | 0.72702989 | 15.5747192 | 0.00041495 | 0.00316124 | RPS1B | XIII | 146482 | 147249 | 1 |
| YML064C | 17.0269822 | -1.7357753 | 0.82761904 | 5.08684574 | 0.07859691 | 0.18409228 | TEM1 | XIII | 145139 | 145876 | -1 |
| YML065W | 66.7441541 | -1.3002093 | 0.7989764 | 5.26666665 | 0.0718386 | 0.17209164 | ORC1 | XIII | 142210 | 144954 | 1 |
| YML066C | 994.627674 | 0.98919088 | 0.48206004 | 4.23724677 | 0.12019698 | 0.24944104 | SMA2 | XIII | 140424 | 141533 | -1 |
| YML067C | 399.78616 | -0.0968396 | 0.49215499 | 0.6427121 | 0.72516501 | 0.83456178 | ERV41 | XIII | 139063 | 140214 | -1 |
| YML068W | 188.4442 | -1.0453439 | 0.60272568 | 6.76661588 | 0.03393501 | 0.09824187 | ITT1 | XIII | 137550 | 138944 | 1 |
| YML069W | 44.6293787 | -0.3043438 | 0.59436436 | 0.99787162 | 0.60717647 | 0.75134682 | POB3 | XIII | 135500 | 137158 | 1 |
| YML070W | 55.4336425 | 0.77072848 | 0.64170292 | 1.98460515 | 0.37072209 | 0.54410163 | DAK1 | XIII | 133475 | 135229 | 1 |
| YML071C | 267.011606 | 1.4066688 | 0.71368648 | 11.1894268 | 0.00371746 | 0.01814182 | COG8 | XIII | 129749 | 131572 | -1 |
| YML072C | 262.166211 | 0.55517754 | 0.49479051 | 1.41290492 | 0.49339142 | 0.65446422 | TCB3 | XIII | 124730 | 129367 | -1 |
| YML073C | 186.67094 | -3.1739337 | 0.62580811 | 22.5512317 | 1.27E-05 | 0.00016205 | RPL6A | XIII | 123227 | 124172 | -1 |
| YML074C | 680.951872 | -0.473745 | 0.6447178 | 5.44109224 | 0.06583879 | 0.16064998 | FPR3 | XIII | 120089 | 121324 | -1 |
| YML075C | 75.2321225 | 0.09812086 | 0.5590611 | 0.24894529 | 0.88296241 | 0.93100373 | HMG1 | XIII | 115734 | 118898 | -1 |
| YML076C | 91.8079686 | 1.99510265 | 0.54222199 | 18.0516511 | 0.00012026 | 0.00109485 | WAR1 | XIII | 112513 | 115347 | -1 |
| YML077W | 48.2922499 | 0.72733272 | 0.58180972 | 1.66060621 | 0.43591714 | 0.60483111 | BET5 | XIII | 111865 | 112344 | 1 |
| YML078W | 106.227883 | 0.30640755 | 0.55622454 | 0.35122854 | 0.83894153 | 0.90540008 | CPR3 | XIII | 111002 | 111550 | 1 |
| YML079W | 24.0588246 | -0.0614078 | 0.72324539 | 0.13638016 | 0.93408291 | 0.95735444 | NA | XIII | 110247 | 110852 | 1 |
| YML080W | 16.1795302 | 0.21895696 | 0.90942383 | 1.0008501 | 0.60627291 | 0.75088539 | DUS1 | XIII | 108806 | 110077 | 1 |
| YML081C-A | 51.1614985 | -0.6832479 | 0.59926217 | 2.3316168 | 0.31167061 | 0.48405923 | ATP18 | XIII | 103983 | 104162 | -1 |
| YML081W | 44.2925372 | 0.57191444 | 0.66516617 | 4.10416492 | 0.1284671 | 0.26218699 | TDA9 | XIII | 104777 | 108532 | 1 |
| YML082W | 28.1193849 | -1.4157447 | 0.67529532 | 4.27259839 | 0.11809107 | 0.24673616 | NA | XIII | 101862 | 103811 | 1 |
| YML083C | 278.671987 | -0.2410427 | 0.47940675 | 0.82459892 | 0.66212597 | 0.78964147 | NA | XIII | 99794 | 101050 | -1 |
| YML085C | 37.3557016 | -0.9368606 | 0.71328287 | 2.59957376 | 0.27258988 | 0.44584616 | TUB1 | XIII | 97941 | 99400 | -1 |
| YML086C | 70.2729105 | 1.08569175 | 0.62383288 | 5.18786936 | 0.07472544 | 0.17681255 | ALO1 | XIII | 95791 | 97371 | -1 |
| YML087C | 1892.93277 | 3.0803924 | 0.47574271 | 35.9987802 | 1.52E-08 | 4.35E-07 | AIM33 | XIII | 94431 | 95369 | -1 |
| YML088W | 205.194481 | 2.66839009 | 0.51672237 | 26.4499081 | 1.80E-06 | 2.99E-05 | UFO1 | XIII | 92235 | 94241 | 1 |
| YML091C | 41.1568071 | 0.56394786 | 0.66641884 | 3.58330463 | 0.16668453 | 0.31570278 | RPM2 | XIII | 87123 | 90731 | -1 |
| YML092C | 36.4034058 | -0.3895483 | 0.81973209 | 0.8802231 | 0.64396458 | 0.77686719 | PRE8 | XIII | 85987 | 86739 | -1 |
| YML093W | 54.8671805 | -3.1886649 | 0.7059565 | 20.4773375 | 3.58E-05 | 0.0003914 | UTP14 | XIII | 83090 | 85789 | 1 |
| YML094W | 13.0292275 | -1.9977304 | 0.87841353 | 4.88886156 | 0.08677551 | 0.19851056 | GIM5 | XIII | 82275 | 82849 | 1 |
| YML095C | 185.069877 | -2.6453558 | 0.51360149 | 24.2487739 | 5.43E-06 | 7.66E-05 | RAD10 | XIII | 81481 | 82113 | -1 |

| | | | | | | | | | | | |
|---|---|---|---|---|---|---|---|---|---|---|---|
| YML096W | 869.395302 | 0.51705483 | 0.46058624 | 1.576424 | 0.454657 | 0.62066785 | NA | XIII | 79909 | 81486 | 1 |
| YML097C | 31.4509663 | -0.4968004 | 0.663296 | 0.70572745 | 0.70267294 | 0.81860691 | VPS9 | XIII | 78335 | 79690 | -1 |
| YML098W | 35.8373151 | 0.20381181 | 0.64002898 | 0.23908611 | 0.8873258 | 0.93215081 | TAF13 | XIII | 77267 | 77770 | 1 |
| YML099C | 15.6251821 | 0.40204919 | 0.92578995 | 0.25261807 | 0.88134244 | 0.9300006 | ARG81 | XIII | 74398 | 77040 | -1 |
| YML100W | 126.786729 | 0.86764766 | 0.50928731 | 3.42928132 | 0.1800284 | 0.33462743 | TSL1 | XIII | 70624 | 73920 | 1 |
| YML100W-A | 129.566306 | 0.71278018 | 0.70622524 | 1.932583 | 0.38049148 | 0.55436479 | NA | XIII | 70138 | 70311 | 1 |
| YML101C | 158.046585 | 0.05776659 | 0.74977986 | 5.63283821 | 0.05981977 | 0.14961402 | CUE4 | XIII | 69735 | 70088 | -1 |
| YML102W | 433.992011 | 0.0151115 | 0.68585515 | 4.02216999 | 0.13384338 | 0.26992447 | CAC2 | XIII | 68294 | 69700 | 1 |
| YML103C | 11.3770154 | -0.5603078 | 1.10586182 | 0.5442316 | 0.76176604 | 0.85526961 | NUP188 | XIII | 62582 | 67549 | -1 |
| YML104C | 89.1669602 | -0.7812278 | 0.51919804 | 2.91446787 | 0.23287954 | 0.40027153 | MDM1 | XIII | 58939 | 62322 | -1 |
| YML105C | 66.6157127 | 2.45776314 | 0.70146471 | 13.6496431 | 0.00108647 | 0.00708137 | SEC65 | XIII | 57866 | 58687 | -1 |
| YML106W | 34.6021929 | -0.7062421 | 0.61803061 | 2.4959741 | 0.2870821 | 0.46133926 | URA5 | XIII | 56773 | 57453 | 1 |
| YML107C | 553.507954 | -1.329314 | 0.57817951 | 5.19781115 | 0.07435491 | 0.17629168 | PML39 | XIII | 55265 | 56269 | -1 |
| YML108W | 182.735696 | -1.0799843 | 0.6725158 | 4.11450406 | 0.12780469 | 0.26147074 | NA | XIII | 54793 | 55110 | 1 |
| YML109W | 46.2871023 | 1.69703879 | 0.627665 | 8.52022721 | 0.0141207 | 0.05110095 | ZDS2 | XIII | 51640 | 54468 | 1 |
| YML110C | 11.1069397 | -0.0980418 | 0.86273895 | 0.10004106 | 0.9512099 | 0.96614462 | COQ5 | XIII | 50031 | 50954 | -1 |
| YML111W | 87.8360469 | 0.45882759 | 0.59921366 | 1.3399216 | 0.51172864 | 0.66867723 | BUL2 | XIII | 46942 | 49704 | 1 |
| YML112W | 40.9382117 | 0.14623731 | 0.74149406 | 4.55162399 | 0.10271347 | 0.2231561 | CTK3 | XIII | 45063 | 45953 | 1 |
| YML113W | 46.447141 | 0.42665076 | 0.59993436 | 0.64075808 | 0.72587385 | 0.83456178 | DAT1 | XIII | 44045 | 44791 | 1 |
| YML114C | 48.9693991 | -0.5069399 | 0.64031167 | 2.92959932 | 0.23112429 | 0.39792139 | TAF8 | XIII | 42043 | 43575 | -1 |
| YML115C | 47.4230925 | -0.4460585 | 0.62187548 | 0.54926024 | 0.75985313 | 0.85451323 | VAN1 | XIII | 40187 | 41794 | -1 |
| YML116W | 244.627165 | 4.23741378 | 0.59471634 | 48.5814827 | 2.82E-11 | 1.49E-09 | ATR1 | XIII | 38196 | 39824 | 1 |
| YML117W | 169.692587 | 0.28600004 | 0.59791508 | 1.3118757 | 0.51895513 | 0.67522582 | NAB6 | XIII | 34243 | 37647 | 1 |
| YML118W | 183.049701 | 3.3082479 | 0.52535951 | 35.15155 | 2.33E-08 | 6.39E-07 | NGL3 | XIII | 32334 | 33851 | 1 |
| YML119W | 245.260817 | 3.34240591 | 0.56862165 | 29.9675796 | 3.11E-07 | 6.43E-06 | NA | XIII | 30611 | 31684 | 1 |
| YML120C | 24.57111 | 0.81102816 | 0.77548594 | 2.60001639 | 0.27252956 | 0.44584616 | NDI1 | XIII | 28266 | 29807 | -1 |
| YML121W | 13.6359108 | -1.5243497 | 0.82983021 | 3.33052931 | 0.18914059 | 0.34524718 | GTR1 | XIII | 26930 | 27862 | 1 |
| YML122C | 254.232888 | -0.6962283 | 0.50601011 | 2.23148788 | 0.32767142 | 0.50039829 | NA | XIII | 26038 | 26418 | -1 |
| YML123C | 1912.46145 | -0.3073352 | 0.4795027 | 0.5934124 | 0.74326235 | 0.8433253 | PHO84 | XIII | 24037 | 25800 | -1 |
| YML124C | 73.3289378 | 1.01653946 | 0.58382247 | 3.04936011 | 0.21769069 | 0.38155469 | TUB3 | XIII | 22048 | 23683 | -1 |
| YML125C | 526.521536 | -1.2910463 | 0.53815094 | 6.01353847 | 0.04945119 | 0.12955763 | PGA3 | XIII | 20761 | 21699 | -1 |
| YML126C | 113.072242 | -0.8991104 | 0.56208519 | 3.79773066 | 0.14973843 | 0.29250523 | ERG13 | XIII | 19060 | 20535 | -1 |
| YML127W | 12.3468812 | 0.12684486 | 0.88768484 | 0.06790372 | 0.96661804 | 0.97486964 | RSC9 | XIII | 17064 | 18809 | 1 |

| | | | | | | | | | | | |
|---|---|---|---|---|---|---|---|---|---|---|---|
| YML128C | 14.8391883 | 1.65095326 | 0.90384718 | 4.43364252 | 0.1089549 | 0.23299267 | MSC1 | XIII | 15135 | 16676 | -1 |
| YML129C | 6.88575157 | 0.25742317 | 1.12927836 | 0.66017479 | 0.71886091 | 0.82938751 | COX14 | XIII | 14541 | 14753 | -1 |
| YML130C | 88.1022797 | 1.77372251 | 0.60735406 | 10.4606207 | 0.00535186 | 0.02418992 | ERO1 | XIII | 11483 | 13174 | -1 |
| YML131W | 14.9524082 | 0.00149679 | 0.77635117 | 0.05551388 | 0.97262475 | 0.97905029 | NA | XIII | 10198 | 11295 | 1 |
| YML133C | 11.5094429 | -0.4788943 | 0.88064655 | 3.58466689 | 0.16657103 | 0.31564276 | NA | XIII | 461 | 4684 | -1 |
| YMR001C | 64.4256951 | 1.55696962 | 0.59804704 | 13.1517357 | 0.0013936 | 0.00862986 | CDC5 | XIII | 269019 | 271136 | -1 |
| YMR002W | 63.783678 | -0.0879196 | 0.57699057 | 0.18597073 | 0.91120684 | 0.94542349 | MIX17 | XIII | 272193 | 272663 | 1 |
| YMR003W | 46.5387994 | 0.4420037 | 0.59381656 | 2.4822426 | 0.28905991 | 0.46297563 | AIM34 | XIII | 273118 | 273714 | 1 |
| YMR004W | 48.8037284 | 1.11525465 | 0.58166032 | 5.02547894 | 0.08104591 | 0.1881539 | MVP1 | XIII | 274017 | 275552 | 1 |
| YMR005W | 236.768225 | 1.70850903 | 0.47700976 | 17.7652578 | 0.00013878 | 0.0012449 | TAF4 | XIII | 276045 | 277211 | 1 |
| YMR006C | 1733.72974 | 2.19527057 | 0.48648381 | 18.8439444 | 8.09E-05 | 0.00078224 | PLB2 | XIII | 277561 | 279681 | -1 |
| YMR007W | 2468.55699 | 0.64572296 | 0.5589992 | 18.8042747 | 8.25E-05 | 0.00079658 | NA | XIII | 279960 | 280340 | 1 |
| YMR008C | 112.197859 | 1.49675761 | 0.78148709 | 5.61451275 | 0.0603704 | 0.15063165 | PLB1 | XIII | 280590 | 282584 | -1 |
| YMR008C-A | 112.497476 | 1.6679144 | 0.53258936 | 10.1311087 | 0.00631041 | 0.02756307 | NA | NA | NA | NA | NA |
| YMR009W | 15.3253306 | 0.19008115 | 0.83050118 | 0.74594413 | 0.68868447 | 0.81049746 | ADI1 | XIII | 284102 | 284641 | 1 |
| YMR010W | 29.6458472 | -0.6525332 | 0.684314 | 1.05350261 | 0.59052028 | 0.73767258 | ANY1 | XIII | 285100 | 286317 | 1 |
| YMR011W | 171.711574 | 1.80516739 | 0.53362347 | 24.4765424 | 4.84E-06 | 6.90E-05 | HXT2 | XIII | 288079 | 289704 | 1 |
| YMR012W | 75.4907809 | -1.7449393 | 0.54660953 | 9.99047116 | 0.00677013 | 0.02923119 | CLU1 | XIII | 291134 | 294967 | 1 |
| YMR014W | 107.202825 | -1.5566092 | 0.62694282 | 6.62125991 | 0.03649318 | 0.10388176 | BUD22 | XIII | 298868 | 300427 | 1 |
| YMR015C | 25.1016391 | 1.47676926 | 0.73173152 | 6.94376112 | 0.03105857 | 0.09182162 | ERG5 | XIII | 300869 | 302485 | -1 |
| YMR016C | 204.248051 | -0.900262 | 0.50845795 | 3.0816558 | 0.21420369 | 0.37813088 | SOK2 | XIII | 303236 | 305593 | -1 |
| YMR017W | 36.7532861 | -1.6816844 | 0.62215551 | 9.14753313 | 0.01031902 | 0.04032978 | SPO20 | XIII | 307489 | 308682 | 1 |
| YMR018W | 59.5061239 | 0.68566083 | 0.92612956 | 6.00634065 | 0.04962948 | 0.12996593 | PEX9 | XIII | 310208 | 311752 | 1 |
| YMR019W | 246.029845 | 4.02241422 | 0.55882525 | 45.2526342 | 1.49E-10 | 6.80E-09 | STB4 | XIII | 312156 | 315005 | 1 |
| YMR020W | 268.222547 | -0.07389 | 0.64647518 | 0.13938119 | 0.93268235 | 0.95731799 | FMS1 | XIII | 315377 | 316903 | 1 |
| YMR021C | 94.7806107 | -0.7814025 | 0.5568991 | 2.42622122 | 0.29727115 | 0.47117436 | MAC1 | XIII | 317165 | 318418 | -1 |
| YMR022W | 11.0636959 | 1.02719368 | 0.96144033 | 1.36769023 | 0.50467273 | 0.66289817 | UBC7 | XIII | 318680 | 319177 | 1 |
| YMR023C | 110.571512 | 1.14436524 | 0.58414875 | 5.19168299 | 0.07458309 | 0.17654787 | MSS1 | XIII | 319437 | 321017 | -1 |
| YMR024W | 6.49353361 | -0.5750947 | 1.10983335 | 0.33730925 | 0.84480062 | 0.90911019 | MRPL3 | XIII | 321875 | 323047 | 1 |
| YMR025W | 188.326141 | 1.70772153 | 0.84487507 | 12.2737722 | 0.00216164 | 0.01184098 | CSI1 | XIII | 323300 | 324187 | 1 |
| YMR026C | 150.884966 | -0.5654496 | 0.51048503 | 1.88534121 | 0.38958601 | 0.56101577 | PEX12 | XIII | 324236 | 325435 | -1 |
| YMR027W | 26.2509223 | 0.47908002 | 0.95962671 | 0.70644384 | 0.70242129 | 0.81847842 | NA | XIII | 325877 | 327289 | 1 |
| YMR028W | 212.914341 | 0.75934148 | 0.93626111 | 8.69412505 | 0.01294478 | 0.04783043 | TAP42 | XIII | 327482 | 328582 | 1 |

| | | | | | | | | | | | |
|---|---|---|---|---|---|---|---|---|---|---|---|
| YMR029C | 123.952837 | 2.8343362 | 0.75286795 | 12.9990031 | 0.00150419 | 0.00913878 | FAR8 | XIII | 328660 | 330231 | -1 |
| YMR030W | 26.0597765 | 0.31096821 | 0.76717106 | 0.17369188 | 0.91681833 | 0.94860224 | RSF1 | XIII | 330793 | 331923 | 1 |
| YMR030W-A | 11.7322247 | 1.66779368 | 0.96154625 | 2.98150187 | 0.22520348 | 0.39051457 | NA | XIII | 337313 | 337603 | 1 |
| YMR031C | 41.3945074 | -0.3627252 | 0.66132962 | 0.4839551 | 0.7850738 | 0.86996695 | EIS1 | XIII | 332212 | 334743 | -1 |
| YMR032W | 15.4277085 | -0.945181 | 0.95217359 | 1.27363345 | 0.52897362 | 0.68319367 | HOF1 | XIII | 335298 | 337307 | 1 |
| YMR033W | 103.899303 | 3.668877 | 0.65082859 | 40.181675 | 1.88E-09 | 6.60E-08 | ARP9 | XIII | 337788 | 339277 | 1 |
| YMR034C | 115.266115 | -0.7329374 | 0.62288341 | 1.37049473 | 0.50396555 | 0.6624201 | RCH1 | XIII | 339418 | 340722 | -1 |
| YMR035W | 53.2344765 | 0.33905174 | 0.60580477 | 0.34738216 | 0.84055652 | 0.90645906 | IMP2 | XIII | 341142 | 341675 | 1 |
| YMR036C | 23.8936685 | 0.28940071 | 0.93299953 | 7.9667577 | 0.01862261 | 0.0626889 | MIH1 | XIII | 341856 | 343520 | -1 |
| YMR037C | 91.9025761 | 2.50858765 | 0.58356003 | 21.4440308 | 2.21E-05 | 0.00025693 | MSN2 | XIII | 344403 | 346517 | -1 |
| YMR038C | 537.351494 | -1.0237156 | 0.48597165 | 4.71678461 | 0.09457214 | 0.21003333 | CCS1 | XIII | 347511 | 348260 | -1 |
| YMR039C | 159.641959 | -0.493586 | 0.50425945 | 3.84924875 | 0.14593056 | 0.28719849 | SUB1 | XIII | 348644 | 349522 | -1 |
| YMR040W | 139.229145 | 2.98388589 | 1.05286437 | 15.5023201 | 0.00043024 | 0.0032521 | YET2 | XIII | 350381 | 350863 | 1 |
| YMR041C | 132.286413 | 1.78732599 | 0.52220007 | 11.1546135 | 0.00378274 | 0.01836234 | ARA2 | XIII | 350966 | 351973 | -1 |
| YMR042W | 177.280757 | -0.8065252 | 0.50597068 | 2.77170394 | 0.25011062 | 0.42121597 | ARG80 | XIII | 352603 | 353136 | 1 |
| YMR043W | 234.040152 | 0.70121273 | 0.747766 | 7.15218968 | 0.02798477 | 0.08430376 | MCM1 | XIII | 353871 | 354731 | 1 |
| YMR044W | 77.8223333 | 1.06779017 | 0.53790256 | 4.90355174 | 0.08614048 | 0.19736975 | IOC4 | XIII | 355384 | 356811 | 1 |
| YMR045C | 7.14014222 | -1.2767855 | 1.19531519 | 1.62796025 | 0.44309099 | 0.61162622 | NA | XIII | 357359 | 362627 | -1 |
| YMR047C | 48.5533361 | -0.9001695 | 0.61909396 | 3.07094727 | 0.21535367 | 0.37922681 | NUP116 | XIII | 363364 | 366705 | -1 |
| YMR048W | 47.6138775 | -3.2500416 | 0.85058842 | 17.3317136 | 0.00017237 | 0.00149406 | CSM3 | XIII | 366981 | 367934 | 1 |
| YMR049C | 47.4267326 | -2.7488846 | 0.64722253 | 17.4012326 | 0.00016648 | 0.00145171 | ERB1 | XIII | 368094 | 370517 | -1 |
| YMR052W | 63.2738524 | 2.22238795 | 0.64850813 | 16.3725746 | 0.00027845 | 0.00229332 | FAR3 | XIII | 379586 | 380200 | 1 |
| YMR053C | 395.186856 | -0.171794 | 0.57316396 | 0.34738535 | 0.84055518 | 0.90645906 | STB2 | XIII | 380346 | 382898 | -1 |
| YMR054W | 323.372774 | -1.9503459 | 0.55118227 | 12.9031458 | 0.00157804 | 0.00949776 | STV1 | XIII | 383303 | 385975 | 1 |
| YMR055C | 90.8357744 | -0.1529989 | 0.54568171 | 0.55486397 | 0.7577271 | 0.85371471 | BUB2 | XIII | 386101 | 387021 | -1 |
| YMR056C | 62.5697369 | 0.9361859 | 0.64987937 | 13.2366635 | 0.00133566 | 0.00831554 | AAC1 | XIII | 387315 | 388244 | -1 |
| YMR058W | 427.046514 | 2.45561701 | 0.57592869 | 25.1894024 | 3.39E-06 | 5.12E-05 | FET3 | XIII | 388822 | 390732 | 1 |
| YMR060C | 183.718682 | -1.6144728 | 0.6229029 | 7.25902376 | 0.02652913 | 0.08104993 | SAM37 | XIII | 391532 | 392515 | -1 |
| YMR061W | 11.7091193 | -0.8045543 | 0.91561019 | 0.8017716 | 0.66972654 | 0.79510827 | RNA14 | XIII | 392755 | 394788 | 1 |
| YMR062C | 66.3844012 | -0.519691 | 0.62515077 | 1.88965463 | 0.3887467 | 0.56060856 | ARG7 | XIII | 395054 | 396379 | -1 |
| YMR063W | 108.81423 | 0.21816283 | 0.52324426 | 0.5111789 | 0.77445986 | 0.86282905 | RIM9 | XIII | 397077 | 397796 | 1 |
| YMR064W | 19.3955902 | -0.0445821 | 0.74725992 | 1.07913714 | 0.58299972 | 0.7314341 | AEP1 | XIII | 397903 | 399459 | 1 |
| YMR065W | 104.896978 | 0.93724617 | 0.52240538 | 7.55224637 | 0.02291134 | 0.07329098 | KAR5 | XIII | 399702 | 401216 | 1 |

| | | | | | | | | | | | |
|---|---|---|---|---|---|---|---|---|---|---|---|
| YMR066W | 52.0377484 | 0.53632658 | 0.94337887 | 4.33566817 | 0.11442518 | 0.24134059 | SOV1 | XIII | 401541 | 404237 | 1 |
| YMR067C | 21.7461241 | 0.4218783 | 0.71224294 | 0.6046635 | 0.73909283 | 0.84040603 | UBX4 | XIII | 404323 | 405573 | -1 |
| YMR068W | 17.0363636 | 0.82570591 | 0.76389607 | 1.64350459 | 0.43966056 | 0.60823052 | AVO2 | XIII | 406304 | 407584 | 1 |
| YMR069W | 25.0370625 | 0.61579909 | 0.79101939 | 0.94622382 | 0.62306034 | 0.76286238 | NAT4 | XIII | 407709 | 408566 | 1 |
| YMR070W | 635.192795 | 5.86425336 | 0.55689909 | 84.1103701 | 5.44E-19 | 1.21E-16 | MOT3 | XIII | 409154 | 410626 | 1 |
| YMR071C | 91.2768935 | 0.09285803 | 0.76587743 | 4.60431893 | 0.10004257 | 0.21919967 | TVP18 | XIII | 410762 | 411265 | -1 |
| YMR072W | 170.847745 | -0.7080514 | 0.5079579 | 2.14006927 | 0.34299664 | 0.51663645 | ABF2 | XIII | 411569 | 412120 | 1 |
| YMR073C | 24.9586269 | -0.3719562 | 0.78978786 | 3.05743229 | 0.21681385 | 0.380987 | IRC21 | XIII | 412268 | 412873 | -1 |
| YMR074C | 319.915176 | -0.3966902 | 0.5227479 | 1.50371684 | 0.47148951 | 0.6333931 | SDD2 | XIII | 413036 | 413473 | -1 |
| YMR075C-A | 1154.45491 | -0.0548646 | 0.61451527 | 1.17137007 | 0.55672436 | 0.70829137 | NA | XIII | 415688 | 416053 | -1 |
| YMR075W | 95.8032476 | -2.2473702 | 0.5990607 | 13.5332329 | 0.00115158 | 0.0073921 | RCO1 | XIII | 413982 | 416036 | 1 |
| YMR076C | 104.874823 | 0.75458254 | 0.50751585 | 3.05146422 | 0.21746179 | 0.38155469 | PDS5 | XIII | 416196 | 420029 | -1 |
| YMR077C | 18.688232 | 0.89615538 | 0.81295222 | 2.85500919 | 0.23990684 | 0.40806716 | VPS20 | XIII | 421484 | 422149 | -1 |
| YMR078C | 322.054751 | 0.7108528 | 0.4909239 | 8.87355536 | 0.01183401 | 0.04466683 | CTF18 | XIII | 422503 | 424728 | -1 |
| YMR079W | 24.2044627 | -0.5176294 | 0.83179469 | 0.44727155 | 0.79960632 | 0.87884032 | SEC14 | XIII | 424989 | 426059 | 1 |
| YMR080C | 20.4186915 | -0.2427238 | 0.74355574 | 0.94533827 | 0.62333628 | 0.76302686 | NAM7 | XIII | 426712 | 429627 | -1 |
| YMR081C | 455.523337 | 0.35137141 | 0.49122341 | 1.29572048 | 0.52316402 | 0.67871828 | ISF1 | XIII | 430079 | 431095 | -1 |
| YMR083W | 88.866678 | -0.5371814 | 1.68791862 | 0.11776601 | 0.94281707 | 0.96176085 | ADH3 | XIII | 434788 | 435915 | 1 |
| YMR085W | 10.9076129 | 1.02495379 | 1.08517877 | 1.00310118 | 0.60559091 | 0.75082898 | NA | XIII | 437491 | 438789 | 1 |
| YMR086W | 442.428445 | -1.3926312 | 0.58675284 | 5.92310874 | 0.05173843 | 0.13421395 | SEG1 | XIII | 439208 | 442090 | 1 |
| YMR087W | 417.279923 | 0.34174421 | 0.59833072 | 2.16077159 | 0.33946454 | 0.51259098 | NA | XIII | 442527 | 443381 | 1 |
| YMR088C | 633.536411 | 3.20436759 | 0.61612147 | 24.5185351 | 4.74E-06 | 6.83E-05 | VBA1 | XIII | 443414 | 445102 | -1 |
| YMR089C | 33.0238837 | -0.1247274 | 0.62319512 | 0.66270647 | 0.71795152 | 0.82923185 | YTA12 | XIII | 445609 | 448086 | -1 |
| YMR090W | 27.7869719 | -0.0453387 | 0.70504916 | 0.21321937 | 0.89887645 | 0.93754851 | NA | XIII | 449245 | 449928 | 1 |
| YMR091C | 75.6216463 | 0.19718391 | 0.6338649 | 0.92617228 | 0.62933842 | 0.76680754 | NPL6 | XIII | 450058 | 451365 | -1 |
| YMR092C | 23.8554022 | 0.42388037 | 0.86880087 | 0.81631602 | 0.66487381 | 0.7916141 | AIP1 | XIII | 451632 | 453479 | -1 |
| YMR093W | 9.83668325 | -2.2897681 | 1.28394559 | 3.53575451 | 0.17069495 | 0.32088433 | UTP15 | XIII | 454015 | 455556 | 1 |
| YMR094W | 600.557516 | 3.52598306 | 0.57966419 | 32.2747037 | 9.81E-08 | 2.40E-06 | CTF13 | XIII | 455825 | 457261 | 1 |
| YMR095C | 600.469589 | -1.1011434 | 0.47948203 | 7.46832016 | 0.02389323 | 0.07547289 | SNO1 | XIII | 457285 | 457959 | -1 |
| YMR096W | 44.5718488 | 2.71353404 | 0.7251227 | 17.4123505 | 0.00016556 | 0.00144584 | SNZ1 | XIII | 458408 | 459301 | 1 |
| YMR097C | 860.395237 | 3.52366924 | 0.74775984 | 22.4058367 | 1.36E-05 | 0.00017236 | MTG1 | XIII | 459424 | 460527 | -1 |
| YMR098C | 88.2131072 | 0.4914527 | 0.63729716 | 2.3461783 | 0.30940965 | 0.48239987 | ATP25 | XIII | 460771 | 462609 | -1 |
| YMR099C | 21.6920342 | 0.22485735 | 0.80008576 | 0.10920212 | 0.94686281 | 0.96418144 | NA | XIII | 463934 | 464827 | -1 |

| | | | | | | | | | | | |
|---|---|---|---|---|---|---|---|---|---|---|---|
| YMR100W | 38.3055422 | 0.68516147 | 0.72978383 | 6.70572308 | 0.0349841 | 0.10042536 | MUB1 | XIII | 466300 | 468162 | 1 |
| YMR101C | 11.2113408 | 0.52411819 | 0.84799423 | 3.42536117 | 0.18038162 | 0.33506648 | SRT1 | XIII | 468445 | 469476 | -1 |
| YMR102C | 63.8429626 | 3.03131082 | 0.60791269 | 23.9829022 | 6.20E-06 | 8.57E-05 | NA | XIII | 469848 | 472352 | -1 |
| YMR103C | 12.1600389 | 2.74686401 | 1.25366456 | 5.02167994 | 0.08120001 | 0.18843608 | NA | XIII | 472902 | 473264 | -1 |
| YMR104C | 83.8299788 | 1.6156484 | 0.66258989 | 7.06702577 | 0.02920215 | 0.08706512 | YPK2 | XIII | 473420 | 475453 | -1 |
| YMR105C | 23.3300621 | 2.39191 | 0.77225628 | 10.3719875 | 0.00559437 | 0.02503202 | PGM2 | XIII | 475897 | 477606 | -1 |
| YMR105W-A | 276.814889 | -0.0820989 | 0.53124008 | 0.02646183 | 0.98685623 | 0.98907696 | NA | XIII | 478064 | 478258 | 1 |
| YMR106C | 21.2639694 | 1.14800664 | 0.78200266 | 2.39093065 | 0.30256313 | 0.47513982 | YKU80 | XIII | 478301 | 480190 | -1 |
| YMR108W | 61.6187542 | -1.2143874 | 0.59308429 | 6.39276525 | 0.04090992 | 0.11317174 | ILV2 | XIII | 484084 | 486147 | 1 |
| YMR109W | 36.0271123 | 0.38757784 | 0.84349356 | 5.14017788 | 0.07652874 | 0.18012252 | MYO5 | XIII | 486587 | 490246 | 1 |
| YMR110C | 44.2777111 | -0.139992 | 0.61951877 | 0.87192461 | 0.6466421 | 0.77855225 | HFD1 | XIII | 490394 | 491992 | -1 |
| YMR111C | 23.6371965 | -0.11735 | 0.69329804 | 0.34438912 | 0.84181537 | 0.90731776 | NA | XIII | 492404 | 493792 | -1 |
| YMR112C | 9.70930219 | 0.14876398 | 0.93123135 | 0.05840111 | 0.97122166 | 0.9781481 | NA | XIII | 494100 | 494495 | -1 |
| YMR113W | 129.627505 | 1.17230585 | 0.85057144 | 4.88649529 | 0.08687824 | 0.19866707 | FOL3 | XIII | 494999 | 496282 | 1 |
| YMR114C | 65.3201365 | 1.29041214 | 0.9463748 | 6.6577345 | 0.03583367 | 0.10225577 | NA | XIII | 496343 | 497449 | -1 |
| YMR115W | 13.0570643 | -1.4386925 | 0.88687257 | 3.00564288 | 0.2225015 | 0.38746266 | MGR3 | XIII | 497704 | 499209 | 1 |
| YMR116C | 56.3146113 | -2.8570985 | 0.70353257 | 15.4107974 | 0.00045039 | 0.00337792 | ASC1 | XIII | 499456 | 500688 | -1 |
| YMR117C | 33.5603311 | -2.1143105 | 0.63117818 | 12.5344517 | 0.00189749 | 0.01073943 | SPC24 | XIII | 501250 | 501891 | -1 |
| YMR118C | 557.329708 | -2.3317541 | 0.50508881 | 20.1290894 | 4.26E-05 | 0.0004497 | SHH3 | XIII | 502144 | 502734 | -1 |
| YMR119W | 20.0257178 | 0.01204408 | 0.82481203 | 0.06615971 | 0.9674613 | 0.97486964 | ASI1 | XIII | 505333 | 507207 | 1 |
| YMR120C | 556.624801 | 0.05977785 | 0.53543216 | 2.82769055 | 0.24320629 | 0.41283037 | ADE17 | XIII | 507502 | 509280 | -1 |
| YMR121C | 33.8845462 | 0.68231442 | 0.72054953 | 3.36556832 | 0.1858558 | 0.34169876 | RPL15B | XIII | 509734 | 510348 | -1 |
| YMR122C | 71.4408725 | -1.8416509 | 0.63593243 | 15.059504 | 0.00053687 | 0.00388561 | NA | XIII | 510701 | 511075 | -1 |
| YMR122W-A | 88.3203857 | 1.12397568 | 0.56513153 | 12.2081162 | 0.00223378 | 0.01216709 | NA | XIII | 511315 | 511569 | 1 |
| YMR123W | 94.8121014 | -0.1068158 | 0.51094219 | 0.05093939 | 0.97485192 | 0.98061025 | PKR1 | XIII | 513593 | 513961 | 1 |
| YMR124W | 92.996837 | 0.75871205 | 0.58321426 | 2.36798912 | 0.30605374 | 0.47867401 | EPO1 | XIII | 514456 | 517287 | 1 |
| YMR125W | 16.2205696 | -1.8593583 | 1.16679158 | 2.54785182 | 0.27973126 | 0.45380891 | STO1 | XIII | 517539 | 520446 | 1 |
| YMR126C | 18.8130016 | 1.26900563 | 0.84464646 | 2.7763635 | 0.2495286 | 0.42048038 | DLT1 | XIII | 520761 | 521789 | -1 |
| YMR127C | 992.383999 | 3.78746693 | 0.46048162 | 66.4345489 | 3.75E-15 | 4.02E-13 | SAS2 | XIII | 522329 | 523345 | -1 |
| YMR128W | 41.5662843 | -1.7936674 | 0.63441377 | 7.86122349 | 0.01963166 | 0.06510155 | ECM16 | XIII | 523696 | 527499 | 1 |
| YMR129W | 58.959327 | 1.26697328 | 0.63965344 | 6.87410442 | 0.03215934 | 0.0942798 | POM152 | XIII | 527804 | 531817 | 1 |
| YMR130W | 179.659093 | -0.3601199 | 0.5974307 | 0.3749221 | 0.82906141 | 0.89791724 | NA | XIII | 532119 | 533027 | 1 |
| YMR131C | 72.1682185 | -1.7546654 | 0.6129604 | 8.51891203 | 0.01412999 | 0.05110095 | RRB1 | XIII | 533163 | 534698 | -1 |

| | | | | | | | | | | | |
|---|---|---|---|---|---|---|---|---|---|---|---|
| YMR132C | 58.6891595 | -3.2033138 | 0.59396259 | 28.4617005 | 6.60E-07 | 1.23E-05 | JLP2 | XIII | 534944 | 535570 | -1 |
| YMR133W | 888.151951 | 0.42399782 | 0.59252352 | 0.56043513 | 0.75561933 | 0.85233507 | REC114 | XIII | 536207 | 537609 | 1 |
| YMR134W | 27.4120029 | 0.13487008 | 0.69703448 | 0.13611666 | 0.93420598 | 0.95735444 | ERG29 | XIII | 537838 | 538551 | 1 |
| YMR135C | 372.328043 | -0.847527 | 0.51213618 | 4.33236104 | 0.11461455 | 0.24149136 | GID8 | XIII | 538689 | 540056 | -1 |
| YMR136W | 32.0518832 | 0.33768719 | 0.69214629 | 0.46459549 | 0.79271006 | 0.87407947 | GAT2 | XIII | 541199 | 542881 | 1 |
| YMR137C | 16.0910266 | 0.36272386 | 0.78060564 | 1.11004653 | 0.57405891 | 0.72398194 | PSO2 | XIII | 542978 | 544963 | -1 |
| YMR138W | 10.4156651 | 2.44749094 | 1.03197997 | 6.3229871 | 0.04236242 | 0.11597089 | CIN4 | XIII | 545155 | 545730 | 1 |
| YMR139W | 14.5939226 | 1.04684942 | 0.80722141 | 1.76998813 | 0.41271662 | 0.58426143 | RIM11 | XIII | 546125 | 547237 | 1 |
| YMR140W | 22.4743501 | 1.28270225 | 0.76931215 | 2.85577264 | 0.23981528 | 0.40803129 | SIP5 | XIII | 547714 | 549183 | 1 |
| YMR141C | 1612.15191 | 1.51590236 | 0.45952421 | 12.204249 | 0.00223811 | 0.01217918 | NA | XIII | 549736 | 550044 | -1 |
| YMR142C | 71.1469457 | -2.8860449 | 0.69726143 | 15.845667 | 0.00036237 | 0.0028442 | RPL13B | XIII | 550206 | 551207 | -1 |
| YMR143W | 30.3857902 | -3.0216083 | 2.15935613 | 1.73008056 | 0.42103459 | 0.59184032 | RPS16A | XIII | 551928 | 552903 | 1 |
| YMR144W | 38.2350813 | 0.88797789 | 0.62107418 | 2.69944716 | 0.25931193 | 0.43094606 | FDO1 | XIII | 553362 | 554390 | 1 |
| YMR145C | 76.1691602 | -0.6160303 | 0.67460077 | 2.75494896 | 0.25221472 | 0.42365049 | NDE1 | XIII | 554793 | 556475 | -1 |
| YMR146C | 84.9429412 | -2.1546394 | 0.71962551 | 8.46331433 | 0.01452829 | 0.0523127 | TIF34 | XIII | 557481 | 558524 | -1 |
| YMR147W | 35.163466 | -1.5587313 | 0.62740168 | 6.48368992 | 0.03909171 | 0.10976769 | NA | XIII | 559199 | 559870 | 1 |
| YMR149W | 74.0052761 | -0.3177933 | 0.8055409 | 8.44156905 | 0.01468712 | 0.05261147 | SWP1 | XIII | 560996 | 561856 | 1 |
| YMR150C | 131.440535 | 0.6626661 | 0.77642561 | 6.25911082 | 0.04373724 | 0.1185574 | IMP1 | XIII | 561956 | 562528 | -1 |
| YMR152W | 19.27489 | -1.3269765 | 0.73549845 | 3.34325046 | 0.18794137 | 0.34376539 | YIM1 | XIII | 563096 | 564193 | 1 |
| YMR153W | 689.723149 | -1.0695186 | 0.51029763 | 5.07005742 | 0.07925944 | 0.18517193 | NUP53 | XIII | 564435 | 565862 | 1 |
| YMR154C | 483.685533 | -1.3824789 | 0.45983889 | 8.88698187 | 0.01175483 | 0.04445491 | RIM13 | XIII | 565999 | 568182 | -1 |
| YMR155W | 50.6163872 | 0.78953033 | 0.57553715 | 2.78594458 | 0.24833608 | 0.41920289 | NA | XIII | 568551 | 570194 | 1 |
| YMR156C | 725.556989 | 0.21106042 | 0.63058487 | 0.52992353 | 0.76723529 | 0.85857988 | TPP1 | XIII | 570300 | 571016 | -1 |
| YMR157C | 180.484619 | -0.1944316 | 0.61529726 | 0.76861878 | 0.68092072 | 0.80384598 | AIM36 | XIII | 571277 | 572044 | -1 |
| YMR158W | 31.1766209 | -1.7439005 | 0.7323508 | 7.3389446 | 0.02548992 | 0.07880632 | MRPS8 | XIII | 572248 | 572715 | 1 |
| YMR160W | 302.71924 | 1.98281155 | 0.50555033 | 20.782502 | 3.07E-05 | 0.00034448 | NA | XIII | 575066 | 577516 | 1 |
| YMR161W | 11.9788389 | -0.2449984 | 0.88587316 | 0.16990984 | 0.91855369 | 0.94954934 | HLJ1 | XIII | 577718 | 578392 | 1 |
| YMR162C | 40.4705831 | 0.70820014 | 0.62552199 | 1.51839345 | 0.46804224 | 0.63108197 | DNF3 | XIII | 578951 | 583921 | -1 |
| YMR163C | 46.2938112 | -2.4407091 | 0.62378436 | 19.8181619 | 4.97E-05 | 0.00051177 | INP2 | XIII | 584271 | 586388 | -1 |
| YMR164C | 102.876209 | 1.8005727 | 0.61891047 | 11.1748687 | 0.00374462 | 0.01821964 | MSS11 | XIII | 587274 | 589550 | -1 |
| YMR165C | 562.182382 | 0.97494158 | 0.46826852 | 4.9449535 | 0.08437562 | 0.19386304 | PAH1 | XIII | 590040 | 592628 | -1 |
| YMR166C | 10.7988574 | -0.3222179 | 0.92309258 | 0.16993789 | 0.91854081 | 0.94954934 | MME1 | XIII | 593367 | 594473 | -1 |
| YMR167W | 852.509593 | -0.6832558 | 0.49743439 | 2.56054845 | 0.27796107 | 0.45261809 | MLH1 | XIII | 594886 | 597195 | 1 |

| | | | | | | | | | | | |
|---|---|---|---|---|---|---|---|---|---|---|---|
| YMR168C | 977.985175 | -0.7459991 | 0.47552489 | 2.43503816 | 0.29596352 | 0.46995614 | CEP3 | XIII | 597332 | 599158 | -1 |
| YMR169C | 35.1993292 | -1.4227975 | 0.62779749 | 5.19738861 | 0.07437062 | 0.17629168 | ALD3 | XIII | 599352 | 600872 | -1 |
| YMR170C | 166.388819 | 1.20710649 | 0.49320061 | 6.05900146 | 0.04833977 | 0.12713301 | ALD2 | XIII | 601562 | 603082 | -1 |
| YMR171C | 44.489071 | 3.16365387 | 0.68136398 | 28.9076869 | 5.28E-07 | 1.02E-05 | EAR1 | XIII | 603868 | 605520 | -1 |
| YMR172W | 118.53845 | -0.171543 | 0.5078876 | 0.30868075 | 0.85698028 | 0.91531374 | HOT1 | XIII | 605981 | 608140 | 1 |
| YMR173W | 79.5212963 | -0.5911388 | 0.53993441 | 1.43737404 | 0.48739177 | 0.64903366 | DDR48 | XIII | 608689 | 609981 | 1 |
| YMR174C | 135.488713 | 1.49534054 | 0.85172464 | 9.21220427 | 0.00999068 | 0.03943154 | PAI3 | XIII | 610159 | 610365 | -1 |
| YMR175W-A | 23.8353227 | 2.16124077 | 0.76241377 | 9.49180201 | 0.00868723 | 0.03535208 | NA | XIII | 611314 | 611508 | 1 |
| YMR176W | 47.8402548 | 0.95569256 | 0.58640246 | 3.26701851 | 0.19524321 | 0.35245914 | ECM5 | XIII | 611740 | 615975 | 1 |
| YMR177W | 42.6302027 | 3.90543725 | 0.73330491 | 34.2748927 | 3.61E-08 | 9.72E-07 | MMT1 | XIII | 616566 | 618098 | 1 |
| YMR178W | 316.059453 | -3.4088466 | 0.5109999 | 48.4470682 | 3.02E-11 | 1.55E-09 | NA | XIII | 618479 | 619303 | 1 |
| YMR179W | 366.546287 | 1.63593728 | 0.56588141 | 12.5875969 | 0.00184773 | 0.01056192 | SPT21 | XIII | 619858 | 622134 | 1 |
| YMR180C | 349.881297 | 3.41749869 | 0.50447752 | 46.1211437 | 9.66E-11 | 4.58E-09 | CTL1 | XIII | 622251 | 623213 | -1 |
| YMR181C | 602.157426 | -1.8231658 | 0.46875534 | 14.4223485 | 0.00073829 | 0.00511327 | NA | XIII | 623616 | 624080 | -1 |
| YMR182C | 789.244588 | 0.13679746 | 0.48280747 | 0.83027263 | 0.66025027 | 0.78805382 | RGM1 | XIII | 624532 | 625167 | -1 |
| YMR182W-A | 30.6548282 | 2.45789923 | 0.73806705 | 11.5989766 | 0.0030291 | 0.01538156 | NA | XIII | 625811 | 625897 | 1 |
| YMR183C | 39.114781 | -1.659632 | 0.75934444 | 5.05197647 | 0.07997924 | 0.18642503 | SSO2 | XIII | 626921 | 627808 | -1 |
| YMR184W | 21.6053344 | 2.05458542 | 0.95104006 | 5.15889535 | 0.07581587 | 0.17873529 | ADD37 | XIII | 628189 | 628785 | 1 |
| YMR185W | 12.4895121 | 0.11674122 | 0.91782995 | 0.08617217 | 0.95782893 | 0.9702291 | RTP1 | XIII | 629025 | 631970 | 1 |
| YMR186W | 1339.60841 | -0.5972858 | 0.52236146 | 2.79046993 | 0.24777481 | 0.41849946 | HSC82 | XIII | 632355 | 634472 | 1 |
| YMR187C | 184.922633 | 1.01408485 | 0.5010577 | 4.19293515 | 0.12288976 | 0.25402775 | NA | XIII | 634689 | 635984 | -1 |
| YMR188C | 17.9843346 | -0.931583 | 1.07287463 | 1.00083122 | 0.60627863 | 0.75088539 | MRPS17 | XIII | 636291 | 637004 | -1 |
| YMR189W | 21.7328492 | 0.30512552 | 0.70422182 | 0.42901673 | 0.80693806 | 0.88421109 | GCV2 | XIII | 637500 | 640604 | 1 |
| YMR190C | 21.2037388 | 0.92368882 | 0.70732675 | 1.69089565 | 0.42936504 | 0.59890573 | SGS1 | XIII | 640915 | 645258 | -1 |
| YMR191W | 78.868164 | 2.69151106 | 0.59091794 | 19.3870398 | 6.17E-05 | 0.00061682 | SPG5 | XIII | 645656 | 646777 | 1 |
| YMR192W | 86.7694842 | 3.13535569 | 0.57945329 | 32.0817225 | 1.08E-07 | 2.58E-06 | GYL1 | XIII | 647118 | 649280 | 1 |
| YMR193W | 44.0970783 | -1.4107608 | 0.59681272 | 5.58599298 | 0.06123744 | 0.15215948 | MRPL24 | XIII | 650036 | 650812 | 1 |
| YMR194C-B | 62.2148507 | -1.4885201 | 0.61947584 | 5.70144038 | 0.05780268 | 0.14608359 | CMC4 | XIII | 652594 | 652887 | -1 |
| YMR194W | 33.430147 | -2.7140112 | 0.7015402 | 14.1816655 | 0.0008327 | 0.00566552 | RPL36A | XIII | 651145 | 651910 | 1 |
| YMR195W | 42.1386467 | 1.33059984 | 1.0774421 | 5.67947817 | 0.05844091 | 0.14688185 | ICY1 | XIII | 654034 | 654417 | 1 |
| YMR196W | 512.657772 | -1.7311709 | 0.488662 | 12.6943449 | 0.00175169 | 0.01015246 | NA | XIII | 655076 | 658342 | 1 |
| YMR197C | 19.3812042 | 1.40098666 | 0.8925597 | 2.6521517 | 0.26551715 | 0.4376158 | VTI1 | XIII | 658545 | 659198 | -1 |
| YMR198W | 10.1150821 | -1.5965707 | 0.91309456 | 3.2304051 | 0.19885039 | 0.35758801 | CIK1 | XIII | 659745 | 661529 | 1 |

| | | | | | | | | | | | |
|---|---|---|---|---|---|---|---|---|---|---|---|
| YMR199W | 177.684583 | 2.74181671 | 0.56676238 | 22.5987904 | 1.24E-05 | 0.00015894 | CLN1 | XIII | 662644 | 664284 | 1 |
| YMR200W | 53.8132285 | 2.12126634 | 0.62302653 | 11.5874959 | 0.00304654 | 0.01538156 | ROT1 | XIII | 664752 | 665522 | 1 |
| YMR201C | 524.234841 | 1.02269851 | 0.88940465 | 9.63336205 | 0.00809361 | 0.03344895 | RAD14 | XIII | 665845 | 667044 | -1 |
| YMR202W | 53.4306376 | 0.23473839 | 0.65372387 | 0.74545519 | 0.68885285 | 0.81049746 | ERG2 | XIII | 667537 | 668205 | 1 |
| YMR203W | 27.731935 | -0.6222053 | 0.65629221 | 1.78056211 | 0.41054035 | 0.58217699 | TOM40 | XIII | 668492 | 669655 | 1 |
| YMR204C | 21.4404617 | 1.48834345 | 0.86613049 | 4.59874427 | 0.10032181 | 0.21944212 | INP1 | XIII | 670063 | 671325 | -1 |
| YMR205C | 70.8931425 | -1.8167983 | 0.82773058 | 5.22131703 | 0.07348614 | 0.17480884 | PFK2 | XIII | 671887 | 674766 | -1 |
| YMR206W | 182.174668 | -0.0400164 | 0.49824365 | 0.01498432 | 0.99253584 | 0.99425303 | NA | XIII | 675896 | 676837 | 1 |
| YMR207C | 49.0408939 | 0.49315551 | 0.60488243 | 0.68432214 | 0.7102338 | 0.82438173 | NA | XIII | 677193 | 683564 | -1 |
| YMR208W | 101.994348 | 0.16229249 | 0.53318828 | 0.55250415 | 0.75862168 | 0.85422394 | ERG12 | XIII | 684467 | 685798 | 1 |
| YMR209C | 24.8871767 | -1.1331391 | 0.75305097 | 6.36132442 | 0.04155813 | 0.11436385 | NA | XIII | 685911 | 687284 | -1 |
| YMR210W | 887.559967 | -0.2916192 | 0.50842127 | 2.69512177 | 0.25987335 | 0.43163129 | MGL2 | XIII | 687516 | 688865 | 1 |
| YMR211W | 80.7322697 | -3.03214 | 0.5840056 | 24.4895548 | 4.81E-06 | 6.90E-05 | DML1 | XIII | 689083 | 690510 | 1 |
| YMR212C | 37.3184876 | -1.4728665 | 0.65483453 | 5.21754312 | 0.07362493 | 0.17499522 | EFR3 | XIII | 690695 | 693043 | -1 |
| YMR213W | 21.9636381 | -0.2846659 | 0.7582941 | 0.27038655 | 0.87354706 | 0.92573532 | CEF1 | XIII | 693381 | 695153 | 1 |
| YMR214W | 82.5827693 | 2.10395451 | 0.56931889 | 13.3792579 | 0.00124374 | 0.00786166 | SCJ1 | XIII | 695350 | 696483 | 1 |
| YMR215W | 144.308261 | -1.2323059 | 0.50302363 | 5.98227498 | 0.05023027 | 0.1311241 | GAS3 | XIII | 696796 | 698370 | 1 |
| YMR216C | 34.5712365 | 0.27703696 | 0.70504299 | 0.78061441 | 0.67684891 | 0.80093096 | SKY1 | XIII | 698811 | 701039 | -1 |
| YMR217W | 77.3674681 | -2.2125216 | 0.71012163 | 9.28234748 | 0.00964637 | 0.03839103 | GUA1 | XIII | 701790 | 703367 | 1 |
| YMR218C | 18.9631284 | -0.0093546 | 0.88050541 | 0.74246825 | 0.6898824 | 0.81104957 | TRS130 | XIII | 703579 | 706887 | -1 |
| YMR219W | 253.529671 | -0.8826343 | 0.48174885 | 3.6037226 | 0.1649915 | 0.31392516 | ESC1 | XIII | 707133 | 712109 | 1 |
| YMR220W | 416.984848 | 3.90430434 | 0.57636301 | 45.8103912 | 1.13E-10 | 5.23E-09 | ERG8 | XIII | 712316 | 713671 | 1 |
| YMR221C | 96.650884 | 1.11015762 | 0.57454249 | 5.62558327 | 0.06003716 | 0.15002811 | FMP42 | XIII | 713931 | 715445 | -1 |
| YMR222C | 15.9101257 | -1.4472612 | 0.79614313 | 3.41353387 | 0.18145149 | 0.33630094 | FSH2 | XIII | 715638 | 716309 | -1 |
| YMR223W | 326.772084 | 5.43935712 | 0.59758529 | 96.492647 | 1.11E-21 | 4.61E-19 | UBP8 | XIII | 716715 | 718130 | 1 |
| YMR224C | 53.2771094 | 2.55001474 | 0.70817878 | 12.5387154 | 0.00189344 | 0.01073788 | MRE11 | XIII | 718575 | 720653 | -1 |
| YMR225C | 65.9524926 | 1.44958923 | 0.57543283 | 9.83937275 | 0.00730142 | 0.03103908 | MRPL44 | XIII | 720960 | 721403 | -1 |
| YMR226C | 87.642058 | 0.0290952 | 0.59857764 | 1.05056873 | 0.59138717 | 0.73843686 | NA | XIII | 721593 | 722396 | -1 |
| YMR227C | 108.018012 | 0.71966011 | 0.60572644 | 2.61704585 | 0.2702189 | 0.44347149 | TAF7 | XIII | 722613 | 724385 | -1 |
| YMR228W | 21.6624428 | -0.4857144 | 0.72226331 | 0.44989323 | 0.79855885 | 0.87821055 | MTF1 | XIII | 724626 | 725651 | 1 |
| YMR229C | 58.2710905 | -0.1431059 | 0.67449495 | 0.67322275 | 0.71418634 | 0.82607012 | RRP5 | XIII | 725934 | 731123 | -1 |
| YMR230W | 62.8559353 | -2.9173021 | 0.67275057 | 18.1907132 | 0.00011219 | 0.0010294 | RPS10B | XIII | 732414 | 733141 | 1 |
| YMR231W | 12.0493044 | 0.13538848 | 0.91940796 | 1.23844777 | 0.53836211 | 0.69284654 | PEP5 | XIII | 733545 | 736634 | 1 |

| | | | | | | | | | | | |
|---|---|---|---|---|---|---|---|---|---|---|---|
| YMR232W | 16.9539949 | 1.14632098 | 0.82556609 | 2.04474196 | 0.35974099 | 0.53284736 | FUS2 | XIII | 736926 | 738959 | 1 |
| YMR233W | 24.966868 | 0.88029016 | 0.70246843 | 1.66247146 | 0.43551078 | 0.60441213 | TRI1 | XIII | 739185 | 739865 | 1 |
| YMR234W | 95.2734434 | 1.15476383 | 0.6525173 | 3.80217509 | 0.14940605 | 0.29203007 | RNH1 | XIII | 740266 | 741312 | 1 |
| YMR235C | 225.586251 | 3.02360439 | 0.56354738 | 38.3802886 | 4.63E-09 | 1.50E-07 | RNA1 | XIII | 741512 | 742735 | -1 |
| YMR236W | 9.33532621 | -0.6780993 | 1.07109616 | 0.89693168 | 0.63860713 | 0.77465312 | TAF9 | XIII | 742971 | 743444 | 1 |
| YMR237W | 167.31406 | 3.59655947 | 0.56202386 | 36.7564837 | 1.04E-08 | 3.11E-07 | BCH1 | XIII | 743749 | 745923 | 1 |
| YMR238W | 53.7888874 | 0.7595551 | 0.74983328 | 2.02996011 | 0.36240966 | 0.53529386 | DFG5 | XIII | 746353 | 747729 | 1 |
| YMR239C | 109.141091 | -0.4313719 | 0.55271 | 0.72284097 | 0.69668599 | 0.81659461 | RNT1 | XIII | 748262 | 749677 | -1 |
| YMR240C | 16.1986197 | 0.41515554 | 0.8871473 | 0.27105016 | 0.87325726 | 0.92569746 | CUS1 | XIII | 749930 | 751240 | -1 |
| YMR241W | 54.6898996 | -1.5980187 | 0.60432775 | 6.89179255 | 0.03187618 | 0.09368684 | YHM2 | XIII | 751961 | 752905 | 1 |
| YMR242C | 61.3521873 | -2.5998911 | 0.57567889 | 19.4729207 | 5.91E-05 | 0.00059708 | NA | XIII | 753225 | 753742 | -1 |
| YMR242W-A | 324.962782 | -2.6391034 | 0.6265961 | 15.9625004 | 0.00034181 | 0.00272586 | NA | XIII | 754297 | 754386 | 1 |
| YMR243C | 11.7024977 | -2.7356393 | 1.1076741 | 6.16840794 | 0.04576645 | 0.12211417 | ZRC1 | XIII | 754838 | 756166 | -1 |
| YMR244C-A | 23.5356352 | -2.0557162 | 0.76612618 | 7.37666335 | 0.0250137 | 0.07790711 | COA6 | XIII | 758517 | 758831 | -1 |
| YMR244W | 16.0120832 | -1.5940599 | 0.82932586 | 5.59323216 | 0.06101619 | 0.15186924 | NA | XIII | 757250 | 758317 | 1 |
| YMR246W | 53.9384973 | -2.1878075 | 0.79423899 | 7.81021392 | 0.0201388 | 0.06640299 | FAA4 | XIII | 759807 | 761891 | 1 |
| YMR247C | 563.075547 | 0.66930472 | 0.48156312 | 1.92934131 | 0.3811087 | 0.55483451 | RKR1 | XIII | 763351 | 768039 | -1 |
| YMR247W-A | 182.409228 | 2.00833356 | 0.51279151 | 14.7870811 | 0.00061521 | 0.00439222 | NA | XIII | 769283 | 769426 | 1 |
| YMR250W | 33.4089057 | 0.50284967 | 0.67059217 | 1.34507349 | 0.51041215 | 0.66786132 | GAD1 | XIII | 770801 | 772558 | 1 |
| YMR251W | 522.415512 | 2.52264582 | 0.50905231 | 25.1356752 | 3.48E-06 | 5.24E-05 | GTO3 | XIII | 772915 | 774015 | 1 |
| YMR251W-A | 15.753937 | 1.42982945 | 0.86791731 | 5.17109151 | 0.07535494 | 0.17808372 | HOR7 | XIII | 774752 | 774931 | 1 |
| YMR252C | 21.1232859 | 0.8144379 | 0.71495865 | 1.36930801 | 0.50426467 | 0.66262751 | NA | XIII | 775315 | 775719 | -1 |
| YMR253C | 389.585196 | 1.06595873 | 0.50625855 | 4.81678072 | 0.08995998 | 0.20354369 | NA | XIII | 775946 | 777190 | -1 |
| YMR254C | 284.69552 | 1.17351941 | 0.73226104 | 2.53421914 | 0.28164452 | 0.45576349 | NA | XIII | 777615 | 777923 | -1 |
| YMR255W | 65.183087 | 0.3500035 | 0.66181935 | 0.42987045 | 0.80659369 | 0.88416839 | GFD1 | XIII | 778001 | 778567 | 1 |
| YMR257C | 20.0722321 | -0.6452785 | 0.75210765 | 1.27936691 | 0.52745936 | 0.68184633 | PET111 | XIII | 779629 | 782031 | -1 |
| YMR258C | 18.0929946 | 0.3124142 | 0.77466512 | 0.22346623 | 0.8942829 | 0.9350658 | ROY1 | XIII | 782620 | 784281 | -1 |
| YMR259C | 243.975792 | 1.66467459 | 0.51367948 | 10.8520785 | 0.00440049 | 0.02061395 | TRM732 | XIII | 784621 | 788883 | -1 |
| YMR260C | 68.4084866 | -2.4992188 | 0.84499427 | 8.17285352 | 0.01679915 | 0.05813116 | TIF11 | XIII | 789378 | 789839 | -1 |
| YMR261C | 27.5113042 | -0.774564 | 1.07082073 | 0.53625819 | 0.76480904 | 0.85692899 | TPS3 | XIII | 790205 | 793369 | -1 |
| YMR262W | 20.9902952 | 0.59561421 | 0.87487828 | 0.53609349 | 0.76487203 | 0.85692899 | NA | XIII | 793726 | 794667 | 1 |
| YMR263W | 55.8552871 | -2.3988848 | 0.58161153 | 16.0030104 | 0.00033496 | 0.00267504 | SAP30 | XIII | 794919 | 795524 | 1 |
| YMR264W | 185.539888 | -2.2164409 | 0.59235239 | 12.8650846 | 0.00160836 | 0.00961569 | CUE1 | XIII | 795805 | 796416 | 1 |

| | | | | | | | | | | | |
|---|---|---|---|---|---|---|---|---|---|---|---|
| YMR265C | 79.7643207 | -1.2505405 | 0.59711003 | 4.48897671 | 0.10598175 | 0.22838155 | NA | XIII | 796540 | 797925 | -1 |
| YMR266W | 46.0305882 | -0.5196404 | 0.58316022 | 0.80175799 | 0.6697311 | 0.79510827 | RSN1 | XIII | 798518 | 801379 | 1 |
| YMR267W | 60.5667982 | -0.6466441 | 0.59697009 | 1.17556657 | 0.55555743 | 0.70703257 | PPA2 | XIII | 801772 | 802704 | 1 |
| YMR268C | 20.5685782 | -1.845896 | 0.84398379 | 5.95977042 | 0.05079866 | 0.13219068 | PRP24 | XIII | 802888 | 804222 | -1 |
| YMR269W | 46.7332342 | -1.0356717 | 0.59662461 | 2.98218304 | 0.22512679 | 0.39049854 | TMA23 | XIII | 804456 | 805091 | 1 |
| YMR270C | 8.39799049 | 0.21976359 | 1.09853786 | 0.16679576 | 0.91998503 | 0.95001129 | RRN9 | XIII | 805326 | 806423 | -1 |
| YMR271C | 47.632561 | -1.6539063 | 0.60146565 | 8.72677008 | 0.01273521 | 0.0472672 | URA10 | XIII | 806865 | 807548 | -1 |
| YMR272C | 9.94260701 | 0.15880736 | 0.99331244 | 0.09644403 | 0.9529222 | 0.96731849 | SCS7 | XIII | 809623 | 810777 | -1 |
| YMR272W-B | 164.572769 | -0.3899991 | 0.70165362 | 0.60835152 | 0.73773119 | 0.84034303 | NA | XIII | 811089 | 811196 | 1 |
| YMR273C | 130.163054 | 1.14519622 | 0.72460081 | 3.86973302 | 0.14444355 | 0.28495484 | ZDS1 | XIII | 811233 | 813980 | -1 |
| YMR274C | 16.6674444 | 0.08326861 | 0.94263856 | 0.10582163 | 0.9484646 | 0.96471939 | RCE1 | XIII | 814364 | 815311 | -1 |
| YMR275C | 71.9229385 | -0.4085912 | 0.54136486 | 0.98450411 | 0.61124828 | 0.75489714 | BUL1 | XIII | 815651 | 818581 | -1 |
| YMR276W | 84.895957 | -0.9670236 | 0.52079109 | 3.63405919 | 0.16250775 | 0.31104789 | DSK2 | XIII | 818827 | 819948 | 1 |
| YMR277W | 76.9720482 | -0.2002901 | 0.52854122 | 1.77860477 | 0.41094233 | 0.58260433 | FCP1 | XIII | 820256 | 822454 | 1 |
| YMR278W | 9.33355823 | 0.24149513 | 1.00124216 | 0.28667924 | 0.86645975 | 0.92150571 | PRM15 | XIII | 822763 | 824631 | 1 |
| YMR279C | 2973.57562 | 0.05932238 | 0.60128131 | 0.13834849 | 0.93316407 | 0.95735444 | NA | XIII | 824729 | 826351 | -1 |
| YMR280C | 1075.89824 | -1.7775583 | 0.50903444 | 11.5294861 | 0.0031362 | 0.01579009 | CAT8 | XIII | 827028 | 831329 | -1 |
| YMR281W | 2186.48818 | 6.03866163 | 0.47406185 | 117.664425 | 2.82E-26 | 2.72E-23 | GPI12 | XIII | 832339 | 833253 | 1 |
| YMR282C | 45.6882923 | 1.53080121 | 0.61761488 | 6.11725021 | 0.04695221 | 0.12464616 | AEP2 | XIII | 833356 | 835098 | -1 |
| YMR283C | 17.457102 | -0.9097185 | 0.79563674 | 1.85120892 | 0.3962918 | 0.56739108 | RIT1 | XIII | 835326 | 836867 | -1 |
| YMR284W | 43.7424105 | -0.8984125 | 0.88877768 | 7.86803276 | 0.01956493 | 0.06501327 | YKU70 | XIII | 838187 | 839995 | 1 |
| YMR285C | 35.939145 | 0.41364108 | 0.62865292 | 0.48039831 | 0.78647122 | 0.87021248 | NGL2 | XIII | 840144 | 841691 | -1 |
| YMR286W | 13.4450905 | -0.3263737 | 0.80328793 | 0.20255045 | 0.90368428 | 0.93937738 | MRPL33 | XIII | 841942 | 842202 | 1 |
| YMR287C | 19.8916551 | 0.36562649 | 0.70580966 | 1.91201309 | 0.38442501 | 0.5574307 | DSS1 | XIII | 842436 | 845345 | -1 |
| YMR288W | 64.6967216 | -0.1838592 | 0.54370125 | 0.25178509 | 0.88170959 | 0.9300006 | HSH155 | XIII | 845571 | 848486 | 1 |
| YMR289W | 116.255289 | 3.63478651 | 0.57346634 | 44.1622714 | 2.57E-10 | 1.09E-08 | ABZ2 | XIII | 848685 | 849809 | 1 |
| YMR290C | 189.258027 | 0.2702188 | 0.48597227 | 0.42848721 | 0.80715174 | 0.88427787 | HAS1 | XIII | 850074 | 851591 | -1 |
| YMR291W | 38.8690368 | -1.7729512 | 0.65494862 | 9.01033048 | 0.01105176 | 0.04244332 | TDA1 | XIII | 852630 | 854390 | 1 |
| YMR292W | 249.383657 | 0.19720844 | 0.63277986 | 0.13707508 | 0.93375841 | 0.95735444 | GOT1 | XIII | 854795 | 855293 | 1 |
| YMR293C | 14.1865154 | -0.8202154 | 0.86139786 | 1.09576028 | 0.57817416 | 0.727271 | HER2 | XIII | 855399 | 856793 | -1 |
| YMR294W | 109.184096 | 1.11278768 | 1.05946489 | 10.7120277 | 0.00471968 | 0.02186157 | JNM1 | XIII | 856966 | 858087 | 1 |
| YMR295C | 136.859378 | 0.66142137 | 0.58081251 | 4.88314329 | 0.08702397 | 0.19878563 | NA | XIII | 858297 | 858890 | -1 |
| YMR296C | 253.459598 | 2.72074052 | 0.50508942 | 28.4700126 | 6.57E-07 | 1.23E-05 | LCB1 | XIII | 859215 | 860891 | -1 |

| | | | | | | | | | | | |
|---|---|---|---|---|---|---|---|---|---|---|---|
| YMR297W | 82.6784122 | 0.65884011 | 0.61478455 | 1.16533406 | 0.55840709 | 0.70949683 | PRC1 | XIII | 861922 | 863520 | 1 |
| YMR298W | 62.3430126 | -0.6278022 | 0.56606983 | 1.2801731 | 0.52724679 | 0.68184633 | LIP1 | XIII | 863819 | 864271 | 1 |
| YMR299C | 94.1038902 | 0.17044948 | 0.53658646 | 0.43412244 | 0.80488069 | 0.88279205 | DYN3 | XIII | 864412 | 865350 | -1 |
| YMR300C | 82.0272278 | 2.96819664 | 0.6160959 | 29.3881051 | 4.15E-07 | 8.21E-06 | ADE4 | XIII | 865559 | 867091 | -1 |
| YMR301C | 40.9017031 | 0.30985927 | 0.62707082 | 3.80344215 | 0.14931142 | 0.29203007 | ATM1 | XIII | 867555 | 869627 | -1 |
| YMR302C | 45.7471963 | 0.74981917 | 0.78917995 | 0.95139521 | 0.62145138 | 0.76184702 | YME2 | XIII | 870073 | 872625 | -1 |
| YMR303C | 589.12651 | -1.6905791 | 0.54531848 | 14.3897347 | 0.00075043 | 0.00517876 | ADH2 | XIII | 873291 | 874337 | -1 |
| YMR304W | 26.9568287 | -0.2072474 | 0.81520231 | 0.25634102 | 0.87970337 | 0.92911891 | UBP15 | XIII | 874987 | 878679 | 1 |
| YMR305C | 89.9559844 | 0.20484631 | 0.60499275 | 5.16232255 | 0.07568606 | 0.17861909 | SCW10 | XIII | 879063 | 880232 | -1 |
| YMR306W | 957.648805 | -1.0227961 | 0.53733233 | 3.5509367 | 0.16940409 | 0.31949501 | FKS3 | XIII | 881159 | 886516 | 1 |
| YMR307W | 84.8820185 | -1.4154501 | 0.72824958 | 3.95580616 | 0.13835906 | 0.27652708 | GAS1 | XIII | 887003 | 888682 | 1 |
| YMR308C | 32.0667848 | -1.335519 | 0.67568115 | 4.53055415 | 0.10380127 | 0.22509714 | PSE1 | XIII | 888952 | 892221 | -1 |
| YMR309C | 29.015788 | -2.246849 | 0.95258955 | 5.83826413 | 0.05398052 | 0.13854043 | NIP1 | XIII | 892988 | 895426 | -1 |
| YMR310C | 20.7709717 | -2.9329843 | 0.7703597 | 14.1577015 | 0.00084274 | 0.00572036 | NA | XIII | 895716 | 896669 | -1 |
| YMR311C | 112.785042 | 0.55855383 | 0.58871213 | 2.48822819 | 0.28819611 | 0.46197549 | GLC8 | XIII | 896914 | 897603 | -1 |
| YMR312W | 116.099661 | -0.3243752 | 0.70498167 | 8.49133962 | 0.01432613 | 0.05171342 | ELP6 | XIII | 898405 | 899226 | 1 |
| YMR313C | 42.9923358 | -1.2941216 | 0.60018618 | 5.89575703 | 0.05245086 | 0.13539478 | TGL3 | XIII | 899379 | 901307 | -1 |
| YMR314W | 51.4940978 | 1.02751269 | 0.72834492 | 4.12895091 | 0.12688483 | 0.2604265 | PRE5 | XIII | 901709 | 902413 | 1 |
| YMR315W | 3358.23331 | 4.44860744 | 0.46828363 | 73.9485637 | 8.76E-17 | 1.17E-14 | NA | XIII | 902800 | 903849 | 1 |
| YMR315W-A | 68.6713206 | 0.25986539 | 0.56587674 | 0.96542466 | 0.61710732 | 0.75846612 | NA | XIII | 904286 | 904393 | 1 |
| YMR316C-A | 7202.28902 | 3.28666933 | 0.56444355 | 45.0256027 | 1.67E-10 | 7.44E-09 | NA | XIII | 905662 | 905973 | -1 |
| YMR316W | 121.020751 | -1.0466283 | 0.53308766 | 4.2528651 | 0.119262 | 0.24840182 | DIA1 | XIII | 904825 | 905835 | 1 |
| YMR317W | 3148.13071 | 0.16179768 | 0.47005179 | 1.01956396 | 0.60062651 | 0.74682758 | NA | XIII | 907364 | 910786 | 1 |
| YMR318C | 200.206853 | 0.68207119 | 0.55475963 | 1.53240165 | 0.46477548 | 0.62728439 | ADH6 | XIII | 911061 | 912143 | -1 |
| YMR319C | 1223.15629 | 1.19303173 | 0.54847958 | 7.77854745 | 0.0204602 | 0.06719418 | FET4 | XIII | 912880 | 914538 | -1 |
| YNCA0001W | 790.335017 | -0.7898134 | 0.5638457 | 2.32191797 | 0.3131857 | 0.48516903 | NA | NA | NA | NA | NA |
| YNCA0003W | 10.0289279 | 3.4422853 | 1.19214841 | 12.7762643 | 0.00168139 | 0.00981378 | NA | NA | NA | NA | NA |
| YNCB0003W | 39.4998415 | -0.4804199 | 0.69512405 | 6.55496383 | 0.03772313 | 0.10638914 | NA | NA | NA | NA | NA |
| YNCB0006W | 5.1851498 | 3.77646147 | 1.47424515 | 7.90514899 | 0.01920519 | 0.06416508 | NA | NA | NA | NA | NA |
| YNCB0008W | 63.4237925 | 0.02500507 | 0.57262016 | 2.27159582 | 0.32116576 | 0.49377316 | NA | NA | NA | NA | NA |
| YNCB0009C | 33.4896672 | 0.22248757 | 0.61693908 | 0.16358129 | 0.92146485 | 0.95035295 | NA | NA | NA | NA | NA |
| YNCB0010W | 1454.72461 | 2.07332609 | 0.89354476 | 7.85973526 | 0.01964627 | 0.06511272 | NA | NA | NA | NA | NA |
| YNCB0014W | 10538.2983 | -3.1999246 | 0.49762974 | 35.8151894 | 1.67E-08 | 4.74E-07 | NA | NA | NA | NA | NA |

| | | | | | | | | | | | |
|---|---|---|---|---|---|---|---|---|---|---|---|
| YNCB0016W | 15.3023853 | 4.16854789 | 1.11363282 | 17.3431793 | 0.00017139 | 0.00148775 | NA | NA | NA | NA | NA |
| YNCB0017C | 81.9735576 | 2.94001131 | 0.66591671 | 22.48513 | 1.31E-05 | 0.00016676 | NA | NA | NA | NA | NA |
| YNCB0019C | 308.399574 | 1.10437527 | 0.55656108 | 6.4987871 | 0.03879773 | 0.10910095 | NA | NA | NA | NA | NA |
| YNCC0002W | 27.8134993 | -0.3116856 | 0.65964304 | 0.86102168 | 0.65017687 | 0.78140989 | NA | NA | NA | NA | NA |
| YNCC0003C | 2675.13139 | -1.1132859 | 0.88409309 | 7.67922124 | 0.02150197 | 0.06990254 | NA | NA | NA | NA | NA |
| YNCC0011W | 60.0694742 | -0.4336773 | 0.60720314 | 0.59676395 | 0.74201785 | 0.84243165 | NA | NA | NA | NA | NA |
| YNCC0013W | 11.0760926 | -1.8202176 | 0.97321282 | 4.74396846 | 0.09329542 | 0.20840297 | NA | NA | NA | NA | NA |
| YNCD0002C | 28.3098886 | 1.86340587 | 0.73163241 | 7.27962801 | 0.02625723 | 0.08048139 | NA | NA | NA | NA | NA |
| YNCD0005C | 6.50321707 | -1.4666974 | 1.12192519 | 2.67849222 | 0.26204315 | 0.43337041 | NA | NA | NA | NA | NA |
| YNCD0008W | 9.86040035 | 5.28933352 | 1.50578748 | 13.5878559 | 0.00112056 | 0.00722498 | NA | NA | NA | NA | NA |
| YNCD0009W | 10.6730292 | 3.78953224 | 1.1829239 | 12.948177 | 0.0015429 | 0.00934751 | NA | NA | NA | NA | NA |
| YNCD0015C | 9.16206783 | 4.91733345 | 1.37272342 | 26.3314364 | 1.92E-06 | 3.15E-05 | NA | NA | NA | NA | NA |
| YNCD0016C | 11.6028954 | 2.29529337 | 0.98588474 | 5.26830682 | 0.07177971 | 0.17202174 | NA | NA | NA | NA | NA |
| YNCD0017W | 101.952971 | 5.70240019 | 0.76454741 | 55.6056023 | 8.42E-13 | 5.95E-11 | NA | NA | NA | NA | NA |
| YNCD0030W | 19.5957293 | -0.075452 | 0.80479256 | 1.98159249 | 0.37128094 | 0.54464572 | NA | NA | NA | NA | NA |
| YNCD0032C | 279.60473 | -0.9070844 | 0.54818433 | 2.6919902 | 0.26028057 | 0.43181218 | NA | NA | NA | NA | NA |
| YNCE0003W | 13.6869103 | -3.1816537 | 0.93777283 | 11.1208852 | 0.00384707 | 0.01862421 | NA | NA | NA | NA | NA |
| YNCE0004C | 15.2555353 | -4.0169255 | 1.08534678 | 20.1625627 | 4.19E-05 | 0.00044467 | NA | NA | NA | NA | NA |
| YNCE0007C | 95.2876892 | 4.15451073 | 0.65193859 | 38.7697246 | 3.81E-09 | 1.26E-07 | NA | NA | NA | NA | NA |
| YNCE0011C | 94.2778616 | -0.3556864 | 0.54901419 | 0.88418798 | 0.64268923 | 0.77621415 | NA | NA | NA | NA | NA |
| YNCE0012W | 48.4629026 | 1.02258923 | 0.67856727 | 5.5145476 | 0.06346455 | 0.15629934 | NA | NA | NA | NA | NA |
| YNCE0013W | 13.2278782 | -1.5630875 | 1.07293336 | 3.30506668 | 0.191564 | 0.34813419 | NA | NA | NA | NA | NA |
| YNCE0014W | 9.31911418 | 3.43632338 | 1.27389805 | 8.8372941 | 0.01205052 | 0.04533628 | NA | NA | NA | NA | NA |
| YNCE0017W | 6.85800097 | 0.16924912 | 1.08971372 | 0.14003858 | 0.93237584 | 0.95731799 | NA | NA | NA | NA | NA |
| YNCE0018W | 17.9646158 | 4.52717135 | 1.04189247 | 23.2295086 | 9.03E-06 | 0.00012049 | NA | NA | NA | NA | NA |
| YNCE0019W | 831.715072 | 3.30121883 | 0.58494249 | 26.8880701 | 1.45E-06 | 2.45E-05 | NA | NA | NA | NA | NA |
| YNCE0020C | 23.2007434 | 2.06907377 | 0.75774453 | 8.24091578 | 0.01623708 | 0.05677094 | NA | NA | NA | NA | NA |
| YNCF0001C | 6782.19328 | -0.1983807 | 0.95720414 | 5.05997144 | 0.07966016 | 0.18583091 | NA | NA | NA | NA | NA |
| YNCF0003C | 1193.16544 | -0.2488482 | 0.59093528 | 5.67034957 | 0.05870826 | 0.14739097 | NA | NA | NA | NA | NA |
| YNCF0010C | 7.29313492 | 2.08773168 | 1.13341002 | 10.2276119 | 0.00601315 | 0.02655695 | NA | NA | NA | NA | NA |
| YNCF0013W | 9.59737856 | -0.3561954 | 0.9185318 | 4.48992906 | 0.1059313 | 0.22838155 | NA | NA | NA | NA | NA |
| YNCG0001C | 137.098384 | 0.59077352 | 0.56426335 | 1.55189179 | 0.46026821 | 0.62448065 | NA | NA | NA | NA | NA |
| YNCG0006C | 14.9671049 | 0.16548754 | 0.8794592 | 0.11768355 | 0.94285594 | 0.96176085 | NA | NA | NA | NA | NA |

| | | | | | | | | | | | |
|---|---|---|---|---|---|---|---|---|---|---|---|
| YNCG0008W | 21.6187333 | 2.66329614 | 0.88365666 | 12.404239 | 0.00202513 | 0.01126371 | NA | NA | NA | NA | NA |
| YNCG0010W | 39.4910692 | 3.94013695 | 1.00463946 | 14.173318 | 0.00083619 | 0.00568253 | NA | NA | NA | NA | NA |
| YNCG0015C | 25.5048131 | 0.67505655 | 0.74139113 | 0.89069905 | 0.64060033 | 0.77561669 | NA | NA | NA | NA | NA |
| YNCG0017W | 23.2246281 | 0.78952166 | 0.76407428 | 1.5687622 | 0.45640208 | 0.62207609 | NA | NA | NA | NA | NA |
| YNCG0018C | 7.07812754 | -0.163242 | 1.03899167 | 0.14385902 | 0.93059649 | 0.95636381 | NA | NA | NA | NA | NA |
| YNCG0024W | 9.72516321 | -3.5464523 | 1.04621322 | 11.5323541 | 0.00313171 | 0.01578119 | NA | NA | NA | NA | NA |
| YNCG0025C | 7.89994494 | 2.70936155 | 1.09221161 | 8.54605347 | 0.01393953 | 0.05066533 | NA | NA | NA | NA | NA |
| YNCG0026W | 45.9295051 | 0.46373109 | 0.77630543 | 2.63102091 | 0.26833731 | 0.44113375 | NA | NA | NA | NA | NA |
| YNCG0037W | 42.4356806 | -1.1890156 | 0.70011889 | 3.72032187 | 0.15564758 | 0.30083183 | NA | NA | NA | NA | NA |
| YNCG0038C | 8.39934761 | 3.37836763 | 1.31213169 | 7.73095988 | 0.02095286 | 0.06842475 | NA | NA | NA | NA | NA |
| YNCH0003C | 11.2755452 | -3.4608434 | 0.99715052 | 12.419671 | 0.00200957 | 0.01119865 | NA | NA | NA | NA | NA |
| YNCH0008W | 28.8518657 | 1.2146895 | 0.86978235 | 4.43930891 | 0.10864664 | 0.23264204 | NA | NA | NA | NA | NA |
| YNCH0009C | 10.9534641 | -0.3033981 | 0.95716413 | 2.68664398 | 0.26097726 | 0.43235173 | NA | NA | NA | NA | NA |
| YNCH0010C | 10.024353 | -0.0308202 | 0.97986306 | 0.53563979 | 0.76504556 | 0.85695759 | NA | NA | NA | NA | NA |
| YNCH0011W | 587.822998 | 3.13080225 | 0.47985271 | 37.2830838 | 8.02E-09 | 2.48E-07 | NA | NA | NA | NA | NA |
| YNCH0012W | 34.5506099 | 0.67919982 | 0.79146719 | 0.74418649 | 0.68928997 | 0.81068229 | NA | NA | NA | NA | NA |
| YNCH0014W | 111.011961 | 0.3373075 | 0.52848332 | 0.45521755 | 0.79643579 | 0.87685172 | NA | NA | NA | NA | NA |
| YNCI0001W | 49.6082987 | 0.45591936 | 0.64226936 | 0.68154361 | 0.71122119 | 0.82438173 | NA | NA | NA | NA | NA |
| YNCI0003C | 5.97890307 | 6.39583385 | 1.80276124 | 24.0921757 | 5.87E-06 | 8.19E-05 | NA | NA | NA | NA | NA |
| YNCI0006W | 50.0742922 | -0.7463261 | 0.62342964 | 2.40270227 | 0.30078753 | 0.47389383 | NA | NA | NA | NA | NA |
| YNCI0012 | 18.7474564 | -0.3803076 | 0.74622815 | 0.68742949 | 0.70913118 | 0.82408359 | NA | NA | NA | NA | NA |
| YNCI0013W | 332.000789 | -0.9733598 | 0.57560928 | 2.79855419 | 0.24677529 | 0.41766412 | NA | NA | NA | NA | NA |
| YNCJ0004C | 10.4359128 | -0.4338758 | 1.08130841 | 4.12323797 | 0.12724779 | 0.26090922 | NA | NA | NA | NA | NA |
| YNCJ0006W | 6.73079234 | 1.50767747 | 1.12265538 | 2.99457915 | 0.22373576 | 0.38936881 | NA | NA | NA | NA | NA |
| YNCJ0007C | 27.55903 | 0.69411924 | 0.72576319 | 1.84760605 | 0.39700634 | 0.56785755 | NA | NA | NA | NA | NA |
| YNCJ0009C | 17.7194969 | 3.35810993 | 0.92289699 | 13.7479762 | 0.00103434 | 0.00679008 | NA | NA | NA | NA | NA |
| YNCJ0028C | 74.1698523 | 0.51503812 | 0.59022235 | 1.71289754 | 0.4246675 | 0.59492495 | NA | NA | NA | NA | NA |
| YNCJ0030W | 400.94239 | -0.1090742 | 0.47423621 | 0.94450726 | 0.62359533 | 0.76318262 | NA | NA | NA | NA | NA |
| YNCK0004W | 6.04427312 | 2.78909439 | 1.27639613 | 9.733855 | 0.00769698 | 0.03215404 | NA | NA | NA | NA | NA |
| YNCK0005C | 20.9076363 | 7.20402113 | 1.64995284 | 26.3837513 | 1.87E-06 | 3.09E-05 | NA | NA | NA | NA | NA |
| YNCK0010W | 35.6133643 | 8.82469562 | 1.59045587 | 60.1010224 | 8.90E-14 | 7.47E-12 | NA | NA | NA | NA | NA |
| YNCK0013W | 82.4758454 | 1.65453557 | 0.65247619 | 6.17336368 | 0.04565319 | 0.12191465 | NA | NA | NA | NA | NA |
| YNCK0015C | 134.642469 | -0.4395261 | 0.54762515 | 0.65360878 | 0.72122481 | 0.83085787 | NA | NA | NA | NA | NA |

| | | | | | | | | | | | |
|---|---|---|---|---|---|---|---|---|---|---|---|
| YNCK0017W | 7.78558252 | -1.7591811 | 1.00008775 | 3.42409234 | 0.18049609 | 0.33506648 | NA | NA | NA | NA | NA |
| YNCK0020C | 463.463479 | 3.44740143 | 0.59397704 | 42.4281448 | 6.12E-10 | 2.32E-08 | NA | NA | NA | NA | NA |
| YNCL0018W | 1392.084 | -3.0808258 | 0.63986149 | 29.5378476 | 3.85E-07 | 7.70E-06 | NA | NA | NA | NA | NA |
| YNCL0032C | 21.0259403 | 6.5864279 | 1.42883066 | 28.8698258 | 5.38E-07 | 1.04E-05 | NA | NA | NA | NA | NA |
| YNCL0036C | 16.7683428 | 0.56653398 | 0.99887892 | 0.40162603 | 0.81806538 | 0.89067292 | NA | NA | NA | NA | NA |
| YNCL0039W | 5.82013184 | 1.53526771 | 1.18202715 | 6.94821515 | 0.03098948 | 0.09168578 | NA | NA | NA | NA | NA |
| YNCL0040W | 14.8728956 | 0.31212495 | 0.84530956 | 0.7743864 | 0.67895991 | 0.8022812 | NA | NA | NA | NA | NA |
| YNCL0041C | 36.5714911 | 3.86592874 | 0.94712917 | 16.1881048 | 0.00030535 | 0.0024866 | NA | NA | NA | NA | NA |
| YNCL0044W | 8.36776444 | 3.26948628 | 1.13564064 | 16.2588412 | 0.00029474 | 0.0024162 | NA | NA | NA | NA | NA |
| YNCL0048W | 11.1850325 | -0.2983829 | 1.08932141 | 6.94321125 | 0.03106711 | 0.09182162 | NA | NA | NA | NA | NA |
| YNCM0001W | 1223.47622 | -0.4447113 | 0.51999395 | 0.84789762 | 0.65445738 | 0.78442961 | NA | NA | NA | NA | NA |
| YNCM0002C | 32.190886 | -0.9725709 | 0.66488414 | 4.05612524 | 0.13159021 | 0.26686772 | NA | NA | NA | NA | NA |
| YNCM0003W | 11.6359297 | 2.73960902 | 1.03873483 | 7.20945311 | 0.02719488 | 0.08256862 | NA | NA | NA | NA | NA |
| YNCM0004C | 50.4566467 | 5.025148 | 1.04215832 | 21.2235051 | 2.46E-05 | 0.00028516 | NA | NA | NA | NA | NA |
| YNCM0008C | 22.0944308 | -0.1954882 | 0.7037344 | 0.42442982 | 0.80879086 | 0.8854035 | NA | NA | NA | NA | NA |
| YNCM0011W | 10.5055136 | -1.2454159 | 0.94137375 | 3.90809521 | 0.14169937 | 0.28087618 | NA | NA | NA | NA | NA |
| YNCM0012W | 12.8602663 | 0.74938097 | 1.0092383 | 1.4140128 | 0.49311819 | 0.65425168 | NA | NA | NA | NA | NA |
| YNCM0019W | 24.9135411 | 1.25914763 | 0.74511922 | 2.82576531 | 0.24344052 | 0.41310685 | NA | NA | NA | NA | NA |
| YNCM0030W | 27.2305026 | -0.4247427 | 0.98879769 | 9.04398076 | 0.01086737 | 0.04192011 | NA | NA | NA | NA | NA |
| YNCM0031W | 260.606947 | 0.42076655 | 0.59551771 | 2.58324754 | 0.27482417 | 0.44836065 | NA | NA | NA | NA | NA |
| YNCM0032C | 80.8060757 | 0.68393006 | 0.56465517 | 3.88379059 | 0.14343185 | 0.28353376 | NA | NA | NA | NA | NA |
| YNCM0033C | 15.8194089 | 2.03423139 | 0.99007445 | 12.2111726 | 0.00223037 | 0.01215995 | NA | NA | NA | NA | NA |
| YNCM0034C | 124.942867 | 1.98981608 | 0.56780967 | 18.7538817 | 8.47E-05 | 0.00081555 | NA | NA | NA | NA | NA |
| YNCN0001W | 76.8726536 | 1.18250222 | 0.55956449 | 4.60859081 | 0.09982912 | 0.21918634 | NA | NA | NA | NA | NA |
| YNCN0002C | 14.0832359 | 2.20351979 | 0.97539064 | 4.98484958 | 0.08270917 | 0.19079127 | NA | NA | NA | NA | NA |
| YNCN0005C | 14.2686022 | -0.9310257 | 0.98213933 | 1.09306821 | 0.57895293 | 0.7279343 | NA | NA | NA | NA | NA |
| YNCN0008C | 26.7858364 | -0.2534247 | 0.83297526 | 0.20708138 | 0.90163933 | 0.93824416 | NA | NA | NA | NA | NA |
| YNCN0013W | 23.5620445 | 0.25345701 | 0.7064932 | 1.29609627 | 0.52306573 | 0.67871828 | NA | NA | NA | NA | NA |
| YNCN0018W | 14.9877506 | -1.7468776 | 0.82785256 | 4.3153395 | 0.11559417 | 0.242911 | NA | NA | NA | NA | NA |
| YNCN0019C | 15.8072341 | 5.4201338 | 1.25674825 | 30.5163193 | 2.36E-07 | 5.05E-06 | NA | NA | NA | NA | NA |
| YNCO0003C | 87.3822961 | -1.7011273 | 0.59574557 | 9.7825461 | 0.00751185 | 0.03164938 | NA | NA | NA | NA | NA |
| YNCO0007W | 10.639455 | 0.27900283 | 0.90110382 | 0.50578351 | 0.77655193 | 0.8639961 | NA | NA | NA | NA | NA |
| YNCO0014C | 727.000133 | 1.163708 | 0.61843176 | 3.93131864 | 0.14006351 | 0.27867477 | NA | NA | NA | NA | NA |

| | | | | | | | | | | | |
|---|---|---|---|---|---|---|---|---|---|---|---|
| YNCO0015C | 172.896618 | 0.54665033 | 0.61682435 | 12.8639649 | 0.00160926 | 0.00961569 | NA | NA | NA | NA | NA |
| YNCO0016W | 32.4083216 | 0.6509327 | 0.65611172 | 0.967192 | 0.61656224 | 0.75817303 | NA | NA | NA | NA | NA |
| YNCO0022C | 76.6236034 | -0.9990216 | 0.56590291 | 4.77128852 | 0.09202967 | 0.20645168 | NA | NA | NA | NA | NA |
| YNCO0024C | 34.3450664 | -1.2912154 | 0.63994597 | 6.27077213 | 0.04348296 | 0.1181447 | NA | NA | NA | NA | NA |
| YNCO0025W | 126.379561 | 3.83425452 | 0.56840372 | 43.2647016 | 4.03E-10 | 1.62E-08 | NA | NA | NA | NA | NA |
| YNCO0026W | 24.7348683 | -0.4130717 | 0.73170813 | 2.44801694 | 0.29404911 | 0.46854198 | NA | NA | NA | NA | NA |
| YNCO0027C | 123.198071 | -1.6050479 | 0.53306979 | 8.90262769 | 0.01166323 | 0.04427675 | NA | NA | NA | NA | NA |
| YNCO0028W | 16.2343628 | -3.27572 | 0.84468328 | 16.4191919 | 0.00027203 | 0.00224367 | NA | NA | NA | NA | NA |
| YNCO0029C | 350.558512 | 4.5065056 | 0.95236666 | 20.8070685 | 3.03E-05 | 0.00034284 | NA | NA | NA | NA | NA |
| YNCP0001C | 30.0954422 | -0.6029027 | 0.74035997 | 3.20985334 | 0.20090429 | 0.36006531 | NA | NA | NA | NA | NA |
| YNCP0002W | 1571.57567 | 1.64895951 | 0.52738819 | 9.29628236 | 0.00957939 | 0.03819881 | NA | NA | NA | NA | NA |
| YNCP0005C | 31.2905589 | -2.2527005 | 0.79155814 | 7.60556033 | 0.02230866 | 0.07187933 | NA | NA | NA | NA | NA |
| YNCP0007C | 10.3773681 | 0.68525543 | 0.99194629 | 1.70517083 | 0.42631131 | 0.59587092 | NA | NA | NA | NA | NA |
| YNCP0013C | 10.4571387 | 1.82423298 | 0.95092649 | 5.6101777 | 0.0605014 | 0.15079771 | NA | NA | NA | NA | NA |
| YNCP0014C | 26.3667168 | 1.39875176 | 0.74095087 | 3.62566143 | 0.16319153 | 0.31184125 | NA | NA | NA | NA | NA |
| YNCP0015C | 22.1066181 | 0.06594733 | 0.71606736 | 1.19674074 | 0.54970673 | 0.70152126 | NA | NA | NA | NA | NA |
| YNCP0018C | 10.9465899 | -2.7661212 | 1.01039346 | 9.43599573 | 0.00893305 | 0.03616947 | NA | NA | NA | NA | NA |
| YNCP0021W | 30.116723 | -0.0336663 | 0.69429457 | 0.36732797 | 0.83221539 | 0.90069191 | NA | NA | NA | NA | NA |
| YNL001W | 7.25780802 | -0.1634545 | 1.03037434 | 0.37966906 | 0.82709598 | 0.89645933 | DOM34 | XIV | 627456 | 628616 | 1 |
| YNL002C | 45.455493 | -0.6535254 | 0.63896221 | 1.11485712 | 0.57267978 | 0.72277589 | RLP7 | XIV | 626174 | 627142 | -1 |
| YNL003C | 111.762781 | -0.2307294 | 0.54766164 | 1.77064369 | 0.41258136 | 0.58426143 | PET8 | XIV | 624975 | 625829 | -1 |
| YNL004W | 890.717428 | -0.0160485 | 0.66218315 | 8.6709681 | 0.01309553 | 0.04820096 | HRB1 | XIV | 622915 | 624621 | 1 |
| YNL005C | 136.723965 | -0.2880444 | 0.56477045 | 0.83735655 | 0.65791583 | 0.78607772 | MRP7 | XIV | 621313 | 622428 | -1 |
| YNL006W | 173.062834 | -0.9910388 | 0.55272638 | 4.61609088 | 0.09945545 | 0.21864382 | LST8 | XIV | 620067 | 620978 | 1 |
| YNL007C | 77.5664949 | -0.4911294 | 0.66246923 | 0.61026608 | 0.73702531 | 0.83986943 | SIS1 | XIV | 618507 | 619565 | -1 |
| YNL008C | 683.128962 | -0.6549809 | 0.46837041 | 2.05971426 | 0.35705797 | 0.53036574 | ASI3 | XIV | 616211 | 618241 | -1 |
| YNL009W | 126.382162 | 0.13447009 | 0.5140943 | 0.76158009 | 0.68332134 | 0.80595449 | IDP3 | XIV | 614821 | 616083 | 1 |
| YNL010W | 10.9832782 | -0.4880086 | 0.97516844 | 0.32008919 | 0.85210579 | 0.91195795 | NA | XIV | 613635 | 614360 | 1 |
| YNL011C | 51.2603742 | -0.0972795 | 0.61323062 | 0.55338021 | 0.75828945 | 0.85403579 | NA | XIV | 611836 | 613170 | -1 |
| YNL012W | 1392.68163 | 1.16753656 | 0.46926121 | 7.47514509 | 0.02381184 | 0.07531616 | SPO1 | XIV | 609686 | 611665 | 1 |
| YNL014W | 66.9920647 | 0.82990168 | 0.57625489 | 2.10389503 | 0.3492569 | 0.52293528 | HEF3 | XIV | 606319 | 609453 | 1 |
| YNL015W | 28.5598836 | 1.23841859 | 0.68382309 | 3.56750512 | 0.16800651 | 0.31779081 | PBI2 | XIV | 605384 | 605611 | 1 |
| YNL016W | 158.658512 | -0.7690619 | 0.61526937 | 10.3765967 | 0.0055815 | 0.02501305 | PUB1 | XIV | 602907 | 604268 | 1 |

| | | | | | | | | | | | |
|---|---|---|---|---|---|---|---|---|---|---|---|
| YNL018C | 137.786783 | 1.07021557 | 0.5576878 | 4.49430725 | 0.10569966 | 0.22827341 | NA | XIV | 599936 | 601774 | -1 |
| YNL019C | 276.146147 | 2.31842624 | 0.53269804 | 17.3587795 | 0.00017005 | 0.0014784 | NA | XIV | 598376 | 599230 | -1 |
| YNL020C | 1420.64948 | -1.967599 | 0.53525258 | 13.4946075 | 0.00117404 | 0.00750302 | ARK1 | XIV | 595623 | 597539 | -1 |
| YNL021W | 36.4262932 | -0.2833269 | 0.61547738 | 1.57264801 | 0.4555162 | 0.62135279 | HDA1 | XIV | 593227 | 595347 | 1 |
| YNL022C | 9.53352065 | -0.280556 | 0.98199637 | 0.09991861 | 0.95126814 | 0.96614462 | RCM1 | XIV | 591427 | 592899 | -1 |
| YNL023C | 61.9186462 | -0.8618042 | 0.7043356 | 1.48944503 | 0.47486605 | 0.6373376 | FAP1 | XIV | 588263 | 591160 | -1 |
| YNL024C | 184.694133 | -1.7358772 | 0.60963902 | 7.69873909 | 0.02129316 | 0.06934048 | EFM6 | XIV | 587107 | 587847 | -1 |
| YNL024C-A | 14.3848552 | -1.5394744 | 0.85391433 | 3.89668496 | 0.14251009 | 0.28204189 | KSH1 | XIV | 586602 | 586820 | -1 |
| YNL025C | 8.12756857 | 0.64226431 | 1.01906565 | 0.41543972 | 0.8124346 | 0.88731158 | SSN8 | XIV | 584320 | 585291 | -1 |
| YNL026W | 91.4750315 | -0.1504125 | 0.55739058 | 1.3013873 | 0.52168378 | 0.67740505 | SAM50 | XIV | 581920 | 583374 | 1 |
| YNL027W | 15.7901544 | 0.92166144 | 0.82995563 | 1.33001602 | 0.51426941 | 0.67150244 | CRZ1 | XIV | 579580 | 581616 | 1 |
| YNL029C | 586.255355 | -0.9389672 | 1.80946857 | 3.86592186 | 0.14471906 | 0.28539624 | KTR5 | XIV | 577205 | 578773 | -1 |
| YNL030W | 132.707423 | -2.4427785 | 0.88474339 | 7.30360603 | 0.02594431 | 0.07981804 | HHF2 | XIV | 576727 | 577038 | 1 |
| YNL031C | 278.397464 | -1.9902417 | 0.49770896 | 15.3335618 | 0.00046812 | 0.00348384 | HHT2 | XIV | 575640 | 576050 | -1 |
| YNL032W | 28.5681931 | -1.3803567 | 0.75139863 | 6.9504792 | 0.03095442 | 0.09162887 | SIW14 | XIV | 574506 | 575351 | 1 |
| YNL033W | 92.8457098 | 0.82302679 | 0.57948861 | 3.69072033 | 0.15796842 | 0.30416931 | NA | XIV | 572999 | 573853 | 1 |
| YNL034W | 77.1518942 | 0.18254649 | 0.5508607 | 0.14256612 | 0.93119827 | 0.95681242 | NA | XIV | 570477 | 572315 | 1 |
| YNL035C | 48.1012349 | -0.1067302 | 0.5908696 | 2.25072135 | 0.32453539 | 0.49763242 | NA | XIV | 568522 | 569691 | -1 |
| YNL036W | 50.607255 | 2.56596728 | 0.75217516 | 12.8940185 | 0.00158526 | 0.00951154 | NCE103 | XIV | 559814 | 560479 | 1 |
| YNL037C | 67.2585874 | 2.19039 | 0.58392242 | 15.5975706 | 0.00041023 | 0.00313357 | IDH1 | XIV | 557920 | 559002 | -1 |
| YNL038W | 44.6053403 | -0.0203625 | 0.96938485 | 15.2031916 | 0.00049965 | 0.00367131 | GPI15 | XIV | 557020 | 557783 | 1 |
| YNL039W | 51.8690704 | -0.5295386 | 0.56676239 | 0.89094646 | 0.64052109 | 0.77561669 | BDP1 | XIV | 555048 | 556832 | 1 |
| YNL040W | 13.6442109 | 0.66333064 | 0.86450904 | 0.74574462 | 0.68875318 | 0.81049746 | NA | XIV | 553380 | 554750 | 1 |
| YNL041C | 17.2744245 | 0.703469 | 0.75864827 | 0.88663657 | 0.64190287 | 0.77608641 | COG6 | XIV | 549468 | 551987 | -1 |
| YNL042W | 125.767646 | 3.11721595 | 0.53972567 | 32.2712568 | 9.83E-08 | 2.40E-06 | BOP3 | XIV | 548100 | 549290 | 1 |
| YNL042W-B | 13.6064407 | 0.48841062 | 1.03595219 | 0.34814523 | 0.84023588 | 0.90645906 | NA | XIV | 547113 | 547370 | 1 |
| YNL044W | 77.7611086 | 0.20055185 | 0.61786849 | 2.88641632 | 0.23616887 | 0.40406514 | YIP3 | XIV | 545268 | 545877 | 1 |
| YNL045W | 715.326555 | 4.33025631 | 0.51824601 | 55.8888897 | 7.31E-13 | 5.29E-11 | LAP2 | XIV | 542963 | 544978 | 1 |
| YNL046W | 285.830051 | 0.9088578 | 0.49205526 | 3.54574225 | 0.16984464 | 0.32001317 | NA | XIV | 542304 | 542822 | 1 |
| YNL047C | 175.311761 | 0.68177371 | 0.72713246 | 4.45839767 | 0.10761461 | 0.23077356 | SLM2 | XIV | 539911 | 541881 | -1 |
| YNL048W | 33.54787 | 0.90622367 | 0.6598626 | 2.47501114 | 0.29010697 | 0.46388272 | ALG11 | XIV | 538173 | 539819 | 1 |
| YNL049C | 69.9526782 | -1.6107506 | 0.64352632 | 7.1193895 | 0.02844751 | 0.08543105 | SFB2 | XIV | 535281 | 537911 | -1 |
| YNL050C | 18.4094928 | -1.8318709 | 0.80937396 | 7.09599358 | 0.02878224 | 0.08625733 | NA | XIV | 534079 | 534982 | -1 |

| | | | | | | | | | | | |
|---|---|---|---|---|---|---|---|---|---|---|---|
| YNL051W | 26.1701317 | 1.2736141 | 0.70574832 | 6.43955116 | 0.03996403 | 0.11135309 | COG5 | XIV | 532659 | 533870 | 1 |
| YNL052W | 41.9136576 | -0.3109974 | 0.64167608 | 0.38349151 | 0.82551672 | 0.8954181 | COX5A | XIV | 531725 | 532186 | 1 |
| YNL053W | 615.515419 | -3.9524542 | 0.51580501 | 59.9851213 | 9.43E-14 | 7.80E-12 | MSG5 | XIV | 529942 | 531411 | 1 |
| YNL054W | 102.289492 | 1.19398432 | 0.53581404 | 4.83563205 | 0.08911603 | 0.2024252 | VAC7 | XIV | 526086 | 529583 | 1 |
| YNL054W-B | 18.947988 | -0.2463253 | 0.7801875 | 0.16378276 | 0.92137203 | 0.95035295 | NA | XIV | 519456 | 524706 | 1 |
| YNL055C | 261.220122 | 0.04330213 | 0.56918838 | 9.52578014 | 0.00854089 | 0.03489891 | POR1 | XIV | 517994 | 518845 | -1 |
| YNL056W | 36.6365605 | -0.4700247 | 0.61412443 | 1.56463602 | 0.45734465 | 0.62262533 | OCA2 | XIV | 517249 | 517842 | 1 |
| YNL058C | 355.288961 | -0.0642155 | 0.48033257 | 0.59907503 | 0.74116092 | 0.84242342 | NA | XIV | 515763 | 516713 | -1 |
| YNL059C | 98.7137308 | 1.3631604 | 0.54108977 | 7.06131431 | 0.02928566 | 0.08719023 | ARP5 | XIV | 512668 | 514935 | -1 |
| YNL061W | 35.3224471 | -2.4610245 | 0.68091282 | 12.5413696 | 0.00189093 | 0.01073788 | NOP2 | XIV | 510540 | 512396 | 1 |
| YNL062C | 65.9047539 | 0.49946408 | 0.63625404 | 1.63050581 | 0.4425274 | 0.61121985 | GCD10 | XIV | 508776 | 510212 | -1 |
| YNL063W | 270.248997 | -1.4593345 | 0.46735364 | 9.42689763 | 0.00897378 | 0.03630899 | MTQ1 | XIV | 507758 | 508702 | 1 |
| YNL064C | 103.463994 | -0.6309331 | 0.73713772 | 0.74075724 | 0.69047285 | 0.81157893 | YDJ1 | XIV | 505868 | 507097 | -1 |
| YNL065W | 49.0477664 | -0.625623 | 0.57480548 | 1.61764083 | 0.44538312 | 0.61297083 | AQR1 | XIV | 503724 | 505484 | 1 |
| YNL066W | 212.457049 | 1.11988432 | 0.49439568 | 5.00540776 | 0.08186335 | 0.18936828 | SUN4 | XIV | 501515 | 502777 | 1 |
| YNL067W | 45.4721521 | -1.2561539 | 0.62200271 | 5.1068029 | 0.07781653 | 0.18263384 | RPL9B | XIV | 499681 | 500256 | 1 |
| YNL068C | 21.8112389 | 0.93698317 | 0.82055837 | 5.25470404 | 0.07226958 | 0.17276666 | FKH2 | XIV | 495701 | 498289 | -1 |
| YNL069C | 85.5661003 | -1.8605233 | 0.64277173 | 8.08302977 | 0.01757083 | 0.05999792 | RPL16B | XIV | 493956 | 495001 | -1 |
| YNL070W | 42.11087 | -2.4932611 | 0.68655815 | 13.2143386 | 0.00135065 | 0.00839084 | TOM7 | XIV | 493366 | 493548 | 1 |
| YNL071W | 40.26343 | -0.8171126 | 0.72304613 | 1.38553636 | 0.50018954 | 0.65970328 | LAT1 | XIV | 491523 | 492971 | 1 |
| YNL072W | 166.894019 | -0.0396135 | 0.61264224 | 1.97157013 | 0.37314617 | 0.54640641 | RNH201 | XIV | 490317 | 491240 | 1 |
| YNL073W | 150.43352 | 1.13405626 | 0.59264165 | 13.3338672 | 0.00127229 | 0.00800643 | MSK1 | XIV | 488386 | 490116 | 1 |
| YNL074C | 415.812097 | 3.31912136 | 0.48153418 | 42.5069732 | 5.88E-10 | 2.24E-08 | MLF3 | XIV | 486766 | 488124 | -1 |
| YNL075W | 59.7274771 | -0.4620604 | 0.72049313 | 1.55931797 | 0.45856236 | 0.62328904 | IMP4 | XIV | 485607 | 486479 | 1 |
| YNL076W | 47.2650051 | -1.7222261 | 0.58805744 | 8.64844587 | 0.01324384 | 0.04862512 | MKS1 | XIV | 483556 | 485310 | 1 |
| YNL077W | 92.9081468 | 1.8685176 | 0.54892158 | 11.8344444 | 0.00269267 | 0.01402028 | APJ1 | XIV | 481391 | 482977 | 1 |
| YNL078W | 68.5134103 | -0.6708714 | 0.6049743 | 1.84518856 | 0.39748651 | 0.56811822 | NIS1 | XIV | 479768 | 480991 | 1 |
| YNL079C | 495.02206 | 0.10290158 | 0.50342952 | 6.06067987 | 0.04829922 | 0.12713301 | TPM1 | XIV | 478566 | 479165 | -1 |
| YNL080C | 75.9093976 | 2.46696962 | 0.56843092 | 26.725215 | 1.57E-06 | 2.63E-05 | EOS1 | XIV | 476932 | 478032 | -1 |
| YNL081C | 13.6338551 | -0.1893149 | 0.87075813 | 0.37804202 | 0.82776912 | 0.897021 | SWS2 | XIV | 476188 | 476619 | -1 |
| YNL082W | 93.7699191 | 0.95940652 | 0.98579811 | 7.8187896 | 0.02005263 | 0.06615655 | PMS1 | XIV | 473391 | 476012 | 1 |
| YNL083W | 32.1224337 | 2.41980223 | 0.69901889 | 14.0779487 | 0.00087703 | 0.00593222 | NA | XIV | 471377 | 472861 | 1 |
| YNL084C | 39.2845386 | -1.7226169 | 0.78766637 | 4.84106223 | 0.0888744 | 0.20203486 | END3 | XIV | 470053 | 471102 | -1 |

| | | | | | | | | | | | |
|---|---|---|---|---|---|---|---|---|---|---|---|
| YNL085W | 59.9361002 | -0.3453601 | 0.91283608 | 0.14910096 | 0.92816062 | 0.95498585 | MKT1 | XIV | 467131 | 469623 | 1 |
| YNL086W | 106.565075 | 0.78077568 | 0.5723987 | 2.93499614 | 0.23050146 | 0.39754218 | SNN1 | XIV | 466334 | 466642 | 1 |
| YNL087W | 431.10816 | 2.05568274 | 0.51575413 | 20.8314609 | 3.00E-05 | 0.00033944 | TCB2 | XIV | 462411 | 465947 | 1 |
| YNL088W | 54.1902413 | -0.5311374 | 0.55850454 | 2.32265586 | 0.31307017 | 0.48516903 | TOP2 | XIV | 457704 | 461990 | 1 |
| YNL090W | 12.2897595 | -1.36117 | 0.95719336 | 2.39785612 | 0.30151725 | 0.47448396 | RHO2 | XIV | 456565 | 457143 | 1 |
| YNL091W | 394.688535 | 0.55561488 | 0.75780211 | 5.64251933 | 0.05953091 | 0.1490203 | NST1 | XIV | 452408 | 456130 | 1 |
| YNL092W | 101.437923 | 1.01340846 | 0.51923493 | 3.93884202 | 0.13953762 | 0.27801887 | NA | XIV | 450871 | 452073 | 1 |
| YNL093W | 42.2184248 | 2.44597383 | 0.7182987 | 10.9946479 | 0.00409772 | 0.0195435 | YPT53 | XIV | 449868 | 450530 | 1 |
| YNL094W | 44.4771965 | 0.88048362 | 0.60398197 | 2.18317352 | 0.33568342 | 0.50893088 | APP1 | XIV | 447611 | 449374 | 1 |
| YNL095C | 994.545445 | -0.4524946 | 0.50664296 | 0.89251773 | 0.64001807 | 0.77557652 | NA | XIV | 444912 | 446840 | -1 |
| YNL096C | 34.0681041 | -1.5950042 | 0.71009509 | 5.14292483 | 0.0764237 | 0.18002165 | RPS7B | XIV | 443398 | 444315 | -1 |
| YNL097C | 34.9026636 | -0.5143472 | 0.6726258 | 0.77193933 | 0.67979115 | 0.80293569 | PHO23 | XIV | 441366 | 442358 | -1 |
| YNL097C-B | 79.995857 | 1.41341036 | 0.57636417 | 11.7030494 | 0.00287551 | 0.01474687 | NA | XIV | 440797 | 440919 | -1 |
| YNL098C | 113.157832 | -1.5872633 | 0.548147 | 8.10748173 | 0.01735732 | 0.05959198 | RAS2 | XIV | 439602 | 440570 | -1 |
| YNL099C | 65.1304785 | 0.54731856 | 0.57039288 | 0.91917889 | 0.63154288 | 0.76880708 | OCA1 | XIV | 438567 | 439283 | -1 |
| YNL100W | 93.552157 | -1.2902934 | 0.60127899 | 4.80174645 | 0.09063877 | 0.20427392 | MIC27 | XIV | 437613 | 438317 | 1 |
| YNL101W | 55.7589256 | -1.7697664 | 0.57017522 | 9.45783893 | 0.00883601 | 0.03582669 | AVT4 | XIV | 434999 | 437140 | 1 |
| YNL102W | 25.0197702 | 0.71674576 | 0.68340554 | 4.30465263 | 0.11621349 | 0.24370198 | POL1 | XIV | 430087 | 434493 | 1 |
| YNL103W | 55.7445089 | 0.76022665 | 0.57572717 | 2.87238664 | 0.23783139 | 0.40584843 | NA | XIV | 427735 | 429753 | 1 |
| YNL104C | 122.894442 | -0.546494 | 0.65143335 | 1.34892316 | 0.50943063 | 0.66733108 | LEU4 | XIV | 424895 | 426754 | -1 |
| YNL106C | 114.247228 | 0.16341825 | 0.74705 | 4.8440894 | 0.08873999 | 0.20185727 | INP52 | XIV | 420944 | 424495 | -1 |
| YNL107W | 55.3082166 | -0.8806924 | 0.57186872 | 2.49042978 | 0.28787904 | 0.46172289 | YAF9 | XIV | 420098 | 420778 | 1 |
| YNL108C | 76.4481733 | -0.0413456 | 0.53390482 | 0.00803877 | 0.99598868 | 0.99658361 | NA | XIV | 419014 | 419826 | -1 |
| YNL110C | 84.5231469 | -3.0216119 | 0.5643415 | 26.138301 | 2.11E-06 | 3.36E-05 | NOP15 | XIV | 417826 | 418488 | -1 |
| YNL111C | 6.86775019 | -2.3129717 | 1.1961143 | 4.14939552 | 0.12559438 | 0.25832734 | CYB5 | XIV | 416940 | 417302 | -1 |
| YNL112W | 122.087146 | 0.46927273 | 0.62956439 | 7.70595481 | 0.02121647 | 0.06916857 | DBP2 | XIV | 413639 | 416281 | 1 |
| YNL113W | 240.808963 | -0.905745 | 0.53441989 | 3.28371509 | 0.19362005 | 0.35033128 | RPC19 | XIV | 412771 | 413199 | 1 |
| YNL115C | 74.0206021 | 1.67419225 | 0.67483491 | 6.00307398 | 0.04971061 | 0.13011953 | NA | XIV | 410119 | 412053 | -1 |
| YNL116W | 26.0244842 | 0.89016319 | 0.68743588 | 2.86005822 | 0.23930196 | 0.40763705 | DMA2 | XIV | 408341 | 409909 | 1 |
| YNL117W | 192.221024 | -0.0349536 | 0.51603294 | 1.36580812 | 0.50514788 | 0.66322137 | MLS1 | XIV | 406358 | 408022 | 1 |
| YNL118C | 98.9233882 | -0.1792706 | 0.68626996 | 3.85797586 | 0.14529517 | 0.28623989 | DCP2 | XIV | 402652 | 405564 | -1 |
| YNL119W | 189.391569 | -1.0138871 | 0.67348363 | 8.26445602 | 0.01604709 | 0.05641325 | NCS2 | XIV | 401040 | 402521 | 1 |
| YNL121C | 38.9229377 | -1.0932491 | 0.66857611 | 2.86008842 | 0.23929834 | 0.40763705 | TOM70 | XIV | 398684 | 400537 | -1 |

| | | | | | | | | | | | |
|---|---|---|---|---|---|---|---|---|---|---|---|
| YNL122C | 24.9515894 | 2.00406472 | 0.75022842 | 8.62409263 | 0.01340609 | 0.04903427 | NA | XIV | 398023 | 398370 | -1 |
| YNL123W | 91.5331075 | 2.3725165 | 0.61246615 | 14.6161484 | 0.00067011 | 0.00472009 | NMA111 | XIV | 394685 | 397678 | 1 |
| YNL124W | 213.548022 | 0.11267952 | 0.58229739 | 0.35323248 | 0.83810135 | 0.90469306 | NAF1 | XIV | 392892 | 394370 | 1 |
| YNL125C | 24.9550021 | 1.34024807 | 0.72229019 | 3.37805898 | 0.18469869 | 0.34046654 | ESBP6 | XIV | 390146 | 392167 | -1 |
| YNL126W | 33.32406 | 1.21749669 | 0.7226932 | 2.90270008 | 0.23425382 | 0.40187545 | SPC98 | XIV | 387227 | 389767 | 1 |
| YNL127W | 59.5018119 | 1.75552872 | 0.56927143 | 14.3716634 | 0.00075724 | 0.00521018 | FAR11 | XIV | 383987 | 386848 | 1 |
| YNL128W | 18.6545304 | 0.7018565 | 0.74675402 | 2.51141352 | 0.28487444 | 0.45855519 | TEP1 | XIV | 382359 | 383663 | 1 |
| YNL129W | 54.0429913 | -0.0554575 | 0.57098054 | 1.41803372 | 0.49212779 | 0.6530873 | NRK1 | XIV | 381480 | 382202 | 1 |
| YNL130C | 26.721403 | -1.0620743 | 0.79050275 | 3.06967394 | 0.21549082 | 0.37935295 | CPT1 | XIV | 379558 | 380831 | -1 |
| YNL130C-A | 43.966705 | -2.1819188 | 0.68498717 | 11.0028733 | 0.0040809 | 0.01952763 | DGR1 | XIV | 381245 | 381391 | -1 |
| YNL131W | 24.6389554 | -1.7245059 | 0.71085343 | 6.40125281 | 0.04073668 | 0.11285424 | TOM22 | XIV | 378767 | 379225 | 1 |
| YNL132W | 26.8956583 | 0.02383711 | 0.75960378 | 1.13161926 | 0.56790017 | 0.71824858 | KRE33 | XIV | 375321 | 378491 | 1 |
| YNL133C | 40.059583 | -0.2708394 | 0.61149663 | 1.22541186 | 0.54188259 | 0.6958306 | FYV6 | XIV | 374171 | 374692 | -1 |
| YNL134C | 583.416392 | -0.818534 | 0.59071365 | 2.15014758 | 0.34127257 | 0.5148432 | NA | XIV | 372451 | 373581 | -1 |
| YNL136W | 696.624233 | 1.03648892 | 0.60848695 | 5.97756824 | 0.05034862 | 0.13131464 | EAF7 | XIV | 370368 | 371645 | 1 |
| YNL137C | 24.0473104 | -0.4105058 | 0.72006212 | 0.34163985 | 0.84297336 | 0.90805874 | NAM9 | XIV | 368595 | 370055 | -1 |
| YNL138W | 68.9820231 | 1.59964966 | 0.57469914 | 7.54001107 | 0.02305194 | 0.0736187 | SRV2 | XIV | 366741 | 368321 | 1 |
| YNL138W-A | 15.4470672 | -0.3443451 | 0.79194475 | 0.25226746 | 0.88149696 | 0.9300006 | YSF3 | XIV | 366033 | 366412 | 1 |
| YNL139C | 74.2041521 | 2.25911552 | 0.57173083 | 24.5971554 | 4.56E-06 | 6.58E-05 | THO2 | XIV | 360924 | 365717 | -1 |
| YNL141W | 795.159069 | -1.0770132 | 0.48635184 | 6.27371126 | 0.04341911 | 0.11809931 | AAH1 | XIV | 359596 | 360639 | 1 |
| YNL142W | 49.3757652 | -0.5113839 | 0.58148468 | 12.3879246 | 0.00204172 | 0.01132334 | MEP2 | XIV | 357453 | 358952 | 1 |
| YNL143C | 185.652425 | -0.3704512 | 0.54009393 | 0.49409776 | 0.78110251 | 0.86705973 | NA | XIV | 356794 | 357186 | -1 |
| YNL144C | 121.66511 | 0.38235704 | 0.53018836 | 2.36465471 | 0.30656442 | 0.47921383 | NA | XIV | 352820 | 355042 | -1 |
| YNL145W | 15.9440573 | -2.4304157 | 0.8062856 | 9.98230154 | 0.00679784 | 0.02932897 | MFA2 | XIV | 352414 | 352530 | 1 |
| YNL146C-A | 313.210433 | -1.6399078 | 0.67157423 | 8.7899673 | 0.01233908 | 0.0461222 | NA | XIV | 351386 | 351580 | -1 |
| YNL147W | 153.439426 | -0.5000732 | 0.59086147 | 0.71761253 | 0.69850966 | 0.81736153 | LSM7 | XIV | 350940 | 351383 | 1 |
| YNL148C | 23.6942946 | -0.6418719 | 0.67368091 | 1.28660785 | 0.52555316 | 0.6809024 | ALF1 | XIV | 349907 | 350671 | -1 |
| YNL149C | 40.3400702 | -0.7163988 | 0.72505385 | 1.00942572 | 0.60367889 | 0.74934389 | PGA2 | XIV | 349367 | 349756 | -1 |
| YNL151C | 32.2450808 | -0.9427142 | 0.68057546 | 1.91872314 | 0.38313741 | 0.55639971 | RPC31 | XIV | 347766 | 348521 | -1 |
| YNL152W | 14.6076563 | 0.90785933 | 0.83214508 | 3.98062346 | 0.13665282 | 0.27406298 | INN1 | XIV | 346313 | 347542 | 1 |
| YNL153C | 56.715367 | -0.3948712 | 0.62356917 | 2.6110715 | 0.27102729 | 0.44454618 | GIM3 | XIV | 345669 | 346058 | -1 |
| YNL154C | 1031.05432 | 3.78360095 | 0.45777228 | 62.737617 | 2.38E-14 | 2.15E-12 | YCK2 | XIV | 343636 | 345276 | -1 |
| YNL155W | 15.9200379 | -0.7428355 | 0.83289731 | 0.97717188 | 0.6134933 | 0.75641529 | CUZ1 | XIV | 342516 | 343340 | 1 |

| | | | | | | | | | | | |
|---|---|---|---|---|---|---|---|---|---|---|---|
| YNL156C | 51.3730688 | 0.16646644 | 0.56645091 | 0.12465728 | 0.93957405 | 0.96037943 | NSG2 | XIV | 341069 | 341968 | -1 |
| YNL157W | 57.7125304 | 1.3903464 | 0.60428742 | 5.40464559 | 0.06704959 | 0.16264366 | IGO1 | XIV | 340352 | 340858 | 1 |
| YNL158W | 114.9382 | 3.24479184 | 0.57360109 | 33.5834895 | 5.10E-08 | 1.33E-06 | PGA1 | XIV | 339612 | 340208 | 1 |
| YNL159C | 80.0657402 | -1.8821016 | 0.5725117 | 12.8036988 | 0.00165849 | 0.00977866 | ASI2 | XIV | 338478 | 339347 | -1 |
| YNL160W | 233.930179 | -1.5399985 | 0.48467653 | 10.2800947 | 0.00585741 | 0.02598806 | YGP1 | XIV | 336545 | 337609 | 1 |
| YNL161W | 85.0185231 | 1.57245317 | 0.58654025 | 7.07495574 | 0.02908659 | 0.08680999 | CBK1 | XIV | 332597 | 334867 | 1 |
| YNL162W | 66.7535243 | -2.8707649 | 0.74476343 | 14.0743575 | 0.0008786 | 0.00593594 | RPL42A | XIV | 331322 | 332154 | 1 |
| YNL162W-A | 78.7536025 | -0.798456 | 0.54970499 | 2.36718795 | 0.30617637 | 0.47873648 | NA | XIV | 330329 | 330547 | 1 |
| YNL163C | 38.0530854 | -0.435298 | 0.66758968 | 0.66930432 | 0.71558695 | 0.8274912 | RIA1 | XIV | 326741 | 330073 | -1 |
| YNL164C | 24.1225233 | 0.09193128 | 1.04778784 | 7.50755807 | 0.02342904 | 0.07454061 | IBD2 | XIV | 325265 | 326320 | -1 |
| YNL165W | 37.9920069 | -0.0049539 | 0.63308512 | 0.25599689 | 0.87985475 | 0.92911891 | NA | XIV | 323832 | 325052 | 1 |
| YNL166C | 151.849359 | 2.46926016 | 0.62089693 | 18.9433808 | 7.70E-05 | 0.00074679 | BNI5 | XIV | 322219 | 323565 | -1 |
| YNL167C | 155.266063 | 1.55910515 | 0.50740622 | 12.0193847 | 0.00245484 | 0.01302799 | SKO1 | XIV | 319416 | 321359 | -1 |
| YNL168C | 97.4701283 | -0.511106 | 0.59204996 | 2.36258194 | 0.30688231 | 0.47958126 | FMP41 | XIV | 318030 | 318809 | -1 |
| YNL169C | 105.557281 | -1.2009713 | 0.69664265 | 6.66133962 | 0.03576914 | 0.10217233 | PSD1 | XIV | 316169 | 317671 | -1 |
| YNL172W | 1184.90243 | 4.82527052 | 0.47505793 | 78.9683086 | 7.12E-18 | 1.21E-15 | APC1 | XIV | 310636 | 315882 | 1 |
| YNL173C | 23.8126649 | -1.6851803 | 0.70474902 | 5.94080522 | 0.05128266 | 0.13339021 | MDG1 | XIV | 308957 | 310057 | -1 |
| YNL175C | 57.6833051 | -1.705675 | 0.64466716 | 6.87576509 | 0.03213265 | 0.0942798 | NOP13 | XIV | 307401 | 308612 | -1 |
| YNL176C | 44.7625102 | -0.3838008 | 0.63790173 | 0.47108729 | 0.79014118 | 0.87290926 | TDA7 | XIV | 305070 | 306980 | -1 |
| YNL177C | 14.6632185 | -1.4604994 | 0.90473532 | 2.70472959 | 0.25862793 | 0.43030337 | MRPL22 | XIV | 303686 | 304615 | -1 |
| YNL178W | 100.034115 | -2.9978725 | 0.5772937 | 26.7607682 | 1.55E-06 | 2.59E-05 | RPS3 | XIV | 302680 | 303402 | 1 |
| YNL180C | 184.802711 | 0.77999011 | 0.83031047 | 5.11040941 | 0.07767633 | 0.18237873 | RHO5 | XIV | 299654 | 300649 | -1 |
| YNL181W | 30.7319179 | -1.3378569 | 0.90431122 | 4.2712764 | 0.11816915 | 0.24673616 | PBR1 | XIV | 298335 | 299558 | 1 |
| YNL182C | 13.6025685 | -1.7348759 | 0.82696174 | 4.28548586 | 0.11733257 | 0.24546419 | IPI3 | XIV | 295960 | 297627 | -1 |
| YNL183C | 23.6767746 | -1.1819033 | 0.70885711 | 2.74510867 | 0.25345871 | 0.42549317 | NPR1 | XIV | 293137 | 295509 | -1 |
| YNL185C | 6.2353873 | -0.569424 | 1.19693967 | 0.42776017 | 0.8074452 | 0.88441988 | MRPL19 | XIV | 292193 | 292669 | -1 |
| YNL186W | 59.8978407 | -1.1651703 | 0.59676842 | 4.11558353 | 0.12773573 | 0.26143156 | UBP10 | XIV | 289498 | 291876 | 1 |
| YNL187W | 66.651575 | -1.229511 | 0.63755699 | 9.33293414 | 0.00940544 | 0.03766079 | SWT21 | XIV | 287994 | 289067 | 1 |
| YNL188W | 74.112174 | 1.9761011 | 0.58126968 | 19.0163206 | 7.42E-05 | 0.00072613 | KAR1 | XIV | 286308 | 287609 | 1 |
| YNL189W | 16.2443157 | -1.5053283 | 0.93559078 | 2.94342805 | 0.22953172 | 0.39623992 | SRP1 | XIV | 284260 | 285888 | 1 |
| YNL190W | 95.6737106 | -0.5935473 | 0.57806048 | 2.88704543 | 0.2360946 | 0.40406514 | NA | XIV | 282395 | 283009 | 1 |
| YNL191W | 32.5600283 | 2.29801483 | 0.68105217 | 16.832576 | 0.00022123 | 0.00185644 | DUG3 | XIV | 280432 | 281505 | 1 |
| YNL192W | 105.75236 | 1.04906122 | 0.68505708 | 12.5435889 | 0.00188884 | 0.01073788 | CHS1 | XIV | 276502 | 279897 | 1 |

| | | | | | | | | | | | |
|---|---|---|---|---|---|---|---|---|---|---|---|
| YNL193W | 200.047279 | 1.91701928 | 0.49055668 | 18.7244955 | 8.59E-05 | 0.00082519 | NA | XIV | 274368 | 276044 | 1 |
| YNL194C | 336.429552 | -1.2746611 | 0.49233332 | 15.413795 | 0.00044971 | 0.00337792 | NA | XIV | 272710 | 273615 | -1 |
| YNL195C | 83.5456041 | -1.728987 | 0.5585921 | 16.9069798 | 0.00021316 | 0.00180434 | NA | XIV | 271523 | 272308 | -1 |
| YNL196C | 255.317109 | -0.6269089 | 0.56593735 | 2.2550751 | 0.32382969 | 0.49734056 | SLZ1 | XIV | 270275 | 271171 | -1 |
| YNL197C | 36.5653046 | -1.3460589 | 0.62902895 | 4.47807891 | 0.10656081 | 0.22939149 | WHI3 | XIV | 267608 | 269593 | -1 |
| YNL199C | 123.154241 | 3.26783879 | 0.59819444 | 32.6680712 | 8.06E-08 | 2.02E-06 | GCR2 | XIV | 264926 | 266530 | -1 |
| YNL200C | 54.468545 | 0.60097647 | 0.57964552 | 2.63670587 | 0.26757565 | 0.44038176 | NNR1 | XIV | 263713 | 264453 | -1 |
| YNL201C | 63.1730495 | 2.48968861 | 0.61724942 | 18.3623209 | 0.00010296 | 0.00095843 | PSY2 | XIV | 260627 | 263203 | -1 |
| YNL202W | 408.132515 | -1.3607552 | 0.52805194 | 7.06082436 | 0.02929284 | 0.08719023 | SPS19 | XIV | 259578 | 260456 | 1 |
| YNL204C | 7.8099756 | -0.6238759 | 0.96819641 | 0.6316781 | 0.7291768 | 0.83668919 | SPS18 | XIV | 258375 | 259277 | -1 |
| YNL206C | 394.769881 | -0.4715265 | 0.46520451 | 5.50024007 | 0.06392019 | 0.15715409 | RTT106 | XIV | 256788 | 258155 | -1 |
| YNL207W | 206.635338 | 3.28082939 | 0.53166651 | 43.2619649 | 4.03E-10 | 1.62E-08 | RIO2 | XIV | 255353 | 256630 | 1 |
| YNL208W | 540.192045 | 0.27760817 | 0.53192879 | 1.8692073 | 0.3927415 | 0.56412138 | NA | XIV | 254418 | 255017 | 1 |
| YNL209W | 139.248198 | -2.0223082 | 0.60380814 | 10.6200701 | 0.00494175 | 0.02269053 | SSB2 | XIV | 252059 | 253900 | 1 |
| YNL210W | 540.133401 | -0.0285653 | 0.56835345 | 0.00785888 | 0.99607827 | 0.99658361 | MER1 | XIV | 250931 | 251743 | 1 |
| YNL212W | 39.2975461 | 0.81877921 | 0.77655197 | 1.13849288 | 0.56595176 | 0.71688048 | VID27 | XIV | 247461 | 249809 | 1 |
| YNL213C | 52.2152757 | -0.1662109 | 0.81238704 | 10.3743424 | 0.00558779 | 0.0250219 | RRG9 | XIV | 246460 | 247104 | -1 |
| YNL215W | 108.784861 | -0.2932118 | 0.6716352 | 1.24534432 | 0.53650888 | 0.69122974 | IES2 | XIV | 244468 | 245430 | 1 |
| YNL216W | 595.612072 | 2.64450892 | 0.48835055 | 30.6985151 | 2.16E-07 | 4.66E-06 | RAP1 | XIV | 241689 | 244172 | 1 |
| YNL217W | 34.9619619 | -0.1127107 | 0.62407227 | 0.03437917 | 0.98295731 | 0.98687755 | NA | XIV | 240331 | 241311 | 1 |
| YNL218W | 14.0170888 | -0.6227118 | 0.780307 | 3.25183478 | 0.19673111 | 0.35456204 | MGS1 | XIV | 238238 | 240001 | 1 |
| YNL219C | 8.39230777 | -0.9215312 | 1.0853346 | 0.80029607 | 0.67022082 | 0.79536351 | ALG9 | XIV | 235996 | 237663 | -1 |
| YNL220W | 34.7808311 | 0.40940664 | 0.64417823 | 1.05440221 | 0.59025472 | 0.73765326 | ADE12 | XIV | 234413 | 235714 | 1 |
| YNL221C | 55.2585846 | 2.28353497 | 0.61061139 | 18.3078154 | 0.00010581 | 0.00098018 | POP1 | XIV | 231068 | 233695 | -1 |
| YNL222W | 24.3789743 | -0.0489818 | 0.68823593 | 0.03108121 | 0.98457953 | 0.98751104 | SSU72 | XIV | 229094 | 229714 | 1 |
| YNL223W | 13.1989844 | 1.60701211 | 1.01042965 | 2.67878102 | 0.26200531 | 0.43337041 | ATG4 | XIV | 227370 | 228854 | 1 |
| YNL224C | 40.383097 | -0.6164972 | 0.61242325 | 4.20402501 | 0.12221023 | 0.25307484 | SQS1 | XIV | 224796 | 227099 | -1 |
| YNL225C | 29.6062206 | 2.56129164 | 0.68791543 | 20.7367585 | 3.14E-05 | 0.00034994 | CNM67 | XIV | 222724 | 224469 | -1 |
| YNL227C | 54.2701139 | 0.46072566 | 0.59338594 | 0.65577074 | 0.7204456 | 0.83045591 | JJJ1 | XIV | 220659 | 222431 | -1 |
| YNL229C | 49.939976 | 0.44187046 | 0.66490524 | 4.65830808 | 0.09737809 | 0.21492841 | URE2 | XIV | 219137 | 220201 | -1 |
| YNL230C | 18.5218307 | 0.72454414 | 0.81971989 | 3.3324652 | 0.1889576 | 0.34502193 | ELA1 | XIV | 217523 | 218662 | -1 |
| YNL231C | 24.571091 | -1.115671 | 0.83154612 | 2.05405247 | 0.3580702 | 0.53120548 | PDR16 | XIV | 215987 | 217042 | -1 |
| YNL232W | 30.6201998 | -0.1594087 | 0.77850899 | 0.10933652 | 0.94679919 | 0.96418144 | CSL4 | XIV | 214923 | 215801 | 1 |

| | | | | | | | | | | | |
|---|---|---|---|---|---|---|---|---|---|---|---|
| YNL233W | 36.0092696 | 1.30567736 | 0.69927519 | 3.7614841 | 0.15247692 | 0.29615611 | BNI4 | XIV | 211922 | 214600 | 1 |
| YNL234W | 10.1462143 | 0.18087705 | 0.96864687 | 2.74905005 | 0.25295972 | 0.42477864 | NA | XIV | 210233 | 211513 | 1 |
| YNL236W | 110.444384 | 1.44105247 | 0.60769734 | 8.28133921 | 0.01591219 | 0.05607523 | SIN4 | XIV | 206930 | 209854 | 1 |
| YNL237W | 1197.98362 | 0.17968747 | 0.54678308 | 0.15462871 | 0.92559884 | 0.95325814 | YTP1 | XIV | 205188 | 206567 | 1 |
| YNL238W | 25.2509688 | 0.22026355 | 0.72683626 | 1.1493231 | 0.56289535 | 0.71410256 | KEX2 | XIV | 202428 | 204872 | 1 |
| YNL239W | 344.374505 | 0.66715429 | 0.4773861 | 2.47665463 | 0.28986867 | 0.46375784 | NA | XIV | 200569 | 201933 | 1 |
| YNL240C | 155.487744 | -2.2609796 | 0.56385601 | 15.2355786 | 0.00049163 | 0.00363486 | NAR1 | XIV | 198502 | 199977 | -1 |
| YNL241C | 166.074698 | 0.16291765 | 0.52754923 | 1.13174532 | 0.56786438 | 0.71824858 | ZWF1 | XIV | 196426 | 197943 | -1 |
| YNL242W | 195.193844 | -0.1678958 | 0.52390439 | 0.87441108 | 0.64583867 | 0.77803336 | ATG2 | XIV | 191324 | 196102 | 1 |
| YNL243W | 62.0550639 | 0.08803437 | 0.57553793 | 1.71330264 | 0.42458149 | 0.59492495 | SLA2 | XIV | 188051 | 190957 | 1 |
| YNL244C | 58.3218267 | -1.020182 | 0.75015108 | 2.86024171 | 0.23928 | 0.40763705 | SUI1 | XIV | 187170 | 187496 | -1 |
| YNL245C | 487.202946 | -1.8111072 | 0.51200525 | 11.9622882 | 0.00252593 | 0.01335631 | CWC25 | XIV | 186346 | 186885 | -1 |
| YNL246W | 2197.14155 | 0.47358453 | 0.45787662 | 1.0868479 | 0.58075636 | 0.72940182 | VPS75 | XIV | 185460 | 186349 | 1 |
| YNL247W | 81.0404292 | -0.6243523 | 0.55580808 | 1.36896077 | 0.50435223 | 0.66262751 | NA | XIV | 182875 | 185178 | 1 |
| YNL248C | 21.8765074 | -0.4998276 | 0.70082304 | 1.35222039 | 0.50859147 | 0.66638258 | RPA49 | XIV | 181361 | 182608 | -1 |
| YNL249C | 385.060768 | -0.4868615 | 0.47012788 | 1.10887591 | 0.57439501 | 0.72409037 | MPA43 | XIV | 179394 | 181022 | -1 |
| YNL250W | 411.788885 | -0.7224048 | 0.46815083 | 4.08696602 | 0.12957661 | 0.26389327 | RAD50 | XIV | 175410 | 179348 | 1 |
| YNL251C | 96.0842937 | -0.9730951 | 0.52526438 | 3.38324113 | 0.18422074 | 0.33979289 | NRD1 | XIV | 172588 | 174315 | -1 |
| YNL252C | 47.8379783 | -0.9308963 | 0.81838889 | 5.42874583 | 0.06624648 | 0.16143398 | MRPL17 | XIV | 171441 | 172286 | -1 |
| YNL253W | 32.7777418 | -0.9719891 | 0.68300873 | 3.17054814 | 0.20489163 | 0.36603596 | TEX1 | XIV | 170018 | 171286 | 1 |
| YNL254C | 166.907524 | 0.8944042 | 0.56804189 | 4.82924801 | 0.08940095 | 0.20267482 | RTC4 | XIV | 168043 | 169248 | -1 |
| YNL255C | 55.0902088 | -1.7082268 | 0.66743234 | 6.5969286 | 0.03693985 | 0.10474131 | GIS2 | XIV | 167329 | 167790 | -1 |
| YNL256W | 988.504187 | 0.18593096 | 0.69101227 | 1.28244423 | 0.5266484 | 0.68176637 | FOL1 | XIV | 164623 | 167097 | 1 |
| YNL257C | 74.709436 | -0.8482813 | 0.55683602 | 4.40817738 | 0.11035104 | 0.23522866 | SIP3 | XIV | 160631 | 164320 | -1 |
| YNL258C | 76.3408158 | 2.17350716 | 0.58706147 | 19.4116893 | 6.09E-05 | 0.00061032 | DSL1 | XIV | 158109 | 160373 | -1 |
| YNL260C | 107.750961 | -2.0685613 | 0.58193211 | 11.7871172 | 0.00275715 | 0.01424075 | LTO1 | XIV | 156859 | 157455 | -1 |
| YNL261W | 119.827604 | 0.08169465 | 0.61458086 | 2.51717331 | 0.28405521 | 0.45768219 | ORC5 | XIV | 155100 | 156539 | 1 |
| YNL262W | 74.9939808 | 0.27705179 | 0.55495321 | 1.21634957 | 0.54434351 | 0.69728958 | POL2 | XIV | 148212 | 154880 | 1 |
| YNL263C | 15.9385297 | -1.250906 | 0.85748268 | 2.38288445 | 0.30378282 | 0.47692586 | YIF1 | XIV | 146896 | 147840 | -1 |
| YNL264C | 5.39301556 | -0.2848715 | 1.23984302 | 0.34706169 | 0.84069122 | 0.90645906 | PDR17 | XIV | 145563 | 146615 | -1 |
| YNL265C | 93.5840283 | -0.6250143 | 0.66796388 | 5.01595415 | 0.08143281 | 0.18867385 | IST1 | XIV | 144280 | 145281 | -1 |
| YNL267W | 44.9170339 | -0.3489572 | 0.6878634 | 3.53054885 | 0.17113982 | 0.32161621 | PIK1 | XIV | 140878 | 144078 | 1 |
| YNL268W | 196.397832 | 1.40761353 | 0.52539905 | 7.19769378 | 0.02735525 | 0.08296851 | LYP1 | XIV | 138550 | 140385 | 1 |

| | | | | | | | | | | | |
|---|---|---|---|---|---|---|---|---|---|---|---|
| YNL269W | 56.4007457 | -0.9801158 | 0.74814759 | 1.88416498 | 0.3898152 | 0.56101577 | BSC4 | XIV | 137699 | 138094 | 1 |
| YNL270C | 73.1100454 | 2.4383201 | 0.65225678 | 21.8033093 | 1.84E-05 | 0.00021999 | ALP1 | XIV | 135940 | 137661 | -1 |
| YNL271C | 64.6654796 | -0.0511847 | 0.6047172 | 0.23893083 | 0.8873947 | 0.93215081 | BNI1 | XIV | 129522 | 135383 | -1 |
| YNL272C | 21.7670439 | -0.090699 | 0.69531905 | 0.03189003 | 0.98418144 | 0.98751104 | SEC2 | XIV | 126805 | 129084 | -1 |
| YNL273W | 398.275851 | -0.5510483 | 0.49380715 | 2.39725054 | 0.30160856 | 0.47448396 | TOF1 | XIV | 122883 | 126599 | 1 |
| YNL274C | 20.7320622 | -0.0729769 | 0.82910567 | 0.27234951 | 0.87269011 | 0.92527295 | GOR1 | XIV | 121118 | 122170 | -1 |
| YNL275W | 36.2056233 | 0.25808263 | 0.77347189 | 0.48622922 | 0.78418163 | 0.86949041 | BOR1 | XIV | 119268 | 120998 | 1 |
| YNL277W | 2021.596 | 0.01196318 | 0.50378055 | 0.34589753 | 0.84118071 | 0.90680252 | MET2 | XIV | 117349 | 118809 | 1 |
| YNL278W | 48.6309448 | 0.25134639 | 0.62160591 | 0.1846427 | 0.91181209 | 0.9456192 | CAF120 | XIV | 113271 | 116453 | 1 |
| YNL279W | 891.428855 | -1.0840757 | 0.50226543 | 9.57327268 | 0.00834046 | 0.03427345 | PRM1 | XIV | 110917 | 112902 | 1 |
| YNL280C | 529.483013 | -0.0984525 | 0.58460973 | 1.53914283 | 0.46321155 | 0.62634163 | ERG24 | XIV | 109095 | 110411 | -1 |
| YNL281W | 70.7021229 | -0.7381991 | 0.54662429 | 2.79478717 | 0.24724054 | 0.4180849 | HCH1 | XIV | 108467 | 108928 | 1 |
| YNL282W | 24.0004457 | -0.8860244 | 0.67534372 | 2.57393812 | 0.27610638 | 0.45007205 | POP3 | XIV | 107687 | 108274 | 1 |
| YNL283C | 58.7146703 | 0.8135715 | 0.62990956 | 1.91634647 | 0.38359298 | 0.55678199 | WSC2 | XIV | 105184 | 106695 | -1 |
| YNL284C | 23.594413 | 0.07444171 | 0.69421672 | 0.04101004 | 0.97970378 | 0.98440542 | MRPL10 | XIV | 103134 | 104102 | -1 |
| YNL286W | 205.347349 | 2.28006225 | 0.48484348 | 22.0703246 | 1.61E-05 | 0.00019822 | CUS2 | XIV | 95223 | 96080 | 1 |
| YNL287W | 33.4801967 | 0.84998564 | 0.73425282 | 1.40676881 | 0.4949075 | 0.65527428 | SEC21 | XIV | 91994 | 94801 | 1 |
| YNL288W | 54.6704749 | 1.30650224 | 0.58408893 | 5.55559836 | 0.06217519 | 0.15384375 | CAF40 | XIV | 90303 | 91424 | 1 |
| YNL289W | 19.7573996 | -2.3494316 | 0.7888162 | 9.30483506 | 0.00953851 | 0.03806203 | PCL1 | XIV | 87897 | 88736 | 1 |
| YNL290W | 27.589636 | -1.69137 | 0.73650225 | 5.34832286 | 0.06896464 | 0.16644654 | RFC3 | XIV | 86218 | 87240 | 1 |
| YNL291C | 22.0747864 | 1.63034995 | 0.80103703 | 6.65781712 | 0.03583219 | 0.10225577 | MID1 | XIV | 84169 | 85815 | -1 |
| YNL292W | 66.2504152 | -0.4828402 | 0.58020497 | 0.83479115 | 0.65876028 | 0.78659971 | PUS4 | XIV | 82806 | 84017 | 1 |
| YNL293W | 51.4356241 | 0.80878966 | 0.57376828 | 2.03625128 | 0.36127146 | 0.53408966 | MSB3 | XIV | 80640 | 82541 | 1 |
| YNL294C | 89.5947016 | 0.50034851 | 0.56799754 | 3.02363181 | 0.22050919 | 0.38525897 | RIM21 | XIV | 78659 | 80260 | -1 |
| YNL295W | 175.44022 | 0.604391 | 0.64170292 | 11.5128383 | 0.00316242 | 0.01590824 | MRX6 | XIV | 76946 | 78520 | 1 |
| YNL297C | 23.5195825 | -0.4534514 | 0.70202538 | 0.59631688 | 0.74218374 | 0.84243165 | MON2 | XIV | 71673 | 76583 | -1 |
| YNL298W | 66.8407858 | 0.11853928 | 0.71189023 | 2.08926492 | 0.3518211 | 0.5255532 | CLA4 | XIV | 68915 | 71443 | 1 |
| YNL299W | 143.682919 | 3.3850624 | 0.57151319 | 51.8950465 | 5.38E-12 | 3.43E-10 | TRF5 | XIV | 66517 | 68445 | 1 |
| YNL300W | 151.888724 | -0.2688162 | 0.6841298 | 1.43177719 | 0.48875761 | 0.65040372 | TOS6 | XIV | 65744 | 66052 | 1 |
| YNL301C | 44.2047003 | -1.19795 | 0.5979457 | 4.01033568 | 0.1346377 | 0.27105433 | RPL18B | XIV | 63570 | 64562 | -1 |
| YNL302C | 122.086514 | -2.8136807 | 0.88173027 | 9.20270099 | 0.01003827 | 0.03953849 | RPS19B | XIV | 61958 | 62943 | -1 |
| YNL304W | 42.5735855 | 0.64292298 | 0.69324228 | 4.24559713 | 0.11969618 | 0.2489022 | YPT11 | XIV | 60297 | 61550 | 1 |
| YNL305C | 108.733107 | 0.11162977 | 0.8189946 | 6.47651791 | 0.03923214 | 0.10991524 | BXI1 | XIV | 58898 | 59791 | -1 |

| | | | | | | | | | | | |
|---|---|---|---|---|---|---|---|---|---|---|---|
| YNL306W | 221.199628 | 1.99548745 | 0.90250493 | 11.4715595 | 0.00322836 | 0.01618375 | MRPS18 | XIV | 58155 | 58808 | 1 |
| YNL307C | 20.381995 | -1.3389287 | 0.79745756 | 2.98955596 | 0.2242984 | 0.38976222 | MCK1 | XIV | 56446 | 57573 | -1 |
| YNL308C | 232.689932 | -1.8863076 | 0.53978508 | 12.0463478 | 0.00242197 | 0.01291271 | KRI1 | XIV | 54122 | 55897 | -1 |
| YNL309W | 17.3113154 | -0.0610249 | 0.7577151 | 0.9067155 | 0.63549075 | 0.77219127 | STB1 | XIV | 52662 | 53924 | 1 |
| YNL310C | 10.8437912 | -1.02954 | 0.88539252 | 1.34525321 | 0.51036629 | 0.66786132 | ZIM17 | XIV | 51907 | 52431 | -1 |
| YNL311C | 37.3035245 | -0.0185015 | 0.77409549 | 2.80980887 | 0.24539051 | 0.41605008 | SKP2 | XIV | 49397 | 51688 | -1 |
| YNL312W | 53.128492 | -0.1320978 | 0.62062905 | 1.97643772 | 0.37223911 | 0.54549847 | RFA2 | XIV | 48287 | 49216 | 1 |
| YNL313C | 63.1044581 | 0.11239799 | 0.64512729 | 2.25286772 | 0.32418729 | 0.49763242 | EMW1 | XIV | 45308 | 48022 | -1 |
| YNL314W | 58.539092 | -1.2724501 | 0.90620244 | 4.76271407 | 0.09242507 | 0.20685781 | DAL82 | XIV | 44447 | 45214 | 1 |
| YNL315C | 16.2354101 | -0.2693423 | 0.75311219 | 0.14718791 | 0.92904885 | 0.95541731 | ATP11 | XIV | 43324 | 44280 | -1 |
| YNL316C | 306.475876 | 1.56589064 | 0.90408029 | 10.1383313 | 0.00628766 | 0.0275011 | PHA2 | XIV | 42071 | 43075 | -1 |
| YNL317W | 34.3889775 | 0.49488688 | 0.85487514 | 5.67297509 | 0.05863124 | 0.14727762 | PFS2 | XIV | 40619 | 42016 | 1 |
| YNL318C | 162.537016 | 0.39913587 | 0.84976606 | 8.47175688 | 0.0144671 | 0.05215721 | HXT14 | XIV | 38707 | 40329 | -1 |
| YNL320W | 27.6964423 | 0.73638902 | 0.83040922 | 2.12713689 | 0.3452217 | 0.51852261 | NA | XIV | 37700 | 38554 | 1 |
| YNL321W | 74.6297752 | -0.2253808 | 0.57210831 | 0.59722675 | 0.74184617 | 0.84243165 | VNX1 | XIV | 34696 | 37422 | 1 |
| YNL322C | 294.874699 | 0.75987667 | 0.8310522 | 9.11358629 | 0.01049566 | 0.04092248 | KRE1 | XIV | 33294 | 34235 | -1 |
| YNL323W | 305.087323 | -0.6018908 | 0.54177 | 3.62638926 | 0.16313216 | 0.3118307 | LEM3 | XIV | 31944 | 33188 | 1 |
| YNL325C | 54.877303 | 0.32912283 | 0.57433487 | 0.32983346 | 0.84796431 | 0.90980423 | FIG4 | XIV | 28739 | 31378 | -1 |
| YNL326C | 8.18216707 | 0.09863135 | 1.11823612 | 0.18286926 | 0.91262097 | 0.94617945 | PFA3 | XIV | 27337 | 28347 | -1 |
| YNL327W | 72.2198397 | -0.1748841 | 0.69015273 | 1.11762453 | 0.57188791 | 0.72218779 | EGT2 | XIV | 24048 | 27173 | 1 |
| YNL328C | 80.1179336 | 1.08600881 | 0.75548856 | 3.7147934 | 0.15607842 | 0.30143231 | MDJ2 | XIV | 22834 | 23274 | -1 |
| YNL329C | 38.7722231 | -1.0550519 | 0.61520389 | 2.99642148 | 0.22352975 | 0.38912726 | PEX6 | XIV | 19541 | 22633 | -1 |
| YNL330C | 13.4291546 | -0.1168191 | 0.93724704 | 0.17296047 | 0.91715368 | 0.94877967 | RPD3 | XIV | 18001 | 19302 | -1 |
| YNL331C | 348.213508 | 0.11725267 | 0.50390351 | 0.18561469 | 0.91136907 | 0.94542349 | AAD14 | XIV | 16118 | 17248 | -1 |
| YNL332W | 7.85211016 | 0.51538306 | 1.04234827 | 1.75671778 | 0.41546418 | 0.58728945 | THI12 | XIV | 14832 | 15854 | 1 |
| YNL333W | 47.5625948 | 1.33239966 | 0.62514179 | 4.3583556 | 0.11313451 | 0.23968124 | SNZ2 | XIV | 13267 | 14163 | 1 |
| YNL334C | 6.48151049 | 0.97699549 | 1.09446733 | 1.06863326 | 0.58606966 | 0.73378964 | SNO2 | XIV | 12208 | 12876 | -1 |
| YNL335W | 134.095756 | -0.9986288 | 0.54212245 | 5.9987314 | 0.04981866 | 0.13021013 | DDI3 | XIV | 11452 | 12129 | 1 |
| YNL336W | 229.973348 | 3.73868833 | 0.60059764 | 36.6014371 | 1.13E-08 | 3.34E-07 | COS1 | XIV | 8330 | 9475 | 1 |
| YNR001C | 48.3517872 | 1.70548 | 0.76112928 | 8.0055985 | 0.01826444 | 0.06184275 | CIT1 | XIV | 629622 | 631061 | -1 |
| YNR002C | 898.140829 | 1.10406156 | 0.50587431 | 4.66269792 | 0.09716459 | 0.21489634 | ATO2 | XIV | 633008 | 633856 | -1 |
| YNR004W | 681.992359 | 1.16109954 | 0.51572398 | 8.73556585 | 0.01267932 | 0.0471202 | SWM2 | XIV | 635943 | 636383 | 1 |
| YNR006W | 43.1013307 | 1.23463516 | 0.66821304 | 7.9242109 | 0.01902302 | 0.06366664 | VPS27 | XIV | 636986 | 638854 | 1 |

| | | | | | | | | | | | |
|---|---|---|---|---|---|---|---|---|---|---|---|
| YNR007C | 18.1083202 | 1.27179323 | 0.97564176 | 1.88372424 | 0.38990112 | 0.56101577 | ATG3 | XIV | 639182 | 640114 | -1 |
| YNR008W | 79.8538422 | -0.0053991 | 0.5348329 | 0.76233051 | 0.683065 | 0.80595449 | LRO1 | XIV | 640396 | 642381 | 1 |
| YNR009W | 195.101111 | 2.49924584 | 0.51074716 | 21.9352424 | 1.73E-05 | 0.00020766 | NRM1 | XIV | 642690 | 643439 | 1 |
| YNR010W | 61.9264056 | 1.558795 | 0.57654032 | 7.17266937 | 0.02769967 | 0.08374992 | CSE2 | XIV | 643744 | 644193 | 1 |
| YNR011C | 464.512499 | 4.23330553 | 0.53400824 | 51.524773 | 6.48E-12 | 4.08E-10 | PRP2 | XIV | 644320 | 646950 | -1 |
| YNR012W | 546.595819 | 0.24748793 | 0.92042914 | 10.8591535 | 0.00438495 | 0.02059113 | URK1 | XIV | 647432 | 648937 | 1 |
| YNR013C | 37.0748061 | -0.8986005 | 0.60328819 | 2.19222677 | 0.33416734 | 0.5079834 | PHO91 | XIV | 649028 | 651712 | -1 |
| YNR014W | 212.26516 | -1.839605 | 0.53212434 | 18.2394484 | 0.00010948 | 0.00100942 | NA | XIV | 652465 | 653103 | 1 |
| YNR015W | 79.8110266 | -0.4578737 | 0.89142888 | 4.5102024 | 0.10486293 | 0.22688953 | SMM1 | XIV | 653387 | 654541 | 1 |
| YNR016C | 85.0057386 | -0.9349526 | 0.625205 | 2.5231267 | 0.28321092 | 0.45678444 | ACC1 | XIV | 654673 | 661374 | -1 |
| YNR017W | 19.5008405 | -1.7696818 | 0.91375044 | 3.81168304 | 0.14869746 | 0.29106095 | TIM23 | XIV | 662913 | 663581 | 1 |
| YNR018W | 112.511025 | -2.0048721 | 0.52854566 | 15.8553076 | 0.00036063 | 0.00284348 | RCF2 | XIV | 664270 | 664944 | 1 |
| YNR019W | 89.5544446 | 0.72679359 | 0.57157127 | 4.42674502 | 0.10933131 | 0.2335036 | ARE2 | XIV | 665339 | 667267 | 1 |
| YNR020C | 23.9278169 | -0.7803113 | 0.70098206 | 1.58437853 | 0.45285229 | 0.61883757 | NA | XIV | 667410 | 668222 | -1 |
| YNR021W | 62.4212231 | -1.4501881 | 0.56264228 | 6.47736626 | 0.0392155 | 0.10991524 | NA | XIV | 668377 | 669591 | 1 |
| YNR022C | 313.694246 | -1.9878992 | 0.8557506 | 11.4370966 | 0.00328448 | 0.01637994 | MRPL50 | XIV | 669774 | 670193 | -1 |
| YNR023W | 35.929788 | 1.96257874 | 0.72275042 | 9.93661107 | 0.00695492 | 0.02989533 | SNF12 | XIV | 670418 | 672118 | 1 |
| YNR024W | 309.290298 | 2.85930319 | 0.80456576 | 11.541699 | 0.00311711 | 0.0157213 | MPP6 | XIV | 672409 | 672969 | 1 |
| YNR026C | 387.891433 | 4.89644276 | 0.532079 | 93.9485265 | 3.97E-21 | 1.35E-18 | SEC12 | XIV | 673274 | 674689 | -1 |
| YNR027W | 6.18698074 | -0.0965738 | 1.44068669 | 0.41736979 | 0.81165095 | 0.88731158 | BUD17 | XIV | 674923 | 675876 | 1 |
| YNR028W | 89.6730653 | 1.71602575 | 0.68026542 | 6.35292379 | 0.04173305 | 0.11462731 | CPR8 | XIV | 676177 | 677103 | 1 |
| YNR029C | 72.3947954 | -0.8544959 | 0.74853485 | 8.13159647 | 0.01714929 | 0.05904684 | NA | XIV | 677199 | 678488 | -1 |
| YNR030W | 11.0227003 | -0.3260907 | 1.03086332 | 0.32881706 | 0.84839536 | 0.90980423 | ALG12 | XIV | 678799 | 680454 | 1 |
| YNR031C | 12.0077344 | -0.40494 | 0.86752507 | 0.75729564 | 0.68478674 | 0.80718958 | SSK2 | XIV | 680694 | 685433 | -1 |
| YNR032C-A | 111.418053 | -0.6831592 | 0.7953782 | 2.88645192 | 0.23616467 | 0.40406514 | HUB1 | XIV | 687243 | 687464 | -1 |
| YNR032W | 44.9744378 | -1.2937401 | 0.64228085 | 4.29069412 | 0.11702741 | 0.24505921 | PPG1 | XIV | 686010 | 687116 | 1 |
| YNR033W | 65.4243211 | 0.82770839 | 0.57472827 | 2.71143318 | 0.25776252 | 0.42975819 | ABZ1 | XIV | 687635 | 689998 | 1 |
| YNR034W | 17.4871431 | 1.37718087 | 0.84854847 | 2.85716543 | 0.23964833 | 0.40786709 | SOL1 | XIV | 690321 | 691286 | 1 |
| YNR034W-A | 38.4801718 | 0.62407133 | 0.62896118 | 1.53432954 | 0.46432768 | 0.62726488 | EGO4 | XIV | 692561 | 692857 | 1 |
| YNR035C | 49.9743691 | 0.41887127 | 0.60029601 | 2.36868452 | 0.30594735 | 0.47867401 | ARC35 | XIV | 693019 | 694047 | -1 |
| YNR036C | 24.030335 | 0.37402197 | 0.72260054 | 0.33557076 | 0.84553529 | 0.90923806 | MRPS12 | XIV | 694361 | 694822 | -1 |
| YNR037C | 8.00364484 | 0.53294189 | 1.15327239 | 0.56848459 | 0.75258428 | 0.85034238 | RSM19 | XIV | 695052 | 695327 | -1 |
| YNR038W | 27.2784937 | -1.5894591 | 0.700916 | 5.58216062 | 0.0613549 | 0.15226954 | DBP6 | XIV | 695595 | 697484 | 1 |

| | | | | | | | | | | | |
|---|---|---|---|---|---|---|---|---|---|---|---|
| YNR039C | 80.2611807 | -1.6360593 | 0.71058857 | 10.9033002 | 0.00428922 | 0.02021351 | ZRG17 | XIV | 697614 | 699431 | -1 |
| YNR040W | 14.1816304 | 0.06130924 | 0.83204875 | 0.12095005 | 0.94131728 | 0.96106984 | NA | XIV | 699690 | 700460 | 1 |
| YNR041C | 56.2393097 | 0.24639807 | 0.5785967 | 0.31349152 | 0.85492139 | 0.91395769 | COQ2 | XIV | 700541 | 701659 | -1 |
| YNR043W | 45.6257504 | -0.5276666 | 0.63712476 | 0.71062027 | 0.70095602 | 0.81801018 | MVD1 | XIV | 701895 | 703085 | 1 |
| YNR044W | 20.5945621 | -0.3372817 | 0.7520393 | 0.95757607 | 0.61953379 | 0.76030111 | AGA1 | XIV | 703699 | 705876 | 1 |
| YNR045W | 58.6279175 | -1.6268211 | 0.57664129 | 10.6812619 | 0.00479285 | 0.02212703 | PET494 | XIV | 706139 | 707608 | 1 |
| YNR046W | 14.5027222 | -1.0410113 | 0.78803794 | 3.08398878 | 0.21395397 | 0.37804141 | TRM112 | XIV | 707788 | 708195 | 1 |
| YNR047W | 53.1910999 | -2.0765211 | 0.61125667 | 12.1547266 | 0.00229422 | 0.01236827 | FPK1 | XIV | 708523 | 711204 | 1 |
| YNR048W | 15.6427688 | 1.29332542 | 0.78642759 | 3.48637707 | 0.17496164 | 0.32678559 | NA | XIV | 711630 | 712811 | 1 |
| YNR049C | 29.2830958 | -0.0730445 | 0.68923681 | 0.01717526 | 0.99144914 | 0.99333631 | MSO1 | XIV | 713023 | 713655 | -1 |
| YNR050C | 34.1369823 | 0.0882798 | 0.74171473 | 0.03251604 | 0.98387343 | 0.98751104 | LYS9 | XIV | 714048 | 715388 | -1 |
| YNR051C | 80.3609361 | -1.1466559 | 0.53266666 | 5.95981619 | 0.0507975 | 0.13219068 | BRE5 | XIV | 716780 | 718327 | -1 |
| YNR052C | 20.3978625 | -0.723809 | 0.79432689 | 1.28012677 | 0.527259 | 0.68184633 | POP2 | XIV | 719346 | 720647 | -1 |
| YNR053C | 134.116212 | 0.28889069 | 0.64831191 | 1.83227671 | 0.40006096 | 0.57095216 | NOG2 | XIV | 721120 | 723112 | -1 |
| YNR054C | 37.3213495 | -2.534206 | 0.72430649 | 11.8080925 | 0.00272838 | 0.01413927 | ESF2 | XIV | 723356 | 724306 | -1 |
| YNR055C | 71.0876633 | 1.04384274 | 0.54829502 | 5.60861605 | 0.06054865 | 0.15085056 | HOL1 | XIV | 728426 | 730186 | -1 |
| YNR056C | 2600.7659 | 0.9379105 | 0.61312692 | 2.35967645 | 0.30732845 | 0.47995819 | BIO5 | XIV | 731618 | 733303 | -1 |
| YNR057C | 70.6033261 | -1.1545699 | 0.60210375 | 4.72988211 | 0.09395484 | 0.2093602 | BIO4 | XIV | 733356 | 734069 | -1 |
| YNR058W | 755.807315 | 6.99517331 | 0.62657523 | 99.6467019 | 2.30E-22 | 1.33E-19 | BIO3 | XIV | 734291 | 735733 | 1 |
| YNR059W | 35.5637298 | 0.46850417 | 0.69444745 | 3.18617346 | 0.20329712 | 0.36352388 | MNT4 | XIV | 736803 | 738545 | 1 |
| YNR060W | 601.466991 | 1.23806304 | 0.46263563 | 9.73618171 | 0.00768803 | 0.03213985 | FRE4 | XIV | 739951 | 742110 | 1 |
| YNR061C | 11.4951043 | -1.4907416 | 0.91469066 | 2.67459945 | 0.26255368 | 0.43396683 | NA | XIV | 742881 | 743540 | -1 |
| YNR062C | 965.310118 | 0.33396015 | 0.52591598 | 0.62617636 | 0.73118543 | 0.83700349 | NA | XIV | 744360 | 745343 | -1 |
| YNR063W | 719.023215 | 5.61898419 | 0.50283449 | 95.4702596 | 1.86E-21 | 6.72E-19 | NA | XIV | 746943 | 748766 | 1 |
| YNR064C | 413.403717 | -1.187711 | 0.49932869 | 7.2612672 | 0.02649939 | 0.08100922 | NA | XIV | 749136 | 750008 | -1 |
| YNR065C | 82.7083224 | 2.56480936 | 0.63881727 | 17.1049429 | 0.00019307 | 0.0016512 | NA | XIV | 750350 | 753700 | -1 |
| YNR066C | 60.9852785 | 0.52678697 | 0.59836784 | 2.29471987 | 0.31747382 | 0.49004889 | NA | XIV | 753725 | 755035 | -1 |
| YNR067C | 1658.65563 | 3.50506617 | 0.46990586 | 46.2706748 | 8.96E-11 | 4.32E-09 | DSE4 | XIV | 755746 | 759099 | -1 |
| YNR068C | 25.5915277 | -2.1250254 | 0.85090849 | 6.25006512 | 0.0439355 | 0.11887223 | NA | XIV | 760067 | 760885 | -1 |
| YNR069C | 53.7557522 | -0.8427551 | 0.6737019 | 2.76257547 | 0.25125479 | 0.42270232 | BSC5 | XIV | 761123 | 762592 | -1 |
| YNR070W | 23.6777157 | 1.38744175 | 0.86252212 | 3.35260168 | 0.18706468 | 0.34297166 | PDR18 | XIV | 765375 | 769376 | 1 |
| YNR072W | 12.4821222 | 1.5038855 | 0.91248861 | 3.09825428 | 0.21243332 | 0.3760284 | HXT17 | XIV | 772657 | 774351 | 1 |
| YNR075C-A | 679.999401 | 0.94799752 | 0.60030227 | 2.70838389 | 0.25815581 | 0.43001213 | NA | XIV | 781511 | 781603 | -1 |

| | | | | | | | | | | | |
|---|---|---|---|---|---|---|---|---|---|---|---|
| YNR075W | 102.688702 | 4.24456908 | 0.71658641 | 32.9628881 | 6.95E-08 | 1.75E-06 | COS10 | XIV | 779916 | 781040 | 1 |
| YOL001W | 63.4947046 | 3.01885494 | 0.65313164 | 21.9847393 | 1.68E-05 | 0.00020343 | PHO80 | XV | 325249 | 326130 | 1 |
| YOL002C | 111.771148 | 2.83061082 | 0.61886345 | 21.1478779 | 2.56E-05 | 0.00029438 | IZH2 | XV | 323411 | 324364 | -1 |
| YOL003C | 425.251481 | 1.50301214 | 0.47036871 | 9.8088094 | 0.00741386 | 0.03137882 | PFA4 | XV | 321858 | 322994 | -1 |
| YOL004W | 92.9126621 | 0.56977458 | 0.5164198 | 1.30229551 | 0.52144694 | 0.67724939 | SIN3 | XV | 316938 | 321548 | 1 |
| YOL005C | 35.2508043 | -0.1143366 | 0.69765777 | 0.08132799 | 0.96015169 | 0.97173192 | RPB11 | XV | 315813 | 316175 | -1 |
| YOL006C | 257.191758 | 0.3258985 | 0.51184196 | 0.60483753 | 0.73902852 | 0.84040603 | TOP1 | XV | 313078 | 315387 | -1 |
| YOL007C | 276.933098 | 2.1133809 | 0.53347047 | 20.9612818 | 2.81E-05 | 0.00031936 | CSI2 | XV | 311342 | 312367 | -1 |
| YOL008W | 73.3903805 | 0.13501437 | 0.59107759 | 0.83965085 | 0.65716153 | 0.78549013 | COQ10 | XV | 310312 | 310935 | 1 |
| YOL009C | 16.340064 | 2.53207895 | 0.96548575 | 7.29754111 | 0.0260231 | 0.07997546 | MDM12 | XV | 309324 | 310139 | -1 |
| YOL010W | 25.0995328 | 0.18271155 | 0.68787612 | 0.07981806 | 0.96087684 | 0.97212597 | RCL1 | XV | 307938 | 309041 | 1 |
| YOL011W | 22.9642181 | 0.47922394 | 0.7025744 | 0.49229392 | 0.78180732 | 0.86750947 | PLB3 | XV | 305349 | 307409 | 1 |
| YOL012C | 232.957717 | 0.20341952 | 0.52695761 | 0.35622844 | 0.83684683 | 0.90381331 | HTZ1 | XV | 303579 | 303983 | -1 |
| YOL013C | 44.7404919 | -0.3713406 | 0.65586152 | 1.05398654 | 0.59037741 | 0.73765326 | HRD1 | XV | 301380 | 303035 | -1 |
| YOL014W | 25.050078 | 1.11487497 | 0.69988966 | 2.53555322 | 0.28145671 | 0.45563342 | NA | XV | 299693 | 300067 | 1 |
| YOL015W | 39.6820177 | 0.46213454 | 0.68322123 | 1.149641 | 0.56280588 | 0.71410256 | IRC10 | XV | 297078 | 298838 | 1 |
| YOL016C | 98.538753 | 1.00186552 | 0.6795068 | 5.80695412 | 0.05483223 | 0.14011689 | CMK2 | XV | 294777 | 296120 | -1 |
| YOL017W | 306.619047 | 1.85224132 | 0.57892193 | 27.2520332 | 1.21E-06 | 2.08E-05 | ESC8 | XV | 292529 | 294673 | 1 |
| YOL018C | 27.0563243 | -0.0977269 | 0.70071356 | 1.47958926 | 0.47721191 | 0.63989277 | TLG2 | XV | 290881 | 292074 | -1 |
| YOL019W | 85.4745499 | 1.47516854 | 0.65709261 | 13.6217769 | 0.00110171 | 0.00712729 | NA | XV | 288898 | 290553 | 1 |
| YOL020W | 89.0652031 | 1.62934041 | 0.55720604 | 9.13180929 | 0.01040047 | 0.040606 | TAT2 | XV | 286172 | 287950 | 1 |
| YOL021C | 26.3072108 | -0.8993624 | 0.67249696 | 1.90264419 | 0.38623005 | 0.55874541 | DIS3 | XV | 282421 | 285426 | -1 |
| YOL022C | 41.3626991 | 0.95998416 | 0.61198862 | 4.67023911 | 0.09679891 | 0.21424529 | TSR4 | XV | 280272 | 281498 | -1 |
| YOL023W | 25.7067295 | 1.26205809 | 0.70101061 | 3.74389427 | 0.15382385 | 0.29821719 | IFM1 | XV | 278056 | 280086 | 1 |
| YOL025W | 26.809326 | 1.52716233 | 0.68401658 | 7.17104983 | 0.02772211 | 0.08377402 | LAG2 | XV | 274957 | 276939 | 1 |
| YOL026C | 14.8414966 | 0.68319021 | 0.83641039 | 0.78142093 | 0.67657602 | 0.8007717 | MIM1 | XV | 274012 | 274353 | -1 |
| YOL027C | 43.7781176 | 0.49958054 | 0.63139194 | 0.64571108 | 0.72407845 | 0.83364768 | MDM38 | XV | 272003 | 273724 | -1 |
| YOL028C | 25.6122913 | 0.05992021 | 0.75013631 | 0.95892285 | 0.61911674 | 0.76021421 | YAP7 | XV | 270633 | 271370 | -1 |
| YOL029C | 390.344253 | 0.85268016 | 1.59405248 | 4.06618957 | 0.13092969 | 0.26590071 | NA | XV | 269815 | 270420 | -1 |
| YOL030W | 38.6674049 | 0.18256521 | 0.6445402 | 0.20802644 | 0.90121338 | 0.93824416 | GAS5 | XV | 268187 | 269641 | 1 |
| YOL031C | 92.7343884 | -0.0583127 | 0.51794386 | 0.66179059 | 0.71828037 | 0.82928083 | SIL1 | XV | 266264 | 267529 | -1 |
| YOL032W | 179.817304 | 2.45176126 | 0.82927543 | 21.4512297 | 2.20E-05 | 0.00025693 | OPI10 | XV | 265429 | 266169 | 1 |
| YOL033W | 94.1959288 | 0.92852299 | 0.53531789 | 7.35746093 | 0.02525502 | 0.07827974 | MSE1 | XV | 263475 | 265085 | 1 |

| | | | | | | | | | | | |
|---|---|---|---|---|---|---|---|---|---|---|---|
| YOL034W | 19.3622168 | -0.6256284 | 0.78724938 | 0.63875136 | 0.72660253 | 0.83481707 | SMC5 | XV | 259923 | 263204 | 1 |
| YOL036W | 201.455001 | 4.09040385 | 0.57178223 | 45.0381059 | 1.66E-10 | 7.44E-09 | NA | XV | 256745 | 259030 | 1 |
| YOL038C-A | 31.5396252 | -1.3370382 | 0.64433571 | 4.34984823 | 0.11361678 | 0.24017566 | NA | XV | 254926 | 255021 | -1 |
| YOL038W | 82.1544553 | -0.024696 | 0.60006692 | 1.97058222 | 0.37333053 | 0.54640641 | PRE6 | XV | 255337 | 256101 | 1 |
| YOL039W | 76.8831998 | -1.9682732 | 0.5869568 | 10.6974152 | 0.00475429 | 0.02196915 | RPP2A | XV | 254297 | 254617 | 1 |
| YOL040C | 165.496314 | -2.6491246 | 0.69654276 | 13.9818714 | 0.00092019 | 0.00615027 | RPS15 | XV | 253149 | 253577 | -1 |
| YOL041C | 60.9819771 | 0.8789264 | 0.60933546 | 3.30304434 | 0.1917578 | 0.34826777 | NOP12 | XV | 251267 | 252646 | -1 |
| YOL042W | 90.1501908 | 4.81874794 | 0.75860216 | 38.741722 | 3.87E-09 | 1.27E-07 | NGL1 | XV | 249825 | 250916 | 1 |
| YOL043C | 142.616706 | 2.77657442 | 0.59522796 | 26.3016207 | 1.94E-06 | 3.18E-05 | NTG2 | XV | 248392 | 249534 | -1 |
| YOL044W | 32.0489979 | 1.04090828 | 0.92803798 | 4.74562512 | 0.09321818 | 0.20831078 | PEX15 | XV | 247150 | 248301 | 1 |
| YOL045W | 31.7394034 | 2.36795347 | 0.74170742 | 12.020465 | 0.00245352 | 0.01302799 | PSK2 | XV | 243497 | 246802 | 1 |
| YOL047C | 406.592469 | 2.26058651 | 0.51915695 | 17.7635719 | 0.0001389 | 0.0012449 | LDS2 | XV | 241614 | 242747 | -1 |
| YOL048C | 156.235201 | 0.18603333 | 0.87801098 | 9.7957388 | 0.00746247 | 0.03156149 | RRT8 | XV | 240204 | 241310 | -1 |
| YOL049W | 125.820516 | 1.26073964 | 0.69945698 | 6.07735615 | 0.04789817 | 0.12634641 | GSH2 | XV | 238619 | 240094 | 1 |
| YOL051W | 83.5437118 | -0.4678179 | 0.59598274 | 2.53140434 | 0.28204119 | 0.45618192 | GAL11 | XV | 234940 | 238185 | 1 |
| YOL052C | 242.187212 | 2.87731602 | 0.50202671 | 32.0652819 | 1.09E-07 | 2.58E-06 | SPE2 | XV | 232446 | 233636 | -1 |
| YOL052C-A | 22.9689517 | 3.92328927 | 0.85295174 | 30.0239551 | 3.02E-07 | 6.30E-06 | DDR2 | XV | 231570 | 231755 | -1 |
| YOL053W | 138.382063 | 5.82915386 | 0.90536042 | 36.454432 | 1.21E-08 | 3.55E-07 | AIM39 | XV | 230085 | 231272 | 1 |
| YOL054W | 34.2377613 | 1.82565218 | 0.67189034 | 7.78070453 | 0.02043815 | 0.06716053 | PSH1 | XV | 228614 | 229834 | 1 |
| YOL055C | 114.80483 | 0.18269443 | 0.52251196 | 1.92400222 | 0.38212744 | 0.55547441 | THI20 | XV | 224420 | 226075 | -1 |
| YOL056W | 8.53704861 | 0.75515974 | 0.95496067 | 1.1683332 | 0.55757035 | 0.70905608 | GPM3 | XV | 223268 | 224179 | 1 |
| YOL057W | 31.3450712 | -0.3994875 | 0.73485341 | 0.55758391 | 0.75669731 | 0.8528864 | NA | XV | 220767 | 222902 | 1 |
| YOL058W | 1927.87114 | 1.30987793 | 0.47165779 | 7.46144673 | 0.02397549 | 0.07565017 | ARG1 | XV | 219211 | 220473 | 1 |
| YOL059W | 56.6939223 | -0.0306481 | 0.58606589 | 0.68813055 | 0.70888266 | 0.82407349 | GPD2 | XV | 217126 | 218448 | 1 |
| YOL060C | 193.292297 | 3.34065747 | 0.5628081 | 41.8093606 | 8.34E-10 | 3.08E-08 | MAM3 | XV | 214017 | 216137 | -1 |
| YOL061W | 85.6035009 | 0.23233064 | 0.72364868 | 1.85721957 | 0.39510261 | 0.5664635 | PRS5 | XV | 212244 | 213734 | 1 |
| YOL062C | 27.7127169 | 0.5387816 | 0.76363871 | 0.53434335 | 0.76554164 | 0.8573474 | APM4 | XV | 210520 | 211995 | -1 |
| YOL063C | 10.6249843 | -0.4465856 | 0.97878232 | 0.30923669 | 0.8567421 | 0.91522818 | CRT10 | XV | 207392 | 210265 | -1 |
| YOL064C | 8.13586319 | -0.9125338 | 1.07138325 | 0.75583236 | 0.68528794 | 0.80761595 | MET22 | XV | 206103 | 207176 | -1 |
| YOL066C | 78.0005346 | 1.36402587 | 0.68836322 | 8.72837003 | 0.01272502 | 0.0472597 | RIB2 | XV | 202696 | 204471 | -1 |
| YOL067C | 35.7252296 | -0.8819345 | 0.64076325 | 6.06460722 | 0.04820447 | 0.12703862 | RTG1 | XV | 201985 | 202518 | -1 |
| YOL068C | 181.704835 | 0.28294947 | 0.50697967 | 2.02583342 | 0.36315821 | 0.53614552 | HST1 | XV | 200368 | 201879 | -1 |
| YOL069W | 451.504409 | -1.0070462 | 0.51177325 | 5.50294205 | 0.06383389 | 0.15707532 | NUF2 | XV | 198942 | 200297 | 1 |

| | | | | | | | | | | | |
|---|---|---|---|---|---|---|---|---|---|---|---|
| YOL070C | 23.2665935 | -0.7639989 | 0.71710188 | 1.55827734 | 0.45880102 | 0.62328904 | NBA1 | XV | 197220 | 198725 | -1 |
| YOL071W | 56.9918828 | 0.97199793 | 0.58464627 | 3.20805348 | 0.20108517 | 0.36023611 | SDH5 | XV | 196507 | 196995 | 1 |
| YOL072W | 7.86451042 | 0.23502789 | 1.07826456 | 0.1134023 | 0.94487641 | 0.96266662 | THP1 | XV | 194970 | 196337 | 1 |
| YOL073C | 22.9062031 | -0.3662519 | 0.70937999 | 0.32963303 | 0.8480493 | 0.90980423 | DSC2 | XV | 193832 | 194800 | -1 |
| YOL075C | 473.147564 | 1.30017606 | 0.47969575 | 8.53421166 | 0.01402231 | 0.05090229 | NA | XV | 189658 | 193542 | -1 |
| YOL076W | 111.015564 | -1.3079588 | 0.72487795 | 5.36731415 | 0.06831287 | 0.16528689 | MDM20 | XV | 187024 | 189414 | 1 |
| YOL077C | 64.4396751 | -1.7609281 | 0.88801259 | 8.37420278 | 0.01519025 | 0.05409075 | BRX1 | XV | 185848 | 186723 | -1 |
| YOL077W-A | 46.5758531 | 0.55728371 | 0.64766313 | 1.35928264 | 0.50679874 | 0.66464916 | ATP19 | XV | 185438 | 185644 | 1 |
| YOL078W | 33.5714347 | 0.47813864 | 0.64607275 | 2.53105213 | 0.28209086 | 0.45618192 | AVO1 | XV | 181682 | 185212 | 1 |
| YOL080C | 39.9091585 | -2.9907386 | 0.6853083 | 18.1549589 | 0.00011421 | 0.00104466 | REX4 | XV | 180558 | 181427 | -1 |
| YOL081W | 59.2170702 | -0.895046 | 0.58509666 | 2.60545949 | 0.27178887 | 0.44516479 | IRA2 | XV | 171070 | 180309 | 1 |
| YOL082W | 96.7384656 | -0.6460618 | 0.54265855 | 3.82102296 | 0.14800467 | 0.2902937 | ATG19 | XV | 168727 | 169974 | 1 |
| YOL083W | 29.9370306 | 0.37146629 | 0.63949626 | 1.39956882 | 0.49669237 | 0.65643662 | ATG34 | XV | 165714 | 166952 | 1 |
| YOL084W | 14.3495785 | 0.69855788 | 0.81736849 | 1.13440139 | 0.56711074 | 0.71787738 | PHM7 | XV | 162356 | 165331 | 1 |
| YOL085C | 103.45469 | -0.6972524 | 0.59374747 | 1.56036579 | 0.45832218 | 0.62328904 | NA | XV | 161673 | 162014 | -1 |
| YOL086C | 551.162842 | -1.8399946 | 0.65198299 | 8.96705991 | 0.01129348 | 0.04310431 | ADH1 | XV | 159548 | 160594 | -1 |
| YOL086W-A | 104.776284 | -0.608668 | 0.78196252 | 16.1751178 | 0.00030734 | 0.00249667 | MHF1 | XV | 159173 | 159445 | 1 |
| YOL087C | 29.944492 | -0.9404243 | 0.8474382 | 1.40614747 | 0.49506128 | 0.65532803 | DUF1 | XV | 155287 | 158637 | -1 |
| YOL088C | 30.4572721 | 0.61732393 | 0.6461526 | 1.44595394 | 0.48530536 | 0.64773825 | MPD2 | XV | 153912 | 154745 | -1 |
| YOL089C | 55.6766668 | -0.6625043 | 0.56576665 | 2.60934262 | 0.27126168 | 0.44467869 | HAL9 | XV | 150398 | 153490 | -1 |
| YOL090W | 76.9468128 | -0.999963 | 0.72993911 | 9.94467736 | 0.00692693 | 0.02979712 | MSH2 | XV | 147382 | 150276 | 1 |
| YOL091W | 57.2099947 | 2.56154314 | 0.60252225 | 17.8202421 | 0.00013502 | 0.00121388 | SPO21 | XV | 145334 | 147163 | 1 |
| YOL093W | 45.1446968 | -0.994984 | 0.65547282 | 2.37900609 | 0.30437249 | 0.47720463 | TRM10 | XV | 142815 | 143696 | 1 |
| YOL094C | 41.0235307 | -0.3622574 | 0.8141145 | 4.87333134 | 0.08745196 | 0.19966358 | RFC4 | XV | 141584 | 142555 | -1 |
| YOL095C | 29.9833478 | 1.93842543 | 0.68385106 | 9.14680993 | 0.01032275 | 0.04032978 | HMI1 | XV | 139227 | 141347 | -1 |
| YOL096C | 8.97811843 | 0.07459903 | 1.09117283 | 2.05529474 | 0.35784785 | 0.53120548 | COQ3 | XV | 138107 | 139045 | -1 |
| YOL097C | 82.4596159 | -2.0633745 | 0.57458922 | 12.2166812 | 0.00222424 | 0.01213793 | WRS1 | XV | 136527 | 137825 | -1 |
| YOL097W-A | 123.63831 | -2.9469371 | 0.83058566 | 20.2855543 | 3.94E-05 | 0.0004228 | NA | XV | 136220 | 136405 | 1 |
| YOL098C | 91.9019144 | 1.01975713 | 0.53982041 | 4.40348551 | 0.11061022 | 0.23545338 | SDD3 | XV | 132725 | 135838 | -1 |
| YOL100W | 207.471582 | 0.19853387 | 0.56895601 | 2.42483306 | 0.29747755 | 0.47136562 | PKH2 | XV | 129237 | 132482 | 1 |
| YOL101C | 87.2250739 | 2.13920697 | 0.55150337 | 17.0003828 | 0.00020343 | 0.00173469 | IZH4 | XV | 126982 | 127920 | -1 |
| YOL102C | 39.0274342 | -1.0583694 | 0.63923403 | 3.52549223 | 0.17157306 | 0.32232576 | TPT1 | XV | 125997 | 126689 | -1 |
| YOL103W | 129.857466 | 1.11348507 | 0.52655648 | 7.08455179 | 0.02894737 | 0.08664034 | ITR2 | XV | 124001 | 125830 | 1 |

| | | | | | | | | | | | |
|---|---|---|---|---|---|---|---|---|---|---|---|
| YOL103W-B | 7.44196863 | -1.5463343 | 1.04143292 | 2.54370611 | 0.28031171 | 0.45449588 | NA | XV | 118000 | 123268 | 1 |
| YOL104C | 11.586194 | 0.18658189 | 0.858325 | 0.23943952 | 0.88716902 | 0.93215081 | NDJ1 | XV | 116396 | 117454 | -1 |
| YOL105C | 15.5387254 | -0.591677 | 0.79517406 | 2.88140581 | 0.23676128 | 0.40473806 | WSC3 | XV | 114138 | 115808 | -1 |
| YOL107W | 15.6360287 | 2.53068098 | 0.87511862 | 13.0502243 | 0.00146615 | 0.00896413 | NA | XV | 112102 | 113130 | 1 |
| YOL108C | 104.766931 | 2.31676359 | 0.60558854 | 23.8347354 | 6.67E-06 | 9.16E-05 | INO4 | XV | 111431 | 111886 | -1 |
| YOL109W | 421.347437 | -2.695581 | 0.5737099 | 21.6306284 | 2.01E-05 | 0.00023746 | ZEO1 | XV | 110297 | 110638 | 1 |
| YOL110W | 16.267897 | -0.4623596 | 0.85389979 | 0.53929088 | 0.7636502 | 0.85622283 | SHR5 | XV | 109176 | 109889 | 1 |
| YOL111C | 175.71969 | 0.72136645 | 0.73368139 | 5.23010941 | 0.07316379 | 0.17432853 | MDY2 | XV | 108258 | 108896 | -1 |
| YOL112W | 111.054547 | 0.2608172 | 0.70242314 | 0.96904193 | 0.61599221 | 0.75801867 | MSB4 | XV | 106710 | 108188 | 1 |
| YOL113W | 43.1805564 | -0.4322053 | 0.65019784 | 0.55157088 | 0.75897576 | 0.85429037 | SKM1 | XV | 104326 | 106293 | 1 |
| YOL114C | 37.6135884 | -0.1875207 | 1.16352054 | 6.79925432 | 0.03338572 | 0.09713733 | PTH4 | XV | 103317 | 103925 | -1 |
| YOL115W | 26.0084112 | -1.5579019 | 0.66202609 | 6.71730722 | 0.03478206 | 0.10004686 | PAP2 | XV | 101475 | 103229 | 1 |
| YOL116W | 36.7293796 | -3.619962 | 0.6783867 | 29.2027946 | 4.56E-07 | 8.94E-06 | MSN1 | XV | 99809 | 100957 | 1 |
| YOL117W | 11.2597536 | 0.7352988 | 0.91050545 | 1.54011979 | 0.46298534 | 0.62618199 | RRI2 | XV | 97551 | 99488 | 1 |
| YOL118C | 625.513705 | -0.3461378 | 0.51470591 | 0.4554138 | 0.79635764 | 0.87685172 | NA | XV | 96608 | 96916 | -1 |
| YOL119C | 22.9022304 | -0.1274676 | 0.71655382 | 0.05312988 | 0.9737848 | 0.97984448 | MCH4 | XV | 94856 | 96361 | -1 |
| YOL120C | 125.677943 | -3.1909127 | 0.79717091 | 14.3726163 | 0.00075688 | 0.00521018 | RPL18B | XV | 93395 | 94402 | -1 |
| YOL121C | 68.976464 | -1.8536166 | 0.62706474 | 12.8812236 | 0.00159543 | 0.00955278 | RPS19A | XV | 92026 | 92850 | -1 |
| YOL122C | 133.055772 | -2.312939 | 0.56379751 | 15.7382166 | 0.00038238 | 0.00297068 | SMF1 | XV | 89692 | 91419 | -1 |
| YOL123W | 42.4202096 | -2.39793 | 0.66545556 | 14.5792764 | 0.00068257 | 0.00479044 | HRP1 | XV | 87844 | 89448 | 1 |
| YOL124C | 104.269007 | 4.0050357 | 0.60117489 | 45.1895727 | 1.54E-10 | 6.96E-09 | TRM11 | XV | 85456 | 86757 | -1 |
| YOL125W | 256.89019 | 1.99294969 | 0.53356409 | 13.8991031 | 0.00095907 | 0.00635691 | TRM13 | XV | 83834 | 85264 | 1 |
| YOL126C | 859.195963 | 3.06084309 | 0.51357333 | 31.160816 | 1.71E-07 | 3.86E-06 | MDH2 | XV | 81787 | 82920 | -1 |
| YOL127W | 64.6320847 | -3.0413269 | 0.85665791 | 11.6218065 | 0.00299472 | 0.01523678 | RPL25 | XV | 80348 | 81190 | 1 |
| YOL128C | 38.8809389 | -1.6341437 | 0.69374308 | 5.82127701 | 0.05444096 | 0.13947484 | YGK3 | XV | 78352 | 79479 | -1 |
| YOL129W | 65.2338704 | -0.2279217 | 0.56192015 | 0.51694053 | 0.77223199 | 0.86150115 | VPS68 | XV | 77560 | 78114 | 1 |
| YOL130W | 35.2755909 | 0.01386389 | 0.6858732 | 0.42238853 | 0.80961677 | 0.8858947 | ALR1 | XV | 74400 | 76979 | 1 |
| YOL131W | 10.9306547 | -0.5183204 | 0.89506967 | 0.68241579 | 0.7109111 | 0.82438173 | NA | XV | 73031 | 73357 | 1 |
| YOL132W | 181.722749 | 0.20611728 | 0.49059732 | 0.37656105 | 0.82838229 | 0.89751749 | GAS4 | XV | 71300 | 72715 | 1 |
| YOL133W | 14.8149122 | -0.769891 | 0.80901188 | 1.05136591 | 0.5911515 | 0.7383018 | HRT1 | XV | 70325 | 70690 | 1 |
| YOL135C | 36.3126098 | -1.0934919 | 0.61231493 | 3.45746522 | 0.17750924 | 0.3305817 | MED7 | XV | 69376 | 70044 | -1 |
| YOL136C | 197.278376 | 4.72580299 | 0.57080629 | 63.3259808 | 1.77E-14 | 1.63E-12 | PFK27 | XV | 67561 | 68754 | -1 |
| YOL137W | 420.923363 | 4.36356885 | 0.64752799 | 39.5916206 | 2.53E-09 | 8.61E-08 | BSC6 | XV | 65621 | 67114 | 1 |

| | | | | | | | | | | | |
|---|---|---|---|---|---|---|---|---|---|---|---|
| YOL138C | 47.8426269 | 0.13989257 | 0.56498299 | 5.03116619 | 0.08081578 | 0.1877702 | RTC1 | XV | 61325 | 65350 | -1 |
| YOL139C | 26.0744634 | -3.340179 | 0.87413283 | 14.0525421 | 0.00088824 | 0.00596624 | CDC33 | XV | 60383 | 61024 | -1 |
| YOL140W | 9.96772976 | -1.6001909 | 0.97447816 | 3.37211504 | 0.18524842 | 0.34115406 | ARG8 | XV | 58759 | 60030 | 1 |
| YOL141W | 9.70493127 | 1.0716201 | 0.97811559 | 2.43926493 | 0.29533869 | 0.46952527 | PPM2 | XV | 56452 | 58539 | 1 |
| YOL142W | 12.8116719 | -1.494731 | 0.83960291 | 3.32693721 | 0.18948061 | 0.34544902 | RRP40 | XV | 55558 | 56280 | 1 |
| YOL143C | 53.10384 | 0.08223919 | 0.58096034 | 0.04583216 | 0.9773445 | 0.98260543 | RIB4 | XV | 54595 | 55104 | -1 |
| YOL144W | 701.166036 | 0.32518318 | 0.66933004 | 5.49506186 | 0.0640859 | 0.15736105 | NOP8 | XV | 53098 | 54552 | 1 |
| YOL145C | 66.0989102 | -0.033462 | 0.55318883 | 0.88434274 | 0.6426395 | 0.77621415 | CTR9 | XV | 49555 | 52788 | -1 |
| YOL146W | 118.11049 | 1.57332317 | 0.64020541 | 6.4323814 | 0.04010755 | 0.11164553 | PSF3 | XV | 48864 | 49448 | 1 |
| YOL147C | 26.9923362 | -1.0178201 | 0.6937767 | 3.02842966 | 0.21998084 | 0.38460377 | PEX11 | XV | 47933 | 48643 | -1 |
| YOL148C | 34.3962596 | 0.61770321 | 0.62555393 | 1.35626421 | 0.50756418 | 0.66521629 | SPT20 | XV | 45760 | 47574 | -1 |
| YOL149W | 14.7914644 | -2.2527108 | 0.93954772 | 6.11761034 | 0.04694375 | 0.12464616 | DCP1 | XV | 44938 | 45633 | 1 |
| YOL151W | 174.413073 | -2.8509408 | 0.5399605 | 29.7749926 | 3.42E-07 | 6.93E-06 | GRE2 | XV | 43694 | 44722 | 1 |
| YOL152W | 638.951866 | 0.62223022 | 0.5096635 | 4.45516274 | 0.10778882 | 0.23089058 | FRE7 | XV | 40748 | 42610 | 1 |
| YOL153C | 1435.49136 | 0.25586909 | 0.49133991 | 1.43278151 | 0.48851224 | 0.65022664 | NA | NA | NA | NA | NA |
| YOL154W | 468.904414 | 1.68394125 | 0.46737619 | 12.8475359 | 0.00162253 | 0.00963534 | ZPS1 | XV | 34658 | 35407 | 1 |
| YOL155C | 6482.0637 | -0.9917942 | 0.46007578 | 8.42038218 | 0.01484353 | 0.05301915 | HPF1 | XV | 28703 | 31606 | -1 |
| YOL155W-A | 199.56082 | 1.44016294 | 0.70615033 | 7.32272561 | 0.02569747 | 0.07926923 | NA | XV | 27084 | 27218 | 1 |
| YOL156W | 526.968956 | 1.35561657 | 0.49236875 | 7.41193502 | 0.02457643 | 0.07694787 | HXT11 | XV | 25273 | 26976 | 1 |
| YOL158C | 146.132403 | 2.97117035 | 0.55367363 | 33.0677791 | 6.60E-08 | 1.67E-06 | ENB1 | XV | 19491 | 21311 | -1 |
| YOL159C | 27.3725643 | 2.38191763 | 0.80619128 | 8.24251641 | 0.01622409 | 0.0567598 | CSS3 | XV | 17281 | 17796 | -1 |
| YOL159C-A | 157.181904 | 3.2100206 | 0.53808492 | 32.1355047 | 1.05E-07 | 2.53E-06 | NA | XV | 15233 | 15505 | -1 |
| YOL160W | 24.9658078 | 0.35308783 | 0.71663624 | 0.26139268 | 0.87748419 | 0.92780012 | NA | XV | 14313 | 14654 | 1 |
| YOL163W | 47.6941897 | 2.83518141 | 0.74636862 | 19.7832217 | 5.06E-05 | 0.00051851 | NA | XV | 9597 | 10106 | 1 |
| YOL164W | 470.54277 | -1.0010952 | 0.59683976 | 3.7317351 | 0.15476189 | 0.29951658 | BDS1 | XV | 6175 | 8115 | 1 |
| YOL164W-A | 186.96233 | 0.73472089 | 0.4812637 | 2.51336859 | 0.2845961 | 0.45823454 | NA | XV | 4130 | 4312 | 1 |
| YOL165C | 13.6286884 | -1.7603499 | 0.94246554 | 6.18537427 | 0.04537985 | 0.12165304 | AAD15 | XV | 1647 | 2078 | -1 |
| YOR001W | 35.6340114 | 0.01928172 | 0.67604718 | 0.49433408 | 0.78101022 | 0.86705973 | RRP6 | XV | 326832 | 329033 | 1 |
| YOR002W | 53.8234793 | 0.98555602 | 0.74935044 | 1.74337839 | 0.41824446 | 0.589459 | ALG6 | XV | 329417 | 331051 | 1 |
| YOR003W | 363.902661 | 1.46553739 | 0.47206442 | 9.63692563 | 0.0080792 | 0.03341947 | YSP3 | XV | 331455 | 332891 | 1 |
| YOR004W | 16.1589526 | -2.1788845 | 0.78920184 | 8.35341998 | 0.01534892 | 0.05445482 | UTP23 | XV | 333592 | 334356 | 1 |
| YOR005C | 998.780796 | 1.4528003 | 0.51061275 | 8.9770079 | 0.01123744 | 0.04297543 | DNL4 | XV | 334509 | 337343 | -1 |
| YOR006C | 794.243636 | 4.13243774 | 0.67000081 | 36.0213919 | 1.51E-08 | 4.32E-07 | TSR3 | XV | 337680 | 338621 | -1 |

| | | | | | | | | | | | |
|---|---|---|---|---|---|---|---|---|---|---|---|
| YOR007C | 81.4950711 | 0.14075836 | 0.65115726 | 0.11739765 | 0.94299073 | 0.96176085 | SGT2 | XV | 338938 | 339978 | -1 |
| YOR008C | 50.2356803 | 0.32971271 | 0.66762311 | 1.00789072 | 0.60414239 | 0.7493774 | SLG1 | XV | 341278 | 342414 | -1 |
| YOR008C-A | 80.2524277 | -3.2407763 | 0.60239838 | 29.905815 | 3.21E-07 | 6.58E-06 | NA | XV | 342857 | 343081 | -1 |
| YOR009W | 16.7794487 | 4.55451611 | 1.10912043 | 19.3443547 | 6.30E-05 | 0.00062796 | TIR4 | XV | 344335 | 345798 | 1 |
| YOR010C | 103.83477 | 1.34644239 | 0.5669089 | 7.30507212 | 0.0259253 | 0.07980195 | TIR2 | XV | 346195 | 346950 | -1 |
| YOR011W | 276.814243 | -0.2307637 | 0.52431718 | 0.20200154 | 0.90393234 | 0.93946657 | AUS1 | XV | 349679 | 353863 | 1 |
| YOR011W-A | 26.3636701 | 1.17467245 | 0.70934322 | 4.30925457 | 0.1159464 | 0.24332354 | NA | XV | 355652 | 355858 | 1 |
| YOR012W | 441.756498 | -1.6227605 | 0.53312568 | 8.97159336 | 0.01126791 | 0.04303508 | NA | XV | 356543 | 356956 | 1 |
| YOR014W | 152.183153 | 2.40835472 | 0.52276116 | 21.8281488 | 1.82E-05 | 0.00021773 | RTS1 | XV | 357674 | 359947 | 1 |
| YOR015W | 514.086934 | 0.21736668 | 0.80690154 | 4.35208785 | 0.11348962 | 0.24016437 | NA | XV | 359993 | 360352 | 1 |
| YOR016C | 109.53205 | 1.22163054 | 0.63288849 | 4.03179106 | 0.13320106 | 0.26928567 | ERP4 | XV | 360461 | 361084 | -1 |
| YOR017W | 1028.19863 | 5.32912555 | 0.48456393 | 98.2593317 | 4.61E-22 | 2.22E-19 | PET127 | XV | 361412 | 363814 | 1 |
| YOR018W | 74.8474412 | -0.7276718 | 0.5467033 | 1.81962936 | 0.40259883 | 0.57358445 | ROD1 | XV | 364369 | 366882 | 1 |
| YOR019W | 356.523054 | -1.2850487 | 0.47357264 | 8.59596015 | 0.01359599 | 0.0495723 | NA | XV | 368127 | 370319 | 1 |
| YOR020C | 56.4426165 | 0.59075778 | 0.58763781 | 2.37931841 | 0.30432496 | 0.47720463 | HSP10 | XV | 370524 | 370844 | -1 |
| YOR020W-A | 30.7133408 | -0.7026596 | 0.66347216 | 1.31621827 | 0.51782955 | 0.67436642 | NA | XV | 371685 | 371957 | 1 |
| YOR021C | 8.92841555 | -1.6149187 | 1.24989796 | 1.82927067 | 0.40066271 | 0.57138844 | SFM1 | XV | 372798 | 373439 | -1 |
| YOR022C | 158.926166 | 1.37231606 | 0.52392063 | 11.8975227 | 0.00260907 | 0.01368683 | DDL1 | XV | 373710 | 375857 | -1 |
| YOR023C | 36.3323203 | 2.10187545 | 0.66493187 | 10.2171509 | 0.00604469 | 0.02665555 | AHC1 | XV | 376012 | 377712 | -1 |
| YOR025W | 29.8815031 | -0.0683755 | 0.66855051 | 0.10005575 | 0.95120291 | 0.96614462 | HST3 | XV | 378219 | 379562 | 1 |
| YOR026W | 71.8919627 | -0.5854033 | 0.63914924 | 4.73268067 | 0.09382346 | 0.20925957 | BUB3 | XV | 379781 | 380806 | 1 |
| YOR027W | 126.038214 | 0.63696875 | 0.6048913 | 1.52445307 | 0.46662631 | 0.62934226 | STI1 | XV | 381053 | 382822 | 1 |
| YOR028C | 10.0234402 | -1.5068854 | 0.8866607 | 3.93495672 | 0.13980896 | 0.27827222 | CIN5 | XV | 383533 | 384420 | -1 |
| YOR029W | 96.3287753 | -2.7279285 | 0.57286037 | 22.9154154 | 1.06E-05 | 0.00013875 | NA | XV | 384600 | 384935 | 1 |
| YOR030W | 93.0652191 | 3.42979209 | 0.58132146 | 37.8747401 | 5.96E-09 | 1.89E-07 | DFG16 | XV | 386825 | 388684 | 1 |
| YOR031W | 23.6088454 | 4.14641715 | 0.92380225 | 22.2659689 | 1.46E-05 | 0.00018246 | NA | NA | NA | NA | NA |
| YOR032C | 543.844122 | 0.83718833 | 0.50502358 | 9.16132162 | 0.01024812 | 0.04019232 | HMS1 | XV | 389771 | 391075 | -1 |
| YOR032W-A | 100.805184 | -0.5190714 | 0.75763184 | 3.12443523 | 0.20967059 | 0.37244958 | NA | XV | 392176 | 392376 | 1 |
| YOR033C | 87.5243077 | 1.23173082 | 0.56894157 | 11.8267849 | 0.002703 | 0.01404881 | EXO1 | XV | 392416 | 394524 | -1 |
| YOR034C | 57.1301806 | -0.4968599 | 0.56261361 | 11.881075 | 0.00263062 | 0.01378395 | AKR2 | XV | 394837 | 397086 | -1 |
| YOR034C-A | 11.0205544 | 1.52958865 | 0.96830516 | 7.71378356 | 0.02113359 | 0.06897602 | NA | XV | 397426 | 397668 | -1 |
| YOR035C | 44.7567629 | 2.51883704 | 0.63683224 | 15.1232077 | 0.00052004 | 0.00378167 | SHE4 | XV | 397735 | 400104 | -1 |
| YOR036W | 387.047932 | -0.7824863 | 0.49595899 | 3.15061344 | 0.20694407 | 0.36890584 | PEP12 | XV | 400348 | 401214 | 1 |

| | | | | | | | | | | | |
|---|---|---|---|---|---|---|---|---|---|---|---|
| YOR037W | 21.1699019 | 0.9330864 | 0.84829245 | 1.25976049 | 0.53265559 | 0.68718267 | CYC2 | XV | 401555 | 402655 | 1 |
| YOR038C | 41.3166425 | 1.65884034 | 0.77788833 | 5.16101052 | 0.07573573 | 0.17861909 | HIR2 | XV | 402761 | 405388 | -1 |
| YOR039W | 22.7492578 | -1.1739993 | 1.02441439 | 1.34197831 | 0.51120267 | 0.66829159 | CKB2 | XV | 405768 | 406544 | 1 |
| YOR040W | 172.030989 | 0.23088134 | 0.5195402 | 0.86181094 | 0.64992034 | 0.78139046 | GLO4 | XV | 407064 | 407921 | 1 |
| YOR042W | 149.347648 | -0.5661049 | 0.51929865 | 1.34635067 | 0.51008631 | 0.66786132 | CUE5 | XV | 408425 | 409660 | 1 |
| YOR043W | 47.0034561 | 0.62368693 | 0.72919215 | 0.9074327 | 0.6352629 | 0.77207645 | WHI2 | XV | 410870 | 412330 | 1 |
| YOR044W | 22.2652151 | 3.33376264 | 0.82793113 | 16.3364178 | 0.00028353 | 0.00232853 | IRC23 | XV | 413007 | 413480 | 1 |
| YOR045W | 41.9821184 | 0.38010065 | 0.67296662 | 4.20277091 | 0.12228689 | 0.25314304 | TOM6 | XV | 413852 | 414037 | 1 |
| YOR046C | 27.8629602 | 0.19764169 | 0.64927067 | 0.16056615 | 0.92285507 | 0.9514478 | DBP5 | XV | 414459 | 415907 | -1 |
| YOR047C | 16.6471161 | -0.7750126 | 0.84274316 | 2.27453001 | 0.32069492 | 0.49318024 | STD1 | XV | 416347 | 417681 | -1 |
| YOR048C | 62.0833798 | -0.5464248 | 0.54122993 | 3.78314289 | 0.15083459 | 0.2938534 | RAT1 | XV | 418630 | 421650 | -1 |
| YOR049C | 1879.43074 | 1.10735556 | 0.49497845 | 4.8955788 | 0.08648456 | 0.19807974 | NA | XV | 422668 | 423732 | -1 |
| YOR051C | 84.2332694 | 1.10405348 | 0.58471646 | 3.58844437 | 0.16625672 | 0.31551177 | ETT1 | XV | 424846 | 426084 | -1 |
| YOR052C | 59.8399416 | -2.3248176 | 0.61979806 | 13.6240747 | 0.00110045 | 0.00712707 | TMC1 | XV | 426772 | 427224 | -1 |
| YOR054C | 129.83011 | 1.26954575 | 0.62329474 | 9.28074133 | 0.00965412 | 0.03839103 | VHS3 | XV | 427833 | 429857 | -1 |
| YOR056C | 79.495714 | -0.3077166 | 0.55857032 | 0.64133662 | 0.72566391 | 0.83456178 | NOB1 | XV | 430247 | 431626 | -1 |
| YOR057W | 52.0609252 | 0.41015619 | 0.7811065 | 1.40382056 | 0.49563759 | 0.65579106 | SGT1 | XV | 432186 | 433373 | 1 |
| YOR058C | 782.967987 | 2.77541111 | 0.55479105 | 22.8769491 | 1.08E-05 | 0.0001408 | ASE1 | XV | 433688 | 436345 | -1 |
| YOR059C | 23.6992291 | 1.73766796 | 0.77833168 | 5.32960872 | 0.06961297 | 0.16759214 | LPL1 | XV | 438906 | 440258 | -1 |
| YOR060C | 19.9926872 | -0.344949 | 0.81030744 | 3.63826325 | 0.16216651 | 0.31078956 | SLD7 | XV | 440390 | 441163 | -1 |
| YOR061W | 30.0412746 | -1.6871998 | 0.73603148 | 6.14931215 | 0.04620552 | 0.12305885 | CKA2 | XV | 441534 | 442553 | 1 |
| YOR062C | 297.309682 | -0.6139441 | 0.64427498 | 5.78109215 | 0.05554587 | 0.14149168 | NA | XV | 442725 | 443531 | -1 |
| YOR063W | 220.199076 | -3.1077426 | 0.75977313 | 14.594028 | 0.00067756 | 0.004761 | RPL3 | XV | 444686 | 445849 | 1 |
| YOR064C | 178.261766 | -1.7242448 | 0.52689666 | 12.7903174 | 0.00166962 | 0.00979022 | YNG1 | XV | 446079 | 446738 | -1 |
| YOR065W | 184.811555 | 1.61840214 | 0.55677033 | 14.655679 | 0.00065699 | 0.00463727 | CYT1 | XV | 447439 | 448368 | 1 |
| YOR066W | 80.4108603 | 1.41159804 | 0.54867459 | 6.4449342 | 0.03985661 | 0.11116077 | MSA1 | XV | 449436 | 451325 | 1 |
| YOR067C | 29.0104386 | 1.47574229 | 0.67875483 | 7.83204282 | 0.01992019 | 0.06575707 | ALG8 | XV | 451729 | 453462 | -1 |
| YOR068C | 1152.90432 | 0.09081128 | 0.53444213 | 0.54484192 | 0.76153362 | 0.85526961 | VAM10 | XV | 453869 | 454213 | -1 |
| YOR069W | 74.5800439 | -2.0585223 | 0.60638416 | 11.0415674 | 0.00400271 | 0.01926491 | VPS5 | XV | 453768 | 455795 | 1 |
| YOR070C | 135.838085 | 0.57777838 | 0.83556422 | 9.66365492 | 0.00797194 | 0.03311157 | GYP1 | XV | 455907 | 457820 | -1 |
| YOR071C | 484.169463 | 1.34218958 | 0.51033738 | 12.5386544 | 0.0018935 | 0.01073788 | NRT1 | XV | 459480 | 461276 | -1 |
| YOR072W | 138.780857 | -1.4638678 | 0.57133955 | 7.3742069 | 0.02504444 | 0.07796092 | NA | XV | 461502 | 461816 | 1 |
| YOR073W | 85.4348211 | 1.15616612 | 0.5783848 | 10.7734562 | 0.00457692 | 0.02123429 | SGO1 | XV | 464771 | 466543 | 1 |

| | | | | | | | | | | | |
|---|---|---|---|---|---|---|---|---|---|---|---|
| YOR074C | 17.9526262 | -1.9958878 | 0.86401808 | 5.78449733 | 0.05545138 | 0.14137538 | CDC21 | XV | 466675 | 467589 | -1 |
| YOR075W | 15.2867584 | -1.248817 | 0.95600405 | 4.09162754 | 0.12927495 | 0.26337155 | UFE1 | XV | 468212 | 469252 | 1 |
| YOR076C | 9.57473995 | -1.2870677 | 0.95952741 | 2.24670402 | 0.32518793 | 0.49807536 | SKI7 | XV | 469377 | 471620 | -1 |
| YOR078W | 172.529836 | -0.0719515 | 0.83236377 | 8.10682954 | 0.01736298 | 0.05959198 | BUD21 | XV | 472725 | 473369 | 1 |
| YOR079C | 41.4516387 | -2.2209188 | 0.61654079 | 12.8141145 | 0.00164987 | 0.00975767 | ATX2 | XV | 473475 | 474416 | -1 |
| YOR080W | 96.4419123 | -1.8295134 | 0.62540972 | 8.82153097 | 0.01214588 | 0.04558315 | DIA2 | XV | 474594 | 476792 | 1 |
| YOR081C | 257.22113 | 0.58169551 | 0.65805828 | 4.7018752 | 0.09527979 | 0.21128685 | TGL5 | XV | 476939 | 479188 | -1 |
| YOR083W | 14.9585073 | -1.7015227 | 0.83386201 | 4.55866132 | 0.10235269 | 0.22279026 | WHI5 | XV | 479533 | 480420 | 1 |
| YOR084W | 15.7058557 | -0.8980748 | 0.77867 | 1.39006286 | 0.49905876 | 0.65860324 | LPX1 | XV | 480587 | 481750 | 1 |
| YOR085W | 58.8530464 | 0.23339726 | 0.60711666 | 2.37319715 | 0.30525781 | 0.47807485 | OST3 | XV | 482033 | 483085 | 1 |
| YOR086C | 116.515581 | -0.4186407 | 0.54002591 | 2.0105741 | 0.36593958 | 0.53926957 | TCB1 | XV | 483219 | 486779 | -1 |
| YOR087W | 26.0652891 | 1.33261593 | 0.73916318 | 4.80103965 | 0.09067081 | 0.20427392 | YVC1 | XV | 487707 | 489734 | 1 |
| YOR089C | 113.666329 | -0.2254304 | 0.55405403 | 1.43544608 | 0.48786184 | 0.64951024 | VPS21 | XV | 490196 | 490828 | -1 |
| YOR090C | 388.983381 | 0.35752828 | 0.49787161 | 5.72116153 | 0.05723551 | 0.14501691 | PTC5 | XV | 491123 | 492841 | -1 |
| YOR091W | 89.5624656 | -1.7038749 | 0.54042727 | 9.62742832 | 0.00811765 | 0.0335244 | TMA46 | XV | 493433 | 494470 | 1 |
| YOR092W | 146.795003 | -0.5950696 | 0.49543738 | 1.89400398 | 0.38790222 | 0.56008824 | ECM3 | XV | 495127 | 496968 | 1 |
| YOR093C | 43.6637967 | 0.64290432 | 0.59972824 | 2.82186295 | 0.24391598 | 0.41379241 | CMR2 | XV | 497506 | 502452 | -1 |
| YOR094W | 281.618257 | -3.9836641 | 0.59554765 | 37.3435535 | 7.78E-09 | 2.43E-07 | ARF3 | XV | 502795 | 503346 | 1 |
| YOR095C | 36.2023375 | -1.6218254 | 0.69131017 | 5.29342558 | 0.07088384 | 0.17022706 | RKI1 | XV | 503552 | 504328 | -1 |
| YOR096W | 58.791797 | -2.1725073 | 0.62240243 | 15.2394004 | 0.00049069 | 0.00363486 | RPS7A | XV | 505794 | 506767 | 1 |
| YOR097C | 353.146468 | 0.06383599 | 0.58269127 | 0.46534496 | 0.79241306 | 0.87392122 | NA | XV | 506978 | 507505 | -1 |
| YOR098C | 185.753282 | 2.78735343 | 0.56730117 | 22.5908543 | 1.24E-05 | 0.00015922 | NUP1 | XV | 507948 | 511178 | -1 |
| YOR099W | 37.9409054 | 1.65869065 | 0.78262367 | 4.73544232 | 0.093694 | 0.20905135 | KTR1 | XV | 511825 | 513006 | 1 |
| YOR101W | 27.7377992 | -0.3265927 | 0.69995689 | 1.1757923 | 0.55549473 | 0.70703257 | RAS1 | XV | 515244 | 516173 | 1 |
| YOR103C | 51.4427627 | -0.7800472 | 0.577963 | 1.99739885 | 0.36835821 | 0.54145571 | OST2 | XV | 516449 | 516841 | -1 |
| YOR104W | 34.7964257 | -0.0231755 | 0.62349235 | 0.01335669 | 0.99334391 | 0.9947183 | PIN2 | XV | 517642 | 518490 | 1 |
| YOR106W | 25.70349 | 2.51268812 | 0.7352623 | 19.8764669 | 4.83E-05 | 0.0005002 | VAM3 | XV | 519121 | 519972 | 1 |
| YOR107W | 115.318134 | -2.8410055 | 0.5380496 | 25.3123462 | 3.19E-06 | 4.88E-05 | RGS2 | XV | 521353 | 522282 | 1 |
| YOR108W | 35.084505 | -1.5612517 | 0.70346776 | 4.78351079 | 0.09146898 | 0.20551237 | LEU9 | XV | 523027 | 524841 | 1 |
| YOR109W | 60.4248504 | 0.50893227 | 0.61830373 | 0.72387426 | 0.69632614 | 0.81659461 | INP53 | XV | 525278 | 528601 | 1 |
| YOR110W | 49.9414387 | 0.95125614 | 0.58112481 | 2.65383197 | 0.26529417 | 0.43749737 | TFC7 | XV | 528941 | 530248 | 1 |
| YOR111W | 8.94155505 | 0.12307369 | 1.08678788 | 1.55959418 | 0.45849904 | 0.62328904 | NA | XV | 530429 | 531127 | 1 |
| YOR112W | 28.2133388 | -0.6428013 | 0.65153637 | 0.97191094 | 0.6151092 | 0.75743987 | CEX1 | XV | 531508 | 533793 | 1 |

| | | | | | | | | | | | |
|---|---|---|---|---|---|---|---|---|---|---|---|
| YOR113W | 47.5948239 | -1.5646581 | 0.69527755 | 5.62678845 | 0.06000099 | 0.15000247 | AZF1 | XV | 534075 | 536819 | 1 |
| YOR114W | 28.0008121 | -0.0207714 | 0.66330163 | 0.07851177 | 0.96150464 | 0.97258112 | NA | XV | 537570 | 538454 | 1 |
| YOR115C | 252.673376 | 2.29138312 | 0.56746653 | 18.7190639 | 8.61E-05 | 0.00082519 | TRS33 | XV | 538659 | 539465 | -1 |
| YOR116C | 259.581436 | 3.78818139 | 0.55296115 | 54.2317599 | 1.67E-12 | 1.14E-10 | RPO31 | XV | 539763 | 544145 | -1 |
| YOR117W | 40.0537699 | -0.4443416 | 1.08343574 | 0.18811669 | 0.91022965 | 0.94499367 | RPT5 | XV | 545029 | 546333 | 1 |
| YOR118W | 1711.36309 | -0.6985271 | 0.55928205 | 3.62447747 | 0.16328817 | 0.31192296 | RTC5 | XV | 546857 | 548560 | 1 |
| YOR119C | 202.664739 | -2.1834007 | 0.54477824 | 16.0287185 | 0.00033068 | 0.00265185 | RIO1 | XV | 548792 | 550246 | -1 |
| YOR120W | 122.857448 | 0.53385076 | 0.83940059 | 12.4948302 | 0.00193545 | 0.01091164 | GCY1 | XV | 551114 | 552052 | 1 |
| YOR122C | 26.8072216 | 0.02865189 | 0.79108421 | 0.2213026 | 0.89525087 | 0.93531262 | PFY1 | XV | 552298 | 552887 | -1 |
| YOR123C | 112.68325 | -0.0734112 | 0.57576553 | 3.57059574 | 0.16774709 | 0.3174038 | LEO1 | XV | 553176 | 554570 | -1 |
| YOR124C | 90.8470071 | -1.4634187 | 0.53425225 | 7.31979215 | 0.02573519 | 0.07934331 | UBP2 | XV | 554824 | 558642 | -1 |
| YOR125C | 71.2024743 | 3.14178417 | 0.6012277 | 27.4432826 | 1.10E-06 | 1.93E-05 | CAT5 | XV | 559030 | 559731 | -1 |
| YOR126C | 90.8834209 | -0.3132593 | 0.60262681 | 0.52385325 | 0.76956749 | 0.85969434 | IAH1 | XV | 559961 | 560677 | -1 |
| YOR127W | 29.9978847 | -0.2146169 | 0.69196029 | 0.79186421 | 0.67305239 | 0.79774275 | RGA1 | XV | 561170 | 564193 | 1 |
| YOR128C | 20.3164846 | 1.23306786 | 0.77051123 | 2.73527202 | 0.25470838 | 0.42697206 | ADE2 | XV | 564476 | 566191 | -1 |
| YOR129C | 80.6224739 | 0.94225282 | 0.59765388 | 2.72461242 | 0.25606954 | 0.42826189 | AFI1 | XV | 566877 | 569558 | -1 |
| YOR130C | 12.4647715 | -1.082393 | 0.83598655 | 1.67138018 | 0.43357517 | 0.60207676 | ORT1 | XV | 569929 | 570807 | -1 |
| YOR131C | 27.9307282 | -0.2726115 | 0.6937319 | 3.37292189 | 0.18517371 | 0.34112496 | NA | XV | 572182 | 572838 | -1 |
| YOR132W | 21.9483489 | 0.80614596 | 0.7371523 | 1.40126455 | 0.49627142 | 0.65603003 | VPS17 | XV | 573175 | 574830 | 1 |
| YOR133W | 40.8188868 | -1.201472 | 0.80351884 | 7.57766831 | 0.02262196 | 0.07268654 | EFT1 | XV | 575098 | 577626 | 1 |
| YOR134W | 280.410025 | 1.66725434 | 0.49604051 | 17.034807 | 0.00019996 | 0.00170761 | BAG7 | XV | 578564 | 579793 | 1 |
| YOR136W | 32.1745453 | 0.74589535 | 0.67091035 | 1.61015094 | 0.44705419 | 0.61424863 | IDH2 | XV | 580250 | 581359 | 1 |
| YOR137C | 32.9294103 | -0.334538 | 0.71192975 | 0.24417685 | 0.8850701 | 0.9315066 | SIA1 | XV | 581813 | 583681 | -1 |
| YOR138C | 56.4637835 | 1.29596773 | 0.6320649 | 4.77753625 | 0.09174263 | 0.20596736 | RUP1 | XV | 584309 | 586324 | -1 |
| YOR140W | 117.391406 | -1.6209351 | 0.5212056 | 9.27597543 | 0.00967715 | 0.0384035 | SFL1 | XV | 586981 | 589281 | 1 |
| YOR141C | 29.8101027 | 0.09771346 | 0.67026625 | 2.32843329 | 0.3121671 | 0.48457038 | ARP8 | XV | 589942 | 592587 | -1 |
| YOR142W | 32.1719218 | 0.57947389 | 0.63022466 | 0.9488558 | 0.62224094 | 0.76265348 | LSC1 | XV | 593057 | 594046 | 1 |
| YOR143C | 25.5927815 | 0.5928594 | 0.74700073 | 2.06479553 | 0.35615197 | 0.52942744 | THI80 | XV | 601383 | 602342 | -1 |
| YOR144C | 9.6568257 | 0.31529031 | 0.89782814 | 0.15607803 | 0.92492834 | 0.95277599 | ELG1 | XV | 602717 | 605092 | -1 |
| YOR145C | 75.9011814 | -1.5565766 | 0.7181665 | 5.38493965 | 0.06771349 | 0.16390515 | PNO1 | XV | 605347 | 606171 | -1 |
| YOR147W | 102.105408 | 3.56934556 | 0.6027572 | 34.6646705 | 2.97E-08 | 8.07E-07 | MDM32 | XV | 606607 | 608475 | 1 |
| YOR148C | 81.8643288 | -0.2181601 | 0.6470747 | 5.32977326 | 0.06960724 | 0.16759214 | SPP2 | XV | 608640 | 609197 | -1 |
| YOR149C | 127.652308 | 3.2715127 | 0.60733381 | 33.307188 | 5.85E-08 | 1.50E-06 | SMP3 | XV | 609838 | 611388 | -1 |

| | | | | | | | | | | | |
|---|---|---|---|---|---|---|---|---|---|---|---|
| YOR150W | 11.9670922 | 1.40369693 | 0.99312189 | 2.67988026 | 0.26186135 | 0.43337041 | MRPL23 | XV | 611999 | 612490 | 1 |
| YOR151C | 47.5063648 | -1.2729929 | 0.69746549 | 3.30203635 | 0.19185447 | 0.34833408 | RPB2 | XV | 612997 | 616671 | -1 |
| YOR152C | 49.7589791 | 0.65952358 | 0.58245321 | 2.3346658 | 0.31119583 | 0.48393923 | ATG40 | XV | 617518 | 618288 | -1 |
| YOR153W | 130.824396 | 0.28724179 | 0.66031885 | 18.4575234 | 9.82E-05 | 0.00092128 | PDR5 | XV | 619840 | 624375 | 1 |
| YOR154W | 42.3355903 | 2.18227114 | 0.61810966 | 15.161946 | 0.00051006 | 0.0037336 | SLP1 | XV | 624729 | 626492 | 1 |
| YOR155C | 104.02862 | 2.95016926 | 0.69825509 | 16.1575686 | 0.00031005 | 0.00251202 | ISN1 | XV | 626628 | 627980 | -1 |
| YOR156C | 36.2573285 | 1.80225589 | 0.63648148 | 9.07738187 | 0.01068739 | 0.04141899 | NFI1 | XV | 628360 | 630540 | -1 |
| YOR157C | 25.6050167 | -0.4174815 | 0.73684786 | 0.56332813 | 0.75452711 | 0.85160078 | PUP1 | XV | 630966 | 631751 | -1 |
| YOR158W | 308.492114 | -2.5432053 | 0.58547755 | 18.2100943 | 0.0001111 | 0.0010215 | PET123 | XV | 632164 | 633120 | 1 |
| YOR159C | 34.9980821 | -2.1436623 | 0.64858985 | 10.5043752 | 0.00523605 | 0.02383391 | SME1 | XV | 633282 | 633566 | -1 |
| YOR160W | 21.6087521 | 0.77301537 | 0.78312119 | 1.02425768 | 0.59921858 | 0.74628427 | MTR10 | XV | 633839 | 636757 | 1 |
| YOR161C | 146.625809 | 1.3105927 | 0.49467048 | 8.9855316 | 0.01118965 | 0.04282094 | PNS1 | XV | 636939 | 638558 | -1 |
| YOR161C-C | 21.9711705 | -1.0549567 | 0.70287379 | 2.40812248 | 0.29997347 | 0.47325515 | NA | XV | 639121 | 639267 | -1 |
| YOR162C | 209.988775 | -1.0237829 | 0.48349392 | 9.86529757 | 0.00720739 | 0.03068439 | YRR1 | XV | 639560 | 641992 | -1 |
| YOR163W | 374.975174 | -1.381944 | 0.52109497 | 9.08936097 | 0.01062357 | 0.04128218 | DDP1 | XV | 642741 | 643307 | 1 |
| YOR164C | 63.7567297 | -0.6982347 | 0.56010196 | 2.07897527 | 0.35363583 | 0.52744756 | GET4 | XV | 643394 | 644332 | -1 |
| YOR165W | 14.8736446 | -0.5904968 | 0.86214651 | 0.57713636 | 0.74933571 | 0.8479727 | SEY1 | XV | 644566 | 646896 | 1 |
| YOR166C | 77.9183439 | -1.2525546 | 0.57444577 | 4.83252469 | 0.0892546 | 0.20266044 | SWT1 | XV | 647126 | 648502 | -1 |
| YOR167C | 72.3060609 | -2.7811682 | 0.66314343 | 16.9620718 | 0.00020736 | 0.00176305 | RPS28A | XV | 648804 | 649007 | -1 |
| YOR168W | 43.6567533 | -2.2018299 | 0.8255783 | 7.42394163 | 0.02442933 | 0.0767476 | GLN4 | XV | 649303 | 651732 | 1 |
| YOR171C | 10.4305168 | -0.2899324 | 1.02481323 | 0.18820841 | 0.91018791 | 0.94499367 | LCB4 | XV | 652010 | 653884 | -1 |
| YOR172W | 26.8654167 | -0.6777512 | 0.69085242 | 0.96913523 | 0.61596347 | 0.75801867 | YRM1 | XV | 654210 | 656570 | 1 |
| YOR173W | 150.63964 | 1.78414655 | 0.50264576 | 18.3341759 | 0.00010442 | 0.0009689 | DCS2 | XV | 657264 | 658325 | 1 |
| YOR174W | 38.9371932 | -0.4090737 | 0.68542748 | 0.5510834 | 0.75916078 | 0.85433253 | MED4 | XV | 658747 | 659601 | 1 |
| YOR175C | 18.6225794 | 0.25982305 | 0.74897872 | 0.16771492 | 0.91956232 | 0.94974417 | ALE1 | XV | 659815 | 661674 | -1 |
| YOR176W | 75.7990219 | -0.9106136 | 0.69637307 | 1.68778762 | 0.43003279 | 0.59923342 | HEM15 | XV | 662401 | 663582 | 1 |
| YOR177C | 16.4096436 | -0.9246381 | 0.78574445 | 1.9248871 | 0.38195841 | 0.55538403 | MPC54 | XV | 665785 | 667179 | -1 |
| YOR178C | 1465.05242 | -1.548436 | 0.45685292 | 11.6192485 | 0.00299856 | 0.01524288 | GAC1 | XV | 667860 | 670241 | -1 |
| YOR179C | 5.6168239 | -1.4566613 | 1.24327482 | 1.85715673 | 0.39511502 | 0.5664635 | SYC1 | XV | 671845 | 672411 | -1 |
| YOR180C | 11.1607266 | -1.7232087 | 0.95064634 | 3.30847263 | 0.19123805 | 0.34765095 | DCI1 | XV | 674352 | 675167 | -1 |
| YOR181W | 63.0818532 | 0.85799323 | 0.76831522 | 5.2636222 | 0.07194804 | 0.17226374 | LAS17 | XV | 675939 | 677840 | 1 |
| YOR182C | 38.9373858 | -2.5137478 | 0.71140879 | 11.9543693 | 0.00253596 | 0.01338485 | RPS30A | XV | 678191 | 678793 | -1 |
| YOR183W | 43.3308802 | 0.71503178 | 0.60662833 | 1.61472266 | 0.44603345 | 0.61342843 | FYV12 | XV | 678872 | 679261 | 1 |

| | | | | | | | | | | | |
|---|---|---|---|---|---|---|---|---|---|---|---|
| YOR184W | 317.66801 | 4.55582421 | 0.60994492 | 56.7473054 | 4.76E-13 | 3.49E-11 | SER1 | XV | 679357 | 680544 | 1 |
| YOR185C | 270.52564 | 0.53778387 | 0.49601899 | 16.1565449 | 0.00031021 | 0.00251202 | GSP2 | XV | 681444 | 682106 | -1 |
| YOR187W | 48.6657299 | -0.5893503 | 0.72031604 | 0.69744246 | 0.7055898 | 0.82101385 | TUF1 | XV | 684030 | 685343 | 1 |
| YOR188W | 46.7652818 | 0.74051053 | 0.65822807 | 2.25225488 | 0.32428665 | 0.49763242 | MSB1 | XV | 685767 | 689180 | 1 |
| YOR189W | 12.7717702 | -0.5821325 | 0.9246988 | 1.32579915 | 0.51535486 | 0.67220199 | IES4 | XV | 689624 | 689974 | 1 |
| YOR190W | 444.087938 | 0.83302391 | 0.51211547 | 3.77644821 | 0.15134033 | 0.2946404 | SPR1 | XV | 690695 | 692032 | 1 |
| YOR191W | 61.0258604 | 0.25915141 | 0.595281 | 0.36367352 | 0.83373743 | 0.9018008 | ULS1 | XV | 692475 | 697334 | 1 |
| YOR192C | 1390.24366 | 2.21427083 | 0.58751301 | 13.2590477 | 0.00132079 | 0.00824961 | THI72 | XV | 698768 | 700567 | -1 |
| YOR192C-B | 7.39366793 | -0.399263 | 1.08634676 | 0.21605929 | 0.89760098 | 0.93705725 | NA | XV | 704419 | 709732 | -1 |
| YOR192C-C | 154.024734 | -0.545134 | 0.54143563 | 1.03532837 | 0.59591086 | 0.7432839 | NA | XV | 703988 | 704224 | -1 |
| YOR193W | 251.973422 | -1.4519679 | 0.48900701 | 8.81994614 | 0.01215551 | 0.04558315 | PEX27 | XV | 710446 | 711576 | 1 |
| YOR194C | 397.306355 | 1.03375314 | 0.70644358 | 7.15594249 | 0.02793231 | 0.08418952 | TOA1 | XV | 711683 | 712543 | -1 |
| YOR195W | 33.1060478 | -0.2713027 | 0.69298737 | 0.26691332 | 0.87506539 | 0.9267658 | SLK19 | XV | 712866 | 715331 | 1 |
| YOR196C | 74.7984565 | -1.0701053 | 0.55826248 | 3.58433929 | 0.16659832 | 0.31564276 | LIP5 | XV | 715593 | 716837 | -1 |
| YOR197W | 101.853367 | 0.61956364 | 0.67461614 | 2.58196599 | 0.27500033 | 0.44841232 | MCA1 | XV | 717086 | 718384 | 1 |
| YOR198C | 157.167293 | 0.47996133 | 0.61934529 | 3.02763963 | 0.22006775 | 0.38460377 | BFR1 | XV | 718653 | 720065 | -1 |
| YOR201C | 130.22359 | 3.10071847 | 0.5772132 | 26.6366356 | 1.64E-06 | 2.74E-05 | MRM1 | XV | 720470 | 721708 | -1 |
| YOR202W | 228.175575 | 1.05705249 | 0.52996501 | 9.04501516 | 0.01086175 | 0.04192011 | HIS3 | XV | 721946 | 722608 | 1 |
| YOR204W | 43.5909786 | -1.6879834 | 0.69887525 | 5.72401536 | 0.0571539 | 0.14493103 | DED1 | XV | 722911 | 724725 | 1 |
| YOR205C | 16.3792919 | -0.5893378 | 0.78193088 | 0.62030254 | 0.73333602 | 0.83764363 | GEP3 | XV | 725564 | 727234 | -1 |
| YOR206W | 52.5328667 | -3.3865803 | 0.73207082 | 20.0899528 | 4.34E-05 | 0.00045692 | NOC2 | XV | 727512 | 729644 | 1 |
| YOR207C | 56.3565503 | 0.7792811 | 0.60510516 | 1.82478297 | 0.40156274 | 0.57224915 | RET1 | XV | 730008 | 733457 | -1 |
| YOR208W | 75.5377254 | 2.95070041 | 0.6288067 | 20.0576829 | 4.41E-05 | 0.00046183 | PTP2 | XV | 733925 | 736177 | 1 |
| YOR209C | 35.15554 | -0.2135067 | 1.06702155 | 1.24109116 | 0.53765103 | 0.69208525 | NPT1 | XV | 736437 | 737726 | -1 |
| YOR211C | 18.9390104 | 0.10254017 | 0.75562298 | 0.27312282 | 0.87235275 | 0.92527295 | MGM1 | XV | 738924 | 741569 | -1 |
| YOR212W | 63.1175787 | -2.1803276 | 0.63363373 | 11.3374463 | 0.00345227 | 0.01701161 | STE4 | XV | 742910 | 744181 | 1 |
| YOR213C | 318.991893 | -1.2460826 | 0.49223038 | 12.0969448 | 0.00236147 | 0.01267182 | SAS5 | XV | 744535 | 745281 | -1 |
| YOR214C | 20.7181688 | -0.9774198 | 0.74545992 | 4.52314219 | 0.10418667 | 0.22564606 | SPR2 | XV | 745593 | 746303 | -1 |
| YOR215C | 11.3516725 | 0.13256508 | 0.92937277 | 0.03750069 | 0.98142435 | 0.98585131 | AIM41 | XV | 746725 | 747282 | -1 |
| YOR216C | 17.9759537 | -0.5644279 | 0.76605467 | 1.89040623 | 0.38860063 | 0.56060856 | RUD3 | XV | 747525 | 748979 | -1 |
| YOR217W | 31.7678766 | -0.2706675 | 0.68511417 | 1.71132191 | 0.42500219 | 0.59510585 | RFC1 | XV | 749301 | 751886 | 1 |
| YOR219C | 51.5677574 | -1.5785276 | 0.6270825 | 6.38027401 | 0.04116623 | 0.11377206 | STE13 | XV | 752214 | 755009 | -1 |
| YOR220W | 23.7660617 | 3.06219877 | 0.95381682 | 12.0795241 | 0.00238213 | 0.01274723 | RCN2 | XV | 755328 | 756125 | 1 |

| | | | | | | | | | | | |
|---|---|---|---|---|---|---|---|---|---|---|---|
| YOR221C | 26.1487751 | 1.86929708 | 0.82630014 | 5.33909458 | 0.06928358 | 0.16707703 | MCT1 | XV | 756476 | 757558 | -1 |
| YOR222W | 47.7326645 | 1.04458134 | 0.67152854 | 2.71464372 | 0.25734907 | 0.42958807 | ODC2 | XV | 758330 | 759253 | 1 |
| YOR223W | 170.241113 | -2.5634289 | 0.56428617 | 18.6655585 | 8.85E-05 | 0.00084395 | DSC3 | XV | 759782 | 760660 | 1 |
| YOR224C | 26.2863965 | -0.8809349 | 0.78125996 | 3.86969881 | 0.14444602 | 0.28495484 | RPB8 | XV | 760825 | 761265 | -1 |
| YOR226C | 25.5473901 | -0.4509636 | 0.70549886 | 0.6346967 | 0.72807708 | 0.83592431 | ISU2 | XV | 761614 | 762084 | -1 |
| YOR227W | 59.8373859 | 0.71016356 | 0.62706083 | 1.31443813 | 0.51829066 | 0.67466343 | HER1 | XV | 762825 | 766565 | 1 |
| YOR228C | 38.6345038 | 1.36128956 | 0.64935709 | 12.6880406 | 0.00175722 | 0.01017432 | MCP1 | XV | 766869 | 767777 | -1 |
| YOR229W | 10.7809174 | -1.2095592 | 0.93351258 | 1.73730867 | 0.4195157 | 0.59032562 | WTM2 | XV | 768409 | 769812 | 1 |
| YOR230W | 395.068118 | -1.0267952 | 0.51876811 | 7.36814153 | 0.02512051 | 0.07811371 | WTM1 | XV | 770800 | 772113 | 1 |
| YOR231W | 1030.94928 | 5.17029079 | 0.49898983 | 119.881288 | 9.29E-27 | 1.17E-23 | MKK1 | XV | 772601 | 774127 | 1 |
| YOR232W | 14.4879354 | -2.3016172 | 1.01665234 | 5.34224231 | 0.06917463 | 0.16688379 | MGE1 | XV | 774573 | 775259 | 1 |
| YOR233W | 80.167473 | 2.44786773 | 0.56350583 | 23.3042466 | 8.70E-06 | 0.00011634 | KIN4 | XV | 775846 | 778248 | 1 |
| YOR234C | 12.0094995 | -1.8283186 | 1.10528949 | 2.90523556 | 0.23395704 | 0.40148525 | RPL33B | XV | 778555 | 779405 | -1 |
| YOR235W | 11.4588188 | 2.92459438 | 1.27638227 | 5.61410393 | 0.06038274 | 0.15063165 | NA | XV | 779870 | 780184 | 1 |
| YOR236W | 196.569903 | 4.3773454 | 0.58810329 | 49.3317793 | 1.94E-11 | 1.07E-09 | DFR1 | XV | 780906 | 781541 | 1 |
| YOR237W | 18.218713 | 2.64559994 | 1.22912928 | 6.08453772 | 0.04772648 | 0.12600836 | HES1 | XV | 781994 | 783298 | 1 |
| YOR238W | 185.911328 | 2.98672438 | 0.5463644 | 27.8491641 | 8.97E-07 | 1.62E-05 | NA | XV | 783677 | 784588 | 1 |
| YOR239W | 87.8929727 | -0.1637091 | 0.7125674 | 0.51618439 | 0.772524 | 0.86150115 | ABP140 | XV | 784857 | 786744 | 1 |
| YOR241W | 21.9665429 | -1.2091745 | 0.81099402 | 3.38472097 | 0.18408448 | 0.33965875 | MET7 | XV | 786995 | 788641 | 1 |
| YOR242C | 381.65876 | -1.5366842 | 0.70690903 | 10.2464273 | 0.00595685 | 0.02634848 | SSP2 | XV | 788742 | 789857 | -1 |
| YOR243C | 58.4888433 | 0.32087014 | 0.6085573 | 4.40760236 | 0.11038278 | 0.23522866 | PUS7 | XV | 790211 | 792241 | -1 |
| YOR244W | 9.56326399 | -0.4258734 | 0.91064889 | 0.24410041 | 0.88510393 | 0.9315066 | ESA1 | XV | 792531 | 793868 | 1 |
| YOR245C | 18.4755076 | -0.2333937 | 0.82060826 | 0.126555 | 0.93868295 | 0.95995205 | DGA1 | XV | 794076 | 795332 | -1 |
| YOR246C | 112.788849 | 1.94944121 | 0.5396375 | 14.859082 | 0.00059346 | 0.00424214 | ENV9 | XV | 795801 | 796793 | -1 |
| YOR247W | 250.324165 | 1.17134959 | 0.51798309 | 7.26854877 | 0.02640309 | 0.08080014 | SRL1 | XV | 797676 | 798308 | 1 |
| YOR249C | 71.9222831 | -1.7220865 | 0.64613983 | 9.83384892 | 0.00732161 | 0.03110209 | APC5 | XV | 798674 | 800731 | -1 |
| YOR250C | 28.1231595 | 0.75052771 | 0.7669975 | 1.00063012 | 0.6063396 | 0.75088539 | CLP1 | XV | 800970 | 802307 | -1 |
| YOR251C | 36.3483662 | 0.6150873 | 0.64611248 | 1.01300249 | 0.60260024 | 0.74824264 | TUM1 | XV | 802550 | 803464 | -1 |
| YOR252W | 14.9084759 | -2.4126406 | 0.86049361 | 7.94474565 | 0.0188287 | 0.06320157 | TMA16 | XV | 803667 | 804203 | 1 |
| YOR253W | 318.265424 | -0.1259446 | 0.47281185 | 0.99936574 | 0.60672304 | 0.75094621 | NAT5 | XV | 804377 | 804907 | 1 |
| YOR254C | 70.5239589 | 0.01208346 | 0.69191337 | 0.01092097 | 0.9945544 | 0.99571207 | SEC63 | XV | 805032 | 807023 | -1 |
| YOR256C | 16.3779748 | 0.24321254 | 0.82106021 | 2.40303043 | 0.30073818 | 0.47389383 | TRE2 | XV | 808254 | 810683 | -1 |
| YOR257W | 24.6494025 | -0.4404989 | 0.74712398 | 0.45136026 | 0.79797331 | 0.8780436 | CDC31 | XV | 811008 | 811493 | 1 |

| | | | | | | | | | | | |
|---|---|---|---|---|---|---|---|---|---|---|---|
| YOR258W | 188.973747 | -0.3046118 | 0.70068073 | 10.4267891 | 0.00544317 | 0.02452601 | HNT3 | XV | 811671 | 812324 | 1 |
| YOR259C | 247.807576 | -0.2189407 | 0.56640106 | 5.25878263 | 0.07212235 | 0.17248591 | RPT4 | XV | 812395 | 813708 | -1 |
| YOR260W | 87.3072357 | -1.3268323 | 0.59620398 | 5.17013394 | 0.07539103 | 0.17809631 | GCD1 | XV | 813984 | 815720 | 1 |
| YOR261C | 132.746662 | -1.7259275 | 0.78212808 | 13.548602 | 0.00114277 | 0.00734365 | RPN8 | XV | 815915 | 816931 | -1 |
| YOR262W | 197.811163 | 1.34082092 | 0.51837058 | 7.45080627 | 0.02410338 | 0.0760123 | GPN2 | XV | 817292 | 818335 | 1 |
| YOR264W | 12.7254128 | 0.90910158 | 0.8872857 | 1.89044291 | 0.38859351 | 0.56060856 | DSE3 | XV | 818866 | 820158 | 1 |
| YOR265W | 8.76054654 | -1.1982921 | 1.01900037 | 1.67117319 | 0.43362005 | 0.60207676 | RBL2 | XV | 820454 | 820774 | 1 |
| YOR266W | 138.966954 | 2.69985194 | 0.5160621 | 29.881675 | 3.25E-07 | 6.64E-06 | PNT1 | XV | 821023 | 822294 | 1 |
| YOR267C | 332.18335 | 1.15609966 | 0.75422823 | 7.08094987 | 0.02899955 | 0.08672903 | HRK1 | XV | 822588 | 824867 | -1 |
| YOR268C | 20.8965296 | 1.17074701 | 0.70781186 | 5.08041632 | 0.07884998 | 0.18438668 | NA | XV | 825534 | 825932 | -1 |
| YOR269W | 16.8205995 | -1.9081083 | 0.78312405 | 5.83397135 | 0.05409651 | 0.13865373 | PAC1 | XV | 826385 | 827869 | 1 |
| YOR270C | 39.2635651 | -1.1373598 | 0.85703901 | 2.37336834 | 0.30523168 | 0.47807485 | VPH1 | XV | 828052 | 830574 | -1 |
| YOR271C | 44.1304952 | -1.7046165 | 0.61518008 | 8.14484705 | 0.01703605 | 0.05876231 | FSF1 | XV | 831059 | 832042 | -1 |
| YOR272W | 77.8258738 | -0.6810753 | 0.95952578 | 8.33359184 | 0.01550185 | 0.05493006 | YTM1 | XV | 832813 | 834195 | 1 |
| YOR273C | 79.80623 | 2.14429693 | 0.63667949 | 11.2397567 | 0.00362508 | 0.01780257 | TPO4 | XV | 834452 | 836431 | -1 |
| YOR274W | 494.837565 | 1.60006875 | 0.54876935 | 9.64921365 | 0.00802971 | 0.03330374 | MOD5 | XV | 837674 | 838960 | 1 |
| YOR275C | 95.9573223 | -1.1245325 | 0.61148441 | 3.35720548 | 0.18663457 | 0.34246214 | RIM20 | XV | 839084 | 841069 | -1 |
| YOR276W | 42.3178799 | -2.1736081 | 0.63548859 | 10.9961238 | 0.0040947 | 0.0195435 | CAF20 | XV | 841333 | 841818 | 1 |
| YOR278W | 90.4495369 | 0.16820173 | 0.67594359 | 1.92133905 | 0.38263662 | 0.55595132 | HEM4 | XV | 842817 | 843644 | 1 |
| YOR279C | 73.7903509 | 0.14982933 | 0.75636471 | 3.96469784 | 0.1377453 | 0.27549061 | RFM1 | XV | 843698 | 844630 | -1 |
| YOR280C | 104.466537 | 1.81546439 | 0.59868274 | 9.00986737 | 0.01105432 | 0.04244332 | FSH3 | XV | 844992 | 845792 | -1 |
| YOR281C | 17.4444953 | -0.7059059 | 0.7657584 | 2.85831193 | 0.23951099 | 0.40781483 | PLP2 | XV | 846269 | 847129 | -1 |
| YOR283W | 11.3482217 | -0.7167879 | 1.13493658 | 0.50652753 | 0.7762631 | 0.8639961 | NA | XV | 847453 | 848145 | 1 |
| YOR284W | 22.5444158 | -1.7660323 | 0.73063247 | 6.18867342 | 0.04530505 | 0.12155526 | HUA2 | XV | 848478 | 849209 | 1 |
| YOR285W | 85.4736968 | 0.43069896 | 0.57485333 | 0.55762178 | 0.75668298 | 0.8528864 | RDL1 | XV | 849635 | 850054 | 1 |
| YOR286W | 6.96077264 | -1.722768 | 1.11151578 | 3.40213661 | 0.18248847 | 0.3376824 | RDL2 | XV | 850280 | 850729 | 1 |
| YOR287C | 162.866501 | -2.3910158 | 0.76715726 | 13.331876 | 0.00127356 | 0.00800643 | RRP36 | XV | 850937 | 851839 | -1 |
| YOR288C | 29.9976978 | -1.5472874 | 0.75578428 | 4.24705879 | 0.11960874 | 0.24884462 | MPD1 | XV | 852121 | 853077 | -1 |
| YOR289W | 158.001314 | 1.26642415 | 0.57349902 | 4.76964316 | 0.09210541 | 0.20646161 | NA | XV | 853357 | 854112 | 1 |
| YOR290C | 52.2205755 | -0.0256276 | 0.6012121 | 1.19288758 | 0.5507668 | 0.70271921 | SNF2 | XV | 855147 | 860258 | -1 |
| YOR291W | 65.0876991 | 0.72244008 | 0.71328914 | 2.3345869 | 0.3112081 | 0.48393923 | YPK9 | XV | 861175 | 865593 | 1 |
| YOR292C | 115.549892 | -1.3437432 | 0.72992077 | 9.81284664 | 0.0073989 | 0.03135777 | NA | XV | 865653 | 866582 | -1 |
| YOR293C-A | 380.830429 | -2.9837602 | 0.99998416 | 15.8534856 | 0.00036096 | 0.00284348 | NA | XV | 867998 | 868147 | -1 |

| | | | | | | | | | | | |
|---|---|---|---|---|---|---|---|---|---|---|---|
| YOR293W | 239.156947 | -3.337524 | 0.56459542 | 30.385615 | 2.52E-07 | 5.35E-06 | RPS10A | XV | 867098 | 867852 | 1 |
| YOR294W | 58.6664118 | -2.8391982 | 0.78002935 | 12.7955128 | 0.00166529 | 0.00978886 | RRS1 | XV | 868340 | 868951 | 1 |
| YOR295W | 78.7493891 | 0.91007897 | 0.56188661 | 3.34675255 | 0.18761256 | 0.34351531 | UAF30 | XV | 869208 | 869894 | 1 |
| YOR296W | 55.8722376 | 0.53537971 | 0.56956175 | 3.03843137 | 0.21888349 | 0.38305402 | NA | XV | 870202 | 874071 | 1 |
| YOR297C | 112.907707 | -0.4048529 | 0.51425373 | 0.71761044 | 0.69851039 | 0.81736153 | TIM18 | XV | 874743 | 875321 | -1 |
| YOR298C-A | 243.404967 | 0.32959476 | 0.49941732 | 0.43757009 | 0.80349442 | 0.88210707 | MBF1 | XV | 877230 | 877685 | -1 |
| YOR298W | 1488.23207 | 0.16116928 | 0.80248574 | 2.06340401 | 0.35639985 | 0.52952403 | MUM3 | XV | 875599 | 877038 | 1 |
| YOR299W | 48.6533455 | -2.1371591 | 0.67987323 | 9.75635815 | 0.00761086 | 0.03195568 | BUD7 | XV | 878435 | 880675 | 1 |
| YOR301W | 241.745519 | 1.07333618 | 0.58478639 | 4.23787242 | 0.12015938 | 0.24944104 | RAX1 | XV | 880965 | 882272 | 1 |
| YOR302W | 11.7856481 | 1.71315642 | 1.00939387 | 2.99242231 | 0.22397717 | 0.38943778 | NA | XV | 882765 | 882842 | 1 |
| YOR303W | 70.9760742 | 1.70671086 | 0.61935482 | 7.38583168 | 0.02489929 | 0.07763431 | CPA1 | XV | 882899 | 884134 | 1 |
| YOR304C-A | 19.6012972 | -0.5586354 | 0.77591872 | 1.30715607 | 0.52018122 | 0.67636408 | BIL1 | XV | 888520 | 888750 | -1 |
| YOR304W | 55.2867461 | 0.48723104 | 0.63376347 | 0.61607526 | 0.73488766 | 0.83825839 | ISW2 | XV | 884514 | 887876 | 1 |
| YOR305W | 86.2857957 | 2.88449404 | 0.63505888 | 19.7467908 | 5.15E-05 | 0.00052711 | RRG7 | XV | 889022 | 889750 | 1 |
| YOR306C | 1233.74031 | -1.8636867 | 0.52788973 | 11.6939112 | 0.00288868 | 0.01477514 | MCH5 | XV | 889867 | 891432 | -1 |
| YOR307C | 77.8204442 | 2.7169635 | 0.59976112 | 24.8414885 | 4.03E-06 | 5.94E-05 | SLY41 | XV | 892731 | 894092 | -1 |
| YOR308C | 30.5567725 | -1.3269205 | 0.70702817 | 3.76824547 | 0.15196231 | 0.29555317 | SNU66 | XV | 894624 | 896387 | -1 |
| YOR310C | 575.962285 | -2.7438374 | 0.69520578 | 14.7777229 | 0.0006181 | 0.00440738 | NOP58 | XV | 896825 | 898360 | -1 |
| YOR311C | 33.3328283 | -1.4218401 | 0.71415073 | 4.0341632 | 0.13304317 | 0.26909961 | DGK1 | XV | 899056 | 899928 | -1 |
| YOR312C | 50.7697352 | -3.1469427 | 0.68546952 | 20.3637945 | 3.78E-05 | 0.00040962 | NA | XV | 900250 | 900767 | -1 |
| YOR314W | 33.5880509 | -2.2809196 | 0.6684391 | 14.9020271 | 0.00058085 | 0.00417263 | NA | XV | 903043 | 903372 | 1 |
| YOR315W | 26.3145683 | -0.0779199 | 0.66558924 | 3.36171565 | 0.18621417 | 0.34184529 | SFG1 | XV | 904760 | 905800 | 1 |
| YOR316C | 36.4264501 | 0.31885119 | 0.74271525 | 3.24616053 | 0.19729006 | 0.35541675 | COT1 | XV | 906236 | 907555 | -1 |
| YOR317W | 347.076466 | -1.9819536 | 0.50001223 | 26.1820149 | 2.06E-06 | 3.32E-05 | FAA1 | XV | 909343 | 911445 | 1 |
| YOR319W | 122.582243 | -2.8316857 | 0.59251135 | 21.1998998 | 2.49E-05 | 0.00028739 | HSH49 | XV | 912822 | 913463 | 1 |
| YOR320C | 318.903594 | -1.2201861 | 0.47356529 | 6.73988442 | 0.03439162 | 0.09926596 | GNT1 | XV | 913619 | 915094 | -1 |
| YOR321W | 107.901295 | -0.8962578 | 0.58809161 | 2.72812802 | 0.25561982 | 0.42788053 | PMT3 | XV | 916030 | 918291 | 1 |
| YOR322C | 233.869053 | 4.20897597 | 0.526271 | 80.5985127 | 3.15E-18 | 5.97E-16 | LDB19 | XV | 918606 | 921062 | -1 |
| YOR323C | 86.0330806 | -0.4514934 | 0.74071973 | 0.38438885 | 0.82514642 | 0.89541781 | PRO2 | XV | 921535 | 922905 | -1 |
| YOR324C | 15.6933135 | -0.0695326 | 0.75052785 | 1.64170385 | 0.4400566 | 0.60824247 | FRT1 | XV | 923232 | 925040 | -1 |
| YOR326W | 110.890262 | 1.58109856 | 0.53527583 | 8.82108914 | 0.01214856 | 0.04558315 | MYO2 | XV | 925721 | 930445 | 1 |
| YOR327C | 123.911574 | -1.07761 | 0.5201315 | 5.3993884 | 0.06722607 | 0.16286148 | SNC2 | XV | 930734 | 931081 | -1 |
| YOR328W | 108.330122 | -1.553644 | 0.57458719 | 7.10075765 | 0.02871376 | 0.08612855 | PDR10 | XV | 931803 | 936497 | 1 |

| | | | | | | | | | | | |
|---|---|---|---|---|---|---|---|---|---|---|---|
| YOR329C | 104.770624 | 0.76473709 | 0.60171452 | 2.67378266 | 0.26266093 | 0.43402019 | SCD5 | XV | 936731 | 939349 | -1 |
| YOR330C | 294.861232 | 4.16277137 | 0.53714072 | 54.2419324 | 1.67E-12 | 1.14E-10 | MIP1 | XV | 939621 | 943385 | -1 |
| YOR332W | 151.369254 | -0.7572987 | 0.54126241 | 4.65524321 | 0.09752743 | 0.21503573 | VMA4 | XV | 943656 | 944357 | 1 |
| YOR334W | 40.8388899 | 0.78956383 | 0.62092008 | 2.12654714 | 0.34532352 | 0.51852261 | MRS2 | XV | 944596 | 946008 | 1 |
| YOR335C | 39.9192451 | -2.2610893 | 0.82818206 | 7.49727574 | 0.0235498 | 0.07474302 | NA | XV | 946233 | 949109 | -1 |
| YOR336W | 59.3633252 | -0.7756252 | 0.59478134 | 1.74555503 | 0.41778952 | 0.58926319 | KRE5 | XV | 949773 | 953870 | 1 |
| YOR337W | 20.7307779 | 1.4041546 | 0.85445402 | 3.50139436 | 0.17365283 | 0.32507061 | TEA1 | XV | 954344 | 956623 | 1 |
| YOR338W | 18.4152278 | 1.22521524 | 0.80979054 | 2.52488256 | 0.28296239 | 0.45678444 | NA | XV | 956898 | 957989 | 1 |
| YOR339C | 178.820893 | 2.84417904 | 0.55074716 | 31.0223664 | 1.83E-07 | 4.09E-06 | UBC11 | XV | 958362 | 958832 | -1 |
| YOR340C | 57.8572901 | -1.7191991 | 0.58125008 | 9.02154112 | 0.01098999 | 0.04228042 | RPA43 | XV | 959202 | 960182 | -1 |
| YOR341W | 58.5309308 | -1.4710514 | 0.72357309 | 5.423599 | 0.06641718 | 0.16169345 | RPA190 | XV | 960987 | 965981 | 1 |
| YOR342C | 38.1283567 | -0.560241 | 0.71060758 | 2.07057768 | 0.35512379 | 0.52857757 | NA | XV | 966666 | 967625 | -1 |
| YOR343C | 24.6685287 | -1.2271261 | 0.74038799 | 3.07749504 | 0.21464978 | 0.37844769 | NA | XV | 968148 | 968474 | -1 |
| YOR343W-B | 13.6096358 | 0.2147465 | 0.87832606 | 0.89944489 | 0.63780515 | 0.77419116 | NA | XV | 970578 | 975891 | 1 |
| YOR344C | 75.0057882 | 2.46640079 | 0.56688803 | 24.0597024 | 5.96E-06 | 8.30E-05 | TYE7 | XV | 977194 | 978069 | -1 |
| YOR346W | 1060.24576 | -0.1272374 | 0.47950654 | 7.24044987 | 0.02677665 | 0.0815554 | REV1 | XV | 981828 | 984785 | 1 |
| YOR347C | 33.9033217 | -0.896723 | 0.70356039 | 2.39408135 | 0.30208686 | 0.4745206 | PYK2 | XV | 984942 | 986462 | -1 |
| YOR348C | 62.7778027 | 0.22832672 | 0.55302346 | 0.18056917 | 0.91367113 | 0.94703828 | PUT4 | XV | 986899 | 988782 | -1 |
| YOR349W | 758.857754 | -2.2969219 | 0.45586028 | 23.4208758 | 8.21E-06 | 0.00011052 | CIN1 | XV | 989789 | 992833 | 1 |
| YOR350C | 88.1724224 | -2.0877188 | 0.62663349 | 10.801742 | 0.00451265 | 0.02098654 | MNE1 | XV | 992864 | 994855 | -1 |
| YOR351C | 207.598597 | -1.411037 | 0.5383893 | 6.96713931 | 0.03069764 | 0.09096178 | MEK1 | XV | 995018 | 996511 | -1 |
| YOR352W | 13.9161607 | 0.01880514 | 0.78818487 | 0.09389455 | 0.9541377 | 0.96767513 | TFB6 | XV | 997213 | 998244 | 1 |
| YOR353C | 23.1342607 | -0.3962254 | 0.82741767 | 0.54282172 | 0.76230323 | 0.85554094 | SOG2 | XV | 998453 | 1E+06 | -1 |
| YOR354C | 7.27177081 | -0.4048756 | 1.06272379 | 0.87780975 | 0.64474211 | 0.77739625 | MSC6 | XV | 1001147 | 1E+06 | -1 |
| YOR355W | 51.322971 | -0.8690163 | 0.62255859 | 8.09194629 | 0.01749267 | 0.05989508 | GDS1 | XV | 1005137 | 1E+06 | 1 |
| YOR356W | 116.45056 | 2.04772261 | 0.59917233 | 12.9454597 | 0.001545 | 0.00934751 | CIR2 | XV | 1007221 | 1E+06 | 1 |
| YOR357C | 294.924605 | 0.50566682 | 1.63537793 | 4.30015413 | 0.11647518 | 0.24407937 | SNX3 | XV | 1009224 | 1E+06 | -1 |
| YOR358W | 17.2642865 | -0.2750434 | 0.77894664 | 0.24512999 | 0.88464841 | 0.9315066 | HAP5 | XV | 1010161 | 1E+06 | 1 |
| YOR359W | 222.713578 | 4.73863775 | 0.5688996 | 79.8053352 | 4.68E-18 | 8.22E-16 | VTS1 | XV | 1011189 | 1E+06 | 1 |
| YOR360C | 6.43066135 | -0.6298504 | 1.22073455 | 0.39793314 | 0.81957729 | 0.89171017 | PDE2 | XV | 1013241 | 1E+06 | -1 |
| YOR361C | 289.58826 | 3.13338928 | 0.63535683 | 37.2423066 | 8.18E-09 | 2.52E-07 | PRT1 | XV | 1015361 | 1E+06 | -1 |
| YOR362C | 40.3953689 | 0.33909006 | 0.62478825 | 0.31499744 | 0.85427791 | 0.91360714 | PRE10 | XV | 1017880 | 1E+06 | -1 |
| YOR363C | 29.6052992 | 1.08678264 | 0.64941519 | 3.59329389 | 0.16585407 | 0.31516084 | PIP2 | XV | 1020222 | 1E+06 | -1 |

| | | | | | | | | | | | |
|---|---|---|---|---|---|---|---|---|---|---|---|
| YOR365C | 802.435808 | 0.68328568 | 0.5027339 | 2.00765264 | 0.36647451 | 0.53971939 | NA | XV | 1023461 | 1E+06 | -1 |
| YOR367W | 32.7706104 | -1.0641933 | 0.63419974 | 3.70404022 | 0.15691985 | 0.30265354 | SCP1 | XV | 1026007 | 1E+06 | 1 |
| YOR368W | 465.952288 | -2.4612938 | 0.48096139 | 23.839668 | 6.66E-06 | 9.16E-05 | RAD17 | XV | 1026843 | 1E+06 | 1 |
| YOR369C | 127.98923 | -3.2712626 | 0.84521622 | 13.0682237 | 0.00145302 | 0.00892151 | RPS12 | XV | 1028194 | 1E+06 | -1 |
| YOR370C | 22.1651221 | 0.41959587 | 0.7719438 | 0.34260887 | 0.84256503 | 0.90781117 | MRS6 | XV | 1029183 | 1E+06 | -1 |
| YOR371C | 128.10914 | -2.0982001 | 0.52843891 | 15.9403662 | 0.00034562 | 0.00274125 | GPB1 | XV | 1031489 | 1E+06 | -1 |
| YOR372C | 55.273807 | 1.32774848 | 0.58359798 | 5.86874346 | 0.05316411 | 0.13694173 | NDD1 | XV | 1034807 | 1E+06 | -1 |
| YOR373W | 51.6882843 | 0.1218721 | 0.60087176 | 0.51624752 | 0.77249962 | 0.86150115 | NUD1 | XV | 1036834 | 1E+06 | 1 |
| YOR374W | 84.471092 | -3.1677022 | 0.57157923 | 39.8132823 | 2.26E-09 | 7.75E-08 | ALD4 | XV | 1039840 | 1E+06 | 1 |
| YOR375C | 46.2371931 | -3.2164726 | 0.89856348 | 12.4598397 | 0.00196961 | 0.01104002 | GDH1 | XV | 1041678 | 1E+06 | -1 |
| YOR376W | 73.1900499 | 0.35202945 | 0.55465726 | 1.64300973 | 0.43976936 | 0.60823052 | NA | XV | 1043191 | 1E+06 | 1 |
| YOR376W-A | 35.65045 | -1.2056368 | 0.6631787 | 4.00025915 | 0.13531775 | 0.27204506 | NA | XV | 1045196 | 1E+06 | 1 |
| YOR377W | 108.561375 | -2.1648175 | 0.61136655 | 13.7262367 | 0.00104565 | 0.0068333 | ATF1 | XV | 1046226 | 1E+06 | 1 |
| YOR378W | 722.839546 | 0.59107041 | 0.52141965 | 3.37091733 | 0.18535939 | 0.34124989 | AMF1 | XV | 1049511 | 1E+06 | 1 |
| YOR380W | 93.8767858 | 0.11882473 | 0.52983318 | 6.19883008 | 0.04507556 | 0.12108273 | RDR1 | XV | 1051290 | 1E+06 | 1 |
| YOR381W | 38.4545213 | -0.6628915 | 0.67207394 | 4.22509366 | 0.12092959 | 0.25060211 | FRE3 | XV | 1055545 | 1E+06 | 1 |
| YOR382W | 41.9240109 | -0.2394417 | 0.62007715 | 0.20619581 | 0.90203865 | 0.93824416 | FIT2 | XV | 1059531 | 1E+06 | 1 |
| YOR384W | 3179.00303 | 0.65962702 | 0.48183295 | 1.91085396 | 0.38464787 | 0.55747464 | FRE5 | XV | 1061564 | 1E+06 | 1 |
| YOR385W | 25.4800499 | 0.26301692 | 0.7303865 | 3.29559466 | 0.1924734 | 0.34902003 | NA | XV | 1065043 | 1E+06 | 1 |
| YOR386W | 13.2884771 | -0.5723978 | 0.81784363 | 0.63629785 | 0.72749444 | 0.83558675 | PHR1 | XV | 1066839 | 1E+06 | 1 |
| YOR387C | 376.852227 | 0.1553302 | 0.50635632 | 0.09395782 | 0.95410752 | 0.96767513 | NA | XV | 1069621 | 1E+06 | -1 |
| YOR388C | 41.9995532 | -0.4986678 | 0.74611654 | 0.45420016 | 0.79684103 | 0.8771311 | FDH1 | XV | 1071793 | 1E+06 | -1 |
| YOR389W | 259.975014 | 1.8159356 | 0.47839223 | 17.4915881 | 0.00015913 | 0.00140024 | NA | XV | 1074213 | 1E+06 | 1 |
| YOR390W | 20.3428834 | 1.49282368 | 0.82229881 | 3.98832565 | 0.13612757 | 0.27319883 | FEX1 | XV | 1076784 | 1E+06 | 1 |
| YOR391C | 7.16797393 | 0.70100523 | 1.07873537 | 1.06769645 | 0.58634424 | 0.73388092 | HSP33 | XV | 1078545 | 1E+06 | -1 |
| YPL001W | 39.8364639 | 2.30119512 | 0.69231303 | 10.8667777 | 0.00436827 | 0.02052944 | HAT1 | XVI | 554605 | 555729 | 1 |
| YPL002C | 480.214076 | 2.15657403 | 0.66692924 | 9.76857287 | 0.00756452 | 0.03183036 | SNF8 | XVI | 553627 | 554328 | -1 |
| YPL003W | 962.45856 | 0.48756657 | 0.56494323 | 2.21491676 | 0.33039764 | 0.50355418 | ULA1 | XVI | 552020 | 553408 | 1 |
| YPL004C | 63.0880644 | -0.5991163 | 0.5971488 | 2.96074017 | 0.22755346 | 0.39362867 | LSP1 | XVI | 550632 | 551657 | -1 |
| YPL005W | 40.8617127 | 0.46161741 | 0.61533589 | 1.93540351 | 0.37995527 | 0.55386228 | AEP3 | XVI | 548486 | 550306 | 1 |
| YPL006W | 623.098925 | 4.04646037 | 0.49952207 | 66.1472076 | 4.33E-15 | 4.56E-13 | NCR1 | XVI | 544631 | 548143 | 1 |
| YPL007C | 25.5416857 | 0.28678218 | 0.70314821 | 2.17462906 | 0.33712061 | 0.51030806 | TFC8 | XVI | 542082 | 543848 | -1 |
| YPL008W | 124.553814 | 2.29169859 | 0.571824 | 21.4140883 | 2.24E-05 | 0.00026028 | CHL1 | XVI | 539385 | 541970 | 1 |

| | | | | | | | | | | | |
|---|---|---|---|---|---|---|---|---|---|---|---|
| YPL009C | 272.474234 | 0.58865933 | 0.79436605 | 1.82564716 | 0.40138927 | 0.57214275 | RQC2 | XVI | 535820 | 538936 | -1 |
| YPL010W | 9.02302525 | -0.8914256 | 1.02501411 | 0.88497912 | 0.64243505 | 0.77621415 | RET3 | XVI | 535018 | 535587 | 1 |
| YPL011C | 13.1722737 | -0.8403942 | 0.89985934 | 1.11384272 | 0.57297032 | 0.72277589 | TAF3 | XVI | 533642 | 534703 | -1 |
| YPL012W | 34.8792974 | -1.7868183 | 0.71696325 | 7.40961616 | 0.02460494 | 0.07694787 | RRP12 | XVI | 529723 | 533409 | 1 |
| YPL013C | 19.0866289 | -0.4812012 | 0.78866369 | 0.52216856 | 0.770216 | 0.86008691 | MRPS16 | XVI | 528985 | 529350 | -1 |
| YPL014W | 78.8043128 | -1.5668922 | 0.54381602 | 8.17193965 | 0.01680683 | 0.05813116 | CIP1 | XVI | 527547 | 528692 | 1 |
| YPL015C | 69.1421886 | 1.24479201 | 0.54517711 | 5.9313571 | 0.05152549 | 0.13385547 | HST2 | XVI | 525810 | 526883 | -1 |
| YPL016W | 39.3259789 | 0.31902412 | 0.62910106 | 0.85209487 | 0.65308536 | 0.78337772 | SWI1 | XVI | 521014 | 524958 | 1 |
| YPL017C | 979.03059 | -0.2165998 | 0.47095318 | 0.26535334 | 0.8757482 | 0.92712351 | IRC15 | XVI | 518734 | 520233 | -1 |
| YPL018W | 1127.89283 | 3.00735504 | 0.55463224 | 28.8905979 | 5.33E-07 | 1.03E-05 | CTF19 | XVI | 517651 | 518760 | 1 |
| YPL019C | 244.541202 | 1.79878781 | 0.48311102 | 15.7207094 | 0.00038574 | 0.00298585 | VTC3 | XVI | 514511 | 517018 | -1 |
| YPL020C | 21.263298 | -0.1736614 | 0.72902414 | 0.61767272 | 0.73430092 | 0.83775415 | ULP1 | XVI | 512313 | 514178 | -1 |
| YPL021W | 8.32123492 | 6.50597024 | 1.69206749 | 19.101024 | 7.12E-05 | 0.00070435 | ECM23 | XVI | 511101 | 511664 | 1 |
| YPL022W | 27.3856249 | 0.2859495 | 0.69946741 | 0.2446333 | 0.88486813 | 0.9315066 | RAD1 | XVI | 506697 | 509999 | 1 |
| YPL023C | 127.830725 | -0.40265 | 0.68834915 | 5.97956542 | 0.05029836 | 0.13124269 | MET12 | XVI | 504339 | 506312 | -1 |
| YPL024W | 315.363001 | -0.738353 | 0.61652953 | 5.30424528 | 0.07050141 | 0.16937889 | RMI1 | XVI | 503517 | 504242 | 1 |
| YPL026C | 183.38368 | 0.6159367 | 0.81843058 | 2.66480448 | 0.26384269 | 0.43572423 | SKS1 | XVI | 500675 | 502183 | -1 |
| YPL027W | 34.8098475 | -0.1228752 | 0.65436979 | 5.84537775 | 0.05378886 | 0.13823236 | SMA1 | XVI | 499665 | 500402 | 1 |
| YPL028W | 74.7152412 | 0.48054855 | 0.72911723 | 0.45082049 | 0.7981887 | 0.87811373 | ERG10 | XVI | 498096 | 499292 | 1 |
| YPL029W | 15.0239213 | 1.13117341 | 0.79402579 | 3.17733354 | 0.20419767 | 0.3649088 | SUV3 | XVI | 495506 | 497719 | 1 |
| YPL030W | 8.09327774 | -0.4899154 | 0.98770268 | 0.27248868 | 0.87262939 | 0.92527295 | TRM44 | XVI | 493543 | 495246 | 1 |
| YPL031C | 244.18573 | -3.0946389 | 0.58831477 | 27.6885378 | 9.72E-07 | 1.73E-05 | PHO85 | XVI | 492018 | 493037 | -1 |
| YPL032C | 65.9095271 | -0.0887485 | 0.64321251 | 2.93150283 | 0.23090442 | 0.3978978 | SVL3 | XVI | 488887 | 491364 | -1 |
| YPL033C | 105.743499 | 3.88344716 | 0.6814492 | 33.2044107 | 6.16E-08 | 1.56E-06 | SRL4 | XVI | 487362 | 488207 | -1 |
| YPL034W | 17.1663606 | 2.4716564 | 0.82643761 | 10.1087207 | 0.00638145 | 0.02778089 | NA | XVI | 486712 | 487209 | 1 |
| YPL036W | 328.364258 | 1.08438154 | 0.60280719 | 4.24438087 | 0.119769 | 0.24890972 | PMA2 | XVI | 482843 | 485686 | 1 |
| YPL037C | 36.9572784 | -2.4495114 | 0.76842021 | 9.8701957 | 0.00718976 | 0.03067501 | EGD1 | XVI | 481428 | 481901 | -1 |
| YPL038W | 27.4989747 | -0.2754149 | 0.67427984 | 0.30469444 | 0.85869007 | 0.91650698 | MET31 | XVI | 480535 | 481068 | 1 |
| YPL039W | 31.9558132 | 0.45883328 | 0.72243794 | 0.41676505 | 0.81189641 | 0.88731158 | NA | XVI | 479225 | 480175 | 1 |
| YPL040C | 15.5684484 | 0.73165872 | 0.82336801 | 1.03072869 | 0.59728294 | 0.74451415 | ISM1 | XVI | 475993 | 479001 | -1 |
| YPL041C | 9.95484099 | -1.5317226 | 1.02267456 | 3.10000623 | 0.21224731 | 0.37592901 | MRX11 | XVI | 475118 | 475741 | -1 |
| YPL042C | 22.1754736 | 1.27906753 | 0.7143721 | 3.3409666 | 0.18815611 | 0.34377528 | SSN3 | XVI | 473040 | 474707 | -1 |
| YPL043W | 257.359002 | 0.34098689 | 0.56880527 | 4.11387037 | 0.12784519 | 0.26147074 | NOP4 | XVI | 469939 | 471996 | 1 |

| | | | | | | | | | | | |
|---|---|---|---|---|---|---|---|---|---|---|---|
| YPL045W | 71.8395613 | -0.0422285 | 0.5313622 | 0.27294112 | 0.872432 | 0.92527295 | VPS16 | XVI | 467260 | 469656 | 1 |
| YPL046C | 83.8585488 | -1.0749763 | 0.6899533 | 5.08363203 | 0.07872331 | 0.18416483 | ELC1 | XVI | 466644 | 466943 | -1 |
| YPL047W | 10.2339223 | -1.908999 | 0.93629075 | 4.12704761 | 0.12700564 | 0.26058208 | SGF11 | XVI | 465962 | 466261 | 1 |
| YPL048W | 50.3629168 | -0.2762857 | 0.722483 | 0.23149807 | 0.89069873 | 0.93317728 | CAM1 | XVI | 464401 | 465648 | 1 |
| YPL049C | 126.681952 | 1.61282318 | 0.55332461 | 9.01596063 | 0.0110207 | 0.04237041 | DIG1 | XVI | 462481 | 463839 | -1 |
| YPL050C | 56.4463265 | 1.25656702 | 0.65313395 | 6.34622041 | 0.04187316 | 0.11493835 | MNN9 | XVI | 460779 | 461966 | -1 |
| YPL051W | 13.1748276 | 0.05681773 | 0.92030071 | 0.18734868 | 0.91057926 | 0.94518714 | ARL3 | XVI | 459963 | 460559 | 1 |
| YPL052W | 137.485912 | 0.75051305 | 0.5334202 | 2.45570972 | 0.29292026 | 0.46721992 | OAZ1 | XVI | 458799 | 459678 | 1 |
| YPL053C | 27.9643379 | 1.01900542 | 0.81853196 | 2.83521528 | 0.24229298 | 0.41140068 | KTR6 | XVI | 457118 | 458458 | -1 |
| YPL054W | 189.300702 | 0.45301177 | 0.52352242 | 1.51560911 | 0.46869429 | 0.63146966 | LEE1 | XVI | 455759 | 456664 | 1 |
| YPL055C | 164.646292 | 0.80376919 | 0.49692664 | 7.60984637 | 0.02226091 | 0.07177998 | LGE1 | XVI | 453992 | 454990 | -1 |
| YPL056C | 313.006661 | 1.63582005 | 0.4774688 | 11.6482708 | 0.00295536 | 0.01506296 | LCL1 | XVI | 453430 | 453735 | -1 |
| YPL057C | 16.9818611 | -0.9115084 | 0.78170467 | 1.42335963 | 0.49081902 | 0.65209778 | SUR1 | XVI | 451909 | 453057 | -1 |
| YPL058C | 38.1879325 | 0.54651144 | 0.63676146 | 1.0089982 | 0.60380795 | 0.74934389 | PDR12 | XVI | 445842 | 450377 | -1 |
| YPL059W | 42.8501553 | 0.37750132 | 0.70871732 | 1.54195058 | 0.46256172 | 0.62590146 | GRX5 | XVI | 444579 | 445031 | 1 |
| YPL060C-A | 12.1185018 | -0.2502382 | 0.9014831 | 0.53144476 | 0.76665193 | 0.85825884 | NA | XVI | 437333 | 442742 | -1 |
| YPL060W | 18.9381274 | 0.5552035 | 0.74986476 | 0.57616003 | 0.7497016 | 0.8479727 | MFM1 | XVI | 434523 | 435764 | 1 |
| YPL061W | 71.694741 | -1.5623365 | 0.6281818 | 7.87011772 | 0.01954455 | 0.06499882 | ALD6 | XVI | 432588 | 434090 | 1 |
| YPL063W | 54.8978124 | -0.8210243 | 0.61151896 | 3.05873509 | 0.21667266 | 0.38085449 | TIM50 | XVI | 429939 | 431369 | 1 |
| YPL064C | 50.4793807 | 0.09380651 | 0.88297872 | 5.09780291 | 0.07816749 | 0.18338253 | CWC27 | XVI | 428711 | 429616 | -1 |
| YPL065W | 83.9920713 | -0.3932514 | 0.75496578 | 5.7609506 | 0.05610809 | 0.14279817 | VPS28 | XVI | 427927 | 428655 | 1 |
| YPL066W | 116.261524 | 0.49650816 | 0.5439085 | 2.87125595 | 0.23796588 | 0.40595829 | RGL1 | XVI | 426233 | 427672 | 1 |
| YPL067C | 92.3994418 | 1.55715558 | 0.53621975 | 10.0257389 | 0.00665179 | 0.02880618 | NA | XVI | 425251 | 425847 | -1 |
| YPL068C | 13.7410094 | -2.288147 | 1.02489798 | 5.19809214 | 0.07434446 | 0.17629168 | NA | XVI | 424215 | 425096 | -1 |
| YPL069C | 259.067028 | 0.81381855 | 0.53178411 | 12.0120734 | 0.00246383 | 0.01306373 | BTS1 | XVI | 422885 | 423892 | -1 |
| YPL070W | 122.815856 | -1.0265395 | 0.61401697 | 8.9265848 | 0.01152436 | 0.04384102 | MUK1 | XVI | 420948 | 422786 | 1 |
| YPL071C | 23.1309044 | -0.5423701 | 0.74092761 | 2.87554199 | 0.23745646 | 0.4053281 | NA | XVI | 420048 | 420518 | -1 |
| YPL072W | 216.458191 | 1.14416383 | 0.73378221 | 6.85153161 | 0.03252436 | 0.09506111 | UBP16 | XVI | 418512 | 420011 | 1 |
| YPL074W | 72.7913575 | 1.28135286 | 0.56565043 | 7.47466104 | 0.0238176 | 0.07531616 | YTA6 | XVI | 415763 | 418027 | 1 |
| YPL075W | 58.3034661 | 1.44707995 | 0.62799756 | 11.2061682 | 0.00368648 | 0.01802762 | GCR1 | XVI | 412254 | 415362 | 1 |
| YPL076W | 150.082019 | 2.76778807 | 0.59854644 | 20.3754236 | 3.76E-05 | 0.00040878 | GPI2 | XVI | 410443 | 411285 | 1 |
| YPL077C | 26.1346564 | 1.41451052 | 0.73324584 | 4.36257592 | 0.11289603 | 0.23943884 | NA | XVI | 409368 | 410090 | -1 |
| YPL078C | 38.0277542 | 0.77789523 | 0.75645336 | 2.72622846 | 0.25586272 | 0.42803963 | ATP4 | XVI | 408010 | 408744 | -1 |

| | | | | | | | | | | | |
|---|---|---|---|---|---|---|---|---|---|---|---|
| YPL079W | 38.5673722 | -3.1019314 | 0.79290856 | 14.5001281 | 0.00071013 | 0.00493003 | RPL21B | XVI | 406636 | 407539 | 1 |
| YPL080C | 557.027376 | -2.7779728 | 0.62447094 | 28.1951653 | 7.54E-07 | 1.40E-05 | NA | XVI | 406172 | 406498 | -1 |
| YPL081W | 46.1188934 | -3.4888565 | 0.90771765 | 13.7141136 | 0.00105201 | 0.00686709 | RPS9A | XVI | 404950 | 406044 | 1 |
| YPL082C | 20.7410377 | -1.439975 | 0.73365252 | 3.90539782 | 0.1418906 | 0.28114371 | MOT1 | XVI | 398480 | 404083 | -1 |
| YPL083C | 55.8367687 | -0.7599828 | 0.91853799 | 4.43851446 | 0.10868981 | 0.23264843 | SEN54 | XVI | 396702 | 398105 | -1 |
| YPL084W | 458.497616 | -2.9605128 | 0.50095434 | 31.5457724 | 1.41E-07 | 3.26E-06 | BRO1 | XVI | 394038 | 396572 | 1 |
| YPL085W | 79.9950468 | -0.302318 | 0.5407224 | 1.5303399 | 0.46525485 | 0.62778504 | SEC16 | XVI | 387067 | 393654 | 1 |
| YPL086C | 19.0850517 | -2.0063026 | 0.81946012 | 6.46665889 | 0.03942601 | 0.11038521 | ELP3 | XVI | 384773 | 386446 | -1 |
| YPL087W | 14.6098457 | 1.00787653 | 1.14421263 | 0.90610901 | 0.63568349 | 0.7722634 | YDC1 | XVI | 383455 | 384408 | 1 |
| YPL088W | 66.7766802 | 3.66302555 | 0.69018948 | 28.5904699 | 6.19E-07 | 1.17E-05 | NA | XVI | 381965 | 382993 | 1 |
| YPL089C | 596.910242 | -1.7701608 | 0.71135211 | 6.01490848 | 0.04941732 | 0.12952753 | RLM1 | XVI | 379120 | 381150 | -1 |
| YPL090C | 58.8007764 | -3.6928557 | 0.93070508 | 13.9270425 | 0.00094576 | 0.0062942 | RPS6A | XVI | 377291 | 378395 | -1 |
| YPL091W | 133.376952 | 0.25388619 | 0.52757145 | 0.54678951 | 0.7607924 | 0.85483951 | GLR1 | XVI | 375502 | 376953 | 1 |
| YPL092W | 19.5299865 | 0.21460434 | 0.73770923 | 2.32593153 | 0.31255783 | 0.4849178 | SSU1 | XVI | 373793 | 375169 | 1 |
| YPL093W | 48.3463212 | -2.7130619 | 0.71976338 | 14.0664303 | 0.00088209 | 0.00595257 | NOG1 | XVI | 370978 | 372921 | 1 |
| YPL094C | 88.8940085 | 0.162566 | 0.57254118 | 1.22294546 | 0.54255125 | 0.69630051 | SEC62 | XVI | 369839 | 370663 | -1 |
| YPL095C | 434.005486 | 0.24121945 | 0.50421949 | 2.21594712 | 0.33022747 | 0.50342734 | EEB1 | XVI | 368071 | 369441 | -1 |
| YPL096C-A | 31.2846611 | 3.38251165 | 2.32710765 | 4.68071113 | 0.09629339 | 0.2132895 | ERI1 | XVI | 366529 | 366735 | -1 |
| YPL096W | 71.0354076 | 0.28305248 | 0.8162894 | 0.87670551 | 0.64509818 | 0.77750176 | PNG1 | XVI | 366927 | 368018 | 1 |
| YPL097W | 371.247 | 1.23186713 | 0.97134796 | 7.59451493 | 0.02243221 | 0.07215694 | MSY1 | XVI | 364952 | 366430 | 1 |
| YPL098C | 13.9549594 | -0.5510853 | 0.81716557 | 0.60610456 | 0.73856048 | 0.84040603 | MGR2 | XVI | 364386 | 364727 | -1 |
| YPL099C | 159.247904 | -0.4689833 | 0.51536769 | 0.82303683 | 0.66264332 | 0.79009572 | INA17 | XVI | 363520 | 364068 | -1 |
| YPL100W | 157.301126 | -0.2171666 | 0.76051226 | 4.47472658 | 0.10673957 | 0.22957732 | ATG21 | XVI | 361872 | 363362 | 1 |
| YPL101W | 23.2508073 | -0.7242223 | 0.69607535 | 1.07561778 | 0.58402652 | 0.7320878 | ELP4 | XVI | 360209 | 361579 | 1 |
| YPL103C | 27.2967526 | 0.98806153 | 0.74907666 | 3.92893625 | 0.14023045 | 0.27872788 | FMP30 | XVI | 358000 | 359406 | -1 |
| YPL104W | 53.6088984 | -2.1398213 | 0.61815938 | 11.4473097 | 0.00326775 | 0.01635285 | MSD1 | XVI | 355700 | 357676 | 1 |
| YPL105C | 121.539729 | -0.1379263 | 0.56240267 | 0.06969875 | 0.96575087 | 0.97467275 | SYH1 | XVI | 352863 | 355412 | -1 |
| YPL106C | 185.594199 | 0.27330171 | 0.63488453 | 0.33532661 | 0.84563851 | 0.90923806 | SSE1 | XVI | 350194 | 352275 | -1 |
| YPL107W | 15.7866973 | 0.53191888 | 0.82426714 | 1.2221154 | 0.54277647 | 0.69636068 | NA | XVI | 349119 | 349865 | 1 |
| YPL108W | 46.960818 | -1.2778701 | 0.62204352 | 4.3043658 | 0.11623016 | 0.24370198 | NA | XVI | 348446 | 348952 | 1 |
| YPL109C | 604.813862 | 3.88073374 | 0.49102789 | 53.9545242 | 1.92E-12 | 1.29E-10 | NA | XVI | 345264 | 347389 | -1 |
| YPL110C | 175.582545 | 0.08308618 | 0.58308504 | 1.56864827 | 0.45642808 | 0.62207609 | GDE1 | XVI | 341068 | 344739 | -1 |
| YPL111W | 68.3600467 | 0.13331548 | 0.61373289 | 1.85854235 | 0.39484138 | 0.56643498 | CAR1 | XVI | 339944 | 340945 | 1 |

| | | | | | | | | | | | |
|---|---|---|---|---|---|---|---|---|---|---|---|
| YPL112C | 42.0847528 | 0.65100731 | 0.73976616 | 3.79529466 | 0.14992092 | 0.29275789 | PEX25 | XVI | 337436 | 338620 | -1 |
| YPL113C | 529.530621 | -0.6885234 | 0.48750775 | 2.09783434 | 0.35031688 | 0.523786 | NA | XVI | 335953 | 337143 | -1 |
| YPL115C | 151.681311 | 3.37453329 | 0.55967836 | 38.1615086 | 5.17E-09 | 1.64E-07 | BEM3 | XVI | 332100 | 335486 | -1 |
| YPL116W | 61.0156048 | -0.421206 | 0.57256662 | 0.54361936 | 0.76199927 | 0.8553656 | HOS3 | XVI | 329606 | 331699 | 1 |
| YPL117C | 41.9580598 | -0.0234018 | 0.67214461 | 0.04078315 | 0.97981493 | 0.98440542 | IDI1 | XVI | 327864 | 328730 | -1 |
| YPL118W | 17.4520826 | -0.4229381 | 0.81136218 | 0.33307667 | 0.84659036 | 0.90975468 | MRP51 | XVI | 326628 | 327662 | 1 |
| YPL119C | 285.037131 | -0.1297192 | 0.50801935 | 6.72521735 | 0.03464476 | 0.09984728 | DBP1 | XVI | 324411 | 326264 | -1 |
| YPL119C-A | 40.1708839 | -2.2804322 | 0.71033046 | 9.93144839 | 0.0069729 | 0.02994378 | NA | XVI | 324024 | 324287 | -1 |
| YPL120W | 13.0196837 | 0.09665595 | 0.86697619 | 0.22742625 | 0.89251396 | 0.93387025 | VPS30 | XVI | 322071 | 323744 | 1 |
| YPL121C | 30.1622048 | -1.2198075 | 0.78476638 | 5.41365784 | 0.06674814 | 0.16211061 | MEI5 | XVI | 320962 | 321630 | -1 |
| YPL122C | 25.2109492 | -1.119339 | 0.68546245 | 3.04954188 | 0.21767091 | 0.38155469 | TFB2 | XVI | 319226 | 320767 | -1 |
| YPL123C | 14.0181933 | 0.38756765 | 0.89668871 | 1.01481781 | 0.60205354 | 0.74772415 | RNY1 | XVI | 317641 | 318945 | -1 |
| YPL124W | 119.181686 | 2.8684353 | 0.9399625 | 10.8268375 | 0.00445638 | 0.02079165 | SPC29 | XVI | 316755 | 317516 | 1 |
| YPL125W | 10.9744936 | -0.3886747 | 0.88830322 | 0.3470276 | 0.84070555 | 0.90645906 | KAP120 | XVI | 313388 | 316486 | 1 |
| YPL126W | 26.3275124 | -0.563551 | 0.72344243 | 1.40857376 | 0.49446106 | 0.65483292 | NAN1 | XVI | 310210 | 312900 | 1 |
| YPL127C | 63.9471792 | -1.9963106 | 0.56213362 | 12.715291 | 0.00173344 | 0.01006684 | HHO1 | XVI | 308828 | 309604 | -1 |
| YPL128C | 121.285395 | 1.89263848 | 0.54445446 | 11.7944723 | 0.00274703 | 0.01420115 | TBF1 | XVI | 306533 | 308221 | -1 |
| YPL129W | 74.3504342 | 0.5480093 | 0.56585453 | 1.28082869 | 0.52707399 | 0.68184633 | TAF14 | XVI | 305298 | 306137 | 1 |
| YPL131W | 298.94375 | -3.0793969 | 0.57348072 | 25.8485221 | 2.44E-06 | 3.83E-05 | RPL5 | XVI | 303121 | 304014 | 1 |
| YPL132W | 40.3992312 | 0.98455953 | 0.59733374 | 3.55820148 | 0.16878986 | 0.3187519 | COX11 | XVI | 301716 | 302618 | 1 |
| YPL133C | 24.2136169 | 1.19838129 | 0.70819579 | 5.0200682 | 0.08126547 | 0.18851244 | RDS2 | XVI | 299888 | 301228 | -1 |
| YPL134C | 16.2367729 | -0.4097165 | 1.01763158 | 3.05020309 | 0.21759896 | 0.38155469 | ODC1 | XVI | 298571 | 299503 | -1 |
| YPL135W | 37.9698232 | 0.25455892 | 0.70892502 | 1.15002421 | 0.56269806 | 0.71410256 | ISU1 | XVI | 297553 | 298050 | 1 |
| YPL137C | 86.4841071 | 1.77351152 | 0.59973681 | 11.3604354 | 0.00341282 | 0.01686024 | GIP3 | XVI | 292817 | 296647 | -1 |
| YPL138C | 12.9784267 | 1.01380101 | 0.85673476 | 1.65316125 | 0.43754285 | 0.60627631 | SPP1 | XVI | 291366 | 292427 | -1 |
| YPL139C | 34.9498958 | 0.91842856 | 0.68446225 | 9.18527645 | 0.01012611 | 0.03983028 | UME1 | XVI | 289669 | 291051 | -1 |
| YPL140C | 129.469642 | 3.90945683 | 0.58931095 | 41.0624305 | 1.21E-09 | 4.38E-08 | MKK2 | XVI | 287514 | 289034 | -1 |
| YPL141C | 291.130431 | -0.1108559 | 0.48298651 | 0.68037438 | 0.7116371 | 0.82438173 | FRK1 | XVI | 283464 | 286061 | -1 |
| YPL143W | 117.819209 | -2.7859797 | 0.68776859 | 16.0649687 | 0.00032474 | 0.00261509 | RPL33A | XVI | 282122 | 282970 | 1 |
| YPL144W | 11.3359409 | -3.8543587 | 1.05839987 | 16.8569941 | 0.00021855 | 0.0018473 | POC4 | XVI | 280480 | 280926 | 1 |
| YPL145C | 55.834204 | -0.9504091 | 0.68222728 | 2.19214421 | 0.33418114 | 0.5079834 | KES1 | XVI | 278395 | 279699 | -1 |
| YPL146C | 54.8597733 | -2.6781966 | 0.7348671 | 13.2606858 | 0.00131971 | 0.00824961 | NOP53 | XVI | 276162 | 277529 | -1 |
| YPL147W | 33.9082953 | -0.8653725 | 0.68844214 | 1.70824284 | 0.425657 | 0.5957336 | PXA1 | XVI | 273255 | 275867 | 1 |

| | | | | | | | | | | | |
|---|---|---|---|---|---|---|---|---|---|---|---|
| YPL148C | 83.0580671 | -0.8050025 | 0.5556023 | 3.93887977 | 0.13953499 | 0.27801887 | PPT2 | XVI | 272294 | 272815 | -1 |
| YPL149W | 99.0589687 | -0.5649513 | 0.53328852 | 1.54592022 | 0.46164453 | 0.62532091 | ATG5 | XVI | 271310 | 272194 | 1 |
| YPL150W | 44.9007843 | 1.32228653 | 0.61489329 | 4.71938334 | 0.09444934 | 0.20984715 | NA | XVI | 268188 | 270893 | 1 |
| YPL151C | 160.083167 | 0.11126032 | 0.62261029 | 1.0913625 | 0.5794469 | 0.72802613 | PRP46 | XVI | 266180 | 267535 | -1 |
| YPL152W | 76.3505619 | -1.1566227 | 0.68271239 | 3.50053647 | 0.17372734 | 0.32507061 | RRD2 | XVI | 265028 | 266104 | 1 |
| YPL153C | 7.45896983 | -0.1192334 | 1.11461324 | 0.18897236 | 0.90984031 | 0.94492832 | RAD53 | XVI | 261727 | 264192 | -1 |
| YPL154C | 75.3691311 | -0.4782233 | 0.80260299 | 0.3672555 | 0.83224555 | 0.90069191 | PEP4 | XVI | 259714 | 260931 | -1 |
| YPL155C | 28.9025403 | 1.512877 | 0.67325627 | 6.72112818 | 0.03471567 | 0.10000186 | KIP2 | XVI | 257216 | 259336 | -1 |
| YPL156C | 749.616911 | -0.5776142 | 0.50412581 | 2.44024757 | 0.29519362 | 0.46951321 | PRM4 | XVI | 255913 | 256767 | -1 |
| YPL157W | 116.662219 | 1.4000123 | 0.61925185 | 6.25140467 | 0.04390609 | 0.11884817 | TGS1 | XVI | 254814 | 255761 | 1 |
| YPL158C | 268.813706 | 0.39105628 | 0.48796814 | 1.71854031 | 0.42347104 | 0.59425528 | AIM44 | XVI | 252034 | 254310 | -1 |
| YPL159C | 119.904526 | 1.88063117 | 0.51955093 | 12.9468447 | 0.00154393 | 0.00934751 | PET20 | XVI | 250907 | 251668 | -1 |
| YPL160W | 52.3162552 | -2.0831405 | 0.67280464 | 9.30705206 | 0.00952795 | 0.03806203 | CDC60 | XVI | 246990 | 250262 | 1 |
| YPL161C | 71.2311932 | 1.39469624 | 0.5794224 | 6.75529053 | 0.03412772 | 0.09875038 | BEM4 | XVI | 244319 | 246220 | -1 |
| YPL162C | 1130.6657 | 1.20290218 | 0.49597341 | 6.17331954 | 0.0456542 | 0.12191465 | NA | XVI | 243206 | 244027 | -1 |
| YPL163C | 30.3554485 | 1.20170852 | 0.89656478 | 7.0600385 | 0.02930435 | 0.08719023 | SVS1 | XVI | 241918 | 242700 | -1 |
| YPL164C | 33.2342481 | 1.07971705 | 0.74926652 | 4.25102025 | 0.11937206 | 0.24853082 | MLH3 | XVI | 239350 | 241497 | -1 |
| YPL165C | 2534.39633 | 1.89538033 | 0.45724875 | 16.703864 | 0.00023594 | 0.00196844 | SET6 | XVI | 237956 | 239077 | -1 |
| YPL166W | 13.1536764 | -1.2378329 | 0.90745547 | 5.02790264 | 0.08094776 | 0.18800141 | ATG29 | XVI | 237338 | 237979 | 1 |
| YPL167C | 41.7186997 | -0.4428653 | 0.60894627 | 0.57555708 | 0.74992765 | 0.84806272 | REV3 | XVI | 232593 | 237107 | -1 |
| YPL168W | 137.827041 | 1.05535333 | 0.57339307 | 4.31528333 | 0.11559742 | 0.242911 | MRX4 | XVI | 231217 | 232509 | 1 |
| YPL169C | 788.258884 | 6.27643712 | 0.52613239 | 134.021168 | 7.90E-30 | 2.29E-26 | MEX67 | XVI | 229039 | 230838 | -1 |
| YPL170W | 63.5742305 | 3.22408706 | 0.62275105 | 28.0599181 | 8.07E-07 | 1.47E-05 | DAP1 | XVI | 228314 | 228772 | 1 |
| YPL171C | 19.9927473 | 2.96286104 | 0.83186934 | 17.3191172 | 0.00017346 | 0.00150125 | OYE3 | XVI | 226169 | 227371 | -1 |
| YPL172C | 43.3699555 | 2.37646516 | 0.65230537 | 14.1836212 | 0.00083189 | 0.00566552 | COX10 | XVI | 224353 | 225741 | -1 |
| YPL173W | 49.524344 | 2.45179859 | 0.61123438 | 22.2894836 | 1.45E-05 | 0.00018111 | MRPL40 | XVI | 223143 | 224036 | 1 |
| YPL174C | 228.2472 | 0.30151199 | 0.55505796 | 3.09510017 | 0.2127686 | 0.37639175 | NIP100 | XVI | 220167 | 222773 | -1 |
| YPL175W | 138.123499 | -2.6305253 | 0.50503799 | 25.0485718 | 3.64E-06 | 5.44E-05 | SPT14 | XVI | 218630 | 220088 | 1 |
| YPL176C | 38.9679815 | 0.82563289 | 0.66453235 | 2.59979834 | 0.27255927 | 0.44584616 | TRE1 | XVI | 216012 | 218363 | -1 |
| YPL177C | 18.3234074 | 0.25748967 | 0.73712485 | 0.14158013 | 0.93165746 | 0.95711439 | CUP9 | XVI | 213042 | 213962 | -1 |
| YPL178W | 13.4110517 | 0.72212141 | 0.81563572 | 0.82035698 | 0.66353181 | 0.79066663 | CBC2 | XVI | 212158 | 212784 | 1 |
| YPL179W | 58.4547875 | 0.46372511 | 0.73343246 | 0.54452832 | 0.76165304 | 0.85526961 | PPQ1 | XVI | 208157 | 209806 | 1 |
| YPL180W | 37.5913026 | -0.769219 | 0.64157001 | 1.44268397 | 0.48609948 | 0.64790884 | TCO89 | XVI | 205248 | 207647 | 1 |

| | | | | | | | | | | | |
|---|---|---|---|---|---|---|---|---|---|---|---|
| YPL181W | 90.0941349 | 0.17763028 | 0.55472104 | 1.56781234 | 0.45661889 | 0.62207609 | CTI6 | XVI | 203421 | 204941 | 1 |
| YPL183C | 252.884567 | 1.58009793 | 0.75365959 | 5.00767266 | 0.0817707 | 0.18922955 | RTT10 | XVI | 199495 | 202536 | -1 |
| YPL183W-A | 17.8925869 | -1.0479656 | 0.7367071 | 4.45569522 | 0.10776012 | 0.23089058 | RTC6 | XVI | 199095 | 199376 | 1 |
| YPL184C | 28.0105598 | 0.3968325 | 0.70370312 | 0.65402682 | 0.72107407 | 0.83085787 | MRN1 | XVI | 195950 | 197788 | -1 |
| YPL186C | 83.2509087 | 0.36378156 | 0.57799802 | 1.98715935 | 0.37024895 | 0.54354498 | UIP4 | XVI | 194512 | 195426 | -1 |
| YPL187W | 125.429403 | -4.7247572 | 0.66465237 | 44.0175255 | 2.77E-10 | 1.16E-08 | MF(ALPHA)1 | XVI | 193648 | 194145 | 1 |
| YPL188W | 571.553037 | -5.6978924 | 0.50513573 | 116.064346 | 6.27E-26 | 5.18E-23 | POS5 | XVI | 191406 | 192650 | 1 |
| YPL189C-A | 20.4366754 | -0.5904408 | 0.70694456 | 3.08880672 | 0.21343918 | 0.37734744 | COA2 | XVI | 188307 | 188513 | -1 |
| YPL189W | 33.4723445 | 0.64297533 | 0.70178687 | 1.58561899 | 0.45257151 | 0.61859987 | GUP2 | XVI | 189154 | 190983 | 1 |
| YPL190C | 385.523114 | 0.01523522 | 0.63105202 | 3.29998135 | 0.1920517 | 0.34838007 | NAB3 | XVI | 185317 | 187725 | -1 |
| YPL191C | 14.5640653 | 0.31688836 | 0.82316274 | 0.54138636 | 0.76285052 | 0.85565759 | NA | XVI | 183597 | 184679 | -1 |
| YPL192C | 469.227848 | -2.3716485 | 0.4658034 | 25.2204051 | 3.34E-06 | 5.07E-05 | PRM3 | XVI | 182655 | 183056 | -1 |
| YPL193W | 15.4447164 | 0.16177971 | 0.79397363 | 0.40983528 | 0.81471441 | 0.88869564 | RSA1 | XVI | 181403 | 182548 | 1 |
| YPL194W | 9.94502394 | 0.71210996 | 0.95592766 | 0.93067435 | 0.62792335 | 0.76540551 | DDC1 | XVI | 179277 | 181115 | 1 |
| YPL195W | 33.4552174 | -0.363784 | 0.69181484 | 0.42254769 | 0.80955235 | 0.8858947 | APL5 | XVI | 176223 | 179021 | 1 |
| YPL196W | 103.082693 | 0.32565725 | 0.50988413 | 0.80492719 | 0.66867068 | 0.79450097 | OXR1 | XVI | 175043 | 175864 | 1 |
| YPL198W | 28.2920328 | -0.4504853 | 0.63906885 | 1.34514401 | 0.51039415 | 0.66786132 | RPL7B | XVI | 173152 | 174702 | 1 |
| YPL199C | 21.4068206 | -0.0731397 | 0.75461753 | 0.03101604 | 0.98461161 | 0.98751104 | NA | XVI | 172033 | 172755 | -1 |
| YPL200W | 44.6024688 | -2.8605388 | 0.77512516 | 12.401916 | 0.00202749 | 0.01126565 | CSM4 | XVI | 171484 | 171954 | 1 |
| YPL201C | 40.4085867 | 1.5619579 | 0.72828281 | 4.60713053 | 0.09990203 | 0.21918634 | YIG1 | XVI | 169770 | 171155 | -1 |
| YPL202C | 124.130738 | 0.84556236 | 0.62864089 | 2.13946059 | 0.34310104 | 0.5166593 | AFT2 | XVI | 168088 | 169338 | -1 |
| YPL203W | 14.5916833 | 1.2042034 | 1.02026651 | 1.57564185 | 0.45483484 | 0.62066785 | TPK2 | XVI | 166256 | 167398 | 1 |
| YPL204W | 47.8322832 | 0.94272193 | 0.60775216 | 3.59862128 | 0.16541288 | 0.31442566 | HRR25 | XVI | 164276 | 165760 | 1 |
| YPL206C | 69.1752654 | -0.3822278 | 0.53214297 | 1.13090904 | 0.56810187 | 0.71834677 | PGC1 | XVI | 162632 | 163597 | -1 |
| YPL207W | 74.6385325 | -0.8175932 | 0.61651269 | 2.61793289 | 0.27009907 | 0.44340052 | TYW1 | XVI | 159909 | 162341 | 1 |
| YPL208W | 10.3747291 | -1.2199463 | 0.91172234 | 3.07150475 | 0.21529365 | 0.37922681 | RKM1 | XVI | 157842 | 159593 | 1 |
| YPL209C | 6.39482843 | -0.7305826 | 1.20809503 | 0.38418133 | 0.82523204 | 0.89541781 | IPL1 | XVI | 156490 | 157593 | -1 |
| YPL210C | 33.9351836 | -2.0245996 | 0.67755313 | 8.97387029 | 0.01125509 | 0.04301449 | SRP72 | XVI | 154290 | 156212 | -1 |
| YPL211W | 17.3917799 | -3.2381664 | 0.85992213 | 14.2862783 | 0.00079027 | 0.00542138 | NIP7 | XVI | 153495 | 154040 | 1 |
| YPL212C | 29.4611048 | -2.1316864 | 0.74715911 | 9.4584718 | 0.00883322 | 0.03582669 | PUS1 | XVI | 151515 | 153149 | -1 |
| YPL213W | 11.2003942 | 0.17051675 | 0.87298508 | 0.46695849 | 0.79177403 | 0.87354642 | LEA1 | XVI | 150614 | 151330 | 1 |
| YPL214C | 250.733721 | -0.1923295 | 0.77157273 | 4.47323406 | 0.10681926 | 0.22957814 | THI6 | XVI | 148569 | 150191 | -1 |
| YPL215W | 23.7834351 | -0.5399485 | 0.69192276 | 0.85571714 | 0.65190361 | 0.78244649 | CBP3 | XVI | 147416 | 148423 | 1 |

| | | | | | | | | | | | |
|---|---|---|---|---|---|---|---|---|---|---|---|
| YPL216W | 1410.0772 | -0.5921273 | 0.49909511 | 1.46072786 | 0.48173364 | 0.64424159 | NA | XVI | 143821 | 147129 | 1 |
| YPL217C | 191.233867 | 0.56922497 | 0.52157947 | 2.72674413 | 0.25579676 | 0.42803963 | BMS1 | XVI | 139620 | 143171 | -1 |
| YPL218W | 41.5656051 | 0.64299384 | 0.59942687 | 1.25800543 | 0.53312321 | 0.68763274 | SAR1 | XVI | 138698 | 139409 | 1 |
| YPL219W | 21.6007373 | 0.87270891 | 0.72639354 | 2.88995673 | 0.23575118 | 0.40372651 | PCL8 | XVI | 136750 | 138228 | 1 |
| YPL220W | 15.1209626 | -2.7504117 | 0.89744779 | 9.74501126 | 0.00765416 | 0.03206773 | RPL1A | XVI | 135790 | 136443 | 1 |
| YPL221W | 80.6127681 | 0.2124499 | 0.63278998 | 1.41028054 | 0.49403927 | 0.65469026 | FLC1 | XVI | 133043 | 135424 | 1 |
| YPL222W | 123.203915 | 2.35419981 | 0.51879943 | 19.662291 | 5.38E-05 | 0.00054696 | FMP40 | XVI | 130162 | 132228 | 1 |
| YPL223C | 107.83558 | 1.20944835 | 0.5762115 | 4.81142069 | 0.0902014 | 0.20366605 | GRE1 | XVI | 128632 | 129138 | -1 |
| YPL224C | 26.664884 | -0.612271 | 0.67557246 | 1.90907768 | 0.38498964 | 0.55783034 | MMT2 | XVI | 126634 | 128088 | -1 |
| YPL225W | 84.5608594 | 0.03519866 | 0.66562771 | 5.40457715 | 0.06705188 | 0.16264366 | NA | XVI | 126006 | 126446 | 1 |
| YPL226W | 59.9064108 | -1.4699198 | 0.62170086 | 5.73912192 | 0.05672383 | 0.14417513 | NEW1 | XVI | 121767 | 125357 | 1 |
| YPL227C | 50.5108446 | -0.6512258 | 0.90545511 | 4.82284733 | 0.08968752 | 0.20308593 | ALG5 | XVI | 120163 | 121167 | -1 |
| YPL228W | 32.459821 | 1.09570971 | 0.67298898 | 2.70689917 | 0.25834753 | 0.43008401 | CET1 | XVI | 118382 | 120031 | 1 |
| YPL229W | 10.3792567 | -0.550857 | 0.99340953 | 0.32881643 | 0.84839562 | 0.90980423 | NA | XVI | 117067 | 117687 | 1 |
| YPL230W | 162.210869 | -0.0165904 | 0.54276211 | 0.9805714 | 0.61245139 | 0.7554524 | USV1 | XVI | 115312 | 116487 | 1 |
| YPL231W | 177.939825 | -0.2867226 | 0.7014745 | 1.10029763 | 0.57686396 | 0.72593834 | FAS2 | XVI | 108652 | 114315 | 1 |
| YPL232W | 125.331993 | 0.20306262 | 0.5266802 | 2.98292368 | 0.22504344 | 0.3904964 | SSO1 | XVI | 107275 | 108147 | 1 |
| YPL233W | 9.23564103 | 1.72485047 | 1.05351923 | 2.93286724 | 0.23074695 | 0.39774482 | NSL1 | XVI | 106172 | 106822 | 1 |
| YPL234C | 169.026772 | -0.313737 | 0.5734323 | 0.36794399 | 0.8319591 | 0.90069191 | VMA11 | XVI | 104946 | 105440 | -1 |
| YPL235W | 36.4765922 | -0.1231228 | 0.73565795 | 0.92241005 | 0.63052339 | 0.76792815 | RVB2 | XVI | 103232 | 104647 | 1 |
| YPL236C | 9.22046165 | 1.61279687 | 1.05228251 | 3.49946836 | 0.17382014 | 0.32507061 | ENV7 | XVI | 101608 | 102702 | -1 |
| YPL237W | 155.52167 | -0.6692125 | 0.48543124 | 3.48697122 | 0.17490967 | 0.32678559 | SUI3 | XVI | 100496 | 101353 | 1 |
| YPL239W | 47.430462 | -1.1633292 | 0.58632763 | 4.05184107 | 0.13187239 | 0.26715926 | YAR1 | XVI | 99484 | 100086 | 1 |
| YPL240C | 628.241469 | 2.63824654 | 0.48786132 | 31.2019876 | 1.68E-07 | 3.81E-06 | HSP82 | XVI | 96496 | 98625 | -1 |
| YPL241C | 480.450821 | 0.18239636 | 0.467308 | 1.12115775 | 0.5708785 | 0.72122769 | CIN2 | XVI | 95372 | 96258 | -1 |
| YPL242C | 925.218825 | 4.46424257 | 0.48006273 | 69.2951204 | 8.97E-16 | 1.06E-13 | IQG1 | XVI | 90622 | 95109 | -1 |
| YPL243W | 351.574726 | 3.78388305 | 0.53302674 | 42.9669903 | 4.68E-10 | 1.82E-08 | SRP68 | XVI | 88517 | 90316 | 1 |
| YPL244C | 42.0135049 | -1.1064156 | 0.79649905 | 2.01797695 | 0.36458758 | 0.53782474 | HUT1 | XVI | 87014 | 88033 | -1 |
| YPL245W | 530.872484 | -0.6075421 | 0.62068109 | 6.79699875 | 0.03342339 | 0.0971981 | NA | XVI | 85586 | 86950 | 1 |
| YPL246C | 88.5673853 | 0.22882101 | 0.57083402 | 1.65825189 | 0.43643058 | 0.60539844 | RBD2 | XVI | 84509 | 85297 | -1 |
| YPL247C | 50.3800452 | 1.21364846 | 0.60028622 | 4.06128528 | 0.13125115 | 0.26636668 | NA | XVI | 82625 | 84196 | -1 |
| YPL248C | 1384.06982 | -0.9548986 | 0.47878019 | 4.48985083 | 0.10593544 | 0.22838155 | GAL4 | XVI | 79711 | 82356 | -1 |
| YPL249C | 562.087254 | -2.4457318 | 0.51530187 | 23.0815831 | 9.73E-06 | 0.00012915 | GYP5 | XVI | 76669 | 79353 | -1 |

| | | | | | | | | | | | |
|---|---|---|---|---|---|---|---|---|---|---|---|
| YPL249C-A | 254.175645 | -2.9643517 | 0.63963988 | 19.5120085 | 5.79E-05 | 0.00058655 | RPL36B | XVI | 75699 | 76239 | -1 |
| YPL250C | 10.6573536 | 0.19478514 | 0.95760859 | 0.12717452 | 0.93839223 | 0.95995205 | ATG41 | XVI | 74309 | 74719 | -1 |
| YPL252C | 27.4685969 | 1.361654 | 0.67446953 | 8.28466083 | 0.01588579 | 0.05601627 | YAH1 | XVI | 73363 | 73881 | -1 |
| YPL253C | 580.071865 | -2.0618783 | 0.48887305 | 16.9365121 | 0.00021003 | 0.0017805 | VIK1 | XVI | 71063 | 73006 | -1 |
| YPL254W | 25.888218 | -0.434175 | 0.74816432 | 0.39616487 | 0.82030223 | 0.8920536 | HFI1 | XVI | 69485 | 70951 | 1 |
| YPL255W | 14.2222383 | -0.5929465 | 0.78446598 | 2.10566082 | 0.34894868 | 0.52261068 | BBP1 | XVI | 67725 | 68882 | 1 |
| YPL256C | 160.696653 | 0.71852016 | 0.49876013 | 5.85472679 | 0.05353801 | 0.13770994 | CLN2 | XVI | 64977 | 66614 | -1 |
| YPL257W | 50.1080659 | 0.57744751 | 0.58933696 | 1.75778382 | 0.41524278 | 0.58726324 | NA | XVI | 63279 | 63860 | 1 |
| YPL258C | 475.460018 | 4.96864183 | 0.53755362 | 75.0910987 | 4.95E-17 | 7.53E-15 | THI21 | XVI | 53498 | 55153 | -1 |
| YPL259C | 23.3456495 | 0.06969664 | 0.72660195 | 2.3682758 | 0.30600988 | 0.47867401 | APM1 | XVI | 51244 | 52671 | -1 |
| YPL260W | 58.9210019 | 1.67726489 | 0.57901814 | 8.76993894 | 0.01246327 | 0.04652632 | CUB1 | XVI | 49303 | 50958 | 1 |
| YPL261C | 1331.73461 | 0.30386086 | 0.55342197 | 0.57814092 | 0.74895943 | 0.8479615 | NA | XVI | 48996 | 49304 | -1 |
| YPL262W | 23.214562 | -0.5638947 | 0.79953535 | 1.60226924 | 0.44881944 | 0.61506853 | FUM1 | XVI | 47336 | 48802 | 1 |
| YPL263C | 58.3671485 | -0.8193805 | 0.62296941 | 2.43059017 | 0.29662247 | 0.47066158 | KEL3 | XVI | 44551 | 46506 | -1 |
| YPL264C | 101.917013 | 0.49451939 | 0.56567156 | 1.93719688 | 0.37961472 | 0.55350522 | NA | XVI | 43283 | 44344 | -1 |
| YPL265W | 733.145839 | -0.791914 | 0.58671387 | 1.90533383 | 0.38571099 | 0.55845627 | DIP5 | XVI | 41043 | 42869 | 1 |
| YPL266W | 38.6004515 | -0.7919865 | 0.72834793 | 6.76745468 | 0.03392078 | 0.09824187 | DIM1 | XVI | 39121 | 40077 | 1 |
| YPL267W | 5.88051903 | -1.8573407 | 1.22008454 | 3.11964615 | 0.21017325 | 0.3730543 | ACM1 | XVI | 38169 | 38798 | 1 |
| YPL268W | 62.4517549 | -2.0599128 | 0.58226445 | 18.4407873 | 9.90E-05 | 0.00092602 | PLC1 | XVI | 35236 | 37845 | 1 |
| YPL269W | 25.2443604 | 1.13860222 | 0.70366034 | 2.59115026 | 0.27374038 | 0.44722258 | KAR9 | XVI | 33013 | 34947 | 1 |
| YPL270W | 26.0190313 | 0.64339212 | 0.66981849 | 1.05666483 | 0.58958734 | 0.73728204 | MDL2 | XVI | 30482 | 32803 | 1 |
| YPL271W | 16.7640561 | -0.1679715 | 0.76219731 | 0.16953419 | 0.91872623 | 0.94955817 | ATP15 | XVI | 30079 | 30267 | 1 |
| YPL272C | 8.06734631 | -0.5344129 | 0.94651729 | 1.81405111 | 0.40372329 | 0.57458523 | PBI1 | XVI | 26611 | 28164 | -1 |
| YPL273W | 11.4630495 | -1.970775 | 0.89220964 | 4.93476028 | 0.08480675 | 0.19477631 | SAM4 | XVI | 25087 | 26064 | 1 |
| YPL274W | 131.473962 | -1.0115966 | 0.53951401 | 3.59201425 | 0.16596023 | 0.31525228 | SAM3 | XVI | 22938 | 24701 | 1 |
| YPL275W | 158.924384 | -1.2364135 | 0.64213767 | 7.49910269 | 0.0235283 | 0.07472784 | NA | XVI | 18369 | 19079 | 1 |
| YPL276W | 158.569452 | -1.9248106 | 0.61873668 | 9.14736417 | 0.01031989 | 0.04032978 | NA | XVI | 17948 | 18385 | 1 |
| YPL277C | 25.7690514 | 2.35044043 | 0.79634572 | 13.6420655 | 0.00109059 | 0.00709251 | NA | XVI | 15405 | 16868 | -1 |
| YPL279C | 13.1864606 | 2.60869483 | 1.02028554 | 8.75707396 | 0.0125437 | 0.04676626 | FEX2 | XVI | 13228 | 14355 | -1 |
| YPL280W | 7.94479807 | 1.52503671 | 1.00375801 | 10.9948512 | 0.00409731 | 0.0195435 | HSP32 | XVI | 11887 | 12600 | 1 |
| YPL283C | 5.03946198 | -0.5734342 | 1.34018777 | 0.23248221 | 0.89026055 | 0.93317728 | YRF1-7 | XVI | 280 | 6007 | -1 |
| YPR001W | 283.284459 | -2.1752516 | 0.5126308 | 17.6048633 | 0.00015037 | 0.00133737 | CIT3 | XVI | 556377 | 557837 | 1 |
| YPR002W | 76.753294 | 1.05026889 | 0.56767167 | 4.27156744 | 0.11815196 | 0.24673616 | PDH1 | XVI | 558385 | 559935 | 1 |

| | | | | | | | | | | | |
|---|---|---|---|---|---|---|---|---|---|---|---|
| YPR003C | 27.2607812 | -0.6765975 | 0.71597771 | 1.40216476 | 0.4960481 | 0.65603003 | NA | XVI | 561504 | 563768 | -1 |
| YPR004C | 302.593763 | 4.43914584 | 0.51690522 | 74.4700943 | 6.75E-17 | 9.79E-15 | AIM45 | XVI | 564007 | 565041 | -1 |
| YPR005C | 106.84206 | 1.49676237 | 0.53384425 | 14.6889129 | 0.00064616 | 0.00457931 | HAL1 | XVI | 565787 | 566671 | -1 |
| YPR006C | 137.461258 | -0.2431556 | 0.50671493 | 0.98443897 | 0.61126819 | 0.75489714 | ICL2 | XVI | 567269 | 568996 | -1 |
| YPR007C | 24.881462 | -0.9993093 | 0.72285514 | 3.57238572 | 0.16759702 | 0.31722352 | REC8 | XVI | 569336 | 571378 | -1 |
| YPR008W | 149.498349 | 1.07840627 | 0.54960464 | 14.8607473 | 0.00059297 | 0.00424214 | HAA1 | XVI | 573018 | 575102 | 1 |
| YPR009W | 226.443113 | -3.181078 | 0.51218773 | 35.1821864 | 2.29E-08 | 6.35E-07 | SUT2 | XVI | 576552 | 577358 | 1 |
| YPR010C | 44.5632116 | -2.9702954 | 0.75117475 | 15.1568779 | 0.00051136 | 0.00373834 | RPA135 | XVI | 577585 | 581196 | -1 |
| YPR010C-A | 12.556251 | -0.522297 | 0.86434346 | 0.63499674 | 0.72796787 | 0.83592431 | NA | XVI | 582373 | 582734 | -1 |
| YPR011C | 226.558345 | 1.234867 | 0.49549814 | 8.20805527 | 0.01650606 | 0.05733167 | NA | XVI | 583062 | 584042 | -1 |
| YPR013C | 35.7681176 | 1.16504922 | 0.65849604 | 3.56513586 | 0.16820565 | 0.31795975 | CMR3 | XVI | 584632 | 585585 | -1 |
| YPR014C | 136.241567 | 0.2427723 | 0.60359205 | 0.2129217 | 0.89901024 | 0.93754851 | NA | XVI | 587189 | 587518 | -1 |
| YPR015C | 57.2397923 | 3.06373926 | 0.71415479 | 24.679608 | 4.37E-06 | 6.35E-05 | NA | XVI | 590283 | 591026 | -1 |
| YPR016C | 21.0717727 | -2.0216517 | 0.89769956 | 5.05524392 | 0.07984868 | 0.18619567 | TIF6 | XVI | 592332 | 593069 | -1 |
| YPR017C | 42.0035366 | -1.1788489 | 0.71621453 | 3.0140964 | 0.22156302 | 0.38636302 | DSS4 | XVI | 593486 | 593917 | -1 |
| YPR018W | 78.6938773 | -0.4261058 | 0.63158939 | 2.07094979 | 0.35505772 | 0.52857757 | RLF2 | XVI | 594476 | 596296 | 1 |
| YPR019W | 19.6066079 | -0.1964024 | 0.84016112 | 0.29119275 | 0.86450657 | 0.9204658 | MCM4 | XVI | 596750 | 599551 | 1 |
| YPR020W | 8.86099299 | 0.38567103 | 1.22642061 | 0.32934998 | 0.84816932 | 0.90980423 | ATP20 | XVI | 599870 | 600217 | 1 |
| YPR021C | 54.3561844 | 2.10710581 | 0.60611594 | 22.8942104 | 1.07E-05 | 0.00013991 | AGC1 | XVI | 600649 | 603357 | -1 |
| YPR022C | 41.0964281 | -0.4293615 | 0.62929186 | 0.55653428 | 0.75709454 | 0.85316805 | SDD4 | XVI | 603911 | 607312 | -1 |
| YPR023C | 69.2204601 | -0.0994115 | 0.54124699 | 0.83405742 | 0.659002 | 0.7867261 | EAF3 | XVI | 608826 | 610031 | -1 |
| YPR024W | 84.2865756 | 2.31974915 | 0.57271643 | 19.5506316 | 5.68E-05 | 0.00057634 | YME1 | XVI | 610481 | 612724 | 1 |
| YPR025C | 52.1863232 | 0.22625881 | 0.71120168 | 1.99925069 | 0.36801729 | 0.5412294 | CCL1 | XVI | 613377 | 614558 | -1 |
| YPR026W | 83.3897458 | -0.6507718 | 0.57447975 | 3.14716305 | 0.2073014 | 0.36942908 | ATH1 | XVI | 615379 | 619014 | 1 |
| YPR027C | 839.67191 | -2.527461 | 0.47148506 | 27.7898133 | 9.24E-07 | 1.66E-05 | NA | XVI | 620425 | 621258 | -1 |
| YPR028W | 23.3804826 | -0.4504105 | 0.93295178 | 0.86624248 | 0.64848187 | 0.78028055 | YOP1 | XVI | 623527 | 624202 | 1 |
| YPR029C | 44.4809343 | -0.748109 | 0.71372664 | 1.12446081 | 0.56993646 | 0.72050919 | APL4 | XVI | 624469 | 626967 | -1 |
| YPR030W | 246.855467 | -3.1866379 | 0.52004509 | 34.2375443 | 3.68E-08 | 9.83E-07 | CSR2 | XVI | 627880 | 631245 | 1 |
| YPR031W | 18.606281 | 0.62490237 | 0.74551046 | 3.01943499 | 0.2209724 | 0.38583539 | NTO1 | XVI | 631515 | 633761 | 1 |
| YPR032W | 19.651125 | -0.3815409 | 0.75257918 | 0.35957453 | 0.83544792 | 0.90300565 | SRO7 | XVI | 634123 | 637224 | 1 |
| YPR033C | 31.5755907 | -1.4014111 | 0.72084209 | 4.08342841 | 0.12980601 | 0.26426638 | HTS1 | XVI | 637379 | 639019 | -1 |
| YPR034W | 38.4524955 | 0.60806944 | 0.6279174 | 1.81567172 | 0.40339629 | 0.57429666 | ARP7 | XVI | 639525 | 640958 | 1 |
| YPR035W | 99.0273959 | 0.69431102 | 0.70252341 | 3.42419692 | 0.18048665 | 0.33506648 | GLN1 | XVI | 642208 | 643320 | 1 |

| | | | | | | | | | | | |
|---|---|---|---|---|---|---|---|---|---|---|---|
| YPR036W | 351.851788 | -1.4510196 | 0.56489347 | 7.36216253 | 0.02519572 | 0.07822156 | VMA13 | XVI | 643836 | 645272 | 1 |
| YPR036W-A | 100.69823 | 2.03843681 | 0.58106797 | 11.9087477 | 0.00259447 | 0.01364393 | SPO24 | XVI | 645950 | 646153 | 1 |
| YPR037C | 36.6514529 | 0.68016708 | 0.75224434 | 5.66510258 | 0.05886249 | 0.14753844 | ERV2 | XVI | 646448 | 647038 | -1 |
| YPR038W | 1219.38698 | 0.05298726 | 0.48986315 | 0.27284522 | 0.87247384 | 0.92527295 | NA | XVI | 646836 | 647195 | 1 |
| YPR040W | 99.1660519 | -0.6026459 | 0.53531353 | 2.09515949 | 0.35078571 | 0.52427705 | TIP41 | XVI | 647305 | 648375 | 1 |
| YPR041W | 36.8095756 | -2.3045603 | 0.86678488 | 6.8435885 | 0.03265379 | 0.09534315 | TIF5 | XVI | 648704 | 649921 | 1 |
| YPR042C | 18.9980674 | 0.77029088 | 0.95674314 | 1.29353474 | 0.52373609 | 0.67900939 | PUF2 | XVI | 650435 | 653662 | -1 |
| YPR043W | 70.3730866 | -2.8642999 | 0.73547175 | 13.9576547 | 0.00093139 | 0.00620573 | RPL43B | XVI | 654166 | 654847 | 1 |
| YPR045C | 48.7943914 | -0.9432754 | 0.58583619 | 2.62272405 | 0.2694528 | 0.44259056 | THP3 | XVI | 655140 | 656552 | -1 |
| YPR046W | 62.1102106 | 0.73250925 | 0.57702676 | 3.20132604 | 0.2017627 | 0.36122636 | MCM16 | XVI | 656799 | 657344 | 1 |
| YPR047W | 55.9062388 | -2.484527 | 0.60922273 | 19.4157262 | 6.08E-05 | 0.00061014 | MSF1 | XVI | 657529 | 658938 | 1 |
| YPR048W | 46.1531846 | 0.17970672 | 0.87461452 | 1.7992975 | 0.40671249 | 0.57811175 | TAH18 | XVI | 659182 | 661053 | 1 |
| YPR049C | 24.7740599 | -0.4578591 | 0.6829353 | 0.71422503 | 0.69969377 | 0.81801018 | ATG11 | XVI | 661137 | 664673 | -1 |
| YPR051W | 48.2897862 | -0.7492269 | 0.76641571 | 8.74855413 | 0.01259725 | 0.04684525 | MAK3 | XVI | 664960 | 665490 | 1 |
| YPR053C | 24.0057033 | -2.8114797 | 0.74859284 | 15.84528 | 0.00036244 | 0.0028442 | NA | XVI | 665788 | 666243 | -1 |
| YPR054W | 1001.74801 | -1.2739999 | 0.54803662 | 7.09211841 | 0.02883806 | 0.08637991 | SMK1 | XVI | 666280 | 667446 | 1 |
| YPR055W | 42.8673356 | 0.02342111 | 0.59648743 | 0.00753545 | 0.99623937 | 0.99658361 | SEC8 | XVI | 667676 | 670873 | 1 |
| YPR056W | 119.951034 | -0.0287744 | 0.56372748 | 0.89354086 | 0.63969075 | 0.77534214 | TFB4 | XVI | 671126 | 672142 | 1 |
| YPR057W | 46.2769565 | -0.8720855 | 0.57986662 | 9.38821 | 0.00914905 | 0.03683798 | BRR1 | XVI | 672471 | 673496 | 1 |
| YPR058W | 85.2992813 | -3.0669603 | 0.56061867 | 27.422304 | 1.11E-06 | 1.94E-05 | YMC1 | XVI | 673751 | 674674 | 1 |
| YPR060C | 45.2808916 | -1.9679017 | 0.59483533 | 10.5120197 | 0.00521608 | 0.02376215 | ARO7 | XVI | 674861 | 675631 | -1 |
| YPR061C | 36.4760078 | -1.1258119 | 0.61872831 | 11.5040644 | 0.00317632 | 0.01596431 | JID1 | XVI | 675977 | 676882 | -1 |
| YPR062W | 482.269379 | -1.7909857 | 0.7876599 | 10.2889377 | 0.00583157 | 0.02592206 | FCY1 | XVI | 677165 | 677641 | 1 |
| YPR063C | 22.2085036 | -0.5761968 | 0.71304006 | 0.66016039 | 0.71886608 | 0.82938751 | NA | XVI | 677812 | 678320 | -1 |
| YPR064W | 65.3512448 | 1.71201784 | 0.63391349 | 7.19936922 | 0.02733234 | 0.08294248 | NA | XVI | 678951 | 679370 | 1 |
| YPR065W | 114.260189 | 1.28003186 | 0.59222678 | 5.06598131 | 0.07942114 | 0.18542275 | ROX1 | XVI | 679693 | 680799 | 1 |
| YPR066W | 177.590596 | 0.48980454 | 0.4940991 | 1.16626555 | 0.55814708 | 0.70932212 | UBA3 | XVI | 681213 | 682112 | 1 |
| YPR067W | 18.7564939 | -0.7116339 | 0.81752036 | 3.76356093 | 0.15231867 | 0.29594801 | ISA2 | XVI | 682220 | 682777 | 1 |
| YPR068C | 132.828873 | -0.4661614 | 0.60866911 | 1.41148979 | 0.49374065 | 0.65469026 | HOS1 | XVI | 682941 | 684353 | -1 |
| YPR069C | 21.5254471 | -2.035507 | 0.92847126 | 4.75842763 | 0.09262337 | 0.20714148 | SPE3 | XVI | 684556 | 685437 | -1 |
| YPR070W | 6.93047238 | -0.0381047 | 1.13817957 | 1.99854322 | 0.3681475 | 0.5412834 | MED1 | XVI | 685898 | 687598 | 1 |
| YPR071W | 6.27428739 | -1.0391835 | 1.23833592 | 0.91314566 | 0.63345087 | 0.77035929 | NA | XVI | 688172 | 688807 | 1 |
| YPR072W | 553.038133 | -0.4460107 | 0.67205807 | 5.61070849 | 0.06048534 | 0.15079771 | NOT5 | XVI | 690107 | 691789 | 1 |

| | | | | | | | | | | | |
|---|---|---|---|---|---|---|---|---|---|---|---|
| YPR073C | 39.5086949 | -1.0343137 | 0.66591066 | 2.41913912 | 0.29832566 | 0.47168367 | LTP1 | XVI | 691933 | 692418 | -1 |
| YPR074C | 111.917052 | -0.6758673 | 0.62284866 | 1.88514156 | 0.38962491 | 0.56101577 | TKL1 | XVI | 692796 | 694838 | -1 |
| YPR075C | 91.5168473 | 0.27794044 | 0.52582095 | 0.33791181 | 0.84454614 | 0.90907644 | OPY2 | XVI | 695737 | 696819 | -1 |
| YPR078C | 3551.47758 | 3.4910593 | 0.51420533 | 40.4220038 | 1.67E-09 | 5.89E-08 | NA | XVI | 697147 | 698265 | -1 |
| YPR079W | 15.5386709 | 0.9637806 | 0.86666516 | 1.39829974 | 0.49700765 | 0.65655356 | MRL1 | XVI | 698869 | 700014 | 1 |
| YPR080W | 108.552734 | -1.908629 | 0.79504676 | 5.84046541 | 0.05392114 | 0.1384494 | TEF1 | XVI | 700594 | 701970 | 1 |
| YPR081C | 300.970113 | 2.64501539 | 0.65274033 | 15.7651245 | 0.00037727 | 0.00293993 | GRS2 | XVI | 702114 | 703970 | -1 |
| YPR082C | 88.0293804 | 3.23441996 | 0.57635577 | 33.4497057 | 5.45E-08 | 1.42E-06 | DIB1 | XVI | 704225 | 704656 | -1 |
| YPR083W | 12.4207663 | -0.5764608 | 0.90715734 | 0.75416996 | 0.68585779 | 0.80812303 | MDM36 | XVI | 704854 | 706593 | 1 |
| YPR084W | 48.6250406 | 1.70866167 | 0.59746897 | 9.25738424 | 0.00976753 | 0.03870798 | NA | XVI | 706972 | 708342 | 1 |
| YPR085C | 281.912116 | -1.8949938 | 0.51557725 | 12.6850173 | 0.00175988 | 0.01017954 | ASA1 | XVI | 708497 | 709828 | -1 |
| YPR086W | 91.3255302 | 0.16437104 | 0.61633943 | 0.15321968 | 0.92625117 | 0.95376032 | SUA7 | XVI | 710101 | 711138 | 1 |
| YPR088C | 189.869246 | -0.069159 | 0.81332736 | 11.1049768 | 0.0038778 | 0.0187416 | SRP54 | XVI | 711403 | 713028 | -1 |
| YPR089W | 151.229419 | -0.3146607 | 0.60830289 | 5.36116047 | 0.06852338 | 0.16558864 | NA | XVI | 713275 | 715941 | 1 |
| YPR091C | 71.4334244 | 1.11580575 | 0.64989325 | 2.88591515 | 0.23622806 | 0.40406514 | NVJ2 | XVI | 716156 | 718468 | -1 |
| YPR093C | 33.5784538 | -0.1721721 | 0.67570443 | 0.28636169 | 0.86659734 | 0.92150571 | ASR1 | XVI | 719558 | 720424 | -1 |
| YPR094W | 115.836428 | -0.1695841 | 0.80347377 | 8.66285808 | 0.01314874 | 0.04830662 | RDS3 | XVI | 720637 | 720960 | 1 |
| YPR095C | 25.6102537 | 0.02672521 | 0.70628382 | 1.74898461 | 0.41707371 | 0.58892811 | SYT1 | XVI | 721037 | 724717 | -1 |
| YPR096C | 13.1761713 | 0.39509813 | 0.83257168 | 1.14581722 | 0.56388293 | 0.71488553 | NA | XVI | 724842 | 725144 | -1 |
| YPR097W | 43.081126 | 0.93214608 | 0.67663425 | 1.93917433 | 0.37923957 | 0.55323686 | NA | XVI | 725394 | 728615 | 1 |
| YPR098C | 106.187201 | 2.48474401 | 0.55848766 | 22.0140575 | 1.66E-05 | 0.00020089 | NA | XVI | 728947 | 729528 | -1 |
| YPR099C | 2962.89302 | -0.9032023 | 0.56328645 | 3.36476788 | 0.1859302 | 0.34169876 | NA | XVI | 729757 | 730113 | -1 |
| YPR100W | 96.8956499 | 0.44133705 | 0.56939889 | 1.5008531 | 0.47216511 | 0.63393528 | MRPL51 | XVI | 729791 | 730213 | 1 |
| YPR101W | 79.8619494 | -1.1521942 | 0.52281232 | 4.98320405 | 0.08277725 | 0.19087227 | SNT309 | XVI | 730492 | 731019 | 1 |
| YPR102C | 105.88572 | -1.5460097 | 0.69160944 | 10.0014825 | 0.00673295 | 0.02911412 | RPL11A | XVI | 731224 | 731748 | -1 |
| YPR103W | 53.2917747 | -0.0800515 | 0.67791424 | 0.46231362 | 0.79361501 | 0.87457764 | PRE2 | XVI | 732349 | 733212 | 1 |
| YPR104C | 53.6749955 | 0.25254938 | 0.61817834 | 0.26431319 | 0.87620377 | 0.92712351 | FHL1 | XVI | 733623 | 736433 | -1 |
| YPR105C | 24.0145628 | -0.268428 | 0.69655863 | 1.68736658 | 0.43012333 | 0.59923342 | COG4 | XVI | 736984 | 739569 | -1 |
| YPR106W | 188.100392 | -0.5320225 | 0.50574321 | 2.51056999 | 0.28499461 | 0.45862113 | ISR1 | XVI | 740061 | 741392 | 1 |
| YPR107C | 129.146495 | -1.1435518 | 0.58033188 | 3.79465473 | 0.1499689 | 0.29275789 | YTH1 | XVI | 741438 | 742064 | -1 |
| YPR108W | 49.8542291 | 0.29206799 | 0.69768231 | 0.42713621 | 0.80769715 | 0.88441988 | RPN7 | XVI | 742454 | 743743 | 1 |
| YPR109W | 22.6336722 | -1.9887515 | 0.73093457 | 7.42683642 | 0.024394 | 0.07671608 | NA | XVI | 744689 | 745573 | 1 |
| YPR110C | 42.6983544 | -1.006619 | 0.83381921 | 1.60861814 | 0.44739694 | 0.61428226 | RPC40 | XVI | 745828 | 746835 | -1 |

| | | | | | | | | | | | |
|---|---|---|---|---|---|---|---|---|---|---|---|
| YPR111W | 14.7592286 | 1.06717329 | 0.81248786 | 2.09988779 | 0.34995738 | 0.52354248 | DBF20 | XVI | 747306 | 749000 | 1 |
| YPR112C | 56.5603051 | -2.3709077 | 0.62458385 | 13.5773329 | 0.00112647 | 0.00725502 | MRD1 | XVI | 749256 | 751919 | -1 |
| YPR113W | 14.006557 | -0.9421659 | 0.96565832 | 0.98375491 | 0.6114773 | 0.75489714 | PIS1 | XVI | 752257 | 752919 | 1 |
| YPR114W | 40.323588 | 0.6932025 | 0.67542862 | 2.39483099 | 0.30197366 | 0.47448396 | NA | XVI | 753301 | 754248 | 1 |
| YPR115W | 34.7210355 | -0.0920993 | 0.71917878 | 1.20104716 | 0.54852436 | 0.70063006 | RGC1 | XVI | 754877 | 758128 | 1 |
| YPR116W | 12.3286806 | -0.950547 | 0.85497925 | 3.08391093 | 0.2139623 | 0.37804141 | RRG8 | XVI | 758648 | 759481 | 1 |
| YPR117W | 56.5279885 | -0.7810832 | 0.59495527 | 1.70689982 | 0.42594293 | 0.59578916 | NA | XVI | 760025 | 767494 | 1 |
| YPR118W | 65.464723 | 1.69398136 | 0.56263876 | 13.0142892 | 0.00149274 | 0.0091027 | MRI1 | XVI | 767752 | 768987 | 1 |
| YPR119W | 39.2633039 | 1.93486086 | 0.64883658 | 13.1900114 | 0.00136718 | 0.00848442 | CLB2 | XVI | 771653 | 773128 | 1 |
| YPR120C | 8.79983662 | -1.1620327 | 0.95970085 | 1.94417218 | 0.37829306 | 0.55223646 | CLB5 | XVI | 773875 | 775182 | -1 |
| YPR122W | 1392.99908 | -1.3493125 | 0.48638664 | 7.63006373 | 0.02203701 | 0.07124193 | AXL1 | XVI | 782045 | 785671 | 1 |
| YPR124W | 320.609035 | 1.37816157 | 0.48612416 | 11.1173075 | 0.00385396 | 0.01864197 | CTR1 | XVI | 786208 | 787428 | 1 |
| YPR125W | 21.0083741 | -0.2702716 | 0.79546796 | 0.29183376 | 0.86422954 | 0.92034008 | YLH47 | XVI | 787961 | 789325 | 1 |
| YPR127W | 261.990283 | -0.0665375 | 0.64981986 | 0.03594328 | 0.98218888 | 0.98644816 | NA | XVI | 790083 | 791120 | 1 |
| YPR128C | 27.7725382 | -0.644035 | 0.67572304 | 2.52239651 | 0.28331434 | 0.45680591 | ANT1 | XVI | 791218 | 792204 | -1 |
| YPR129W | 346.507596 | 0.11882834 | 0.80272129 | 7.8518501 | 0.01972388 | 0.06522061 | SCD6 | XVI | 792687 | 793736 | 1 |
| YPR131C | 28.9309338 | -0.3796391 | 2.14699855 | 0.20658367 | 0.90186374 | 0.93824416 | NAT3 | XVI | 793911 | 794498 | -1 |
| YPR132W | 165.67757 | -2.9756873 | 0.72046984 | 15.1212461 | 0.00052055 | 0.00378167 | RPS23B | XVI | 794965 | 795767 | 1 |
| YPR133C | 82.2538506 | -0.4442523 | 0.59587042 | 1.54520921 | 0.46180867 | 0.62532091 | SPN1 | XVI | 795978 | 797210 | -1 |
| YPR133W-A | 18.2827308 | -2.2530433 | 0.95133081 | 7.01011563 | 0.03004504 | 0.08916492 | TOM5 | XVI | 797557 | 797709 | 1 |
| YPR134W | 167.716243 | -1.338422 | 0.48841067 | 9.20529446 | 0.01002526 | 0.03951413 | MSS18 | XVI | 798051 | 798857 | 1 |
| YPR135W | 231.991345 | 1.0799166 | 0.49484751 | 5.78534621 | 0.05542785 | 0.14137538 | CTF4 | XVI | 799234 | 802017 | 1 |
| YPR137W | 42.3265204 | -1.132838 | 0.64933291 | 3.54201691 | 0.1701613 | 0.32050551 | RRP9 | XVI | 802359 | 804080 | 1 |
| YPR138C | 2332.41412 | 1.84060705 | 0.71625457 | 6.22058673 | 0.04458787 | 0.12002035 | MEP3 | XVI | 810984 | 812453 | -1 |
| YPR139C | 38.116452 | -0.2842211 | 0.65375526 | 0.97459565 | 0.61428406 | 0.75689745 | LOA1 | XVI | 813156 | 814058 | -1 |
| YPR140W | 228.030671 | 4.42295959 | 0.58354393 | 73.9224345 | 8.87E-17 | 1.17E-14 | TAZ1 | XVI | 814391 | 815536 | 1 |
| YPR141C | 104.041312 | 1.5114326 | 0.55774221 | 13.1248784 | 0.00141244 | 0.00873719 | KAR3 | XVI | 815734 | 817923 | -1 |
| YPR143W | 47.2086741 | -2.2490426 | 0.69754006 | 10.1380085 | 0.00628868 | 0.0275011 | RRP15 | XVI | 818323 | 819075 | 1 |
| YPR144C | 96.6849167 | 2.25611378 | 0.53742174 | 23.4467957 | 8.10E-06 | 0.00010935 | NOC4 | XVI | 819765 | 821423 | -1 |
| YPR145C-A | 17.3632703 | 6.10885972 | 1.65700967 | 26.3143147 | 1.93E-06 | 3.17E-05 | NA | XVI | 824690 | 824926 | -1 |
| YPR145W | 65.2262466 | -1.5417464 | 0.64247 | 7.56338517 | 0.02278409 | 0.07300493 | ASN1 | XVI | 822620 | 824338 | 1 |
| YPR147C | 17.2630117 | 1.37918955 | 0.82417305 | 2.76979909 | 0.25034895 | 0.42149474 | NA | XVI | 825645 | 826559 | -1 |
| YPR148C | 104.848504 | 0.62739291 | 0.60689424 | 4.78554442 | 0.09137602 | 0.20538321 | NA | XVI | 826833 | 828140 | -1 |

| | | | | | | | | | | | |
|---|---|---|---|---|---|---|---|---|---|---|---|
| YPR149W | 294.377135 | 1.20174044 | 0.47588797 | 6.59436646 | 0.03698721 | 0.10476085 | NCE102 | XVI | 829918 | 830439 | 1 |
| YPR151C | 76.4446537 | 1.98086546 | 0.69115606 | 8.28478312 | 0.01588482 | 0.05601627 | SUE1 | XVI | 831055 | 831675 | -1 |
| YPR152C | 47.3845413 | 2.06399995 | 0.63552278 | 13.6359354 | 0.00109394 | 0.00709286 | URN1 | XVI | 832061 | 833458 | -1 |
| YPR153W | 109.381204 | 4.30329622 | 0.62610823 | 49.5328177 | 1.75E-11 | 9.77E-10 | 24-May | XVI | 833689 | 834245 | 1 |
| YPR154W | 40.8861089 | 2.02621521 | 0.74745439 | 8.61677264 | 0.01345524 | 0.049183 | PIN3 | XVI | 834565 | 835212 | 1 |
| YPR155C | 55.1345794 | 1.53369966 | 0.60085803 | 6.30600471 | 0.04272366 | 0.11657399 | NCA2 | XVI | 835563 | 837413 | -1 |
| YPR156C | 49.1105503 | 2.95549184 | 0.84208979 | 13.0697735 | 0.00145189 | 0.00892151 | TPO3 | XVI | 837909 | 839777 | -1 |
| YPR157W | 29.4432781 | 3.4605127 | 0.75255245 | 24.7698062 | 4.18E-06 | 6.13E-05 | TDA6 | XVI | 841266 | 842669 | 1 |
| YPR158W | 21.5599466 | 1.94732679 | 0.7985166 | 5.86424349 | 0.05328386 | 0.13711714 | CUR1 | XVI | 843262 | 844020 | 1 |
| YPR159W | 75.5596377 | 1.63559453 | 0.64652504 | 8.07101308 | 0.01767672 | 0.06020484 | KRE6 | XVI | 857583 | 859745 | 1 |
| YPR160W | 250.266328 | -1.2859906 | 0.49666713 | 7.50605936 | 0.0234466 | 0.07454061 | GPH1 | XVI | 861306 | 864014 | 1 |
| YPR161C | 475.691265 | 2.68021914 | 0.52544222 | 30.9123856 | 1.94E-07 | 4.27E-06 | SGV1 | XVI | 864449 | 866422 | -1 |
| YPR162C | 122.055498 | 1.76699436 | 0.52169752 | 14.8643199 | 0.00059191 | 0.00424214 | ORC4 | XVI | 866715 | 868304 | -1 |
| YPR163C | 90.8924626 | -1.9739848 | 0.62232221 | 9.78853969 | 0.00748938 | 0.03162131 | TIF3 | XVI | 868645 | 869955 | -1 |
| YPR164W | 35.2820128 | 1.66596742 | 0.66052528 | 6.61789132 | 0.03655469 | 0.10400574 | MMS1 | XVI | 870703 | 874926 | 1 |
| YPR165W | 102.945771 | 0.18902098 | 0.64013073 | 2.11010353 | 0.3481744 | 0.52188793 | RHO1 | XVI | 875368 | 875997 | 1 |
| YPR166C | 288.045616 | 2.67619204 | 0.49040078 | 30.8417151 | 2.01E-07 | 4.39E-06 | MRP2 | XVI | 876282 | 876629 | -1 |
| YPR167C | 96.1612302 | 1.8661734 | 0.55826657 | 12.1754355 | 0.00227058 | 0.01229812 | MET16 | XVI | 876847 | 877632 | -1 |
| YPR169W | 37.9062945 | -0.5040095 | 0.72244134 | 0.58581143 | 0.74609248 | 0.84537681 | JIP5 | XVI | 878690 | 880168 | 1 |
| YPR170W-B | 49.3001327 | 1.82471809 | 0.66478085 | 7.86136649 | 0.01963026 | 0.06510155 | NA | XVI | 883239 | 883595 | 1 |
| YPR171W | 224.073624 | 0.31118826 | 0.50618991 | 3.30318183 | 0.19174462 | 0.34826777 | BSP1 | XVI | 883828 | 885558 | 1 |
| YPR172W | 510.264979 | -1.0196814 | 0.60960648 | 3.54759931 | 0.16968701 | 0.31992439 | NA | XVI | 885796 | 886398 | 1 |
| YPR173C | 86.2053314 | -0.1971209 | 0.54456644 | 0.95865268 | 0.61920038 | 0.76021421 | VPS4 | XVI | 886524 | 887837 | -1 |
| YPR174C | 9.10928838 | 0.64183116 | 0.93434959 | 0.60505586 | 0.73894785 | 0.84040603 | NA | XVI | 888043 | 888708 | -1 |
| YPR175W | 730.649462 | 0.21057183 | 0.74631672 | 7.55983192 | 0.02282461 | 0.0730943 | DPB2 | XVI | 888974 | 891043 | 1 |
| YPR176C | 158.503041 | -1.9235683 | 0.56341551 | 11.0143749 | 0.0040575 | 0.01946391 | BET2 | XVI | 891101 | 892078 | -1 |
| YPR178W | 60.5813449 | 0.42079058 | 0.5735503 | 0.77860068 | 0.67753075 | 0.8011991 | PRP4 | XVI | 892332 | 893729 | 1 |
| YPR179C | 69.6040016 | -0.9150216 | 0.71575977 | 5.99677514 | 0.04986741 | 0.13023559 | HDA3 | XVI | 893797 | 895764 | -1 |
| YPR180W | 26.4959684 | 0.26268314 | 0.71838746 | 0.17658108 | 0.91549485 | 0.94807998 | AOS1 | XVI | 895961 | 897004 | 1 |
| YPR181C | 333.261303 | 4.153908 | 0.54933489 | 57.8245057 | 2.78E-13 | 2.12E-11 | SEC23 | XVI | 897361 | 899667 | -1 |
| YPR182W | 17.8289841 | -1.3520115 | 0.88723935 | 2.46294141 | 0.29186302 | 0.46616698 | SMX3 | XVI | 900194 | 900454 | 1 |
| YPR183W | 71.2488422 | 0.39669563 | 0.61459549 | 0.9146666 | 0.63296933 | 0.76993539 | DPM1 | XVI | 900755 | 901558 | 1 |
| YPR184W | 29.7395926 | 1.25529297 | 0.83471192 | 2.92046462 | 0.23218233 | 0.39938672 | GDB1 | XVI | 902044 | 906654 | 1 |

| | | | | | | | | | | | |
|---|---|---|---|---|---|---|---|---|---|---|---|
| YPR185W | 32.4305316 | 1.26340976 | 0.69535695 | 3.5864862 | 0.16641958 | 0.31561394 | ATG13 | XVI | 907218 | 909434 | 1 |
| YPR186C | 57.2341255 | 1.91193953 | 0.67115923 | 8.06810298 | 0.01770246 | 0.06025706 | PZF1 | XVI | 909733 | 911022 | -1 |
| YPR187W | 61.8118719 | -2.8575831 | 0.70004076 | 15.5476264 | 0.00042061 | 0.00320015 | RPO26 | XVI | 911257 | 911800 | 1 |
| YPR188C | 164.438319 | -3.3084618 | 0.50821451 | 37.393422 | 7.59E-09 | 2.39E-07 | MLC2 | XVI | 911988 | 912479 | -1 |
| YPR189W | 73.763073 | -1.5193506 | 0.56796427 | 7.90556774 | 0.01920117 | 0.06416508 | SKI3 | XVI | 912664 | 916962 | 1 |
| YPR190C | 64.3540037 | -0.8860722 | 0.69659558 | 6.14213605 | 0.0463716 | 0.123331 | RPC82 | XVI | 917077 | 919041 | -1 |
| YPR191W | 69.768793 | 1.11145903 | 0.88541705 | 1.68268579 | 0.43113117 | 0.59977162 | QCR2 | XVI | 919381 | 920487 | 1 |
| YPR192W | 1226.10006 | 2.53147401 | 0.52367359 | 22.1499196 | 1.55E-05 | 0.0001913 | AQY1 | XVI | 921860 | 922777 | 1 |
| YPR193C | 9.43111939 | 0.7177187 | 0.98684767 | 0.62824486 | 0.7304296 | 0.83678999 | HPA2 | XVI | 922909 | 923379 | -1 |
| YPR194C | 1639.24605 | 0.24976233 | 0.49194663 | 0.97986154 | 0.61266881 | 0.75555962 | OPT2 | XVI | 924304 | 926937 | -1 |
| YPR195C | 312.947221 | 0.51736031 | 0.52315954 | 0.98315896 | 0.61165953 | 0.75495815 | NA | XVI | 927965 | 928294 | -1 |
| YPR196W | 180.560311 | 2.24563789 | 0.51845501 | 24.389874 | 5.06E-06 | 7.19E-05 | NA | XVI | 931376 | 932788 | 1 |
| YPR198W | 52.9415312 | 0.51370279 | 0.58170356 | 0.80669905 | 0.66807855 | 0.79444823 | SGE1 | XVI | 934034 | 935665 | 1 |
| YPR199C | 558.329676 | 4.0976535 | 0.50369249 | 54.2806462 | 1.63E-12 | 1.14E-10 | ARR1 | XVI | 938148 | 939032 | -1 |
| YPR200C | 50.9232182 | -0.6022793 | 0.64714361 | 1.51007944 | 0.46999195 | 0.63255541 | ARR2 | XVI | 939279 | 939671 | -1 |
| YPR201W | 359.508137 | 0.02461976 | 0.49854698 | 0.40614167 | 0.81622042 | 0.88933313 | ARR3 | XVI | 939922 | 941136 | 1 |
| YPR202W | 21.6403239 | -1.7160761 | 0.74019643 | 6.71137743 | 0.03488534 | 0.10019152 | NA | XVI | 943032 | 943896 | 1 |
